## Supplemental Table S6c for "Leaf movements as a quantitative metric for early stress detection"

**Supplementary Table S6.** 1 h integrated motion data in **(c)** 100 mM KCl treatment under dimGday/night and RGBday/dimGnight with their respective controls.

**Supplementary Table S6.** 1 h integrated motion data in **(c)** 100 mM KCl treatment under dimGday/night with their respective controls.

| IntervalMid | Control1 | Control2 | Control3 | Control4 | Control5 | Control6 | Control7 | Control8 | Control9 | Stress1 | Stress2 | Stress3 | Stress4 | Stress5 | Stress6 | Stress7 | Stress8 | Stress9 | Stress10 |
|---|---|---|---|---|---|---|---|---|---|---|---|---|---|---|---|---|---|---|---|
| 18.1111111 | 0.00551175 | 0.02524301 | 0.0064888 | 0.0081837 | 0.01429981 | 0.01117075 | 0.00256811 | 0.02888294 | 0.05418874 | 0.00760157 | 0.00923628 | 0.00696484 | 0.04740143 | 0.0052615 | 0.0785469 | 0.01904748 | 0.01869099 | 0.01619429 | 0.00252306 |
| 18.125 | 0.00461097 | 0.02133304 | 0.00364931 | 0.0045687 | 0.01237378 | 0.01310632 | 0.00051808 | 0.02117326 | 0.05002349 | 0.00337395 | 0.00226006 | 0.00870087 | 0.03292852 | 0.00555243 | 0.07046169 | 0.02233832 | 0.02900732 | 0.01362587 | 0.00197823 |
| 18.1388889 | 0.00464432 | 0.01045982 | 0.00106283 | 0.00144431 | 0.00534633 | 0.00821231 | 0.00084424 | 0.01034543 | 0.03155305 | 0.00209343 | 0.00143327 | 0.00793685 | 0.01674154 | 0.00328707 | 0.04623718 | 0.01597558 | 0.02840111 | 0.0085713 | 0.00226027 |
| 18.1527778 | 0.00371663 | 0.00360124 | 0.00029997 | 0.00030297 | 0.00121931 | 0.00388395 | 0.0021379 | 0.00381766 | 0.01613575 | 0.00340957 | 0.00135808 | 0.00457004 | 0.00791446 | 0.00144211 | 0.02759954 | 0.00952008 | 0.01912345 | 0.00518254 | 0.00183296 |
| 18.1666667 | 0.00331441 | 0.00191205 | 0.00054738 | 0.00033313 | 0.00018176 | 0.00213052 | 0.00319395 | 0.0011959 | 0.00695094 | 0.00635556 | 0.00110558 | 0.00174489 | 0.00389004 | 0.0019102 | 0.01691957 | 0.00547736 | 0.01015508 | 0.00367902 | 0.00154795 |
| 18.1805556 | 0.00400898 | 0.00160741 | 0.00115819 | 0.001361 | 0.00033091 | 0.00138196 | 0.00176203 | 0.0006336 | 0.0033204 | 0.00995092 | 0.00130609 | 0.00130088 | 0.00245522 | 0.00321259 | 0.0092121 | 0.00347417 | 0.00511767 | 0.00291324 | 0.00177359 |
| 18.1944444 | 0.00557449 | 0.00151386 | 0.00158324 | 0.00198901 | 0.0009708 | 0.00160316 | 0.00936338 | 0.00119452 | 0.00450616 | 0.01559183 | 0.00325611 | 0.00336895 | 0.0036814 | 0.00449493 | 0.00430372 | 0.0048066 | 0.0032755 | 0.00439863 | 0.00406284 |
| 18.2083333 | 0.00625641 | 0.00193123 | 0.00625048 | 0.00820224 | 0.00136976 | 0.00249863 | 0.01003346 | 0.00186252 | 0.00492818 | 0.0211578 | 0.00464493 | 0.00493805 | 0.00576716 | 0.00812017 | 0.0037385 | 0.0071914 | 0.00320651 | 0.00741876 | 0.0071518 |
| 18.2222222 | 0.00429455 | 0.00375459 | 0.01154222 | 0.0181313 | 0.00116451 | 0.00291395 | 0.01683211 | 0.0017629 | 0.00278036 | 0.0232392 | 0.00310991 | 0.00370301 | 0.00575921 | 0.01039798 | 0.00398976 | 0.00558518 | 0.00368081 | 0.00678534 | 0.00661789 |
| 18.2361111 | 0.0024739 | 0.00557801 | 0.00726695 | 0.01487607 | 0.00102933 | 0.00311076 | 0.00792467 | 0.00185961 | 0.00316472 | 0.02423894 | 0.00318829 | 0.00261193 | 0.00458514 | 0.00651398 | 0.004861 | 0.00265619 | 0.00677289 | 0.00335695 | 0.00328902 |
| 18.25 | 0.0029234 | 0.00581946 | 0.00184588 | 0.00521015 | 0.00080805 | 0.00322425 | 0.00220939 | 0.00237071 | 0.00697709 | 0.02644879 | 0.00837182 | 0.0039815 | 0.00497061 | 0.00376013 | 0.00740111 | 0.00292856 | 0.01247091 | 0.00191365 | 0.00154256 |
| 18.2638889 | 0.00286702 | 0.00406145 | 0.00116578 | 0.0021093 | 0.00037399 | 0.00257329 | 0.00190271 | 0.00187719 | 0.008721 | 0.02670316 | 0.01382552 | 0.00558901 | 0.00539593 | 0.00499677 | 0.00751585 | 0.00403121 | 0.01457929 | 0.00189733 | 0.00158767 |
| 18.2777778 | 0.00227753 | 0.00238627 | 0.00071051 | 0.00128921 | 0.00064281 | 0.0017266 | 0.0015198 | 0.00091467 | 0.0063006 | 0.02411607 | 0.01457056 | 0.00574303 | 0.00362927 | 0.00557259 | 0.0050408 | 0.00344077 | 0.01118795 | 0.00123371 | 0.00138202 |
| 18.2916667 | 0.00230459 | 0.00229434 | 0.00048999 | 0.0006649 | 0.00172008 | 0.00170044 | 0.00165808 | 0.00107093 | 0.00459493 | 0.02153023 | 0.0121079 | 0.0048663 | 0.00123113 | 0.00468389 | 0.00474258 | 0.00220186 | 0.00617462 | 0.00050977 | 0.0007564 |
| 18.3055556 | 0.00328189 | 0.00284777 | 0.00073998 | 0.00095323 | 0.00230025 | 0.00211765 | 0.00199977 | 0.00251109 | 0.00807527 | 0.01937292 | 0.0097803 | 0.00339147 | 0.00077443 | 0.00322779 | 0.00486356 | 0.00163434 | 0.0027056 | 0.00035829 | 0.00038247 |
| 18.3194444 | 0.00428782 | 0.00426348 | 0.00071977 | 0.00128594 | 0.00169643 | 0.00197352 | 0.00169259 | 0.00378223 | 0.01296346 | 0.01620711 | 0.00818743 | 0.00228226 | 0.00127707 | 0.00221378 | 0.00370604 | 0.00159079 | 0.00171431 | 0.00037114 | 0.00051682 |
| 18.3333333 | 0.00320187 | 0.0041813 | 0.00038222 | 0.00101843 | 0.00101199 | 0.00136497 | 0.00112422 | 0.00300043 | 0.01183155 | 0.01367359 | 0.00727413 | 0.00262347 | 0.00318875 | 0.00180082 | 0.0065454 | 0.00159947 | 0.00181816 | 0.0012641 | 0.00200189 |
| 18.3472222 | 0.00189498 | 0.00239449 | 0.0002698 | 0.00051901 | 0.00120312 | 0.00105268 | 0.00124266 | 0.00124626 | 0.00624978 | 0.01380139 | 0.00795803 | 0.00318853 | 0.00903788 | 0.00127806 | 0.01357656 | 0.00236518 | 0.00260672 | 0.0033141 | 0.00541048 |
| 18.3611111 | 0.0026257 | 0.00255334 | 0.00018749 | 0.00027221 | 0.00145088 | 0.0008643 | 0.00150673 | 0.00060872 | 0.00311383 | 0.01241471 | 0.0070172 | 0.00526104 | 0.01088763 | 0.002226 | 0.01384482 | 0.00525288 | 0.00376816 | 0.00381707 | 0.00777512 |
| 18.375 | 0.00308638 | 0.00340011 | 0.00012155 | 0.00018478 | 0.00099681 | 0.00067765 | 0.00105147 | 0.0012127 | 0.00245051 | 0.01624103 | 0.01293695 | 0.01362201 | 0.02073261 | 0.00909102 | 0.02030116 | 0.02074981 | 0.00366998 | 0.00637093 | 0.01714276 |
| 18.3888889 | 0.00274151 | 0.00361975 | 0.00024355 | 0.00027356 | 0.00047092 | 0.0009033 | 0.00063931 | 0.00268207 | 0.00261665 | 0.0339338 | 0.02543275 | 0.02492005 | 0.05510746 | 0.01850692 | 0.04699548 | 0.04715611 | 0.00691626 | 0.01322115 | 0.02909013 |
| 18.4027778 | 0.0039726 | 0.00474993 | 0.00046119 | 0.00077333 | 0.00025653 | 0.0009216 | 0.00101608 | 0.00375157 | 0.00351332 | 0.04051632 | 0.02137986 | 0.0264234 | 0.07112945 | 0.01845458 | 0.05221272 | 0.04939888 | 0.01205036 | 0.01290817 | 0.02118315 |
| 18.4166667 | 0.00400916 | 0.0040433 | 0.00049868 | 0.00102474 | 0.0002661 | 0.00042575 | 0.00140395 | 0.00287663 | 0.00294498 | 0.02241689 | 0.00786912 | 0.01570248 | 0.04265079 | 0.00915501 | 0.02413441 | 0.0235395 | 0.00904535 | 0.00503568 | 0.00661485 |
| 18.4305556 | 0.00270364 | 0.00247626 | 0.00037514 | 0.00083535 | 0.00039742 | 0.00026372 | 0.00113275 | 0.0011651 | 0.00198613 | 0.00544793 | 0.00209984 | 0.00592928 | 0.0116919 | 0.00220886 | 0.0046602 | 0.00522266 | 0.00274815 | 0.00089379 | 0.00183665 |
| 18.4444444 | 0.00254497 | 0.00230671 | 0.00060037 | 0.00093985 | 0.00067971 | 0.00073301 | 0.00061976 | 0.00078092 | 0.00169244 | 0.00191166 | 0.00099896 | 0.00281243 | 0.00284067 | 0.00058016 | 0.00293969 | 0.00141929 | 0.00126547 | 0.00089322 | 0.00071905 |
| 18.4583333 | 0.0030862 | 0.00281527 | 0.00116668 | 0.00122629 | 0.0016969 | 0.00176301 | 0.00050578 | 0.00250752 | 0.00221506 | 0.00239359 | 0.00075524 | 0.00187079 | 0.00375427 | 0.00084484 | 0.00625229 | 0.00138739 | 0.00272695 | 0.00212336 | 0.00121115 |
| 18.4722222 | 0.00547881 | 0.00552264 | 0.00238097 | 0.00226252 | 0.00357656 | 0.00395518 | 0.00136913 | 0.00587004 | 0.00576303 | 0.00191403 | 0.00093202 | 0.00214395 | 0.00546309 | 0.00115113 | 0.01113391 | 0.00336603 | 0.00503942 | 0.00490341 | 0.00224113 |
| 18.4861111 | 0.00717741 | 0.00794571 | 0.00415518 | 0.00448614 | 0.00545107 | 0.00698503 | 0.00305218 | 0.00869491 | 0.00979515 | 0.00173273 | 0.00223224 | 0.00469622 | 0.006404 | 0.00221247 | 0.01435409 | 0.00537795 | 0.00742912 | 0.00686394 | 0.00231342 |
| 18.5 | 0.00655423 | 0.00786761 | 0.00493181 | 0.00580064 | 0.00641166 | 0.00859763 | 0.00423969 | 0.00920617 | 0.01072298 | 0.00303404 | 0.00452389 | 0.00777547 | 0.0060903 | 0.00518582 | 0.01337932 | 0.00578338 | 0.00874318 | 0.00621628 | 0.00239597 |
| 18.5138889 | 0.00879113 | 0.01049511 | 0.00401781 | 0.00468281 | 0.00608303 | 0.00760349 | 0.0037907 | 0.00912699 | 0.01245404 | 0.00466514 | 0.00830247 | 0.00934069 | 0.00654994 | 0.00817932 | 0.01211811 | 0.00629262 | 0.00923694 | 0.00561065 | 0.00397386 |
| 18.5277778 | 0.02003915 | 0.02349991 | 0.00226381 | 0.00237899 | 0.00479618 | 0.00562379 | 0.00291338 | 0.01221899 | 0.02077977 | 0.00476001 | 0.01674103 | 0.01123944 | 0.01063206 | 0.01144804 | 0.01577996 | 0.00871513 | 0.01219338 | 0.00775199 | 0.00669912 |
| 18.5416667 | 0.05057455 | 0.04180542 | 0.00182155 | 0.00117192 | 0.0050098 | 0.00663868 | 0.0023674 | 0.01896781 | 0.03813388 | 0.00397036 | 0.04250254 | 0.01789746 | 0.01860293 | 0.0182435 | 0.02580998 | 0.01385901 | 0.02208546 | 0.01600421 | 0.01013862 |
| 18.5555556 | 0.09224539 | 0.05694923 | 0.00367235 | 0.00303223 | 0.00841967 | 0.01263523 | 0.003248 | 0.02223446 | 0.05718832 | 0.01023053 | 0.07956235 | 0.03041586 | 0.0267441 | 0.02602365 | 0.03629638 | 0.02165847 | 0.03550432 | 0.0290814 | 0.01419189 |
| 18.5694444 | 0.10140166 | 0.0598849 | 0.00467073 | 0.00882253 | 0.01367461 | 0.01675934 | 0.00613175 | 0.01564484 | 0.05595705 | 0.02303753 | 0.08618357 | 0.03829771 | 0.03011402 | 0.02805633 | 0.03620299 | 0.02986895 | 0.03798889 | 0.03389927 | 0.01743467 |
| 18.5833333 | 0.07048372 | 0.04187014 | 0.00293709 | 0.01379187 | 0.01763836 | 0.0129072 | 0.00623562 | 0.00626096 | 0.03429447 | 0.02738872 | 0.05593381 | 0.03119076 | 0.02745854 | 0.02445686 | 0.02693563 | 0.03645984 | 0.02670011 | 0.02656019 | 0.01658665 |
| 18.5972222 | 0.0398483 | 0.02034027 | 0.00098363 | 0.01270418 | 0.01663994 | 0.0066413 | 0.00338009 | 0.00251547 | 0.01725491 | 0.02114169 | 0.02627874 | 0.01700931 | 0.02077747 | 0.021789 | 0.01822515 | 0.04037493 | 0.0150863 | 0.01759824 | 0.0117614 |
| 18.6111111 | 0.02199696 | 0.00855453 | 0.0005183 | 0.00920736 | 0.01186341 | 0.00378209 | 0.00163424 | 0.00195303 | 0.00886867 | 0.01399584 | 0.0119318 | 0.00744403 | 0.01361679 | 0.02230569 | 0.01236145 | 0.04256351 | 0.00893672 | 0.01108793 | 0.00755515 |
| 18.625 | 0.01171175 | 0.00234524 | 0.00076269 | 0.0067301 | 0.00758118 | 0.00326373 | 0.00091601 | 0.00127922 | 0.00416648 | 0.01068699 | 0.00713923 | 0.00355872 | 0.00975383 | 0.02190249 | 0.00889139 | 0.04401328 | 0.00598256 | 0.00617774 | 0.00506572 |
| 18.6388889 | 0.01066521 | 0.00102194 | 0.0007716 | 0.00463742 | 0.00578139 | 0.00340528 | 0.00045676 | 0.00133137 | 0.00645814 | 0.01013376 | 0.00621841 | 0.00204359 | 0.00887325 | 0.01868041 | 0.00687871 | 0.0405718 | 0.00489114 | 0.0028624 | 0.00304729 |
| 18.6527778 | 0.01244659 | 0.00139688 | 0.00060444 | 0.00376306 | 0.00597306 | 0.00358241 | 0.00039431 | 0.00212831 | 0.0125758 | 0.00931739 | 0.00459107 | 0.0014211 | 0.0079087 | 0.0147669 | 0.00563789 | 0.03218619 | 0.00443039 | 0.00112361 | 0.00165997 |
| 18.6666667 | 0.00894244 | 0.00193201 | 0.0006949 | 0.0044597 | 0.00602913 | 0.00262702 | 0.00054618 | 0.00197174 | 0.01643093 | 0.00722876 | 0.00256025 | 0.00241859 | 0.00601318 | 0.01129643 | 0.00583373 | 0.02577813 | 0.00400836 | 0.00104645 | 0.00128411 |
| 18.6805556 | 0.005531 | 0.00420892 | 0.00074637 | 0.00533408 | 0.00452262 | 0.00148334 | 0.00070786 | 0.00169263 | 0.01598349 | 0.00414061 | 0.00263079 | 0.00398667 | 0.00464084 | 0.00837271 | 0.00665416 | 0.02222623 | 0.00378005 | 0.00251221 | 0.00122601 |
| 18.6944444 | 0.00899526 | 0.00625374 | 0.00068528 | 0.00515807 | 0.00220473 | 0.00196824 | 0.00084839 | 0.00187938 | 0.01109591 | 0.00196164 | 0.00460649 | 0.00441471 | 0.00498891 | 0.00603293 | 0.0067448 | 0.01768793 | 0.00382862 | 0.00560985 | 0.00079958 |
| 18.7083333 | 0.01075815 | 0.00532939 | 0.00117779 | 0.00387774 | 0.00080088 | 0.00243159 | 0.00088722 | 0.0019072 | 0.00814533 | 0.0022784 | 0.00588956 | 0.00399294 | 0.00648868 | 0.00410321 | 0.00553605 | 0.01214759 | 0.00375668 | 0.0086444 | 0.00065494 |
| 18.7222222 | 0.00672562 | 0.00257531 | 0.00185072 | 0.00267551 | 0.00067011 | 0.00169887 | 0.00071115 | 0.0020724 | 0.01395205 | 0.00304671 | 0.00526568 | 0.00304714 | 0.00721716 | 0.00272768 | 0.00333186 | 0.00655678 | 0.00331693 | 0.01012237 | 0.00073122 |
| 18.7361111 | 0.0035845 | 0.00127199 | 0.00160777 | 0.00267854 | 0.00082206 | 0.00088004 | 0.00050395 | 0.00192299 | 0.01898844 | 0.00255231 | 0.00372136 | 0.00177239 | 0.00587259 | 0.00210841 | 0.00152292 | 0.00349816 | 0.00279684 | 0.0105634 | 0.00056458 |
| 18.75 | 0.00235993 | 0.00298142 | 0.00083422 | 0.0027536 | 0.00069535 | 0.00086354 | 0.00061354 | 0.00174177 | 0.01397336 | 0.00142491 | 0.00245093 | 0.00076784 | 0.00332796 | 0.00224529 | 0.00159629 | 0.00587594 | 0.00245928 | 0.01008652 | 0.00047098 |
| 18.7638889 | 0.00189847 | 0.00664001 | 0.00072795 | 0.00191009 | 0.00044097 | 0.00166483 | 0.00065781 | 0.00150922 | 0.00560877 | 0.00053633 | 0.00160697 | 0.00031105 | 0.00130086 | 0.00296853 | 0.00237958 | 0.00981717 | 0.00229304 | 0.0080376 | 0.00053924 |
| 18.7777778 | 0.00372083 | 0.00976205 | 0.00125276 | 0.00179856 | 0.0005674 | 0.00237499 | 0.00041231 | 0.00176947 | 0.00194852 | 0.0001876 | 0.00098197 | 0.00063782 | 0.00043685 | 0.00361545 | 0.00264288 | 0.01007148 | 0.00205588 | 0.00475181 | 0.00047302 |
| 18.7916667 | 0.0053901 | 0.01054421 | 0.00155871 | 0.00240175 | 0.00118286 | 0.00244361 | 0.00028528 | 0.0025429 | 0.00480631 | 0.00020358 | 0.00066363 | 0.00182904 | 0.00028896 | 0.00350861 | 0.00450337 | 0.00652796 | 0.00133083 | 0.00183363 | 0.00039738 |
| 18.8055556 | 0.00503615 | 0.00840522 | 0.00112494 | 0.00203086 | 0.00154209 | 0.0018338 | 0.00056927 | 0.00271051 | 0.01236659 | 0.00049428 | 0.00066651 | 0.00305633 | 0.00045794 | 0.00285814 | 0.00793535 | 0.00255778 | 0.00068403 | 0.0004382 | 0.00047213 |
| 18.8194444 | 0.00631268 | 0.00470086 | 0.00058259 | 0.00196252 | 0.00117795 | 0.00097573 | 0.00128733 | 0.00351393 | 0.01896216 | 0.00078002 | 0.00077946 | 0.00345639 | 0.0011303 | 0.00240009 | 0.00902382 | 0.00064227 | 0.00069075 | 0.00041551 | 0.00047907 |
| 18.8333333 | 0.00875042 | 0.00260686 | 0.00063533 | 0.00289763 | 0.0005306 | 0.00078172 | 0.00218096 | 0.00437329 | 0.01770943 | 0.00077145 | 0.0007822 | 0.00291868 | 0.00164713 | 0.00257109 | 0.00664697 | 0.00039229 | 0.00058304 | 0.00179883 | 0.00030176 |
| 18.8472222 | 0.00760269 | 0.00372861 | 0.00057014 | 0.00280503 | 0.00024929 | 0.00137929 | 0.00266553 | 0.00324215 | 0.0109107 | 0.00067217 | 0.00076038 | 0.0018856 | 0.00250843 | 0.00308027 | 0.00328545 | 0.00111081 | 0.00035758 | 0.00482804 | 0.00019292 |
| 18.8611111 | 0.00424105 | 0.00706787 | 0.00046502 | 0.00171629 | 0.00027378 | 0.0020159 | 0.00217134 | 0.00238167 | 0.00909544 | 0.0008491 | 0.00102563 | 0.00143424 | 0.00435087 | 0.00329716 | 0.00161554 | 0.0023783 | 0.00073574 | 0.00703046 | 0.00047525 |
| 18.875 | 0.00330263 | 0.0101815 | 0.00103866 | 0.00143328 | 0.00021844 | 0.00209914 | 0.00107375 | 0.00362087 | 0.01164552 | 0.00089113 | 0.00142595 | 0.00204249 | 0.00418568 | 0.00305009 | 0.00261497 | 0.00369708 | 0.00160772 | 0.00548814 | 0.00091204 |
| 18.8888889 | 0.00323524 | 0.01149841 | 0.00141748 | 0.00180347 | 0.00032469 | 0.00139457 | 0.00041755 | 0.00382507 | 0.01000311 | 0.00073964 | 0.00151267 | 0.00279075 | 0.0037797 | 0.0025679 | 0.005706 | 0.00545781 | 0.00234779 | 0.00441936 | 0.00082731 |
| 18.9027778 | 0.00362982 | 0.01131033 | 0.001288 | 0.00209684 | 0.00101931 | 0.0007116 | 0.00043358 | 0.00243573 | 0.00960756 | 0.00070688 | 0.00154986 | 0.00313961 | 0.00708541 | 0.00205906 | 0.01022055 | 0.00783796 | 0.00285397 | 0.00884224 | 0.00034767 |
| 18.9166667 | 0.00831276 | 0.01092954 | 0.0018001 | 0.00219603 | 0.00184638 | 0.00082781 | 0.00061779 | 0.00326325 | 0.01603705 | 0.00097165 | 0.00200106 | 0.00362835 | 0.01105172 | 0.00163025 | 0.01340321 | 0.00955188 | 0.00377754 | 0.01405244 | 0.00035332 |
| 18.9305556 | 0.01391162 | 0.01074058 | 0.00250518 | 0.00190727 | 0.00212217 | 0.00080877 | 0.00087489 | 0.0060088 | 0.02092736 | 0.00222521 | 0.00240117 | 0.00415425 | 0.01368912 | 0.00146111 | 0.01275721 | 0.00970243 | 0.00506407 | 0.01311291 | 0.00101385 |
| 18.9444444 | 0.01464933 | 0.01216721 | 0.00212977 | 0.00216451 | 0.00203056 | 0.00047301 | 0.00100844 | 0.0060789 | 0.02053712 | 0.00385007 | 0.00218576 | 0.00381149 | 0.01673592 | 0.00175568 | 0.0106045 | 0.00894617 | 0.00569707 | 0.00785679 | 0.00159927 |
| 18.9583333 | 0.01315623 | 0.01545175 | 0.00166401 | 0.00336152 | 0.00237329 | 0.00112586 | 0.00107331 | 0.00334063 | 0.02009794 | 0.00523605 | 0.00174911 | 0.0027543 | 0.01980687 | 0.00217188 | 0.01008736 | 0.00816578 | 0.0057252 | 0.00526773 | 0.00155536 |
| 18.9722222 | 0.01478905 | 0.0164536 | 0.00291611 | 0.00434246 | 0.00383792 | 0.00325725 | 0.00174802 | 0.00224477 | 0.02233422 | 0.0066105 | 0.0016005 | 0.00225172 | 0.01889583 | 0.00205376 | 0.01043829 | 0.00767655 | 0.00633437 | 0.00442073 | 0.00100427 |
| 18.9861111 | 0.01695456 | 0.01576564 | 0.00539186 | 0.00514421 | 0.00548772 | 0.00524786 | 0.00275881 | 0.00309203 | 0.02273818 | 0.00774426 | 0.00165008 | 0.00284618 | 0.01177964 | 0.00147285 | 0.00940912 | 0.00715951 | 0.00765291 | 0.00411542 | 0.0004766 |
| 19 | 0.01631124 | 0.01763789 | 0.00688546 | 0.0053266 | 0.00549184 | 0.00504856 | 0.00282727 | 0.00384652 | 0.01737546 | 0.0086762 | 0.00177853 | 0.00396346 | 0.00386345 | 0.00101036 | 0.00738542 | 0.00637838 | 0.00871119 | 0.00861545 | 0.00024384 |
| 19.0138889 | 0.01806408 | 0.01911212 | 0.00679723 | 0.00359193 | 0.00453641 | 0.00323597 | 0.00220406 | 0.00667919 | 0.01216809 | 0.00949243 | 0.00211931 | 0.00520557 | 0.00218038 | 0.00067017 | 0.00709199 | 0.00619432 | 0.00949605 | 0.01474718 | 0.00040448 |
| 19.0277778 | 0.02549885 | 0.01518332 | 0.00649841 | 0.0027281 | 0.00516332 | 0.00256556 | 0.0022569 | 0.01345121 | 0.01868498 | 0.01169563 | 0.00451823 | 0.00781817 | 0.00878026 | 0.00126963 | 0.01183157 | 0.00944988 | 0.0116637 | 0.0195238 | 0.00244058 |
| 19.0416667 | 0.03828099 | 0.01635075 | 0.00715229 | 0.00725434 | 0.00829779 | 0.00630014 | 0.00295327 | 0.02078976 | 0.04130388 | 0.02493212 | 0.01602498 | 0.01550845 | 0.0252581 | 0.00559482 | 0.02873845 | 0.02427101 | 0.01981579 | 0.03358064 | 0.01081801 |
| 19.0555556 | 0.0514405 | 0.0276109 | 0.0084829 | 0.01641136 | 0.01295479 | 0.01300567 | 0.00540053 | 0.02289731 | 0.06082356 | 0.04885288 | 0.04696474 | 0.02869332 | 0.03525307 | 0.01271065 | 0.05321395 | 0.04952493 | 0.03103201 | 0.05160848 | 0.02913638 |
| 19.0694444 | 0.04714631 | 0.02616381 | 0.00787017 | 0.02143935 | 0.01654519 | 0.0150954 | 0.00857452 | 0.01568705 | 0.05111287 | 0.06561088 | 0.09508023 | 0.03924039 | 0.03823473 | 0.01325069 | 0.05088234 | 0.05738582 | 0.0284337 | 0.04347766 | 0.05101089 |
| 19.0833333 | 0.02412246 | 0.01331673 | 0.00458986 | 0.01476372 | 0.01450327 | 0.00987691 | 0.00774049 | 0.00877067 | 0.02723507 | 0.06480827 | 0.11948331 | 0.03902734 | 0.05613815 | 0.01209147 | 0.02981495 | 0.03492898 | 0.01435315 | 0.02409327 | 0.06251337 |
| 19.0972222 | 0.00627825 | 0.012861 | 0.00277859 | 0.00478361 | 0.00809221 | 0.00379507 | 0.00417475 | 0.00921057 | 0.01928459 | 0.05497199 | 0.08200925 | 0.02715895 | 0.0598251 | 0.01662551 | 0.02798879 | 0.01117859 | 0.00722162 | 0.02337319 | 0.05431104 |
| 19.1111111 | 0.00200881 | 0.01845062 | 0.00257656 | 0.00165596 | 0.00453942 | 0.0013277 | 0.00190199 | 0.00689452 | 0.01609126 | 0.03862418 | 0.0265004 | 0.01144364 | 0.03675387 | 0.01311225 | 0.02632337 | 0.00430329 | 0.00505914 | 0.02348288 | 0.0301126 |
| 19.125 | 0.00336132 | 0.02471849 | 0.00259415 | 0.00157702 | 0.00404301 | 0.00088278 | 0.00139979 | 0.00367393 | 0.01088491 | 0.02708065 | 0.01155811 | 0.0023574 | 0.02309355 | 0.0046038 | 0.01342498 | 0.00556937 | 0.00229986 | 0.0173156 | 0.01110341 |
| 19.1388889 | 0.00515455 | 0.03147973 | 0.0036995 | 0.00135905 | 0.00297545 | 0.00112554 | 0.00201745 | 0.00504761 | 0.01375754 | 0.02735289 | 0.01708335 | 0.00041288 | 0.02137187 | 0.00199283 | 0.0044814 | 0.00780711 | 0.00100492 | 0.01333893 | 0.00788333 |
| 19.1527778 | 0.0046375 | 0.02693716 | 0.0031818 | 0.00154282 | 0.00179313 | 0.00127232 | 0.00193136 | 0.00479216 | 0.01185282 | 0.01454183 | 0.0122127 | 0.00044017 | 0.01671627 | 0.00120198 | 0.00177494 | 0.00721967 | 0.00057231 | 0.00975047 | 0.00614257 |
| 19.1666667 | 0.0026864 | 0.01667931 | 0.00121547 | 0.00123505 | 0.00151344 | 0.00105886 | 0.00121969 | 0.00194625 | 0.00402066 | 0.00130546 | 0.0053725 | 0.0003281 | 0.01152764 | 0.00073738 | 0.00352676 | 0.0038802 | 0.00044136 | 0.00801883 | 0.00171574 |
| 19.1805556 | 0.00154932 | 0.01013934 | 0.00030764 | 0.00100039 | 0.00138542 | 0.00099384 | 0.00110102 | 0.00075202 | 0.00175063 | 0.00111405 | 0.00282004 | 0.00028651 | 0.00918591 | 0.00059257 | 0.00527363 | 0.00122953 | 0.00041982 | 0.00837409 | 0.0004857 |
| 19.1944444 | 0.00146589 | 0.00674955 | 0.00018217 | 0.00105774 | 0.00099799 | 0.00087578 | 0.00107446 | 0.0012515 | 0.00208108 | 0.00146289 | 0.00157198 | 0.00074101 | 0.00787086 | 0.00030079 | 0.00491239 | 0.00035821 | 0.00088658 | 0.00837711 | 0.00084211 |
| 19.2083333 | 0.00159347 | 0.00728072 | 0.00022189 | 0.00074608 | 0.00088975 | 0.00060956 | 0.00085913 | 0.00146117 | 0.00202158 | 0.00239381 | 0.00102413 | 0.00115924 | 0.00740638 | 0.00022274 | 0.0038471 | 0.0006879 | 0.00168972 | 0.00754224 | 0.00073183 |
| 19.2222222 | 0.00138332 | 0.00886233 | 0.00019999 | 0.00043069 | 0.00101006 | 0.00029697 | 0.00052785 | 0.00101999 | 0.00167713 | 0.0026413 | 0.00068363 | 0.00110018 | 0.00775045 | 0.00026063 | 0.00331122 | 0.0011455 | 0.00173247 | 0.00636205 | 0.00049364 |
| 19.2361111 | 0.00176499 | 0.00686108 | 0.00032302 | 0.00056924 | 0.00094422 | 8.77E-05 | 0.0002839 | 0.00050163 | 0.00083279 | 0.00186115 | 0.00042208 | 0.00119446 | 0.00769712 | 0.00055058 | 0.00283385 | 0.00141871 | 0.00240688 | 0.00497319 | 0.00038615 |
| 19.25 | 0.00332763 | 0.00287462 | 0.00075805 | 0.00079985 | 0.00060581 | 8.50E-05 | 0.00050355 | 0.00036063 | 0.00057229 | 0.00185262 | 0.00051145 | 0.00164386 | 0.00654212 | 0.00109461 | 0.00179747 | 0.00194452 | 0.00418625 | 0.00327173 | 0.0004376 |
| 19.2638889 | 0.00481623 | 0.00056909 | 0.00120401 | 0.0007807 | 0.00047445 | 0.00017217 | 0.00091587 | 0.00034222 | 0.00136229 | 0.00226737 | 0.00105695 | 0.00155064 | 0.0049172 | 0.00133183 | 0.0008609 | 0.00236819 | 0.00430627 | 0.00196993 | 0.0007786 |
| 19.2777778 | 0.0044323 | 0.00188788 | 0.00102593 | 0.00079131 | 0.0008652 | 0.00025673 | 0.00085438 | 0.00037204 | 0.00167354 | 0.00175696 | 0.00163988 | 0.00083349 | 0.00394079 | 0.00099847 | 0.00055579 | 0.00210669 | 0.00258106 | 0.0023842 | 0.00112547 |
| 19.2916667 | 0.00251284 | 0.00705526 | 0.00040737 | 0.00119049 | 0.00133194 | 0.00052179 | 0.00070374 | 0.00055608 | 0.00152031 | 0.00105277 | 0.00193844 | 0.00034099 | 0.00384691 | 0.00082529 | 0.000478 | 0.00175397 | 0.00112878 | 0.00367666 | 0.00120646 |
| 19.3055556 | 0.00156521 | 0.01186409 | 0.00014444 | 0.00133199 | 0.00151533 | 0.00093688 | 0.00104809 | 0.00070914 | 0.0022958 | 0.00062841 | 0.00200421 | 0.00034556 | 0.00385334 | 0.00130314 | 0.00063414 | 0.00190065 | 0.00049892 | 0.00387651 | 0.0013648 |
| 19.3194444 | 0.00139332 | 0.01383831 | 0.00050549 | 0.00087148 | 0.00118388 | 0.00110084 | 0.00122983 | 0.00062838 | 0.00298669 | 0.00063548 | 0.00197063 | 0.00053606 | 0.00286608 | 0.00196388 | 0.00072536 | 0.00211223 | 0.00029502 | 0.00338552 | 0.0017807 |
| 19.3333333 | 0.00102453 | 0.01617468 | 0.00135281 | 0.00046399 | 0.00057978 | 0.00082857 | 0.00094951 | 0.00068915 | 0.00340359 | 0.00127586 | 0.00169941 | 0.00076267 | 0.00140547 | 0.00283945 | 0.00090524 | 0.00200136 | 0.00016676 | 0.00433437 | 0.00167795 |
| 19.3472222 | 0.0018247 | 0.02065732 | 0.00215099 | 0.00048222 | 0.00039581 | 0.00039578 | 0.00083771 | 0.00111038 | 0.00490265 | 0.00182984 | 0.00122522 | 0.00073799 | 0.00102183 | 0.00370013 | 0.00214331 | 0.00152518 | 0.00042163 | 0.00678667 | 0.0009519 |
| 19.3611111 | 0.00432761 | 0.02580219 | 0.00250989 | 0.0009339 | 0.00067687 | 0.00025825 | 0.00109366 | 0.00205364 | 0.00772462 | 0.00182868 | 0.00107459 | 0.00074254 | 0.00127776 | 0.00300291 | 0.00405339 | 0.00106633 | 0.00145714 | 0.00825753 | 0.00080792 |
| 19.375 | 0.00722359 | 0.0287485 | 0.00269148 | 0.00135467 | 0.00112188 | 0.00052646 | 0.00136237 | 0.00341616 | 0.0106826 | 0.00198556 | 0.00127797 | 0.00131342 | 0.00140029 | 0.0013529 | 0.00635946 | 0.00152793 | 0.00256525 | 0.00821176 | 0.00152943 |
| 19.3888889 | 0.00739944 | 0.02760163 | 0.0027946 | 0.0012514 | 0.00122024 | 0.00071347 | 0.00141666 | 0.00376565 | 0.01159776 | 0.00193634 | 0.00138915 | 0.00167512 | 0.00275736 | 0.00084223 | 0.00934595 | 0.00276447 | 0.00287724 | 0.00855231 | 0.00208077 |
| 19.4027778 | 0.00431683 | 0.02529324 | 0.00258107 | 0.00079874 | 0.00093341 | 0.00064678 | 0.0013359 | 0.00246952 | 0.01048349 | 0.00108021 | 0.0014022 | 0.00143087 | 0.00633981 | 0.00089767 | 0.011925 | 0.00377509 | 0.00250685 | 0.00994615 | 0.00172002 |
| 19.4166667 | 0.00182099 | 0.02595864 | 0.0024722 | 0.00092843 | 0.00106029 | 0.00092256 | 0.00102843 | 0.00127046 | 0.01015197 | 0.00051701 | 0.00110875 | 0.00101458 | 0.01082968 | 0.0007696 | 0.01333979 | 0.00425185 | 0.00218578 | 0.01229338 | 0.00114296 |
| 19.4305556 | 0.00161607 | 0.0277921 | 0.0030797 | 0.00247916 | 0.00157611 | 0.00173213 | 0.00097202 | 0.00131979 | 0.01152975 | 0.00059821 | 0.00078205 | 0.00086263 | 0.01406585 | 0.00063603 | 0.01428593 | 0.0045027 | 0.00248587 | 0.01501861 | 0.00134318 |
| 19.4444444 | 0.00193008 | 0.02660181 | 0.00371213 | 0.00417201 | 0.00189306 | 0.00236017 | 0.00141837 | 0.00211084 | 0.01259323 | 0.00107635 | 0.00097744 | 0.00154364 | 0.01420096 | 0.00103355 | 0.01417995 | 0.00465905 | 0.00313115 | 0.01596212 | 0.00148952 |
| 19.4583333 | 0.00242747 | 0.02443053 | 0.00385722 | 0.00457833 | 0.00256006 | 0.00281263 | 0.00162144 | 0.00361492 | 0.01296791 | 0.00208473 | 0.00112908 | 0.0026666 | 0.01161695 | 0.00252139 | 0.01239175 | 0.0044985 | 0.00352417 | 0.01486835 | 0.00101074 |
| 19.4722222 | 0.00464536 | 0.02525178 | 0.00444722 | 0.00537149 | 0.00449115 | 0.00447137 | 0.00241074 | 0.00587223 | 0.01394321 | 0.00266147 | 0.00121214 | 0.00303833 | 0.00904892 | 0.00325552 | 0.01075864 | 0.00401177 | 0.00360776 | 0.01423556 | 0.00087647 |
| 19.4861111 | 0.00750614 | 0.02677985 | 0.00556048 | 0.00768064 | 0.0070478 | 0.0072618 | 0.00437815 | 0.00796602 | 0.01521352 | 0.00249506 | 0.00152267 | 0.00253568 | 0.00847053 | 0.00238263 | 0.01032799 | 0.00356736 | 0.00353424 | 0.01475659 | 0.0015069 |
| 19.5 | 0.00877256 | 0.02627286 | 0.00596202 | 0.00897631 | 0.00826018 | 0.00872022 | 0.00560606 | 0.00904101 | 0.0163108 | 0.0030913 | 0.00195493 | 0.00252531 | 0.00966029 | 0.00196226 | 0.01087493 | 0.00391686 | 0.00383038 | 0.01526928 | 0.00252897 |
| 19.5138889 | 0.01051419 | 0.02751594 | 0.00528639 | 0.00671657 | 0.0071063 | 0.00744726 | 0.00438085 | 0.00955924 | 0.01868283 | 0.00553075 | 0.0027652 | 0.00408596 | 0.01146465 | 0.00349525 | 0.01259019 | 0.00558626 | 0.00529059 | 0.01556622 | 0.00366588 |
| 19.5277778 | 0.01822131 | 0.0342672 | 0.00504336 | 0.00408448 | 0.00523928 | 0.00494186 | 0.00270663 | 0.01103931 | 0.0264804 | 0.00893054 | 0.00442975 | 0.00654121 | 0.01400567 | 0.0072772 | 0.01416271 | 0.00772296 | 0.00892765 | 0.01723265 | 0.00508463 |
| 19.5416667 | 0.03757551 | 0.04270936 | 0.00658182 | 0.00348189 | 0.00490316 | 0.00475954 | 0.00229029 | 0.013054 | 0.04800449 | 0.01490202 | 0.00890735 | 0.01016856 | 0.01897105 | 0.0121471 | 0.01401068 | 0.00937484 | 0.0167681 | 0.0256467 | 0.00805649 |
| 19.5555556 | 0.05898741 | 0.03732659 | 0.00778033 | 0.00542224 | 0.00817293 | 0.00958776 | 0.00388105 | 0.01277171 | 0.06826368 | 0.0229628 | 0.02145317 | 0.01282912 | 0.02584821 | 0.01977693 | 0.01245687 | 0.01144626 | 0.02630733 | 0.03766368 | 0.01435412 |
| 19.5694444 | 0.06158161 | 0.0197058 | 0.00548798 | 0.010173 | 0.0148639 | 0.01625187 | 0.00854947 | 0.00975299 | 0.05787264 | 0.02330008 | 0.04148982 | 0.01226657 | 0.02633664 | 0.02649202 | 0.00844143 | 0.01482508 | 0.02748119 | 0.03542165 | 0.02234437 |
| 19.5833333 | 0.04555821 | 0.01420446 | 0.00230788 | 0.01180881 | 0.01767612 | 0.01808413 | 0.01167463 | 0.00699042 | 0.02905868 | 0.01573592 | 0.05362876 | 0.01022782 | 0.01602885 | 0.02295175 | 0.00352059 | 0.01957161 | 0.01823289 | 0.01801238 | 0.02751387 |

| | | | | | | | | | | | | | | | | | | | |
|---|---|---|---|---|---|---|---|---|---|---|---|---|---|---|---|---|---|---|---|
| 19.5972222 | 0.02632671 | 0.02524724 | 0.00239178 | 0.00974122 | 0.01330616 | 0.01393737 | 0.01058732 | 0.00654805 | 0.01096154 | 0.01126969 | 0.04760928 | 0.00607564 | 0.00881167 | 0.01559157 | 0.00368811 | 0.02495295 | 0.0085144 | 0.00648878 | 0.02551482 |
| 19.6111111 | 0.0148345 | 0.04074155 | 0.00377134 | 0.00704746 | 0.00744978 | 0.00846944 | 0.00796509 | 0.0067168 | 0.00585633 | 0.01147611 | 0.03249033 | 0.00235122 | 0.01299423 | 0.01271634 | 0.00868357 | 0.02626972 | 0.00355195 | 0.00722287 | 0.01643838 |
| 19.625 | 0.01285818 | 0.04823877 | 0.00363153 | 0.00371284 | 0.0033134 | 0.00500359 | 0.00535637 | 0.00515846 | 0.00508743 | 0.01240419 | 0.01939854 | 0.0021865 | 0.01744196 | 0.01065583 | 0.00939608 | 0.02201199 | 0.00199063 | 0.00923662 | 0.00742117 |
| 19.6388889 | 0.01279523 | 0.04229424 | 0.00251352 | 0.00153179 | 0.00119664 | 0.00349345 | 0.00343058 | 0.00363528 | 0.00500835 | 0.01338095 | 0.0116578 | 0.00327704 | 0.01304123 | 0.00713914 | 0.00783748 | 0.01807008 | 0.00230141 | 0.006664 | 0.00276463 |
| 19.6527778 | 0.00922608 | 0.02747518 | 0.00258081 | 0.000835 | 0.00065748 | 0.00277575 | 0.00218617 | 0.00510223 | 0.00611053 | 0.01519686 | 0.00878973 | 0.00338686 | 0.01052161 | 0.00413836 | 0.0151526 | 0.01719439 | 0.00311525 | 0.00479698 | 0.00097501 |
| 19.6666667 | 0.00450453 | 0.01238541 | 0.00362998 | 0.00075775 | 0.00100672 | 0.00183186 | 0.00123404 | 0.00823008 | 0.01328105 | 0.01723206 | 0.00876588 | 0.00234169 | 0.02164418 | 0.00306902 | 0.01984535 | 0.01717528 | 0.00404266 | 0.01001941 | 0.00068656 |
| 19.6805556 | 0.00255344 | 0.00581169 | 0.00307657 | 0.00106896 | 0.00253791 | 0.0011102 | 0.00050231 | 0.00926057 | 0.02683208 | 0.01729418 | 0.00914907 | 0.0021723 | 0.02899316 | 0.00302462 | 0.01240053 | 0.01697388 | 0.00525522 | 0.01767888 | 0.00101371 |
| 19.6944444 | 0.00335975 | 0.00798805 | 0.00274567 | 0.00091231 | 0.00410436 | 0.00136237 | 0.0002539 | 0.0063632 | 0.03990062 | 0.01471813 | 0.00880629 | 0.00355976 | 0.02000419 | 0.0025096 | 0.0070109 | 0.01833359 | 0.00536318 | 0.01946724 | 0.00135369 |
| 19.7083333 | 0.00593262 | 0.01105165 | 0.00504602 | 0.0007111 | 0.00336165 | 0.00194788 | 0.00043357 | 0.00300872 | 0.0421437 | 0.01231594 | 0.00835624 | 0.00422354 | 0.01333439 | 0.00161246 | 0.0094877 | 0.02070373 | 0.00365623 | 0.01426926 | 0.00152022 |
| 19.7222222 | 0.00907554 | 0.01114472 | 0.00677196 | 0.00138747 | 0.0023008 | 0.00199762 | 0.00088034 | 0.00280023 | 0.02673348 | 0.01423243 | 0.00818628 | 0.00284698 | 0.01978581 | 0.00179438 | 0.01360985 | 0.02033778 | 0.00286092 | 0.00716873 | 0.00139861 |
| 19.7361111 | 0.00831568 | 0.01434755 | 0.00566523 | 0.0016295 | 0.00427138 | 0.00156395 | 0.00160227 | 0.00291224 | 0.01114162 | 0.0199867 | 0.00781004 | 0.00199755 | 0.02848925 | 0.00279497 | 0.01339441 | 0.0168564 | 0.00402277 | 0.00495895 | 0.00117373 |
| 19.75 | 0.0063596 | 0.02175514 | 0.00323288 | 0.00160247 | 0.00668837 | 0.00153112 | 0.00179913 | 0.0017982 | 0.01292373 | 0.02327894 | 0.00650741 | 0.00407983 | 0.03141862 | 0.00314874 | 0.01111191 | 0.0133798 | 0.00470984 | 0.00942331 | 0.00135789 |
| 19.7638889 | 0.00719308 | 0.022993 | 0.00210066 | 0.00205009 | 0.00666463 | 0.00141434 | 0.00122499 | 0.00122006 | 0.02030254 | 0.01981362 | 0.00403374 | 0.00598129 | 0.02338236 | 0.00281434 | 0.01753278 | 0.01157884 | 0.00428951 | 0.01562837 | 0.00208278 |
| 19.7777778 | 0.00622468 | 0.01521165 | 0.00331511 | 0.00154994 | 0.00477777 | 0.0012163 | 0.00112303 | 0.00133799 | 0.02020022 | 0.01184627 | 0.00176786 | 0.00430911 | 0.01900215 | 0.00260491 | 0.0307379 | 0.01064903 | 0.00401722 | 0.0163353 | 0.00275195 |
| 19.7916667 | 0.00488392 | 0.01024386 | 0.00340693 | 0.00099488 | 0.00283929 | 0.00194415 | 0.00166435 | 0.00124253 | 0.01256574 | 0.00640046 | 0.00171239 | 0.00159529 | 0.03441087 | 0.00275152 | 0.03518943 | 0.00891601 | 0.00404617 | 0.01265054 | 0.0025487 |
| 19.8055556 | 0.00587263 | 0.0146782 | 0.00208758 | 0.0012968 | 0.00280441 | 0.00260986 | 0.00183963 | 0.00111359 | 0.00569107 | 0.00645224 | 0.00338408 | 0.00074834 | 0.040458 | 0.00296115 | 0.02591361 | 0.00574199 | 0.00295668 | 0.01530429 | 0.0016712 |
| 19.8194444 | 0.00621568 | 0.0205996 | 0.00253922 | 0.00109802 | 0.00385394 | 0.0024348 | 0.00127164 | 0.00245509 | 0.00713631 | 0.00677705 | 0.00415363 | 0.00091215 | 0.02442224 | 0.00295987 | 0.01347009 | 0.00263675 | 0.00099445 | 0.02016001 | 0.00139486 |
| 19.8333333 | 0.00522787 | 0.02289096 | 0.00354497 | 0.00147499 | 0.00369585 | 0.00157151 | 0.00065407 | 0.00418336 | 0.01212548 | 0.00532632 | 0.00336821 | 0.00140234 | 0.01669421 | 0.00253545 | 0.00934866 | 0.00136186 | 0.00061782 | 0.01522932 | 0.00204157 |
| 19.8472222 | 0.00469363 | 0.02383423 | 0.00393445 | 0.0039499 | 0.00224535 | 0.00087664 | 0.00062631 | 0.00336917 | 0.01113302 | 0.00331057 | 0.00194 | 0.00172088 | 0.02817718 | 0.00162077 | 0.01393326 | 0.00169531 | 0.00220194 | 0.01189357 | 0.00252878 |
| 19.8611111 | 0.006064 | 0.01869506 | 0.00429452 | 0.00628837 | 0.00131973 | 0.00114411 | 0.00096884 | 0.00182302 | 0.01322171 | 0.0018912 | 0.00124485 | 0.00120501 | 0.04160438 | 0.001065 | 0.01880196 | 0.00245315 | 0.00452703 | 0.02150996 | 0.00221704 |
| 19.875 | 0.01088485 | 0.01174202 | 0.00365726 | 0.00538716 | 0.00283673 | 0.0013178 | 0.00119935 | 0.00351222 | 0.03271937 | 0.00299889 | 0.00241221 | 0.00075033 | 0.04562196 | 0.00134675 | 0.01935873 | 0.00325959 | 0.00616247 | 0.03185366 | 0.00154544 |
| 19.8888889 | 0.01537255 | 0.01536693 | 0.00201474 | 0.00373775 | 0.00622397 | 0.00153511 | 0.00112226 | 0.00646473 | 0.04875465 | 0.00491983 | 0.0037222 | 0.0010266 | 0.03790075 | 0.00135901 | 0.01462482 | 0.00360713 | 0.00672948 | 0.0308104 | 0.00123674 |
| 19.9027778 | 0.01273929 | 0.02322088 | 0.00094631 | 0.00639769 | 0.00915935 | 0.0030927 | 0.00156117 | 0.00741237 | 0.04266346 | 0.00533643 | 0.00362608 | 0.00129126 | 0.02134352 | 0.00156388 | 0.0083566 | 0.0026132 | 0.00742468 | 0.01976203 | 0.0013719 |
| 19.9166667 | 0.008773 | 0.03001782 | 0.00096351 | 0.01107468 | 0.0108009 | 0.00544591 | 0.00276296 | 0.00865887 | 0.02668008 | 0.00395706 | 0.00281487 | 0.00164065 | 0.00971991 | 0.00317295 | 0.00859835 | 0.00179249 | 0.00775739 | 0.01051768 | 0.00120501 |
| 19.9305556 | 0.01122708 | 0.03573942 | 0.00195548 | 0.01182956 | 0.01113016 | 0.0067113 | 0.00309835 | 0.01296377 | 0.01359226 | 0.00178792 | 0.00233768 | 0.00227817 | 0.00661197 | 0.00504573 | 0.01302218 | 0.00297436 | 0.00666401 | 0.01161425 | 0.00056758 |
| 19.9444444 | 0.01686412 | 0.03324661 | 0.00319177 | 0.00907369 | 0.01039504 | 0.00608169 | 0.00188215 | 0.01853126 | 0.00927999 | 0.00064754 | 0.00290045 | 0.00209347 | 0.00578387 | 0.00603825 | 0.01328823 | 0.00394813 | 0.00517472 | 0.01320505 | 0.00017908 |
| 19.9583333 | 0.02221041 | 0.02400663 | 0.00364193 | 0.00716928 | 0.00958848 | 0.00578958 | 0.00069681 | 0.02107861 | 0.01238158 | 0.00091801 | 0.00413341 | 0.00113546 | 0.0099777 | 0.00639114 | 0.00871367 | 0.00284349 | 0.00522698 | 0.00862793 | 0.00018526 |
| 19.9722222 | 0.02521953 | 0.01630556 | 0.00373509 | 0.00902861 | 0.00954815 | 0.00789233 | 0.00142842 | 0.01910199 | 0.0154994 | 0.00183586 | 0.00469391 | 0.00062099 | 0.01751346 | 0.0068304 | 0.00486072 | 0.00168665 | 0.00723894 | 0.00718528 | 0.00040108 |
| 19.9861111 | 0.02444124 | 0.01303905 | 0.00400101 | 0.01371736 | 0.01024875 | 0.0107134 | 0.00392231 | 0.01544055 | 0.01602388 | 0.0026823 | 0.0038368 | 0.00069694 | 0.02209752 | 0.00807269 | 0.00690518 | 0.00286849 | 0.00952548 | 0.01261312 | 0.00105663 |
| 20 | 0.02273852 | 0.01399041 | 0.00348586 | 0.01548147 | 0.01105829 | 0.0112888 | 0.00549164 | 0.01264087 | 0.01548045 | 0.0024844 | 0.00212749 | 0.00075335 | 0.02314869 | 0.00986598 | 0.01400075 | 0.00636968 | 0.01013228 | 0.01795315 | 0.00194839 |
| 20.0138889 | 0.02349489 | 0.01998482 | 0.00189606 | 0.01110644 | 0.01209439 | 0.01000511 | 0.00438634 | 0.01250281 | 0.01455268 | 0.00214433 | 0.0010196 | 0.00073922 | 0.02177875 | 0.01124485 | 0.02082336 | 0.01097528 | 0.00911217 | 0.01820363 | 0.00196013 |
| 20.0277778 | 0.02699831 | 0.03373666 | 0.00080138 | 0.00513712 | 0.01369096 | 0.0102178 | 0.00202674 | 0.01708131 | 0.01796935 | 0.00723259 | 0.00313235 | 0.00224229 | 0.01842695 | 0.01216879 | 0.02533346 | 0.01603421 | 0.00866424 | 0.01498605 | 0.00230548 |
| 20.0416667 | 0.03113418 | 0.05551113 | 0.00124848 | 0.00420936 | 0.01555244 | 0.01336354 | 0.00136805 | 0.02442191 | 0.03271548 | 0.02565334 | 0.01489373 | 0.00987189 | 0.01521401 | 0.01453055 | 0.0328831 | 0.02391821 | 0.0155141 | 0.01650494 | 0.00816065 |
| 20.0555556 | 0.03625441 | 0.07022164 | 0.00282225 | 0.01033268 | 0.01806058 | 0.01641281 | 0.0035472 | 0.02769325 | 0.05090611 | 0.0516847 | 0.04000028 | 0.02203537 | 0.01620587 | 0.01858044 | 0.04753111 | 0.03532167 | 0.03301588 | 0.02702974 | 0.02137781 |
| 20.0694444 | 0.03388984 | 0.05742231 | 0.00403601 | 0.01649596 | 0.01804727 | 0.01485157 | 0.00531923 | 0.02242115 | 0.05395098 | 0.05622574 | 0.05600561 | 0.02244544 | 0.0209292 | 0.01668733 | 0.06046653 | 0.04210125 | 0.03737596 | 0.03332913 | 0.02814383 |
| 20.0833333 | 0.01764256 | 0.02816738 | 0.00412149 | 0.01439966 | 0.01118049 | 0.00899567 | 0.0039521 | 0.01350123 | 0.04096778 | 0.03468647 | 0.04195812 | 0.01373453 | 0.02523896 | 0.00922933 | 0.06132986 | 0.03723752 | 0.02346979 | 0.02379489 | 0.02176758 |
| 20.0972222 | 0.00417791 | 0.01361765 | 0.00426572 | 0.00718317 | 0.00354998 | 0.00339304 | 0.00158288 | 0.0084136 | 0.02564858 | 0.01889404 | 0.02395553 | 0.01499801 | 0.0230125 | 0.00911819 | 0.04819958 | 0.02245639 | 0.02176966 | 0.00935497 | 0.01729822 |
| 20.1111111 | 0.00160429 | 0.01556424 | 0.00368101 | 0.00294482 | 0.00219751 | 0.00169852 | 0.00128726 | 0.00559043 | 0.01297456 | 0.01314636 | 0.01963846 | 0.01784849 | 0.01632971 | 0.0163301 | 0.02685286 | 0.01065866 | 0.02679486 | 0.0033284 | 0.01505896 |
| 20.125 | 0.00178734 | 0.01401079 | 0.00238956 | 0.00317364 | 0.00398686 | 0.0037255 | 0.00331102 | 0.00315485 | 0.00469516 | 0.00683403 | 0.01479752 | 0.01256723 | 0.0141436 | 0.01892895 | 0.01115242 | 0.01044973 | 0.01945393 | 0.0028392 | 0.00933232 |
| 20.1388889 | 0.0020049 | 0.00752101 | 0.00225284 | 0.00368836 | 0.00443705 | 0.00503319 | 0.00453359 | 0.00290032 | 0.00235935 | 0.00478716 | 0.00920465 | 0.00631126 | 0.0109336 | 0.01175978 | 0.0055672 | 0.01035079 | 0.00965057 | 0.00231811 | 0.0058874 |
| 20.1527778 | 0.00238889 | 0.00439617 | 0.00165842 | 0.00234713 | 0.0029643 | 0.00346313 | 0.00301159 | 0.00218672 | 0.00116107 | 0.00473897 | 0.00535835 | 0.00333075 | 0.00540393 | 0.00370334 | 0.00340331 | 0.0049988 | 0.00792241 | 0.0025412 | 0.00385521 |
| 20.1666667 | 0.00295815 | 0.00467302 | 0.00080324 | 0.00113512 | 0.00177581 | 0.0017703 | 0.00122958 | 0.00145754 | 0.00115413 | 0.0029748 | 0.00185029 | 0.00219604 | 0.00503748 | 0.00192119 | 0.00238159 | 0.00106488 | 0.00798943 | 0.00255417 | 0.00158888 |
| 20.1805556 | 0.00391397 | 0.00764338 | 0.00148348 | 0.00104443 | 0.00080361 | 0.00131618 | 0.00097186 | 0.00255141 | 0.00309709 | 0.00198169 | 0.0006448 | 0.00098276 | 0.00508576 | 0.00217325 | 0.00322321 | 0.00046649 | 0.00463297 | 0.00214704 | 0.00084975 |
| 20.1944444 | 0.00376137 | 0.00856085 | 0.00220892 | 0.00105609 | 0.00183078 | 0.0009507 | 0.00108647 | 0.00255415 | 0.00287783 | 0.00175775 | 0.00036684 | 0.0004236 | 0.00452619 | 0.00243807 | 0.00502841 | 0.00155759 | 0.00157639 | 0.00374404 | 0.00085507 |
| 20.2083333 | 0.00308875 | 0.00530204 | 0.00162983 | 0.00075513 | 0.00102081 | 0.00060234 | 0.000975 | 0.00123467 | 0.00121917 | 0.00140499 | 0.0005529 | 0.00054147 | 0.00682426 | 0.00344016 | 0.00493294 | 0.00210444 | 0.00084469 | 0.00526291 | 0.00097037 |
| 20.2222222 | 0.00271843 | 0.00291224 | 0.00079033 | 0.00066333 | 0.0005941 | 0.00061381 | 0.00109181 | 0.0007606 | 0.00113763 | 0.00156887 | 0.00158163 | 0.00084464 | 0.00852555 | 0.00428645 | 0.00313079 | 0.00130908 | 0.00167422 | 0.0047686 | 0.00148183 |
| 20.2361111 | 0.00260503 | 0.00263893 | 0.000518 | 0.00084793 | 0.00059634 | 0.00051426 | 0.00131474 | 0.0014437 | 0.00187246 | 0.0018794 | 0.00214849 | 0.00101583 | 0.00866686 | 0.00416139 | 0.00316616 | 0.00057639 | 0.00208356 | 0.00438711 | 0.00217052 |
| 20.25 | 0.00339242 | 0.00285089 | 0.00051362 | 0.00070824 | 0.00059321 | 0.0004379 | 0.00165281 | 0.00260094 | 0.00196999 | 0.00149049 | 0.00170838 | 0.00089379 | 0.00902352 | 0.00328239 | 0.00578381 | 0.00095031 | 0.00135618 | 0.0051234 | 0.00191408 |
| 20.2638889 | 0.0030021 | 0.00371641 | 0.00081199 | 0.0003374 | 0.00069961 | 0.00078717 | 0.0019546 | 0.00263418 | 0.00249565 | 0.00095651 | 0.00109023 | 0.00076941 | 0.00934047 | 0.00254927 | 0.00910797 | 0.00188955 | 0.00058397 | 0.00650584 | 0.00092036 |
| 20.2777778 | 0.00161175 | 0.00622286 | 0.00119395 | 0.00032497 | 0.00138827 | 0.0012409 | 0.00161302 | 0.00255295 | 0.00502824 | 0.00063985 | 0.00077102 | 0.00057292 | 0.00924357 | 0.00193966 | 0.01140066 | 0.00270153 | 0.00067758 | 0.00795826 | 0.00038217 |
| 20.2916667 | 0.00179983 | 0.00834881 | 0.00160227 | 0.00088827 | 0.00219477 | 0.00163538 | 0.00121098 | 0.00470475 | 0.00872176 | 0.00046667 | 0.00050872 | 0.00034006 | 0.00901913 | 0.00106354 | 0.01213235 | 0.00306254 | 0.00130578 | 0.0079669 | 0.0003095 |
| 20.3055556 | 0.00258484 | 0.00768812 | 0.00213017 | 0.00197792 | 0.00244214 | 0.00219396 | 0.00165674 | 0.00677553 | 0.01082856 | 0.00067191 | 0.00048562 | 0.0004275 | 0.00929145 | 0.00100561 | 0.01235856 | 0.00329701 | 0.00153495 | 0.00599755 | 0.00107024 |
| 20.3194444 | 0.00204505 | 0.00525523 | 0.0018918 | 0.00233202 | 0.0020151 | 0.00207192 | 0.00202215 | 0.00547416 | 0.01041442 | 0.00352217 | 0.00422868 | 0.00219824 | 0.01303399 | 0.00330885 | 0.01527343 | 0.00486571 | 0.00106283 | 0.00352129 | 0.00549673 |
| 20.3333333 | 0.00156806 | 0.0033267 | 0.00176276 | 0.00181322 | 0.00210714 | 0.00161364 | 0.00173603 | 0.0032182 | 0.00936697 | 0.01634558 | 0.02096566 | 0.0096704 | 0.02790406 | 0.00771585 | 0.02764616 | 0.01064002 | 0.0004251 | 0.00353545 | 0.01727727 |
| 20.3472222 | 0.00260774 | 0.00344731 | 0.00234759 | 0.00155074 | 0.00248677 | 0.00184563 | 0.00184111 | 0.00271899 | 0.01030039 | 0.03815929 | 0.0482887 | 0.01907573 | 0.05224086 | 0.0139595 | 0.05041351 | 0.02230907 | 0.00018644 | 0.00994743 | 0.03127988 |
| 20.3611111 | 0.0028019 | 0.00682819 | 0.00281283 | 0.00245156 | 0.00261533 | 0.00324509 | 0.00328277 | 0.00339313 | 0.01466284 | 0.04114457 | 0.04964864 | 0.01655826 | 0.05890264 | 0.01756745 | 0.06316306 | 0.0313593 | 0.00218451 | 0.01919656 | 0.03068616 |
| 20.375 | 0.00207849 | 0.01256082 | 0.00428166 | 0.00487146 | 0.00411472 | 0.00601494 | 0.00516544 | 0.00611857 | 0.02033441 | 0.01864861 | 0.01938077 | 0.00607119 | 0.03879303 | 0.01278391 | 0.05016265 | 0.02677813 | 0.005389 | 0.02125207 | 0.014452 |
| 20.3888889 | 0.00324959 | 0.01650842 | 0.00566173 | 0.00610011 | 0.00548515 | 0.00722443 | 0.00471188 | 0.00856529 | 0.02292455 | 0.003071 | 0.0013227 | 0.00203057 | 0.02042217 | 0.0052653 | 0.02815491 | 0.01430534 | 0.00537498 | 0.01597155 | 0.0026613 |
| 20.4027778 | 0.00587853 | 0.0173797 | 0.00540653 | 0.00576375 | 0.00578198 | 0.00677484 | 0.00291695 | 0.00861305 | 0.02259718 | 0.00173011 | 0.00091999 | 0.00224868 | 0.01437973 | 0.00258858 | 0.01615256 | 0.00627735 | 0.00398322 | 0.01130957 | 0.00111706 |
| 20.4166667 | 0.00876403 | 0.01932461 | 0.00481355 | 0.00663316 | 0.00641716 | 0.00732729 | 0.00260462 | 0.00950036 | 0.02454877 | 0.00373327 | 0.00198307 | 0.00296224 | 0.01332852 | 0.00416382 | 0.01552818 | 0.00485698 | 0.00480246 | 0.01077262 | 0.00227741 |
| 20.4305556 | 0.01126073 | 0.02396752 | 0.0048333 | 0.00859667 | 0.00705655 | 0.00834128 | 0.00310449 | 0.01297322 | 0.03007387 | 0.00583216 | 0.00322734 | 0.00447735 | 0.01619043 | 0.00617621 | 0.02025545 | 0.00570759 | 0.00649256 | 0.01306711 | 0.00326165 |
| 20.4444444 | 0.01138274 | 0.02572549 | 0.00522347 | 0.01002692 | 0.00743731 | 0.00866958 | 0.00324109 | 0.01587187 | 0.03325114 | 0.00503591 | 0.00366368 | 0.00526706 | 0.02174088 | 0.00597158 | 0.02302873 | 0.00579625 | 0.00783215 | 0.01583487 | 0.0037681 |
| 20.4583333 | 0.00966621 | 0.02074684 | 0.00558372 | 0.01088042 | 0.00789654 | 0.00830221 | 0.00332963 | 0.01472171 | 0.02980475 | 0.00296235 | 0.00409243 | 0.00611382 | 0.02802196 | 0.00487194 | 0.02194077 | 0.0056048 | 0.00943049 | 0.01760299 | 0.00437092 |
| 20.4722222 | 0.01018894 | 0.01444182 | 0.00590878 | 0.0118079 | 0.00849199 | 0.00787308 | 0.00379697 | 0.01108012 | 0.02369526 | 0.00240827 | 0.00507339 | 0.00781696 | 0.0323622 | 0.00542932 | 0.02068543 | 0.00624793 | 0.01156919 | 0.01851845 | 0.00540454 |
| 20.4861111 | 0.01318282 | 0.01311427 | 0.00606167 | 0.01222836 | 0.00927844 | 0.00788552 | 0.004807 | 0.00901755 | 0.01968714 | 0.0025342 | 0.00597974 | 0.00893995 | 0.03188098 | 0.00697238 | 0.0206133 | 0.00700038 | 0.01331203 | 0.0184666 | 0.00611064 |
| 20.5 | 0.01496169 | 0.01632106 | 0.00625995 | 0.01208507 | 0.01024732 | 0.00838919 | 0.00528333 | 0.01021765 | 0.01650968 | 0.00328755 | 0.00618507 | 0.00870406 | 0.02587395 | 0.00810751 | 0.01941668 | 0.0072302 | 0.01371537 | 0.01599095 | 0.00621155 |
| 20.5138889 | 0.01618871 | 0.0219914 | 0.00697465 | 0.01174314 | 0.01049835 | 0.00919658 | 0.00582853 | 0.01217818 | 0.01395651 | 0.00752103 | 0.00626293 | 0.00901496 | 0.01716822 | 0.00991338 | 0.01695234 | 0.00821887 | 0.01368534 | 0.01173342 | 0.00674267 |
| 20.5277778 | 0.02410962 | 0.03123008 | 0.00761952 | 0.00978303 | 0.00966888 | 0.01031383 | 0.0055949 | 0.01264239 | 0.01881856 | 0.01395943 | 0.00824212 | 0.01068358 | 0.01082169 | 0.01261133 | 0.01786256 | 0.01196821 | 0.01635628 | 0.0099504 | 0.00884506 |
| 20.5416667 | 0.04787059 | 0.0485543 | 0.00764327 | 0.00932475 | 0.01287923 | 0.01333832 | 0.00815854 | 0.01438421 | 0.0390488 | 0.02167292 | 0.01573224 | 0.01310578 | 0.01273868 | 0.01682344 | 0.03077723 | 0.02243713 | 0.02743247 | 0.01494335 | 0.01582381 |
| 20.5555556 | 0.07815698 | 0.06930101 | 0.00775844 | 0.01768124 | 0.02469278 | 0.02086917 | 0.01843623 | 0.01930318 | 0.07584874 | 0.03495098 | 0.02957968 | 0.01660074 | 0.02844504 | 0.02109283 | 0.06177071 | 0.04217433 | 0.04434662 | 0.02420689 | 0.02490505 |
| 20.5694444 | 0.08091482 | 0.07140166 | 0.00746753 | 0.02705704 | 0.03290812 | 0.02684449 | 0.02741621 | 0.02289553 | 0.09986597 | 0.05012397 | 0.04124723 | 0.01881219 | 0.05262565 | 0.0189718 | 0.07988973 | 0.05239905 | 0.04981873 | 0.02787125 | 0.02487976 |
| 20.5833333 | 0.04945304 | 0.04766458 | 0.00499517 | 0.02451673 | 0.02715542 | 0.02304522 | 0.02418054 | 0.01920187 | 0.07605944 | 0.05473792 | 0.04038721 | 0.01582223 | 0.06370776 | 0.01275949 | 0.05390376 | 0.03726982 | 0.03843502 | 0.02070391 | 0.01691794 |
| 20.5972222 | 0.01957664 | 0.02099264 | 0.00402012 | 0.0157335 | 0.01715466 | 0.01479163 | 0.01635851 | 0.01116981 | 0.03873653 | 0.0447257 | 0.02870543 | 0.00875725 | 0.04580914 | 0.00921939 | 0.02206862 | 0.01568727 | 0.02319589 | 0.0101633 | 0.01049237 |
| 20.6111111 | 0.01215174 | 0.01089779 | 0.0085834 | 0.00898324 | 0.00919599 | 0.00838833 | 0.01159429 | 0.0093006 | 0.0386052 | 0.02880227 | 0.01607155 | 0.00300831 | 0.02290359 | 0.00696723 | 0.022313 | 0.00632206 | 0.0115692 | 0.00723486 | 0.00731873 |
| 20.625 | 0.01700542 | 0.01175006 | 0.01238193 | 0.00407212 | 0.00317281 | 0.00451107 | 0.00833601 | 0.01195384 | 0.05987617 | 0.01743575 | 0.00912407 | 0.00233018 | 0.02661442 | 0.00492893 | 0.03919495 | 0.00863544 | 0.00458212 | 0.01132775 | 0.00632127 |
| 20.6388889 | 0.01933747 | 0.0110937 | 0.00928423 | 0.00158514 | 0.00094629 | 0.00423926 | 0.00595321 | 0.01023262 | 0.0652795 | 0.01424187 | 0.0082686 | 0.00582984 | 0.03994629 | 0.0043911 | 0.04747489 | 0.01426412 | 0.00261312 | 0.01290582 | 0.00658418 |
| 20.6527778 | 0.01484452 | 0.01558885 | 0.00423078 | 0.00296329 | 0.0013259 | 0.00593567 | 0.00378041 | 0.01007012 | 0.04504022 | 0.01591774 | 0.00930697 | 0.00895146 | 0.03757509 | 0.00438264 | 0.04037228 | 0.01885997 | 0.00384642 | 0.00861822 | 0.0057844 |
| 20.6666667 | 0.01669638 | 0.03111484 | 0.0039669 | 0.00764178 | 0.00207607 | 0.00541307 | 0.00270271 | 0.01686981 | 0.02342565 | 0.01644728 | 0.00797227 | 0.00781874 | 0.02391222 | 0.00324627 | 0.02639901 | 0.02277154 | 0.00615959 | 0.00639296 | 0.00375644 |
| 20.6805556 | 0.03023443 | 0.04090499 | 0.00639862 | 0.01232738 | 0.00180771 | 0.00361985 | 0.00471584 | 0.02171174 | 0.02814047 | 0.01151092 | 0.00473969 | 0.00383677 | 0.01224324 | 0.00198474 | 0.01363154 | 0.02402802 | 0.00806872 | 0.0113245 | 0.00367019 |
| 20.6944444 | 0.03175351 | 0.02921678 | 0.00635445 | 0.01268008 | 0.00169267 | 0.00447117 | 0.00630286 | 0.01826096 | 0.04311906 | 0.0059966 | 0.0035521 | 0.0012516 | 0.01507638 | 0.00228638 | 0.01025305 | 0.02129654 | 0.0074467 | 0.0145608 | 0.00549346 |
| 20.7083333 | 0.01811197 | 0.01425287 | 0.00441543 | 0.00925101 | 0.00360538 | 0.00539738 | 0.00439882 | 0.01086427 | 0.03786836 | 0.00650835 | 0.00450559 | 0.00144933 | 0.03138711 | 0.00254489 | 0.01603053 | 0.01466817 | 0.00638955 | 0.0110551 | 0.00580063 |
| 20.7222222 | 0.01182676 | 0.01648592 | 0.00440286 | 0.00547946 | 0.00541218 | 0.00363684 | 0.00265676 | 0.00870778 | 0.02065224 | 0.00703748 | 0.00395477 | 0.00390684 | 0.03871182 | 0.00276513 | 0.01488822 | 0.00749878 | 0.00995086 | 0.01057752 | 0.00408306 |
| 20.7361111 | 0.01175573 | 0.01707133 | 0.00561926 | 0.00258597 | 0.00454849 | 0.00200105 | 0.00321934 | 0.01155602 | 0.01908986 | 0.00505159 | 0.00225524 | 0.00739598 | 0.02650774 | 0.00354722 | 0.00701171 | 0.0038373 | 0.0130562 | 0.01694982 | 0.00214463 |
| 20.75 | 0.00906406 | 0.01095815 | 0.00419421 | 0.00145242 | 0.00284092 | 0.00322288 | 0.00398504 | 0.01040241 | 0.03713024 | 0.00791233 | 0.00197899 | 0.0079591 | 0.01767403 | 0.00266838 | 0.00611782 | 0.00227765 | 0.00976174 | 0.01856558 | 0.00097906 |
| 20.7638889 | 0.00622012 | 0.01491775 | 0.00214527 | 0.00164599 | 0.00341993 | 0.0058963 | 0.00466665 | 0.00845267 | 0.05412851 | 0.01405764 | 0.00241222 | 0.00482318 | 0.02598709 | 0.00102298 | 0.01077693 | 0.00173486 | 0.00699317 | 0.01266809 | 0.00087086 |
| 20.7777778 | 0.00388561 | 0.02268743 | 0.00182983 | 0.00259725 | 0.00541614 | 0.00617198 | 0.0048416 | 0.00916418 | 0.05054052 | 0.01499713 | 0.00199993 | 0.00351194 | 0.03426635 | 0.00115637 | 0.01571467 | 0.0034228 | 0.00912933 | 0.01215909 | 0.00177629 |
| 20.7916667 | 0.00174945 | 0.02242536 | 0.00241376 | 0.00655873 | 0.00626655 | 0.00404077 | 0.00344325 | 0.00664403 | 0.0286712 | 0.00916474 | 0.00118735 | 0.00575687 | 0.03358978 | 0.00285916 | 0.01950254 | 0.0047926 | 0.00938978 | 0.01754562 | 0.00259997 |
| 20.8055556 | 0.00103127 | 0.01391875 | 0.00481144 | 0.01234654 | 0.00544773 | 0.00316896 | 0.00244973 | 0.00549102 | 0.01098871 | 0.00490915 | 0.00098691 | 0.00685023 | 0.02730842 | 0.00480015 | 0.01895296 | 0.00375306 | 0.00620941 | 0.01642174 | 0.00223068 |
| 20.8194444 | 0.00116518 | 0.00865979 | 0.00519163 | 0.01373453 | 0.00341598 | 0.00296933 | 0.00209115 | 0.00900607 | 0.00865919 | 0.00551529 | 0.00113347 | 0.00507003 | 0.01562785 | 0.00591779 | 0.01353185 | 0.00263224 | 0.00536041 | 0.00857852 | 0.00240876 |
| 20.8333333 | 0.00091203 | 0.01424736 | 0.00322601 | 0.00908215 | 0.00139308 | 0.0022399 | 0.00220856 | 0.00909796 | 0.00941758 | 0.00566723 | 0.00089508 | 0.00419542 | 0.00897481 | 0.00600798 | 0.00731276 | 0.00476571 | 0.00632658 | 0.00397164 | 0.00441256 |
| 20.8472222 | 0.00084789 | 0.01764067 | 0.00435536 | 0.00345595 | 0.00068507 | 0.00302777 | 0.00402927 | 0.00546927 | 0.00558654 | 0.00762884 | 0.00046044 | 0.00470905 | 0.01625244 | 0.00536033 | 0.00348254 | 0.00981205 | 0.00540413 | 0.00757474 | 0.00518002 |
| 20.8611111 | 0.00224336 | 0.01264193 | 0.00487515 | 0.00241877 | 0.00144299 | 0.00423055 | 0.00442272 | 0.00439095 | 0.00356363 | 0.01271622 | 0.00084038 | 0.00365772 | 0.02603733 | 0.00420393 | 0.00286116 | 0.0144223 | 0.00423059 | 0.01681826 | 0.00365446 |
| 20.875 | 0.00362236 | 0.01052279 | 0.00407634 | 0.00675473 | 0.00360456 | 0.00353959 | 0.00268903 | 0.00494418 | 0.01064594 | 0.01402505 | 0.00231801 | 0.00332601 | 0.02404478 | 0.00296997 | 0.00453895 | 0.01671879 | 0.00496513 | 0.02522033 | 0.00340506 |
| 20.8888889 | 0.00844606 | 0.01063573 | 0.00762678 | 0.01167484 | 0.00679134 | 0.00256343 | 0.00224359 | 0.00800915 | 0.02692933 | 0.00879886 | 0.00344711 | 0.00577485 | 0.01757903 | 0.00239716 | 0.00951886 | 0.01678391 | 0.00686451 | 0.02674261 | 0.00544844 |
| 20.9027778 | 0.02205741 | 0.01293346 | 0.01214508 | 0.0140055 | 0.00945409 | 0.00416082 | 0.00309387 | 0.01667447 | 0.04368042 | 0.00524253 | 0.00394366 | 0.00718756 | 0.02575067 | 0.00335143 | 0.01863767 | 0.01551847 | 0.0093633 | 0.02240873 | 0.0074457 |
| 20.9166667 | 0.03614123 | 0.02916531 | 0.0127658 | 0.01490288 | 0.01087229 | 0.00747251 | 0.00411459 | 0.02481097 | 0.04996324 | 0.01023817 | 0.00542278 | 0.00505609 | 0.04459188 | 0.00622966 | 0.02735109 | 0.01430658 | 0.01130989 | 0.01767238 | 0.0082458 |
| 20.9305556 | 0.04036254 | 0.05008955 | 0.01039561 | 0.01485993 | 0.0119388 | 0.00924534 | 0.00439878 | 0.02689744 | 0.04337396 | 0.0154962 | 0.00814353 | 0.00238196 | 0.05599847 | 0.01032991 | 0.03505315 | 0.01499168 | 0.0110036 | 0.01360082 | 0.00844757 |
| 20.9444444 | 0.03384796 | 0.05750196 | 0.00613255 | 0.01328598 | 0.01274865 | 0.00782188 | 0.00308349 | 0.02350317 | 0.03159968 | 0.01315554 | 0.01052112 | 0.0020453 | 0.05504633 | 0.0140237 | 0.04292393 | 0.01742242 | 0.00920705 | 0.00829887 | 0.0081768 |
| 20.9583333 | 0.02189279 | 0.04767494 | 0.00293715 | 0.01078317 | 0.01292409 | 0.00420974 | 0.00155038 | 0.01754399 | 0.02283955 | 0.0068772 | 0.01113636 | 0.00189214 | 0.04317598 | 0.01587617 | 0.04460225 | 0.01794461 | 0.00854156 | 0.00570837 | 0.00568767 |
| 20.9722222 | 0.01304315 | 0.02802821 | 0.00239606 | 0.00843412 | 0.01271949 | 0.00135065 | 0.00143061 | 0.012723 | 0.01829889 | 0.0023227 | 0.01030198 | 0.00101439 | 0.02348791 | 0.01590317 | 0.03528768 | 0.01468666 | 0.0103546 | 0.00935879 | 0.00237334 |
| 20.9861111 | 0.01464484 | 0.0125507 | 0.00345565 | 0.00792518 | 0.01231868 | 0.00208847 | 0.00159217 | 0.0119085 | 0.01650808 | 0.00215493 | 0.00966185 | 0.00125856 | 0.0105087 | 0.01529222 | 0.02028591 | 0.01062338 | 0.01302844 | 0.01199581 | 0.00130017 |
| 21 | 0.02620751 | 0.00950794 | 0.00683632 | 0.01114117 | 0.0122421 | 0.00561313 | 0.00119933 | 0.0153971 | 0.01795792 | 0.00675201 | 0.00982519 | 0.00179984 | 0.01180304 | 0.01463655 | 0.00979993 | 0.00922512 | 0.01374927 | 0.00847139 | 0.00236985 |
| 21.0138889 | 0.03795006 | 0.01540667 | 0.01007463 | 0.0152416 | 0.01193132 | 0.00767269 | 0.00226631 | 0.01932395 | 0.02179852 | 0.01482153 | 0.00986414 | 0.00131636 | 0.01406477 | 0.01333019 | 0.00946771 | 0.01131125 | 0.01242805 | 0.00418327 | 0.00538329 |
| 21.0277778 | 0.0431499 | 0.0259061 | 0.01150868 | 0.01589464 | 0.01095889 | 0.00815887 | 0.00537298 | 0.02222006 | 0.02877382 | 0.02385094 | 0.01025038 | 0.00121772 | 0.01108491 | 0.01053408 | 0.01992961 | 0.01685741 | 0.01245962 | 0.00527353 | 0.01011043 |
| 21.0416667 | 0.04543702 | 0.04306249 | 0.01349991 | 0.0153258 | 0.01265052 | 0.0129318 | 0.00719467 | 0.02941079 | 0.04422472 | 0.03747372 | 0.01523103 | 0.00411272 | 0.00950476 | 0.00718639 | 0.0453912 | 0.02791124 | 0.01886119 | 0.01334641 | 0.01950024 |
| 21.0555556 | 0.0491569 | 0.06323793 | 0.01674009 | 0.01683652 | 0.01795779 | 0.02077855 | 0.00893295 | 0.0424423 | 0.06524856 | 0.05135192 | 0.02426402 | 0.00869802 | 0.01839972 | 0.00580008 | 0.07333622 | 0.03790022 | 0.02972381 | 0.02684803 | 0.03019175 |
| 21.0694444 | 0.0466851 | 0.07139193 | 0.01855061 | 0.017203 | 0.02017765 | 0.02162628 | 0.01226378 | 0.0506869 | 0.07078464 | 0.04498358 | 0.02540794 | 0.00990004 | 0.03398018 | 0.0061393 | 0.07012412 | 0.03267703 | 0.03211043 | 0.03691638 | 0.02789325 |
| 21.0833333 | 0.03315697 | 0.05939246 | 0.01500959 | 0.01350057 | 0.01538753 | 0.01357075 | 0.01106547 | 0.0439291 | 0.04878003 | 0.02140429 | 0.01531599 | 0.00692064 | 0.03837573 | 0.00470793 | 0.03663484 | 0.01619623 | 0.02123945 | 0.03560567 | 0.01469487 |
| 21.0972222 | 0.02096211 | 0.03827827 | 0.00801862 | 0.0087506 | 0.00898544 | 0.00554466 | 0.00535272 | 0.02540917 | 0.02091887 | 0.00597825 | 0.00787449 | 0.00345294 | 0.02954591 | 0.00189268 | 0.01183891 | 0.00469238 | 0.00955572 | 0.02733917 | 0.0071119 |

| | | | | | | | | | | | | | | | | | | | |
|---|---|---|---|---|---|---|---|---|---|---|---|---|---|---|---|---|---|---|---|
| 21.1111111 | 0.0158836 | 0.0209377 | 0.00513232 | 0.00510364 | 0.00581272 | 0.00313282 | 0.00178979 | 0.01201893 | 0.0105733 | 0.00332479 | 0.00770102 | 0.00137095 | 0.0200147 | 0.00080394 | 0.00920839 | 0.00182589 | 0.00712028 | 0.01855989 | 0.00636115 |
| 21.125 | 0.01067811 | 0.01105677 | 0.00708529 | 0.00255531 | 0.00452161 | 0.00296502 | 0.00254348 | 0.01322083 | 0.01299002 | 0.00235549 | 0.00652992 | 0.00108597 | 0.01503076 | 0.00097871 | 0.00964042 | 0.00154686 | 0.00923538 | 0.01181414 | 0.00449362 |
| 21.1388889 | 0.00530197 | 0.00718587 | 0.00754228 | 0.00180806 | 0.00299544 | 0.00282088 | 0.0058829 | 0.01512902 | 0.01254248 | 0.00082832 | 0.00318577 | 0.00307147 | 0.01086443 | 0.00106356 | 0.0054002 | 0.00126217 | 0.00788581 | 0.00861377 | 0.00200291 |
| 21.1527778 | 0.00464455 | 0.00451843 | 0.00546597 | 0.00184183 | 0.0018474 | 0.00393026 | 0.00898724 | 0.00915469 | 0.00739268 | 0.00057336 | 0.00119863 | 0.00527519 | 0.00571243 | 0.0010138 | 0.00174921 | 0.00138172 | 0.00368913 | 0.00743233 | 0.00157807 |
| 21.1666667 | 0.00572659 | 0.00180002 | 0.00308322 | 0.00119278 | 0.00181487 | 0.00525721 | 0.00967372 | 0.00497464 | 0.00669453 | 0.00094525 | 0.00070156 | 0.00475814 | 0.00250545 | 0.00102785 | 0.00050785 | 0.0011522 | 0.00184993 | 0.00601753 | 0.00215922 |
| 21.1805556 | 0.00387986 | 0.00122013 | 0.00142862 | 0.00064962 | 0.00176401 | 0.00574469 | 0.00817613 | 0.00676741 | 0.01064997 | 0.00182274 | 0.00121665 | 0.00232551 | 0.00292318 | 0.00122518 | 0.00177171 | 0.00053848 | 0.00373176 | 0.00536799 | 0.00256449 |
| 21.1944444 | 0.0020554 | 0.00158962 | 0.00105628 | 0.0008602 | 0.00120593 | 0.0059814 | 0.00672304 | 0.00734829 | 0.01156098 | 0.00211419 | 0.00244511 | 0.00078964 | 0.00498532 | 0.00201523 | 0.00467065 | 0.00037567 | 0.00529341 | 0.00605091 | 0.00392849 |
| 21.2083333 | 0.00380293 | 0.00182047 | 0.00151182 | 0.00085747 | 0.00111841 | 0.00606205 | 0.00630789 | 0.00497618 | 0.00767478 | 0.00130293 | 0.00268503 | 0.00056796 | 0.00487826 | 0.00227204 | 0.00562604 | 0.00061574 | 0.00381154 | 0.00506647 | 0.00567482 |
| 21.2222222 | 0.00665301 | 0.00329879 | 0.00254793 | 0.0007903 | 0.00149357 | 0.00550031 | 0.00576417 | 0.00534182 | 0.00550477 | 0.00068412 | 0.00156356 | 0.00124903 | 0.00389504 | 0.00150158 | 0.00454234 | 0.00073024 | 0.00208683 | 0.00359506 | 0.00460748 |
| 21.2361111 | 0.00790733 | 0.00530178 | 0.00287703 | 0.00130626 | 0.00183592 | 0.00452639 | 0.00459334 | 0.00827822 | 0.00719707 | 0.00064541 | 0.0007978 | 0.002552 | 0.00645101 | 0.00159633 | 0.00586389 | 0.00058479 | 0.00205153 | 0.00566756 | 0.00221468 |
| 21.25 | 0.00791774 | 0.00451009 | 0.00182623 | 0.00127654 | 0.00219055 | 0.00359929 | 0.00415216 | 0.0091384 | 0.00604132 | 0.00086624 | 0.00099688 | 0.00255248 | 0.00965739 | 0.00282486 | 0.00802876 | 0.00050581 | 0.00226022 | 0.00808535 | 0.00153254 |
| 21.2638889 | 0.00764857 | 0.00228579 | 0.00069306 | 0.00051204 | 0.00228686 | 0.00287314 | 0.00433835 | 0.00704352 | 0.00272275 | 0.00140949 | 0.00149109 | 0.00182418 | 0.00841162 | 0.00398 | 0.00818417 | 0.00048331 | 0.00259243 | 0.00773824 | 0.00124956 |
| 21.2777778 | 0.00810106 | 0.00196092 | 0.00016765 | 0.00069953 | 0.0020964 | 0.00269343 | 0.0032413 | 0.00388435 | 0.00196289 | 0.00203538 | 0.0013937 | 0.00189031 | 0.00472914 | 0.00379787 | 0.00780028 | 0.00057708 | 0.00290404 | 0.00611332 | 0.00099236 |
| 21.2916667 | 0.00837161 | 0.00402314 | 0.00070026 | 0.00164929 | 0.00231776 | 0.003776 | 0.00129275 | 0.00153549 | 0.00247606 | 0.00193204 | 0.00075195 | 0.00152536 | 0.00317377 | 0.00214785 | 0.00741178 | 0.00106545 | 0.00229112 | 0.00409565 | 0.00181658 |
| 21.3055556 | 0.00646909 | 0.00685075 | 0.00221168 | 0.00197299 | 0.00315549 | 0.0055961 | 0.00127849 | 0.00111871 | 0.00286157 | 0.00173486 | 0.00062741 | 0.00154758 | 0.00273906 | 0.00088647 | 0.0074685 | 0.00111264 | 0.00212239 | 0.00280862 | 0.00210429 |
| 21.3194444 | 0.00557656 | 0.00779153 | 0.00260361 | 0.00281605 | 0.00380189 | 0.00606756 | 0.00414044 | 0.00163825 | 0.00389003 | 0.00223699 | 0.00148389 | 0.00276877 | 0.00210333 | 0.00114719 | 0.01026094 | 0.00093047 | 0.00365023 | 0.00417497 | 0.00202977 |
| 21.3333333 | 0.00755361 | 0.00770046 | 0.00154324 | 0.00459424 | 0.0041641 | 0.00548399 | 0.00771663 | 0.00226132 | 0.005761 | 0.00249498 | 0.00271755 | 0.00307807 | 0.00363502 | 0.00183129 | 0.01512299 | 0.00171588 | 0.00535073 | 0.00855384 | 0.00469954 |
| 21.3472222 | 0.00774108 | 0.00901138 | 0.001203 | 0.00541605 | 0.00521021 | 0.00716 | 0.01016793 | 0.00446793 | 0.00595165 | 0.00205806 | 0.00301822 | 0.00231343 | 0.00650048 | 0.00207691 | 0.01743846 | 0.00301291 | 0.00595055 | 0.01333497 | 0.00929692 |
| 21.3611111 | 0.0081554 | 0.01309679 | 0.00249301 | 0.00539991 | 0.00674583 | 0.01159804 | 0.01204069 | 0.00801233 | 0.00562907 | 0.00194795 | 0.00216463 | 0.00281343 | 0.00860769 | 0.002727 | 0.01575754 | 0.00407541 | 0.00655144 | 0.01479451 | 0.01144781 |
| 21.375 | 0.01160386 | 0.01488155 | 0.00452662 | 0.00501284 | 0.00722265 | 0.0150181 | 0.01353293 | 0.00935399 | 0.00598646 | 0.0028241 | 0.00129196 | 0.00470136 | 0.00949421 | 0.00395348 | 0.01334398 | 0.0042463 | 0.00781404 | 0.01238021 | 0.00961303 |
| 21.3888889 | 0.01329132 | 0.01145401 | 0.00491446 | 0.00365932 | 0.006137 | 0.01515264 | 0.01340068 | 0.00766801 | 0.00500709 | 0.00348591 | 0.00086549 | 0.00565529 | 0.008747 | 0.0043156 | 0.01135144 | 0.00307939 | 0.00861424 | 0.00895978 | 0.00660454 |
| 21.4027778 | 0.01360127 | 0.01184056 | 0.00341236 | 0.00289361 | 0.00510421 | 0.01419161 | 0.0120332 | 0.00850287 | 0.00723506 | 0.00311042 | 0.00068423 | 0.00561017 | 0.00701568 | 0.00379466 | 0.00843869 | 0.00183653 | 0.00844467 | 0.00645943 | 0.00571228 |
| 21.4166667 | 0.01617143 | 0.02144792 | 0.00308414 | 0.00541941 | 0.00594431 | 0.01487845 | 0.01194272 | 0.01672491 | 0.01568617 | 0.00280587 | 0.00114828 | 0.00674167 | 0.00711823 | 0.00430692 | 0.00675971 | 0.00190821 | 0.0082107 | 0.00539866 | 0.00765651 |
| 21.4305556 | 0.02062649 | 0.03470194 | 0.00575447 | 0.01057706 | 0.00816457 | 0.01669342 | 0.01346309 | 0.02718681 | 0.02332389 | 0.00342595 | 0.00294414 | 0.00887142 | 0.01163011 | 0.00701582 | 0.01004139 | 0.00280618 | 0.00868386 | 0.00684623 | 0.01080257 |
| 21.4444444 | 0.02562999 | 0.04305939 | 0.00945644 | 0.01342121 | 0.00928232 | 0.01774761 | 0.01439742 | 0.02950881 | 0.02176125 | 0.00471626 | 0.00536434 | 0.00980694 | 0.01790224 | 0.01089689 | 0.0168385 | 0.00321203 | 0.00971789 | 0.01057378 | 0.01304365 |
| 21.4583333 | 0.02861847 | 0.04072064 | 0.01012994 | 0.01127835 | 0.0083407 | 0.0175576 | 0.01372064 | 0.02045103 | 0.01436624 | 0.00576064 | 0.0065018 | 0.00866056 | 0.01977572 | 0.01318551 | 0.02082269 | 0.00271995 | 0.01033036 | 0.01384885 | 0.0135307 |
| 21.4722222 | 0.0287141 | 0.03040981 | 0.00674153 | 0.00749381 | 0.007252 | 0.01619161 | 0.01178122 | 0.00921985 | 0.01331805 | 0.00539327 | 0.00564048 | 0.00620021 | 0.01558346 | 0.01241125 | 0.01752179 | 0.00209204 | 0.00963163 | 0.01445845 | 0.01238657 |
| 21.4861111 | 0.02993966 | 0.02234966 | 0.00287175 | 0.00586882 | 0.0078858 | 0.01437555 | 0.01014599 | 0.00453572 | 0.01404762 | 0.00403573 | 0.00452906 | 0.00416017 | 0.01068349 | 0.01055831 | 0.01030997 | 0.00253186 | 0.00853772 | 0.01307685 | 0.01020729 |
| 21.5 | 0.03495167 | 0.02354073 | 0.00252547 | 0.00703132 | 0.01011567 | 0.01395049 | 0.01106309 | 0.0084863 | 0.01376539 | 0.00349235 | 0.00565262 | 0.00396511 | 0.01230601 | 0.01049763 | 0.00831157 | 0.00483049 | 0.00932985 | 0.01380082 | 0.00865569 |
| 21.5138889 | 0.04252246 | 0.03413603 | 0.00673448 | 0.00929712 | 0.01229713 | 0.01531966 | 0.01421154 | 0.02279234 | 0.02851644 | 0.00574239 | 0.00886316 | 0.00573069 | 0.02190034 | 0.01315794 | 0.01470266 | 0.00841053 | 0.01241882 | 0.01909809 | 0.00882657 |
| 21.5277778 | 0.04989205 | 0.04878164 | 0.01392166 | 0.01088934 | 0.01364225 | 0.01688242 | 0.01699667 | 0.03561211 | 0.05397158 | 0.01401032 | 0.01242 | 0.00839174 | 0.03203333 | 0.01719886 | 0.02248423 | 0.01158911 | 0.01511978 | 0.02527426 | 0.00922281 |
| 21.5416667 | 0.06230269 | 0.06375716 | 0.01959844 | 0.01653535 | 0.01873633 | 0.01767683 | 0.01870401 | 0.03417076 | 0.08250267 | 0.03142334 | 0.0176332 | 0.00970778 | 0.03749435 | 0.02161291 | 0.02924288 | 0.01539144 | 0.02008068 | 0.03211998 | 0.00910694 |
| 21.5555556 | 0.08248523 | 0.07896904 | 0.01844317 | 0.0342265 | 0.03193775 | 0.01814025 | 0.02144307 | 0.02600393 | 0.11237401 | 0.05776233 | 0.02634894 | 0.00961954 | 0.0473798 | 0.0264663 | 0.04611977 | 0.02038006 | 0.03248506 | 0.0349885 | 0.01097897 |
| 21.5694444 | 0.08698545 | 0.08270716 | 0.01205409 | 0.05036664 | 0.04022475 | 0.01480869 | 0.02027057 | 0.0203793 | 0.11752173 | 0.07403064 | 0.03271313 | 0.01136579 | 0.063196 | 0.0269585 | 0.06429525 | 0.01996875 | 0.03917252 | 0.02612396 | 0.01493833 |
| 21.5833333 | 0.06200756 | 0.06202295 | 0.00569731 | 0.04358277 | 0.032485 | 0.00900276 | 0.01298115 | 0.01592287 | 0.08757153 | 0.05945821 | 0.02856519 | 0.01302766 | 0.06356172 | 0.01865763 | 0.05872493 | 0.01218118 | 0.02810845 | 0.01705278 | 0.01702386 |
| 21.5972222 | 0.03293391 | 0.03460925 | 0.00370872 | 0.0223558 | 0.0205818 | 0.01000457 | 0.01198529 | 0.01032322 | 0.04454223 | 0.02999697 | 0.01650262 | 0.00911619 | 0.03870766 | 0.00955891 | 0.03327292 | 0.00468621 | 0.01274872 | 0.01343884 | 0.0149726 |
| 21.6111111 | 0.02844073 | 0.03283929 | 0.00909825 | 0.00808203 | 0.01299313 | 0.01490887 | 0.01743503 | 0.00851502 | 0.02206701 | 0.00986654 | 0.00605392 | 0.00375966 | 0.0165284 | 0.00938901 | 0.01468932 | 0.00208762 | 0.00487927 | 0.00784715 | 0.00994126 |
| 21.625 | 0.04321723 | 0.05136994 | 0.01529685 | 0.0058586 | 0.00790119 | 0.01505047 | 0.01699589 | 0.00945927 | 0.03754196 | 0.00276855 | 0.00213409 | 0.00231333 | 0.01950865 | 0.01380955 | 0.01688974 | 0.00348844 | 0.00280934 | 0.00287988 | 0.00631934 |
| 21.6388889 | 0.04756427 | 0.05474179 | 0.01268875 | 0.00749214 | 0.00429686 | 0.00954257 | 0.01013237 | 0.0088181 | 0.06242597 | 0.00480232 | 0.00372837 | 0.00370242 | 0.02924902 | 0.01274194 | 0.02028284 | 0.00633075 | 0.00282124 | 0.00159775 | 0.0067208 |
| 21.6527778 | 0.03423008 | 0.03501026 | 0.00897085 | 0.00893097 | 0.0032873 | 0.00479312 | 0.00615962 | 0.01566103 | 0.06712057 | 0.00992618 | 0.00712501 | 0.00783908 | 0.02815977 | 0.00755356 | 0.01334558 | 0.00837405 | 0.00304726 | 0.00229185 | 0.00713828 |
| 21.6666667 | 0.02859036 | 0.02850073 | 0.01514678 | 0.00961363 | 0.00593494 | 0.00622098 | 0.00728554 | 0.02903 | 0.05225428 | 0.01112308 | 0.00737687 | 0.0115882 | 0.01908956 | 0.00729914 | 0.01274332 | 0.00811249 | 0.0062085 | 0.00301542 | 0.00522605 |
| 21.6805556 | 0.03846942 | 0.05264934 | 0.02038159 | 0.00772366 | 0.00898162 | 0.00781081 | 0.00589989 | 0.0315886 | 0.02856748 | 0.00754542 | 0.00476974 | 0.01077739 | 0.01120509 | 0.01247181 | 0.01795992 | 0.00632565 | 0.0129069 | 0.00233859 | 0.00543276 |
| 21.6944444 | 0.04125784 | 0.07086128 | 0.01582368 | 0.0055977 | 0.00903641 | 0.00556126 | 0.00253074 | 0.01885732 | 0.0135052 | 0.00606442 | 0.00431245 | 0.00656563 | 0.01108184 | 0.01535694 | 0.01724646 | 0.00615774 | 0.01803342 | 0.00171971 | 0.00959436 |
| 21.7083333 | 0.02666306 | 0.05714914 | 0.00848044 | 0.0041488 | 0.00566164 | 0.00339008 | 0.00301818 | 0.00919291 | 0.01896072 | 0.00790429 | 0.00560964 | 0.00441957 | 0.01285216 | 0.01086844 | 0.0230205 | 0.00807515 | 0.01919503 | 0.00372286 | 0.01369522 |
| 21.7222222 | 0.01830711 | 0.03379896 | 0.00509754 | 0.00235422 | 0.00219846 | 0.00332743 | 0.00530291 | 0.01485769 | 0.03386485 | 0.00692561 | 0.00521881 | 0.00584609 | 0.01376219 | 0.00575033 | 0.03883019 | 0.00890526 | 0.01784062 | 0.00769451 | 0.01454162 |
| 21.7361111 | 0.03552887 | 0.03429376 | 0.007348 | 0.00121686 | 0.00176161 | 0.00390679 | 0.00519055 | 0.0222268 | 0.04278518 | 0.00576637 | 0.00310291 | 0.00614784 | 0.02062974 | 0.00826036 | 0.04081211 | 0.00652286 | 0.01391899 | 0.010044 | 0.01072042 |
| 21.75 | 0.05539046 | 0.05071031 | 0.00840247 | 0.00100092 | 0.00340611 | 0.00311386 | 0.00327168 | 0.01920371 | 0.03940298 | 0.00881333 | 0.00168292 | 0.00741849 | 0.0264959 | 0.01193647 | 0.02468671 | 0.00435688 | 0.00762705 | 0.00826213 | 0.00579403 |
| 21.7638889 | 0.05357001 | 0.0530759 | 0.00624073 | 0.0024496 | 0.00521046 | 0.00175657 | 0.00338742 | 0.01319149 | 0.02430172 | 0.01022202 | 0.00255223 | 0.01267792 | 0.02293734 | 0.00961422 | 0.01452826 | 0.00695494 | 0.0025583 | 0.00433909 | 0.00585231 |
| 21.7777778 | 0.03190205 | 0.0351876 | 0.00652981 | 0.00453908 | 0.00474015 | 0.0018368 | 0.00600392 | 0.01725858 | 0.01108849 | 0.00659497 | 0.00385245 | 0.01677519 | 0.01641774 | 0.00471987 | 0.01709188 | 0.01162236 | 0.00084495 | 0.00203811 | 0.00847791 |
| 21.7916667 | 0.01281535 | 0.01747835 | 0.00698085 | 0.0045265 | 0.00238996 | 0.00446041 | 0.00787209 | 0.02685448 | 0.01403766 | 0.0039369 | 0.00322615 | 0.01517064 | 0.01204912 | 0.00215304 | 0.01466428 | 0.01297816 | 0.00161935 | 0.00208429 | 0.00697989 |
| 21.8055556 | 0.00847459 | 0.01474572 | 0.00719355 | 0.00571394 | 0.0013846 | 0.00894386 | 0.00749444 | 0.0240594 | 0.03158806 | 0.00543595 | 0.00299857 | 0.01059265 | 0.01718266 | 0.00141814 | 0.01259651 | 0.01075148 | 0.00467291 | 0.00439457 | 0.00508568 |
| 21.8194444 | 0.01144228 | 0.01684285 | 0.00947093 | 0.01245691 | 0.00223945 | 0.01126757 | 0.00494705 | 0.01434946 | 0.05142757 | 0.00595263 | 0.00579517 | 0.01098612 | 0.03361672 | 0.00126922 | 0.02483153 | 0.00738452 | 0.00910652 | 0.00841798 | 0.00993326 |
| 21.8333333 | 0.02752694 | 0.02632116 | 0.01042859 | 0.01997127 | 0.00360444 | 0.00843764 | 0.00295117 | 0.0170539 | 0.06480387 | 0.00393114 | 0.00838733 | 0.01419237 | 0.0466655 | 0.00332711 | 0.03598837 | 0.00408773 | 0.01270893 | 0.01106574 | 0.01510883 |
| 21.8472222 | 0.05235317 | 0.04618933 | 0.00733383 | 0.02115271 | 0.00424532 | 0.00379731 | 0.00381678 | 0.02462973 | 0.06831863 | 0.00254242 | 0.00785369 | 0.01057469 | 0.04330445 | 0.00705989 | 0.0330211 | 0.00370302 | 0.01454265 | 0.01026722 | 0.01356821 |
| 21.8611111 | 0.06589366 | 0.05592481 | 0.00370226 | 0.01700701 | 0.00331029 | 0.00318563 | 0.00584974 | 0.0204839 | 0.05718287 | 0.00352562 | 0.0058614 | 0.00638022 | 0.0268087 | 0.01021798 | 0.02335717 | 0.00836108 | 0.01545712 | 0.00704851 | 0.00875224 |
| 21.875 | 0.05881612 | 0.04646672 | 0.00376962 | 0.01153077 | 0.00155113 | 0.00706828 | 0.00847372 | 0.01146162 | 0.03321121 | 0.00783441 | 0.00535694 | 0.01091872 | 0.01763857 | 0.01217086 | 0.01727785 | 0.01410616 | 0.01605955 | 0.00653351 | 0.00516634 |
| 21.8888889 | 0.04115019 | 0.02972975 | 0.00420564 | 0.00590194 | 0.00082482 | 0.01172502 | 0.01047819 | 0.00697779 | 0.01327141 | 0.0148025 | 0.00742533 | 0.01552992 | 0.02025732 | 0.01371552 | 0.01871961 | 0.01672846 | 0.01684846 | 0.0102468 | 0.00529889 |
| 21.9027778 | 0.03067373 | 0.01766658 | 0.00581478 | 0.00312283 | 0.00230404 | 0.01443257 | 0.01003888 | 0.00892381 | 0.00688606 | 0.02067995 | 0.01075543 | 0.01128374 | 0.01694565 | 0.01529146 | 0.025546 | 0.01601142 | 0.01823199 | 0.00998154 | 0.01007829 |
| 21.9166667 | 0.03371789 | 0.01386519 | 0.01196083 | 0.00657083 | 0.00676176 | 0.01441558 | 0.00858681 | 0.02178675 | 0.00750199 | 0.02297271 | 0.0137773 | 0.00714858 | 0.01504798 | 0.01634555 | 0.03638959 | 0.01321604 | 0.01921961 | 0.00555774 | 0.01724081 |
| 21.9305556 | 0.04629167 | 0.01675403 | 0.01774452 | 0.01332935 | 0.01383915 | 0.01292396 | 0.00735229 | 0.03619246 | 0.0158247 | 0.02700536 | 0.01524955 | 0.01201453 | 0.02896299 | 0.0164825 | 0.04391375 | 0.00955722 | 0.01910558 | 0.00448392 | 0.02115265 |
| 21.9444444 | 0.06183613 | 0.02601006 | 0.01991438 | 0.0171945 | 0.02034629 | 0.01187264 | 0.00614929 | 0.04095726 | 0.030371 | 0.03290755 | 0.01386911 | 0.01656764 | 0.04371558 | 0.01578145 | 0.04058836 | 0.00697679 | 0.01771293 | 0.00458894 | 0.01873141 |
| 21.9583333 | 0.07234871 | 0.03970739 | 0.01938643 | 0.01531769 | 0.022447 | 0.01134822 | 0.00504368 | 0.03665037 | 0.03862888 | 0.03060546 | 0.01019678 | 0.01332348 | 0.04285492 | 0.0145532 | 0.03330967 | 0.00600704 | 0.01435874 | 0.00249397 | 0.01342859 |
| 21.9722222 | 0.07043485 | 0.05044898 | 0.0166588 | 0.01108237 | 0.01962544 | 0.01016817 | 0.00458562 | 0.02533134 | 0.03386347 | 0.0202709 | 0.00640216 | 0.00724505 | 0.02766808 | 0.01321529 | 0.02608406 | 0.00420274 | 0.00983065 | 0.00106863 | 0.00900475 |
| 21.9861111 | 0.05893462 | 0.0547032 | 0.01176275 | 0.0088001 | 0.0155714 | 0.00802942 | 0.00501137 | 0.01210986 | 0.02061614 | 0.01015257 | 0.00327557 | 0.00312614 | 0.01251431 | 0.01251839 | 0.01598868 | 0.00205654 | 0.00626632 | 0.00153188 | 0.00676653 |
| 22 | 0.04642525 | 0.05319382 | 0.00625041 | 0.00782145 | 0.01292578 | 0.00537193 | 0.0051388 | 0.00440869 | 0.00943505 | 0.00545685 | 0.0016593 | 0.00191523 | 0.00770731 | 0.01313943 | 0.00613292 | 0.00200754 | 0.00439809 | 0.00235297 | 0.0077644 |
| 22.0138889 | 0.03740044 | 0.04493774 | 0.00323109 | 0.00680409 | 0.01081401 | 0.00308961 | 0.00371049 | 0.00734267 | 0.00641064 | 0.00828231 | 0.00242349 | 0.00292254 | 0.00674451 | 0.01415403 | 0.00268756 | 0.00262545 | 0.00355209 | 0.00342413 | 0.01171412 |
| 22.0277778 | 0.03747635 | 0.03647119 | 0.00351513 | 0.0069692 | 0.00939089 | 0.00371478 | 0.00181088 | 0.02184721 | 0.00689808 | 0.01636405 | 0.00418474 | 0.00568282 | 0.00813051 | 0.01423576 | 0.00911131 | 0.00239321 | 0.00408554 | 0.00435689 | 0.01687585 |
| 22.0416667 | 0.04687664 | 0.0417307 | 0.00877039 | 0.00961045 | 0.0102661 | 0.00898543 | 0.00257534 | 0.04493431 | 0.01280519 | 0.02678441 | 0.00521647 | 0.01086806 | 0.01784908 | 0.01431001 | 0.02566883 | 0.00423257 | 0.0075955 | 0.00358879 | 0.02212179 |
| 22.0555556 | 0.04963109 | 0.05815329 | 0.01959918 | 0.01344416 | 0.01268358 | 0.01565924 | 0.00695837 | 0.06976321 | 0.03484573 | 0.03800772 | 0.00464875 | 0.01901823 | 0.03295417 | 0.01497219 | 0.04690233 | 0.00953428 | 0.01223026 | 0.00171407 | 0.02626697 |
| 22.0694444 | 0.03422469 | 0.06826177 | 0.0285823 | 0.01484312 | 0.01278331 | 0.01796053 | 0.0112792 | 0.08229185 | 0.05326781 | 0.05263578 | 0.00296661 | 0.0262032 | 0.04385731 | 0.01323013 | 0.06034208 | 0.01261306 | 0.01182664 | 0.00075784 | 0.02654702 |
| 22.0833333 | 0.01588499 | 0.06404379 | 0.02891026 | 0.01273623 | 0.00877331 | 0.0140416 | 0.01206385 | 0.0685848 | 0.0406918 | 0.06427722 | 0.00284924 | 0.02576977 | 0.03905478 | 0.00820839 | 0.05236568 | 0.00978728 | 0.00667253 | 0.00143922 | 0.02271848 |
| 22.0972222 | 0.01106064 | 0.04737498 | 0.02228916 | 0.00963495 | 0.00376228 | 0.00788888 | 0.00946685 | 0.03839477 | 0.01588326 | 0.05610855 | 0.00437671 | 0.01817297 | 0.02239782 | 0.00384591 | 0.0276309 | 0.00490169 | 0.00384317 | 0.00266821 | 0.01806061 |
| 22.1111111 | 0.01095378 | 0.02692283 | 0.0144194 | 0.00659085 | 0.00106402 | 0.00398123 | 0.00526856 | 0.01472603 | 0.00421093 | 0.0317715 | 0.0041065 | 0.01010015 | 0.01418773 | 0.00210863 | 0.01659694 | 0.00183382 | 0.00362369 | 0.00287641 | 0.01361613 |
| 22.125 | 0.00981024 | 0.01224936 | 0.00834428 | 0.00330179 | 0.00054252 | 0.00215049 | 0.00286393 | 0.00647609 | 0.00151682 | 0.01170277 | 0.00206756 | 0.00461331 | 0.01754506 | 0.00161228 | 0.03088566 | 0.00129633 | 0.00235136 | 0.0019209 | 0.0086814 |
| 22.1388889 | 0.00963286 | 0.01005972 | 0.00418643 | 0.00167363 | 0.00128517 | 0.0020659 | 0.00391486 | 0.01318934 | 0.00463129 | 0.00732614 | 0.00078636 | 0.00170081 | 0.01979665 | 0.00103563 | 0.04552134 | 0.00243695 | 0.00081012 | 0.00154402 | 0.00479709 |
| 22.1527778 | 0.00652123 | 0.02033723 | 0.0018283 | 0.00292116 | 0.00306581 | 0.00413123 | 0.00548274 | 0.02871327 | 0.01549167 | 0.01477723 | 0.00061852 | 0.00149917 | 0.02010467 | 0.0006225 | 0.0468351 | 0.00354759 | 0.00073056 | 0.00401456 | 0.00481622 |
| 22.1666667 | 0.00516077 | 0.02856506 | 0.00241418 | 0.00468193 | 0.00429487 | 0.00501621 | 0.00583955 | 0.04207839 | 0.02716356 | 0.0245258 | 0.00077333 | 0.00294025 | 0.02117859 | 0.00077365 | 0.04128338 | 0.00389051 | 0.00173941 | 0.00902113 | 0.00736315 |
| 22.1805556 | 0.01047869 | 0.03063879 | 0.00720472 | 0.00473281 | 0.00434897 | 0.00376917 | 0.00506687 | 0.04909929 | 0.03461359 | 0.0306991 | 0.00102685 | 0.00405265 | 0.01831637 | 0.00067718 | 0.03085628 | 0.00321954 | 0.00379851 | 0.01370998 | 0.00815344 |
| 22.1944444 | 0.02067967 | 0.0358269 | 0.01293976 | 0.00359215 | 0.00417335 | 0.00310609 | 0.00313887 | 0.05079862 | 0.0382994 | 0.03250528 | 0.00114804 | 0.0035532 | 0.01004622 | 0.00024309 | 0.01790021 | 0.00167503 | 0.00739212 | 0.01660928 | 0.00775955 |
| 22.2083333 | 0.02969615 | 0.04209367 | 0.01485473 | 0.00297878 | 0.00349613 | 0.00327738 | 0.00155274 | 0.04545018 | 0.0371865 | 0.0313324 | 0.00116379 | 0.00206564 | 0.00415208 | 0.00044613 | 0.00825538 | 0.00083141 | 0.01040444 | 0.01806858 | 0.00921476 |
| 22.2222222 | 0.030755 | 0.04280187 | 0.01260404 | 0.00347193 | 0.00223529 | 0.0033472 | 0.00150027 | 0.03270527 | 0.03005785 | 0.02699176 | 0.00176504 | 0.00125488 | 0.00526155 | 0.00170383 | 0.00349475 | 0.00156312 | 0.01099531 | 0.01830417 | 0.01174133 |
| 22.2361111 | 0.02797161 | 0.04175437 | 0.01053682 | 0.00451941 | 0.00190612 | 0.00400284 | 0.00191077 | 0.01969001 | 0.02011527 | 0.02097114 | 0.0020955 | 0.00248593 | 0.00603793 | 0.003027 | 0.00161097 | 0.00280606 | 0.01070117 | 0.01681142 | 0.01156509 |
| 22.25 | 0.02665984 | 0.04223859 | 0.01178651 | 0.00560887 | 0.00311143 | 0.005901 | 0.00168566 | 0.01359411 | 0.01222538 | 0.01590144 | 0.00145485 | 0.00574293 | 0.00404279 | 0.00329951 | 0.00156711 | 0.00291227 | 0.01045204 | 0.01268375 | 0.0071573 |
| 22.2638889 | 0.02172414 | 0.03349832 | 0.01280795 | 0.00589482 | 0.00376288 | 0.00684035 | 0.00109879 | 0.01239279 | 0.0073894 | 0.01072571 | 0.00069132 | 0.00817017 | 0.00471721 | 0.0025304 | 0.00477864 | 0.00175828 | 0.00873178 | 0.00696167 | 0.00305879 |
| 22.2777778 | 0.01268534 | 0.02134095 | 0.00912096 | 0.00511049 | 0.00254631 | 0.00477687 | 0.00053817 | 0.00950062 | 0.0060181 | 0.00591589 | 0.00050883 | 0.00719296 | 0.0070634 | 0.00137314 | 0.01044509 | 0.00078499 | 0.00495984 | 0.00267498 | 0.00238132 |
| 22.2916667 | 0.00909377 | 0.0267825 | 0.00612653 | 0.00401657 | 0.00135825 | 0.00324653 | 0.00025573 | 0.00634083 | 0.00730626 | 0.00527131 | 0.00061992 | 0.00419836 | 0.00622668 | 0.00094645 | 0.01195477 | 0.00169261 | 0.00169879 | 0.00097108 | 0.00290626 |
| 22.3055556 | 0.00909452 | 0.03711369 | 0.00929578 | 0.00248866 | 0.00122983 | 0.00481266 | 0.00023025 | 0.0104072 | 0.00894051 | 0.00628845 | 0.00088861 | 0.00212237 | 0.00377364 | 0.00149586 | 0.00841341 | 0.00484453 | 0.0012517 | 0.00072729 | 0.00500845 |
| 22.3194444 | 0.00714391 | 0.02862661 | 0.01139765 | 0.00124816 | 0.00201546 | 0.00528765 | 0.00036428 | 0.01907952 | 0.01214859 | 0.00474343 | 0.00137418 | 0.00200168 | 0.00310972 | 0.00175839 | 0.00596522 | 0.00701153 | 0.00360954 | 0.00149306 | 0.00775075 |
| 22.3333333 | 0.00823024 | 0.01367537 | 0.00754021 | 0.00106801 | 0.0037192 | 0.00311106 | 0.00061494 | 0.02242189 | 0.01788409 | 0.00382618 | 0.00224747 | 0.00266208 | 0.0039459 | 0.00143981 | 0.00634754 | 0.00541476 | 0.00832774 | 0.00428055 | 0.00626787 |
| 22.3472222 | 0.01235308 | 0.01196404 | 0.00340186 | 0.00140295 | 0.00449319 | 0.00195986 | 0.00087715 | 0.01558996 | 0.02036411 | 0.00501043 | 0.0043272 | 0.00274166 | 0.00785794 | 0.00164367 | 0.01110229 | 0.00388288 | 0.01319575 | 0.0080079 | 0.00289697 |
| 22.3611111 | 0.01709965 | 0.01566838 | 0.00345113 | 0.00248914 | 0.00394416 | 0.00317717 | 0.00086779 | 0.00726883 | 0.01534205 | 0.0044235 | 0.00533618 | 0.00243612 | 0.01267183 | 0.00224541 | 0.01674273 | 0.00527209 | 0.01522847 | 0.00928687 | 0.00254718 |
| 22.375 | 0.02341855 | 0.0150646 | 0.00498602 | 0.00335879 | 0.00391653 | 0.00480763 | 0.0004119 | 0.00852987 | 0.01570085 | 0.00274164 | 0.00375767 | 0.00322962 | 0.01200611 | 0.0026915 | 0.01506814 | 0.00566198 | 0.01381639 | 0.00775436 | 0.00390457 |
| 22.3888889 | 0.02826479 | 0.01309923 | 0.00557984 | 0.00365623 | 0.00466495 | 0.00572805 | 0.00047649 | 0.01373685 | 0.02987536 | 0.00456601 | 0.00264754 | 0.00648704 | 0.00663217 | 0.00378708 | 0.0085385 | 0.0036196 | 0.01119239 | 0.00568705 | 0.00763014 |
| 22.4027778 | 0.02936812 | 0.01493253 | 0.00652473 | 0.00465407 | 0.00492365 | 0.00633778 | 0.00157654 | 0.01488656 | 0.03796403 | 0.01078972 | 0.00319218 | 0.01150725 | 0.00296107 | 0.00552644 | 0.00455171 | 0.00246789 | 0.00985933 | 0.00410997 | 0.01366396 |
| 22.4166667 | 0.02649226 | 0.02015263 | 0.00752158 | 0.0067618 | 0.00370707 | 0.00622779 | 0.00262645 | 0.0141735 | 0.02500319 | 0.01683189 | 0.00291515 | 0.01560916 | 0.00264003 | 0.00631209 | 0.0050148 | 0.00315051 | 0.00943529 | 0.00244513 | 0.01608676 |
| 22.4305556 | 0.01870516 | 0.02116094 | 0.00685294 | 0.0091685 | 0.00189717 | 0.00512522 | 0.00283522 | 0.0160526 | 0.01172472 | 0.0163934 | 0.00139449 | 0.01620615 | 0.00522004 | 0.00530895 | 0.00825055 | 0.00355856 | 0.00789023 | 0.00131861 | 0.01256803 |
| 22.4444444 | 0.01100802 | 0.01989581 | 0.00564974 | 0.01020113 | 0.00186107 | 0.00498578 | 0.00284675 | 0.01912152 | 0.01304178 | 0.00981398 | 0.00070028 | 0.013646 | 0.01147282 | 0.00353516 | 0.01038899 | 0.0024656 | 0.00454629 | 0.00204199 | 0.00695295 |
| 22.4583333 | 0.00956539 | 0.02471716 | 0.00675857 | 0.00959334 | 0.00401728 | 0.00693341 | 0.00380027 | 0.02033249 | 0.01365787 | 0.00490156 | 0.00103562 | 0.01132028 | 0.01479442 | 0.00235793 | 0.00794195 | 0.00128983 | 0.00137344 | 0.00268028 | 0.00268488 |
| 22.4722222 | 0.01143174 | 0.03029618 | 0.00943915 | 0.0084366 | 0.00613605 | 0.00855451 | 0.00505762 | 0.02194568 | 0.0173689 | 0.00605053 | 0.00149251 | 0.01048347 | 0.0100497 | 0.00218664 | 0.00501405 | 0.00159433 | 0.00139162 | 0.00175081 | 0.00179841 |
| 22.4861111 | 0.00924854 | 0.02430267 | 0.00957615 | 0.00781389 | 0.00764923 | 0.00763083 | 0.0048446 | 0.02313372 | 0.0323556 | 0.010863 | 0.00184152 | 0.00901406 | 0.00529414 | 0.00288384 | 0.0046418 | 0.00224813 | 0.00569139 | 0.00200007 | 0.00627138 |
| 22.5 | 0.00418151 | 0.01074852 | 0.00633907 | 0.00755587 | 0.00983017 | 0.00603051 | 0.00359413 | 0.01684056 | 0.03461314 | 0.01597878 | 0.00242844 | 0.00587675 | 0.00587011 | 0.00410845 | 0.00471214 | 0.00192402 | 0.01092731 | 0.00456652 | 0.01296987 |
| 22.5138889 | 0.00734587 | 0.01129102 | 0.00491546 | 0.00549813 | 0.00958163 | 0.00704246 | 0.00294297 | 0.00794318 | 0.01967753 | 0.01843953 | 0.00381551 | 0.00329981 | 0.0062763 | 0.00597219 | 0.00838615 | 0.00114488 | 0.01200014 | 0.00661623 | 0.01431403 |
| 22.5277778 | 0.02100138 | 0.03557615 | 0.00986547 | 0.00272878 | 0.00560232 | 0.01018169 | 0.00356407 | 0.00526126 | 0.01712852 | 0.01797859 | 0.00777908 | 0.00414373 | 0.00953729 | 0.01050514 | 0.0163024 | 0.00197711 | 0.00975485 | 0.00677153 | 0.01010852 |
| 22.5416667 | 0.03329162 | 0.07083392 | 0.02204157 | 0.00460377 | 0.00716076 | 0.01392648 | 0.00867684 | 0.01062447 | 0.02192616 | 0.02069145 | 0.01721084 | 0.01111821 | 0.02328329 | 0.02209848 | 0.02739667 | 0.00587775 | 0.01121614 | 0.00741374 | 0.00856651 |
| 22.5555556 | 0.04256441 | 0.09957738 | 0.03807645 | 0.01634821 | 0.01913126 | 0.02035828 | 0.02089388 | 0.02823292 | 0.03331649 | 0.03247525 | 0.02723035 | 0.02172626 | 0.04368349 | 0.04150975 | 0.04131273 | 0.00814988 | 0.01951868 | 0.0104495 | 0.01307969 |
| 22.5694444 | 0.04902102 | 0.09958914 | 0.04618738 | 0.03301728 | 0.02983202 | 0.02558867 | 0.03006022 | 0.04270598 | 0.0687389 | 0.04582051 | 0.02663083 | 0.02625829 | 0.05873831 | 0.05188458 | 0.04784371 | 0.00532441 | 0.0263005 | 0.01404493 | 0.01959914 |
| 22.5833333 | 0.04051009 | 0.06400382 | 0.03716742 | 0.03734346 | 0.02817454 | 0.02099049 | 0.02457262 | 0.03205848 | 0.08361798 | 0.0442003 | 0.01515332 | 0.0207795 | 0.05931031 | 0.03905727 | 0.04463161 | 0.00291168 | 0.02233462 | 0.01673365 | 0.02287257 |
| 22.5972222 | 0.02070208 | 0.02903789 | 0.01985983 | 0.02398282 | 0.01992162 | 0.01096817 | 0.01262292 | 0.01502626 | 0.05629791 | 0.02605533 | 0.0080531 | 0.01169567 | 0.04615667 | 0.01818575 | 0.04133317 | 0.00551807 | 0.01240764 | 0.01728043 | 0.01793964 |
| 22.6111111 | 0.0106474 | 0.02977956 | 0.01377874 | 0.01118595 | 0.01427084 | 0.00867361 | 0.01039722 | 0.02038186 | 0.0267763 | 0.01401657 | 0.01255101 | 0.01088072 | 0.02540287 | 0.00966931 | 0.03796078 | 0.01303171 | 0.0094966 | 0.01298699 | 0.01294721 |

| | | | | | | | | | | | | | | | | | | | |
|---|---|---|---|---|---|---|---|---|---|---|---|---|---|---|---|---|---|---|---|
| 22.625 | 0.01764333 | 0.0501246 | 0.02187414 | 0.01193586 | 0.0109212 | 0.0158682 | 0.01771273 | 0.03718518 | 0.02222739 | 0.02323931 | 0.01319787 | 0.02115603 | 0.01255304 | 0.01417638 | 0.03036727 | 0.02262528 | 0.01578346 | 0.00651749 | 0.01679012 |
| 22.6388889 | 0.02604903 | 0.05353607 | 0.02667228 | 0.0145208 | 0.00740174 | 0.01948252 | 0.02108416 | 0.04687572 | 0.04443382 | 0.03370858 | 0.01052768 | 0.02302908 | 0.01134608 | 0.01751295 | 0.02254875 | 0.02804921 | 0.01726917 | 0.00343004 | 0.01578206 |
| 22.6527778 | 0.02303737 | 0.03349426 | 0.01886982 | 0.01151819 | 0.00401435 | 0.0133868 | 0.01430544 | 0.03850132 | 0.060972 | 0.0278984 | 0.01721987 | 0.0170078 | 0.00928805 | 0.01274281 | 0.02384705 | 0.02232193 | 0.01230225 | 0.00438568 | 0.00939691 |
| 22.6666667 | 0.01484729 | 0.02182173 | 0.01046699 | 0.0128251 | 0.0028696 | 0.00885229 | 0.00820929 | 0.02984698 | 0.04463276 | 0.02279563 | 0.01959916 | 0.02420898 | 0.00847547 | 0.00623914 | 0.03072683 | 0.01262324 | 0.01637437 | 0.00662774 | 0.01247584 |
| 22.6805556 | 0.01461118 | 0.0361117 | 0.01517531 | 0.01708575 | 0.00469808 | 0.0097355 | 0.01089033 | 0.05146402 | 0.03165551 | 0.02518225 | 0.01120827 | 0.02921486 | 0.01191662 | 0.00360498 | 0.025621 | 0.01378119 | 0.02360429 | 0.00586107 | 0.01894766 |
| 22.6944444 | 0.01946746 | 0.04552131 | 0.02284476 | 0.01339414 | 0.00652902 | 0.00759609 | 0.01164885 | 0.06250798 | 0.03870795 | 0.01991071 | 0.00606399 | 0.01780935 | 0.01991433 | 0.00353025 | 0.01933875 | 0.02280605 | 0.0180772 | 0.00407724 | 0.01626934 |
| 22.7083333 | 0.01532127 | 0.02997633 | 0.01844981 | 0.0080054 | 0.00628954 | 0.00618913 | 0.00686688 | 0.04238652 | 0.03172232 | 0.00972437 | 0.00799485 | 0.00838339 | 0.03259295 | 0.00260412 | 0.03465583 | 0.03074652 | 0.0083879 | 0.0062033 | 0.01132922 |
| 22.7222222 | 0.00778821 | 0.01716309 | 0.01120752 | 0.01270773 | 0.00524769 | 0.01268779 | 0.00642132 | 0.04056701 | 0.02523946 | 0.0106281 | 0.00979105 | 0.01129256 | 0.04039853 | 0.00228213 | 0.05892347 | 0.03585304 | 0.00880303 | 0.00756162 | 0.01686903 |
| 22.7361111 | 0.00850117 | 0.03070028 | 0.01261596 | 0.02197446 | 0.00527241 | 0.01941107 | 0.01227884 | 0.05977971 | 0.0355688 | 0.02499274 | 0.00712285 | 0.02000034 | 0.03768343 | 0.00462861 | 0.07766824 | 0.03651831 | 0.01497078 | 0.00540914 | 0.02517596 |
| 22.75 | 0.01128395 | 0.05066853 | 0.01596233 | 0.02245388 | 0.00674978 | 0.01705685 | 0.017623 | 0.06067123 | 0.0334541 | 0.02917809 | 0.00509972 | 0.02246416 | 0.02431831 | 0.00639125 | 0.08096066 | 0.03058989 | 0.01353264 | 0.00617049 | 0.02050668 |
| 22.7638889 | 0.01008097 | 0.05105133 | 0.01257406 | 0.0130784 | 0.00871421 | 0.00913785 | 0.01756856 | 0.03740344 | 0.02031032 | 0.02107845 | 0.0076398 | 0.01768648 | 0.01274527 | 0.00497825 | 0.05523639 | 0.01943539 | 0.01033904 | 0.01382967 | 0.01271413 |
| 22.7777778 | 0.00721356 | 0.03176486 | 0.00584813 | 0.00699238 | 0.00964498 | 0.00462617 | 0.01240964 | 0.0244776 | 0.02207646 | 0.02356284 | 0.00854657 | 0.02059277 | 0.01911178 | 0.00253305 | 0.02485067 | 0.00921982 | 0.0164746 | 0.02255732 | 0.01620387 |
| 22.7916667 | 0.00556978 | 0.01831915 | 0.00298667 | 0.00955992 | 0.00797594 | 0.00376563 | 0.00612348 | 0.03437261 | 0.03155146 | 0.02203998 | 0.00612148 | 0.02663538 | 0.03143333 | 0.00202688 | 0.01532351 | 0.0050022 | 0.0172026 | 0.02399742 | 0.0153589 |
| 22.8055556 | 0.00597293 | 0.01952813 | 0.00517889 | 0.01115132 | 0.00425238 | 0.00460725 | 0.00331293 | 0.03179243 | 0.02779861 | 0.01319614 | 0.00731451 | 0.0203504 | 0.03005121 | 0.00394144 | 0.01040894 | 0.00556795 | 0.01308657 | 0.01752395 | 0.00904822 |
| 22.8194444 | 0.00698988 | 0.01833604 | 0.01406655 | 0.00791132 | 0.00170164 | 0.00909493 | 0.0029213 | 0.02314266 | 0.02896116 | 0.02353461 | 0.0126647 | 0.0105766 | 0.01789774 | 0.00657475 | 0.00588882 | 0.00645531 | 0.02081956 | 0.00870963 | 0.01511273 |
| 22.8333333 | 0.01008412 | 0.02573403 | 0.02596826 | 0.00723528 | 0.00136751 | 0.01376209 | 0.0020924 | 0.04528824 | 0.05664961 | 0.04604763 | 0.01486505 | 0.01200603 | 0.01397995 | 0.00937646 | 0.01622589 | 0.00603636 | 0.03046044 | 0.0044042 | 0.02776674 |
| 22.8472222 | 0.01742997 | 0.05241272 | 0.03215376 | 0.01313968 | 0.00135086 | 0.01449726 | 0.00231907 | 0.06869024 | 0.08249031 | 0.0567912 | 0.01078186 | 0.0203074 | 0.02548551 | 0.01308732 | 0.04041887 | 0.00576796 | 0.02984356 | 0.00667558 | 0.02935853 |
| 22.8611111 | 0.02484958 | 0.07743865 | 0.03346377 | 0.01892467 | 0.00158663 | 0.01260263 | 0.00464825 | 0.05732884 | 0.08057979 | 0.04663488 | 0.00481436 | 0.02116386 | 0.03706441 | 0.01667076 | 0.05960728 | 0.00472737 | 0.02254476 | 0.01040143 | 0.01812061 |
| 22.875 | 0.02610315 | 0.08191708 | 0.03217804 | 0.01910072 | 0.00281856 | 0.01048213 | 0.00732 | 0.03165389 | 0.06341833 | 0.02332528 | 0.00264628 | 0.01289432 | 0.03773414 | 0.01720131 | 0.05576038 | 0.003055 | 0.01507632 | 0.01166009 | 0.01119297 |
| 22.8888889 | 0.0201839 | 0.06801764 | 0.02712106 | 0.01588671 | 0.00503025 | 0.01016079 | 0.00897428 | 0.01985106 | 0.04864349 | 0.0061129 | 0.00509642 | 0.00646286 | 0.03151365 | 0.01447706 | 0.03613688 | 0.00403203 | 0.01088114 | 0.01047336 | 0.01190116 |
| 22.9027778 | 0.01205855 | 0.04863493 | 0.02069166 | 0.01398376 | 0.00852758 | 0.01206122 | 0.00891251 | 0.02511784 | 0.03859111 | 0.00390975 | 0.00905308 | 0.00600121 | 0.02402321 | 0.01258853 | 0.01742815 | 0.00603241 | 0.01047685 | 0.00874674 | 0.00937865 |
| 22.9166667 | 0.00746753 | 0.02898486 | 0.01846895 | 0.01712954 | 0.012375 | 0.01471395 | 0.00790338 | 0.03823323 | 0.0296481 | 0.01215696 | 0.01260547 | 0.00612728 | 0.01647185 | 0.01394779 | 0.01166491 | 0.00556696 | 0.01220005 | 0.00821752 | 0.010096 |
| 22.9305556 | 0.00697654 | 0.01385595 | 0.02170099 | 0.02415092 | 0.01507644 | 0.01780603 | 0.00802976 | 0.05094821 | 0.0231724 | 0.02385939 | 0.01458429 | 0.00921232 | 0.01122045 | 0.01616697 | 0.02265095 | 0.00335373 | 0.01239961 | 0.00713656 | 0.02320406 |
| 22.9444444 | 0.00868458 | 0.00976734 | 0.02617338 | 0.02931261 | 0.01644308 | 0.02076714 | 0.00888289 | 0.0540309 | 0.01788808 | 0.03388684 | 0.01398603 | 0.01709286 | 0.01044513 | 0.01631244 | 0.03433211 | 0.00151674 | 0.00889914 | 0.00412789 | 0.03960403 |
| 22.9583333 | 0.01060954 | 0.01011997 | 0.0264485 | 0.0275529 | 0.01634396 | 0.02001667 | 0.00806431 | 0.040816 | 0.01131659 | 0.04141482 | 0.0115772 | 0.0202839 | 0.01150959 | 0.01436585 | 0.02985525 | 0.00174458 | 0.00417455 | 0.00185415 | 0.04539067 |
| 22.9722222 | 0.0100896 | 0.00912148 | 0.0201995 | 0.01977698 | 0.01389291 | 0.01452527 | 0.00500005 | 0.0209042 | 0.00968106 | 0.04078475 | 0.00863726 | 0.01383424 | 0.01199568 | 0.01162947 | 0.01594942 | 0.00504354 | 0.00208379 | 0.00187824 | 0.03756169 |
| 22.9861111 | 0.00650964 | 0.00871411 | 0.01121443 | 0.01043872 | 0.00894653 | 0.00779867 | 0.00209522 | 0.01275349 | 0.0127887 | 0.02536456 | 0.00610158 | 0.00647025 | 0.01019684 | 0.00855149 | 0.01288509 | 0.01024356 | 0.00194674 | 0.00174511 | 0.02257163 |
| 23 | 0.00454055 | 0.0067353 | 0.00437718 | 0.00471968 | 0.00390997 | 0.0035165 | 0.00119348 | 0.01502005 | 0.01131272 | 0.01248011 | 0.00453313 | 0.00579367 | 0.00726526 | 0.00509313 | 0.02720745 | 0.01680644 | 0.00133247 | 0.00133197 | 0.01078084 |
| 23.0138889 | 0.00546501 | 0.00810853 | 0.00182902 | 0.0055047 | 0.00117647 | 0.0040394 | 0.00102052 | 0.012511 | 0.01087225 | 0.01697132 | 0.00361077 | 0.00544565 | 0.01006899 | 0.00228156 | 0.0438156 | 0.02309934 | 0.00135419 | 0.00230049 | 0.00869935 |
| 23.0277778 | 0.00732646 | 0.02407459 | 0.00285492 | 0.00700882 | 0.00042474 | 0.00925508 | 0.00111383 | 0.01248243 | 0.01899866 | 0.01781379 | 0.00364579 | 0.00530757 | 0.01776655 | 0.00238846 | 0.04744598 | 0.02405085 | 0.00430196 | 0.00355963 | 0.00824908 |
| 23.0416667 | 0.01415907 | 0.05043876 | 0.01004363 | 0.00633019 | 0.0008878 | 0.01646412 | 0.00487116 | 0.02824581 | 0.0304897 | 0.01569353 | 0.00665692 | 0.01434385 | 0.01676674 | 0.00703112 | 0.04058867 | 0.02015796 | 0.00954214 | 0.0046663 | 0.00833088 |
| 23.0555556 | 0.0222625 | 0.07296118 | 0.0252384 | 0.00986219 | 0.00224198 | 0.02255222 | 0.01449346 | 0.04836649 | 0.04038714 | 0.0413138 | 0.01300619 | 0.02707599 | 0.01564569 | 0.01422894 | 0.03237101 | 0.01799044 | 0.01371016 | 0.00701643 | 0.01852588 |
| 23.0694444 | 0.02201089 | 0.0789366 | 0.03738411 | 0.01656551 | 0.00398803 | 0.02480624 | 0.02540131 | 0.05510133 | 0.03971555 | 0.08665911 | 0.01849074 | 0.03026174 | 0.03284444 | 0.01753371 | 0.02476385 | 0.01951095 | 0.01225234 | 0.009753 | 0.02926833 |
| 23.0833333 | 0.01622898 | 0.06620623 | 0.0360792 | 0.01987013 | 0.00513647 | 0.01940524 | 0.02911271 | 0.04284324 | 0.02404106 | 0.10982684 | 0.01808705 | 0.02369707 | 0.04958605 | 0.01351761 | 0.01621545 | 0.02043303 | 0.00706862 | 0.009865 | 0.02835202 |
| 23.0972222 | 0.01595937 | 0.0498562 | 0.02562213 | 0.02025822 | 0.00565744 | 0.00901644 | 0.02286082 | 0.02467442 | 0.01006042 | 0.09923223 | 0.01249776 | 0.01584133 | 0.04658499 | 0.00683483 | 0.00826453 | 0.01759006 | 0.00521559 | 0.0066598 | 0.02377319 |
| 23.1111111 | 0.02230059 | 0.04110091 | 0.01561313 | 0.01923492 | 0.00646157 | 0.0027746 | 0.01310393 | 0.01742374 | 0.00710466 | 0.08001632 | 0.00822497 | 0.00932565 | 0.02977273 | 0.00287311 | 0.00505683 | 0.01239729 | 0.00570932 | 0.0033657 | 0.02387529 |
| 23.125 | 0.02355759 | 0.03506313 | 0.00905419 | 0.01469204 | 0.00655185 | 0.00146994 | 0.00613847 | 0.01825292 | 0.00473204 | 0.05864244 | 0.00843611 | 0.00395941 | 0.01298964 | 0.00210917 | 0.00559916 | 0.00725629 | 0.00442378 | 0.00341292 | 0.02350423 |
| 23.1388889 | 0.0148231 | 0.02504861 | 0.00429032 | 0.00741778 | 0.00477273 | 0.00271206 | 0.00274532 | 0.01607471 | 0.00130068 | 0.03150707 | 0.00835266 | 0.00116126 | 0.00925703 | 0.00216348 | 0.0055205 | 0.00399508 | 0.00244506 | 0.00562656 | 0.01779824 |
| 23.1527778 | 0.00533823 | 0.01248653 | 0.00191352 | 0.00322633 | 0.00290845 | 0.00740112 | 0.00136961 | 0.00972942 | 0.00123701 | 0.0137663 | 0.00600717 | 0.00066659 | 0.01721632 | 0.00141688 | 0.00630698 | 0.00553017 | 0.00139129 | 0.00590198 | 0.00995488 |
| 23.1666667 | 0.00334597 | 0.00529103 | 0.0026969 | 0.00452239 | 0.00412724 | 0.01391891 | 0.00142283 | 0.00489665 | 0.00417049 | 0.01598768 | 0.00708991 | 0.0012387 | 0.02636443 | 0.0008654 | 0.01168846 | 0.01179921 | 0.00189699 | 0.00399775 | 0.00808074 |
| 23.1805556 | 0.00703717 | 0.00675199 | 0.00445516 | 0.00707049 | 0.00743852 | 0.01781775 | 0.00276412 | 0.00612296 | 0.00910599 | 0.02912388 | 0.01198855 | 0.0013943 | 0.03155469 | 0.00154603 | 0.02279809 | 0.01968139 | 0.0046177 | 0.00389275 | 0.01445514 |
| 23.1944444 | 0.01027759 | 0.00803587 | 0.00491073 | 0.00767892 | 0.00849265 | 0.0154362 | 0.00482039 | 0.01288228 | 0.01518675 | 0.04147149 | 0.01475551 | 0.00092507 | 0.03259222 | 0.00232106 | 0.03721231 | 0.02707011 | 0.00871739 | 0.00634129 | 0.02120043 |
| 23.2083333 | 0.0095694 | 0.00408867 | 0.00457509 | 0.00728869 | 0.00711523 | 0.00842047 | 0.00707167 | 0.02287596 | 0.02304737 | 0.05155868 | 0.01526903 | 0.00098745 | 0.02946787 | 0.00256889 | 0.04590367 | 0.03081849 | 0.01257501 | 0.00898902 | 0.02546345 |
| 23.2222222 | 0.0084009 | 0.00926626 | 0.00642436 | 0.00856717 | 0.00641891 | 0.00261252 | 0.00819906 | 0.03562381 | 0.03160902 | 0.05759963 | 0.01584347 | 0.00129572 | 0.02409948 | 0.00333011 | 0.04230664 | 0.03076804 | 0.01546927 | 0.01182687 | 0.02885676 |
| 23.2361111 | 0.01033714 | 0.03068819 | 0.01127817 | 0.01161673 | 0.00610926 | 0.00143127 | 0.0075219 | 0.04575785 | 0.03731272 | 0.05358115 | 0.01426923 | 0.00165553 | 0.01906117 | 0.0048764 | 0.03139602 | 0.03006539 | 0.01695083 | 0.01431583 | 0.02815788 |
| 23.25 | 0.01272531 | 0.04925792 | 0.01585272 | 0.0135313 | 0.0043147 | 0.003395 | 0.00641994 | 0.04803306 | 0.03894239 | 0.04068987 | 0.0098742 | 0.00288209 | 0.01486196 | 0.00641141 | 0.02027597 | 0.02666373 | 0.01701812 | 0.01455823 | 0.02305755 |
| 23.2638889 | 0.0134364 | 0.05409852 | 0.01737096 | 0.01314251 | 0.00199831 | 0.00593423 | 0.00668581 | 0.04571504 | 0.03730851 | 0.02638433 | 0.00517945 | 0.00440644 | 0.01044461 | 0.00711662 | 0.01088413 | 0.01765124 | 0.01632608 | 0.01259645 | 0.01732216 |
| 23.2777778 | 0.01254401 | 0.05050132 | 0.01582004 | 0.01122811 | 0.00140697 | 0.00777054 | 0.00793488 | 0.04106376 | 0.03264796 | 0.01562398 | 0.00855745 | 0.00594858 | 0.00720235 | 0.00615894 | 0.00646753 | 0.01066884 | 0.01177545 | 0.01003232 | 0.01306982 |
| 23.2916667 | 0.01100744 | 0.04169333 | 0.01319484 | 0.00903364 | 0.0025049 | 0.00798197 | 0.00822341 | 0.03179619 | 0.02686973 | 0.01870055 | 0.03133656 | 0.01425192 | 0.01022231 | 0.00851222 | 0.01152297 | 0.0174953 | 0.01074698 | 0.01493785 | 0.02087945 |
| 23.3055556 | 0.00921594 | 0.03143392 | 0.01108387 | 0.00749136 | 0.00250686 | 0.00609752 | 0.00711255 | 0.01871702 | 0.02139796 | 0.03258041 | 0.05915909 | 0.02795668 | 0.01645069 | 0.01618612 | 0.02000726 | 0.0316344 | 0.02216956 | 0.02776455 | 0.04043929 |
| 23.3194444 | 0.00566011 | 0.01920788 | 0.00798692 | 0.00556708 | 0.00184606 | 0.00323031 | 0.00581215 | 0.01072456 | 0.0171541 | 0.03169724 | 0.05910525 | 0.02744972 | 0.01465065 | 0.01761932 | 0.0202774 | 0.03383806 | 0.02792227 | 0.02969653 | 0.04376446 |
| 23.3333333 | 0.0019555 | 0.01006748 | 0.00440194 | 0.00271999 | 0.00281634 | 0.00159416 | 0.00464258 | 0.01775147 | 0.02107273 | 0.01763731 | 0.03277983 | 0.01361409 | 0.00877774 | 0.01050091 | 0.01195978 | 0.02125877 | 0.01912916 | 0.01678994 | 0.02578498 |
| 23.3472222 | 0.00127299 | 0.01313612 | 0.00375188 | 0.00075373 | 0.00342574 | 0.00150699 | 0.00283684 | 0.03373262 | 0.03021822 | 0.01277875 | 0.01166603 | 0.00600582 | 0.01008088 | 0.0057296 | 0.00451491 | 0.00769716 | 0.01005492 | 0.00598914 | 0.01096888 |
| 23.3611111 | 0.00311787 | 0.01825985 | 0.00466589 | 0.0007655 | 0.00231927 | 0.00155642 | 0.00100354 | 0.04240278 | 0.02869602 | 0.01401623 | 0.00632713 | 0.00495543 | 0.01116924 | 0.00588081 | 0.00412757 | 0.00192658 | 0.005062 | 0.00354912 | 0.00670344 |
| 23.375 | 0.00549712 | 0.01335589 | 0.0041014 | 0.0022247 | 0.00146952 | 0.00241134 | 0.00041371 | 0.03749091 | 0.01675692 | 0.01146666 | 0.00516898 | 0.00481971 | 0.00725114 | 0.0059365 | 0.0086044 | 0.00256695 | 0.00365466 | 0.00299075 | 0.00450488 |
| 23.3888889 | 0.00765468 | 0.00932986 | 0.00249745 | 0.00468193 | 0.00140374 | 0.00455452 | 0.00090751 | 0.02230096 | 0.01163356 | 0.01394128 | 0.0040062 | 0.006005 | 0.00580819 | 0.00377245 | 0.01219098 | 0.00626889 | 0.00906527 | 0.00352679 | 0.00288738 |
| 23.4027778 | 0.01059449 | 0.02022657 | 0.00235347 | 0.00771668 | 0.0022664 | 0.00650271 | 0.00189218 | 0.01361039 | 0.02027513 | 0.02546146 | 0.00692203 | 0.00761017 | 0.00564969 | 0.00264508 | 0.01152134 | 0.0101564 | 0.01792904 | 0.00747776 | 0.00488302 |
| 23.4166667 | 0.0143743 | 0.03545313 | 0.00502855 | 0.01047443 | 0.00483639 | 0.00759978 | 0.00326134 | 0.02725811 | 0.03153407 | 0.03358738 | 0.01135653 | 0.00861892 | 0.00453143 | 0.00545746 | 0.00757237 | 0.01147473 | 0.02192533 | 0.00996704 | 0.00802267 |
| 23.4305556 | 0.01647155 | 0.03774985 | 0.00835503 | 0.01182741 | 0.0066281 | 0.0079922 | 0.00497097 | 0.04827804 | 0.0344251 | 0.03217724 | 0.01243922 | 0.00862664 | 0.00757602 | 0.00961465 | 0.00428833 | 0.00936215 | 0.01814803 | 0.00911799 | 0.01022198 |
| 23.4444444 | 0.01488868 | 0.02433906 | 0.00992556 | 0.01103006 | 0.00563261 | 0.00767378 | 0.00619676 | 0.05681527 | 0.02803971 | 0.02441424 | 0.01036568 | 0.00831582 | 0.01464952 | 0.01199575 | 0.00541005 | 0.00620358 | 0.01091328 | 0.0071882 | 0.01355229 |
| 23.4583333 | 0.01179759 | 0.00969137 | 0.01025417 | 0.00944004 | 0.00357141 | 0.00690757 | 0.00651673 | 0.0519197 | 0.01917421 | 0.01474505 | 0.00787719 | 0.00916652 | 0.0190737 | 0.01262633 | 0.01140695 | 0.00568982 | 0.00493259 | 0.00495009 | 0.01976765 |
| 23.4722222 | 0.01229087 | 0.0046568 | 0.01109404 | 0.00943671 | 0.00213291 | 0.00681308 | 0.0071568 | 0.03954477 | 0.01429123 | 0.01097878 | 0.00842282 | 0.01168338 | 0.01854833 | 0.01291367 | 0.01953747 | 0.00823884 | 0.00183548 | 0.00410286 | 0.02467375 |
| 23.4861111 | 0.01730342 | 0.00497134 | 0.01193656 | 0.01079243 | 0.00127443 | 0.00797533 | 0.00860228 | 0.02468542 | 0.01138499 | 0.01611372 | 0.01111329 | 0.01460995 | 0.01454618 | 0.01322136 | 0.0242853 | 0.01036828 | 0.00105999 | 0.00490327 | 0.02009653 |
| 23.5 | 0.02094144 | 0.0100726 | 0.01068902 | 0.01098868 | 0.00071719 | 0.00882196 | 0.00930791 | 0.01367148 | 0.00790758 | 0.01968866 | 0.0106046 | 0.01510411 | 0.00957492 | 0.01207693 | 0.01912047 | 0.00871447 | 0.00083181 | 0.00534992 | 0.01190779 |
| 23.5138889 | 0.01930193 | 0.02088191 | 0.00716675 | 0.01038057 | 0.00160581 | 0.00947966 | 0.00773472 | 0.01260761 | 0.00771103 | 0.01420659 | 0.00667151 | 0.01176129 | 0.00924461 | 0.00960649 | 0.01056057 | 0.00458991 | 0.00062917 | 0.00450144 | 0.01263362 |
| 23.5277778 | 0.01709289 | 0.04362816 | 0.00533996 | 0.01223231 | 0.00565791 | 0.01329185 | 0.00425976 | 0.01384557 | 0.01167028 | 0.0109279 | 0.00580712 | 0.00801238 | 0.01079847 | 0.00833701 | 0.00910459 | 0.00183077 | 0.00160367 | 0.00374541 | 0.01378082 |
| 23.5416667 | 0.02245732 | 0.08458871 | 0.01371034 | 0.01614737 | 0.0107741 | 0.01959132 | 0.00161556 | 0.0231365 | 0.01653646 | 0.02428207 | 0.01164847 | 0.01018897 | 0.01181071 | 0.01123247 | 0.01086974 | 0.00170863 | 0.00794799 | 0.00669506 | 0.01405555 |
| 23.5555556 | 0.03680646 | 0.12788155 | 0.03452525 | 0.02195879 | 0.01175349 | 0.02405646 | 0.0051435 | 0.05767256 | 0.03318963 | 0.05528426 | 0.02081436 | 0.02156367 | 0.02500413 | 0.02036856 | 0.02054942 | 0.00292203 | 0.02385763 | 0.01553975 | 0.02758646 |
| 23.5694444 | 0.05401264 | 0.14052056 | 0.05370714 | 0.03077171 | 0.01099077 | 0.02434859 | 0.01266437 | 0.09092917 | 0.06081104 | 0.0857947 | 0.02895752 | 0.04141124 | 0.047664 | 0.03248645 | 0.04203894 | 0.00278289 | 0.04228574 | 0.02297943 | 0.04356407 |
| 23.5833333 | 0.06192115 | 0.11313672 | 0.0558367 | 0.03497183 | 0.01560526 | 0.01873601 | 0.01416304 | 0.08852039 | 0.07148147 | 0.08783612 | 0.03274851 | 0.0528182 | 0.05777151 | 0.03651014 | 0.05248204 | 0.00180777 | 0.04825085 | 0.02184732 | 0.04198828 |
| 23.5972222 | 0.04861725 | 0.06777711 | 0.04177628 | 0.02847645 | 0.02357332 | 0.01020062 | 0.00857867 | 0.05737387 | 0.05895039 | 0.0550811 | 0.02633314 | 0.03773317 | 0.05057255 | 0.02492394 | 0.03942805 | 0.00358143 | 0.04022316 | 0.01649424 | 0.02699479 |
| 23.6111111 | 0.02589492 | 0.03507456 | 0.02145013 | 0.01552323 | 0.027965 | 0.00918825 | 0.0050886 | 0.0309605 | 0.03403443 | 0.03128026 | 0.01570783 | 0.01759092 | 0.03536556 | 0.01171468 | 0.02161628 | 0.00956824 | 0.02951839 | 0.01021752 | 0.02373553 |
| 23.625 | 0.02043538 | 0.03820428 | 0.00942095 | 0.00822637 | 0.02735596 | 0.01819049 | 0.00943693 | 0.03884434 | 0.01810542 | 0.05165883 | 0.01566226 | 0.01893244 | 0.01758558 | 0.01101246 | 0.01694452 | 0.01708059 | 0.02098723 | 0.00508 | 0.03905452 |
| 23.6388889 | 0.03201827 | 0.0598199 | 0.01277957 | 0.01420473 | 0.02482131 | 0.02509445 | 0.01505088 | 0.0700476 | 0.03043657 | 0.08100462 | 0.01975059 | 0.03192292 | 0.00534232 | 0.01102736 | 0.02233014 | 0.02163707 | 0.01261023 | 0.00459987 | 0.05124 |
| 23.6527778 | 0.03352625 | 0.06730447 | 0.01884159 | 0.02111577 | 0.02272115 | 0.01925933 | 0.01372204 | 0.0969543 | 0.05205758 | 0.07428267 | 0.01507437 | 0.0361702 | 0.00356961 | 0.00645313 | 0.02292589 | 0.02094354 | 0.005357 | 0.00451509 | 0.04172317 |
| 23.6666667 | 0.02376835 | 0.04754681 | 0.01699554 | 0.01658347 | 0.02002678 | 0.01194695 | 0.0074383 | 0.09762085 | 0.05362371 | 0.04269902 | 0.01103456 | 0.02454462 | 0.00443087 | 0.00622838 | 0.02378894 | 0.01680438 | 0.00242307 | 0.00293792 | 0.02712407 |
| 23.6805556 | 0.02805588 | 0.03076853 | 0.01032155 | 0.01072641 | 0.01420775 | 0.01456483 | 0.00455653 | 0.06701498 | 0.03311458 | 0.02826396 | 0.01592665 | 0.01526991 | 0.00267597 | 0.00906637 | 0.03596922 | 0.01586047 | 0.00265864 | 0.00311232 | 0.0353384 |
| 23.6944444 | 0.0418699 | 0.05411345 | 0.0101737 | 0.01782742 | 0.00725379 | 0.01444523 | 0.00720671 | 0.05309572 | 0.02178763 | 0.02841875 | 0.01863967 | 0.02875987 | 0.00615808 | 0.01078453 | 0.04067132 | 0.01833039 | 0.00436161 | 0.00296722 | 0.04562297 |
| 23.7083333 | 0.0422316 | 0.08059459 | 0.01994504 | 0.02487993 | 0.00589969 | 0.00771475 | 0.0097586 | 0.07193792 | 0.03098298 | 0.02116089 | 0.01234251 | 0.03942471 | 0.02472163 | 0.00863801 | 0.02558485 | 0.01620563 | 0.00722702 | 0.00274081 | 0.03186353 |
| 23.7222222 | 0.02787288 | 0.07354853 | 0.02995127 | 0.0168746 | 0.01191268 | 0.00554733 | 0.00920984 | 0.06842044 | 0.03010329 | 0.02142432 | 0.00715096 | 0.02468521 | 0.0507101 | 0.00590943 | 0.01309422 | 0.01021978 | 0.00911406 | 0.00495384 | 0.01650001 |
| 23.7361111 | 0.01418547 | 0.04467108 | 0.03263588 | 0.00894876 | 0.01913285 | 0.01094482 | 0.00599018 | 0.04249318 | 0.02317943 | 0.03956172 | 0.01156309 | 0.01137275 | 0.06280911 | 0.01108788 | 0.02184643 | 0.00995276 | 0.00850673 | 0.0070251 | 0.02382914 |
| 23.75 | 0.01236309 | 0.01982986 | 0.02463585 | 0.01301694 | 0.02109176 | 0.01494637 | 0.00265806 | 0.04502395 | 0.03709188 | 0.05559966 | 0.0152542 | 0.01362305 | 0.05618546 | 0.02027477 | 0.03821368 | 0.01559433 | 0.00607252 | 0.00798278 | 0.04123703 |
| 23.7638889 | 0.01366178 | 0.01965553 | 0.01433855 | 0.01878015 | 0.01595376 | 0.01016781 | 0.00222153 | 0.07632396 | 0.05853869 | 0.05494039 | 0.01095778 | 0.01748152 | 0.03874446 | 0.0249794 | 0.05008042 | 0.02098117 | 0.00325572 | 0.00984779 | 0.04930419 |
| 23.7777778 | 0.00989192 | 0.03296378 | 0.01534789 | 0.01637587 | 0.00789574 | 0.00380842 | 0.00370903 | 0.10430785 | 0.07052847 | 0.0370982 | 0.00784054 | 0.0256649 | 0.01741492 | 0.0223583 | 0.05248538 | 0.02170896 | 0.00117391 | 0.01025587 | 0.0367078 |
| 23.7916667 | 0.01021452 | 0.0457029 | 0.02222825 | 0.01064525 | 0.00402659 | 0.00243121 | 0.00477003 | 0.10352054 | 0.06200224 | 0.01530439 | 0.0069275 | 0.03856014 | 0.00638617 | 0.01297208 | 0.03647325 | 0.01356698 | 0.00104595 | 0.00695289 | 0.02308742 |
| 23.8055556 | 0.01422268 | 0.05048404 | 0.02101684 | 0.01242171 | 0.004939 | 0.00449886 | 0.00653439 | 0.06657317 | 0.03411411 | 0.00823595 | 0.00513295 | 0.03466822 | 0.01086909 | 0.00525787 | 0.01585628 | 0.00564549 | 0.00336439 | 0.00615044 | 0.02863241 |
| 23.8194444 | 0.01162621 | 0.03576144 | 0.01233527 | 0.01606097 | 0.00519992 | 0.01304897 | 0.0097174 | 0.04117181 | 0.01863925 | 0.00823192 | 0.0079663 | 0.0199472 | 0.02063361 | 0.00501648 | 0.00716826 | 0.00451213 | 0.00659166 | 0.00993224 | 0.02409786 |
| 23.8333333 | 0.00792844 | 0.02057052 | 0.0088769 | 0.01226234 | 0.00358668 | 0.02654671 | 0.01033017 | 0.06049095 | 0.02368801 | 0.01481878 | 0.0149888 | 0.02270306 | 0.02825587 | 0.00551743 | 0.00594423 | 0.004771 | 0.00624056 | 0.01102158 | 0.01359828 |
| 23.8472222 | 0.01344114 | 0.02575222 | 0.01475209 | 0.01008349 | 0.00442406 | 0.03297285 | 0.00638043 | 0.07301 | 0.02208532 | 0.03292637 | 0.01970318 | 0.03416774 | 0.02783822 | 0.00418321 | 0.00650298 | 0.00624558 | 0.00634164 | 0.00709835 | 0.02854267 |
| 23.8611111 | 0.02036043 | 0.0261968 | 0.01705618 | 0.01601681 | 0.00966489 | 0.02745376 | 0.00234244 | 0.04867654 | 0.01585532 | 0.04699689 | 0.01854719 | 0.02953225 | 0.01832895 | 0.00656529 | 0.00743083 | 0.00686076 | 0.01569507 | 0.0043943 | 0.05173316 |
| 23.875 | 0.01831215 | 0.01647487 | 0.01466821 | 0.02314866 | 0.01492432 | 0.01648553 | 0.00130167 | 0.03445301 | 0.02759824 | 0.05546497 | 0.01219073 | 0.0159442 | 0.01535895 | 0.01252708 | 0.00697392 | 0.00499987 | 0.02752349 | 0.00835855 | 0.05683363 |
| 23.8888889 | 0.01106913 | 0.02358962 | 0.02316093 | 0.02665073 | 0.01641132 | 0.00891325 | 0.0013032 | 0.06247209 | 0.04860013 | 0.05648859 | 0.00542259 | 0.01379725 | 0.02784571 | 0.0178607 | 0.00687576 | 0.00526603 | 0.03098758 | 0.01449373 | 0.0458004 |
| 23.9027778 | 0.00522827 | 0.04546706 | 0.0387276 | 0.02602742 | 0.01571944 | 0.008661 | 0.00233642 | 0.09170075 | 0.05556502 | 0.04476162 | 0.00262666 | 0.02478376 | 0.0409147 | 0.02151818 | 0.00906313 | 0.00982983 | 0.02492728 | 0.01429257 | 0.03385769 |
| 23.9166667 | 0.0027091 | 0.06554402 | 0.04723593 | 0.02125157 | 0.0151048 | 0.01240985 | 0.00599985 | 0.08591939 | 0.04245459 | 0.03530018 | 0.00418566 | 0.03498294 | 0.04321587 | 0.02335185 | 0.01265618 | 0.01628752 | 0.01539022 | 0.00802481 | 0.02846996 |
| 23.9305556 | 0.00309588 | 0.07544639 | 0.04774575 | 0.01392112 | 0.01431337 | 0.01334034 | 0.00970829 | 0.05900837 | 0.02345939 | 0.03463428 | 0.00769267 | 0.03483861 | 0.03808166 | 0.02147726 | 0.01604967 | 0.02098299 | 0.00876245 | 0.00349144 | 0.02083436 |
| 23.9444444 | 0.00377241 | 0.06818944 | 0.04233726 | 0.0065041 | 0.01239302 | 0.00833371 | 0.01022577 | 0.0296299 | 0.01071222 | 0.03333707 | 0.01088503 | 0.02539638 | 0.03126549 | 0.01622698 | 0.01618428 | 0.01999966 | 0.00538517 | 0.00375742 | 0.01040414 |
| 23.9583333 | 0.00360054 | 0.04594666 | 0.03395806 | 0.00195372 | 0.00985586 | 0.00255381 | 0.00911672 | 0.01099232 | 0.00490981 | 0.02688759 | 0.01259644 | 0.01307372 | 0.02552992 | 0.01150496 | 0.01230693 | 0.01216552 | 0.00363459 | 0.00522796 | 0.0083786 |
| 23.9722222 | 0.00288202 | 0.02235284 | 0.02339619 | 0.00053989 | 0.00793073 | 0.00208749 | 0.00756614 | 0.00868318 | 0.00567228 | 0.01893622 | 0.01349298 | 0.00794808 | 0.01925097 | 0.00917988 | 0.00824056 | 0.00491019 | 0.00249245 | 0.00481602 | 0.00948549 |
| 23.9861111 | 0.0016274 | 0.01330416 | 0.01284589 | 0.00192313 | 0.00640815 | 0.00686901 | 0.00471413 | 0.01620126 | 0.0103909 | 0.01529927 | 0.01512507 | 0.01108821 | 0.01212877 | 0.00717317 | 0.00649891 | 0.00327304 | 0.00141904 | 0.00303538 | 0.00867099 |
| 24 | 0.00213424 | 0.02098175 | 0.0128806 | 0.00690282 | 0.0050408 | 0.01341125 | 0.00409486 | 0.02515627 | 0.01273304 | 0.01525803 | 0.01653504 | 0.01387536 | 0.0114191 | 0.00415145 | 0.00597229 | 0.00485159 | 0.0009359 | 0.00258888 | 0.01193329 |
| 24.0138889 | 0.00471966 | 0.02559398 | 0.02326653 | 0.01215252 | 0.00591494 | 0.01968861 | 0.00843886 | 0.03141281 | 0.01103832 | 0.01280225 | 0.0148835 | 0.01352896 | 0.01878762 | 0.00220285 | 0.00485773 | 0.00830154 | 0.00157218 | 0.00354843 | 0.01591022 |
| 24.0277778 | 0.01000215 | 0.02326725 | 0.03004282 | 0.01272277 | 0.01011765 | 0.02843694 | 0.01202429 | 0.02831257 | 0.00766747 | 0.00846419 | 0.01176729 | 0.01065274 | 0.02158661 | 0.00209413 | 0.00487318 | 0.01092124 | 0.00183597 | 0.00360713 | 0.01722299 |
| 24.0416667 | 0.02265189 | 0.0350494 | 0.02358048 | 0.01333264 | 0.01299841 | 0.03982784 | 0.00971302 | 0.02174273 | 0.00915892 | 0.01508221 | 0.01363184 | 0.00841253 | 0.01624946 | 0.00210686 | 0.01158795 | 0.01377748 | 0.00231889 | 0.00272211 | 0.02063293 |
| 24.0555556 | 0.03805215 | 0.05755763 | 0.01860528 | 0.02322531 | 0.01172484 | 0.04742508 | 0.00845553 | 0.02745518 | 0.02116731 | 0.03950115 | 0.02202754 | 0.01587314 | 0.0174312 | 0.00346148 | 0.02292783 | 0.02051071 | 0.00554953 | 0.00255837 | 0.03016762 |
| 24.0694444 | 0.0423738 | 0.06701848 | 0.02961432 | 0.03663874 | 0.01009991 | 0.05037299 | 0.01254528 | 0.03292103 | 0.03359811 | 0.06673927 | 0.03095596 | 0.02957511 | 0.02553486 | 0.00667899 | 0.02617693 | 0.02701916 | 0.00820401 | 0.00302203 | 0.04167389 |
| 24.0833333 | 0.03213874 | 0.0614969 | 0.03948985 | 0.03954808 | 0.00972728 | 0.05013216 | 0.01394192 | 0.02211616 | 0.0331853 | 0.0830175 | 0.03563208 | 0.0323569 | 0.02318035 | 0.00834222 | 0.01809057 | 0.02734995 | 0.00740714 | 0.00342363 | 0.0441467 |
| 24.0972222 | 0.01735137 | 0.05094318 | 0.03735909 | 0.03048477 | 0.00873783 | 0.04188524 | 0.01081867 | 0.00778485 | 0.02233618 | 0.08065306 | 0.03339017 | 0.02197559 | 0.01194557 | 0.00714136 | 0.01008064 | 0.02126704 | 0.00534136 | 0.0028961 | 0.03866341 |
| 24.1111111 | 0.00742897 | 0.03772225 | 0.02877495 | 0.01799283 | 0.0062212 | 0.0256576 | 0.00682351 | 0.00253801 | 0.01131143 | 0.05380801 | 0.02255333 | 0.01011967 | 0.00478805 | 0.00441236 | 0.00791698 | 0.01260511 | 0.00342712 | 0.00219625 | 0.0324138 |
| 24.125 | 0.00452061 | 0.02431843 | 0.01699477 | 0.00844759 | 0.00328277 | 0.01078059 | 0.00382165 | 0.00221276 | 0.00469551 | 0.02076183 | 0.00976146 | 0.0055318 | 0.00201398 | 0.00182702 | 0.00858418 | 0.0065021 | 0.00211499 | 0.00462032 | 0.02332421 |

| | | | | | | | | | | | | | | | | | | | |
|---|---|---|---|---|---|---|---|---|---|---|---|---|---|---|---|---|---|---|---|
| 24.1388889 | 0.01008616 | 0.012468 | 0.00758899 | 0.00632208 | 0.00150799 | 0.00641821 | 0.00374627 | 0.00437174 | 0.0020886 | 0.00835504 | 0.00985706 | 0.0067574 | 0.00497204 | 0.0017973 | 0.01837928 | 0.01085628 | 0.0068642 | 0.01209018 | 0.01821626 |
| 24.1527778 | 0.02593548 | 0.01228969 | 0.00530371 | 0.01355825 | 0.00358341 | 0.01396055 | 0.0045408 | 0.01033561 | 0.00358012 | 0.03117587 | 0.03660524 | 0.02127541 | 0.02877895 | 0.0085211 | 0.06046865 | 0.03843228 | 0.03111582 | 0.03140443 | 0.03478256 |
| 24.1666667 | 0.0515144 | 0.0390635 | 0.00600204 | 0.02774386 | 0.01594135 | 0.03136619 | 0.00630195 | 0.01325509 | 0.00634391 | 0.07079074 | 0.07139083 | 0.05784244 | 0.07356197 | 0.02461994 | 0.11103722 | 0.07303437 | 0.06570387 | 0.05536079 | 0.06544225 |
| 24.1805556 | 0.07354177 | 0.07272864 | 0.00551517 | 0.04276041 | 0.03515208 | 0.0499261 | 0.01061322 | 0.01106379 | 0.00939569 | 0.07022545 | 0.0667862 | 0.07241516 | 0.09315856 | 0.03526637 | 0.10496484 | 0.0712084 | 0.06999939 | 0.05569207 | 0.06620164 |
| 24.1944444 | 0.06923641 | 0.07319333 | 0.0077493 | 0.04838483 | 0.04204241 | 0.05885767 | 0.01333041 | 0.0133005 | 0.02173462 | 0.03714933 | 0.03305929 | 0.03995352 | 0.0631521 | 0.0249155 | 0.06487629 | 0.04364748 | 0.04289003 | 0.03432829 | 0.03781462 |
| 24.2083333 | 0.04272353 | 0.04332141 | 0.01220696 | 0.04061225 | 0.03087319 | 0.05670121 | 0.01267263 | 0.01851558 | 0.03866755 | 0.02575117 | 0.01632173 | 0.00839881 | 0.03234741 | 0.01108536 | 0.05148138 | 0.03128211 | 0.02410598 | 0.01918155 | 0.02545818 |
| 24.2222222 | 0.02047695 | 0.01845336 | 0.01341076 | 0.02561893 | 0.01581705 | 0.04442153 | 0.01084513 | 0.01629339 | 0.03964135 | 0.02163666 | 0.01284126 | 0.00568532 | 0.02188436 | 0.007612 | 0.04813475 | 0.02783401 | 0.01837976 | 0.01587667 | 0.02080791 |
| 24.2361111 | 0.01047121 | 0.00879824 | 0.00954917 | 0.01185622 | 0.0060519 | 0.02843286 | 0.00710051 | 0.00820697 | 0.02407251 | 0.01271713 | 0.01167636 | 0.01501775 | 0.01368762 | 0.00532419 | 0.04398129 | 0.02642276 | 0.01492599 | 0.01890486 | 0.01109178 |
| 24.25 | 0.00527976 | 0.00408551 | 0.00479181 | 0.00644885 | 0.00242193 | 0.01731837 | 0.00329529 | 0.00223915 | 0.00915042 | 0.01375291 | 0.01373723 | 0.0182219 | 0.01243545 | 0.00352802 | 0.05101493 | 0.03129656 | 0.01501746 | 0.02357582 | 0.01067193 |
| 24.2638889 | 0.00442799 | 0.00169876 | 0.00313793 | 0.00689512 | 0.00341722 | 0.01199268 | 0.00217248 | 0.00115681 | 0.00272202 | 0.01379692 | 0.01172633 | 0.01493271 | 0.01212404 | 0.00293311 | 0.04089847 | 0.02568499 | 0.01275009 | 0.02105346 | 0.01249885 |
| 24.2777778 | 0.01041861 | 0.00241341 | 0.00315277 | 0.00631315 | 0.00532292 | 0.00885423 | 0.00242953 | 0.00237352 | 0.0029497 | 0.00814692 | 0.0067296 | 0.01289139 | 0.01075771 | 0.00217213 | 0.01623787 | 0.01237998 | 0.0086226 | 0.0142399 | 0.01045435 |
| 24.2916667 | 0.01781488 | 0.00474507 | 0.00460122 | 0.00678186 | 0.00505625 | 0.00586838 | 0.00379634 | 0.0024554 | 0.00617743 | 0.00456258 | 0.00433979 | 0.01400129 | 0.01531137 | 0.00305266 | 0.00479263 | 0.00741048 | 0.00818223 | 0.00925742 | 0.00985347 |
| 24.3055556 | 0.01767196 | 0.00511438 | 0.00510959 | 0.00780027 | 0.00279405 | 0.0060586 | 0.00382398 | 0.00128925 | 0.00800992 | 0.00497195 | 0.00524787 | 0.01610958 | 0.01719721 | 0.00475753 | 0.0043311 | 0.00916282 | 0.0100697 | 0.00660525 | 0.0115958 |
| 24.3194444 | 0.01105446 | 0.00316858 | 0.00396899 | 0.00597244 | 0.00082895 | 0.01127675 | 0.00242584 | 0.00057458 | 0.00711927 | 0.00608726 | 0.00654785 | 0.01613183 | 0.01479243 | 0.00521942 | 0.00889608 | 0.01076952 | 0.01101804 | 0.0103299 | 0.01306562 |
| 24.3333333 | 0.00802492 | 0.00196366 | 0.00566034 | 0.00592416 | 0.00088356 | 0.01621199 | 0.00370014 | 0.00072065 | 0.00571475 | 0.00479558 | 0.00590713 | 0.01219787 | 0.01211558 | 0.00369811 | 0.01305527 | 0.00978569 | 0.00952742 | 0.01713843 | 0.01280997 |
| 24.3472222 | 0.00962651 | 0.00135838 | 0.00772066 | 0.0089198 | 0.00311105 | 0.01700193 | 0.00682222 | 0.00131073 | 0.00545304 | 0.00338811 | 0.00486218 | 0.00589074 | 0.00806703 | 0.00242148 | 0.01429507 | 0.00840209 | 0.00773635 | 0.01739702 | 0.01245109 |
| 24.3611111 | 0.00797625 | 0.00076772 | 0.00562371 | 0.00900772 | 0.00524537 | 0.0155078 | 0.00839408 | 0.00160836 | 0.0059138 | 0.00528664 | 0.00539823 | 0.00156267 | 0.00378069 | 0.00332051 | 0.01547436 | 0.00830612 | 0.00783695 | 0.01136752 | 0.01248351 |
| 24.375 | 0.00580713 | 0.00120372 | 0.00369329 | 0.00653624 | 0.00478825 | 0.01152779 | 0.00604735 | 0.00189725 | 0.00641791 | 0.00806376 | 0.00613468 | 0.00082942 | 0.00249236 | 0.00465041 | 0.01627737 | 0.00831006 | 0.00825481 | 0.00590763 | 0.01122796 |
| 24.3888889 | 0.0119617 | 0.00307102 | 0.00641614 | 0.00925642 | 0.00254181 | 0.00740755 | 0.00345082 | 0.00269594 | 0.00656473 | 0.00747939 | 0.00498338 | 0.00311859 | 0.00232211 | 0.0036975 | 0.01406853 | 0.00665608 | 0.005968 | 0.00503259 | 0.00824148 |
| 24.4027778 | 0.02163014 | 0.00534494 | 0.00856558 | 0.01722847 | 0.00118386 | 0.00910288 | 0.00517744 | 0.00313082 | 0.00554899 | 0.00482772 | 0.00261911 | 0.00657361 | 0.00222109 | 0.00153417 | 0.00896096 | 0.00373127 | 0.00242518 | 0.00781824 | 0.00432986 |
| 24.4166667 | 0.02218748 | 0.00546756 | 0.00578457 | 0.02075575 | 0.0017077 | 0.01363943 | 0.00731854 | 0.00383043 | 0.00321878 | 0.00308433 | 0.00126063 | 0.00825205 | 0.00466803 | 0.00049529 | 0.00501563 | 0.00204499 | 0.0014601 | 0.01145827 | 0.00239218 |
| 24.4305556 | 0.01599869 | 0.00409474 | 0.00297821 | 0.01856882 | 0.00225301 | 0.01611117 | 0.00789019 | 0.0090594 | 0.00378138 | 0.00361744 | 0.00278877 | 0.01086141 | 0.01046198 | 0.00148408 | 0.00812879 | 0.00426956 | 0.00397461 | 0.01572641 | 0.00559896 |
| 24.4444444 | 0.01928638 | 0.00491832 | 0.00709693 | 0.01778567 | 0.00156147 | 0.01599104 | 0.00957378 | 0.02064966 | 0.01333411 | 0.008145 | 0.00709711 | 0.01761711 | 0.01806022 | 0.00473066 | 0.01833455 | 0.0096996 | 0.009014 | 0.02149585 | 0.01165955 |
| 24.4583333 | 0.0317616 | 0.00964531 | 0.01652878 | 0.01791226 | 0.00080303 | 0.0114173 | 0.01229288 | 0.02931989 | 0.02642842 | 0.01347756 | 0.0108292 | 0.02253679 | 0.0217508 | 0.00768565 | 0.02582758 | 0.01356001 | 0.01386835 | 0.02479283 | 0.01470749 |
| 24.4722222 | 0.03832208 | 0.01490204 | 0.02134055 | 0.01714895 | 0.00063848 | 0.00609823 | 0.01362334 | 0.02527293 | 0.02878055 | 0.01415921 | 0.01141359 | 0.02016054 | 0.01819224 | 0.00792528 | 0.02430054 | 0.01329583 | 0.01490646 | 0.02146708 | 0.01289553 |
| 24.4861111 | 0.03176325 | 0.01335175 | 0.01637637 | 0.01697793 | 0.00091323 | 0.00765303 | 0.01140982 | 0.01348079 | 0.01838639 | 0.01169831 | 0.01014287 | 0.01476107 | 0.01213943 | 0.00659681 | 0.01834458 | 0.01144308 | 0.01216293 | 0.01535921 | 0.00965181 |
| 24.5 | 0.01593695 | 0.00649702 | 0.00763486 | 0.01360675 | 0.00161122 | 0.01330546 | 0.00667174 | 0.00439963 | 0.00731189 | 0.00969206 | 0.00886071 | 0.01147464 | 0.00810539 | 0.00565455 | 0.01366143 | 0.00999973 | 0.00925897 | 0.01031006 | 0.00744478 |
| 24.5138889 | 0.00889524 | 0.00925397 | 0.00324777 | 0.00811928 | 0.00243069 | 0.01303837 | 0.00330201 | 0.00141293 | 0.0046736 | 0.00813864 | 0.00809256 | 0.0097727 | 0.00614994 | 0.00519551 | 0.01121793 | 0.00868857 | 0.00778708 | 0.00641362 | 0.00604665 |
| 24.5277778 | 0.02580052 | 0.03102991 | 0.00260429 | 0.00720858 | 0.00222565 | 0.01416549 | 0.00326004 | 0.00634905 | 0.01313117 | 0.00751704 | 0.00951267 | 0.00911535 | 0.00686008 | 0.00537571 | 0.01348568 | 0.00969563 | 0.00901096 | 0.00633107 | 0.00677729 |
| 24.5416667 | 0.05491852 | 0.05875817 | 0.00847753 | 0.00806368 | 0.00141424 | 0.02408946 | 0.00689785 | 0.02774254 | 0.03151764 | 0.01095208 | 0.01522196 | 0.01315275 | 0.0142101 | 0.00883189 | 0.03136213 | 0.01914547 | 0.01735497 | 0.01538714 | 0.01308481 |
| 24.5555556 | 0.07245769 | 0.07295972 | 0.03151182 | 0.01492072 | 0.00365027 | 0.02919078 | 0.01622301 | 0.07326152 | 0.05351868 | 0.01818596 | 0.0223472 | 0.02289898 | 0.02885187 | 0.01519216 | 0.07146721 | 0.04110606 | 0.03106047 | 0.03363385 | 0.02424715 |
| 24.5694444 | 0.07031867 | 0.07030522 | 0.0582804 | 0.03361827 | 0.00876572 | 0.02212474 | 0.02539346 | 0.12873659 | 0.07519538 | 0.02266551 | 0.02405558 | 0.03214103 | 0.0415132 | 0.01868146 | 0.103542 | 0.06115122 | 0.03773624 | 0.04428134 | 0.03099221 |
| 24.5833333 | 0.05701364 | 0.05913928 | 0.06138831 | 0.04545372 | 0.01222273 | 0.01027109 | 0.02426069 | 0.1552936 | 0.08629209 | 0.02044894 | 0.01862825 | 0.03413087 | 0.04058393 | 0.01665999 | 0.09093368 | 0.0576599 | 0.03052433 | 0.03150821 | 0.0266465 |
| 24.5972222 | 0.03891775 | 0.04683884 | 0.04465645 | 0.03623613 | 0.01305276 | 0.00572356 | 0.01655804 | 0.12860046 | 0.07251759 | 0.01498445 | 0.01094611 | 0.02771891 | 0.02787962 | 0.01216671 | 0.05317213 | 0.03785557 | 0.01783404 | 0.01083695 | 0.01654 |
| 24.6111111 | 0.02419665 | 0.0332163 | 0.02660933 | 0.01902321 | 0.01287941 | 0.01124527 | 0.00957353 | 0.07058642 | 0.04162883 | 0.01029302 | 0.00506755 | 0.01799838 | 0.01581092 | 0.00804774 | 0.02659454 | 0.02207869 | 0.00923694 | 0.00591302 | 0.00797894 |
| 24.625 | 0.02949673 | 0.01777771 | 0.01272492 | 0.01449881 | 0.01191675 | 0.02071653 | 0.0054541 | 0.03040811 | 0.01972815 | 0.0070606 | 0.00194181 | 0.01057301 | 0.00885373 | 0.00498097 | 0.01463903 | 0.01391868 | 0.00565335 | 0.01545768 | 0.00310197 |
| 24.6388889 | 0.04653979 | 0.00870202 | 0.00764012 | 0.02486242 | 0.01078615 | 0.02733015 | 0.00632166 | 0.03650424 | 0.02272576 | 0.00482759 | 0.00148186 | 0.00669741 | 0.00475628 | 0.00299233 | 0.00902963 | 0.00964851 | 0.0048923 | 0.02417298 | 0.00177887 |
| 24.6527778 | 0.04672917 | 0.0110201 | 0.01097795 | 0.03512304 | 0.01195933 | 0.02225734 | 0.0086373 | 0.07650575 | 0.0420245 | 0.00350994 | 0.00302979 | 0.00520528 | 0.00254552 | 0.00214286 | 0.00641419 | 0.00793282 | 0.00545912 | 0.02441919 | 0.00285554 |
| 24.6666667 | 0.02720895 | 0.01757323 | 0.0112538 | 0.03261404 | 0.01729905 | 0.01292801 | 0.0067501 | 0.12101319 | 0.05908821 | 0.00338862 | 0.00554968 | 0.00460099 | 0.00171012 | 0.00217392 | 0.00536737 | 0.00784333 | 0.0062283 | 0.02016603 | 0.0044877 |
| 24.6805556 | 0.01893775 | 0.02108019 | 0.00761909 | 0.02058376 | 0.02468358 | 0.01314758 | 0.00264199 | 0.13988558 | 0.05852461 | 0.00443908 | 0.00763096 | 0.00386182 | 0.00180304 | 0.00237122 | 0.00487154 | 0.00809609 | 0.00662038 | 0.01584878 | 0.00535001 |
| 24.6944444 | 0.03740421 | 0.02244684 | 0.00951578 | 0.01429269 | 0.02705064 | 0.0142641 | 0.00224537 | 0.11084834 | 0.03708845 | 0.00571665 | 0.007543 | 0.00279273 | 0.00209128 | 0.00216591 | 0.00452441 | 0.00784744 | 0.00619624 | 0.01175733 | 0.00496574 |
| 24.7083333 | 0.05098751 | 0.02410498 | 0.01492599 | 0.01160125 | 0.02015374 | 0.00947357 | 0.00415172 | 0.06247999 | 0.01694487 | 0.006202 | 0.00500021 | 0.00160952 | 0.00201329 | 0.00150774 | 0.00425505 | 0.00706133 | 0.00470271 | 0.00814545 | 0.00353884 |
| 24.7222222 | 0.03798677 | 0.0237523 | 0.01751887 | 0.00739539 | 0.00944867 | 0.00777626 | 0.00404365 | 0.05910027 | 0.01928336 | 0.00585897 | 0.00216627 | 0.00101322 | 0.00169656 | 0.0007839 | 0.00388358 | 0.00625891 | 0.00283866 | 0.00629171 | 0.0022329 |
| 24.7361111 | 0.01948871 | 0.01755905 | 0.02097654 | 0.00978391 | 0.00380657 | 0.01223152 | 0.00397433 | 0.09638159 | 0.03524257 | 0.00527461 | 0.00157466 | 0.00375521 | 0.00148214 | 0.00047303 | 0.002965 | 0.00580495 | 0.00151638 | 0.00663801 | 0.00225002 |
| 24.75 | 0.02180705 | 0.00892113 | 0.02365973 | 0.01543242 | 0.00439291 | 0.02000708 | 0.00554397 | 0.1189931 | 0.04899244 | 0.00481697 | 0.003577 | 0.01566772 | 0.0013842 | 0.00059983 | 0.00151313 | 0.00556816 | 0.00099356 | 0.00761171 | 0.00341916 |
| 24.7638889 | 0.04364799 | 0.00449361 | 0.01890196 | 0.0184843 | 0.00572791 | 0.02584272 | 0.0066983 | 0.0994355 | 0.04753415 | 0.00447359 | 0.00597011 | 0.0268507 | 0.0013539 | 0.00092104 | 0.00049179 | 0.00518453 | 0.00114072 | 0.00730035 | 0.00451133 |
| 24.7777778 | 0.05707507 | 0.0068856 | 0.01074245 | 0.02050083 | 0.00447108 | 0.01994684 | 0.00569428 | 0.05788497 | 0.02909839 | 0.00430193 | 0.00588466 | 0.0212472 | 0.00147191 | 0.00148504 | 0.00034065 | 0.0045481 | 0.00177207 | 0.00532937 | 0.00474475 |
| 24.7916667 | 0.04668736 | 0.01755444 | 0.00506947 | 0.01955201 | 0.00215792 | 0.01213421 | 0.00605012 | 0.05440208 | 0.02277009 | 0.00459636 | 0.00352207 | 0.01089915 | 0.00182512 | 0.00228739 | 0.00084134 | 0.00396995 | 0.00261194 | 0.00267217 | 0.00437787 |
| 24.8055556 | 0.02497435 | 0.03822818 | 0.00522078 | 0.01376501 | 0.00161921 | 0.01627817 | 0.00963076 | 0.09784038 | 0.04101793 | 0.00526623 | 0.00196061 | 0.00801211 | 0.0022466 | 0.00260941 | 0.00183753 | 0.00345185 | 0.00308571 | 0.00133535 | 0.00383316 |
| 24.8194444 | 0.00784735 | 0.05578553 | 0.01093609 | 0.00757488 | 0.00159356 | 0.01567758 | 0.00995699 | 0.12897174 | 0.05365473 | 0.00548359 | 0.00145679 | 0.00680137 | 0.00222512 | 0.00180101 | 0.00231682 | 0.00245105 | 0.0025461 | 0.00240416 | 0.00310771 |
| 24.8333333 | 0.00200283 | 0.04383442 | 0.01740217 | 0.00446355 | 0.00157833 | 0.01129201 | 0.0095709 | 0.11537773 | 0.04453824 | 0.00468067 | 0.0015083 | 0.00468403 | 0.00156075 | 0.00112783 | 0.0017209 | 0.001283 | 0.00154133 | 0.00420645 | 0.00229586 |
| 24.8472222 | 0.00259496 | 0.02474483 | 0.0203612 | 0.00429992 | 0.00362742 | 0.02203659 | 0.01559537 | 0.07453397 | 0.02844255 | 0.00338168 | 0.00387191 | 0.00300599 | 0.00071101 | 0.00184583 | 0.00080156 | 0.00091839 | 0.00149557 | 0.00460687 | 0.00190258 |
| 24.8611111 | 0.00637921 | 0.03710568 | 0.01628085 | 0.00486125 | 0.00675652 | 0.03849654 | 0.02266207 | 0.03565683 | 0.01755828 | 0.00232617 | 0.00666092 | 0.00205154 | 0.00024745 | 0.00277399 | 0.000383 | 0.00093075 | 0.0023184 | 0.0030097 | 0.00223355 |
| 24.875 | 0.01847794 | 0.05571442 | 0.00944886 | 0.0083577 | 0.00920931 | 0.04808239 | 0.02351889 | 0.01256634 | 0.01009058 | 0.00175055 | 0.00560612 | 0.00134562 | 0.00027211 | 0.00288366 | 0.00069858 | 0.00078639 | 0.00289376 | 0.0013138 | 0.00288602 |
| 24.8888889 | 0.04370473 | 0.05466818 | 0.00820268 | 0.02456455 | 0.01162847 | 0.05105924 | 0.01767287 | 0.00341752 | 0.00447332 | 0.00189549 | 0.00292934 | 0.00065363 | 0.00046944 | 0.00218543 | 0.00132535 | 0.00126547 | 0.00292673 | 0.00134602 | 0.0031875 |
| 24.9027778 | 0.07788944 | 0.04664493 | 0.00932063 | 0.04954484 | 0.01820872 | 0.04920564 | 0.00927955 | 0.00162382 | 0.00286688 | 0.00307802 | 0.00280023 | 0.00022917 | 0.00046516 | 0.00133391 | 0.00165161 | 0.00237259 | 0.00248154 | 0.00216016 | 0.00300754 |
| 24.9166667 | 0.10019456 | 0.04108979 | 0.00855544 | 0.06557609 | 0.03038302 | 0.04107764 | 0.00338032 | 0.00198215 | 0.00236436 | 0.00497171 | 0.00293691 | 0.00042628 | 0.00060283 | 0.00119812 | 0.00120275 | 0.00295574 | 0.00163316 | 0.00287831 | 0.00271602 |
| 24.9305556 | 0.09652661 | 0.03566442 | 0.01136017 | 0.06309494 | 0.04185576 | 0.02612324 | 0.00215545 | 0.00267932 | 0.00200473 | 0.00650361 | 0.0033051 | 0.0009997 | 0.00131557 | 0.00117347 | 0.00069393 | 0.00252102 | 0.00100745 | 0.00354589 | 0.00241995 |
| 24.9444444 | 0.07529788 | 0.03006498 | 0.01735321 | 0.0485371 | 0.04665481 | 0.01228548 | 0.00415979 | 0.00265171 | 0.00393542 | 0.00648451 | 0.00624463 | 0.00110223 | 0.00162769 | 0.00143757 | 0.00101557 | 0.00165297 | 0.00134007 | 0.00387259 | 0.00201431 |
| 24.9583333 | 0.04819968 | 0.02523454 | 0.0193167 | 0.03068021 | 0.04101585 | 0.01083593 | 0.00548084 | 0.00333293 | 0.00598927 | 0.00453793 | 0.00803113 | 0.00077367 | 0.0013363 | 0.00299247 | 0.00115682 | 0.00113101 | 0.00261327 | 0.00342757 | 0.00181205 |
| 24.9722222 | 0.02380736 | 0.01978167 | 0.01707517 | 0.01711299 | 0.02514433 | 0.0215203 | 0.00384899 | 0.00574084 | 0.00575529 | 0.00203339 | 0.00612388 | 0.00087022 | 0.00201668 | 0.00417438 | 0.00075633 | 0.00109632 | 0.00411411 | 0.00254852 | 0.00232681 |
| 24.9861111 | 0.01193402 | 0.0144141 | 0.01514046 | 0.01708094 | 0.01072121 | 0.03073831 | 0.00198411 | 0.00688139 | 0.00356502 | 0.00121871 | 0.0029648 | 0.00124061 | 0.00369748 | 0.00327577 | 0.00078917 | 0.00109746 | 0.00464502 | 0.0027222 | 0.00344143 |
| 25 | 0.01249478 | 0.01000229 | 0.01564956 | 0.02254643 | 0.00678714 | 0.03151036 | 0.00260421 | 0.00584404 | 0.00217794 | 0.00308345 | 0.0012187 | 0.00118636 | 0.00428844 | 0.00151502 | 0.000956 | 0.00087835 | 0.00324829 | 0.00456439 | 0.00435464 |
| 25.0138889 | 0.01144838 | 0.00718317 | 0.01746836 | 0.02006996 | 0.00644252 | 0.02846743 | 0.00371577 | 0.00461775 | 0.00377276 | 0.00620305 | 0.00161266 | 0.00092018 | 0.0030714 | 0.00069015 | 0.0007814 | 0.0006436 | 0.00168688 | 0.00675067 | 0.00416638 |
| 25.0277778 | 0.01329784 | 0.01120875 | 0.01896042 | 0.01277619 | 0.00917542 | 0.01980906 | 0.00347127 | 0.00419107 | 0.00487093 | 0.00660371 | 0.00307086 | 0.00216656 | 0.00144662 | 0.00123637 | 0.00203964 | 0.00336613 | 0.00213004 | 0.00782418 | 0.00462548 |
| 25.0416667 | 0.03853098 | 0.02428646 | 0.02081688 | 0.01946839 | 0.02263375 | 0.01545049 | 0.00615909 | 0.00697174 | 0.00797415 | 0.00760237 | 0.00717543 | 0.01002542 | 0.0053277 | 0.00525112 | 0.01605455 | 0.01844047 | 0.00835328 | 0.00936635 | 0.01425682 |
| 25.0555556 | 0.07370644 | 0.03339065 | 0.02211111 | 0.04747418 | 0.0360611 | 0.03251562 | 0.01608151 | 0.01229244 | 0.0195411 | 0.02266388 | 0.01946818 | 0.03195256 | 0.02308833 | 0.01581408 | 0.05706149 | 0.05191723 | 0.02465404 | 0.01520547 | 0.03711208 |
| 25.0694444 | 0.08342536 | 0.02811569 | 0.01997436 | 0.06455148 | 0.03364627 | 0.05598565 | 0.02670892 | 0.01222105 | 0.03142945 | 0.04426991 | 0.03453668 | 0.05368318 | 0.03886373 | 0.02565276 | 0.09226007 | 0.07720124 | 0.03795631 | 0.01917042 | 0.0523817 |
| 25.0833333 | 0.06405413 | 0.0184734 | 0.01522143 | 0.05104973 | 0.02082026 | 0.06759105 | 0.02901751 | 0.00675935 | 0.03307575 | 0.04170251 | 0.03145435 | 0.04328707 | 0.0308318 | 0.02130954 | 0.06952322 | 0.05768175 | 0.02885869 | 0.01422886 | 0.03992974 |
| 25.0972222 | 0.03855543 | 0.01404816 | 0.01020542 | 0.02970771 | 0.01135904 | 0.06121913 | 0.0232035 | 0.0043337 | 0.02515015 | 0.02064488 | 0.01463616 | 0.01730365 | 0.02022431 | 0.01076079 | 0.02997422 | 0.02403443 | 0.01267128 | 0.01055697 | 0.01738505 |
| 25.1111111 | 0.01939602 | 0.01012662 | 0.00544623 | 0.01589158 | 0.00670362 | 0.03711241 | 0.01439945 | 0.00555474 | 0.01379842 | 0.0129079 | 0.00761273 | 0.01235521 | 0.02523243 | 0.01044471 | 0.02893974 | 0.0197829 | 0.01126584 | 0.01339598 | 0.00817605 |
| 25.125 | 0.01562398 | 0.00725623 | 0.0039243 | 0.00730044 | 0.00343697 | 0.01485971 | 0.00740918 | 0.00598248 | 0.00627446 | 0.01640159 | 0.00765239 | 0.01806565 | 0.02829895 | 0.01332522 | 0.03796401 | 0.02545828 | 0.01326824 | 0.01465315 | 0.00875603 |
| 25.1388889 | 0.03084707 | 0.01380272 | 0.00803596 | 0.00414354 | 0.00240763 | 0.00996488 | 0.00664873 | 0.00533881 | 0.00698415 | 0.01723586 | 0.00863337 | 0.02020609 | 0.02230262 | 0.01072013 | 0.03822645 | 0.02583765 | 0.01134693 | 0.01210357 | 0.00975622 |
| 25.1527778 | 0.0450017 | 0.02408658 | 0.01324034 | 0.00790537 | 0.00384518 | 0.01468239 | 0.01081361 | 0.00548061 | 0.01349213 | 0.01569437 | 0.01242357 | 0.0205258 | 0.01534095 | 0.00717489 | 0.03581756 | 0.0256336 | 0.01086486 | 0.00971331 | 0.01072369 |
| 25.1666667 | 0.04350271 | 0.02505736 | 0.01514355 | 0.01582437 | 0.00588537 | 0.01723474 | 0.01386678 | 0.00652965 | 0.01920056 | 0.01441911 | 0.01406443 | 0.01964682 | 0.01050001 | 0.00618821 | 0.02710276 | 0.02136397 | 0.0113633 | 0.0082706 | 0.01040745 |
| 25.1805556 | 0.03266172 | 0.01607514 | 0.0137279 | 0.02398179 | 0.00652309 | 0.01563872 | 0.01368299 | 0.00641694 | 0.0183532 | 0.01192277 | 0.01024457 | 0.01899898 | 0.00616832 | 0.00676588 | 0.01427895 | 0.01281359 | 0.01145029 | 0.00632417 | 0.00799672 |
| 25.1944444 | 0.01939825 | 0.00602257 | 0.00911458 | 0.02835431 | 0.00546307 | 0.01288633 | 0.01042027 | 0.00461673 | 0.01049903 | 0.00644564 | 0.00652706 | 0.0173323 | 0.00308801 | 0.00621794 | 0.00550874 | 0.00462093 | 0.00986775 | 0.00533265 | 0.00765176 |
| 25.2083333 | 0.00976661 | 0.00165234 | 0.0046238 | 0.02720638 | 0.00443642 | 0.01324446 | 0.00543645 | 0.00335622 | 0.00274765 | 0.00218013 | 0.00575364 | 0.01246536 | 0.00325284 | 0.00390256 | 0.00436877 | 0.0024828 | 0.00625922 | 0.00617246 | 0.0072329 |
| 25.2222222 | 0.01177582 | 0.002431 | 0.00416825 | 0.02420734 | 0.00611994 | 0.01912994 | 0.00248363 | 0.00343329 | 0.00307514 | 0.00296024 | 0.00633688 | 0.00624642 | 0.00653391 | 0.00231622 | 0.00774817 | 0.00703778 | 0.00269779 | 0.00978479 | 0.00619019 |
| 25.2361111 | 0.02485603 | 0.00435489 | 0.00426619 | 0.02296503 | 0.0121358 | 0.02770336 | 0.00394789 | 0.00376994 | 0.01229791 | 0.0067281 | 0.00923218 | 0.00292072 | 0.01126575 | 0.00222117 | 0.01252892 | 0.01239002 | 0.00123996 | 0.01405021 | 0.00894777 |
| 25.25 | 0.03977087 | 0.00624027 | 0.00460989 | 0.02364501 | 0.02002859 | 0.03481709 | 0.00833751 | 0.00406024 | 0.02322425 | 0.00940243 | 0.00893899 | 0.00297051 | 0.01259089 | 0.001532 | 0.01296673 | 0.0132737 | 0.00091433 | 0.0122702 | 0.01025075 |
| 25.2638889 | 0.04397668 | 0.00834543 | 0.00801498 | 0.02322063 | 0.02296028 | 0.03723675 | 0.01159 | 0.00452413 | 0.02709053 | 0.01028734 | 0.00443232 | 0.00445274 | 0.00973207 | 0.00060163 | 0.00855213 | 0.01057853 | 0.00119948 | 0.00644471 | 0.00731313 |
| 25.2777778 | 0.02931519 | 0.00909974 | 0.00941463 | 0.01716007 | 0.01558187 | 0.03277675 | 0.01059956 | 0.00480048 | 0.02193628 | 0.01103333 | 0.00387314 | 0.00587818 | 0.00795956 | 0.00050421 | 0.005811 | 0.00857054 | 0.00223939 | 0.0048328 | 0.0051908 |
| 25.2916667 | 0.01284534 | 0.00838677 | 0.00582671 | 0.01008017 | 0.00772819 | 0.02492888 | 0.00601471 | 0.00475364 | 0.0131115 | 0.012202 | 0.00974165 | 0.00472526 | 0.01008436 | 0.00113529 | 0.00730481 | 0.00931408 | 0.0026857 | 0.00929289 | 0.0059137 |
| 25.3055556 | 0.01080955 | 0.00671233 | 0.0026324 | 0.01036681 | 0.00924635 | 0.01755234 | 0.00255956 | 0.00453593 | 0.0065504 | 0.01270604 | 0.01478365 | 0.00228693 | 0.0126665 | 0.00273268 | 0.00894866 | 0.01060689 | 0.00337268 | 0.01283787 | 0.00669605 |
| 25.3194444 | 0.01006386 | 0.00430878 | 0.00269934 | 0.01068848 | 0.01017681 | 0.01097468 | 0.00201311 | 0.00402369 | 0.0051468 | 0.01204993 | 0.01273491 | 0.00270414 | 0.01121485 | 0.00398744 | 0.00690435 | 0.01005514 | 0.00471543 | 0.01039159 | 0.00521824 |
| 25.3333333 | 0.00921369 | 0.00334075 | 0.00337278 | 0.00795083 | 0.00596191 | 0.00587939 | 0.0028218 | 0.00337718 | 0.00814979 | 0.01131438 | 0.00804485 | 0.00518223 | 0.00643312 | 0.0032695 | 0.00388318 | 0.00914055 | 0.00413133 | 0.00667637 | 0.00373713 |
| 25.3472222 | 0.01723622 | 0.00343855 | 0.00596259 | 0.01119324 | 0.0051338 | 0.0025589 | 0.0064391 | 0.00344221 | 0.01238232 | 0.0107717 | 0.0061285 | 0.00656751 | 0.00435202 | 0.00266145 | 0.00287079 | 0.00994789 | 0.00270105 | 0.00546084 | 0.00344529 |
| 25.3611111 | 0.02428527 | 0.00431055 | 0.01028685 | 0.01573061 | 0.00856078 | 0.00166471 | 0.01092721 | 0.00454677 | 0.01510724 | 0.01069573 | 0.00594079 | 0.00533378 | 0.00800733 | 0.0044122 | 0.00286772 | 0.01113665 | 0.0040085 | 0.00438049 | 0.00253547 |
| 25.375 | 0.01966424 | 0.00640305 | 0.01013018 | 0.01253062 | 0.0086896 | 0.00492601 | 0.011549 | 0.00546607 | 0.01431721 | 0.01180108 | 0.00517372 | 0.00614906 | 0.01043121 | 0.00650233 | 0.00273511 | 0.00955782 | 0.00722617 | 0.00373875 | 0.00250693 |
| 25.3888889 | 0.00917474 | 0.00595947 | 0.00589189 | 0.0054105 | 0.00498211 | 0.01006588 | 0.00832928 | 0.00515767 | 0.01036056 | 0.01291189 | 0.00719861 | 0.01301449 | 0.00700941 | 0.00753389 | 0.00441393 | 0.00533614 | 0.00966355 | 0.00731155 | 0.00543025 |
| 25.4027778 | 0.00372098 | 0.00320218 | 0.00355911 | 0.00278835 | 0.00265801 | 0.01127427 | 0.00469841 | 0.00432882 | 0.00760789 | 0.01308648 | 0.0109165 | 0.01689139 | 0.00475357 | 0.007381 | 0.00650305 | 0.00290009 | 0.00924088 | 0.00986315 | 0.00644126 |
| 25.4166667 | 0.00522775 | 0.00187199 | 0.0032747 | 0.00643023 | 0.00236549 | 0.00729401 | 0.00345095 | 0.0039159 | 0.01004786 | 0.01420406 | 0.01041102 | 0.0121519 | 0.00740947 | 0.00543115 | 0.00618753 | 0.00647202 | 0.00564119 | 0.00919687 | 0.00652146 |
| 25.4305556 | 0.01040399 | 0.00165917 | 0.00408669 | 0.0113237 | 0.00440128 | 0.00258163 | 0.00461904 | 0.00326338 | 0.01529648 | 0.01582947 | 0.00924465 | 0.00765297 | 0.00838536 | 0.00373041 | 0.00570289 | 0.01263548 | 0.00219875 | 0.01129513 | 0.0095775 |
| 25.4444444 | 0.01227456 | 0.00197207 | 0.00770231 | 0.01251756 | 0.00866377 | 0.0007206 | 0.00632054 | 0.00198385 | 0.01611695 | 0.01584473 | 0.00947502 | 0.00682707 | 0.00778853 | 0.00444746 | 0.00499418 | 0.01432953 | 0.00182807 | 0.01208243 | 0.00974477 |
| 25.4583333 | 0.00781605 | 0.00234333 | 0.01089981 | 0.00877975 | 0.01232059 | 0.00104447 | 0.00678536 | 0.00092386 | 0.01028986 | 0.0176921 | 0.0130763 | 0.00500006 | 0.00781183 | 0.0059156 | 0.00674673 | 0.01459289 | 0.00310765 | 0.01351034 | 0.00994504 |
| 25.4722222 | 0.00263569 | 0.00343457 | 0.0093766 | 0.00429651 | 0.01264655 | 0.00328216 | 0.00566875 | 0.00126369 | 0.00415131 | 0.02290528 | 0.02068092 | 0.0034902 | 0.00837585 | 0.00631875 | 0.01245055 | 0.01675297 | 0.00381204 | 0.01946095 | 0.01619763 |
| 25.4861111 | 0.00166638 | 0.00655392 | 0.00553852 | 0.00232249 | 0.0090476 | 0.00814519 | 0.00366054 | 0.00317672 | 0.00242086 | 0.02637659 | 0.02211523 | 0.00585235 | 0.01363041 | 0.00656119 | 0.01420622 | 0.01511769 | 0.00505596 | 0.01991115 | 0.02153941 |
| 25.5 | 0.00521207 | 0.00686951 | 0.00405867 | 0.00294076 | 0.00427759 | 0.01298908 | 0.00222311 | 0.00426354 | 0.00315787 | 0.02548756 | 0.01571926 | 0.01003597 | 0.01855274 | 0.00715886 | 0.01069338 | 0.00956407 | 0.00721753 | 0.01337005 | 0.02095516 |
| 25.5138889 | 0.0113364 | 0.00348982 | 0.00384 | 0.00743985 | 0.00159379 | 0.01631465 | 0.00316436 | 0.00375406 | 0.00739771 | 0.02203475 | 0.01116308 | 0.01101835 | 0.01617074 | 0.00706915 | 0.00901687 | 0.00635273 | 0.00867219 | 0.00966858 | 0.01837949 |
| 25.5277778 | 0.01922104 | 0.00402347 | 0.00549074 | 0.01298948 | 0.0026482 | 0.02051529 | 0.00711998 | 0.00338366 | 0.01658505 | 0.0212958 | 0.01369953 | 0.00955097 | 0.00940531 | 0.00681473 | 0.01099284 | 0.00762508 | 0.01031169 | 0.01430054 | 0.02093033 |
| 25.5416667 | 0.02990177 | 0.01232448 | 0.01609181 | 0.01720244 | 0.01011186 | 0.02772947 | 0.01448634 | 0.00382715 | 0.02431485 | 0.03157658 | 0.02221423 | 0.01882218 | 0.00748755 | 0.01033722 | 0.01727401 | 0.01395711 | 0.01682715 | 0.03128975 | 0.03598607 |
| 25.5555556 | 0.0444426 | 0.02477015 | 0.03761565 | 0.02578041 | 0.02338925 | 0.03579952 | 0.02354438 | 0.00459534 | 0.02605981 | 0.04819443 | 0.02852006 | 0.04508172 | 0.01690086 | 0.01973009 | 0.03222782 | 0.0255981 | 0.02911623 | 0.05061296 | 0.05643362 |
| 25.5694444 | 0.05929313 | 0.03778774 | 0.06005213 | 0.03994558 | 0.03049829 | 0.03782948 | 0.02936765 | 0.00831193 | 0.02371006 | 0.05416052 | 0.02488241 | 0.06416972 | 0.03136255 | 0.02732255 | 0.04751288 | 0.03343019 | 0.0384172 | 0.05197238 | 0.06165985 |
| 25.5833333 | 0.06140576 | 0.04941496 | 0.06689643 | 0.04703407 | 0.02308348 | 0.03108287 | 0.02953498 | 0.01964656 | 0.02058054 | 0.04714666 | 0.01536562 | 0.05818907 | 0.04102908 | 0.02577605 | 0.05188876 | 0.030142 | 0.037477 | 0.04206894 | 0.0496768 |
| 25.5972222 | 0.03971114 | 0.05097429 | 0.05242697 | 0.03477838 | 0.01165637 | 0.01814075 | 0.02247819 | 0.03418441 | 0.01628685 | 0.03407582 | 0.00667134 | 0.04077368 | 0.03984923 | 0.01966049 | 0.05060822 | 0.0233313 | 0.02771356 | 0.03577379 | 0.03453246 |
| 25.6111111 | 0.02367291 | 0.03882657 | 0.0268859 | 0.02134631 | 0.01144615 | 0.01053796 | 0.01197865 | 0.04287864 | 0.00985068 | 0.01870742 | 0.00199371 | 0.02347647 | 0.02700958 | 0.01458991 | 0.04524395 | 0.01820938 | 0.01514996 | 0.02914078 | 0.02178834 |
| 25.625 | 0.04087308 | 0.02387501 | 0.01270536 | 0.03026307 | 0.02159223 | 0.01874817 | 0.00851851 | 0.04571699 | 0.00726531 | 0.01215779 | 0.00282685 | 0.01116584 | 0.01286902 | 0.00988788 | 0.03244619 | 0.01281518 | 0.0084921 | 0.0188684 | 0.01175306 |
| 25.6388889 | 0.05821491 | 0.01299843 | 0.02053176 | 0.04439758 | 0.02656294 | 0.02808061 | 0.01080181 | 0.04699477 | 0.00922558 | 0.01702237 | 0.00734624 | 0.00861026 | 0.00970804 | 0.00573053 | 0.01683258 | 0.00672841 | 0.00683079 | 0.01049002 | 0.00501114 |

| | | | | | | | | | | | | | | | | | | | |
|---|---|---|---|---|---|---|---|---|---|---|---|---|---|---|---|---|---|---|---|
| 25.6527778 | 0.05370742 | 0.00855651 | 0.02710161 | 0.04763814 | 0.0205092 | 0.02729813 | 0.00908806 | 0.04513447 | 0.00840104 | 0.01935406 | 0.01093019 | 0.00832176 | 0.01717925 | 0.00331476 | 0.00899699 | 0.00449613 | 0.00747524 | 0.00659446 | 0.00309964 |
| 25.6666667 | 0.03553122 | 0.01556003 | 0.01689364 | 0.03982891 | 0.01079537 | 0.02433456 | 0.00402854 | 0.03762964 | 0.01061563 | 0.01522291 | 0.00975951 | 0.00720011 | 0.02332581 | 0.00277434 | 0.01622998 | 0.01020476 | 0.0147202 | 0.00411129 | 0.00768449 |
| 25.6805556 | 0.02241622 | 0.02941125 | 0.00539947 | 0.02454 | 0.0068239 | 0.01940691 | 0.00186433 | 0.02521989 | 0.02236074 | 0.01057408 | 0.00964641 | 0.01187933 | 0.01908542 | 0.00312887 | 0.02840431 | 0.01796122 | 0.01773947 | 0.00343596 | 0.01408505 |
| 25.6944444 | 0.03588145 | 0.0406805 | 0.0055077 | 0.02149871 | 0.01211125 | 0.01023161 | 0.00209649 | 0.02195215 | 0.02971955 | 0.00859507 | 0.01530845 | 0.01703582 | 0.0104599 | 0.00360517 | 0.03342659 | 0.02235147 | 0.01064439 | 0.00639614 | 0.01652561 |
| 25.7083333 | 0.04994926 | 0.04335101 | 0.00820144 | 0.02817685 | 0.01806692 | 0.0061146 | 0.00167908 | 0.03857569 | 0.02067883 | 0.01177443 | 0.01819053 | 0.01335072 | 0.00863183 | 0.00798912 | 0.03062416 | 0.02191886 | 0.00887352 | 0.00922265 | 0.01618072 |
| 25.7222222 | 0.03490888 | 0.03363294 | 0.0106097 | 0.02219475 | 0.01644174 | 0.00770948 | 0.00154262 | 0.05670394 | 0.01072952 | 0.01871046 | 0.01306118 | 0.00748767 | 0.01087747 | 0.01543471 | 0.02107325 | 0.01456653 | 0.01898823 | 0.00855849 | 0.01543368 |
| 25.7361111 | 0.01862992 | 0.0175794 | 0.02095818 | 0.01216582 | 0.00902416 | 0.00510339 | 0.00170365 | 0.0617518 | 0.01697233 | 0.02255718 | 0.0063542 | 0.01295131 | 0.00838745 | 0.02065552 | 0.01126833 | 0.00935885 | 0.02878113 | 0.00561682 | 0.01288391 |
| 25.75 | 0.03097107 | 0.0122262 | 0.02619349 | 0.01686978 | 0.00450342 | 0.00299758 | 0.00138373 | 0.05257744 | 0.02787583 | 0.01696413 | 0.00451036 | 0.0236608 | 0.00583812 | 0.01943965 | 0.01106196 | 0.01470457 | 0.0229579 | 0.00288688 | 0.00882542 |
| 25.7638889 | 0.05540488 | 0.02593054 | 0.01660533 | 0.03082694 | 0.01069754 | 0.01028766 | 0.00161801 | 0.03249243 | 0.02991014 | 0.00973533 | 0.00508468 | 0.02565077 | 0.00898468 | 0.01191553 | 0.01695208 | 0.02128254 | 0.01209675 | 0.00122901 | 0.00996407 |
| 25.7777778 | 0.06787408 | 0.04381306 | 0.00923138 | 0.04229657 | 0.02385383 | 0.02051109 | 0.00336788 | 0.01458672 | 0.02079397 | 0.01405791 | 0.00394875 | 0.01738758 | 0.0103727 | 0.00869209 | 0.01854093 | 0.02108884 | 0.01720998 | 0.00047712 | 0.01671431 |
| 25.7916667 | 0.05520991 | 0.04932712 | 0.01361258 | 0.04244981 | 0.02824283 | 0.02925972 | 0.00626948 | 0.00964071 | 0.01189191 | 0.0234888 | 0.00217501 | 0.01019885 | 0.00628991 | 0.01304695 | 0.01381215 | 0.0146743 | 0.02868122 | 0.00063302 | 0.01805119 |
| 25.8055556 | 0.02920004 | 0.04074542 | 0.01648215 | 0.03042857 | 0.01801175 | 0.03993598 | 0.00695097 | 0.01467386 | 0.01773066 | 0.02638732 | 0.0023758 | 0.01454292 | 0.00417883 | 0.0143186 | 0.00737986 | 0.00715305 | 0.02674611 | 0.0008361 | 0.01124309 |
| 25.8194444 | 0.02234888 | 0.0227668 | 0.01230296 | 0.01723647 | 0.00952209 | 0.04345772 | 0.00464455 | 0.01652233 | 0.02697072 | 0.021754 | 0.00513011 | 0.02092936 | 0.00677459 | 0.00876924 | 0.0056591 | 0.00618931 | 0.01425794 | 0.00054051 | 0.00459831 |
| 25.8333333 | 0.02662213 | 0.011855 | 0.01239741 | 0.01030118 | 0.01369966 | 0.03474993 | 0.00278159 | 0.01010461 | 0.02492235 | 0.01243614 | 0.00688977 | 0.01917325 | 0.00774165 | 0.00537825 | 0.01071644 | 0.01165649 | 0.00920647 | 0.00058504 | 0.00294401 |
| 25.8472222 | 0.01985254 | 0.02223422 | 0.01883792 | 0.01158818 | 0.02023702 | 0.0248842 | 0.00578079 | 0.00541156 | 0.01549914 | 0.00818231 | 0.00582371 | 0.0114565 | 0.00529416 | 0.00877467 | 0.01895656 | 0.01768065 | 0.01487308 | 0.00148231 | 0.00441721 |
| 25.8611111 | 0.01444833 | 0.04114569 | 0.02544866 | 0.0173271 | 0.02038577 | 0.02348771 | 0.01692011 | 0.00993929 | 0.01474438 | 0.01619299 | 0.00727314 | 0.00913395 | 0.00517242 | 0.01139296 | 0.0275132 | 0.02336642 | 0.01736538 | 0.00246095 | 0.00643966 |
| 25.875 | 0.01812963 | 0.04771686 | 0.03054417 | 0.01999443 | 0.01559303 | 0.02733488 | 0.02782618 | 0.01623587 | 0.02870875 | 0.02787169 | 0.01308812 | 0.01958392 | 0.00884109 | 0.00910029 | 0.03414631 | 0.03031885 | 0.0107405 | 0.00239657 | 0.0093354 |
| 25.8888889 | 0.0223238 | 0.0360619 | 0.03128818 | 0.01854949 | 0.00907634 | 0.02641219 | 0.02637405 | 0.01573637 | 0.03944678 | 0.03543154 | 0.01680242 | 0.0284153 | 0.0114211 | 0.00636199 | 0.03592526 | 0.03481773 | 0.0035302 | 0.00180709 | 0.01309565 |
| 25.9027778 | 0.0271284 | 0.01788224 | 0.02911277 | 0.01606065 | 0.01004221 | 0.02211708 | 0.01752576 | 0.00911994 | 0.03415333 | 0.03917884 | 0.01457887 | 0.02559952 | 0.01038311 | 0.0083696 | 0.03283028 | 0.03202603 | 0.00320731 | 0.00282867 | 0.01573068 |
| 25.9166667 | 0.03249526 | 0.00614616 | 0.02802273 | 0.01579286 | 0.02628051 | 0.02045311 | 0.01220894 | 0.00315233 | 0.01920383 | 0.0391458 | 0.01020356 | 0.01894879 | 0.00761866 | 0.01523098 | 0.02760515 | 0.02219793 | 0.00721964 | 0.00469384 | 0.01651773 |
| 25.9305556 | 0.03212991 | 0.00279211 | 0.02323433 | 0.01883238 | 0.04114618 | 0.01863852 | 0.01147961 | 0.00168426 | 0.00810626 | 0.03527653 | 0.00956701 | 0.01323933 | 0.00491104 | 0.02061452 | 0.02201257 | 0.01056207 | 0.01039124 | 0.00397566 | 0.01737467 |
| 25.9444444 | 0.02605514 | 0.00495162 | 0.01255239 | 0.02325971 | 0.0342876 | 0.01204944 | 0.01188421 | 0.00147981 | 0.00739415 | 0.02769536 | 0.0137057 | 0.00845902 | 0.00280301 | 0.02044236 | 0.01586489 | 0.00364782 | 0.01040677 | 0.00271265 | 0.01901188 |
| 25.9583333 | 0.02077321 | 0.00846235 | 0.00547771 | 0.02664211 | 0.01786279 | 0.00773596 | 0.0101081 | 0.00111808 | 0.0089196 | 0.01716735 | 0.01800701 | 0.00506525 | 0.00191251 | 0.01671898 | 0.00868389 | 0.00199262 | 0.00805538 | 0.00506562 | 0.01970488 |
| 25.9722222 | 0.01756226 | 0.01052697 | 0.00686938 | 0.02445158 | 0.01322202 | 0.0155236 | 0.00579687 | 0.00235386 | 0.00702372 | 0.00818008 | 0.01576859 | 0.00260346 | 0.00212972 | 0.01255434 | 0.00337179 | 0.00231304 | 0.00533851 | 0.00799654 | 0.01772432 |
| 25.9861111 | 0.01135292 | 0.01085434 | 0.00872119 | 0.01504667 | 0.01623285 | 0.02843014 | 0.00338843 | 0.00469359 | 0.00529555 | 0.00586814 | 0.00956157 | 0.00168915 | 0.00202015 | 0.0089844 | 0.00280669 | 0.00495216 | 0.00388397 | 0.007571 | 0.01244 |
| 26 | 0.00751481 | 0.01071852 | 0.00640969 | 0.00714427 | 0.01393295 | 0.03560749 | 0.00475214 | 0.00618025 | 0.00636525 | 0.00757818 | 0.00881038 | 0.00252275 | 0.00117135 | 0.0054613 | 0.0041985 | 0.00813952 | 0.00325269 | 0.00497432 | 0.00592199 |
| 26.0138889 | 0.01480801 | 0.01105765 | 0.004135 | 0.00832019 | 0.00867626 | 0.03509505 | 0.00604893 | 0.00556819 | 0.00934754 | 0.00782504 | 0.01044199 | 0.00233124 | 0.00087806 | 0.00241903 | 0.00406891 | 0.00911513 | 0.00246351 | 0.00292964 | 0.0023936 |
| 26.0277778 | 0.02058522 | 0.00991634 | 0.00425499 | 0.01066664 | 0.00575099 | 0.02648363 | 0.00445641 | 0.00348 | 0.00938401 | 0.00570213 | 0.0078544 | 0.00295756 | 0.00361477 | 0.00221895 | 0.00476582 | 0.01143845 | 0.00177765 | 0.00494983 | 0.00326319 |
| 26.0416667 | 0.02144854 | 0.00797625 | 0.00529218 | 0.01253476 | 0.00880601 | 0.01809826 | 0.0050557 | 0.00318522 | 0.01084268 | 0.01013272 | 0.0084984 | 0.01240865 | 0.01655409 | 0.00885194 | 0.01471696 | 0.02238775 | 0.00524595 | 0.01891782 | 0.01265591 |
| 26.0555556 | 0.04131326 | 0.0118006 | 0.00883069 | 0.02738089 | 0.02068397 | 0.03015288 | 0.01445997 | 0.00779458 | 0.02846967 | 0.02829661 | 0.01785862 | 0.03154045 | 0.04065289 | 0.02122392 | 0.03578139 | 0.04179223 | 0.01687529 | 0.04527544 | 0.03685186 |
| 26.0694444 | 0.07368585 | 0.01924864 | 0.01343341 | 0.04739076 | 0.03707121 | 0.05368671 | 0.02440126 | 0.01530723 | 0.05642474 | 0.04686968 | 0.02480029 | 0.04324405 | 0.0539594 | 0.02686771 | 0.04386529 | 0.04939595 | 0.0256757 | 0.06096427 | 0.05433559 |
| 26.0833333 | 0.07749174 | 0.02023968 | 0.01490025 | 0.05213059 | 0.04799693 | 0.0555107 | 0.02624068 | 0.01832321 | 0.07195114 | 0.04147604 | 0.02096102 | 0.03100982 | 0.03863114 | 0.01868156 | 0.02935941 | 0.03128221 | 0.01962368 | 0.05000317 | 0.04032596 |
| 26.0972222 | 0.04821891 | 0.01443814 | 0.01840348 | 0.046562 | 0.0422985 | 0.03589131 | 0.03093575 | 0.01323756 | 0.06438785 | 0.01956536 | 0.01128677 | 0.01201016 | 0.01831179 | 0.00762952 | 0.02079722 | 0.01106106 | 0.01079534 | 0.02589627 | 0.01682881 |
| 26.1111111 | 0.01835582 | 0.0077888 | 0.02386179 | 0.03527226 | 0.02479947 | 0.01526173 | 0.03065741 | 0.0070423 | 0.03743759 | 0.01248593 | 0.00657378 | 0.00750858 | 0.01866369 | 0.00391918 | 0.02431717 | 0.00784406 | 0.00991733 | 0.01218681 | 0.0109249 |
| 26.125 | 0.00696673 | 0.00398128 | 0.01872832 | 0.01700693 | 0.01862895 | 0.00468066 | 0.01333123 | 0.00688598 | 0.01760377 | 0.01304537 | 0.00840047 | 0.01041161 | 0.02519349 | 0.00426589 | 0.0246968 | 0.00899572 | 0.00875473 | 0.01428275 | 0.01353748 |
| 26.1388889 | 0.01158929 | 0.00387277 | 0.00778817 | 0.00651166 | 0.02928883 | 0.00492487 | 0.00424845 | 0.00940949 | 0.02564674 | 0.0091251 | 0.00932843 | 0.01383468 | 0.02238512 | 0.00442103 | 0.01651903 | 0.00622205 | 0.00673903 | 0.01488899 | 0.01168361 |
| 26.1527778 | 0.02119092 | 0.00406877 | 0.00616594 | 0.00721898 | 0.03607448 | 0.00777521 | 0.01029005 | 0.00897773 | 0.03999671 | 0.01027511 | 0.01044281 | 0.01489514 | 0.01581985 | 0.00414929 | 0.00716033 | 0.00421616 | 0.0076323 | 0.01267738 | 0.01018868 |
| 26.1666667 | 0.02855408 | 0.00305456 | 0.00862978 | 0.01221838 | 0.03059423 | 0.00828467 | 0.01509473 | 0.00627155 | 0.04167275 | 0.01104479 | 0.0104331 | 0.00974922 | 0.01199145 | 0.00272446 | 0.00520057 | 0.00347819 | 0.00646127 | 0.01155967 | 0.00909787 |
| 26.1805556 | 0.03096136 | 0.0028355 | 0.00715421 | 0.01802402 | 0.02280645 | 0.00670913 | 0.01520859 | 0.00431682 | 0.03449303 | 0.00680086 | 0.00873796 | 0.00513163 | 0.00913293 | 0.00114717 | 0.00685481 | 0.00264515 | 0.00439853 | 0.00862394 | 0.00610265 |
| 26.1944444 | 0.02871582 | 0.00231559 | 0.0042577 | 0.0202597 | 0.01967716 | 0.00411466 | 0.01209634 | 0.00378237 | 0.02463829 | 0.00382058 | 0.00723442 | 0.00578614 | 0.00666756 | 0.00082771 | 0.00810373 | 0.0026079 | 0.0060499 | 0.00751469 | 0.0049157 |
| 26.2083333 | 0.02540506 | 0.00140954 | 0.00317161 | 0.01858857 | 0.0182121 | 0.0025339 | 0.00839816 | 0.00344692 | 0.01727787 | 0.00339024 | 0.00359479 | 0.00670694 | 0.00792165 | 0.00082299 | 0.00596228 | 0.00181022 | 0.00712937 | 0.00679251 | 0.00302793 |
| 26.2222222 | 0.0227662 | 0.002878 | 0.00574369 | 0.01553819 | 0.01301529 | 0.00381883 | 0.00745904 | 0.00306529 | 0.01463027 | 0.00323337 | 0.00157857 | 0.00536425 | 0.0086641 | 0.00054174 | 0.00223434 | 0.00048389 | 0.00503466 | 0.00566664 | 0.00177237 |
| 26.2361111 | 0.02168861 | 0.00732748 | 0.01114123 | 0.01352512 | 0.00631308 | 0.00920154 | 0.01114725 | 0.00343566 | 0.01687369 | 0.00253802 | 0.00137611 | 0.00375559 | 0.00514277 | 0.00063254 | 0.00185064 | 0.00026555 | 0.0043175 | 0.00600015 | 0.00206574 |
| 26.25 | 0.02212325 | 0.0117042 | 0.01538478 | 0.01329346 | 0.00324289 | 0.01653255 | 0.01639801 | 0.00443523 | 0.02295309 | 0.0041108 | 0.00154702 | 0.00317639 | 0.00213366 | 0.00140606 | 0.00289168 | 0.0008945 | 0.00479028 | 0.00817111 | 0.00200783 |
| 26.2638889 | 0.02151618 | 0.01198684 | 0.01514681 | 0.01415483 | 0.00516926 | 0.01987926 | 0.01800486 | 0.00523537 | 0.02957361 | 0.00874668 | 0.00588437 | 0.00326459 | 0.0026355 | 0.00197416 | 0.00417959 | 0.00282529 | 0.00433434 | 0.00873624 | 0.0035367 |
| 26.2777778 | 0.01885162 | 0.00891803 | 0.01157045 | 0.01543896 | 0.00883552 | 0.01802229 | 0.01556562 | 0.00561347 | 0.03229714 | 0.01291417 | 0.0120367 | 0.00432152 | 0.00348496 | 0.00211805 | 0.00792119 | 0.00503788 | 0.00495324 | 0.0082066 | 0.00990381 |
| 26.2916667 | 0.01615821 | 0.00711894 | 0.00781785 | 0.01606801 | 0.00901404 | 0.01609883 | 0.01166224 | 0.00583828 | 0.0294228 | 0.0125965 | 0.01166492 | 0.0060866 | 0.00265623 | 0.00277486 | 0.01019601 | 0.00509475 | 0.0043042 | 0.00864489 | 0.01359992 |
| 26.3055556 | 0.01430299 | 0.00759469 | 0.00516092 | 0.01483178 | 0.00525381 | 0.01630269 | 0.00719102 | 0.00558233 | 0.0222996 | 0.0071868 | 0.00665295 | 0.00616815 | 0.00262271 | 0.00344596 | 0.0070148 | 0.002901 | 0.00196311 | 0.00636077 | 0.01056436 |
| 26.3194444 | 0.01275501 | 0.00669809 | 0.00392313 | 0.0122022 | 0.00267098 | 0.01588327 | 0.0033931 | 0.0043041 | 0.01433315 | 0.00253177 | 0.00534177 | 0.00397363 | 0.00419006 | 0.00297595 | 0.00304699 | 0.00096713 | 0.00207498 | 0.00509802 | 0.00498772 |
| 26.3333333 | 0.01137739 | 0.0041499 | 0.00467302 | 0.00975481 | 0.00350891 | 0.01273341 | 0.00248238 | 0.00252606 | 0.01087866 | 0.00464016 | 0.00589909 | 0.00174511 | 0.00429708 | 0.00166202 | 0.00203722 | 0.00215377 | 0.00420234 | 0.00674954 | 0.00204611 |
| 26.3472222 | 0.00805408 | 0.00375499 | 0.00549181 | 0.0073122 | 0.00435407 | 0.00810794 | 0.00402945 | 0.00150892 | 0.01467319 | 0.01161637 | 0.00659432 | 0.00223554 | 0.00355223 | 0.00104801 | 0.00228398 | 0.00605869 | 0.00535302 | 0.00599822 | 0.00293935 |
| 26.3611111 | 0.00544644 | 0.00413652 | 0.00433768 | 0.00466467 | 0.0032933 | 0.00451587 | 0.00456291 | 0.00214685 | 0.01904402 | 0.01424112 | 0.00727843 | 0.00504548 | 0.00355552 | 0.00212928 | 0.00268205 | 0.0077998 | 0.00665561 | 0.00563734 | 0.00323362 |
| 26.375 | 0.00944068 | 0.00340656 | 0.00362798 | 0.00482981 | 0.00387134 | 0.00249864 | 0.00305766 | 0.00402629 | 0.01457532 | 0.0094106 | 0.00556241 | 0.00592449 | 0.00221264 | 0.00373681 | 0.00268506 | 0.00539187 | 0.00877183 | 0.00703289 | 0.00166521 |
| 26.3888889 | 0.01340939 | 0.00498375 | 0.00341988 | 0.00763024 | 0.00823396 | 0.00172872 | 0.00190491 | 0.00555007 | 0.00881571 | 0.00442914 | 0.0053443 | 0.00397952 | 0.00049082 | 0.00416036 | 0.00196429 | 0.00246595 | 0.00708113 | 0.00585054 | 0.00142546 |
| 26.4027778 | 0.00985193 | 0.0085346 | 0.00389322 | 0.00722375 | 0.01105959 | 0.00280494 | 0.00220544 | 0.00574598 | 0.00862983 | 0.00592121 | 0.00752478 | 0.00285257 | 0.00031335 | 0.00357443 | 0.00205295 | 0.00142036 | 0.00286 | 0.00555595 | 0.00172654 |
| 26.4166667 | 0.00728887 | 0.01005941 | 0.00940851 | 0.00478452 | 0.00865791 | 0.00551929 | 0.00470172 | 0.00466254 | 0.0099575 | 0.01454923 | 0.00995551 | 0.00245686 | 0.00215514 | 0.00273215 | 0.00271059 | 0.00283521 | 0.00087384 | 0.00621165 | 0.00110445 |
| 26.4305556 | 0.0131599 | 0.00721241 | 0.01628285 | 0.00745408 | 0.00428056 | 0.00888536 | 0.00955498 | 0.00261852 | 0.01955633 | 0.02373935 | 0.01272942 | 0.00206874 | 0.00574723 | 0.00212067 | 0.00373321 | 0.00650774 | 0.00311616 | 0.00520129 | 0.00151507 |
| 26.4444444 | 0.01880627 | 0.00366663 | 0.01717247 | 0.01194583 | 0.00229912 | 0.01161462 | 0.0144178 | 0.00114468 | 0.03158088 | 0.02463723 | 0.01355565 | 0.00376775 | 0.00822645 | 0.00288781 | 0.00685048 | 0.00995369 | 0.01041643 | 0.00839018 | 0.0036787 |
| 26.4583333 | 0.01661237 | 0.00300648 | 0.01262009 | 0.01161862 | 0.00371727 | 0.01077146 | 0.01763608 | 0.00094399 | 0.02937434 | 0.01837688 | 0.01259391 | 0.00730189 | 0.00804174 | 0.00541854 | 0.00975586 | 0.01153618 | 0.01808376 | 0.0142051 | 0.00630667 |
| 26.4722222 | 0.01104854 | 0.00417572 | 0.00845895 | 0.00826748 | 0.00634148 | 0.00775796 | 0.01850714 | 0.00190163 | 0.01622037 | 0.01387786 | 0.0107903 | 0.01070711 | 0.00637786 | 0.00858233 | 0.00885631 | 0.0116677 | 0.02101229 | 0.01338901 | 0.00696025 |
| 26.4861111 | 0.00731185 | 0.00772124 | 0.0087796 | 0.00485885 | 0.0080836 | 0.00874506 | 0.01601498 | 0.00442927 | 0.0068519 | 0.01558444 | 0.00979532 | 0.01159068 | 0.00570353 | 0.01041433 | 0.00783198 | 0.01216318 | 0.01891221 | 0.00851879 | 0.0066032 |
| 26.5 | 0.00524995 | 0.00931317 | 0.01179199 | 0.0024026 | 0.00697955 | 0.01210017 | 0.01053151 | 0.00580518 | 0.00469691 | 0.02023279 | 0.01305218 | 0.00770321 | 0.00788742 | 0.00908994 | 0.01194669 | 0.01425271 | 0.01326316 | 0.00982515 | 0.00978644 |
| 26.5138889 | 0.00436256 | 0.00585895 | 0.01252027 | 0.0023563 | 0.00418395 | 0.01262214 | 0.00471081 | 0.00401329 | 0.00724244 | 0.02257455 | 0.01532667 | 0.00253345 | 0.01053165 | 0.00602494 | 0.01609899 | 0.01555632 | 0.00854553 | 0.01409173 | 0.01375982 |
| 26.5277778 | 0.00773183 | 0.00277811 | 0.00947556 | 0.00723778 | 0.00266907 | 0.01312827 | 0.00375207 | 0.00222399 | 0.020606 | 0.02303275 | 0.01203262 | 0.00274957 | 0.0132215 | 0.00619598 | 0.01679001 | 0.01719425 | 0.01010434 | 0.01525411 | 0.01656877 |
| 26.5416667 | 0.02128523 | 0.00312286 | 0.00902447 | 0.02084889 | 0.00355279 | 0.02162843 | 0.01363723 | 0.00529059 | 0.03812404 | 0.03017972 | 0.01087366 | 0.01294887 | 0.02413605 | 0.01550967 | 0.02601183 | 0.0295006 | 0.01884668 | 0.02092289 | 0.02700176 |
| 26.5555556 | 0.0463547 | 0.0076533 | 0.02112717 | 0.0437671 | 0.00736698 | 0.03748943 | 0.03379469 | 0.01354049 | 0.04589069 | 0.04537051 | 0.01538972 | 0.0312019 | 0.04406043 | 0.03211572 | 0.0453037 | 0.05030175 | 0.03027277 | 0.03140401 | 0.042902 |
| 26.5694444 | 0.06879345 | 0.01577836 | 0.04464882 | 0.06690468 | 0.00803995 | 0.04776017 | 0.05559865 | 0.01952329 | 0.04563796 | 0.05331455 | 0.01912167 | 0.0436841 | 0.05742535 | 0.04435659 | 0.05394779 | 0.05836094 | 0.03815887 | 0.03380745 | 0.04790414 |
| 26.5833333 | 0.07550535 | 0.02311066 | 0.0639333 | 0.07909613 | 0.0073936 | 0.0428569 | 0.06862661 | 0.01876565 | 0.04096335 | 0.04493818 | 0.01691086 | 0.0432104 | 0.05674291 | 0.04756377 | 0.04791965 | 0.04959232 | 0.04084405 | 0.02585517 | 0.03806518 |
| 26.5972222 | 0.06581432 | 0.02427913 | 0.06287527 | 0.07396733 | 0.0170061 | 0.0265092 | 0.06191971 | 0.01497862 | 0.02931753 | 0.02595962 | 0.00995434 | 0.03621286 | 0.04644721 | 0.04306953 | 0.04334926 | 0.03823784 | 0.03724735 | 0.01666464 | 0.02318698 |
| 26.6111111 | 0.04309806 | 0.01650673 | 0.03892992 | 0.04926807 | 0.03104792 | 0.01142734 | 0.03518824 | 0.01013179 | 0.01593009 | 0.01367391 | 0.00694378 | 0.02556303 | 0.02798119 | 0.03001747 | 0.04177694 | 0.02777227 | 0.02492712 | 0.01017882 | 0.01065525 |
| 26.625 | 0.01980842 | 0.00619457 | 0.01874686 | 0.0239994 | 0.03779973 | 0.00846187 | 0.01911449 | 0.00532692 | 0.01471488 | 0.02234227 | 0.01226375 | 0.01326634 | 0.01387964 | 0.01560259 | 0.03490643 | 0.01581255 | 0.01360996 | 0.00545348 | 0.00501034 |
| 26.6388889 | 0.00910644 | 0.00102818 | 0.02559778 | 0.02028677 | 0.03789625 | 0.01719933 | 0.03457192 | 0.00336872 | 0.02771472 | 0.03619417 | 0.01629567 | 0.00610642 | 0.01647924 | 0.0125332 | 0.02117725 | 0.01203635 | 0.01487128 | 0.00325769 | 0.00846994 |
| 26.6527778 | 0.01221893 | 0.00035111 | 0.03907507 | 0.02596874 | 0.03410365 | 0.02770746 | 0.05396744 | 0.00280307 | 0.03700337 | 0.03580195 | 0.01134322 | 0.00629224 | 0.0245902 | 0.02001016 | 0.0116491 | 0.02299798 | 0.01878871 | 0.00377263 | 0.01534806 |
| 26.6666667 | 0.01969082 | 0.0012002 | 0.0397163 | 0.02198093 | 0.02529981 | 0.03850586 | 0.04998327 | 0.00211095 | 0.02688551 | 0.02201225 | 0.00695462 | 0.0065411 | 0.02406786 | 0.02365001 | 0.01804833 | 0.03409228 | 0.01393966 | 0.00398502 | 0.02035818 |
| 26.6805556 | 0.02124995 | 0.00168005 | 0.03430382 | 0.01155415 | 0.01418201 | 0.04962389 | 0.03044391 | 0.00215523 | 0.01445929 | 0.00675362 | 0.00864487 | 0.00338072 | 0.01400692 | 0.01742189 | 0.0311458 | 0.03400973 | 0.01039748 | 0.00400401 | 0.0242572 |
| 26.6944444 | 0.01443703 | 0.00133599 | 0.02277238 | 0.00706984 | 0.00630682 | 0.0529935 | 0.02231916 | 0.00242392 | 0.02041539 | 0.00515233 | 0.00844877 | 0.0036232 | 0.00409321 | 0.00892928 | 0.03797677 | 0.02811622 | 0.01262912 | 0.00823752 | 0.02669461 |
| 26.7083333 | 0.01376241 | 0.00150222 | 0.01591256 | 0.01392255 | 0.00718548 | 0.04462572 | 0.03599631 | 0.00354326 | 0.02542406 | 0.01791551 | 0.01043886 | 0.00639205 | 0.00526166 | 0.00695534 | 0.03752114 | 0.02264832 | 0.01189485 | 0.01440424 | 0.02535481 |
| 26.7222222 | 0.03049716 | 0.00237449 | 0.02702679 | 0.02640656 | 0.01728543 | 0.02575702 | 0.04984963 | 0.00535394 | 0.01506787 | 0.03320755 | 0.01943456 | 0.00704842 | 0.01648744 | 0.01212354 | 0.0317737 | 0.01748942 | 0.01468519 | 0.01462823 | 0.02014717 |
| 26.7361111 | 0.04421079 | 0.00239215 | 0.02926236 | 0.0298989 | 0.02505233 | 0.01583281 | 0.03822368 | 0.00632645 | 0.0071664 | 0.03953873 | 0.02415362 | 0.00797645 | 0.02480076 | 0.01781304 | 0.02355225 | 0.0102557 | 0.02549552 | 0.00900576 | 0.01237535 |
| 26.75 | 0.03496661 | 0.00496706 | 0.01350118 | 0.0181494 | 0.0231013 | 0.02593368 | 0.0193806 | 0.00552211 | 0.01100179 | 0.02827854 | 0.01687558 | 0.00949879 | 0.01976395 | 0.0178991 | 0.01527815 | 0.00526399 | 0.03094284 | 0.00532398 | 0.00664842 |
| 26.7638889 | 0.01633367 | 0.01305062 | 0.00537545 | 0.00684567 | 0.01481191 | 0.03516083 | 0.02654974 | 0.00299094 | 0.01884975 | 0.0190444 | 0.00874284 | 0.00976534 | 0.01010882 | 0.01153696 | 0.0077075 | 0.00807162 | 0.02252041 | 0.00709521 | 0.01060738 |
| 26.7777778 | 0.00869046 | 0.01747314 | 0.00982845 | 0.00436793 | 0.00643845 | 0.03448016 | 0.04523013 | 0.0008866 | 0.02363819 | 0.03435292 | 0.01021349 | 0.00777279 | 0.00893676 | 0.00547836 | 0.0037264 | 0.01328789 | 0.01233613 | 0.01211145 | 0.02365746 |
| 26.7916667 | 0.00939276 | 0.01152345 | 0.01525951 | 0.00284779 | 0.00452639 | 0.02882285 | 0.04999516 | 0.00045944 | 0.02240921 | 0.04865555 | 0.01106376 | 0.00772528 | 0.01029881 | 0.00554628 | 0.00435214 | 0.01256841 | 0.01461007 | 0.01620961 | 0.03478596 |
| 26.8055556 | 0.01418999 | 0.00435096 | 0.01469813 | 0.0052896 | 0.01141765 | 0.01858188 | 0.03672344 | 0.00136229 | 0.01687545 | 0.03829901 | 0.00867514 | 0.00866788 | 0.00731047 | 0.0071731 | 0.00350574 | 0.00911503 | 0.01775541 | 0.01320267 | 0.03464734 |
| 26.8194444 | 0.03029518 | 0.00297835 | 0.01213154 | 0.0179002 | 0.02029755 | 0.01694235 | 0.01629498 | 0.00340983 | 0.00933451 | 0.02211814 | 0.01351602 | 0.0068633 | 0.00540272 | 0.00549262 | 0.00388187 | 0.01334886 | 0.01261758 | 0.00595747 | 0.02400228 |
| 26.8333333 | 0.04553158 | 0.00296828 | 0.01423619 | 0.02733036 | 0.02033287 | 0.03235968 | 0.00362648 | 0.00520672 | 0.00486895 | 0.02152402 | 0.01885829 | 0.00945577 | 0.00737624 | 0.00421775 | 0.01014512 | 0.02478288 | 0.01253454 | 0.00266295 | 0.01293216 |
| 26.8472222 | 0.03789395 | 0.00415615 | 0.01709519 | 0.0202087 | 0.0126727 | 0.04954819 | 0.00072088 | 0.00526789 | 0.007847 | 0.02555299 | 0.01502867 | 0.01912891 | 0.01290391 | 0.00694196 | 0.01957904 | 0.03375347 | 0.02262994 | 0.002448 | 0.01338917 |
| 26.8611111 | 0.02055499 | 0.00930551 | 0.01247868 | 0.01179875 | 0.00671977 | 0.05973497 | 0.00128916 | 0.00361409 | 0.01025637 | 0.01913906 | 0.00909988 | 0.02772386 | 0.02406039 | 0.01154883 | 0.02676329 | 0.03510522 | 0.02938091 | 0.00462452 | 0.02466915 |
| 26.875 | 0.01470739 | 0.01392913 | 0.00549682 | 0.01808747 | 0.00401822 | 0.06592165 | 0.00291659 | 0.00168667 | 0.00923585 | 0.01390105 | 0.01042687 | 0.02803504 | 0.0336933 | 0.01581721 | 0.02800455 | 0.03028738 | 0.0239853 | 0.00824607 | 0.03527642 |
| 26.8888889 | 0.01560848 | 0.01226623 | 0.00464064 | 0.02909165 | 0.00643208 | 0.0626319 | 0.00851277 | 0.0008864 | 0.01354656 | 0.0222243 | 0.01884502 | 0.02228143 | 0.03199222 | 0.0187242 | 0.0220129 | 0.02473627 | 0.01384531 | 0.00772706 | 0.03823571 |
| 26.9027778 | 0.01471191 | 0.00671371 | 0.005305 | 0.03216913 | 0.01664829 | 0.04535845 | 0.01869167 | 0.0011955 | 0.02319535 | 0.03740151 | 0.02642744 | 0.01691625 | 0.0238121 | 0.01972729 | 0.0126393 | 0.02219968 | 0.00854486 | 0.0041111 | 0.03314779 |
| 26.9166667 | 0.01212215 | 0.00447752 | 0.00711219 | 0.02829889 | 0.02298007 | 0.02686204 | 0.0290382 | 0.00292978 | 0.02910434 | 0.04689629 | 0.02566674 | 0.0143703 | 0.01734122 | 0.02050724 | 0.00536741 | 0.02198605 | 0.01142277 | 0.00289763 | 0.02432677 |
| 26.9305556 | 0.01170708 | 0.00459491 | 0.01272552 | 0.02410437 | 0.01744378 | 0.017042 | 0.03407226 | 0.00536536 | 0.02755556 | 0.04354243 | 0.01778974 | 0.01262729 | 0.013846 | 0.02135041 | 0.00203918 | 0.02110783 | 0.01861986 | 0.00588902 | 0.01767717 |
| 26.9444444 | 0.01581056 | 0.00381922 | 0.01560383 | 0.02471869 | 0.00884529 | 0.01052999 | 0.03056504 | 0.00731548 | 0.02277059 | 0.02959228 | 0.01114101 | 0.00944361 | 0.01286728 | 0.01982406 | 0.00177239 | 0.0171553 | 0.02344555 | 0.01254466 | 0.01500656 |
| 26.9583333 | 0.02083768 | 0.00449766 | 0.0118137 | 0.03034417 | 0.00675261 | 0.00525569 | 0.02010004 | 0.00950824 | 0.0188647 | 0.01669545 | 0.01127649 | 0.00554984 | 0.01323052 | 0.01543384 | 0.00295685 | 0.0111807 | 0.02307035 | 0.01831136 | 0.014756 |
| 26.9722222 | 0.01920852 | 0.00522827 | 0.00675881 | 0.03303267 | 0.01210234 | 0.00432274 | 0.01011949 | 0.01227457 | 0.01462121 | 0.01269009 | 0.01568212 | 0.00246219 | 0.01101254 | 0.01024502 | 0.00418584 | 0.00642733 | 0.01741419 | 0.01866736 | 0.01449568 |
| 26.9861111 | 0.01139743 | 0.00397056 | 0.00676601 | 0.02569435 | 0.01793135 | 0.00589151 | 0.00716051 | 0.0139389 | 0.00891003 | 0.0159327 | 0.01786406 | 0.00115779 | 0.00627104 | 0.00645511 | 0.00441906 | 0.00416266 | 0.00905297 | 0.01385123 | 0.01290346 |
| 27 | 0.0080369 | 0.00454266 | 0.00812944 | 0.01303363 | 0.01932565 | 0.00647915 | 0.00922419 | 0.01346085 | 0.00795481 | 0.02147192 | 0.01444597 | 0.00115499 | 0.00402782 | 0.00479443 | 0.00321727 | 0.00275 | 0.00431295 | 0.00712871 | 0.01130412 |
| 27.0138889 | 0.01179453 | 0.00774821 | 0.0055891 | 0.00654623 | 0.01795206 | 0.00668115 | 0.01028036 | 0.01117609 | 0.01486244 | 0.02376384 | 0.00791754 | 0.00096162 | 0.00341 | 0.00428916 | 0.00173285 | 0.00154281 | 0.00460327 | 0.00400602 | 0.0109544 |
| 27.0277778 | 0.0132075 | 0.00862964 | 0.00375436 | 0.00718969 | 0.01643051 | 0.0068203 | 0.00805465 | 0.00859585 | 0.01862208 | 0.0206837 | 0.00340983 | 0.00213151 | 0.00768199 | 0.00517806 | 0.00591587 | 0.00398362 | 0.00613602 | 0.01017778 | 0.01465531 |
| 27.0416667 | 0.01142643 | 0.00791703 | 0.00885497 | 0.01009902 | 0.01864854 | 0.00695502 | 0.01146527 | 0.01001316 | 0.01681549 | 0.02008131 | 0.00777389 | 0.00995481 | 0.03050629 | 0.01012655 | 0.02769282 | 0.01915587 | 0.01310134 | 0.02702315 | 0.02781525 |
| 27.0555556 | 0.01884605 | 0.01477407 | 0.02105318 | 0.02168526 | 0.02289776 | 0.01146206 | 0.02795419 | 0.01646707 | 0.02704311 | 0.03241385 | 0.02130404 | 0.02241839 | 0.06591746 | 0.01712302 | 0.05568905 | 0.04244735 | 0.02592033 | 0.0440334 | 0.04361109 |
| 27.0694444 | 0.03535119 | 0.02836408 | 0.03250685 | 0.04026164 | 0.01998873 | 0.01882452 | 0.04138875 | 0.02021355 | 0.04910301 | 0.04366592 | 0.02796979 | 0.02431053 | 0.08157953 | 0.01728423 | 0.05401341 | 0.04496611 | 0.03118858 | 0.03888068 | 0.04061894 |
| 27.0833333 | 0.04385629 | 0.03601338 | 0.03442512 | 0.04699029 | 0.01084274 | 0.01989147 | 0.03648236 | 0.01509778 | 0.0621389 | 0.03278207 | 0.01935451 | 0.01408492 | 0.06055807 | 0.01063672 | 0.0286779 | 0.02686408 | 0.02123427 | 0.01631983 | 0.01953032 |
| 27.0972222 | 0.03792838 | 0.03397278 | 0.02745679 | 0.03512108 | 0.00419944 | 0.0142405 | 0.0216888 | 0.00787228 | 0.0570278 | 0.02622371 | 0.01387632 | 0.00957476 | 0.02681558 | 0.0075167 | 0.01661615 | 0.0200869 | 0.00933485 | 0.00542391 | 0.00539453 |
| 27.1111111 | 0.02403221 | 0.02293512 | 0.01671344 | 0.01905384 | 0.00388343 | 0.00821883 | 0.01051238 | 0.00605726 | 0.04357215 | 0.03624281 | 0.0192729 | 0.01071718 | 0.01324747 | 0.00707508 | 0.02029906 | 0.02544499 | 0.00814028 | 0.0063133 | 0.00401268 |
| 27.125 | 0.01178272 | 0.01564239 | 0.00899708 | 0.00853712 | 0.00963197 | 0.00869485 | 0.00566909 | 0.00560553 | 0.03994404 | 0.02741315 | 0.01812258 | 0.00728791 | 0.01944011 | 0.00442532 | 0.02474474 | 0.02590081 | 0.0099433 | 0.01257767 | 0.00835967 |
| 27.1388889 | 0.01231564 | 0.02959484 | 0.00975685 | 0.00412362 | 0.02066235 | 0.01984535 | 0.0071098 | 0.0082057 | 0.03942467 | 0.01310013 | 0.01402057 | 0.00299762 | 0.0252601 | 0.0029158 | 0.02502418 | 0.0208152 | 0.00883119 | 0.02143912 | 0.01257104 |
| 27.1527778 | 0.0257202 | 0.05269031 | 0.0153425 | 0.00531477 | 0.03326755 | 0.02667165 | 0.01399283 | 0.02138316 | 0.02695256 | 0.01603421 | 0.01443514 | 0.0015988 | 0.02441452 | 0.00232637 | 0.0209726 | 0.01430405 | 0.00905694 | 0.01899709 | 0.01030812 |

| | | | | | | | | | | | | | | | | | | | |
|---|---|---|---|---|---|---|---|---|---|---|---|---|---|---|---|---|---|---|---|
| 27.1666667 | 0.02845943 | 0.05116169 | 0.03095174 | 0.01167754 | 0.03543881 | 0.04036833 | 0.01739876 | 0.03424426 | 0.01184656 | 0.01633946 | 0.00969829 | 0.00151442 | 0.02190241 | 0.00168242 | 0.01579553 | 0.00832831 | 0.01534153 | 0.00899158 | 0.00888343 |
| 27.1805556 | 0.02438776 | 0.03895818 | 0.038743 | 0.02128228 | 0.02517753 | 0.06604453 | 0.01295325 | 0.04253363 | 0.00917218 | 0.00988646 | 0.00566457 | 0.00245063 | 0.02262903 | 0.00360114 | 0.01478598 | 0.00632316 | 0.02720573 | 0.00496554 | 0.00923567 |
| 27.1944444 | 0.03156963 | 0.04329447 | 0.01765869 | 0.02711439 | 0.01595755 | 0.06100938 | 0.00918738 | 0.04323504 | 0.02653654 | 0.00812261 | 0.01047949 | 0.00575987 | 0.02651813 | 0.0083736 | 0.0188346 | 0.00954409 | 0.03641753 | 0.00781787 | 0.00808705 |
| 27.2083333 | 0.02626717 | 0.03141609 | 0.00534921 | 0.02911691 | 0.01721028 | 0.04580857 | 0.01514941 | 0.02761392 | 0.05022214 | 0.01097006 | 0.01505723 | 0.00893703 | 0.02544491 | 0.01184196 | 0.01933648 | 0.01228288 | 0.03560347 | 0.00890283 | 0.00999143 |
| 27.2222222 | 0.01295177 | 0.00968129 | 0.01049643 | 0.02778242 | 0.02445969 | 0.04206927 | 0.02308662 | 0.01512532 | 0.05987612 | 0.0138043 | 0.01410062 | 0.00704071 | 0.01728106 | 0.00990101 | 0.01257498 | 0.00901528 | 0.02349556 | 0.00794469 | 0.00969987 |
| 27.2361111 | 0.00875413 | 0.00445678 | 0.0119497 | 0.01927909 | 0.02390587 | 0.02573156 | 0.02222383 | 0.01479739 | 0.05810898 | 0.01753808 | 0.01636028 | 0.00283299 | 0.01460253 | 0.00467573 | 0.00848712 | 0.00516368 | 0.01022474 | 0.01160173 | 0.00789949 |
| 27.25 | 0.00889505 | 0.00939977 | 0.01950866 | 0.00945719 | 0.01609365 | 0.01078784 | 0.01996234 | 0.0231659 | 0.0662478 | 0.0160285 | 0.01600085 | 0.00241438 | 0.01850695 | 0.00198211 | 0.00869635 | 0.00475956 | 0.00514308 | 0.01171397 | 0.0071383 |
| 27.2638889 | 0.01602671 | 0.02394998 | 0.03474487 | 0.00480266 | 0.01685137 | 0.00911718 | 0.02265214 | 0.03577969 | 0.0871204 | 0.01677446 | 0.01758544 | 0.00786989 | 0.01484403 | 0.00579139 | 0.0093049 | 0.00653928 | 0.00624671 | 0.01191537 | 0.00907131 |
| 27.2777778 | 0.01709601 | 0.02861975 | 0.04373681 | 0.00898528 | 0.01893198 | 0.01112373 | 0.025685 | 0.0354745 | 0.08883411 | 0.0307145 | 0.03558048 | 0.01589119 | 0.00852989 | 0.01250569 | 0.01502461 | 0.01326373 | 0.01247123 | 0.0203541 | 0.01972057 |
| 27.2916667 | 0.01108625 | 0.01792683 | 0.04212962 | 0.02378173 | 0.00952304 | 0.01065363 | 0.02567112 | 0.02252218 | 0.0624689 | 0.03240139 | 0.04312233 | 0.01739803 | 0.00796471 | 0.01308768 | 0.01999847 | 0.01829355 | 0.01576103 | 0.01958853 | 0.02478602 |
| 27.3055556 | 0.01521444 | 0.01515379 | 0.03515635 | 0.03337173 | 0.00392445 | 0.00707516 | 0.02111002 | 0.01008321 | 0.033887 | 0.01883327 | 0.02532783 | 0.01084544 | 0.00700464 | 0.00745702 | 0.0143627 | 0.01315656 | 0.01182815 | 0.00983447 | 0.01478538 |
| 27.3194444 | 0.02028401 | 0.01873923 | 0.0225375 | 0.02345267 | 0.00819743 | 0.00359623 | 0.01288685 | 0.00327731 | 0.01527456 | 0.01545114 | 0.00863309 | 0.00452325 | 0.00799104 | 0.00449568 | 0.00634593 | 0.00615282 | 0.01272471 | 0.00750716 | 0.00458578 |
| 27.3333333 | 0.01449494 | 0.01368485 | 0.00954514 | 0.01071324 | 0.01424311 | 0.00282133 | 0.00930091 | 0.00124961 | 0.00816513 | 0.01444109 | 0.00446907 | 0.00225388 | 0.01247646 | 0.00627144 | 0.00573579 | 0.00653214 | 0.01911384 | 0.00692944 | 0.00146934 |
| 27.3472222 | 0.00648518 | 0.00670401 | 0.00473721 | 0.01004647 | 0.01647592 | 0.00370842 | 0.011183 | 0.00113956 | 0.01020552 | 0.01147221 | 0.00671503 | 0.00222345 | 0.0143886 | 0.00679644 | 0.00827081 | 0.00897104 | 0.01879593 | 0.00346695 | 0.00073116 |
| 27.3611111 | 0.00299401 | 0.00697951 | 0.00508227 | 0.01100207 | 0.01358167 | 0.00304701 | 0.01010044 | 0.00209306 | 0.00944849 | 0.01994255 | 0.01441667 | 0.00266039 | 0.01262707 | 0.00415105 | 0.0080239 | 0.00634344 | 0.01156743 | 0.00238967 | 0.00087573 |
| 27.375 | 0.00405426 | 0.00777505 | 0.00969624 | 0.01478181 | 0.00870533 | 0.00333094 | 0.01120556 | 0.00517197 | 0.00906678 | 0.02735838 | 0.02011266 | 0.00266468 | 0.00955838 | 0.00254763 | 0.00528557 | 0.00349461 | 0.00771072 | 0.00241189 | 0.00221392 |
| 27.3888889 | 0.01055875 | 0.00871116 | 0.01704099 | 0.02540632 | 0.0053157 | 0.00768562 | 0.01856081 | 0.0083634 | 0.01839247 | 0.02030357 | 0.0176084 | 0.00231331 | 0.00589242 | 0.00368302 | 0.00270166 | 0.00465095 | 0.00781027 | 0.00215067 | 0.00275806 |
| 27.4027778 | 0.01642665 | 0.01235573 | 0.01932121 | 0.02943063 | 0.00694087 | 0.01065529 | 0.02156141 | 0.0087137 | 0.02748039 | 0.00875965 | 0.01250101 | 0.00268893 | 0.00357111 | 0.00497079 | 0.00251371 | 0.00512865 | 0.0060761 | 0.002404 | 0.00212755 |
| 27.4166667 | 0.01435234 | 0.01098366 | 0.01520999 | 0.02207297 | 0.01001618 | 0.00822843 | 0.01601946 | 0.00636079 | 0.02607371 | 0.00316872 | 0.01034119 | 0.00441755 | 0.00605177 | 0.00489429 | 0.00409979 | 0.00348747 | 0.00449593 | 0.00273563 | 0.00228406 |
| 27.4305556 | 0.00744487 | 0.00572632 | 0.00976854 | 0.011156 | 0.01259697 | 0.00382194 | 0.00794235 | 0.00368472 | 0.01811728 | 0.0075869 | 0.0120265 | 0.00716304 | 0.01072388 | 0.00405887 | 0.00480815 | 0.00220658 | 0.00724193 | 0.00426855 | 0.00238957 |
| 27.4444444 | 0.00219075 | 0.00370747 | 0.00680946 | 0.00545449 | 0.01658847 | 0.00178019 | 0.00394574 | 0.00212296 | 0.01059688 | 0.02092085 | 0.01484241 | 0.00959321 | 0.01364537 | 0.00366402 | 0.00419172 | 0.00178502 | 0.01306663 | 0.0052756 | 0.00195017 |
| 27.4583333 | 0.00178804 | 0.00406787 | 0.00669091 | 0.00519576 | 0.01984223 | 0.00295348 | 0.00430789 | 0.00127739 | 0.00609516 | 0.02986861 | 0.01425128 | 0.0095542 | 0.01406768 | 0.00444655 | 0.00444188 | 0.00192003 | 0.01705285 | 0.00458962 | 0.00291136 |
| 27.4722222 | 0.00510344 | 0.00306776 | 0.00826449 | 0.0054903 | 0.02056255 | 0.00440931 | 0.00378718 | 0.00102746 | 0.00628744 | 0.02343805 | 0.00913799 | 0.00689669 | 0.01295654 | 0.00634682 | 0.00690671 | 0.00313654 | 0.01716316 | 0.00632014 | 0.00376145 |
| 27.4861111 | 0.00855269 | 0.00163616 | 0.00989898 | 0.0081386 | 0.01928731 | 0.00364314 | 0.00348868 | 0.00171042 | 0.01205186 | 0.01075744 | 0.00463966 | 0.00423193 | 0.01340665 | 0.00841281 | 0.01055286 | 0.00528087 | 0.01614956 | 0.00740811 | 0.00419611 |
| 27.5 | 0.00914281 | 0.00155252 | 0.01033851 | 0.0121402 | 0.01618232 | 0.00182665 | 0.00630593 | 0.00273038 | 0.02046376 | 0.00550294 | 0.00495427 | 0.00368675 | 0.01661133 | 0.00842153 | 0.0129716 | 0.0071273 | 0.01577415 | 0.00637777 | 0.00786122 |
| 27.5138889 | 0.00778646 | 0.00150177 | 0.01079713 | 0.01231498 | 0.01258459 | 0.00178527 | 0.00868666 | 0.00376642 | 0.02562268 | 0.00539933 | 0.00733738 | 0.0052358 | 0.01866647 | 0.00595666 | 0.01181923 | 0.00704739 | 0.01546814 | 0.0088635 | 0.01187164 |
| 27.5277778 | 0.00890669 | 0.00112444 | 0.013804 | 0.01114338 | 0.01125259 | 0.0049579 | 0.01034235 | 0.00670815 | 0.02430929 | 0.00705501 | 0.00805001 | 0.00761103 | 0.01824842 | 0.00431274 | 0.00890441 | 0.00585487 | 0.01567844 | 0.01341105 | 0.01149807 |
| 27.5416667 | 0.01624654 | 0.00165274 | 0.02157095 | 0.01886138 | 0.0141125 | 0.01533609 | 0.01985542 | 0.01492446 | 0.02299257 | 0.01814104 | 0.00962515 | 0.0128334 | 0.02281793 | 0.007282 | 0.01248313 | 0.00985685 | 0.01935098 | 0.02372802 | 0.01293789 |
| 27.5555556 | 0.03052426 | 0.00254744 | 0.03511479 | 0.04190333 | 0.02060113 | 0.03762229 | 0.0438439 | 0.02664128 | 0.02937319 | 0.03749058 | 0.01620809 | 0.02719695 | 0.03563182 | 0.01476873 | 0.02429802 | 0.02031596 | 0.02712006 | 0.04118646 | 0.02304199 |
| 27.5694444 | 0.04568569 | 0.00424706 | 0.05143614 | 0.0727471 | 0.02343706 | 0.06304529 | 0.07578573 | 0.03141234 | 0.04112654 | 0.05351418 | 0.02679423 | 0.03856351 | 0.05004544 | 0.02184765 | 0.03224951 | 0.02741032 | 0.03358243 | 0.04853325 | 0.03404039 |
| 27.5833333 | 0.05231623 | 0.00783307 | 0.06286961 | 0.09205085 | 0.01854463 | 0.07315835 | 0.09754963 | 0.02608792 | 0.04812305 | 0.05638423 | 0.03333039 | 0.0291927 | 0.06176924 | 0.02433698 | 0.0285064 | 0.024913 | 0.03523173 | 0.04060394 | 0.03481509 |
| 27.5972222 | 0.04769184 | 0.0103091 | 0.06134049 | 0.08170425 | 0.01163065 | 0.06286771 | 0.09147221 | 0.01796317 | 0.04063947 | 0.04220389 | 0.0253697 | 0.01289916 | 0.06453141 | 0.02228446 | 0.01869368 | 0.01819413 | 0.03205545 | 0.02969279 | 0.02811511 |
| 27.6111111 | 0.03461817 | 0.00996517 | 0.04589703 | 0.04643676 | 0.00657256 | 0.04329667 | 0.05972241 | 0.00973954 | 0.02161843 | 0.02252638 | 0.01640901 | 0.00636732 | 0.04951142 | 0.01695935 | 0.00911659 | 0.01304932 | 0.02402417 | 0.02021732 | 0.02097273 |
| 27.625 | 0.01850304 | 0.0083801 | 0.02366603 | 0.0207437 | 0.0063068 | 0.0251515 | 0.02677183 | 0.00358995 | 0.01034483 | 0.02113018 | 0.03042573 | 0.00797298 | 0.0239271 | 0.00954666 | 0.00523418 | 0.01048352 | 0.01415387 | 0.01159131 | 0.01551157 |
| 27.6388889 | 0.0076733 | 0.00788648 | 0.01045368 | 0.02663374 | 0.01036208 | 0.01161242 | 0.01447005 | 0.00171792 | 0.01719031 | 0.03651052 | 0.04659046 | 0.01114345 | 0.00785364 | 0.00342785 | 0.0095565 | 0.0095587 | 0.0069551 | 0.00528859 | 0.01280393 |
| 27.6527778 | 0.00614514 | 0.00875493 | 0.01444904 | 0.04301015 | 0.01312491 | 0.00513401 | 0.02076004 | 0.00176752 | 0.02825588 | 0.03528729 | 0.03443171 | 0.01109987 | 0.00708876 | 0.00137023 | 0.01402864 | 0.0078084 | 0.00679673 | 0.00221477 | 0.01278128 |
| 27.6666667 | 0.0099487 | 0.00958542 | 0.01967081 | 0.04704188 | 0.01368965 | 0.00830634 | 0.03040983 | 0.00279121 | 0.0291957 | 0.02435901 | 0.02213007 | 0.00980492 | 0.00977667 | 0.00307679 | 0.01254996 | 0.00576543 | 0.01118243 | 0.00157982 | 0.01427844 |
| 27.6805556 | 0.01304797 | 0.01095087 | 0.01551594 | 0.03754866 | 0.01395242 | 0.01955779 | 0.03678317 | 0.00453458 | 0.01846411 | 0.038423 | 0.03573539 | 0.0096442 | 0.01022181 | 0.00823467 | 0.00699946 | 0.00665316 | 0.00987201 | 0.0019256 | 0.01486894 |
| 27.6944444 | 0.01124908 | 0.0103734 | 0.00841953 | 0.02028582 | 0.01211386 | 0.03714498 | 0.0392454 | 0.0049636 | 0.01057369 | 0.05618358 | 0.04155661 | 0.0090766 | 0.01081508 | 0.01431292 | 0.00243241 | 0.00872378 | 0.00709148 | 0.00233782 | 0.01287537 |
| 27.7083333 | 0.00605401 | 0.00863429 | 0.00372001 | 0.00898028 | 0.00866446 | 0.05624889 | 0.03314897 | 0.00324543 | 0.0157441 | 0.04406804 | 0.0254739 | 0.00661801 | 0.01237852 | 0.01805817 | 0.00132765 | 0.00692935 | 0.01206836 | 0.00191907 | 0.00989776 |
| 27.7222222 | 0.00341625 | 0.01130192 | 0.00357288 | 0.01076509 | 0.01113178 | 0.06531802 | 0.01993347 | 0.00215687 | 0.01804354 | 0.0226407 | 0.01875941 | 0.00336818 | 0.01393432 | 0.01854576 | 0.00130937 | 0.00408957 | 0.01586596 | 0.00166801 | 0.00806462 |
| 27.7361111 | 0.00835656 | 0.01346993 | 0.00757024 | 0.014031 | 0.0181443 | 0.05644427 | 0.01715681 | 0.00415919 | 0.01348658 | 0.02454464 | 0.02811035 | 0.00179037 | 0.01324346 | 0.01402015 | 0.00117866 | 0.0061795 | 0.0130778 | 0.00251822 | 0.00802356 |
| 27.75 | 0.0201665 | 0.00898619 | 0.01208068 | 0.01449684 | 0.02293437 | 0.03511665 | 0.03286615 | 0.00715036 | 0.0156429 | 0.04111921 | 0.03132653 | 0.00213497 | 0.00865481 | 0.0078355 | 0.00125672 | 0.01043556 | 0.00836652 | 0.00325286 | 0.00924146 |
| 27.7638889 | 0.02803471 | 0.00351634 | 0.01204257 | 0.01296339 | 0.02542888 | 0.0168122 | 0.05027415 | 0.00790737 | 0.01897406 | 0.04498879 | 0.02055935 | 0.00285968 | 0.00527857 | 0.00810142 | 0.00178696 | 0.01204734 | 0.00488692 | 0.00463957 | 0.00930504 |
| 27.7777778 | 0.02054152 | 0.00194252 | 0.00714722 | 0.01184241 | 0.01947903 | 0.01497819 | 0.05414549 | 0.00563468 | 0.01312428 | 0.02877868 | 0.00927515 | 0.00504897 | 0.00998121 | 0.013724 | 0.00210758 | 0.00969431 | 0.00711717 | 0.00817616 | 0.00612713 |
| 27.7916667 | 0.01061033 | 0.00270683 | 0.00349493 | 0.0211443 | 0.01229306 | 0.02887438 | 0.04180247 | 0.00368045 | 0.00704456 | 0.01351747 | 0.00745035 | 0.00974115 | 0.01775335 | 0.01717645 | 0.00202492 | 0.00512743 | 0.01396832 | 0.01148312 | 0.00235585 |
| 27.8055556 | 0.01486772 | 0.00560539 | 0.00309173 | 0.03400381 | 0.01941016 | 0.04294321 | 0.02105389 | 0.00460021 | 0.00868468 | 0.014256 | 0.00708235 | 0.01348715 | 0.01935485 | 0.01604031 | 0.00166928 | 0.00172219 | 0.01716772 | 0.01192032 | 0.00177125 |
| 27.8194444 | 0.01982434 | 0.00968166 | 0.00449228 | 0.03684833 | 0.03193373 | 0.04566092 | 0.01167319 | 0.00521416 | 0.00874537 | 0.01351355 | 0.00672129 | 0.01239697 | 0.01369027 | 0.01263162 | 0.00145215 | 0.00043739 | 0.01337397 | 0.01041087 | 0.00445143 |
| 27.8333333 | 0.01398541 | 0.01111924 | 0.00513334 | 0.02907461 | 0.03696722 | 0.03805385 | 0.02423339 | 0.00336784 | 0.00532828 | 0.01209979 | 0.0164656 | 0.00700398 | 0.01292085 | 0.01047299 | 0.00271387 | 0.00045685 | 0.00711993 | 0.00762705 | 0.0090605 |
| 27.8472222 | 0.00984763 | 0.00767308 | 0.00715869 | 0.01657584 | 0.03259625 | 0.02223322 | 0.04655784 | 0.00125746 | 0.00904699 | 0.02797949 | 0.03122646 | 0.00205631 | 0.02805002 | 0.01270545 | 0.00466803 | 0.00168282 | 0.00468432 | 0.00399857 | 0.01488163 |
| 27.8611111 | 0.01393354 | 0.00353475 | 0.01614094 | 0.0062 | 0.02237173 | 0.01228355 | 0.05747609 | 0.00062631 | 0.01793457 | 0.04622979 | 0.03737441 | 0.00224689 | 0.04745595 | 0.01938477 | 0.00629607 | 0.00463201 | 0.0096363 | 0.00154276 | 0.02161575 |
| 27.875 | 0.01513594 | 0.00285397 | 0.02455783 | 0.00391009 | 0.0124603 | 0.02534111 | 0.04591865 | 0.00147508 | 0.02078144 | 0.0447014 | 0.02974944 | 0.00760974 | 0.05367766 | 0.02590896 | 0.00745624 | 0.00776422 | 0.02004706 | 0.00174008 | 0.02751447 |
| 27.8888889 | 0.01776688 | 0.00562919 | 0.03105075 | 0.01382212 | 0.01106361 | 0.05032935 | 0.02668024 | 0.00325666 | 0.01506609 | 0.02609058 | 0.01471669 | 0.01250047 | 0.04340162 | 0.02709086 | 0.00679649 | 0.00822798 | 0.02970983 | 0.00506021 | 0.02970954 |
| 27.9027778 | 0.0300494 | 0.01462266 | 0.04228879 | 0.03056714 | 0.01820824 | 0.06462996 | 0.02163075 | 0.0034776 | 0.00836683 | 0.00935095 | 0.00397151 | 0.01119944 | 0.02433432 | 0.02357872 | 0.00415408 | 0.00618391 | 0.03430028 | 0.01000011 | 0.02750611 |
| 27.9166667 | 0.03676794 | 0.0272584 | 0.04997297 | 0.04153763 | 0.02371002 | 0.05850566 | 0.03057524 | 0.00319666 | 0.0066131 | 0.00505331 | 0.003394 | 0.00563019 | 0.00859379 | 0.01933711 | 0.00163332 | 0.00466341 | 0.0325448 | 0.01489789 | 0.0224865 |
| 27.9305556 | 0.0333153 | 0.0352771 | 0.0435133 | 0.04475329 | 0.02174035 | 0.04226597 | 0.0408381 | 0.00630397 | 0.00880523 | 0.01109966 | 0.01095686 | 0.00193396 | 0.00230217 | 0.01590922 | 0.00093127 | 0.00543522 | 0.02448053 | 0.01944306 | 0.01726986 |
| 27.9444444 | 0.02996973 | 0.03448874 | 0.02482858 | 0.04634311 | 0.01326093 | 0.03036984 | 0.04177372 | 0.01140757 | 0.01454481 | 0.02684293 | 0.02000987 | 0.0025583 | 0.00298252 | 0.01271161 | 0.00241517 | 0.00796417 | 0.0134848 | 0.02231447 | 0.01424086 |
| 27.9583333 | 0.02478827 | 0.02440783 | 0.01173696 | 0.04362545 | 0.00699828 | 0.02187197 | 0.03007864 | 0.01603801 | 0.01963879 | 0.04066739 | 0.02292748 | 0.00534909 | 0.00627337 | 0.00824176 | 0.00536143 | 0.01080395 | 0.0052393 | 0.02287793 | 0.0132396 |
| 27.9722222 | 0.01435998 | 0.01302146 | 0.01771149 | 0.0311948 | 0.01011073 | 0.0131427 | 0.0148567 | 0.0179312 | 0.01578976 | 0.0360224 | 0.01646065 | 0.00624751 | 0.00823037 | 0.00451443 | 0.00784448 | 0.01147843 | 0.0024988 | 0.02272348 | 0.01181351 |
| 27.9861111 | 0.00835462 | 0.01304368 | 0.027993 | 0.01684243 | 0.01740938 | 0.00603819 | 0.01042493 | 0.01435974 | 0.00860302 | 0.01987838 | 0.00723703 | 0.00419087 | 0.00787074 | 0.005047 | 0.00814323 | 0.00819945 | 0.00350546 | 0.02277505 | 0.00854912 |
| 28 | 0.01241988 | 0.02166318 | 0.0280365 | 0.01539403 | 0.02205962 | 0.00381752 | 0.02007663 | 0.00763936 | 0.00970446 | 0.01136522 | 0.00294532 | 0.00295994 | 0.00723955 | 0.00573184 | 0.0066066 | 0.00398505 | 0.00533519 | 0.02271348 | 0.0046935 |
| 28.0138889 | 0.01700777 | 0.02555793 | 0.02223338 | 0.02516039 | 0.02134631 | 0.00659653 | 0.03289008 | 0.00365769 | 0.01937191 | 0.00867283 | 0.00384592 | 0.00439827 | 0.0075039 | 0.00296973 | 0.00582398 | 0.00220822 | 0.00596775 | 0.02150036 | 0.00212977 |
| 28.0277778 | 0.01382335 | 0.01891375 | 0.01371612 | 0.02615155 | 0.01797219 | 0.00881726 | 0.03648037 | 0.00519405 | 0.02857286 | 0.00794251 | 0.00824228 | 0.00446145 | 0.01111337 | 0.00257575 | 0.01263383 | 0.0076053 | 0.00608757 | 0.0196255 | 0.00346313 |
| 28.0416667 | 0.0132343 | 0.01501218 | 0.01457011 | 0.02419896 | 0.02055345 | 0.00824093 | 0.02840093 | 0.01193346 | 0.02663868 | 0.02301821 | 0.01527483 | 0.00922033 | 0.02792834 | 0.01064194 | 0.03784542 | 0.03207283 | 0.01092519 | 0.02371181 | 0.01287799 |
| 28.0555556 | 0.03390738 | 0.03625732 | 0.04447139 | 0.04841033 | 0.02865485 | 0.01189711 | 0.02903788 | 0.02079602 | 0.03072806 | 0.05667351 | 0.02746195 | 0.02883921 | 0.05327443 | 0.02458529 | 0.07327751 | 0.06778897 | 0.02440396 | 0.03376116 | 0.02876143 |
| 28.0694444 | 0.06694954 | 0.07533061 | 0.09176905 | 0.08398845 | 0.02842569 | 0.01996785 | 0.04469397 | 0.02592971 | 0.05454702 | 0.08233541 | 0.0424493 | 0.04074753 | 0.05640421 | 0.0310768 | 0.07941061 | 0.07806381 | 0.03844307 | 0.03276842 | 0.03349087 |
| 28.0833333 | 0.07976831 | 0.10077501 | 0.11336497 | 0.08762738 | 0.01571287 | 0.02051103 | 0.04501782 | 0.02278718 | 0.06717064 | 0.06845976 | 0.04237394 | 0.02718534 | 0.03397952 | 0.02143759 | 0.04690799 | 0.04944945 | 0.03723888 | 0.0173514 | 0.02218002 |
| 28.0972222 | 0.06182613 | 0.09301316 | 0.08614567 | 0.05695179 | 0.00618852 | 0.01145781 | 0.02595797 | 0.012435 | 0.06050674 | 0.03343409 | 0.02590706 | 0.01495007 | 0.0206585 | 0.00920401 | 0.02174016 | 0.01929817 | 0.02089102 | 0.00803468 | 0.01647251 |
| 28.1111111 | 0.03212715 | 0.05755498 | 0.03938862 | 0.02561433 | 0.00805714 | 0.00383723 | 0.01020682 | 0.00567257 | 0.05278322 | 0.01986054 | 0.02078182 | 0.01630777 | 0.02525749 | 0.00749474 | 0.02091938 | 0.0155531 | 0.00793528 | 0.01096087 | 0.01782485 |
| 28.125 | 0.01293902 | 0.02596498 | 0.01764822 | 0.01751329 | 0.01118983 | 0.00228613 | 0.00347827 | 0.00689004 | 0.04106052 | 0.02546158 | 0.03007882 | 0.01564189 | 0.02803291 | 0.00996522 | 0.02074416 | 0.02105817 | 0.00733992 | 0.01461883 | 0.01277168 |
| 28.1388889 | 0.01321886 | 0.02404226 | 0.02927118 | 0.02861493 | 0.00927338 | 0.00245824 | 0.00183036 | 0.00669243 | 0.02271909 | 0.02806411 | 0.03311678 | 0.00931917 | 0.02392187 | 0.00971335 | 0.01617292 | 0.02099735 | 0.01122319 | 0.01386905 | 0.00547525 |
| 28.1527778 | 0.02406432 | 0.04290548 | 0.04946758 | 0.04137353 | 0.0063366 | 0.00290583 | 0.00287841 | 0.00336397 | 0.00957105 | 0.02640131 | 0.02927159 | 0.00423594 | 0.01690612 | 0.00721579 | 0.01259382 | 0.01791476 | 0.01214716 | 0.00960129 | 0.00289373 |
| 28.1666667 | 0.03414862 | 0.06470408 | 0.05978178 | 0.04843892 | 0.00653791 | 0.00302911 | 0.00324378 | 0.00177425 | 0.0081471 | 0.02779432 | 0.02841418 | 0.00350634 | 0.00898253 | 0.00461904 | 0.00933542 | 0.0151618 | 0.01027584 | 0.00516347 | 0.002768 |
| 28.1805556 | 0.04052425 | 0.07875377 | 0.05639489 | 0.04973449 | 0.00840703 | 0.00338707 | 0.00418664 | 0.00177897 | 0.00975141 | 0.03214343 | 0.03095187 | 0.00445653 | 0.00439287 | 0.00403179 | 0.00653187 | 0.0140078 | 0.0098393 | 0.00615867 | 0.00315098 |
| 28.1944444 | 0.04190664 | 0.07748899 | 0.04462851 | 0.04343762 | 0.00836528 | 0.00706828 | 0.01045452 | 0.00181609 | 0.00926031 | 0.03402683 | 0.0298874 | 0.00799492 | 0.0027533 | 0.00549685 | 0.00501948 | 0.01301812 | 0.01422689 | 0.01265036 | 0.00659494 |
| 28.2083333 | 0.03691984 | 0.06307109 | 0.03473876 | 0.03165797 | 0.00664169 | 0.01219913 | 0.01829422 | 0.00218131 | 0.01424639 | 0.03097838 | 0.02387175 | 0.01366996 | 0.0022623 | 0.00663544 | 0.00534505 | 0.01196189 | 0.02034338 | 0.01408418 | 0.01047748 |
| 28.2222222 | 0.02823849 | 0.04531505 | 0.02895329 | 0.02012434 | 0.0051439 | 0.0149255 | 0.02223927 | 0.00415795 | 0.02936206 | 0.02494697 | 0.02000597 | 0.01294613 | 0.00480825 | 0.00576784 | 0.00627849 | 0.01128167 | 0.02125583 | 0.0108951 | 0.00894882 |
| 28.2361111 | 0.021468 | 0.0317571 | 0.02482212 | 0.01246307 | 0.00353853 | 0.01393678 | 0.02174618 | 0.0064453 | 0.04434667 | 0.01733765 | 0.01697717 | 0.00816904 | 0.00690528 | 0.0046245 | 0.00562521 | 0.00895039 | 0.01606667 | 0.01284944 | 0.00640061 |
| 28.25 | 0.01795552 | 0.0218215 | 0.02266739 | 0.00875192 | 0.00169305 | 0.01131946 | 0.01951196 | 0.00672808 | 0.0463482 | 0.00866031 | 0.01057679 | 0.00830088 | 0.00592828 | 0.005088 | 0.00348332 | 0.00527825 | 0.0090347 | 0.01184322 | 0.00637508 |
| 28.2638889 | 0.01587517 | 0.01347159 | 0.02429187 | 0.00820846 | 0.00194474 | 0.00998765 | 0.01931072 | 0.00494582 | 0.03860983 | 0.0029257 | 0.00712843 | 0.00840254 | 0.0072688 | 0.00603407 | 0.00256334 | 0.0041249 | 0.00491125 | 0.00763647 | 0.00581198 |
| 28.2777778 | 0.01484569 | 0.00820738 | 0.02802397 | 0.00987815 | 0.00452188 | 0.00990021 | 0.0210512 | 0.00249511 | 0.03234715 | 0.00210975 | 0.00791151 | 0.00833117 | 0.01039343 | 0.00526985 | 0.00322306 | 0.00424061 | 0.00697968 | 0.0102019 | 0.00945032 |
| 28.2916667 | 0.01503578 | 0.00701768 | 0.0302145 | 0.01081727 | 0.00680414 | 0.00964135 | 0.02153818 | 0.00105583 | 0.03062128 | 0.00207795 | 0.00736044 | 0.0131454 | 0.00952753 | 0.00356598 | 0.00304872 | 0.00286353 | 0.01091102 | 0.01116989 | 0.01546252 |
| 28.3055556 | 0.0160837 | 0.00935849 | 0.02872493 | 0.0084622 | 0.00713855 | 0.00848823 | 0.01896844 | 0.00158187 | 0.02994502 | 0.00121962 | 0.00915088 | 0.01353447 | 0.00549823 | 0.0042261 | 0.00264644 | 0.00209395 | 0.00954984 | 0.00940633 | 0.01384865 |
| 28.3194444 | 0.01703692 | 0.01302693 | 0.02313041 | 0.00423187 | 0.00599907 | 0.00662461 | 0.01442724 | 0.00396887 | 0.02480205 | 0.00156645 | 0.01452857 | 0.00769161 | 0.00273093 | 0.00619386 | 0.00396476 | 0.00230858 | 0.00723086 | 0.0137168 | 0.01028248 |
| 28.3333333 | 0.01551376 | 0.01302066 | 0.01532401 | 0.0012588 | 0.00498179 | 0.00475678 | 0.01029986 | 0.00567841 | 0.01488753 | 0.00393296 | 0.01304225 | 0.00548071 | 0.00191754 | 0.00510595 | 0.00438184 | 0.00206474 | 0.01013707 | 0.01529489 | 0.01054136 |
| 28.3472222 | 0.00968001 | 0.00906026 | 0.00803319 | 0.00117736 | 0.00447441 | 0.00314247 | 0.00755097 | 0.0042755 | 0.01183795 | 0.00577146 | 0.00595044 | 0.00517796 | 0.00173228 | 0.00327302 | 0.00282216 | 0.00265056 | 0.01238748 | 0.00915288 | 0.00767387 |
| 28.3611111 | 0.00372114 | 0.00818607 | 0.00330165 | 0.00469685 | 0.00401144 | 0.00178385 | 0.00531548 | 0.00308743 | 0.02010985 | 0.00502296 | 0.00240884 | 0.00459394 | 0.00139149 | 0.00391719 | 0.00268809 | 0.00379911 | 0.00938925 | 0.00418261 | 0.00341587 |
| 28.375 | 0.00254263 | 0.00903465 | 0.00155272 | 0.01208415 | 0.00416029 | 0.0012413 | 0.00386739 | 0.00520032 | 0.02506281 | 0.00723828 | 0.00409193 | 0.00892038 | 0.00157338 | 0.00363166 | 0.00355162 | 0.00313926 | 0.00491371 | 0.00209502 | 0.00346784 |
| 28.3888889 | 0.0057714 | 0.00712523 | 0.002675 | 0.02128111 | 0.00530214 | 0.0010504 | 0.00496786 | 0.00625028 | 0.01754585 | 0.01659195 | 0.01190435 | 0.01393934 | 0.00406918 | 0.00213081 | 0.00338613 | 0.00289034 | 0.00314575 | 0.0008702 | 0.00388442 |
| 28.4027778 | 0.0129002 | 0.00936583 | 0.00912731 | 0.02801333 | 0.00642186 | 0.00121198 | 0.00900552 | 0.00471515 | 0.01127509 | 0.02509021 | 0.02078341 | 0.01304809 | 0.00801376 | 0.00230891 | 0.00462846 | 0.00618297 | 0.00326619 | 0.00084223 | 0.00298827 |
| 28.4166667 | 0.02149358 | 0.01883427 | 0.02026875 | 0.02767034 | 0.0064103 | 0.00252326 | 0.01326578 | 0.00492455 | 0.01549946 | 0.02460866 | 0.02202047 | 0.00840937 | 0.01073452 | 0.00340795 | 0.00810608 | 0.0098467 | 0.00290245 | 0.00097286 | 0.00421995 |
| 28.4305556 | 0.02599547 | 0.02688825 | 0.02949532 | 0.01890081 | 0.0049954 | 0.00345078 | 0.01359281 | 0.00516256 | 0.01499658 | 0.01735001 | 0.0145016 | 0.00458007 | 0.01030034 | 0.00358569 | 0.00995913 | 0.01048078 | 0.00312739 | 0.00129836 | 0.00635032 |
| 28.4444444 | 0.0212869 | 0.02340866 | 0.02919956 | 0.00980326 | 0.00272283 | 0.00288946 | 0.00869233 | 0.00418502 | 0.01078996 | 0.00998607 | 0.00609845 | 0.00440033 | 0.0069877 | 0.00272696 | 0.00862409 | 0.00848748 | 0.00515929 | 0.00193667 | 0.00683184 |
| 28.4583333 | 0.01297418 | 0.01505543 | 0.01770749 | 0.01051088 | 0.00139731 | 0.00204918 | 0.00454538 | 0.00666407 | 0.01980599 | 0.00700601 | 0.0027748 | 0.00757514 | 0.00333587 | 0.00157309 | 0.00622323 | 0.00538665 | 0.00843331 | 0.00338269 | 0.0067902 |
| 28.4722222 | 0.01377593 | 0.01850563 | 0.00953978 | 0.0156743 | 0.0018119 | 0.00183752 | 0.00539628 | 0.01066609 | 0.03249064 | 0.00970527 | 0.00493037 | 0.01147911 | 0.00175911 | 0.0009886 | 0.00473225 | 0.00276643 | 0.01251773 | 0.0065306 | 0.00680701 |
| 28.4861111 | 0.01878508 | 0.02733736 | 0.01437096 | 0.01282809 | 0.00206197 | 0.00211848 | 0.00519659 | 0.01097394 | 0.03147696 | 0.01603146 | 0.01433325 | 0.01317438 | 0.00313015 | 0.00153631 | 0.0046153 | 0.00185346 | 0.01528326 | 0.00686255 | 0.00584261 |
| 28.5 | 0.01528335 | 0.02632399 | 0.01633155 | 0.00833417 | 0.00158491 | 0.00273092 | 0.00306869 | 0.00696646 | 0.01835414 | 0.02037722 | 0.02532119 | 0.0105219 | 0.00581401 | 0.00218089 | 0.00518268 | 0.00277298 | 0.01341408 | 0.00423093 | 0.00386168 |
| 28.5138889 | 0.00807053 | 0.01599984 | 0.01216465 | 0.01453586 | 0.00171533 | 0.00337922 | 0.00406541 | 0.00326052 | 0.00782001 | 0.01779524 | 0.02644165 | 0.00546419 | 0.0073378 | 0.00195309 | 0.00625046 | 0.00458619 | 0.00842368 | 0.0040789 | 0.00296447 |
| 28.5277778 | 0.00855766 | 0.00809601 | 0.01933045 | 0.02608953 | 0.00335941 | 0.00448058 | 0.00853136 | 0.00558396 | 0.00763156 | 0.01052601 | 0.01613622 | 0.0027765 | 0.0082819 | 0.00289655 | 0.00985699 | 0.00789698 | 0.00693342 | 0.00568678 | 0.00678099 |
| 28.5416667 | 0.01750428 | 0.01184426 | 0.03774128 | 0.04093393 | 0.00993314 | 0.00865823 | 0.0209703 | 0.01465929 | 0.01736903 | 0.00957045 | 0.00701929 | 0.0098473 | 0.01231293 | 0.00926574 | 0.01920272 | 0.01549167 | 0.01757951 | 0.00826406 | 0.01553487 |
| 28.5555556 | 0.02741846 | 0.02112711 | 0.05555668 | 0.06471947 | 0.02483656 | 0.01595838 | 0.0424695 | 0.02747177 | 0.03861417 | 0.02076491 | 0.00708855 | 0.02852871 | 0.02109193 | 0.02077834 | 0.03034955 | 0.02472644 | 0.03444846 | 0.01355631 | 0.02374728 |
| 28.5694444 | 0.03061086 | 0.02769992 | 0.07122884 | 0.09739984 | 0.04086971 | 0.02188286 | 0.06215839 | 0.03599251 | 0.06344638 | 0.03683135 | 0.01637872 | 0.04087894 | 0.03019539 | 0.02843405 | 0.03222709 | 0.02686187 | 0.03771052 | 0.01761243 | 0.02898723 |
| 28.5833333 | 0.03044977 | 0.03343348 | 0.08506787 | 0.12868028 | 0.04952785 | 0.02525009 | 0.07231364 | 0.03387787 | 0.07930971 | 0.05080384 | 0.03399259 | 0.03330392 | 0.03509626 | 0.02688568 | 0.02400827 | 0.02195534 | 0.02832854 | 0.01432619 | 0.03099 |
| 28.5972222 | 0.03311222 | 0.03738325 | 0.08466386 | 0.12786263 | 0.05062276 | 0.02688297 | 0.07054381 | 0.02430328 | 0.07463864 | 0.05488604 | 0.04123124 | 0.0170684 | 0.03512432 | 0.02208836 | 0.01515569 | 0.01717517 | 0.0202989 | 0.00709964 | 0.02711329 |
| 28.6111111 | 0.03310802 | 0.03210062 | 0.06423159 | 0.08591 | 0.04139851 | 0.02440738 | 0.05789831 | 0.01308544 | 0.04621587 | 0.04198204 | 0.02836406 | 0.00688957 | 0.02906179 | 0.01743889 | 0.00998501 | 0.01266931 | 0.01364652 | 0.00386923 | 0.01824353 |
| 28.625 | 0.02795108 | 0.01890268 | 0.03766814 | 0.03746218 | 0.02284606 | 0.01930365 | 0.04212847 | 0.00748463 | 0.02004814 | 0.02266968 | 0.02468894 | 0.00426793 | 0.01776635 | 0.01024713 | 0.00713474 | 0.00709118 | 0.00766463 | 0.00513961 | 0.00945958 |
| 28.6388889 | 0.02410728 | 0.00799294 | 0.01961168 | 0.01757299 | 0.00898782 | 0.01749675 | 0.02851414 | 0.01025021 | 0.01549032 | 0.01800506 | 0.04694896 | 0.00394145 | 0.00875574 | 0.00511905 | 0.00454771 | 0.00420927 | 0.00718905 | 0.00735097 | 0.00535749 |
| 28.6527778 | 0.02347141 | 0.00617864 | 0.01956626 | 0.02839929 | 0.00859154 | 0.02056668 | 0.01879693 | 0.01120818 | 0.01724344 | 0.02955409 | 0.06360767 | 0.00523496 | 0.00751428 | 0.00671815 | 0.00193906 | 0.00337315 | 0.00947251 | 0.0088409 | 0.00385854 |
| 28.6666667 | 0.02250404 | 0.01158302 | 0.02936679 | 0.03799196 | 0.01476289 | 0.02422694 | 0.01514298 | 0.00578778 | 0.01620518 | 0.03450473 | 0.0479081 | 0.00748633 | 0.00837993 | 0.00833001 | 0.00175915 | 0.00308069 | 0.00819476 | 0.01013175 | 0.00535276 |

| | | | | | | | | | | | | | | | | | | | |
|---|---|---|---|---|---|---|---|---|---|---|---|---|---|---|---|---|---|---|---|
| 28.6805556 | 0.02098142 | 0.01672165 | 0.02697581 | 0.02956113 | 0.02077321 | 0.02351575 | 0.01443218 | 0.00130383 | 0.01721537 | 0.02245483 | 0.02586602 | 0.00625202 | 0.00540508 | 0.00561196 | 0.00458029 | 0.00609941 | 0.00428688 | 0.01185027 | 0.01312534 |
| 28.6944444 | 0.01901457 | 0.01725412 | 0.01480402 | 0.01557752 | 0.02518734 | 0.01650025 | 0.01102514 | 0.00032097 | 0.01922823 | 0.01533133 | 0.03125504 | 0.00409833 | 0.00213037 | 0.00333471 | 0.0072962 | 0.00951069 | 0.0025325 | 0.01287215 | 0.01790249 |
| 28.7083333 | 0.01270779 | 0.01450378 | 0.01251558 | 0.00727189 | 0.0260192 | 0.01010382 | 0.00792435 | 0.00070585 | 0.02220583 | 0.02890793 | 0.04358427 | 0.00570403 | 0.00168086 | 0.00492112 | 0.00783255 | 0.00975677 | 0.00297016 | 0.01306834 | 0.01244577 |
| 28.7222222 | 0.00824003 | 0.01117002 | 0.01869324 | 0.00978607 | 0.0211095 | 0.01546346 | 0.01509909 | 0.00199376 | 0.02782188 | 0.04235795 | 0.03164896 | 0.00744537 | 0.00283432 | 0.00706574 | 0.00683796 | 0.00744476 | 0.00225626 | 0.01384738 | 0.00551925 |
| 28.7361111 | 0.01794046 | 0.00876799 | 0.0203909 | 0.02214866 | 0.0116025 | 0.02971336 | 0.03269598 | 0.00322245 | 0.0309105 | 0.03626616 | 0.01555535 | 0.00650336 | 0.00489668 | 0.00779585 | 0.00580823 | 0.0036902 | 0.00179741 | 0.01395532 | 0.00423399 |
| 28.75 | 0.03251401 | 0.00735854 | 0.01478545 | 0.03217766 | 0.00644762 | 0.03915146 | 0.04648827 | 0.0028247 | 0.02526829 | 0.01954002 | 0.02018641 | 0.0044141 | 0.00543684 | 0.00815337 | 0.00531246 | 0.00112695 | 0.00415881 | 0.01193088 | 0.00403244 |
| 28.7638889 | 0.03298362 | 0.00649523 | 0.0089073 | 0.02459642 | 0.01391573 | 0.03425553 | 0.04337424 | 0.00217893 | 0.01334402 | 0.01285089 | 0.02981921 | 0.00258011 | 0.00350181 | 0.00637189 | 0.0054321 | 0.00150734 | 0.00746846 | 0.00947602 | 0.00377614 |
| 28.7777778 | 0.02094495 | 0.00634975 | 0.00970386 | 0.01190377 | 0.02915805 | 0.01805822 | 0.02486384 | 0.00327749 | 0.00428116 | 0.02122375 | 0.02901931 | 0.0028961 | 0.00191079 | 0.00447027 | 0.00640253 | 0.00330613 | 0.00801144 | 0.00813504 | 0.00635121 |
| 28.7916667 | 0.00933298 | 0.00710429 | 0.01493207 | 0.0147141 | 0.03934394 | 0.00633337 | 0.01159891 | 0.00457234 | 0.00699865 | 0.02967536 | 0.0237059 | 0.00568385 | 0.00288252 | 0.00509912 | 0.00704617 | 0.00389779 | 0.00671742 | 0.00805297 | 0.00837836 |
| 28.8055556 | 0.00541789 | 0.00862316 | 0.02688293 | 0.01961687 | 0.03815211 | 0.0058116 | 0.01815869 | 0.00533897 | 0.01743899 | 0.0300918 | 0.02143753 | 0.0081587 | 0.00588059 | 0.00409447 | 0.00603213 | 0.0026014 | 0.0099032 | 0.00986236 | 0.00616735 |
| 28.8194444 | 0.00936924 | 0.01050659 | 0.0463307 | 0.01281169 | 0.02853494 | 0.00677723 | 0.03117648 | 0.00745137 | 0.02194111 | 0.02522214 | 0.02319614 | 0.00827406 | 0.00794087 | 0.00134879 | 0.00382803 | 0.00193296 | 0.01713056 | 0.01469369 | 0.00493096 |
| 28.8333333 | 0.0144481 | 0.01347162 | 0.04826319 | 0.00748705 | 0.01627283 | 0.00455816 | 0.0364339 | 0.00960423 | 0.01779725 | 0.01983213 | 0.0257273 | 0.00725473 | 0.0077726 | 0.00196236 | 0.00301154 | 0.00386229 | 0.02180638 | 0.02113084 | 0.01087755 |
| 28.8472222 | 0.01666334 | 0.01875324 | 0.02796156 | 0.01361776 | 0.01354697 | 0.00353153 | 0.03294797 | 0.00792878 | 0.01144238 | 0.0188382 | 0.0282183 | 0.00776533 | 0.008244 | 0.0063616 | 0.00490302 | 0.0061576 | 0.02278138 | 0.0245159 | 0.01896884 |
| 28.8611111 | 0.01626809 | 0.02375551 | 0.01970447 | 0.02115887 | 0.02580819 | 0.00527985 | 0.02255806 | 0.00467027 | 0.00928549 | 0.02377957 | 0.03213475 | 0.01183824 | 0.01246141 | 0.01074613 | 0.00677065 | 0.00665312 | 0.02277257 | 0.02194896 | 0.02408505 |
| 28.875 | 0.01201853 | 0.02435639 | 0.02845977 | 0.022638 | 0.03735313 | 0.00739843 | 0.01233087 | 0.00588439 | 0.01381946 | 0.03125988 | 0.03756584 | 0.01909264 | 0.01785424 | 0.01344577 | 0.00659112 | 0.00609991 | 0.0215707 | 0.01612496 | 0.02652572 |
| 28.8888889 | 0.01239043 | 0.02239739 | 0.03495265 | 0.02575774 | 0.03807435 | 0.00893445 | 0.01661493 | 0.00971826 | 0.02403375 | 0.03454426 | 0.04163376 | 0.02623224 | 0.01862248 | 0.01740689 | 0.00547823 | 0.00624234 | 0.01844063 | 0.01246396 | 0.0256409 |
| 28.9027778 | 0.02757633 | 0.02394696 | 0.02789332 | 0.03709344 | 0.03593931 | 0.01097618 | 0.04297394 | 0.01009842 | 0.03773386 | 0.02945453 | 0.04047188 | 0.0278288 | 0.0134878 | 0.02389673 | 0.00543116 | 0.00785623 | 0.01673594 | 0.0131483 | 0.02242776 |
| 28.9166667 | 0.04796051 | 0.02942496 | 0.02115045 | 0.0508887 | 0.03397794 | 0.0135055 | 0.07555815 | 0.00702883 | 0.05053979 | 0.01858501 | 0.0352179 | 0.02022897 | 0.00633607 | 0.02696173 | 0.00648946 | 0.00966523 | 0.01822264 | 0.01578754 | 0.0190317 |
| 28.9305556 | 0.05554025 | 0.03308898 | 0.04142656 | 0.0563795 | 0.02730486 | 0.01456806 | 0.08290782 | 0.00905837 | 0.05956598 | 0.00805753 | 0.02886437 | 0.0097625 | 0.00184003 | 0.02213482 | 0.00618703 | 0.00902834 | 0.01862521 | 0.01605454 | 0.01535034 |
| 28.9444444 | 0.04565076 | 0.03132345 | 0.06998778 | 0.05005305 | 0.01814905 | 0.01387745 | 0.05814241 | 0.01910345 | 0.06169262 | 0.0026817 | 0.02093158 | 0.00550639 | 0.00145159 | 0.01246 | 0.00355694 | 0.00522928 | 0.01502196 | 0.01215296 | 0.01005459 |
| 28.9583333 | 0.02630038 | 0.02452979 | 0.07218473 | 0.03864761 | 0.01233058 | 0.01271964 | 0.02681385 | 0.02710639 | 0.0536544 | 0.00304532 | 0.01218408 | 0.00443109 | 0.00443314 | 0.00487439 | 0.00135097 | 0.00148667 | 0.01086215 | 0.00727434 | 0.00485255 |
| 28.9722222 | 0.01208006 | 0.01468224 | 0.04611593 | 0.02648479 | 0.00979659 | 0.00986613 | 0.00935816 | 0.02775558 | 0.03732034 | 0.0057577 | 0.00666759 | 0.00364304 | 0.01048301 | 0.00360769 | 0.00160376 | 0.00059275 | 0.00976077 | 0.005629 | 0.00313395 |
| 28.9861111 | 0.01199897 | 0.00652928 | 0.02242035 | 0.01469018 | 0.00676043 | 0.00509046 | 0.00732661 | 0.02268368 | 0.01964195 | 0.00787065 | 0.00642103 | 0.00597954 | 0.0158108 | 0.00404291 | 0.00256413 | 0.00143246 | 0.0100275 | 0.00933975 | 0.00353699 |
| 29 | 0.01699179 | 0.00383059 | 0.03031491 | 0.01150395 | 0.00625843 | 0.00272971 | 0.01171565 | 0.01375299 | 0.01426687 | 0.00938548 | 0.00851501 | 0.00625824 | 0.01491475 | 0.00352643 | 0.00268538 | 0.00176451 | 0.00821349 | 0.01710806 | 0.00340148 |
| 29.0138889 | 0.01588712 | 0.00327425 | 0.05749371 | 0.01705435 | 0.01193559 | 0.00506465 | 0.01538041 | 0.00629598 | 0.02844309 | 0.01114609 | 0.00922325 | 0.00400207 | 0.00887869 | 0.00492153 | 0.00198702 | 0.00138835 | 0.00464826 | 0.02444794 | 0.00297595 |
| 29.0277778 | 0.01180785 | 0.00512292 | 0.0738197 | 0.01582818 | 0.01386759 | 0.00673978 | 0.01502135 | 0.00448306 | 0.04338626 | 0.01511982 | 0.01043615 | 0.0045725 | 0.00405103 | 0.00891554 | 0.00494751 | 0.00465027 | 0.00232934 | 0.03229796 | 0.00394795 |
| 29.0416667 | 0.02129142 | 0.02185256 | 0.05679363 | 0.01903208 | 0.01340477 | 0.00671351 | 0.01287014 | 0.01440721 | 0.03943846 | 0.03036 | 0.02001289 | 0.0149173 | 0.01308175 | 0.01738143 | 0.02091777 | 0.01986986 | 0.00816358 | 0.05769582 | 0.01385059 |
| 29.0555556 | 0.05411651 | 0.06354613 | 0.04203131 | 0.0535287 | 0.02716884 | 0.01156466 | 0.01949752 | 0.04356417 | 0.04457377 | 0.05639873 | 0.03867392 | 0.04231648 | 0.03884814 | 0.02995611 | 0.04470091 | 0.04299113 | 0.02627945 | 0.10768797 | 0.03409592 |
| 29.0694444 | 0.08568373 | 0.11309477 | 0.08170806 | 0.09249283 | 0.04518028 | 0.0187768 | 0.03060104 | 0.06836669 | 0.07796484 | 0.06454779 | 0.05005693 | 0.06454832 | 0.05230676 | 0.03682508 | 0.04884316 | 0.04857865 | 0.0385461 | 0.13633762 | 0.04210376 |
| 29.0833333 | 0.08727297 | 0.13614888 | 0.12506647 | 0.09496949 | 0.04300394 | 0.02182598 | 0.03026435 | 0.06189327 | 0.09612272 | 0.04002475 | 0.0407548 | 0.05178485 | 0.03643656 | 0.02808539 | 0.02845888 | 0.02953511 | 0.02797988 | 0.10429048 | 0.02552063 |
| 29.0972222 | 0.06586469 | 0.12172997 | 0.12251997 | 0.0681537 | 0.02356402 | 0.01979009 | 0.02064711 | 0.04086101 | 0.08483602 | 0.01328093 | 0.02052095 | 0.02299183 | 0.02189655 | 0.01344263 | 0.01291117 | 0.01377714 | 0.01134316 | 0.04746978 | 0.00877047 |
| 29.1111111 | 0.03624381 | 0.0822935 | 0.1000335 | 0.036652 | 0.00919323 | 0.0122259 | 0.0118915 | 0.02397873 | 0.07070414 | 0.00531241 | 0.00909032 | 0.01312711 | 0.02717968 | 0.00864549 | 0.01059395 | 0.01310686 | 0.00591181 | 0.01436065 | 0.00801953 |
| 29.125 | 0.01672384 | 0.03959411 | 0.07467455 | 0.01540432 | 0.00980434 | 0.00530651 | 0.0058723 | 0.01109274 | 0.05729792 | 0.00390753 | 0.01239051 | 0.01824328 | 0.03316958 | 0.01077522 | 0.0096905 | 0.01450847 | 0.00557005 | 0.01046522 | 0.01296502 |
| 29.1388889 | 0.02298314 | 0.02060744 | 0.04667372 | 0.01210728 | 0.01576073 | 0.00515516 | 0.00460083 | 0.00454418 | 0.03950032 | 0.00236839 | 0.01888154 | 0.02418084 | 0.03253099 | 0.01264252 | 0.00758483 | 0.01154164 | 0.00412612 | 0.01701187 | 0.01582489 |
| 29.1527778 | 0.03899663 | 0.02768072 | 0.02262399 | 0.02103971 | 0.01939934 | 0.00537707 | 0.00900204 | 0.00855578 | 0.02159977 | 0.00187197 | 0.01802663 | 0.02727235 | 0.02796538 | 0.01301766 | 0.00467311 | 0.00666117 | 0.00232249 | 0.01950025 | 0.01386934 |
| 29.1666667 | 0.05078953 | 0.04029656 | 0.01538408 | 0.02855211 | 0.02214548 | 0.00251177 | 0.0123587 | 0.01954229 | 0.00870356 | 0.00204129 | 0.01278358 | 0.02412787 | 0.01871633 | 0.01186722 | 0.00272615 | 0.00452615 | 0.00112262 | 0.01512712 | 0.00716486 |
| 29.1805556 | 0.05767642 | 0.04599022 | 0.02907573 | 0.02736184 | 0.0233882 | 0.00174318 | 0.01015412 | 0.03128012 | 0.00335405 | 0.00317102 | 0.01521068 | 0.01730805 | 0.0111441 | 0.01074623 | 0.00276642 | 0.00435212 | 0.00144072 | 0.01049398 | 0.00479396 |
| 29.1944444 | 0.0568558 | 0.04293908 | 0.05616474 | 0.02285742 | 0.02082588 | 0.00427337 | 0.00843309 | 0.04056125 | 0.00486022 | 0.00553882 | 0.0310025 | 0.01243146 | 0.00991324 | 0.01123401 | 0.00460089 | 0.0047628 | 0.00339584 | 0.01207974 | 0.01247456 |
| 29.2083333 | 0.04931023 | 0.0382022 | 0.087694 | 0.02214136 | 0.01608884 | 0.0071466 | 0.0130952 | 0.04257456 | 0.00813145 | 0.01001326 | 0.04659279 | 0.01061374 | 0.01148241 | 0.01235257 | 0.0076999 | 0.00651453 | 0.00497619 | 0.01166043 | 0.02020364 |
| 29.2222222 | 0.03791318 | 0.04084282 | 0.1096805 | 0.02093984 | 0.01143761 | 0.00839634 | 0.02138373 | 0.03763947 | 0.01149615 | 0.01488134 | 0.04654937 | 0.00982061 | 0.01054021 | 0.01150164 | 0.00795253 | 0.00562962 | 0.00380816 | 0.0095504 | 0.01666323 |
| 29.2361111 | 0.02655488 | 0.05441433 | 0.11330058 | 0.01567474 | 0.00660557 | 0.00811446 | 0.02807545 | 0.02972763 | 0.01601934 | 0.01635711 | 0.03424817 | 0.00849761 | 0.00657166 | 0.00732007 | 0.00497035 | 0.00246818 | 0.00212272 | 0.01641649 | 0.00905832 |
| 29.25 | 0.01872229 | 0.07348531 | 0.10436033 | 0.00941725 | 0.00248772 | 0.00780758 | 0.02676505 | 0.02065824 | 0.02076024 | 0.01369163 | 0.01980031 | 0.00541145 | 0.0029783 | 0.00363877 | 0.00219174 | 0.0009302 | 0.00248175 | 0.02653961 | 0.01057956 |
| 29.2638889 | 0.01470912 | 0.08182645 | 0.08974574 | 0.00430029 | 0.00080912 | 0.00825049 | 0.01719641 | 0.01229288 | 0.02422976 | 0.01013518 | 0.00876083 | 0.00324252 | 0.00289113 | 0.00417286 | 0.0009728 | 0.00071813 | 0.00298845 | 0.03286148 | 0.0177237 |
| 29.2777778 | 0.01202087 | 0.07101554 | 0.06685278 | 0.00178597 | 0.00116056 | 0.00870324 | 0.00876874 | 0.00796462 | 0.02550684 | 0.0081424 | 0.00402746 | 0.00538056 | 0.00509652 | 0.00479265 | 0.00088879 | 0.00081282 | 0.00357869 | 0.03432749 | 0.01864822 |
| 29.2916667 | 0.01070782 | 0.05212024 | 0.03893865 | 0.0017306 | 0.00136867 | 0.0095671 | 0.00510359 | 0.00765529 | 0.02779856 | 0.00836252 | 0.00298418 | 0.00664744 | 0.00553788 | 0.00380045 | 0.00168607 | 0.00107767 | 0.00415812 | 0.02813693 | 0.01176448 |
| 29.3055556 | 0.01337577 | 0.0387432 | 0.0198986 | 0.00456644 | 0.00177448 | 0.01176721 | 0.0032064 | 0.00691671 | 0.03469396 | 0.01129671 | 0.00269737 | 0.00582395 | 0.00354679 | 0.00508183 | 0.00350162 | 0.00176919 | 0.00354664 | 0.01704081 | 0.00705854 |
| 29.3194444 | 0.01782871 | 0.03190666 | 0.01221989 | 0.01239719 | 0.00371599 | 0.01349274 | 0.00288377 | 0.00680482 | 0.04194262 | 0.01437466 | 0.00295567 | 0.00830416 | 0.00226862 | 0.00770086 | 0.00459589 | 0.00288395 | 0.00348808 | 0.00895639 | 0.00725875 |
| 29.3333333 | 0.01996796 | 0.02564599 | 0.00705157 | 0.01987686 | 0.00466126 | 0.01249102 | 0.00435583 | 0.00706555 | 0.04358035 | 0.01262136 | 0.0054734 | 0.00946968 | 0.00289156 | 0.00698307 | 0.00336311 | 0.00338234 | 0.00405944 | 0.00739325 | 0.00763243 |
| 29.3472222 | 0.01928346 | 0.01792474 | 0.00252181 | 0.02077479 | 0.00361335 | 0.00931197 | 0.00581049 | 0.00426278 | 0.04006139 | 0.00711427 | 0.01210107 | 0.00639129 | 0.00405111 | 0.00351668 | 0.00136186 | 0.00279754 | 0.0043182 | 0.0081787 | 0.0116823 |
| 29.3611111 | 0.01706681 | 0.01085927 | 0.00131349 | 0.01604653 | 0.00244859 | 0.00595479 | 0.00642769 | 0.00167832 | 0.03493575 | 0.00405211 | 0.01574489 | 0.00613017 | 0.00489208 | 0.00116826 | 0.0005247 | 0.00211334 | 0.00495624 | 0.0082539 | 0.01663203 |
| 29.375 | 0.01499655 | 0.006072 | 0.00123364 | 0.00912767 | 0.00177864 | 0.00395015 | 0.00603912 | 0.00147624 | 0.03070467 | 0.00477039 | 0.01085716 | 0.00832442 | 0.00473742 | 0.00167329 | 0.00093584 | 0.00211489 | 0.00497139 | 0.00892183 | 0.01287674 |
| 29.3888889 | 0.01519036 | 0.00564643 | 0.00088385 | 0.00458268 | 0.00118082 | 0.00399782 | 0.00511657 | 0.00217626 | 0.02957325 | 0.00611928 | 0.00677017 | 0.00726704 | 0.00318375 | 0.00503584 | 0.00229458 | 0.00239925 | 0.00336247 | 0.00922638 | 0.00665103 |
| 29.4027778 | 0.01795932 | 0.0106698 | 0.001333 | 0.00512608 | 0.00204218 | 0.00468016 | 0.00436955 | 0.00423377 | 0.03103213 | 0.0068996 | 0.0108049 | 0.00711846 | 0.00253178 | 0.00831517 | 0.00329929 | 0.00211748 | 0.00226767 | 0.00705712 | 0.00769374 |
| 29.4166667 | 0.01967368 | 0.0168871 | 0.00220881 | 0.00822339 | 0.00693029 | 0.00389189 | 0.00379794 | 0.00570732 | 0.03177591 | 0.00880243 | 0.01629884 | 0.01226381 | 0.0035038 | 0.00801402 | 0.00291381 | 0.0015901 | 0.00244247 | 0.00375607 | 0.01020192 |
| 29.4305556 | 0.01608536 | 0.01867687 | 0.0042352 | 0.01083569 | 0.01349381 | 0.0020565 | 0.0045799 | 0.00462585 | 0.03077314 | 0.01338521 | 0.01854994 | 0.01717317 | 0.00450279 | 0.00496815 | 0.00221879 | 0.00196528 | 0.00457229 | 0.00196937 | 0.00907856 |
| 29.4444444 | 0.00871976 | 0.01553973 | 0.0097391 | 0.0133602 | 0.01616413 | 0.00096727 | 0.00829974 | 0.00302396 | 0.02854555 | 0.01886019 | 0.01827968 | 0.01784914 | 0.00854633 | 0.00252975 | 0.002392 | 0.00306774 | 0.01047398 | 0.00157415 | 0.00735225 |
| 29.4583333 | 0.0046157 | 0.01035366 | 0.01800238 | 0.0158705 | 0.01333409 | 0.00076433 | 0.01419123 | 0.00181519 | 0.02569678 | 0.02163406 | 0.01553512 | 0.01508032 | 0.0148933 | 0.00251231 | 0.00321978 | 0.0036823 | 0.01586215 | 0.00098387 | 0.00737177 |
| 29.4722222 | 0.00461381 | 0.00575023 | 0.02374934 | 0.01733494 | 0.00817939 | 0.00064247 | 0.01884332 | 0.0013681 | 0.02272779 | 0.02081338 | 0.01216512 | 0.01029866 | 0.01643554 | 0.00515683 | 0.00362173 | 0.00320163 | 0.01624745 | 0.00081674 | 0.00938756 |
| 29.4861111 | 0.00335906 | 0.00288464 | 0.02252618 | 0.01741908 | 0.00460511 | 0.00083497 | 0.01868225 | 0.00253045 | 0.01822417 | 0.01845679 | 0.01063077 | 0.00595747 | 0.01149043 | 0.00889251 | 0.00306163 | 0.00182988 | 0.01335625 | 0.00258078 | 0.01189444 |
| 29.5 | 0.00274873 | 0.00134885 | 0.01609043 | 0.01581955 | 0.00368381 | 0.00096025 | 0.01359131 | 0.00336896 | 0.01166989 | 0.01718315 | 0.01201405 | 0.00487457 | 0.00483314 | 0.01106618 | 0.00234636 | 0.00073342 | 0.01123384 | 0.00570653 | 0.01297348 |
| 29.5138889 | 0.00695053 | 0.00066508 | 0.01088465 | 0.01429146 | 0.00310225 | 0.00101632 | 0.00810485 | 0.00254097 | 0.00710751 | 0.01868457 | 0.0153761 | 0.00868514 | 0.00209909 | 0.01141939 | 0.00270972 | 0.0015665 | 0.01311603 | 0.00752238 | 0.01167788 |
| 29.5277778 | 0.01456739 | 0.00218938 | 0.0123325 | 0.01800344 | 0.00582294 | 0.00157877 | 0.00803038 | 0.00307397 | 0.01046897 | 0.0242556 | 0.0199332 | 0.01621589 | 0.00600176 | 0.01246621 | 0.00475393 | 0.00480681 | 0.01928957 | 0.00781393 | 0.01044227 |
| 29.5416667 | 0.0237067 | 0.00805778 | 0.02258079 | 0.03217221 | 0.02018477 | 0.00396242 | 0.01730347 | 0.0086788 | 0.02667213 | 0.03566692 | 0.02738408 | 0.02372527 | 0.01455566 | 0.01731997 | 0.0078793 | 0.00917544 | 0.02891443 | 0.01257302 | 0.01516234 |
| 29.5555556 | 0.0334496 | 0.01616098 | 0.04127766 | 0.0532113 | 0.04303761 | 0.01134329 | 0.03555951 | 0.01809655 | 0.05061062 | 0.04991029 | 0.03728445 | 0.02793934 | 0.02207365 | 0.02549914 | 0.00980164 | 0.0118874 | 0.03713967 | 0.0291386 | 0.02452719 |
| 29.5694444 | 0.04142505 | 0.02171344 | 0.06251219 | 0.06932243 | 0.05800013 | 0.02105256 | 0.05526443 | 0.02608091 | 0.06574696 | 0.05464575 | 0.04424897 | 0.03017963 | 0.02512511 | 0.03190346 | 0.00811899 | 0.01067874 | 0.03604027 | 0.05206345 | 0.02986025 |
| 29.5833333 | 0.04635481 | 0.02399483 | 0.08140438 | 0.0744085 | 0.05634152 | 0.02525594 | 0.0656806 | 0.02828504 | 0.07077967 | 0.04630129 | 0.04488222 | 0.03443623 | 0.02630674 | 0.03350905 | 0.00400513 | 0.00775544 | 0.02808408 | 0.06032148 | 0.03198242 |
| 29.5972222 | 0.04511402 | 0.02446552 | 0.08940277 | 0.06778872 | 0.0458296 | 0.02262534 | 0.06085604 | 0.02672972 | 0.07354027 | 0.03686022 | 0.03945798 | 0.03628266 | 0.02566006 | 0.02938667 | 0.00104379 | 0.00533796 | 0.01995056 | 0.04623461 | 0.03262344 |
| 29.6111111 | 0.03323385 | 0.02359857 | 0.06970311 | 0.05115514 | 0.0368477 | 0.01789647 | 0.04524491 | 0.02502175 | 0.06384357 | 0.02981325 | 0.02510491 | 0.02485879 | 0.01972263 | 0.01835064 | 0.00080607 | 0.00341586 | 0.01145835 | 0.02865702 | 0.02585253 |
| 29.625 | 0.01694791 | 0.02280899 | 0.03276495 | 0.03229363 | 0.0346406 | 0.01412284 | 0.03029625 | 0.01797572 | 0.03835193 | 0.0201687 | 0.01723742 | 0.01369503 | 0.0106369 | 0.00937435 | 0.00222404 | 0.00398084 | 0.00679305 | 0.02329884 | 0.01423352 |
| 29.6388889 | 0.0090356 | 0.02202016 | 0.0169339 | 0.0165188 | 0.0366781 | 0.01210014 | 0.02113215 | 0.0097348 | 0.0158752 | 0.00937773 | 0.03839417 | 0.02176839 | 0.00486909 | 0.01228151 | 0.00378942 | 0.00606393 | 0.00953522 | 0.02702125 | 0.00997372 |
| 29.6527778 | 0.01207253 | 0.01864915 | 0.03487948 | 0.00568211 | 0.0350424 | 0.01192264 | 0.01708898 | 0.00858736 | 0.00797446 | 0.00292393 | 0.06710988 | 0.03351287 | 0.00506825 | 0.01963109 | 0.00475041 | 0.00638362 | 0.01228229 | 0.03026235 | 0.01513991 |
| 29.6666667 | 0.0176276 | 0.01138984 | 0.05743434 | 0.0019617 | 0.02713082 | 0.01371318 | 0.01771092 | 0.00716206 | 0.00812699 | 0.00154528 | 0.0663764 | 0.03357266 | 0.00566163 | 0.02165818 | 0.00404784 | 0.00434398 | 0.01185822 | 0.02784399 | 0.01657884 |
| 29.6805556 | 0.02131504 | 0.00887745 | 0.06242812 | 0.00278712 | 0.01514642 | 0.01366886 | 0.01944067 | 0.00468158 | 0.0103987 | 0.001463 | 0.03769266 | 0.02362277 | 0.0037538 | 0.01748654 | 0.00224451 | 0.00204694 | 0.01344003 | 0.02126408 | 0.0101233 |
| 29.6944444 | 0.02203183 | 0.01959506 | 0.05269812 | 0.00551328 | 0.00823389 | 0.0131867 | 0.01808493 | 0.00557724 | 0.01298136 | 0.00332338 | 0.02021304 | 0.01190998 | 0.00292956 | 0.00960548 | 0.00116434 | 0.00146941 | 0.01876212 | 0.01436268 | 0.00542492 |
| 29.7083333 | 0.01545648 | 0.03247479 | 0.03441019 | 0.01017361 | 0.01487725 | 0.01458823 | 0.01273073 | 0.00694046 | 0.01043977 | 0.00905669 | 0.03012089 | 0.0097826 | 0.00253993 | 0.00475666 | 0.00181933 | 0.00261977 | 0.02289353 | 0.00789888 | 0.00427239 |
| 29.7222222 | 0.00995587 | 0.0362719 | 0.02738356 | 0.01440192 | 0.02692605 | 0.01014832 | 0.00669505 | 0.00709083 | 0.00519912 | 0.01444695 | 0.03628906 | 0.0146915 | 0.00197125 | 0.0067657 | 0.00395937 | 0.00501887 | 0.01856101 | 0.00359638 | 0.00386792 |
| 29.7361111 | 0.01496389 | 0.03360621 | 0.0421505 | 0.01623612 | 0.03655765 | 0.00514706 | 0.00583149 | 0.00634468 | 0.005573 | 0.01485602 | 0.02369391 | 0.01744531 | 0.00326618 | 0.00957893 | 0.00472402 | 0.00537234 | 0.01170827 | 0.00346234 | 0.00578495 |
| 29.75 | 0.01958524 | 0.029884 | 0.05609157 | 0.0170996 | 0.04384935 | 0.00964513 | 0.01086671 | 0.00495646 | 0.01076921 | 0.01126926 | 0.01556417 | 0.01607805 | 0.0035068 | 0.00955781 | 0.00415731 | 0.00383016 | 0.0155439 | 0.00367382 | 0.0058573 |
| 29.7638889 | 0.01681284 | 0.02684743 | 0.05642718 | 0.01913495 | 0.04311522 | 0.01546384 | 0.01754672 | 0.00304814 | 0.01669218 | 0.00733885 | 0.02529844 | 0.01194967 | 0.00246621 | 0.00830785 | 0.00539825 | 0.00481104 | 0.02345903 | 0.00469886 | 0.00438255 |
| 29.7777778 | 0.01089565 | 0.02200782 | 0.04872606 | 0.02226123 | 0.02837626 | 0.01144748 | 0.02431952 | 0.0012834 | 0.01837771 | 0.00450009 | 0.03481401 | 0.00652909 | 0.00394074 | 0.00559296 | 0.00657388 | 0.00649146 | 0.02264528 | 0.00975011 | 0.00645352 |
| 29.7916667 | 0.00653767 | 0.01494605 | 0.03457219 | 0.02228872 | 0.01439439 | 0.00318833 | 0.02927655 | 0.00042003 | 0.01345758 | 0.00263268 | 0.03312095 | 0.00430661 | 0.00570078 | 0.00473637 | 0.00465436 | 0.00491764 | 0.01460161 | 0.01508518 | 0.0067424 |
| 29.8055556 | 0.00936954 | 0.01038515 | 0.01849091 | 0.01636109 | 0.01997055 | 0.00045722 | 0.02895657 | 0.0012191 | 0.00981471 | 0.00334494 | 0.02915248 | 0.00825167 | 0.0050264 | 0.00801203 | 0.00292962 | 0.0034027 | 0.00652059 | 0.01759124 | 0.00610468 |
| 29.8194444 | 0.01989095 | 0.01426075 | 0.01661005 | 0.00950426 | 0.03374164 | 0.00203724 | 0.02427085 | 0.00634448 | 0.01373343 | 0.00874273 | 0.03104238 | 0.01407857 | 0.00378779 | 0.008684 | 0.00361128 | 0.00600699 | 0.00396373 | 0.0170398 | 0.01373693 |
| 29.8333333 | 0.03445116 | 0.02668035 | 0.03454269 | 0.00981331 | 0.03823177 | 0.0075253 | 0.01706632 | 0.01910236 | 0.02067329 | 0.01808682 | 0.03999712 | 0.01681677 | 0.00623997 | 0.00461957 | 0.00324925 | 0.00854187 | 0.00760429 | 0.01264076 | 0.02541867 |
| 29.8472222 | 0.04716113 | 0.04028299 | 0.05656888 | 0.01733819 | 0.03312529 | 0.01185155 | 0.00864858 | 0.03589899 | 0.02375497 | 0.02730683 | 0.05077674 | 0.01550979 | 0.01470895 | 0.0020762 | 0.0014539 | 0.00760521 | 0.01472839 | 0.00651071 | 0.02975104 |
| 29.8611111 | 0.05107594 | 0.04767327 | 0.06857192 | 0.02497765 | 0.02112725 | 0.00859687 | 0.00291054 | 0.04478864 | 0.02445685 | 0.03199256 | 0.05800692 | 0.0127139 | 0.02397051 | 0.0041479 | 0.00109452 | 0.0050568 | 0.02407105 | 0.00416104 | 0.02473751 |
| 29.875 | 0.04473531 | 0.04674325 | 0.06424093 | 0.02832702 | 0.01245598 | 0.00553299 | 0.00123071 | 0.04080958 | 0.0266348 | 0.03104921 | 0.06059791 | 0.01251902 | 0.02704526 | 0.0096027 | 0.00223102 | 0.00270485 | 0.03342683 | 0.00656664 | 0.01603299 |
| 29.8888889 | 0.0299117 | 0.03985569 | 0.04966537 | 0.02742492 | 0.01959934 | 0.00484925 | 0.00221053 | 0.03130128 | 0.02438833 | 0.02728113 | 0.05866076 | 0.01670258 | 0.02367389 | 0.01720751 | 0.00388446 | 0.00203088 | 0.03893702 | 0.00954663 | 0.00958246 |
| 29.9027778 | 0.01287989 | 0.03166067 | 0.03315567 | 0.02571822 | 0.03445931 | 0.00352268 | 0.0066015 | 0.02251415 | 0.01402381 | 0.02350699 | 0.05243763 | 0.02203771 | 0.01656399 | 0.02540048 | 0.00522583 | 0.00287895 | 0.03761511 | 0.01090151 | 0.00609759 |
| 29.9166667 | 0.00421394 | 0.02498985 | 0.01945486 | 0.02677812 | 0.04221606 | 0.00340287 | 0.01414971 | 0.01570135 | 0.00728268 | 0.0195414 | 0.04272593 | 0.02252942 | 0.00852515 | 0.03015755 | 0.00421975 | 0.00268738 | 0.02893834 | 0.01099794 | 0.00367429 |
| 29.9305556 | 0.00554298 | 0.01851243 | 0.01479428 | 0.02898678 | 0.04450742 | 0.00319242 | 0.02000554 | 0.0090515 | 0.0104369 | 0.01423996 | 0.03095098 | 0.01635581 | 0.00463646 | 0.02791023 | 0.0033077 | 0.00206076 | 0.01748344 | 0.01033317 | 0.00338357 |
| 29.9444444 | 0.00997487 | 0.01209117 | 0.01160984 | 0.02738955 | 0.04975229 | 0.00217556 | 0.01722081 | 0.00606573 | 0.01474428 | 0.00788863 | 0.0195163 | 0.008151 | 0.00584366 | 0.01949345 | 0.00587856 | 0.00430588 | 0.00938939 | 0.00825992 | 0.005024 |
| 29.9583333 | 0.01317075 | 0.0071951 | 0.01009211 | 0.0188284 | 0.04824476 | 0.00311687 | 0.0100396 | 0.01131096 | 0.01791094 | 0.00527775 | 0.0099021 | 0.00439566 | 0.00714553 | 0.00941151 | 0.00823501 | 0.0080599 | 0.00592544 | 0.00452938 | 0.00554891 |
| 29.9722222 | 0.01307373 | 0.00347312 | 0.02263441 | 0.00828878 | 0.03169253 | 0.00607096 | 0.01119277 | 0.01624135 | 0.02037278 | 0.00911949 | 0.00351745 | 0.00466476 | 0.00620754 | 0.00368952 | 0.00734244 | 0.0099607 | 0.00454937 | 0.00176298 | 0.00408252 |
| 29.9861111 | 0.00845719 | 0.00220793 | 0.04068194 | 0.00487417 | 0.01413537 | 0.00588842 | 0.01896084 | 0.01493169 | 0.01804695 | 0.01244661 | 0.00095395 | 0.00388367 | 0.00401281 | 0.00312793 | 0.00479591 | 0.00882803 | 0.00321848 | 0.00118202 | 0.00203556 |
| 30 | 0.00377148 | 0.00444971 | 0.04981782 | 0.00798646 | 0.00828991 | 0.00338971 | 0.02456025 | 0.01056454 | 0.01254907 | 0.0102674 | 0.00038435 | 0.00344316 | 0.00189674 | 0.00252027 | 0.00258529 | 0.0054881 | 0.00298911 | 0.0015022 | 0.00085651 |
| 30.0138889 | 0.00343563 | 0.00673111 | 0.04764905 | 0.01050865 | 0.01148319 | 0.00372386 | 0.0264746 | 0.0065572 | 0.00849931 | 0.00537357 | 0.00026246 | 0.00530341 | 0.00124344 | 0.00118819 | 0.00152517 | 0.00248719 | 0.00598221 | 0.00307167 | 0.00061449 |
| 30.0277778 | 0.00785791 | 0.00883651 | 0.03368452 | 0.01085435 | 0.01145459 | 0.00405945 | 0.02264884 | 0.01047686 | 0.01469553 | 0.00445596 | 0.00207629 | 0.00998054 | 0.00297833 | 0.00259752 | 0.00133565 | 0.00384101 | 0.00699455 | 0.00584567 | 0.00194917 |
| 30.0416667 | 0.02424242 | 0.02549116 | 0.0251477 | 0.02145436 | 0.01566093 | 0.00784171 | 0.017788 | 0.03148313 | 0.04746831 | 0.01563993 | 0.01196479 | 0.02622366 | 0.00966454 | 0.01259406 | 0.001853 | 0.01478 | 0.01310336 | 0.01658637 | 0.00949612 |
| 30.0555556 | 0.05709104 | 0.06444043 | 0.05311373 | 0.04891773 | 0.03865016 | 0.02030262 | 0.02827781 | 0.05712839 | 0.10048481 | 0.03729029 | 0.03003831 | 0.05649714 | 0.0236901 | 0.0347645 | 0.00517874 | 0.03306639 | 0.04019451 | 0.03744315 | 0.02409565 |
| 30.0694444 | 0.08667441 | 0.09835453 | 0.10904709 | 0.07568849 | 0.05614894 | 0.03091972 | 0.0565759 | 0.06001563 | 0.13250546 | 0.04740804 | 0.04050258 | 0.07667823 | 0.03521742 | 0.05267509 | 0.00807083 | 0.04232978 | 0.0614627 | 0.04222712 | 0.03013238 |
| 30.0833333 | 0.09024829 | 0.09860552 | 0.14356251 | 0.08162461 | 0.04895371 | 0.03273978 | 0.07975816 | 0.0391601 | 0.12126465 | 0.0314336 | 0.03171936 | 0.06205723 | 0.02893656 | 0.0446905 | 0.00621416 | 0.03161526 | 0.04670079 | 0.02773631 | 0.02139486 |
| 30.0972222 | 0.0745833 | 0.07729608 | 0.13194915 | 0.06071987 | 0.03350981 | 0.0267637 | 0.07307485 | 0.01563246 | 0.08866897 | 0.01357866 | 0.01416403 | 0.02967448 | 0.01486537 | 0.02207738 | 0.00607722 | 0.01566189 | 0.01995407 | 0.02506492 | 0.0161598 |
| 30.1111111 | 0.05718716 | 0.06044211 | 0.09475928 | 0.03083675 | 0.01984963 | 0.01628547 | 0.03998268 | 0.00339792 | 0.06212843 | 0.01170644 | 0.00396077 | 0.01552156 | 0.01385521 | 0.01216898 | 0.00924314 | 0.0129302 | 0.00980412 | 0.02951997 | 0.01672423 |
| 30.125 | 0.03808308 | 0.04560907 | 0.05229187 | 0.01485025 | 0.00859481 | 0.00768955 | 0.02035347 | 0.00216365 | 0.04713257 | 0.01073206 | 0.00312059 | 0.02337152 | 0.02192061 | 0.01678081 | 0.01140409 | 0.01945132 | 0.01261598 | 0.0228521 | 0.01264709 |
| 30.1388889 | 0.02056975 | 0.0254376 | 0.02007484 | 0.01710122 | 0.00434631 | 0.00559837 | 0.03525927 | 0.00421079 | 0.03499874 | 0.00560552 | 0.00446644 | 0.03471508 | 0.02612063 | 0.02167569 | 0.01357478 | 0.02311602 | 0.01700269 | 0.01222556 | 0.00700766 |
| 30.1527778 | 0.02233171 | 0.01758866 | 0.01044669 | 0.03054551 | 0.00829817 | 0.01015267 | 0.05509127 | 0.00402432 | 0.01962209 | 0.00320338 | 0.00445814 | 0.04186845 | 0.02394344 | 0.02231744 | 0.01533288 | 0.02152586 | 0.01903163 | 0.00571003 | 0.00418289 |
| 30.1666667 | 0.0415371 | 0.03098852 | 0.01202244 | 0.04316103 | 0.01524776 | 0.01572305 | 0.06134684 | 0.00249793 | 0.01054963 | 0.00323912 | 0.00472462 | 0.04885675 | 0.01974648 | 0.02239124 | 0.0146546 | 0.01746184 | 0.0190814 | 0.00334391 | 0.00332828 |
| 30.1805556 | 0.05562394 | 0.0459722 | 0.01452852 | 0.04690998 | 0.01873961 | 0.01781136 | 0.0602665 | 0.00225329 | 0.01681844 | 0.0049479 | 0.00750298 | 0.05966371 | 0.0178489 | 0.02380653 | 0.01206154 | 0.01243382 | 0.01875218 | 0.00311166 | 0.0029571 |

| | | | | | | | | | | | | | | | | | | | |
|---|---|---|---|---|---|---|---|---|---|---|---|---|---|---|---|---|---|---|---|
| 30.1944444 | 0.05594933 | 0.04925849 | 0.01850915 | 0.04038448 | 0.01940745 | 0.01630421 | 0.05232965 | 0.00210955 | 0.02357749 | 0.00640632 | 0.0117989 | 0.06854231 | 0.01757017 | 0.02498594 | 0.00800087 | 0.00730329 | 0.01760532 | 0.00302703 | 0.00292555 |
| 30.2083333 | 0.05634722 | 0.04532314 | 0.02366291 | 0.0272369 | 0.022324 | 0.01571351 | 0.03799304 | 0.00190458 | 0.02222745 | 0.00516996 | 0.01448778 | 0.0665151 | 0.01659681 | 0.02460146 | 0.00401327 | 0.00393694 | 0.01437903 | 0.00272484 | 0.00410343 |
| 30.2222222 | 0.06205667 | 0.03354694 | 0.03053765 | 0.01656628 | 0.0257758 | 0.02094681 | 0.02081678 | 0.00555836 | 0.02705896 | 0.00424194 | 0.01513331 | 0.05309419 | 0.01374962 | 0.02266755 | 0.00248372 | 0.00438857 | 0.01066417 | 0.00333385 | 0.00587071 |
| 30.2361111 | 0.06259436 | 0.01535472 | 0.0406509 | 0.01346925 | 0.02610246 | 0.0280725 | 0.00715465 | 0.0144155 | 0.04089551 | 0.00391159 | 0.01645138 | 0.03293829 | 0.00951142 | 0.01883002 | 0.0020304 | 0.00744487 | 0.00760789 | 0.0053366 | 0.00593842 |
| 30.25 | 0.06081443 | 0.00647583 | 0.05535805 | 0.01364173 | 0.02314725 | 0.0290432 | 0.00312575 | 0.02375371 | 0.05039311 | 0.00495853 | 0.02144481 | 0.01477149 | 0.00560251 | 0.01283553 | 0.00147989 | 0.00875542 | 0.00446087 | 0.00777951 | 0.00379628 |
| 30.2638889 | 0.06092123 | 0.01465323 | 0.07244354 | 0.01840015 | 0.01867229 | 0.02481037 | 0.00644359 | 0.03081244 | 0.05023204 | 0.01160792 | 0.02994057 | 0.00604934 | 0.00343747 | 0.00612294 | 0.00164086 | 0.00659762 | 0.00283636 | 0.00957763 | 0.00189763 |
| 30.2777778 | 0.05693747 | 0.03048584 | 0.08471235 | 0.02961242 | 0.01488516 | 0.02348553 | 0.01381096 | 0.03570038 | 0.04417528 | 0.01894263 | 0.03665471 | 0.00447186 | 0.003643 | 0.00163715 | 0.00186474 | 0.00396882 | 0.00272397 | 0.0098789 | 0.0012942 |
| 30.2916667 | 0.04868437 | 0.04373215 | 0.08634427 | 0.03630556 | 0.01163214 | 0.02516579 | 0.02368752 | 0.03708204 | 0.038713 | 0.0199472 | 0.03509231 | 0.00731326 | 0.00538855 | 0.00074217 | 0.00282753 | 0.00419289 | 0.00297765 | 0.0080412 | 0.00118815 |
| 30.3055556 | 0.0354516 | 0.05085032 | 0.07753879 | 0.03212702 | 0.00924201 | 0.02432024 | 0.03034481 | 0.03481552 | 0.03987553 | 0.01625807 | 0.02536244 | 0.01442685 | 0.00719239 | 0.00214787 | 0.00513043 | 0.00774925 | 0.0065488 | 0.00512778 | 0.0017549 |
| 30.3194444 | 0.01857129 | 0.05239697 | 0.06405925 | 0.02091289 | 0.00896728 | 0.02160689 | 0.02943521 | 0.0296234 | 0.05142392 | 0.01626655 | 0.017174 | 0.02015129 | 0.00918318 | 0.00440174 | 0.00863241 | 0.01320599 | 0.01432254 | 0.00373723 | 0.00226265 |
| 30.3333333 | 0.00779738 | 0.04879367 | 0.05516195 | 0.01062504 | 0.01140232 | 0.01976806 | 0.02600216 | 0.02330906 | 0.06979861 | 0.02552389 | 0.01755257 | 0.01852885 | 0.01108138 | 0.0054995 | 0.00732034 | 0.01809483 | 0.02244584 | 0.00497713 | 0.00369475 |
| 30.3472222 | 0.0071157 | 0.0398391 | 0.05202116 | 0.00969863 | 0.01628767 | 0.01870279 | 0.02835222 | 0.02118486 | 0.07937351 | 0.03570279 | 0.02108963 | 0.01056808 | 0.011919 | 0.00672835 | 0.00866039 | 0.01939238 | 0.024415 | 0.00698359 | 0.00796415 |
| 30.3611111 | 0.01428834 | 0.03005908 | 0.0479503 | 0.01698699 | 0.02183177 | 0.01797985 | 0.0353167 | 0.02672497 | 0.07707366 | 0.03689547 | 0.01929819 | 0.00519098 | 0.01330638 | 0.01171682 | 0.00943631 | 0.01673436 | 0.0191583 | 0.00773897 | 0.01250691 |
| 30.375 | 0.02397721 | 0.02491528 | 0.03896856 | 0.02308025 | 0.02631671 | 0.01706958 | 0.03711001 | 0.03405408 | 0.07150112 | 0.02943673 | 0.01345333 | 0.00922592 | 0.01617955 | 0.01679094 | 0.00735651 | 0.01207722 | 0.0135358 | 0.00721137 | 0.01393505 |
| 30.3888889 | 0.02713386 | 0.02309949 | 0.02369307 | 0.02244727 | 0.02805429 | 0.0131888 | 0.02714243 | 0.03192802 | 0.05301254 | 0.01756294 | 0.01279505 | 0.01835572 | 0.01771227 | 0.01667358 | 0.00378129 | 0.00638549 | 0.01040293 | 0.00596891 | 0.01111326 |
| 30.4027778 | 0.02100716 | 0.01933298 | 0.01383966 | 0.01724126 | 0.02519203 | 0.00652316 | 0.01247888 | 0.0189793 | 0.02616606 | 0.00989377 | 0.01539597 | 0.02552776 | 0.01494378 | 0.01305708 | 0.00158775 | 0.00186954 | 0.00769162 | 0.004309 | 0.00614052 |
| 30.4166667 | 0.01321816 | 0.0118416 | 0.01548985 | 0.01248167 | 0.01835362 | 0.0023142 | 0.00384627 | 0.00929835 | 0.01275868 | 0.01272835 | 0.01345466 | 0.02533817 | 0.00873979 | 0.01042165 | 0.0015208 | 0.00065318 | 0.00506832 | 0.00259916 | 0.00287988 |
| 30.4305556 | 0.01493116 | 0.00496193 | 0.0136459 | 0.01283114 | 0.01056338 | 0.00273402 | 0.00344516 | 0.00765689 | 0.01036743 | 0.02120648 | 0.01647343 | 0.01655091 | 0.00338718 | 0.01024495 | 0.00380376 | 0.00206587 | 0.00410759 | 0.00137323 | 0.00292413 |
| 30.4444444 | 0.02917413 | 0.00334448 | 0.00673047 | 0.01986973 | 0.00536363 | 0.0058934 | 0.00931536 | 0.00536601 | 0.01522389 | 0.025152 | 0.02955833 | 0.00767021 | 0.00170356 | 0.01095095 | 0.00664137 | 0.00526903 | 0.00582356 | 0.00121172 | 0.00683852 |
| 30.4583333 | 0.04725859 | 0.00725144 | 0.00487477 | 0.03057819 | 0.0041357 | 0.00917329 | 0.01626882 | 0.00309942 | 0.02881404 | 0.02039911 | 0.03884182 | 0.00520569 | 0.003374 | 0.010998 | 0.00726392 | 0.00803981 | 0.008662 | 0.00190993 | 0.01447309 |
| 30.4722222 | 0.05765132 | 0.0128414 | 0.00667825 | 0.03894348 | 0.00465947 | 0.00976983 | 0.01870115 | 0.00243269 | 0.03843319 | 0.01227356 | 0.03736904 | 0.00536027 | 0.00752293 | 0.01029257 | 0.00486134 | 0.00732247 | 0.00972895 | 0.00213751 | 0.02191899 |
| 30.4861111 | 0.05506147 | 0.01612639 | 0.00604288 | 0.04009112 | 0.00393432 | 0.00680771 | 0.01609171 | 0.00297446 | 0.03044809 | 0.00720728 | 0.02999066 | 0.00873783 | 0.01172958 | 0.00997497 | 0.00172382 | 0.00402823 | 0.00796832 | 0.00133002 | 0.02371716 |
| 30.5 | 0.0437018 | 0.01763025 | 0.00361437 | 0.03515338 | 0.00318135 | 0.00311161 | 0.01151796 | 0.00692302 | 0.01754801 | 0.00836719 | 0.02206589 | 0.01923049 | 0.01306429 | 0.01122829 | 0.00148656 | 0.00209039 | 0.00488611 | 0.00047144 | 0.01829122 |
| 30.5138889 | 0.03218473 | 0.02094155 | 0.0034322 | 0.02964393 | 0.00507968 | 0.00146789 | 0.00836998 | 0.01019679 | 0.02186122 | 0.01336442 | 0.01629874 | 0.02867501 | 0.01086485 | 0.01350525 | 0.00445105 | 0.00338247 | 0.00350009 | 0.00027629 | 0.00915824 |
| 30.5277778 | 0.02595928 | 0.02619999 | 0.00676391 | 0.02760947 | 0.00622328 | 0.00123042 | 0.00861592 | 0.00822792 | 0.03018922 | 0.01679074 | 0.01530328 | 0.0283271 | 0.00627578 | 0.01446297 | 0.00686341 | 0.00603589 | 0.00621335 | 0.00052338 | 0.00286798 |
| 30.5416667 | 0.02877643 | 0.03050968 | 0.01588672 | 0.03302395 | 0.01213847 | 0.00106786 | 0.01396887 | 0.00621227 | 0.02463735 | 0.01922148 | 0.02345463 | 0.02201403 | 0.00222722 | 0.01337788 | 0.00566646 | 0.00769608 | 0.01273811 | 0.0016599 | 0.00264327 |
| 30.5555556 | 0.04447528 | 0.03341744 | 0.03534938 | 0.04947822 | 0.03533149 | 0.00237579 | 0.027118 | 0.01318522 | 0.02351758 | 0.02352012 | 0.03823446 | 0.01930951 | 0.0023077 | 0.01319772 | 0.00382933 | 0.00919679 | 0.02195038 | 0.00599581 | 0.00938317 |
| 30.5694444 | 0.06761589 | 0.03681969 | 0.06718618 | 0.06993298 | 0.06235593 | 0.00697442 | 0.04724921 | 0.02831997 | 0.03942084 | 0.02834571 | 0.04713048 | 0.02525188 | 0.00882401 | 0.017318 | 0.00664167 | 0.01369054 | 0.031223 | 0.01470358 | 0.02451404 |
| 30.5833333 | 0.08204252 | 0.0440439 | 0.09991883 | 0.0794637 | 0.07041893 | 0.01043436 | 0.06292036 | 0.04606529 | 0.05523413 | 0.03292845 | 0.04359657 | 0.03611275 | 0.01981906 | 0.02466884 | 0.01294464 | 0.02162177 | 0.03580082 | 0.0213157 | 0.04093711 |
| 30.5972222 | 0.07943516 | 0.05138459 | 0.10903641 | 0.07004724 | 0.06335519 | 0.01155304 | 0.06118543 | 0.05662291 | 0.05987383 | 0.03612791 | 0.03159319 | 0.04221292 | 0.02759961 | 0.02928342 | 0.01632535 | 0.0269229 | 0.03392315 | 0.0185212 | 0.04536297 |
| 30.6111111 | 0.06121497 | 0.04928194 | 0.08782438 | 0.04874085 | 0.0553849 | 0.0148527 | 0.04443829 | 0.05204453 | 0.05505184 | 0.03321476 | 0.01671013 | 0.03524749 | 0.02607614 | 0.02613117 | 0.01229739 | 0.02138623 | 0.0263406 | 0.01047281 | 0.03159409 |
| 30.625 | 0.03395979 | 0.03973839 | 0.05417352 | 0.02811102 | 0.05124294 | 0.01768502 | 0.0231373 | 0.04239292 | 0.04780584 | 0.02520446 | 0.01055781 | 0.02009427 | 0.01744553 | 0.01606223 | 0.00745433 | 0.0129695 | 0.01486373 | 0.00485995 | 0.01727485 |
| 30.6388889 | 0.02043002 | 0.03097545 | 0.02431818 | 0.01254866 | 0.05101306 | 0.01785853 | 0.00998388 | 0.03464343 | 0.03936011 | 0.01778058 | 0.02035851 | 0.01482353 | 0.00829903 | 0.00692877 | 0.01020593 | 0.01631018 | 0.00691273 | 0.00336802 | 0.02251907 |
| 30.6527778 | 0.02545471 | 0.02521666 | 0.01188652 | 0.00407404 | 0.05016808 | 0.01701627 | 0.01074674 | 0.0239978 | 0.02569122 | 0.01015918 | 0.03726443 | 0.02571952 | 0.00276851 | 0.00541096 | 0.01416009 | 0.02422606 | 0.00737559 | 0.00308496 | 0.03647202 |
| 30.6666667 | 0.03022564 | 0.02037214 | 0.02222061 | 0.00230199 | 0.04380829 | 0.01412174 | 0.01184634 | 0.01248442 | 0.01973251 | 0.00346668 | 0.05069617 | 0.03498168 | 0.00126774 | 0.00736204 | 0.01238257 | 0.02415049 | 0.00925539 | 0.00206521 | 0.04443145 |
| 30.6805556 | 0.0327623 | 0.01315048 | 0.04054019 | 0.00291146 | 0.03138296 | 0.00818168 | 0.00632914 | 0.01093456 | 0.02639612 | 0.0052229 | 0.05385597 | 0.02759582 | 0.00259664 | 0.00839294 | 0.00693138 | 0.01602072 | 0.00732631 | 0.00276379 | 0.04059449 |
| 30.6944444 | 0.03299779 | 0.01091933 | 0.05051512 | 0.00558076 | 0.01679905 | 0.00514713 | 0.00203222 | 0.02108077 | 0.03179448 | 0.01420067 | 0.0456833 | 0.01468937 | 0.00521691 | 0.01055532 | 0.00272688 | 0.00626679 | 0.00452306 | 0.00660225 | 0.02379182 |
| 30.7083333 | 0.02700547 | 0.02021889 | 0.0544845 | 0.00915702 | 0.01227779 | 0.00997165 | 0.00128659 | 0.03015771 | 0.0343811 | 0.01885637 | 0.03073171 | 0.01475346 | 0.00744752 | 0.01303484 | 0.00173496 | 0.00238328 | 0.00316506 | 0.01142115 | 0.01146241 |
| 30.7222222 | 0.02551145 | 0.02860418 | 0.05875767 | 0.01015964 | 0.02302843 | 0.01525269 | 0.00215821 | 0.03233227 | 0.03502439 | 0.01286034 | 0.01513152 | 0.02351449 | 0.00827925 | 0.01243739 | 0.00253147 | 0.00544587 | 0.002723 | 0.01423187 | 0.01268423 |
| 30.7361111 | 0.0310752 | 0.0271294 | 0.04529685 | 0.00714153 | 0.03507308 | 0.01590428 | 0.00416732 | 0.03113821 | 0.02503553 | 0.00572442 | 0.00711423 | 0.02639889 | 0.00784824 | 0.00818276 | 0.00446243 | 0.00946463 | 0.00287327 | 0.01364818 | 0.01133582 |
| 30.75 | 0.0315908 | 0.01860717 | 0.02339921 | 0.00515482 | 0.03945711 | 0.01662828 | 0.00532845 | 0.03071564 | 0.01705727 | 0.0065223 | 0.00577062 | 0.02173034 | 0.0056708 | 0.0044061 | 0.00542818 | 0.00936903 | 0.00427035 | 0.01079636 | 0.0062213 |
| 30.7638889 | 0.02778368 | 0.00913597 | 0.02763599 | 0.01100736 | 0.04344556 | 0.01737582 | 0.00381561 | 0.03497163 | 0.02733411 | 0.00731097 | 0.00437764 | 0.01389616 | 0.00330954 | 0.00535275 | 0.00444586 | 0.00634144 | 0.00613797 | 0.00735781 | 0.00780167 |
| 30.7777778 | 0.02201198 | 0.00640447 | 0.03921062 | 0.02090905 | 0.04935748 | 0.01197397 | 0.002509 | 0.04097123 | 0.04404962 | 0.00633055 | 0.00417129 | 0.00822404 | 0.00390534 | 0.0081072 | 0.00339716 | 0.00386334 | 0.00683373 | 0.00449628 | 0.01016841 |
| 30.7916667 | 0.01278081 | 0.01146663 | 0.030078 | 0.02985777 | 0.04818013 | 0.00409848 | 0.00468805 | 0.03963271 | 0.04857606 | 0.0111672 | 0.00455096 | 0.01139618 | 0.00560382 | 0.00795712 | 0.00303091 | 0.00245395 | 0.00560525 | 0.00295637 | 0.00830051 |
| 30.8055556 | 0.00736011 | 0.01832533 | 0.01244435 | 0.03506341 | 0.03608244 | 0.00124296 | 0.00749628 | 0.03017048 | 0.0343317 | 0.01749094 | 0.00371264 | 0.02003223 | 0.00502733 | 0.00483624 | 0.00263911 | 0.00149595 | 0.00308818 | 0.00292898 | 0.00529762 |
| 30.8194444 | 0.00736824 | 0.02196252 | 0.00715365 | 0.03174683 | 0.0194542 | 0.00142941 | 0.00714259 | 0.02153578 | 0.0151034 | 0.01639384 | 0.00648941 | 0.02485357 | 0.00317544 | 0.0028568 | 0.00206656 | 0.00098942 | 0.00193974 | 0.00379801 | 0.00427777 |
| 30.8333333 | 0.00678327 | 0.02172171 | 0.012647 | 0.01983541 | 0.00915813 | 0.00171124 | 0.00534404 | 0.01653053 | 0.00866923 | 0.01027861 | 0.01630864 | 0.02587683 | 0.00410252 | 0.00583238 | 0.00193322 | 0.00160077 | 0.00245108 | 0.00451311 | 0.00701313 |
| 30.8472222 | 0.00556471 | 0.01859731 | 0.01651394 | 0.01092671 | 0.01147909 | 0.00299228 | 0.00726394 | 0.01201925 | 0.01865099 | 0.01021061 | 0.03140257 | 0.02689524 | 0.00989732 | 0.01235217 | 0.00271221 | 0.00496923 | 0.00250459 | 0.0047607 | 0.01045611 |
| 30.8611111 | 0.00732704 | 0.01462594 | 0.01798449 | 0.01547148 | 0.01788584 | 0.00824066 | 0.01445203 | 0.00750564 | 0.0348026 | 0.01522746 | 0.04556988 | 0.02678304 | 0.01739272 | 0.0181781 | 0.003696 | 0.00942577 | 0.00406893 | 0.00516993 | 0.00897608 |
| 30.875 | 0.01337075 | 0.01473282 | 0.02687519 | 0.02627114 | 0.01744458 | 0.01604157 | 0.02072638 | 0.00479201 | 0.04263308 | 0.01568153 | 0.05257101 | 0.02417474 | 0.02216423 | 0.01963681 | 0.00421632 | 0.01147663 | 0.00806163 | 0.0062446 | 0.00511616 |
| 30.8888889 | 0.02002047 | 0.02307287 | 0.05362137 | 0.03580206 | 0.01166454 | 0.0170587 | 0.02388165 | 0.00367219 | 0.03901135 | 0.0134859 | 0.05352026 | 0.02263754 | 0.02258011 | 0.01674791 | 0.00490718 | 0.01117178 | 0.01213855 | 0.00791314 | 0.00504237 |
| 30.9027778 | 0.02335471 | 0.03699308 | 0.08774847 | 0.04243249 | 0.01271432 | 0.00922287 | 0.02899049 | 0.00277877 | 0.03129634 | 0.01509233 | 0.05261117 | 0.02357866 | 0.01915165 | 0.01263844 | 0.00528893 | 0.00930397 | 0.0164009 | 0.01004441 | 0.00625807 |
| 30.9166667 | 0.0234668 | 0.04656335 | 0.10607058 | 0.04412076 | 0.02667422 | 0.00226157 | 0.03475908 | 0.00334388 | 0.02686821 | 0.02143606 | 0.048017 | 0.02202733 | 0.01318059 | 0.00926249 | 0.00414737 | 0.00606032 | 0.02035704 | 0.01174092 | 0.00602143 |
| 30.9305556 | 0.01885525 | 0.04421836 | 0.10349958 | 0.04193192 | 0.04287948 | 0.00186747 | 0.03673373 | 0.00668094 | 0.02715307 | 0.02996964 | 0.03821553 | 0.01587596 | 0.00693452 | 0.00669197 | 0.00400892 | 0.00558244 | 0.0220226 | 0.01147271 | 0.00631668 |
| 30.9444444 | 0.01188184 | 0.032889 | 0.08946444 | 0.0370358 | 0.05074679 | 0.0062086 | 0.03262857 | 0.01342699 | 0.02678879 | 0.03470194 | 0.02782776 | 0.00913059 | 0.00255486 | 0.0040064 | 0.00757863 | 0.00986371 | 0.02162425 | 0.00930643 | 0.00783502 |
| 30.9583333 | 0.01360995 | 0.01875986 | 0.06890753 | 0.02614099 | 0.04880406 | 0.01089274 | 0.01982943 | 0.02077584 | 0.0208229 | 0.03051179 | 0.01954827 | 0.00456939 | 0.00074297 | 0.00242323 | 0.01125413 | 0.01292713 | 0.01951543 | 0.00717089 | 0.00892806 |
| 30.9722222 | 0.03025969 | 0.00877425 | 0.04151696 | 0.01403287 | 0.03844388 | 0.01191332 | 0.01233364 | 0.02250543 | 0.01236415 | 0.01930312 | 0.01266854 | 0.00179105 | 0.00065649 | 0.0040697 | 0.01098836 | 0.01052949 | 0.01536926 | 0.00619177 | 0.00806329 |
| 30.9861111 | 0.05370063 | 0.0062055 | 0.02194971 | 0.01422448 | 0.02309804 | 0.00999901 | 0.02632327 | 0.01774539 | 0.0111034 | 0.00801278 | 0.00732796 | 0.00095873 | 0.00058328 | 0.00665813 | 0.00706912 | 0.00592428 | 0.01107038 | 0.00573694 | 0.00589077 |
| 31 | 0.07114242 | 0.0070391 | 0.02925242 | 0.02174915 | 0.01122516 | 0.0079257 | 0.04744512 | 0.01302985 | 0.01949363 | 0.00181015 | 0.00350291 | 0.00258455 | 0.00041815 | 0.00649298 | 0.0029435 | 0.00422053 | 0.00921194 | 0.00511607 | 0.00378325 |
| 31.0138889 | 0.07491006 | 0.00975833 | 0.05114245 | 0.02431636 | 0.01066407 | 0.00663098 | 0.05661014 | 0.01300627 | 0.02729038 | 0.001514 | 0.00127851 | 0.00588946 | 0.00130311 | 0.00422792 | 0.00130629 | 0.00413759 | 0.00886189 | 0.00411045 | 0.00217522 |
| 31.0277778 | 0.0605962 | 0.01093945 | 0.06244709 | 0.02271667 | 0.01302152 | 0.0084759 | 0.05048533 | 0.0150384 | 0.02575019 | 0.00554701 | 0.00284327 | 0.01032336 | 0.00517727 | 0.00658725 | 0.00168315 | 0.00545383 | 0.00954235 | 0.00427704 | 0.00239749 |
| 31.0416667 | 0.04279456 | 0.01109224 | 0.0468375 | 0.01879458 | 0.01584098 | 0.0169491 | 0.03214326 | 0.01986193 | 0.01992678 | 0.01213503 | 0.0131837 | 0.019553 | 0.01647088 | 0.02103031 | 0.00595628 | 0.01757521 | 0.01693433 | 0.01145862 | 0.00886393 |
| 31.0555556 | 0.05551321 | 0.0183867 | 0.03300611 | 0.02333863 | 0.03312516 | 0.0266839 | 0.02518466 | 0.02682355 | 0.02719698 | 0.01994352 | 0.03168442 | 0.03482182 | 0.03567183 | 0.04091213 | 0.01461929 | 0.04001808 | 0.0322174 | 0.02357168 | 0.02140542 |
| 31.0694444 | 0.08646975 | 0.02638648 | 0.05917429 | 0.04194756 | 0.05315946 | 0.02474487 | 0.05056552 | 0.02340965 | 0.04508912 | 0.02250292 | 0.04126717 | 0.04236575 | 0.04649911 | 0.04728523 | 0.01874865 | 0.04957962 | 0.04187771 | 0.0243623 | 0.02564397 |
| 31.0833333 | 0.09102847 | 0.02758189 | 0.08877101 | 0.05295943 | 0.05350133 | 0.01278055 | 0.08743818 | 0.01333452 | 0.05374705 | 0.01589702 | 0.02933891 | 0.03113099 | 0.0340311 | 0.03275494 | 0.01204343 | 0.03288451 | 0.03601742 | 0.01500251 | 0.01778976 |
| 31.0972222 | 0.07307524 | 0.02669909 | 0.08800709 | 0.0484688 | 0.03818946 | 0.00661223 | 0.10531022 | 0.01123934 | 0.05312809 | 0.01195456 | 0.01082645 | 0.01381139 | 0.01524425 | 0.01373028 | 0.00388911 | 0.01154505 | 0.02239305 | 0.01282894 | 0.01441425 |
| 31.1111111 | 0.05639092 | 0.02696199 | 0.07125304 | 0.03870956 | 0.02187214 | 0.00813303 | 0.08995348 | 0.01240466 | 0.04888587 | 0.01833797 | 0.00205718 | 0.00731537 | 0.01022146 | 0.00725087 | 0.00246735 | 0.00477084 | 0.01030967 | 0.01489381 | 0.01556966 |
| 31.125 | 0.04811014 | 0.03244905 | 0.05273254 | 0.0258515 | 0.01185699 | 0.00940071 | 0.04935454 | 0.01137101 | 0.0385307 | 0.02585217 | 0.00162734 | 0.01346316 | 0.01328858 | 0.00890745 | 0.00375837 | 0.00669615 | 0.00567021 | 0.01331493 | 0.01181274 |
| 31.1388889 | 0.04420971 | 0.03679241 | 0.03451407 | 0.01274577 | 0.00615667 | 0.0113066 | 0.02825541 | 0.01095665 | 0.02377088 | 0.02756713 | 0.00559972 | 0.02563794 | 0.0167512 | 0.01145554 | 0.00573531 | 0.01289685 | 0.01306896 | 0.01402432 | 0.00792426 |
| 31.1527778 | 0.02956435 | 0.02562468 | 0.02170252 | 0.01033925 | 0.00376603 | 0.01756137 | 0.05704023 | 0.01274943 | 0.01280239 | 0.02586464 | 0.01303011 | 0.03662235 | 0.01980872 | 0.0156687 | 0.00925163 | 0.02249235 | 0.02614849 | 0.01930565 | 0.00860152 |
| 31.1666667 | 0.01725094 | 0.01445266 | 0.02468244 | 0.0238082 | 0.00663515 | 0.02579142 | 0.1015277 | 0.01424032 | 0.01459604 | 0.02346341 | 0.01845666 | 0.03906769 | 0.02017818 | 0.02018188 | 0.01313361 | 0.02987634 | 0.03286896 | 0.02153019 | 0.00956564 |
| 31.1805556 | 0.02879054 | 0.02386936 | 0.03338377 | 0.03699785 | 0.00995403 | 0.02840447 | 0.1202794 | 0.01069479 | 0.02416558 | 0.01782284 | 0.01625056 | 0.0309701 | 0.01834531 | 0.02181089 | 0.01642557 | 0.03159629 | 0.02945384 | 0.01667584 | 0.00712436 |
| 31.1944444 | 0.04202651 | 0.03650301 | 0.02918014 | 0.035684 | 0.01007061 | 0.02212045 | 0.10792571 | 0.00451325 | 0.02617524 | 0.01031461 | 0.00874189 | 0.01823965 | 0.01713075 | 0.02022081 | 0.01595679 | 0.02469501 | 0.02059296 | 0.00998051 | 0.00599317 |
| 31.2083333 | 0.0380308 | 0.03649543 | 0.01534494 | 0.02628438 | 0.00831083 | 0.01145629 | 0.07694689 | 0.00124749 | 0.01652396 | 0.00843562 | 0.00375261 | 0.00727771 | 0.01544817 | 0.01553792 | 0.01009858 | 0.01443817 | 0.01178623 | 0.00550037 | 0.00841871 |
| 31.2222222 | 0.02825672 | 0.03465656 | 0.00717803 | 0.01834399 | 0.00503584 | 0.00421832 | 0.04464795 | 0.00165318 | 0.00730546 | 0.00908615 | 0.00400568 | 0.00353876 | 0.01120635 | 0.00844699 | 0.00641582 | 0.01410994 | 0.00597454 | 0.00269343 | 0.00855994 |
| 31.2361111 | 0.0252199 | 0.04108678 | 0.0059923 | 0.0140128 | 0.00177899 | 0.0025662 | 0.02251537 | 0.00457099 | 0.00765791 | 0.0067091 | 0.00426897 | 0.00889153 | 0.00640689 | 0.00326776 | 0.00782907 | 0.01979349 | 0.00285505 | 0.00152965 | 0.00490928 |
| 31.25 | 0.03042711 | 0.04977381 | 0.01024058 | 0.01319775 | 0.00095844 | 0.0026729 | 0.01258082 | 0.00696667 | 0.00905158 | 0.00545464 | 0.00261715 | 0.0170113 | 0.00367492 | 0.00285279 | 0.0070213 | 0.01912924 | 0.00218478 | 0.00195988 | 0.00163703 |
| 31.2638889 | 0.04024261 | 0.05432902 | 0.02298402 | 0.01470773 | 0.00266775 | 0.00418329 | 0.01156968 | 0.00559481 | 0.00656432 | 0.00471099 | 0.00184122 | 0.020369 | 0.00244238 | 0.00427583 | 0.00381215 | 0.01251225 | 0.00399152 | 0.00276385 | 0.00052459 |
| 31.2777778 | 0.05392565 | 0.05588938 | 0.03713571 | 0.01675174 | 0.00582841 | 0.00635151 | 0.0160436 | 0.0027405 | 0.00881834 | 0.0029725 | 0.00240899 | 0.0169966 | 0.00282385 | 0.0037571 | 0.00262339 | 0.00584418 | 0.00486199 | 0.0034888 | 0.0002338 |
| 31.2916667 | 0.06754072 | 0.05202537 | 0.04326154 | 0.01790798 | 0.00874996 | 0.00747207 | 0.02222809 | 0.00241635 | 0.01717364 | 0.00375341 | 0.00329549 | 0.00959362 | 0.00553516 | 0.00239319 | 0.00242702 | 0.00325615 | 0.00349693 | 0.00408782 | 9.87E-05 |
| 31.3055556 | 0.07347662 | 0.04182884 | 0.03914045 | 0.01640721 | 0.00943895 | 0.00759212 | 0.02408077 | 0.00435809 | 0.02152525 | 0.00566054 | 0.0047783 | 0.00564914 | 0.00840312 | 0.00275057 | 0.002596 | 0.00681584 | 0.00289337 | 0.00431063 | 0.00016507 |
| 31.3194444 | 0.06922037 | 0.02681325 | 0.02789047 | 0.01245949 | 0.00810854 | 0.00679987 | 0.01860531 | 0.00544991 | 0.01606516 | 0.0052959 | 0.0060771 | 0.0073011 | 0.00931679 | 0.00593286 | 0.00542285 | 0.01468062 | 0.00400644 | 0.00414836 | 0.00019662 |
| 31.3333333 | 0.05539595 | 0.01380475 | 0.01721825 | 0.00805001 | 0.00649964 | 0.00516637 | 0.00958349 | 0.00405225 | 0.00727699 | 0.00295834 | 0.00486633 | 0.0065716 | 0.00780365 | 0.00973972 | 0.00876161 | 0.0195449 | 0.0044469 | 0.003545 | 0.00064769 |
| 31.3472222 | 0.03390617 | 0.01207254 | 0.01270837 | 0.00412314 | 0.00536648 | 0.00341134 | 0.00336436 | 0.00229706 | 0.00346046 | 0.0014605 | 0.00312488 | 0.00410598 | 0.00495501 | 0.00933282 | 0.00885688 | 0.01723028 | 0.00337549 | 0.00252365 | 0.00165219 |
| 31.3611111 | 0.01459707 | 0.01360069 | 0.01370822 | 0.00190631 | 0.00401748 | 0.00421484 | 0.00183927 | 0.00291461 | 0.0033875 | 0.00135252 | 0.00494597 | 0.00586461 | 0.00253418 | 0.00561591 | 0.00611652 | 0.01044031 | 0.00361197 | 0.00142847 | 0.0023147 |
| 31.375 | 0.00776192 | 0.00971229 | 0.01836347 | 0.00177647 | 0.00252225 | 0.00840238 | 0.00226747 | 0.0053958 | 0.00262201 | 0.00233168 | 0.00893088 | 0.00896262 | 0.00162916 | 0.00439571 | 0.00345539 | 0.00558105 | 0.00601098 | 0.00136036 | 0.00229891 |
| 31.3888889 | 0.00858163 | 0.00984771 | 0.02419373 | 0.00184773 | 0.00177652 | 0.01310755 | 0.00520894 | 0.00794168 | 0.00253264 | 0.00545264 | 0.01137406 | 0.00968288 | 0.00247777 | 0.00665857 | 0.00345566 | 0.00706354 | 0.00732946 | 0.00267679 | 0.00207575 |
| 31.4027778 | 0.008385 | 0.01666351 | 0.02860275 | 0.00154077 | 0.00119834 | 0.01548291 | 0.00960488 | 0.00910647 | 0.00518009 | 0.00902645 | 0.01127746 | 0.00802254 | 0.00283945 | 0.00840045 | 0.00457404 | 0.00994681 | 0.0066023 | 0.00360499 | 0.00173947 |
| 31.4166667 | 0.00942181 | 0.01652954 | 0.0305915 | 0.00113453 | 0.0009019 | 0.01357196 | 0.00895305 | 0.00817978 | 0.00775043 | 0.0105039 | 0.01022714 | 0.00549205 | 0.00184552 | 0.00796941 | 0.00363566 | 0.00832524 | 0.00612302 | 0.00302365 | 0.00148112 |
| 31.4305556 | 0.01575349 | 0.01060768 | 0.02977029 | 0.00207021 | 0.00203755 | 0.00786368 | 0.00749703 | 0.00644898 | 0.00774011 | 0.00949009 | 0.01064974 | 0.00458819 | 0.00152283 | 0.00707076 | 0.00214086 | 0.00471584 | 0.00692941 | 0.0017183 | 0.00172115 |
| 31.4444444 | 0.02554955 | 0.01217514 | 0.02684125 | 0.0053166 | 0.00373647 | 0.00269671 | 0.01260767 | 0.00601528 | 0.0062771 | 0.00686032 | 0.01275905 | 0.00672189 | 0.00153193 | 0.0071693 | 0.00268904 | 0.00382404 | 0.00794705 | 0.00066441 | 0.00228933 |
| 31.4583333 | 0.0312098 | 0.01446844 | 0.0225937 | 0.0078843 | 0.00453161 | 0.00204455 | 0.01537305 | 0.00749349 | 0.00494059 | 0.00462342 | 0.01473016 | 0.01065006 | 0.00118796 | 0.00865187 | 0.0030779 | 0.00420214 | 0.00880197 | 0.00036033 | 0.00284884 |
| 31.4722222 | 0.02696118 | 0.00878594 | 0.01823424 | 0.00608512 | 0.00401641 | 0.00529651 | 0.00957533 | 0.00954585 | 0.00392618 | 0.00504569 | 0.01541272 | 0.01360657 | 0.0019637 | 0.01154882 | 0.00200535 | 0.003417 | 0.01101779 | 0.00057444 | 0.00341513 |
| 31.4861111 | 0.01928843 | 0.00665312 | 0.01650195 | 0.00406098 | 0.00379599 | 0.00628698 | 0.00390504 | 0.00988674 | 0.00252348 | 0.00791398 | 0.01493605 | 0.01323389 | 0.00379695 | 0.01504688 | 0.00135708 | 0.00421432 | 0.01496327 | 0.00080181 | 0.00358027 |
| 31.5 | 0.01646183 | 0.01542135 | 0.01802155 | 0.00737603 | 0.00648018 | 0.004475 | 0.00478607 | 0.00756541 | 0.00092458 | 0.01001336 | 0.01384471 | 0.00939045 | 0.0050853 | 0.01806356 | 0.00121908 | 0.00757343 | 0.01826363 | 0.00131054 | 0.00260959 |
| 31.5138889 | 0.01298275 | 0.01942507 | 0.01976623 | 0.01152726 | 0.01156165 | 0.00361861 | 0.00625869 | 0.0045654 | 0.00096504 | 0.00882135 | 0.01279395 | 0.00480509 | 0.00593853 | 0.02078319 | 0.00072214 | 0.01029301 | 0.01895172 | 0.00186507 | 0.00113245 |
| 31.5277778 | 0.01270503 | 0.01624596 | 0.02300583 | 0.01463787 | 0.01548606 | 0.00542006 | 0.00674348 | 0.00396702 | 0.00514952 | 0.00723236 | 0.01461157 | 0.00432624 | 0.00896987 | 0.02493739 | 0.00085733 | 0.01247694 | 0.0191562 | 0.00197408 | 0.00119663 |
| 31.5416667 | 0.02888888 | 0.02078071 | 0.03975943 | 0.0216518 | 0.01539171 | 0.01255195 | 0.01462882 | 0.00697894 | 0.01596637 | 0.01194279 | 0.02617763 | 0.01139442 | 0.01721039 | 0.03403867 | 0.00298032 | 0.01823136 | 0.02357213 | 0.00371575 | 0.00415269 |
| 31.5555556 | 0.05176978 | 0.02324101 | 0.07143153 | 0.03436947 | 0.01130431 | 0.01924888 | 0.03165477 | 0.01013266 | 0.03043767 | 0.02213771 | 0.04775056 | 0.02443735 | 0.02754895 | 0.04746215 | 0.00620216 | 0.02700295 | 0.0314153 | 0.00744533 | 0.00934135 |
| 31.5694444 | 0.07110995 | 0.01837323 | 0.09333693 | 0.05229979 | 0.01292 | 0.02098337 | 0.05749155 | 0.00802693 | 0.03862467 | 0.02872234 | 0.05978903 | 0.04121463 | 0.0311489 | 0.05636719 | 0.00876532 | 0.03235515 | 0.03551093 | 0.00810485 | 0.01550932 |
| 31.5833333 | 0.0882252 | 0.01952662 | 0.0910713 | 0.0693814 | 0.0259754 | 0.01862286 | 0.0815848 | 0.00406723 | 0.03940538 | 0.02913361 | 0.04814396 | 0.05947 | 0.02698045 | 0.05491602 | 0.00889412 | 0.03230825 | 0.03377192 | 0.00561754 | 0.02309634 |
| 31.5972222 | 0.09714138 | 0.02223298 | 0.07586589 | 0.07417866 | 0.03536017 | 0.01118833 | 0.07986547 | 0.00336793 | 0.03664794 | 0.02777223 | 0.02548481 | 0.06479066 | 0.02200068 | 0.04465115 | 0.00576996 | 0.028696 | 0.02827832 | 0.00651676 | 0.02802997 |
| 31.6111111 | 0.08986254 | 0.01592288 | 0.058218 | 0.06212479 | 0.0329193 | 0.00408338 | 0.05129124 | 0.00323877 | 0.0288822 | 0.02246657 | 0.01174863 | 0.04406976 | 0.01767491 | 0.02704396 | 0.00244453 | 0.01995527 | 0.01793728 | 0.01076976 | 0.02229844 |
| 31.625 | 0.06830767 | 0.00742356 | 0.04236767 | 0.03976427 | 0.03097624 | 0.00345921 | 0.02206098 | 0.00485639 | 0.02508985 | 0.01367558 | 0.01589189 | 0.02346903 | 0.01202977 | 0.01327318 | 0.00295261 | 0.01068526 | 0.00913664 | 0.01343623 | 0.01206346 |
| 31.6388889 | 0.04251081 | 0.00605487 | 0.03091433 | 0.01806725 | 0.04027057 | 0.00736017 | 0.01381714 | 0.01057416 | 0.03036705 | 0.01154693 | 0.02916627 | 0.02994733 | 0.00681648 | 0.01621852 | 0.00627696 | 0.01041494 | 0.00964179 | 0.01293892 | 0.01077861 |
| 31.6527778 | 0.021845 | 0.00821166 | 0.02042537 | 0.00545739 | 0.05282063 | 0.01019986 | 0.02521885 | 0.01677708 | 0.03019393 | 0.01364141 | 0.03963309 | 0.04136843 | 0.003849 | 0.0254094 | 0.01188796 | 0.01774861 | 0.0121247 | 0.01002918 | 0.0162507 |
| 31.6666667 | 0.01598751 | 0.00871115 | 0.00909863 | 0.00285766 | 0.05301791 | 0.0089934 | 0.0367984 | 0.01959068 | 0.01757767 | 0.00977964 | 0.04594847 | 0.03794405 | 0.00224824 | 0.02651755 | 0.01430045 | 0.02405894 | 0.01217293 | 0.00625438 | 0.0196882 |
| 31.6805556 | 0.02683404 | 0.00743943 | 0.00379318 | 0.00464724 | 0.0444329 | 0.00591669 | 0.03742293 | 0.0163973 | 0.01175987 | 0.00393877 | 0.04944607 | 0.02852084 | 0.00177161 | 0.02309448 | 0.01237539 | 0.02333883 | 0.01432163 | 0.00305129 | 0.0200966 |
| 31.6944444 | 0.03925883 | 0.00722757 | 0.00451861 | 0.00857364 | 0.03791772 | 0.00377413 | 0.03393431 | 0.01048623 | 0.02883521 | 0.00275112 | 0.04784766 | 0.02032683 | 0.00385471 | 0.02287257 | 0.00696646 | 0.01596882 | 0.01749892 | 0.00131641 | 0.01892899 |

| | | | | | | | | | | | | | | | | | | | |
|---|---|---|---|---|---|---|---|---|---|---|---|---|---|---|---|---|---|---|---|
| 31.7083333 | 0.04111348 | 0.01190436 | 0.0065781 | 0.01176861 | 0.03097392 | 0.00500059 | 0.03377437 | 0.01136011 | 0.04970935 | 0.00288572 | 0.03909584 | 0.01439841 | 0.00754567 | 0.02517462 | 0.00380075 | 0.00764436 | 0.0187317 | 0.00102942 | 0.01618168 |
| 31.7222222 | 0.03169412 | 0.01719799 | 0.00910388 | 0.00859956 | 0.01939997 | 0.01026127 | 0.03562256 | 0.01978843 | 0.05275842 | 0.00194842 | 0.02457342 | 0.01054897 | 0.00987602 | 0.02595048 | 0.00407214 | 0.00315587 | 0.01617111 | 0.00092954 | 0.0120577 |
| 31.7361111 | 0.0168726 | 0.01673676 | 0.00902715 | 0.00515577 | 0.01150387 | 0.01311431 | 0.03371676 | 0.02653515 | 0.04194557 | 0.00243036 | 0.01365746 | 0.00659784 | 0.00926585 | 0.02323516 | 0.00403923 | 0.00341999 | 0.01011999 | 0.00073536 | 0.00747651 |
| 31.75 | 0.01362648 | 0.01421153 | 0.00515105 | 0.00900132 | 0.020352 | 0.01127948 | 0.02614459 | 0.02521883 | 0.02638325 | 0.00424899 | 0.01836196 | 0.004295 | 0.00643117 | 0.01537178 | 0.00641137 | 0.00640006 | 0.00771003 | 0.0007916 | 0.00397393 |
| 31.7638889 | 0.02744346 | 0.01508248 | 0.00195049 | 0.01578185 | 0.04154626 | 0.00995372 | 0.01577114 | 0.0166381 | 0.01632082 | 0.00644991 | 0.03275787 | 0.00669538 | 0.00326457 | 0.00865782 | 0.01168668 | 0.0085285 | 0.01179329 | 0.0008184 | 0.00388432 |
| 31.7777778 | 0.04241864 | 0.01856854 | 0.00246246 | 0.01937098 | 0.05758842 | 0.01159784 | 0.00790733 | 0.00749117 | 0.02327091 | 0.01003962 | 0.04030052 | 0.0098949 | 0.00157805 | 0.01214177 | 0.01313012 | 0.00687282 | 0.01367439 | 0.0014552 | 0.00729574 |
| 31.7916667 | 0.0515539 | 0.02096895 | 0.00577924 | 0.01904092 | 0.06001099 | 0.01770829 | 0.00722928 | 0.00562291 | 0.04314912 | 0.01390587 | 0.03504122 | 0.01102114 | 0.00209895 | 0.01957443 | 0.0095861 | 0.00356248 | 0.00965858 | 0.0030757 | 0.01054096 |
| 31.8055556 | 0.059355 | 0.01847689 | 0.01108482 | 0.01479111 | 0.05310331 | 0.02782027 | 0.00992966 | 0.01419355 | 0.05774608 | 0.01439543 | 0.02103622 | 0.01087078 | 0.00359765 | 0.02160073 | 0.00535918 | 0.00289196 | 0.00850548 | 0.00435192 | 0.00991534 |
| 31.8194444 | 0.05745494 | 0.01234791 | 0.01606383 | 0.00682633 | 0.04604156 | 0.03393009 | 0.01267708 | 0.0244447 | 0.05614417 | 0.01065274 | 0.00983139 | 0.00925051 | 0.00526314 | 0.01739915 | 0.00234226 | 0.00356941 | 0.01428688 | 0.0044095 | 0.00700196 |
| 31.8333333 | 0.03717186 | 0.01130013 | 0.01422275 | 0.00373963 | 0.0427972 | 0.03353525 | 0.01393345 | 0.02615886 | 0.04059965 | 0.00550413 | 0.01086423 | 0.00605672 | 0.00687614 | 0.01088599 | 0.00092319 | 0.00270123 | 0.0185009 | 0.00375473 | 0.0089076 |
| 31.8472222 | 0.01645963 | 0.01461241 | 0.00717906 | 0.00994953 | 0.04221249 | 0.03596533 | 0.01183447 | 0.02478001 | 0.02294686 | 0.00193825 | 0.01883316 | 0.00585476 | 0.00746407 | 0.01138834 | 0.00125782 | 0.00147909 | 0.01597624 | 0.00313054 | 0.01594692 |
| 31.8611111 | 0.01879106 | 0.01489511 | 0.00661493 | 0.01944654 | 0.04260243 | 0.04511442 | 0.01621629 | 0.0257406 | 0.02118835 | 0.00244938 | 0.02742142 | 0.01287901 | 0.00582 | 0.0216413 | 0.00206302 | 0.00148843 | 0.01077328 | 0.00275188 | 0.01914561 |
| 31.875 | 0.03806668 | 0.02033413 | 0.01209007 | 0.02551163 | 0.04077108 | 0.05345963 | 0.02916999 | 0.02425323 | 0.03304323 | 0.00820975 | 0.03556562 | 0.02205434 | 0.0035613 | 0.03317488 | 0.0026178 | 0.00222687 | 0.00829785 | 0.00228734 | 0.01477548 |
| 31.8888889 | 0.05449538 | 0.03542254 | 0.01747919 | 0.02540391 | 0.03663468 | 0.05291613 | 0.03941542 | 0.01988354 | 0.04392858 | 0.0159144 | 0.0399344 | 0.02512455 | 0.00441828 | 0.03856452 | 0.00311808 | 0.00284725 | 0.00998776 | 0.00150043 | 0.00753184 |
| 31.9027778 | 0.06281483 | 0.04763295 | 0.02540428 | 0.02420836 | 0.03367767 | 0.04626979 | 0.04636338 | 0.01772671 | 0.05245704 | 0.01985125 | 0.03775459 | 0.02027818 | 0.00777736 | 0.03601173 | 0.00335561 | 0.00265159 | 0.01201977 | 0.00078151 | 0.00334066 |
| 31.9166667 | 0.06631234 | 0.05125722 | 0.03776224 | 0.02531871 | 0.03172523 | 0.04314299 | 0.0569155 | 0.01945681 | 0.05882108 | 0.01768801 | 0.03025952 | 0.01382888 | 0.01077143 | 0.02828184 | 0.00355231 | 0.00170215 | 0.01084854 | 0.00073622 | 0.00380491 |
| 31.9305556 | 0.06555905 | 0.0510714 | 0.05120157 | 0.02663281 | 0.02837346 | 0.04592927 | 0.06872129 | 0.02380926 | 0.05982211 | 0.01109084 | 0.02140862 | 0.00985386 | 0.01282977 | 0.01883019 | 0.00487686 | 0.00149158 | 0.0070329 | 0.00132429 | 0.00420071 |
| 31.9444444 | 0.0609213 | 0.05533322 | 0.0581554 | 0.02638272 | 0.02312325 | 0.04714572 | 0.06964326 | 0.02932212 | 0.05352667 | 0.00512448 | 0.01533839 | 0.0063401 | 0.01420733 | 0.00935691 | 0.00735265 | 0.00241048 | 0.00371666 | 0.00235493 | 0.00230682 |
| 31.9583333 | 0.05026758 | 0.05937716 | 0.05138673 | 0.02343754 | 0.01654954 | 0.04021199 | 0.05646495 | 0.03322194 | 0.04483691 | 0.00352749 | 0.01379497 | 0.00284633 | 0.01470948 | 0.00311893 | 0.00948729 | 0.00322352 | 0.00412333 | 0.00390257 | 0.0010877 |
| 31.9722222 | 0.02967363 | 0.04747703 | 0.03398537 | 0.01535778 | 0.00948627 | 0.02806367 | 0.03864736 | 0.03074616 | 0.03726771 | 0.00392959 | 0.0142683 | 0.0007562 | 0.01383164 | 0.00140736 | 0.00999518 | 0.00267395 | 0.00782237 | 0.00577904 | 0.00121471 |
| 31.9861111 | 0.01354498 | 0.02587913 | 0.01642694 | 0.00894962 | 0.00450863 | 0.01547001 | 0.02108161 | 0.021599 | 0.02386126 | 0.00560521 | 0.0136045 | 0.00016852 | 0.01092467 | 0.00126767 | 0.00821965 | 0.0011184 | 0.01143661 | 0.00754867 | 0.00157768 |
| 32 | 0.01715657 | 0.01365917 | 0.00984721 | 0.01497339 | 0.00453605 | 0.0060734 | 0.01080824 | 0.01245182 | 0.01306577 | 0.00851692 | 0.01073439 | 0.00056785 | 0.00641858 | 0.0034587 | 0.00505168 | 0.00051879 | 0.01287354 | 0.00860187 | 0.00333652 |
| 32.0138889 | 0.02792731 | 0.01815278 | 0.01833365 | 0.0265046 | 0.00776827 | 0.00313855 | 0.01660112 | 0.00752174 | 0.02120759 | 0.00811023 | 0.0071478 | 0.00122794 | 0.00253854 | 0.00917434 | 0.00231429 | 0.0011884 | 0.01136795 | 0.00773501 | 0.00637541 |
| 32.0277778 | 0.02873061 | 0.02657543 | 0.0276009 | 0.02841359 | 0.00775882 | 0.00459758 | 0.02682025 | 0.00701113 | 0.03475251 | 0.00510093 | 0.00735724 | 0.00388668 | 0.00387207 | 0.01046069 | 0.00157534 | 0.00478275 | 0.00884773 | 0.00770269 | 0.00616425 |
| 32.0416667 | 0.02314345 | 0.02514401 | 0.02803257 | 0.02699197 | 0.00840767 | 0.0122575 | 0.02446346 | 0.01045664 | 0.03287807 | 0.00979807 | 0.01751981 | 0.01568751 | 0.01634469 | 0.01575755 | 0.00514854 | 0.02007781 | 0.01582731 | 0.02055943 | 0.0078824 |
| 32.0555556 | 0.03363544 | 0.03359484 | 0.04176234 | 0.04954113 | 0.02072653 | 0.02770298 | 0.02833677 | 0.01540833 | 0.03774345 | 0.02809694 | 0.03491928 | 0.03287941 | 0.03315475 | 0.04419319 | 0.01188409 | 0.04286123 | 0.03597874 | 0.04520062 | 0.01918865 |
| 32.0694444 | 0.06302663 | 0.06382107 | 0.07027762 | 0.0853192 | 0.03498799 | 0.03793611 | 0.0541068 | 0.01887028 | 0.06699748 | 0.04557917 | 0.03908595 | 0.03519616 | 0.03415011 | 0.06922788 | 0.01304521 | 0.04566211 | 0.0479653 | 0.04997819 | 0.02497187 |
| 32.0833333 | 0.08384741 | 0.08708228 | 0.07934833 | 0.0951831 | 0.03713001 | 0.03954213 | 0.06983363 | 0.01925462 | 0.08464134 | 0.04228882 | 0.02378523 | 0.01970905 | 0.0201465 | 0.05643767 | 0.00855593 | 0.02761815 | 0.03665647 | 0.02927591 | 0.0168114 |
| 32.0972222 | 0.08247756 | 0.09320569 | 0.07316622 | 0.08048072 | 0.03214752 | 0.04380891 | 0.06553984 | 0.01587013 | 0.0769939 | 0.02191327 | 0.01059274 | 0.00629819 | 0.01210577 | 0.02606936 | 0.00759601 | 0.01843449 | 0.01694624 | 0.01767406 | 0.01154508 |
| 32.1111111 | 0.06611525 | 0.09341279 | 0.07073083 | 0.05808567 | 0.02731431 | 0.04679744 | 0.05792012 | 0.01052593 | 0.05803596 | 0.00898685 | 0.0059166 | 0.0022987 | 0.01260501 | 0.01321601 | 0.00830322 | 0.02071609 | 0.00796422 | 0.01972178 | 0.01397612 |
| 32.125 | 0.04210982 | 0.08349925 | 0.06053847 | 0.03130606 | 0.02177336 | 0.03754024 | 0.04187558 | 0.00601876 | 0.0299645 | 0.0163279 | 0.00213154 | 0.00314995 | 0.01253923 | 0.01905799 | 0.00714187 | 0.02082753 | 0.01074746 | 0.01844988 | 0.01591988 |
| 32.1388889 | 0.02142175 | 0.05913021 | 0.03793185 | 0.01280508 | 0.01304252 | 0.02220154 | 0.01902025 | 0.00398586 | 0.01189561 | 0.0304943 | 0.00055564 | 0.00990624 | 0.01133731 | 0.02649135 | 0.00620255 | 0.01850976 | 0.01578267 | 0.01437505 | 0.01702953 |
| 32.1527778 | 0.01831444 | 0.03008594 | 0.01611358 | 0.01295159 | 0.0078711 | 0.02041932 | 0.00711542 | 0.00585187 | 0.01739977 | 0.03711141 | 0.0012786 | 0.02104422 | 0.00968351 | 0.0275444 | 0.00556261 | 0.01490035 | 0.01757981 | 0.01160963 | 0.01751259 |
| 32.1666667 | 0.0287815 | 0.01046589 | 0.00663303 | 0.02100311 | 0.01208868 | 0.03437271 | 0.01060016 | 0.00954217 | 0.02930569 | 0.03718439 | 0.00336042 | 0.02977954 | 0.00659666 | 0.02356398 | 0.00476226 | 0.00931858 | 0.01528806 | 0.00998371 | 0.0166434 |
| 32.1805556 | 0.03775584 | 0.00394022 | 0.00952622 | 0.02734558 | 0.01877058 | 0.04555954 | 0.01945002 | 0.01156063 | 0.03550108 | 0.03483482 | 0.0073016 | 0.03295904 | 0.00310879 | 0.01882129 | 0.00447473 | 0.00390138 | 0.01119555 | 0.00858975 | 0.01546295 |
| 32.1944444 | 0.04277633 | 0.00568187 | 0.01487544 | 0.03188046 | 0.02300496 | 0.04870915 | 0.02646814 | 0.01278448 | 0.03863547 | 0.02910376 | 0.01009991 | 0.03359969 | 0.00177124 | 0.01589396 | 0.00444288 | 0.00167912 | 0.01049431 | 0.00714005 | 0.01277169 |
| 32.2083333 | 0.04883189 | 0.01318402 | 0.01693529 | 0.03285367 | 0.02485067 | 0.04943519 | 0.02875075 | 0.01577998 | 0.04143988 | 0.01932144 | 0.00873699 | 0.03285885 | 0.00389561 | 0.01554007 | 0.00323707 | 0.0024452 | 0.01668974 | 0.00603035 | 0.00772274 |
| 32.2222222 | 0.05461938 | 0.02408029 | 0.01776781 | 0.03023486 | 0.02385999 | 0.04682659 | 0.02722162 | 0.01945795 | 0.04744668 | 0.00906851 | 0.00469336 | 0.02676313 | 0.00834233 | 0.01916305 | 0.00182426 | 0.00439014 | 0.02615244 | 0.00475187 | 0.00303293 |
| 32.2361111 | 0.05505801 | 0.03211385 | 0.01900401 | 0.02800967 | 0.02086624 | 0.03940238 | 0.02663033 | 0.02193198 | 0.05816965 | 0.0036873 | 0.00207429 | 0.01528298 | 0.01296529 | 0.02498644 | 0.00142765 | 0.00637516 | 0.02985091 | 0.00283046 | 0.0012061 |
| 32.25 | 0.05165533 | 0.03632427 | 0.01979006 | 0.02755981 | 0.01746281 | 0.03002542 | 0.02898556 | 0.02395564 | 0.06739478 | 0.00394225 | 0.00183002 | 0.00614553 | 0.01593553 | 0.0272942 | 0.00099922 | 0.00639515 | 0.02400382 | 0.00106422 | 0.00119606 |
| 32.2638889 | 0.04771606 | 0.03933637 | 0.01946627 | 0.02650851 | 0.01469443 | 0.02265882 | 0.03528916 | 0.02346825 | 0.07002249 | 0.00588355 | 0.00142065 | 0.00405053 | 0.01603928 | 0.0231055 | 0.00044082 | 0.0042047 | 0.01302915 | 0.00035688 | 0.00158592 |
| 32.2777778 | 0.04219061 | 0.0401293 | 0.01844588 | 0.02305381 | 0.01240168 | 0.01841095 | 0.04645745 | 0.01850592 | 0.06787906 | 0.00810503 | 0.00093819 | 0.00444405 | 0.01276009 | 0.0134163 | 0.00054255 | 0.00224172 | 0.00547417 | 0.00072822 | 0.00317286 |
| 32.2916667 | 0.03617609 | 0.0412712 | 0.01655346 | 0.01632328 | 0.01112051 | 0.01619017 | 0.05744675 | 0.01346585 | 0.06312774 | 0.01023262 | 0.00188551 | 0.00430792 | 0.00751287 | 0.00597672 | 0.00094563 | 0.00183163 | 0.00516299 | 0.00110927 | 0.00586985 |
| 32.3055556 | 0.03115182 | 0.04594009 | 0.01611369 | 0.008517 | 0.01145571 | 0.01409328 | 0.06464123 | 0.01163893 | 0.05802401 | 0.00969273 | 0.00265457 | 0.00396039 | 0.00388096 | 0.00647899 | 0.0010228 | 0.00195751 | 0.0050606 | 0.00135892 | 0.00601158 |
| 32.3194444 | 0.0273019 | 0.05076502 | 0.02102496 | 0.00406206 | 0.01263436 | 0.01036792 | 0.06958871 | 0.01121971 | 0.05258273 | 0.00618782 | 0.00234976 | 0.00304389 | 0.00383915 | 0.00698598 | 0.00123318 | 0.00293375 | 0.00406342 | 0.00267058 | 0.00387974 |
| 32.3333333 | 0.027269 | 0.05199099 | 0.02857804 | 0.0032592 | 0.01347322 | 0.00684001 | 0.07086842 | 0.01022142 | 0.04462758 | 0.00460121 | 0.0029103 | 0.00375942 | 0.00543634 | 0.00724839 | 0.00174718 | 0.00523828 | 0.00843533 | 0.00414031 | 0.00433603 |
| 32.3472222 | 0.03072571 | 0.04885787 | 0.03277305 | 0.00480019 | 0.01354786 | 0.00812346 | 0.06882596 | 0.00835148 | 0.03393018 | 0.00697423 | 0.00424327 | 0.00733203 | 0.00569915 | 0.01346903 | 0.00170195 | 0.00674504 | 0.01449494 | 0.00413427 | 0.00726908 |
| 32.3611111 | 0.03539121 | 0.04525559 | 0.03265692 | 0.00786103 | 0.01397131 | 0.01513551 | 0.06752646 | 0.00734808 | 0.02341672 | 0.00924393 | 0.0050785 | 0.0110582 | 0.00407299 | 0.01900733 | 0.00125059 | 0.0067828 | 0.01555342 | 0.00351954 | 0.00887047 |
| 32.375 | 0.0402265 | 0.04599207 | 0.03000916 | 0.00895753 | 0.01558096 | 0.02204145 | 0.065445 | 0.00861056 | 0.01726822 | 0.01059468 | 0.00637066 | 0.01388942 | 0.00269954 | 0.01846199 | 0.00137528 | 0.00694661 | 0.0113807 | 0.00398316 | 0.00755104 |
| 32.3888889 | 0.04057394 | 0.04859894 | 0.0273058 | 0.00655501 | 0.01582333 | 0.02256117 | 0.05794115 | 0.00941047 | 0.01432736 | 0.01247597 | 0.00876524 | 0.01733876 | 0.00342208 | 0.01432222 | 0.00215634 | 0.00801792 | 0.00565348 | 0.00550507 | 0.00425212 |
| 32.4027778 | 0.03494609 | 0.04822247 | 0.02656449 | 0.00364375 | 0.01217161 | 0.01694721 | 0.04449479 | 0.00681562 | 0.00980264 | 0.01346677 | 0.01068622 | 0.02198268 | 0.00520222 | 0.00980456 | 0.00253946 | 0.00901705 | 0.00247041 | 0.00634855 | 0.00192425 |
| 32.4166667 | 0.03257035 | 0.04597452 | 0.02708343 | 0.00412455 | 0.00825756 | 0.01148912 | 0.02777578 | 0.00366454 | 0.00430361 | 0.01204036 | 0.01103991 | 0.02593357 | 0.00584309 | 0.00715141 | 0.00212349 | 0.00879438 | 0.00366162 | 0.00559751 | 0.00205086 |
| 32.4305556 | 0.04100619 | 0.04650302 | 0.02789692 | 0.00898469 | 0.00921038 | 0.01215301 | 0.01244359 | 0.00515779 | 0.00387468 | 0.00822559 | 0.01108505 | 0.02550653 | 0.00519747 | 0.00667627 | 0.00210492 | 0.00782763 | 0.00766918 | 0.00419561 | 0.00191809 |
| 32.4444444 | 0.05254252 | 0.04873729 | 0.02963495 | 0.01456681 | 0.01297216 | 0.01697103 | 0.00359722 | 0.00935745 | 0.00951787 | 0.00370634 | 0.01188107 | 0.01904592 | 0.00467388 | 0.00750985 | 0.00278107 | 0.00666425 | 0.01078801 | 0.00281174 | 0.00136249 |
| 32.4583333 | 0.05374593 | 0.0471052 | 0.03040841 | 0.01574937 | 0.01378721 | 0.01769901 | 0.00232879 | 0.00926429 | 0.01418423 | 0.00129157 | 0.01307575 | 0.01000309 | 0.00468904 | 0.008766 | 0.00259002 | 0.00447561 | 0.01082425 | 0.00139096 | 0.00218424 |
| 32.4722222 | 0.04367117 | 0.04032902 | 0.02352641 | 0.01027619 | 0.01098748 | 0.01171891 | 0.00857421 | 0.00481327 | 0.01289154 | 0.00291051 | 0.01401246 | 0.00541738 | 0.0045963 | 0.00938875 | 0.00132127 | 0.00215508 | 0.00821951 | 0.00048992 | 0.00292977 |
| 32.4861111 | 0.03068748 | 0.0304815 | 0.01218299 | 0.00540085 | 0.00746005 | 0.00940547 | 0.02134498 | 0.00188604 | 0.00772949 | 0.00791136 | 0.01394631 | 0.00497113 | 0.00460172 | 0.00841633 | 0.00064963 | 0.00116456 | 0.00454629 | 0.00040495 | 0.0028894 |
| 32.5 | 0.02144366 | 0.02169264 | 0.00784471 | 0.00735964 | 0.00457256 | 0.01747767 | 0.03418515 | 0.00201641 | 0.00341394 | 0.01301439 | 0.01233333 | 0.00511978 | 0.00515668 | 0.00725001 | 0.00092328 | 0.00136238 | 0.00210663 | 0.00058722 | 0.00273379 |
| 32.5138889 | 0.02046675 | 0.02123478 | 0.00854465 | 0.00907908 | 0.0031102 | 0.02657788 | 0.0409276 | 0.00197788 | 0.00311668 | 0.01561058 | 0.00958831 | 0.00949697 | 0.00543013 | 0.00878857 | 0.00117744 | 0.00234895 | 0.00298637 | 0.00055026 | 0.00191677 |
| 32.5277778 | 0.02698355 | 0.03056459 | 0.00897262 | 0.00677271 | 0.00408396 | 0.02723713 | 0.03853568 | 0.00178104 | 0.00381904 | 0.01751971 | 0.00939887 | 0.01787264 | 0.00582832 | 0.01583757 | 0.00142334 | 0.00405463 | 0.00829167 | 0.00151031 | 0.00095125 |
| 32.5416667 | 0.03741202 | 0.04340874 | 0.02145728 | 0.01107399 | 0.00844692 | 0.01906658 | 0.02634369 | 0.00417046 | 0.00721696 | 0.02395188 | 0.01835738 | 0.02988018 | 0.00999904 | 0.03213077 | 0.00170486 | 0.01033699 | 0.01963217 | 0.00686547 | 0.00192804 |
| 32.5555556 | 0.05345973 | 0.052164 | 0.04889676 | 0.02943739 | 0.01583037 | 0.01767765 | 0.02516608 | 0.00869282 | 0.02066914 | 0.03418965 | 0.03849326 | 0.04577603 | 0.01832458 | 0.05098236 | 0.00241811 | 0.02092723 | 0.0342557 | 0.01462527 | 0.00338865 |
| 32.5694444 | 0.07239041 | 0.05077185 | 0.06622683 | 0.04999142 | 0.02083537 | 0.03278979 | 0.0556463 | 0.01453022 | 0.03786668 | 0.04056734 | 0.05781048 | 0.05725163 | 0.02338421 | 0.05689217 | 0.00440571 | 0.02640988 | 0.04228648 | 0.01604387 | 0.0030929 |
| 32.5833333 | 0.08418427 | 0.04241457 | 0.06721146 | 0.06809886 | 0.02040271 | 0.04621944 | 0.09520308 | 0.02251357 | 0.05026889 | 0.04274402 | 0.06369595 | 0.06389308 | 0.02029661 | 0.05038104 | 0.00633658 | 0.0246143 | 0.03967269 | 0.00967022 | 0.00517664 |
| 32.5972222 | 0.08409332 | 0.03411828 | 0.06804849 | 0.08851407 | 0.02053766 | 0.04737759 | 0.11420839 | 0.02996877 | 0.05604338 | 0.0417116 | 0.05789948 | 0.067563 | 0.01296503 | 0.04113665 | 0.00762024 | 0.02091449 | 0.03222716 | 0.00358383 | 0.01279019 |
| 32.6111111 | 0.0739244 | 0.02748845 | 0.06901209 | 0.09716019 | 0.02465867 | 0.04167239 | 0.10799511 | 0.03179717 | 0.05085433 | 0.02947819 | 0.04370283 | 0.054774 | 0.00633638 | 0.0280536 | 0.00902102 | 0.01421159 | 0.02033484 | 0.00299002 | 0.02215823 |
| 32.625 | 0.0572294 | 0.02212562 | 0.06439504 | 0.08149557 | 0.03074455 | 0.02928013 | 0.08688265 | 0.02857293 | 0.03727556 | 0.01545306 | 0.02461848 | 0.03256449 | 0.00383572 | 0.01445156 | 0.0077368 | 0.00804374 | 0.01240302 | 0.00584927 | 0.02900385 |
| 32.6388889 | 0.03781795 | 0.01613483 | 0.05652116 | 0.05156473 | 0.03922462 | 0.01387332 | 0.06645835 | 0.02370271 | 0.02509286 | 0.01664883 | 0.01028538 | 0.03185992 | 0.00676381 | 0.01129297 | 0.00430809 | 0.00914157 | 0.01953927 | 0.00767149 | 0.03300855 |
| 32.6527778 | 0.02258225 | 0.01003326 | 0.04732732 | 0.02403957 | 0.04759868 | 0.00723985 | 0.05254401 | 0.01834844 | 0.01572975 | 0.02166373 | 0.00722836 | 0.04805032 | 0.00942638 | 0.01499203 | 0.00444971 | 0.01109492 | 0.02453035 | 0.00736573 | 0.03240487 |
| 32.6666667 | 0.01580832 | 0.00684649 | 0.03825927 | 0.01072297 | 0.04969506 | 0.00671434 | 0.04226437 | 0.01441078 | 0.00752536 | 0.01918927 | 0.01174091 | 0.05409515 | 0.00664873 | 0.01643246 | 0.00631133 | 0.00923663 | 0.01707243 | 0.00675253 | 0.02250859 |
| 32.6805556 | 0.01239567 | 0.00516563 | 0.03011882 | 0.01297412 | 0.04416306 | 0.00583891 | 0.03373568 | 0.01138552 | 0.00692089 | 0.01368752 | 0.01775202 | 0.04843868 | 0.00336692 | 0.01686905 | 0.0060903 | 0.00756028 | 0.01042563 | 0.00528041 | 0.01218754 |
| 32.6944444 | 0.00753064 | 0.00358444 | 0.02182428 | 0.01831212 | 0.03385696 | 0.00776542 | 0.02148935 | 0.00729907 | 0.01724282 | 0.00744129 | 0.02388723 | 0.03980135 | 0.00319451 | 0.02027554 | 0.00527222 | 0.00764838 | 0.0130725 | 0.00319352 | 0.01572476 |
| 32.7083333 | 0.00497103 | 0.00420752 | 0.01436604 | 0.01894338 | 0.02314829 | 0.00725314 | 0.01654533 | 0.00373929 | 0.03127625 | 0.00357666 | 0.02974742 | 0.03126582 | 0.00311145 | 0.02693665 | 0.00411042 | 0.00761011 | 0.02267355 | 0.00318754 | 0.028853 |
| 32.7222222 | 0.00871187 | 0.00639848 | 0.00814344 | 0.01714837 | 0.0144921 | 0.00882577 | 0.04047569 | 0.0017762 | 0.0443576 | 0.00266646 | 0.03485789 | 0.02597819 | 0.00447506 | 0.03282845 | 0.00221283 | 0.00570522 | 0.03039252 | 0.00359794 | 0.0385012 |
| 32.7361111 | 0.01444131 | 0.00762133 | 0.00490823 | 0.01733713 | 0.00793858 | 0.02613303 | 0.07732183 | 0.00099791 | 0.05506891 | 0.00300386 | 0.03619534 | 0.02354843 | 0.00873322 | 0.03306024 | 0.00111888 | 0.0038537 | 0.03033648 | 0.00253229 | 0.03935838 |
| 32.75 | 0.0173354 | 0.00785465 | 0.00680737 | 0.01980501 | 0.00477905 | 0.05250398 | 0.09914825 | 0.00219877 | 0.05631518 | 0.00635079 | 0.0298705 | 0.01849165 | 0.01080777 | 0.02716604 | 0.00242951 | 0.00569705 | 0.0218847 | 0.00191731 | 0.0342282 |
| 32.7638889 | 0.01613249 | 0.0081166 | 0.01139492 | 0.01774006 | 0.00562587 | 0.06736872 | 0.09352471 | 0.00665624 | 0.04056696 | 0.00991371 | 0.01832279 | 0.01134325 | 0.00784477 | 0.0178693 | 0.00596762 | 0.01016487 | 0.0113298 | 0.00188867 | 0.02503117 |
| 32.7777778 | 0.01052447 | 0.00969796 | 0.01600173 | 0.01205084 | 0.0079738 | 0.05996546 | 0.0617774 | 0.012404 | 0.02148188 | 0.0101064 | 0.00804908 | 0.01063394 | 0.00564043 | 0.00871038 | 0.00913583 | 0.01226557 | 0.00964737 | 0.00132663 | 0.01290363 |
| 32.7916667 | 0.00552638 | 0.01250481 | 0.01888366 | 0.01459614 | 0.00802612 | 0.03961799 | 0.02646705 | 0.0137221 | 0.0226059 | 0.00973588 | 0.00373539 | 0.01598306 | 0.00727569 | 0.00507323 | 0.00970686 | 0.00976145 | 0.01534612 | 0.00086945 | 0.00401223 |
| 32.8055556 | 0.00713147 | 0.01371023 | 0.01861695 | 0.02849735 | 0.00560793 | 0.01923867 | 0.00825877 | 0.00895294 | 0.03565715 | 0.01216703 | 0.0037432 | 0.01861632 | 0.00748624 | 0.00866633 | 0.00780955 | 0.00506555 | 0.01813453 | 0.00144164 | 0.00179135 |
| 32.8194444 | 0.0106537 | 0.01308903 | 0.01626244 | 0.04204557 | 0.00739464 | 0.01001799 | 0.00672029 | 0.00562501 | 0.04502992 | 0.0175858 | 0.00321049 | 0.01416999 | 0.0051979 | 0.01461 | 0.00441997 | 0.00164055 | 0.01385631 | 0.00261595 | 0.00284891 |
| 32.8333333 | 0.00977992 | 0.01347736 | 0.01346943 | 0.04512411 | 0.01789844 | 0.0145705 | 0.00890287 | 0.00814211 | 0.05322617 | 0.02503285 | 0.00260072 | 0.00916558 | 0.00317434 | 0.01826431 | 0.00239862 | 0.00058679 | 0.00873208 | 0.00303707 | 0.00320912 |
| 32.8472222 | 0.01102361 | 0.01363873 | 0.01047083 | 0.03387836 | 0.03166594 | 0.02003872 | 0.00927592 | 0.00946911 | 0.0550354 | 0.03256147 | 0.00501767 | 0.0134347 | 0.00161524 | 0.01795297 | 0.00293481 | 0.00154217 | 0.01159728 | 0.00222727 | 0.00232577 |
| 32.8611111 | 0.02026086 | 0.01128201 | 0.00630014 | 0.02210314 | 0.04312964 | 0.01969581 | 0.00928547 | 0.00591505 | 0.04237165 | 0.03540346 | 0.00851846 | 0.02049793 | 0.00111425 | 0.01382949 | 0.00324727 | 0.00420139 | 0.01745563 | 0.00123027 | 0.00147967 |
| 32.875 | 0.03128735 | 0.00871255 | 0.00430487 | 0.03221508 | 0.05237647 | 0.0129329 | 0.00864201 | 0.00327229 | 0.02293369 | 0.02893959 | 0.00994654 | 0.02027252 | 0.0031104 | 0.00773625 | 0.00405623 | 0.0072546 | 0.01703696 | 0.00145444 | 0.00076008 |
| 32.8888889 | 0.03952099 | 0.00846007 | 0.00936835 | 0.05331663 | 0.05873944 | 0.00777137 | 0.00627457 | 0.00505289 | 0.01734392 | 0.01599436 | 0.01082495 | 0.01505335 | 0.00652425 | 0.00598629 | 0.00693942 | 0.00868165 | 0.01170541 | 0.003759 | 0.00049241 |
| 32.9027778 | 0.04714584 | 0.01078688 | 0.01819839 | 0.06389902 | 0.05935616 | 0.01525727 | 0.00688043 | 0.00657239 | 0.02919961 | 0.00896733 | 0.01252593 | 0.00906652 | 0.0077763 | 0.01321975 | 0.00808748 | 0.00763692 | 0.00884624 | 0.00675971 | 0.00130063 |
| 32.9166667 | 0.0545081 | 0.01530721 | 0.02508489 | 0.06308874 | 0.05312454 | 0.02745751 | 0.01854883 | 0.00642953 | 0.04280876 | 0.01258071 | 0.01355449 | 0.0044268 | 0.00557903 | 0.02268016 | 0.00593253 | 0.00559879 | 0.0116464 | 0.00829368 | 0.00181165 |
| 32.9305556 | 0.05878269 | 0.02099801 | 0.02711508 | 0.05619194 | 0.04062203 | 0.03584397 | 0.03596011 | 0.01096559 | 0.05814903 | 0.01733851 | 0.01205248 | 0.00370428 | 0.00247579 | 0.02610229 | 0.00430555 | 0.00490515 | 0.01420005 | 0.00844544 | 0.00138587 |
| 32.9444444 | 0.05684323 | 0.0253913 | 0.02421374 | 0.04546867 | 0.02343507 | 0.04343105 | 0.04828025 | 0.01851788 | 0.07326404 | 0.01770671 | 0.008193 | 0.0075055 | 0.00122678 | 0.02289436 | 0.00486472 | 0.00544061 | 0.01112315 | 0.00782337 | 0.0013231 |
| 32.9583333 | 0.04679724 | 0.02519425 | 0.01910831 | 0.03225466 | 0.01087958 | 0.04926816 | 0.0494431 | 0.02165467 | 0.07962986 | 0.0140749 | 0.00444039 | 0.01348801 | 0.00183625 | 0.0162009 | 0.00588013 | 0.00529632 | 0.00571537 | 0.0064006 | 0.00146451 |
| 32.9722222 | 0.03091008 | 0.02010243 | 0.01455459 | 0.01960365 | 0.0127707 | 0.04638218 | 0.04025112 | 0.02100044 | 0.08029152 | 0.00742504 | 0.00238477 | 0.01959966 | 0.00208033 | 0.00883584 | 0.0069718 | 0.00467393 | 0.00415251 | 0.00518959 | 0.00151219 |
| 32.9861111 | 0.01480366 | 0.01350424 | 0.00979491 | 0.01105843 | 0.02193326 | 0.03196314 | 0.02833009 | 0.01790018 | 0.06809067 | 0.0024279 | 0.00154187 | 0.02342554 | 0.00269948 | 0.00365876 | 0.00962421 | 0.00570614 | 0.00648893 | 0.00572565 | 0.00249506 |
| 33 | 0.00732366 | 0.00745332 | 0.00787576 | 0.01434696 | 0.0283431 | 0.01714167 | 0.01580446 | 0.01138202 | 0.04154767 | 0.00160508 | 0.00156311 | 0.02268248 | 0.00580109 | 0.00272737 | 0.01416798 | 0.00903647 | 0.00886578 | 0.00759453 | 0.00431238 |
| 33.0138889 | 0.01239737 | 0.00312784 | 0.01506174 | 0.03216561 | 0.03277744 | 0.0175373 | 0.0110733 | 0.00525964 | 0.0328093 | 0.00157967 | 0.00204886 | 0.01766255 | 0.00810103 | 0.0034712 | 0.01780279 | 0.01226227 | 0.0094146 | 0.00823606 | 0.00540123 |
| 33.0277778 | 0.01986088 | 0.00272533 | 0.02182018 | 0.04890122 | 0.03457752 | 0.02482939 | 0.0195453 | 0.00327313 | 0.05115067 | 0.00257154 | 0.0037809 | 0.0119795 | 0.00726873 | 0.00511563 | 0.01655079 | 0.01055039 | 0.00934236 | 0.0076105 | 0.00416831 |
| 33.0416667 | 0.01797169 | 0.00931269 | 0.02025391 | 0.04096996 | 0.02644657 | 0.02525001 | 0.02441839 | 0.00348934 | 0.05253787 | 0.0129667 | 0.01375134 | 0.01958595 | 0.01142904 | 0.01722619 | 0.01082519 | 0.0097352 | 0.02276958 | 0.01508714 | 0.00440475 |
| 33.0555556 | 0.02283867 | 0.03099532 | 0.02696057 | 0.02995074 | 0.01785091 | 0.03816031 | 0.03507981 | 0.00721674 | 0.04046566 | 0.03752859 | 0.03465326 | 0.04569558 | 0.02694453 | 0.04378933 | 0.00696999 | 0.01786596 | 0.05843851 | 0.03367491 | 0.00952129 |
| 33.0694444 | 0.05655466 | 0.07079289 | 0.04493734 | 0.05516789 | 0.02759677 | 0.06919586 | 0.06321761 | 0.01706059 | 0.06032368 | 0.05868774 | 0.04628927 | 0.05995076 | 0.03818672 | 0.06006314 | 0.00627277 | 0.0201756 | 0.08558858 | 0.03917141 | 0.01068971 |
| 33.0833333 | 0.10073992 | 0.10854492 | 0.0520984 | 0.0867989 | 0.04641183 | 0.08595745 | 0.07401761 | 0.02543789 | 0.09070731 | 0.05273392 | 0.03269055 | 0.04080156 | 0.02782383 | 0.04607456 | 0.00510102 | 0.01149336 | 0.07533408 | 0.02364973 | 0.00700852 |
| 33.0972222 | 0.12129587 | 0.12326626 | 0.04592942 | 0.08767397 | 0.0577435 | 0.0824089 | 0.05851889 | 0.0265289 | 0.1050923 | 0.02859063 | 0.01401202 | 0.01614294 | 0.01261614 | 0.02057534 | 0.00524651 | 0.00617928 | 0.04356774 | 0.01255018 | 0.00863411 |
| 33.1111111 | 0.10111779 | 0.11874732 | 0.03787355 | 0.06755365 | 0.05576827 | 0.0752566 | 0.04231676 | 0.02355628 | 0.11053739 | 0.01733658 | 0.00990594 | 0.01154463 | 0.01313208 | 0.00843153 | 0.00709853 | 0.00486107 | 0.01916869 | 0.01053487 | 0.01255235 |
| 33.125 | 0.05740495 | 0.10173928 | 0.03079789 | 0.04270592 | 0.04119698 | 0.07006683 | 0.03061666 | 0.02024258 | 0.10001489 | 0.02590492 | 0.01330821 | 0.01592304 | 0.0191276 | 0.01215498 | 0.00771359 | 0.0047274 | 0.01439941 | 0.00845587 | 0.01262904 |
| 33.1388889 | 0.02300639 | 0.07368882 | 0.02495692 | 0.02122757 | 0.02201748 | 0.06711002 | 0.01966731 | 0.01644599 | 0.07652087 | 0.03336664 | 0.01427524 | 0.01835046 | 0.01946071 | 0.01918845 | 0.00694559 | 0.00565196 | 0.02246418 | 0.00790287 | 0.01034524 |
| 33.1527778 | 0.0113484 | 0.03961031 | 0.01897799 | 0.01005602 | 0.01005926 | 0.06164 | 0.00971833 | 0.00969509 | 0.05295302 | 0.02937671 | 0.01287403 | 0.02175755 | 0.01653778 | 0.0224306 | 0.00665078 | 0.0048319 | 0.03011115 | 0.00799453 | 0.00905281 |
| 33.1666667 | 0.01637292 | 0.01501728 | 0.01170513 | 0.01352715 | 0.01262217 | 0.05050442 | 0.00419883 | 0.0031527 | 0.03408563 | 0.02222067 | 0.01136975 | 0.02778849 | 0.01204647 | 0.02247467 | 0.00736911 | 0.00370843 | 0.03323398 | 0.00613846 | 0.01117048 |
| 33.1805556 | 0.02199247 | 0.01103131 | 0.00547609 | 0.02415591 | 0.02081055 | 0.03516138 | 0.00681852 | 0.00099401 | 0.02239447 | 0.0203145 | 0.008867 | 0.03254119 | 0.00605299 | 0.01961695 | 0.00715201 | 0.00503402 | 0.0329518 | 0.00385724 | 0.01282272 |
| 33.1944444 | 0.02452661 | 0.01474862 | 0.00513831 | 0.03378978 | 0.026771 | 0.0210244 | 0.01425513 | 0.00138818 | 0.01488517 | 0.02560919 | 0.00674776 | 0.03339106 | 0.00145935 | 0.01650998 | 0.00554104 | 0.00664005 | 0.03082588 | 0.0028394 | 0.00928312 |
| 33.2083333 | 0.03095813 | 0.01830436 | 0.01252459 | 0.04391656 | 0.03051397 | 0.02191244 | 0.02175349 | 0.00401319 | 0.01072913 | 0.03132643 | 0.00600413 | 0.03334646 | 0.00084363 | 0.01572328 | 0.00447409 | 0.00585918 | 0.03012898 | 0.00243662 | 0.00417011 |

| | | | | | | | | | | | | | | | | | | | |
|---|---|---|---|---|---|---|---|---|---|---|---|---|---|---|---|---|---|---|---|
| 33.2222222 | 0.04159803 | 0.02167652 | 0.0225175 | 0.05110965 | 0.03226465 | 0.04166744 | 0.02686337 | 0.00878095 | 0.01854686 | 0.02801104 | 0.0048255 | 0.03491719 | 0.00272082 | 0.01514707 | 0.00385514 | 0.00418757 | 0.03192911 | 0.00318979 | 0.00177688 |
| 33.2361111 | 0.05242897 | 0.02125029 | 0.02797574 | 0.04717177 | 0.03383908 | 0.06276729 | 0.02930928 | 0.01349745 | 0.03968553 | 0.01757667 | 0.00536973 | 0.03493461 | 0.00511616 | 0.01199628 | 0.00256718 | 0.00335892 | 0.0345074 | 0.00464092 | 0.00191521 |
| 33.25 | 0.05945851 | 0.01771494 | 0.02787733 | 0.03648593 | 0.0343055 | 0.07339262 | 0.03011852 | 0.01757665 | 0.06410404 | 0.00862503 | 0.00754902 | 0.02640438 | 0.00735743 | 0.00732847 | 0.00151804 | 0.00506877 | 0.0348343 | 0.00418991 | 0.00369132 |
| 33.2638889 | 0.05904514 | 0.0109071 | 0.02551421 | 0.02755309 | 0.03100125 | 0.0745579 | 0.02867646 | 0.02079512 | 0.08388637 | 0.00363015 | 0.0084521 | 0.01527946 | 0.00815378 | 0.00431527 | 0.00112894 | 0.00803119 | 0.02780406 | 0.00213587 | 0.00543149 |
| 33.2777778 | 0.05152119 | 0.00531771 | 0.02207948 | 0.02082184 | 0.02364225 | 0.06776917 | 0.02478887 | 0.02328809 | 0.09338361 | 0.00257118 | 0.00879797 | 0.01601194 | 0.00651839 | 0.00587239 | 0.00156221 | 0.00781501 | 0.01510581 | 0.00083289 | 0.0057975 |
| 33.2916667 | 0.04161309 | 0.00511802 | 0.01735906 | 0.01364586 | 0.01494638 | 0.05721394 | 0.02014042 | 0.02639751 | 0.08902568 | 0.00666972 | 0.00907367 | 0.02353068 | 0.00415998 | 0.01161059 | 0.00237367 | 0.00437559 | 0.00856031 | 0.00104484 | 0.00574496 |
| 33.3055556 | 0.0305516 | 0.00517334 | 0.01119506 | 0.00715682 | 0.00722906 | 0.0453329 | 0.01479755 | 0.02941782 | 0.07818543 | 0.01247283 | 0.00887682 | 0.02562799 | 0.00325382 | 0.01736021 | 0.00195289 | 0.00181767 | 0.01139719 | 0.00192652 | 0.00588255 |
| 33.3194444 | 0.01662859 | 0.0066623 | 0.00483564 | 0.00559517 | 0.00273005 | 0.0284616 | 0.00804276 | 0.02807924 | 0.07319473 | 0.01484605 | 0.00871564 | 0.02244984 | 0.00372886 | 0.01873978 | 0.00087176 | 0.00169191 | 0.01348802 | 0.00318762 | 0.00550849 |
| 33.3333333 | 0.00568866 | 0.0089876 | 0.001424 | 0.00732333 | 0.00247102 | 0.01306627 | 0.00370549 | 0.02268596 | 0.07429327 | 0.01232717 | 0.00713457 | 0.01585534 | 0.00495175 | 0.01343694 | 0.00080256 | 0.00281668 | 0.01048986 | 0.00388833 | 0.00520513 |
| 33.3472222 | 0.00295568 | 0.01122926 | 0.00272554 | 0.00618244 | 0.00281787 | 0.00870135 | 0.00376327 | 0.01883674 | 0.07233313 | 0.00725093 | 0.00485627 | 0.00788415 | 0.0067426 | 0.00783328 | 0.0017658 | 0.00574088 | 0.01147355 | 0.00311359 | 0.00561028 |
| 33.3611111 | 0.00434532 | 0.0197185 | 0.00681607 | 0.00318682 | 0.00345194 | 0.00877778 | 0.00381528 | 0.01628027 | 0.05994765 | 0.00554148 | 0.0061717 | 0.00400454 | 0.0082134 | 0.01159436 | 0.00214868 | 0.00885192 | 0.02034463 | 0.00301375 | 0.00597623 |
| 33.375 | 0.00823835 | 0.02640836 | 0.01018859 | 0.00364124 | 0.00680935 | 0.00684968 | 0.0033597 | 0.01267873 | 0.04201742 | 0.0081976 | 0.00994619 | 0.00532993 | 0.00883158 | 0.02096629 | 0.00160802 | 0.00944851 | 0.02686416 | 0.00456693 | 0.00614689 |
| 33.3888889 | 0.01573817 | 0.02976053 | 0.01183273 | 0.00720875 | 0.01210113 | 0.00484333 | 0.00504907 | 0.01034136 | 0.03237748 | 0.01012292 | 0.01214543 | 0.00766899 | 0.0087831 | 0.02656221 | 0.00223614 | 0.00847944 | 0.02394419 | 0.00688058 | 0.00628605 |
| 33.4027778 | 0.02373669 | 0.03832786 | 0.01215082 | 0.01008108 | 0.0168408 | 0.00641086 | 0.00620444 | 0.00928599 | 0.02948334 | 0.00974853 | 0.01035412 | 0.0092332 | 0.00767478 | 0.02529947 | 0.00440681 | 0.00659603 | 0.01419709 | 0.00946322 | 0.0056435 |
| 33.4166667 | 0.02637099 | 0.04291635 | 0.01228329 | 0.00981324 | 0.01779854 | 0.01509327 | 0.00454296 | 0.00739524 | 0.02135279 | 0.00974698 | 0.0059695 | 0.01068504 | 0.00579734 | 0.01883969 | 0.00554832 | 0.00420935 | 0.00607868 | 0.01094642 | 0.00450661 |
| 33.4305556 | 0.02391185 | 0.03758468 | 0.01485926 | 0.00806842 | 0.01474338 | 0.02723592 | 0.00285204 | 0.00536164 | 0.01085547 | 0.01201131 | 0.00377824 | 0.01227727 | 0.00444215 | 0.01154615 | 0.00447486 | 0.00323953 | 0.00485996 | 0.01066613 | 0.0039048 |
| 33.4444444 | 0.0228492 | 0.0318485 | 0.01986263 | 0.00786761 | 0.01103054 | 0.03531857 | 0.00224339 | 0.00402093 | 0.00836616 | 0.01515358 | 0.00290084 | 0.01351148 | 0.00366357 | 0.00720468 | 0.00301878 | 0.00373475 | 0.00483743 | 0.00893287 | 0.0034705 |
| 33.4583333 | 0.02443618 | 0.02990055 | 0.02428503 | 0.00827222 | 0.00816702 | 0.03787959 | 0.00286522 | 0.00299534 | 0.01100392 | 0.01636248 | 0.00232526 | 0.01355959 | 0.00334495 | 0.00715244 | 0.00227409 | 0.00404622 | 0.00453127 | 0.0062149 | 0.00292851 |
| 33.4722222 | 0.02610483 | 0.02842774 | 0.02646202 | 0.00814413 | 0.00595968 | 0.03542614 | 0.00575965 | 0.00250976 | 0.01104482 | 0.0149095 | 0.00486288 | 0.01250042 | 0.00418987 | 0.01059019 | 0.00147559 | 0.00397444 | 0.00938558 | 0.00394643 | 0.00288165 |
| 33.4861111 | 0.02747091 | 0.02661235 | 0.02593437 | 0.00781665 | 0.00467105 | 0.0312967 | 0.00805133 | 0.00201065 | 0.0090777 | 0.01198695 | 0.00851387 | 0.01028235 | 0.00559699 | 0.01439965 | 0.00070742 | 0.00398523 | 0.01629636 | 0.00294293 | 0.00274053 |
| 33.5 | 0.02697948 | 0.02292656 | 0.02213183 | 0.00596728 | 0.00325602 | 0.03375887 | 0.00601475 | 0.00151783 | 0.00864129 | 0.00918314 | 0.01033392 | 0.00820194 | 0.0065804 | 0.01548576 | 0.00043806 | 0.00418359 | 0.01897651 | 0.00265409 | 0.00184576 |
| 33.5138889 | 0.02214954 | 0.01584727 | 0.01723923 | 0.00286443 | 0.00225252 | 0.04266361 | 0.00309052 | 0.00201693 | 0.01081846 | 0.00828249 | 0.01149979 | 0.00884812 | 0.00732512 | 0.01417866 | 0.00079442 | 0.00501401 | 0.01815906 | 0.00295884 | 0.00144646 |
| 33.5277778 | 0.01555548 | 0.01453819 | 0.01641205 | 0.00377158 | 0.00259464 | 0.04726975 | 0.00242177 | 0.0025187 | 0.01117354 | 0.010363 | 0.01531328 | 0.01313444 | 0.00795508 | 0.01487083 | 0.00214347 | 0.00730921 | 0.02016785 | 0.00460853 | 0.00220533 |
| 33.5416667 | 0.01359464 | 0.03221689 | 0.0243213 | 0.01471216 | 0.00414201 | 0.03618952 | 0.00581584 | 0.00508809 | 0.00991004 | 0.01656526 | 0.02588127 | 0.02192606 | 0.00950597 | 0.02494368 | 0.00460057 | 0.01233103 | 0.03499356 | 0.0095086 | 0.00382018 |
| 33.5555556 | 0.02195241 | 0.06509104 | 0.04092806 | 0.03499183 | 0.00992426 | 0.01885742 | 0.01912863 | 0.01229423 | 0.01883578 | 0.0266572 | 0.04045787 | 0.03504763 | 0.01403319 | 0.04382923 | 0.00722127 | 0.01850332 | 0.05822374 | 0.01766148 | 0.0064185 |
| 33.5694444 | 0.03993025 | 0.08541508 | 0.05700668 | 0.05628215 | 0.01928571 | 0.01854482 | 0.03804958 | 0.01987648 | 0.03901774 | 0.03548967 | 0.04663028 | 0.04773564 | 0.02042301 | 0.05739534 | 0.00769843 | 0.02017595 | 0.07108599 | 0.02260769 | 0.00777875 |
| 33.5833333 | 0.05922523 | 0.07864583 | 0.06280586 | 0.07052261 | 0.02509655 | 0.03143909 | 0.04845189 | 0.02342021 | 0.06000733 | 0.0377696 | 0.04107734 | 0.05483313 | 0.02305613 | 0.05843402 | 0.00508873 | 0.01596712 | 0.0739547 | 0.02006677 | 0.00589008 |
| 33.5972222 | 0.07189392 | 0.06716856 | 0.05789572 | 0.07287504 | 0.02720922 | 0.03991908 | 0.04613593 | 0.02372428 | 0.07603534 | 0.03172412 | 0.03377408 | 0.0543649 | 0.01898906 | 0.05351102 | 0.00273775 | 0.00932907 | 0.0737832 | 0.01445338 | 0.00409104 |
| 33.6111111 | 0.07570577 | 0.06715579 | 0.04724744 | 0.064609 | 0.03052523 | 0.0431027 | 0.03784287 | 0.02279661 | 0.07902624 | 0.01816617 | 0.02991247 | 0.04355433 | 0.01135048 | 0.04579635 | 0.00354962 | 0.00623148 | 0.06305236 | 0.00906071 | 0.00378357 |
| 33.625 | 0.07265092 | 0.0707111 | 0.03663621 | 0.0539082 | 0.03195096 | 0.04459786 | 0.02466876 | 0.02096684 | 0.06587356 | 0.00841968 | 0.02853203 | 0.02309564 | 0.00532297 | 0.03485258 | 0.00570753 | 0.01102236 | 0.0416092 | 0.00801987 | 0.00635392 |
| 33.6388889 | 0.06301891 | 0.07128773 | 0.02821552 | 0.04379554 | 0.03050688 | 0.04120427 | 0.01105036 | 0.01895997 | 0.04681431 | 0.00973373 | 0.02841659 | 0.00864776 | 0.00578287 | 0.02328655 | 0.00674379 | 0.01837481 | 0.02078479 | 0.01278192 | 0.01473797 |
| 33.6527778 | 0.04722743 | 0.06872501 | 0.0216026 | 0.03277135 | 0.02986419 | 0.0350756 | 0.00644745 | 0.01846788 | 0.030232 | 0.00999061 | 0.02751046 | 0.00981475 | 0.00913994 | 0.01310789 | 0.00499623 | 0.01981213 | 0.00834232 | 0.01547374 | 0.02417173 |
| 33.6666667 | 0.0289392 | 0.0590504 | 0.01666022 | 0.02286025 | 0.02768895 | 0.03159429 | 0.00570893 | 0.01784833 | 0.01594864 | 0.00486058 | 0.02459639 | 0.0153046 | 0.00740431 | 0.00688951 | 0.00256964 | 0.01290799 | 0.0062072 | 0.01472475 | 0.02863665 |
| 33.6805556 | 0.01373372 | 0.03799362 | 0.01313204 | 0.01494848 | 0.02031441 | 0.02844452 | 0.00515916 | 0.01233396 | 0.01045903 | 0.00184551 | 0.02026979 | 0.01827463 | 0.00311541 | 0.00852503 | 0.00272179 | 0.00456426 | 0.01123414 | 0.01332774 | 0.025208 |
| 33.6944444 | 0.01149438 | 0.01807136 | 0.01163587 | 0.0090619 | 0.01215946 | 0.02065508 | 0.00954556 | 0.0084543 | 0.02189005 | 0.00347448 | 0.01524188 | 0.01867422 | 0.00127813 | 0.0155483 | 0.00273346 | 0.00103775 | 0.0169828 | 0.0097627 | 0.01518757 |
| 33.7083333 | 0.02236568 | 0.0173506 | 0.01127366 | 0.00967651 | 0.01358357 | 0.01110011 | 0.01425095 | 0.01626101 | 0.04027596 | 0.00839106 | 0.00999216 | 0.01609103 | 0.00158068 | 0.02193221 | 0.00170732 | 0.00086145 | 0.02109401 | 0.00469555 | 0.00762839 |
| 33.7222222 | 0.03165748 | 0.03090816 | 0.01002057 | 0.01563094 | 0.02364986 | 0.00978258 | 0.01450955 | 0.03067755 | 0.05702008 | 0.01318597 | 0.00498861 | 0.01300393 | 0.00468898 | 0.02682729 | 0.00300139 | 0.00236716 | 0.02503962 | 0.00117197 | 0.01023252 |
| 33.7361111 | 0.03217919 | 0.04267799 | 0.00819495 | 0.01838801 | 0.03042386 | 0.01681281 | 0.01224055 | 0.04483254 | 0.07364543 | 0.01341574 | 0.00190434 | 0.0096093 | 0.0093969 | 0.03088657 | 0.00543668 | 0.00428896 | 0.02826386 | 0.00029283 | 0.01798663 |
| 33.75 | 0.02852532 | 0.04668208 | 0.00771388 | 0.0143148 | 0.02949866 | 0.02187256 | 0.00914288 | 0.0534842 | 0.08493196 | 0.00881626 | 0.00145468 | 0.00649598 | 0.01051508 | 0.03076003 | 0.0046443 | 0.00395991 | 0.02828456 | 0.00043135 | 0.02093719 |
| 33.7638889 | 0.02499974 | 0.04696586 | 0.00993542 | 0.00847981 | 0.02391164 | 0.02064468 | 0.00810222 | 0.04959775 | 0.08374877 | 0.00679285 | 0.00126278 | 0.00880165 | 0.00725193 | 0.02430307 | 0.00317847 | 0.00227652 | 0.02312765 | 0.00037088 | 0.01766949 |
| 33.7777778 | 0.02011154 | 0.04633261 | 0.01558446 | 0.01139963 | 0.01614666 | 0.0172862 | 0.0111037 | 0.03395747 | 0.06777066 | 0.01030108 | 0.00088661 | 0.01674739 | 0.00522342 | 0.01510585 | 0.00539903 | 0.002467 | 0.01322605 | 0.0003481 | 0.01220283 |
| 33.7916667 | 0.01480947 | 0.04145676 | 0.02311426 | 0.02368619 | 0.00840449 | 0.01412062 | 0.01543863 | 0.01598468 | 0.03925955 | 0.01271536 | 0.00119404 | 0.02342364 | 0.00600892 | 0.00694751 | 0.00804849 | 0.00415128 | 0.00623 | 0.00080846 | 0.00657428 |
| 33.8055556 | 0.01413433 | 0.03254577 | 0.02903595 | 0.03358215 | 0.00337571 | 0.00931712 | 0.01787489 | 0.00473096 | 0.02422054 | 0.01111519 | 0.00112643 | 0.02495911 | 0.00575801 | 0.00317012 | 0.00921056 | 0.00658891 | 0.00821531 | 0.00202728 | 0.00338074 |
| 33.8194444 | 0.01686952 | 0.02392112 | 0.03139065 | 0.03453902 | 0.00316889 | 0.00572491 | 0.01789283 | 0.0014446 | 0.04197036 | 0.00793192 | 0.0004838 | 0.0218054 | 0.0038504 | 0.00437837 | 0.01009958 | 0.00899133 | 0.01122356 | 0.00304812 | 0.00383451 |
| 33.8333333 | 0.02006991 | 0.0168384 | 0.02915791 | 0.02808289 | 0.00907074 | 0.00908222 | 0.01523121 | 0.00247357 | 0.06385427 | 0.00457241 | 0.00102966 | 0.01542746 | 0.00204103 | 0.00463264 | 0.0101389 | 0.0088808 | 0.00791036 | 0.00242536 | 0.00372832 |
| 33.8472222 | 0.02760899 | 0.0130725 | 0.02185149 | 0.01669019 | 0.02161616 | 0.016839 | 0.00973937 | 0.00507153 | 0.06670899 | 0.00414347 | 0.0034507 | 0.00812323 | 0.00113172 | 0.00258869 | 0.00936481 | 0.00620189 | 0.00488343 | 0.00191369 | 0.00303661 |
| 33.8611111 | 0.04664285 | 0.02247846 | 0.01273919 | 0.01298462 | 0.03961976 | 0.02227916 | 0.00746971 | 0.00702666 | 0.05551752 | 0.00919913 | 0.0062633 | 0.0057728 | 0.00163829 | 0.00157167 | 0.00732442 | 0.00338619 | 0.00885831 | 0.00355053 | 0.00383846 |
| 33.875 | 0.07452897 | 0.04953425 | 0.01133649 | 0.02749415 | 0.05997335 | 0.02319941 | 0.01233565 | 0.00552207 | 0.03327246 | 0.01708017 | 0.00942656 | 0.01266982 | 0.00283895 | 0.0018464 | 0.00397506 | 0.00174724 | 0.01578487 | 0.00549437 | 0.00341296 |
| 33.8888889 | 0.09866056 | 0.07858563 | 0.02085175 | 0.04686147 | 0.08086207 | 0.02102921 | 0.01534194 | 0.0042965 | 0.0184454 | 0.02466945 | 0.01533539 | 0.0236018 | 0.0027798 | 0.002985 | 0.00139317 | 0.00143452 | 0.0204442 | 0.00663898 | 0.00209557 |
| 33.9027778 | 0.10765793 | 0.0918756 | 0.03063776 | 0.05989676 | 0.10129969 | 0.01753069 | 0.01134054 | 0.00920267 | 0.03482547 | 0.0295045 | 0.02356859 | 0.03344584 | 0.00295444 | 0.00300263 | 0.0003843 | 0.00188354 | 0.02081487 | 0.00725378 | 0.00289327 |
| 33.9166667 | 0.10289895 | 0.09602863 | 0.03480232 | 0.06870076 | 0.11381142 | 0.0129555 | 0.00546164 | 0.01608066 | 0.07051379 | 0.02867072 | 0.02910506 | 0.03736118 | 0.00550153 | 0.00321311 | 0.00088423 | 0.00206154 | 0.01606707 | 0.00589475 | 0.00515294 |
| 33.9305556 | 0.09415168 | 0.10489607 | 0.03674494 | 0.072075 | 0.11063767 | 0.00715893 | 0.00215849 | 0.01942887 | 0.10351474 | 0.02182904 | 0.02888917 | 0.03005761 | 0.00739206 | 0.00750463 | 0.00199667 | 0.00202036 | 0.00832771 | 0.00374545 | 0.00646287 |
| 33.9444444 | 0.08010988 | 0.11362786 | 0.03621353 | 0.0674793 | 0.09284618 | 0.00376773 | 0.00200028 | 0.01901498 | 0.12345777 | 0.01222014 | 0.02522523 | 0.01674006 | 0.00663842 | 0.0137844 | 0.00335068 | 0.00156706 | 0.00350287 | 0.00427581 | 0.00551411 |
| 33.9583333 | 0.05942017 | 0.11099771 | 0.03019031 | 0.05593543 | 0.06390599 | 0.00664016 | 0.00334132 | 0.0167091 | 0.12644079 | 0.00557284 | 0.02116429 | 0.00827628 | 0.00455366 | 0.01779024 | 0.00567736 | 0.00079198 | 0.00507503 | 0.00655214 | 0.00298433 |
| 33.9722222 | 0.03875407 | 0.09684664 | 0.01930668 | 0.0388417 | 0.03350667 | 0.01033961 | 0.00423894 | 0.01440721 | 0.1200226 | 0.00583428 | 0.01703187 | 0.01144976 | 0.00308945 | 0.01746363 | 0.00568462 | 0.00111159 | 0.00898706 | 0.00808415 | 0.00112189 |
| 33.9861111 | 0.01969899 | 0.08017443 | 0.01019814 | 0.02113041 | 0.0267897 | 0.00790283 | 0.00461886 | 0.01112056 | 0.10825754 | 0.01030251 | 0.01299179 | 0.02182171 | 0.00421159 | 0.01384258 | 0.00331838 | 0.00223234 | 0.01173634 | 0.00876204 | 0.00181002 |
| 34 | 0.01021432 | 0.06424025 | 0.01130482 | 0.01772445 | 0.04995356 | 0.00691324 | 0.00850391 | 0.00641562 | 0.0866144 | 0.01389389 | 0.00956889 | 0.0294482 | 0.0060786 | 0.00992155 | 0.00315822 | 0.00228096 | 0.01287321 | 0.00883928 | 0.00486305 |
| 34.0138889 | 0.01614419 | 0.04776993 | 0.01986186 | 0.0329762 | 0.06530733 | 0.0174882 | 0.0173588 | 0.00463069 | 0.05854888 | 0.01425264 | 0.00683866 | 0.03035957 | 0.0053085 | 0.00791578 | 0.00340809 | 0.0031189 | 0.01119011 | 0.00770935 | 0.0076538 |
| 34.0277778 | 0.01950597 | 0.03743371 | 0.02204514 | 0.04691554 | 0.04939404 | 0.026171 | 0.02345846 | 0.00493832 | 0.03941284 | 0.011162 | 0.00876458 | 0.02148384 | 0.00318204 | 0.01073133 | 0.00410658 | 0.01032673 | 0.01036631 | 0.00803761 | 0.00688989 |
| 34.0416667 | 0.02095136 | 0.04851384 | 0.02122148 | 0.0399567 | 0.0249192 | 0.02747494 | 0.01945543 | 0.00786496 | 0.052352 | 0.01800544 | 0.02408059 | 0.01982043 | 0.00806725 | 0.02466058 | 0.01165411 | 0.0288375 | 0.02523598 | 0.02295199 | 0.00943983 |
| 34.0555556 | 0.05004824 | 0.08846646 | 0.03577464 | 0.03930071 | 0.02456552 | 0.03764363 | 0.01721352 | 0.02042512 | 0.09425219 | 0.04733027 | 0.05440742 | 0.04755632 | 0.02469975 | 0.05030538 | 0.02347133 | 0.05322618 | 0.05888374 | 0.05617232 | 0.02290702 |
| 34.0694444 | 0.09449717 | 0.13642931 | 0.05734366 | 0.06970693 | 0.05110946 | 0.04933448 | 0.02471868 | 0.03350558 | 0.12597382 | 0.07344241 | 0.07024096 | 0.0702945 | 0.03500541 | 0.06425389 | 0.02592988 | 0.05875579 | 0.07673993 | 0.07336652 | 0.03070903 |
| 34.0833333 | 0.11338629 | 0.15252959 | 0.06706728 | 0.08428503 | 0.07262278 | 0.05074154 | 0.02709496 | 0.03538629 | 0.13403229 | 0.06510059 | 0.04713361 | 0.04964972 | 0.0250203 | 0.04424072 | 0.01676708 | 0.03622286 | 0.05310651 | 0.04890825 | 0.02152793 |
| 34.0972222 | 0.10913884 | 0.12962525 | 0.06387174 | 0.06411377 | 0.07335077 | 0.04701136 | 0.02085742 | 0.03104692 | 0.12993793 | 0.03320545 | 0.01839681 | 0.02019555 | 0.01521813 | 0.01759808 | 0.01027616 | 0.01688251 | 0.02640287 | 0.02233148 | 0.01411166 |
| 34.1111111 | 0.09914668 | 0.1000477 | 0.05128544 | 0.04218717 | 0.06156506 | 0.04012088 | 0.01289488 | 0.02784309 | 0.11474513 | 0.01517878 | 0.01451729 | 0.01980753 | 0.01808515 | 0.0149047 | 0.0115522 | 0.01651596 | 0.02964378 | 0.02602687 | 0.01732101 |
| 34.125 | 0.07869466 | 0.07939546 | 0.03497442 | 0.02632832 | 0.04145782 | 0.03284376 | 0.00602172 | 0.02517435 | 0.09094077 | 0.02534858 | 0.02100332 | 0.03198237 | 0.02410284 | 0.02090495 | 0.01528898 | 0.01816797 | 0.03905827 | 0.03639307 | 0.01984712 |
| 34.1388889 | 0.04875475 | 0.0629967 | 0.02093591 | 0.01216523 | 0.01942989 | 0.02962138 | 0.00558181 | 0.02025403 | 0.06477121 | 0.03836976 | 0.02276058 | 0.03906236 | 0.03362021 | 0.02145358 | 0.01925448 | 0.01613146 | 0.03926126 | 0.03896048 | 0.01739558 |
| 34.1527778 | 0.02288819 | 0.04737283 | 0.01129176 | 0.00404232 | 0.0080041 | 0.02915365 | 0.01168052 | 0.01387093 | 0.04166519 | 0.03707345 | 0.02008499 | 0.03822569 | 0.04050653 | 0.02008634 | 0.02048815 | 0.01448819 | 0.03558036 | 0.03794032 | 0.01248574 |
| 34.1666667 | 0.00879188 | 0.03277048 | 0.00693653 | 0.00531296 | 0.01231982 | 0.02684399 | 0.01808313 | 0.00790835 | 0.02444948 | 0.02940097 | 0.01825908 | 0.02875232 | 0.03322201 | 0.02029661 | 0.01864077 | 0.01401908 | 0.03154299 | 0.03354022 | 0.00822755 |
| 34.1805556 | 0.00801885 | 0.02043326 | 0.00944684 | 0.01218108 | 0.0219519 | 0.02051044 | 0.02209364 | 0.00383675 | 0.01154332 | 0.0236404 | 0.01974453 | 0.01798547 | 0.02172095 | 0.02309803 | 0.01687667 | 0.01468639 | 0.03067086 | 0.02862274 | 0.00654402 |
| 34.1944444 | 0.02022381 | 0.00992425 | 0.01630281 | 0.02118615 | 0.02360523 | 0.01117213 | 0.02277085 | 0.0022094 | 0.00474149 | 0.02292404 | 0.02024597 | 0.01011058 | 0.01592621 | 0.02646551 | 0.01403239 | 0.01396185 | 0.03122753 | 0.02472233 | 0.00541781 |
| 34.2083333 | 0.03744824 | 0.00361263 | 0.02254926 | 0.03171384 | 0.01792603 | 0.00683819 | 0.0207222 | 0.00176653 | 0.00728101 | 0.02755796 | 0.01662302 | 0.00584989 | 0.01142684 | 0.02956767 | 0.00935487 | 0.01064606 | 0.02916038 | 0.01920108 | 0.00351193 |
| 34.2222222 | 0.04842937 | 0.00517602 | 0.02849498 | 0.04132834 | 0.01743392 | 0.01615655 | 0.02160227 | 0.00254704 | 0.01579025 | 0.03646509 | 0.01255301 | 0.01019543 | 0.00569523 | 0.03080639 | 0.00576578 | 0.00800381 | 0.02519401 | 0.01088136 | 0.00306306 |
| 34.2361111 | 0.04938721 | 0.01348945 | 0.03380503 | 0.04865371 | 0.02539972 | 0.03498832 | 0.02828215 | 0.00607917 | 0.02657738 | 0.04566394 | 0.0104844 | 0.02164673 | 0.00179204 | 0.02743714 | 0.00408364 | 0.00782456 | 0.02236318 | 0.00530053 | 0.0048454 |
| 34.25 | 0.04343456 | 0.019731 | 0.0343105 | 0.05528612 | 0.03363113 | 0.05922264 | 0.03521972 | 0.00901032 | 0.03505388 | 0.04701518 | 0.0073168 | 0.02862438 | 0.00079751 | 0.01884228 | 0.00255611 | 0.00719452 | 0.01925443 | 0.00692136 | 0.00572894 |
| 34.2638889 | 0.0334422 | 0.0165907 | 0.02916713 | 0.05974428 | 0.03490319 | 0.08387026 | 0.03582154 | 0.0077871 | 0.03502702 | 0.03604336 | 0.004279 | 0.02418698 | 0.00058472 | 0.00960731 | 0.0010318 | 0.0042798 | 0.01389646 | 0.012037 | 0.00455283 |
| 34.2777778 | 0.02086048 | 0.01426153 | 0.02252676 | 0.05496631 | 0.02508872 | 0.09461173 | 0.0304693 | 0.00469212 | 0.03057992 | 0.01852953 | 0.00482836 | 0.0124161 | 0.00096032 | 0.00734497 | 0.00097506 | 0.00229735 | 0.00758933 | 0.01689176 | 0.00274746 |
| 34.2916667 | 0.0128036 | 0.03666737 | 0.01776714 | 0.04010327 | 0.01710413 | 0.0845423 | 0.02190821 | 0.00306275 | 0.03176848 | 0.00627698 | 0.00579776 | 0.00449813 | 0.00309173 | 0.00951495 | 0.00192066 | 0.0028766 | 0.00290583 | 0.01853841 | 0.00226812 |
| 34.3055556 | 0.01505769 | 0.07687175 | 0.01343171 | 0.02296792 | 0.02865773 | 0.06263534 | 0.01238986 | 0.00396708 | 0.04066025 | 0.00355397 | 0.00524485 | 0.00470441 | 0.00612342 | 0.00847898 | 0.00292483 | 0.00290037 | 0.00121855 | 0.01484625 | 0.00463183 |
| 34.3194444 | 0.01623881 | 0.09041631 | 0.00798065 | 0.01084574 | 0.04443156 | 0.04300041 | 0.00656398 | 0.00649545 | 0.05093332 | 0.00362209 | 0.00524034 | 0.00585066 | 0.00772467 | 0.00560236 | 0.00377868 | 0.00249098 | 0.00151998 | 0.0086519 | 0.00749119 |
| 34.3333333 | 0.00963765 | 0.06899851 | 0.00520323 | 0.00646058 | 0.0452941 | 0.02893879 | 0.00623523 | 0.00881048 | 0.05523432 | 0.00348952 | 0.00700758 | 0.00369738 | 0.00764779 | 0.00384778 | 0.0036672 | 0.00247083 | 0.00239839 | 0.00368394 | 0.00775065 |
| 34.3472222 | 0.00516264 | 0.04923177 | 0.00583041 | 0.00516006 | 0.04019991 | 0.01576857 | 0.00790465 | 0.00968213 | 0.05322213 | 0.00539168 | 0.00978321 | 0.0014853 | 0.00804148 | 0.00269897 | 0.00302721 | 0.0016728 | 0.00223291 | 0.00220941 | 0.00669768 |
| 34.3611111 | 0.00440573 | 0.04493406 | 0.00580757 | 0.00592955 | 0.03752257 | 0.01007979 | 0.0091684 | 0.00925799 | 0.05028359 | 0.00843581 | 0.01095594 | 0.00211955 | 0.01055469 | 0.00381264 | 0.00363254 | 0.00223968 | 0.00197839 | 0.00403156 | 0.00664217 |
| 34.375 | 0.00249101 | 0.04392197 | 0.0050409 | 0.00897982 | 0.0344562 | 0.02339229 | 0.00930765 | 0.00785297 | 0.04935028 | 0.00997043 | 0.00906794 | 0.00334663 | 0.01426401 | 0.00818225 | 0.0052091 | 0.00514464 | 0.0041472 | 0.00644085 | 0.00705012 |
| 34.3888889 | 0.00088146 | 0.04009863 | 0.00453117 | 0.0085635 | 0.03066961 | 0.04764328 | 0.00746801 | 0.00521223 | 0.05155217 | 0.00789561 | 0.00528173 | 0.00408564 | 0.01655984 | 0.01279407 | 0.00521237 | 0.00742096 | 0.00720412 | 0.00855949 | 0.00672009 |
| 34.4027778 | 0.003588 | 0.0383195 | 0.00363348 | 0.00589859 | 0.02881206 | 0.06513039 | 0.00496536 | 0.0025642 | 0.05693627 | 0.00559842 | 0.0043155 | 0.00879922 | 0.0175494 | 0.0153756 | 0.00350806 | 0.00783086 | 0.00945305 | 0.0115767 | 0.00647807 |
| 34.4166667 | 0.01279689 | 0.04709755 | 0.00325268 | 0.00836574 | 0.03292448 | 0.06776267 | 0.00449732 | 0.00226287 | 0.06478107 | 0.00954255 | 0.0086813 | 0.01719274 | 0.0177363 | 0.01575063 | 0.00263295 | 0.00763344 | 0.01066333 | 0.01521992 | 0.00751092 |
| 34.4305556 | 0.02201022 | 0.0599251 | 0.00581813 | 0.01494122 | 0.04099633 | 0.05513628 | 0.00505979 | 0.00403308 | 0.07035435 | 0.01654776 | 0.01266801 | 0.02228934 | 0.01400726 | 0.01275165 | 0.00235222 | 0.00663209 | 0.00974973 | 0.0169705 | 0.00829491 |
| 34.4444444 | 0.01960352 | 0.0566149 | 0.00976619 | 0.01483657 | 0.03966682 | 0.03304196 | 0.00563119 | 0.00455635 | 0.065355 | 0.01886715 | 0.01194702 | 0.02062298 | 0.00849922 | 0.00742451 | 0.00151711 | 0.00411748 | 0.0070189 | 0.01450661 | 0.00708077 |
| 34.4583333 | 0.00866197 | 0.03750593 | 0.01063072 | 0.00724893 | 0.02995193 | 0.01387798 | 0.00959701 | 0.00259031 | 0.05404737 | 0.01734691 | 0.00954606 | 0.01573606 | 0.00775276 | 0.00497342 | 0.00102983 | 0.00230084 | 0.00485616 | 0.00932995 | 0.00540807 |
| 34.4722222 | 0.00388556 | 0.02272488 | 0.00733237 | 0.00260743 | 0.03076536 | 0.00763789 | 0.01483091 | 0.00124711 | 0.04669922 | 0.01681801 | 0.00900983 | 0.01288236 | 0.00835138 | 0.00568464 | 0.00134068 | 0.00307001 | 0.00436215 | 0.00473489 | 0.00460725 |
| 34.4861111 | 0.00828383 | 0.01765998 | 0.0041725 | 0.00386093 | 0.04201701 | 0.01305313 | 0.01509146 | 0.00299891 | 0.03995405 | 0.01804776 | 0.00963119 | 0.01220682 | 0.005194 | 0.00475956 | 0.00211196 | 0.00457821 | 0.00495888 | 0.00226318 | 0.00344435 |
| 34.5 | 0.01107259 | 0.01509086 | 0.00458386 | 0.00627109 | 0.0441249 | 0.02088831 | 0.01158612 | 0.00603517 | 0.02937587 | 0.02047369 | 0.00944314 | 0.01228988 | 0.0025 | 0.00326908 | 0.0028518 | 0.0052223 | 0.00689853 | 0.00160739 | 0.00161212 |
| 34.5138889 | 0.00715396 | 0.01085749 | 0.00519718 | 0.00697728 | 0.03026234 | 0.02194466 | 0.01021263 | 0.00680073 | 0.02098135 | 0.02397785 | 0.00866956 | 0.01468489 | 0.00358122 | 0.00578123 | 0.0042901 | 0.00585843 | 0.01050264 | 0.00167817 | 0.0008304 |
| 34.5277778 | 0.00653649 | 0.00896514 | 0.00839747 | 0.00948499 | 0.012874 | 0.01450331 | 0.01412472 | 0.00498759 | 0.02001927 | 0.02739083 | 0.00965962 | 0.01999888 | 0.00510925 | 0.01240501 | 0.00638636 | 0.00750562 | 0.01505654 | 0.00370584 | 0.00112667 |
| 34.5416667 | 0.01725382 | 0.02129514 | 0.0220555 | 0.01925699 | 0.00724853 | 0.00971331 | 0.02291089 | 0.00284973 | 0.02991774 | 0.03161078 | 0.01769197 | 0.02938426 | 0.00489653 | 0.02361837 | 0.00741757 | 0.01087068 | 0.0225607 | 0.01046257 | 0.00244901 |
| 34.5555556 | 0.03538381 | 0.04834603 | 0.04478293 | 0.03783036 | 0.01738237 | 0.02377085 | 0.03536581 | 0.00373258 | 0.05352901 | 0.03760833 | 0.0321947 | 0.04129373 | 0.0061379 | 0.04107347 | 0.00786476 | 0.01561791 | 0.03890702 | 0.0217327 | 0.00708717 |
| 34.5694444 | 0.0562793 | 0.07754194 | 0.06984192 | 0.06076351 | 0.03600926 | 0.05124537 | 0.0492958 | 0.00982752 | 0.0813993 | 0.0409285 | 0.04111035 | 0.04705614 | 0.01106228 | 0.05670809 | 0.00916226 | 0.01851937 | 0.0553162 | 0.03160356 | 0.01257747 |
| 34.5833333 | 0.07638729 | 0.1016799 | 0.08833351 | 0.08017802 | 0.0524563 | 0.07018377 | 0.06013422 | 0.01588278 | 0.10127714 | 0.03842609 | 0.03870556 | 0.04413906 | 0.01590462 | 0.06017049 | 0.01081874 | 0.01639333 | 0.05455365 | 0.03307344 | 0.01341909 |
| 34.5972222 | 0.08727667 | 0.12396436 | 0.08898372 | 0.08883297 | 0.05924451 | 0.07496962 | 0.0620771 | 0.01576376 | 0.11226145 | 0.03224016 | 0.0332058 | 0.03876911 | 0.01574914 | 0.05453116 | 0.01112577 | 0.01091136 | 0.04281985 | 0.02691031 | 0.00961836 |
| 34.6111111 | 0.08161065 | 0.1451907 | 0.0711164 | 0.07981853 | 0.05690795 | 0.0754601 | 0.0527422 | 0.01330638 | 0.1185645 | 0.02315465 | 0.03288259 | 0.0333107 | 0.01123517 | 0.04630893 | 0.00777562 | 0.00542121 | 0.03233618 | 0.01740805 | 0.00520356 |
| 34.625 | 0.06278157 | 0.1571487 | 0.04549431 | 0.05402738 | 0.04872309 | 0.07592248 | 0.03640292 | 0.01501135 | 0.12252644 | 0.01239639 | 0.03642252 | 0.02533738 | 0.00596232 | 0.03749881 | 0.00494068 | 0.00304685 | 0.021675 | 0.01113006 | 0.00446386 |
| 34.6388889 | 0.04421456 | 0.15428322 | 0.02109123 | 0.02484276 | 0.03624069 | 0.07101254 | 0.02134791 | 0.01870141 | 0.12441361 | 0.00739294 | 0.03754902 | 0.01412181 | 0.00277468 | 0.03155914 | 0.00865394 | 0.00640319 | 0.01167983 | 0.01810522 | 0.00529189 |
| 34.6527778 | 0.03586327 | 0.13681994 | 0.00693304 | 0.00897377 | 0.02543897 | 0.06134192 | 0.01510833 | 0.0177091 | 0.1201727 | 0.01013067 | 0.03418653 | 0.00579181 | 0.00234024 | 0.03006878 | 0.01290238 | 0.01021853 | 0.00812183 | 0.03150741 | 0.00435849 |
| 34.6666667 | 0.031272 | 0.1084015 | 0.00630926 | 0.00887259 | 0.02046509 | 0.05046873 | 0.01896734 | 0.01057272 | 0.09955907 | 0.01230659 | 0.02922288 | 0.00505046 | 0.00190494 | 0.03018113 | 0.01176323 | 0.00792016 | 0.01029647 | 0.03510115 | 0.00533084 |
| 34.6805556 | 0.01977995 | 0.07054261 | 0.01028937 | 0.01620818 | 0.01472609 | 0.03237907 | 0.02663145 | 0.00533641 | 0.06280118 | 0.01251245 | 0.02595894 | 0.0080886 | 0.00070065 | 0.0272547 | 0.00703297 | 0.00376016 | 0.01115458 | 0.03022069 | 0.00709023 |
| 34.6944444 | 0.01208036 | 0.03669295 | 0.01541923 | 0.02879268 | 0.01492487 | 0.0251578 | 0.03081566 | 0.00867325 | 0.03160227 | 0.01458299 | 0.02427069 | 0.01236864 | 0.00106024 | 0.01808124 | 0.00309001 | 0.00254488 | 0.00814677 | 0.02554678 | 0.00567403 |
| 34.7083333 | 0.02254225 | 0.03557295 | 0.02244079 | 0.03580001 | 0.03269224 | 0.04983126 | 0.03293559 | 0.01576071 | 0.03279575 | 0.01681592 | 0.02250107 | 0.01653651 | 0.00350121 | 0.00847074 | 0.00192917 | 0.00187804 | 0.0044911 | 0.02297991 | 0.0047483 |
| 34.7222222 | 0.0389919 | 0.06225413 | 0.0267174 | 0.02726724 | 0.05432362 | 0.07889896 | 0.03779767 | 0.02195399 | 0.06495436 | 0.01483736 | 0.02153067 | 0.01743004 | 0.00471223 | 0.00623547 | 0.00277345 | 0.00122636 | 0.00418948 | 0.02044736 | 0.00595473 |

| | | | | | | | | | | | | | | | | | | | |
|---|---|---|---|---|---|---|---|---|---|---|---|---|---|---|---|---|---|---|---|
| 34.7361111 | 0.04561422 | 0.08483459 | 0.02543202 | 0.01442203 | 0.06862309 | 0.08809311 | 0.04249049 | 0.02634972 | 0.09824318 | 0.00927919 | 0.02274196 | 0.01473792 | 0.00368369 | 0.00874353 | 0.00776569 | 0.00258447 | 0.00773251 | 0.01575529 | 0.00508807 |
| 34.75 | 0.04217555 | 0.08995269 | 0.02024224 | 0.00649569 | 0.07268581 | 0.07871945 | 0.04456217 | 0.02627953 | 0.11357479 | 0.00835194 | 0.02349982 | 0.01049288 | 0.00443323 | 0.00924538 | 0.01464789 | 0.00516571 | 0.01102842 | 0.00881598 | 0.00361034 |
| 34.7638889 | 0.0327099 | 0.08050423 | 0.01335536 | 0.00390931 | 0.06100534 | 0.05576532 | 0.0461589 | 0.02246893 | 0.11616926 | 0.0158391 | 0.02039654 | 0.00603285 | 0.00655779 | 0.00613645 | 0.01690781 | 0.00608558 | 0.01079995 | 0.00588632 | 0.0038167 |
| 34.7777778 | 0.02097908 | 0.06367029 | 0.00853758 | 0.00694639 | 0.03835716 | 0.02783625 | 0.04733561 | 0.01907636 | 0.11429889 | 0.02536504 | 0.01395786 | 0.00347381 | 0.0080945 | 0.004669 | 0.01268797 | 0.00423239 | 0.00748203 | 0.01221718 | 0.00286491 |
| 34.7916667 | 0.01156608 | 0.0460161 | 0.01342761 | 0.01394192 | 0.01681793 | 0.01204221 | 0.04753402 | 0.01626569 | 0.1034397 | 0.02977446 | 0.0068476 | 0.00582775 | 0.01088367 | 0.00696704 | 0.00668101 | 0.00211811 | 0.00401797 | 0.01950811 | 0.00112702 |
| 34.8055556 | 0.005803 | 0.03063118 | 0.0274199 | 0.02166411 | 0.00543815 | 0.01702193 | 0.04654873 | 0.01188378 | 0.07887827 | 0.02706767 | 0.00386442 | 0.01126723 | 0.01368346 | 0.00756352 | 0.00549811 | 0.00171243 | 0.00280758 | 0.01701776 | 0.00107855 |
| 34.8194444 | 0.00234876 | 0.01779898 | 0.03801894 | 0.02471224 | 0.00246001 | 0.02221195 | 0.04397219 | 0.0079569 | 0.04924725 | 0.01799978 | 0.00790285 | 0.01433542 | 0.01296426 | 0.0049484 | 0.00818227 | 0.00233034 | 0.0034068 | 0.01080948 | 0.00215895 |
| 34.8333333 | 0.00089299 | 0.00819103 | 0.03764708 | 0.02175912 | 0.00253914 | 0.01451892 | 0.04067036 | 0.00863866 | 0.02482601 | 0.01033766 | 0.01483436 | 0.01341388 | 0.00923771 | 0.00357464 | 0.00876314 | 0.00313661 | 0.00425226 | 0.01408034 | 0.00258855 |
| 34.8472222 | 0.00086402 | 0.00459037 | 0.03012255 | 0.01626358 | 0.0061215 | 0.00788359 | 0.03634419 | 0.01292784 | 0.01061092 | 0.01311514 | 0.01972845 | 0.01079893 | 0.00577205 | 0.00425801 | 0.00583185 | 0.00396478 | 0.00469736 | 0.0219242 | 0.00451603 |
| 34.8611111 | 0.00182649 | 0.00613976 | 0.02004956 | 0.00897496 | 0.01395746 | 0.01428908 | 0.02952708 | 0.01615925 | 0.00916658 | 0.0201705 | 0.02222 | 0.00851528 | 0.00427511 | 0.00329211 | 0.00298433 | 0.00442059 | 0.00404734 | 0.02315651 | 0.01045745 |
| 34.875 | 0.00210577 | 0.00671031 | 0.01061844 | 0.00379738 | 0.02507284 | 0.02762451 | 0.02058802 | 0.01746209 | 0.01404047 | 0.02428488 | 0.02500958 | 0.00599797 | 0.00423278 | 0.00189856 | 0.0041742 | 0.00384668 | 0.00227996 | 0.01857101 | 0.01564511 |
| 34.8888889 | 0.00272089 | 0.00548278 | 0.01016144 | 0.00565316 | 0.03540481 | 0.03958873 | 0.01068415 | 0.0191302 | 0.01348986 | 0.02780645 | 0.03093599 | 0.00380282 | 0.00462574 | 0.0036002 | 0.00816075 | 0.00271826 | 0.00092185 | 0.01262856 | 0.01444179 |
| 34.9027778 | 0.00870247 | 0.00968732 | 0.02066126 | 0.01059056 | 0.04052864 | 0.04980874 | 0.00425775 | 0.02159794 | 0.01136506 | 0.03076867 | 0.03994838 | 0.00434211 | 0.00544713 | 0.0065022 | 0.01008323 | 0.00257495 | 0.00067424 | 0.00685543 | 0.00917686 |
| 34.9166667 | 0.01859773 | 0.02221915 | 0.03133198 | 0.01418917 | 0.04625495 | 0.06123697 | 0.00425777 | 0.02279187 | 0.0229543 | 0.0304752 | 0.04867661 | 0.00561694 | 0.00673787 | 0.00831999 | 0.00791924 | 0.00387711 | 0.0008621 | 0.00397125 | 0.00573536 |
| 34.9305556 | 0.02695775 | 0.0346053 | 0.03569508 | 0.01537199 | 0.05909923 | 0.07493154 | 0.00513885 | 0.02117265 | 0.04506033 | 0.02627086 | 0.05340857 | 0.00426688 | 0.00796672 | 0.00987343 | 0.00409298 | 0.00576476 | 0.00131052 | 0.00685991 | 0.00599684 |
| 34.9444444 | 0.03327222 | 0.03750179 | 0.03809542 | 0.0146997 | 0.072849 | 0.08278797 | 0.00461326 | 0.01764201 | 0.06479827 | 0.01997863 | 0.05222492 | 0.00268749 | 0.00934377 | 0.01171253 | 0.00212367 | 0.00707157 | 0.00247712 | 0.00943487 | 0.00736195 |
| 34.9583333 | 0.03508217 | 0.02958135 | 0.04331817 | 0.0152486 | 0.07671437 | 0.0790214 | 0.005684 | 0.01381527 | 0.07951334 | 0.01451884 | 0.04724934 | 0.00365477 | 0.01075313 | 0.01292811 | 0.00200943 | 0.00717403 | 0.0041015 | 0.0063593 | 0.00698287 |
| 34.9722222 | 0.02689451 | 0.01712101 | 0.04317706 | 0.01525753 | 0.06192801 | 0.06901771 | 0.00696625 | 0.00905041 | 0.08754062 | 0.01080678 | 0.04192244 | 0.00452651 | 0.01114976 | 0.01276421 | 0.00199727 | 0.0066171 | 0.00541029 | 0.00237772 | 0.00602214 |
| 34.9861111 | 0.01463695 | 0.00820106 | 0.02762686 | 0.01061852 | 0.03957582 | 0.05511697 | 0.00564245 | 0.00775042 | 0.08302923 | 0.00727773 | 0.03691716 | 0.00478562 | 0.01053206 | 0.01028297 | 0.0019004 | 0.00636828 | 0.00563919 | 0.00379218 | 0.00649274 |
| 35 | 0.01313205 | 0.00791643 | 0.01287852 | 0.00629628 | 0.04695506 | 0.03814477 | 0.00599736 | 0.01541533 | 0.06553432 | 0.00372913 | 0.03120207 | 0.00670369 | 0.00888553 | 0.00673462 | 0.00233297 | 0.00647469 | 0.00442081 | 0.0112383 | 0.0073189 |
| 35.0138889 | 0.0221599 | 0.01861833 | 0.01860246 | 0.00924272 | 0.09023498 | 0.02319361 | 0.01264631 | 0.02587187 | 0.04524911 | 0.00150265 | 0.0250046 | 0.00942026 | 0.00543957 | 0.00837917 | 0.00395571 | 0.00569005 | 0.00230831 | 0.01805682 | 0.00698033 |
| 35.0277778 | 0.02600644 | 0.02936426 | 0.03378445 | 0.01392018 | 0.12274946 | 0.01481373 | 0.02031297 | 0.03125421 | 0.0322975 | 0.00443576 | 0.02469637 | 0.00839828 | 0.00350098 | 0.01674812 | 0.0050217 | 0.00479179 | 0.00266301 | 0.0149835 | 0.00488641 |
| 35.0416667 | 0.0182451 | 0.02768367 | 0.03577659 | 0.01219198 | 0.11443929 | 0.01862844 | 0.02002827 | 0.02669459 | 0.0358104 | 0.02029166 | 0.03926171 | 0.01117133 | 0.00733655 | 0.03354262 | 0.005814 | 0.01121114 | 0.01364987 | 0.01735858 | 0.00744494 |
| 35.0555556 | 0.01429214 | 0.03407651 | 0.02882896 | 0.01159574 | 0.08259378 | 0.03574484 | 0.012327 | 0.01745438 | 0.06008683 | 0.04937549 | 0.07140421 | 0.03342151 | 0.01744796 | 0.06644438 | 0.01319779 | 0.02855724 | 0.03236861 | 0.04855572 | 0.02184789 |
| 35.0694444 | 0.02393131 | 0.06273538 | 0.03616089 | 0.01754825 | 0.05068365 | 0.05066416 | 0.0080894 | 0.01929262 | 0.07956371 | 0.07012989 | 0.09311033 | 0.05420569 | 0.0284545 | 0.09152325 | 0.02205628 | 0.03947679 | 0.04181379 | 0.0780481 | 0.03574869 |
| 35.0833333 | 0.03291625 | 0.0857293 | 0.05159787 | 0.01761399 | 0.0251445 | 0.04855183 | 0.00823917 | 0.02802117 | 0.07463527 | 0.05831532 | 0.06785051 | 0.04068407 | 0.02691524 | 0.07401628 | 0.01743511 | 0.02822518 | 0.03278206 | 0.06541674 | 0.03094635 |
| 35.0972222 | 0.03639342 | 0.09242897 | 0.05411022 | 0.0101677 | 0.01302881 | 0.03842868 | 0.00642713 | 0.02794085 | 0.06755077 | 0.02629822 | 0.02631636 | 0.01733399 | 0.01538583 | 0.03310444 | 0.0063726 | 0.01392738 | 0.01626664 | 0.03451849 | 0.01531441 |
| 35.1111111 | 0.03834228 | 0.09682138 | 0.04646013 | 0.00649739 | 0.02006744 | 0.03367505 | 0.00510093 | 0.02107754 | 0.06784428 | 0.00740905 | 0.01214174 | 0.0156272 | 0.01400555 | 0.01014654 | 0.00222728 | 0.01399657 | 0.01192577 | 0.02972484 | 0.00610922 |
| 35.125 | 0.03585858 | 0.09846667 | 0.03861257 | 0.00987743 | 0.03141459 | 0.0353449 | 0.00718728 | 0.01391689 | 0.06358767 | 0.0103612 | 0.00949894 | 0.02112295 | 0.02503588 | 0.01147541 | 0.00202264 | 0.01563779 | 0.01837751 | 0.04689265 | 0.00251799 |
| 35.1388889 | 0.03104125 | 0.09235659 | 0.03014157 | 0.01616612 | 0.03397109 | 0.03918716 | 0.01070916 | 0.00747587 | 0.05730143 | 0.02081167 | 0.00479593 | 0.02715611 | 0.03577311 | 0.01808184 | 0.00384027 | 0.0124425 | 0.02258393 | 0.05880514 | 0.00188814 |
| 35.1527778 | 0.02686737 | 0.08024077 | 0.01890723 | 0.01789077 | 0.02820026 | 0.04044582 | 0.01126414 | 0.00467634 | 0.05694373 | 0.0262104 | 0.0036362 | 0.03740825 | 0.03890033 | 0.01969599 | 0.00780505 | 0.00947821 | 0.02092464 | 0.05948694 | 0.00503649 |
| 35.1666667 | 0.02183775 | 0.06477768 | 0.00857778 | 0.01248912 | 0.01581681 | 0.03822059 | 0.00703983 | 0.00741769 | 0.06170373 | 0.02373118 | 0.00376084 | 0.04044194 | 0.03487467 | 0.01820235 | 0.00997174 | 0.00643883 | 0.01494113 | 0.05460083 | 0.00721996 |
| 35.1805556 | 0.01510982 | 0.04892961 | 0.0045857 | 0.00568267 | 0.00979638 | 0.03402673 | 0.0035862 | 0.01187622 | 0.06327502 | 0.01698684 | 0.00379711 | 0.03112641 | 0.02748474 | 0.01561146 | 0.00853959 | 0.00357688 | 0.00904739 | 0.0449059 | 0.0052046 |
| 35.1944444 | 0.00861765 | 0.03397676 | 0.00773243 | 0.0053557 | 0.02519033 | 0.0290815 | 0.00736227 | 0.0119882 | 0.05419004 | 0.01048811 | 0.00339659 | 0.01816993 | 0.01914246 | 0.01436932 | 0.00600788 | 0.0024504 | 0.00583347 | 0.02903836 | 0.00220236 |
| 35.2083333 | 0.00939292 | 0.01871853 | 0.01526035 | 0.01566851 | 0.05370816 | 0.02228474 | 0.01866451 | 0.00811868 | 0.033898 | 0.00734404 | 0.00282487 | 0.01026238 | 0.01044221 | 0.01974714 | 0.00566373 | 0.00226671 | 0.00436041 | 0.01338593 | 0.00191008 |
| 35.2222222 | 0.02074866 | 0.01221656 | 0.02571425 | 0.03211381 | 0.07616151 | 0.01250154 | 0.0328049 | 0.00810874 | 0.01721935 | 0.00834551 | 0.00483866 | 0.00886576 | 0.00499354 | 0.03120523 | 0.00731727 | 0.00284618 | 0.00431462 | 0.0077626 | 0.00401165 |
| 35.2361111 | 0.0328363 | 0.02115188 | 0.03650126 | 0.04313715 | 0.07932642 | 0.00787509 | 0.04281822 | 0.01408905 | 0.01940611 | 0.01100119 | 0.00836667 | 0.01030534 | 0.00657005 | 0.03857586 | 0.00821942 | 0.00375685 | 0.00619346 | 0.01590108 | 0.00631816 |
| 35.25 | 0.03832554 | 0.03187411 | 0.04366983 | 0.04205762 | 0.06576575 | 0.011264 | 0.04476315 | 0.01867624 | 0.02879611 | 0.01203773 | 0.00977481 | 0.01071573 | 0.0101322 | 0.03538601 | 0.00674865 | 0.00329859 | 0.00863581 | 0.02983875 | 0.0063334 |
| 35.2638889 | 0.0403203 | 0.03370306 | 0.0468782 | 0.03551504 | 0.05524416 | 0.01131422 | 0.04106442 | 0.01902466 | 0.02815515 | 0.01020774 | 0.00807446 | 0.00937224 | 0.0101016 | 0.02291678 | 0.00449639 | 0.00181845 | 0.0094229 | 0.04221219 | 0.00491081 |
| 35.2777778 | 0.04264825 | 0.03064645 | 0.04636236 | 0.03402999 | 0.05669263 | 0.00641538 | 0.0361935 | 0.01659575 | 0.01693892 | 0.00733954 | 0.00496775 | 0.00619596 | 0.0069368 | 0.0112267 | 0.00405594 | 0.00091719 | 0.00781215 | 0.05009655 | 0.00446721 |
| 35.2916667 | 0.04396146 | 0.02716949 | 0.03979203 | 0.039612 | 0.06157967 | 0.0055417 | 0.0335406 | 0.01148461 | 0.0088325 | 0.00546025 | 0.00236529 | 0.00373077 | 0.00335375 | 0.01154445 | 0.00527782 | 0.00127217 | 0.00535593 | 0.04886634 | 0.00455332 |
| 35.3055556 | 0.0398081 | 0.02203319 | 0.02843791 | 0.04192242 | 0.05953828 | 0.01074007 | 0.02991087 | 0.00941179 | 0.01528885 | 0.00385146 | 0.00119234 | 0.00530083 | 0.00228976 | 0.01857732 | 0.00614214 | 0.00227589 | 0.00332564 | 0.04066001 | 0.00404539 |
| 35.3194444 | 0.03006371 | 0.01382011 | 0.01638663 | 0.02979967 | 0.04558429 | 0.01804999 | 0.01998682 | 0.01275261 | 0.0272703 | 0.0017976 | 0.00161798 | 0.00733211 | 0.00353642 | 0.02217018 | 0.00660536 | 0.00304685 | 0.00179965 | 0.03203417 | 0.0031476 |
| 35.3333333 | 0.02011499 | 0.00718727 | 0.00717213 | 0.01330073 | 0.02448312 | 0.02234087 | 0.0096979 | 0.0145463 | 0.03243702 | 0.00093692 | 0.00208826 | 0.00563196 | 0.00436098 | 0.01992087 | 0.00803458 | 0.00409874 | 0.00121154 | 0.02546758 | 0.00227324 |
| 35.3472222 | 0.01498324 | 0.00617729 | 0.00438291 | 0.00659542 | 0.01199694 | 0.0215908 | 0.00850508 | 0.01333259 | 0.02977315 | 0.00214125 | 0.00146037 | 0.00416384 | 0.00328188 | 0.01310784 | 0.0096282 | 0.00481453 | 0.00121431 | 0.01926312 | 0.00153091 |
| 35.3611111 | 0.01488827 | 0.00668771 | 0.00547673 | 0.00493564 | 0.01790415 | 0.01908538 | 0.0139212 | 0.01193741 | 0.02486217 | 0.00347152 | 0.00077312 | 0.0055956 | 0.00150115 | 0.00610739 | 0.0089091 | 0.00343382 | 0.00142652 | 0.01196708 | 0.00124311 |
| 35.375 | 0.01604926 | 0.00459992 | 0.00525982 | 0.00577423 | 0.03250744 | 0.01706897 | 0.01295947 | 0.00966976 | 0.02482462 | 0.00299409 | 0.00148215 | 0.00644225 | 0.00070703 | 0.00266267 | 0.00555362 | 0.00188797 | 0.00167885 | 0.005043 | 0.00099921 |
| 35.3888889 | 0.01598466 | 0.00340727 | 0.00638801 | 0.00953485 | 0.04407795 | 0.01464852 | 0.00521555 | 0.00648504 | 0.03025955 | 0.00167667 | 0.00361434 | 0.00581755 | 0.00107682 | 0.00211258 | 0.00220081 | 0.00206002 | 0.00122467 | 0.00121553 | 0.00079072 |
| 35.4027778 | 0.01586223 | 0.00534782 | 0.01150466 | 0.01038497 | 0.05103539 | 0.01197957 | 0.00157301 | 0.00432899 | 0.0373815 | 0.00116956 | 0.00548311 | 0.00557886 | 0.00270506 | 0.00171419 | 0.00068154 | 0.00204771 | 0.00110528 | 0.00088019 | 0.00134942 |
| 35.4166667 | 0.01669562 | 0.0073449 | 0.01584108 | 0.00773351 | 0.05447181 | 0.01117624 | 0.00107999 | 0.00386125 | 0.04284883 | 0.001332 | 0.00529602 | 0.00646614 | 0.00465773 | 0.00168595 | 0.00057387 | 0.00122833 | 0.00314281 | 0.00284747 | 0.0019234 |
| 35.4305556 | 0.01699675 | 0.00909012 | 0.01667235 | 0.00408196 | 0.05003211 | 0.01361884 | 0.00137478 | 0.00356687 | 0.04710778 | 0.00241099 | 0.00347747 | 0.0081414 | 0.00525601 | 0.00315738 | 0.00091924 | 0.00063489 | 0.00616938 | 0.00588428 | 0.00179611 |
| 35.4444444 | 0.01592113 | 0.01122695 | 0.01665071 | 0.00320254 | 0.03613825 | 0.01638185 | 0.00288837 | 0.00220252 | 0.05541696 | 0.00453775 | 0.00212092 | 0.0094633 | 0.00404015 | 0.00457616 | 0.00084944 | 0.00042587 | 0.00757071 | 0.00779377 | 0.00122473 |
| 35.4583333 | 0.01498669 | 0.01194378 | 0.01740604 | 0.00848209 | 0.01876603 | 0.01447622 | 0.00439439 | 0.00073008 | 0.0652678 | 0.00617789 | 0.00174079 | 0.00928967 | 0.00219076 | 0.00580841 | 0.00041326 | 0.00047236 | 0.00701512 | 0.00746752 | 0.00076147 |
| 35.4722222 | 0.01475956 | 0.01160413 | 0.01696146 | 0.01669741 | 0.00757942 | 0.0096725 | 0.00867072 | 0.00210332 | 0.07400945 | 0.00631988 | 0.00145016 | 0.00807866 | 0.00159694 | 0.00744118 | 0.00099401 | 0.0006706 | 0.00611745 | 0.00616479 | 0.00081415 |
| 35.4861111 | 0.01358012 | 0.01210704 | 0.01302311 | 0.02210425 | 0.0060873 | 0.00851478 | 0.01476957 | 0.00683469 | 0.08409039 | 0.00604194 | 0.00112273 | 0.00752728 | 0.00236099 | 0.00787312 | 0.00296168 | 0.0014545 | 0.00619682 | 0.00538291 | 0.00105454 |
| 35.5 | 0.01016344 | 0.01352742 | 0.00709037 | 0.02259972 | 0.00612623 | 0.01261264 | 0.0178688 | 0.01086634 | 0.08719476 | 0.00658203 | 0.00107602 | 0.00798943 | 0.00364934 | 0.00608934 | 0.00370652 | 0.00261913 | 0.00721446 | 0.00485412 | 0.00167617 |
| 35.5138889 | 0.00561591 | 0.01409727 | 0.00340432 | 0.02097338 | 0.00638703 | 0.01849661 | 0.01757845 | 0.01234976 | 0.07575639 | 0.00779195 | 0.00113627 | 0.00918592 | 0.00501416 | 0.00351559 | 0.003291 | 0.00247022 | 0.00865027 | 0.00359119 | 0.00259647 |
| 35.5277778 | 0.00309991 | 0.01035729 | 0.00426425 | 0.02139799 | 0.00962433 | 0.01899044 | 0.01888445 | 0.01254038 | 0.05430147 | 0.01208217 | 0.00447316 | 0.01591028 | 0.00601525 | 0.00457302 | 0.00573053 | 0.00278679 | 0.01319251 | 0.00534672 | 0.00230058 |
| 35.5416667 | 0.00768601 | 0.00964464 | 0.01172938 | 0.02868427 | 0.01077743 | 0.01846576 | 0.03375742 | 0.01040608 | 0.03094718 | 0.02758471 | 0.02197704 | 0.03752801 | 0.00694839 | 0.01578444 | 0.01063634 | 0.00815199 | 0.02865607 | 0.01828393 | 0.00342549 |
| 35.5555556 | 0.02316191 | 0.02449081 | 0.02579809 | 0.04584535 | 0.01600395 | 0.0331404 | 0.06587551 | 0.00707268 | 0.0255345 | 0.05327669 | 0.04847017 | 0.06966642 | 0.00857396 | 0.03623347 | 0.01262737 | 0.01717748 | 0.04839863 | 0.03940892 | 0.01096521 |
| 35.5694444 | 0.04301988 | 0.04336331 | 0.03594897 | 0.06446978 | 0.03480255 | 0.05351214 | 0.09155116 | 0.0078079 | 0.04619832 | 0.06535123 | 0.05536994 | 0.08290701 | 0.01148663 | 0.0539356 | 0.01112997 | 0.02245122 | 0.05168609 | 0.0524081 | 0.02069737 |
| 35.5833333 | 0.05713624 | 0.0519813 | 0.040848 | 0.07495952 | 0.05725487 | 0.05912085 | 0.0912048 | 0.0093519 | 0.06925133 | 0.05205601 | 0.03844573 | 0.06457357 | 0.01532835 | 0.0602017 | 0.01003579 | 0.02021177 | 0.04005932 | 0.05223082 | 0.02418559 |
| 35.5972222 | 0.06124994 | 0.05625967 | 0.04594925 | 0.07663592 | 0.06945915 | 0.05640695 | 0.07578478 | 0.00789593 | 0.08280658 | 0.03309605 | 0.02066692 | 0.04071077 | 0.01637948 | 0.05374377 | 0.00833234 | 0.01330661 | 0.02850052 | 0.04505688 | 0.02012718 |
| 35.6111111 | 0.05282145 | 0.05741412 | 0.04409479 | 0.07021797 | 0.0660828 | 0.05633793 | 0.06049343 | 0.00951677 | 0.090402 | 0.02096056 | 0.01162001 | 0.02827015 | 0.01223991 | 0.03411737 | 0.00490189 | 0.00817056 | 0.01989818 | 0.03073025 | 0.01202661 |
| 35.625 | 0.03490359 | 0.04743607 | 0.03660292 | 0.05825496 | 0.05128938 | 0.05551045 | 0.047889 | 0.01356812 | 0.09124919 | 0.01252252 | 0.00782865 | 0.02258719 | 0.00623681 | 0.01875322 | 0.00374475 | 0.01113736 | 0.01294634 | 0.01581952 | 0.00629877 |
| 35.6388889 | 0.01596192 | 0.02769815 | 0.03033961 | 0.0445189 | 0.03304122 | 0.04880097 | 0.03598626 | 0.01334311 | 0.08654885 | 0.0082506 | 0.00715089 | 0.01982381 | 0.00295232 | 0.02724234 | 0.0056167 | 0.01820331 | 0.01039186 | 0.01572348 | 0.00753652 |
| 35.6527778 | 0.0056153 | 0.01059228 | 0.0249605 | 0.03367464 | 0.01805531 | 0.0379454 | 0.02739085 | 0.00823051 | 0.07384525 | 0.00996742 | 0.00751503 | 0.01832935 | 0.00277557 | 0.03963423 | 0.00581374 | 0.01902127 | 0.0113976 | 0.02777787 | 0.01143985 |
| 35.6666667 | 0.00915822 | 0.00401884 | 0.01762243 | 0.02772982 | 0.00969829 | 0.02877562 | 0.0259412 | 0.00513 | 0.05563326 | 0.01407638 | 0.00611864 | 0.01517578 | 0.00237839 | 0.03285618 | 0.00372264 | 0.01182366 | 0.01236921 | 0.03631386 | 0.01132549 |
| 35.6805556 | 0.01930853 | 0.00620161 | 0.01183591 | 0.02153641 | 0.00606363 | 0.02231841 | 0.02774819 | 0.00877968 | 0.04118144 | 0.01595661 | 0.00311905 | 0.00984532 | 0.00217167 | 0.01610431 | 0.0018745 | 0.00464538 | 0.0153085 | 0.03895232 | 0.0069883 |
| 35.6944444 | 0.02622757 | 0.01430038 | 0.01383143 | 0.01550369 | 0.0080779 | 0.01502498 | 0.02733319 | 0.01422709 | 0.02837614 | 0.01477795 | 0.00092254 | 0.00544201 | 0.00370085 | 0.00441398 | 0.00071554 | 0.00239142 | 0.01871991 | 0.04152338 | 0.00455754 |
| 35.7083333 | 0.02853705 | 0.02537772 | 0.01703251 | 0.01813248 | 0.01971087 | 0.00747183 | 0.02458387 | 0.01611305 | 0.0163504 | 0.01057896 | 0.00046307 | 0.00265757 | 0.0043685 | 0.00159475 | 0.00029129 | 0.00240759 | 0.01801258 | 0.04628931 | 0.00540793 |
| 35.7222222 | 0.02900284 | 0.03625157 | 0.01454619 | 0.02638791 | 0.0393978 | 0.00732607 | 0.02122472 | 0.01339083 | 0.00859769 | 0.00609454 | 0.00153258 | 0.00163766 | 0.00387694 | 0.00532891 | 0.00087368 | 0.00385685 | 0.01181036 | 0.05182212 | 0.00479097 |
| 35.7361111 | 0.02952094 | 0.04602436 | 0.00874503 | 0.02966967 | 0.06232189 | 0.01757712 | 0.01696249 | 0.01076001 | 0.00676579 | 0.00695179 | 0.00360118 | 0.00365482 | 0.005756 | 0.01159472 | 0.00287207 | 0.00601096 | 0.00656625 | 0.05366959 | 0.00482768 |
| 35.75 | 0.029783 | 0.05346825 | 0.00375054 | 0.02625817 | 0.08260715 | 0.03285332 | 0.01295948 | 0.01377883 | 0.01313257 | 0.01211018 | 0.00514357 | 0.00812707 | 0.00994929 | 0.01498701 | 0.00582725 | 0.00591003 | 0.00974045 | 0.04974196 | 0.00882636 |
| 35.7638889 | 0.02763147 | 0.05674166 | 0.00142627 | 0.01800963 | 0.08815602 | 0.04516893 | 0.01435082 | 0.01669284 | 0.02274216 | 0.01513329 | 0.00523839 | 0.01467894 | 0.01262641 | 0.01207939 | 0.0086634 | 0.0039405 | 0.012817 | 0.0392339 | 0.01250743 |
| 35.7777778 | 0.02198937 | 0.05558811 | 0.00180993 | 0.00925792 | 0.07360728 | 0.04671964 | 0.0206885 | 0.01148774 | 0.03264996 | 0.0136999 | 0.00436841 | 0.02168249 | 0.01154113 | 0.00762564 | 0.01102668 | 0.00428366 | 0.00866237 | 0.02181382 | 0.01234295 |
| 35.7916667 | 0.01518287 | 0.05088653 | 0.00414537 | 0.00778521 | 0.04887743 | 0.04026881 | 0.0270854 | 0.00553779 | 0.04306015 | 0.0095899 | 0.00317884 | 0.02469442 | 0.00689276 | 0.00992282 | 0.01274702 | 0.00763729 | 0.00507446 | 0.01059255 | 0.00882241 |
| 35.8055556 | 0.01028468 | 0.04449653 | 0.00768746 | 0.01455244 | 0.02388678 | 0.03199612 | 0.0314885 | 0.00782706 | 0.05014278 | 0.00606051 | 0.00193538 | 0.02180319 | 0.00399127 | 0.01866155 | 0.01310355 | 0.01005093 | 0.00540884 | 0.01816547 | 0.00442686 |
| 35.8194444 | 0.00808145 | 0.03885927 | 0.00815217 | 0.02302474 | 0.01027799 | 0.02380027 | 0.03333715 | 0.01246011 | 0.0519098 | 0.00574794 | 0.00116665 | 0.01687739 | 0.00710685 | 0.02808508 | 0.0124846 | 0.00952624 | 0.00445613 | 0.03266303 | 0.00194985 |
| 35.8333333 | 0.00858797 | 0.03659417 | 0.00889396 | 0.02828451 | 0.01308232 | 0.01809412 | 0.03239199 | 0.01425526 | 0.05078345 | 0.00833623 | 0.00112257 | 0.01360973 | 0.01202365 | 0.03427783 | 0.01213961 | 0.00662335 | 0.00266989 | 0.03944955 | 0.00153383 |
| 35.8472222 | 0.01164888 | 0.03865232 | 0.01824192 | 0.02881933 | 0.013869 | 0.01860822 | 0.02732716 | 0.01639846 | 0.04940292 | 0.01181554 | 0.00137394 | 0.01106558 | 0.013891 | 0.03484065 | 0.01238543 | 0.00324304 | 0.00318259 | 0.03625592 | 0.00174947 |
| 35.8611111 | 0.01622011 | 0.04226329 | 0.02415942 | 0.02480739 | 0.0125947 | 0.02976936 | 0.01630635 | 0.01612772 | 0.04984027 | 0.01405966 | 0.00172392 | 0.00797243 | 0.01127329 | 0.02866825 | 0.0121893 | 0.00155283 | 0.00507729 | 0.02312713 | 0.00146761 |
| 35.875 | 0.02089707 | 0.04409289 | 0.01742072 | 0.01774677 | 0.02786258 | 0.04951492 | 0.0082021 | 0.01318653 | 0.05677373 | 0.01374863 | 0.00212744 | 0.00493297 | 0.00618839 | 0.01781619 | 0.01026777 | 0.00188222 | 0.00797646 | 0.01236324 | 0.00100359 |
| 35.8888889 | 0.02477149 | 0.04164275 | 0.01686262 | 0.00963242 | 0.05112819 | 0.06594244 | 0.01253867 | 0.02059451 | 0.0720477 | 0.01147677 | 0.00214515 | 0.00302862 | 0.00265031 | 0.00798817 | 0.00724035 | 0.00319079 | 0.01135411 | 0.01565882 | 0.00163865 |
| 35.9027778 | 0.02845268 | 0.03463463 | 0.03369762 | 0.00327881 | 0.06972564 | 0.07002617 | 0.02129823 | 0.03629572 | 0.09417633 | 0.00969034 | 0.00180169 | 0.00285377 | 0.00176747 | 0.00371667 | 0.00424538 | 0.00458348 | 0.01321637 | 0.02019972 | 0.00320983 |
| 35.9166667 | 0.03219492 | 0.02477847 | 0.05069536 | 0.0015887 | 0.08214956 | 0.06815787 | 0.02787795 | 0.04595818 | 0.11840152 | 0.01076304 | 0.00263352 | 0.00385313 | 0.00243192 | 0.00407146 | 0.00190665 | 0.00511106 | 0.01338111 | 0.0165297 | 0.00535968 |
| 35.9305556 | 0.03463916 | 0.01374434 | 0.05521621 | 0.00459809 | 0.08492984 | 0.06649642 | 0.03312336 | 0.04777401 | 0.12945722 | 0.01443911 | 0.00553151 | 0.00501463 | 0.00351406 | 0.00573145 | 0.00096855 | 0.00368365 | 0.01282706 | 0.00988162 | 0.00866592 |
| 35.9444444 | 0.03624049 | 0.00970099 | 0.0530881 | 0.0094316 | 0.08438943 | 0.06159992 | 0.03535378 | 0.04580343 | 0.11690012 | 0.01799926 | 0.0096874 | 0.00593637 | 0.00404329 | 0.00636123 | 0.00136391 | 0.00183586 | 0.01178807 | 0.00556176 | 0.01239996 |
| 35.9583333 | 0.03681906 | 0.01792701 | 0.05091887 | 0.01187675 | 0.08978903 | 0.05771308 | 0.03285003 | 0.04387513 | 0.09078333 | 0.01936152 | 0.01436224 | 0.00620748 | 0.00347346 | 0.00539269 | 0.00205069 | 0.00134876 | 0.01073379 | 0.00807659 | 0.01413467 |
| 35.9722222 | 0.03188307 | 0.02950074 | 0.04635325 | 0.0090986 | 0.09119769 | 0.0575013 | 0.02603542 | 0.04207819 | 0.06522761 | 0.01846207 | 0.01885854 | 0.00549474 | 0.00209513 | 0.00335638 | 0.00176619 | 0.00183396 | 0.00977751 | 0.01589197 | 0.01227854 |
| 35.9861111 | 0.01896571 | 0.03702045 | 0.03598536 | 0.00454227 | 0.07573771 | 0.0506655 | 0.01679212 | 0.0342541 | 0.0461499 | 0.01574746 | 0.02165587 | 0.00403181 | 0.00114514 | 0.00257506 | 0.00142564 | 0.00436227 | 0.00799711 | 0.02389955 | 0.00857734 |
| 36 | 0.01096609 | 0.04238612 | 0.0203838 | 0.00373498 | 0.0459695 | 0.03364923 | 0.00820764 | 0.0205201 | 0.03315885 | 0.01248903 | 0.02195233 | 0.00251087 | 0.00217165 | 0.00533487 | 0.0024002 | 0.00802478 | 0.00500141 | 0.03076571 | 0.00675341 |
| 36.0138889 | 0.01990998 | 0.0482111 | 0.00979244 | 0.00599016 | 0.02348262 | 0.01630278 | 0.00473505 | 0.01489819 | 0.02197193 | 0.01113234 | 0.02077854 | 0.00233732 | 0.00538626 | 0.00813669 | 0.0026659 | 0.00930644 | 0.0024094 | 0.03413018 | 0.00837403 |
| 36.0277778 | 0.02640759 | 0.04556484 | 0.01118646 | 0.00638698 | 0.02378472 | 0.00664389 | 0.00425246 | 0.01646031 | 0.01819864 | 0.01527333 | 0.02135426 | 0.00889377 | 0.00651601 | 0.00850297 | 0.00323838 | 0.00787917 | 0.00583615 | 0.02742901 | 0.00818589 |
| 36.0416667 | 0.02891959 | 0.03742245 | 0.01647062 | 0.01161502 | 0.03007788 | 0.01406989 | 0.0093084 | 0.02004792 | 0.04422047 | 0.03331314 | 0.03440915 | 0.03699956 | 0.01300418 | 0.02064155 | 0.01137468 | 0.01460082 | 0.02604803 | 0.03605232 | 0.01186895 |
| 36.0555556 | 0.05614105 | 0.0515337 | 0.03719191 | 0.02727297 | 0.05114392 | 0.04718432 | 0.0259228 | 0.0419491 | 0.1069478 | 0.06621619 | 0.06453941 | 0.08623751 | 0.04007296 | 0.0518804 | 0.02934267 | 0.03868922 | 0.05987816 | 0.08156542 | 0.03124423 |
| 36.0694444 | 0.08438096 | 0.07169883 | 0.06622688 | 0.03357642 | 0.09152754 | 0.08286589 | 0.03740551 | 0.07195638 | 0.15726522 | 0.07817599 | 0.07859955 | 0.09458631 | 0.06181951 | 0.06290385 | 0.03942711 | 0.05336573 | 0.06598975 | 0.10487788 | 0.04433182 |
| 36.0833333 | 0.07410842 | 0.06059335 | 0.06467509 | 0.02108985 | 0.10512704 | 0.08825728 | 0.03119181 | 0.08744985 | 0.14660274 | 0.05316547 | 0.05697664 | 0.05490992 | 0.04470272 | 0.04182294 | 0.03009368 | 0.04092294 | 0.04529838 | 0.06588962 | 0.03228289 |
| 36.0972222 | 0.04475395 | 0.03622818 | 0.04307928 | 0.00971977 | 0.08912903 | 0.06987469 | 0.01745785 | 0.09307561 | 0.10558463 | 0.033238 | 0.03907852 | 0.04402522 | 0.02357585 | 0.0351232 | 0.02457201 | 0.03332315 | 0.04786773 | 0.0244138 | 0.02183414 |
| 36.1111111 | 0.02353995 | 0.02409655 | 0.02987856 | 0.00945109 | 0.07436451 | 0.050378 | 0.00674942 | 0.08310444 | 0.0797567 | 0.02701924 | 0.03407934 | 0.04805314 | 0.03494605 | 0.04372236 | 0.03443692 | 0.03812241 | 0.05419224 | 0.02214766 | 0.02778081 |
| 36.125 | 0.010449 | 0.02337072 | 0.02317656 | 0.01044882 | 0.06434719 | 0.03547154 | 0.00258977 | 0.04848541 | 0.0609749 | 0.01586235 | 0.01913386 | 0.03182759 | 0.05191489 | 0.04054837 | 0.04164909 | 0.03330383 | 0.03588498 | 0.03048218 | 0.03161931 |
| 36.1388889 | 0.00364961 | 0.02781837 | 0.01711995 | 0.00884777 | 0.05683395 | 0.02665551 | 0.00176772 | 0.02354757 | 0.03361228 | 0.00818943 | 0.00645998 | 0.01841651 | 0.05250648 | 0.02937071 | 0.03797431 | 0.02192965 | 0.01449889 | 0.03260914 | 0.02867698 |
| 36.1527778 | 0.00361634 | 0.03084546 | 0.00979567 | 0.00873803 | 0.05003544 | 0.02280622 | 0.00156045 | 0.04005371 | 0.01452233 | 0.00647094 | 0.00429613 | 0.01491915 | 0.04415682 | 0.01669478 | 0.0292324 | 0.01357287 | 0.00391686 | 0.03278583 | 0.0236763 |
| 36.1666667 | 0.00752342 | 0.03113608 | 0.00527108 | 0.01077806 | 0.03687614 | 0.01784056 | 0.00334319 | 0.07420643 | 0.01427346 | 0.00481282 | 0.00350447 | 0.01246595 | 0.03400931 | 0.00566325 | 0.02029752 | 0.00785715 | 0.00214652 | 0.0329841 | 0.01498486 |
| 36.1805556 | 0.01391647 | 0.03107447 | 0.00910738 | 0.01333133 | 0.02008646 | 0.00986991 | 0.00710151 | 0.09841501 | 0.0197932 | 0.00420178 | 0.00248334 | 0.00717242 | 0.02562314 | 0.00096983 | 0.01303016 | 0.00407213 | 0.00326583 | 0.03636982 | 0.00695173 |
| 36.1944444 | 0.02432616 | 0.02866026 | 0.01753417 | 0.01590238 | 0.01806646 | 0.0047786 | 0.01131709 | 0.10882856 | 0.02497091 | 0.00444544 | 0.00207049 | 0.00355633 | 0.01878448 | 0.000463 | 0.00734539 | 0.00234837 | 0.0055448 | 0.04128643 | 0.00587714 |
| 36.2083333 | 0.03782266 | 0.02087573 | 0.02615642 | 0.0189537 | 0.03585179 | 0.00889192 | 0.01496066 | 0.10367645 | 0.02660756 | 0.00630544 | 0.00301123 | 0.0040025 | 0.01323078 | 0.00090928 | 0.00301061 | 0.00184654 | 0.00849721 | 0.04497429 | 0.00883487 |
| 36.2222222 | 0.05036021 | 0.01042709 | 0.03574847 | 0.02207671 | 0.05813581 | 0.02178689 | 0.01657999 | 0.08648899 | 0.02194449 | 0.00971665 | 0.00572457 | 0.00352269 | 0.01017523 | 0.0030497 | 0.00090406 | 0.00136731 | 0.00883928 | 0.04587101 | 0.00757369 |
| 36.2361111 | 0.05901405 | 0.00518393 | 0.04699061 | 0.02480151 | 0.07501007 | 0.04127165 | 0.01599934 | 0.06680304 | 0.0146476 | 0.01091239 | 0.00753794 | 0.00354972 | 0.00907224 | 0.00645987 | 0.00109093 | 0.00095345 | 0.00604483 | 0.04146325 | 0.00411573 |

| | | | | | | | | | | | | | | | | | | | |
|---|---|---|---|---|---|---|---|---|---|---|---|---|---|---|---|---|---|---|---|
| 36.25 | 0.06216738 | 0.00873603 | 0.0590786 | 0.02715491 | 0.08430998 | 0.05169474 | 0.01570408 | 0.05006907 | 0.01067193 | 0.0105624 | 0.00840649 | 0.00754102 | 0.0082671 | 0.00891792 | 0.00305886 | 0.00125256 | 0.00332354 | 0.0291845 | 0.00449948 |
| 36.2638889 | 0.06111576 | 0.01692046 | 0.06783843 | 0.0289943 | 0.08475497 | 0.0470917 | 0.01742304 | 0.04127785 | 0.01422397 | 0.01149683 | 0.01044914 | 0.01105369 | 0.00622182 | 0.00960571 | 0.00525603 | 0.00229683 | 0.00356604 | 0.01564639 | 0.00499139 |
| 36.2777778 | 0.06091182 | 0.02646138 | 0.06867907 | 0.03001214 | 0.08010792 | 0.04029484 | 0.019931 | 0.03965119 | 0.02254908 | 0.01291126 | 0.01263976 | 0.01185088 | 0.00397735 | 0.0092079 | 0.00510327 | 0.00339182 | 0.00592131 | 0.01491239 | 0.00344143 |
| 36.2916667 | 0.06249475 | 0.03374259 | 0.06125529 | 0.03021663 | 0.0815405 | 0.02807276 | 0.02108143 | 0.03736311 | 0.03138053 | 0.01297317 | 0.0131935 | 0.01043556 | 0.00528449 | 0.00860062 | 0.00290975 | 0.00413233 | 0.00752249 | 0.026633 | 0.00367359 |
| 36.3055556 | 0.06250696 | 0.03388595 | 0.04880611 | 0.02989163 | 0.09237494 | 0.01484255 | 0.02061217 | 0.03576663 | 0.03702693 | 0.01214136 | 0.01231583 | 0.00634076 | 0.0099595 | 0.00958225 | 0.0012691 | 0.00514378 | 0.00687164 | 0.03681234 | 0.0052575 |
| 36.3194444 | 0.06096066 | 0.02804251 | 0.03687037 | 0.02993198 | 0.10370911 | 0.01181937 | 0.0197748 | 0.03493354 | 0.03461284 | 0.01199031 | 0.01156608 | 0.00280298 | 0.01350726 | 0.01352845 | 0.00126969 | 0.00675119 | 0.00409891 | 0.04012618 | 0.00481447 |
| 36.3333333 | 0.06108196 | 0.02341537 | 0.02916156 | 0.03150033 | 0.10759802 | 0.01532923 | 0.01997096 | 0.03148681 | 0.02474912 | 0.01278809 | 0.01179552 | 0.00233762 | 0.0126801 | 0.01852449 | 0.00323132 | 0.00846684 | 0.00143279 | 0.04008655 | 0.00504577 |
| 36.3472222 | 0.06210909 | 0.02531331 | 0.02591734 | 0.0345741 | 0.09663586 | 0.01596557 | 0.02162568 | 0.02798947 | 0.01426025 | 0.01374472 | 0.01257472 | 0.00257304 | 0.0074669 | 0.02074082 | 0.00684275 | 0.01011367 | 0.00158502 | 0.03648666 | 0.00875399 |
| 36.3611111 | 0.05930846 | 0.03101821 | 0.02469545 | 0.03745533 | 0.06999979 | 0.00961325 | 0.02350405 | 0.01993588 | 0.0067243 | 0.0142233 | 0.0125067 | 0.00323041 | 0.00317455 | 0.01865216 | 0.0099186 | 0.0108842 | 0.00418623 | 0.02778183 | 0.01119199 |
| 36.375 | 0.05257327 | 0.03452481 | 0.02259269 | 0.03852088 | 0.03690873 | 0.00580165 | 0.02448819 | 0.01396592 | 0.00324107 | 0.01383284 | 0.01045719 | 0.00486564 | 0.00341705 | 0.01376133 | 0.01011659 | 0.00951079 | 0.00685081 | 0.0169994 | 0.0091171 |
| 36.3888889 | 0.04446911 | 0.03495823 | 0.02030972 | 0.03794757 | 0.01552134 | 0.00884609 | 0.02522884 | 0.02421454 | 0.00401961 | 0.01245324 | 0.00715649 | 0.00649161 | 0.00415688 | 0.00867427 | 0.00783721 | 0.00679692 | 0.00834351 | 0.00781336 | 0.004997 |
| 36.4027778 | 0.0366089 | 0.03504426 | 0.01972727 | 0.03684156 | 0.01736892 | 0.01263383 | 0.027357 | 0.04114763 | 0.00578759 | 0.01149699 | 0.00512648 | 0.00841424 | 0.00356221 | 0.00625458 | 0.00613685 | 0.00488998 | 0.00876076 | 0.00404148 | 0.00283609 |
| 36.4166667 | 0.03091882 | 0.03541037 | 0.02085841 | 0.03568016 | 0.02851179 | 0.01561761 | 0.03085343 | 0.05294848 | 0.00645577 | 0.01265892 | 0.0059581 | 0.01047031 | 0.00363373 | 0.00725745 | 0.00641176 | 0.0044188 | 0.00867255 | 0.00682478 | 0.0027925 |
| 36.4305556 | 0.0279626 | 0.03387846 | 0.02235181 | 0.0340598 | 0.03442602 | 0.01911534 | 0.03395188 | 0.0565155 | 0.00700715 | 0.01533071 | 0.00853641 | 0.01165588 | 0.00487993 | 0.00958091 | 0.00682406 | 0.00399024 | 0.00853814 | 0.01231802 | 0.00226706 |
| 36.4444444 | 0.02654075 | 0.02908698 | 0.02275966 | 0.03208096 | 0.03554328 | 0.02100793 | 0.03513771 | 0.05162211 | 0.00856385 | 0.01734027 | 0.01033721 | 0.01223939 | 0.00651963 | 0.01152445 | 0.00537977 | 0.00249895 | 0.00802899 | 0.01807211 | 0.00217257 |
| 36.4583333 | 0.02593709 | 0.02306458 | 0.02231578 | 0.03088907 | 0.03711749 | 0.02177141 | 0.03401178 | 0.0445817 | 0.01080302 | 0.01735474 | 0.00987313 | 0.01294472 | 0.00777351 | 0.01228992 | 0.00266134 | 0.00096453 | 0.00697607 | 0.02352746 | 0.00299616 |
| 36.4722222 | 0.02752092 | 0.01854915 | 0.02208057 | 0.03095256 | 0.04405293 | 0.02555894 | 0.03217535 | 0.04143564 | 0.01386319 | 0.0159007 | 0.00806586 | 0.01354309 | 0.00872986 | 0.01080806 | 0.00086758 | 0.00043781 | 0.00617403 | 0.02745737 | 0.00254409 |
| 36.4861111 | 0.02975617 | 0.0156886 | 0.02200126 | 0.03126246 | 0.05344069 | 0.03203892 | 0.03124103 | 0.04318521 | 0.0194765 | 0.01394805 | 0.00651744 | 0.01383426 | 0.00879982 | 0.00740751 | 0.00054783 | 0.00066897 | 0.0062212 | 0.02888425 | 0.00206173 |
| 36.5 | 0.02650565 | 0.01284246 | 0.02128418 | 0.030833 | 0.05819529 | 0.0382216 | 0.03037213 | 0.04737364 | 0.0265339 | 0.01172444 | 0.00567486 | 0.01362824 | 0.00620555 | 0.00401811 | 0.00147065 | 0.00092905 | 0.00642184 | 0.02756774 | 0.00391809 |
| 36.5138889 | 0.01981835 | 0.01067163 | 0.01980861 | 0.03069738 | 0.0572503 | 0.04169865 | 0.02891716 | 0.05100155 | 0.030538 | 0.01112787 | 0.00570739 | 0.01315374 | 0.00339269 | 0.00230311 | 0.00376153 | 0.00084544 | 0.00598694 | 0.02537446 | 0.0056127 |
| 36.5277778 | 0.0218759 | 0.016947 | 0.02092436 | 0.03335015 | 0.04713627 | 0.03860627 | 0.02973755 | 0.05361861 | 0.02798278 | 0.01898508 | 0.01004663 | 0.01710903 | 0.0035766 | 0.00423645 | 0.00525353 | 0.00148172 | 0.00811418 | 0.02999617 | 0.00452841 |
| 36.5416667 | 0.04260748 | 0.04461084 | 0.03041244 | 0.04063228 | 0.0319134 | 0.02615747 | 0.03761882 | 0.05237315 | 0.01973371 | 0.04514514 | 0.02807811 | 0.03306315 | 0.00430158 | 0.01310382 | 0.0053672 | 0.00539729 | 0.01976917 | 0.05022415 | 0.00651654 |
| 36.5555556 | 0.07643162 | 0.09190232 | 0.04445194 | 0.04900116 | 0.03450852 | 0.01852358 | 0.05092358 | 0.03595153 | 0.02090136 | 0.0812874 | 0.05146634 | 0.05672616 | 0.00804528 | 0.02851789 | 0.00945508 | 0.01323734 | 0.0390089 | 0.07598466 | 0.01889147 |
| 36.5694444 | 0.09450616 | 0.12112644 | 0.04853906 | 0.04815092 | 0.05618827 | 0.02516446 | 0.0588033 | 0.01920857 | 0.03921964 | 0.09154964 | 0.05289324 | 0.06582366 | 0.01845943 | 0.04272241 | 0.01834158 | 0.01928995 | 0.04883108 | 0.08031806 | 0.03729112 |
| 36.5833333 | 0.07873949 | 0.10340865 | 0.0386582 | 0.03603096 | 0.07910244 | 0.02814894 | 0.05512632 | 0.02398817 | 0.04941122 | 0.06577291 | 0.0343464 | 0.05105162 | 0.02937865 | 0.04763878 | 0.02443022 | 0.01940554 | 0.04025849 | 0.0633177 | 0.04871839 |
| 36.5972222 | 0.05316736 | 0.0669798 | 0.02559121 | 0.02469939 | 0.096367 | 0.02374669 | 0.04580195 | 0.03064638 | 0.04145789 | 0.03562462 | 0.02110007 | 0.03140048 | 0.03371743 | 0.04245562 | 0.02399362 | 0.0160174 | 0.02558826 | 0.04938862 | 0.04382638 |
| 36.6111111 | 0.04522048 | 0.04686435 | 0.01738522 | 0.02436164 | 0.10378718 | 0.02303614 | 0.03848998 | 0.03135539 | 0.03780513 | 0.01790225 | 0.0180738 | 0.02064615 | 0.02829588 | 0.03166581 | 0.01893739 | 0.00994681 | 0.01663187 | 0.04034 | 0.02592422 |
| 36.625 | 0.05203335 | 0.04434448 | 0.01451745 | 0.03614706 | 0.10839105 | 0.02785734 | 0.03848933 | 0.03194985 | 0.04197393 | 0.00774258 | 0.01359912 | 0.01576064 | 0.01539732 | 0.01882122 | 0.01175852 | 0.00650326 | 0.0129312 | 0.02468994 | 0.01418501 |
| 36.6388889 | 0.06004483 | 0.04738756 | 0.01499981 | 0.05394062 | 0.11021197 | 0.03215966 | 0.04640497 | 0.02199707 | 0.03717622 | 0.00259729 | 0.00601799 | 0.01242329 | 0.00670141 | 0.008922 | 0.00570587 | 0.01135531 | 0.0108113 | 0.0146535 | 0.02320814 |
| 36.6527778 | 0.06212461 | 0.04805709 | 0.01702286 | 0.07094306 | 0.10093664 | 0.02968889 | 0.05313147 | 0.01264522 | 0.02028077 | 0.00310277 | 0.00240752 | 0.00957153 | 0.00649551 | 0.00711707 | 0.00396511 | 0.01636325 | 0.00890839 | 0.02266287 | 0.0390453 |
| 36.6666667 | 0.05803625 | 0.04423979 | 0.02084006 | 0.08531419 | 0.08840319 | 0.02652499 | 0.05579699 | 0.02082771 | 0.01064587 | 0.00857502 | 0.00303538 | 0.00807876 | 0.00753681 | 0.01077793 | 0.00731085 | 0.01437629 | 0.00767904 | 0.03428996 | 0.04293901 |
| 36.6805556 | 0.05046112 | 0.03528057 | 0.02744435 | 0.09186703 | 0.07277801 | 0.02659322 | 0.05901104 | 0.03037226 | 0.01977168 | 0.01395341 | 0.00444205 | 0.00878087 | 0.01111984 | 0.01297617 | 0.01361779 | 0.0080683 | 0.00723067 | 0.04182378 | 0.03296933 |
| 36.6944444 | 0.04096301 | 0.02232238 | 0.03464023 | 0.07885545 | 0.04962985 | 0.0218275 | 0.05150618 | 0.02743891 | 0.02828723 | 0.01395549 | 0.00454193 | 0.01025007 | 0.01685078 | 0.01282828 | 0.01832121 | 0.00361686 | 0.00672585 | 0.05056737 | 0.01878656 |
| 36.7083333 | 0.02872902 | 0.01097054 | 0.0352341 | 0.04460144 | 0.02528899 | 0.01503075 | 0.03048982 | 0.02278379 | 0.02046281 | 0.00853012 | 0.00302907 | 0.00977985 | 0.01472784 | 0.01594747 | 0.01706685 | 0.00306856 | 0.00570096 | 0.05996126 | 0.01478072 |
| 36.7222222 | 0.01470589 | 0.00944095 | 0.02702631 | 0.02225146 | 0.01582798 | 0.02031499 | 0.02305581 | 0.02965166 | 0.00770333 | 0.00533193 | 0.00275883 | 0.00667832 | 0.0100191 | 0.02284753 | 0.01130126 | 0.00292998 | 0.00452565 | 0.06757505 | 0.02197737 |
| 36.7361111 | 0.00600933 | 0.01823229 | 0.0180614 | 0.03304469 | 0.03237912 | 0.03475729 | 0.03560151 | 0.05301804 | 0.00287895 | 0.01038639 | 0.00423753 | 0.00341313 | 0.01460573 | 0.02882067 | 0.00560953 | 0.00428215 | 0.00373138 | 0.06933492 | 0.02818115 |
| 36.75 | 0.00688539 | 0.0293637 | 0.01418066 | 0.04835552 | 0.06482207 | 0.04128899 | 0.03885623 | 0.08514192 | 0.00646682 | 0.0182542 | 0.01737881 | 0.00279801 | 0.01958548 | 0.03180261 | 0.00322275 | 0.00805557 | 0.00307784 | 0.05946344 | 0.02639954 |
| 36.7638889 | 0.01166888 | 0.03729447 | 0.01562198 | 0.04956135 | 0.09300007 | 0.03818679 | 0.02383014 | 0.10437341 | 0.01152537 | 0.02234343 | 0.03341333 | 0.00343054 | 0.01526532 | 0.03094696 | 0.00516699 | 0.00962847 | 0.00287514 | 0.04431198 | 0.01662234 |
| 36.7777778 | 0.01178055 | 0.03985882 | 0.021808 | 0.04070987 | 0.10718709 | 0.03626148 | 0.0106922 | 0.08917274 | 0.00901628 | 0.0219853 | 0.031347 | 0.0025148 | 0.00912172 | 0.02518878 | 0.00970847 | 0.00653839 | 0.00328493 | 0.03386315 | 0.00888292 |
| 36.7916667 | 0.00674437 | 0.03652315 | 0.03028752 | 0.02742941 | 0.11321883 | 0.03950209 | 0.00931246 | 0.05317306 | 0.00543736 | 0.01870732 | 0.03159896 | 0.00133361 | 0.01055454 | 0.01634523 | 0.014442 | 0.00359523 | 0.00307152 | 0.02494829 | 0.01090016 |
| 36.8055556 | 0.00321397 | 0.02945238 | 0.03529673 | 0.01522914 | 0.11580278 | 0.04687857 | 0.01366133 | 0.04414378 | 0.00779325 | 0.01456416 | 0.03137475 | 0.00247943 | 0.01658665 | 0.00842966 | 0.01582624 | 0.00452683 | 0.00373731 | 0.01420856 | 0.01212125 |
| 36.8194444 | 0.00250409 | 0.02280874 | 0.03199346 | 0.0084536 | 0.1131177 | 0.05064425 | 0.01708564 | 0.07372312 | 0.01069031 | 0.01111147 | 0.01595121 | 0.00547635 | 0.01703157 | 0.00423246 | 0.01202253 | 0.00488279 | 0.00654222 | 0.01122924 | 0.01262531 |
| 36.8333333 | 0.00241434 | 0.01946745 | 0.02312574 | 0.01292378 | 0.09507463 | 0.04760162 | 0.01697891 | 0.09903741 | 0.01105384 | 0.00878715 | 0.00671582 | 0.00791351 | 0.01100932 | 0.00359749 | 0.00630582 | 0.00296564 | 0.00829403 | 0.02306821 | 0.01837816 |
| 36.8472222 | 0.00374316 | 0.01855717 | 0.01521262 | 0.02676848 | 0.06422518 | 0.04286612 | 0.01293998 | 0.09785103 | 0.01013622 | 0.00813382 | 0.00744549 | 0.00851956 | 0.00787487 | 0.00741426 | 0.00538976 | 0.00133779 | 0.00747125 | 0.03901698 | 0.01892874 |
| 36.8611111 | 0.00450275 | 0.01834229 | 0.00872553 | 0.04170683 | 0.0401791 | 0.04041504 | 0.0078042 | 0.08541858 | 0.01320841 | 0.01068579 | 0.00927451 | 0.00804939 | 0.00800874 | 0.01611689 | 0.01213647 | 0.00110997 | 0.0055528 | 0.04944607 | 0.01323472 |
| 36.875 | 0.00490182 | 0.01932719 | 0.00869847 | 0.05246775 | 0.02641613 | 0.04038608 | 0.00776974 | 0.0680526 | 0.0253677 | 0.01635092 | 0.01637913 | 0.00782467 | 0.00760033 | 0.02408401 | 0.01969349 | 0.00161671 | 0.00408476 | 0.05263302 | 0.01018468 |
| 36.8888889 | 0.00675701 | 0.02329471 | 0.01854748 | 0.05713391 | 0.0147527 | 0.03776313 | 0.01419352 | 0.04071028 | 0.05549235 | 0.0218236 | 0.0336386 | 0.00804689 | 0.01348943 | 0.02700715 | 0.02107776 | 0.00328012 | 0.00363376 | 0.04760618 | 0.00895084 |
| 36.9027778 | 0.00704534 | 0.02935747 | 0.02549564 | 0.05503822 | 0.00677276 | 0.03209583 | 0.0243673 | 0.02746102 | 0.08951585 | 0.02391323 | 0.04369656 | 0.00815821 | 0.02467594 | 0.02583974 | 0.01629046 | 0.00881587 | 0.00406721 | 0.03673914 | 0.00640829 |
| 36.9166667 | 0.01012415 | 0.03360919 | 0.02708836 | 0.04669847 | 0.00926124 | 0.02846015 | 0.03650773 | 0.05348738 | 0.09647844 | 0.02249966 | 0.03996634 | 0.00792521 | 0.0317538 | 0.02251891 | 0.0092289 | 0.01617 | 0.00544294 | 0.02309532 | 0.0047651 |
| 36.9305556 | 0.01937853 | 0.03241522 | 0.03219137 | 0.03515465 | 0.02352431 | 0.03332588 | 0.04648207 | 0.09096407 | 0.08250394 | 0.01959261 | 0.04101905 | 0.00733249 | 0.03302434 | 0.0172538 | 0.00433574 | 0.02018666 | 0.00708933 | 0.01030938 | 0.00667801 |
| 36.9444444 | 0.02583966 | 0.02538345 | 0.04261442 | 0.02589783 | 0.04501887 | 0.04734171 | 0.05130288 | 0.11682054 | 0.06718796 | 0.01692048 | 0.03558467 | 0.00629188 | 0.03097126 | 0.01056919 | 0.00321246 | 0.01848609 | 0.00824115 | 0.00411976 | 0.00850876 |
| 36.9583333 | 0.02461483 | 0.01515992 | 0.05221122 | 0.02206991 | 0.06219795 | 0.056822 | 0.05217535 | 0.14131256 | 0.05350958 | 0.01484091 | 0.01698926 | 0.00490827 | 0.02545748 | 0.00473177 | 0.00428987 | 0.01288693 | 0.00853664 | 0.00575565 | 0.00761179 |
| 36.9722222 | 0.01751359 | 0.00787485 | 0.05149581 | 0.02130492 | 0.06530289 | 0.05128824 | 0.0486218 | 0.16126121 | 0.04196661 | 0.01273401 | 0.00624142 | 0.00527581 | 0.01664169 | 0.00146164 | 0.00520666 | 0.00753576 | 0.00811842 | 0.00977038 | 0.01235391 |
| 36.9861111 | 0.01049231 | 0.01026319 | 0.03936686 | 0.02008645 | 0.05752641 | 0.03759626 | 0.03973786 | 0.15722507 | 0.03158906 | 0.01003897 | 0.00522371 | 0.01026588 | 0.00877926 | 0.0008472 | 0.0046143 | 0.005456 | 0.00771115 | 0.01341058 | 0.02476273 |
| 37 | 0.01310703 | 0.0210776 | 0.02090394 | 0.01774526 | 0.04602054 | 0.02435969 | 0.02742556 | 0.11460263 | 0.02080957 | 0.00710601 | 0.00470597 | 0.01585199 | 0.00578702 | 0.00225982 | 0.00276047 | 0.00755101 | 0.00692292 | 0.0148476 | 0.03323971 |
| 37.0138889 | 0.0235761 | 0.03285298 | 0.00801375 | 0.01426162 | 0.03361926 | 0.01439367 | 0.01464189 | 0.0629857 | 0.02247892 | 0.00480109 | 0.00347049 | 0.01480099 | 0.00657492 | 0.00552762 | 0.00166482 | 0.01190338 | 0.00542723 | 0.01142994 | 0.0359367 |
| 37.0277778 | 0.02630222 | 0.0301265 | 0.00570825 | 0.00937451 | 0.02632331 | 0.01114739 | 0.00679342 | 0.05287679 | 0.03841654 | 0.00587984 | 0.00383235 | 0.01197234 | 0.00581048 | 0.00664224 | 0.00361446 | 0.01353136 | 0.00800317 | 0.01167004 | 0.03394055 |
| 37.0416667 | 0.02641468 | 0.03212503 | 0.01240327 | 0.0051991 | 0.03557547 | 0.0210154 | 0.01026211 | 0.05259347 | 0.03996333 | 0.01629578 | 0.00811853 | 0.02237846 | 0.00942357 | 0.01624536 | 0.01479098 | 0.01364099 | 0.02337785 | 0.03350397 | 0.02512333 |
| 37.0555556 | 0.04725183 | 0.06655875 | 0.03125965 | 0.00310547 | 0.06002633 | 0.04136904 | 0.02246779 | 0.05289826 | 0.03815238 | 0.03752177 | 0.01089051 | 0.04596358 | 0.03238528 | 0.04702525 | 0.04027262 | 0.025639 | 0.0498926 | 0.07578952 | 0.02328275 |
| 37.0694444 | 0.06583505 | 0.0809516 | 0.04155901 | 0.00600883 | 0.07502558 | 0.05262565 | 0.02969347 | 0.09863429 | 0.04807424 | 0.04937959 | 0.02428529 | 0.05165868 | 0.0614166 | 0.0616257 | 0.05413938 | 0.04114073 | 0.05941252 | 0.09622823 | 0.03147581 |
| 37.0833333 | 0.04922318 | 0.04547428 | 0.0290825 | 0.01875627 | 0.06974359 | 0.04423713 | 0.02663813 | 0.13394676 | 0.03524047 | 0.03658958 | 0.06208748 | 0.03413393 | 0.06024799 | 0.04350187 | 0.04020537 | 0.03262319 | 0.04300381 | 0.06598068 | 0.02803697 |
| 37.0972222 | 0.02010582 | 0.01067747 | 0.01602808 | 0.03456296 | 0.06786923 | 0.02718809 | 0.02256373 | 0.11759537 | 0.01419789 | 0.02246704 | 0.08489093 | 0.02757196 | 0.03159339 | 0.03575005 | 0.03115544 | 0.0158744 | 0.03416334 | 0.03182363 | 0.01951909 |
| 37.1111111 | 0.00695873 | 0.00332462 | 0.01684772 | 0.04009044 | 0.07845276 | 0.01258493 | 0.02238851 | 0.08985809 | 0.0191845 | 0.0194577 | 0.06309019 | 0.02840325 | 0.01564151 | 0.04298817 | 0.03338583 | 0.01705242 | 0.03428478 | 0.02985484 | 0.02492723 |
| 37.125 | 0.00995327 | 0.01013637 | 0.0231301 | 0.03474257 | 0.07934647 | 0.00456523 | 0.02304234 | 0.06939367 | 0.03405947 | 0.01451617 | 0.02800167 | 0.01926622 | 0.02410455 | 0.04507991 | 0.0267125 | 0.02365795 | 0.02284651 | 0.033215 | 0.03267946 |
| 37.1388889 | 0.01873546 | 0.02010597 | 0.02656654 | 0.02832312 | 0.06774373 | 0.0024752 | 0.02285288 | 0.05210963 | 0.04204895 | 0.0082135 | 0.01289102 | 0.01029176 | 0.03998342 | 0.04252081 | 0.01695717 | 0.02613595 | 0.00887081 | 0.0330766 | 0.03776566 |
| 37.1527778 | 0.02559296 | 0.0293656 | 0.02785005 | 0.02449644 | 0.06281904 | 0.0029154 | 0.0228943 | 0.03607775 | 0.04381993 | 0.00428124 | 0.01163816 | 0.00628428 | 0.04889975 | 0.03386371 | 0.0124823 | 0.02572135 | 0.00246188 | 0.03639561 | 0.04365826 |
| 37.1666667 | 0.02764312 | 0.03477798 | 0.02965861 | 0.02162491 | 0.06545732 | 0.0039842 | 0.02244212 | 0.02018069 | 0.03960783 | 0.00226127 | 0.00867839 | 0.00429207 | 0.04441895 | 0.02154251 | 0.00923817 | 0.02027638 | 0.00120357 | 0.03676366 | 0.04274937 |
| 37.1805556 | 0.02334736 | 0.03448794 | 0.03023395 | 0.01892736 | 0.06358686 | 0.00427888 | 0.01942468 | 0.01047214 | 0.0268388 | 0.0026634 | 0.00503165 | 0.00495235 | 0.03142176 | 0.0112815 | 0.00561844 | 0.01165825 | 0.00174677 | 0.03206352 | 0.03022448 |
| 37.1944444 | 0.01463273 | 0.0280689 | 0.02496455 | 0.0166178 | 0.0513742 | 0.0030599 | 0.01456294 | 0.0143986 | 0.01686216 | 0.00251121 | 0.00523193 | 0.00797705 | 0.01800177 | 0.00662889 | 0.00427011 | 0.00457003 | 0.00213964 | 0.02623965 | 0.01400805 |
| 37.2083333 | 0.00748592 | 0.01844804 | 0.01514403 | 0.01642574 | 0.03143978 | 0.00137321 | 0.00918983 | 0.02308223 | 0.02659791 | 0.00193499 | 0.00654764 | 0.01128402 | 0.00807815 | 0.00804407 | 0.0051377 | 0.00158882 | 0.00217924 | 0.02138781 | 0.00860308 |
| 37.2222222 | 0.00713507 | 0.01156675 | 0.00628617 | 0.01832444 | 0.0130447 | 0.00042991 | 0.00446606 | 0.03223443 | 0.04869207 | 0.0040228 | 0.00537893 | 0.01312086 | 0.00283602 | 0.01008377 | 0.00600662 | 0.00127721 | 0.0018352 | 0.01535255 | 0.01629051 |
| 37.2361111 | 0.01182352 | 0.00981927 | 0.00256263 | 0.02017831 | 0.00555275 | 0.0001615 | 0.00232384 | 0.04493508 | 0.06891635 | 0.00746429 | 0.00359412 | 0.01295811 | 0.00138583 | 0.00884822 | 0.00577725 | 0.00278995 | 0.00170353 | 0.0096407 | 0.02591008 |
| 37.25 | 0.01833026 | 0.00849448 | 0.004708 | 0.02071541 | 0.0097429 | 0.00051015 | 0.00269997 | 0.05749005 | 0.08416555 | 0.0101637 | 0.00310677 | 0.01233253 | 0.00240601 | 0.00605778 | 0.00490076 | 0.00524998 | 0.00342905 | 0.00658117 | 0.02813187 |
| 37.2638889 | 0.02513303 | 0.00450967 | 0.00885217 | 0.01980678 | 0.01734039 | 0.00158291 | 0.0034063 | 0.06468191 | 0.09554073 | 0.01218116 | 0.00263852 | 0.01257145 | 0.00480897 | 0.00809358 | 0.00400139 | 0.00620147 | 0.00639194 | 0.005243 | 0.02314582 |
| 37.2777778 | 0.02867879 | 0.00162828 | 0.01154874 | 0.01708502 | 0.02227244 | 0.00261916 | 0.0028695 | 0.06081144 | 0.10268805 | 0.01326891 | 0.00148014 | 0.01313057 | 0.0070691 | 0.015907 | 0.00337918 | 0.00620346 | 0.00848302 | 0.00744214 | 0.01579234 |
| 37.2916667 | 0.0280114 | 0.00127763 | 0.01205679 | 0.01255986 | 0.02417901 | 0.00272911 | 0.0020513 | 0.05322979 | 0.09453522 | 0.0122506 | 0.00119945 | 0.01248284 | 0.00641631 | 0.0224762 | 0.0024053 | 0.00681319 | 0.00840162 | 0.01427876 | 0.00886876 |
| 37.3055556 | 0.02410769 | 0.0031649 | 0.00994283 | 0.00780485 | 0.02330456 | 0.00175042 | 0.00187193 | 0.05603063 | 0.07081014 | 0.00817795 | 0.00161609 | 0.00959228 | 0.00551283 | 0.02248373 | 0.00193288 | 0.00733115 | 0.00551185 | 0.02216938 | 0.00372966 |
| 37.3194444 | 0.016306 | 0.00972973 | 0.00581736 | 0.0065311 | 0.02026867 | 0.00068908 | 0.00273078 | 0.06949377 | 0.05006581 | 0.00329187 | 0.00255606 | 0.00585662 | 0.01129883 | 0.01628938 | 0.00385902 | 0.00812017 | 0.00280452 | 0.02809358 | 0.00156062 |
| 37.3333333 | 0.00747435 | 0.01691021 | 0.00269341 | 0.01134656 | 0.01711826 | 0.00037374 | 0.00501868 | 0.08606446 | 0.03482238 | 0.00184759 | 0.00325536 | 0.00391791 | 0.02021949 | 0.00920287 | 0.0056219 | 0.01027307 | 0.00263659 | 0.02919466 | 0.00139765 |
| 37.3472222 | 0.00250325 | 0.01721615 | 0.00295838 | 0.01960498 | 0.0148747 | 0.00031874 | 0.00659308 | 0.09348447 | 0.01729644 | 0.0052215 | 0.00315981 | 0.00441193 | 0.02374946 | 0.00750821 | 0.00483706 | 0.0111251 | 0.00283338 | 0.02268503 | 0.00329059 |
| 37.3611111 | 0.00325496 | 0.01084437 | 0.00545414 | 0.02682582 | 0.0131522 | 0.00074963 | 0.00651128 | 0.08527578 | 0.00517097 | 0.01087706 | 0.00479736 | 0.00518832 | 0.02049757 | 0.01301179 | 0.00434336 | 0.00733802 | 0.00372705 | 0.01150609 | 0.00783403 |
| 37.375 | 0.00965691 | 0.00507991 | 0.007721 | 0.03174612 | 0.01144311 | 0.00231977 | 0.00603018 | 0.06686642 | 0.00437871 | 0.01460634 | 0.00687179 | 0.00476164 | 0.01387051 | 0.02232917 | 0.00656359 | 0.00386491 | 0.00540057 | 0.00514217 | 0.0104411 |
| 37.3888889 | 0.01807788 | 0.00497066 | 0.00906484 | 0.03544519 | 0.00924481 | 0.00394545 | 0.00461981 | 0.04939914 | 0.01284998 | 0.01437139 | 0.00591193 | 0.0043498 | 0.00792192 | 0.02889932 | 0.00949976 | 0.00548086 | 0.00529443 | 0.00766885 | 0.00766333 |
| 37.4027778 | 0.02340805 | 0.00771045 | 0.0101293 | 0.03817388 | 0.00565208 | 0.00377833 | 0.00262132 | 0.04266581 | 0.03180649 | 0.01188123 | 0.00315645 | 0.00461982 | 0.00417546 | 0.02839218 | 0.01008869 | 0.00620077 | 0.00353885 | 0.01150183 | 0.00548311 |
| 37.4166667 | 0.02428998 | 0.00997547 | 0.01106879 | 0.03875 | 0.0046813 | 0.0019446 | 0.0028523 | 0.04454267 | 0.05484622 | 0.01016649 | 0.00162063 | 0.00480551 | 0.00179789 | 0.02334964 | 0.00809335 | 0.00393678 | 0.00173423 | 0.01106733 | 0.00771402 |
| 37.4305556 | 0.02299003 | 0.01222981 | 0.01191063 | 0.03719443 | 0.01145327 | 0.00059277 | 0.00611174 | 0.04618207 | 0.06856047 | 0.01060498 | 0.00204426 | 0.00474311 | 0.00055258 | 0.01771682 | 0.00547659 | 0.00331659 | 0.00071285 | 0.00847089 | 0.00818618 |
| 37.4444444 | 0.0214797 | 0.01403429 | 0.01342582 | 0.03634304 | 0.02044105 | 0.0006149 | 0.01114141 | 0.04105542 | 0.06564924 | 0.0119199 | 0.00304887 | 0.00487692 | 0.00076014 | 0.01299616 | 0.00408839 | 0.00307523 | 0.00059849 | 0.00796871 | 0.00660668 |
| 37.4583333 | 0.01887638 | 0.01286704 | 0.01566984 | 0.03832966 | 0.02576502 | 0.00088881 | 0.01587927 | 0.02546401 | 0.04510832 | 0.01181295 | 0.00317846 | 0.00534829 | 0.00292151 | 0.01037681 | 0.00413622 | 0.00249624 | 0.00092901 | 0.01152604 | 0.01006213 |
| 37.4722222 | 0.01415511 | 0.00873665 | 0.01696188 | 0.04090558 | 0.02951069 | 0.00064053 | 0.01676807 | 0.01566606 | 0.0244479 | 0.00986002 | 0.00285576 | 0.00631476 | 0.00613488 | 0.01060899 | 0.00479726 | 0.00482945 | 0.00152472 | 0.01623722 | 0.01807157 |
| 37.4861111 | 0.00910076 | 0.00549445 | 0.0159582 | 0.03986641 | 0.0338554 | 0.00042951 | 0.01244867 | 0.03015858 | 0.02727627 | 0.00797701 | 0.00262965 | 0.00754795 | 0.00680076 | 0.01253376 | 0.00551864 | 0.00804131 | 0.00221783 | 0.0185254 | 0.02254281 |
| 37.5 | 0.00547007 | 0.00529783 | 0.01341463 | 0.03501806 | 0.03691036 | 0.00099163 | 0.00681343 | 0.05356397 | 0.0465895 | 0.00698313 | 0.00213114 | 0.00739774 | 0.00454599 | 0.01303223 | 0.0062131 | 0.0089438 | 0.00217144 | 0.01844647 | 0.01768688 |
| 37.5138889 | 0.00332493 | 0.00667426 | 0.01146387 | 0.03051722 | 0.03643868 | 0.00144366 | 0.00569567 | 0.07100214 | 0.06436063 | 0.00670272 | 0.00198339 | 0.00559749 | 0.00408454 | 0.01151659 | 0.00660022 | 0.00814583 | 0.00178008 | 0.01903545 | 0.00958228 |
| 37.5277778 | 0.00662808 | 0.01317839 | 0.01432614 | 0.02944516 | 0.02973899 | 0.00337824 | 0.01438902 | 0.08330125 | 0.07635262 | 0.01074718 | 0.0038693 | 0.00613454 | 0.00833093 | 0.01332972 | 0.0061279 | 0.00541342 | 0.00506685 | 0.02707862 | 0.0071025 |
| 37.5416667 | 0.02623152 | 0.03778174 | 0.02739738 | 0.03363691 | 0.02241143 | 0.01252491 | 0.0399588 | 0.07919263 | 0.07109936 | 0.02662153 | 0.01391191 | 0.01403632 | 0.01817039 | 0.02524286 | 0.01003664 | 0.00476988 | 0.01672327 | 0.04932157 | 0.01119738 |
| 37.5555556 | 0.06340735 | 0.08075126 | 0.04780738 | 0.04223535 | 0.03696528 | 0.02599684 | 0.07736656 | 0.05504529 | 0.0509033 | 0.05076037 | 0.03094893 | 0.02761206 | 0.03094854 | 0.04287168 | 0.02044277 | 0.01151733 | 0.03255339 | 0.07726926 | 0.03325353 |
| 37.5694445 | 0.09308707 | 0.11283607 | 0.06179808 | 0.04676322 | 0.0676518 | 0.03056293 | 0.09958831 | 0.04703199 | 0.0601253 | 0.06238526 | 0.03930247 | 0.03564227 | 0.03928403 | 0.05036783 | 0.0263651 | 0.02035661 | 0.03917888 | 0.08429191 | 0.0743153 |
| 37.5833333 | 0.09551579 | 0.11228884 | 0.06292497 | 0.0397692 | 0.09190474 | 0.02266997 | 0.09356616 | 0.06791124 | 0.1078008 | 0.05045664 | 0.03837866 | 0.03052682 | 0.04092782 | 0.04150595 | 0.02321342 | 0.02426924 | 0.03080084 | 0.06803868 | 0.09858788 |

**Supplementary Table S6.** 1 h integrated motion data in **(c)** 100 mM KCl treatment under RGBday/dimGnight with their respective controls.

| IntervalMid | Control1 | Control2 | Control3 | Control4 | Control5 | Control6 | Control7 | Control8 | Control9 | Stress1 | Stress2 | Stress3 | Stress4 | Stress5 | Stress6 | Stress7 | Stress8 | Stress9 | Stress10 |
|---|---|---|---|---|---|---|---|---|---|---|---|---|---|---|---|---|---|---|---|
| 24.11 | 0.0052362 | 0.0287335 | 0.0215677 | 0.0142466 | 0.0065964 | 0.0206155 | 0.0057566 | 0.0014438 | 0.005802 | 0.0278187 | 0.0135663 | 0.0053148 | 0.0042876 | 0.002741 | 0.0076536 | 0.0097685 | 0.000763 | 0.0023791 | 0.0355702 |
| 24.13 | 0.0058917 | 0.0196451 | 0.0141213 | 0.0092929 | 0.0043701 | 0.0107492 | 0.0037518 | 0.0021811 | 0.0028295 | 0.0158846 | 0.008583 | 0.004948 | 0.0061542 | 0.0035458 | 0.0078696 | 0.0061076 | 0.0012409 | 0.001793 | 0.0288186 |
| 24.14 | 0.0113859 | 0.0062784 | 0.0032548 | 0.0086924 | 0.0014741 | 0.0073616 | 0.0024803 | 0.0042329 | 0.0012965 | 0.0123742 | 0.0116647 | 0.0051118 | 0.0089191 | 0.0075122 | 0.0105451 | 0.0043481 | 0.0044776 | 0.0047996 | 0.0188841 |
| 24.15 | 0.0372951 | 0.0214486 | 0.0031418 | 0.0282384 | 0.0108756 | 0.0252017 | 0.005369 | 0.0094922 | 0.0034518 | 0.0518874 | 0.0507688 | 0.0301842 | 0.0383093 | 0.0204469 | 0.0711363 | 0.0399516 | 0.0333937 | 0.0370815 | 0.0573134 |
| 24.17 | 0.0516964 | 0.0414661 | 0.0047436 | 0.0382519 | 0.0237581 | 0.0379511 | 0.0073721 | 0.0094236 | 0.0052536 | 0.0686114 | 0.0655624 | 0.0514075 | 0.067851 | 0.0307302 | 0.1112957 | 0.0678876 | 0.0508627 | 0.0520484 | 0.0756638 |
| 24.18 | 0.0475292 | 0.0512966 | 0.0044385 | 0.0347279 | 0.0335476 | 0.044229 | 0.0085308 | 0.0085411 | 0.0113724 | 0.0371809 | 0.0346438 | 0.0396882 | 0.0582956 | 0.0310858 | 0.0715489 | 0.0471582 | 0.0327204 | 0.0268974 | 0.0442442 |
| 24.19 | 0.0584342 | 0.0672784 | 0.0068601 | 0.0492122 | 0.0507663 | 0.0776849 | 0.0137379 | 0.0208553 | 0.0356686 | 0.022146 | 0.0187782 | 0.0221386 | 0.0432555 | 0.0258671 | 0.0535357 | 0.0344397 | 0.0223132 | 0.0164276 | 0.0273463 |
| 24.21 | 0.0401989 | 0.0456658 | 0.0083408 | 0.0395198 | 0.0378434 | 0.0675809 | 0.0122646 | 0.0204412 | 0.0383692 | 0.0174969 | 0.012689 | 0.0083393 | 0.02924 | 0.0140574 | 0.0504061 | 0.0308157 | 0.016855 | 0.0143668 | 0.0209042 |
| 24.22 | 0.0121314 | 0.0122374 | 0.0102178 | 0.0155477 | 0.0112458 | 0.0337096 | 0.0087595 | 0.0103444 | 0.023966 | 0.0116 | 0.0083653 | 0.0054258 | 0.0192795 | 0.0117384 | 0.0358782 | 0.0226542 | 0.0113991 | 0.0104905 | 0.0164019 |
| 24.24 | 0.007761 | 0.0074473 | 0.008021 | 0.0084144 | 0.0054265 | 0.0241041 | 0.006276 | 0.0066167 | 0.0162214 | 0.0055216 | 0.0048519 | 0.007417 | 0.0120936 | 0.009826 | 0.0219535 | 0.0152331 | 0.0073152 | 0.0085418 | 0.0093574 |
| 24.25 | 0.0056371 | 0.0043223 | 0.0020365 | 0.0042207 | 0.0021377 | 0.0119722 | 0.0012615 | 0.0026061 | 0.0052093 | 0.0029737 | 0.0033248 | 0.0065718 | 0.0068001 | 0.0024233 | 0.0144157 | 0.0116334 | 0.0049562 | 0.0079455 | 0.0036327 |
| 24.26 | 0.0141303 | 0.006076 | 0.0022945 | 0.0056279 | 0.0043065 | 0.0135097 | 0.0014877 | 0.0025516 | 0.0039732 | 0.007619 | 0.0074926 | 0.0077806 | 0.0178633 | 0.0030251 | 0.0232985 | 0.02142 | 0.009327 | 0.0133442 | 0.0099202 |
| 24.28 | 0.0177066 | 0.0063048 | 0.003161 | 0.0051307 | 0.0050129 | 0.0100699 | 0.0026006 | 0.0019904 | 0.0045359 | 0.0082872 | 0.0083311 | 0.0068158 | 0.0206536 | 0.0037736 | 0.0253663 | 0.0234344 | 0.0095195 | 0.011919 | 0.012683 |
| 24.29 | 0.0183624 | 0.0064347 | 0.0060356 | 0.0082449 | 0.0043507 | 0.0057324 | 0.0047101 | 0.0016463 | 0.0090123 | 0.0070918 | 0.0082055 | 0.0081344 | 0.019215 | 0.0042001 | 0.03417 | 0.0261807 | 0.0098786 | 0.0081366 | 0.0164337 |
| 24.31 | 0.0138027 | 0.0050367 | 0.0060076 | 0.0078204 | 0.0030806 | 0.0083517 | 0.0042069 | 0.0016498 | 0.0100293 | 0.005691 | 0.0073657 | 0.0098764 | 0.0164934 | 0.0040823 | 0.0351355 | 0.0245747 | 0.0098693 | 0.0109249 | 0.0176107 |
| 24.32 | 0.0043788 | 0.0017892 | 0.0040358 | 0.003298 | 0.0012076 | 0.0116583 | 0.0022502 | 0.0008168 | 0.0051128 | 0.0026037 | 0.0041015 | 0.0080323 | 0.0066678 | 0.0027787 | 0.0201191 | 0.0124412 | 0.0061676 | 0.0134797 | 0.0130232 |
| 24.33 | 0.011438 | 0.0045882 | 0.0104047 | 0.0098463 | 0.0041775 | 0.0216873 | 0.0073818 | 0.0004626 | 0.003279 | 0.0047167 | 0.0064195 | 0.0102063 | 0.0041568 | 0.0044608 | 0.0257818 | 0.0145268 | 0.0077592 | 0.0173888 | 0.0202695 |
| 24.35 | 0.0131513 | 0.0048866 | 0.0103116 | 0.0116804 | 0.0056004 | 0.0209271 | 0.0091358 | 0.0006742 | 0.0024731 | 0.0057004 | 0.0065248 | 0.0072477 | 0.0030864 | 0.0043267 | 0.0243681 | 0.0132283 | 0.0068698 | 0.0112853 | 0.0190427 |
| 24.36 | 0.0090823 | 0.0029896 | 0.005484 | 0.0076996 | 0.0054063 | 0.0148375 | 0.0064903 | 0.0026549 | 0.0056026 | 0.007129 | 0.0058435 | 0.0014173 | 0.0043955 | 0.003739 | 0.0210761 | 0.0110827 | 0.0060712 | 0.0025199 | 0.0142707 |
| 24.38 | 0.0104343 | 0.0035945 | 0.0057658 | 0.0078788 | 0.0044026 | 0.0107127 | 0.0042723 | 0.0030688 | 0.0071864 | 0.0077881 | 0.0059062 | 0.0003338 | 0.0053103 | 0.0037984 | 0.0194126 | 0.0102553 | 0.0060283 | 0.0017716 | 0.0126688 |
| 24.39 | 0.0129337 | 0.0041027 | 0.0058384 | 0.0102445 | 0.0015399 | 0.0052524 | 0.0027122 | 0.0014445 | 0.0044812 | 0.0060965 | 0.0046374 | 0.0005526 | 0.0044811 | 0.0027379 | 0.0114922 | 0.0061295 | 0.003659 | 0.0016799 | 0.0077397 |
| 24.4 | 0.0213889 | 0.0056137 | 0.0069493 | 0.0195513 | 0.0007155 | 0.0134642 | 0.0065363 | 0.0023544 | 0.0037342 | 0.0086061 | 0.0061903 | 0.0029321 | 0.0051141 | 0.0029511 | 0.0128544 | 0.0064898 | 0.0032783 | 0.0020224 | 0.0089932 |
| 24.42 | 0.0183346 | 0.0042466 | 0.0045217 | 0.0186512 | 0.0005783 | 0.0168286 | 0.0072035 | 0.0046719 | 0.0035003 | 0.0064256 | 0.0044264 | 0.0046653 | 0.0041827 | 0.0019372 | 0.009527 | 0.004714 | 0.0021739 | 0.0026388 | 0.0063733 |
| 24.43 | 0.0110602 | 0.0015838 | 0.0029576 | 0.0120797 | 0.0008267 | 0.0120358 | 0.0053309 | 0.009881 | 0.0069807 | 0.0024048 | 0.0017927 | 0.0057877 | 0.0052323 | 0.0010746 | 0.0065069 | 0.0037912 | 0.0022093 | 0.0056692 | 0.0029563 |
| 24.44 | 0.0244586 | 0.0038406 | 0.0151138 | 0.0191899 | 0.00151 | 0.0115192 | 0.0105041 | 0.0261176 | 0.0248781 | 0.0084017 | 0.0067948 | 0.0136617 | 0.0145362 | 0.0040601 | 0.0203192 | 0.0127593 | 0.0085361 | 0.0172484 | 0.009533 |
| 24.46 | 0.0292612 | 0.0070931 | 0.021809 | 0.0186188 | 0.0012609 | 0.0073086 | 0.0122752 | 0.0266225 | 0.0296353 | 0.0113398 | 0.0096822 | 0.015219 | 0.0160653 | 0.0058214 | 0.0248094 | 0.016005 | 0.0112146 | 0.0193034 | 0.0121726 |
| 24.47 | 0.0327245 | 0.0131302 | 0.0231939 | 0.0196415 | 0.000736 | 0.0063991 | 0.0133436 | 0.0150542 | 0.0231319 | 0.0125114 | 0.0121919 | 0.0140444 | 0.0138746 | 0.008093 | 0.0246749 | 0.0170203 | 0.012578 | 0.0147605 | 0.0116987 |
| 24.49 | 0.0267295 | 0.0117795 | 0.016798 | 0.0186273 | 0.0005837 | 0.0102543 | 0.0112847 | 0.0088388 | 0.0152244 | 0.0115877 | 0.0114786 | 0.0127411 | 0.0113274 | 0.0079578 | 0.0218316 | 0.0154407 | 0.0116612 | 0.0108895 | 0.0094625 |
| 24.5 | 0.0061184 | 0.0038745 | 0.0033717 | 0.0071214 | 0.0001514 | 0.0096393 | 0.0034163 | 0.0025491 | 0.003072 | 0.0055078 | 0.0053433 | 0.0062316 | 0.0046089 | 0.0036225 | 0.0101572 | 0.0074059 | 0.0055307 | 0.0046446 | 0.0037534 |
| 24.51 | 0.0106823 | 0.0179308 | 0.0036953 | 0.008148 | 0.0011498 | 0.0136338 | 0.0031243 | 0.0041091 | 0.0086618 | 0.0054909 | 0.0065635 | 0.0077329 | 0.0104416 | 0.0029759 | 0.0138839 | 0.0104483 | 0.0035226 | 0.0177762 | 0.0098088 |
| 24.53 | 0.0286875 | 0.0421904 | 0.0036486 | 0.0103927 | 0.0017814 | 0.0206302 | 0.0061564 | 0.0057584 | 0.0212411 | 0.0064609 | 0.010405 | 0.0129838 | 0.0192652 | 0.0039129 | 0.020558 | 0.0158057 | 0.0016594 | 0.0366445 | 0.0196069 |
| 24.54 | 0.0872815 | 0.1031442 | 0.0097129 | 0.0102837 | 0.0013652 | 0.0457427 | 0.0156079 | 0.0312561 | 0.0592207 | 0.0190352 | 0.0338269 | 0.0248168 | 0.0492385 | 0.0081311 | 0.0517447 | 0.0415218 | 0.003326 | 0.0889339 | 0.0484195 |
| 24.56 | 0.1042295 | 0.1137108 | 0.025383 | 0.0136615 | 0.0025681 | 0.0503766 | 0.0203905 | 0.0666991 | 0.0736213 | 0.0285169 | 0.0458694 | 0.0238178 | 0.0623423 | 0.0091195 | 0.0785769 | 0.058828 | 0.0105482 | 0.1032545 | 0.0574856 |
| 24.57 | 0.0628232 | 0.0624157 | 0.0405002 | 0.0243797 | 0.0057755 | 0.0243846 | 0.0188822 | 0.0960671 | 0.0603821 | 0.0235766 | 0.0304863 | 0.0166789 | 0.0442618 | 0.0079846 | 0.0806546 | 0.0526902 | 0.0192407 | 0.0589767 | 0.0358348 |
| 24.58 | 0.0633602 | 0.0648684 | 0.0634294 | 0.0495539 | 0.0132648 | 0.0121868 | 0.025061 | 0.1700552 | 0.0952992 | 0.0246029 | 0.0242446 | 0.0329815 | 0.0471474 | 0.0148733 | 0.0961253 | 0.0629596 | 0.0288017 | 0.0391176 | 0.0322518 |
| 24.6 | 0.0467918 | 0.053351 | 0.0518193 | 0.0426245 | 0.0140601 | 0.0063444 | 0.0193999 | 0.1489212 | 0.0837766 | 0.0180036 | 0.014368 | 0.0311173 | 0.0339172 | 0.0136251 | 0.065647 | 0.0458581 | 0.0213284 | 0.0166312 | 0.0209137 |
| 24.61 | 0.0245269 | 0.0330763 | 0.0258558 | 0.0177789 | 0.0138374 | 0.0136989 | 0.009207 | 0.0650348 | 0.0389096 | 0.0104308 | 0.0046557 | 0.0177452 | 0.0155106 | 0.0080454 | 0.025763 | 0.021929 | 0.0090014 | 0.0074847 | 0.0073944 |
| 24.63 | 0.028343 | 0.0219605 | 0.0158591 | 0.0145214 | 0.0131443 | 0.0205183 | 0.0062411 | 0.0361065 | 0.0231412 | 0.0081035 | 0.0024052 | 0.012399 | 0.0105014 | 0.0058373 | 0.0171551 | 0.0160538 | 0.0064212 | 0.0143569 | 0.003883 |
| 24.64 | 0.0335086 | 0.0062655 | 0.0055009 | 0.0179009 | 0.007766 | 0.0196777 | 0.0045515 | 0.026283 | 0.0163626 | 0.0034759 | 0.0010669 | 0.0048221 | 0.0034245 | 0.0021545 | 0.0065013 | 0.0069469 | 0.0035225 | 0.0174045 | 0.0012808 |
| 24.65 | 0.0479137 | 0.0132429 | 0.0127723 | 0.0390331 | 0.0134714 | 0.0216863 | 0.0094989 | 0.0915664 | 0.0494202 | 0.0036422 | 0.003687 | 0.0054996 | 0.002489 | 0.0022665 | 0.0066602 | 0.0084654 | 0.0060583 | 0.0256612 | 0.003415 |
| 24.67 | 0.0338592 | 0.0179611 | 0.0129709 | 0.0372266 | 0.017316 | 0.0149693 | 0.0081402 | 0.1236117 | 0.061706 | 0.0035549 | 0.0055013 | 0.0050645 | 0.0018875 | 0.0022889 | 0.0059065 | 0.008401 | 0.0066121 | 0.0225989 | 0.0045902 |
| 24.68 | 0.0136353 | 0.0151778 | 0.0054857 | 0.0148203 | 0.0177723 | 0.0094664 | 0.0019022 | 0.1007179 | 0.0421378 | 0.0031961 | 0.0054943 | 0.0027805 | 0.0012982 | 0.0017073 | 0.0035075 | 0.0058292 | 0.0047667 | 0.0114111 | 0.003852 |
| 24.69 | 0.0455486 | 0.0244901 | 0.0114516 | 0.0154988 | 0.0287756 | 0.0147212 | 0.002911 | 0.1095515 | 0.0346508 | 0.0063606 | 0.0078579 | 0.0028019 | 0.0022884 | 0.0022532 | 0.0048302 | 0.0083646 | 0.0065126 | 0.0119596 | 0.0051708 |
| 24.71 | 0.0553285 | 0.0256838 | 0.0153469 | 0.0135609 | 0.0238097 | 0.0112719 | 0.0042835 | 0.074726 | 0.0201472 | 0.0067101 | 0.0060271 | 0.00195 | 0.0022322 | 0.0017793 | 0.0046362 | 0.0077985 | 0.0054372 | 0.009359 | 0.0041434 |
| 24.72 | 0.0366075 | 0.025176 | 0.0190807 | 0.0075045 | 0.0083944 | 0.0088278 | 0.004128 | 0.0698592 | 0.0235146 | 0.0062161 | 0.0019462 | 0.0010913 | 0.0017761 | 0.0007369 | 0.0041013 | 0.0066431 | 0.002753 | 0.0066896 | 0.0022629 |
| 24.74 | 0.0232889 | 0.020717 | 0.0215703 | 0.0092243 | 0.0043321 | 0.0120357 | 0.0040782 | 0.0967017 | 0.0350053 | 0.0057953 | 0.0015202 | 0.0030656 | 0.0016217 | 0.0005131 | 0.0034399 | 0.0063162 | 0.001801 | 0.0069389 | 0.0022752 |
| 24.75 | 0.0157011 | 0.0064233 | 0.017035 | 0.0111114 | 0.0031629 | 0.0144052 | 0.0039917 | 0.0856749 | 0.0352745 | 0.0034682 | 0.0025754 | 0.0112808 | 0.0009966 | 0.0004319 | 0.0010895 | 0.0040091 | 0.0007154 | 0.0054804 | 0.0024618 |
| 24.76 | 0.0519269 | 0.0046158 | 0.0188197 | 0.0203706 | 0.0062348 | 0.0283863 | 0.0072988 | 0.0995998 | 0.0489186 | 0.0047738 | 0.0067688 | 0.0297262 | 0.0014655 | 0.0010625 | 0.0004484 | 0.0054996 | 0.0013099 | 0.0076518 | 0.0049994 |
| 24.78 | 0.0615943 | 0.006338 | 0.0129449 | 0.0218225 | 0.0053299 | 0.0241412 | 0.0065759 | 0.0696834 | 0.0356448 | 0.0046502 | 0.0067073 | 0.0256916 | 0.0015505 | 0.0014685 | 0.0003396 | 0.0050413 | 0.0017645 | 0.0062327 | 0.0051674 |
| 24.79 | 0.046636 | 0.0217011 | 0.0047759 | 0.0204563 | 0.0019989 | 0.0128909 | 0.0070883 | 0.0646681 | 0.026532 | 0.0050742 | 0.0033051 | 0.0107905 | 0.0020591 | 0.0026239 | 0.0010606 | 0.0042088 | 0.0029739 | 0.0024064 | 0.0046391 |
| 24.81 | 0.0310027 | 0.0365864 | 0.0048849 | 0.0162897 | 0.001611 | 0.0158747 | 0.0096664 | 0.095944 | 0.0396706 | 0.0055565 | 0.0021809 | 0.0087118 | 0.0023626 | 0.0028558 | 0.0017779 | 0.0038358 | 0.003315 | 0.0014439 | 0.0042469 |
| 24.82 | 0.0056501 | 0.0401657 | 0.007874 | 0.0054539 | 0.0011474 | 0.011288 | 0.0071691 | 0.0928599 | 0.0386315 | 0.0039482 | 0.0010489 | 0.004897 | 0.0016021 | 0.0012967 | 0.0016681 | 0.0017648 | 0.0018332 | 0.001731 | 0.0022376 |
| 24.83 | 0.0021944 | 0.0414183 | 0.0198184 | 0.0045055 | 0.0018462 | 0.0125386 | 0.010815 | 0.1182306 | 0.044899 | 0.0048188 | 0.0017654 | 0.0046737 | 0.0015179 | 0.0012291 | 0.0016782 | 0.0012177 | 0.0015147 | 0.0048053 | 0.0023413 |
| 24.85 | 0.0026208 | 0.0276739 | 0.0219489 | 0.0043878 | 0.0033215 | 0.0202746 | 0.0151525 | 0.0888231 | 0.0333101 | 0.0038836 | 0.0034672 | 0.0034655 | 0.0009061 | 0.0017461 | 0.0010163 | 0.0009551 | 0.0014818 | 0.0050936 | 0.0020582 |
| 24.86 | 0.00807 | 0.0454734 | 0.016073 | 0.0051961 | 0.0078273 | 0.0439769 | 0.0253285 | 0.0328401 | 0.0176367 | 0.0023636 | 0.0074834 | 0.0020911 | 0.0002386 | 0.0030826 | 0.0004135 | 0.0009729 | 0.0026667 | 0.0027937 | 0.0025347 |
| 24.88 | 0.0167811 | 0.0588718 | 0.011154 | 0.0077136 | 0.0095933 | 0.0508788 | 0.0259455 | 0.016215 | 0.01226 | 0.0019492 | 0.006724 | 0.0015828 | 0.0002563 | 0.0031616 | 0.0006407 | 0.000869 | 0.0030809 | 0.0015727 | 0.0030045 |
| 24.89 | 0.0314673 | 0.0393611 | 0.0059059 | 0.0176864 | 0.0083725 | 0.0367626 | 0.0127245 | 0.0024607 | 0.0032208 | 0.0013647 | 0.0021091 | 0.0004706 | 0.000338 | 0.0015735 | 0.0009542 | 0.0009111 | 0.0021072 | 0.0009691 | 0.002295 |
| 24.9 | 0.0905205 | 0.0491647 | 0.0099681 | 0.0579797 | 0.0215161 | 0.0524533 | 0.0085598 | 0.0017583 | 0.003178 | 0.0036392 | 0.0032721 | 0.0002143 | 0.0004674 | 0.0013382 | 0.0017893 | 0.0027688 | 0.0025523 | 0.0024607 | 0.0031872 |
| 24.92 | 0.1065802 | 0.045165 | 0.0094172 | 0.0695222 | 0.0302816 | 0.0466011 | 0.0043123 | 0.0020163 | 0.0028162 | 0.0050027 | 0.0033705 | 0.0003562 | 0.0005665 | 0.0012404 | 0.0014661 | 0.0031887 | 0.0019414 | 0.0029778 | 0.0029773 |
| 24.93 | 0.0694993 | 0.0256785 | 0.0081794 | 0.0454284 | 0.0301363 | 0.0188088 | 0.0015519 | 0.0019292 | 0.0014434 | 0.0046826 | 0.0023797 | 0.0007198 | 0.0009472 | 0.0008449 | 0.0004996 | 0.0018151 | 0.0007254 | 0.002553 | 0.0017424 |
| 24.94 | 0.07663 | 0.0316198 | 0.019712 | 0.0493726 | 0.0505169 | 0.011433 | 0.0049935 | 0.0027274 | 0.004812 | 0.0068117 | 0.0074713 | 0.0011251 | 0.0016983 | 0.0017101 | 0.0012287 | 0.0016327 | 0.0016147 | 0.0041905 | 0.002092 |
| 24.96 | 0.0571191 | 0.0281417 | 0.0211256 | 0.0365155 | 0.0464567 | 0.0103892 | 0.0059446 | 0.0032178 | 0.0062908 | 0.0054101 | 0.0087576 | 0.0008885 | 0.0014885 | 0.0028296 | 0.0013303 | 0.0012569 | 0.0025314 | 0.0038701 | 0.0019464 |
| 24.97 | 0.0219941 | 0.0202895 | 0.0178979 | 0.0169788 | 0.0236018 | 0.0257747 | 0.0035947 | 0.0066853 | 0.0059539 | 0.0018036 | 0.0060111 | 0.0010275 | 0.0024682 | 0.0045315 | 0.0007536 | 0.001207 | 0.0046926 | 0.0026042 | 0.0026876 |
| 24.99 | 0.0134453 | 0.0164249 | 0.016505 | 0.0169557 | 0.0132172 | 0.0324117 | 0.0022521 | 0.0075066 | 0.0043769 | 0.001217 | 0.0037366 | 0.0012942 | 0.0036784 | 0.0038846 | 0.0007773 | 0.0012078 | 0.0050748 | 0.0027293 | 0.0034902 |
| 25 | 0.0086242 | 0.0074703 | 0.0113963 | 0.0163109 | 0.0047975 | 0.022572 | 0.0018673 | 0.0041098 | 0.0014689 | 0.0018838 | 0.0013331 | 0.0007404 | 0.0035314 | 0.0012324 | 0.0011186 | 0.0006808 | 0.0025036 | 0.0022047 | 0.0027988 |
| 25.01 | 0.0138732 | 0.011774 | 0.0213091 | 0.0214576 | 0.0077699 | 0.0269726 | 0.0035325 | 0.00333 | 0.0021594 | 0.005569 | 0.0109271 | 0.001153 | 0.0107405 | 0.0030241 | 0.0087879 | 0.005314 | 0.0040621 | 0.013351 | 0.0064225 |
| 25.03 | 0.0238262 | 0.0216485 | 0.0252875 | 0.0161722 | 0.0120525 | 0.0229329 | 0.0038234 | 0.0040708 | 0.0047398 | 0.0116938 | 0.0224599 | 0.0014479 | 0.0217669 | 0.0049931 | 0.0241294 | 0.0170597 | 0.0052475 | 0.0327974 | 0.0152816 |
| 25.04 | 0.0798241 | 0.0575268 | 0.0329953 | 0.0232006 | 0.0321827 | 0.0379975 | 0.0130586 | 0.0159388 | 0.0245156 | 0.0341576 | 0.046809 | 0.0088304 | 0.0602315 | 0.006986 | 0.0818194 | 0.0666469 | 0.0154113 | 0.0780496 | 0.0547531 |
| 25.06 | 0.1006705 | 0.063617 | 0.0328738 | 0.039872 | 0.03662 | 0.0582302 | 0.022653 | 0.0207908 | 0.0361647 | 0.0411707 | 0.0505052 | 0.0185293 | 0.0711283 | 0.0090464 | 0.1018358 | 0.0857866 | 0.0256351 | 0.0777312 | 0.068593 |
| 25.07 | 0.0591838 | 0.0288775 | 0.0173053 | 0.0374184 | 0.0175297 | 0.0569149 | 0.0234222 | 0.0107023 | 0.0281267 | 0.0243745 | 0.024801 | 0.017202 | 0.034625 | 0.0093701 | 0.0531331 | 0.0466168 | 0.0205173 | 0.0261175 | 0.0365221 |
| 25.08 | 0.0466451 | 0.0197889 | 0.0166157 | 0.0364393 | 0.0132323 | 0.0799104 | 0.0312345 | 0.006764 | 0.0298818 | 0.0178173 | 0.0151948 | 0.0127842 | 0.0209934 | 0.0096828 | 0.0338777 | 0.0286234 | 0.0155904 | 0.0097696 | 0.0184185 |
| 25.1 | 0.027125 | 0.0132377 | 0.0111517 | 0.0237955 | 0.0091391 | 0.0630832 | 0.0243435 | 0.006295 | 0.0197592 | 0.0121273 | 0.0092359 | 0.0108363 | 0.0187293 | 0.0079133 | 0.0313504 | 0.0232555 | 0.0119626 | 0.0075332 | 0.0076287 |
| 25.11 | 0.0097742 | 0.006707 | 0.0039064 | 0.008075 | 0.0035757 | 0.0209957 | 0.0091694 | 0.0082203 | 0.0057507 | 0.0152733 | 0.0086468 | 0.0177797 | 0.0261289 | 0.0126997 | 0.0388733 | 0.0286787 | 0.0148806 | 0.0109298 | 0.0072614 |
| 25.13 | 0.0121768 | 0.0056146 | 0.0039375 | 0.0041642 | 0.0018592 | 0.0089949 | 0.0054084 | 0.0063724 | 0.0053796 | 0.0164507 | 0.0086452 | 0.020392 | 0.0244718 | 0.0134923 | 0.0332405 | 0.0262566 | 0.0146925 | 0.0101049 | 0.0082535 |
| 25.14 | 0.0130477 | 0.0065264 | 0.0041496 | 0.0030914 | 0.0010505 | 0.003139 | 0.0049614 | 0.0023015 | 0.0067423 | 0.0106158 | 0.0060714 | 0.016019 | 0.0112445 | 0.0068802 | 0.0161655 | 0.0149797 | 0.0087035 | 0.0056764 | 0.0054679 |
| 25.15 | 0.0185327 | 0.0139602 | 0.0072769 | 0.0123722 | 0.0022459 | 0.0051732 | 0.0117959 | 0.0047494 | 0.0138967 | 0.015306 | 0.0103074 | 0.0238458 | 0.0092857 | 0.0077587 | 0.0188128 | 0.0201761 | 0.0131155 | 0.0082089 | 0.0081987 |
| 25.17 | 0.0177465 | 0.0117799 | 0.0079193 | 0.0184949 | 0.0028331 | 0.0048053 | 0.0129041 | 0.0043122 | 0.0143306 | 0.0137125 | 0.0100768 | 0.0205042 | 0.0069938 | 0.0064811 | 0.0156035 | 0.0170426 | 0.0121637 | 0.0079889 | 0.0072255 |
| 25.18 | 0.010505 | 0.0029016 | 0.0051537 | 0.0188713 | 0.0019598 | 0.0050016 | 0.0079396 | 0.0011098 | 0.0069893 | 0.0050339 | 0.0052518 | 0.0068997 | 0.0052552 | 0.0022051 | 0.0089722 | 0.0076668 | 0.0048655 | 0.0060282 | 0.0036082 |
| 25.19 | 0.0107622 | 0.002408 | 0.0046129 | 0.030683 | 0.0013072 | 0.0134701 | 0.0069659 | 0.0015691 | 0.0041238 | 0.0021297 | 0.0048899 | 0.0024834 | 0.0110593 | 0.0014777 | 0.0165367 | 0.0115847 | 0.0023625 | 0.0124038 | 0.0067095 |
| 25.21 | 0.0111967 | 0.003059 | 0.0034416 | 0.0290188 | 0.0014329 | 0.0180742 | 0.0041911 | 0.0021353 | 0.0027326 | 0.0013694 | 0.0032543 | 0.0012836 | 0.0105229 | 0.0018389 | 0.0147035 | 0.0114969 | 0.0012013 | 0.011786 | 0.0063981 |
| 25.22 | 0.0269947 | 0.0080834 | 0.0036236 | 0.0254953 | 0.0075828 | 0.033241 | 0.0041566 | 0.0049244 | 0.0100222 | 0.003894 | 0.0017275 | 0.0031315 | 0.0056594 | 0.0028237 | 0.0047346 | 0.0078809 | 0.0010656 | 0.0043883 | 0.0023313 |
| 25.24 | 0.0387984 | 0.0132807 | 0.0050824 | 0.0255753 | 0.0139024 | 0.0417053 | 0.0075396 | 0.0076237 | 0.0181221 | 0.0061286 | 0.0023863 | 0.0033489 | 0.005176 | 0.0026923 | 0.0031773 | 0.007601 | 0.0012556 | 0.002327 | 0.0024139 |
| 25.25 | 0.0331569 | 0.0132265 | 0.0067216 | 0.0175485 | 0.015721 | 0.0325028 | 0.0091715 | 0.007309 | 0.0195333 | 0.0061914 | 0.002914 | 0.0012899 | 0.0049519 | 0.0011568 | 0.0038704 | 0.007525 | 0.0011417 | 0.0026717 | 0.0046037 |
| 25.26 | 0.0318446 | 0.0124312 | 0.0101585 | 0.020026 | 0.0188584 | 0.039334 | 0.0119534 | 0.0075234 | 0.0234025 | 0.0103528 | 0.0049355 | 0.0026931 | 0.0072411 | 0.0009277 | 0.0070445 | 0.0131484 | 0.0029388 | 0.0064476 | 0.0079352 |
| 25.28 | 0.0200517 | 0.007291 | 0.0072403 | 0.0142797 | 0.0130201 | 0.0299637 | 0.0081583 | 0.0045455 | 0.0152051 | 0.0104702 | 0.0046241 | 0.0048829 | 0.0058456 | 0.0010728 | 0.0062779 | 0.0115778 | 0.0045054 | 0.0058791 | 0.0064165 |
| 25.29 | 0.01912 | 0.0080797 | 0.004018 | 0.0110275 | 0.0105303 | 0.0127828 | 0.003503 | 0.002215 | 0.004123 | 0.0101226 | 0.0030214 | 0.0099622 | 0.0033266 | 0.0010293 | 0.006688 | 0.0055756 | 0.0054007 | 0.0027669 | 0.0058189 |
| 25.31 | 0.0191609 | 0.0092284 | 0.0043704 | 0.0122287 | 0.0115561 | 0.0079623 | 0.0033268 | 0.0025667 | 0.0036326 | 0.0102534 | 0.0026006 | 0.0098018 | 0.0035195 | 0.0008272 | 0.0073829 | 0.0044106 | 0.004116 | 0.0020255 | 0.0060367 |
| 25.32 | 0.0085766 | 0.0044972 | 0.0031282 | 0.0070455 | 0.0050356 | 0.0040033 | 0.0025925 | 0.0034692 | 0.0055368 | 0.0073915 | 0.0026381 | 0.0039368 | 0.0031272 | 0.0006092 | 0.0033578 | 0.0050095 | 0.0016759 | 0.0012347 | 0.0027668 |
| 25.33 | 0.0215845 | 0.0077474 | 0.0078024 | 0.0132867 | 0.0034811 | 0.0066169 | 0.008554 | 0.0091802 | 0.0175093 | 0.0111013 | 0.0059139 | 0.0061231 | 0.0049399 | 0.0013765 | 0.0029619 | 0.0121263 | 0.0021188 | 0.0033233 | 0.0038919 |
| 25.35 | 0.0272695 | 0.0087584 | 0.010051 | 0.0171501 | 0.0047455 | 0.0059974 | 0.01137 | 0.0093624 | 0.0199196 | 0.0106324 | 0.0063095 | 0.0064624 | 0.0069704 | 0.0024514 | 0.0029105 | 0.0135206 | 0.0026386 | 0.0039113 | 0.0036433 |
| 25.36 | 0.0205703 | 0.0062255 | 0.0078793 | 0.0156173 | 0.0069705 | 0.0078877 | 0.0108483 | 0.0045079 | 0.0127012 | 0.0108827 | 0.0062281 | 0.0042005 | 0.0106576 | 0.0049908 | 0.0035922 | 0.0133068 | 0.0058592 | 0.0038452 | 0.0010744 |
| 25.38 | 0.0127118 | 0.0049384 | 0.0049806 | 0.010222 | 0.0055767 | 0.0118018 | 0.0083378 | 0.0022368 | 0.0070794 | 0.0125002 | 0.006703 | 0.003498 | 0.0103636 | 0.0052139 | 0.0057802 | 0.013083 | 0.00619 | 0.0048971 | 0.0017338 |
| 25.39 | 0.0030364 | 0.0030328 | 0.0018447 | 0.0019982 | 0.0021834 | 0.0095968 | 0.0030769 | 0.0011083 | 0.0020086 | 0.0113717 | 0.0064953 | 0.0027761 | 0.01026 | 0.0025967 | 0.0099271 | 0.010816 | 0.0025767 | 0.0070282 | 0.0067332 |
| 25.4 | 0.0068066 | 0.0071445 | 0.0029057 | 0.0046772 | 0.002926 | 0.0072033 | 0.0048801 | 0.0049815 | 0.0107078 | 0.0197513 | 0.0124916 | 0.0073667 | 0.0222652 | 0.0022877 | 0.0212009 | 0.0209869 | 0.0028633 | 0.0156983 | 0.0199956 |
| 25.42 | 0.0109225 | 0.0090278 | 0.0039558 | 0.0078746 | 0.0038173 | 0.0037233 | 0.0061012 | 0.0068185 | 0.0168056 | 0.0184808 | 0.011278 | 0.0081677 | 0.0191904 | 0.0024858 | 0.017627 | 0.020185 | 0.0025749 | 0.0137971 | 0.0194137 |
| 25.43 | 0.010553 | 0.0059125 | 0.0055116 | 0.0080959 | 0.0057342 | 0.0015408 | 0.005436 | 0.0049696 | 0.014548 | 0.0089327 | 0.0046596 | 0.0042648 | 0.0075955 | 0.0033704 | 0.0036768 | 0.0097784 | 0.0016036 | 0.0036583 | 0.0066979 |
| 25.44 | 0.0131726 | 0.009118 | 0.0126121 | 0.0106161 | 0.0146397 | 0.0041938 | 0.0098141 | 0.0072355 | 0.0187405 | 0.0122272 | 0.0070362 | 0.004474 | 0.0125036 | 0.0080295 | 0.0028469 | 0.0127403 | 0.0056496 | 0.0030279 | 0.0052279 |

| | | | | | | | | | | | | | | | | | | | |
|---|---|---|---|---|---|---|---|---|---|---|---|---|---|---|---|---|---|---|---|
| 25.46 | 0.0098809 | 0.0104205 | 0.0130932 | 0.0075293 | 0.0157248 | 0.0058304 | 0.0101869 | 0.007175 | 0.0146834 | 0.0144032 | 0.0070565 | 0.0064642 | 0.011179 | 0.0094599 | 0.0044963 | 0.011154 | 0.0086445 | 0.0038785 | 0.0042396 |
| 25.47 | 0.0031865 | 0.0096249 | 0.0074495 | 0.0021521 | 0.0091145 | 0.0073354 | 0.0071894 | 0.0048104 | 0.0095938 | 0.0214745 | 0.0040884 | 0.0193425 | 0.006145 | 0.0116258 | 0.0084596 | 0.0039684 | 0.0130078 | 0.0066783 | 0.0030538 |
| 25.49 | 0.0023123 | 0.0089448 | 0.0048994 | 0.0026745 | 0.0051976 | 0.0077964 | 0.0047403 | 0.0037203 | 0.0114574 | 0.023386 | 0.003393 | 0.0229551 | 0.0079794 | 0.0114986 | 0.0086126 | 0.0016897 | 0.0127151 | 0.0064697 | 0.0054468 |
| 25.5 | 0.0047496 | 0.0052639 | 0.0028337 | 0.0052692 | 0.002686 | 0.0082378 | 0.0022371 | 0.0021588 | 0.0090473 | 0.0136228 | 0.0030038 | 0.0114911 | 0.0089885 | 0.0057258 | 0.0042249 | 0.0017538 | 0.0065454 | 0.0025647 | 0.0090057 |
| 25.51 | 0.0182924 | 0.012995 | 0.0061189 | 0.0096532 | 0.0035794 | 0.0221137 | 0.0085991 | 0.0047831 | 0.021766 | 0.0134892 | 0.0095353 | 0.0100736 | 0.0148984 | 0.0041325 | 0.0196186 | 0.0123884 | 0.0092514 | 0.0141838 | 0.0243639 |
| 25.53 | 0.0276185 | 0.0289741 | 0.0118153 | 0.0084366 | 0.0047815 | 0.0304999 | 0.0140349 | 0.0101714 | 0.0367401 | 0.0137406 | 0.0188724 | 0.0122691 | 0.0231364 | 0.0023124 | 0.040815 | 0.0247512 | 0.0106172 | 0.0352499 | 0.0400619 |
| 25.54 | 0.0544329 | 0.076926 | 0.0310894 | 0.0069064 | 0.0195986 | 0.0518294 | 0.0246859 | 0.0270415 | 0.0692911 | 0.0357513 | 0.0573427 | 0.0180165 | 0.0588532 | 0.0042956 | 0.0806112 | 0.05134 | 0.0176127 | 0.1082251 | 0.0884485 |
| 25.56 | 0.0675292 | 0.0872645 | 0.0428351 | 0.0106982 | 0.0292317 | 0.0576115 | 0.0287288 | 0.0293809 | 0.0710777 | 0.0524342 | 0.0722632 | 0.0254963 | 0.0695341 | 0.0101717 | 0.0820269 | 0.055987 | 0.0243663 | 0.1293765 | 0.1012077 |
| 25.57 | 0.0529339 | 0.0506307 | 0.0438416 | 0.0205815 | 0.0235117 | 0.0342604 | 0.0226545 | 0.0154187 | 0.0331217 | 0.0434618 | 0.0405549 | 0.0364048 | 0.0426026 | 0.0153986 | 0.0456279 | 0.0328096 | 0.0247535 | 0.0680363 | 0.0587456 |
| 25.58 | 0.0688503 | 0.0630113 | 0.0712729 | 0.0473698 | 0.0225299 | 0.0340401 | 0.0322118 | 0.0271781 | 0.0272473 | 0.0514698 | 0.0238485 | 0.0583462 | 0.0517008 | 0.0250581 | 0.0584916 | 0.0337298 | 0.0382881 | 0.0537167 | 0.0555956 |
| 25.6 | 0.0500832 | 0.0579281 | 0.0608578 | 0.0412163 | 0.0141588 | 0.0229523 | 0.0264264 | 0.0351494 | 0.0197914 | 0.0401671 | 0.010334 | 0.0472183 | 0.045699 | 0.0219657 | 0.0554219 | 0.0267608 | 0.0321566 | 0.0411462 | 0.0408729 |
| 25.61 | 0.0259972 | 0.0388054 | 0.024129 | 0.0224284 | 0.0138463 | 0.0114288 | 0.0112354 | 0.047074 | 0.0093862 | 0.0174534 | 0.0018169 | 0.0222864 | 0.0259878 | 0.0150026 | 0.0471276 | 0.0187684 | 0.0141933 | 0.029786 | 0.0214125 |
| 25.63 | 0.0383814 | 0.0280403 | 0.0139191 | 0.0288484 | 0.0211515 | 0.0173402 | 0.0087442 | 0.0491177 | 0.0075248 | 0.0127376 | 0.0024167 | 0.0134227 | 0.0158066 | 0.0116172 | 0.0379133 | 0.0148846 | 0.0093997 | 0.0223899 | 0.0141872 |
| 25.64 | 0.0419147 | 0.0093589 | 0.0147829 | 0.0319662 | 0.0191252 | 0.0202181 | 0.0077773 | 0.0338361 | 0.0066424 | 0.0122561 | 0.0052893 | 0.0061994 | 0.0069898 | 0.004126 | 0.0121194 | 0.0048445 | 0.0049182 | 0.0075528 | 0.003608 |
| 25.65 | 0.0558999 | 0.0093023 | 0.0286202 | 0.051363 | 0.0203492 | 0.0287487 | 0.008832 | 0.0478721 | 0.0083923 | 0.0203833 | 0.0121041 | 0.00844 | 0.0204731 | 0.0032686 | 0.0096454 | 0.0051992 | 0.0086751 | 0.0067238 | 0.0035927 |
| 25.67 | 0.042813 | 0.0143448 | 0.0212704 | 0.0457404 | 0.0133553 | 0.0266149 | 0.0051892 | 0.0424687 | 0.0099868 | 0.0174089 | 0.0112612 | 0.0076324 | 0.0248986 | 0.0028795 | 0.0148532 | 0.0093094 | 0.0138915 | 0.0049359 | 0.0068938 |
| 25.68 | 0.0161397 | 0.0211762 | 0.0038876 | 0.0176688 | 0.0049132 | 0.013973 | 0.0013423 | 0.0181584 | 0.0160999 | 0.0076134 | 0.0069454 | 0.0085532 | 0.0137415 | 0.0022528 | 0.0204512 | 0.0129321 | 0.0127724 | 0.0024739 | 0.0101413 |
| 25.69 | 0.0438816 | 0.0454656 | 0.0071001 | 0.0249026 | 0.014717 | 0.0091876 | 0.0022886 | 0.0251708 | 0.0319962 | 0.0092757 | 0.0178912 | 0.0187142 | 0.0100113 | 0.0040542 | 0.0362012 | 0.0246395 | 0.0098611 | 0.0077209 | 0.0178275 |
| 25.71 | 0.0540104 | 0.0473882 | 0.0090395 | 0.0297107 | 0.019005 | 0.0062232 | 0.001988 | 0.0371396 | 0.0254868 | 0.0115648 | 0.0199663 | 0.0160609 | 0.0086951 | 0.0072106 | 0.0341834 | 0.024328 | 0.0085861 | 0.009756 | 0.0175792 |
| 25.72 | 0.0323523 | 0.0333942 | 0.0125073 | 0.0210965 | 0.0166648 | 0.0085198 | 0.0017915 | 0.0632442 | 0.0111598 | 0.0215527 | 0.01258 | 0.0076181 | 0.0119288 | 0.0179322 | 0.0204301 | 0.0137406 | 0.0231512 | 0.0087996 | 0.0165034 |
| 25.74 | 0.0206315 | 0.0218357 | 0.0199582 | 0.0138758 | 0.0113242 | 0.0066435 | 0.0019075 | 0.0668787 | 0.0156547 | 0.0243223 | 0.007751 | 0.0115519 | 0.010136 | 0.0216912 | 0.0133706 | 0.0099911 | 0.0302021 | 0.0066816 | 0.0146676 |
| 25.75 | 0.0222992 | 0.0088029 | 0.0188593 | 0.0121463 | 0.0032425 | 0.0021583 | 0.0009963 | 0.0378557 | 0.0200706 | 0.0122142 | 0.0032475 | 0.0170358 | 0.0042034 | 0.0139965 | 0.0079646 | 0.0105873 | 0.0165297 | 0.0020786 | 0.0063543 |
| 25.76 | 0.0636796 | 0.0320158 | 0.0152739 | 0.0358875 | 0.0138327 | 0.0133613 | 0.0018795 | 0.0309771 | 0.0317855 | 0.0099647 | 0.0054735 | 0.0269993 | 0.0106764 | 0.0112145 | 0.0193022 | 0.0235452 | 0.0121323 | 0.0010978 | 0.0118792 |
| 25.78 | 0.0726573 | 0.0448911 | 0.0099646 | 0.0441456 | 0.0233051 | 0.0207221 | 0.0031393 | 0.018085 | 0.0252218 | 0.0130769 | 0.0046556 | 0.0210498 | 0.0116758 | 0.0088936 | 0.0204461 | 0.0234057 | 0.0158138 | 0.0005564 | 0.0167393 |
| 25.79 | 0.0544208 | 0.0528313 | 0.0162735 | 0.0443663 | 0.0296252 | 0.0333202 | 0.0073176 | 0.0109628 | 0.0124029 | 0.0266574 | 0.0021014 | 0.010698 | 0.0057966 | 0.0151646 | 0.0136367 | 0.0141688 | 0.0323324 | 0.0007809 | 0.0186645 |
| 25.81 | 0.0356938 | 0.0466527 | 0.0183397 | 0.0357111 | 0.0222589 | 0.0410073 | 0.0078083 | 0.0145868 | 0.0166069 | 0.0287444 | 0.002246 | 0.0138258 | 0.0042766 | 0.0160803 | 0.0090055 | 0.0087535 | 0.0309392 | 0.0009272 | 0.0137626 |
| 25.82 | 0.0160912 | 0.0163922 | 0.0088582 | 0.0124104 | 0.0068559 | 0.0312896 | 0.0033441 | 0.0118962 | 0.019419 | 0.0156629 | 0.0036937 | 0.0150692 | 0.0048777 | 0.0063139 | 0.0040746 | 0.0044563 | 0.0102658 | 0.0003892 | 0.0033108 |
| 25.83 | 0.0285989 | 0.0126998 | 0.0143888 | 0.0105296 | 0.0165496 | 0.035251 | 0.0028625 | 0.0091983 | 0.0253946 | 0.0114985 | 0.0074027 | 0.0196564 | 0.0080111 | 0.0059694 | 0.0129786 | 0.0138759 | 0.0105668 | 0.0007301 | 0.003356 |
| 25.85 | 0.0237247 | 0.0201728 | 0.019026 | 0.0114561 | 0.0212564 | 0.0281477 | 0.0050218 | 0.0058194 | 0.0186099 | 0.0084358 | 0.0066352 | 0.0141003 | 0.006249 | 0.0084148 | 0.0189115 | 0.0182133 | 0.0146468 | 0.0013761 | 0.0044167 |
| 25.86 | 0.0159427 | 0.0474054 | 0.0285453 | 0.0197202 | 0.0214706 | 0.0261367 | 0.0210624 | 0.0123393 | 0.0175229 | 0.0199711 | 0.0087223 | 0.0107559 | 0.0060736 | 0.0121851 | 0.0312331 | 0.0264008 | 0.0182357 | 0.0027854 | 0.007329 |
| 25.88 | 0.0185933 | 0.0521367 | 0.0322141 | 0.0216408 | 0.0180199 | 0.0289067 | 0.0289149 | 0.0168729 | 0.0275773 | 0.0283798 | 0.0129091 | 0.0182798 | 0.0087151 | 0.0105344 | 0.0360089 | 0.0314066 | 0.0134658 | 0.002739 | 0.009414 |
| 25.89 | 0.0160731 | 0.0259646 | 0.0225274 | 0.0133557 | 0.006535 | 0.0190168 | 0.0189893 | 0.0113302 | 0.0284016 | 0.0255107 | 0.0120977 | 0.020459 | 0.0082232 | 0.0045806 | 0.0258662 | 0.0250687 | 0.0025417 | 0.0013011 | 0.0094289 |
| 25.9 | 0.0305051 | 0.0163613 | 0.0311644 | 0.016923 | 0.0123598 | 0.0232874 | 0.0174025 | 0.0083093 | 0.0345423 | 0.0426622 | 0.0148336 | 0.0263253 | 0.0107445 | 0.0100416 | 0.0345711 | 0.0333012 | 0.0040773 | 0.0034842 | 0.0172523 |
| 25.92 | 0.0343692 | 0.0079114 | 0.0303795 | 0.0167302 | 0.0240513 | 0.0220893 | 0.0135745 | 0.0040127 | 0.0237786 | 0.0426384 | 0.0116834 | 0.0215367 | 0.0087541 | 0.0149818 | 0.0308091 | 0.026225 | 0.0069662 | 0.0048271 | 0.017819 |
| 25.93 | 0.0231336 | 0.0020103 | 0.0167287 | 0.0135594 | 0.0296253 | 0.0134197 | 0.0082653 | 0.0012127 | 0.0058365 | 0.0253992 | 0.0068883 | 0.0095323 | 0.003536 | 0.0148425 | 0.0158491 | 0.0076047 | 0.0074817 | 0.0028625 | 0.0125098 |
| 25.94 | 0.0268098 | 0.0060591 | 0.0113201 | 0.0259463 | 0.0335415 | 0.0112947 | 0.0128543 | 0.0014449 | 0.0086859 | 0.0282138 | 0.0158577 | 0.0083982 | 0.0027171 | 0.0215454 | 0.0159388 | 0.0033882 | 0.0109342 | 0.0029757 | 0.0208339 |
| 25.96 | 0.0230067 | 0.0085867 | 0.0062262 | 0.0283815 | 0.0217155 | 0.0081889 | 0.0115754 | 0.0011845 | 0.0097842 | 0.0206335 | 0.0189546 | 0.0059547 | 0.0020759 | 0.0188645 | 0.0107684 | 0.0021964 | 0.0092412 | 0.0046699 | 0.0213328 |
| 25.97 | 0.018383 | 0.0115598 | 0.0082302 | 0.0252271 | 0.0150328 | 0.0194031 | 0.0053673 | 0.0029641 | 0.0070541 | 0.0076919 | 0.0157313 | 0.0024097 | 0.0023887 | 0.0128674 | 0.003119 | 0.0026578 | 0.0053289 | 0.0088904 | 0.0184808 |
| 25.99 | 0.0139124 | 0.0117956 | 0.0095636 | 0.0184557 | 0.0172006 | 0.0286959 | 0.0036333 | 0.0046487 | 0.0058099 | 0.0060273 | 0.0112623 | 0.0017514 | 0.0023098 | 0.0102971 | 0.0027121 | 0.004558 | 0.0042816 | 0.008584 | 0.0146761 |
| 26 | 0.0050059 | 0.0070851 | 0.0044791 | 0.0052369 | 0.0098934 | 0.0250086 | 0.0032237 | 0.0048627 | 0.0042154 | 0.0048992 | 0.0055418 | 0.0017331 | 0.0015384 | 0.0039194 | 0.0025214 | 0.006171 | 0.0022324 | 0.0044131 | 0.0045266 |
| 26.01 | 0.01014 | 0.0143135 | 0.0027583 | 0.0110553 | 0.007864 | 0.0261302 | 0.0053158 | 0.0125118 | 0.0069805 | 0.0098972 | 0.0154575 | 0.0031611 | 0.0136698 | 0.0022155 | 0.0089744 | 0.0148968 | 0.0039426 | 0.0175625 | 0.0069694 |
| 26.03 | 0.0155484 | 0.0309626 | 0.0040678 | 0.0124878 | 0.0098093 | 0.0229232 | 0.0094869 | 0.019006 | 0.0118033 | 0.019558 | 0.0291138 | 0.0052361 | 0.0315508 | 0.0027219 | 0.0239541 | 0.02373 | 0.0049065 | 0.0378455 | 0.0165724 |
| 26.04 | 0.0519505 | 0.0883775 | 0.0166671 | 0.0159238 | 0.0317643 | 0.0594453 | 0.0361466 | 0.0310976 | 0.0411884 | 0.0487002 | 0.0607553 | 0.0179645 | 0.0769863 | 0.0149002 | 0.077724 | 0.0546998 | 0.0105741 | 0.0936847 | 0.0559095 |
| 26.06 | 0.0784394 | 0.0966099 | 0.0220178 | 0.0285904 | 0.0431101 | 0.0832539 | 0.0467534 | 0.0310363 | 0.0578091 | 0.0554369 | 0.0589626 | 0.0251309 | 0.0832243 | 0.0227971 | 0.0852879 | 0.0618468 | 0.0172652 | 0.1005364 | 0.0686998 |
| 26.07 | 0.06596 | 0.0408657 | 0.0148754 | 0.0365632 | 0.0369646 | 0.0602511 | 0.0285439 | 0.0158075 | 0.049799 | 0.0292117 | 0.0209516 | 0.0183556 | 0.0345716 | 0.0159511 | 0.0357575 | 0.0281883 | 0.0148422 | 0.0391929 | 0.0349209 |
| 26.08 | 0.0652249 | 0.0219241 | 0.0226074 | 0.0531113 | 0.0550776 | 0.0549061 | 0.0358747 | 0.014174 | 0.0714233 | 0.0194294 | 0.0150932 | 0.0142773 | 0.0207667 | 0.0098497 | 0.0437566 | 0.019309 | 0.0116606 | 0.0188261 | 0.0263102 |
| 26.1 | 0.0394682 | 0.012431 | 0.0260642 | 0.0417954 | 0.0431666 | 0.033552 | 0.0316777 | 0.0091669 | 0.0587178 | 0.0156183 | 0.0131589 | 0.007937 | 0.0220134 | 0.0042593 | 0.0469479 | 0.0163294 | 0.0087541 | 0.0150406 | 0.0237198 |
| 26.11 | 0.0100958 | 0.0083852 | 0.022936 | 0.0186102 | 0.0175257 | 0.0084509 | 0.0101919 | 0.0047229 | 0.0249576 | 0.0121088 | 0.0093823 | 0.0105605 | 0.0277454 | 0.0023592 | 0.02713 | 0.0120145 | 0.0110772 | 0.0123563 | 0.0177395 |
| 26.13 | 0.0088405 | 0.0064611 | 0.0144524 | 0.0120842 | 0.0196145 | 0.004394 | 0.0053147 | 0.0060992 | 0.0211412 | 0.0074103 | 0.0086016 | 0.0149696 | 0.0223838 | 0.0023917 | 0.0126376 | 0.0092918 | 0.0108599 | 0.0076341 | 0.0125648 |
| 26.14 | 0.0135707 | 0.0022143 | 0.0036653 | 0.0051557 | 0.0231334 | 0.0040342 | 0.0068288 | 0.0058314 | 0.023834 | 0.0043743 | 0.0053599 | 0.0100448 | 0.0114214 | 0.0013393 | 0.0033797 | 0.005077 | 0.0056731 | 0.0037072 | 0.0064445 |
| 26.15 | 0.0327109 | 0.0024652 | 0.0076459 | 0.0120801 | 0.0343238 | 0.0080625 | 0.0148221 | 0.0063773 | 0.0414114 | 0.0056131 | 0.0047502 | 0.0058673 | 0.0193871 | 0.0019116 | 0.0101831 | 0.0055639 | 0.0064001 | 0.0079472 | 0.0079282 |
| 26.17 | 0.0355588 | 0.0023653 | 0.0082525 | 0.0159829 | 0.0297605 | 0.0078392 | 0.015877 | 0.0050651 | 0.0373782 | 0.0039366 | 0.004398 | 0.0021852 | 0.0167037 | 0.0020889 | 0.0109068 | 0.0052803 | 0.0047522 | 0.0088833 | 0.0079794 |
| 26.18 | 0.0230395 | 0.0012866 | 0.0043523 | 0.0136808 | 0.0164307 | 0.0040651 | 0.0093127 | 0.0034399 | 0.0181049 | 0.0021912 | 0.0036123 | 0.0007269 | 0.0058566 | 0.0016168 | 0.0071185 | 0.0042104 | 0.0019056 | 0.0081019 | 0.0070759 |
| 26.19 | 0.0326306 | 0.0035068 | 0.0051165 | 0.020994 | 0.0229918 | 0.005605 | 0.0101478 | 0.006427 | 0.0205834 | 0.0049567 | 0.0045618 | 0.003808 | 0.0106426 | 0.001776 | 0.0098389 | 0.0052992 | 0.0054053 | 0.0124031 | 0.0096223 |
| 26.21 | 0.0319999 | 0.0072768 | 0.0063909 | 0.0196149 | 0.020237 | 0.0072408 | 0.0089244 | 0.0069376 | 0.0190769 | 0.0045774 | 0.0036014 | 0.0059142 | 0.011039 | 0.0011812 | 0.0085023 | 0.0036674 | 0.0057962 | 0.0083446 | 0.0077117 |
| 26.22 | 0.0293201 | 0.0149984 | 0.0134354 | 0.0161808 | 0.0091051 | 0.0133828 | 0.0115206 | 0.0068749 | 0.0201373 | 0.0039874 | 0.0052462 | 0.0054117 | 0.00506 | 0.0007799 | 0.0062459 | 0.0035314 | 0.0041697 | 0.0010641 | 0.0087214 |
| 26.24 | 0.0264948 | 0.0137135 | 0.016364 | 0.0155857 | 0.0052268 | 0.0157713 | 0.0143512 | 0.0060203 | 0.0221766 | 0.0066949 | 0.0077125 | 0.0030154 | 0.0035656 | 0.0008901 | 0.0064655 | 0.0055404 | 0.0048499 | 0.001467 | 0.0109529 |
| 26.25 | 0.0138748 | 0.0039527 | 0.0108287 | 0.0106667 | 0.0049724 | 0.0113765 | 0.011661 | 0.0025302 | 0.0182137 | 0.007444 | 0.0067866 | 0.0004678 | 0.0035755 | 0.0006188 | 0.0055756 | 0.0056953 | 0.0034043 | 0.0024659 | 0.0085356 |
| 26.26 | 0.0184367 | 0.0042068 | 0.011371 | 0.0166298 | 0.0133255 | 0.0174161 | 0.0160732 | 0.0034895 | 0.031287 | 0.0084164 | 0.0075442 | 0.0009888 | 0.0063996 | 0.0014493 | 0.0090043 | 0.0071645 | 0.0037075 | 0.0071055 | 0.0096956 |
| 26.28 | 0.0184037 | 0.0066028 | 0.0086228 | 0.0167051 | 0.0135825 | 0.018746 | 0.0140315 | 0.0045369 | 0.0308951 | 0.0055034 | 0.0066636 | 0.0014314 | 0.0079863 | 0.002172 | 0.0105261 | 0.0064535 | 0.0033708 | 0.0120427 | 0.0082102 |
| 26.29 | 0.0160502 | 0.0106471 | 0.0040609 | 0.0150556 | 0.0079409 | 0.0209709 | 0.0084722 | 0.005919 | 0.022836 | 0.0019257 | 0.0076123 | 0.0044877 | 0.0125958 | 0.0030558 | 0.0103612 | 0.0052805 | 0.0023953 | 0.0158313 | 0.0058162 |
| 26.31 | 0.0132196 | 0.0087143 | 0.0023914 | 0.0135765 | 0.0048466 | 0.019622 | 0.0057625 | 0.005091 | 0.0165721 | 0.001606 | 0.0063498 | 0.0067851 | 0.0118349 | 0.0025851 | 0.0077694 | 0.0045054 | 0.0036941 | 0.0113654 | 0.0048791 |
| 26.32 | 0.0067471 | 0.0017762 | 0.0016173 | 0.0076251 | 0.0019359 | 0.0098946 | 0.0025044 | 0.001965 | 0.0060521 | 0.0027637 | 0.0029775 | 0.0056072 | 0.0045203 | 0.0007814 | 0.0056152 | 0.0054281 | 0.0054813 | 0.0023106 | 0.0055336 |
| 26.33 | 0.0106331 | 0.0021939 | 0.0058457 | 0.0093992 | 0.0057102 | 0.0120507 | 0.0057682 | 0.0034241 | 0.0146574 | 0.0124685 | 0.0091729 | 0.0061809 | 0.0055712 | 0.001562 | 0.0106383 | 0.0139453 | 0.006997 | 0.0029602 | 0.0108852 |
| 26.35 | 0.0093724 | 0.0042297 | 0.0067456 | 0.0070599 | 0.0063998 | 0.0115463 | 0.0068129 | 0.0050207 | 0.0207017 | 0.0165295 | 0.011671 | 0.005999 | 0.0072605 | 0.0028314 | 0.0086346 | 0.0146147 | 0.0066093 | 0.0034245 | 0.0094143 |
| 26.36 | 0.0065451 | 0.0078992 | 0.0043871 | 0.0035367 | 0.0058419 | 0.0088924 | 0.0050332 | 0.0078615 | 0.0206895 | 0.0161075 | 0.0130768 | 0.006529 | 0.0099943 | 0.005353 | 0.0065156 | 0.0132396 | 0.0124465 | 0.0043375 | 0.008269 |
| 26.38 | 0.0092878 | 0.0068511 | 0.0033613 | 0.0052162 | 0.0086886 | 0.0065223 | 0.0032687 | 0.0072187 | 0.0156582 | 0.0122578 | 0.0124604 | 0.0051844 | 0.0086038 | 0.0054405 | 0.0071259 | 0.0116063 | 0.0128245 | 0.0047673 | 0.008917 |
| 26.39 | 0.0094386 | 0.0015539 | 0.0020565 | 0.0060688 | 0.011265 | 0.0039084 | 0.0011138 | 0.002225 | 0.008967 | 0.0041764 | 0.0069664 | 0.0032474 | 0.0023522 | 0.0022696 | 0.0030359 | 0.0041448 | 0.0046225 | 0.0021378 | 0.00449 |
| 26.4 | 0.0096846 | 0.0005423 | 0.0054001 | 0.0060386 | 0.0172494 | 0.011064 | 0.0024528 | 0.000625 | 0.0115163 | 0.0108553 | 0.0126306 | 0.0087605 | 0.0022297 | 0.0015238 | 0.0029753 | 0.0044163 | 0.0039854 | 0.0011852 | 0.0038727 |
| 26.42 | 0.009654 | 0.0009419 | 0.009211 | 0.0049512 | 0.0129408 | 0.0134649 | 0.0046059 | 0.0004379 | 0.0118394 | 0.0168459 | 0.0142089 | 0.0083644 | 0.003766 | 0.0013148 | 0.0037015 | 0.0059953 | 0.0042454 | 0.0011854 | 0.0031 |
| 26.43 | 0.0132389 | 0.0027841 | 0.0106257 | 0.0064193 | 0.0035396 | 0.0087223 | 0.0067799 | 0.0013557 | 0.017956 | 0.0157457 | 0.0081807 | 0.0037863 | 0.0053743 | 0.0017734 | 0.0022552 | 0.0061287 | 0.0063959 | 0.0012678 | 0.0009053 |
| 26.44 | 0.0278047 | 0.0112953 | 0.0163417 | 0.0137437 | 0.0063623 | 0.0068329 | 0.0151401 | 0.0048868 | 0.038413 | 0.0206206 | 0.0080038 | 0.0123492 | 0.0100792 | 0.0081913 | 0.0017827 | 0.0106861 | 0.0218498 | 0.0023973 | 0.0013518 |
| 26.46 | 0.0254205 | 0.0134911 | 0.0147999 | 0.0127134 | 0.0078646 | 0.0036336 | 0.0172648 | 0.0054451 | 0.0332781 | 0.0179784 | 0.0074499 | 0.0166365 | 0.0093339 | 0.0114949 | 0.0012656 | 0.0112509 | 0.0267011 | 0.0033568 | 0.002556 |
| 26.47 | 0.0109819 | 0.0102272 | 0.0114564 | 0.0062449 | 0.0068807 | 0.0020109 | 0.0164729 | 0.0031884 | 0.0161908 | 0.0168395 | 0.0106096 | 0.0152097 | 0.0069524 | 0.0126771 | 0.0059458 | 0.0146306 | 0.0246895 | 0.0063895 | 0.0070136 |
| 26.49 | 0.0056712 | 0.009007 | 0.0105138 | 0.003392 | 0.0052292 | 0.0022631 | 0.0129116 | 0.0022274 | 0.0145043 | 0.0177639 | 0.0110357 | 0.0106234 | 0.0061236 | 0.010648 | 0.0093233 | 0.0163197 | 0.0193948 | 0.0066913 | 0.0080326 |
| 26.5 | 0.0020239 | 0.0048341 | 0.0066548 | 0.0012051 | 0.001967 | 0.0035162 | 0.003477 | 0.0010692 | 0.0099551 | 0.0122365 | 0.0047085 | 0.0022361 | 0.0030139 | 0.0042324 | 0.0055704 | 0.0091871 | 0.0066373 | 0.0038685 | 0.0036854 |
| 26.51 | 0.0078658 | 0.0135377 | 0.009525 | 0.0052114 | 0.0028891 | 0.0169787 | 0.0026508 | 0.0031459 | 0.022575 | 0.0176375 | 0.0060173 | 0.0042083 | 0.0073769 | 0.0033657 | 0.0146403 | 0.0146471 | 0.0096279 | 0.0138162 | 0.0116922 |
| 26.53 | 0.0171136 | 0.0347811 | 0.0097653 | 0.0099645 | 0.0054863 | 0.0317747 | 0.0071838 | 0.0088473 | 0.0400729 | 0.0189907 | 0.0123114 | 0.0083868 | 0.0151951 | 0.0023095 | 0.0366673 | 0.0264329 | 0.0142888 | 0.032406 | 0.029388 |
| 26.54 | 0.0470652 | 0.0944865 | 0.0153116 | 0.0229305 | 0.0190715 | 0.0785886 | 0.03198 | 0.0324531 | 0.0830205 | 0.0348766 | 0.0446759 | 0.0111834 | 0.044926 | 0.0101732 | 0.1016771 | 0.0656316 | 0.0263772 | 0.0965228 | 0.0846559 |
| 26.56 | 0.0634563 | 0.1004186 | 0.0238821 | 0.0331682 | 0.0235365 | 0.0931043 | 0.0472767 | 0.0423197 | 0.0901039 | 0.0480523 | 0.0583513 | 0.0143631 | 0.0594669 | 0.0208353 | 0.1126821 | 0.077862 | 0.0319936 | 0.1110518 | 0.0975622 |
| 26.57 | 0.0563668 | 0.0441992 | 0.0328149 | 0.04074 | 0.0110194 | 0.0546832 | 0.0457519 | 0.0273594 | 0.0498931 | 0.0400509 | 0.0351423 | 0.0233135 | 0.046451 | 0.0265994 | 0.057785 | 0.0501383 | 0.0276084 | 0.0536808 | 0.0533907 |
| 26.58 | 0.0842733 | 0.0357547 | 0.0711446 | 0.0832656 | 0.0096489 | 0.0500163 | 0.0772244 | 0.0244914 | 0.0490828 | 0.0465575 | 0.0254252 | 0.0446967 | 0.0619434 | 0.0490713 | 0.0552898 | 0.0543 | 0.0448099 | 0.0349767 | 0.044809 |
| 26.6 | 0.075083 | 0.0285872 | 0.0705023 | 0.082119 | 0.015636 | 0.033336 | 0.0708486 | 0.0177981 | 0.035795 | 0.0323518 | 0.0139131 | 0.0407924 | 0.0532582 | 0.0476556 | 0.0479885 | 0.0438076 | 0.0418558 | 0.0210159 | 0.0290489 |
| 26.61 | 0.0415482 | 0.0156194 | 0.03601 | 0.0470467 | 0.0356005 | 0.0104003 | 0.0318998 | 0.0099087 | 0.0152478 | 0.0143364 | 0.0078749 | 0.0252818 | 0.0261517 | 0.0292985 | 0.044589 | 0.0279332 | 0.0238862 | 0.0100078 | 0.0096172 |
| 26.63 | 0.0247797 | 0.0081947 | 0.0214782 | 0.0288533 | 0.0404618 | 0.0082652 | 0.0203267 | 0.0064492 | 0.0143729 | 0.0205776 | 0.0117053 | 0.0164282 | 0.0159986 | 0.0189198 | 0.0396423 | 0.0193223 | 0.0157378 | 0.0066056 | 0.0055529 |
| 26.64 | 0.0065567 | 0.0007404 | 0.0184303 | 0.0146064 | 0.0272853 | 0.0123835 | 0.0248918 | 0.0024254 | 0.0199545 | 0.0260598 | 0.0117329 | 0.0043966 | 0.011865 | 0.0090239 | 0.0152476 | 0.0086662 | 0.0107073 | 0.0023455 | 0.0060983 |
| 26.65 | 0.014783 | 0.0004909 | 0.0433549 | 0.0284943 | 0.0357937 | 0.0314351 | 0.0600195 | 0.0028325 | 0.0401467 | 0.0370903 | 0.0105559 | 0.0073395 | 0.0275329 | 0.0233386 | 0.0121501 | 0.0278987 | 0.0202089 | 0.0043345 | 0.0176648 |
| 26.67 | 0.0201627 | 0.0011021 | 0.0438166 | 0.0256231 | 0.0294549 | 0.03921 | 0.0571509 | 0.0023342 | 0.0328618 | 0.0271617 | 0.0073961 | 0.0075187 | 0.0271568 | 0.0259593 | 0.0167575 | 0.0358866 | 0.0167176 | 0.0044874 | 0.0212721 |
| 26.68 | 0.0153 | 0.0012096 | 0.0246988 | 0.008319 | 0.010211 | 0.0357293 | 0.0219196 | 0.0015518 | 0.0104106 | 0.0048626 | 0.0062243 | 0.0024341 | 0.010085 | 0.0125438 | 0.0224251 | 0.0244871 | 0.0074862 | 0.0028829 | 0.0174652 |
| 26.69 | 0.0136814 | 0.0013403 | 0.0214684 | 0.0078911 | 0.0058477 | 0.0568201 | 0.025045 | 0.0026444 | 0.0248407 | 0.0070927 | 0.0085605 | 0.0046838 | 0.0033102 | 0.0084111 | 0.0414937 | 0.0292067 | 0.0140553 | 0.0102302 | 0.0290188 |
| 26.71 | 0.0131956 | 0.00146 | 0.0165292 | 0.0128249 | 0.0064802 | 0.0507952 | 0.0348924 | 0.0034502 | 0.0284469 | 0.0162822 | 0.0099933 | 0.0066774 | 0.0041514 | 0.0069898 | 0.0411656 | 0.0252697 | 0.0135265 | 0.0146702 | 0.028054 |
| 26.72 | 0.0372936 | 0.0027162 | 0.0325338 | 0.0308018 | 0.0210241 | 0.0236242 | 0.0548412 | 0.0060576 | 0.0135341 | 0.0383027 | 0.0229471 | 0.0076225 | 0.0203165 | 0.014432 | 0.0328893 | 0.0178409 | 0.0175446 | 0.0149927 | 0.0205551 |
| 26.74 | 0.0471674 | 0.0027289 | 0.0341435 | 0.0333163 | 0.0266163 | 0.0164788 | 0.0464706 | 0.0067579 | 0.0078451 | 0.0428611 | 0.0263449 | 0.0082907 | 0.0263021 | 0.0185285 | 0.0269698 | 0.0126326 | 0.0253281 | 0.0109446 | 0.0149594 |
| 26.75 | 0.0251759 | 0.0035763 | 0.0097209 | 0.0130675 | 0.0166329 | 0.0186723 | 0.0139541 | 0.0039759 | 0.0079213 | 0.0203605 | 0.0121504 | 0.0068391 | 0.01423 | 0.0128873 | 0.0110003 | 0.0037901 | 0.0222788 | 0.0038333 | 0.0047869 |
| 26.76 | 0.0148406 | 0.0159325 | 0.0060143 | 0.0065282 | 0.0141552 | 0.0382799 | 0.0328491 | 0.0026833 | 0.021697 | 0.021138 | 0.0088756 | 0.0104183 | 0.0097376 | 0.0108482 | 0.0070439 | 0.0099686 | 0.021536 | 0.0084076 | 0.0133489 |
| 26.78 | 0.0093375 | 0.0191167 | 0.0092204 | 0.0047443 | 0.0081263 | 0.0377898 | 0.0462989 | 0.0011681 | 0.0251446 | 0.0321601 | 0.0099345 | 0.0089837 | 0.0088937 | 0.006486 | 0.0041775 | 0.0137243 | 0.0142033 | 0.0120192 | 0.0227449 |
| 26.79 | 0.0103488 | 0.0105491 | 0.0170411 | 0.0025553 | 0.005373 | 0.0296433 | 0.0531606 | 0.000553 | 0.0233749 | 0.0527559 | 0.0111588 | 0.0089153 | 0.0108551 | 0.0065838 | 0.0045026 | 0.0126883 | 0.0174778 | 0.0179206 | 0.03877 |
| 26.81 | 0.0138029 | 0.0053849 | 0.016637 | 0.0043134 | 0.0103348 | 0.0222699 | 0.0436051 | 0.0012031 | 0.0193907 | 0.0452992 | 0.0094391 | 0.009594 | 0.0087035 | 0.0077551 | 0.0041812 | 0.0102019 | 0.0197425 | 0.0157556 | 0.0386702 |
| 26.82 | 0.0218125 | 0.0021444 | 0.0087348 | 0.0128881 | 0.0146143 | 0.0121985 | 0.0117325 | 0.0024551 | 0.006721 | 0.0159251 | 0.0097315 | 0.0049416 | 0.00389 | 0.0039547 | 0.0027949 | 0.0096112 | 0.0090847 | 0.0042894 | 0.0172817 |

| | | | | | | | | | | | | | | | | | | | |
|---|---|---|---|---|---|---|---|---|---|---|---|---|---|---|---|---|---|---|---|
| 26.83 | 0.0503837 | 0.0030104 | 0.0162939 | 0.0297712 | 0.0207718 | 0.0388876 | 0.0027665 | 0.0058012 | 0.0052006 | 0.0249209 | 0.0204775 | 0.0115625 | 0.0085554 | 0.004693 | 0.0126811 | 0.0289999 | 0.0147904 | 0.0028015 | 0.0125626 |
| 26.85 | 0.0448847 | 0.0038656 | 0.0183523 | 0.0246436 | 0.0152565 | 0.0512633 | 0.0006744 | 0.0058452 | 0.0073447 | 0.0278217 | 0.0177202 | 0.0185272 | 0.0125353 | 0.0066545 | 0.0194735 | 0.0354588 | 0.0220591 | 0.0026468 | 0.0128917 |
| 26.86 | 0.0199165 | 0.0112806 | 0.0117045 | 0.0125334 | 0.0066843 | 0.0659323 | 0.0014927 | 0.0034637 | 0.0108141 | 0.0185857 | 0.0091813 | 0.0309039 | 0.0286007 | 0.0132883 | 0.0298188 | 0.0374053 | 0.0317805 | 0.0057809 | 0.0291348 |
| 26.88 | 0.0157063 | 0.0146097 | 0.0066776 | 0.0170613 | 0.0047392 | 0.0703867 | 0.0026645 | 0.002076 | 0.0100794 | 0.0148143 | 0.0101375 | 0.0311279 | 0.0355364 | 0.0163616 | 0.0307125 | 0.0339364 | 0.0278957 | 0.0083884 | 0.0367721 |
| 26.89 | 0.0112381 | 0.0088317 | 0.0033412 | 0.020946 | 0.0046311 | 0.0450949 | 0.0061291 | 0.0006382 | 0.0097535 | 0.0160015 | 0.0135684 | 0.0160426 | 0.0230344 | 0.0134814 | 0.0158493 | 0.0178101 | 0.0099686 | 0.0055635 | 0.0275297 |
| 26.9 | 0.0154195 | 0.0062843 | 0.0055358 | 0.0343624 | 0.0203896 | 0.0447616 | 0.0223222 | 0.0014362 | 0.0268681 | 0.0430598 | 0.0293643 | 0.0175593 | 0.024237 | 0.0213411 | 0.0119653 | 0.0238348 | 0.0089791 | 0.0038424 | 0.034229 |
| 26.92 | 0.0135549 | 0.0046743 | 0.006837 | 0.0315757 | 0.0249484 | 0.0314443 | 0.0297717 | 0.0026848 | 0.0311226 | 0.0498961 | 0.0288166 | 0.0157262 | 0.019578 | 0.0219026 | 0.0067295 | 0.023681 | 0.0110512 | 0.0029687 | 0.0278779 |
| 26.93 | 0.0084292 | 0.0033083 | 0.0091625 | 0.0173551 | 0.0125595 | 0.0122703 | 0.0245321 | 0.0038631 | 0.01984 | 0.0313506 | 0.0128086 | 0.0090917 | 0.0099691 | 0.0153723 | 0.0014682 | 0.0151977 | 0.0134063 | 0.0042401 | 0.0127276 |
| 26.94 | 0.0182393 | 0.0039648 | 0.0167119 | 0.0274146 | 0.0083207 | 0.0101302 | 0.0315251 | 0.0082054 | 0.0237375 | 0.028897 | 0.011378 | 0.0094742 | 0.0139532 | 0.0208022 | 0.0020836 | 0.0174849 | 0.0257529 | 0.0150281 | 0.0160345 |
| 26.96 | 0.0218587 | 0.0044532 | 0.0139829 | 0.0314648 | 0.0068139 | 0.0063326 | 0.0239901 | 0.0097841 | 0.0209252 | 0.0196112 | 0.0114755 | 0.0066707 | 0.0142147 | 0.0176412 | 0.0029364 | 0.0131832 | 0.0254827 | 0.0191801 | 0.0158541 |
| 26.97 | 0.0195128 | 0.0055052 | 0.0067038 | 0.0353361 | 0.0145175 | 0.0049891 | 0.0096181 | 0.0137676 | 0.0148145 | 0.0138434 | 0.0178196 | 0.0022245 | 0.0110208 | 0.0101693 | 0.0047061 | 0.0063383 | 0.0173378 | 0.0196928 | 0.0154792 |
| 26.99 | 0.0138889 | 0.0045997 | 0.006709 | 0.0300526 | 0.0187144 | 0.0061186 | 0.0074877 | 0.014966 | 0.0107025 | 0.0161781 | 0.0193906 | 0.0012853 | 0.0076069 | 0.0074405 | 0.004874 | 0.0047078 | 0.0113178 | 0.0162252 | 0.0143328 |
| 27 | 0.0054033 | 0.002473 | 0.0056671 | 0.0095106 | 0.0141721 | 0.0040463 | 0.0064887 | 0.0101773 | 0.0055685 | 0.01579 | 0.0109947 | 0.000931 | 0.0024405 | 0.0034373 | 0.0021652 | 0.0020513 | 0.0031298 | 0.0059948 | 0.0088121 |
| 27.01 | 0.0083509 | 0.0124411 | 0.0075057 | 0.0090674 | 0.0244868 | 0.008873 | 0.0073102 | 0.0192076 | 0.0142343 | 0.0312617 | 0.0161588 | 0.0024049 | 0.0088978 | 0.004702 | 0.0067773 | 0.0019944 | 0.0049033 | 0.0196008 | 0.0239162 |
| 27.03 | 0.0146859 | 0.0313976 | 0.010846 | 0.0105635 | 0.0315507 | 0.0187754 | 0.0084341 | 0.0261044 | 0.0152199 | 0.0365145 | 0.0237235 | 0.0031148 | 0.0236206 | 0.0055779 | 0.0203433 | 0.0065273 | 0.0079253 | 0.0418488 | 0.0423659 |
| 27.04 | 0.0507156 | 0.0857335 | 0.0280916 | 0.0153912 | 0.0546597 | 0.0489013 | 0.0339342 | 0.0420135 | 0.022756 | 0.0473161 | 0.0441607 | 0.0111519 | 0.0774378 | 0.012605 | 0.072668 | 0.0401966 | 0.0278066 | 0.0912545 | 0.0905727 |
| 27.06 | 0.0647761 | 0.0934109 | 0.0360291 | 0.0246398 | 0.0571024 | 0.0539316 | 0.047863 | 0.0419414 | 0.0370517 | 0.0504295 | 0.0454767 | 0.0177007 | 0.0958142 | 0.0157446 | 0.0860234 | 0.0559044 | 0.0372164 | 0.0915763 | 0.095541 |
| 27.07 | 0.0430561 | 0.0453482 | 0.0265371 | 0.0304545 | 0.0221985 | 0.0261656 | 0.0333068 | 0.0186246 | 0.0416253 | 0.0272591 | 0.0214003 | 0.0121654 | 0.0566269 | 0.0079038 | 0.0381756 | 0.0296643 | 0.0220107 | 0.0337798 | 0.0402631 |
| 27.08 | 0.0490104 | 0.0371956 | 0.0315527 | 0.0442348 | 0.0082414 | 0.0184476 | 0.0318776 | 0.0159164 | 0.0676692 | 0.0218487 | 0.0177657 | 0.0101479 | 0.0371672 | 0.0046787 | 0.0201214 | 0.0187588 | 0.0130617 | 0.0149011 | 0.0188074 |
| 27.1 | 0.0379904 | 0.023678 | 0.0248534 | 0.0347827 | 0.0052774 | 0.0104462 | 0.0216754 | 0.0134362 | 0.0639342 | 0.0226936 | 0.0176928 | 0.0091433 | 0.0183623 | 0.0039416 | 0.0183195 | 0.0202771 | 0.0070448 | 0.011235 | 0.0092373 |
| 27.11 | 0.0172028 | 0.0065173 | 0.0104972 | 0.0126973 | 0.008912 | 0.0046215 | 0.0077256 | 0.0110847 | 0.0450977 | 0.0288401 | 0.0233104 | 0.0076866 | 0.0163518 | 0.0027998 | 0.0228785 | 0.0269448 | 0.009737 | 0.0116849 | 0.0080935 |
| 27.13 | 0.0102846 | 0.0040403 | 0.0085166 | 0.0066241 | 0.0084153 | 0.0054945 | 0.0059902 | 0.0103651 | 0.0317712 | 0.0235277 | 0.0193786 | 0.0055639 | 0.0179731 | 0.0014439 | 0.0163316 | 0.0209865 | 0.0110418 | 0.008844 | 0.0097424 |
| 27.14 | 0.0024536 | 0.0024047 | 0.0167085 | 0.0046941 | 0.0029166 | 0.0043875 | 0.0103208 | 0.0092645 | 0.0086987 | 0.0086328 | 0.0048911 | 0.0017955 | 0.0105847 | 0.0004451 | 0.0025212 | 0.0060802 | 0.0067904 | 0.0069434 | 0.0109826 |
| 27.15 | 0.0032841 | 0.0026249 | 0.0396779 | 0.0128461 | 0.0047287 | 0.0061619 | 0.0288077 | 0.015969 | 0.0183053 | 0.0084015 | 0.0028339 | 0.0011974 | 0.0201687 | 0.0009572 | 0.0068423 | 0.0070356 | 0.0159534 | 0.0109272 | 0.0149578 |
| 27.17 | 0.0065431 | 0.0029929 | 0.0320497 | 0.0180707 | 0.0107637 | 0.0081495 | 0.0303715 | 0.0135293 | 0.0310872 | 0.0067707 | 0.0053623 | 0.0014489 | 0.025827 | 0.0019704 | 0.0155169 | 0.0104456 | 0.0239931 | 0.0092438 | 0.0114762 |
| 27.18 | 0.0096993 | 0.0041067 | 0.0077486 | 0.0175792 | 0.0168377 | 0.0089836 | 0.0170586 | 0.0068903 | 0.0345788 | 0.0044265 | 0.0099238 | 0.0037881 | 0.0225685 | 0.0045569 | 0.019321 | 0.0114473 | 0.0256968 | 0.0064207 | 0.007469 |
| 27.19 | 0.0167948 | 0.0073831 | 0.0103092 | 0.0226777 | 0.028053 | 0.0111063 | 0.021224 | 0.0082899 | 0.0533809 | 0.0137957 | 0.0159221 | 0.0133731 | 0.0276399 | 0.0133989 | 0.0204612 | 0.0143284 | 0.0410194 | 0.0092434 | 0.0097066 |
| 27.21 | 0.0136748 | 0.0075901 | 0.0123393 | 0.016376 | 0.0228229 | 0.0070721 | 0.0184355 | 0.0068125 | 0.0513108 | 0.0149819 | 0.0126287 | 0.014673 | 0.0189192 | 0.0142355 | 0.0114637 | 0.0096604 | 0.034959 | 0.0094524 | 0.0070079 |
| 27.22 | 0.012769 | 0.0228911 | 0.0237821 | 0.0044075 | 0.0125342 | 0.0015654 | 0.0158396 | 0.0187357 | 0.0695722 | 0.0112149 | 0.0122001 | 0.0071899 | 0.0085319 | 0.0070123 | 0.0036473 | 0.0025691 | 0.0138285 | 0.0102495 | 0.0045024 |
| 27.24 | 0.0194457 | 0.0361862 | 0.0355369 | 0.0027206 | 0.0152996 | 0.001809 | 0.019109 | 0.0295319 | 0.0874014 | 0.0100896 | 0.0119949 | 0.0033307 | 0.0114146 | 0.002918 | 0.0031813 | 0.0019698 | 0.0066097 | 0.0076819 | 0.0042392 |
| 27.25 | 0.0147857 | 0.0258072 | 0.0331879 | 0.0077292 | 0.0128765 | 0.0042049 | 0.0187995 | 0.0223934 | 0.061085 | 0.007682 | 0.0077365 | 0.0027353 | 0.0108911 | 0.0015975 | 0.0019074 | 0.0022072 | 0.0032287 | 0.00328 | 0.0043703 |
| 27.26 | 0.014682 | 0.0182955 | 0.0430396 | 0.0295305 | 0.007178 | 0.0093822 | 0.0291293 | 0.0183968 | 0.0526833 | 0.023911 | 0.0258261 | 0.012495 | 0.0088051 | 0.0092684 | 0.004698 | 0.0071107 | 0.0122029 | 0.0080725 | 0.0110883 |
| 27.28 | 0.0167098 | 0.0148372 | 0.033863 | 0.0312104 | 0.0049369 | 0.0082916 | 0.0229389 | 0.0104497 | 0.0312005 | 0.0257014 | 0.028951 | 0.0156815 | 0.0042946 | 0.0120636 | 0.0053986 | 0.0102499 | 0.0157342 | 0.008511 | 0.0105448 |
| 27.29 | 0.0169585 | 0.0149104 | 0.012474 | 0.015456 | 0.0161451 | 0.0036818 | 0.0100409 | 0.0025474 | 0.0097833 | 0.0129098 | 0.0136817 | 0.0131902 | 0.004358 | 0.0096324 | 0.0031299 | 0.0107005 | 0.0113521 | 0.0094133 | 0.0063551 |
| 27.31 | 0.0111434 | 0.0101037 | 0.0059414 | 0.011456 | 0.020858 | 0.0023114 | 0.0101679 | 0.001528 | 0.0084586 | 0.0110976 | 0.0078831 | 0.0098173 | 0.0077829 | 0.006727 | 0.0028472 | 0.0082682 | 0.0092365 | 0.0112912 | 0.0063206 |
| 27.32 | 0.0031996 | 0.0021099 | 0.0029101 | 0.0067495 | 0.0143938 | 0.0023182 | 0.0072695 | 0.002024 | 0.005816 | 0.0095274 | 0.0046147 | 0.0033411 | 0.0098942 | 0.0036275 | 0.0044641 | 0.0057529 | 0.0107983 | 0.007526 | 0.0039095 |
| 27.33 | 0.0100378 | 0.0060599 | 0.0115356 | 0.0108223 | 0.0141435 | 0.011081 | 0.0121157 | 0.0077452 | 0.0092763 | 0.0119928 | 0.0084425 | 0.0032028 | 0.0185921 | 0.0062004 | 0.009949 | 0.0131958 | 0.0202717 | 0.0062632 | 0.0026536 |
| 27.35 | 0.0143932 | 0.0086779 | 0.0170061 | 0.0177298 | 0.0088242 | 0.0136327 | 0.0173143 | 0.0098064 | 0.0149362 | 0.0144369 | 0.0128939 | 0.0035597 | 0.0175977 | 0.0052144 | 0.009064 | 0.0115303 | 0.0159137 | 0.0050932 | 0.0016019 |
| 27.36 | 0.0165252 | 0.0077311 | 0.019898 | 0.0313124 | 0.004313 | 0.0075972 | 0.0234872 | 0.007736 | 0.0275585 | 0.027582 | 0.0251052 | 0.0059244 | 0.0101348 | 0.0048488 | 0.0042183 | 0.0066491 | 0.0075098 | 0.0075739 | 0.0022968 |
| 27.38 | 0.0139479 | 0.0052187 | 0.0163437 | 0.0296315 | 0.0062774 | 0.0033386 | 0.0205618 | 0.0047637 | 0.0279491 | 0.0278446 | 0.0258487 | 0.0064667 | 0.0060318 | 0.005982 | 0.003659 | 0.010425 | 0.0081648 | 0.0073572 | 0.0031729 |
| 27.39 | 0.0045338 | 0.0014619 | 0.005705 | 0.0109512 | 0.00845 | 0.0002485 | 0.0068574 | 0.0007344 | 0.0131885 | 0.0099456 | 0.0119595 | 0.0036305 | 0.0015143 | 0.004288 | 0.0032171 | 0.010851 | 0.0053839 | 0.0030813 | 0.002212 |
| 27.4 | 0.0015004 | 0.0032314 | 0.0053082 | 0.0058095 | 0.0192157 | 0.0007319 | 0.0033869 | 0.0004233 | 0.0100832 | 0.0036921 | 0.0080142 | 0.0040073 | 0.0037368 | 0.0045343 | 0.0033422 | 0.0091594 | 0.0031245 | 0.0051434 | 0.0032309 |
| 27.42 | 0.0014102 | 0.0038541 | 0.0058856 | 0.0046718 | 0.0223595 | 0.0011326 | 0.0031294 | 0.0009889 | 0.0068845 | 0.0032156 | 0.0061198 | 0.0040333 | 0.0058296 | 0.0032301 | 0.0021111 | 0.0062804 | 0.002918 | 0.0051463 | 0.0036613 |
| 27.43 | 0.0050258 | 0.0028588 | 0.0065506 | 0.0033467 | 0.0171737 | 0.0011086 | 0.002228 | 0.0025009 | 0.0045145 | 0.0080172 | 0.0080812 | 0.0035012 | 0.0080366 | 0.0009291 | 0.0021025 | 0.0035454 | 0.0059743 | 0.002994 | 0.0028685 |
| 27.44 | 0.018194 | 0.009178 | 0.0143446 | 0.0060334 | 0.0258428 | 0.0030261 | 0.0027171 | 0.0071972 | 0.014319 | 0.0247273 | 0.0213007 | 0.0060996 | 0.0207463 | 0.0015627 | 0.0125899 | 0.0065201 | 0.0169433 | 0.007882 | 0.0090333 |
| 27.46 | 0.02163 | 0.0115621 | 0.0150932 | 0.0090198 | 0.0238831 | 0.0043851 | 0.0043336 | 0.0078866 | 0.0200307 | 0.0263389 | 0.0227174 | 0.0060638 | 0.0246498 | 0.0031845 | 0.0192172 | 0.0120546 | 0.0196636 | 0.0085956 | 0.0130622 |
| 27.47 | 0.0188476 | 0.009601 | 0.0126804 | 0.013717 | 0.0165498 | 0.0060598 | 0.0079039 | 0.0063327 | 0.0279552 | 0.0148293 | 0.0152231 | 0.0053913 | 0.0268271 | 0.0087731 | 0.0239143 | 0.0225748 | 0.021143 | 0.0043766 | 0.0152683 |
| 27.49 | 0.0206436 | 0.0115108 | 0.0131059 | 0.0137539 | 0.0137976 | 0.0092788 | 0.0081043 | 0.007322 | 0.0282405 | 0.0091872 | 0.0101454 | 0.005555 | 0.0240925 | 0.0106793 | 0.0188464 | 0.0192172 | 0.0206938 | 0.0038811 | 0.0126202 |
| 27.5 | 0.0320526 | 0.0234181 | 0.0152311 | 0.0096815 | 0.0123837 | 0.0191987 | 0.0053466 | 0.0130181 | 0.0181378 | 0.004552 | 0.0037632 | 0.0048141 | 0.0099363 | 0.0065649 | 0.0048738 | 0.0051454 | 0.0120192 | 0.0050854 | 0.0056466 |
| 27.51 | 0.0935202 | 0.073338 | 0.0427681 | 0.0209768 | 0.0346629 | 0.0657937 | 0.0160055 | 0.0473502 | 0.039549 | 0.0063358 | 0.007837 | 0.0071983 | 0.013538 | 0.0053829 | 0.010793 | 0.0074566 | 0.0158426 | 0.0122632 | 0.0177985 |
| 27.53 | 0.1044773 | 0.0774438 | 0.0531112 | 0.0327307 | 0.040651 | 0.0851384 | 0.0310385 | 0.0609708 | 0.0506226 | 0.0077033 | 0.0137359 | 0.0053092 | 0.0237988 | 0.0041165 | 0.0225475 | 0.0141126 | 0.0167632 | 0.0229085 | 0.0376604 |
| 27.54 | 0.0844908 | 0.0392448 | 0.0663027 | 0.0776843 | 0.0351782 | 0.0985645 | 0.081542 | 0.0558039 | 0.0622067 | 0.0245761 | 0.0351257 | 0.0057733 | 0.0626571 | 0.009704 | 0.0621939 | 0.0399661 | 0.0215827 | 0.0698501 | 0.0921016 |
| 27.56 | 0.0703895 | 0.022072 | 0.0699089 | 0.0949716 | 0.0266506 | 0.0918176 | 0.1007656 | 0.0411683 | 0.0599838 | 0.0375779 | 0.0432069 | 0.0152536 | 0.0764263 | 0.0155535 | 0.0731345 | 0.0487939 | 0.0251886 | 0.0885682 | 0.1007052 |
| 27.57 | 0.0351451 | 0.0082379 | 0.0443331 | 0.0586269 | 0.0087949 | 0.0459208 | 0.0657791 | 0.0135976 | 0.0298489 | 0.0382884 | 0.0308835 | 0.0232229 | 0.0526462 | 0.0162891 | 0.0403902 | 0.0296824 | 0.0224217 | 0.0556309 | 0.0518235 |
| 27.58 | 0.0344338 | 0.010521 | 0.0457497 | 0.0425479 | 0.0063678 | 0.0430771 | 0.0572665 | 0.0089517 | 0.01946 | 0.0600698 | 0.0405189 | 0.026225 | 0.0746835 | 0.0264612 | 0.0342679 | 0.028805 | 0.0377409 | 0.0493829 | 0.0454453 |
| 27.6 | 0.022831 | 0.0093797 | 0.0297434 | 0.024049 | 0.0061764 | 0.0300126 | 0.0335421 | 0.0045241 | 0.0113431 | 0.0498684 | 0.0314177 | 0.0158481 | 0.0719559 | 0.0248251 | 0.023057 | 0.0215365 | 0.0357948 | 0.0352808 | 0.033561 |
| 27.61 | 0.0072362 | 0.0086688 | 0.0107177 | 0.0325305 | 0.012016 | 0.0105164 | 0.0158276 | 0.0018283 | 0.0212005 | 0.0212949 | 0.0176287 | 0.006652 | 0.0489401 | 0.0171017 | 0.008256 | 0.0133984 | 0.0241031 | 0.0201312 | 0.0214591 |
| 27.63 | 0.0061359 | 0.0092941 | 0.0135944 | 0.0443216 | 0.0140053 | 0.005852 | 0.0203564 | 0.0018641 | 0.0291678 | 0.0202897 | 0.0277208 | 0.0078081 | 0.0305195 | 0.0117646 | 0.0054607 | 0.0115511 | 0.0169965 | 0.0139204 | 0.0175271 |
| 27.64 | 0.0071631 | 0.0069014 | 0.014163 | 0.0338701 | 0.0098565 | 0.0059806 | 0.0218951 | 0.0020097 | 0.0210209 | 0.0262876 | 0.0335451 | 0.0080233 | 0.0056546 | 0.0024681 | 0.0068807 | 0.0068823 | 0.0050077 | 0.0038078 | 0.0092188 |
| 27.65 | 0.0143392 | 0.0121808 | 0.0153923 | 0.037813 | 0.0151295 | 0.0237239 | 0.040611 | 0.0052015 | 0.0170881 | 0.0346638 | 0.0321288 | 0.0116335 | 0.0084887 | 0.0014877 | 0.0153971 | 0.007907 | 0.0081647 | 0.0021525 | 0.0139958 |
| 27.67 | 0.013044 | 0.0117651 | 0.0102829 | 0.0253838 | 0.0138058 | 0.0363866 | 0.0423838 | 0.0055103 | 0.011407 | 0.0267955 | 0.0232716 | 0.0107011 | 0.010424 | 0.0027164 | 0.0143325 | 0.0064361 | 0.0113224 | 0.0016954 | 0.0150737 |
| 27.68 | 0.0043589 | 0.0062167 | 0.0026784 | 0.0064658 | 0.0062384 | 0.0404992 | 0.0238673 | 0.0023367 | 0.0113358 | 0.0276647 | 0.0257296 | 0.0069438 | 0.0073597 | 0.005929 | 0.0050396 | 0.0047903 | 0.0071079 | 0.0013864 | 0.0107057 |
| 27.69 | 0.0037154 | 0.0132309 | 0.0043078 | 0.0128639 | 0.0134296 | 0.0706609 | 0.0189668 | 0.0024065 | 0.0184661 | 0.0615213 | 0.0427078 | 0.0095444 | 0.0118684 | 0.0163958 | 0.0022025 | 0.0096381 | 0.0078842 | 0.0025596 | 0.0133221 |
| 27.71 | 0.0072723 | 0.0147918 | 0.0071859 | 0.0152153 | 0.0184785 | 0.0642717 | 0.0169675 | 0.0038481 | 0.015185 | 0.052798 | 0.0311281 | 0.0077742 | 0.012994 | 0.0190924 | 0.0014071 | 0.0083461 | 0.0114675 | 0.0022581 | 0.0111782 |
| 27.72 | 0.0242785 | 0.0083352 | 0.013681 | 0.0158861 | 0.0253763 | 0.03356 | 0.0395889 | 0.0081554 | 0.0182901 | 0.0219427 | 0.0214593 | 0.0030945 | 0.0152344 | 0.0197945 | 0.0013808 | 0.0042568 | 0.016936 | 0.0018966 | 0.0085385 |
| 27.74 | 0.0299437 | 0.004397 | 0.0136536 | 0.0147821 | 0.0271724 | 0.0203809 | 0.0521228 | 0.0087006 | 0.0206886 | 0.0233135 | 0.0281919 | 0.0019584 | 0.014737 | 0.0165361 | 0.0012867 | 0.0057616 | 0.0149286 | 0.0025087 | 0.0085089 |
| 27.75 | 0.0147899 | 0.0013987 | 0.005146 | 0.0085266 | 0.0140249 | 0.0107844 | 0.0389848 | 0.004057 | 0.0094495 | 0.0296058 | 0.0225551 | 0.0015372 | 0.0062315 | 0.0056416 | 0.0009048 | 0.0075136 | 0.0060239 | 0.0023421 | 0.0066538 |
| 27.76 | 0.0109152 | 0.0031658 | 0.0035746 | 0.0258066 | 0.0131223 | 0.0346236 | 0.041483 | 0.003929 | 0.0073287 | 0.0472087 | 0.0193373 | 0.0032004 | 0.0057033 | 0.0096774 | 0.002079 | 0.0129715 | 0.0049167 | 0.0054564 | 0.0097904 |
| 27.78 | 0.0139806 | 0.0052527 | 0.0032842 | 0.0350654 | 0.0182466 | 0.0447532 | 0.026544 | 0.0045912 | 0.0085096 | 0.0355375 | 0.0112127 | 0.0047767 | 0.0090892 | 0.0137256 | 0.0023098 | 0.0112773 | 0.0065225 | 0.0080028 | 0.0075023 |
| 27.79 | 0.0211181 | 0.0111225 | 0.0053009 | 0.0391615 | 0.0362805 | 0.0486667 | 0.0130357 | 0.0054753 | 0.008597 | 0.0135613 | 0.008585 | 0.0115335 | 0.0202811 | 0.0186972 | 0.0021908 | 0.004701 | 0.0163882 | 0.0127648 | 0.0020716 |
| 27.81 | 0.016914 | 0.0121576 | 0.0057625 | 0.0335644 | 0.0399046 | 0.0431895 | 0.022079 | 0.004146 | 0.0061367 | 0.0140931 | 0.00832 | 0.0142306 | 0.0214342 | 0.0178792 | 0.0019347 | 0.0022492 | 0.0186918 | 0.0130796 | 0.0016507 |
| 27.82 | 0.0070902 | 0.0055246 | 0.0051542 | 0.0119346 | 0.0234693 | 0.016008 | 0.0335216 | 0.0009055 | 0.0065138 | 0.0097298 | 0.0048393 | 0.0089259 | 0.009857 | 0.0090948 | 0.0010455 | 0.0003149 | 0.0096293 | 0.0074958 | 0.003205 |
| 27.83 | 0.0161989 | 0.0034194 | 0.0197182 | 0.0052964 | 0.0220742 | 0.0132886 | 0.0620611 | 0.0007004 | 0.020848 | 0.0141295 | 0.0207497 | 0.0063508 | 0.0151859 | 0.0112052 | 0.0033314 | 0.0005737 | 0.0066532 | 0.0075728 | 0.0107419 |
| 27.85 | 0.0170646 | 0.0029293 | 0.0257783 | 0.0036477 | 0.014938 | 0.02269 | 0.0537397 | 0.0013114 | 0.0228977 | 0.0255628 | 0.0313775 | 0.0027885 | 0.0260789 | 0.0128126 | 0.0047384 | 0.0014564 | 0.0048996 | 0.0049603 | 0.0149332 |
| 27.86 | 0.0208532 | 0.0069255 | 0.0351281 | 0.0179211 | 0.0129267 | 0.058673 | 0.0260363 | 0.0038768 | 0.0146675 | 0.0518433 | 0.0402305 | 0.0030629 | 0.0540234 | 0.0224308 | 0.0070828 | 0.0057032 | 0.0120304 | 0.0014239 | 0.0246303 |
| 27.88 | 0.0296966 | 0.0134009 | 0.0432195 | 0.0299775 | 0.0180708 | 0.0689694 | 0.0224007 | 0.004036 | 0.0098442 | 0.0507428 | 0.0347406 | 0.0069242 | 0.058503 | 0.0271282 | 0.0079181 | 0.0079584 | 0.0195262 | 0.001566 | 0.0288774 |
| 27.89 | 0.0264729 | 0.019626 | 0.0359805 | 0.029907 | 0.0170711 | 0.0421241 | 0.0220141 | 0.0023016 | 0.0047615 | 0.0187852 | 0.010596 | 0.0090003 | 0.0312491 | 0.0195054 | 0.0048935 | 0.0059241 | 0.0213911 | 0.0036433 | 0.0213909 |
| 27.9 | 0.0350758 | 0.0388289 | 0.0443551 | 0.0485771 | 0.022384 | 0.0428368 | 0.0455405 | 0.007748 | 0.0101224 | 0.0083073 | 0.0032215 | 0.0111607 | 0.022585 | 0.0246286 | 0.0039072 | 0.0062027 | 0.0373429 | 0.0116965 | 0.0289614 |
| 27.92 | 0.032667 | 0.0382613 | 0.0309021 | 0.0497218 | 0.0162789 | 0.0342716 | 0.0462142 | 0.0114226 | 0.0142548 | 0.005213 | 0.0028057 | 0.0071508 | 0.0112519 | 0.0215746 | 0.0020923 | 0.005108 | 0.0360789 | 0.0152229 | 0.0253473 |
| 27.93 | 0.0178476 | 0.0175736 | 0.0084505 | 0.0314103 | 0.0050387 | 0.0157478 | 0.0216566 | 0.0115474 | 0.0141399 | 0.0079918 | 0.007889 | 0.0013925 | 0.0016576 | 0.0114547 | 0.0006705 | 0.0039134 | 0.017626 | 0.013999 | 0.0124343 |
| 27.94 | 0.0133205 | 0.01258 | 0.0219051 | 0.0305228 | 0.0123376 | 0.0126389 | 0.0136773 | 0.0192781 | 0.0152871 | 0.0327258 | 0.0230069 | 0.0032481 | 0.0037605 | 0.0131209 | 0.003067 | 0.0091624 | 0.0125777 | 0.0243639 | 0.0151059 |
| 27.96 | 0.0089966 | 0.0125959 | 0.0293077 | 0.0201891 | 0.0175926 | 0.0075236 | 0.0104864 | 0.0167066 | 0.0101126 | 0.0426794 | 0.0251075 | 0.0052575 | 0.0061299 | 0.0099026 | 0.0051883 | 0.011207 | 0.0066409 | 0.0247696 | 0.0143849 |
| 27.97 | 0.0133194 | 0.0235314 | 0.0292074 | 0.0180988 | 0.0259058 | 0.0031843 | 0.0239309 | 0.0082975 | 0.010878 | 0.0359906 | 0.0157953 | 0.0065512 | 0.0089479 | 0.0045352 | 0.0087639 | 0.0121783 | 0.0026946 | 0.0245471 | 0.0123313 |
| 27.99 | 0.0165879 | 0.0237431 | 0.0236143 | 0.023845 | 0.0319072 | 0.0052928 | 0.0306924 | 0.0107845 | 0.0138133 | 0.0243669 | 0.0091543 | 0.0050704 | 0.008689 | 0.0049187 | 0.008979 | 0.0098174 | 0.0034194 | 0.0245843 | 0.009981 |
| 28 | 0.0265424 | 0.0249498 | 0.0111779 | 0.0181925 | 0.0396463 | 0.0162882 | 0.0190477 | 0.0224485 | 0.0123488 | 0.0077398 | 0.0027636 | 0.0021694 | 0.0058479 | 0.0041054 | 0.0055038 | 0.0024212 | 0.0037503 | 0.0172134 | 0.0042894 |
| 28.01 | 0.0994761 | 0.0946115 | 0.0448694 | 0.0445041 | 0.098317 | 0.0614632 | 0.0342424 | 0.0614655 | 0.0452348 | 0.0135908 | 0.0160002 | 0.0056121 | 0.0187658 | 0.0037266 | 0.0194647 | 0.0083534 | 0.0051899 | 0.0388717 | 0.0182043 |
| 28.03 | 0.1221198 | 0.1197137 | 0.0770735 | 0.0671932 | 0.0987169 | 0.0706066 | 0.0478339 | 0.0662523 | 0.0670739 | 0.0241284 | 0.0341155 | 0.005734 | 0.0338666 | 0.0058399 | 0.0420734 | 0.0241168 | 0.0061245 | 0.058015 | 0.0410793 |
| 28.04 | 0.1135014 | 0.1277632 | 0.1242167 | 0.0900937 | 0.0466346 | 0.0450831 | 0.0554879 | 0.0472157 | 0.0794375 | 0.0644628 | 0.072509 | 0.010747 | 0.0783004 | 0.0198891 | 0.1149717 | 0.081922 | 0.0180562 | 0.0962964 | 0.1031458 |
| 28.06 | 0.0948136 | 0.1180726 | 0.1163652 | 0.0769851 | 0.023058 | 0.0270146 | 0.0431403 | 0.0321986 | 0.069866 | 0.0795897 | 0.0772792 | 0.0204578 | 0.0881044 | 0.0249281 | 0.1306493 | 0.1017213 | 0.0264641 | 0.0906498 | 0.1119357 |
| 28.07 | 0.0357672 | 0.0537859 | 0.0416983 | 0.0232745 | 0.0060978 | 0.0051008 | 0.0127879 | 0.0087316 | 0.0366779 | 0.0495019 | 0.0419798 | 0.0180262 | 0.0392445 | 0.0145434 | 0.0564467 | 0.0555597 | 0.0237556 | 0.030598 | 0.0460658 |
| 28.08 | 0.0191735 | 0.0338527 | 0.0219429 | 0.0129524 | 0.0109177 | 0.0020643 | 0.008024 | 0.0125551 | 0.0436242 | 0.0339925 | 0.0313225 | 0.0137242 | 0.0220253 | 0.0102498 | 0.0255111 | 0.0320321 | 0.0283034 | 0.012774 | 0.0197215 |
| 28.1 | 0.0124721 | 0.0221898 | 0.0210927 | 0.0169764 | 0.0115663 | 0.0021902 | 0.0058697 | 0.015157 | 0.0320093 | 0.0185857 | 0.0193799 | 0.01273 | 0.0203906 | 0.0070152 | 0.0182463 | 0.0182159 | 0.0183805 | 0.007698 | 0.0139139 |
| 28.11 | 0.0248627 | 0.0438532 | 0.046553 | 0.033871 | 0.0114321 | 0.0041859 | 0.0043144 | 0.0148378 | 0.0115415 | 0.020465 | 0.0291485 | 0.0173027 | 0.0261442 | 0.0090254 | 0.0202296 | 0.0202778 | 0.0068806 | 0.0132877 | 0.0154795 |
| 28.13 | 0.0331283 | 0.0585615 | 0.0571327 | 0.0389959 | 0.011136 | 0.0041346 | 0.0032901 | 0.0117241 | 0.0099618 | 0.0229255 | 0.0356462 | 0.0151769 | 0.0239252 | 0.0097898 | 0.0165253 | 0.0216561 | 0.0093299 | 0.015101 | 0.0102528 |
| 28.14 | 0.0275057 | 0.0472436 | 0.0406265 | 0.0302563 | 0.0055365 | 0.0013573 | 0.002648 | 0.0042318 | 0.0062689 | 0.0146673 | 0.0228882 | 0.0053533 | 0.0133525 | 0.0057865 | 0.0083741 | 0.0139451 | 0.009517 | 0.0076432 | 0.0021204 |
| 28.15 | 0.0435792 | 0.0735162 | 0.0488019 | 0.0444172 | 0.0041398 | 0.0023594 | 0.0122384 | 0.0071944 | 0.00327 | 0.0249537 | 0.0341406 | 0.0025262 | 0.0138315 | 0.0067613 | 0.0116503 | 0.0207536 | 0.011677 | 0.0048488 | 0.0024067 |
| 28.17 | 0.0417519 | 0.0703106 | 0.0409143 | 0.0392459 | 0.0032256 | 0.0056376 | 0.0172991 | 0.0069736 | 0.0036317 | 0.0283933 | 0.0377175 | 0.0019439 | 0.0098715 | 0.0058329 | 0.0118675 | 0.022031 | 0.00997 | 0.0029716 | 0.0024881 |
| 28.18 | 0.0216569 | 0.0356042 | 0.0212413 | 0.0183744 | 0.0028924 | 0.0087143 | 0.0154455 | 0.0032989 | 0.0116633 | 0.021461 | 0.029865 | 0.0029573 | 0.0044455 | 0.0030066 | 0.0087294 | 0.0171448 | 0.0067055 | 0.0024917 | 0.0029 |
| 28.19 | 0.0237769 | 0.0332608 | 0.0278428 | 0.0218352 | 0.0055284 | 0.014855 | 0.0254465 | 0.0092423 | 0.0408543 | 0.0280569 | 0.0437321 | 0.009248 | 0.0099461 | 0.0026154 | 0.0149233 | 0.0265071 | 0.0126376 | 0.0097206 | 0.0091018 |

| | | | | | | | | | | | | | | | | | | | |
|---|---|---|---|---|---|---|---|---|---|---|---|---|---|---|---|---|---|---|---|
| 28.21 | 0.0211976 | 0.025115 | 0.0266725 | 0.0208448 | 0.0050374 | 0.0136413 | 0.0248179 | 0.0109784 | 0.0481291 | 0.0232554 | 0.0376261 | 0.0101998 | 0.0116751 | 0.0016218 | 0.0147401 | 0.0237239 | 0.0140258 | 0.0134187 | 0.0094675 |
| 28.22 | 0.0176523 | 0.0129858 | 0.0263153 | 0.0191356 | 0.002762 | 0.011117 | 0.0220135 | 0.0067882 | 0.0409421 | 0.0143332 | 0.0169013 | 0.0059 | 0.008145 | 0.0027225 | 0.0091523 | 0.0112841 | 0.0170485 | 0.0133434 | 0.0107377 |
| 28.24 | 0.0159057 | 0.0083945 | 0.0271194 | 0.0182059 | 0.0029018 | 0.011049 | 0.0224475 | 0.0040087 | 0.0338604 | 0.0111176 | 0.0104906 | 0.0047606 | 0.0059415 | 0.0049589 | 0.0072806 | 0.0070637 | 0.017617 | 0.0139312 | 0.015412 |
| 28.25 | 0.0097589 | 0.0037167 | 0.0194135 | 0.011039 | 0.0034352 | 0.0081078 | 0.0169946 | 0.0011427 | 0.0188627 | 0.0039216 | 0.0072153 | 0.0037338 | 0.0055119 | 0.0053877 | 0.005363 | 0.0053618 | 0.0093911 | 0.0116658 | 0.0119268 |
| 28.26 | 0.0170993 | 0.0093252 | 0.0299254 | 0.012779 | 0.0059236 | 0.0118774 | 0.0245168 | 0.0023945 | 0.0310041 | 0.0032928 | 0.0094634 | 0.005909 | 0.0095064 | 0.0061764 | 0.0057484 | 0.0065833 | 0.0087368 | 0.0123376 | 0.00881 |
| 28.28 | 0.0191012 | 0.0126378 | 0.0283832 | 0.0093392 | 0.0054228 | 0.0104538 | 0.021551 | 0.0043503 | 0.0310747 | 0.0033247 | 0.010057 | 0.0082922 | 0.0078387 | 0.0042881 | 0.0053625 | 0.0054133 | 0.0078738 | 0.0119296 | 0.0074664 |
| 28.29 | 0.0193047 | 0.0156658 | 0.0195539 | 0.0031529 | 0.0040881 | 0.0061568 | 0.0139928 | 0.0075417 | 0.0190752 | 0.0025393 | 0.0197427 | 0.0136842 | 0.0066657 | 0.0041697 | 0.0102258 | 0.0100394 | 0.0080854 | 0.0197324 | 0.0115475 |
| 28.31 | 0.0158038 | 0.0126009 | 0.0138849 | 0.0023499 | 0.0035766 | 0.0043258 | 0.0105917 | 0.006408 | 0.0130899 | 0.0017725 | 0.0187824 | 0.0121413 | 0.0066943 | 0.0051038 | 0.0099503 | 0.0102926 | 0.0064173 | 0.0182602 | 0.0121826 |
| 28.32 | 0.0046342 | 0.003957 | 0.003715 | 0.0017645 | 0.002018 | 0.0014873 | 0.0038062 | 0.0024193 | 0.0120883 | 0.0012687 | 0.0061973 | 0.0043268 | 0.0047313 | 0.0027021 | 0.004198 | 0.0049685 | 0.0021923 | 0.0078562 | 0.0107789 |
| 28.33 | 0.0020913 | 0.0069722 | 0.0019337 | 0.0051829 | 0.0034368 | 0.0008657 | 0.0038989 | 0.0061633 | 0.0276771 | 0.0046793 | 0.0110577 | 0.0058409 | 0.0098968 | 0.0019398 | 0.0082944 | 0.0102322 | 0.0059648 | 0.0149964 | 0.0191142 |
| 28.35 | 0.0030497 | 0.0070127 | 0.0023018 | 0.0106999 | 0.0042905 | 0.0003778 | 0.0048316 | 0.0074765 | 0.023929 | 0.0055698 | 0.0113466 | 0.0060108 | 0.0091198 | 0.0024122 | 0.0089331 | 0.0105472 | 0.0075127 | 0.0140152 | 0.0154065 |
| 28.36 | 0.0129904 | 0.0074439 | 0.009568 | 0.0263078 | 0.0061412 | 0.0004402 | 0.009328 | 0.0045377 | 0.0116387 | 0.0063229 | 0.0083682 | 0.0093394 | 0.0050588 | 0.0040676 | 0.0049524 | 0.0057029 | 0.0047626 | 0.0068806 | 0.0051869 |
| 28.38 | 0.0186384 | 0.0112303 | 0.0162214 | 0.0309133 | 0.005879 | 0.0011415 | 0.0119271 | 0.0029833 | 0.0132215 | 0.0105487 | 0.0127167 | 0.0140983 | 0.0066914 | 0.0039727 | 0.0036195 | 0.0049255 | 0.0031461 | 0.0072883 | 0.00359 |
| 28.39 | 0.0161851 | 0.012954 | 0.0183513 | 0.0193234 | 0.0020718 | 0.0022033 | 0.0102209 | 0.0025194 | 0.0103367 | 0.0142981 | 0.0163763 | 0.0116231 | 0.0083104 | 0.0022574 | 0.0043404 | 0.0060669 | 0.0017745 | 0.0057259 | 0.0033656 |
| 28.4 | 0.021581 | 0.0203691 | 0.0294415 | 0.0156939 | 0.0010263 | 0.0055094 | 0.0129769 | 0.0036952 | 0.0118685 | 0.0221091 | 0.0246008 | 0.008478 | 0.0132545 | 0.0027153 | 0.0096334 | 0.0124372 | 0.0015493 | 0.0041529 | 0.0051747 |
| 28.42 | 0.0161933 | 0.0163007 | 0.0241639 | 0.0096681 | 0.0012168 | 0.0059806 | 0.0092624 | 0.0041237 | 0.0178011 | 0.0176328 | 0.0191501 | 0.0037769 | 0.0120271 | 0.0025899 | 0.0095552 | 0.0123129 | 0.0011169 | 0.0027746 | 0.0047172 |
| 28.43 | 0.0064475 | 0.0084038 | 0.0080176 | 0.0067553 | 0.0020904 | 0.003374 | 0.003229 | 0.0062085 | 0.0249733 | 0.0060786 | 0.0052158 | 0.0017886 | 0.0061991 | 0.0023949 | 0.0055441 | 0.0071523 | 0.0020226 | 0.0019587 | 0.0032061 |
| 28.44 | 0.0150111 | 0.0228326 | 0.0136041 | 0.0103849 | 0.0054075 | 0.0038786 | 0.0050861 | 0.0139367 | 0.041963 | 0.0077369 | 0.0029521 | 0.0097632 | 0.0059153 | 0.0051896 | 0.006767 | 0.0080585 | 0.010119 | 0.0020436 | 0.0044908 |
| 28.46 | 0.015811 | 0.0260902 | 0.0161685 | 0.0084484 | 0.0058408 | 0.0037052 | 0.0047996 | 0.0131238 | 0.0337006 | 0.0103335 | 0.0030297 | 0.015432 | 0.00391 | 0.0055941 | 0.0053147 | 0.0056999 | 0.014968 | 0.0014203 | 0.0034018 |
| 28.47 | 0.0083844 | 0.0176678 | 0.0120817 | 0.0129491 | 0.0034642 | 0.0046185 | 0.0055677 | 0.0069149 | 0.0119509 | 0.0181633 | 0.0100754 | 0.0210642 | 0.0021697 | 0.0047787 | 0.0021049 | 0.0012878 | 0.0196764 | 0.0011316 | 0.001138 |
| 28.49 | 0.0115389 | 0.0137032 | 0.0131302 | 0.0190268 | 0.0037514 | 0.0087581 | 0.0084218 | 0.0071612 | 0.0083699 | 0.020386 | 0.0170435 | 0.0180477 | 0.0033479 | 0.0038718 | 0.0017549 | 0.0009361 | 0.0178395 | 0.0016726 | 0.0010059 |
| 28.5 | 0.0290314 | 0.017966 | 0.0209411 | 0.0217347 | 0.0107182 | 0.0204996 | 0.0128268 | 0.0106064 | 0.0099297 | 0.0123222 | 0.017448 | 0.0054371 | 0.0052943 | 0.0017296 | 0.0031041 | 0.0028017 | 0.0070474 | 0.0013696 | 0.0024518 |
| 28.51 | 0.0888056 | 0.0655402 | 0.0554275 | 0.0469349 | 0.043159 | 0.0633838 | 0.0435324 | 0.0321734 | 0.038236 | 0.0099691 | 0.0236704 | 0.0020415 | 0.0175581 | 0.0017898 | 0.0140796 | 0.0123782 | 0.0059053 | 0.0065441 | 0.0191937 |
| 28.53 | 0.0970437 | 0.0759625 | 0.0659365 | 0.0577258 | 0.0566339 | 0.0693162 | 0.0582749 | 0.0441185 | 0.0574867 | 0.0052293 | 0.0216473 | 0.0030512 | 0.0289098 | 0.0016454 | 0.0250053 | 0.0186786 | 0.0053239 | 0.0166614 | 0.0414997 |
| 28.54 | 0.0642709 | 0.0540729 | 0.08433 | 0.1032472 | 0.0636955 | 0.0460071 | 0.0743995 | 0.0537173 | 0.0877724 | 0.0055211 | 0.0230633 | 0.017954 | 0.0552587 | 0.005779 | 0.0566895 | 0.0323307 | 0.0066364 | 0.0346263 | 0.0986344 |
| 28.56 | 0.0464752 | 0.0438609 | 0.0924302 | 0.129858 | 0.0608411 | 0.0350201 | 0.0778859 | 0.0460535 | 0.0926228 | 0.0138338 | 0.0244923 | 0.0287251 | 0.0596232 | 0.012797 | 0.0663888 | 0.037627 | 0.0122866 | 0.0302081 | 0.1070974 |
| 28.57 | 0.0247399 | 0.0275148 | 0.0612187 | 0.0917121 | 0.0370335 | 0.0197954 | 0.0507728 | 0.0179592 | 0.0543945 | 0.0248455 | 0.0200901 | 0.027098 | 0.0348988 | 0.018046 | 0.0378343 | 0.0247394 | 0.0178998 | 0.0106043 | 0.0528237 |
| 28.58 | 0.035036 | 0.0325583 | 0.0642918 | 0.0826148 | 0.0418063 | 0.0253937 | 0.0595092 | 0.0121599 | 0.0430481 | 0.0571011 | 0.0442336 | 0.0317622 | 0.0431552 | 0.0277854 | 0.0293518 | 0.0251513 | 0.0268052 | 0.011367 | 0.0444149 |
| 28.6 | 0.031323 | 0.0230557 | 0.0451662 | 0.0477324 | 0.0284486 | 0.0217191 | 0.0481549 | 0.0081273 | 0.0242074 | 0.0601336 | 0.0463913 | 0.0211861 | 0.0396118 | 0.0246139 | 0.0186722 | 0.01978 | 0.0227166 | 0.0083354 | 0.0327632 |
| 28.61 | 0.0257879 | 0.0073457 | 0.0191649 | 0.0190919 | 0.0084927 | 0.0191171 | 0.0285502 | 0.0121097 | 0.0176441 | 0.0414505 | 0.0270924 | 0.0065494 | 0.0296396 | 0.0178786 | 0.0101533 | 0.0126674 | 0.0133298 | 0.0041929 | 0.0178417 |
| 28.63 | 0.0253301 | 0.0060395 | 0.0191323 | 0.0268868 | 0.0082075 | 0.0213275 | 0.0215538 | 0.0127995 | 0.0189064 | 0.0275456 | 0.0244464 | 0.0046618 | 0.0215069 | 0.0127005 | 0.0081011 | 0.0086511 | 0.0090228 | 0.0051076 | 0.0115172 |
| 28.64 | 0.0162029 | 0.0083398 | 0.0211441 | 0.0273541 | 0.0106293 | 0.0174434 | 0.0109029 | 0.0041672 | 0.0116677 | 0.0129636 | 0.0338032 | 0.0028378 | 0.0063041 | 0.0036857 | 0.0032743 | 0.0030307 | 0.0051761 | 0.0052927 | 0.0038574 |
| 28.65 | 0.0224793 | 0.0185974 | 0.0267443 | 0.0292219 | 0.0234487 | 0.0247079 | 0.0153932 | 0.0010506 | 0.0190467 | 0.034693 | 0.0688974 | 0.0061845 | 0.0085931 | 0.0080438 | 0.0016908 | 0.0033868 | 0.0104785 | 0.0097316 | 0.0038034 |
| 28.67 | 0.0210631 | 0.0189808 | 0.0179807 | 0.0191538 | 0.0266268 | 0.0196568 | 0.0129401 | 0.000343 | 0.020496 | 0.0382574 | 0.0575938 | 0.0078055 | 0.0092163 | 0.0092043 | 0.0015612 | 0.0031763 | 0.0095585 | 0.010661 | 0.0048793 |
| 28.68 | 0.0091496 | 0.0104427 | 0.0090112 | 0.0052357 | 0.0187338 | 0.0072747 | 0.0057055 | 0.0005083 | 0.0159882 | 0.0161675 | 0.0186236 | 0.0045015 | 0.0038917 | 0.0040406 | 0.0032978 | 0.0043916 | 0.0030866 | 0.0085322 | 0.0094503 |
| 28.69 | 0.0090341 | 0.0115074 | 0.0212443 | 0.0121342 | 0.021307 | 0.0190963 | 0.0189733 | 0.0024821 | 0.0312681 | 0.017339 | 0.0377776 | 0.0043749 | 0.0020269 | 0.0035523 | 0.0081807 | 0.0105638 | 0.0028022 | 0.0139324 | 0.0191581 |
| 28.71 | 0.0160184 | 0.0097779 | 0.0224665 | 0.0210352 | 0.0144618 | 0.0293562 | 0.031643 | 0.0033667 | 0.0334918 | 0.0271141 | 0.0466546 | 0.005531 | 0.0017032 | 0.0046945 | 0.0085668 | 0.0107409 | 0.0031173 | 0.0140736 | 0.0152292 |
| 28.72 | 0.0370042 | 0.0077484 | 0.0143947 | 0.0354769 | 0.0070388 | 0.0429471 | 0.0515039 | 0.002794 | 0.0253671 | 0.0464822 | 0.0294678 | 0.0080386 | 0.0033642 | 0.0077827 | 0.0071346 | 0.0074116 | 0.0020892 | 0.0151954 | 0.005255 |
| 28.74 | 0.0373423 | 0.0071269 | 0.0101625 | 0.0300185 | 0.012416 | 0.0394221 | 0.0492618 | 0.0023291 | 0.0167817 | 0.0420961 | 0.0178804 | 0.0073603 | 0.0048491 | 0.0083084 | 0.0063932 | 0.0047083 | 0.0017589 | 0.0152732 | 0.0043296 |
| 28.75 | 0.0150804 | 0.0045719 | 0.0069868 | 0.0085708 | 0.0209939 | 0.0130019 | 0.017902 | 0.0023599 | 0.0030825 | 0.0140688 | 0.0145342 | 0.0031782 | 0.0039145 | 0.0058704 | 0.003825 | 0.0008114 | 0.0029943 | 0.0085902 | 0.0029034 |
| 28.76 | 0.0085133 | 0.0078848 | 0.0171795 | 0.018018 | 0.0432842 | 0.0060241 | 0.0119976 | 0.0050331 | 0.0094292 | 0.0145636 | 0.0329247 | 0.0025417 | 0.0033302 | 0.0062227 | 0.0059679 | 0.0019647 | 0.0085896 | 0.0098777 | 0.0043016 |
| 28.78 | 0.0056944 | 0.0089784 | 0.0257842 | 0.021548 | 0.0424261 | 0.0056484 | 0.0167206 | 0.005585 | 0.0169463 | 0.0205921 | 0.0323488 | 0.0027692 | 0.0021847 | 0.0048535 | 0.0066666 | 0.0032598 | 0.0089806 | 0.0089122 | 0.0061557 |
| 28.79 | 0.0113842 | 0.0116905 | 0.053789 | 0.0117486 | 0.0285774 | 0.007039 | 0.0354661 | 0.0086269 | 0.0234295 | 0.0327624 | 0.0246975 | 0.0067836 | 0.003584 | 0.0057591 | 0.0075934 | 0.0041118 | 0.0071172 | 0.008794 | 0.0090747 |
| 28.81 | 0.015041 | 0.0138253 | 0.0551803 | 0.0079148 | 0.0197487 | 0.0054413 | 0.0392514 | 0.0101771 | 0.020446 | 0.0330623 | 0.0230643 | 0.0085655 | 0.0057426 | 0.0050357 | 0.0068634 | 0.0031784 | 0.009411 | 0.0100968 | 0.0074828 |
| 28.82 | 0.0119976 | 0.0135023 | 0.0201324 | 0.0098048 | 0.0097538 | 0.0025427 | 0.0237225 | 0.0057088 | 0.0082386 | 0.01816 | 0.0167013 | 0.0059573 | 0.0057174 | 0.0009711 | 0.0027562 | 0.0013917 | 0.012334 | 0.0105795 | 0.0035503 |
| 28.83 | 0.0171927 | 0.0262795 | 0.0225609 | 0.0235306 | 0.0310942 | 0.0061829 | 0.0219744 | 0.0046697 | 0.0103148 | 0.0206683 | 0.0281518 | 0.0076725 | 0.0082006 | 0.002715 | 0.0034034 | 0.0047376 | 0.0239168 | 0.023954 | 0.0135115 |
| 28.85 | 0.014133 | 0.0267121 | 0.0288646 | 0.0245956 | 0.0394066 | 0.0077083 | 0.0146109 | 0.0055438 | 0.0135792 | 0.0199526 | 0.0299453 | 0.0080402 | 0.00854 | 0.0058824 | 0.0047652 | 0.0063902 | 0.0246188 | 0.0263912 | 0.0193373 |
| 28.86 | 0.0146685 | 0.0238911 | 0.0382143 | 0.0290879 | 0.0404451 | 0.0099201 | 0.0207798 | 0.0110646 | 0.0282877 | 0.0270682 | 0.035585 | 0.0139051 | 0.0146131 | 0.0121961 | 0.0074546 | 0.0071037 | 0.0246061 | 0.0226948 | 0.0267046 |
| 28.88 | 0.0256024 | 0.0250068 | 0.0331316 | 0.0372497 | 0.0389079 | 0.0113902 | 0.0397583 | 0.0113383 | 0.0381518 | 0.032454 | 0.0394954 | 0.0191283 | 0.0184959 | 0.0141399 | 0.0073254 | 0.0067054 | 0.0237408 | 0.0185015 | 0.0284619 |
| 28.89 | 0.0345315 | 0.021186 | 0.0152283 | 0.0366399 | 0.0244641 | 0.0097239 | 0.0544018 | 0.0050608 | 0.0363886 | 0.0248718 | 0.0299763 | 0.0188872 | 0.0134082 | 0.0125329 | 0.0039443 | 0.0044945 | 0.0132772 | 0.008974 | 0.0184614 |
| 28.9 | 0.0595543 | 0.0359391 | 0.0514609 | 0.0607117 | 0.0276997 | 0.0156697 | 0.0871098 | 0.0111869 | 0.0655715 | 0.0301028 | 0.0429025 | 0.0295513 | 0.0132284 | 0.0271003 | 0.0060424 | 0.0088969 | 0.0181435 | 0.0146779 | 0.0235697 |
| 28.92 | 0.0524339 | 0.0346679 | 0.072025 | 0.0561567 | 0.0211075 | 0.0151725 | 0.0692788 | 0.0184193 | 0.0671027 | 0.0222768 | 0.0391197 | 0.0240794 | 0.0080792 | 0.0293071 | 0.0068044 | 0.0101994 | 0.0192139 | 0.0165781 | 0.0211245 |
| 28.93 | 0.0189363 | 0.0176614 | 0.0519729 | 0.0278263 | 0.008878 | 0.0091581 | 0.019306 | 0.0195166 | 0.0386311 | 0.0058014 | 0.0207824 | 0.007029 | 0.0013248 | 0.0159371 | 0.0044547 | 0.0065004 | 0.0134102 | 0.0115593 | 0.0110523 |
| 28.94 | 0.0117626 | 0.013919 | 0.0431916 | 0.0262319 | 0.0102563 | 0.0098603 | 0.0086165 | 0.0295412 | 0.0368451 | 0.0025459 | 0.0209533 | 0.0059505 | 0.0018102 | 0.0115231 | 0.0032078 | 0.0047483 | 0.0152482 | 0.0120969 | 0.0097075 |
| 28.96 | 0.0117041 | 0.0080488 | 0.0261308 | 0.0177397 | 0.0080702 | 0.0064218 | 0.0071537 | 0.0258894 | 0.0241166 | 0.0028077 | 0.014655 | 0.0051762 | 0.0039569 | 0.0060614 | 0.0016195 | 0.0020536 | 0.0122531 | 0.0085843 | 0.005962 |
| 28.97 | 0.0173393 | 0.0028987 | 0.0382538 | 0.0131529 | 0.0064174 | 0.0020757 | 0.013663 | 0.0146135 | 0.015775 | 0.0067135 | 0.0067113 | 0.0040414 | 0.0126687 | 0.0042374 | 0.0019835 | 0.0007663 | 0.0106671 | 0.006215 | 0.0034872 |
| 28.99 | 0.019371 | 0.0082565 | 0.0599533 | 0.0162827 | 0.007001 | 0.0040505 | 0.0166866 | 0.0152428 | 0.0227123 | 0.0082348 | 0.0065337 | 0.0057237 | 0.0165048 | 0.0045507 | 0.002675 | 0.0013709 | 0.0108591 | 0.0088867 | 0.0037774 |
| 29 | 0.029773 | 0.0315874 | 0.0559157 | 0.0130995 | 0.0106333 | 0.0148037 | 0.0100395 | 0.0304663 | 0.0196714 | 0.0070305 | 0.0066652 | 0.0048919 | 0.0113344 | 0.0026527 | 0.0017987 | 0.0018117 | 0.0057736 | 0.0134619 | 0.0021536 |
| 29.01 | 0.1107921 | 0.1244292 | 0.0614406 | 0.0445044 | 0.0520339 | 0.0595297 | 0.0170832 | 0.0983561 | 0.0407604 | 0.0170628 | 0.0195308 | 0.0068405 | 0.0174396 | 0.0085378 | 0.0103329 | 0.0113551 | 0.0053746 | 0.0485053 | 0.0130961 |
| 29.03 | 0.143691 | 0.1665251 | 0.0613014 | 0.0712025 | 0.0695331 | 0.0760486 | 0.0319009 | 0.1135922 | 0.0664717 | 0.0251026 | 0.0341059 | 0.0142362 | 0.0269141 | 0.0162388 | 0.0286548 | 0.0278441 | 0.0056547 | 0.085623 | 0.0325882 |
| 29.04 | 0.1460906 | 0.1988093 | 0.12779 | 0.0978614 | 0.0654026 | 0.0731622 | 0.059179 | 0.0990192 | 0.1108863 | 0.0521698 | 0.0727042 | 0.0506901 | 0.0652399 | 0.0382496 | 0.0806049 | 0.0777442 | 0.0171724 | 0.1869588 | 0.0835557 |
| 29.06 | 0.1186612 | 0.1843397 | 0.1495765 | 0.0877229 | 0.0495761 | 0.0564997 | 0.054108 | 0.0785064 | 0.1097778 | 0.06284 | 0.0792489 | 0.0682052 | 0.0765928 | 0.0453275 | 0.087091 | 0.0882103 | 0.0272194 | 0.2102187 | 0.091902 |
| 29.07 | 0.0402228 | 0.0849483 | 0.0869397 | 0.0346174 | 0.013537 | 0.0159958 | 0.0206214 | 0.0256504 | 0.0613956 | 0.0355887 | 0.0445572 | 0.0471587 | 0.0399343 | 0.0286291 | 0.0375625 | 0.0432726 | 0.022448 | 0.1190297 | 0.0448496 |
| 29.08 | 0.024658 | 0.0543905 | 0.0841309 | 0.0185056 | 0.0114621 | 0.0083124 | 0.0123733 | 0.01469 | 0.0704841 | 0.0208033 | 0.043636 | 0.0341221 | 0.0260154 | 0.0216381 | 0.0184316 | 0.0251544 | 0.0167266 | 0.1013585 | 0.0263578 |
| 29.1 | 0.0235539 | 0.0277402 | 0.0597163 | 0.0100138 | 0.0151059 | 0.0082962 | 0.0073244 | 0.0091393 | 0.0532837 | 0.009985 | 0.0288484 | 0.0178663 | 0.0195905 | 0.0117818 | 0.0082723 | 0.0140225 | 0.0081315 | 0.0589778 | 0.0126667 |
| 29.11 | 0.0527286 | 0.0336515 | 0.0216766 | 0.015193 | 0.0246484 | 0.0155758 | 0.0127431 | 0.013902 | 0.0173743 | 0.0050121 | 0.0122501 | 0.0217128 | 0.0263348 | 0.0137859 | 0.0051728 | 0.0140249 | 0.0073644 | 0.0097268 | 0.009263 |
| 29.13 | 0.0684488 | 0.0468037 | 0.0135204 | 0.0196651 | 0.0269056 | 0.01587 | 0.0171861 | 0.0219465 | 0.007734 | 0.0043148 | 0.0131969 | 0.0292463 | 0.0278392 | 0.0185274 | 0.0052703 | 0.012051 | 0.0094307 | 0.0037818 | 0.0091359 |
| 29.14 | 0.0548042 | 0.0368895 | 0.0171565 | 0.0149987 | 0.0181717 | 0.0065395 | 0.0135485 | 0.0231585 | 0.0040216 | 0.0032973 | 0.0126311 | 0.0235274 | 0.0180036 | 0.0152286 | 0.0035515 | 0.0022722 | 0.0056345 | 0.0046778 | 0.0056573 |
| 29.15 | 0.0731611 | 0.0435479 | 0.0566375 | 0.0232163 | 0.022462 | 0.0033302 | 0.0220973 | 0.0398868 | 0.009156 | 0.0049541 | 0.0261318 | 0.0329332 | 0.0261001 | 0.0212797 | 0.004332 | 0.0023911 | 0.0034912 | 0.0189099 | 0.0155321 |
| 29.17 | 0.0631522 | 0.0357979 | 0.077993 | 0.0248895 | 0.0180902 | 0.0036645 | 0.0259849 | 0.0410915 | 0.0100244 | 0.0041977 | 0.0319712 | 0.0286705 | 0.0247817 | 0.0188324 | 0.0053779 | 0.0052917 | 0.0016749 | 0.0248206 | 0.020356 |
| 29.18 | 0.0314611 | 0.0230085 | 0.0768662 | 0.0188009 | 0.0070408 | 0.0074555 | 0.0249168 | 0.0271622 | 0.0080746 | 0.0029332 | 0.0280363 | 0.0125082 | 0.0123561 | 0.0102388 | 0.004805 | 0.0055771 | 0.0019462 | 0.0151825 | 0.0144601 |
| 29.19 | 0.0347165 | 0.0518655 | 0.1316657 | 0.0244354 | 0.004681 | 0.0168714 | 0.0438121 | 0.0343739 | 0.0178863 | 0.0088346 | 0.0451315 | 0.0095841 | 0.0077771 | 0.0139758 | 0.0046684 | 0.0065371 | 0.0057272 | 0.0245476 | 0.0111757 |
| 29.21 | 0.027629 | 0.0640657 | 0.1276711 | 0.0187012 | 0.0022251 | 0.0173437 | 0.0423432 | 0.0291117 | 0.0226894 | 0.0115724 | 0.042982 | 0.0063305 | 0.0032187 | 0.0129954 | 0.0037322 | 0.006531 | 0.005267 | 0.0323877 | 0.0057374 |
| 29.22 | 0.0144193 | 0.0766569 | 0.1045481 | 0.0065104 | 0.0005889 | 0.0158331 | 0.0288411 | 0.0199811 | 0.0345612 | 0.0133785 | 0.0346916 | 0.0051062 | 0.001701 | 0.0077551 | 0.0024181 | 0.0045756 | 0.0028636 | 0.0386749 | 0.0041395 |
| 29.24 | 0.0093525 | 0.0740719 | 0.0894515 | 0.0039249 | 0.0007547 | 0.0150477 | 0.0213696 | 0.016553 | 0.0395216 | 0.01259 | 0.0293559 | 0.0054412 | 0.0025809 | 0.0052112 | 0.0019789 | 0.0028153 | 0.0035335 | 0.0354148 | 0.0070393 |
| 29.25 | 0.0029571 | 0.0392753 | 0.0366857 | 0.0030733 | 0.0008959 | 0.0087661 | 0.0092355 | 0.006674 | 0.029095 | 0.0068937 | 0.0123224 | 0.0027266 | 0.0022168 | 0.00335 | 0.002427 | 0.0029442 | 0.0028812 | 0.0194269 | 0.0093151 |
| 29.26 | 0.0081469 | 0.0430221 | 0.0246621 | 0.0052529 | 0.0038077 | 0.0125609 | 0.0102776 | 0.0044379 | 0.0482031 | 0.0081719 | 0.0094063 | 0.0031852 | 0.0029968 | 0.0058042 | 0.0067454 | 0.0085987 | 0.0038075 | 0.0227248 | 0.0143069 |
| 29.28 | 0.0135485 | 0.0376802 | 0.0157393 | 0.007562 | 0.0064247 | 0.0136331 | 0.0092767 | 0.0051882 | 0.0508317 | 0.0078169 | 0.0057345 | 0.0043816 | 0.0029377 | 0.0048262 | 0.0067409 | 0.0082547 | 0.0047334 | 0.0210625 | 0.0112715 |
| 29.29 | 0.0240513 | 0.0307852 | 0.0077159 | 0.0183517 | 0.0101758 | 0.0139699 | 0.0095976 | 0.011062 | 0.048947 | 0.0110097 | 0.0019513 | 0.005208 | 0.0043811 | 0.0048017 | 0.0028179 | 0.0036437 | 0.0048162 | 0.0227926 | 0.0044694 |
| 29.31 | 0.0265674 | 0.0263461 | 0.0042451 | 0.0219466 | 0.0095129 | 0.0128457 | 0.0101174 | 0.0109656 | 0.0465207 | 0.0127422 | 0.0020531 | 0.0054488 | 0.0044909 | 0.0066879 | 0.0017989 | 0.0025577 | 0.0032819 | 0.022219 | 0.0025016 |
| 29.32 | 0.0178341 | 0.0119496 | 0.0006481 | 0.0138117 | 0.0038736 | 0.0069112 | 0.007038 | 0.0041378 | 0.029134 | 0.0075064 | 0.0036066 | 0.0055387 | 0.0020769 | 0.004973 | 0.0017899 | 0.0014749 | 0.0009416 | 0.0117946 | 0.0020622 |
| 29.33 | 0.0235888 | 0.0115158 | 0.001083 | 0.0137584 | 0.0024635 | 0.005521 | 0.0084631 | 0.002368 | 0.0389913 | 0.0064701 | 0.0106972 | 0.0070807 | 0.0063136 | 0.0030611 | 0.0051402 | 0.0052263 | 0.0033035 | 0.0124275 | 0.0084015 |
| 29.35 | 0.0212555 | 0.0091972 | 0.0014383 | 0.0099275 | 0.0021093 | 0.003446 | 0.0061936 | 0.0014686 | 0.0363353 | 0.0049089 | 0.0110264 | 0.0052809 | 0.0101612 | 0.0012776 | 0.0073459 | 0.0086206 | 0.0054578 | 0.0086558 | 0.0108396 |
| 29.36 | 0.0192361 | 0.0094878 | 0.0025873 | 0.0068554 | 0.0066447 | 0.0023651 | 0.0019197 | 0.0012707 | 0.0348223 | 0.0042178 | 0.0045361 | 0.0075362 | 0.0139689 | 0.0019462 | 0.0099969 | 0.0113571 | 0.0071717 | 0.0034701 | 0.0088954 |
| 29.38 | 0.0191188 | 0.0120212 | 0.0027747 | 0.0075226 | 0.0107232 | 0.0019032 | 0.0009963 | 0.0015461 | 0.0348894 | 0.0043076 | 0.0017667 | 0.0093421 | 0.0114454 | 0.0038327 | 0.0083881 | 0.0092107 | 0.0057145 | 0.0061459 | 0.0062813 |
| 29.39 | 0.0106782 | 0.0110571 | 0.0025578 | 0.0069271 | 0.0112462 | 0.0007357 | 0.0021186 | 0.0015788 | 0.0234025 | 0.003019 | 0.0007257 | 0.0051837 | 0.0048101 | 0.0053693 | 0.0031843 | 0.0040534 | 0.0020261 | 0.0114528 | 0.0021279 |
| 29.4 | 0.0079164 | 0.0172351 | 0.0112048 | 0.0147909 | 0.0170476 | 0.0022119 | 0.0112811 | 0.0043213 | 0.0362986 | 0.0048709 | 0.0053977 | 0.0090835 | 0.0097684 | 0.0093236 | 0.0035147 | 0.0056214 | 0.0030944 | 0.0203639 | 0.0022394 |
| 29.42 | 0.0048293 | 0.0151832 | 0.0170243 | 0.0171643 | 0.0150886 | 0.0032516 | 0.016386 | 0.0044248 | 0.0354838 | 0.0079228 | 0.0104453 | 0.0144131 | 0.009712 | 0.0080789 | 0.0031388 | 0.005205 | 0.0047187 | 0.0155643 | 0.00346 |
| 29.43 | 0.0038844 | 0.0054266 | 0.0164391 | 0.0123733 | 0.0073345 | 0.0021532 | 0.014791 | 0.0020684 | 0.0193625 | 0.0130057 | 0.0133875 | 0.0165683 | 0.0066781 | 0.0032164 | 0.0031559 | 0.0046461 | 0.0089483 | 0.0034267 | 0.005201 |
| 29.44 | 0.0050782 | 0.002476 | 0.025059 | 0.0160938 | 0.0072998 | 0.0021958 | 0.0227788 | 0.0048073 | 0.0198203 | 0.0262994 | 0.0206971 | 0.0235653 | 0.0178783 | 0.00236 | 0.0051263 | 0.0076817 | 0.0223764 | 0.0031459 | 0.0099892 |
| 29.46 | 0.0037987 | 0.0017643 | 0.0219873 | 0.0141603 | 0.0052929 | 0.0022376 | 0.0201849 | 0.0047654 | 0.014668 | 0.0243314 | 0.0167602 | 0.0172711 | 0.0203087 | 0.0025592 | 0.0040941 | 0.0061154 | 0.0220929 | 0.0029397 | 0.0101771 |
| 29.47 | 0.0096315 | 0.0041756 | 0.0140857 | 0.0124083 | 0.0024552 | 0.002193 | 0.0111119 | 0.0022909 | 0.0060794 | 0.0165621 | 0.0099926 | 0.0049567 | 0.015818 | 0.0095141 | 0.0038482 | 0.004024 | 0.0138144 | 0.0033439 | 0.0118234 |
| 29.49 | 0.0197278 | 0.0091841 | 0.0117724 | 0.0155444 | 0.0042365 | 0.0066227 | 0.0085077 | 0.0058928 | 0.0045273 | 0.0153307 | 0.0105678 | 0.0038842 | 0.0120856 | 0.0142529 | 0.0046512 | 0.0040613 | 0.0108093 | 0.0049802 | 0.0133554 |
| 29.5 | 0.0360835 | 0.0210438 | 0.0069639 | 0.0222465 | 0.0128347 | 0.0233935 | 0.0049938 | 0.0203273 | 0.0073676 | 0.0112794 | 0.0104258 | 0.0064961 | 0.0055765 | 0.0118826 | 0.0039876 | 0.0030168 | 0.0070234 | 0.0042904 | 0.0103146 |
| 29.51 | 0.0938856 | 0.0668312 | 0.0171449 | 0.0592453 | 0.0510341 | 0.0690419 | 0.0084297 | 0.0705236 | 0.0372826 | 0.0180235 | 0.0236279 | 0.019979 | 0.0163979 | 0.0132874 | 0.0115409 | 0.0084425 | 0.0139786 | 0.0090623 | 0.0238607 |
| 29.53 | 0.1012826 | 0.0767562 | 0.0298349 | 0.068105 | 0.0694234 | 0.0712219 | 0.0172225 | 0.0818869 | 0.0582146 | 0.0180871 | 0.0327647 | 0.0247972 | 0.0307584 | 0.0106092 | 0.0222414 | 0.0144603 | 0.0151605 | 0.0168086 | 0.0389402 |
| 29.54 | 0.0784953 | 0.055214 | 0.0671933 | 0.074982 | 0.0810544 | 0.0473303 | 0.0538042 | 0.0624908 | 0.0825117 | 0.0241319 | 0.0591168 | 0.0304073 | 0.0708885 | 0.0167192 | 0.0542152 | 0.0276085 | 0.0156442 | 0.0255461 | 0.0801425 |
| 29.56 | 0.0645221 | 0.0397577 | 0.0835986 | 0.0777383 | 0.07226 | 0.0365732 | 0.0667605 | 0.0461934 | 0.0825611 | 0.0337799 | 0.0673085 | 0.0320752 | 0.0774786 | 0.0255684 | 0.0589824 | 0.0289875 | 0.018979 | 0.0257362 | 0.087134 |
| 29.57 | 0.0333133 | 0.0184441 | 0.0644117 | 0.0487122 | 0.0339477 | 0.0167959 | 0.0436703 | 0.0201222 | 0.0534528 | 0.0335333 | 0.0431592 | 0.0225152 | 0.0374691 | 0.0251296 | 0.0251007 | 0.0146696 | 0.0186421 | 0.031106 | 0.0453923 |

| | | | | | | | | | | | | | | | | | | | |
|---|---|---|---|---|---|---|---|---|---|---|---|---|---|---|---|---|---|---|---|
| 29.58 | 0.032718 | 0.0252304 | 0.0685217 | 0.0515755 | 0.038642 | 0.0185165 | 0.0455754 | 0.0267029 | 0.0651789 | 0.0466812 | 0.0520363 | 0.0384929 | 0.0345216 | 0.0374183 | 0.0101631 | 0.0111791 | 0.0272011 | 0.0624052 | 0.0437774 |
| 29.6 | 0.0209922 | 0.0246619 | 0.0419262 | 0.0379952 | 0.0370529 | 0.0157995 | 0.0348123 | 0.0216297 | 0.046825 | 0.0409838 | 0.0449399 | 0.0396706 | 0.0294421 | 0.0334333 | 0.0027633 | 0.0069933 | 0.0227812 | 0.0537712 | 0.0372802 |
| 29.61 | 0.0095984 | 0.023526 | 0.0191294 | 0.0152795 | 0.0399868 | 0.0128869 | 0.0216161 | 0.0097626 | 0.0144585 | 0.0308142 | 0.0234512 | 0.0231711 | 0.0196751 | 0.0171991 | 0.00107 | 0.0034304 | 0.0107062 | 0.0287085 | 0.025776 |
| 29.63 | 0.011785 | 0.0210989 | 0.0320502 | 0.0074771 | 0.0388092 | 0.0127591 | 0.0187051 | 0.0089365 | 0.00877 | 0.0238701 | 0.0177867 | 0.0151332 | 0.0131334 | 0.0107362 | 0.0020909 | 0.0038372 | 0.0073472 | 0.0248506 | 0.0174103 |
| 29.64 | 0.0126918 | 0.0082007 | 0.0413527 | 0.0014124 | 0.0195342 | 0.0098735 | 0.0127519 | 0.0051567 | 0.0058514 | 0.006752 | 0.0276438 | 0.0156732 | 0.0035057 | 0.0088427 | 0.0027284 | 0.004366 | 0.0068654 | 0.0194553 | 0.0071811 |
| 29.65 | 0.0236014 | 0.0103388 | 0.0667202 | 0.0033063 | 0.0139285 | 0.0142193 | 0.021146 | 0.005014 | 0.011836 | 0.0026313 | 0.0760296 | 0.0371729 | 0.0059763 | 0.0221889 | 0.0052239 | 0.0067216 | 0.013256 | 0.0328222 | 0.0177063 |
| 29.67 | 0.0241175 | 0.0180554 | 0.0597146 | 0.0052691 | 0.0089515 | 0.0138722 | 0.0201698 | 0.0056588 | 0.0136956 | 0.0016387 | 0.0755015 | 0.037216 | 0.0064035 | 0.0236484 | 0.004718 | 0.0052531 | 0.0129506 | 0.0310809 | 0.0187424 |
| 29.68 | 0.0111286 | 0.0233818 | 0.0247753 | 0.007325 | 0.0107117 | 0.0105035 | 0.009166 | 0.004997 | 0.0075166 | 0.0010534 | 0.0271387 | 0.0170084 | 0.0027027 | 0.0125903 | 0.001616 | 0.0014738 | 0.0096769 | 0.0153102 | 0.0072888 |
| 29.69 | 0.0108163 | 0.0390112 | 0.0308281 | 0.0160838 | 0.0309673 | 0.0092268 | 0.0063473 | 0.007586 | 0.0049678 | 0.0043188 | 0.0220486 | 0.0112706 | 0.0032651 | 0.0087809 | 0.0011715 | 0.0016444 | 0.0214659 | 0.0143865 | 0.0056543 |
| 29.71 | 0.0144221 | 0.0370919 | 0.0414602 | 0.0174044 | 0.037902 | 0.0056259 | 0.0057255 | 0.0070488 | 0.005237 | 0.0084468 | 0.0291822 | 0.0097389 | 0.0029845 | 0.0052898 | 0.0016431 | 0.0024727 | 0.0244404 | 0.0097326 | 0.0048244 |
| 29.72 | 0.0210769 | 0.031732 | 0.0615352 | 0.0186478 | 0.048402 | 0.0121032 | 0.0129762 | 0.0050498 | 0.0127195 | 0.0162088 | 0.0380643 | 0.0168077 | 0.0021656 | 0.0081828 | 0.0047247 | 0.0059274 | 0.0182016 | 0.003537 | 0.004242 |
| 29.74 | 0.0190808 | 0.0295457 | 0.0617769 | 0.0201133 | 0.0478734 | 0.0162927 | 0.0177859 | 0.0036758 | 0.016984 | 0.0165033 | 0.0289958 | 0.0187904 | 0.003098 | 0.0102083 | 0.0052753 | 0.0061819 | 0.0132676 | 0.0034405 | 0.0056223 |
| 29.75 | 0.0078449 | 0.0158456 | 0.0350827 | 0.0160281 | 0.0204309 | 0.0082422 | 0.01751 | 0.0009241 | 0.0132319 | 0.0081139 | 0.0112062 | 0.0115762 | 0.0025249 | 0.0068816 | 0.0029933 | 0.0027577 | 0.0111916 | 0.0026452 | 0.0042173 |
| 29.76 | 0.0066992 | 0.0147914 | 0.0341794 | 0.0235087 | 0.0145095 | 0.0023244 | 0.0321295 | 0.0003944 | 0.0131996 | 0.0073434 | 0.0303469 | 0.0119621 | 0.0026318 | 0.008667 | 0.0063075 | 0.0057245 | 0.0261816 | 0.0056665 | 0.0048017 |
| 29.78 | 0.0087382 | 0.0115075 | 0.0226009 | 0.0192408 | 0.0185244 | 0.0003579 | 0.0318991 | 0.0009697 | 0.0105767 | 0.0052995 | 0.0371981 | 0.0080593 | 0.0036934 | 0.0067123 | 0.007154 | 0.0069344 | 0.0255957 | 0.0093034 | 0.0062928 |
| 29.79 | 0.0238889 | 0.0170884 | 0.0198925 | 0.009367 | 0.0383548 | 0.0026331 | 0.0251517 | 0.0083502 | 0.0162196 | 0.0026105 | 0.0347601 | 0.0048589 | 0.0062279 | 0.0054788 | 0.0043899 | 0.0046681 | 0.0140128 | 0.0169859 | 0.0067088 |
| 29.81 | 0.0343722 | 0.0260306 | 0.032804 | 0.0095895 | 0.0415876 | 0.0065846 | 0.0199645 | 0.0175359 | 0.0212163 | 0.0031233 | 0.0319029 | 0.0076994 | 0.0057423 | 0.0078373 | 0.0031481 | 0.0035773 | 0.0081945 | 0.0187903 | 0.0062496 |
| 29.82 | 0.0339559 | 0.0290037 | 0.0407295 | 0.0124835 | 0.0238502 | 0.0085331 | 0.006227 | 0.0258473 | 0.0171035 | 0.0062948 | 0.0223506 | 0.0101366 | 0.0027272 | 0.0062525 | 0.0026001 | 0.004325 | 0.0028539 | 0.0122687 | 0.0098906 |
| 29.83 | 0.0548307 | 0.0520348 | 0.0749515 | 0.028026 | 0.020039 | 0.0080983 | 0.0025752 | 0.0486199 | 0.0266674 | 0.0214437 | 0.045322 | 0.0182626 | 0.007754 | 0.0040997 | 0.0031494 | 0.0092978 | 0.0092914 | 0.0124751 | 0.0292534 |
| 29.85 | 0.0502655 | 0.0513652 | 0.0718332 | 0.0304376 | 0.0137958 | 0.0058923 | 0.0013658 | 0.045755 | 0.0282356 | 0.0280821 | 0.0530834 | 0.0173216 | 0.0138517 | 0.0022685 | 0.0018567 | 0.0086234 | 0.0144208 | 0.0080615 | 0.0323727 |
| 29.86 | 0.0289195 | 0.0414471 | 0.0508443 | 0.0291385 | 0.0240822 | 0.0049077 | 0.0027766 | 0.032063 | 0.0250005 | 0.0348295 | 0.0634948 | 0.0132955 | 0.0271883 | 0.0052517 | 0.0013392 | 0.0049537 | 0.0278742 | 0.004678 | 0.0251532 |
| 29.88 | 0.0166567 | 0.0355468 | 0.0389573 | 0.0279097 | 0.0347815 | 0.0039526 | 0.0059382 | 0.0257362 | 0.0175381 | 0.0341503 | 0.0653603 | 0.0131552 | 0.0294022 | 0.0091792 | 0.0021575 | 0.0032603 | 0.0346103 | 0.00641 | 0.018886 |
| 29.89 | 0.0030341 | 0.0179927 | 0.0140075 | 0.0192802 | 0.0303955 | 0.0024501 | 0.0101878 | 0.0113049 | 0.0052435 | 0.0196424 | 0.0422357 | 0.0120258 | 0.0170452 | 0.0123894 | 0.0027968 | 0.0014622 | 0.0280346 | 0.0068736 | 0.0068994 |
| 29.9 | 0.0068384 | 0.0187098 | 0.0160831 | 0.0315855 | 0.0485353 | 0.0032673 | 0.0221883 | 0.0084133 | 0.0124419 | 0.0247682 | 0.055175 | 0.0245806 | 0.0163499 | 0.0288359 | 0.0057835 | 0.0033181 | 0.0396554 | 0.0118705 | 0.0060827 |
| 29.92 | 0.0100294 | 0.0140865 | 0.0137903 | 0.0304355 | 0.0523116 | 0.0025351 | 0.0201832 | 0.0062636 | 0.0155432 | 0.021913 | 0.0481826 | 0.0249347 | 0.010562 | 0.032261 | 0.0050592 | 0.0031802 | 0.0334081 | 0.01194 | 0.0043379 |
| 29.93 | 0.0094829 | 0.0051804 | 0.0072664 | 0.0135565 | 0.0347362 | 0.0022442 | 0.0072284 | 0.0081439 | 0.0128959 | 0.0102528 | 0.0222847 | 0.0117762 | 0.0033383 | 0.0200954 | 0.0023816 | 0.0014838 | 0.0125881 | 0.0074399 | 0.0024362 |
| 29.94 | 0.0136778 | 0.0031457 | 0.0282553 | 0.0073852 | 0.030233 | 0.0070959 | 0.0135039 | 0.0177147 | 0.0221296 | 0.0073335 | 0.0191615 | 0.007469 | 0.0067909 | 0.0191411 | 0.0071009 | 0.0054064 | 0.0091958 | 0.0083515 | 0.0057515 |
| 29.96 | 0.0103539 | 0.0022347 | 0.0412495 | 0.0049266 | 0.0175918 | 0.0069644 | 0.0190969 | 0.0167717 | 0.020455 | 0.0054537 | 0.0122392 | 0.0047651 | 0.0077282 | 0.011882 | 0.0087975 | 0.0081092 | 0.0067017 | 0.0056655 | 0.0061294 |
| 29.97 | 0.0047616 | 0.002716 | 0.0545906 | 0.0092234 | 0.0087584 | 0.0029849 | 0.0285634 | 0.0086688 | 0.0115605 | 0.0109858 | 0.0030454 | 0.0051526 | 0.0063775 | 0.0036424 | 0.0075122 | 0.0108464 | 0.0047202 | 0.0016819 | 0.0040178 |
| 29.99 | 0.0099707 | 0.011503 | 0.0537144 | 0.0096315 | 0.0088834 | 0.0031973 | 0.0345927 | 0.0118733 | 0.013245 | 0.0133813 | 0.0011996 | 0.0045902 | 0.0047973 | 0.003238 | 0.0056787 | 0.0100309 | 0.0037619 | 0.0012636 | 0.002544 |
| 30 | 0.0303797 | 0.0494139 | 0.0253571 | 0.0112456 | 0.0097034 | 0.012432 | 0.0313795 | 0.0392038 | 0.0326702 | 0.0074828 | 0.0009432 | 0.0026086 | 0.0020587 | 0.0019368 | 0.0022421 | 0.0038925 | 0.0022997 | 0.0011414 | 0.0012816 |
| 30.01 | 0.1128422 | 0.1861464 | 0.0315442 | 0.06359 | 0.0495408 | 0.0580989 | 0.0356913 | 0.1357086 | 0.1191021 | 0.0054015 | 0.0120474 | 0.0099741 | 0.0139557 | 0.0040029 | 0.0153863 | 0.0093146 | 0.0081721 | 0.0029114 | 0.0132419 |
| 30.03 | 0.1400186 | 0.2239875 | 0.0575646 | 0.0947803 | 0.0737045 | 0.0710567 | 0.0319069 | 0.1486173 | 0.1479597 | 0.0044473 | 0.0302161 | 0.0201635 | 0.0340843 | 0.0101565 | 0.0389334 | 0.0241389 | 0.0097846 | 0.0096711 | 0.0307388 |
| 30.04 | 0.1330015 | 0.1891339 | 0.1406956 | 0.1132475 | 0.0859278 | 0.053624 | 0.0752041 | 0.0852016 | 0.1498148 | 0.0146597 | 0.0783826 | 0.057994 | 0.0907483 | 0.0395943 | 0.0977032 | 0.0735211 | 0.01973 | 0.0565369 | 0.0734205 |
| 30.06 | 0.1134172 | 0.1467129 | 0.154039 | 0.0966188 | 0.074192 | 0.0401093 | 0.0909292 | 0.0477828 | 0.1300855 | 0.0219046 | 0.0863649 | 0.0751273 | 0.1002312 | 0.0559894 | 0.1002198 | 0.0861874 | 0.0342409 | 0.0766591 | 0.0774282 |
| 30.07 | 0.0531843 | 0.0591555 | 0.0810425 | 0.0348287 | 0.030552 | 0.0171551 | 0.0423853 | 0.0097767 | 0.0579421 | 0.0141512 | 0.0424405 | 0.0498386 | 0.0456462 | 0.0412949 | 0.035676 | 0.0446576 | 0.0318982 | 0.0400396 | 0.0313329 |
| 30.08 | 0.0478285 | 0.0549765 | 0.0643213 | 0.0221276 | 0.0203813 | 0.0120596 | 0.0272794 | 0.0085966 | 0.059425 | 0.0118157 | 0.025385 | 0.0352646 | 0.0243856 | 0.0301976 | 0.014498 | 0.0283949 | 0.0241023 | 0.0318234 | 0.0181385 |
| 30.1 | 0.0310045 | 0.0365969 | 0.0362772 | 0.0152357 | 0.0118929 | 0.007151 | 0.0281942 | 0.0073401 | 0.0486412 | 0.0123774 | 0.0108771 | 0.0189349 | 0.0137472 | 0.0150982 | 0.0120778 | 0.0205071 | 0.0122443 | 0.0322894 | 0.015125 |
| 30.11 | 0.0228093 | 0.0194251 | 0.0132092 | 0.0238662 | 0.0162328 | 0.0130286 | 0.0553113 | 0.004077 | 0.0230608 | 0.0119048 | 0.0018817 | 0.0224483 | 0.0191727 | 0.0166885 | 0.0206928 | 0.0332478 | 0.0118163 | 0.0297689 | 0.0126437 |
| 30.13 | 0.0350087 | 0.0305863 | 0.0145507 | 0.0329874 | 0.0244115 | 0.0187125 | 0.0625746 | 0.0046625 | 0.016527 | 0.0074684 | 0.0025267 | 0.0305689 | 0.0246121 | 0.0230563 | 0.0231963 | 0.0394814 | 0.0162667 | 0.0212666 | 0.0088188 |
| 30.14 | 0.0413852 | 0.0397694 | 0.0124173 | 0.0286026 | 0.0235794 | 0.016356 | 0.0423712 | 0.0049972 | 0.019084 | 0.0013896 | 0.0054389 | 0.0291499 | 0.0200281 | 0.0200024 | 0.018105 | 0.0278185 | 0.0154644 | 0.0075754 | 0.0045253 |
| 30.15 | 0.0753751 | 0.0744229 | 0.0220421 | 0.0406365 | 0.0359705 | 0.0258239 | 0.058961 | 0.0059431 | 0.0434906 | 0.0013836 | 0.0119685 | 0.055211 | 0.0282871 | 0.0323352 | 0.0237793 | 0.0329625 | 0.0244013 | 0.0079796 | 0.007306 |
| 30.17 | 0.076539 | 0.0773247 | 0.0270887 | 0.034719 | 0.034325 | 0.0269175 | 0.053537 | 0.0051795 | 0.0444512 | 0.0022163 | 0.0123766 | 0.0611243 | 0.0259305 | 0.0329489 | 0.0173126 | 0.0239903 | 0.0224307 | 0.0059738 | 0.0068299 |
| 30.18 | 0.0478034 | 0.0438361 | 0.0271672 | 0.0124664 | 0.022394 | 0.0213382 | 0.0261788 | 0.0055113 | 0.0296296 | 0.002815 | 0.0105299 | 0.0469124 | 0.0153974 | 0.0232618 | 0.0042368 | 0.0060386 | 0.0119698 | 0.0014992 | 0.0041826 |
| 30.19 | 0.0640657 | 0.0299153 | 0.0525422 | 0.0064432 | 0.0339622 | 0.0380635 | 0.0221575 | 0.0158833 | 0.0515157 | 0.0034386 | 0.0253327 | 0.0703167 | 0.0205491 | 0.0361505 | 0.0055131 | 0.0035273 | 0.0142222 | 0.00165 | 0.0091917 |
| 30.21 | 0.0627868 | 0.0143631 | 0.0573496 | 0.0070837 | 0.0320006 | 0.0395045 | 0.0145598 | 0.0217303 | 0.0540104 | 0.002818 | 0.0296083 | 0.0625331 | 0.0173762 | 0.0346758 | 0.0066292 | 0.0030765 | 0.0117625 | 0.0026657 | 0.0108045 |
| 30.22 | 0.0710212 | 0.0172727 | 0.0725685 | 0.0243484 | 0.023269 | 0.0392339 | 0.0121266 | 0.0361325 | 0.0554238 | 0.0049166 | 0.0302955 | 0.0344736 | 0.0080512 | 0.0248651 | 0.0058033 | 0.0041562 | 0.0054615 | 0.0048317 | 0.0080787 |
| 30.24 | 0.0735847 | 0.028128 | 0.0835058 | 0.0367931 | 0.0182934 | 0.0392608 | 0.0180222 | 0.0423623 | 0.0551164 | 0.0060698 | 0.030743 | 0.0204618 | 0.004421 | 0.0178228 | 0.0043641 | 0.0048819 | 0.0030106 | 0.0058208 | 0.0053298 |
| 30.25 | 0.0440648 | 0.0303745 | 0.0631436 | 0.0332599 | 0.0078681 | 0.0251843 | 0.0219773 | 0.0313382 | 0.0332971 | 0.0070829 | 0.0264144 | 0.0039505 | 0.0017217 | 0.004413 | 0.0020135 | 0.0027199 | 0.0014285 | 0.0057259 | 0.0029611 |
| 30.26 | 0.0479505 | 0.0534675 | 0.0867722 | 0.0392334 | 0.0094376 | 0.0321789 | 0.0406624 | 0.0437413 | 0.0451866 | 0.0201926 | 0.0504388 | 0.0032479 | 0.0044381 | 0.002631 | 0.0044391 | 0.003323 | 0.003916 | 0.0106056 | 0.007338 |
| 30.28 | 0.0344739 | 0.0539972 | 0.0791056 | 0.0271021 | 0.0091871 | 0.0280144 | 0.0397929 | 0.0395705 | 0.048195 | 0.0216897 | 0.0495352 | 0.0036087 | 0.0055266 | 0.0030259 | 0.0044992 | 0.0034355 | 0.0041089 | 0.0098903 | 0.0086149 |
| 30.29 | 0.0098851 | 0.0522151 | 0.0682097 | 0.0056082 | 0.0110347 | 0.0214444 | 0.0310386 | 0.029853 | 0.0669375 | 0.0150137 | 0.0333807 | 0.0112448 | 0.0084006 | 0.0039278 | 0.0044729 | 0.0050909 | 0.0055873 | 0.006287 | 0.0081707 |
| 30.31 | 0.0066333 | 0.0504225 | 0.0648159 | 0.0028622 | 0.0139111 | 0.020372 | 0.0285117 | 0.0274166 | 0.0761005 | 0.0129868 | 0.0253893 | 0.0168482 | 0.01056 | 0.0047903 | 0.0065735 | 0.0088434 | 0.0094388 | 0.0052685 | 0.0062717 |
| 30.32 | 0.0107692 | 0.0280205 | 0.0393316 | 0.0066031 | 0.0147624 | 0.0139417 | 0.0219807 | 0.02043 | 0.0561144 | 0.0119301 | 0.0130046 | 0.0130517 | 0.0085247 | 0.0042625 | 0.0060745 | 0.0103833 | 0.0121758 | 0.0035357 | 0.0015752 |
| 30.33 | 0.0311413 | 0.032901 | 0.0519067 | 0.0200911 | 0.0328933 | 0.021949 | 0.0404924 | 0.0393243 | 0.0850219 | 0.0310828 | 0.0218182 | 0.0111529 | 0.0116933 | 0.006484 | 0.0084241 | 0.0161314 | 0.0244587 | 0.0067403 | 0.0023317 |
| 30.35 | 0.0362943 | 0.031149 | 0.0431493 | 0.0238292 | 0.0383743 | 0.0210445 | 0.0390092 | 0.0402317 | 0.0760674 | 0.0367493 | 0.0214887 | 0.0065864 | 0.011943 | 0.0093835 | 0.0094321 | 0.0159945 | 0.0249144 | 0.0075972 | 0.0053987 |
| 30.36 | 0.0320066 | 0.0293094 | 0.0186185 | 0.0225897 | 0.0399457 | 0.0117991 | 0.0210373 | 0.0236242 | 0.0357665 | 0.0337381 | 0.0123798 | 0.0088555 | 0.0173149 | 0.0193478 | 0.0119506 | 0.0155565 | 0.0190792 | 0.0088122 | 0.013976 |
| 30.38 | 0.0257659 | 0.0253154 | 0.0133118 | 0.0191797 | 0.0336201 | 0.0066136 | 0.0112344 | 0.0140419 | 0.0231186 | 0.0259004 | 0.0105044 | 0.0152343 | 0.0206592 | 0.0211096 | 0.0100059 | 0.0130525 | 0.0158217 | 0.009218 | 0.0155391 |
| 30.39 | 0.0143697 | 0.0098906 | 0.0094151 | 0.0108792 | 0.0109823 | 0.0019021 | 0.0022629 | 0.0063137 | 0.0118103 | 0.0082665 | 0.0108717 | 0.0184882 | 0.0150088 | 0.0118185 | 0.0025097 | 0.0039126 | 0.0079149 | 0.0057996 | 0.0081843 |
| 30.4 | 0.0346582 | 0.0075184 | 0.0087016 | 0.022642 | 0.0047241 | 0.0041868 | 0.0092664 | 0.0077403 | 0.0143272 | 0.011922 | 0.0129365 | 0.0315196 | 0.0165737 | 0.0137746 | 0.0020677 | 0.0020774 | 0.0093165 | 0.0056518 | 0.006423 |
| 30.42 | 0.0460566 | 0.006711 | 0.0049048 | 0.0286983 | 0.0028821 | 0.0070743 | 0.0148001 | 0.0048637 | 0.0208464 | 0.0168668 | 0.0109069 | 0.0268373 | 0.0113111 | 0.0128543 | 0.0031457 | 0.0018009 | 0.0083583 | 0.0035254 | 0.0037378 |
| 30.43 | 0.0402441 | 0.0063101 | 0.0027126 | 0.026401 | 0.0029994 | 0.0079521 | 0.0139052 | 0.001172 | 0.0264889 | 0.0166593 | 0.015847 | 0.0082056 | 0.0022781 | 0.0080051 | 0.0050874 | 0.0027002 | 0.0051726 | 0.0008056 | 0.0018955 |
| 30.44 | 0.0575031 | 0.0126777 | 0.005722 | 0.0450097 | 0.0047311 | 0.0104345 | 0.0193878 | 0.0008751 | 0.0373405 | 0.0212938 | 0.0395191 | 0.0051364 | 0.0019143 | 0.0107773 | 0.0107501 | 0.0085821 | 0.0089732 | 0.0015294 | 0.0112311 |
| 30.46 | 0.0509473 | 0.0152778 | 0.0053059 | 0.0438805 | 0.0045014 | 0.0082032 | 0.016347 | 0.000793 | 0.0278217 | 0.0156817 | 0.0408416 | 0.004287 | 0.0044658 | 0.0105511 | 0.0092851 | 0.0094361 | 0.0099044 | 0.0022452 | 0.0193776 |
| 30.47 | 0.0389729 | 0.0268274 | 0.002974 | 0.0372854 | 0.0075303 | 0.0059226 | 0.0098875 | 0.0020001 | 0.0241611 | 0.0075291 | 0.032412 | 0.0067042 | 0.014036 | 0.0119404 | 0.0031713 | 0.005909 | 0.0106046 | 0.0033972 | 0.032853 |
| 30.49 | 0.0414343 | 0.0420067 | 0.0024281 | 0.0375076 | 0.0092545 | 0.0109132 | 0.0093567 | 0.0068409 | 0.033331 | 0.0075112 | 0.026979 | 0.0133322 | 0.0178039 | 0.0129693 | 0.0017418 | 0.003922 | 0.0099766 | 0.0032953 | 0.033038 |
| 30.5 | 0.0499705 | 0.0661433 | 0.0039078 | 0.037188 | 0.0067946 | 0.0264656 | 0.0120104 | 0.0243396 | 0.0253481 | 0.0080278 | 0.0126588 | 0.0187363 | 0.0127011 | 0.0095238 | 0.0024669 | 0.0021715 | 0.0049171 | 0.0014183 | 0.0149112 |
| 30.51 | 0.1287343 | 0.1716548 | 0.0205318 | 0.0879514 | 0.0290237 | 0.0699141 | 0.0355417 | 0.0797502 | 0.0177015 | 0.011514 | 0.0171114 | 0.0285203 | 0.0235561 | 0.0141465 | 0.0187207 | 0.0082779 | 0.0049573 | 0.0052472 | 0.016842 |
| 30.53 | 0.1403787 | 0.1803437 | 0.036053 | 0.0963942 | 0.0524556 | 0.0678358 | 0.0455373 | 0.0874384 | 0.0144136 | 0.0097639 | 0.0225597 | 0.0216145 | 0.0368403 | 0.0132195 | 0.0383886 | 0.0133964 | 0.0037501 | 0.0108459 | 0.028476 |
| 30.54 | 0.1162114 | 0.110701 | 0.0824767 | 0.0931977 | 0.0880248 | 0.0352342 | 0.0654319 | 0.0663426 | 0.0426215 | 0.0134207 | 0.049098 | 0.0077631 | 0.0727723 | 0.0079079 | 0.0808656 | 0.0181304 | 0.0016584 | 0.0170794 | 0.065923 |
| 30.56 | 0.1026204 | 0.0766117 | 0.1049831 | 0.0902478 | 0.086388 | 0.0222936 | 0.0717105 | 0.0601235 | 0.0569374 | 0.0156925 | 0.062133 | 0.006151 | 0.0738344 | 0.006005 | 0.0799045 | 0.01567 | 0.0043758 | 0.0146863 | 0.0699899 |
| 30.57 | 0.0578316 | 0.0385731 | 0.0786703 | 0.0507029 | 0.0463578 | 0.0089345 | 0.0444245 | 0.0414727 | 0.0431728 | 0.0135389 | 0.0446917 | 0.0096355 | 0.0317679 | 0.0083854 | 0.0297484 | 0.0100102 | 0.0129744 | 0.0105126 | 0.0385862 |
| 30.58 | 0.0614547 | 0.0516637 | 0.0888926 | 0.0484404 | 0.0586867 | 0.0169349 | 0.0441918 | 0.0540618 | 0.0578087 | 0.0330263 | 0.0498222 | 0.0364691 | 0.031429 | 0.0266349 | 0.0220238 | 0.0250888 | 0.0348517 | 0.0229846 | 0.0537464 |
| 30.6 | 0.0418309 | 0.0447923 | 0.064664 | 0.033587 | 0.0557045 | 0.0189742 | 0.0288552 | 0.0471127 | 0.0525916 | 0.0381285 | 0.0377418 | 0.0445137 | 0.0306218 | 0.031093 | 0.0188447 | 0.0290219 | 0.0367214 | 0.0213923 | 0.0515803 |
| 30.61 | 0.0223481 | 0.0321737 | 0.0216721 | 0.011177 | 0.0553733 | 0.0192239 | 0.0099143 | 0.0359533 | 0.0404297 | 0.0345438 | 0.0153557 | 0.0354127 | 0.026955 | 0.0268435 | 0.0117875 | 0.0208248 | 0.0265879 | 0.0098788 | 0.0300005 |
| 30.63 | 0.0259659 | 0.0280274 | 0.0127214 | 0.0050753 | 0.054765 | 0.0186175 | 0.0104636 | 0.0282885 | 0.0305881 | 0.0287764 | 0.010926 | 0.0245024 | 0.020741 | 0.0195939 | 0.0083005 | 0.0147647 | 0.0183246 | 0.0058376 | 0.0196907 |
| 30.64 | 0.0217624 | 0.014668 | 0.0159988 | 0.0016574 | 0.0315419 | 0.0101676 | 0.0085294 | 0.0089888 | 0.0142073 | 0.012802 | 0.0146581 | 0.0106729 | 0.0059753 | 0.0049887 | 0.0073483 | 0.0117433 | 0.0049772 | 0.002425 | 0.0162137 |
| 30.65 | 0.0361044 | 0.0125533 | 0.0467744 | 0.0033263 | 0.0311266 | 0.0074998 | 0.0055965 | 0.0130254 | 0.0302104 | 0.0093452 | 0.0432798 | 0.0307965 | 0.0023919 | 0.0063388 | 0.015476 | 0.0270535 | 0.0087228 | 0.0031781 | 0.041491 |
| 30.67 | 0.0362739 | 0.0109469 | 0.0539563 | 0.0052482 | 0.0206262 | 0.005315 | 0.0025027 | 0.0203306 | 0.0340973 | 0.0045266 | 0.0529507 | 0.0374652 | 0.0013114 | 0.0077435 | 0.0141962 | 0.0269991 | 0.0100763 | 0.0024439 | 0.0472218 |
| 30.68 | 0.0194439 | 0.0145576 | 0.0392288 | 0.0065931 | 0.00884 | 0.0071796 | 0.0009263 | 0.0217135 | 0.0247544 | 0.0037605 | 0.0387764 | 0.019869 | 0.0018696 | 0.0060429 | 0.0049906 | 0.0115349 | 0.0052749 | 0.0019899 | 0.0292281 |
| 30.69 | 0.0285356 | 0.0314094 | 0.0641678 | 0.0108249 | 0.0278854 | 0.0168812 | 0.0026054 | 0.0348706 | 0.0373678 | 0.0171539 | 0.0470595 | 0.0141921 | 0.006157 | 0.0120146 | 0.0025708 | 0.0052669 | 0.0044898 | 0.0080473 | 0.0217917 |
| 30.71 | 0.0325415 | 0.0303475 | 0.054476 | 0.0086518 | 0.0365575 | 0.0173503 | 0.0040519 | 0.0340108 | 0.0301758 | 0.020506 | 0.0362943 | 0.0142382 | 0.007763 | 0.0137998 | 0.0018566 | 0.0024707 | 0.003512 | 0.0115169 | 0.0129144 |
| 30.72 | 0.033284 | 0.0180755 | 0.0229434 | 0.005791 | 0.0428705 | 0.018393 | 0.0057239 | 0.0334023 | 0.0186871 | 0.0118717 | 0.01383 | 0.0268541 | 0.0089607 | 0.0129395 | 0.0030036 | 0.0068809 | 0.0028754 | 0.0155644 | 0.0146739 |
| 30.74 | 0.0305429 | 0.0112563 | 0.0259939 | 0.0100049 | 0.0457423 | 0.0189313 | 0.0046347 | 0.0364667 | 0.0260864 | 0.0067338 | 0.0080576 | 0.0289308 | 0.0086503 | 0.0098761 | 0.0043939 | 0.0097744 | 0.0029836 | 0.0151441 | 0.013703 |
| 30.75 | 0.0158486 | 0.0046112 | 0.0282317 | 0.0150545 | 0.0355374 | 0.0086213 | 0.0018065 | 0.0294993 | 0.0317157 | 0.0046961 | 0.0041548 | 0.0156458 | 0.004083 | 0.0031724 | 0.0039083 | 0.0067457 | 0.0030747 | 0.0077734 | 0.0044793 |
| 30.76 | 0.0120213 | 0.0138002 | 0.0288827 | 0.0337864 | 0.0506612 | 0.0033706 | 0.0058041 | 0.0414642 | 0.0515085 | 0.0073411 | 0.0043359 | 0.0134482 | 0.0033339 | 0.0064426 | 0.0045257 | 0.0062407 | 0.0069179 | 0.0073311 | 0.0093756 |
| 30.78 | 0.0081184 | 0.0187385 | 0.0161864 | 0.0375345 | 0.0419509 | 0.0013145 | 0.007826 | 0.0346514 | 0.0412526 | 0.0066352 | 0.0041873 | 0.0093642 | 0.0037629 | 0.0084258 | 0.0037707 | 0.0044564 | 0.0074189 | 0.0052708 | 0.0110797 |
| 30.79 | 0.0081301 | 0.0239761 | 0.0085677 | 0.0327587 | 0.0180063 | 0.001583 | 0.0073167 | 0.0222175 | 0.0135639 | 0.0135999 | 0.004734 | 0.0139321 | 0.0062087 | 0.0082097 | 0.0032433 | 0.0024269 | 0.0056498 | 0.0030534 | 0.0083899 |
| 30.81 | 0.007709 | 0.0238027 | 0.0125229 | 0.0241825 | 0.0105932 | 0.001786 | 0.0060218 | 0.0186137 | 0.0089313 | 0.0181531 | 0.0041305 | 0.0201501 | 0.0057936 | 0.0059627 | 0.0029612 | 0.0017372 | 0.0038376 | 0.0030337 | 0.0062278 |
| 30.82 | 0.0040065 | 0.0133901 | 0.01189 | 0.0078673 | 0.008265 | 0.0021544 | 0.00523 | 0.0086539 | 0.0134287 | 0.0118036 | 0.0046724 | 0.0178946 | 0.0022863 | 0.0020569 | 0.0014879 | 0.0007124 | 0.0013966 | 0.0027346 | 0.00308 |
| 30.83 | 0.0086032 | 0.0151479 | 0.0196963 | 0.0187924 | 0.0200853 | 0.0103903 | 0.0172545 | 0.0073445 | 0.0402395 | 0.0100315 | 0.0200278 | 0.0280644 | 0.0050671 | 0.0073433 | 0.0021351 | 0.0020097 | 0.0027424 | 0.0049484 | 0.0084196 |
| 30.85 | 0.0129547 | 0.0152248 | 0.0260976 | 0.0265681 | 0.0197676 | 0.0160069 | 0.021772 | 0.0053907 | 0.0458774 | 0.0099826 | 0.0308954 | 0.0287977 | 0.0092393 | 0.0120375 | 0.0026959 | 0.004435 | 0.0027809 | 0.0051267 | 0.0108986 |
| 30.86 | 0.0225303 | 0.0272344 | 0.0646121 | 0.0402926 | 0.0114443 | 0.0174716 | 0.0263155 | 0.0037781 | 0.0406621 | 0.0172068 | 0.0512963 | 0.0286361 | 0.0200327 | 0.0204268 | 0.0041126 | 0.0108524 | 0.0050335 | 0.005716 | 0.0087673 |
| 30.88 | 0.024931 | 0.037257 | 0.0891837 | 0.0450665 | 0.0122002 | 0.0118298 | 0.029994 | 0.0031349 | 0.0351073 | 0.0175338 | 0.0563372 | 0.0267581 | 0.0234682 | 0.0214771 | 0.0044872 | 0.0123291 | 0.0079083 | 0.0064897 | 0.0059882 |
| 30.89 | 0.016896 | 0.0335255 | 0.0763708 | 0.0317669 | 0.0192054 | 0.0016283 | 0.0250265 | 0.0024075 | 0.019345 | 0.0097098 | 0.0385346 | 0.016299 | 0.0162577 | 0.0120585 | 0.0035332 | 0.0080437 | 0.0087369 | 0.0056975 | 0.0036305 |
| 30.9 | 0.0189553 | 0.0461619 | 0.1098432 | 0.0446515 | 0.0486497 | 0.0025119 | 0.0396226 | 0.0080795 | 0.0297148 | 0.0170707 | 0.0565905 | 0.0257077 | 0.019769 | 0.0129407 | 0.0056672 | 0.0095513 | 0.0185809 | 0.011269 | 0.0067602 |
| 30.92 | 0.0139345 | 0.0380048 | 0.0997379 | 0.0411263 | 0.0543142 | 0.0056375 | 0.0366669 | 0.0129366 | 0.0294526 | 0.0216382 | 0.0532827 | 0.0245908 | 0.0154698 | 0.0105101 | 0.0048452 | 0.0072159 | 0.0214293 | 0.0124905 | 0.0065898 |
| 30.93 | 0.0097992 | 0.0135071 | 0.0496134 | 0.0188215 | 0.0351389 | 0.0078428 | 0.0142772 | 0.0149586 | 0.0149925 | 0.0215782 | 0.0275152 | 0.0114307 | 0.0049929 | 0.0048182 | 0.0028864 | 0.0040194 | 0.0158563 | 0.0082604 | 0.004548 |
| 30.94 | 0.0369502 | 0.0084834 | 0.0392625 | 0.0135914 | 0.0389817 | 0.0126153 | 0.0138017 | 0.0238474 | 0.0119353 | 0.0376224 | 0.0284186 | 0.008846 | 0.0021964 | 0.0037789 | 0.0091538 | 0.0116954 | 0.0231675 | 0.0095574 | 0.0088155 |

| | | | | | | | | | | | | | | | | | | | |
|---|---|---|---|---|---|---|---|---|---|---|---|---|---|---|---|---|---|---|---|
| 30.96 | 0.0538276 | 0.0066338 | 0.025174 | 0.0137293 | 0.0279326 | 0.0112369 | 0.0238742 | 0.0204202 | 0.0110275 | 0.0346054 | 0.0224573 | 0.0055619 | 0.0008918 | 0.002639 | 0.0118001 | 0.013901 | 0.0216491 | 0.0080198 | 0.0096025 |
| 30.97 | 0.078316 | 0.0058038 | 0.0358543 | 0.0244877 | 0.0103395 | 0.0092953 | 0.0542683 | 0.0149164 | 0.0226834 | 0.0184649 | 0.0125505 | 0.0015611 | 0.0007673 | 0.0050099 | 0.0114174 | 0.0104484 | 0.0156786 | 0.0066098 | 0.0083572 |
| 30.99 | 0.0773278 | 0.0093593 | 0.0507773 | 0.0250866 | 0.0074449 | 0.0127592 | 0.0594071 | 0.0207035 | 0.0252156 | 0.0103359 | 0.0087053 | 0.0009618 | 0.0007145 | 0.0068736 | 0.0085956 | 0.0071327 | 0.0125835 | 0.0062824 | 0.006793 |
| 31 | 0.0398481 | 0.0268226 | 0.0356523 | 0.0134312 | 0.0078715 | 0.0230617 | 0.0288319 | 0.0327832 | 0.0158192 | 0.0013339 | 0.0030309 | 0.00159 | 0.001076 | 0.0047222 | 0.0030116 | 0.003526 | 0.0062698 | 0.0034767 | 0.0026278 |
| 31.01 | 0.0880315 | 0.0914324 | 0.0400825 | 0.042155 | 0.0405321 | 0.0620433 | 0.0343662 | 0.0791435 | 0.0425501 | 0.0013881 | 0.0132188 | 0.0031699 | 0.0163312 | 0.0051085 | 0.0183755 | 0.0143987 | 0.0083349 | 0.0056421 | 0.0119476 |
| 31.03 | 0.1212885 | 0.1039876 | 0.0519931 | 0.0693308 | 0.0624235 | 0.063405 | 0.049805 | 0.0789006 | 0.057649 | 0.0042859 | 0.0327344 | 0.0022352 | 0.0393435 | 0.0098898 | 0.0443377 | 0.0314077 | 0.0073212 | 0.0141185 | 0.0317699 |
| 31.04 | 0.1270699 | 0.0816629 | 0.0921771 | 0.0988311 | 0.0771904 | 0.0359216 | 0.0901092 | 0.039637 | 0.0681533 | 0.015641 | 0.0853318 | 0.0116766 | 0.099434 | 0.0423348 | 0.1143268 | 0.0926497 | 0.0153503 | 0.0589279 | 0.091876 |
| 31.06 | 0.1011678 | 0.066485 | 0.0966662 | 0.089246 | 0.0666234 | 0.02111 | 0.0989788 | 0.0203257 | 0.0647768 | 0.0191417 | 0.0939396 | 0.0238155 | 0.1089875 | 0.0583234 | 0.1235715 | 0.1123916 | 0.0275702 | 0.0737083 | 0.1010667 |
| 31.07 | 0.043918 | 0.0318089 | 0.0542058 | 0.0387396 | 0.0247203 | 0.0041317 | 0.0598082 | 0.0036213 | 0.037701 | 0.0098443 | 0.0461354 | 0.0222729 | 0.0509433 | 0.0381757 | 0.0536739 | 0.0615024 | 0.027524 | 0.0338583 | 0.0405975 |
| 31.08 | 0.0534512 | 0.0368484 | 0.0559225 | 0.033986 | 0.0119837 | 0.0062266 | 0.0508399 | 0.0095187 | 0.041261 | 0.0089855 | 0.0280128 | 0.0166531 | 0.0273348 | 0.0262275 | 0.0246071 | 0.0353765 | 0.0283671 | 0.016992 | 0.0183308 |
| 31.1 | 0.0488285 | 0.0356391 | 0.0425048 | 0.0246869 | 0.0047628 | 0.0093927 | 0.032564 | 0.0146167 | 0.0283288 | 0.0126113 | 0.0129968 | 0.0089425 | 0.0132755 | 0.014218 | 0.0089009 | 0.0152075 | 0.0170651 | 0.0151966 | 0.0137619 |
| 31.11 | 0.0293268 | 0.0249026 | 0.0216244 | 0.0134258 | 0.0049039 | 0.0177021 | 0.0452498 | 0.021887 | 0.0100418 | 0.0288175 | 0.0037941 | 0.0135731 | 0.0149716 | 0.0130653 | 0.0078154 | 0.0122032 | 0.0056876 | 0.020731 | 0.0156333 |
| 31.13 | 0.0208907 | 0.0181133 | 0.0208675 | 0.0149951 | 0.0076047 | 0.0235784 | 0.0716087 | 0.0243163 | 0.0128291 | 0.0333497 | 0.0043121 | 0.021461 | 0.0170266 | 0.0143274 | 0.0118287 | 0.0170863 | 0.0090558 | 0.0200807 | 0.0134386 |
| 31.14 | 0.0189144 | 0.0165414 | 0.0231761 | 0.0193232 | 0.0079172 | 0.0215839 | 0.0729566 | 0.0163141 | 0.0207364 | 0.021058 | 0.0090363 | 0.0241231 | 0.0109387 | 0.0103931 | 0.0134664 | 0.0191435 | 0.0151986 | 0.0149337 | 0.0086329 |
| 31.15 | 0.0402904 | 0.0334196 | 0.0379747 | 0.0336174 | 0.0138284 | 0.0272976 | 0.1086344 | 0.0149343 | 0.0371615 | 0.0246676 | 0.0250766 | 0.0430067 | 0.02016 | 0.0215362 | 0.0291461 | 0.0449923 | 0.0342148 | 0.025103 | 0.0150665 |
| 31.17 | 0.0393556 | 0.0328917 | 0.031707 | 0.0292034 | 0.0133022 | 0.0194808 | 0.0955768 | 0.0085665 | 0.0300327 | 0.0202976 | 0.0254727 | 0.0401215 | 0.0246292 | 0.0253586 | 0.0334687 | 0.0523008 | 0.0360982 | 0.0228116 | 0.0135415 |
| 31.18 | 0.023392 | 0.0243107 | 0.0101343 | 0.0129074 | 0.006397 | 0.005074 | 0.0411029 | 0.0023743 | 0.0079715 | 0.0075345 | 0.0101218 | 0.017484 | 0.0213702 | 0.019912 | 0.0235514 | 0.0345587 | 0.0195661 | 0.0092271 | 0.0050378 |
| 31.19 | 0.0316978 | 0.0434697 | 0.0048532 | 0.013634 | 0.0044207 | 0.0040732 | 0.0298232 | 0.0041599 | 0.0072919 | 0.0074475 | 0.0057441 | 0.010479 | 0.0300305 | 0.0272036 | 0.0207292 | 0.0275386 | 0.0146603 | 0.0061612 | 0.0070794 |
| 31.21 | 0.0277065 | 0.04196 | 0.0053908 | 0.0119698 | 0.0026807 | 0.0043344 | 0.0181989 | 0.003605 | 0.0107187 | 0.0088991 | 0.0046309 | 0.0047033 | 0.0239376 | 0.0211336 | 0.0130709 | 0.0174569 | 0.0074959 | 0.0033805 | 0.0079598 |
| 31.22 | 0.026051 | 0.0382979 | 0.0184013 | 0.0124846 | 0.0022425 | 0.004642 | 0.0110492 | 0.0012956 | 0.0109848 | 0.0072977 | 0.0052062 | 0.0062231 | 0.0100246 | 0.0065246 | 0.0190772 | 0.0294824 | 0.0010469 | 0.0011709 | 0.0053659 |
| 31.24 | 0.0332644 | 0.0426911 | 0.0281137 | 0.0147257 | 0.0031371 | 0.0051267 | 0.0142297 | 0.0008769 | 0.0084942 | 0.004647 | 0.0042321 | 0.0119797 | 0.0059784 | 0.003195 | 0.0245452 | 0.0368126 | 0.0004012 | 0.0014732 | 0.0031321 |
| 31.25 | 0.034995 | 0.0336589 | 0.0277962 | 0.012405 | 0.0046407 | 0.0060279 | 0.0162344 | 0.0008183 | 0.0097595 | 0.0022009 | 0.0015907 | 0.0138277 | 0.0019281 | 0.0027884 | 0.0139679 | 0.0203725 | 0.0006319 | 0.0019331 | 0.0008212 |
| 31.26 | 0.0658455 | 0.0448236 | 0.0416271 | 0.0174558 | 0.0095879 | 0.0117667 | 0.0293873 | 0.0024266 | 0.0225138 | 0.002958 | 0.0024887 | 0.019315 | 0.0021434 | 0.0039236 | 0.0122935 | 0.0152885 | 0.0018428 | 0.0043115 | 0.0006584 |
| 31.28 | 0.0690849 | 0.0382426 | 0.0360144 | 0.0148068 | 0.0094316 | 0.0121401 | 0.0277272 | 0.0028467 | 0.0215473 | 0.0035952 | 0.0032407 | 0.0142827 | 0.003325 | 0.0027663 | 0.0126917 | 0.0106882 | 0.0020253 | 0.0049456 | 0.0003334 |
| 31.29 | 0.0688789 | 0.0282371 | 0.0216027 | 0.0090446 | 0.0084263 | 0.0153032 | 0.0204034 | 0.0028889 | 0.0150106 | 0.0062149 | 0.0060136 | 0.0063739 | 0.0079416 | 0.0031683 | 0.0123068 | 0.0110263 | 0.0034837 | 0.0054365 | 0.0002698 |
| 31.31 | 0.065153 | 0.027467 | 0.0175556 | 0.0071105 | 0.0083363 | 0.0187417 | 0.018596 | 0.0045022 | 0.0133308 | 0.0066956 | 0.0072564 | 0.0074962 | 0.009219 | 0.0053857 | 0.0114082 | 0.0165105 | 0.0052776 | 0.0051756 | 0.0003325 |
| 31.32 | 0.0330908 | 0.018911 | 0.0112689 | 0.0027205 | 0.0049659 | 0.0158342 | 0.0116625 | 0.006735 | 0.0079644 | 0.0036083 | 0.0038528 | 0.0064249 | 0.0050922 | 0.0075726 | 0.0127658 | 0.0206987 | 0.0046301 | 0.002481 | 0.0004969 |
| 31.33 | 0.0229029 | 0.0193681 | 0.0194013 | 0.0022796 | 0.0042894 | 0.0215221 | 0.0128635 | 0.0120378 | 0.0066738 | 0.004656 | 0.0039721 | 0.0058197 | 0.0038837 | 0.0119238 | 0.0259929 | 0.0346912 | 0.0046484 | 0.0016153 | 0.0027105 |
| 31.35 | 0.0128231 | 0.0132309 | 0.0207464 | 0.0034381 | 0.0026154 | 0.0171491 | 0.0085275 | 0.0101928 | 0.0039159 | 0.0054569 | 0.0063998 | 0.0054261 | 0.003 | 0.0087684 | 0.0241802 | 0.0296102 | 0.0045063 | 0.0007561 | 0.0035596 |
| 31.36 | 0.0188755 | 0.0152513 | 0.0224774 | 0.0070418 | 0.0023295 | 0.006316 | 0.002623 | 0.0041565 | 0.0041944 | 0.0060007 | 0.0130319 | 0.0097354 | 0.003703 | 0.0043097 | 0.0110506 | 0.0133355 | 0.0081782 | 0.0005568 | 0.0026734 |
| 31.38 | 0.0245205 | 0.0199483 | 0.0231762 | 0.0075076 | 0.0034231 | 0.0026895 | 0.0032656 | 0.0026046 | 0.004208 | 0.0052231 | 0.0142325 | 0.0104027 | 0.0040412 | 0.0053591 | 0.008106 | 0.0109637 | 0.0086428 | 0.0008304 | 0.0020343 |
| 31.39 | 0.0168302 | 0.0142119 | 0.0163315 | 0.0045897 | 0.0035354 | 0.0008884 | 0.0047815 | 0.0030171 | 0.0018324 | 0.0031987 | 0.0093887 | 0.0049824 | 0.0028769 | 0.0055442 | 0.0107868 | 0.0131355 | 0.0040823 | 0.0011552 | 0.0009909 |
| 31.4 | 0.0209296 | 0.013089 | 0.0255037 | 0.0059532 | 0.0064174 | 0.0037971 | 0.0086565 | 0.0108254 | 0.0059206 | 0.0074035 | 0.0134501 | 0.0046343 | 0.003973 | 0.0087225 | 0.0213923 | 0.0242594 | 0.0051227 | 0.003685 | 0.0010021 |
| 31.42 | 0.0189145 | 0.0078908 | 0.0253914 | 0.0056357 | 0.0069212 | 0.0060803 | 0.0073671 | 0.0143017 | 0.0084526 | 0.0092367 | 0.0142533 | 0.0048861 | 0.0034698 | 0.0087902 | 0.0169572 | 0.019279 | 0.0065448 | 0.0042815 | 0.0014797 |
| 31.43 | 0.0117752 | 0.0023707 | 0.0144792 | 0.0035012 | 0.005081 | 0.006058 | 0.0022272 | 0.0109559 | 0.0054962 | 0.0072811 | 0.0124638 | 0.0060721 | 0.0020521 | 0.0069024 | 0.0049778 | 0.0055247 | 0.00727 | 0.0024546 | 0.0021322 |
| 31.44 | 0.016511 | 0.0062125 | 0.0154827 | 0.0038024 | 0.0083757 | 0.007178 | 0.0020927 | 0.0121964 | 0.0041288 | 0.0098401 | 0.0231241 | 0.0149254 | 0.0036866 | 0.0134353 | 0.0122806 | 0.0106616 | 0.0136185 | 0.0023066 | 0.0035013 |
| 31.46 | 0.0128579 | 0.0070488 | 0.0146593 | 0.0043017 | 0.0090548 | 0.006233 | 0.0039908 | 0.0099887 | 0.0030169 | 0.0090378 | 0.0231211 | 0.0169124 | 0.0037231 | 0.0151758 | 0.0151598 | 0.0127301 | 0.0141679 | 0.0016191 | 0.0026714 |
| 31.47 | 0.0078144 | 0.0081146 | 0.0188688 | 0.0112114 | 0.0124207 | 0.0101428 | 0.012848 | 0.0115705 | 0.0044924 | 0.0100879 | 0.0181158 | 0.0161756 | 0.003315 | 0.0193738 | 0.0097819 | 0.0092999 | 0.0159073 | 0.0009551 | 0.0008565 |
| 31.49 | 0.0192366 | 0.018071 | 0.020503 | 0.0164873 | 0.0157181 | 0.0166984 | 0.0221352 | 0.0155607 | 0.0083056 | 0.0113054 | 0.0161881 | 0.0140052 | 0.0041465 | 0.0222611 | 0.0086858 | 0.0103529 | 0.017427 | 0.0011494 | 0.0013432 |
| 31.5 | 0.0488679 | 0.0413955 | 0.0157852 | 0.0214379 | 0.0146005 | 0.0271254 | 0.0416292 | 0.0183122 | 0.0135789 | 0.0070511 | 0.0111531 | 0.0059665 | 0.0052981 | 0.0183416 | 0.0115207 | 0.0140483 | 0.0121174 | 0.0018433 | 0.0028618 |
| 31.51 | 0.1393843 | 0.1151443 | 0.0427783 | 0.0521061 | 0.0218302 | 0.0698285 | 0.1292142 | 0.0339622 | 0.0347996 | 0.0095414 | 0.0221054 | 0.0064184 | 0.0203129 | 0.0314373 | 0.030543 | 0.0274044 | 0.0128105 | 0.0089294 | 0.0123644 |
| 31.53 | 0.1542878 | 0.1212917 | 0.0654472 | 0.057804 | 0.018664 | 0.0721032 | 0.1468594 | 0.0293323 | 0.0415839 | 0.0147739 | 0.0307328 | 0.0087698 | 0.0383285 | 0.0332981 | 0.0430881 | 0.0278296 | 0.0092741 | 0.019082 | 0.0236809 |
| 31.54 | 0.1302144 | 0.0714781 | 0.1043236 | 0.0661294 | 0.0201736 | 0.0431913 | 0.120712 | 0.0110084 | 0.0460241 | 0.0238356 | 0.0683144 | 0.0213554 | 0.0859937 | 0.0433373 | 0.0719138 | 0.0295488 | 0.0081679 | 0.0452813 | 0.0463002 |
| 31.56 | 0.1162312 | 0.0456437 | 0.1037589 | 0.0738863 | 0.0281201 | 0.0286275 | 0.1069587 | 0.005369 | 0.0447191 | 0.0208393 | 0.0878135 | 0.02798 | 0.093127 | 0.0523541 | 0.0705991 | 0.0309902 | 0.0105338 | 0.0464548 | 0.0445529 |
| 31.57 | 0.0714769 | 0.0175459 | 0.0549791 | 0.0534166 | 0.0257831 | 0.0083603 | 0.058779 | 0.0023621 | 0.0266187 | 0.0132119 | 0.0606931 | 0.0281923 | 0.0448903 | 0.0420412 | 0.027481 | 0.0221062 | 0.013136 | 0.017696 | 0.0199563 |
| 31.58 | 0.094033 | 0.0151301 | 0.0596856 | 0.0634929 | 0.0343935 | 0.0036239 | 0.0487981 | 0.0033193 | 0.0292819 | 0.0277987 | 0.0545235 | 0.0660948 | 0.0347481 | 0.0585936 | 0.0155338 | 0.0340745 | 0.0305549 | 0.0089977 | 0.0286998 |
| 31.6 | 0.0785135 | 0.0090107 | 0.0482733 | 0.0473933 | 0.0329945 | 0.0031744 | 0.0277523 | 0.0044839 | 0.0265514 | 0.0295554 | 0.0330764 | 0.0711235 | 0.0256134 | 0.0508471 | 0.008195 | 0.0320834 | 0.030903 | 0.0070023 | 0.0305455 |
| 31.61 | 0.0413659 | 0.0070397 | 0.0317804 | 0.0161496 | 0.0463784 | 0.0087918 | 0.0158812 | 0.0127174 | 0.0341506 | 0.0225735 | 0.0117825 | 0.0414449 | 0.0181915 | 0.0253843 | 0.0022602 | 0.0193618 | 0.0169677 | 0.0124139 | 0.021811 |
| 31.63 | 0.0264866 | 0.0085927 | 0.0242284 | 0.0070705 | 0.0554145 | 0.0108365 | 0.0240905 | 0.0171836 | 0.034026 | 0.016244 | 0.0147657 | 0.0266124 | 0.014127 | 0.0154694 | 0.002626 | 0.0126874 | 0.0106312 | 0.0143338 | 0.0144418 |
| 31.64 | 0.011511 | 0.006272 | 0.006551 | 0.0020575 | 0.0381729 | 0.0064752 | 0.0264949 | 0.0141053 | 0.0126559 | 0.0083138 | 0.0209997 | 0.0215621 | 0.0049079 | 0.0116773 | 0.0051765 | 0.0074987 | 0.0069421 | 0.009316 | 0.0077606 |
| 31.65 | 0.0321005 | 0.0076729 | 0.0040953 | 0.0054228 | 0.046458 | 0.005896 | 0.0396191 | 0.0164824 | 0.0137317 | 0.0147786 | 0.0441905 | 0.0451914 | 0.0038306 | 0.0285562 | 0.0135679 | 0.0207586 | 0.013043 | 0.0101404 | 0.01857 |
| 31.67 | 0.0410463 | 0.0075203 | 0.0046176 | 0.0082497 | 0.041767 | 0.0043535 | 0.0371073 | 0.0122264 | 0.0260259 | 0.0119982 | 0.0487376 | 0.0427258 | 0.0026781 | 0.0293541 | 0.0153049 | 0.0253021 | 0.0130777 | 0.0074225 | 0.021045 |
| 31.68 | 0.0296016 | 0.0085711 | 0.0047362 | 0.0084734 | 0.0223012 | 0.0036005 | 0.0243175 | 0.0081793 | 0.0357908 | 0.0028359 | 0.0356013 | 0.020535 | 0.0012756 | 0.0166281 | 0.0089103 | 0.016804 | 0.0103116 | 0.0021969 | 0.0144696 |
| 31.69 | 0.0315244 | 0.0190813 | 0.0102626 | 0.0080064 | 0.0184694 | 0.0122571 | 0.0387373 | 0.02319 | 0.0556783 | 0.0032102 | 0.0507277 | 0.0206492 | 0.0048346 | 0.0251397 | 0.0064092 | 0.015534 | 0.0193382 | 0.0012825 | 0.0200942 |
| 31.71 | 0.0208529 | 0.0187492 | 0.0102073 | 0.0055268 | 0.0127842 | 0.0143112 | 0.0373651 | 0.0280476 | 0.0478929 | 0.0033071 | 0.0444263 | 0.0163807 | 0.0074921 | 0.0267971 | 0.0041299 | 0.0095403 | 0.0202257 | 0.0010759 | 0.0181161 |
| 31.72 | 0.0161786 | 0.0151129 | 0.0046257 | 0.0110643 | 0.0255256 | 0.011669 | 0.0264612 | 0.026152 | 0.0254831 | 0.0019574 | 0.023514 | 0.0108375 | 0.0108478 | 0.0279144 | 0.0045207 | 0.0030769 | 0.0165385 | 0.000932 | 0.0121776 |
| 31.74 | 0.0261269 | 0.01574 | 0.0023213 | 0.0159463 | 0.0407855 | 0.0107145 | 0.0189924 | 0.0199739 | 0.0182382 | 0.0023044 | 0.0156545 | 0.0079927 | 0.0104085 | 0.0259593 | 0.004497 | 0.0032671 | 0.0121817 | 0.0007922 | 0.0088792 |
| 31.75 | 0.0305415 | 0.0133693 | 0.001773 | 0.0139471 | 0.0414637 | 0.0083504 | 0.0056933 | 0.0053936 | 0.016755 | 0.0030593 | 0.0132206 | 0.0030924 | 0.0046304 | 0.0110677 | 0.0046162 | 0.004608 | 0.0055512 | 0.00057 | 0.0028612 |
| 31.76 | 0.057088 | 0.0228046 | 0.0070366 | 0.0202514 | 0.0639933 | 0.0207683 | 0.008435 | 0.0066882 | 0.0506433 | 0.0074239 | 0.0379629 | 0.0080571 | 0.0030048 | 0.0088989 | 0.0135321 | 0.0093307 | 0.0138222 | 0.0008752 | 0.0046192 |
| 31.78 | 0.0627049 | 0.0210104 | 0.0108566 | 0.0171915 | 0.0590198 | 0.028049 | 0.0103792 | 0.0128591 | 0.0611533 | 0.0100085 | 0.0433936 | 0.0103607 | 0.0017905 | 0.0114073 | 0.0145714 | 0.0081386 | 0.0151766 | 0.0013337 | 0.0070754 |
| 31.79 | 0.0599688 | 0.0121847 | 0.0180264 | 0.0057221 | 0.0488509 | 0.0368471 | 0.0143356 | 0.0276208 | 0.0588466 | 0.0155759 | 0.0358202 | 0.0119825 | 0.0025406 | 0.0221103 | 0.009468 | 0.0033717 | 0.0093938 | 0.0036672 | 0.0117485 |
| 31.81 | 0.0453649 | 0.0114303 | 0.0167008 | 0.0034997 | 0.0465149 | 0.0365628 | 0.0152403 | 0.028855 | 0.0476544 | 0.0159284 | 0.0257367 | 0.0118743 | 0.0036197 | 0.0235693 | 0.0064247 | 0.002889 | 0.0085636 | 0.0045861 | 0.0112981 |
| 31.82 | 0.0118509 | 0.0105209 | 0.0051689 | 0.0071637 | 0.0303929 | 0.025895 | 0.0085209 | 0.0178416 | 0.0165218 | 0.00767 | 0.0070786 | 0.0066604 | 0.0037895 | 0.0125274 | 0.0016864 | 0.00257 | 0.0102866 | 0.0031748 | 0.0050414 |
| 31.83 | 0.0233708 | 0.0157699 | 0.0081366 | 0.0227247 | 0.0460675 | 0.0507977 | 0.0196548 | 0.0280387 | 0.0247986 | 0.005057 | 0.0131214 | 0.0058409 | 0.0076731 | 0.0106842 | 0.0009355 | 0.0025953 | 0.020039 | 0.0039126 | 0.0107655 |
| 31.85 | 0.0372492 | 0.0196859 | 0.0120788 | 0.0270915 | 0.0447489 | 0.0568062 | 0.0289814 | 0.0269677 | 0.0333341 | 0.0024896 | 0.018859 | 0.0056955 | 0.0080964 | 0.0110459 | 0.0011765 | 0.0017153 | 0.0182213 | 0.0034632 | 0.0158338 |
| 31.86 | 0.0609065 | 0.0414132 | 0.019839 | 0.0269364 | 0.0387925 | 0.0560236 | 0.0437348 | 0.0206753 | 0.0490089 | 0.0032256 | 0.0312261 | 0.0158104 | 0.0057289 | 0.0258102 | 0.0023359 | 0.0017288 | 0.010729 | 0.0029179 | 0.0204399 |
| 31.88 | 0.0668965 | 0.0502048 | 0.0255451 | 0.0260757 | 0.0366635 | 0.0512383 | 0.0487373 | 0.0191224 | 0.0551495 | 0.0073731 | 0.0370899 | 0.0224167 | 0.0041027 | 0.0341143 | 0.0027354 | 0.0022604 | 0.0089467 | 0.0025835 | 0.0172934 |
| 31.89 | 0.0477448 | 0.0369051 | 0.0271887 | 0.0182295 | 0.0228421 | 0.0310629 | 0.0409791 | 0.0140089 | 0.0423512 | 0.0114584 | 0.0287527 | 0.0180897 | 0.0031812 | 0.0277664 | 0.002245 | 0.00205 | 0.0071912 | 0.0010803 | 0.0054229 |
| 31.9 | 0.0702173 | 0.0550245 | 0.0575788 | 0.0289064 | 0.0298481 | 0.0504656 | 0.0760059 | 0.0266806 | 0.0641817 | 0.0215764 | 0.0397384 | 0.0204524 | 0.0090438 | 0.0377479 | 0.0036213 | 0.0027145 | 0.0131195 | 0.0007554 | 0.0033709 |
| 31.92 | 0.0668781 | 0.058093 | 0.0625856 | 0.0287263 | 0.026068 | 0.0513414 | 0.0766697 | 0.0306498 | 0.0596489 | 0.0200189 | 0.0343419 | 0.0158089 | 0.0111996 | 0.0321824 | 0.0037629 | 0.0020309 | 0.0122762 | 0.0007228 | 0.0037051 |
| 31.93 | 0.0361926 | 0.0427515 | 0.0369984 | 0.016875 | 0.0119156 | 0.0289526 | 0.0406547 | 0.0239197 | 0.0322825 | 0.0079854 | 0.0154142 | 0.0070948 | 0.0092375 | 0.0135578 | 0.0035114 | 0.0010739 | 0.0050637 | 0.0009535 | 0.0030245 |
| 31.94 | 0.0273863 | 0.0472024 | 0.033099 | 0.0145052 | 0.0089742 | 0.0280076 | 0.0386144 | 0.0319287 | 0.0388127 | 0.0048457 | 0.0159049 | 0.0061396 | 0.0155324 | 0.0084597 | 0.0084289 | 0.0028173 | 0.003634 | 0.002775 | 0.0020713 |
| 31.96 | 0.0157736 | 0.0316519 | 0.0204569 | 0.0098912 | 0.0053903 | 0.0189401 | 0.025967 | 0.0253428 | 0.02916 | 0.0036959 | 0.0147936 | 0.0036241 | 0.0158939 | 0.0039683 | 0.0099658 | 0.0034027 | 0.0039268 | 0.0038893 | 0.0011935 |
| 31.97 | 0.0184689 | 0.0122938 | 0.0112559 | 0.0178322 | 0.0044586 | 0.0045843 | 0.0109012 | 0.0138155 | 0.0134024 | 0.0042749 | 0.0154752 | 0.0006143 | 0.0146299 | 0.0015037 | 0.0106972 | 0.0026113 | 0.0092534 | 0.0065914 | 0.0013764 |
| 31.99 | 0.0229256 | 0.0088577 | 0.0176093 | 0.0223648 | 0.005843 | 0.0042328 | 0.0128808 | 0.0170805 | 0.0177924 | 0.0054813 | 0.0149972 | 0.0001912 | 0.0125368 | 0.0014031 | 0.0094188 | 0.0014913 | 0.0118557 | 0.0078656 | 0.0016377 |
| 32 | 0.0337128 | 0.0122461 | 0.0167631 | 0.019998 | 0.0119398 | 0.0159625 | 0.0111603 | 0.0325365 | 0.0163825 | 0.0059904 | 0.0085731 | 0.0006614 | 0.0052064 | 0.0026384 | 0.0034488 | 0.0003766 | 0.0096442 | 0.0057216 | 0.0019896 |
| 32.01 | 0.1321196 | 0.0750377 | 0.0319513 | 0.0737202 | 0.0473146 | 0.0657121 | 0.0275 | 0.0990079 | 0.0311733 | 0.0066928 | 0.0215327 | 0.0057623 | 0.0130632 | 0.0118752 | 0.0080321 | 0.0070752 | 0.016904 | 0.0106499 | 0.0067918 |
| 32.03 | 0.167456 | 0.1118734 | 0.0513961 | 0.105148 | 0.0604316 | 0.0873005 | 0.0490662 | 0.1099767 | 0.0539811 | 0.0042755 | 0.038119 | 0.0075485 | 0.0272795 | 0.016492 | 0.0204007 | 0.0237842 | 0.0174524 | 0.0212402 | 0.0160231 |
| 32.04 | 0.1554482 | 0.134285 | 0.0830666 | 0.1215989 | 0.0647354 | 0.1017817 | 0.084255 | 0.0714686 | 0.0966484 | 0.0138532 | 0.0751664 | 0.0156799 | 0.0678213 | 0.0454952 | 0.0545421 | 0.0846521 | 0.0213122 | 0.0665679 | 0.0561519 |
| 32.06 | 0.1256656 | 0.124178 | 0.0823054 | 0.103762 | 0.0586514 | 0.0956675 | 0.082329 | 0.0451004 | 0.0958038 | 0.025326 | 0.0760498 | 0.0254145 | 0.0734394 | 0.0725957 | 0.0570776 | 0.0994554 | 0.0302225 | 0.0787001 | 0.0674959 |
| 32.07 | 0.0537834 | 0.0718732 | 0.0514174 | 0.0422989 | 0.0297919 | 0.051637 | 0.0443909 | 0.0130536 | 0.0503274 | 0.0230201 | 0.0313725 | 0.0208652 | 0.0312089 | 0.061363 | 0.023555 | 0.0457511 | 0.0288737 | 0.0364464 | 0.0327026 |
| 32.08 | 0.0509744 | 0.0941316 | 0.0668853 | 0.0324519 | 0.0345036 | 0.0538811 | 0.0432486 | 0.0099925 | 0.042059 | 0.0184869 | 0.0143456 | 0.0162165 | 0.0136917 | 0.0494438 | 0.0152338 | 0.0229686 | 0.0279486 | 0.0181932 | 0.0171403 |
| 32.1 | 0.0332296 | 0.0773757 | 0.0531453 | 0.0187183 | 0.0269231 | 0.0362482 | 0.0268153 | 0.0062672 | 0.0250711 | 0.0109038 | 0.0050383 | 0.0091884 | 0.0061193 | 0.026998 | 0.0097144 | 0.0148187 | 0.0161189 | 0.0122005 | 0.0083054 |
| 32.11 | 0.011491 | 0.0385626 | 0.0199136 | 0.0065832 | 0.0103647 | 0.0155607 | 0.0100811 | 0.0038413 | 0.0198619 | 0.0180473 | 0.0034318 | 0.0045107 | 0.0084725 | 0.024309 | 0.0043667 | 0.0201934 | 0.0115928 | 0.0166968 | 0.011004 |
| 32.13 | 0.0126588 | 0.0264845 | 0.0102158 | 0.0077369 | 0.0079816 | 0.0226379 | 0.0142784 | 0.0036426 | 0.0269239 | 0.0265643 | 0.0041992 | 0.0072817 | 0.0117749 | 0.0336806 | 0.0045163 | 0.0211936 | 0.0169015 | 0.01644 | 0.0154997 |
| 32.14 | 0.0155844 | 0.0091408 | 0.0066088 | 0.0084948 | 0.0092468 | 0.0300129 | 0.0171247 | 0.002756 | 0.0252105 | 0.0251847 | 0.0020083 | 0.0147392 | 0.0115407 | 0.0293069 | 0.0058946 | 0.012853 | 0.0174598 | 0.0086901 | 0.0136318 |
| 32.15 | 0.0389663 | 0.0114653 | 0.0164273 | 0.0161581 | 0.0216198 | 0.0580786 | 0.0283285 | 0.0092057 | 0.0433556 | 0.0422087 | 0.0031251 | 0.0378268 | 0.0171664 | 0.0403991 | 0.0112522 | 0.0141783 | 0.0269826 | 0.0101343 | 0.0216156 |
| 32.17 | 0.0484048 | 0.0179733 | 0.0181558 | 0.0162698 | 0.0246327 | 0.0629838 | 0.0288768 | 0.0135998 | 0.0469041 | 0.0414544 | 0.0044286 | 0.0426559 | 0.0135905 | 0.0347075 | 0.0107147 | 0.0098619 | 0.023875 | 0.0080691 | 0.0206045 |
| 32.18 | 0.0389344 | 0.0211947 | 0.0139386 | 0.0090491 | 0.0178298 | 0.0441139 | 0.0209148 | 0.0143821 | 0.0378937 | 0.0222471 | 0.0036686 | 0.0304335 | 0.0047138 | 0.0159387 | 0.0054906 | 0.002434 | 0.0121445 | 0.0025158 | 0.010333 |
| 32.19 | 0.0622945 | 0.0400238 | 0.0260896 | 0.0152739 | 0.0260023 | 0.059171 | 0.033659 | 0.0302321 | 0.0733379 | 0.0194659 | 0.0048557 | 0.0433821 | 0.0077665 | 0.018839 | 0.0068 | 0.0041484 | 0.0207989 | 0.0014005 | 0.0085145 |
| 32.21 | 0.0642282 | 0.0452179 | 0.0275092 | 0.0189578 | 0.0243683 | 0.052107 | 0.0336886 | 0.0347268 | 0.081731 | 0.0117973 | 0.0045921 | 0.0387083 | 0.0109438 | 0.0204415 | 0.0067579 | 0.0070498 | 0.026001 | 0.0019371 | 0.0055162 |
| 32.22 | 0.0613055 | 0.0511846 | 0.0246177 | 0.0244926 | 0.0182599 | 0.0382847 | 0.0342163 | 0.0349565 | 0.0864813 | 0.0025519 | 0.0035058 | 0.0219538 | 0.0183739 | 0.0321692 | 0.0048448 | 0.0121933 | 0.0343568 | 0.0043956 | 0.0029828 |
| 32.24 | 0.0564493 | 0.0508109 | 0.0221756 | 0.0242901 | 0.0156982 | 0.0336632 | 0.0373637 | 0.0308495 | 0.086042 | 0.0020837 | 0.0027889 | 0.0136295 | 0.0204209 | 0.0375412 | 0.0034634 | 0.011245 | 0.0321454 | 0.0041538 | 0.0026386 |
| 32.25 | 0.0328259 | 0.0344912 | 0.0131946 | 0.0133984 | 0.0094857 | 0.0193185 | 0.0308999 | 0.0160995 | 0.0560417 | 0.0038322 | 0.0018877 | 0.0051038 | 0.013314 | 0.0253712 | 0.002313 | 0.0035139 | 0.0127657 | 0.0012714 | 0.001328 |
| 32.26 | 0.041045 | 0.0540235 | 0.0212185 | 0.011174 | 0.0137443 | 0.0231229 | 0.0531007 | 0.0194069 | 0.073105 | 0.0099738 | 0.0028744 | 0.0089789 | 0.0163343 | 0.0249136 | 0.0028028 | 0.0013747 | 0.0067114 | 0.0018482 | 0.0022154 |
| 32.28 | 0.0348536 | 0.05367 | 0.0242752 | 0.0061328 | 0.0130317 | 0.0180161 | 0.0560294 | 0.0156865 | 0.063759 | 0.010458 | 0.0019537 | 0.0086121 | 0.0136315 | 0.0159125 | 0.001847 | 0.0009141 | 0.0038611 | 0.0025398 | 0.0033576 |
| 32.29 | 0.0269854 | 0.0511593 | 0.0345437 | 0.0020675 | 0.0129112 | 0.0093853 | 0.059232 | 0.0088976 | 0.0462247 | 0.007699 | 0.0012844 | 0.0065306 | 0.0064977 | 0.0090626 | 0.0021327 | 0.0021013 | 0.0048333 | 0.0023698 | 0.0059896 |
| 32.31 | 0.0273984 | 0.0487593 | 0.0393612 | 0.0026637 | 0.0143383 | 0.0093403 | 0.0580413 | 0.0079146 | 0.0392797 | 0.0064837 | 0.0022503 | 0.0057962 | 0.0033368 | 0.0094006 | 0.0034939 | 0.0029049 | 0.0046014 | 0.0018462 | 0.0060138 |
| 32.32 | 0.0214916 | 0.0293124 | 0.0299745 | 0.0040198 | 0.0119738 | 0.0112893 | 0.0372172 | 0.0060388 | 0.019516 | 0.0032397 | 0.0019023 | 0.0027822 | 0.0011027 | 0.0057834 | 0.0032984 | 0.0019761 | 0.0031156 | 0.0011416 | 0.0030221 |

| | | | | | | | | | | | | | | | | | | | |
|---|---|---|---|---|---|---|---|---|---|---|---|---|---|---|---|---|---|---|---|
| 32.33 | 0.0340745 | 0.0418165 | 0.0458309 | 0.0074436 | 0.0191953 | 0.0253979 | 0.056672 | 0.0093798 | 0.0210962 | 0.0062692 | 0.0030248 | 0.0049338 | 0.0042381 | 0.0145431 | 0.0027095 | 0.0036026 | 0.0115288 | 0.00332 | 0.0067392 |
| 32.35 | 0.0317173 | 0.0410684 | 0.0424855 | 0.0058091 | 0.017639 | 0.0262514 | 0.0543153 | 0.0082889 | 0.0169954 | 0.009628 | 0.0051516 | 0.0068979 | 0.0060608 | 0.0202851 | 0.0014233 | 0.0054583 | 0.0148951 | 0.0045553 | 0.0092513 |
| 32.36 | 0.0280251 | 0.0440371 | 0.0311121 | 0.0021089 | 0.0122355 | 0.0204722 | 0.0399695 | 0.0070791 | 0.0131279 | 0.0145605 | 0.0094651 | 0.0102569 | 0.0064972 | 0.023361 | 0.0009013 | 0.0076462 | 0.0132602 | 0.0049037 | 0.0094353 |
| 32.38 | 0.0317253 | 0.0490299 | 0.0281684 | 0.0027585 | 0.0106171 | 0.0173177 | 0.0308818 | 0.0090439 | 0.0145332 | 0.0140188 | 0.0094645 | 0.012219 | 0.0052947 | 0.019638 | 0.0009713 | 0.0065183 | 0.0097126 | 0.0038975 | 0.0064957 |
| 32.39 | 0.0287237 | 0.0370321 | 0.0203441 | 0.0048313 | 0.0069523 | 0.0101458 | 0.011272 | 0.0091455 | 0.0118857 | 0.0078226 | 0.0062518 | 0.0137877 | 0.0034531 | 0.0077109 | 0.0011743 | 0.0034254 | 0.0035614 | 0.0022418 | 0.0010566 |
| 32.4 | 0.0476991 | 0.0502598 | 0.0365378 | 0.0113449 | 0.0118112 | 0.0180311 | 0.0089226 | 0.0123907 | 0.016281 | 0.0112377 | 0.0134868 | 0.0331936 | 0.0094163 | 0.0088321 | 0.0049404 | 0.0096185 | 0.0087878 | 0.0067423 | 0.0017894 |
| 32.42 | 0.0475639 | 0.0465476 | 0.0384135 | 0.0125524 | 0.0126122 | 0.0196119 | 0.0060221 | 0.0100585 | 0.0154821 | 0.0106996 | 0.0159388 | 0.0364469 | 0.0110611 | 0.0094124 | 0.0062956 | 0.012287 | 0.0123811 | 0.0079285 | 0.0028199 |
| 32.43 | 0.0317265 | 0.0322182 | 0.0225178 | 0.0078555 | 0.0089828 | 0.0112698 | 0.0029839 | 0.0060138 | 0.0119251 | 0.0044104 | 0.0110314 | 0.0201245 | 0.0060506 | 0.0074297 | 0.0030705 | 0.00752 | 0.0104767 | 0.0035384 | 0.0016853 |
| 32.44 | 0.0443777 | 0.0519111 | 0.0167201 | 0.0069302 | 0.0123493 | 0.0114603 | 0.0141103 | 0.0099721 | 0.0185638 | 0.0015276 | 0.0149673 | 0.0143977 | 0.0051647 | 0.011703 | 0.0011731 | 0.0051394 | 0.0139777 | 0.0013101 | 0.0008933 |
| 32.46 | 0.0390977 | 0.0478145 | 0.0097347 | 0.0051245 | 0.0104037 | 0.0151177 | 0.0227494 | 0.0083157 | 0.0146839 | 0.0016799 | 0.015718 | 0.007513 | 0.0050012 | 0.0112556 | 0.0005665 | 0.0030421 | 0.0123087 | 0.000691 | 0.0011717 |
| 32.47 | 0.0297421 | 0.0335182 | 0.0117545 | 0.0061001 | 0.005516 | 0.0345636 | 0.0356139 | 0.0034468 | 0.00524 | 0.0093769 | 0.0193627 | 0.0039206 | 0.0080812 | 0.0089113 | 0.0005863 | 0.003277 | 0.0086392 | 0.001555 | 0.0029498 |
| 32.49 | 0.0361164 | 0.0343133 | 0.0127889 | 0.0057244 | 0.0039244 | 0.0379934 | 0.0347495 | 0.0064283 | 0.0036044 | 0.0136235 | 0.0193362 | 0.0043877 | 0.0088767 | 0.0079944 | 0.0005184 | 0.003104 | 0.0067929 | 0.001618 | 0.0028481 |
| 32.5 | 0.0582269 | 0.0481714 | 0.005089 | 0.0044089 | 0.0043303 | 0.0164525 | 0.0168063 | 0.0213974 | 0.0040797 | 0.0110981 | 0.0106658 | 0.0060821 | 0.0060543 | 0.0060276 | 0.0010221 | 0.0017208 | 0.0037167 | 0.00134 | 0.0014747 |
| 32.51 | 0.1747654 | 0.1577172 | 0.0137302 | 0.0198557 | 0.0220615 | 0.016144 | 0.0338615 | 0.0749492 | 0.0220912 | 0.0112337 | 0.0183199 | 0.0170746 | 0.0171619 | 0.0167733 | 0.0107154 | 0.0127307 | 0.0071866 | 0.012391 | 0.0078823 |
| 32.53 | 0.20033 | 0.1869989 | 0.0313331 | 0.0320207 | 0.0318798 | 0.0256794 | 0.050795 | 0.0846414 | 0.0351209 | 0.0104532 | 0.0275897 | 0.0177633 | 0.0303689 | 0.0255081 | 0.0245277 | 0.0292641 | 0.0066273 | 0.0278586 | 0.0156295 |
| 32.54 | 0.1596054 | 0.1363179 | 0.068462 | 0.0572997 | 0.0343649 | 0.0518003 | 0.0908563 | 0.0559334 | 0.0530325 | 0.0171637 | 0.057579 | 0.0087186 | 0.0662814 | 0.0486397 | 0.0584803 | 0.0708355 | 0.0056483 | 0.0625687 | 0.0270071 |
| 32.56 | 0.126469 | 0.0890114 | 0.0733503 | 0.070214 | 0.0286364 | 0.0556504 | 0.1075706 | 0.040248 | 0.0571281 | 0.0194503 | 0.0696366 | 0.0115828 | 0.0730563 | 0.0579386 | 0.0592627 | 0.0754401 | 0.0132473 | 0.064857 | 0.0222296 |
| 32.57 | 0.0627156 | 0.0270757 | 0.0491302 | 0.0637933 | 0.0153001 | 0.0346978 | 0.0828928 | 0.0225381 | 0.0405726 | 0.0191036 | 0.0519336 | 0.0268189 | 0.0368201 | 0.041652 | 0.0208213 | 0.0349111 | 0.0231476 | 0.0278725 | 0.0043099 |
| 32.58 | 0.077255 | 0.0286294 | 0.0742008 | 0.1043061 | 0.027621 | 0.0435566 | 0.1134086 | 0.0339751 | 0.0527359 | 0.0431689 | 0.0719924 | 0.0674359 | 0.0271404 | 0.053249 | 0.0110916 | 0.029898 | 0.0397254 | 0.0142463 | 0.0059308 |
| 32.6 | 0.0652346 | 0.0247682 | 0.0708765 | 0.0930276 | 0.0320028 | 0.0346342 | 0.0982077 | 0.0316537 | 0.0429592 | 0.0453122 | 0.0652356 | 0.0729441 | 0.0167132 | 0.0462618 | 0.0086826 | 0.0240256 | 0.0358087 | 0.0056927 | 0.0117903 |
| 32.61 | 0.0372822 | 0.0161415 | 0.0595381 | 0.0499271 | 0.0442141 | 0.0124484 | 0.0687967 | 0.0246406 | 0.0254162 | 0.0281471 | 0.043842 | 0.0544663 | 0.0058444 | 0.0272719 | 0.0100046 | 0.0135998 | 0.0191043 | 0.0036538 | 0.0255054 |
| 32.63 | 0.0263126 | 0.0117484 | 0.0529186 | 0.0301091 | 0.0502434 | 0.0076724 | 0.0587783 | 0.0207855 | 0.0186747 | 0.018049 | 0.0301013 | 0.0384755 | 0.0040439 | 0.0174785 | 0.00908 | 0.0091589 | 0.0133934 | 0.0057124 | 0.0304343 |
| 32.64 | 0.011382 | 0.0049295 | 0.0275467 | 0.0077205 | 0.0357804 | 0.0048343 | 0.0304303 | 0.0103757 | 0.0054182 | 0.0119871 | 0.0074055 | 0.0229391 | 0.0048699 | 0.0081309 | 0.0031018 | 0.0065819 | 0.0140683 | 0.0055235 | 0.0237661 |
| 32.65 | 0.0126739 | 0.0052087 | 0.0310574 | 0.0154702 | 0.0461133 | 0.0063269 | 0.0350342 | 0.0116558 | 0.0085241 | 0.023412 | 0.0083526 | 0.0546707 | 0.0101835 | 0.0167893 | 0.0053667 | 0.0118188 | 0.0257224 | 0.0077937 | 0.0342281 |
| 32.67 | 0.0091711 | 0.0040702 | 0.0250853 | 0.0193136 | 0.0386929 | 0.007714 | 0.0262169 | 0.0087136 | 0.0159558 | 0.0216303 | 0.0116017 | 0.0590229 | 0.0081836 | 0.0178264 | 0.0067071 | 0.0104808 | 0.0203527 | 0.0073522 | 0.0271028 |
| 32.68 | 0.0035792 | 0.0030294 | 0.0103435 | 0.0136392 | 0.0166667 | 0.0052223 | 0.0119126 | 0.0026923 | 0.0225189 | 0.009855 | 0.0127815 | 0.0348759 | 0.0024242 | 0.0121457 | 0.004385 | 0.0054434 | 0.0075065 | 0.0038019 | 0.008775 |
| 32.69 | 0.0106391 | 0.007276 | 0.0077247 | 0.0181104 | 0.0142632 | 0.0109551 | 0.0511003 | 0.0016626 | 0.05023 | 0.0068626 | 0.0269214 | 0.0413122 | 0.0035665 | 0.0228874 | 0.0055882 | 0.0083902 | 0.0155496 | 0.0031682 | 0.0191782 |
| 32.71 | 0.0147643 | 0.0081565 | 0.0053955 | 0.0182463 | 0.0095447 | 0.0234163 | 0.0776295 | 0.0011023 | 0.0579421 | 0.0040801 | 0.0311407 | 0.0351665 | 0.0035067 | 0.0276833 | 0.0047517 | 0.0083626 | 0.0224623 | 0.0031639 | 0.0286305 |
| 32.72 | 0.0189318 | 0.0084924 | 0.0081303 | 0.0218 | 0.004995 | 0.0613904 | 0.1090596 | 0.0028316 | 0.0597367 | 0.0027694 | 0.0384998 | 0.0275523 | 0.0054703 | 0.0361675 | 0.0020192 | 0.0056196 | 0.0335776 | 0.0037549 | 0.0425239 |
| 32.74 | 0.0180657 | 0.0086809 | 0.0114333 | 0.0203132 | 0.0056047 | 0.072093 | 0.1050107 | 0.006041 | 0.048398 | 0.0030123 | 0.0394627 | 0.0258029 | 0.0085361 | 0.0363344 | 0.0012315 | 0.0042865 | 0.0335372 | 0.0029876 | 0.0431411 |
| 32.75 | 0.0075776 | 0.0069825 | 0.0115212 | 0.0086766 | 0.0057411 | 0.0431751 | 0.0444797 | 0.0089309 | 0.0154669 | 0.0045726 | 0.0215067 | 0.013314 | 0.0077816 | 0.0195595 | 0.0017492 | 0.0041019 | 0.015757 | 0.0013805 | 0.0246443 |
| 32.76 | 0.0055455 | 0.0140315 | 0.0206467 | 0.0175743 | 0.0083131 | 0.0387926 | 0.0232977 | 0.014373 | 0.027104 | 0.0110164 | 0.0175366 | 0.0111271 | 0.0075546 | 0.0174445 | 0.0072238 | 0.0117984 | 0.0107052 | 0.0019981 | 0.0249355 |
| 32.78 | 0.0067011 | 0.0148994 | 0.0204547 | 0.0275832 | 0.006572 | 0.0241195 | 0.0101879 | 0.0109392 | 0.036501 | 0.0111551 | 0.0101395 | 0.0106164 | 0.0059674 | 0.0108501 | 0.0095049 | 0.0133109 | 0.0094938 | 0.0015934 | 0.0162037 |
| 32.79 | 0.0117731 | 0.0140049 | 0.0170642 | 0.0470926 | 0.0091449 | 0.0108417 | 0.0079725 | 0.006147 | 0.0499978 | 0.0106216 | 0.0039278 | 0.0183191 | 0.0083222 | 0.0056126 | 0.0103272 | 0.0097049 | 0.0177 | 0.0009351 | 0.0034001 |
| 32.81 | 0.011144 | 0.0142845 | 0.0150532 | 0.049309 | 0.0167076 | 0.0141195 | 0.0095439 | 0.0079593 | 0.055899 | 0.012372 | 0.0039334 | 0.0202151 | 0.0084738 | 0.0081997 | 0.0089611 | 0.0063239 | 0.0197077 | 0.0013471 | 0.0018011 |
| 32.82 | 0.0079369 | 0.0098199 | 0.007539 | 0.0243924 | 0.0227994 | 0.0144278 | 0.0066787 | 0.0068178 | 0.0396255 | 0.0126618 | 0.0023116 | 0.0102024 | 0.0037425 | 0.0105192 | 0.0031824 | 0.0011812 | 0.0099766 | 0.0018835 | 0.0020512 |
| 32.83 | 0.0241776 | 0.0115493 | 0.0058759 | 0.0231057 | 0.0483734 | 0.0207961 | 0.0099687 | 0.0054652 | 0.0421364 | 0.028534 | 0.0028903 | 0.0095885 | 0.0031349 | 0.0200612 | 0.0025563 | 0.0006343 | 0.0090037 | 0.0032188 | 0.0033484 |
| 32.85 | 0.0321167 | 0.0096993 | 0.0044393 | 0.0303863 | 0.0550311 | 0.0159268 | 0.0095054 | 0.0035625 | 0.0281411 | 0.0339546 | 0.0046305 | 0.0126622 | 0.0020124 | 0.0198371 | 0.0029423 | 0.0013222 | 0.0110666 | 0.0026357 | 0.0027124 |
| 32.86 | 0.0439732 | 0.0093357 | 0.0117308 | 0.0607725 | 0.0641937 | 0.0084943 | 0.0060963 | 0.0061266 | 0.0200918 | 0.0380471 | 0.0096716 | 0.0228333 | 0.001291 | 0.0138993 | 0.0033923 | 0.0051514 | 0.0195026 | 0.0012 | 0.001494 |
| 32.88 | 0.0494631 | 0.011011 | 0.0180884 | 0.0683919 | 0.0646377 | 0.0138841 | 0.0065325 | 0.0072206 | 0.0286278 | 0.0333931 | 0.0106998 | 0.022671 | 0.0027282 | 0.0095121 | 0.0039747 | 0.0073498 | 0.0192012 | 0.0013614 | 0.0009759 |
| 32.89 | 0.0392458 | 0.0110212 | 0.0180611 | 0.0454239 | 0.0382496 | 0.0197694 | 0.0133551 | 0.0046292 | 0.0308223 | 0.0115159 | 0.007794 | 0.0108384 | 0.0046975 | 0.0043101 | 0.0049964 | 0.0062508 | 0.0084279 | 0.0027065 | 0.0003545 |
| 32.9 | 0.0638285 | 0.0237383 | 0.029102 | 0.0591155 | 0.0410812 | 0.0399326 | 0.041803 | 0.0134049 | 0.0663322 | 0.0096769 | 0.0138665 | 0.0087388 | 0.0082512 | 0.0163543 | 0.0085158 | 0.0078433 | 0.0096427 | 0.0077604 | 0.001637 |
| 32.92 | 0.062432 | 0.0269014 | 0.0270131 | 0.0513947 | 0.0287066 | 0.0453953 | 0.0506735 | 0.0188425 | 0.077215 | 0.0122785 | 0.0146071 | 0.0053982 | 0.0066692 | 0.0231658 | 0.0069643 | 0.0063758 | 0.0116588 | 0.0088649 | 0.0020049 |
| 32.93 | 0.0336939 | 0.0181398 | 0.013758 | 0.0232233 | 0.0078332 | 0.0354731 | 0.035599 | 0.0155913 | 0.0573335 | 0.0124838 | 0.0086778 | 0.0026671 | 0.0017826 | 0.0187937 | 0.0031 | 0.0035317 | 0.0102241 | 0.0060807 | 0.0009978 |
| 32.94 | 0.0302389 | 0.0204698 | 0.0150556 | 0.0190862 | 0.0155965 | 0.048463 | 0.0412129 | 0.0224726 | 0.0865719 | 0.0188517 | 0.0080315 | 0.0091172 | 0.0013486 | 0.0236679 | 0.0054838 | 0.0059892 | 0.0109053 | 0.0082629 | 0.0015113 |
| 32.96 | 0.0186424 | 0.0157191 | 0.0116286 | 0.0129337 | 0.0221935 | 0.0380813 | 0.0326298 | 0.0202404 | 0.0777872 | 0.0162367 | 0.0053296 | 0.0134246 | 0.0017874 | 0.0188485 | 0.0062148 | 0.0058853 | 0.0070117 | 0.0072385 | 0.0016131 |
| 32.97 | 0.0052352 | 0.0108547 | 0.0092468 | 0.017378 | 0.0308268 | 0.0167362 | 0.0142878 | 0.012673 | 0.0385884 | 0.0066609 | 0.0023695 | 0.0222254 | 0.0021389 | 0.0082428 | 0.0077868 | 0.0049891 | 0.0047612 | 0.0054947 | 0.0016818 |
| 32.99 | 0.0103677 | 0.0235376 | 0.0158365 | 0.0297997 | 0.0313304 | 0.0156096 | 0.0086827 | 0.0168558 | 0.0276536 | 0.003063 | 0.0017626 | 0.02498 | 0.0025847 | 0.0045153 | 0.0096966 | 0.0057323 | 0.0064434 | 0.0058806 | 0.0023894 |
| 33 | 0.0371613 | 0.0658431 | 0.0178052 | 0.0297362 | 0.0173106 | 0.0161756 | 0.0051937 | 0.0402693 | 0.0190964 | 0.0010856 | 0.0007144 | 0.0163022 | 0.003654 | 0.0019901 | 0.0099193 | 0.0058785 | 0.0066848 | 0.0047028 | 0.0031349 |
| 33.01 | 0.1340108 | 0.2009802 | 0.0304959 | 0.0352832 | 0.0323193 | 0.0569858 | 0.0295841 | 0.1285467 | 0.0525803 | 0.0077789 | 0.0094325 | 0.015771 | 0.0106161 | 0.0030545 | 0.0143376 | 0.0109186 | 0.0140207 | 0.0147887 | 0.0053177 |
| 33.03 | 0.1611464 | 0.2252775 | 0.0407396 | 0.0406293 | 0.0469003 | 0.0843403 | 0.0571906 | 0.1420773 | 0.0872548 | 0.0170207 | 0.0267996 | 0.0151498 | 0.0236867 | 0.0093535 | 0.0158095 | 0.02031 | 0.0147106 | 0.0352121 | 0.0136214 |
| 33.04 | 0.1597646 | 0.1955352 | 0.0563777 | 0.0815756 | 0.0729442 | 0.1023499 | 0.0886399 | 0.0947662 | 0.1468086 | 0.026475 | 0.0794099 | 0.0483478 | 0.0794734 | 0.052301 | 0.0375431 | 0.0639739 | 0.0393101 | 0.1027145 | 0.0576791 |
| 33.06 | 0.1534748 | 0.1772652 | 0.0508711 | 0.0927073 | 0.0773013 | 0.0944307 | 0.0802568 | 0.0636219 | 0.1499381 | 0.0283004 | 0.0928828 | 0.0697317 | 0.0955809 | 0.0749925 | 0.0449626 | 0.0738128 | 0.069608 | 0.1145725 | 0.0689914 |
| 33.07 | 0.0908416 | 0.1056436 | 0.0236356 | 0.0537626 | 0.0510661 | 0.0556743 | 0.0384978 | 0.0214528 | 0.090136 | 0.0289923 | 0.0475574 | 0.0476268 | 0.0465404 | 0.0520221 | 0.0211348 | 0.0308569 | 0.0646369 | 0.0470017 | 0.0291092 |
| 33.08 | 0.080264 | 0.1207917 | 0.025657 | 0.0528755 | 0.0585269 | 0.0741425 | 0.0394106 | 0.0179625 | 0.1117758 | 0.0318319 | 0.0265562 | 0.0320261 | 0.0248229 | 0.0360099 | 0.0113644 | 0.0129341 | 0.0619979 | 0.0196223 | 0.0133384 |
| 33.1 | 0.0451693 | 0.0891509 | 0.0200971 | 0.0375038 | 0.0412344 | 0.0682177 | 0.0286974 | 0.0107806 | 0.0929139 | 0.0187588 | 0.0150452 | 0.016533 | 0.0165396 | 0.0170861 | 0.0086063 | 0.0070725 | 0.0376646 | 0.0129908 | 0.0116876 |
| 33.11 | 0.0119549 | 0.0381533 | 0.0104367 | 0.0140367 | 0.0123546 | 0.0624371 | 0.0092589 | 0.0035175 | 0.055253 | 0.0248714 | 0.0162937 | 0.017516 | 0.0291248 | 0.0163609 | 0.0122587 | 0.005274 | 0.0132705 | 0.0143491 | 0.017162 |
| 33.13 | 0.0094119 | 0.0287422 | 0.0098012 | 0.0111062 | 0.0081816 | 0.0611539 | 0.0048862 | 0.0061348 | 0.0447657 | 0.0333445 | 0.015805 | 0.0217831 | 0.0350782 | 0.0237189 | 0.0127274 | 0.0055149 | 0.0168821 | 0.0119621 | 0.0152546 |
| 33.14 | 0.0046246 | 0.0168663 | 0.0093241 | 0.0128132 | 0.0100401 | 0.0348573 | 0.0060234 | 0.0098969 | 0.0251939 | 0.0257061 | 0.0081911 | 0.0177387 | 0.0228188 | 0.0223653 | 0.00801 | 0.0059334 | 0.0218907 | 0.005292 | 0.0063612 |
| 33.15 | 0.0098071 | 0.0161882 | 0.0129296 | 0.0296357 | 0.0301381 | 0.0275937 | 0.0188449 | 0.0137015 | 0.0274384 | 0.0400906 | 0.0130574 | 0.0349082 | 0.0213391 | 0.0379406 | 0.0101897 | 0.0117748 | 0.0421422 | 0.0041575 | 0.0070915 |
| 33.17 | 0.018212 | 0.0149644 | 0.0110032 | 0.0360636 | 0.0394011 | 0.0199783 | 0.0243112 | 0.0100867 | 0.0188706 | 0.0416513 | 0.0160212 | 0.0424629 | 0.013891 | 0.0380005 | 0.0095243 | 0.0135274 | 0.0452527 | 0.0033912 | 0.0083704 |
| 33.18 | 0.0249337 | 0.0173869 | 0.0104243 | 0.0317921 | 0.0327847 | 0.0282959 | 0.0219722 | 0.0064473 | 0.0090598 | 0.0302186 | 0.0137546 | 0.0380263 | 0.0042743 | 0.0228971 | 0.0064647 | 0.0090378 | 0.0315931 | 0.0021458 | 0.0075562 |
| 33.19 | 0.0492679 | 0.0307061 | 0.026615 | 0.0476009 | 0.0474085 | 0.0775901 | 0.0372591 | 0.0148627 | 0.0332092 | 0.0444322 | 0.0161576 | 0.0657949 | 0.0041849 | 0.0282515 | 0.0085448 | 0.0087521 | 0.0460242 | 0.0021299 | 0.0091626 |
| 33.21 | 0.0530952 | 0.0259591 | 0.0304248 | 0.0422501 | 0.0446872 | 0.0883565 | 0.0376527 | 0.01641 | 0.0515328 | 0.0390786 | 0.0110409 | 0.0676297 | 0.0045224 | 0.0240423 | 0.00696 | 0.0062278 | 0.0452306 | 0.0018764 | 0.0067565 |
| 33.22 | 0.0576373 | 0.0124123 | 0.0327614 | 0.0314936 | 0.0379201 | 0.0974366 | 0.0372882 | 0.0197902 | 0.0965977 | 0.0206538 | 0.0071156 | 0.061422 | 0.0079019 | 0.0165192 | 0.004454 | 0.0035186 | 0.04432 | 0.0032078 | 0.0046845 |
| 33.24 | 0.0578092 | 0.0086325 | 0.032951 | 0.0275252 | 0.0329453 | 0.0976381 | 0.036854 | 0.0235955 | 0.1172379 | 0.0122911 | 0.0103534 | 0.0502888 | 0.0091714 | 0.0136216 | 0.0034935 | 0.0039451 | 0.0429293 | 0.003845 | 0.0049624 |
| 33.25 | 0.0355131 | 0.0039735 | 0.0206383 | 0.0128796 | 0.0143002 | 0.0590778 | 0.0223296 | 0.0205319 | 0.0879319 | 0.0036949 | 0.0107705 | 0.0168324 | 0.0061072 | 0.0050618 | 0.0009637 | 0.0045977 | 0.0237651 | 0.0027206 | 0.0037155 |
| 33.26 | 0.0388127 | 0.0072999 | 0.0218539 | 0.0086281 | 0.0087622 | 0.070522 | 0.0254327 | 0.0311081 | 0.118563 | 0.0072153 | 0.0130192 | 0.0216297 | 0.0083945 | 0.0054027 | 0.0004358 | 0.0084471 | 0.0183497 | 0.0029231 | 0.00686 |
| 33.28 | 0.0266206 | 0.011457 | 0.0144526 | 0.0052647 | 0.0048367 | 0.0563136 | 0.0189361 | 0.0277973 | 0.1087926 | 0.0102619 | 0.0101515 | 0.0302249 | 0.007309 | 0.0100953 | 0.000682 | 0.0074751 | 0.0110999 | 0.0022711 | 0.0075942 |
| 33.29 | 0.0067447 | 0.0131238 | 0.0029489 | 0.0070075 | 0.0065659 | 0.0210994 | 0.00725 | 0.0229638 | 0.1006899 | 0.0192528 | 0.0096586 | 0.0458987 | 0.0037094 | 0.0256446 | 0.0014873 | 0.0033589 | 0.0175661 | 0.0033116 | 0.0074277 |
| 33.31 | 0.0037614 | 0.0106863 | 0.0014829 | 0.0069484 | 0.0073206 | 0.0094184 | 0.0053878 | 0.0233382 | 0.1006672 | 0.0215124 | 0.0105038 | 0.0464682 | 0.0033855 | 0.0291752 | 0.0015362 | 0.0022036 | 0.0227178 | 0.0040414 | 0.0072143 |
| 33.32 | 0.0039365 | 0.0098113 | 0.0034348 | 0.002415 | 0.0032486 | 0.0038383 | 0.0037425 | 0.0149657 | 0.0633615 | 0.0123562 | 0.0060729 | 0.0247409 | 0.0041934 | 0.0152877 | 0.0012653 | 0.0010811 | 0.0118404 | 0.0021565 | 0.0050424 |
| 33.33 | 0.0117493 | 0.0236066 | 0.0115816 | 0.0045824 | 0.0052887 | 0.0050144 | 0.0039609 | 0.0173323 | 0.0786649 | 0.0106944 | 0.0049621 | 0.0219053 | 0.0093962 | 0.0109276 | 0.0026493 | 0.0027046 | 0.010131 | 0.0016499 | 0.007465 |
| 33.35 | 0.0165535 | 0.0282172 | 0.0139483 | 0.0068973 | 0.0083573 | 0.0036684 | 0.00392 | 0.014443 | 0.067801 | 0.0075388 | 0.0047634 | 0.0136185 | 0.0106895 | 0.0107463 | 0.002738 | 0.004419 | 0.0147306 | 0.0022667 | 0.0072914 |
| 33.36 | 0.0238778 | 0.0349543 | 0.0155171 | 0.0109024 | 0.0156865 | 0.0054325 | 0.0066692 | 0.0089458 | 0.0476704 | 0.0097273 | 0.0114943 | 0.0044233 | 0.0126349 | 0.0260859 | 0.0022745 | 0.0074033 | 0.0300735 | 0.0061335 | 0.0075802 |
| 33.38 | 0.0237693 | 0.0348783 | 0.0162971 | 0.0113563 | 0.0173112 | 0.0105051 | 0.0062879 | 0.0061784 | 0.0370457 | 0.0118181 | 0.0139102 | 0.0052438 | 0.0122554 | 0.0332849 | 0.0027516 | 0.007154 | 0.0317385 | 0.0077659 | 0.0075619 |
| 33.39 | 0.0133338 | 0.0203755 | 0.0120128 | 0.006293 | 0.0104042 | 0.017942 | 0.0026694 | 0.0015479 | 0.0119537 | 0.0079005 | 0.0081529 | 0.0072316 | 0.0062596 | 0.0238625 | 0.0033871 | 0.003204 | 0.0160954 | 0.0063658 | 0.004575 |
| 33.4 | 0.0199527 | 0.0275099 | 0.0205596 | 0.0069164 | 0.0117145 | 0.0429869 | 0.0053213 | 0.0013428 | 0.0123095 | 0.0106887 | 0.0064601 | 0.0173079 | 0.0068691 | 0.0271249 | 0.0054095 | 0.0034805 | 0.0100006 | 0.0108308 | 0.006129 |
| 33.42 | 0.0227872 | 0.0282265 | 0.023334 | 0.0065162 | 0.0105957 | 0.0477645 | 0.0048591 | 0.0020633 | 0.0146134 | 0.0102493 | 0.0045583 | 0.0204777 | 0.005686 | 0.0204301 | 0.0047125 | 0.0031011 | 0.0054608 | 0.0105896 | 0.0054322 |
| 33.43 | 0.0193828 | 0.0206118 | 0.0190506 | 0.0048957 | 0.0061879 | 0.0330772 | 0.0013779 | 0.0022515 | 0.0116543 | 0.0072029 | 0.0033071 | 0.0158167 | 0.0026781 | 0.0074035 | 0.0024715 | 0.0017717 | 0.0040131 | 0.0051316 | 0.0025227 |
| 33.44 | 0.0315207 | 0.0298819 | 0.0291235 | 0.008035 | 0.0057996 | 0.0455137 | 0.003288 | 0.0028737 | 0.016338 | 0.0150697 | 0.0053393 | 0.025984 | 0.0048881 | 0.0107103 | 0.0023076 | 0.0041537 | 0.0063063 | 0.0050458 | 0.0036658 |
| 33.46 | 0.0313751 | 0.0278737 | 0.0273518 | 0.0076363 | 0.0037833 | 0.0427966 | 0.00434 | 0.0023925 | 0.0153919 | 0.0172895 | 0.0070405 | 0.0261598 | 0.0062154 | 0.0143831 | 0.0017117 | 0.0055188 | 0.0075581 | 0.0048028 | 0.0043489 |
| 33.47 | 0.032071 | 0.0275606 | 0.0233377 | 0.0054892 | 0.0018864 | 0.0450104 | 0.0035139 | 0.0032157 | 0.0125759 | 0.0169804 | 0.0127661 | 0.0211768 | 0.0081895 | 0.0235367 | 0.0022347 | 0.0065666 | 0.0164893 | 0.0056247 | 0.0054421 |
| 33.49 | 0.0399086 | 0.0387817 | 0.0242348 | 0.0081541 | 0.0025222 | 0.0453875 | 0.0055494 | 0.0078322 | 0.0122534 | 0.015787 | 0.0144878 | 0.0174675 | 0.0090031 | 0.0259857 | 0.0026818 | 0.0064775 | 0.0202406 | 0.0061439 | 0.0055958 |
| 33.5 | 0.0574046 | 0.0661167 | 0.0227916 | 0.0206831 | 0.0079563 | 0.0238308 | 0.0165811 | 0.0199996 | 0.0201523 | 0.0096505 | 0.0107262 | 0.0088927 | 0.0077074 | 0.0169554 | 0.0022107 | 0.0044513 | 0.0146964 | 0.0052087 | 0.0038511 |
| 33.51 | 0.1614644 | 0.1882655 | 0.0452361 | 0.0729538 | 0.0356553 | 0.0302522 | 0.059639 | 0.0594075 | 0.0786682 | 0.0103518 | 0.0212677 | 0.0121419 | 0.0244147 | 0.0248253 | 0.0112965 | 0.0102262 | 0.0220709 | 0.0194157 | 0.0116729 |
| 33.53 | 0.1776381 | 0.2099675 | 0.048674 | 0.0903055 | 0.0450913 | 0.0318103 | 0.0722212 | 0.0644375 | 0.0986622 | 0.0065023 | 0.0322455 | 0.01039 | 0.0467448 | 0.0300598 | 0.0262658 | 0.0187036 | 0.0267156 | 0.0390999 | 0.021392 |
| 33.54 | 0.1237096 | 0.1573505 | 0.058583 | 0.0934052 | 0.0408914 | 0.0344578 | 0.0704597 | 0.0449075 | 0.0932962 | 0.0024491 | 0.0696594 | 0.0043371 | 0.1066026 | 0.044414 | 0.0742626 | 0.0460638 | 0.0612208 | 0.0945075 | 0.0313751 |
| 33.56 | 0.094518 | 0.1159564 | 0.0637939 | 0.0881746 | 0.0346153 | 0.0371652 | 0.0634828 | 0.0343253 | 0.0838867 | 0.0069691 | 0.080884 | 0.0077158 | 0.1134604 | 0.0459209 | 0.0840196 | 0.0505899 | 0.0807331 | 0.1031512 | 0.0221954 |
| 33.57 | 0.0535998 | 0.0499792 | 0.041427 | 0.0533048 | 0.0199418 | 0.0290562 | 0.0338792 | 0.0175816 | 0.0559137 | 0.0170997 | 0.048055 | 0.0210421 | 0.0522964 | 0.032277 | 0.0365816 | 0.0239727 | 0.0592119 | 0.0473661 | 0.0038154 |
| 33.58 | 0.0816948 | 0.0734155 | 0.048861 | 0.0676502 | 0.0337648 | 0.0470174 | 0.0391962 | 0.024392 | 0.0841463 | 0.0380608 | 0.0476893 | 0.0550423 | 0.0375665 | 0.0571886 | 0.0145443 | 0.018554 | 0.0823341 | 0.0304602 | 0.0039086 |
| 33.6 | 0.0794953 | 0.0759753 | 0.0412209 | 0.0599456 | 0.0347914 | 0.0480939 | 0.0297108 | 0.0230746 | 0.0746764 | 0.0358655 | 0.0385617 | 0.0586037 | 0.0243863 | 0.0578728 | 0.0047686 | 0.0119543 | 0.0806058 | 0.0186172 | 0.0040302 |
| 33.61 | 0.0656253 | 0.0768056 | 0.0290951 | 0.0453833 | 0.032712 | 0.0432072 | 0.0101003 | 0.0201847 | 0.0472311 | 0.0166092 | 0.0318723 | 0.0439517 | 0.0107806 | 0.0477104 | 0.0042659 | 0.0068903 | 0.0647159 | 0.0088033 | 0.0040146 |
| 33.63 | 0.0542555 | 0.0749605 | 0.0243338 | 0.0374459 | 0.0322493 | 0.0387946 | 0.0067862 | 0.0198304 | 0.0352918 | 0.0095917 | 0.0308784 | 0.0292214 | 0.0064408 | 0.0398309 | 0.0058195 | 0.0103397 | 0.0492768 | 0.0080539 | 0.0058653 |
| 33.64 | 0.0208362 | 0.0425162 | 0.0119953 | 0.0164594 | 0.019936 | 0.0227479 | 0.0041104 | 0.0128508 | 0.011483 | 0.0070083 | 0.0204599 | 0.0062264 | 0.0041637 | 0.0167663 | 0.0048555 | 0.0132298 | 0.014965 | 0.009203 | 0.0106113 |
| 33.65 | 0.0126924 | 0.0361981 | 0.0136889 | 0.0148114 | 0.020072 | 0.0299266 | 0.005817 | 0.0116735 | 0.0118204 | 0.0099108 | 0.0293628 | 0.0119431 | 0.0102221 | 0.0124658 | 0.0047383 | 0.0208321 | 0.0076883 | 0.0165739 | 0.0274803 |
| 33.67 | 0.01108 | 0.0218541 | 0.0126117 | 0.0105731 | 0.0142005 | 0.0243182 | 0.0089751 | 0.0088802 | 0.0200507 | 0.0062172 | 0.0272647 | 0.0158958 | 0.0089725 | 0.0079886 | 0.0029912 | 0.0158611 | 0.006151 | 0.0160346 | 0.030695 |
| 33.68 | 0.0161032 | 0.0124924 | 0.008117 | 0.0069671 | 0.0097801 | 0.007992 | 0.0102607 | 0.0117079 | 0.0289987 | 0.0013288 | 0.0145943 | 0.0131578 | 0.0022431 | 0.006138 | 0.0019597 | 0.0032863 | 0.0080886 | 0.009596 | 0.0181498 |
| 33.69 | 0.0349553 | 0.036184 | 0.0104568 | 0.0180177 | 0.0275848 | 0.0115201 | 0.0154036 | 0.0357619 | 0.0645813 | 0.0044565 | 0.0155082 | 0.019876 | 0.0012871 | 0.0181838 | 0.0026856 | 0.0009345 | 0.019154 | 0.0096313 | 0.0142621 |

| | | | | | | | | | | | | | | | | | | | |
|---|---|---|---|---|---|---|---|---|---|---|---|---|---|---|---|---|---|---|---|
| 33.71 | 0.035331 | 0.0446583 | 0.0091424 | 0.0200028 | 0.0324621 | 0.0165818 | 0.0137699 | 0.0459534 | 0.0765516 | 0.0079965 | 0.0117283 | 0.0180161 | 0.001505 | 0.0227802 | 0.0019468 | 0.0008076 | 0.0221141 | 0.0059829 | 0.0088194 |
| 33.72 | 0.0300694 | 0.0504243 | 0.0084496 | 0.0141374 | 0.0311299 | 0.0238679 | 0.0093093 | 0.0584117 | 0.0928042 | 0.0149011 | 0.0045232 | 0.0135706 | 0.0059809 | 0.0298291 | 0.0038102 | 0.0029758 | 0.0277957 | 0.000909 | 0.0125694 |
| 33.74 | 0.027531 | 0.0506287 | 0.0100491 | 0.0099362 | 0.0271073 | 0.0229838 | 0.00856 | 0.0556135 | 0.0919523 | 0.0150666 | 0.0023025 | 0.0111265 | 0.0093706 | 0.0327518 | 0.0055636 | 0.0043595 | 0.0301172 | 0.000276 | 0.0181523 |
| 33.75 | 0.0144803 | 0.0333595 | 0.0112209 | 0.0082077 | 0.0116256 | 0.012446 | 0.0079947 | 0.0244493 | 0.0487948 | 0.0063477 | 0.0010474 | 0.0046771 | 0.0075709 | 0.0221472 | 0.0033439 | 0.0028511 | 0.0203649 | 0.0003106 | 0.0150748 |
| 33.76 | 0.015314 | 0.0433405 | 0.0263218 | 0.0281752 | 0.0078024 | 0.0146322 | 0.0173442 | 0.0143758 | 0.0363848 | 0.0076861 | 0.0012107 | 0.0107081 | 0.0070328 | 0.024574 | 0.0034983 | 0.0022437 | 0.0235217 | 0.0003692 | 0.0181177 |
| 33.78 | 0.0148279 | 0.0369246 | 0.0305855 | 0.0353003 | 0.0041818 | 0.0111737 | 0.0190983 | 0.0062731 | 0.0255568 | 0.010212 | 0.0009399 | 0.0164291 | 0.0055723 | 0.017952 | 0.0050971 | 0.0023808 | 0.0163925 | 0.0003528 | 0.0141817 |
| 33.79 | 0.0187666 | 0.0245051 | 0.0339761 | 0.0366782 | 0.003884 | 0.0060009 | 0.0191241 | 0.0015407 | 0.0511165 | 0.0136886 | 0.0013826 | 0.0260429 | 0.0067034 | 0.006204 | 0.0089123 | 0.0048062 | 0.0063182 | 0.0010157 | 0.0060766 |
| 33.81 | 0.0210709 | 0.0194055 | 0.0323684 | 0.0320297 | 0.0081332 | 0.0084181 | 0.0172077 | 0.0022816 | 0.0668728 | 0.0125365 | 0.0013339 | 0.0271485 | 0.0065228 | 0.0034843 | 0.009749 | 0.0065613 | 0.0077477 | 0.0018932 | 0.0037772 |
| 33.82 | 0.0198785 | 0.0094121 | 0.0157331 | 0.0120169 | 0.0155636 | 0.012124 | 0.0070123 | 0.0036515 | 0.0480304 | 0.005711 | 0.0003483 | 0.0156999 | 0.0027723 | 0.0031524 | 0.0072717 | 0.0064738 | 0.008081 | 0.0021947 | 0.0027608 |
| 33.83 | 0.055117 | 0.0277985 | 0.0122303 | 0.0150905 | 0.0464175 | 0.0246181 | 0.0084693 | 0.0077473 | 0.0569919 | 0.0043427 | 0.0014225 | 0.0153083 | 0.0019356 | 0.0045992 | 0.0108425 | 0.0092413 | 0.0073924 | 0.0023239 | 0.003699 |
| 33.85 | 0.075195 | 0.0472786 | 0.0112203 | 0.0255373 | 0.0610721 | 0.0252808 | 0.0119727 | 0.0066639 | 0.0409755 | 0.0040339 | 0.0031657 | 0.0100493 | 0.0012809 | 0.0031276 | 0.0102852 | 0.0073125 | 0.005213 | 0.0019555 | 0.0032009 |
| 33.86 | 0.1099445 | 0.0891127 | 0.0249222 | 0.0534264 | 0.0911518 | 0.0221558 | 0.0163018 | 0.0048991 | 0.0196246 | 0.011314 | 0.0072158 | 0.0066647 | 0.0020103 | 0.0016631 | 0.007347 | 0.0032579 | 0.0109368 | 0.0043126 | 0.0043556 |
| 33.88 | 0.1164226 | 0.0986815 | 0.0319682 | 0.0628118 | 0.1058668 | 0.0196368 | 0.0134209 | 0.0084316 | 0.0314182 | 0.0169883 | 0.0094934 | 0.0116305 | 0.0028748 | 0.0018609 | 0.0049355 | 0.0020779 | 0.0159239 | 0.0057121 | 0.0040492 |
| 33.89 | 0.0740872 | 0.0691406 | 0.0250577 | 0.0494645 | 0.0819441 | 0.009328 | 0.0039324 | 0.0115781 | 0.0507698 | 0.017762 | 0.0110415 | 0.0169933 | 0.0020015 | 0.0021492 | 0.0010031 | 0.0010328 | 0.0147198 | 0.0047801 | 0.0015088 |
| 33.9 | 0.1000963 | 0.1155297 | 0.0400055 | 0.0777634 | 0.1173792 | 0.0065885 | 0.0020855 | 0.0212199 | 0.1170223 | 0.0323315 | 0.0269781 | 0.0377827 | 0.0034662 | 0.0029725 | 0.0003389 | 0.0021176 | 0.0220599 | 0.0078651 | 0.0035229 |
| 33.92 | 0.0899862 | 0.1218165 | 0.0396228 | 0.0744545 | 0.1045693 | 0.0041469 | 0.0019715 | 0.0209219 | 0.1313812 | 0.0317312 | 0.0309644 | 0.0406017 | 0.0053001 | 0.0031241 | 0.0006989 | 0.0022458 | 0.0186414 | 0.0068866 | 0.0051499 |
| 33.93 | 0.0427824 | 0.0799182 | 0.021737 | 0.0402735 | 0.0460122 | 0.0047809 | 0.0024057 | 0.0120305 | 0.0910372 | 0.0157169 | 0.0208003 | 0.0216415 | 0.0053223 | 0.0054034 | 0.0014376 | 0.0014547 | 0.005996 | 0.0026967 | 0.0046533 |
| 33.94 | 0.0383956 | 0.1013489 | 0.0185895 | 0.0383745 | 0.031367 | 0.0113869 | 0.0046072 | 0.0151757 | 0.1281935 | 0.0114347 | 0.026471 | 0.0156626 | 0.0068591 | 0.015949 | 0.0039618 | 0.0015247 | 0.0035433 | 0.0050623 | 0.0055574 |
| 33.96 | 0.0246761 | 0.0893449 | 0.0120314 | 0.0256224 | 0.0265307 | 0.0096324 | 0.0048808 | 0.0128093 | 0.1197226 | 0.0066486 | 0.0235471 | 0.0095686 | 0.0053581 | 0.0188332 | 0.005637 | 0.0009667 | 0.0046753 | 0.0067012 | 0.003736 |
| 33.97 | 0.0077162 | 0.0703034 | 0.0131669 | 0.0197718 | 0.0580323 | 0.0071787 | 0.0096758 | 0.0059327 | 0.091657 | 0.0070251 | 0.0175964 | 0.0141384 | 0.0032496 | 0.0183629 | 0.0056475 | 0.0014269 | 0.0103086 | 0.0089073 | 0.0010541 |
| 33.99 | 0.0148452 | 0.0766895 | 0.0162922 | 0.0259528 | 0.0605529 | 0.0089013 | 0.0112536 | 0.0062393 | 0.0837149 | 0.0102422 | 0.0146876 | 0.0216062 | 0.0040576 | 0.0157557 | 0.0039438 | 0.0022339 | 0.0122881 | 0.0093954 | 0.0015495 |
| 34 | 0.057624 | 0.0978822 | 0.0150475 | 0.0206263 | 0.0211687 | 0.0117457 | 0.0107755 | 0.0167759 | 0.0713635 | 0.0103995 | 0.0073601 | 0.0215877 | 0.0033482 | 0.0073684 | 0.001834 | 0.002061 | 0.0091903 | 0.0054125 | 0.0036916 |
| 34.01 | 0.2275941 | 0.2706587 | 0.0552173 | 0.0548433 | 0.0374048 | 0.0626161 | 0.050356 | 0.0633248 | 0.1724999 | 0.0213368 | 0.0165348 | 0.0348882 | 0.017035 | 0.0133171 | 0.0091248 | 0.0119059 | 0.0101985 | 0.019425 | 0.0085504 |
| 34.03 | 0.273715 | 0.3140201 | 0.0765914 | 0.083196 | 0.0580043 | 0.0958622 | 0.0685888 | 0.0765155 | 0.2003833 | 0.0222653 | 0.0339809 | 0.0312149 | 0.0416338 | 0.023831 | 0.020557 | 0.0286566 | 0.0191595 | 0.0479864 | 0.0133416 |
| 34.04 | 0.2322868 | 0.2844027 | 0.0781273 | 0.1047946 | 0.0981587 | 0.1172961 | 0.0596237 | 0.063117 | 0.1930742 | 0.0250223 | 0.0931564 | 0.0410233 | 0.0997831 | 0.0635761 | 0.0578537 | 0.0852144 | 0.0775794 | 0.1358407 | 0.0528666 |
| 34.06 | 0.1927147 | 0.2431694 | 0.0627761 | 0.088744 | 0.1045029 | 0.106051 | 0.0430386 | 0.0501903 | 0.1743859 | 0.0387337 | 0.109355 | 0.059337 | 0.1035648 | 0.0767453 | 0.0679875 | 0.1003422 | 0.0970687 | 0.1532255 | 0.0715654 |
| 34.07 | 0.0958001 | 0.1162574 | 0.025174 | 0.0358676 | 0.058241 | 0.0543584 | 0.0132404 | 0.0261646 | 0.0975535 | 0.043139 | 0.0582594 | 0.0421161 | 0.0429941 | 0.0420277 | 0.0318898 | 0.0463663 | 0.0482767 | 0.0703467 | 0.0361738 |
| 34.08 | 0.1023971 | 0.1152392 | 0.020043 | 0.0307024 | 0.0533945 | 0.0585242 | 0.0069773 | 0.0347714 | 0.1178842 | 0.0426976 | 0.0362805 | 0.0288867 | 0.025512 | 0.0268456 | 0.0152857 | 0.022171 | 0.0313871 | 0.038237 | 0.0177794 |
| 34.1 | 0.0776059 | 0.0896916 | 0.0123179 | 0.0203328 | 0.035104 | 0.0482295 | 0.0049387 | 0.0303262 | 0.0982849 | 0.0252668 | 0.0236152 | 0.0226441 | 0.0228637 | 0.0193171 | 0.0114887 | 0.0161357 | 0.0319172 | 0.0274743 | 0.0158419 |
| 34.11 | 0.0367227 | 0.0574754 | 0.0036553 | 0.0067111 | 0.0103244 | 0.0400171 | 0.011789 | 0.0176231 | 0.056099 | 0.02868 | 0.0316659 | 0.0354444 | 0.0410872 | 0.0265672 | 0.0202148 | 0.0230451 | 0.0508476 | 0.045065 | 0.0260859 |
| 34.13 | 0.0242213 | 0.046954 | 0.003231 | 0.006372 | 0.0064368 | 0.0389364 | 0.0167941 | 0.0122295 | 0.0381099 | 0.0383356 | 0.0315077 | 0.0410529 | 0.047211 | 0.0284489 | 0.0223231 | 0.0221565 | 0.0503936 | 0.0514381 | 0.024523 |
| 34.14 | 0.0078262 | 0.0194955 | 0.0062569 | 0.009229 | 0.0103073 | 0.0224454 | 0.0154133 | 0.0046674 | 0.01089 | 0.0298641 | 0.0128202 | 0.0301921 | 0.0322935 | 0.0199855 | 0.0134529 | 0.0115124 | 0.0280649 | 0.0323768 | 0.0093501 |
| 34.15 | 0.0150102 | 0.0142634 | 0.0182841 | 0.0252845 | 0.0361862 | 0.0184165 | 0.025585 | 0.0038008 | 0.0071735 | 0.0410699 | 0.0125872 | 0.0385183 | 0.0413695 | 0.03464 | 0.0145827 | 0.0155013 | 0.0360287 | 0.038524 | 0.0052657 |
| 34.17 | 0.0276567 | 0.0137544 | 0.0227034 | 0.0330667 | 0.0457601 | 0.0114798 | 0.0264068 | 0.0029914 | 0.0080532 | 0.0367532 | 0.0143869 | 0.030678 | 0.0360491 | 0.037654 | 0.0131124 | 0.0163518 | 0.0346404 | 0.0358184 | 0.0041304 |
| 34.18 | 0.0376426 | 0.0157145 | 0.0195066 | 0.0309401 | 0.0337106 | 0.0103934 | 0.0194385 | 0.0024558 | 0.0117702 | 0.020954 | 0.0134302 | 0.0123696 | 0.0179761 | 0.0282491 | 0.0085906 | 0.0115667 | 0.0220765 | 0.0231983 | 0.0036711 |
| 34.19 | 0.0698529 | 0.0300917 | 0.0332161 | 0.0560447 | 0.0454892 | 0.0372463 | 0.0341761 | 0.003462 | 0.0288612 | 0.0388027 | 0.0213904 | 0.0158114 | 0.0178544 | 0.0458666 | 0.0118136 | 0.0158431 | 0.0308066 | 0.0270818 | 0.0056114 |
| 34.21 | 0.0698563 | 0.0300467 | 0.034182 | 0.0600877 | 0.0433042 | 0.0535949 | 0.0371744 | 0.0030061 | 0.0340853 | 0.0487004 | 0.0208974 | 0.0190395 | 0.0120498 | 0.0460156 | 0.0104862 | 0.014364 | 0.0295731 | 0.0184292 | 0.0048954 |
| 34.22 | 0.0591307 | 0.0238309 | 0.0332624 | 0.0670624 | 0.0418792 | 0.0911718 | 0.041681 | 0.0029358 | 0.0397201 | 0.0655342 | 0.0175193 | 0.0340086 | 0.0050916 | 0.0389206 | 0.0067727 | 0.0110286 | 0.0274564 | 0.0067843 | 0.0037876 |
| 34.24 | 0.0472659 | 0.0196013 | 0.0309454 | 0.0666019 | 0.0370349 | 0.1041741 | 0.0414728 | 0.0026784 | 0.0393832 | 0.0652495 | 0.0142853 | 0.0401447 | 0.0056802 | 0.0310581 | 0.0057583 | 0.0099583 | 0.0260436 | 0.0088822 | 0.004548 |
| 34.25 | 0.0163345 | 0.0194476 | 0.0169475 | 0.0339619 | 0.0159826 | 0.0687934 | 0.0226837 | 0.0012625 | 0.0252743 | 0.033643 | 0.0050366 | 0.0248823 | 0.0044359 | 0.0099352 | 0.004284 | 0.0058559 | 0.0144687 | 0.0118487 | 0.0044488 |
| 34.26 | 0.0196624 | 0.0704156 | 0.0214038 | 0.0261132 | 0.0325184 | 0.0785623 | 0.0202657 | 0.0036334 | 0.0443191 | 0.0253976 | 0.0046614 | 0.0186592 | 0.0039813 | 0.0096163 | 0.0066807 | 0.0052504 | 0.0146965 | 0.0251311 | 0.005729 |
| 34.28 | 0.0201249 | 0.0818897 | 0.0188742 | 0.0161856 | 0.0448216 | 0.062706 | 0.0137552 | 0.0054089 | 0.0507386 | 0.0188843 | 0.0065421 | 0.0093009 | 0.0034718 | 0.0118401 | 0.00651 | 0.00309 | 0.0107215 | 0.0265123 | 0.0048882 |
| 34.29 | 0.0143408 | 0.0621544 | 0.0109827 | 0.0080278 | 0.0581075 | 0.0383238 | 0.0084027 | 0.0075989 | 0.0592994 | 0.0082742 | 0.0103578 | 0.0030925 | 0.0077997 | 0.0130431 | 0.0094941 | 0.0042942 | 0.0060194 | 0.0198545 | 0.0070924 |
| 34.31 | 0.0104959 | 0.0528523 | 0.0077641 | 0.0060708 | 0.057476 | 0.0292099 | 0.0088095 | 0.008776 | 0.0593878 | 0.0099433 | 0.0107753 | 0.0027696 | 0.0109562 | 0.0103445 | 0.0116962 | 0.0066048 | 0.0050461 | 0.0147535 | 0.0086825 |
| 34.32 | 0.0043457 | 0.0330238 | 0.0041708 | 0.0018044 | 0.0348177 | 0.0103005 | 0.008109 | 0.007825 | 0.0382248 | 0.0086583 | 0.0086167 | 0.0013783 | 0.0097711 | 0.0038515 | 0.0077965 | 0.0050462 | 0.0026788 | 0.0045823 | 0.0056893 |
| 34.33 | 0.005734 | 0.0498448 | 0.0051831 | 0.0021927 | 0.0442149 | 0.01674 | 0.0137745 | 0.0108908 | 0.0564827 | 0.0163002 | 0.0154812 | 0.002677 | 0.0179976 | 0.0033845 | 0.0081732 | 0.0050502 | 0.004819 | 0.0035128 | 0.0078269 |
| 34.35 | 0.0063806 | 0.0495189 | 0.0035903 | 0.0031617 | 0.0389857 | 0.0290559 | 0.0122334 | 0.0084002 | 0.0563269 | 0.0178271 | 0.0157592 | 0.0039546 | 0.0202836 | 0.0041988 | 0.0069117 | 0.0045067 | 0.0065698 | 0.0047897 | 0.0083946 |
| 34.36 | 0.0108362 | 0.0474029 | 0.0015634 | 0.0047715 | 0.0323769 | 0.0623308 | 0.0060547 | 0.0032984 | 0.0607828 | 0.0192264 | 0.0136379 | 0.0077449 | 0.0230172 | 0.0115526 | 0.0056796 | 0.0059124 | 0.0114179 | 0.0082867 | 0.0086429 |
| 34.38 | 0.0160613 | 0.050981 | 0.0014393 | 0.0068742 | 0.0346331 | 0.0728815 | 0.0045197 | 0.0030883 | 0.0682864 | 0.018038 | 0.010794 | 0.0079457 | 0.0236496 | 0.0162772 | 0.0049671 | 0.0070171 | 0.0137882 | 0.0093713 | 0.0076591 |
| 34.39 | 0.0181125 | 0.0459344 | 0.0015433 | 0.0089429 | 0.0294196 | 0.0470025 | 0.0038902 | 0.004588 | 0.056264 | 0.0072422 | 0.0034492 | 0.004215 | 0.0165327 | 0.0155419 | 0.0024353 | 0.00576 | 0.0116079 | 0.0085861 | 0.0038776 |
| 34.4 | 0.0257643 | 0.0732334 | 0.0082854 | 0.011552 | 0.0396057 | 0.0396256 | 0.0072305 | 0.0062046 | 0.0819658 | 0.0065664 | 0.0055153 | 0.0123109 | 0.0227757 | 0.0250281 | 0.0021426 | 0.0074275 | 0.0186437 | 0.0170228 | 0.0045373 |
| 34.42 | 0.0186084 | 0.0614516 | 0.012908 | 0.0077458 | 0.0308189 | 0.021006 | 0.0093178 | 0.0037808 | 0.0723343 | 0.0107067 | 0.0080696 | 0.0185346 | 0.0189795 | 0.0227356 | 0.0013719 | 0.0059084 | 0.0178979 | 0.0187527 | 0.0046722 |
| 34.43 | 0.007171 | 0.0258669 | 0.0103203 | 0.0041018 | 0.0188483 | 0.0034054 | 0.0099586 | 0.0015875 | 0.0380527 | 0.0153809 | 0.0092351 | 0.0179501 | 0.0065792 | 0.0097034 | 0.0011614 | 0.0034742 | 0.0100642 | 0.0137295 | 0.0050179 |
| 34.44 | 0.0227314 | 0.0430492 | 0.008198 | 0.0057371 | 0.0582569 | 0.0092034 | 0.0129273 | 0.0050251 | 0.0469876 | 0.0285104 | 0.0169902 | 0.0266354 | 0.0077754 | 0.0074116 | 0.0040602 | 0.0062889 | 0.0105798 | 0.0177636 | 0.0093251 |
| 34.46 | 0.0307079 | 0.0495585 | 0.0046024 | 0.0038289 | 0.072671 | 0.0146246 | 0.0088136 | 0.0071665 | 0.0397911 | 0.0283298 | 0.0162158 | 0.0236446 | 0.010719 | 0.0072203 | 0.0042404 | 0.0056215 | 0.0077012 | 0.0128388 | 0.0078348 |
| 34.47 | 0.0277813 | 0.040981 | 0.0042967 | 0.0022595 | 0.0589215 | 0.0202349 | 0.0053493 | 0.0081458 | 0.0272886 | 0.0259361 | 0.0132247 | 0.0174318 | 0.0109915 | 0.010418 | 0.0029373 | 0.0031513 | 0.0052632 | 0.0042859 | 0.0032105 |
| 34.49 | 0.0219225 | 0.0325814 | 0.0105852 | 0.0054011 | 0.0388062 | 0.0176563 | 0.0129987 | 0.0077109 | 0.0269219 | 0.0259265 | 0.0131984 | 0.0162941 | 0.0081666 | 0.0089289 | 0.0036581 | 0.0036984 | 0.007015 | 0.003272 | 0.0025748 |
| 34.5 | 0.0229687 | 0.0364971 | 0.0248653 | 0.0143491 | 0.0077084 | 0.0125967 | 0.0319571 | 0.0093029 | 0.0323251 | 0.0181486 | 0.0098121 | 0.0121139 | 0.0070267 | 0.0044858 | 0.00512 | 0.0052953 | 0.0099562 | 0.0026383 | 0.0024365 |
| 34.51 | 0.0893068 | 0.1210415 | 0.0547696 | 0.0502377 | 0.0061883 | 0.0483111 | 0.0949586 | 0.0270134 | 0.1165728 | 0.025171 | 0.0231647 | 0.0207053 | 0.0325748 | 0.0217098 | 0.0206941 | 0.0177696 | 0.0345295 | 0.0183129 | 0.0100237 |
| 34.53 | 0.1134128 | 0.1431449 | 0.0556616 | 0.066319 | 0.0117629 | 0.0682848 | 0.1094056 | 0.0310449 | 0.1501354 | 0.0220217 | 0.0377255 | 0.0196264 | 0.0634871 | 0.0376075 | 0.0408791 | 0.027395 | 0.0560634 | 0.0394506 | 0.0185687 |
| 34.54 | 0.1076078 | 0.1324816 | 0.073449 | 0.0866632 | 0.0387357 | 0.0875951 | 0.0958098 | 0.028713 | 0.1434846 | 0.0160775 | 0.0725209 | 0.0098837 | 0.143733 | 0.0610709 | 0.1029074 | 0.0381014 | 0.1026774 | 0.0884588 | 0.0248802 |
| 34.56 | 0.0985211 | 0.1252633 | 0.0887344 | 0.0922047 | 0.0526913 | 0.0870226 | 0.0829056 | 0.0250812 | 0.1277428 | 0.0178937 | 0.0771673 | 0.0081415 | 0.1513704 | 0.0619318 | 0.1159274 | 0.0334657 | 0.1075294 | 0.0926652 | 0.0167318 |
| 34.57 | 0.063965 | 0.0905929 | 0.0637979 | 0.0645069 | 0.042706 | 0.0547823 | 0.0460343 | 0.0119884 | 0.0817362 | 0.0199793 | 0.0432137 | 0.0147484 | 0.0634139 | 0.0405979 | 0.0537638 | 0.0154907 | 0.057049 | 0.0434755 | 0.0047424 |
| 34.58 | 0.0854826 | 0.1602598 | 0.0723376 | 0.0826779 | 0.060458 | 0.0817271 | 0.0543193 | 0.014155 | 0.128975 | 0.037379 | 0.0454222 | 0.03929 | 0.0362563 | 0.0630365 | 0.0285723 | 0.0169408 | 0.0592639 | 0.0398636 | 0.0104849 |
| 34.6 | 0.0719257 | 0.1688696 | 0.0538897 | 0.0641083 | 0.0545649 | 0.08206 | 0.042555 | 0.0153825 | 0.1318276 | 0.0355683 | 0.0377791 | 0.0414147 | 0.0213729 | 0.0602277 | 0.0151827 | 0.0127368 | 0.0484619 | 0.0314674 | 0.010091 |
| 34.61 | 0.0451892 | 0.1647433 | 0.0187493 | 0.0220823 | 0.0365903 | 0.0749896 | 0.0208758 | 0.0206962 | 0.1345451 | 0.023046 | 0.036118 | 0.0348957 | 0.0111216 | 0.048393 | 0.0073408 | 0.0049525 | 0.0332262 | 0.0168879 | 0.0049738 |
| 34.63 | 0.0391763 | 0.1521697 | 0.0085554 | 0.0106567 | 0.0288131 | 0.0680267 | 0.0163833 | 0.0199818 | 0.1314916 | 0.0153 | 0.0386667 | 0.029155 | 0.0073252 | 0.0420497 | 0.0052996 | 0.003243 | 0.0255502 | 0.0123678 | 0.0044412 |
| 34.64 | 0.0225158 | 0.078049 | 0.0045427 | 0.0063883 | 0.0147349 | 0.0363374 | 0.0136564 | 0.0076123 | 0.0716824 | 0.0053229 | 0.0270353 | 0.0101677 | 0.0019978 | 0.0227226 | 0.0062308 | 0.0046103 | 0.0084095 | 0.0130357 | 0.0038102 |
| 34.65 | 0.0184342 | 0.0683834 | 0.0117023 | 0.0192172 | 0.0142899 | 0.030594 | 0.0300656 | 0.0055915 | 0.0603081 | 0.011816 | 0.035975 | 0.0055033 | 0.0026327 | 0.0325249 | 0.0141635 | 0.0112489 | 0.0089478 | 0.0359944 | 0.0044528 |
| 34.67 | 0.0128905 | 0.0440117 | 0.0153958 | 0.028278 | 0.014433 | 0.0253948 | 0.0330782 | 0.007994 | 0.037845 | 0.0133827 | 0.0324011 | 0.0049696 | 0.0023193 | 0.0326058 | 0.0133434 | 0.009594 | 0.0105135 | 0.0385819 | 0.0051529 |
| 34.68 | 0.0162304 | 0.0256125 | 0.0161574 | 0.025776 | 0.0235384 | 0.0358784 | 0.0237136 | 0.0113477 | 0.0236129 | 0.009009 | 0.0186905 | 0.0058238 | 0.0005045 | 0.0196234 | 0.0050637 | 0.0027073 | 0.0080313 | 0.0217589 | 0.005105 |
| 34.69 | 0.0444231 | 0.0728598 | 0.0291236 | 0.0266724 | 0.0617464 | 0.0885685 | 0.0420114 | 0.0247615 | 0.0775837 | 0.0163294 | 0.0259392 | 0.0142902 | 0.0014461 | 0.0172686 | 0.0029816 | 0.0027683 | 0.0080135 | 0.0269922 | 0.0055892 |
| 34.71 | 0.0491912 | 0.0891176 | 0.0281981 | 0.0174239 | 0.072042 | 0.0951883 | 0.0453903 | 0.0279264 | 0.1015516 | 0.017937 | 0.024665 | 0.017291 | 0.0032036 | 0.010349 | 0.0024158 | 0.0022882 | 0.0053814 | 0.025144 | 0.0049226 |
| 34.72 | 0.0442395 | 0.0964877 | 0.0207248 | 0.0060099 | 0.0778924 | 0.0816902 | 0.0483735 | 0.0278255 | 0.1236292 | 0.0150879 | 0.0232676 | 0.0185777 | 0.0049474 | 0.0071302 | 0.0033615 | 0.0013497 | 0.0049471 | 0.0214972 | 0.0066599 |
| 34.74 | 0.0374242 | 0.0896849 | 0.0157663 | 0.0041478 | 0.0694825 | 0.0651633 | 0.0495231 | 0.0250819 | 0.1254973 | 0.011086 | 0.0241397 | 0.0166394 | 0.0042069 | 0.008936 | 0.0069559 | 0.0023276 | 0.007498 | 0.0181189 | 0.0060359 |
| 34.75 | 0.0151049 | 0.0458426 | 0.006147 | 0.0050014 | 0.0276171 | 0.0200421 | 0.0340816 | 0.013735 | 0.0822952 | 0.0060134 | 0.0169199 | 0.0075549 | 0.0031919 | 0.0066567 | 0.0105465 | 0.0037193 | 0.0079405 | 0.0063475 | 0.0025994 |
| 34.76 | 0.0112282 | 0.0466618 | 0.0166541 | 0.0165567 | 0.0149659 | 0.0124375 | 0.0513131 | 0.016961 | 0.1084445 | 0.0190709 | 0.0210598 | 0.0057157 | 0.0073489 | 0.0058345 | 0.0180274 | 0.006438 | 0.0112482 | 0.0066975 | 0.0041875 |
| 34.78 | 0.0070788 | 0.0355847 | 0.0267286 | 0.0221167 | 0.0067725 | 0.0160229 | 0.0506038 | 0.0138061 | 0.0907604 | 0.0259296 | 0.0164239 | 0.0038732 | 0.0084554 | 0.0047779 | 0.0149891 | 0.0051037 | 0.0088593 | 0.0112557 | 0.0035022 |
| 34.79 | 0.0021046 | 0.0171063 | 0.0419913 | 0.026506 | 0.002484 | 0.0234965 | 0.0468936 | 0.0081515 | 0.0480916 | 0.0322791 | 0.0061852 | 0.0072628 | 0.0124718 | 0.0080871 | 0.0063391 | 0.0020591 | 0.0038711 | 0.0217547 | 0.0009997 |
| 34.81 | 0.0010565 | 0.0101886 | 0.0417234 | 0.0243797 | 0.002541 | 0.0179575 | 0.0445162 | 0.0086423 | 0.0305083 | 0.0303303 | 0.0040373 | 0.0111793 | 0.0144876 | 0.0085166 | 0.0054874 | 0.001767 | 0.0029996 | 0.0199617 | 0.0009648 |
| 34.82 | 0.0006221 | 0.003305 | 0.0216882 | 0.0117098 | 0.0044075 | 0.0056761 | 0.0261678 | 0.009308 | 0.0076399 | 0.0129599 | 0.0056901 | 0.0103215 | 0.0093343 | 0.0035629 | 0.0058912 | 0.0016778 | 0.0024529 | 0.0077829 | 0.0015544 |
| 34.83 | 0.0022376 | 0.0071427 | 0.0196715 | 0.0081676 | 0.0162599 | 0.0178498 | 0.0304004 | 0.0177586 | 0.0108013 | 0.0105736 | 0.0171989 | 0.01404 | 0.0091846 | 0.0039595 | 0.0091575 | 0.0035378 | 0.0047137 | 0.0170368 | 0.0027614 |
| 34.85 | 0.0024387 | 0.0075535 | 0.0128812 | 0.0044397 | 0.0248097 | 0.0274513 | 0.0239643 | 0.0186967 | 0.0143105 | 0.0125734 | 0.0207226 | 0.0121572 | 0.0066893 | 0.0044516 | 0.007047 | 0.004134 | 0.0050341 | 0.0226844 | 0.0041492 |
| 34.86 | 0.0033244 | 0.0057859 | 0.0122231 | 0.007112 | 0.0396878 | 0.0446798 | 0.0097026 | 0.0210988 | 0.0135176 | 0.0227331 | 0.0243105 | 0.0088688 | 0.0045337 | 0.0030607 | 0.0029466 | 0.0047904 | 0.0040981 | 0.0244589 | 0.0127048 |
| 34.88 | 0.0076311 | 0.0088132 | 0.0197829 | 0.010667 | 0.043377 | 0.0520382 | 0.0050757 | 0.0228756 | 0.0119877 | 0.0256954 | 0.026319 | 0.0070563 | 0.0045033 | 0.0020573 | 0.0038033 | 0.0043772 | 0.0028256 | 0.0211573 | 0.0164399 |
| 34.89 | 0.0133904 | 0.0159978 | 0.022559 | 0.0102162 | 0.0333035 | 0.0440905 | 0.0030656 | 0.0164101 | 0.0165271 | 0.0200206 | 0.0222739 | 0.002738 | 0.0033305 | 0.0025921 | 0.0058757 | 0.0019571 | 0.0006637 | 0.0090926 | 0.0103981 |
| 34.9 | 0.0303294 | 0.0389707 | 0.0388115 | 0.016503 | 0.0667839 | 0.0834975 | 0.00539 | 0.022304 | 0.052738 | 0.033561 | 0.0449465 | 0.0050953 | 0.0061053 | 0.0074476 | 0.0108517 | 0.0029128 | 0.0007647 | 0.0063642 | 0.0089056 |
| 34.92 | 0.0348758 | 0.0410561 | 0.0405397 | 0.0160189 | 0.0766837 | 0.0891541 | 0.0050116 | 0.0197619 | 0.0669493 | 0.0333497 | 0.0512309 | 0.0060132 | 0.0070346 | 0.0087564 | 0.0092936 | 0.0038504 | 0.0009 | 0.0042876 | 0.0064277 |
| 34.93 | 0.0252591 | 0.0212985 | 0.0311891 | 0.010979 | 0.0552343 | 0.0568953 | 0.0040925 | 0.0099469 | 0.0572496 | 0.0189151 | 0.0384543 | 0.0030722 | 0.0057361 | 0.0071089 | 0.0029469 | 0.0041506 | 0.0009436 | 0.0049392 | 0.0043177 |
| 34.94 | 0.0265051 | 0.0163023 | 0.0453038 | 0.0161406 | 0.062161 | 0.072551 | 0.0075707 | 0.0087281 | 0.0951693 | 0.0203888 | 0.0556132 | 0.002873 | 0.0103869 | 0.0129815 | 0.0022212 | 0.0077651 | 0.0029649 | 0.0100781 | 0.0081147 |
| 34.96 | 0.0176797 | 0.0098801 | 0.0341077 | 0.0128005 | 0.0460674 | 0.0625425 | 0.0066176 | 0.0077922 | 0.0919211 | 0.0164577 | 0.0520307 | 0.0035694 | 0.0114016 | 0.0138567 | 0.002139 | 0.0078389 | 0.0041345 | 0.0078637 | 0.0078417 |
| 34.97 | 0.0137723 | 0.0079226 | 0.0131594 | 0.0056579 | 0.0557604 | 0.0385034 | 0.0049546 | 0.0180782 | 0.0693312 | 0.0111123 | 0.04436 | 0.0048248 | 0.0119728 | 0.0136207 | 0.0021215 | 0.0070369 | 0.0060191 | 0.0021302 | 0.0064583 |
| 34.99 | 0.016285 | 0.008745 | 0.0171173 | 0.0048012 | 0.0828133 | 0.0301872 | 0.0097104 | 0.0210074 | 0.0655162 | 0.0085714 | 0.0407562 | 0.0050113 | 0.0115281 | 0.0118342 | 0.0020517 | 0.0068578 | 0.0061839 | 0.0031486 | 0.0067971 |
| 35 | 0.0224026 | 0.0140077 | 0.020218 | 0.0079711 | 0.0705109 | 0.0216584 | 0.0263965 | 0.0113312 | 0.0638755 | 0.0031259 | 0.0231707 | 0.0049913 | 0.0055987 | 0.0046604 | 0.0013956 | 0.0046105 | 0.0036017 | 0.0074682 | 0.0052113 |
| 35.01 | 0.0840951 | 0.0783068 | 0.0536158 | 0.0361505 | 0.0695822 | 0.0657467 | 0.0851769 | 0.0202904 | 0.1482681 | 0.0080677 | 0.0404539 | 0.0099676 | 0.0218209 | 0.0091802 | 0.0155767 | 0.0057805 | 0.0098945 | 0.0154959 | 0.0057426 |
| 35.03 | 0.1000618 | 0.1133318 | 0.071748 | 0.0439901 | 0.0479769 | 0.0868106 | 0.0925945 | 0.0267614 | 0.1576357 | 0.0104562 | 0.0626036 | 0.0152396 | 0.0487334 | 0.0247634 | 0.0389598 | 0.0097077 | 0.0281024 | 0.0312368 | 0.0093373 |
| 35.04 | 0.0764361 | 0.1330878 | 0.0635349 | 0.0282245 | 0.0194044 | 0.0901968 | 0.0547269 | 0.0370733 | 0.1206013 | 0.0198839 | 0.1384577 | 0.060802 | 0.1074432 | 0.0948347 | 0.1003697 | 0.0488191 | 0.1019773 | 0.1185065 | 0.0479756 |
| 35.06 | 0.0624786 | 0.1269772 | 0.0448424 | 0.0168869 | 0.0117057 | 0.075288 | 0.0321581 | 0.0358743 | 0.1000352 | 0.0353776 | 0.1564049 | 0.0862673 | 0.1117625 | 0.1232612 | 0.1092428 | 0.0688202 | 0.1243167 | 0.1554113 | 0.0709524 |
| 35.07 | 0.0369605 | 0.0812228 | 0.0159831 | 0.0051098 | 0.0102811 | 0.0342123 | 0.0074318 | 0.017734 | 0.0558377 | 0.0398095 | 0.0798196 | 0.0535532 | 0.052113 | 0.076884 | 0.0472782 | 0.0397419 | 0.0667845 | 0.0986182 | 0.0501943 |

| | | | | | | | | | | | | | | | | | | | |
|---|---|---|---|---|---|---|---|---|---|---|---|---|---|---|---|---|---|---|---|
| 35.08 | 0.0507085 | 0.1186532 | 0.0118264 | 0.0085119 | 0.0260269 | 0.0412574 | 0.0048586 | 0.0150701 | 0.0739446 | 0.045059 | 0.0453831 | 0.0343359 | 0.0310336 | 0.0571862 | 0.021801 | 0.0234757 | 0.0391685 | 0.0670173 | 0.0343854 |
| 35.1 | 0.0457183 | 0.1121699 | 0.0071204 | 0.0109111 | 0.0299841 | 0.0399988 | 0.0033805 | 0.0087718 | 0.066435 | 0.0279394 | 0.0192758 | 0.0219253 | 0.0163442 | 0.0303124 | 0.0108013 | 0.015549 | 0.0189756 | 0.0348211 | 0.0151238 |
| 35.11 | 0.0382979 | 0.0972287 | 0.0028259 | 0.0157195 | 0.0331755 | 0.0454739 | 0.0041423 | 0.0041135 | 0.0598608 | 0.0121433 | 0.0134799 | 0.0304379 | 0.0281019 | 0.0153637 | 0.0125153 | 0.0201154 | 0.0243909 | 0.055987 | 0.0068487 |
| 35.13 | 0.0368371 | 0.091401 | 0.0033692 | 0.0153668 | 0.0300208 | 0.0490746 | 0.0044656 | 0.0070521 | 0.0645476 | 0.0191288 | 0.0160237 | 0.0382416 | 0.0418463 | 0.0202153 | 0.0142143 | 0.0211518 | 0.0324793 | 0.0813012 | 0.0068791 |
| 35.14 | 0.0221348 | 0.0533831 | 0.0054437 | 0.0061671 | 0.0116242 | 0.0341728 | 0.0024604 | 0.0095469 | 0.0491079 | 0.0232662 | 0.009587 | 0.0329946 | 0.0371573 | 0.0187478 | 0.0090943 | 0.0117239 | 0.0243941 | 0.0706748 | 0.0034453 |
| 35.15 | 0.0211247 | 0.0588362 | 0.0150798 | 0.0065559 | 0.0186959 | 0.0457172 | 0.0081067 | 0.0145596 | 0.0641155 | 0.0337333 | 0.0079972 | 0.0524756 | 0.0518636 | 0.0282457 | 0.0087443 | 0.0112034 | 0.0333524 | 0.1077241 | 0.0063463 |
| 35.17 | 0.0147955 | 0.0412405 | 0.0197167 | 0.0122095 | 0.0337451 | 0.0383964 | 0.0147137 | 0.0113187 | 0.0506759 | 0.0286454 | 0.0047355 | 0.04791 | 0.0453643 | 0.0282476 | 0.0055885 | 0.0070595 | 0.0308076 | 0.1029481 | 0.0063964 |
| 35.18 | 0.0113374 | 0.0103501 | 0.0194613 | 0.0207585 | 0.0486188 | 0.0139509 | 0.020001 | 0.0042925 | 0.0154624 | 0.0133484 | 0.0020706 | 0.0216562 | 0.020644 | 0.0197341 | 0.0021245 | 0.0014065 | 0.0165329 | 0.0554122 | 0.0025662 |
| 35.19 | 0.0293196 | 0.0171087 | 0.0382718 | 0.0478881 | 0.1012828 | 0.0081661 | 0.0432273 | 0.0131698 | 0.010951 | 0.0160298 | 0.0030072 | 0.0181318 | 0.0137697 | 0.0342027 | 0.0047191 | 0.0019635 | 0.0165992 | 0.047538 | 0.0018648 |
| 35.21 | 0.0348136 | 0.0250249 | 0.0424835 | 0.0539939 | 0.1088499 | 0.0077309 | 0.0479191 | 0.0201601 | 0.01569 | 0.0160265 | 0.003792 | 0.0134678 | 0.0080447 | 0.0398767 | 0.0049817 | 0.0019313 | 0.0122063 | 0.0299986 | 0.0017915 |
| 35.22 | 0.0452426 | 0.0408011 | 0.0497451 | 0.0602011 | 0.1078338 | 0.0158644 | 0.051577 | 0.0285952 | 0.0300532 | 0.019044 | 0.0091504 | 0.0112056 | 0.0120965 | 0.0499147 | 0.0027778 | 0.0023433 | 0.0070194 | 0.0291571 | 0.0039434 |
| 35.24 | 0.0512297 | 0.0452372 | 0.0514766 | 0.0630623 | 0.1009989 | 0.0163337 | 0.0514477 | 0.0270825 | 0.0286824 | 0.0201277 | 0.0115267 | 0.011444 | 0.0157405 | 0.0480307 | 0.0024155 | 0.0040268 | 0.0075336 | 0.0435953 | 0.0055535 |
| 35.25 | 0.0390414 | 0.0292019 | 0.0318377 | 0.0462835 | 0.0582536 | 0.0063579 | 0.0326235 | 0.0109951 | 0.0114734 | 0.0131019 | 0.0083728 | 0.0077177 | 0.010365 | 0.0225004 | 0.0035159 | 0.0042881 | 0.0074352 | 0.0447432 | 0.0047823 |
| 35.26 | 0.0560487 | 0.0337187 | 0.0375116 | 0.0674477 | 0.0755389 | 0.0076206 | 0.0438582 | 0.011418 | 0.0170632 | 0.0163955 | 0.0096999 | 0.0084645 | 0.0097356 | 0.0156135 | 0.0073549 | 0.0046481 | 0.0105251 | 0.0747396 | 0.0063024 |
| 35.28 | 0.0490276 | 0.0268476 | 0.0303685 | 0.0563506 | 0.0649119 | 0.0110703 | 0.038686 | 0.0156954 | 0.0224646 | 0.0134748 | 0.0069026 | 0.005532 | 0.0065338 | 0.0113554 | 0.0073007 | 0.0029881 | 0.0078786 | 0.0722546 | 0.00498 |
| 35.29 | 0.028323 | 0.0139238 | 0.0147317 | 0.0245857 | 0.0340597 | 0.0194994 | 0.0257363 | 0.025986 | 0.0305987 | 0.0074664 | 0.0037668 | 0.0048214 | 0.0019845 | 0.0197488 | 0.0095664 | 0.0030524 | 0.0026298 | 0.0554226 | 0.0029151 |
| 35.31 | 0.0198683 | 0.0086286 | 0.0081386 | 0.0190783 | 0.0217132 | 0.0220434 | 0.0214982 | 0.0278971 | 0.0309598 | 0.0057482 | 0.0038117 | 0.0062381 | 0.0008319 | 0.0233897 | 0.0129602 | 0.0053861 | 0.0015644 | 0.046749 | 0.0032921 |
| 35.32 | 0.0087417 | 0.0020099 | 0.0014245 | 0.0167936 | 0.0145952 | 0.0155716 | 0.0137939 | 0.0175447 | 0.0172927 | 0.003123 | 0.0017757 | 0.0037466 | 0.0002847 | 0.0140299 | 0.0126012 | 0.0065803 | 0.0010972 | 0.0241347 | 0.0030709 |
| 35.33 | 0.0164782 | 0.0033218 | 0.0048407 | 0.0262803 | 0.0393602 | 0.0186785 | 0.0214185 | 0.0168458 | 0.0170106 | 0.0037896 | 0.001544 | 0.0053679 | 0.0004519 | 0.01355 | 0.0187478 | 0.0097384 | 0.002102 | 0.027906 | 0.0028807 |
| 35.35 | 0.0200176 | 0.0037651 | 0.0094811 | 0.0259407 | 0.0455329 | 0.0149717 | 0.0195319 | 0.012166 | 0.0150551 | 0.0033918 | 0.0025464 | 0.0067166 | 0.000539 | 0.0089015 | 0.0160493 | 0.0075572 | 0.0023942 | 0.0224473 | 0.0013975 |
| 35.36 | 0.0218571 | 0.0057429 | 0.0177561 | 0.0258721 | 0.0512626 | 0.0120399 | 0.0140849 | 0.0115036 | 0.0249481 | 0.0038438 | 0.0053853 | 0.0091823 | 0.0007965 | 0.0021768 | 0.0088437 | 0.0024892 | 0.003998 | 0.0093105 | 0.0006235 |
| 35.38 | 0.0201634 | 0.0091317 | 0.0193503 | 0.0229687 | 0.0534303 | 0.0125904 | 0.0126762 | 0.0120474 | 0.0339163 | 0.0038651 | 0.0062551 | 0.0096921 | 0.000994 | 0.0010321 | 0.0055755 | 0.0012137 | 0.0052288 | 0.0045898 | 0.0015673 |
| 35.39 | 0.0125526 | 0.008051 | 0.0139673 | 0.0089893 | 0.0324227 | 0.0087626 | 0.0079287 | 0.0055026 | 0.0297688 | 0.0017974 | 0.0053195 | 0.0061861 | 0.0015622 | 0.0006355 | 0.0011539 | 0.000579 | 0.0047686 | 0.0020485 | 0.0028926 |
| 35.4 | 0.0203692 | 0.0094376 | 0.0226437 | 0.005819 | 0.0316086 | 0.0128272 | 0.0088306 | 0.0046811 | 0.0469063 | 0.0017799 | 0.0081943 | 0.0087223 | 0.0034414 | 0.0028831 | 0.002252 | 0.0018446 | 0.0094199 | 0.0053892 | 0.0045723 |
| 35.42 | 0.0200877 | 0.0094809 | 0.0230008 | 0.0059549 | 0.0215091 | 0.012001 | 0.0073863 | 0.0052278 | 0.0498334 | 0.0021599 | 0.0062373 | 0.009669 | 0.0028698 | 0.005452 | 0.0039644 | 0.0034209 | 0.0115587 | 0.0073879 | 0.003311 |
| 35.43 | 0.0116488 | 0.0077381 | 0.0149204 | 0.0088763 | 0.0060263 | 0.0057647 | 0.006627 | 0.0042222 | 0.0395953 | 0.0034797 | 0.0011847 | 0.0081161 | 0.0007741 | 0.006894 | 0.0034923 | 0.0039181 | 0.0100346 | 0.0074966 | 0.0010243 |
| 35.44 | 0.015145 | 0.0124333 | 0.019603 | 0.0225111 | 0.0067243 | 0.0064085 | 0.0170609 | 0.0041265 | 0.0742563 | 0.0078173 | 0.0017149 | 0.0112909 | 0.0009066 | 0.0119694 | 0.0032859 | 0.0037716 | 0.0144841 | 0.0116282 | 0.0017633 |
| 35.46 | 0.0139031 | 0.0125039 | 0.0170577 | 0.0258882 | 0.0067749 | 0.006715 | 0.0206739 | 0.0043776 | 0.0836322 | 0.0072708 | 0.0030173 | 0.0095466 | 0.0012154 | 0.011793 | 0.002925 | 0.0019105 | 0.0129304 | 0.0101494 | 0.0019186 |
| 35.47 | 0.0120827 | 0.0108215 | 0.0133012 | 0.0251414 | 0.0053904 | 0.0102213 | 0.0245493 | 0.0109214 | 0.0863582 | 0.005808 | 0.0030463 | 0.0078551 | 0.0028366 | 0.0117867 | 0.0026687 | 0.0006833 | 0.0109853 | 0.0074336 | 0.0016185 |
| 35.49 | 0.0179788 | 0.0162196 | 0.017763 | 0.0198488 | 0.0069376 | 0.0115766 | 0.0274035 | 0.0131165 | 0.0745035 | 0.0072962 | 0.0021857 | 0.0088932 | 0.004564 | 0.0120326 | 0.0019717 | 0.0009728 | 0.0114282 | 0.0075537 | 0.0016233 |
| 35.5 | 0.0382986 | 0.0426954 | 0.0269772 | 0.0070567 | 0.0064619 | 0.007587 | 0.0269538 | 0.009362 | 0.0272665 | 0.0072939 | 0.0021001 | 0.0077349 | 0.00569 | 0.0077125 | 0.0019124 | 0.0010732 | 0.0092141 | 0.0053912 | 0.0007465 |
| 35.51 | 0.1233948 | 0.1396391 | 0.0500221 | 0.0084775 | 0.0059321 | 0.0208496 | 0.0728947 | 0.0205928 | 0.0346865 | 0.0088738 | 0.0199016 | 0.0134554 | 0.0279775 | 0.0148123 | 0.0148012 | 0.0032614 | 0.0237632 | 0.0107022 | 0.0110113 |
| 35.53 | 0.1402902 | 0.1537819 | 0.0452601 | 0.0144126 | 0.0104818 | 0.0402913 | 0.0997389 | 0.023164 | 0.0518402 | 0.0059155 | 0.0480355 | 0.0172182 | 0.0620454 | 0.0241046 | 0.0348834 | 0.0058007 | 0.0412378 | 0.0199918 | 0.0336093 |
| 35.54 | 0.103952 | 0.1059777 | 0.0418789 | 0.0536371 | 0.0424483 | 0.0797777 | 0.1286476 | 0.0182821 | 0.0816174 | 0.0088357 | 0.1164903 | 0.0481973 | 0.1563395 | 0.0534694 | 0.0971321 | 0.0098985 | 0.1005081 | 0.0548445 | 0.09422 |
| 35.56 | 0.0823336 | 0.0794257 | 0.0450126 | 0.0723441 | 0.0593346 | 0.0782006 | 0.1172767 | 0.0140275 | 0.0847671 | 0.0239491 | 0.1233213 | 0.0727991 | 0.1675168 | 0.060101 | 0.1083314 | 0.0125945 | 0.116184 | 0.0660017 | 0.1010998 |
| 35.57 | 0.0452788 | 0.0418589 | 0.0334823 | 0.0549237 | 0.0502963 | 0.0415213 | 0.0559162 | 0.0058045 | 0.0605301 | 0.0375845 | 0.0579293 | 0.0617819 | 0.0697135 | 0.0390693 | 0.0468768 | 0.0142405 | 0.0606083 | 0.0426851 | 0.0468995 |
| 35.58 | 0.0542852 | 0.0615277 | 0.0461732 | 0.0739604 | 0.0692751 | 0.0611192 | 0.0629351 | 0.0112015 | 0.0984026 | 0.0504958 | 0.0418307 | 0.0656445 | 0.0366874 | 0.0623836 | 0.0234388 | 0.0210131 | 0.0472471 | 0.0553744 | 0.0368206 |
| 35.6 | 0.0413843 | 0.0543435 | 0.040779 | 0.0653471 | 0.0586239 | 0.0605235 | 0.0538599 | 0.0141185 | 0.0990126 | 0.0382659 | 0.0255523 | 0.0479054 | 0.021213 | 0.0599185 | 0.0117356 | 0.0159625 | 0.0333981 | 0.0502065 | 0.0255309 |
| 35.61 | 0.0141084 | 0.0259209 | 0.0317599 | 0.0455672 | 0.0323465 | 0.0508619 | 0.0367199 | 0.0137945 | 0.0922441 | 0.0209237 | 0.0115559 | 0.0291862 | 0.0120625 | 0.032109 | 0.0046351 | 0.008428 | 0.020073 | 0.0297641 | 0.0113446 |
| 35.63 | 0.0066589 | 0.0136047 | 0.027887 | 0.0377593 | 0.0215567 | 0.0430458 | 0.0305312 | 0.0101134 | 0.0830975 | 0.0148483 | 0.0088261 | 0.0250945 | 0.0077403 | 0.0210468 | 0.0038019 | 0.0105641 | 0.0150677 | 0.0190284 | 0.0072206 |
| 35.64 | 0.0065939 | 0.0028935 | 0.0126881 | 0.0199655 | 0.0069828 | 0.0207184 | 0.0186776 | 0.0036936 | 0.0400559 | 0.0059404 | 0.0051486 | 0.0142731 | 0.0021257 | 0.0196145 | 0.004044 | 0.0131064 | 0.0074821 | 0.0113209 | 0.0054263 |
| 35.65 | 0.0226377 | 0.0076849 | 0.0124117 | 0.0217793 | 0.0061921 | 0.0230157 | 0.0301915 | 0.0106878 | 0.0424852 | 0.0115968 | 0.0081201 | 0.0195522 | 0.0031119 | 0.0434166 | 0.0059183 | 0.0197579 | 0.0125055 | 0.0324309 | 0.0128967 |
| 35.67 | 0.0276194 | 0.0135159 | 0.0138485 | 0.0174358 | 0.0076423 | 0.0177645 | 0.0298927 | 0.0146099 | 0.0332654 | 0.0145552 | 0.0071147 | 0.0172816 | 0.0028259 | 0.0385364 | 0.0044127 | 0.0145757 | 0.013205 | 0.0385768 | 0.0128144 |
| 35.68 | 0.0205467 | 0.0182719 | 0.0122634 | 0.0130554 | 0.0141918 | 0.0053797 | 0.0177004 | 0.0116014 | 0.0117723 | 0.0114888 | 0.0022457 | 0.0070886 | 0.0015636 | 0.0115951 | 0.0013496 | 0.0033447 | 0.0110222 | 0.0280457 | 0.0050316 |
| 35.69 | 0.0313658 | 0.0410069 | 0.0147519 | 0.029833 | 0.0465311 | 0.0090415 | 0.0223306 | 0.0135586 | 0.0082047 | 0.0155063 | 0.0007789 | 0.0053194 | 0.0043646 | 0.0035375 | 0.0006287 | 0.0025494 | 0.0205412 | 0.0454632 | 0.0050256 |
| 35.71 | 0.0317389 | 0.0480433 | 0.010575 | 0.0321958 | 0.0630364 | 0.0164222 | 0.0192617 | 0.0116644 | 0.0068858 | 0.012483 | 0.0004481 | 0.0033146 | 0.0048453 | 0.0015076 | 0.0003232 | 0.002561 | 0.0200319 | 0.0488946 | 0.0056379 |
| 35.72 | 0.0321163 | 0.0587625 | 0.003383 | 0.0272831 | 0.0922471 | 0.03842 | 0.0134973 | 0.0160115 | 0.016054 | 0.0060254 | 0.0019882 | 0.0018459 | 0.0041375 | 0.0068848 | 0.0011618 | 0.0046479 | 0.0111007 | 0.0568618 | 0.0047494 |
| 35.74 | 0.0305671 | 0.0611194 | 0.0017096 | 0.0213442 | 0.0962423 | 0.0472872 | 0.0144991 | 0.0181095 | 0.0229729 | 0.0066426 | 0.0034775 | 0.0032983 | 0.0054904 | 0.0113962 | 0.0026007 | 0.0061989 | 0.007325 | 0.058192 | 0.0047759 |
| 35.75 | 0.0158323 | 0.0400234 | 0.0013031 | 0.0066657 | 0.0529972 | 0.0336381 | 0.0148957 | 0.0082711 | 0.023508 | 0.0087193 | 0.0037034 | 0.0058515 | 0.0071635 | 0.0107906 | 0.0041956 | 0.0042552 | 0.0070131 | 0.0358142 | 0.006355 |
| 35.76 | 0.0152532 | 0.0538418 | 0.0050105 | 0.0091347 | 0.0480885 | 0.042105 | 0.0302732 | 0.0056313 | 0.0482744 | 0.0164406 | 0.005548 | 0.0172508 | 0.0138168 | 0.0119038 | 0.0098148 | 0.003876 | 0.0135738 | 0.0397748 | 0.0138317 |
| 35.78 | 0.0117266 | 0.0492411 | 0.0075608 | 0.0140071 | 0.0300953 | 0.0361487 | 0.0334436 | 0.0072796 | 0.053374 | 0.0154085 | 0.0049217 | 0.0222934 | 0.0130354 | 0.0086972 | 0.0115163 | 0.004123 | 0.0105825 | 0.0272323 | 0.0137133 |
| 35.79 | 0.0085539 | 0.0411235 | 0.00823 | 0.0262072 | 0.0104987 | 0.0243467 | 0.0361057 | 0.0139514 | 0.0559186 | 0.0094901 | 0.0032087 | 0.0266306 | 0.0063417 | 0.0120051 | 0.0139936 | 0.0089523 | 0.0054159 | 0.0110737 | 0.0086437 |
| 35.81 | 0.0089186 | 0.0394925 | 0.0087641 | 0.0299942 | 0.0125178 | 0.0202382 | 0.035425 | 0.0152439 | 0.0551075 | 0.006949 | 0.0023134 | 0.0245489 | 0.0042527 | 0.018297 | 0.0142504 | 0.0106901 | 0.0056566 | 0.0165263 | 0.0054789 |
| 35.82 | 0.0083872 | 0.0278296 | 0.0131341 | 0.0207499 | 0.0099857 | 0.0133979 | 0.0196755 | 0.0118069 | 0.03557 | 0.0041385 | 0.00084 | 0.0121518 | 0.0051169 | 0.0202213 | 0.0089889 | 0.0068589 | 0.0032084 | 0.0235174 | 0.0014039 |
| 35.83 | 0.018418 | 0.0462327 | 0.0256986 | 0.0257083 | 0.0144715 | 0.0352873 | 0.0150249 | 0.0167009 | 0.0542206 | 0.0096438 | 0.0012501 | 0.014294 | 0.0136919 | 0.0377144 | 0.0131109 | 0.006462 | 0.0027683 | 0.042922 | 0.0017006 |
| 35.85 | 0.0217854 | 0.04755 | 0.0208468 | 0.0206246 | 0.0254643 | 0.049504 | 0.0091899 | 0.0145832 | 0.0592127 | 0.0121489 | 0.001431 | 0.0124622 | 0.0150364 | 0.0381197 | 0.0132879 | 0.0040282 | 0.0031375 | 0.0406226 | 0.0018559 |
| 35.86 | 0.0273645 | 0.0440217 | 0.0200432 | 0.0089398 | 0.0589668 | 0.0731596 | 0.0154346 | 0.0252506 | 0.0812072 | 0.0153805 | 0.001935 | 0.008008 | 0.0112669 | 0.0292314 | 0.0130054 | 0.0015943 | 0.0058659 | 0.0217172 | 0.00145 |
| 35.88 | 0.0300149 | 0.0389759 | 0.0321643 | 0.0043653 | 0.0723569 | 0.0760999 | 0.0217415 | 0.0365555 | 0.0971398 | 0.0151565 | 0.0022256 | 0.0058196 | 0.0076057 | 0.0214179 | 0.0116219 | 0.0018314 | 0.0079533 | 0.0139672 | 0.0011159 |
| 35.89 | 0.0231803 | 0.0178405 | 0.0365007 | 0.0011438 | 0.0591477 | 0.0490735 | 0.0200721 | 0.0330898 | 0.085249 | 0.0082633 | 0.0015445 | 0.0021806 | 0.0019082 | 0.0057515 | 0.005213 | 0.0022974 | 0.0081749 | 0.0112743 | 0.0011798 |
| 35.9 | 0.0376572 | 0.0128494 | 0.0593713 | 0.0058336 | 0.0911997 | 0.0714191 | 0.0365537 | 0.0514152 | 0.139644 | 0.0103323 | 0.0018879 | 0.0032127 | 0.0019425 | 0.0038379 | 0.0040639 | 0.0051818 | 0.0143894 | 0.0216491 | 0.0037716 |
| 35.92 | 0.0388102 | 0.0099382 | 0.057839 | 0.0093137 | 0.0908106 | 0.0678937 | 0.0381596 | 0.0499964 | 0.1306029 | 0.0111047 | 0.0024868 | 0.0039322 | 0.0024209 | 0.0040933 | 0.0023801 | 0.0055617 | 0.0145081 | 0.0190067 | 0.0053195 |
| 35.93 | 0.0265097 | 0.0129074 | 0.0366616 | 0.0085513 | 0.064648 | 0.0415533 | 0.0236519 | 0.03159 | 0.0653639 | 0.0103962 | 0.0039827 | 0.0036105 | 0.0025301 | 0.0041267 | 0.0006974 | 0.0026522 | 0.0092355 | 0.0071148 | 0.0062395 |
| 35.94 | 0.0327783 | 0.0336049 | 0.0486952 | 0.0088185 | 0.0973871 | 0.0620589 | 0.026589 | 0.0449632 | 0.0662976 | 0.0199169 | 0.0112726 | 0.0065542 | 0.0043733 | 0.0068031 | 0.0016322 | 0.0017896 | 0.0125001 | 0.0056902 | 0.0140149 |
| 35.96 | 0.0234778 | 0.0390191 | 0.0412303 | 0.005538 | 0.086256 | 0.0571372 | 0.0199339 | 0.0393299 | 0.0525617 | 0.0208977 | 0.0146385 | 0.0067493 | 0.003963 | 0.0061058 | 0.0021267 | 0.0014389 | 0.011741 | 0.0075009 | 0.0152639 |
| 35.97 | 0.0101189 | 0.0448979 | 0.0190745 | 0.0035568 | 0.0442509 | 0.033253 | 0.0083643 | 0.0211639 | 0.035172 | 0.0196025 | 0.0211132 | 0.005723 | 0.0019782 | 0.0031735 | 0.001717 | 0.0021376 | 0.010364 | 0.018755 | 0.0125911 |
| 35.99 | 0.0107695 | 0.0402032 | 0.0102796 | 0.0051209 | 0.0244575 | 0.0222149 | 0.0136381 | 0.0214293 | 0.034173 | 0.017648 | 0.0231272 | 0.0046697 | 0.0012942 | 0.0026109 | 0.0014718 | 0.003958 | 0.0090821 | 0.0245205 | 0.0099262 |
| 36 | 0.0209748 | 0.031288 | 0.0063385 | 0.012615 | 0.0118603 | 0.0164632 | 0.0417529 | 0.0251001 | 0.0470482 | 0.0087874 | 0.0167964 | 0.0017841 | 0.0008674 | 0.0040476 | 0.0015528 | 0.0059784 | 0.0040838 | 0.0223706 | 0.0048019 |
| 36.01 | 0.0997375 | 0.1429744 | 0.0375939 | 0.0470086 | 0.0683799 | 0.0693318 | 0.1337963 | 0.0400065 | 0.1689429 | 0.0103537 | 0.0412206 | 0.0026532 | 0.0112089 | 0.010078 | 0.0046211 | 0.0124499 | 0.0121615 | 0.034059 | 0.006491 |
| 36.03 | 0.1354053 | 0.190152 | 0.0584314 | 0.0526567 | 0.1087469 | 0.0983154 | 0.1454378 | 0.054189 | 0.212152 | 0.0148857 | 0.0723194 | 0.0115647 | 0.0394818 | 0.0147418 | 0.0123738 | 0.013134 | 0.0345111 | 0.0377873 | 0.012405 |
| 36.04 | 0.1294481 | 0.1463202 | 0.0597862 | 0.0276914 | 0.1434316 | 0.1128893 | 0.0915331 | 0.1389253 | 0.1981148 | 0.050131 | 0.1689899 | 0.0626288 | 0.1451938 | 0.0487221 | 0.0425622 | 0.0200661 | 0.111798 | 0.1053069 | 0.050739 |
| 36.06 | 0.0956978 | 0.0935306 | 0.0147066 | 0.0149232 | 0.129083 | 0.096903 | 0.0574409 | 0.1684587 | 0.1629623 | 0.0686351 | 0.1897391 | 0.0785936 | 0.1744043 | 0.0646034 | 0.0509088 | 0.0275641 | 0.1242914 | 0.1473093 | 0.0676692 |
| 36.07 | 0.0260687 | 0.0265232 | 0.0103076 | 0.0077754 | 0.0638436 | 0.0414794 | 0.0130481 | 0.1007527 | 0.0740763 | 0.0404173 | 0.08625 | 0.0384967 | 0.0867532 | 0.0357341 | 0.0252265 | 0.0201394 | 0.055366 | 0.0922964 | 0.0369836 |
| 36.08 | 0.0140348 | 0.0308125 | 0.0049121 | 0.0133349 | 0.0779052 | 0.0405037 | 0.0048348 | 0.0800794 | 0.0754718 | 0.0289882 | 0.0405876 | 0.044274 | 0.0496891 | 0.0372642 | 0.0241216 | 0.0274539 | 0.0601165 | 0.0571575 | 0.0278047 |
| 36.1 | 0.0075508 | 0.0308001 | 0.0025277 | 0.0117574 | 0.067734 | 0.0328272 | 0.0020643 | 0.0465663 | 0.0513001 | 0.0251398 | 0.0242129 | 0.0486072 | 0.0359158 | 0.0434188 | 0.0299392 | 0.032501 | 0.0640349 | 0.0335624 | 0.0314353 |
| 36.11 | 0.0040893 | 0.0305898 | 0.0037763 | 0.0097192 | 0.050028 | 0.0245723 | 0.002345 | 0.0430803 | 0.0211815 | 0.0192425 | 0.0180158 | 0.0394348 | 0.0616723 | 0.0515513 | 0.0390187 | 0.0324189 | 0.0502525 | 0.0423579 | 0.0421634 |
| 36.13 | 0.0059693 | 0.0303913 | 0.0059604 | 0.0101005 | 0.0428157 | 0.0214338 | 0.0035716 | 0.0661444 | 0.0300181 | 0.0136563 | 0.0113904 | 0.0306599 | 0.0743255 | 0.0457419 | 0.0366706 | 0.0252349 | 0.0347708 | 0.0530497 | 0.0384985 |
| 36.14 | 0.0079777 | 0.0219352 | 0.0072115 | 0.0080832 | 0.0160771 | 0.0096075 | 0.0046752 | 0.0766864 | 0.0360716 | 0.0097175 | 0.0063123 | 0.0164799 | 0.0527356 | 0.0203011 | 0.0215684 | 0.0107551 | 0.0111075 | 0.0463082 | 0.021741 |
| 36.15 | 0.0204204 | 0.0368499 | 0.0167827 | 0.0156009 | 0.0153198 | 0.0063285 | 0.0113069 | 0.1587981 | 0.0577235 | 0.0134813 | 0.0122111 | 0.0201175 | 0.0621454 | 0.013972 | 0.022667 | 0.0099633 | 0.0072168 | 0.0731265 | 0.0219451 |
| 36.17 | 0.0276959 | 0.0356313 | 0.0204219 | 0.0179174 | 0.0256181 | 0.0049437 | 0.013718 | 0.175079 | 0.0573314 | 0.0098549 | 0.0111613 | 0.0159518 | 0.0493888 | 0.0070113 | 0.0156359 | 0.0062115 | 0.003977 | 0.0721723 | 0.0141804 |
| 36.18 | 0.0294695 | 0.0162228 | 0.0180543 | 0.014821 | 0.0382383 | 0.0091163 | 0.0115511 | 0.1187842 | 0.0374605 | 0.0026938 | 0.0031935 | 0.0052483 | 0.0196665 | 0.0009351 | 0.0038439 | 0.0010923 | 0.0017414 | 0.050824 | 0.0048162 |
| 36.19 | 0.0595468 | 0.0103064 | 0.0349091 | 0.0270593 | 0.0886008 | 0.0307821 | 0.0179277 | 0.1654555 | 0.0514335 | 0.0037801 | 0.0024853 | 0.0057966 | 0.0144642 | 0.0008834 | 0.0023284 | 0.0008686 | 0.0088038 | 0.0740037 | 0.0115668 |
| 36.21 | 0.0652083 | 0.0083987 | 0.0402094 | 0.0295105 | 0.1088296 | 0.0421994 | 0.0167443 | 0.1551678 | 0.0483912 | 0.0070469 | 0.0044034 | 0.0080727 | 0.0095561 | 0.0023764 | 0.00204 | 0.0010437 | 0.0125502 | 0.0653386 | 0.0121105 |
| 36.22 | 0.06832 | 0.0189668 | 0.0497109 | 0.0321337 | 0.1465775 | 0.062763 | 0.0147533 | 0.1248401 | 0.0453409 | 0.0147608 | 0.0097238 | 0.0078151 | 0.0084215 | 0.0096886 | 0.0019597 | 0.002515 | 0.0096318 | 0.0475165 | 0.0068105 |
| 36.24 | 0.0672564 | 0.0250488 | 0.0516819 | 0.0321379 | 0.1544858 | 0.064538 | 0.0150136 | 0.1133753 | 0.0478238 | 0.0163619 | 0.0116985 | 0.0060597 | 0.0093582 | 0.0136357 | 0.0028097 | 0.0034441 | 0.00617 | 0.0400238 | 0.0064748 |
| 36.25 | 0.0432268 | 0.0208112 | 0.0330023 | 0.020723 | 0.0996304 | 0.0337146 | 0.010892 | 0.0699369 | 0.0381692 | 0.0114179 | 0.0096325 | 0.0054244 | 0.005999 | 0.0110526 | 0.0033353 | 0.0022976 | 0.0044333 | 0.0146469 | 0.005336 |
| 36.26 | 0.0631028 | 0.0324999 | 0.0429163 | 0.0306997 | 0.1393534 | 0.0308097 | 0.0181379 | 0.0930534 | 0.0596962 | 0.0164975 | 0.0145955 | 0.0099185 | 0.0061204 | 0.0131633 | 0.0044965 | 0.0014617 | 0.0082394 | 0.0144697 | 0.0049538 |
| 36.28 | 0.0620405 | 0.0315857 | 0.0389459 | 0.0313035 | 0.1359378 | 0.0259723 | 0.0194865 | 0.0844252 | 0.0563945 | 0.0157273 | 0.013674 | 0.0091298 | 0.006271 | 0.0105592 | 0.0036244 | 0.0012432 | 0.0070205 | 0.022658 | 0.0035832 |
| 36.29 | 0.0610672 | 0.0285842 | 0.0323872 | 0.0340033 | 0.1257231 | 0.0272092 | 0.0217927 | 0.0707557 | 0.0464086 | 0.016437 | 0.0146728 | 0.0070674 | 0.0121615 | 0.0105406 | 0.003501 | 0.0055929 | 0.0054822 | 0.0432974 | 0.0038926 |
| 36.31 | 0.0606236 | 0.0279722 | 0.0302124 | 0.0357312 | 0.113566 | 0.0247974 | 0.0224803 | 0.0619415 | 0.0393083 | 0.0184271 | 0.0166829 | 0.0056202 | 0.0154706 | 0.0137286 | 0.0040545 | 0.0086479 | 0.0058628 | 0.0468031 | 0.0036157 |
| 36.32 | 0.0377688 | 0.0196714 | 0.0191832 | 0.0257089 | 0.0548146 | 0.008714 | 0.0156886 | 0.0240155 | 0.0139364 | 0.0142518 | 0.0119796 | 0.0016739 | 0.0126154 | 0.0132619 | 0.0031278 | 0.0077015 | 0.0035836 | 0.0321028 | 0.0028492 |
| 36.33 | 0.0505074 | 0.0342092 | 0.0297827 | 0.0395349 | 0.0467375 | 0.0053969 | 0.0256034 | 0.0146121 | 0.0061084 | 0.0198019 | 0.0155381 | 0.0009869 | 0.0144597 | 0.0210141 | 0.0058326 | 0.0111821 | 0.0033063 | 0.0478325 | 0.0100124 |
| 36.35 | 0.0472453 | 0.0372212 | 0.0305685 | 0.0389055 | 0.0283513 | 0.0059786 | 0.027273 | 0.0156823 | 0.0041164 | 0.0176465 | 0.0142508 | 0.0013492 | 0.0101121 | 0.0195358 | 0.0071794 | 0.0106622 | 0.0033106 | 0.0441225 | 0.0133177 |
| 36.36 | 0.0396284 | 0.0409636 | 0.0307757 | 0.0365074 | 0.0106647 | 0.010429 | 0.0305156 | 0.0385488 | 0.0094706 | 0.0147427 | 0.0127139 | 0.005647 | 0.0087159 | 0.0149956 | 0.0101095 | 0.0098828 | 0.0065961 | 0.0276522 | 0.0151982 |
| 36.38 | 0.0356149 | 0.0415587 | 0.0306431 | 0.0356044 | 0.0147685 | 0.0130582 | 0.0319791 | 0.0500824 | 0.0119714 | 0.0146569 | 0.0120894 | 0.0085069 | 0.0104777 | 0.0130992 | 0.0108611 | 0.0093861 | 0.0087247 | 0.0179037 | 0.0129495 |
| 36.39 | 0.0202834 | 0.0269102 | 0.021138 | 0.0227903 | 0.0181011 | 0.0135156 | 0.0227084 | 0.0405169 | 0.0085586 | 0.0099879 | 0.0070749 | 0.0079914 | 0.0069122 | 0.0073769 | 0.0066962 | 0.0050242 | 0.0074455 | 0.0040564 | 0.0047533 |
| 36.4 | 0.0280418 | 0.0347645 | 0.033386 | 0.032099 | 0.0295368 | 0.0266572 | 0.0337634 | 0.0557881 | 0.0115519 | 0.0141589 | 0.0074313 | 0.0126118 | 0.005929 | 0.0101384 | 0.0069308 | 0.0041412 | 0.0102583 | 0.0063341 | 0.0070606 |
| 36.42 | 0.0278472 | 0.0312301 | 0.0337023 | 0.0313325 | 0.0298549 | 0.0277177 | 0.0328906 | 0.0505051 | 0.0116602 | 0.0139533 | 0.0056923 | 0.012204 | 0.0038839 | 0.0102299 | 0.0055748 | 0.0024684 | 0.0088176 | 0.0093054 | 0.0075557 |
| 36.43 | 0.0198149 | 0.0186751 | 0.0219348 | 0.021228 | 0.0236185 | 0.0175959 | 0.0215234 | 0.0316167 | 0.0107913 | 0.0103135 | 0.0037066 | 0.0079782 | 0.003088 | 0.0080421 | 0.0035352 | 0.0011633 | 0.0047379 | 0.0108758 | 0.0037042 |
| 36.44 | 0.0308343 | 0.0247277 | 0.0306089 | 0.0326502 | 0.0470087 | 0.0270689 | 0.0329908 | 0.0539327 | 0.0258364 | 0.0184529 | 0.0093947 | 0.0137939 | 0.0106033 | 0.0147073 | 0.0057264 | 0.0027799 | 0.0080592 | 0.0259105 | 0.0057693 |

| | | | | | | | | | | | | | | | | | | | |
|---|---|---|---|---|---|---|---|---|---|---|---|---|---|---|---|---|---|---|---|
| 36.46 | 0.0277345 | 0.0199924 | 0.0291101 | 0.0322011 | 0.0525385 | 0.0291757 | 0.0328336 | 0.057679 | 0.0312939 | 0.0197382 | 0.0117806 | 0.0153936 | 0.0140893 | 0.0151246 | 0.0050783 | 0.0027477 | 0.0091417 | 0.0314627 | 0.0061911 |
| 36.47 | 0.0178088 | 0.0108264 | 0.028278 | 0.0307748 | 0.0554319 | 0.0337414 | 0.031883 | 0.059733 | 0.0355701 | 0.0186874 | 0.01243 | 0.0166725 | 0.0158474 | 0.0120763 | 0.0024634 | 0.0013449 | 0.0099711 | 0.0379563 | 0.0043731 |
| 36.49 | 0.0203243 | 0.0191273 | 0.0336034 | 0.0314075 | 0.0489895 | 0.0345874 | 0.0365906 | 0.057253 | 0.0297136 | 0.015627 | 0.0098026 | 0.0153215 | 0.0131408 | 0.009615 | 0.002004 | 0.0008628 | 0.0092305 | 0.0369693 | 0.0046899 |
| 36.5 | 0.0325176 | 0.0523063 | 0.042381 | 0.025602 | 0.0181527 | 0.0212961 | 0.0436711 | 0.0304434 | 0.0123619 | 0.0068352 | 0.0039411 | 0.0078615 | 0.0041531 | 0.0049684 | 0.0019041 | 0.0006669 | 0.0057677 | 0.0210313 | 0.0042888 |
| 36.51 | 0.0999308 | 0.1790922 | 0.0969744 | 0.0608983 | 0.036301 | 0.0228516 | 0.1150881 | 0.0361831 | 0.038797 | 0.0167214 | 0.0267544 | 0.0092589 | 0.0126239 | 0.0134134 | 0.0061772 | 0.0022965 | 0.0212868 | 0.0376883 | 0.0085903 |
| 36.53 | 0.1244816 | 0.2124825 | 0.096393 | 0.0745832 | 0.0604731 | 0.0223894 | 0.1331835 | 0.0305892 | 0.054744 | 0.0297127 | 0.0628254 | 0.0077561 | 0.0325633 | 0.0197196 | 0.0117223 | 0.0034397 | 0.0445445 | 0.0549253 | 0.0170481 |
| 36.54 | 0.1269366 | 0.1763159 | 0.0636664 | 0.069715 | 0.0933483 | 0.0343824 | 0.1121815 | 0.0208475 | 0.067309 | 0.0663177 | 0.1451969 | 0.0190129 | 0.0898572 | 0.0289037 | 0.0243012 | 0.0130817 | 0.1221999 | 0.1083634 | 0.0423887 |
| 36.56 | 0.1031709 | 0.1340862 | 0.0480143 | 0.0523343 | 0.0967347 | 0.0344891 | 0.0855263 | 0.0252671 | 0.0630177 | 0.0817738 | 0.1484679 | 0.0386 | 0.0978385 | 0.0319502 | 0.0225622 | 0.02203 | 0.1454588 | 0.1214621 | 0.0487291 |
| 36.57 | 0.0391589 | 0.0490766 | 0.0187724 | 0.0185052 | 0.0704935 | 0.0173529 | 0.0344778 | 0.0223862 | 0.030611 | 0.0599552 | 0.0623725 | 0.0446706 | 0.0464431 | 0.0277564 | 0.0101676 | 0.0202786 | 0.0793026 | 0.0684522 | 0.0360916 |
| 36.58 | 0.0494772 | 0.0491042 | 0.0178445 | 0.0274439 | 0.1123157 | 0.0252547 | 0.040663 | 0.0344476 | 0.0416892 | 0.0623975 | 0.039352 | 0.0514924 | 0.0435521 | 0.0492838 | 0.0231931 | 0.0234085 | 0.0556169 | 0.0673231 | 0.0567503 |
| 36.6 | 0.0543825 | 0.0472899 | 0.0157797 | 0.0359293 | 0.1156305 | 0.0287259 | 0.0406626 | 0.0348756 | 0.0446907 | 0.0426751 | 0.0249342 | 0.0369933 | 0.0384055 | 0.0470549 | 0.0259868 | 0.0186882 | 0.0328504 | 0.0555111 | 0.0507875 |
| 36.61 | 0.065919 | 0.0516719 | 0.0164743 | 0.0615522 | 0.1182526 | 0.0349213 | 0.0520611 | 0.0205436 | 0.0377821 | 0.0170421 | 0.0193922 | 0.0211683 | 0.0285991 | 0.0319938 | 0.0192284 | 0.0093161 | 0.0170336 | 0.0416009 | 0.0241009 |
| 36.63 | 0.0674164 | 0.0521539 | 0.0179309 | 0.0737939 | 0.1115744 | 0.0331424 | 0.0569041 | 0.0138104 | 0.0256174 | 0.0097272 | 0.0161704 | 0.0176507 | 0.0193121 | 0.0227457 | 0.0140596 | 0.0068368 | 0.0143692 | 0.0303328 | 0.0156487 |
| 36.64 | 0.041786 | 0.0318526 | 0.0150049 | 0.0614261 | 0.0636502 | 0.019098 | 0.0401738 | 0.0149959 | 0.007665 | 0.00187 | 0.0043329 | 0.0089448 | 0.004825 | 0.0064238 | 0.0041082 | 0.0081758 | 0.0077841 | 0.0105505 | 0.0167098 |
| 36.65 | 0.0529211 | 0.0359997 | 0.0310973 | 0.0992269 | 0.0749246 | 0.0288062 | 0.0644075 | 0.0331391 | 0.0243933 | 0.0040201 | 0.0025462 | 0.0098828 | 0.0073886 | 0.0081815 | 0.0044742 | 0.0179151 | 0.0092957 | 0.0273047 | 0.044293 |
| 36.67 | 0.0460825 | 0.0266698 | 0.0362783 | 0.0898586 | 0.0582579 | 0.0253748 | 0.059004 | 0.0310272 | 0.0305245 | 0.0079601 | 0.0029982 | 0.008808 | 0.0081383 | 0.0108174 | 0.0068832 | 0.0164845 | 0.0084106 | 0.0356762 | 0.0470965 |
| 36.68 | 0.0206848 | 0.0078987 | 0.0253685 | 0.032113 | 0.018208 | 0.0108221 | 0.0219526 | 0.0164044 | 0.0147332 | 0.0100465 | 0.0031983 | 0.0063222 | 0.0080063 | 0.0093429 | 0.0098048 | 0.0058092 | 0.0052061 | 0.0301132 | 0.023738 |
| 36.69 | 0.013396 | 0.0111707 | 0.0271532 | 0.0236228 | 0.0177935 | 0.0244884 | 0.0264128 | 0.0347793 | 0.0065689 | 0.0144374 | 0.0047148 | 0.0112632 | 0.0189736 | 0.0138328 | 0.0202222 | 0.0036164 | 0.0071264 | 0.0564053 | 0.0180504 |
| 36.71 | 0.0071345 | 0.0175004 | 0.0206984 | 0.0313938 | 0.0297103 | 0.0348869 | 0.0354457 | 0.051603 | 0.0030954 | 0.0105311 | 0.0036255 | 0.0109247 | 0.017445 | 0.0160786 | 0.019319 | 0.0032216 | 0.0063885 | 0.0631688 | 0.0151662 |
| 36.72 | 0.0083961 | 0.0335298 | 0.0151102 | 0.0534415 | 0.0764282 | 0.0442925 | 0.0401326 | 0.0978612 | 0.0082619 | 0.0059257 | 0.0032364 | 0.0064585 | 0.0106644 | 0.0260146 | 0.0110166 | 0.0030221 | 0.004696 | 0.0741326 | 0.0255293 |
| 36.74 | 0.0118402 | 0.0392399 | 0.016148 | 0.0543097 | 0.0967163 | 0.0420589 | 0.0293138 | 0.1117079 | 0.011904 | 0.0095649 | 0.0043011 | 0.0041075 | 0.0139668 | 0.0303153 | 0.0069186 | 0.0039956 | 0.0041241 | 0.0753998 | 0.029996 |
| 36.75 | 0.008482 | 0.0286983 | 0.0157017 | 0.0293111 | 0.0771747 | 0.0261082 | 0.0076984 | 0.0642043 | 0.0064917 | 0.013143 | 0.0125127 | 0.0020146 | 0.0141015 | 0.0228979 | 0.0023204 | 0.0058 | 0.002216 | 0.0428136 | 0.0190077 |
| 36.76 | 0.006193 | 0.0383592 | 0.03426 | 0.0272458 | 0.1230641 | 0.0437048 | 0.0107133 | 0.0509546 | 0.0060307 | 0.024423 | 0.0374034 | 0.0037328 | 0.0149459 | 0.0328522 | 0.0062792 | 0.0102356 | 0.0031732 | 0.045438 | 0.0155133 |
| 36.78 | 0.0036512 | 0.0332683 | 0.0378668 | 0.0184616 | 0.1249248 | 0.049016 | 0.0138445 | 0.0444537 | 0.0077269 | 0.0241651 | 0.0359156 | 0.0030735 | 0.0105226 | 0.0287063 | 0.0095491 | 0.0080108 | 0.0034683 | 0.0379149 | 0.0099409 |
| 36.79 | 0.002629 | 0.0237015 | 0.0331101 | 0.0087048 | 0.1210583 | 0.0544386 | 0.0187755 | 0.0870479 | 0.0118987 | 0.0194448 | 0.0362785 | 0.0013725 | 0.0125858 | 0.0159716 | 0.0163654 | 0.0037726 | 0.0031609 | 0.0250775 | 0.0128457 |
| 36.81 | 0.0025644 | 0.0212957 | 0.0267252 | 0.0119233 | 0.108067 | 0.0522477 | 0.0186986 | 0.105274 | 0.0121605 | 0.0164618 | 0.0361171 | 0.0021975 | 0.016929 | 0.0102724 | 0.0173621 | 0.0044434 | 0.0036403 | 0.0173449 | 0.0137249 |
| 36.82 | 0.0026951 | 0.0133611 | 0.0109531 | 0.0192732 | 0.0462421 | 0.0308636 | 0.0093167 | 0.0704527 | 0.0072981 | 0.0080003 | 0.0114849 | 0.003943 | 0.0122628 | 0.0030474 | 0.0086563 | 0.0035156 | 0.0047104 | 0.0080851 | 0.0090902 |
| 36.83 | 0.0047013 | 0.0197893 | 0.0081876 | 0.0474714 | 0.0404562 | 0.0436327 | 0.0076723 | 0.0898899 | 0.0154792 | 0.0091639 | 0.0072485 | 0.0088253 | 0.0105465 | 0.0039993 | 0.0058519 | 0.0028302 | 0.0089744 | 0.0282486 | 0.021104 |
| 36.85 | 0.0049886 | 0.0204984 | 0.0081681 | 0.0552192 | 0.0305469 | 0.0436119 | 0.0076475 | 0.0773864 | 0.0242338 | 0.0086935 | 0.0077739 | 0.0092617 | 0.0082897 | 0.0067474 | 0.0051923 | 0.0016581 | 0.008382 | 0.0397317 | 0.0215004 |
| 36.86 | 0.0076601 | 0.0262116 | 0.0223506 | 0.0619384 | 0.0137516 | 0.0398348 | 0.0168977 | 0.0380134 | 0.0676133 | 0.0123683 | 0.0104076 | 0.0085846 | 0.0084825 | 0.0192792 | 0.0149076 | 0.0013301 | 0.0056195 | 0.0547496 | 0.013301 |
| 36.88 | 0.0078677 | 0.0305768 | 0.0273533 | 0.0604296 | 0.0080061 | 0.0357543 | 0.0242228 | 0.028474 | 0.0921101 | 0.0164472 | 0.015523 | 0.0084228 | 0.0081884 | 0.0250155 | 0.0203486 | 0.0016949 | 0.0045625 | 0.0570442 | 0.011105 |
| 36.89 | 0.0072894 | 0.0241986 | 0.0195036 | 0.0336229 | 0.0066681 | 0.0204912 | 0.0262855 | 0.0385108 | 0.0694643 | 0.015713 | 0.0242198 | 0.0057938 | 0.0097124 | 0.0194451 | 0.015176 | 0.0023617 | 0.0026163 | 0.0342764 | 0.0064446 |
| 36.9 | 0.0228934 | 0.0341775 | 0.0362125 | 0.0358254 | 0.0289539 | 0.0379618 | 0.0515148 | 0.1035477 | 0.0861341 | 0.0258202 | 0.0467662 | 0.0087987 | 0.028577 | 0.0274295 | 0.0162922 | 0.0109663 | 0.0045601 | 0.0373752 | 0.0063443 |
| 36.92 | 0.0275454 | 0.0291146 | 0.0437172 | 0.0291605 | 0.0444299 | 0.0480532 | 0.0549858 | 0.1221644 | 0.0751066 | 0.0248024 | 0.0440804 | 0.008631 | 0.033673 | 0.0250385 | 0.0112079 | 0.0162613 | 0.0055506 | 0.0275517 | 0.0051612 |
| 36.93 | 0.0177226 | 0.0109151 | 0.0375921 | 0.0158903 | 0.0447825 | 0.0409118 | 0.0375662 | 0.1017449 | 0.0385268 | 0.0141067 | 0.0295338 | 0.0052794 | 0.0237776 | 0.0124228 | 0.0031217 | 0.0145344 | 0.0051043 | 0.0074228 | 0.0048082 |
| 36.94 | 0.0171606 | 0.0078623 | 0.0542097 | 0.0229213 | 0.069651 | 0.0530622 | 0.0513398 | 0.1765286 | 0.0435494 | 0.017856 | 0.0351794 | 0.0065622 | 0.0328736 | 0.0101856 | 0.0035859 | 0.0191091 | 0.0090502 | 0.0042153 | 0.009145 |
| 36.96 | 0.0121054 | 0.0095819 | 0.0454768 | 0.022044 | 0.0640519 | 0.043204 | 0.0449433 | 0.1736226 | 0.0360776 | 0.0163586 | 0.0217906 | 0.005566 | 0.0289036 | 0.0059826 | 0.0043616 | 0.0150777 | 0.0092629 | 0.0053931 | 0.0084992 |
| 36.97 | 0.0141901 | 0.0239372 | 0.0202623 | 0.0153377 | 0.04835 | 0.0235703 | 0.0315152 | 0.1124413 | 0.0203482 | 0.0133247 | 0.0063137 | 0.0060953 | 0.0162088 | 0.0012737 | 0.0056818 | 0.007372 | 0.0086607 | 0.0111743 | 0.0153278 |
| 36.99 | 0.0166275 | 0.0249001 | 0.0130057 | 0.0204923 | 0.0432155 | 0.0152296 | 0.0399909 | 0.0710068 | 0.0207631 | 0.0113843 | 0.0055809 | 0.0096881 | 0.0105479 | 0.0008313 | 0.0052553 | 0.0058746 | 0.0083675 | 0.0137953 | 0.0242622 |
