## Supplemental Table S7 for "Leaf movements as a quantitative metric for early stress detection"

**Supplementary Table S7.** Quantification of 100 mM NaCl stress-induced leaf movement dynamics in **(a-d)** lettuce, **(e-h)** Amaranth, **(i-l)** Tomato plant arrays and the respective controls. **(a,e,i)** Direct motion under RGBday dimGnight lighting conditions **(b,f,j)** Direct motion under dimG day/night lighting conditions **(c,g,k)** Mean day-to-day motion rate in measured under dimGday/night and RGBday dimGnight, calculated using either 24 h motion integration or 22 h motion integration excluding day/night transitions. **(d,h,l)** Mean day-to-day motion rate of individual plants under dimGday/night and RGBday dimGnight. Data represent means of 9–10 plants.

**Supplementary Table S7 (a)** Direct motion in 100 mM NaCl stressed lettuce and their respective controls under RGBday dimGnight lighting condition

| Time | RGBControl1 | RGBControl2 | RGBControl3 | RGBControl4 | RGBControl5 | RGBControl6 | RGBControl7 | RGBControl8 | RGBControl9 | RGBControlnoplant | RGBControltank | RGBStress1 |
|---|---|---|---|---|---|---|---|---|---|---|---|---|
| 24.0069 | -0.329619 | -1.522532 | -1.194629 | -0.950083 | -0.299372 | -1.593943 | -0.329171 | 0.084531 | -0.426306 | -0.016431 | -0.849782 | -2.159448 |
| 24.0208 | -0.182888 | -1.029178 | -0.795705 | -0.550584 | -0.265909 | -0.805003 | -0.255711 | 0.024104 | -0.230874 | -0.059594 | -0.443154 | -0.92932 |
| 24.0347 | 0.084475 | -0.568318 | -0.365886 | -0.123747 | -0.111088 | -0.057082 | -0.027511 | -0.06597 | -0.020736 | -0.029984 | -0.122628 | -0.308167 |
| 24.0486 | 0.559122 | -0.165689 | -0.135156 | 0.386158 | 0.008925 | 0.368591 | 0.156402 | -0.190097 | 0.041936 | 0.058521 | 0.0833 | -0.393779 |
| 24.0625 | 1.07447 | 0.355972 | -0.014746 | 0.842416 | 0.165868 | 0.678065 | 0.15575 | -0.400421 | -0.154678 | 0.531586 | 0.376187 | -1.379117 |
| 24.0764 | 1.838242 | 1.528063 | 0.22447 | 1.367131 | 0.83429 | 1.318769 | 0.225072 | -0.253539 | -0.092186 | 2.572286 | 1.147689 | -2.923172 |
| 24.0903 | 2.675608 | 2.997102 | 0.245818 | 1.864147 | 1.917017 | 2.277775 | 0.487711 | 0.323246 | 0.464785 | 6.245913 | 2.011624 | -2.194534 |
| 24.1042 | 2.316379 | 2.737055 | 0.171586 | 1.850151 | 2.041176 | 2.971455 | 0.505664 | 0.808175 | 1.252712 | 4.137238 | 2.125608 | 0.123932 |
| 24.1181 | 1.031037 | 1.123699 | 0.305707 | 1.213923 | 1.056346 | 2.172413 | 0.371435 | 0.711632 | 1.455397 | 0.786846 | 1.326946 | 0.931431 |
| 24.1319 | 0.313906 | 0.287936 | 0.407853 | 0.561913 | 0.317282 | 1.056025 | 0.327277 | 0.336987 | 0.802229 | 0.108939 | 0.68041 | 0.457231 |
| 24.1458 | 0.226764 | 0.229979 | 0.257145 | 0.10495 | 0.120488 | 0.700708 | 0.199305 | 0.173137 | 0.465563 | 0.052657 | 0.476525 | 0.008442 |
| 24.1597 | 0.244096 | 0.223632 | -0.017054 | -0.244202 | 0.08926 | 0.606629 | 0.002697 | 0.129926 | 0.243133 | 0.042455 | 0.427845 | -0.164522 |
| 24.1736 | 0.412447 | 0.187211 | -0.116037 | -0.250777 | 0.128523 | 0.480472 | -0.047585 | 0.088214 | 0.090026 | -0.030275 | 0.340443 | -0.257262 |
| 24.1875 | 0.6723 | 0.214973 | 0.046864 | 0.064518 | 0.193239 | 0.326954 | 0.076026 | 0.05788 | 0.140708 | -0.06712 | 0.196912 | -0.293743 |
| 24.2014 | 0.699515 | 0.24164 | 0.259552 | 0.331023 | 0.16583 | 0.072186 | 0.196862 | 0.048021 | 0.305003 | -0.084309 | -0.007294 | -0.261678 |
| 24.2153 | 0.450814 | 0.173529 | 0.190811 | 0.295696 | 0.089712 | -0.301479 | 0.137709 | 0.063659 | 0.373378 | -0.066257 | -0.19924 | -0.169787 |
| 24.2292 | 0.009068 | -0.003472 | -0.109976 | 0.014931 | 0.01532 | -0.62152 | -0.034519 | 0.042926 | 0.275711 | -0.055689 | -0.32358 | -0.10549 |
| 24.2431 | -0.406812 | -0.177383 | -0.396399 | -0.334039 | -0.121169 | -0.787143 | -0.24329 | -0.013832 | 0.097752 | -0.089427 | -0.35517 | -0.139978 |
| 24.2569 | -0.529054 | -0.191009 | -0.390884 | -0.476095 | -0.228325 | -0.677261 | -0.355978 | 0.004045 | 0.043417 | -0.094193 | -0.297602 | -0.208962 |
| 24.2708 | -0.229489 | -0.049525 | -0.096879 | -0.239687 | -0.192737 | -0.579346 | -0.272638 | 0.095027 | 0.168848 | -0.066996 | -0.19048 | -0.233236 |
| 24.2847 | 0.29947 | 0.129161 | 0.207642 | 0.172378 | -0.137361 | -0.276099 | -0.062079 | 0.12092 | 0.28528 | -0.077618 | -0.050128 | -0.258567 |
| 24.2986 | 0.729468 | 0.232354 | 0.336468 | 0.579241 | -0.082947 | 0.155542 | 0.122942 | 0.060944 | 0.243737 | -0.091698 | 0.099778 | -0.315444 |
| 24.3125 | 0.828339 | 0.226694 | 0.287108 | 0.718048 | 0.004713 | 0.463313 | 0.234491 | 0.046104 | 0.123483 | -0.076323 | 0.270602 | -0.329855 |
| 24.3264 | 0.53164 | 0.105156 | 0.096034 | 0.588428 | 0.023005 | 0.573677 | 0.240418 | 0.135821 | 0.066328 | -0.07244 | 0.450872 | -0.208108 |
| 24.3403 | 0.415305 | -0.080454 | 0.042021 | 0.528851 | -0.037212 | 0.718056 | 0.250461 | 0.434717 | 0.25411 | -0.124924 | 0.608189 | 0.001082 |
| 24.3542 | 0.849783 | -0.050713 | 0.411441 | 0.76982 | -0.067905 | 0.397357 | 0.324851 | 0.970932 | 0.821592 | -0.142456 | 0.621061 | 0.270693 |
| 24.3681 | 0.963525 | 0.267773 | 0.873207 | 0.499829 | -0.030526 | 0.196851 | 0.437899 | 0.937995 | 1.128372 | -0.019261 | 0.394099 | 0.433589 |
| 24.3819 | 1.262091 | 0.513885 | 0.919304 | 0.684634 | 0.024777 | -0.076456 | 0.491642 | 0.568649 | 0.89877 | 0.042514 | 0.131739 | 0.425808 |
| 24.3958 | 0.897547 | 0.4083 | 0.503844 | 0.687286 | 0.021167 | -0.426802 | 0.377954 | 0.183446 | 0.400848 | 0.085732 | -0.107774 | 0.39563 |
| 24.4097 | 0.12906 | 0.103205 | 0.009483 | 0.294332 | -0.00366 | -0.61753 | 0.139795 | -0.080096 | -0.044777 | 0.090883 | -0.189551 | 0.279038 |
| 24.4236 | -0.068006 | 0.160175 | -0.151532 | -0.148334 | -0.001794 | -0.265985 | 0.025706 | -0.166156 | -0.123975 | 0.051771 | 0.110556 | 0.147859 |
| 24.4375 | 0.901685 | 1.501212 | -0.137486 | -0.445135 | -0.110466 | 0.655626 | 0.203964 | -0.121619 | 0.657823 | 0.000376 | 1.438168 | 0.187783 |
| 24.4514 | 2.896407 | 3.614088 | 0.014195 | -0.446458 | 0.021292 | 1.627179 | 0.567985 | 0.454496 | 2.026799 | -0.169315 | 4.797808 | 0.433108 |
| 24.4653 | 3.932692 | 4.142684 | 0.881251 | 0.136111 | 0.049351 | 1.805691 | 0.606826 | 2.383045 | 2.552956 | 1.694308 | 7.756726 | 1.159967 |
| 24.4792 | 3.255 | 3.085943 | 2.404454 | 1.431045 | 0.322986 | 1.403878 | 1.121616 | 5.28167 | 3.011769 | 1.637958 | 3.854534 | 1.357258 |
| 24.4931 | 2.121942 | 2.168577 | 2.4099 | 1.877731 | 0.459767 | 0.263473 | 0.92637 | 6.267035 | 3.499927 | -0.614886 | 0.72227 | 0.840832 |
| 24.5 | 1.525606 | 1.690997 | 1.525858 | 1.423272 | 0.475325 | -0.121538 | 0.555734 | 4.964117 | 2.774143 | -0.286785 | 1.248236 | 0.520417 |
| 24.5139 | 0.537652 | 1.260037 | 0.966465 | 0.499618 | 0.466485 | -0.377309 | 0.360036 | 2.357789 | 1.433037 | 0.244879 | 1.978703 | 0.357801 |
| 24.5278 | -0.883405 | 0.572073 | 0.372952 | -0.316832 | 0.446338 | -0.743161 | 0.102781 | 0.484741 | 0.35433 | 0.35168 | 1.284942 | 0.246174 |
| 24.5417 | -1.943064 | 0.155805 | -0.120018 | -0.954545 | 0.356849 | -1.119547 | -0.219792 | -1.051494 | -0.699155 | 0.137919 | -0.482557 | 0.166576 |
| 24.5556 | -1.932192 | 0.369407 | -0.487188 | -1.354264 | 0.393169 | -0.953283 | -0.367953 | -2.668877 | -1.519867 | 0.147545 | -0.971515 | 0.115846 |
| 24.5694 | -0.921547 | 0.692274 | -0.486429 | -1.394641 | 0.578955 | -0.178942 | -0.288072 | -4.627571 | -2.312634 | 0.180857 | -0.663143 | 0.107163 |
| 24.5833 | 0.142799 | 0.776588 | -0.1605 | -0.552872 | 0.939982 | 0.550465 | -0.027916 | -5.501867 | -2.36356 | -0.075833 | -0.851213 | 0.157789 |
| 24.5972 | 1.519901 | 0.790106 | 0.289722 | 0.463682 | 1.095528 | 0.613386 | 0.036542 | -4.512274 | -1.38781 | -0.392854 | -1.090948 | 0.216488 |
| 24.6111 | 2.203613 | 0.875554 | 0.630331 | 0.577917 | 0.764265 | 0.2768 | 0.222334 | -1.435806 | -0.20158 | -0.45092 | -0.997345 | 0.232435 |
| 24.625 | 1.415069 | 0.9299 | 0.598957 | 0.051062 | 0.278078 | -0.197207 | 0.116637 | 1.613226 | 0.649089 | -0.02888 | 0.712172 | 0.21173 |
| 24.6389 | 0.43634 | 0.684975 | 0.69447 | -0.384895 | -0.040187 | -0.448567 | 0.126677 | 3.848174 | 1.277044 | 0.427044 | 2.773693 | 0.187796 |
| 24.6528 | -0.518623 | 0.228651 | 1.032723 | -0.58803 | -0.189693 | -0.666996 | 0.202312 | 4.569364 | 1.871749 | 0.341852 | 2.323296 | 0.172221 |
| 24.6667 | -1.666626 | -0.142365 | 0.647083 | -0.661312 | -0.213005 | -1.098458 | -0.26703 | 4.14809 | 2.034364 | -0.133688 | -0.125532 | 0.161405 |
| 24.6806 | -2.43343 | -0.133698 | 0.394991 | -0.751083 | -0.209116 | -0.857436 | -0.228182 | 1.453157 | 0.904435 | -0.352851 | -1.612105 | 0.149166 |
| 24.6944 | -1.685317 | 0.581765 | 0.109847 | -0.788639 | 0.012598 | 0.059013 | 0.096581 | -1.281025 | -0.346931 | -0.082575 | -1.191516 | 0.159741 |
| 24.7083 | -0.91891 | 1.230609 | -0.115318 | -0.487126 | 0.076428 | 0.771863 | 0.449873 | -3.818686 | -1.680594 | 0.331429 | -0.15548 | 0.193227 |
| 24.7222 | -0.073174 | 2.461897 | -0.411312 | -0.219274 | 0.067713 | 0.741326 | 0.390507 | -5.170671 | -2.198486 | 0.433932 | 0.38398 | 0.212145 |
| 24.7361 | -0.064766 | 1.878733 | -0.636857 | -0.165113 | -0.01762 | 0.003066 | -0.202925 | -4.411953 | -1.648737 | 0.115146 | -0.569296 | 0.172122 |
| 24.75 | -0.085701 | 0.092789 | -0.820885 | 0.09325 | -0.124327 | -0.87859 | -0.581852 | -2.619803 | -0.917541 | -0.28923 | -2.918578 | 0.117627 |
| 24.7639 | -0.137506 | -1.498942 | -0.653383 | 0.267575 | -0.256074 | -1.413021 | -0.879102 | -1.081324 | -0.611905 | -0.402504 | -3.962773 | 0.079585 |
| 24.7778 | -0.557892 | -2.25254 | -0.21679 | -0.07162 | -0.336463 | -1.838865 | -0.923279 | -0.352129 | -0.387039 | -0.075539 | -2.717731 | 0.058171 |
| 24.7917 | -1.407531 | -2.018848 | 0.273672 | -0.792693 | -0.397139 | -1.833108 | -0.661058 | 0.023969 | -0.067059 | 0.436314 | -0.710398 | 0.056152 |
| 24.8056 | -2.920522 | -1.581975 | 0.417051 | -1.880286 | -0.543758 | -1.847443 | -0.299496 | 0.092056 | 0.123 | 0.449055 | -0.291269 | 0.102475 |
| 24.8194 | -3.967496 | -1.534067 | 0.234395 | -2.581186 | -1.137398 | -1.557613 | -0.076204 | -0.025749 | 0.099772 | 0.05213 | -0.816753 | 0.182132 |
| 24.8333 | -3.572491 | -1.266816 | -0.346142 | -2.400292 | -1.556598 | -0.952506 | -0.034861 | -0.141875 | -0.017923 | 0.080152 | -0.584046 | 0.249186 |
| 24.8472 | -2.787388 | -1.06799 | -0.709192 | -1.703924 | -1.776655 | -0.299128 | -0.16446 | -0.076324 | -0.153064 | 0.246353 | -0.084852 | 0.256019 |
| 24.8611 | -1.695665 | -0.926575 | -0.734342 | -1.181227 | -1.608403 | 0.218354 | -0.235229 | 0.087324 | -0.24265 | 0.17902 | 0.280366 | 0.172548 |
| 24.875 | -0.76203 | -0.712631 | -0.603727 | -0.351568 | -0.912816 | 0.824537 | -0.154323 | 0.22897 | -0.22409 | 0.091424 | 0.675298 | 0.052347 |
| 24.8889 | 0.208532 | -0.496721 | -0.517027 | 0.579906 | -0.186746 | 1.231493 | 0.010379 | 0.281416 | -0.137931 | -0.229031 | 0.480621 | 0.015566 |
| 24.9028 | 0.539404 | -0.369556 | -0.542444 | 0.948274 | 0.257545 | 1.13879 | 0.11063 | 0.199117 | 0.01341 | -0.64107 | -0.385591 | 0.092015 |
| 24.9167 | 0.437501 | -0.258221 | -0.677344 | 0.785727 | 0.257651 | 1.005323 | 0.141812 | 0.142291 | 0.129017 | -0.687956 | -0.552821 | 0.177164 |
| 24.9306 | -0.461365 | -0.605286 | -0.843678 | 0.493029 | -0.261743 | 0.619879 | 0.08522 | -0.020003 | 0.0157 | -2.062765 | -2.390349 | -0.245148 |
| 24.9444 | -2.506135 | -1.997398 | -1.171783 | -0.183897 | -1.109993 | -0.716309 | -0.225406 | -0.469591 | -0.642824 | 0.47217 | -6.570839 | -1.248997 |
| 24.9583 | -3.992522 | -2.453937 | -1.120023 | -1.797699 | -1.422424 | -2.415345 | -0.925144 | -0.879503 | -1.479476 | -0.850361 | -6.550688 | -1.419694 |
| 24.9722 | -3.14426 | -1.366773 | -0.871133 | -2.213502 | -0.836748 | -2.967356 | -1.304354 | -0.580484 | -1.508846 | -1.135988 | -3.646739 | -1.477067 |
| 24.9861 | -1.555724 | -0.588008 | -0.598775 | -1.258992 | -0.410012 | -3.032924 | -1.150605 | -0.099991 | -1.128168 | -0.248929 | -1.517891 | -0.501075 |
| 25.0069 | -0.758797 | -0.41349 | -0.327826 | -0.648699 | -0.289841 | -1.957975 | -0.745356 | 0.236184 | -0.541503 | -0.087422 | -0.648315 | 0.291583 |
| 25.0208 | -0.037153 | -0.231041 | -0.049291 | -0.295011 | -0.138266 | -0.676846 | -0.331402 | 0.35049 | -0.070994 | 0.010256 | -0.117875 | 0.548559 |
| 25.0347 | 0.542301 | -0.1174 | 0.152791 | -0.040644 | 0.005248 | -0.105312 | 0.047162 | 0.178211 | 0.197839 | -0.014368 | 0.078556 | 0.558698 |
| 25.0486 | 0.734148 | -0.308642 | 0.230538 | 0.120896 | 0.079546 | -0.129342 | 0.277394 | -0.038867 | 0.340419 | -0.251619 | -0.107672 | 0.516936 |
| 25.0625 | 0.598954 | -0.570607 | 0.216053 | 0.335848 | 0.045763 | -0.263809 | 0.390347 | -0.204361 | 0.469785 | -0.346985 | -0.36646 | 0.530583 |
| 25.0764 | 0.587771 | -0.385784 | 0.288337 | 0.672985 | 0.116166 | -0.056934 | 0.455372 | -0.148957 | 0.514767 | -0.207668 | -0.310906 | 0.476253 |
| 25.0903 | 0.588082 | -0.057405 | 0.286732 | 1.008315 | 0.126578 | 0.262641 | 0.434012 | -0.006837 | 0.382222 | -0.128172 | 0.058179 | 0.252576 |
| 25.1042 | 0.337083 | 0.079718 | 0.168936 | 1.084632 | 0.022638 | 0.418105 | 0.264533 | 0.059331 | 0.118664 | -0.019203 | 0.241717 | 0.025376 |
| 25.1181 | 0.276556 | 0.092509 | -0.064507 | 0.937185 | -0.033468 | 0.588515 | 0.082792 | 0.064491 | -0.043278 | 0.00725 | 0.340928 | -0.04862 |
| 25.1319 | 0.735509 | 0.174842 | -0.156908 | 0.822944 | 0.140895 | 1.013059 | 0.060801 | 0.114929 | 0.179368 | -0.018755 | 0.701255 | 0.077267 |
| 25.1458 | 1.457932 | 0.499834 | 0.094743 | 0.846532 | 0.530197 | 1.510253 | 0.283074 | 0.287801 | 0.71153 | 0.036632 | 1.418081 | 0.249189 |
| 25.1597 | 1.914867 | 0.806968 | 0.418444 | 0.929589 | 0.908084 | 1.784179 | 0.537102 | 0.431476 | 1.131083 | 0.201615 | 2.078999 | 0.32361 |
| 25.1736 | 1.343709 | 0.531529 | 0.412702 | 0.803982 | 0.797842 | 1.421949 | 0.477023 | 0.31103 | 0.932973 | 0.163546 | 1.652432 | 0.34188 |
| 25.1875 | 0.211464 | 0.042338 | 0.187584 | 0.331825 | 0.235041 | 0.908378 | 0.211255 | 0.070052 | 0.375246 | -0.056685 | 0.494809 | 0.360652 |
| 25.2014 | -0.688542 | -0.268126 | -0.059929 | -0.252486 | -0.30323 | 0.423704 | -0.043849 | -0.076904 | -0.049331 | -0.152025 | -0.482919 | 0.328331 |
| 25.2153 | -0.773461 | -0.384606 | -0.21812 | -0.558104 | -0.480674 | 0.188542 | -0.178896 | -0.071134 | -0.15068 | -0.14056 | -0.780772 | 0.339435 |
| 25.2292 | -0.13472 | -0.155581 | -0.099856 | -0.26645 | -0.262736 | 0.17944 | -0.040945 | 0.146139 | 0.175816 | -0.054952 | -0.264501 | 0.375719 |
| 25.2431 | 0.67243 | 0.203666 | 0.20781 | 0.318101 | -0.000969 | 0.253239 | 0.257707 | 0.330442 | 0.60504 | 0.052487 | 0.495506 | 0.387419 |
| 25.2569 | 1.082442 | 0.391443 | 0.418586 | 0.718288 | 0.215279 | 0.192107 | 0.448358 | 0.34929 | 0.755458 | 0.152674 | 0.810007 | 0.341145 |
| 25.2708 | 0.861721 | 0.170017 | 0.33881 | 0.638936 | 0.30133 | -0.116663 | 0.413497 | 0.163727 | 0.50746 | 0.066426 | 0.295892 | 0.328199 |
| 25.2847 | 0.223231 | -0.171798 | 0.070424 | 0.244179 | 0.137421 | -0.517728 | 0.240396 | -0.030552 | 0.131197 | -0.100721 | -0.519122 | 0.425397 |
| 25.2986 | -0.149233 | -0.132225 | -0.107239 | -0.033974 | -0.100411 | -0.573836 | 0.119345 | -0.028249 | -0.007363 | -0.086745 | -0.606618 | 0.567445 |
| 25.3125 | 0.085587 | 0.170302 | -0.084047 | 0.087512 | -0.098439 | -0.253944 | 0.136265 | 0.134616 | 0.255812 | -0.021782 | 0.103343 | 0.714049 |
| 25.3264 | 0.477661 | 0.392876 | 0.110872 | 0.31947 | 0.094725 | 0.052499 | 0.223932 | 0.282086 | 0.683378 | 0.02814 | 0.665509 | 0.620334 |
| 25.3403 | 0.580344 | 0.286315 | 0.290458 | 0.441501 | 0.2824 | 0.079278 | 0.275901 | 0.248184 | 0.79936 | -0.007327 | 0.561951 | 0.407068 |
| 25.3542 | 0.472263 | 0.216981 | 0.410524 | 0.416707 | 0.487312 | 0.097096 | 0.311481 | 0.215465 | 0.627512 | 0.011862 | 0.552432 | 0.352079 |
| 25.3681 | 0.318704 | 0.443174 | 0.500199 | 0.215789 | 0.591795 | 0.23409 | 0.37624 | 0.276251 | 0.533104 | 0.044635 | 0.866802 | 0.491068 |
| 25.3819 | 0.075571 | 0.320633 | 0.297004 | 0.025071 | 0.394939 | 0.26962 | 0.285702 | 0.175324 | 0.082366 | 0.081673 | 0.367722 | 0.703277 |
| 25.3958 | -0.045946 | -0.259959 | -0.04935 | 0.069714 | 0.023137 | 0.212072 | 0.102267 | -0.079942 | -0.569279 | -0.132846 | -0.606251 | 0.846999 |
| 25.4097 | 0.173455 | -0.36781 | -0.237818 | 0.275625 | -0.180224 | 0.383833 | 0.069838 | -0.153601 | -0.418886 | -0.18384 | -0.516828 | 0.729654 |
| 25.4236 | 0.557054 | 0.057204 | -0.041736 | 0.432857 | -0.153615 | 0.66783 | 0.205472 | 0.044618 | 0.402392 | 0.028178 | 0.591784 | 0.418256 |
| 25.4375 | 0.906928 | 1.029793 | 0.430371 | 0.178208 | 0.0374 | 1.017709 | 0.516777 | 0.33458 | 1.363569 | 0.862338 | 3.056671 | 0.356708 |
| 25.4514 | 1.688822 | 2.59106 | 0.988474 | -0.286974 | 0.536874 | 1.711187 | 0.845713 | 0.941302 | 2.452527 | 1.703132 | 7.705987 | 0.841485 |
| 25.4653 | 2.456602 | 3.291118 | 1.411046 | -0.171078 | 1.135856 | 2.107298 | 0.941639 | 1.124911 | 2.568529 | -0.211418 | 9.127565 | 2.134546 |
| 25.4792 | 2.758401 | 2.397412 | 2.356929 | 1.052458 | 1.365017 | 1.778017 | 1.229409 | 0.618847 | 1.60488 | 1.364791 | 5.828229 | 2.400224 |
| 25.4931 | 2.613381 | 2.040203 | 2.643409 | 1.840299 | 0.836443 | 1.188726 | 1.130452 | 0.721122 | 0.846037 | 0.219223 | 2.200577 | 1.757364 |
| 25.5 | 1.585759 | 2.04212 | 1.98371 | 1.450296 | 0.315823 | 0.731895 | 0.910804 | 1.3267 | 0.653234 | -0.018379 | 1.322904 | 1.310821 |
| 25.5139 | 0.03259 | 1.378856 | 0.933613 | 0.211758 | -0.211416 | 0.017297 | 0.291966 | 1.60753 | 0.3065 | 0.024956 | 0.431553 | 0.586237 |
| 25.5278 | -1.757958 | 0.79119 | -0.02063 | -1.200054 | -0.90959 | -0.750976 | -0.230188 | 1.632759 | -0.152263 | 0.030573 | -0.302161 | -0.210571 |
| 25.5417 | -2.337213 | 0.476762 | -0.854697 | -1.746013 | -1.078683 | -1.180485 | -0.474322 | 1.710194 | -0.435178 | 0.016393 | -0.42987 | -0.743341 |
| 25.5556 | -1.950556 | 0.127058 | -1.226547 | -1.701166 | -0.758104 | -0.931658 | -0.376627 | 1.714094 | -0.305863 | -0.001812 | -0.025653 | -0.753967 |
| 25.5694 | -1.495537 | -0.501259 | -0.594838 | -1.711541 | -0.358431 | -0.887126 | -0.081104 | 1.360976 | 0.162845 | -0.03623 | 0.331406 | -0.535707 |
| 25.5833 | -0.174862 | -1.111067 | -0.016459 | -0.611111 | 0.079566 | -0.798264 | 0.041275 | 0.982618 | 0.897096 | -0.053977 | 0.547121 | -0.366717 |
| 25.5972 | 1.382683 | -1.511842 | 0.149768 | 0.600003 | 0.465082 | -0.310947 | 0.104811 | -0.305459 | 1.262923 | -0.01835 | 0.36867 | -0.25353 |
| 25.6111 | 2.226708 | -1.723255 | 0.477115 | 1.284704 | 0.734293 | -0.053201 | 0.050998 | -1.567581 | 0.856686 | 0.010083 | 0.028635 | -0.363916 |
| 25.625 | 1.356588 | -1.284189 | -0.077009 | 0.888052 | 0.667966 | 0.463153 | -0.03498 | -2.114274 | -0.001454 | 0.018726 | -0.030197 | -0.714154 |
| 25.6389 | -0.086991 | -0.551506 | -0.896662 | -0.135234 | 0.297383 | 0.130657 | -0.101179 | -2.369232 | -0.685455 | 0.049729 | -0.642314 | -0.90208 |
| 25.6528 | -1.152136 | 0.14423 | -1.147642 | -0.593357 | -0.036746 | 0.01042 | 0.007988 | -2.039514 | -1.071649 | 0.086713 | -1.688884 | -0.729917 |
| 25.6667 | -2.068568 | 0.920606 | -0.579914 | -1.107299 | -0.277617 | -0.280154 | 0.082102 | -1.122885 | -1.185363 | 0.050119 | -2.115257 | -0.080919 |
| 25.6806 | -2.689025 | 1.748553 | 0.083695 | -1.631121 | -0.948464 | -0.910694 | 0.060801 | -0.393622 | -0.864681 | -0.032611 | -1.480796 | 0.510131 |
| 25.6944 | -2.327241 | 1.891364 | 0.582013 | -1.72116 | -1.260404 | -0.852052 | 0.28127 | 0.190357 | -0.079605 | -0.039408 | 0.012667 | 0.923157 |
| 25.7083 | -0.606713 | 1.571819 | 0.712489 | -1.039327 | -0.597692 | -1.598598 | 0.279464 | 0.613925 | 0.688542 | 0.010801 | 0.965109 | 1.025939 |
| 25.7222 | 0.664145 | 0.832351 | 0.366444 | -0.581908 | 0.137907 | -1.701552 | 0.160744 | 0.694837 | 1.096537 | 0.040925 | 1.147518 | 0.824749 |
| 25.7361 | 1.283243 | 0.041913 | -0.326255 | -0.278919 | 0.497679 | -1.256228 | -0.067866 | 0.375624 | 1.002178 | 0.051074 | 0.555177 | 0.45715 |
| 25.75 | 0.602953 | -0.790942 | -0.766271 | -0.343623 | 0.839484 | -0.789978 | -0.104072 | 0.008977 | 0.487922 | 0.045114 | -0.246187 | -0.051754 |
| 25.7639 | -0.369614 | -1.577928 | -0.853861 | -0.702531 | 0.737481 | -0.747138 | -0.556423 | -0.385686 | -0.253852 | 0.028096 | -1.429899 | -0.617593 |
| 25.7778 | -0.738377 | -1.978177 | -1.190611 | -0.746415 | 0.621102 | -1.097973 | -1.219576 | -0.650908 | -1.127563 | -0.007538 | -2.545313 | -1.044848 |
| 25.7917 | -0.764297 | -1.33694 | -1.163274 | -0.683835 | 0.265709 | -0.993135 | -1.011391 | -0.650462 | -1.625079 | -0.032757 | -2.666984 | -1.306231 |
| 25.8056 | -0.947654 | -0.540853 | -0.988335 | -0.557039 | -0.154472 | -0.71911 | -0.555502 | -0.314204 | -1.302611 | -0.006768 | -1.958123 | -1.444827 |
| 25.8194 | -1.246881 | -0.156394 | -1.052292 | -0.514819 | -0.871424 | -0.753502 | -0.401313 | -0.0344 | -0.687774 | 0.010464 | -1.513557 | -1.445863 |
| 25.8333 | -1.237898 | 0.031406 | -0.942351 | -0.687493 | -1.88707 | -0.719132 | -0.399958 | 0.070931 | -0.08719 | -0.003492 | -1.15949 | -1.300438 |
| 25.8472 | -0.904042 | 0.182859 | -0.408757 | -0.822067 | -1.279504 | -0.492188 | -0.45184 | 0.066272 | 0.30515 | -0.008291 | -0.676751 | -1.033094 |
| 25.8611 | -0.70597 | 0.31591 | -0.047688 | -1.01778 | -0.491355 | -0.031619 | -0.407693 | 0.009618 | 0.367269 | -0.016979 | -0.217546 | -0.621519 |
| 25.875 | -0.675358 | 0.403899 | 0.284656 | -0.978834 | 0.310025 | 0.558551 | -0.188339 | -0.075496 | 0.244734 | -0.023009 | 0.141694 | -0.195964 |
| 25.8889 | -0.472277 | 0.392175 | 0.37219 | -0.545576 | 0.792564 | 1.086675 | 0.050378 | -0.178345 | 0.154678 | 0.035153 | 0.311123 | 0.164483 |
| 25.9028 | 0.014907 | 0.374775 | 0.226815 | -0.096733 | 0.442375 | 1.362036 | 0.198839 | -0.243691 | 0.208469 | 0.071205 | 0.416312 | 0.320082 |
| 25.9167 | 0.499078 | 0.275267 | 0.069979 | 0.30834 | 0.301363 | 1.19096 | 0.196681 | -0.306811 | 0.271462 | -0.207556 | 0.260529 | 0.175199 |
| 25.9306 | 0.257919 | -0.831045 | -0.057448 | 0.594652 | -0.113171 | 0.145559 | -0.137143 | -0.669116 | -0.18662 | -0.921884 | -2.558207 | -0.56688 |
| 25.9444 | -1.337757 | -3.148828 | -0.501247 | 0.096934 | -1.019629 | -1.766045 | -1.092614 | -1.18022 | -1.257887 | 1.086224 | -7.611001 | -1.860562 |
| 25.9583 | -3.059454 | -3.698982 | -0.886113 | -1.14966 | -1.590404 | -3.222678 | -1.907179 | -1.004866 | -2.138696 | -0.783455 | -8.841981 | -1.795741 |
| 25.9722 | -3.834012 | -1.927382 | -0.742729 | -2.075238 | -1.811562 | -3.450644 | -1.443914 | -0.832213 | -2.629935 | -0.654799 | -5.625247 | -1.979023 |
| 25.9861 | -2.464539 | -0.619395 | -0.603508 | -2.0125 | -2.179401 | -1.926267 | -0.913783 | -0.492198 | -2.561234 | -0.089691 | -1.410816 | -0.088555 |
| 26.0069 | -0.908682 | -0.299619 | -0.984111 | -1.254757 | -1.33288 | -0.875886 | -1.49484 | -0.263004 | -1.985507 | -0.069689 | 0.100266 | 0.820601 |
| 26.0208 | -0.238513 | -0.348527 | -1.053489 | -0.569785 | -0.275879 | -0.230182 | 0.023464 | 0.059981 | -0.66456 | -0.090668 | -1.891923 | 0.484752 |
| 26.0347 | 0.19748 | -0.16592 | -0.221308 | -0.378963 | 0.672309 | 0.061878 | -0.236582 | 0.250824 | 0.506168 | -0.013231 | -0.149799 | -0.073449 |
| 26.0486 | 0.698642 | 0.097387 | 0.119572 | 0.140381 | 1.354208 | 0.232794 | 0.306271 | 0.347413 | 1.380595 | 0.019352 | 1.25444 | -0.2922 |
| 26.0625 | 1.119388 | 0.082148 | 0.27262 | 0.371429 | 1.245964 | 0.279394 | 0.516637 | 0.220641 | 1.499426 | -0.026103 | 1.402376 | -0.217008 |
| 26.0764 | 1.242677 | -0.074598 | 0.295874 | 0.580668 | 0.886334 | 0.270773 | 0.554113 | 0.133065 | 1.201687 | -0.072655 | 0.839485 | -0.089702 |
| 26.0903 | 1.144936 | -0.084955 | 0.217642 | 0.711703 | 0.801191 | 0.196743 | 0.479779 | 0.172528 | 0.871698 | -0.059128 | 0.564371 | -0.084195 |
| 26.1042 | 1.07535 | 0.012795 | 0.13932 | 0.732097 | 0.79743 | 0.148777 | 0.348865 | 0.20986 | 0.675903 | -0.017833 | 0.599205 | -0.180155 |
| 26.1181 | 1.077564 | 0.288999 | 0.193847 | 0.645396 | 0.702439 | 0.238963 | 0.251577 | 0.241641 | 0.608641 | -0.029574 | 0.5163 | -0.183335 |
| 26.1319 | 1.016613 | 0.562769 | 0.417987 | 0.52259 | 0.345233 | 0.433732 | 0.332395 | 0.242882 | 0.637673 | -0.033381 | 0.396096 | 0.005161 |
| 26.1458 | 0.868322 | 0.511188 | 0.619632 | 0.511502 | 0.041452 | 0.568206 | 0.527671 | 0.202345 | 0.752906 | -0.031759 | 0.386681 | 0.299335 |
| 26.1597 | 0.660765 | 0.083586 | 0.577932 | 0.537461 | -0.23055 | 0.582174 | 0.622416 | 0.111249 | 0.901249 | -0.000745 | 0.493546 | 0.430653 |
| 26.1736 | 0.585102 | -0.112161 | 0.390242 | 0.546906 | -0.491917 | 0.542743 | 0.559688 | 0.081185 | 1.087318 | 0.058229 | 0.619201 | 0.328168 |
| 26.1875 | 0.635632 | 0.210639 | 0.262766 | 0.573878 | -0.479388 | 0.636406 | 0.456566 | 0.17154 | 1.04709 | 0.049766 | 0.659824 | 0.134058 |
| 26.2014 | 0.603647 | 0.464324 | 0.155768 | 0.525564 | -0.294496 | 0.754486 | 0.317205 | 0.230393 | 0.863124 | -0.012657 | 0.678209 | 0.042165 |
| 26.2153 | 0.381733 | 0.262906 | 0.041046 | 0.430284 | -0.112646 | 0.647153 | 0.14312 | 0.160548 | 0.465378 | 0.001073 | 0.576475 | -0.062295 |
| 26.2292 | 0.29142 | -0.025478 | 0.043576 | 0.373344 | 0.042356 | 0.473956 | 0.088712 | 0.075769 | 0.192033 | 0.013272 | 0.422556 | 0.049987 |
| 26.2431 | 0.384852 | -0.04137 | 0.195259 | 0.348056 | 0.189815 | 0.383837 | 0.180346 | 0.080914 | 0.360977 | -0.01637 | 0.403896 | 0.390469 |
| 26.2569 | 0.34032 | 0.144588 | 0.269888 | 0.231157 | 0.265108 | 0.392328 | 0.26194 | 0.183163 | 0.82825 | -0.029821 | 0.526107 | 0.636149 |
| 26.2708 | 0.083813 | 0.329437 | 0.148931 | 0.022372 | 0.104976 | 0.364582 | 0.216056 | 0.292843 | 0.93363 | -0.012923 | 0.367592 | 0.616511 |
| 26.2847 | -0.358636 | 0.223465 | -0.080372 | -0.204539 | -0.294169 | 0.146214 | 0.048263 | 0.255305 | 0.254382 | -0.015018 | -0.217707 | 0.367913 |
| 26.2986 | -0.614006 | 0.040932 | -0.133285 | -0.384875 | -0.64524 | -0.136662 | -0.076841 | 0.092115 | -0.501227 | -0.01329 | -0.700971 | 0.114789 |
| 26.3125 | -0.301068 | 0.005444 | 0.064354 | -0.239473 | -0.668354 | -0.362155 | -0.020808 | -0.005441 | -0.536574 | 0.026171 | -0.580895 | 0.237787 |
| 26.3264 | 0.209848 | 0.025597 | 0.376835 | 0.052223 | -0.399786 | -0.494834 | 0.175636 | -0.008276 | 0.096161 | 0.011439 | 0.06819 | 0.671451 |
| 26.3403 | 0.699111 | 0.100708 | 0.582986 | 0.354644 | -0.05208 | -0.48417 | 0.352648 | 0.051872 | 1.006258 | -0.010068 | 0.630353 | 0.87926 |
| 26.3542 | 1.039714 | 0.329786 | 0.58231 | 0.522362 | 0.203971 | -0.28129 | 0.475031 | 0.159112 | 1.48252 | 0.014334 | 0.682636 | 0.719172 |
| 26.3681 | 0.871345 | 0.584485 | 0.469209 | 0.413506 | 0.304228 | 0.019272 | 0.625135 | 0.224076 | 1.114385 | 0.009279 | 0.327297 | 0.543667 |
| 26.3819 | 0.390633 | 0.285174 | 0.387733 | 0.25744 | 0.252287 | 0.106327 | 0.627493 | 0.098559 | 0.347423 | -0.032486 | -0.109504 | 0.526327 |
| 26.3958 | -0.076573 | -0.302718 | 0.329444 | 0.031575 | 0.157519 | 0.03196 | 0.393486 | -0.058964 | -0.54076 | -0.008399 | -0.452196 | 0.622626 |
| 26.4097 | -0.123274 | -0.322887 | 0.338967 | -0.06877 | 0.068695 | 0.144074 | 0.146873 | -0.071478 | -0.583739 | -0.044057 | -0.348824 | 0.632247 |
| 26.4236 | 0.081672 | 0.01831 | 0.323579 | 0.071899 | -0.098483 | 0.383131 | 0.008163 | -0.011915 | 0.282779 | -0.043485 | 0.203082 | 0.560187 |
| 26.4375 | 0.60244 | 1.164859 | 0.29764 | 0.3789 | -0.106841 | 1.051136 | 0.179399 | 0.260971 | 1.541458 | 0.340228 | 1.724872 | 0.60735 |
| 26.4514 | 1.452067 | 3.42503 | 0.412989 | 0.712405 | 0.57979 | 2.435268 | 0.882241 | 0.962713 | 2.824635 | 2.160158 | 3.811692 | 0.855724 |
| 26.4653 | 2.227199 | 3.728679 | 0.762857 | 1.03499 | 0.984047 | 3.680387 | 1.784937 | 1.670759 | 3.294357 | -1.118312 | 5.588824 | 1.9004 |
| 26.4792 | 3.105515 | 2.047609 | 1.66186 | 2.103847 | 0.59557 | 2.709153 | 2.336027 | 1.526239 | 2.488409 | 1.104781 | 5.133405 | 2.197805 |
| 26.4931 | 2.835116 | 1.015938 | 2.476399 | 2.905318 | 0.028688 | 1.83794 | 2.693378 | 0.748634 | 1.707438 | 0.166384 | 3.644705 | 1.714161 |
| 26.5 | 2.621891 | 1.027748 | 2.568933 | 2.916647 | -0.624066 | 0.890152 | 2.514345 | 0.563083 | 1.102909 | -0.016102 | 2.779415 | 0.985609 |
| 26.5139 | 1.480692 | 0.602299 | 1.44087 | 1.863209 | -1.197614 | 0.320856 | 1.239909 | 0.37312 | 0.435382 | -0.003251 | 1.293871 | 0.085203 |
| 26.5278 | 0.622841 | 0.144621 | 0.155231 | 0.451531 | -1.451603 | -0.113672 | -0.07294 | 0.149653 | -0.320258 | -0.00417 | -0.447359 | -0.813027 |
| 26.5417 | 0.126036 | -0.000476 | -0.948213 | -0.68964 | -1.342337 | -0.670309 | -1.366695 | -0.09465 | -1.043043 | 0.001274 | -1.687042 | -1.506028 |
| 26.5556 | -0.436413 | 0.002485 | -1.63442 | -1.090481 | -1.320778 | -1.022411 | -2.172022 | -0.146142 | -1.584573 | -0.023745 | -1.878115 | -1.386874 |
| 26.5694 | -0.760663 | 0.045114 | -1.409752 | -0.868893 | -0.927029 | -1.27474 | -2.060566 | -0.016708 | -1.116292 | -0.034636 | -1.360041 | -0.8757 |
| 26.5833 | -0.877737 | 0.080115 | -1.265219 | -0.336985 | -0.468334 | -1.972948 | -0.904431 | 0.124419 | -0.054354 | 0.053165 | -0.699786 | -0.031488 |

| Time | RGBStress2 | RGBStress3 | RGBStress4 | RGBStress5 | RGBStress6 | RGBStress7 | RGBStress8 | RGBStress9 | RGBStress10 | RGBStressnoplant | RGBStresstank |
|---|---|---|---|---|---|---|---|---|---|---|---|
| 24.0069 | -1.08029 | -0.290082 | -0.248771 | -0.140491 | -0.23776 | -0.637132 | 0.014491 | -0.096975 | -1.634475 | 0.011077 | -1.363231 |
| 24.0208 | -0.443798 | -0.163099 | -0.142001 | -0.13051 | -0.248656 | -0.322919 | -0.025512 | -0.095398 | -1.282105 | -0.01888 | -1.116381 |
| 24.0347 | -0.150791 | 0.141787 | 0.091377 | 0.008086 | -0.273498 | -0.173908 | 0.030784 | -0.046329 | -0.816621 | -0.059074 | -0.722277 |
| 24.0486 | -0.376637 | 0.238034 | 0.672711 | 0.423888 | -0.256118 | -0.053958 | 0.1048 | 0.028816 | -0.749655 | 0.076431 | -0.888302 |
| 24.0625 | -1.428882 | -0.404508 | 0.347012 | 0.646569 | -1.323279 | -0.587804 | -0.655127 | -0.855967 | -1.460895 | 0.615076 | -3.291267 |
| 24.0764 | -2.745241 | -2.292642 | -2.974059 | -0.862892 | -5.000655 | -3.086477 | -2.304281 | -2.409792 | -3.165173 | -1.32824 | -6.92291 |
| 24.0903 | -2.018984 | -2.594558 | -3.954 | -2.264225 | -4.610614 | -3.142473 | -1.94945 | -1.468377 | -2.715459 | -0.437249 | -6.098551 |
| 24.1042 | -0.145547 | -0.455883 | -0.777061 | -0.825816 | 0.087898 | 0.060208 | -0.340899 | 0.032925 | -0.252746 | 0.279462 | -0.888705 |
| 24.1181 | 0.650007 | 0.232753 | 1.182957 | 0.215872 | 2.916416 | 1.78242 | 0.745251 | 0.85918 | 0.997786 | -0.247954 | 1.632764 |
| 24.1319 | 0.296695 | -0.057827 | 0.745012 | 0.512235 | 1.156171 | 0.695466 | 0.453458 | 0.229962 | 0.681985 | 0.283205 | 0.806726 |
| 24.1458 | -0.066484 | -0.339467 | 0.218953 | 0.297547 | -0.395353 | -0.331013 | -0.087099 | -0.274516 | 0.118317 | 0.118649 | -0.185736 |
| 24.1597 | -0.177536 | -0.349336 | 0.271844 | 0.048083 | -0.892294 | -0.662649 | -0.294822 | -0.424885 | -0.152565 | 0.053973 | -0.406946 |
| 24.1736 | -0.2434 | -0.276223 | 0.597382 | -0.090938 | -0.703197 | -0.670369 | -0.314496 | -0.464821 | -0.303096 | -0.093634 | -0.270812 |
| 24.1875 | -0.295394 | -0.18906 | 0.754339 | -0.142063 | -0.828909 | -0.805119 | -0.313542 | -0.424744 | -0.461094 | -0.196132 | -0.288807 |
| 24.2014 | -0.293244 | -0.243535 | 0.672949 | -0.14665 | -1.180136 | -0.930792 | -0.33428 | -0.179928 | -0.547128 | -0.15362 | -0.349801 |
| 24.2153 | -0.232987 | -0.353605 | 0.534904 | -0.129008 | -1.232342 | -0.819323 | -0.32967 | 0.331399 | -0.592132 | -0.076275 | -0.176778 |
| 24.2292 | -0.187698 | -0.427395 | 0.348064 | -0.136133 | -0.969797 | -0.595396 | -0.316284 | 0.830922 | -0.651484 | -0.10352 | -0.094765 |
| 24.2431 | -0.21191 | -0.398072 | 0.102526 | -0.154475 | -0.851893 | -0.478131 | -0.271286 | 0.702696 | -0.709539 | -0.200669 | -0.255938 |
| 24.2569 | -0.230238 | -0.209829 | -0.087857 | -0.146304 | -0.815442 | -0.437789 | -0.210844 | 0.269372 | -0.636899 | -0.195218 | -0.488193 |
| 24.2708 | -0.190422 | -0.023918 | -0.148651 | -0.11748 | -0.723508 | -0.376024 | -0.198863 | 0.043549 | -0.478764 | -0.069277 | -0.591479 |
| 24.2847 | -0.183602 | -0.00094 | -0.17265 | -0.124531 | -0.614626 | -0.325338 | -0.203402 | -0.051983 | -0.390476 | -0.049448 | -0.555869 |
| 24.2986 | -0.24959 | 0.016112 | -0.246816 | -0.151135 | -0.591619 | -0.322993 | -0.19772 | -0.111275 | -0.404795 | -0.160291 | -0.425641 |
| 24.3125 | -0.244699 | 0.077351 | -0.229938 | -0.120786 | -0.500567 | -0.254577 | -0.132961 | -0.061444 | -0.347872 | -0.144718 | -0.276271 |
| 24.3264 | -0.127186 | 0.169132 | -0.043095 | -0.038364 | -0.238778 | -0.105615 | -0.029528 | 0.054438 | -0.175111 | 0.147753 | -0.054324 |
| 24.3403 | 0.014319 | 0.270962 | 0.247378 | 0.030784 | 0.187926 | 0.116792 | 0.080281 | 0.241554 | 0.053579 | 0.723593 | 0.245665 |
| 24.3542 | 0.20271 | 0.446492 | 0.50861 | 0.114988 | 0.686757 | 0.419035 | 0.25176 | 0.596288 | 0.308982 | -0.049542 | 0.578649 |
| 24.3681 | 0.368256 | 0.560952 | 0.567016 | 0.21814 | 0.907823 | 0.58898 | 0.435831 | 0.709628 | 0.46304 | -0.540566 | 0.65944 |
| 24.3819 | 0.426247 | 0.46797 | 0.494801 | 0.279891 | 0.864901 | 0.595985 | 0.43191 | 0.539218 | 0.42744 | -0.030678 | 0.512771 |
| 24.3958 | 0.395693 | 0.422009 | 0.361752 | 0.280391 | 0.716223 | 0.513565 | 0.392015 | 0.312413 | 0.297195 | 0.188645 | 0.469682 |
| 24.4097 | 0.256148 | 0.346262 | 0.190629 | 0.1906 | 0.487853 | 0.355481 | 0.292375 | 0.162228 | 0.146026 | 0.129555 | 0.370753 |
| 24.4236 | 0.160676 | 0.131789 | 0.178777 | 0.062908 | 0.339519 | 0.25666 | 0.129375 | 0.29206 | 0.161425 | 0.098285 | -0.110243 |
| 24.4375 | 0.287257 | -0.402472 | 0.680681 | -0.107923 | 0.635181 | 0.482103 | -0.012012 | 1.258413 | 0.665727 | 0.353923 | -1.749272 |
| 24.4514 | 0.897877 | -1.125758 | 1.45345 | -0.333002 | 1.527022 | 1.201519 | -0.03709 | 2.969536 | 1.601341 | 1.296627 | -3.796706 |
| 24.4653 | 1.892243 | -0.591807 | 2.403334 | -0.25965 | 2.56635 | 2.108846 | 0.270961 | 3.809884 | 2.107074 | 1.770757 | -2.121362 |
| 24.4792 | 1.701388 | 0.736631 | 2.351565 | 0.378948 | 4.962287 | 3.036031 | 1.222568 | 3.282169 | 1.973209 | 0.86487 | 1.518531 |
| 24.4931 | 0.802239 | 1.270706 | 1.745905 | 0.579373 | 3.639989 | 2.357129 | 1.132045 | 1.421115 | 1.113467 | 0.145095 | 2.07157 |
| 24.5 | 0.370409 | 1.02378 | 0.920104 | 0.428795 | 1.671408 | 1.242251 | 0.570816 | 0.139002 | 0.568375 | 0.06621 | 0.979126 |
| 24.5139 | 0.157898 | 0.610702 | 0.531319 | 0.27988 | 0.835207 | 0.730603 | 0.289143 | -0.066997 | 0.250671 | 0.062534 | 0.388889 |
| 24.5278 | 0.043521 | 0.346581 | 0.294029 | 0.170319 | 0.487789 | 0.475872 | 0.181017 | 0.578478 | 0.079249 | 0.017216 | 0.281061 |
| 24.5417 | 0.034681 | 0.218648 | 0.155559 | 0.096741 | 0.297234 | 0.321942 | 0.162905 | 1.001951 | 0.037514 | -0.015566 | 0.284948 |
| 24.5556 | 0.100504 | 0.180549 | 0.079757 | 0.067094 | 0.218008 | 0.269628 | 0.197664 | 0.898526 | 0.10188 | -0.000526 | 0.288099 |
| 24.5694 | 0.200601 | 0.169814 | 0.051481 | 0.077643 | 0.190393 | 0.281127 | 0.227879 | 0.717357 | 0.169923 | 0.036057 | 0.257713 |
| 24.5833 | 0.297447 | 0.142365 | 0.063538 | 0.090664 | 0.174106 | 0.297556 | 0.243452 | 0.570666 | 0.204502 | 0.088581 | 0.219785 |
| 24.5972 | 0.303366 | 0.101559 | 0.081081 | 0.082486 | 0.162898 | 0.289601 | 0.238555 | 0.423541 | 0.191476 | 0.120385 | 0.190525 |
| 24.6111 | 0.182017 | 0.056672 | 0.075445 | 0.056256 | 0.151615 | 0.253277 | 0.171699 | 0.275314 | 0.127615 | 0.072389 | 0.173046 |
| 24.625 | 0.052629 | 0.016867 | 0.057942 | 0.022117 | 0.146599 | 0.220675 | 0.095237 | 0.198775 | 0.062887 | -0.022379 | 0.163635 |
| 24.6389 | 0.024667 | 0.055497 | 0.052976 | 0.012391 | 0.114422 | 0.206655 | 0.046594 | 0.233142 | 0.068149 | -0.071104 | 0.175007 |
| 24.6528 | 0.124788 | 0.412889 | 0.049534 | 0.021217 | 0.051517 | 0.201927 | 0.029934 | 0.290813 | 0.124817 | -0.042089 | 0.229763 |
| 24.6667 | 0.240844 | 1.374875 | 0.047281 | 0.031551 | -0.000435 | 0.191305 | 0.036611 | 0.281318 | 0.174575 | 0.001755 | 0.32931 |
| 24.6806 | 0.253219 | 0.703858 | 0.050865 | 0.04831 | -0.018431 | 0.162035 | 0.061107 | 0.197801 | 0.175664 | 0.016839 | 0.323903 |
| 24.6944 | 0.100108 | 0.277003 | 0.062944 | 0.085675 | 0.011756 | 0.139551 | 0.096353 | 0.090509 | 0.157341 | 0.043098 | 0.218012 |
| 24.7083 | -0.053743 | 0.311613 | 0.086064 | 0.109724 | 0.07921 | 0.130536 | 0.122306 | -0.005973 | 0.140067 | 0.10305 | 0.172658 |
| 24.7222 | -0.074735 | 0.253518 | 0.088439 | 0.070633 | 0.094429 | 0.096444 | 0.103379 | -0.089836 | 0.114501 | 0.09281 | 0.164011 |
| 24.7361 | -0.006566 | 0.160751 | 0.057476 | -0.008356 | 0.065555 | 0.029529 | 0.037576 | -0.160555 | 0.078443 | -0.009012 | 0.137469 |
| 24.75 | 0.129328 | 0.09948 | 0.021357 | -0.075063 | 0.022271 | -0.02925 | -0.043421 | -0.194783 | 0.059216 | -0.073098 | 0.1255 |
| 24.7639 | 0.292332 | 0.073151 | -0.002195 | -0.107318 | -0.005328 | -0.044219 | -0.090944 | -0.113268 | 0.077096 | -0.088527 | 0.143753 |
| 24.7778 | 0.245179 | 0.04964 | -0.009886 | -0.109755 | -0.022225 | -0.01634 | -0.108541 | -0.012077 | 0.108223 | -0.093994 | 0.147239 |
| 24.7917 | 0.02459 | 0.021338 | -0.017217 | -0.088419 | -0.050817 | 0.036341 | -0.108675 | 0.051765 | 0.122045 | -0.072619 | 0.140491 |
| 24.8056 | -0.127465 | -0.001807 | -0.023279 | -0.028109 | -0.066991 | 0.093205 | -0.095558 | 0.07822 | 0.106686 | -0.064435 | 0.172332 |
| 24.8194 | -0.123713 | 0.008049 | 0.003208 | 0.047446 | -0.053033 | 0.118902 | -0.057551 | 0.102858 | 0.097669 | -0.073309 | 0.268027 |
| 24.8333 | 0.048024 | 0.043479 | 0.057113 | 0.049528 | 0.000139 | 0.094617 | -0.024515 | 0.13054 | 0.089083 | -0.049796 | 0.36074 |
| 24.8472 | 0.256175 | 0.04895 | 0.072009 | -0.022478 | 0.046615 | 0.054892 | -0.038492 | 0.146672 | 0.072638 | -0.01419 | 0.388982 |
| 24.8611 | 0.338854 | 0.017342 | 0.033257 | -0.112527 | 0.052873 | 0.033627 | -0.091472 | 0.13377 | 0.055703 | -0.013375 | 0.31878 |
| 24.875 | 0.222599 | -0.027775 | -0.053904 | -0.183383 | 0.014221 | 0.040719 | -0.154875 | 0.079357 | 0.076891 | -0.035348 | 0.192494 |
| 24.8889 | 0.097786 | -0.052419 | -0.149336 | -0.121817 | -0.027597 | 0.042805 | -0.191209 | 0.074503 | 0.125576 | -0.038285 | 0.142045 |
| 24.9028 | 0.00876 | -0.046034 | -0.179861 | -0.044693 | -0.044226 | 0.031705 | -0.131589 | 0.163634 | 0.167523 | -0.044255 | 0.235404 |
| 24.9167 | -0.151308 | -0.003601 | -0.197221 | 0.035267 | -0.107675 | -0.029954 | 0.046342 | 0.039174 | 0.099141 | -0.181891 | 0.651452 |
| 24.9306 | -0.861369 | 0.08688 | -0.688283 | 0.227163 | -0.695164 | -0.47062 | 0.270899 | -1.194518 | -0.409775 | -0.853616 | 2.342168 |
| 24.9444 | -1.605255 | 0.021732 | -1.894082 | 0.246598 | -2.525045 | -1.940179 | -0.14411 | -2.897569 | -1.628721 | -1.477978 | 3.654744 |
| 24.9583 | -1.842332 | -0.807001 | -2.837889 | -0.215122 | -4.046789 | -3.519116 | -1.189516 | -2.861345 | -2.827338 | -1.106877 | 1.432836 |
| 24.9722 | -1.363534 | -1.241383 | -1.92383 | -0.670711 | -3.176708 | -2.71961 | -1.270245 | -1.167073 | -1.97914 | -0.418936 | -1.192401 |
| 24.9861 | -0.390802 | -0.150625 | -0.239449 | -0.317481 | -0.226407 | -0.365016 | -0.373453 | 0.028002 | -0.518807 | -0.049506 | -0.741832 |
| 25.0069 | 0.251509 | 0.431793 | 0.778252 | 0.156704 | 1.459802 | 0.955007 | 0.363734 | 0.351422 | 0.074068 | -0.006672 | 0.460361 |
| 25.0208 | 0.311058 | 0.570622 | 0.932198 | 0.477579 | 1.437853 | 1.056047 | 0.554414 | 0.396368 | 0.265915 | 0.024317 | 0.793938 |
| 25.0347 | 0.27234 | 0.706141 | 0.825472 | 0.475246 | 1.000648 | 0.806295 | 0.474576 | 0.322714 | 0.290229 | 0.120173 | 0.887975 |
| 25.0486 | 0.28991 | 0.818725 | 0.553547 | 0.317567 | 0.770447 | 0.720331 | 0.40594 | 0.260099 | 0.248183 | 0.40016 | 1.178613 |
| 25.0625 | 0.362128 | 0.860214 | 0.316334 | 0.265661 | 0.691562 | 0.748987 | 0.454242 | 0.292359 | 0.306983 | 0.312991 | 1.419334 |
| 25.0764 | 0.342597 | 0.684899 | 0.177294 | 0.21859 | 0.458712 | 0.533967 | 0.427217 | 0.2523 | 0.235305 | 0.019341 | 1.087528 |
| 25.0903 | 0.263312 | 0.325717 | 0.234215 | 0.109133 | 0.361449 | 0.308635 | 0.242614 | 0.256157 | 0.125211 | -0.154565 | 0.063508 |
| 25.1042 | 0.18113 | 0.043602 | 0.405321 | -0.004172 | 0.612827 | 0.382131 | 0.060657 | 0.441023 | 0.235923 | -0.166404 | -0.871757 |
| 25.1181 | 0.085643 | -0.020082 | 0.380836 | -0.086948 | 0.553705 | 0.430954 | 0.023957 | 0.450769 | 0.254461 | 0.173673 | -0.851395 |
| 25.1319 | 0.031678 | 0.109269 | 0.151436 | -0.098594 | 0.053928 | 0.242262 | 0.04708 | 0.122849 | 0.044383 | -0.131567 | 0.110345 |
| 25.1458 | 0.08241 | 0.139204 | 0.148364 | -0.091008 | -0.115718 | 0.209225 | -0.023958 | -0.029168 | 0.039328 | -0.269973 | 0.707418 |
| 25.1597 | 0.166664 | 0.033622 | 0.287237 | -0.068251 | 0.180441 | 0.407996 | -0.085093 | 0.132199 | 0.276022 | -0.090075 | 0.409879 |
| 25.1736 | 0.167068 | 0.05153 | 0.267532 | 0.003851 | 0.297472 | 0.479779 | 0.0342 | 0.240764 | 0.32936 | 0.029321 | 0.096307 |
| 25.1875 | 0.15962 | 0.175208 | 0.179197 | 0.052864 | 0.168017 | 0.391174 | 0.200035 | 0.217514 | 0.161473 | 0.002853 | 0.11692 |
| 25.2014 | 0.111819 | 0.347901 | 0.096468 | 0.048265 | -0.156592 | 0.182476 | 0.232576 | 0.041741 | -0.140153 | -0.086888 | 0.686971 |
| 25.2153 | 0.0546 | 0.386763 | 0.098357 | -0.004098 | -0.349905 | 0.088264 | 0.091187 | -0.105321 | -0.290921 | -0.061717 | 1.191264 |
| 25.2292 | 0.132686 | 0.104265 | 0.215895 | -0.056617 | -0.113681 | 0.252367 | -0.084496 | 0.005652 | -0.05428 | 0.017422 | 0.708603 |
| 25.2431 | 0.207642 | -0.19206 | 0.095286 | -0.004506 | 0.094301 | 0.408899 | -0.074996 | 0.13031 | 0.153881 | 0.059697 | 0.167345 |
| 25.2569 | 0.213585 | -0.272086 | -0.243116 | 0.102579 | 0.097899 | 0.473094 | 0.057138 | 0.13404 | 0.131231 | 0.010766 | 0.055595 |
| 25.2708 | 0.20586 | -0.092111 | -0.474165 | 0.173529 | 0.0998 | 0.467032 | 0.206314 | 0.119515 | 0.027213 | -0.024334 | 0.054499 |
| 25.2847 | 0.207224 | 0.145842 | -0.23295 | 0.187497 | 0.160569 | 0.393589 | 0.247881 | 0.138231 | 0.001005 | 0.031515 | 0.037303 |
| 25.2986 | 0.307204 | 0.077861 | 0.425541 | 0.133236 | 0.47473 | 0.501772 | 0.081726 | 0.3257 | 0.289309 | 0.373355 | -0.396033 |
| 25.3125 | 0.477425 | -0.253651 | 0.967968 | 0.065376 | 0.875396 | 0.766063 | -0.104016 | 0.616004 | 0.767013 | -0.094323 | -1.096252 |
| 25.3264 | 0.379421 | -0.317261 | 0.561796 | 0.064092 | 0.569632 | 0.698709 | -0.089442 | 0.482977 | 0.704382 | 0.027759 | -0.636101 |
| 25.3403 | 0.162492 | -0.174453 | -0.216748 | 0.174131 | -0.06572 | 0.408741 | 0.03862 | 0.078476 | 0.235566 | -0.0876 | 0.413102 |
| 25.3542 | 0.227518 | -0.186787 | -0.523799 | 0.261726 | 0.034284 | 0.439491 | 0.154031 | 0.091739 | 0.164057 | -0.306455 | 0.311067 |
| 25.3681 | 0.281098 | 0.079994 | -0.35629 | 0.323296 | 0.2004 | 0.41042 | 0.314604 | 0.125945 | 0.158925 | 0.006583 | 0.337685 |
| 25.3819 | 0.112964 | 0.679294 | -0.119332 | 0.398643 | -0.229495 | 0.120564 | 0.477031 | -0.214389 | -0.043022 | -0.130223 | 1.518675 |
| 25.3958 | 0.098845 | 0.875313 | 0.257358 | 0.402969 | -0.401516 | -0.010788 | 0.427935 | -0.283271 | 0.161386 | -0.003047 | 1.942599 |
| 25.4097 | 0.156134 | 0.606502 | 0.585141 | 0.293523 | -0.058984 | 0.06444 | 0.305145 | -0.015355 | 0.519166 | -0.059404 | 1.186818 |
| 25.4236 | 0.189639 | 0.209901 | 0.370051 | 0.15515 | 0.325487 | 0.211091 | 0.270848 | 0.198959 | 0.60143 | -0.071499 | 0.165479 |
| 25.4375 | 0.591007 | -0.389261 | 0.642191 | 0.03376 | 1.444133 | 0.889982 | 0.366299 | 1.112261 | 1.27466 | 0.18066 | -2.277056 |
| 25.4514 | 1.622174 | -0.866232 | 1.960597 | 0.050894 | 3.179636 | 1.858119 | 0.482853 | 3.315281 | 2.985544 | 1.575696 | -5.132402 |
| 25.4653 | 3.005507 | 0.307676 | 2.623326 | 0.336343 | 2.569599 | 1.901829 | 0.853845 | 5.293456 | 3.729206 | 1.9552 | -2.114478 |
| 25.4792 | 2.191768 | 2.44376 | 2.115649 | 0.923527 | 2.337312 | 1.775677 | 1.346024 | 3.364377 | 3.082843 | 0.870047 | 2.685205 |
| 25.4931 | 0.72194 | 2.085772 | 1.665904 | 0.896331 | 1.88135 | 1.108735 | 1.404807 | 1.585057 | 1.854233 | -0.005092 | 3.082954 |
| 25.5 | 0.206057 | 1.484083 | 1.613397 | 0.699552 | 1.858477 | 0.822038 | 1.048591 | 1.3119 | 1.236791 | -0.07499 | 2.158808 |
| 25.5139 | 0.004757 | 0.819862 | 0.9678 | 0.501815 | 1.74624 | 0.668569 | 0.475927 | 1.123429 | 0.755737 | -0.025426 | 1.580097 |
| 25.5278 | 0.071523 | 0.256802 | 0.34038 | 0.397764 | 1.164166 | 0.462972 | -0.181147 | 0.637511 | 0.389253 | 0.011555 | 1.057313 |
| 25.5417 | 0.259263 | -0.274413 | -0.204577 | 0.12651 | 0.597676 | 0.250871 | -0.38464 | 0.318597 | 0.158197 | 0.038384 | 0.438724 |
| 25.5556 | 0.467808 | -0.434249 | -0.648423 | -0.174411 | 0.064371 | -0.004176 | 0.033206 | 0.235857 | 0.015956 | 0.097979 | -0.147635 |
| 25.5694 | 0.379067 | -0.055422 | -0.972387 | 0.001993 | -0.569147 | -0.388219 | 0.625381 | 0.159291 | -0.256239 | 0.109787 | -0.331505 |
| 25.5833 | -0.179426 | 0.491722 | -0.765717 | 0.221107 | -1.13445 | -0.68887 | 0.835739 | -0.037586 | -0.578132 | 0.003298 | -0.423156 |
| 25.5972 | -0.651169 | 0.671765 | -0.244485 | 0.006352 | -1.252187 | -0.820466 | 0.25763 | -0.260317 | -0.615751 | -0.075621 | -0.593454 |
| 25.6111 | -0.722657 | 0.617913 | 0.251544 | -0.285335 | -1.174618 | -0.888818 | -0.1818 | -0.362825 | -0.570061 | -0.073509 | -0.695273 |
| 25.625 | -0.522951 | 0.014911 | 0.49541 | -0.573411 | -0.808452 | -0.558212 | -0.656556 | -0.342094 | -0.574149 | -0.036981 | -0.681653 |
| 25.6389 | -0.11225 | -0.430489 | 0.32399 | -0.79044 | -0.243024 | 0.092337 | -1.239391 | -0.185408 | -0.504089 | 0.007278 | -0.67123 |
| 25.6528 | 0.167553 | -0.989098 | -0.064402 | -0.820102 | 0.328138 | 0.604787 | -1.009141 | -0.095911 | -0.272954 | -0.01127 | -0.430748 |
| 25.6667 | 0.202136 | -0.998468 | -0.387895 | -0.368664 | 0.693621 | 0.815545 | -0.048262 | -0.038481 | 0.220862 | -0.060634 | -0.091481 |
| 25.6806 | 0.160369 | -0.707674 | -0.453601 | 0.158405 | 0.725717 | 0.828806 | 0.636266 | 0.004104 | 0.720147 | -0.044004 | 0.232725 |
| 25.6944 | -0.045745 | -0.089993 | -0.198571 | 0.566186 | 0.524837 | 0.563633 | 1.157441 | -0.022016 | 0.745702 | -0.021726 | 0.570458 |
| 25.7083 | -0.061342 | 0.580973 | 0.055003 | 0.587983 | 0.213557 | 0.157026 | 1.178944 | -0.043019 | 0.387819 | -0.027897 | 0.81709 |
| 25.7222 | 0.173688 | 0.84225 | 0.293177 | 0.319732 | -0.110751 | -0.152355 | 0.336118 | -0.012345 | 0.097667 | -0.012156 | 0.721427 |
| 25.7361 | 0.330019 | 0.748363 | 0.334186 | -0.035329 | -0.379853 | -0.429526 | -0.201973 | -0.010125 | -0.079007 | 0.003074 | 0.391928 |
| 25.75 | 0.1584 | 0.42197 | 0.153247 | -0.384078 | -0.672709 | -0.667126 | -0.585666 | -0.051651 | -0.168256 | 0.000916 | -0.226663 |
| 25.7639 | -0.191794 | -0.057431 | -0.121679 | -0.460067 | -1.004471 | -0.782233 | -0.768417 | -0.100026 | -0.220559 | -0.002598 | -0.969074 |
| 25.7778 | -0.505343 | -0.778456 | -0.348223 | -0.336373 | -1.280246 | -1.13317 | -0.378113 | -0.102673 | -0.317936 | -0.012298 | -1.389697 |
| 25.7917 | -0.682208 | -1.205739 | -0.454991 | -0.177627 | -1.352101 | -1.317338 | -0.021988 | -0.039734 | -0.487865 | -0.018756 | -1.324809 |
| 25.8056 | -0.549787 | -0.901866 | -0.386432 | -0.224499 | -1.188785 | -1.245903 | -0.08626 | 0.07808 | -0.592106 | 0.014043 | -1.074406 |
| 25.8194 | -0.317574 | -0.676857 | -0.267311 | -0.578597 | -0.997886 | -0.802601 | -0.267344 | 0.211434 | -0.593139 | 0.061431 | -0.905534 |
| 25.8333 | -0.284376 | -0.473044 | -0.176032 | -0.811566 | -0.790582 | -0.345395 | -0.41868 | 0.174965 | -0.600167 | 0.070118 | -0.741378 |
| 25.8472 | -0.491328 | -0.283522 | -0.087816 | -0.766771 | -0.590767 | -0.027548 | -0.391639 | -0.011132 | -0.708487 | 0.041685 | -0.523961 |
| 25.8611 | -0.706593 | -0.178015 | -0.051971 | -0.598601 | -0.312436 | 0.124797 | -0.296621 | -0.193394 | -0.720577 | 0.004911 | -0.38914 |
| 25.875 | -0.688494 | -0.089843 | -0.083643 | -0.443558 | -0.03484 | 0.009795 | -0.175093 | -0.331529 | -0.68786 | -0.023467 | -0.164437 |
| 25.8889 | -0.187094 | 0.017197 | -0.087422 | -0.322106 | 0.103422 | -0.18869 | -0.121938 | -0.295049 | -0.456003 | 0.003241 | 0.055631 |
| 25.9028 | 0.314183 | 0.119001 | -0.032415 | -0.205982 | 0.162479 | -0.325935 | -0.140322 | -0.168596 | -0.191493 | 0.063743 | 0.163106 |
| 25.9167 | 0.292909 | 0.091412 | -0.155427 | -0.04981 | 0.075891 | -0.393639 | -0.043905 | -0.250385 | -0.066331 | -0.065156 | 0.26554 |
| 25.9306 | -0.949298 | -0.11949 | -1.117629 | 0.043843 | -0.702359 | -0.736258 | 0.258246 | -1.296376 | -0.501699 | -1.001075 | 0.810773 |
| 25.9444 | -2.389048 | -0.505927 | -2.646419 | -0.313222 | -2.531955 | -1.749949 | 0.118759 | -3.179197 | -1.676717 | -2.063944 | -0.131833 |
| 25.9583 | -2.017311 | -0.97781 | -3.170192 | -0.998268 | -3.623289 | -2.476925 | -0.750149 | -3.951483 | -2.825025 | -1.486496 | -1.436453 |
| 25.9722 | -0.9625 | -1.080887 | -1.841455 | -0.938178 | -1.1238 | -1.438688 | -0.965066 | -1.894433 | -1.911442 | -0.285425 | -1.467108 |
| 25.9861 | 0.248018 | -0.531544 | 0.061213 | -0.315594 | 1.280611 | 0.28336 | -0.28815 | -0.098223 | 0.336274 | 0.137779 | -0.633053 |
| 26.0069 | 0.656047 | -0.089971 | 1.064125 | 0.042488 | 1.892842 | 0.78652 | 0.251306 | 0.788059 | 1.170892 | 0.297237 | -0.047162 |
| 26.0208 | 0.167594 | 0.254402 | 1.087877 | 0.093202 | 1.149827 | 0.339605 | 0.420601 | 0.447489 | 0.558728 | 0.069744 | 0.278648 |
| 26.0347 | -0.358815 | 0.629466 | 0.610659 | 0.074878 | 0.041838 | -0.298779 | 0.351166 | -0.170367 | -0.350414 | -0.20836 | 0.883866 |
| 26.0486 | -0.30868 | 0.596726 | 0.469026 | 0.067297 | -0.102347 | -0.28281 | 0.277285 | -0.12075 | -0.344263 | -0.121755 | 0.556676 |
| 26.0625 | 0.095811 | 0.186041 | 0.735564 | 0.058387 | 0.429411 | 0.15101 | 0.228894 | 0.329581 | 0.249966 | 0.174814 | -0.42682 |
| 26.0764 | 0.176967 | 0.009302 | 0.600848 | 0.069926 | 0.32302 | 0.188474 | 0.158022 | 0.239969 | 0.245736 | 0.117074 | -0.404216 |
| 26.0903 | -0.157084 | 0.037047 | -0.067077 | 0.091233 | -0.353485 | -0.188613 | 0.018578 | -0.397597 | -0.358739 | -0.155913 | 0.545118 |
| 26.1042 | -0.231322 | -0.061992 | -0.436328 | 0.070976 | -0.393715 | -0.276382 | -0.185951 | -0.585208 | -0.451961 | -0.183373 | 0.594962 |
| 26.1181 | 0.030651 | -0.269286 | -0.376234 | 0.025524 | 0.216576 | 0.021043 | -0.252313 | -0.163703 | 0.104919 | 0.113826 | -0.263263 |
| 26.1319 | 0.165684 | -0.250998 | -0.206447 | 0.017708 | 0.263294 | 0.117593 | -0.030405 | -0.007387 | 0.323671 | 0.172414 | -0.314622 |
| 26.1458 | 0.260772 | -0.030034 | -0.008213 | 0.038667 | 0.121364 | 0.166228 | 0.245206 | 0.013483 | 0.334173 | 0.043423 | -0.030075 |
| 26.1597 | 0.428821 | -0.010985 | 0.27558 | 0.0266 | 0.40241 | 0.374217 | 0.17356 | 0.238173 | 0.540713 | 0.05359 | -0.311827 |
| 26.1736 | 0.238901 | -0.041547 | 0.155729 | 0.031889 | 0.188932 | 0.2244 | -0.088537 | 0.003342 | 0.291525 | -0.049525 | -0.115274 |
| 26.1875 | -0.181658 | -0.031062 | -0.268573 | 0.083795 | -0.41583 | -0.192746 | -0.151161 | -0.586447 | -0.261919 | -0.211857 | 0.685672 |
| 26.2014 | -0.309947 | -0.110002 | -0.480091 | 0.119238 | -0.479862 | -0.263662 | -0.043613 | -0.64251 | -0.273234 | -0.28469 | 0.679697 |
| 26.2153 | -0.205485 | -0.268233 | -0.405156 | 0.084827 | -0.10841 | -0.036185 | -0.098533 | -0.32614 | 0.040813 | -0.084842 | 0.045675 |
| 26.2292 | 0.033602 | -0.332326 | -0.191333 | 0.02406 | 0.268932 | 0.265203 | -0.355986 | -0.009636 | 0.306904 | 0.072808 | -0.434855 |
| 26.2431 | 0.322819 | -0.188558 | 0.116243 | 0.023333 | 0.476771 | 0.519029 | -0.285764 | 0.116703 | 0.4521 | 0.21107 | -0.40182 |
| 26.2569 | 0.416257 | 0.16909 | 0.287085 | 0.109173 | 0.21421 | 0.483381 | 0.093057 | -0.116143 | 0.256918 | 0.111658 | 0.376825 |
| 26.2708 | 0.439794 | 0.315412 | 0.358352 | 0.192091 | 0.13074 | 0.434742 | 0.471153 | -0.103059 | 0.207897 | 0.074548 | 0.647197 |
| 26.2847 | 0.439863 | 0.095237 | 0.318358 | 0.192573 | 0.348344 | 0.430157 | 0.491388 | 0.221091 | 0.386597 | 0.089227 | 0.010497 |
| 26.2986 | 0.292907 | -0.120342 | 0.04899 | 0.126192 | 0.118673 | 0.161307 | 0.148429 | 0.084078 | 0.207813 | -0.012979 | -0.24252 |
| 26.3125 | 0.367605 | -0.313553 | -0.054106 | 0.008953 | 0.021494 | 0.076193 | -0.136249 | -0.038321 | 0.095786 | -0.056601 | -0.364436 |
| 26.3264 | 0.565197 | -0.345545 | 0.117312 | -0.075856 | 0.205956 | 0.24669 | -0.119951 | 0.018999 | 0.139686 | -0.025051 | -0.316903 |
| 26.3403 | 0.410985 | 0.009146 | 0.293044 | 0.017488 | 0.079449 | 0.306606 | 0.216724 | -0.103433 | -0.009367 | -0.05183 | 0.44563 |
| 26.3542 | 0.248964 | 0.393431 | 0.37146 | 0.243857 | 0.086179 | 0.365834 | 0.725788 | 0.027702 | 0.022636 | 0.022722 | 0.935135 |
| 26.3681 | 0.221443 | 0.640199 | 0.304204 | 0.444601 | 0.009278 | 0.366552 | 0.987934 | 0.146464 | 0.096542 | 0.026336 | 1.127401 |
| 26.3819 | 0.327718 | 0.61984 | 0.211254 | 0.477465 | 0.052922 | 0.418847 | 0.924845 | 0.176564 | 0.176309 | -0.051291 | 0.975133 |
| 26.3958 | 0.45866 | 0.271109 | 0.226256 | 0.329212 | 0.510563 | 0.651514 | 0.587715 | 0.30087 | 0.388622 | -0.047901 | 0.359179 |
| 26.4097 | 0.17572 | 0.071672 | 0.123442 | 0.187036 | 0.227117 | 0.471708 | 0.266133 | -0.082518 | 0.088028 | -0.09337 | 0.449086 |
| 26.4236 | -0.13159 | 0.032771 | 0.129647 | 0.143187 | -0.149286 | 0.242485 | 0.207474 | -0.307793 | -0.172392 | -0.095208 | 0.681513 |
| 26.4375 | 0.31648 | -0.335839 | 0.481494 | 0.028267 | 1.126541 | 0.865133 | 0.518511 | 0.878674 | 0.866614 | 0.318309 | -0.527134 |
| 26.4514 | 1.249657 | -0.604298 | 1.335249 | 0.09072 | 3.655263 | 2.206205 | 0.887288 | 3.229602 | 2.932979 | 1.55505 | -1.452449 |
| 26.4653 | 2.434868 | -0.043977 | 2.248976 | 0.867107 | 4.121547 | 2.821282 | 1.047532 | 4.368538 | 3.632814 | 2.001046 | 2.221281 |
| 26.4792 | 1.866351 | 1.532092 | 2.475954 | 1.384307 | 2.86125 | 2.69614 | 1.481546 | 2.645537 | 2.748044 | 0.846768 | 4.352613 |
| 26.4931 | 0.860895 | 1.554544 | 2.089316 | 1.684169 | 1.712957 | 1.814105 | 1.511055 | 1.076543 | 1.54924 | -0.010656 | 3.894209 |
| 26.5 | 0.318 | 1.371808 | 1.822391 | 1.688726 | 1.528921 | 1.360776 | 1.473635 | 0.560084 | 0.783016 | -0.066552 | 2.948534 |
| 26.5139 | -0.106829 | 0.933974 | 1.005842 | 1.096602 | 1.585025 | 1.050982 | 0.928139 | 0.369849 | 0.362245 | 0.001301 | 1.695473 |
| 26.5278 | -0.468246 | 0.441318 | 0.195213 | 0.440582 | 1.316903 | 0.536463 | 0.25959 | 0.165967 | 0.026849 | 0.028158 | 0.414118 |
| 26.5417 | -0.722658 | 0.093742 | -0.602398 | -0.269005 | 0.807691 | -0.153097 | -0.512514 | 0.083517 | -0.305545 | 0.006127 | -0.713584 |
| 26.5556 | -0.433013 | -0.250522 | -0.973 | -0.826188 | 0.117236 | -0.890576 | -0.856845 | 0.136106 | -0.581731 | -0.014142 | -1.391688 |
| 26.5694 | 0.044738 | -0.311296 | -0.992588 | -0.96008 | -0.635306 | -1.377458 | -0.479368 | 0.18753 | -0.741112 | -0.012604 | -1.42665 |
| 26.5833 | 0.478976 | -0.068804 | -0.507593 | -0.65925 | -1.21111 | -1.263793 | 0.191729 | 0.062676 | -0.867621 | -0.01596 | -1.064368 |

| Time | Mean RGB_control | | Time | Mean RGB_stress |
|---|---|---|---|---|
| 24.0069 | -0.849782 | | 24 | -1.363231 |
| 24.0208 | -0.443154 | | 24.0139 | -1.116381 |
| 24.0347 | -0.122628 | | 24.0278 | -0.722277 |
| 24.0486 | 0.0833 | | 24.0417 | -0.888302 |
| 24.0625 | 0.376187 | | 24.0556 | -3.291267 |
| 24.0764 | 1.147689 | | 24.0694 | -6.92291 |
| 24.0903 | 2.011624 | | 24.0833 | -6.098551 |
| 24.1042 | 2.125608 | | 24.0972 | -0.888705 |
| 24.1181 | 1.326946 | | 24.1111 | 1.632764 |
| 24.1319 | 0.68041 | | 24.125 | 0.806726 |
| 24.1458 | 0.476525 | | 24.1389 | -0.185736 |
| 24.1597 | 0.427845 | | 24.1528 | -0.406946 |
| 24.1736 | 0.340443 | | 24.1667 | -0.270812 |
| 24.1875 | 0.196912 | | 24.1806 | -0.288807 |
| 24.2014 | -0.007294 | | 24.1944 | -0.349801 |
| 24.2153 | -0.19924 | | 24.2083 | -0.176778 |
| 24.2292 | -0.32358 | | 24.2222 | -0.094765 |
| 24.2431 | -0.35517 | | 24.2361 | -0.255938 |
| 24.2569 | -0.297602 | | 24.25 | -0.488193 |
| 24.2708 | -0.19048 | | 24.2639 | -0.591479 |
| 24.2847 | -0.050128 | | 24.2778 | -0.555869 |
| 24.2986 | 0.099778 | | 24.2917 | -0.425641 |
| 24.3125 | 0.270602 | | 24.3056 | -0.276271 |
| 24.3264 | 0.450872 | | 24.3194 | -0.054324 |
| 24.3403 | 0.608189 | | 24.3333 | 0.245665 |
| 24.3542 | 0.621061 | | 24.3472 | 0.578649 |
| 24.3681 | 0.394099 | | 24.3611 | 0.65944 |
| 24.3819 | 0.131739 | | 24.375 | 0.512771 |
| 24.3958 | -0.107774 | | 24.3889 | 0.469682 |
| 24.4097 | -0.189551 | | 24.4028 | 0.370753 |
| 24.4236 | 0.110556 | | 24.4167 | -0.110243 |
| 24.4375 | 1.438168 | | 24.4306 | -1.749272 |
| 24.4514 | 4.797808 | | 24.4444 | -3.796706 |
| 24.4653 | 7.756726 | | 24.4583 | -2.121362 |
| 24.4792 | 3.854534 | | 24.4722 | 1.518531 |
| 24.4931 | 0.72227 | | 24.4861 | 2.07157 |
| 24.5 | 1.248236 | | 24.5 | 0.979126 |
| 24.5139 | 1.978703 | | 24.5139 | 0.388889 |
| 24.5278 | 1.284942 | | 24.5278 | 0.281061 |
| 24.5417 | -0.482557 | | 24.5417 | 0.284948 |
| 24.5556 | -0.971515 | | 24.5556 | 0.288099 |
| 24.5694 | -0.663143 | | 24.5694 | 0.257713 |
| 24.5833 | -0.851213 | | 24.5833 | 0.219785 |
| 24.5972 | -1.090948 | | 24.5972 | 0.190525 |
| 24.6111 | -0.997345 | | 24.6111 | 0.173046 |
| 24.625 | 0.712172 | | 24.625 | 0.163635 |
| 24.6389 | 2.773693 | | 24.6389 | 0.175007 |
| 24.6528 | 2.323296 | | 24.6528 | 0.229763 |
| 24.6667 | -0.125532 | | 24.6667 | 0.32931 |
| 24.6806 | -1.612105 | | 24.6806 | 0.323903 |
| 24.6944 | -1.191516 | | 24.6944 | 0.218012 |
| 24.7083 | -0.15548 | | 24.7083 | 0.172658 |
| 24.7222 | 0.38398 | | 24.7222 | 0.164011 |
| 24.7361 | -0.569296 | | 24.7361 | 0.137469 |
| 24.75 | -2.918578 | | 24.75 | 0.1255 |
| 24.7639 | -3.962773 | | 24.7639 | 0.143753 |
| 24.7778 | -2.717731 | | 24.7778 | 0.147239 |
| 24.7917 | -0.710398 | | 24.7917 | 0.140491 |
| 24.8056 | -0.291269 | | 24.8056 | 0.172332 |
| 24.8194 | -0.816753 | | 24.8194 | 0.268027 |
| 24.8333 | -0.584046 | | 24.8333 | 0.36074 |
| 24.8472 | -0.084852 | | 24.8472 | 0.388982 |
| 24.8611 | 0.280366 | | 24.8611 | 0.31878 |
| 24.875 | 0.675298 | | 24.875 | 0.192494 |
| 24.8889 | 0.480621 | | 24.8889 | 0.142045 |
| 24.9028 | -0.385591 | | 24.9028 | 0.235404 |
| 24.9167 | -0.552821 | | 24.9167 | 0.651452 |
| 24.9306 | -2.390349 | | 24.9306 | 2.342168 |
| 24.9444 | -6.570839 | | 24.9444 | 3.654744 |
| 24.9583 | -6.550688 | | 24.9583 | 1.432836 |
| 24.9722 | -3.646739 | | 24.9722 | -1.192401 |
| 24.9861 | -1.517891 | | 24.9861 | -0.741832 |
| 25.0069 | -0.648315 | | 25 | 0.460361 |
| 25.0208 | -0.117875 | | 25.0139 | 0.793938 |
| 25.0347 | 0.078556 | | 25.0278 | 0.887975 |
| 25.0486 | -0.107672 | | 25.0417 | 1.178613 |
| 25.0625 | -0.36646 | | 25.0556 | 1.419334 |
| 25.0764 | -0.310906 | | 25.0694 | 1.087528 |
| 25.0903 | 0.058179 | | 25.0833 | 0.063508 |
| 25.1042 | 0.241717 | | 25.0972 | -0.871757 |
| 25.1181 | 0.340928 | | 25.1111 | -0.851395 |
| 25.1319 | 0.701255 | | 25.125 | 0.110345 |
| 25.1458 | 1.418081 | | 25.1389 | 0.707418 |
| 25.1597 | 2.078999 | | 25.1528 | 0.409879 |
| 25.1736 | 1.652432 | | 25.1667 | 0.096307 |
| 25.1875 | 0.494809 | | 25.1806 | 0.11692 |
| 25.2014 | -0.482919 | | 25.1944 | 0.686971 |
| 25.2153 | -0.780772 | | 25.2083 | 1.191264 |
| 25.2292 | -0.264501 | | 25.2222 | 0.708603 |
| 25.2431 | 0.495506 | | 25.2361 | 0.167345 |
| 25.2569 | 0.810007 | | 25.25 | 0.055595 |
| 25.2708 | 0.295892 | | 25.2639 | 0.054499 |
| 25.2847 | -0.519122 | | 25.2778 | 0.037303 |
| 25.2986 | -0.606618 | | 25.2917 | -0.396033 |
| 25.3125 | 0.103343 | | 25.3056 | -1.096252 |
| 25.3264 | 0.665509 | | 25.3194 | -0.636101 |
| 25.3403 | 0.561951 | | 25.3333 | 0.413102 |
| 25.3542 | 0.552432 | | 25.3472 | 0.311067 |
| 25.3681 | 0.866802 | | 25.3611 | 0.337685 |
| 25.3819 | 0.367722 | | 25.375 | 1.518675 |
| 25.3958 | -0.606251 | | 25.3889 | 1.942599 |
| 25.4097 | -0.516828 | | 25.4028 | 1.186818 |
| 25.4236 | 0.591784 | | 25.4167 | 0.165479 |
| 25.4375 | 3.056671 | | 25.4306 | -2.277056 |
| 25.4514 | 7.705987 | | 25.4444 | -5.132402 |
| 25.4653 | 9.127565 | | 25.4583 | -2.114478 |
| 25.4792 | 5.828229 | | 25.4722 | 2.685205 |
| 25.4931 | 2.200577 | | 25.4861 | 3.082954 |
| 25.5 | 1.322904 | | 25.5 | 2.158808 |
| 25.5139 | 0.431553 | | 25.5139 | 1.580097 |
| 25.5278 | -0.302161 | | 25.5278 | 1.057313 |
| 25.5417 | -0.42987 | | 25.5417 | 0.438724 |
| 25.5556 | -0.025653 | | 25.5556 | -0.147635 |
| 25.5694 | 0.331406 | | 25.5694 | -0.331505 |
| 25.5833 | 0.547121 | | 25.5833 | -0.423156 |
| 25.5972 | 0.36867 | | 25.5972 | -0.593454 |
| 25.6111 | 0.028635 | | 25.6111 | -0.695273 |
| 25.625 | -0.030197 | | 25.625 | -0.681653 |
| 25.6389 | -0.642314 | | 25.6389 | -0.67123 |
| 25.6528 | -1.688884 | | 25.6528 | -0.430748 |
| 25.6667 | -2.115257 | | 25.6667 | -0.091481 |
| 25.6806 | -1.480796 | | 25.6806 | 0.232725 |
| 25.6944 | 0.012667 | | 25.6944 | 0.570458 |
| 25.7083 | 0.965109 | | 25.7083 | 0.81709 |
| 25.7222 | 1.147518 | | 25.7222 | 0.721427 |
| 25.7361 | 0.555177 | | 25.7361 | 0.391928 |
| 25.75 | -0.246187 | | 25.75 | -0.226663 |
| 25.7639 | -1.429899 | | 25.7639 | -0.969074 |
| 25.7778 | -2.545313 | | 25.7778 | -1.389697 |
| 25.7917 | -2.666984 | | 25.7917 | -1.324809 |
| 25.8056 | -1.958123 | | 25.8056 | -1.074406 |
| 25.8194 | -1.513557 | | 25.8194 | -0.905534 |
| 25.8333 | -1.15949 | | 25.8333 | -0.741378 |
| 25.8472 | -0.676751 | | 25.8472 | -0.523961 |
| 25.8611 | -0.217546 | | 25.8611 | -0.38914 |
| 25.875 | 0.141694 | | 25.875 | -0.164437 |
| 25.8889 | 0.311123 | | 25.8889 | 0.055631 |
| 25.9028 | 0.416312 | | 25.9028 | 0.163106 |
| 25.9167 | 0.260529 | | 25.9167 | 0.26554 |
| 25.9306 | -2.558207 | | 25.9306 | 0.810773 |
| 25.9444 | -7.611001 | | 25.9444 | -0.131833 |
| 25.9583 | -8.841981 | | 25.9583 | -1.436453 |
| 25.9722 | -5.625247 | | 25.9722 | -1.467108 |
| 25.9861 | -1.410816 | | 25.9861 | -0.633053 |
| 26.0069 | 0.100266 | | 26 | -0.047162 |
| 26.0208 | -1.891923 | | 26.0139 | 0.278648 |
| 26.0347 | -0.149799 | | 26.0278 | 0.883866 |
| 26.0486 | 1.25444 | | 26.0417 | 0.556676 |
| 26.0625 | 1.402376 | | 26.0556 | -0.42682 |
| 26.0764 | 0.839485 | | 26.0694 | -0.404216 |
| 26.0903 | 0.564371 | | 26.0833 | 0.545118 |
| 26.1042 | 0.599205 | | 26.0972 | 0.594962 |
| 26.1181 | 0.5163 | | 26.1111 | -0.263263 |
| 26.1319 | 0.396096 | | 26.125 | -0.314622 |
| 26.1458 | 0.386681 | | 26.1389 | -0.030075 |
| 26.1597 | 0.493546 | | 26.1528 | -0.311827 |
| 26.1736 | 0.619201 | | 26.1667 | -0.115274 |
| 26.1875 | 0.659824 | | 26.1806 | 0.685672 |
| 26.2014 | 0.678209 | | 26.1944 | 0.679697 |
| 26.2153 | 0.576475 | | 26.2083 | 0.045675 |
| 26.2292 | 0.422556 | | 26.2222 | -0.434855 |
| 26.2431 | 0.403896 | | 26.2361 | -0.40182 |
| 26.2569 | 0.526107 | | 26.25 | 0.376825 |
| 26.2708 | 0.367592 | | 26.2639 | 0.647197 |
| 26.2847 | -0.217707 | | 26.2778 | 0.010497 |
| 26.2986 | -0.700971 | | 26.2917 | -0.24252 |
| 26.3125 | -0.580895 | | 26.3056 | -0.364436 |
| 26.3264 | 0.06819 | | 26.3194 | -0.316903 |
| 26.3403 | 0.630353 | | 26.3333 | 0.44563 |
| 26.3542 | 0.682636 | | 26.3472 | 0.935135 |
| 26.3681 | 0.327297 | | 26.3611 | 1.127401 |
| 26.3819 | -0.109504 | | 26.375 | 0.975133 |
| 26.3958 | -0.452196 | | 26.3889 | 0.359179 |
| 26.4097 | -0.348824 | | 26.4028 | 0.449086 |
| 26.4236 | 0.203082 | | 26.4167 | 0.681513 |
| 26.4375 | 1.724872 | | 26.4306 | -0.527134 |
| 26.4514 | 3.811692 | | 26.4444 | -1.452449 |
| 26.4653 | 5.588824 | | 26.4583 | 2.221281 |
| 26.4792 | 5.133405 | | 26.4722 | 4.352613 |
| 26.4931 | 3.644705 | | 26.4861 | 3.894209 |
| 26.5 | 2.779415 | | 26.5 | 2.948534 |
| 26.5139 | 1.293871 | | 26.5139 | 1.695473 |
| 26.5278 | -0.447359 | | 26.5278 | 0.414118 |
| 26.5417 | -1.687042 | | 26.5417 | -0.713584 |
| 26.5556 | -1.878115 | | 26.5556 | -1.391688 |
| 26.5694 | -1.360041 | | 26.5694 | -1.42665 |
| 26.5833 | -0.699786 | | 26.5833 | -1.064368 |

| | | | | | | | | | | | | | | | | | | | | | | | |
|---|---|---|---|---|---|---|---|---|---|---|---|---|---|---|---|---|---|---|---|---|---|---|---|
| 26.5972 | -0.543864 | 0.036584 | -0.999572 | 0.120936 | -0.178515 | -1.925226 | 0.514502 | 0.044808 | 0.857143 | 0.107379 | -0.312712 | 0.033845 | 0.242176 | 0.03792 | -0.009224 | -0.230177 | -1.427485 | -0.99237 | 0.634416 | -0.263698 | -1.016693 | -0.030773 | -0.599435 |
| 26.6111 | 0.113476 | 0.0391 | -0.014874 | 0.439203 | 0.082822 | -1.807686 | 1.280531 | -0.135011 | 1.171181 | 0.03795 | -0.175006 | -0.642757 | -0.253299 | 0.377097 | -0.063385 | -0.166217 | -1.40259 | -0.800216 | 0.358037 | -0.596133 | -0.943027 | -0.011309 | -0.482277 |
| 26.625 | 1.210969 | 0.101535 | 1.262087 | 1.005504 | 0.690549 | -0.885507 | 2.1079 | -0.195399 | 0.461557 | -0.051025 | 0.010739 | -1.260473 | -0.75442 | 0.128341 | -0.621685 | -0.438957 | -1.170375 | -0.668553 | -0.362367 | -0.618245 | -0.748342 | 0.041082 | -0.840107 |
| 26.6389 | 1.856173 | 0.099757 | 1.352807 | 1.35233 | 1.02518 | 0.130308 | 1.68201 | -0.245158 | -0.075477 | -0.063384 | 0.359717 | -1.618181 | -1.036435 | -0.381192 | -1.067435 | -0.701657 | -0.832069 | -0.381153 | -1.031895 | -0.27384 | -0.461479 | 0.078089 | -1.115229 |
| 26.6528 | 1.443034 | -0.04342 | 0.246075 | 0.595274 | 0.866623 | 1.1338 | 0.032285 | -0.225293 | -0.41945 | -0.011899 | 0.543655 | -1.196737 | -0.650828 | -0.257883 | -0.814752 | -0.722805 | -0.557009 | -0.04596 | -1.245195 | -0.130903 | -0.110749 | 0.052292 | -0.906543 |
| 26.6667 | 0.292947 | -0.528659 | -0.099211 | -0.070633 | 0.568158 | 1.336539 | -1.044224 | -0.099439 | -0.66988 | 0.032333 | 0.535226 | 0.060451 | -0.091991 | -0.470866 | -0.149067 | -0.430201 | -0.253965 | 0.284941 | -0.933481 | -0.231007 | 0.274395 | -0.018912 | -0.220062 |
| 26.6806 | -0.323118 | -0.77855 | -0.329567 | -0.249236 | 0.129974 | 1.256277 | -1.702426 | -0.006524 | -0.955152 | 0.034105 | 0.282307 | 1.424753 | 0.424158 | -0.206593 | 0.342783 | -0.078114 | 0.04494 | 0.54647 | -0.130779 | -0.428793 | 0.867922 | -0.030478 | 0.563846 |
| 26.6944 | -0.312242 | -0.430371 | -0.65695 | 0.059876 | -0.09903 | 1.116046 | -2.064057 | 0.015183 | -0.823713 | 0.021183 | -0.013065 | 2.036859 | 0.530435 | 0.235229 | 0.452259 | 0.202454 | 0.192756 | 0.535573 | 0.581363 | -0.655455 | 1.396432 | 0.019721 | 1.172301 |
| 26.7083 | -0.404955 | 0.020083 | -0.5539 | -0.041093 | -0.323766 | 0.662118 | -1.368758 | -0.029269 | -0.624345 | 0.007885 | -0.4927 | 1.507923 | 0.108152 | 0.435389 | 0.235727 | 0.315641 | 0.138656 | 0.192233 | 0.810343 | -0.594479 | 1.348388 | 0.054734 | 1.320527 |
| 26.7222 | -0.921214 | 0.156002 | -0.351785 | -0.619644 | -0.897587 | -0.235513 | -0.486611 | -0.12245 | -0.357667 | -0.004449 | -1.13567 | 0.462362 | -0.502484 | 0.142171 | -0.128996 | 0.199194 | -0.03476 | -0.392528 | 0.354737 | -0.056774 | 0.896021 | 0.058246 | 0.891098 |
| 26.7361 | -2.115129 | 0.096796 | -0.489477 | -1.297251 | -0.803914 | -1.306559 | -0.004509 | -0.216847 | -0.004496 | -0.000865 | -1.476234 | -0.752377 | -0.83319 | -0.268587 | -0.284275 | -0.076911 | -0.350814 | -0.94495 | -0.297122 | 0.149853 | 0.315911 | 0.071939 | 0.033368 |
| 26.75 | -1.40507 | -0.077838 | -0.71927 | -0.721422 | -0.422516 | -1.811159 | -0.026583 | -0.193623 | 0.334469 | 0.011143 | -1.539493 | -1.132341 | -0.546727 | -0.682285 | -0.364631 | -0.25434 | -0.724508 | -1.286304 | -0.855991 | 0.026985 | -0.334387 | 0.079454 | -0.773418 |
| 26.7639 | -0.531456 | -0.346013 | -0.533688 | -0.169956 | -0.175922 | -2.206057 | -0.046132 | -0.154482 | 0.456533 | -0.004158 | -1.570938 | -0.662568 | -0.237483 | -1.121404 | -0.844625 | -0.414051 | -1.01955 | -1.342938 | -1.249605 | -0.148689 | -0.943354 | 0.028448 | -1.22176 |
| 26.7778 | -0.491934 | -0.570128 | -0.010283 | -0.637685 | -0.193286 | -2.378556 | -0.066792 | -0.017841 | 0.229381 | 0.00814 | -1.536587 | -0.298546 | -0.288544 | -1.067139 | -1.410813 | -0.580588 | -1.090303 | -1.082967 | -0.875646 | -0.341567 | -1.33126 | -0.061655 | -1.516609 |
| 26.7917 | -0.602538 | -0.519524 | 0.237288 | -1.159267 | -0.016129 | -2.529541 | -0.240273 | 0.052717 | -0.414667 | 0.016835 | -1.556245 | -0.74209 | -0.686897 | -0.78136 | -1.185581 | -0.70245 | -0.832496 | -0.852507 | -0.452983 | -0.35561 | -1.473926 | -0.067782 | -1.532317 |
| 26.8056 | -0.550609 | -0.15716 | 0.183393 | -1.232975 | 0.700675 | -1.581349 | -0.6785 | -0.004367 | -0.891988 | 0.012468 | -1.519636 | -1.417571 | -1.051342 | -0.578664 | -0.824901 | -0.710794 | -0.41456 | -0.774039 | -0.212111 | -0.059909 | -1.226826 | 0.028135 | -1.366992 |
| 26.8194 | -0.414757 | 0.13293 | -0.159845 | -1.007134 | 0.979873 | -0.839372 | -1.094326 | -0.110701 | -1.141485 | -0.023287 | -1.281653 | -1.808581 | -1.015968 | -0.49725 | -0.593556 | -0.71669 | -0.158442 | -0.796167 | -0.353254 | 0.11657 | -0.8457 | 0.102068 | -1.30216 |
| 26.8333 | -0.365465 | 0.221742 | -0.521071 | -0.827794 | 0.648707 | -0.608038 | -1.314345 | -0.196119 | -1.016064 | -0.045965 | -1.168657 | -1.718328 | -0.612729 | -0.496158 | -0.485121 | -0.808866 | -0.041462 | -0.799615 | -0.726258 | -0.12421 | -0.584826 | 0.0916 | -1.287941 |
| 26.8472 | -0.540138 | 0.085255 | -0.630495 | -0.808316 | 0.234622 | -0.398605 | -1.183403 | -0.269676 | -0.794398 | -0.021471 | -1.030056 | -1.024888 | -0.320301 | -0.328768 | -0.430031 | -0.740046 | 0.052276 | -0.644138 | -0.875499 | -0.483033 | -0.530167 | 0.107744 | -1.261327 |
| 26.8611 | -0.830984 | -0.157717 | -0.464897 | -1.115075 | -0.155776 | -0.111077 | -0.720229 | -0.317961 | -0.674116 | 0.019783 | -0.768746 | -0.4932 | -0.35098 | -0.206191 | -0.50771 | -0.565716 | 0.10921 | -0.38248 | -0.898912 | -0.716158 | -0.515791 | 0.119648 | -1.144603 |
| 26.875 | -0.798517 | -0.246973 | -0.140883 | -1.331091 | -0.426201 | 0.13606 | -0.270543 | -0.463588 | -0.573884 | 0.016794 | -0.366442 | -0.392855 | -0.601552 | -0.058027 | -0.459742 | -0.350993 | 0.15509 | -0.200922 | -0.648805 | -0.721485 | -0.563113 | 0.045593 | -0.91726 |
| 26.8889 | -0.338006 | -0.101207 | 0.226605 | -0.979444 | -0.734558 | 0.239277 | 0.19589 | -0.522399 | -0.283569 | 0.001844 | 0.149997 | -0.548462 | -0.70414 | 0.03231 | -0.158611 | -0.207579 | 0.183371 | -0.141211 | -0.31112 | -0.528969 | -0.445359 | -0.049722 | -0.618672 |
| 26.9028 | 0.166699 | 0.122373 | 0.380211 | -0.410005 | -0.686795 | 0.233763 | 0.368789 | -0.498814 | 0.142022 | -0.002088 | 0.470904 | -0.804528 | -0.56259 | 0.044074 | 0.126065 | -0.163384 | 0.114512 | -0.116078 | 0.032582 | -0.215153 | -0.404266 | -0.096224 | -0.40829 |

| | | | | |
|---|---|---|---|---|
| 26.5972 | -0.312712 | | 26.5972 | -0.599435 |
| 26.6111 | -0.175006 | | 26.6111 | -0.482277 |
| 26.625 | 0.010739 | | 26.625 | -0.840107 |
| 26.6389 | 0.359717 | | 26.6389 | -1.115229 |
| 26.6528 | 0.543655 | | 26.6528 | -0.906543 |
| 26.6667 | 0.535226 | | 26.6667 | -0.220062 |
| 26.6806 | 0.282307 | | 26.6806 | 0.563846 |
| 26.6944 | -0.013065 | | 26.6944 | 1.172301 |
| 26.7083 | -0.4927 | | 26.7083 | 1.320527 |
| 26.7222 | -1.13567 | | 26.7222 | 0.891098 |
| 26.7361 | -1.476234 | | 26.7361 | 0.033368 |
| 26.75 | -1.539493 | | 26.75 | -0.773418 |
| 26.7639 | -1.570938 | | 26.7639 | -1.22176 |
| 26.7778 | -1.536587 | | 26.7778 | -1.516609 |
| 26.7917 | -1.556245 | | 26.7917 | -1.532317 |
| 26.8056 | -1.519636 | | 26.8056 | -1.366992 |
| 26.8194 | -1.281653 | | 26.8194 | -1.30216 |
| 26.8333 | -1.168657 | | 26.8333 | -1.287941 |
| 26.8472 | -1.030056 | | 26.8472 | -1.261327 |
| 26.8611 | -0.768746 | | 26.8611 | -1.144603 |
| 26.875 | -0.366442 | | 26.875 | -0.91726 |
| 26.8889 | 0.149997 | | 26.8889 | -0.618672 |
| 26.9028 | 0.470904 | | 26.9028 | -0.40829 |

| | | | | | | | | | | | | | | | | | | | | | | | |
|---|---|---|---|---|---|---|---|---|---|---|---|---|---|---|---|---|---|---|---|---|---|---|---|
| 29.2292 | 0.905534 | 0.575021 | -0.020713 | 0.729972 | 0.190793 | 0.424747 | 0.371474 | 0.213415 | 1.496572 | 0.284254 | 0.882126 | 0.404879 | -0.13503 | 0.35102 | -0.036082 | 0.295815 | 0.079754 | -0.026432 | -0.023506 | -0.560962 | 0.091008 | 0.348168 | 0.120248 |
| 29.2431 | 0.829091 | 0.402363 | -0.035044 | 0.495317 | 0.080128 | 0.167795 | 0.323158 | 0.042076 | 1.337538 | 0.152021 | 0.667301 | 0.196739 | -0.380747 | 0.293072 | 0.147227 | 0.104547 | 0.146268 | 0.111648 | 0.05759 | -0.479954 | -0.218149 | -0.174574 | 0.011515 |
| 29.2569 | 0.662484 | 0.260533 | -0.04538 | 0.26787 | -0.030763 | 0.088031 | 0.175831 | -0.066971 | 1.144531 | 0.062703 | 0.469028 | 0.149467 | -0.431081 | 0.092953 | 0.392482 | -0.001384 | 0.254744 | 0.341939 | 0.232209 | -0.242342 | -0.467427 | 0.032769 | -0.004519 |
| 29.2708 | 0.605469 | 0.253126 | -0.081429 | 0.191241 | -0.169318 | 0.0895 | 0.064905 | 0.008636 | 1.158375 | 0.051468 | 0.280801 | 0.139897 | -0.15054 | -0.191169 | 0.560339 | 0.042273 | 0.412417 | 0.475004 | 0.279754 | 0.044229 | -0.32934 | 0.090603 | -0.097993 |
| 29.2847 | 0.684168 | 0.438817 | -0.113896 | 0.264753 | -0.39511 | 0.058247 | -0.006687 | 0.080625 | 1.172395 | -0.065663 | 0.273789 | 0.137199 | 0.012258 | -0.420393 | 0.360138 | 0.130517 | 0.253692 | 0.252237 | 0.181434 | -0.159491 | -0.161816 | 0.039635 | -0.15448 |
| 29.2986 | 0.560308 | 0.581512 | -0.05038 | 0.327787 | -0.620669 | -0.02689 | 0.049371 | 0.050043 | 1.143922 | -0.171546 | 0.19917 | 0.170136 | 0.015477 | -0.224895 | -0.087985 | 0.290449 | -0.091703 | -0.135424 | 0.03908 | -0.662252 | -0.112141 | -0.201642 | -0.071849 |
| 29.3125 | 0.330844 | 0.609586 | 0.296896 | 0.465101 | -0.612803 | 0.035103 | 0.318291 | 0.135053 | 1.220265 | -0.128775 | 0.248298 | 0.126333 | 0.101924 | 0.166553 | -0.425908 | 0.362448 | -0.199771 | -0.2876 | -0.145635 | -0.806566 | -0.039488 | -0.22486 | 0.033527 |
| 29.3264 | 0.015232 | 0.51837 | 0.649941 | 0.617537 | -0.47522 | 0.155088 | 0.625988 | 0.204526 | 1.227491 | -0.026761 | 0.625305 | 0.212526 | 0.379142 | 0.546074 | -0.293983 | 0.243467 | 0.005957 | -0.06303 | -0.07145 | -0.495419 | 0.108637 | -0.105074 | 0.117861 |
| 29.3403 | -0.258477 | 0.262336 | 0.861242 | 0.650033 | -0.366454 | 0.114848 | 0.762653 | -0.00126 | 0.97374 | 0.000289 | 0.680728 | 0.694312 | 0.747781 | 0.911294 | 0.208631 | 0.160478 | 0.210342 | 0.276719 | 0.438129 | -0.013471 | 0.287107 | 0.120662 | 0.268761 |
| 29.3542 | -0.24468 | 0.042273 | 0.915413 | 0.557061 | -0.258778 | 0.045845 | 0.806924 | -0.206644 | 0.697531 | -0.068535 | 0.31314 | 0.999989 | 0.802796 | 0.945 | 0.624382 | 0.078865 | 0.204534 | 0.312748 | 0.841949 | 0.162977 | 0.357347 | 0.216945 | 0.293717 |
| 29.3681 | -0.011561 | 0.053024 | 0.702162 | 0.448767 | -0.158577 | 0.09338 | 0.686167 | -0.170142 | 0.448863 | -0.03675 | 0.274008 | 0.782803 | 0.491622 | 0.483384 | 0.746942 | 0.037459 | 0.100658 | 0.160688 | 0.755651 | 0.063388 | 0.319341 | 0.034692 | 0.153899 |
| 29.3819 | 0.235709 | 0.109273 | 0.460171 | 0.371637 | -0.065284 | 0.075679 | 0.364642 | -0.029388 | 0.191963 | 0.081348 | 0.384583 | 0.517885 | 0.271583 | 0.114061 | 0.572143 | 0.241731 | 0.107779 | 0.115527 | 0.467723 | -0.071821 | 0.362704 | -0.059786 | 0.045853 |
| 29.3958 | 0.603809 | 0.227153 | 0.367235 | 0.458989 | 0.068309 | 0.059088 | 0.221151 | 0.099939 | 0.095569 | 0.146112 | 1.139616 | 0.487984 | 0.346071 | 0.08289 | 0.350117 | 0.57008 | 0.172823 | 0.148768 | 0.302035 | -0.194966 | 0.478612 | 0.043916 | 0.041398 |
| 29.4097 | 1.662704 | 0.936816 | 0.332422 | 0.988633 | 0.511809 | 0.995901 | 0.308809 | 0.819962 | 0.234816 | 0.76615 | 7.38184 | 0.566427 | 0.532184 | 0.30005 | 0.22249 | 0.656871 | 0.196211 | 0.134896 | 0.320308 | -0.267498 | 0.509831 | 0.090137 | 0.047808 |
| 29.4236 | 3.287478 | 2.107967 | 0.360698 | 2.013041 | 1.475009 | 2.627809 | 0.159985 | 2.325598 | 0.908313 | 2.593562 | 18.24158 | 0.635049 | 0.674721 | 0.616231 | 0.320211 | 0.492699 | 0.232276 | 0.184807 | 0.462032 | -0.128112 | 0.564655 | 0.002655 | 0.145807 |
| 29.4375 | 3.625992 | 3.052749 | 1.007227 | 2.410649 | 2.634991 | 2.464517 | 0.448575 | 3.153978 | 2.248376 | 0.31551 | 19.169591 | 0.566562 | 1.084617 | 0.923531 | 1.048233 | 0.261252 | 0.707571 | 0.499589 | 0.544657 | 0.580313 | 1.284167 | 0.053237 | 0.162184 |
| 29.4514 | 2.803588 | 1.977051 | 2.223898 | 2.362979 | 3.02969 | 1.59521 | 1.778784 | 2.26425 | 3.081173 | -0.578581 | 10.226039 | 0.647633 | 1.949649 | 0.956389 | 2.366564 | 0.352696 | 1.861011 | 1.023678 | 0.474059 | 1.364565 | 2.806235 | 2.348863 | 0.500808 |
| 29.4653 | 1.831147 | 1.029453 | 2.879868 | 2.748408 | 2.243409 | 1.107958 | 2.487956 | 1.275722 | 2.505229 | -0.001388 | 3.725783 | 1.158458 | 2.444901 | 1.144385 | 3.014885 | 1.012251 | 2.276216 | 0.975538 | 0.581007 | -0.217605 | 3.162815 | 3.136849 | -0.316894 |
| 29.4792 | 1.803777 | 0.909079 | 3.632305 | 2.576469 | 1.662379 | 0.800344 | 2.330648 | 0.883006 | 2.834189 | -0.106968 | 2.840394 | 1.985338 | 2.227689 | 1.069101 | 1.717943 | 1.371538 | 1.245801 | 0.82352 | 1.104664 | -1.765296 | 2.325876 | 1.488152 | 1.095185 |
| 29.4931 | 1.223963 | 0.841222 | 2.737848 | 1.841104 | 1.221357 | 0.650534 | 1.5848 | 0.982696 | 2.516954 | 0.131526 | 2.109937 | 1.577521 | 1.731552 | 1.220459 | 1.043056 | 1.270602 | 0.252317 | 0.311344 | 0.938093 | -2.473007 | 1.263894 | 0.080343 | 1.564364 |
| 29.5 | 0.533968 | 0.806669 | 0.929239 | 1.107657 | 1.200971 | 0.475678 | 1.015014 | 0.754731 | 1.32537 | 0.151917 | 1.280231 | 1.309168 | 1.492456 | 1.484046 | 1.028607 | 1.150153 | 0.014934 | 0.239136 | 0.760643 | -1.648358 | 1.318964 | -0.019064 | 1.313617 |
| 29.5139 | -0.148598 | 0.829932 | -0.121823 | 0.593861 | 1.364943 | 0.431797 | 0.747828 | 0.096343 | 0.354979 | 0.112141 | 0.432297 | 1.114211 | 1.047502 | 1.041067 | 0.716482 | 0.695002 | 0.003214 | 0.058554 | 0.398126 | -0.86316 | 0.971363 | 0.026974 | 1.113898 |
| 29.5278 | -0.469962 | 0.704367 | -1.265594 | 0.083327 | 1.350785 | 0.403147 | 0.532357 | -0.454393 | -0.250699 | 0.08616 | -0.460737 | 0.755515 | -0.027644 | 0.013483 | 0.378485 | 0.102333 | -0.094712 | -0.13564 | -0.093106 | -0.75193 | 0.461071 | 0.013696 | 0.938101 |
| 29.5417 | -0.649921 | 0.446809 | -2.36963 | -0.057708 | 0.979588 | 0.478768 | 0.648269 | -0.23145 | -0.291944 | 0.014022 | -1.418666 | 0.27905 | -1.379397 | -0.90405 | 0.058261 | -0.450238 | -0.127623 | -0.243406 | -0.39386 | -0.98801 | -0.15612 | -0.034286 | 0.557314 |
| 29.5556 | -0.768569 | -0.04215 | -2.265685 | -0.083742 | 0.596874 | 0.614013 | 0.721476 | 0.114043 | -0.335698 | 0.15461 | -2.368124 | -0.036777 | -2.742318 | -1.313062 | -0.206142 | -0.765727 | -0.195718 | -0.250753 | -0.492245 | -1.163107 | -0.662903 | -0.062369 | 0.282745 |
| 29.5694 | -0.882316 | -0.747249 | -2.088674 | 0.176155 | -0.007751 | -0.261527 | 0.708244 | 0.214614 | -0.534078 | -0.183522 | -2.379636 | -0.068442 | -2.799782 | -1.295675 | -0.259282 | -0.845182 | -0.164999 | -0.174328 | -0.390298 | -1.043551 | -0.69822 | -0.044225 | 0.153781 |
| 29.5833 | -0.639392 | -1.285044 | -1.145522 | 0.357861 | -0.573307 | -0.761822 | 0.466274 | 0.259854 | -0.465467 | 0.010146 | -1.949029 | -0.048859 | -1.216312 | -0.930047 | -0.090568 | -0.662684 | -0.057173 | -0.026124 | -0.434742 | -0.759322 | -0.328008 | -0.042549 | 0.224034 |
| 29.5972 | 0.064631 | -1.359029 | 0.575346 | 0.573122 | -0.987958 | -0.315533 | 0.192431 | 0.265104 | 0.038314 | -0.124651 | -0.834901 | 0.044512 | 0.195343 | -0.245916 | 0.10013 | -0.347517 | 0.043865 | 0.068184 | -0.675589 | -0.499839 | 0.103521 | -0.052371 | 0.29734 |
| 29.6111 | 0.66499 | -1.220048 | 1.647015 | 0.56977 | -1.328125 | -0.068469 | -0.112951 | 0.231016 | 0.206578 | -0.133395 | 0.111051 | 0.340684 | 1.303688 | 0.293165 | 0.131029 | 0.025477 | -0.022763 | -0.049104 | -0.915831 | -0.309234 | 0.246141 | -0.03151 | 0.351288 |
| 29.625 | 0.760187 | -1.040165 | 2.200291 | 0.625338 | -1.62009 | 0.288705 | -0.421401 | 0.186497 | 0.351042 | -0.090952 | 0.764243 | 0.578282 | 1.534686 | 0.576447 | -0.003562 | 0.286487 | -0.172592 | -0.210854 | -0.789414 | -0.019131 | 0.01942 | -0.020727 | 0.372204 |
| 29.6389 | 0.634908 | -1.002915 | 2.029582 | 0.641894 | -1.745997 | 0.743019 | -0.609052 | 0.10972 | 0.642103 | -0.080798 | 0.715467 | 0.583111 | 0.85256 | 0.669516 | -0.145707 | 0.375809 | -0.202202 | -0.251905 | -0.178125 | 0.170383 | -0.271999 | 0.006438 | 0.288413 |
| 29.6528 | 0.391044 | -0.820032 | 1.866052 | 0.846305 | -1.096502 | 0.452048 | -0.887221 | 0.032995 | 0.768424 | -0.134189 | 0.132647 | 0.394761 | -0.172115 | 0.596644 | -0.175354 | 0.34126 | -0.103262 | -0.058952 | 0.540325 | 0.13868 | -0.269614 | 0.063235 | 0.166636 |
| 29.6667 | 0.151977 | -0.526144 | 1.254861 | 0.87111 | -0.147177 | 0.001321 | -1.118514 | 0.0091 | 0.467437 | -0.01081 | -0.041365 | 0.250139 | -1.044448 | 0.452433 | -0.008564 | 0.317995 | 0.189926 | 0.181733 | 0.979545 | -0.081287 | 0.032224 | 0.05818 | 0.056731 |
| 29.6806 | -0.246426 | -0.27991 | 0.602617 | 0.621048 | 0.681934 | -0.004429 | -1.091571 | -0.009289 | -0.234592 | 0.005325 | 0.047915 | 0.161755 | -1.381961 | 0.219241 | 0.162652 | 0.21908 | 0.294234 | 0.270371 | 0.878682 | -0.375381 | 0.297024 | 0.018301 | -0.011515 |
| 29.6944 | -0.704391 | -0.409501 | -0.202599 | 0.242795 | 1.364726 | 0.055661 | -0.868101 | -0.147874 | -0.476701 | -0.030769 | -0.240768 | 0.074364 | -1.204844 | -0.049273 | 0.233598 | -0.04923 | 0.168244 | 0.212294 | 0.524008 | -0.571966 | 0.303034 | 0.000393 | -0.005018 |
| 29.7083 | -1.209094 | -0.95464 | -1.384036 | -0.261981 | 1.447421 | -0.177612 | -0.66724 | -0.608569 | -0.789623 | -0.030537 | -2.133119 | -0.068623 | -0.977764 | -0.302365 | 0.191064 | -0.364497 | -0.039505 | 0.01318 | 0.175932 | -0.652951 | 0.067813 | -0.041863 | 0.05159 |
| 29.7222 | -1.838384 | -1.523186 | -2.003473 | -0.646359 | 1.245802 | -0.672757 | -0.254967 | -1.385726 | -0.921004 | -0.335785 | -3.821782 | -0.270063 | -1.037596 | -0.534243 | 0.108077 | -0.375513 | -0.174613 | -0.251336 | -0.063094 | -0.655278 | -0.440417 | -0.10882 | 0.068935 |
| 29.7361 | -1.905336 | -1.799734 | -2.75493 | -0.941999 | 0.831013 | -0.183496 | -0.068221 | -1.789429 | -0.789082 | -0.181953 | -4.52176 | -0.650206 | -1.417158 | -0.656467 | -0.138226 | -0.134978 | -0.131294 | -0.349156 | -0.268659 | -0.490232 | -1.029473 | -0.11652 | -0.045635 |
| 29.75 | -1.705874 | -1.742292 | -2.361016 | -1.066422 | 0.134493 | -0.198199 | 0.027709 | -1.484975 | -1.022616 | -0.08257 | -4.295792 | -1.034025 | -1.887669 | -0.574436 | -0.514026 | -0.019749 | -0.03069 | -0.280381 | -0.494605 | -0.184526 | -1.160923 | -0.068247 | -0.347625 |
| 29.7639 | -1.124796 | -1.446704 | -1.773724 | -1.004245 | -0.693661 | -0.216856 | 0.053583 | -1.117196 | -1.001094 | -0.088436 | -3.009614 | -1.213924 | -2.119348 | -0.428069 | -0.951809 | -0.124497 | -0.016687 | -0.185231 | -0.863018 | 0.078258 | -0.932827 | -0.043 | -0.684986 |
| 29.7778 | -0.351816 | -1.103515 | -1.243344 | -0.874274 | -1.300488 | -0.06638 | -0.183441 | -0.788013 | -0.487113 | -0.032201 | -2.332369 | -1.145052 | -2.226625 | -0.400226 | -1.034107 | -0.328554 | -0.093547 | -0.077335 | -1.245587 | 0.258148 | -0.535621 | -0.051396 | -1.036684 |
| 29.7917 | 0.026275 | -0.905399 | -0.514 | -0.950627 | -1.6675 | 0.157649 | -0.53015 | -0.548813 | 0.044107 | 0.003392 | -2.116019 | -0.967055 | -2.153494 | -0.574216 | -0.874491 | -0.601182 | -0.117485 | -0.049597 | -1.459267 | 0.351042 | -0.304679 | -0.068181 | -1.374541 |
| 29.8056 | 0.202441 | -0.684223 | -0.530154 | -1.080518 | -1.44362 | 0.108335 | -0.793816 | -0.375354 | 0.473378 | 0.03196 | -1.647551 | -0.849318 | -1.91353 | -0.856511 | -0.625948 | -0.946962 | -0.230845 | -0.115917 | -1.442805 | 0.414482 | -0.234894 | -0.063535 | -1.435727 |
| 29.8194 | 0.367035 | -0.391951 | -0.556075 | -1.062446 | -1.854347 | 0.085391 | -0.763024 | 0.003897 | 0.512056 | 0.034768 | -0.906405 | -0.719313 | -1.570459 | -0.886189 | -0.258825 | -1.16256 | -0.173344 | -0.133137 | -1.071694 | 0.38981 | -0.103585 | 0.011042 | -1.127427 |
| 29.8333 | 0.499873 | -0.27301 | 0.029519 | -0.738697 | -2.012034 | -0.034165 | -0.159942 | 0.490319 | 0.625692 | 0.161258 | -0.05371 | -0.526016 | -1.098081 | -0.615345 | 0.084023 | -1.070602 | 0.03011 | -0.004792 | -0.580924 | 0.3896 | 0.087034 | 0.104706 | -0.67704 |
| 29.8472 | 0.529806 | -0.098122 | 0.838149 | -0.171448 | -1.068825 | -0.295108 | 0.362793 | 0.644242 | 0.815734 | 0.188066 | 0.581129 | -0.279214 | -0.690329 | -0.238362 | 0.240781 | -0.715319 | 0.242747 | 0.15403 | -0.284077 | 0.31897 | 0.209583 | 0.11152 | -0.302966 |
| 29.8611 | 0.32313 | -0.030875 | 1.553535 | 0.11199 | -0.414037 | -0.249837 | 0.72623 | 0.559949 | 0.676518 | 0.114906 | 0.710095 | 0.051523 | -0.331619 | 0.08168 | 0.275903 | -0.305828 | 0.33091 | 0.307195 | -0.202999 | 0.161893 | 0.217257 | 0.060171 | -0.141307 |
| 29.875 | 0.041769 | 0.158069 | 1.912976 | 0.306452 | 0.138593 | 0.05315 | 0.915106 | 0.386022 | 0.429989 | 0.105346 | 0.694289 | 0.377735 | -0.072334 | 0.231253 | 0.236368 | 0.028282 | 0.281274 | 0.392204 | -0.163188 | -0.009474 | 0.154945 | 0.018204 | -0.033189 |
| 29.8889 | -0.251936 | 0.019062 | 1.812823 | 0.406671 | 0.460943 | 0.093845 | 1.120559 | 0.007884 | 0.172811 | 0.09214 | -0.699323 | 0.506212 | 0.030226 | 0.127539 | 0.145246 | 0.168899 | 0.163852 | 0.342736 | -0.125734 | -0.073028 | 0.060735 | -0.008405 | 0.04093 |
| 29.9028 | -1.154779 | -1.927274 | 1.478451 | 0.093385 | 0.11667 | -0.251609 | 1.726637 | -1.57898 | -1.270577 | 0.670095 | -9.101848 | 0.402152 | -0.004583 | -0.072917 | 0.050984 | 0.084341 | 0.081632 | 0.19356 | -0.048805 | -0.014681 | -0.016705 | -0.039091 | -0.027601 |
| 29.9167 | -3.514438 | -6.00917 | 0.3017 | -1.655663 | -1.246403 | -1.889324 | 1.702053 | -4.674912 | -3.820057 | 0.830261 | -23.695503 | 0.186052 | -0.149257 | -0.248339 | -0.164521 | -0.049781 | -0.121294 | -0.048636 | 0.236601 | 0.125894 | -0.162169 | -0.288109 | -0.057064 |
| 29.9306 | -5.435169 | -8.637253 | -1.962641 | -3.828813 | -3.026144 | -2.8501 | -0.152734 | -5.768996 | -5.544827 | 0.049646 | -26.43405 | 0.059991 | -0.978559 | -0.588443 | -1.123297 | -0.283446 | -1.315878 | -0.76173 | 0.437909 | 0.094962 | -1.072587 | -1.637573 | -0.270922 |
| 29.9444 | -4.669327 | -6.867415 | -5.021597 | -4.158321 | -3.169691 | -1.8824 | -2.519102 | -3.066653 | -5.31733 | 0.395614 | -15.224804 | -0.337379 | -2.809038 | -1.7707 | -3.117933 | -1.081398 | -3.596459 | -2.445325 | -0.169987 | -1.39754 | -2.605662 | -2.968275 | -1.900025 |
| 29.9583 | -3.578572 | -4.293638 | -5.55584 | -3.172874 | -2.325184 | -1.113749 | -3.665399 | -1.035691 | -4.233065 | -0.225442 | -7.117487 | -0.929911 | -3.135419 | -2.84913 | -3.836279 | -2.195613 | -3.717688 | -3.303258 | -1.499068 | -3.509897 | -2.89726 | -2.029583 | -3.348642 |
| 29.9722 | -2.609056 | -2.726815 | -4.097073 | -1.488501 | -1.523483 | -0.915805 | -2.151427 | -0.316193 | -2.683216 | -0.290751 | -4.402052 | -0.916423 | -2.248648 | -2.936661 | -2.301529 | -2.448254 | -1.697975 | -2.437057 | -2.011001 | -2.007053 | -1.58253 | -0.287739 | -2.687283 |
| 29.9861 | -1.840157 | -2.083823 | -2.458507 | -0.815872 | -0.738246 | -0.485644 | -0.50879 | -0.287261 | -1.988898 | -0.303365 | -2.321934 | 0.067481 | -0.855393 | -1.245264 | -0.689905 | -1.166866 | 0.021562 | -0.754152 | -0.858564 | 0.483912 | 0.204267 | -0.007649 | -0.675096 |
| 30.0069 | -0.718082 | -1.008512 | -0.695832 | -0.244713 | -0.169023 | 0.019598 | 0.889042 | -0.270665 | -1.617536 | -0.190885 | -0.836908 | 0.622136 | -0.131082 | 0.190237 | 0.252942 | 0.045333 | 0.568469 | 0.48973 | 0.116881 | 1.452949 | 0.716183 | -0.020345 | 0.603625 |
| 30.0208 | 0.366339 | 0.126199 | 0.250921 | 0.681266 | 0.395011 | 0.400118 | 1.954136 | -0.081492 | -0.897806 | -0.06243 | 0.113461 | 0.474102 | -0.071381 | 0.725743 | 0.610798 | 0.549899 | 0.667026 | 1.123923 | 0.381083 | 1.131212 | 0.467926 | -0.048444 | 0.778827 |
| 30.0347 | 1.372376 | 1.248961 | 0.596626 | 1.242356 | 0.946258 | 0.692884 | 2.155403 | 0.150125 | 0.150602 | 0.005306 | 1.167848 | 0.168257 | 0.01556 | 1.061102 | 0.874606 | 0.821331 | 0.78451 | 1.394028 | 0.55157 | 0.553596 | 0.204385 | -0.035453 | 0.671312 |
| 30.0486 | 2.15192 | 2.117974 | 0.618396 | 1.507847 | 1.2535 | 0.863617 | 2.140442 | 0.319125 | 1.057983 | -0.008448 | 1.990376 | 0.040711 | 0.315888 | 1.478991 | 1.077008 | 1.047462 | 0.951671 | 1.466359 | 0.836457 | 0.330107 | 0.214759 | 0.014219 | 0.72573 |
| 30.0625 | 2.600825 | 2.46898 | 0.650043 | 1.462466 | 1.262617 | 0.851079 | 2.03797 | 0.211045 | 1.550237 | -0.100867 | 2.275282 | 0.028247 | 0.440442 | 1.810899 | 0.976999 | 1.084224 | 0.933142 | 1.236954 | 0.868399 | 0.301267 | 0.271159 | 0.048181 | 0.757923 |
| 30.0764 | 2.5603 | 2.679828 | 0.919947 | 1.11602 | 1.07637 | 0.873962 | 1.768924 | 0.118171 | 1.494705 | -0.207482 | 2.474898 | 0.07563 | 0.378239 | 2.065251 | 0.824712 | 1.083457 | 0.502385 | 0.707635 | 0.719303 | 0.181009 | 0.21648 | -0.028497 | 0.62602 |
| 30.0903 | 2.425225 | 2.487482 | 1.357801 | 0.670794 | 1.110642 | 1.05945 | 1.324691 | 0.25929 | 1.305621 | -0.192463 | 2.90443 | 0.159883 | 0.45972 | 2.407313 | 0.757964 | 1.170969 | 0.089088 | 0.201239 | 0.591434 | 0.049206 | 0.174224 | -0.078613 | 0.536785 |
| 30.1042 | 2.149928 | 1.112435 | 1.797884 | -0.035694 | 1.181162 | 1.274786 | 0.817464 | 0.465504 | 1.819971 | -0.027395 | 2.513131 | 0.167597 | 0.808293 | 2.502607 | 0.738849 | 1.226964 | -0.166793 | -0.097615 | 0.491797 | 0.020421 | 0.271584 | -0.036297 | 0.521794 |
| 30.1181 | 1.969056 | 0.079125 | 1.878501 | -0.255394 | 1.069159 | 1.364438 | 0.327212 | 0.760427 | 1.768802 | 0.038271 | 2.106381 | 0.012524 | 1.090972 | 2.0741 | 0.567655 | 1.189121 | -0.256573 | -0.105684 | 0.388812 | 0.109762 | 0.424684 | 0.01786 | 0.477053 |
| 30.1319 | 2.350329 | 0.410363 | 2.238655 | 0.579195 | 0.830329 | 1.258006 | 0.295409 | 1.133784 | 1.855746 | -0.193389 | 2.339837 | -0.190837 | 1.032155 | 1.279104 | 0.294573 | 0.91386 | -0.222537 | 0.103025 | 0.202462 | 0.163013 | 0.343686 | -0.031426 | 0.308316 |
| 30.1458 | 2.5921 | 1.072173 | 2.959614 | 1.438345 | 0.546826 | 1.354162 | 0.605942 | 1.532371 | 1.874359 | -0.394705 | 2.661147 | -0.199144 | 0.935832 | 0.491652 | 0.079433 | 0.521157 | -0.118237 | 0.208241 | 0.04805 | 0.183769 | 0.080003 | -0.071635 | 0.05627 |
| 30.1597 | 2.239975 | 1.588365 | 3.345007 | 1.885158 | 0.357518 | 1.27366 | 1.195792 | 1.607782 | 1.688702 | -0.645359 | 2.602834 | 0.23486 | 1.276798 | -0.059195 | 0.056752 | 0.173372 | -0.073037 | 0.145603 | -0.053046 | 0.286836 | -0.138586 | -0.007531 | 0.008004 |
| 30.1736 | 1.740893 | 1.825981 | 2.979079 | 1.443309 | 0.311743 | 1.135374 | 1.397933 | 1.51971 | 1.407658 | -0.455977 | 2.149192 | 0.74772 | 1.79346 | -0.180057 | 0.151397 | -0.014702 | -0.138394 | -0.044532 | -0.131549 | 0.387731 | -0.235054 | 0.081971 | 0.115197 |
| 30.1875 | 1.063716 | 1.813594 | 2.536087 | 0.815787 | 0.297386 | 0.877998 | 1.371448 | 1.290673 | 1.562816 | 0.022501 | 1.219728 | 0.780245 | 1.715287 | -0.02511 | 0.188337 | -0.154356 | -0.199796 | -0.192696 | -0.167754 | 0.335545 | -0.311923 | 0.046586 | 0.185551 |
| 30.2014 | 0.221841 | 1.706708 | 2.254687 | 0.086434 | 0.321765 | 0.698135 | 1.048384 | 1.002675 | 2.135234 | 0.234798 | 0.652977 | 0.50176 | 1.174267 | 0.286292 | 0.237514 | -0.1127 | 0.023674 | -0.019946 | 0.05542 | 0.195957 | -0.317107 | 0.062798 | 0.223157 |
| 30.2153 | 0.123885 | 1.754645 | 2.170887 | 0.023284 | 0.472122 | 0.658226 | 0.845573 | 0.835954 | 2.709502 | 0.297165 | 0.867766 | 0.358605 | 0.719025 | 0.682488 | 0.38962 | 0.146555 | 0.31188 | 0.38282 | 0.391724 | 0.166992 | -0.185444 | 0.137102 | 0.362797 |
| 30.2292 | 0.511305 | 1.407121 | 1.941117 | 0.329735 | 0.722313 | 0.703415 | 1.068869 | 0.970104 | 2.821131 | 0.361966 | 1.128836 | 0.518025 | 0.572255 | 0.74959 | 0.468111 | 0.292989 | 0.331719 | 0.580326 | 0.601748 | 0.160803 | -0.063803 | 0.455056 | 0.472406 |
| 30.2431 | 1.007339 | 1.035212 | 1.813193 | 0.637866 | 1.03575 | 0.723273 | 1.412841 | 1.309859 | 2.871125 | 0.575493 | 1.135159 | 0.991364 | 0.737394 | 0.42868 | 0.379091 | 0.119959 | 0.239591 | 0.53318 | 0.839945 | 0.218537 | 0.00199 | 0.14391 | 0.525874 |
| 30.2569 | 1.347462 | 1.03372 | 1.500318 | 0.887425 | 1.374551 | 0.758284 | 1.395542 | 1.482593 | 2.784931 | 0.573856 | 1.326357 | 1.362222 | 0.789606 | 0.097473 | 0.366633 | 0.32197 | 0.317164 | 0.523203 | 0.885083 | 0.265819 | 0.198282 | -0.052214 | 0.516823 |
| 30.2708 | 1.201793 | 1.058208 | 0.502512 | 0.827795 | 1.479712 | 0.487514 | 0.789534 | 0.97884 | 1.11642 | 0.374431 | 1.058515 | 1.280458 | 0.45413 | 0.167491 | 0.530509 | 0.67088 | 0.455751 | 0.562933 | 0.647731 | 0.284915 | 0.480913 | 0.309454 | 0.442129 |
| 30.2847 | 0.724245 | 0.826769 | -0.357928 | 0.573557 | 1.087733 | 0.069508 | 0.221668 | 0.152872 | -0.509557 | 0.434209 | 0.356837 | 0.772012 | -0.161983 | 0.585574 | 0.752408 | 0.767472 | 0.35285 | 0.449647 | 0.49378 | 0.320942 | 0.577085 | 0.20214 | 0.330464 |
| 30.2986 | 0.576838 | 0.466344 | -0.583537 | 0.459652 | 0.50141 | -0.139168 | -0.002802 | -0.392084 | -0.862197 | 0.523362 | 0.057022 | 0.303134 | -0.706671 | 0.955146 | 0.8136 | 0.577737 | 0.031135 | 0.135696 | 0.387545 | 0.310934 | 0.414747 | 0.249289 | 0.180349 |
| 30.3125 | 0.996013 | 0.218663 | -0.35802 | 0.682979 | 0.105899 | 0.032573 | 0.225324 | -0.325724 | -0.128099 | 0.385864 | 0.23765 | 0.275016 | -0.599011 | 1.201776 | 0.622149 | 0.440759 | -0.086826 | -0.061474 | 0.314115 | 0.217112 | 0.230271 | 0.277197 | 0.191807 |
| 30.3264 | 1.683376 | 0.190681 | -0.04136 | 1.009898 | -0.062858 | 0.300242 | 0.58726 | -0.089408 | 0.809467 | 0.230198 | 0.782457 | 0.628109 | 0.041799 | 0.871726 | 0.31735 | 0.427452 | 0.060968 | -0.047679 | 0.266708 | 0.084042 | 0.08952 | 0.222849 | 0.342746 |
| 30.3403 | 2.169153 | 0.332833 | 0.140839 | 1.344001 | -0.176036 | 0.448997 | 0.747607 | 0.053065 | 1.484683 | -0.023419 | 1.212041 | 0.938999 | 0.85797 | 0.34873 | 0.065876 | 0.406995 | 0.272579 | 0.094204 | 0.243311 | 0.018737 | 0.018452 | 0.0995 | 0.424883 |
| 30.3542 | 2.027153 | 0.405682 | 0.219468 | 1.582298 | -0.184941 | 0.39218 | 0.698565 | 0.038853 | 1.518942 | -0.102668 | 1.133346 | 0.825746 | 1.411657 | -0.071925 | 0.006511 | 0.359587 | 0.411361 | 0.303948 | 0.281193 | 0.039597 | 0.252668 | 0.054292 | 0.398137 |
| 30.3681 | 1.625005 | 0.492829 | 0.172569 | 1.455515 | -0.107682 | 0.230675 | 0.517121 | 0.002704 | 0.713295 | -0.152 | 0.721067 | 0.421257 | 1.405443 | -0.23139 | 0.148723 | 0.33485 | 0.321677 | 0.355189 | 0.353872 | 0.084172 | 0.734885 | 0.068998 | 0.312059 |
| 30.3819 | 1.260127 | 0.765719 | 0.104544 | 1.244937 | -0.207365 | 0.164248 | 0.32091 | -0.0728 | -0.386561 | -0.338485 | 0.249902 | 0.221085 | 1.128672 | 0.019385 | 0.433962 | 0.386498 | 0.069273 | 0.213753 | 0.356877 | 0.117322 | 1.165531 | 0.117669 | 0.274222 |
| 30.3958 | 1.194592 | 1.287754 | 0.054305 | 1.133365 | -0.388139 | 0.23055 | 0.248834 | -0.089459 | -1.479624 | -0.211959 | 0.500715 | 0.210653 | 0.845467 | 0.525652 | 0.678299 | 0.446864 | -0.052387 | 0.092679 | 0.348212 | 0.121653 | 1.169563 | 0.163433 | 0.336271 |
| 30.4097 | 2.182831 | 3.006637 | 0.113597 | 1.611521 | -0.271734 | 1.16249 | 0.483019 | 0.954196 | -1.454216 | -0.025198 | 5.515168 | 0.428107 | 0.602058 | 1.05071 | 0.657951 | 0.480269 | 0.05266 | 0.078857 | 0.236935 | 0.059466 | 0.76784 | 0.120718 | 0.380159 |
| 30.4236 | 4.433841 | 5.927634 | 0.50004 | 3.081187 | 0.427307 | 2.737585 | 1.187207 | 2.870052 | -0.681561 | 2.402339 | 14.572382 | 0.538684 | 0.482173 | 1.12018 | 0.546028 | 0.497362 | 0.335669 | 0.183899 | 0.16134 | 0.043076 | 0.276997 | 0.057035 | 0.502199 |
| 30.4375 | 5.131249 | 6.770707 | 1.246316 | 3.367593 | 2.125537 | 2.303783 | 1.531839 | 3.192843 | 0.020704 | 0.894214 | 15.656138 | 0.129323 | 0.686851 | 0.646409 | 1.207592 | 0.42847 | 1.342237 | 0.512508 | 0.13525 | 0.43037 | 0.884783 | 0.223343 | 1.812336 |
| 30.4514 | 3.81431 | 3.901362 | 2.471562 | 3.013695 | 3.259879 | 1.180636 | 2.137529 | 2.169057 | 1.4575 | NaN | 7.848529 | -0.48745 | 1.487041 | 0.143319 | 2.65322 | 0.363772 | 2.979677 | 0.773962 | -0.0216 | 0.791925 | 2.486861 | 1.453377 | 3.712123 |
| 30.4653 | 3.334053 | 1.832714 | 3.91717 | 3.115426 | 2.849891 | 0.600565 | 2.570978 | 1.78425 | 2.065554 | NaN | 2.487136 | -0.546237 | 2.335809 | -0.238129 | 2.6936 | 0.030896 | 2.945925 | 0.395839 | -0.065818 | 0.30487 | 2.419614 | 2.821479 | 5.063236 |
| 30.4792 | 2.893307 | 2.021494 | 4.301991 | 2.67587 | 2.20816 | 0.295726 | 2.354091 | 2.333832 | 2.275819 | NaN | 1.605812 | 0.524892 | 2.445081 | 0.229165 | 1.379639 | 0.380762 | 1.183769 | 0.398551 | 0.634503 | -0.438791 | 1.765421 | 1.688746 | 2.860019 |
| 30.4931 | 2.445659 | 1.838943 | 3.212917 | 1.673411 | 2.005351 | 0.594871 | 1.605754 | 1.842627 | 2.017377 | NaN | 0.998268 | 1.111755 | 1.712375 | 1.230647 | 0.900707 | 0.884667 | 0.636207 | 0.809097 | 1.260049 | -0.920068 | 1.766787 | 0.139548 | 1.759623 |
| 30.5 | 1.030323 | 1.397214 | 1.918875 | 0.995985 | 1.756557 | 0.653319 | 0.83351 | 1.47532 | 1.616886 | 0.071819 | 0.781523 | 1.372556 | 1.191121 | 1.68636 | 1.102015 | 1.146104 | 0.656188 | 1.095959 | 1.277842 | -0.698959 | 1.841748 | 0.03781 | 1.830771 |
| 30.5139 | -0.383901 | 1.088952 | 0.750314 | 0.382603 | 1.860512 | 0.645132 | 0.058995 | 1.311309 | 1.632887 | 0.128451 | 0.981829 | 1.28453 | 0.550251 | 1.382365 | 1.016907 | 1.010961 | 0.527961 | 0.898313 | 0.990655 | -0.341443 | 1.25724 | 0.145577 | 1.64594 |
| 30.5278 | -1.143796 | 0.885343 | -0.082312 | 0.045814 | 1.868294 | 0.628043 | -0.486178 | 0.890712 | 0.785192 | 0.109578 | 1.258323 | 0.841306 | -0.114634 | 0.624544 | 0.619132 | 0.59486 | 0.058713 | 0.187029 | 0.533891 | -0.126239 | 0.193318 | 0.134194 | 1.267043 |
| 30.5417 | -0.993983 | 0.771558 | -0.79672 | -0.112431 | 1.627098 | 0.549123 | -0.516178 | 0.362948 | -0.496262 | NaN | 1.585401 | 0.662297 | -0.740805 | -0.26212 | 0.256984 | 0.112278 | -0.428036 | -0.595235 | 0.081938 | -0.105911 | -0.8437 | 0.032504 | 0.935123 |
| 30.5556 | -1.220727 | 0.505127 | -1.524013 | -0.06081 | 1.185898 | 0.30724 | -0.187337 | -0.181147 | -1.063763 | NaN | 1.083849 | 0.394502 | -1.335379 | -0.985802 | 0.06196 | -0.178326 | -0.554868 | -0.971165 | -0.297666 | -0.146934 | -1.362026 | -0.011962 | 0.41117 |
| 30.5694 | -1.282349 | -0.11186 | -1.993062 | -0.185201 | 0.520261 | 0.014561 | -0.020545 | -0.849341 | -1.177266 | NaN | -0.93042 | -0.01162 | -1.95451 | -1.469884 | -0.017761 | -0.310247 | -0.501279 | -0.950985 | -0.384814 | -0.044455 | -1.684215 | -0.016974 | -0.018755 |
| 30.5833 | -0.966271 | -0.843539 | -1.76406 | -0.372418 | -0.192653 | -0.404826 | 0.064214 | -1.155807 | -1.160122 | NaN | -2.013025 | 0.081459 | -2.055844 | -1.111788 | -0.085073 | -0.261313 | -0.225663 | -0.604533 | -0.265481 | 0.061546 | -1.667668 | -0.034512 | -0.236287 |
| 30.5972 | 0.673893 | -1.11258 | -2.32458 | -0.388573 | -0.862433 | -0.611703 | -0.036296 | -1.181752 | -1.453365 | NaN | -1.347135 | 0.577564 | -1.689083 | -0.280345 | -0.186011 | -0.375714 | 0.045515 | -0.146936 | -0.139214 | 0.23044 | -0.826069 | -0.005448 | -0.180425 |
| 30.6111 | 1.359589 | -1.0503 | -2.047877 | -0.313423 | -1.398572 | -0.568153 | -0.173976 | -1.136532 | -0.976656 | NaN | -0.992642 | 0.808312 | -1.144407 | 0.442799 | -0.294142 | -0.507229 | 0.075979 | -0.004018 | -0.107414 | 0.4283 | 0.106231 | 0.00892 | -0.091527 |
| 30.625 | 1.081754 | -0.693451 | -0.102407 | -0.012961 | -1.390943 | -0.542205 | -0.215845 | -1.029083 | 0.198437 | NaN | -0.467919 | 0.521127 | -0.447465 | 0.958553 | -0.298147 | -0.486844 | -0.052361 | -0.18822 | -0.101726 | 0.557604 | 0.612054 | -0.027943 | -0.104069 |
| 30.6389 | 1.025975 | -0.242228 | 1.116792 | 0.402948 | -1.501361 | -0.741907 | -0.16163 | -1.228351 | 1.082715 | NaN | -0.35894 | 0.001322 | 0.1396 | 1.026179 | -0.301775 | -0.310066 | -0.183831 | -0.403747 | -0.081246 | 0.50588 | 0.496189 | -0.062626 | -0.246051 |
| 30.6528 | 0.867142 | -0.137672 | 1.643589 | 0.766201 | -1.862491 | -0.476097 | 0.010342 | -1.550126 | 1.572241 | NaN | -1.205049 | -0.300545 | 0.297784 | 0.790526 | -0.228448 | -0.071341 | -0.222566 | -0.367191 | -0.149533 | 0.395973 | 0.027925 | -0.08118 | -0.346417 |
| 30.6667 | 0.409464 | -0.404671 | 1.242355 | 1.075551 | -1.881129 | -0.03015 | 0.178982 | -1.571249 | 2.115943 | NaN | -2.074724 | -0.336798 | 0.0958 | 0.521936 | -0.057923 | 0.18173 | -0.152695 | -0.21101 | -0.234618 | 0.256848 | -0.343827 | -0.068555 | -0.31036 |
| 30.6806 | -0.154365 | -0.70418 | 0.20293 | 1.382212 | -1.313184 | -0.053784 | 0.306772 | -1.014481 | 1.19082 | NaN | -2.215155 | -0.078638 | 0.140996 | 0.166648 | 0.13228 | 0.335995 | -0.112247 | -0.123956 | -0.265098 | 0.149855 | -0.407861 | -0.019012 | -0.109957 |
| 30.6944 | -0.341664 | -0.825825 | -0.143772 | 1.209171 | -0.688384 | 0.041269 | 0.286942 | -0.744348 | 0.446189 | NaN | -1.561717 | 0.417524 | 0.222873 | -0.329029 | 0.239886 | 0.313716 | -0.112001 | -0.097398 | -0.219243 | 0.090906 | -0.304701 | 0.063697 | 0.100576 |
| 30.7083 | -0.223336 | -0.806782 | -0.539653 | 0.771005 | -0.111465 | 0.069513 | 0.147881 | -0.597986 | -0.091702 | NaN | -0.567869 | 0.69439 | 0.068595 | -0.816342 | 0.194898 | 0.182397 | -0.100202 | -0.034617 | -0.103571 | 0.09405 | -0.178009 | 0.146012 | 0.220571 |
| 30.7222 | 0.188454 | -0.688535 | -0.598089 | 0.105116 | 0.407455 | 0.066123 | -0.186837 | -0.440074 | -0.618775 | NaN | 0.251398 | 0.712399 | -0.174559 | -0.922939 | 0.094255 | 0.01791 | -0.067628 | 0.048785 | 0.018314 | 0.142768 | -0.102139 | 0.137726 | 0.273807 |
| 30.7361 | 0.201074 | -0.494158 | -0.642174 | -0.592208 | 0.726612 | 0.229129 | -0.524451 | -0.252637 | -1.356489 | NaN | 0.902502 | 0.241533 | -0.516763 | -0.916704 | -0.073857 | -0.193163 | -0.062127 | 0.01029 | 0.139124 | 0.167328 | -0.233714 | 0.038519 | 0.285835 |
| 30.75 | -0.464491 | -0.429282 | -0.707328 | -0.938359 | 0.71488 | 0.662273 | -0.845351 | 0.135464 | -1.679817 | NaN | 1.38843 | -0.284654 | -1.140358 | -0.969914 | -0.348793 | -0.435626 | -0.086501 | -0.161145 | 0.056393 | 0.172464 | -0.440323 | -0.067128 | 0.316475 |
| 30.7639 | -0.795331 | -0.768803 | -1.813194 | -1.314115 | 0.355645 | 0.756309 | -0.769443 | 0.166484 | -1.423036 | NaN | 1.604467 | -0.659485 | -1.724488 | -1.01638 | -0.65377 | -0.714296 | -0.155429 | -0.382988 | -0.10875 | 0.173284 | -0.391318 | -0.123615 | 0.190261 |
| 30.7778 | -0.827792 | -1.355603 | -3.387755 | -1.588904 | -0.253523 | 0.28156 | -1.054718 | 0.060363 | -1.091741 | NaN | -0.865609 | -0.589128 | -1.972724 | -0.854081 | -0.848217 | -0.753426 | -0.134864 | -0.430189 | -0.312033 | 0.225438 | -0.069595 | -0.106003 | -0.05128 |
| 30.7917 | -0.912161 | -1.84699 | -4.047065 | -1.61835 | -0.968169 | 0.008663 | -1.295748 | -0.112933 | -0.900152 | NaN | -2.330573 | -0.420397 | -1.900283 | -0.756618 | -0.841443 | -0.60655 | -0.181993 | -0.409267 | -0.428058 | 0.275061 | 0.206219 | -0.008003 | -0.427674 |
| 30.8056 | -0.727102 | -1.655534 | -3.792266 | -1.52778 | -1.651224 | -0.02678 | -1.359089 | -0.195289 | -0.976974 | -1.099725 | -1.933311 | -0.512046 | -1.933621 | -0.892486 | -0.72043 | -0.445171 | -0.207784 | -0.360012 | -0.579225 | 0.363931 | 0.244068 | 0.090917 | -0.808081 |
| 30.8194 | -0.348796 | -1.209402 | -3.272382 | -1.364303 | -1.904049 | -0.206694 | -1.275746 | -0.458549 | -1.055954 | -0.579912 | -1.97327 | -0.728805 | -1.808477 | -0.853734 | -0.475533 | -0.323042 | -0.164044 | -0.210479 | -0.77522 | 0.443471 | 0.206806 | 0.083778 | -1.015278 |
| 30.8333 | 0.286297 | -0.661695 | -2.54589 | -1.076786 | -1.848236 | -0.453871 | -0.787948 | -0.821103 | -0.768715 | -0.261752 | -2.35617 | -1.117134 | -1.363868 | -0.571979 | -0.226508 | -0.242542 | 0.061349 | 0.091715 | -0.801747 | 0.439818 | 0.209405 | -0.031618 | -0.981001 |
| 30.8472 | 1.038441 | 0.168626 | -1.558515 | -0.246425 | -1.427258 | -0.454118 | 0.003793 | -0.890964 | -0.405111 | 0.016017 | -1.727736 | -1.352569 | -0.966836 | -0.288451 | -0.070023 | -0.15552 | 0.290544 | 0.409965 | -0.79255 | 0.328967 | 0.283989 | -0.122588 | -0.787515 |
| 30.8611 | 1.994213 | 0.264544 | -0.315518 | 0.451095 | -0.833163 | -0.353409 | 0.980509 | -0.637749 | 0.201499 | 0.143091 | -1.068412 | -1.174821 | -0.709671 | -0.165929 | 0.001348 | -0.023342 | 0.448888 | 0.508732 | -0.727054 | 0.242379 | 0.350863 | -0.103964 | -0.487253 |
| 30.875 | 2.706017 | 0.195878 | 0.971204 | 0.899709 | -0.232531 | -0.278921 | 1.825737 | -0.388872 | 0.790779 | 0.56459 | -0.645935 | -0.691483 | -0.42877 | -0.037682 | 0.034269 | 0.146741 | 0.432274 | 0.434076 | -0.563561 | 0.218882 | 0.299924 | -0.071335 | -0.209296 |
| 30.8889 | 2.775473 | 0.154285 | 1.970867 | 0.873661 | 0.268581 | -0.334446 | 2.197972 | -0.589462 | 0.981421 | 0.804651 | -1.369465 | -0.22186 | -0.257057 | 0.016618 | 0.024648 | 0.269212 | 0.268886 | 0.139361 | -0.358996 | 0.211471 | 0.210402 | -0.096567 | -0.017283 |
| 30.9028 | 1.727106 | -1.017224 | 2.300435 | 0.59691 | 0.218181 | -0.990646 | 1.636027 | -1.594569 | 0.517291 | -0.855043 | -8.641385 | 0.018636 | -0.112342 | -0.067139 | 0.000427 | 0.273605 | 0.047906 | -0.140297 | -0.312581 | 0.184294 | 0.127542 | -0.085954 | 0.057879 |
| 30.9167 | -1.739915 | -3.175783 | 0.558697 | -0.618764 | -0.869352 | -2.296585 | -0.296344 | -2.778035 | -1.147828 | -4.227585 | -19.057704 | 0.007646 | -0.124433 | -0.167095 | -0.189689 | 0.128012 | -0.23762 | -0.285236 | -0.26981 | 0.115273 | -0.060072 | -0.288895 | -0.276194 |
| 30.9306 | -5.329712 | -3.870959 | -2.019983 | -2.988903 | -2.640092 | -2.264118 | -2.174085 | -2.95 | -2.274621 | -3.875751 | -18.287773 | -0.11802 | -1.079063 | -0.032777 | -1.348378 | -0.182028 | -1.457169 | -0.941865 | 0.272481 | -0.299156 | -1.04088 | -1.76931 | -2.724853 |
| 30.9444 | -4.78874 | -2.833093 | -3.22128 | -3.661938 | -2.825735 | -1.295469 | -2.846046 | -1.374044 | -2.410281 | -0.313591 | -9.102933 | -0.487823 | -3.060829 | 0.014525 | -3.444193 | -1.175844 | -4.020794 | -2.885473 | 0.107182 | -1.695678 | -3.111201 | -3.842315 | -7.795961 |
| 30.9583 | -2.859032 | -1.98118 | -3.375798 | -2.895747 | -2.16039 | -0.516899 | -3.654796 | -0.427638 | -2.062604 | 0.551902 | -3.47783 | -0.77336 | -3.402409 | -1.129482 | -4.102926 | -2.378697 | -4.672905 | -4.465828 | -1.238019 | -3.199698 | -4.000361 | -2.924024 | -8.803018 |
| 30.9722 | -2.078979 | -1.687274 | -2.76327 | -1.838063 | -1.192022 | 0.080355 | -3.139248 | -0.03413 | -1.947002 | 0.442043 | -2.005733 | -0.532606 | -2.459029 | -1.345596 | -2.600343 | -2.176905 | -2.654 | -3.309898 | -1.585616 | -1.751207 | -1.989859 | -0.53056 | -3.866396 |
| 30.9861 | -1.766602 | -1.006054 | -1.938809 | -1.17605 | -0.399618 | 0.148711 | -2.028332 | 0.228352 | -1.583601 | 0.386361 | -1.305463 | -0.130279 | -0.906615 | -0.633908 | -0.885057 | -0.902628 | -0.753879 | -1.214861 | -1.095548 | -0.069544 | -0.139413 | -0.130203 | -0.668364 |
| 31.0069 | -1.655726 | -1.332134 | -1.302395 | -0.715485 | -0.002926 | 0.359411 | -0.471871 | 0.592278 | -0.7772 | 0.142641 | -0.114057 | 0.436831 | -0.211839 | 0.041655 | 0.10572 | 0.180357 | 0.002893 | 0.060411 | -0.400579 | 0.719284 | 0.629028 | -0.007438 | 0.807325 |
| 31.0208 | -1.265314 | -1.119322 | -0.716127 | -0.438798 | 0.144527 | 0.502888 | 1.040515 | 0.689802 | -0.166952 | 0.239986 | 0.852779 | 0.9512 | -0.150614 | 0.395723 | 0.567827 | 0.497277 | 0.263736 | 0.430538 | 0.020187 | 0.742386 | 0.586478 | 0.412366 | 1.191014 |
| 31.0347 | -0.20685 | -0.145218 | 0.437053 | 0.326637 | 0.272133 | 0.836174 | 2.72827 | 0.857855 | 0.365158 | 0.227396 | 1.98428 | 1.236536 | -0.047568 | 0.742897 | 0.59256 | 0.47044 | 0.384494 | 0.544303 | 0.338194 | 0.599878 | 0.369095 | -1.127124 | 1.140025 |
| 31.0486 | 1.026963 | 0.948363 | 1.430075 | 1.141499 | 0.415493 | 1.201381 | 4.055885 | 0.910071 | 1.136427 | 0.197529 | 3.128179 | 1.057933 | 0.418485 | 1.272064 | 0.491992 | 0.459599 | 0.684792 | 0.92327 | 0.7562 | 0.731731 | 0.407481 | -0.243125 | 1.229742 |
| 31.0625 | 1.522124 | 1.266343 | 1.338007 | 1.255012 | 0.480335 | 1.077858 | 3.75128 | 0.584819 | 1.509269 | 0.221721 | 2.571239 | 0.859191 | 0.922721 | 1.537598 | 0.611201 | 0.688973 | 0.939203 | 1.437856 | 1.189129 | 0.923402 | 0.542523 | 0.269419 | 1.439057 |
| 31.0764 | 1.232376 | 0.968267 | 1.075407 | 0.908468 | 0.454576 | 0.512282 | 3.208574 | 0.161165 | 0.884027 | 0.212579 | 0.44424 | 0.649011 | 0.914334 | 1.358242 | 0.853201 | 0.890358 | 1.163414 | 1.88081 | 1.25969 | 0.759336 | 0.489122 | 1.811677 | 1.421576 |
| 31.0903 | 1.129729 | 1.140854 | 0.439146 | 0.605235 | 0.336008 | 0.20359 | 1.987085 | 0.06015 | 0.335898 | 0.21428 | 0.18737 | 0.394103 | 0.506094 | 0.874383 | 1.143844 | 1.022693 | 1.325091 | 1.942016 | 1.06232 | 0.437508 | 0.155896 | -0.169289 | 1.026557 |
| 31.1042 | 1.186572 | 1.612173 | 0.07316 | 0.462535 | 0.152802 | 0.095348 | 1.037833 | 0.193394 | -0.038475 | -0.09417 | 1.170767 | -0.069681 | 0.097846 | 0.389798 | 1.133145 | 1.046651 | 0.896692 | 1.146903 | 0.528888 | 0.211077 | -0.206646 | 1.615049 | 0.325751 |
| 31.1181 | 0.825056 | 1.35829 | 0.156 | 0.366974 | -0.04486 | -0.162501 | 0.432029 | 0.095832 | -0.503516 | 0.064562 | -0.308383 | -0.443173 | -0.174589 | -0.026015 | 0.731415 | 0.639036 | 0.065335 | -0.062495 | 0.141539 | 0.080753 | -0.320019 | -0.186842 | -0.403829 |
| 31.1319 | 0.692992 | 1.092802 | 0.463844 | 0.367981 | -0.09597 | -0.188688 | 0.26803 | -0.046736 | -0.556232 | 0.193814 | -1.913059 | -0.310945 | -0.206722 | 0.083025 | 0.326987 | 0.216305 | -0.624793 | -1.01008 | 0.016791 | -0.011683 | -0.21862 | -2.006949 | -0.709131 |
| 31.1458 | 1.153094 | 1.51145 | 1.066366 | 0.513009 | 0.057868 | 0.09992 | 0.486858 | 0.011536 | -0.012382 | 0.008835 | -0.828823 | 0.041766 | -0.123243 | 0.48477 | 0.146277 | -0.008488 | -0.937863 | -1.401874 | 0.008733 | -0.05919 | -0.048651 | -0.08353 | -0.290454 |
| 31.1597 | 1.807633 | 1.81827 | 1.495952 | 0.670152 | 0.261861 | 0.348203 | 0.849288 | 0.047274 | 0.549232 | -0.086845 | 0.59294 | 0.18221 | -0.063252 | 0.775583 | 0.102869 | -0.18092 | -0.845863 | -1.136738 | -0.020449 | -0.093698 | 0.043219 | -0.09542 | 0.156906 |
| 31.1736 | 2.230635 | 1.583776 | 1.500966 | 0.627679 | 0.346553 | 0.409263 | 1.061443 | -0.057573 | 0.841063 | -0.046806 | 1.046464 | 0.034003 | -0.0684 | 0.729594 | 0.0336 | -0.187352 | -0.163987 | -0.399141 | -0.076756 | -0.140025 | 0.029152 | -0.140163 | 0.251994 |
| 31.1875 | 2.334774 | 1.197553 | 1.163283 | 0.46898 | 0.308032 | 0.388678 | 0.921924 | -0.132671 | 0.715174 | -0.211603 | 1.019743 | -0.153686 | -0.114642 | 0.449313 | -0.112507 | 0.020875 | 0.560436 | 0.361766 | -0.05892 | -0.174261 | -0.000506 | -0.176653 | 0.166479 |
| 31.2014 | 2.351401 | 0.888306 | 0.711108 | 0.299058 | 0.269131 | 0.443468 | 0.656813 | -0.056201 | 0.481837 | -0.367236 | 1.306652 | -0.189975 | -0.167406 | 0.049411 | -0.260281 | 0.115925 | 0.529053 | 0.254067 | -0.074788 | -0.18629 | -0.006863 | -0.180568 | -0.089886 |
| 31.2153 | 2.193407 | 0.892471 | 0.511971 | 0.221381 | 0.284915 | 0.670779 | 0.594166 | 0.138259 | 0.420723 | -0.68838 | 2.183354 | -0.259684 | -0.292928 | -0.338616 | -0.347484 | -0.132501 | -0.156882 | -0.540647 | -0.206725 | -0.177085 | -0.016433 | -0.168744 | -0.416085 |
| 31.2292 | 1.747566 | 1.039364 | 0.552988 | 0.159828 | 0.274231 | 0.849056 | 0.623067 | 0.373253 | 0.440314 | -0.814689 | 3.171806 | -0.180117 | -0.237693 | -0.384576 | -0.280783 | -0.447783 | -0.752884 | -1.172031 | -0.285769 | -0.131276 | -0.003467 | -0.100753 | -0.564322 |
| 31.2431 | 0.929606 | 0.811012 | 0.635842 | -0.003061 | 0.159791 | 0.797943 | 0.492191 | 0.462239 | 0.291551 | -0.93086 | 2.770137 | 0.10174 | 0.002247 | -0.177217 | -0.109386 | -0.486456 | -0.89051 | -1.255024 | -0.147755 | -0.056554 | 0.07601 | 0.551953 | -0.335644 |
| 31.2569 | 0.022089 | 0.200616 | 0.709869 | -0.143451 | 0.052135 | 0.530765 | 0.236527 | 0.323788 | 0.009895 | -0.514882 | 1.01918 | 0.22293 | 0.274994 | 0.123858 | 0.083904 | -0.238804 | -0.887086 | -1.000569 | 0.100325 | 0.011061 | 0.155305 | 0.516867 | 0.075914 |
| 31.2708 | -0.639126 | -0.422295 | 0.747979 | -0.251156 | -0.047075 | 0.24162 | 0.011526 | 0.142695 | -0.17037 | -0.063892 | -0.701063 | 0.219109 | 0.450477 | 0.359222 | 0.129523 | 0.064361 | -0.39376 | -0.410286 | 0.304783 | -0.004919 | 0.093994 | -0.471333 | 0.261102 |
| 31.2847 | -0.917805 | -0.791372 | 0.770835 | -0.255722 | -0.136258 | -0.003785 | -0.126735 | 0.039714 | -0.158943 | 0.504754 | -1.712862 | 0.15994 | 0.48998 | 0.37278 | 0.134068 | 0.215028 | 0.070878 | 0.217837 | 0.310483 | -0.042771 | -0.048699 | -0.55444 | 0.241584 |
| 31.2986 | -0.872405 | -0.770943 | 0.826753 | -0.232035 | -0.181682 | -0.043292 | -0.249827 | 0.116379 | -0.024041 | 0.588436 | -1.473526 | 0.126579 | 0.475598 | 0.243757 | 0.146986 | 0.298022 | 0.621038 | 0.718655 | 0.187546 | 0.027937 | -0.072775 | 0.094892 | 0.258532 |
| 31.3125 | -0.703405 | -0.509137 | 0.841973 | -0.198148 | -0.207469 | 0.08732 | -0.32991 | 0.330955 | 0.159458 | 0.38919 | -0.356621 | 0.226642 | 0.436562 | 0.136195 | 0.147331 | 0.297765 | 0.844414 | 0.971957 | 0.13089 | 0.132391 | -0.003924 | 0.740425 | 0.271559 |
| 31.3264 | -0.601644 | -0.159721 | 0.874037 | -0.182072 | -0.239688 | 0.227079 | -0.245873 | 0.527913 | 0.340845 | -0.072556 | 0.817556 | 0.337094 | 0.452364 | 0.137255 | 0.102813 | 0.276594 | 0.562092 | 0.60868 | 0.222165 | 0.155947 | 0.057941 | 0.851572 | 0.178661 |
| 31.3403 | -0.571146 | 0.080713 | 0.772219 | -0.184627 | -0.250221 | 0.360176 | -0.075983 | 0.614683 | 0.349052 | -0.554066 | 1.667453 | 0.382773 | 0.620435 | 0.293074 | 0.090861 | 0.332726 | 0.016327 | 0.085836 | 0.380688 | 0.123671 | 0.114273 | 0.285294 | 0.175913 |
| 31.3542 | -0.611108 | -0.152995 | 0.477357 | -0.148906 | -0.276065 | 0.264156 | -0.047607 | 0.433907 | 0.060288 | -0.128663 | 0.604393 | 0.353579 | 0.799527 | 0.49101 | 0.125887 | 0.438429 | -0.400811 | -0.324587 | 0.47046 | 0.087633 | 0.139952 | -0.735966 | 0.335208 |
| 31.3681 | -0.448864 | -0.351386 | 0.446119 | 0.064564 | -0.29835 | 0.141466 | 0.099857 | 0.261427 | -0.147923 | 0.363703 | -0.347739 | 0.26225 | 0.8029 | 0.609487 | 0.134401 | 0.51952 | -0.618675 | -0.536366 | 0.465817 | 0.037375 | 0.08307 | -1.861612 | 0.442455 |
| 31.3819 | -0.002777 | 0.049595 | 0.638784 | 0.368848 | -0.36378 | 0.293867 | 0.403363 | 0.34564 | 0.039044 | -0.189128 | 1.045292 | 0.313661 | 0.61646 | 0.571993 | 0.089651 | 0.599734 | -0.374315 | -0.287739 | 0.4952 | -0.036427 | 0.005161 | -0.090119 | 0.410469 |
| 31.3958 | 0.552843 | 0.561371 | 0.705654 | 0.53559 | -0.547226 | 0.502741 | 0.629825 | 0.507879 | 0.316717 | -0.852553 | 3.049228 | 0.407175 | 0.485438 | 0.454829 | 0.129822 | 0.778021 | 0.107381 | 0.230198 | 0.615023 | -0.022185 | 0.036371 | 0.553506 | 0.433746 |
| 31.4097 | 2.183226 | 1.830883 | 0.706078 | 0.952985 | -0.774363 | 1.243714 | 1.730832 | 0.897212 | 0.632875 | -1.092379 | 8.211416 | 0.412226 | 0.548381 | 0.29256 | 0.260536 | 0.918698 | 0.666393 | 0.804504 | 0.64994 | 0.090473 | 0.149207 | 1.53998 | 0.47965 |
| 31.4236 | 4.854266 | 4.055964 | 1.039222 | 1.845996 | -0.824145 | 2.434924 | 4.234347 | 1.36013 | 1.133294 | -1.863863 | 16.712343 | 0.178598 | 0.648429 | 0.153359 | 0.408718 | 1.052906 | 0.863979 | 0.970446 | 0.508576 | 0.165526 | 0.237583 | 3.919611 | 0.597731 |
| 31.4375 | 5.565405 | 4.515049 | 2.365956 | 1.965105 | -0.559609 | 2.708613 | 5.704477 | 0.90742 | 1.463582 | NaN | 17.580144 | -0.480133 | 0.963701 | 0.265521 | 1.287942 | 1.105026 | 1.425134 | 0.882517 | 0.193215 | 0.599413 | 0.80546 | 3.044354 | 2.843373 |
| 31.4514 | 4.4528 | 2.601277 | 3.913118 | 1.973475 | 0.405684 | 1.424979 | 3.789742 | 0.423949 | 1.659464 | NaN | 7.008536 | -1.157191 | 1.95859 | 0.650675 | 2.984442 | 1.238739 | 2.614256 | 0.977485 | -0.25802 | 1.687108 | 1.844644 | 2.077437 | 7.430816 |
| 31.4653 | 3.559538 | 0.988371 | 3.3797 | 2.67017 | 1.12904 | 0.827358 | 3.534352 | -0.01121 | 1.381428 | NaN | 2.162687 | -0.407706 | 3.411709 | 1.026772 | 3.47874 | 1.923104 | 2.557428 | 1.047395 | -0.333157 | 1.697761 | 1.460329 | 3.082128 | 8.04982 |
| 31.4792 | 3.662005 | 0.961074 | 2.78643 | 2.819789 | 1.425544 | 0.393447 | 3.209204 | -0.192265 | 1.480611 | NaN | 1.898928 | 0.630351 | 3.410395 | 1.214329 | 2.220299 | 2.224034 | 1.219022 | 1.100886 | 0.591239 | 0.823469 | 0.840314 | 3.955198 | 4.020705 |
| 31.4931 | 3.411817 | 0.598663 | 2.043255 | 2.373578 | 1.176499 | 0.057788 | 1.803038 | -0.076666 | 0.981085 | NaN | 0.869776 | 0.973978 | 1.906119 | 2.183026 | 1.058731 | 2.037059 | 0.500726 | 1.172077 | 1.111556 | 0.194508 | 0.850298 | 1.564408 | 0.929729 |
| 31.5 | 2.454556 | 0.134493 | 1.510446 | 1.379017 | 0.961834 | -0.078983 | 0.570653 | 0.120785 | 0.716253 | NaN | 0.077889 | 1.003731 | 0.887972 | 2.727781 | 0.776565 | 1.691755 | 0.192777 | 1.098888 | 1.092541 | -0.19627 | 1.174374 | -0.124394 | 0.475126 |
| 31.5139 | 1.515367 | 0.201343 | 1.036792 | 0.594438 | 1.360408 | -0.282372 | -0.232434 | 0.381083 | 1.199345 | NaN | 0.483269 | 0.951933 | 0.139124 | 1.675313 | 0.675514 | 1.020037 | 0.059215 | 0.775871 | 0.679858 | -0.422631 | 0.885378 | 1.681575 | 0.215945 |
| 31.5278 | 0.636262 | 0.334722 | 0.86763 | 0.033789 | 2.116308 | -0.416136 | -0.954145 | 0.639727 | 1.257909 | NaN | 0.481307 | 0.327586 | -0.525581 | 0.267633 | 0.417592 | 0.162498 | -0.040805 | 0.222926 | 0.130709 | -0.509312 | 0.265843 | 0.210279 | -0.148799 |
| 31.5417 | -0.357786 | 0.311691 | 0.169199 | -0.123847 | 2.01314 | -0.354134 | -1.490788 | 0.755361 | 0.63276 | NaN | -0.317373 | -0.362177 | -1.098145 | -1.168958 | 0.221588 | -0.566304 | -0.284351 | -0.316953 | -0.374399 | -0.49356 | -0.320073 | -1.131658 | -0.426414 |
| 31.5556 | -0.950366 | 0.2963 | -0.104174 | -0.13002 | 1.491985 | -0.170644 | -1.363245 | 0.670607 | 0.007753 | NaN | -0.357637 | -0.610816 | -1.478069 | -1.706863 | 0.120805 | -1.040359 | -0.425803 | -0.642918 | -0.50891 | -0.366771 | -0.64613 | 0.906963 | -0.556128 |
| 31.5694 | -1.605599 | 0.16699 | -0.168672 | -0.285318 | 1.401243 | 0.156583 | -1.171638 | 0.264641 | -1.045162 | NaN | -0.393389 | -0.380552 | -1.652878 | -1.374365 | 0.091058 | -1.011928 | -0.575908 | -0.953009 | -0.353737 | -0.217099 | -0.727766 | -0.16232 | -0.575194 |
| 31.5833 | -1.491722 | -0.410493 | -0.209164 | -0.53395 | 1.165697 | -0.059666 | -1.180034 | -0.310133 | -2.054197 | NaN | -1.324479 | -0.036347 | -1.83275 | -1.008346 | 0.020826 | -0.754309 | -0.481644 | -0.915548 | -0.536517 | -0.099661 | -0.733437 | -1.667312 | -0.485582 |
| 31.5972 | -1.331293 | -0.726251 | -0.360245 | -0.34146 | 0.727604 | -0.444169 | -1.3318 | -0.750947 | -2.004585 | NaN | -1.484402 | 0.113937 | -1.801877 | -0.715953 | -0.122403 | -0.805068 | -0.242864 | -0.576694 | -0.635551 | -0.022965 | -0.699279 | 0.994994 | -0.332894 |
| 31.6111 | -0.409642 | -0.613514 | -0.381303 | 0.021465 | 0.172688 | -0.529618 | -1.28601 | -1.037505 | -1.53384 | NaN | -0.516215 | 0.131941 | -1.45358 | -0.486822 | -0.289447 | -0.929214 | 0.035803 | -0.230583 | -0.712232 | 0.043973 | -0.593787 | 6.299546 | -0.196977 |
| 31.625 | 0.279076 | -0.456812 | -0.177057 | 0.358041 | -0.583576 | -0.385054 | -0.95139 | -0.995101 | -0.967892 | NaN | 0.635136 | 0.037725 | -0.920759 | -0.383772 | -0.385278 | -0.961647 | 0.232838 | -0.062927 | -0.637348 | 0.037325 | -0.443307 | 6.369974 | -0.136739 |
| 31.6389 | 0.994417 | -0.519321 | 0.006333 | 0.558642 | -1.590846 | -0.324518 | -0.576028 | -0.603801 | -0.329561 | NaN | 1.165322 | -0.073181 | -0.243472 | -0.264685 | -0.362143 | -0.884358 | 0.084908 | -0.098008 | -0.34171 | -0.01523 | -0.255907 | 5.399834 | -0.171056 |
| 31.6528 | 1.683945 | -0.676422 | 0.091147 | 0.79726 | -2.217388 | -0.399244 | -0.167597 | -0.193181 | 0.723182 | NaN | 0.874007 | -0.165884 | 0.558969 | -0.036946 | -0.224718 | -0.615498 | -0.178994 | -0.233535 | 0.136509 | -0.038107 | -0.121496 | 5.471953 | -0.142693 |
| 31.6667 | 1.745972 | -0.801703 | 0.165967 | 0.636257 | -2.267104 | -0.547081 | 0.227432 | -0.088564 | 1.575083 | NaN | 0.043296 | -0.206905 | 1.28271 | 0.279903 | -0.114509 | -0.098181 | -0.48034 | -0.35653 | 0.495516 | -0.022547 | 0.073347 | 3.089149 | 0.004331 |
| 31.6806 | 2.247867 | -0.739699 | 0.409129 | 0.672117 | -1.889979 | -1.056586 | 0.418554 | -0.439389 | 2.34012 | NaN | -0.934179 | -0.349092 | 1.59274 | 0.367382 | -0.016361 | 0.434865 | -0.543206 | -0.281508 | 0.570691 | 0.034648 | 0.291152 | -1.224133 | 0.114972 |
| 31.6944 | 2.30544 | -0.379577 | 0.611995 | 0.149433 | -1.599833 | -1.34588 | 0.365336 | -1.076538 | 2.060138 | NaN | -1.942465 | -0.540615 | 1.33508 | 0.410198 | 0.080008 | 0.780502 | -0.323984 | -0.07014 | 0.332213 | 0.117706 | 0.394934 | -4.70046 | 0.064216 |
| 31.7083 | 1.41479 | 0.279252 | 0.680078 | 0.012014 | -1.540359 | -1.1376 | 0.676277 | -0.92758 | 1.62439 | NaN | -1.631063 | -0.572121 | 0.783031 | 0.399265 | 0.125871 | 0.822847 | -0.189223 | 0.091209 | -0.155718 | 0.172841 | 0.436877 | -7.752094 | -0.081493 |
| 31.7222 | 0.217723 | 0.689136 | 0.075923 | -0.365046 | -1.482241 | -1.207992 | 0.288525 | -0.835175 | 0.537427 | NaN | -1.248653 | -0.38809 | 0.128078 | 0.356669 | 0.186313 | 0.684317 | -0.069293 | 0.163886 | -0.581145 | 0.16329 | 0.159124 | -6.854621 | -0.281414 |
| 31.7361 | -0.519949 | 0.446662 | -0.20186 | -0.690625 | -1.573753 | -1.62542 | -0.450836 | -0.970389 | -0.605101 | NaN | -1.962756 | -0.185699 | -0.376539 | 0.219475 | 0.259397 | 0.314005 | 0.009478 | 0.095016 | -0.739306 | 0.135549 | -0.253161 | -8.500046 | -0.53659 |
| 31.75 | -1.448289 | -0.562434 | -0.472906 | -1.054004 | -1.504998 | -2.03764 | -1.144947 | -0.93069 | -1.303491 | NaN | -3.152301 | 0.033106 | -0.683292 | -0.076548 | 0.285056 | -0.255255 | 0.04469 | 0.035059 | -0.60437 | 0.106293 | -0.617248 | -8.428834 | -0.578628 |
| 31.7639 | -2.065071 | -1.356583 | -0.593297 | -0.875039 | -1.287282 | -1.997481 | -1.459745 | -0.660694 | -1.546139 | NaN | -3.043534 | 0.027197 | -0.96885 | -0.470514 | 0.245316 | -0.815404 | 0.082268 | 0.047855 | -0.352531 | 0.102663 | -0.808698 | -7.994129 | -0.579173 |
| 31.7778 | -2.268898 | -1.825241 | -0.857502 | -0.854078 | -1.195827 | -1.587315 | -1.611379 | -0.611149 | -1.929942 | NaN | -2.068242 | -0.265211 | -1.32769 | -0.837 | 0.062391 | -1.230282 | 0.087848 | 0.083565 | -0.241919 | 0.084651 | -0.522321 | -5.556495 | -0.55585 |
| 31.7917 | -2.442461 | -1.852074 | -1.349913 | -0.902806 | -1.170644 | -1.490733 | -1.993819 | -0.669652 | -2.147785 | NaN | -1.621107 | -0.624584 | -1.497215 | -1.031308 | -0.142729 | -1.501211 | 0.118999 | 0.105684 | -0.35852 | 0.057411 | -0.274328 | -0.179114 | -0.491274 |
| 31.8056 | -2.39515 | -1.851645 | -1.88043 | -0.986202 | -1.031314 | -1.643804 | -2.596809 | -0.851325 | -2.244717 | NaN | -1.518484 | -0.777293 | -1.428429 | -0.718316 | -0.288383 | -1.320582 | 0.123157 | 0.11507 | -0.479277 | 0.016589 | 0.013607 | -0.159755 | -0.303622 |
| 31.8194 | -2.207768 | -1.798938 | -2.262277 | -0.959925 | -0.852517 | -1.835494 | -2.708459 | -1.056243 | -1.977189 | NaN | -1.166771 | -0.679408 | -1.082584 | -0.452116 | -0.400444 | -1.04331 | 0.117894 | 0.046005 | -0.413771 | 0.021948 | 0.179513 | -0.072913 | -0.149639 |
| 31.8333 | -1.962007 | -2.518483 | -1.969417 | -0.89307 | -0.593411 | -1.474211 | -2.014933 | -1.258584 | -1.508769 | NaN | -1.276094 | -0.410963 | -0.763771 | -0.368809 | -0.461814 | -0.665379 | 0.152587 | 0.038029 | -0.255369 | 0.04553 | 0.175273 | 7.99828 | 0.00285 |

| | | | |
|---|---|---|---|
| 29.2292 | 0.882126 | 29.2222 | 0.120248 |
| 29.2431 | 0.667301 | 29.2361 | 0.011515 |
| 29.2569 | 0.469028 | 29.25 | -0.004519 |
| 29.2708 | 0.280801 | 29.2639 | -0.097993 |
| 29.2847 | 0.273789 | 29.2778 | -0.15448 |
| 29.2986 | 0.19917 | 29.2917 | -0.071849 |
| 29.3125 | 0.248298 | 29.3056 | 0.033527 |
| 29.3264 | 0.625305 | 29.3194 | 0.117861 |
| 29.3403 | 0.680728 | 29.3333 | 0.268761 |
| 29.3542 | 0.31314 | 29.3472 | 0.293717 |
| 29.3681 | 0.274008 | 29.3611 | 0.153899 |
| 29.3819 | 0.384583 | 29.375 | 0.045853 |
| 29.3958 | 1.139616 | 29.3889 | 0.041398 |
| 29.4097 | 7.38184 | 29.4028 | 0.047808 |
| 29.4236 | 18.24158 | 29.4167 | 0.145807 |
| 29.4375 | 19.169591 | 29.4306 | 0.162184 |
| 29.4514 | 10.226039 | 29.4444 | 0.500808 |
| 29.4653 | 3.725783 | 29.4583 | -0.316894 |
| 29.4792 | 2.840394 | 29.4722 | 1.095185 |
| 29.4931 | 2.109937 | 29.4861 | 1.564364 |
| 29.5 | 1.280231 | 29.5 | 1.313617 |
| 29.5139 | 0.432297 | 29.5139 | 1.113898 |
| 29.5278 | -0.460737 | 29.5278 | 0.938101 |
| 29.5417 | -1.418666 | 29.5417 | 0.557314 |
| 29.5556 | -2.368124 | 29.5556 | 0.282745 |
| 29.5694 | -2.379636 | 29.5694 | 0.153781 |
| 29.5833 | -1.949029 | 29.5833 | 0.224034 |
| 29.5972 | -0.834901 | 29.5972 | 0.29734 |
| 29.6111 | 0.111051 | 29.6111 | 0.351288 |
| 29.625 | 0.764243 | 29.625 | 0.372204 |
| 29.6389 | 0.715467 | 29.6389 | 0.288413 |
| 29.6528 | 0.132647 | 29.6528 | 0.166636 |
| 29.6667 | -0.041365 | 29.6667 | 0.056731 |
| 29.6806 | 0.047915 | 29.6806 | -0.011515 |
| 29.6944 | -0.240768 | 29.6944 | -0.005018 |
| 29.7083 | -2.133119 | 29.7083 | 0.05159 |
| 29.7222 | -3.821782 | 29.7222 | 0.068935 |
| 29.7361 | -4.52176 | 29.7361 | -0.045635 |
| 29.75 | -4.295792 | 29.75 | -0.347625 |
| 29.7639 | -3.009614 | 29.7639 | -0.684986 |
| 29.7778 | -2.332369 | 29.7778 | -1.036684 |
| 29.7917 | -2.116019 | 29.7917 | -1.374541 |
| 29.8056 | -1.647551 | 29.8056 | -1.435727 |
| 29.8194 | -0.906405 | 29.8194 | -1.127427 |
| 29.8333 | -0.05371 | 29.8333 | -0.67704 |
| 29.8472 | 0.581129 | 29.8472 | -0.302966 |
| 29.8611 | 0.710095 | 29.8611 | -0.141307 |
| 29.875 | 0.694289 | 29.875 | -0.033189 |
| 29.8889 | -0.699323 | 29.8889 | 0.04093 |
| 29.9028 | -9.101848 | 29.9028 | -0.027601 |
| 29.9167 | -23.695503 | 29.9167 | -0.057064 |
| 29.9306 | -26.43405 | 29.9306 | -0.270922 |
| 29.9444 | -15.224804 | 29.9444 | -1.900025 |
| 29.9583 | -7.117487 | 29.9583 | -3.348642 |
| 29.9722 | -4.402052 | 29.9722 | -2.687283 |
| 29.9861 | -2.321934 | 29.9861 | -0.675096 |
| 30.0069 | -0.836908 | 30 | 0.603625 |
| 30.0208 | 0.113461 | 30.0139 | 0.778827 |
| 30.0347 | 1.167848 | 30.0278 | 0.671312 |
| 30.0486 | 1.990376 | 30.0417 | 0.72573 |
| 30.0625 | 2.275282 | 30.0556 | 0.757923 |
| 30.0764 | 2.474898 | 30.0694 | 0.62602 |
| 30.0903 | 2.90443 | 30.0833 | 0.536785 |
| 30.1042 | 2.513131 | 30.0972 | 0.521794 |
| 30.1181 | 2.106381 | 30.1111 | 0.477053 |
| 30.1319 | 2.339837 | 30.125 | 0.308316 |
| 30.1458 | 2.661147 | 30.1389 | 0.05627 |
| 30.1597 | 2.602834 | 30.1528 | 0.008004 |
| 30.1736 | 2.149192 | 30.1667 | 0.115197 |
| 30.1875 | 1.219728 | 30.1806 | 0.185551 |
| 30.2014 | 0.652977 | 30.1944 | 0.223157 |
| 30.2153 | 0.867766 | 30.2083 | 0.362797 |
| 30.2292 | 1.128836 | 30.2222 | 0.472406 |
| 30.2431 | 1.135159 | 30.2361 | 0.525874 |
| 30.2569 | 1.326357 | 30.25 | 0.516823 |
| 30.2708 | 1.058515 | 30.2639 | 0.442129 |
| 30.2847 | 0.356837 | 30.2778 | 0.330464 |
| 30.2986 | 0.057022 | 30.2917 | 0.180349 |
| 30.3125 | 0.23765 | 30.3056 | 0.191807 |
| 30.3264 | 0.782457 | 30.3194 | 0.342746 |
| 30.3403 | 1.212041 | 30.3333 | 0.424883 |
| 30.3542 | 1.133346 | 30.3472 | 0.398137 |
| 30.3681 | 0.721067 | 30.3611 | 0.312059 |
| 30.3819 | 0.249902 | 30.375 | 0.274222 |
| 30.3958 | 0.500715 | 30.3889 | 0.336271 |
| 30.4097 | 5.515168 | 30.4028 | 0.380159 |
| 30.4236 | 14.572382 | 30.4167 | 0.502199 |
| 30.4375 | 15.656138 | 30.4306 | 1.812336 |
| 30.4514 | 7.848529 | 30.4444 | 3.712123 |
| 30.4653 | 2.487136 | 30.4583 | 5.063236 |
| 30.4792 | 1.605812 | 30.4722 | 2.860019 |
| 30.4931 | 0.998268 | 30.4861 | 1.759623 |
| 30.5 | 0.781523 | 30.5 | 1.830771 |
| 30.5139 | 0.981829 | 30.5139 | 1.64594 |
| 30.5278 | 1.258323 | 30.5278 | 1.267043 |
| 30.5417 | 1.585401 | 30.5417 | 0.935123 |
| 30.5556 | 1.083849 | 30.5556 | 0.41117 |
| 30.5694 | -0.93042 | 30.5694 | -0.018755 |
| 30.5833 | -2.013025 | 30.5833 | -0.236287 |
| 30.5972 | -1.347135 | 30.5972 | -0.180425 |
| 30.6111 | -0.992642 | 30.6111 | -0.091527 |
| 30.625 | -0.467919 | 30.625 | -0.104069 |
| 30.6389 | -0.35894 | 30.6389 | -0.246051 |
| 30.6528 | -1.205049 | 30.6528 | -0.346417 |
| 30.6667 | -2.074724 | 30.6667 | -0.31036 |
| 30.6806 | -2.215155 | 30.6806 | -0.109957 |
| 30.6944 | -1.561717 | 30.6944 | 0.100576 |
| 30.7083 | -0.567869 | 30.7083 | 0.220571 |
| 30.7222 | 0.251398 | 30.7222 | 0.273807 |
| 30.7361 | 0.902502 | 30.7361 | 0.285835 |
| 30.75 | 1.38843 | 30.75 | 0.316475 |
| 30.7639 | 1.604467 | 30.7639 | 0.190261 |
| 30.7778 | -0.865609 | 30.7778 | -0.05128 |
| 30.7917 | -2.330573 | 30.7917 | -0.427674 |
| 30.8056 | -1.933311 | 30.8056 | -0.808081 |
| 30.8194 | -1.97327 | 30.8194 | -1.015278 |
| 30.8333 | -2.35617 | 30.8333 | -0.981001 |
| 30.8472 | -1.727736 | 30.8472 | -0.787515 |
| 30.8611 | -1.068412 | 30.8611 | -0.487253 |
| 30.875 | -0.645935 | 30.875 | -0.209296 |
| 30.8889 | -1.369465 | 30.8889 | -0.017283 |
| 30.9028 | -8.641385 | 30.9028 | 0.057879 |
| 30.9167 | -19.057704 | 30.9167 | -0.276194 |
| 30.9306 | -18.287773 | 30.9306 | -2.724853 |
| 30.9444 | -9.102933 | 30.9444 | -7.795961 |
| 30.9583 | -3.47783 | 30.9583 | -8.803018 |
| 30.9722 | -2.005733 | 30.9722 | -3.866396 |
| 30.9861 | -1.305463 | 30.9861 | -0.668364 |
| 31.0069 | -0.114057 | 31 | 0.807325 |
| 31.0208 | 0.852779 | 31.0139 | 1.191014 |
| 31.0347 | 1.98428 | 31.0278 | 1.140025 |
| 31.0486 | 3.128179 | 31.0417 | 1.229742 |
| 31.0625 | 2.571239 | 31.0556 | 1.439057 |
| 31.0764 | 0.44424 | 31.0694 | 1.421576 |
| 31.0903 | 0.18737 | 31.0833 | 1.026557 |
| 31.1042 | 1.170767 | 31.0972 | 0.325751 |
| 31.1181 | -0.308383 | 31.1111 | -0.403829 |
| 31.1319 | -1.913059 | 31.125 | -0.709131 |
| 31.1458 | -0.828823 | 31.1389 | -0.290454 |
| 31.1597 | 0.59294 | 31.1528 | 0.156906 |
| 31.1736 | 1.046464 | 31.1667 | 0.251994 |
| 31.1875 | 1.019743 | 31.1806 | 0.166479 |
| 31.2014 | 1.306652 | 31.1944 | -0.089886 |
| 31.2153 | 2.183354 | 31.2083 | -0.416085 |
| 31.2292 | 3.171806 | 31.2222 | -0.564322 |
| 31.2431 | 2.770137 | 31.2361 | -0.335644 |
| 31.2569 | 1.01918 | 31.25 | 0.075914 |
| 31.2708 | -0.701063 | 31.2639 | 0.261102 |
| 31.2847 | -1.712862 | 31.2778 | 0.241584 |
| 31.2986 | -1.473526 | 31.2917 | 0.258532 |
| 31.3125 | -0.356621 | 31.3056 | 0.271559 |
| 31.3264 | 0.817556 | 31.3194 | 0.178661 |
| 31.3403 | 1.667453 | 31.3333 | 0.175913 |
| 31.3542 | 0.604393 | 31.3472 | 0.335208 |
| 31.3681 | -0.347739 | 31.3611 | 0.442455 |
| 31.3819 | 1.045292 | 31.375 | 0.410469 |
| 31.3958 | 3.049228 | 31.3889 | 0.433746 |
| 31.4097 | 8.211416 | 31.4028 | 0.47965 |
| 31.4236 | 16.712343 | 31.4167 | 0.597731 |
| 31.4375 | 17.580144 | 31.4306 | 2.843373 |
| 31.4514 | 7.008536 | 31.4444 | 7.430816 |
| 31.4653 | 2.162687 | 31.4583 | 8.04982 |
| 31.4792 | 1.898928 | 31.4722 | 4.020705 |
| 31.4931 | 0.869776 | 31.4861 | 0.929729 |
| 31.5 | 0.077889 | 31.5 | 0.475126 |
| 31.5139 | 0.483269 | 31.5139 | 0.215945 |
| 31.5278 | 0.481307 | 31.5278 | -0.148799 |
| 31.5417 | -0.317373 | 31.5417 | -0.426414 |
| 31.5556 | -0.357637 | 31.5556 | -0.556128 |
| 31.5694 | -0.393389 | 31.5694 | -0.575194 |
| 31.5833 | -1.324479 | 31.5833 | -0.485582 |
| 31.5972 | -1.484402 | 31.5972 | -0.332894 |
| 31.6111 | -0.516215 | 31.6111 | -0.196977 |
| 31.625 | 0.635136 | 31.625 | -0.136739 |
| 31.6389 | 1.165322 | 31.6389 | -0.171056 |
| 31.6528 | 0.874007 | 31.6528 | -0.142693 |
| 31.6667 | 0.043296 | 31.6667 | 0.004331 |
| 31.6806 | -0.934179 | 31.6806 | 0.114972 |
| 31.6944 | -1.942465 | 31.6944 | 0.064216 |
| 31.7083 | -1.631063 | 31.7083 | -0.081493 |
| 31.7222 | -1.248653 | 31.7222 | -0.281414 |
| 31.7361 | -1.962756 | 31.7361 | -0.53659 |
| 31.75 | -3.152301 | 31.75 | -0.578628 |
| 31.7639 | -3.043534 | 31.7639 | -0.579173 |
| 31.7778 | -2.068242 | 31.7778 | -0.55585 |
| 31.7917 | -1.621107 | 31.7917 | -0.491274 |
| 31.8056 | -1.518484 | 31.8056 | -0.303622 |
| 31.8194 | -1.166771 | 31.8194 | -0.149639 |
| 31.8333 | -1.276094 | 31.8333 | 0.00285 |

| | | | | | | | | | | |
|---|---|---|---|---|---|---|---|---|---|---|
| 31.8472 | -1.106744 | -1.7144 | -1.198572 | -0.628938 | -0.343793 | -1.006606 | -1.392622 | -1.210544 | -1.461783 | NaN |
| 31.8611 | -0.097505 | -0.889404 | -0.527328 | -0.060573 | -0.085025 | -0.553742 | -0.765039 | -0.747772 | -0.934211 | NaN |
| 31.875 | 0.648721 | -0.233384 | 0.112249 | 0.53866 | 0.135399 | -0.113589 | -0.113049 | -0.404165 | -0.105814 | NaN |
| 31.8889 | 0.825064 | 0.434607 | 0.693554 | 0.944945 | 0.200249 | 0.011178 | 0.538034 | -0.401409 | 0.714419 | NaN |
| 31.9028 | -0.812494 | -0.000938 | 1.216709 | 0.560781 | -0.297046 | -0.585844 | 0.735957 | -1.403515 | 1.203594 | NaN |
| 31.9167 | -4.292531 | -2.012739 | 0.225649 | -1.933099 | -1.593623 | -2.00965 | -0.222096 | -3.298859 | 0.154903 | -5.790423 |
| 31.9306 | -6.366915 | -4.484189 | -2.248302 | -4.191982 | -2.192499 | -3.263809 | -2.048868 | -4.250741 | -2.283175 | -9.553457 |
| 31.9444 | -5.805382 | -4.896633 | -3.083016 | -4.587446 | -2.321565 | -3.649367 | -3.222429 | -2.645395 | -3.636879 | -0.4815 |
| 31.9583 | -3.653289 | -3.841461 | -2.557984 | -3.182717 | -1.894934 | -3.07221 | -2.567424 | -1.053392 | -3.067927 | -0.028446 |
| 31.9722 | -2.56249 | -3.517948 | -2.621087 | -2.029343 | -1.402335 | -2.618124 | -2.318676 | -0.569114 | -2.683201 | -0.396788 |
| 31.9861 | -1.978421 | -3.497291 | -2.483326 | -1.218384 | -1.258779 | -2.018952 | -1.673413 | -0.419107 | -1.631155 | -0.103316 |
| 32.0069 | -0.848571 | -2.40825 | -1.652987 | -0.402945 | -0.86102 | -1.050615 | -0.655406 | -0.086026 | -0.417558 | -0.056338 |
| 32.0208 | 0.143367 | -1.255808 | -0.703425 | 0.09804 | -0.283999 | 0.05987 | 0.050014 | 0.15713 | 0.499219 | -0.176682 |
| 32.0347 | 0.509758 | -0.76842 | -0.109724 | 0.309777 | 0.179957 | 0.940954 | 0.60539 | 0.10542 | 1.028577 | 0.164507 |
| 32.0486 | 0.715739 | -0.480013 | 0.307136 | 0.421973 | 0.488427 | 1.584992 | 0.939499 | 0.098001 | 1.300592 | 0.469974 |
| 32.0625 | 1.175647 | 0.099732 | 0.59776 | 0.545237 | 0.692538 | 1.891661 | 0.94055 | 0.249781 | 1.412344 | 0.290541 |
| 32.0764 | 1.775276 | 0.756926 | 0.592521 | 0.586956 | 0.878946 | 2.177868 | 0.952269 | 0.496901 | 1.566754 | -0.252728 |
| 32.0903 | 2.003821 | 1.124418 | 0.658077 | 0.402599 | 0.904614 | 2.279829 | 1.037451 | 0.729712 | 1.855889 | -0.191983 |

[illegible]

| | | | | | | | | | | | | | | | | | | | | | | | |
|---|---|---|---|---|---|---|---|---|---|---|---|---|---|---|---|---|---|---|---|---|---|---|---|
| 34.4792 | 3.329201 | 4.414964 | 3.412571 | 3.367244 | 2.232727 | 2.776269 | 2.364773 | 0.600496 | 4.138336 | 0.096331 | 2.64924 | 1.087654 | 2.247141 | 0.748746 | 2.979479 | 1.983609 | 2.742561 | 0.796007 | 2.774283 | 1.936649 | -0.177379 | 1.617455 | 8.225337 |
| 34.4931 | 3.078135 | 5.347975 | 2.601438 | 3.084925 | 2.065163 | 2.610835 | 1.96112 | 0.400751 | 4.213203 | 0.172831 | 3.007158 | 1.333817 | 1.528064 | 1.440653 | 1.008605 | 2.236505 | 0.706589 | 0.615586 | 2.053692 | 1.378517 | 0.40526 | 0.265185 | 1.854185 |
| 34.5 | 2.266453 | 5.79653 | 1.625306 | 1.956765 | 1.831685 | 2.868307 | 1.307858 | 0.514119 | 4.508533 | 0.234649 | 3.075496 | 1.193346 | 1.126557 | 1.393652 | 0.622984 | 1.957092 | 0.426068 | 0.372695 | 1.458712 | 0.950259 | 0.35191 | 0.095964 | 0.949355 |
| 34.5139 | 1.429498 | 5.68836 | 0.699126 | 0.781481 | 1.287587 | 2.585379 | 0.66518 | 0.732643 | 4.413524 | 0.223424 | 2.656514 | 0.865996 | 1.120026 | 1.220674 | 0.388417 | 1.701263 | 0.345155 | 0.198111 | 1.208857 | 0.685676 | 0.151954 | 0.221586 | 0.653242 |
| 34.5278 | 1.241442 | 5.043516 | 0.013577 | -0.057627 | 0.811796 | 2.186733 | 0.435878 | 0.713596 | 4.579964 | 0.212859 | 2.100327 | 0.408929 | 1.36848 | 0.961737 | 0.218045 | 1.308862 | -0.00331 | 0.011737 | 0.77998 | 0.185146 | -0.093494 | 0.330246 | 0.109922 |
| 34.5417 | 1.251925 | 3.926664 | -0.272077 | -0.395487 | 0.75203 | 1.874384 | 0.638662 | 0.390273 | 3.731403 | 0.16526 | 1.552476 | -0.101225 | 1.387853 | 0.504432 | 0.034067 | 1.080837 | -0.359683 | -0.217161 | 0.352381 | -0.54676 | -0.303854 | 0.231709 | -0.400336 |
| 34.5556 | 0.757873 | 2.712959 | -0.350802 | -0.429051 | 0.631115 | 1.33199 | 1.018093 | 0.028328 | 2.293724 | 0.049137 | 1.056215 | -0.453203 | 1.262868 | 0.062938 | -0.113374 | 1.073976 | -0.523491 | -0.476 | 0.197152 | -1.328483 | -0.060829 | 0.048921 | -0.64666 |
| 34.5694 | 0.080645 | 0.805571 | -0.507993 | -1.080398 | -0.106301 | 0.124229 | 1.160093 | -0.321516 | 0.724533 | -0.008577 | 0.112288 | -0.451185 | 1.009267 | -0.203712 | -0.07618 | 1.101112 | -0.451276 | -0.302306 | 0.422857 | -1.333337 | 0.20211 | 0.001804 | -0.638341 |
| 34.5833 | -0.820413 | -0.959705 | -0.853586 | -1.556306 | -1.305468 | -2.042283 | 1.099188 | -0.57759 | -0.807955 | -0.135135 | -1.100043 | -0.416575 | 0.92669 | -0.256904 | -0.008577 | 1.069879 | -0.267861 | -0.05989 | 0.439824 | -1.059404 | 0.302592 | 0.056792 | -0.680568 |
| 34.5972 | -1.524611 | -2.397519 | -1.016306 | -0.962188 | -1.990442 | -2.966901 | 1.384252 | -0.792844 | -2.382141 | 0.026425 | -2.151501 | -0.517463 | 0.875451 | -0.447242 | 0.007559 | 0.683817 | -0.025751 | 0.119378 | 0.303758 | -0.899645 | 0.213702 | 0.108163 | -0.7029 |
| 34.6111 | -1.745194 | -3.209844 | -0.961104 | -0.445798 | -2.536243 | -3.385356 | 1.575168 | -0.998094 | -3.781184 | -0.0606 | -2.911751 | -0.648454 | 0.817397 | -0.6297 | -0.128979 | 0.166196 | 0.1256 | 0.067817 | 0.1258 | -0.820054 | -0.087068 | 0.046136 | -0.906638 |
| 34.625 | -1.553444 | -3.398964 | -0.723694 | -0.222987 | -2.818789 | -2.947784 | 1.584038 | -1.005325 | -4.202497 | -0.104617 | -3.124337 | -0.607119 | 0.729907 | -0.674613 | -0.238657 | -0.203576 | 0.000849 | -0.015425 | -0.09136 | -0.769351 | -0.295917 | -0.061894 | -0.965825 |
| 34.6389 | -1.221193 | -2.945405 | -0.506388 | -0.043606 | -2.292927 | -2.054668 | 1.673703 | -0.775505 | -4.168578 | -0.036892 | -2.787518 | -0.273886 | 0.823203 | -0.530998 | -0.072268 | -0.324559 | -0.272079 | -0.077928 | -0.294765 | -0.585662 | -0.178578 | -0.049758 | -0.732936 |
| 34.6528 | -0.714392 | -2.302826 | -0.1867 | 0.252741 | -1.380119 | -0.97308 | 1.715432 | -0.679188 | -4.188707 | 0.07441 | -2.592479 | 0.181311 | 0.898526 | -0.38565 | 0.147258 | -0.406374 | -0.573252 | -0.200882 | -0.432591 | -0.328085 | 0.079608 | 0.024308 | -0.346541 |
| 34.6667 | -0.371008 | -1.617457 | 0.349623 | 0.451192 | -0.470262 | -0.007589 | 1.711755 | -0.613113 | -3.913035 | 0.058028 | -2.108469 | 0.56617 | 0.763717 | -0.208675 | 0.2716 | -0.194027 | -0.690711 | -0.26417 | -0.428144 | 0.027668 | 0.182095 | 0.005312 | 0.080443 |
| 34.6806 | -0.209106 | -1.088573 | 1.047628 | 0.85251 | 0.101137 | 0.745819 | 1.705951 | -0.436843 | -2.880528 | -0.06371 | -1.572614 | 0.967177 | 0.51114 | -0.06573 | 0.253863 | 0.08922 | -0.480049 | -0.147101 | -0.266312 | 0.464208 | 0.105806 | -0.040822 | 0.47359 |
| 34.6944 | -0.046413 | -0.616295 | 1.503602 | 0.963431 | 0.110558 | 0.951941 | 1.579379 | -0.224469 | -1.684404 | -0.103228 | -0.980691 | 1.152039 | 0.223933 | 0.160093 | 0.386281 | 0.299869 | -0.176257 | -0.051092 | -0.116644 | 0.803188 | 0.01884 | -0.0083 | 0.828116 |
| 34.7083 | -0.036291 | -0.241901 | 1.41991 | 0.779201 | -0.03199 | 0.548829 | 1.467305 | -0.260017 | -0.842295 | -0.021561 | -0.647378 | 1.016264 | -0.027047 | 0.45328 | 0.540822 | 0.314295 | 0.129501 | 0.055722 | -0.078987 | 0.738581 | -0.018805 | 0.082713 | 0.876795 |
| 34.7222 | -0.009596 | 0.07941 | 1.077753 | 0.611478 | -0.191098 | -0.041124 | 1.342538 | -0.499463 | -0.205948 | 0.041735 | -0.556335 | 0.713187 | -0.278451 | 0.555834 | 0.502499 | 0.16069 | 0.356471 | 0.084055 | -0.129675 | 0.170212 | -0.098862 | 0.11525 | 0.624789 |
| 34.7361 | 0.068936 | 0.260292 | 0.762228 | 0.339797 | -0.46731 | -0.504159 | 1.081178 | -0.602664 | 0.27378 | -0.020915 | -0.621668 | 0.149341 | -0.554062 | 0.499358 | 0.321039 | -0.076898 | 0.335806 | 0.111737 | -0.152244 | -0.477566 | -0.09436 | 0.003013 | 0.147135 |
| 34.75 | 0.115547 | 0.284131 | 0.284925 | -0.001321 | -0.884154 | -1.008184 | 0.747002 | -0.622139 | 0.566478 | -0.109326 | -0.677233 | -0.476753 | -0.749571 | 0.377047 | 0.185652 | -0.200262 | 0.233808 | 0.144143 | -0.178162 | -0.902224 | 0.085169 | -0.07558 | -0.290198 |
| 34.7639 | -0.003201 | 0.13773 | -0.196976 | -0.204383 | -1.374868 | -1.457399 | 0.389491 | -0.667597 | 0.61509 | -0.103115 | -0.892824 | -0.785732 | -0.787691 | 0.301592 | 0.138832 | -0.135731 | 0.036364 | 0.170905 | -0.167852 | -0.875068 | 0.38561 | 0.053083 | -0.54711 |
| 34.7778 | -0.269859 | -0.229929 | -0.784369 | -0.403968 | -1.464399 | -1.777791 | -0.012532 | -0.797414 | 0.14588 | 0.007781 | -1.314763 | -0.856333 | -0.874724 | 0.245969 | 0.1523 | 0.002339 | -0.123207 | 0.150612 | -0.06895 | -0.682175 | 0.649483 | 0.197828 | -0.647206 |
| 34.7917 | -0.710236 | -0.797385 | -1.209505 | -0.51272 | -1.532454 | -2.159472 | -0.198561 | -0.847676 | -0.729718 | 0.024282 | -1.859646 | -0.998622 | -1.064238 | 0.070177 | 0.166088 | 0.132983 | -0.318306 | 0.081792 | 0.022562 | -0.434806 | 0.568318 | 0.113568 | -0.649315 |
| 34.8056 | -0.98774 | -1.374856 | -1.308422 | -0.613831 | -2.1314 | -2.721382 | -0.203465 | -0.789261 | -1.7001 | -0.150904 | -2.410037 | -1.150549 | -1.451579 | -0.161282 | 0.18163 | 0.250124 | -0.415328 | 0.077233 | 0.018673 | -0.266724 | 0.293497 | -0.099888 | -0.618593 |
| 34.8194 | -1.196213 | -1.436082 | -1.313757 | -0.473191 | -2.715063 | -3.18794 | -0.134505 | -0.622672 | -2.358794 | -0.108991 | -2.829004 | -1.130965 | -1.785166 | -0.232522 | 0.255038 | 0.303085 | -0.303022 | 0.134535 | -0.037183 | -0.018927 | 0.166155 | -0.167521 | -0.562321 |
| 34.8333 | -1.411046 | -1.153254 | -1.549819 | -0.55655 | -2.928757 | -2.824239 | -0.191836 | -0.505852 | -2.913285 | -0.055361 | -2.867256 | -0.975947 | -1.987515 | -0.182512 | 0.278546 | 0.341783 | -0.118997 | 0.212 | -0.031103 | 0.267281 | 0.200084 | -0.132976 | -0.476819 |
| 34.8472 | -1.033523 | -0.517121 | -1.824416 | -0.609506 | -2.474283 | -2.542654 | -0.300319 | -0.355022 | -3.264548 | 0.058305 | -2.979838 | -0.700154 | -1.93066 | 0.016886 | 0.335081 | 0.435128 | 0.048374 | 0.271593 | -0.089327 | 0.434342 | 0.297225 | -0.00788 | -0.286863 |
| 34.8611 | -0.394714 | -0.277928 | -1.01884 | -0.42152 | -1.040304 | -2.028996 | -0.210665 | 0.087362 | -3.163458 | 0.037575 | -2.273506 | -0.500677 | -1.671575 | 0.170716 | 0.396798 | 0.47457 | 0.090065 | 0.263123 | -0.146949 | 0.22266 | 0.26559 | -0.021273 | -0.116479 |
| 34.875 | 0.284768 | 0.107975 | -0.116168 | -0.076519 | 1.144022 | -1.336191 | 0.090863 | 0.586314 | -2.364735 | -0.102097 | -0.929198 | -0.389204 | -1.530089 | 0.167968 | 0.419772 | 0.477378 | 0.060853 | 0.23522 | -0.20739 | -0.036076 | 0.177128 | 0.018426 | -0.104235 |
| 34.8889 | 0.752715 | 0.491319 | 0.63226 | 0.240252 | 3.33986 | -0.831552 | 0.253834 | 0.884669 | -1.804276 | -0.187322 | -1.153821 | -0.27709 | -1.305074 | 0.145165 | 0.369222 | 0.408719 | 0.075835 | 0.219299 | -0.217352 | 0.047579 | 0.247342 | 0.015495 | -0.075794 |
| 34.9028 | -0.429298 | -0.059076 | 1.294343 | -0.086453 | 4.255043 | -0.870406 | -0.998877 | 0.374838 | -2.960922 | -0.043072 | -11.794476 | -0.104608 | -1.17583 | 0.230831 | 0.3584 | 0.18593 | 0.061134 | 0.243213 | -0.169949 | 0.414839 | 0.263142 | NaN | 0.033593 |
| 34.9167 | -2.869203 | -2.19207 | -0.822672 | -1.181069 | 2.252243 | -1.759311 | -3.027721 | -0.631888 | -5.048989 | 2.078531 | -28.284357 | 0.138867 | -0.977397 | 0.391435 | -0.033717 | -0.151499 | -0.08101 | 0.216374 | -0.163087 | 0.61638 | 0.268641 | NaN | -1.074569 |
| 34.9306 | -3.891057 | -4.513029 | -3.480397 | -1.800221 | 1.452337 | -3.500501 | -3.476674 | -0.893778 | -5.772858 | 2.603192 | -24.647908 | 0.546163 | -1.991382 | 0.294188 | -1.952312 | -0.597804 | -1.405587 | -0.131878 | -0.659843 | -0.417605 | 0.039731 | -1.825019 | -9.344092 |
| 34.9444 | -2.600779 | -4.743129 | -2.263063 | -1.035215 | 0.733873 | -3.341993 | -2.033448 | -1.407161 | -4.110573 | 0.942545 | -10.611492 | 0.175017 | -4.591783 | -1.382493 | -3.822309 | -2.856266 | -3.413173 | -1.113167 | -3.314786 | -3.761791 | -1.210984 | -3.512745 | -20.09496 |
| 34.9583 | -1.79691 | -3.937566 | -1.218703 | -0.369507 | 0.113471 | -2.256442 | -0.684176 | -1.1497 | -3.007842 | 0.042642 | -2.747789 | -1.452782 | -5.962405 | -3.859362 | -4.0347 | -4.90017 | -4.214421 | -3.14622 | -4.895628 | -5.999165 | -2.961221 | -1.835783 | -18.822915 |
| 34.9722 | -1.903422 | -4.096466 | -0.786256 | -0.198403 | -0.532987 | -1.604285 | -0.312175 | -0.911431 | -2.762374 | -0.025562 | -1.8744 | -2.367138 | -4.210205 | -2.910381 | -2.60379 | -4.239205 | -2.378951 | -2.099052 | -3.546891 | -5.561189 | -2.884853 | -0.493612 | -8.652979 |
| 34.9861 | -1.788347 | -4.114053 | -0.405408 | -0.25565 | -0.876781 | -1.37744 | -0.177821 | -0.574249 | -2.634967 | -0.030136 | -2.240505 | -1.774837 | -1.581104 | -1.030508 | -1.180324 | -1.998216 | -0.483311 | -0.604047 | -1.367484 | -2.602099 | -1.307933 | -0.141306 | -2.690254 |
| 35.0069 | -1.436605 | -3.646005 | -0.181399 | -0.368512 | -1.021022 | -1.25743 | -0.063041 | -0.189919 | -2.060818 | -0.100726 | -2.131781 | -0.660078 | 0.061553 | 0.432638 | -0.030981 | -0.601697 | 0.265655 | 0.391972 | -0.092181 | -0.017984 | -0.034217 | 0.034988 | -0.574871 |
| 35.0208 | -1.257122 | -3.26785 | -0.069061 | -0.565387 | -1.180193 | -1.472586 | -0.131328 | -0.036099 | -1.834592 | -0.191505 | -1.891135 | 0.057979 | 0.50829 | 1.026129 | 0.815243 | 0.260961 | 0.39668 | 0.725797 | 0.783608 | 1.706065 | 0.306239 | 0.156172 | 0.645469 |
| 35.0347 | -1.225804 | -2.998091 | 0.088299 | -0.539606 | -1.03675 | -1.709789 | -0.185711 | -0.262238 | -2.203777 | -0.20496 | -1.888718 | 0.797326 | 0.55478 | 1.288281 | 1.571839 | 0.844082 | 0.52368 | 0.726855 | 1.217584 | 3.030608 | 0.208404 | 0.189949 | 1.558157 |
| 35.0486 | -1.175776 | -2.750224 | 0.270822 | -0.323735 | -0.533526 | -1.740369 | -0.073315 | -0.551496 | -2.628993 | -0.311946 | -1.786363 | 1.317831 | 0.523813 | 1.731228 | 2.023261 | 0.988899 | 0.478465 | 0.598194 | 1.275896 | 3.75629 | 0.138134 | 0.088442 | 1.95174 |
| 35.0625 | -0.849605 | -2.178089 | 0.458814 | -0.046344 | 0.221027 | -1.64403 | 0.159735 | -0.544146 | -2.359832 | -0.304192 | -1.430535 | 1.220259 | 0.314987 | 1.848185 | 1.813109 | 0.927683 | 0.33826 | 0.421529 | 1.109435 | 3.591781 | 0.204395 | -0.041091 | 2.001013 |
| 35.0764 | -0.250169 | -1.241372 | 0.684347 | 0.424207 | 1.271276 | -1.235492 | 0.534404 | -0.363981 | -1.557311 | -0.200246 | -0.58866 | 0.884024 | 0.065337 | 1.609673 | 1.455065 | 0.938598 | 0.127806 | 0.188953 | 1.033143 | 3.506591 | 0.258972 | -0.038464 | 1.868549 |
| 35.0903 | 0.509365 | -0.167807 | 0.972673 | 1.076597 | 2.493119 | -0.684741 | 1.020059 | -0.083487 | -0.743585 | -0.003114 | 0.903519 | 0.636481 | -0.121112 | 1.056617 | 1.08145 | 0.978352 | -0.057754 | -0.00198 | 0.843226 | 2.88629 | 0.093569 | 0.052318 | 1.387982 |
| 35.1042 | 0.998565 | 0.493039 | 1.262579 | 1.574288 | 3.466241 | -0.185205 | 1.425678 | 0.327547 | -0.047991 | 0.092461 | 1.669049 | 0.512701 | -0.106558 | 0.608342 | 0.510836 | 1.051521 | -0.181581 | -0.088386 | 0.586993 | 1.803265 | -0.06713 | 0.066614 | 0.776888 |
| 35.1181 | 1.179419 | 0.887401 | 1.446968 | 1.889074 | 3.682356 | 0.196427 | 1.67418 | 0.783913 | 0.651317 | 0.042161 | 1.41135 | 0.515686 | 0.080326 | 0.372354 | 0.06999 | 1.353812 | -0.170658 | -0.062779 | 0.357821 | 0.60576 | -0.038996 | -0.041806 | 0.587575 |
| 35.1319 | 1.427316 | 1.356698 | 1.630632 | 1.982979 | 3.79042 | 0.586474 | 1.709909 | 1.025197 | 1.088068 | 0.012748 | 1.390897 | 0.632844 | 0.3043 | 0.359812 | -0.377302 | 1.810866 | -0.121202 | 0.021142 | 0.193805 | -0.575909 | 0.107045 | -0.126522 | 0.465885 |
| 35.1458 | 1.793574 | 1.601357 | 1.805081 | 2.101104 | 3.256567 | 0.608516 | 1.755743 | 0.929758 | 1.047551 | 0.019096 | 1.56059 | 0.697294 | 0.418427 | 0.394661 | -0.608701 | 1.598262 | 0.010647 | 0.17123 | 0.232321 | -1.748962 | 0.214278 | -0.103919 | 0.407857 |
| 35.1597 | 2.010589 | 1.529961 | 1.618815 | 2.343185 | 2.849222 | 0.268235 | 1.602497 | 0.576809 | 0.417677 | -0.057578 | 1.352013 | 0.667965 | 0.441464 | 0.385188 | -0.567396 | 1.189612 | 0.275051 | 0.249391 | 0.42214 | -2.320365 | 0.253779 | 0.004638 | 0.358975 |
| 35.1736 | 1.993543 | 1.179104 | 1.324817 | 2.469221 | 2.695714 | -0.126576 | 1.563959 | -0.115649 | -0.411773 | -0.145128 | 1.095802 | 0.587157 | 0.373214 | 0.378513 | -0.3295 | 0.522598 | 0.275025 | 0.187613 | 0.410445 | -2.558951 | 0.234615 | 0.036386 | 0.395818 |
| 35.1875 | 1.609262 | 0.838237 | 0.954515 | 1.869335 | 2.085705 | -0.43807 | 1.238632 | -0.679916 | -0.879817 | -0.218737 | 0.700999 | 0.42483 | 0.189435 | 0.086084 | -0.195612 | -0.182643 | 0.218578 | 0.058692 | 0.225776 | -2.475348 | 0.151432 | -0.046189 | 0.382015 |
| 35.2014 | 0.990648 | 0.49659 | 0.560501 | 0.723647 | 1.333824 | -0.646685 | 0.893386 | -0.868025 | -1.029915 | -0.170109 | 0.046692 | 0.246165 | -0.065778 | -0.091121 | -0.060919 | -0.677951 | 0.25436 | 0.046308 | 0.077503 | -1.90697 | 0.072466 | -0.032564 | 0.227951 |
| 35.2153 | 0.5417 | 0.228383 | 0.155156 | -0.438426 | 0.36239 | -0.760869 | 0.61423 | -0.956592 | -1.075081 | -0.011367 | -0.362331 | 0.16498 | -0.183292 | -0.302412 | -0.009267 | -0.866635 | 0.465806 | 0.206427 | 0.033834 | -1.444134 | 0.107093 | 0.057424 | 0.062697 |
| 35.2292 | 0.367257 | 0.047403 | 0.041227 | -1.053072 | -0.587833 | -0.832693 | 0.670168 | -0.973593 | -0.906876 | 0.010464 | -0.602017 | 0.16237 | -0.08091 | -0.158148 | 0.016732 | -0.722141 | 0.685906 | 0.365314 | 0.056373 | -1.20346 | 0.192208 | 0.025347 | 0.333722 |
| 35.2431 | 0.472119 | 0.078797 | 0.047291 | -0.814149 | -1.381002 | -0.688077 | 0.804222 | -0.605165 | -0.569701 | 0.048044 | -0.409645 | 0.134884 | 0.010028 | 0.130621 | 0.014202 | -0.495055 | 0.682614 | 0.379001 | 0.072866 | -0.975892 | 0.122668 | -0.068763 | 0.516638 |
| 35.2569 | 0.75601 | 0.190284 | 0.392524 | -0.880271 | -1.570595 | -0.419382 | 0.600429 | -0.29003 | -0.393069 | 0.046613 | -0.127827 | 0.095449 | -0.098902 | 0.261789 | 0.015523 | -0.251342 | 0.507901 | 0.222681 | 0.07271 | -0.725033 | 0.00796 | -0.090936 | 0.285805 |
| 35.2708 | 0.791246 | -0.024572 | 0.624087 | -0.919033 | -1.632779 | -0.360044 | 0.496651 | -0.352768 | -0.692412 | -0.168694 | -0.380713 | 0.122252 | -0.192526 | 0.296705 | 0.032818 | -0.036163 | 0.320938 | 0.085388 | 0.114978 | -0.362573 | -0.010285 | -0.041361 | 0.032527 |
| 35.2847 | 0.620827 | -0.442293 | 0.643214 | -0.768531 | -1.891793 | -0.454244 | 0.363301 | -0.476185 | -1.259663 | -0.374835 | -1.17077 | 0.153282 | -0.200285 | 0.342274 | 0.022661 | 0.037763 | 0.149015 | 0.009498 | 0.190978 | -0.024674 | -0.042947 | -0.025718 | 0.010482 |
| 35.2986 | 0.587682 | -0.450326 | 0.692247 | -0.450083 | -1.745126 | -0.435284 | 0.452025 | -0.275367 | -1.619463 | -0.299054 | -1.046222 | 0.068678 | -0.265104 | 0.318192 | -0.065184 | 0.020805 | 0.02618 | -0.043473 | 0.242877 | 0.118782 | -0.164054 | -0.078578 | 0.020264 |
| 35.3125 | 0.714339 | -0.267253 | 0.765746 | -0.129177 | -1.102488 | -0.427731 | 0.318385 | 0.073592 | -1.455181 | -0.032994 | -0.866945 | -0.06885 | -0.333399 | 0.258572 | -0.15942 | 0.047718 | 0.029415 | -0.019348 | 0.276988 | 0.147466 | -0.207472 | -0.125763 | 0.085165 |
| 35.3264 | 0.671568 | -0.317721 | 0.769623 | 0.163094 | -0.634559 | -0.423478 | 0.179461 | 0.220037 | -1.698127 | 0.01734 | -1.361167 | -0.040376 | -0.186776 | 0.325278 | -0.072418 | 0.206333 | 0.167987 | 0.133701 | 0.405073 | 0.25833 | -0.063993 | 0.187905 | 0.397888 |
| 35.3403 | 0.57415 | -0.408524 | 0.759801 | 0.443347 | -0.193146 | -0.274294 | 0.302096 | 0.238252 | -1.961933 | -0.043524 | -1.393386 | 0.173153 | 0.020336 | 0.440815 | 0.037865 | 0.37596 | 0.230097 | 0.244381 | 0.542553 | 0.407651 | 0.055276 | 0.558263 | 0.606228 |
| 35.3542 | 0.509886 | -0.412872 | 0.694857 | 0.725466 | 0.184396 | -0.18089 | 0.541735 | 0.147903 | -2.297082 | -0.098682 | -1.17586 | 0.309261 | -0.009495 | 0.416318 | -0.00668 | 0.420556 | 0.070276 | 0.161167 | 0.516741 | 0.425692 | -0.03031 | -0.044625 | 0.397539 |
| 35.3681 | 0.462594 | -0.419764 | 0.538111 | 0.941234 | 0.29925 | -0.232372 | 0.742425 | -0.071676 | -2.999039 | -0.190548 | -1.336504 | 0.231263 | -0.147084 | 0.284203 | -0.06171 | 0.378126 | -0.108125 | 0.013223 | 0.402022 | 0.320454 | -0.103225 | -0.345662 | 0.171813 |
| 35.3819 | 0.373061 | -0.415761 | 0.407491 | 0.903011 | 0.088416 | -0.284106 | 0.824006 | -0.36468 | -3.135156 | -0.24561 | -1.378513 | 0.141851 | -0.143226 | 0.224066 | 0.044595 | 0.383105 | -0.120064 | -0.020096 | 0.346527 | 0.217127 | -0.01343 | 0.009003 | 0.22862 |
| 35.3958 | 0.41738 | -0.248633 | 0.449832 | 0.689014 | -0.256783 | -0.479775 | 0.847695 | -0.509276 | -2.433556 | -0.077184 | -0.500408 | 0.252395 | 0.016254 | 0.307305 | 0.185909 | 0.414946 | -0.032716 | 0.031573 | 0.377726 | 0.257446 | 0.090091 | 0.716197 | 0.33657 |
| 35.4097 | 1.597506 | 1.666516 | 1.388141 | 0.096713 | -0.440346 | -0.307745 | 1.28957 | -0.366094 | -1.329689 | 0.834835 | 6.697633 | 0.418353 | 0.10193 | 0.408585 | 0.265885 | 0.395477 | 0.056064 | 0.059659 | 0.45941 | 0.304155 | 0.027534 | 0.421024 | 0.310475 |
| 35.4236 | 4.04733 | 4.957419 | 2.169326 | -0.528919 | -0.154897 | 0.422124 | 1.963921 | -0.630952 | 0.360374 | 1.409057 | 19.941489 | 0.369675 | 0.199915 | 0.422497 | 0.420323 | 0.336593 | 0.237634 | 0.063744 | 0.546284 | 0.212488 | -0.004137 | -0.556986 | 0.765873 |
| 35.4375 | 5.469727 | 5.69452 | 1.054139 | 0.006011 | 0.140689 | 1.297899 | 3.698799 | -0.929814 | 2.263248 | -1.3696 | 23.785753 | 0.123689 | 1.639325 | 0.507505 | 2.034318 | 0.7786 | 1.09564 | 0.200694 | 1.327184 | 0.599408 | 1.081156 | -0.934089 | 6.148671 |
| 35.4514 | 3.554187 | 3.757982 | 1.550927 | 1.806648 | 1.364488 | 3.320329 | 4.923533 | 0.581469 | 2.857564 | 0.397002 | 10.120124 | -0.072673 | 4.289287 | 1.076126 | 5.465483 | 1.811937 | 3.214486 | 0.430572 | 3.17345 | 1.775152 | 3.470114 | 0.715744 | 16.232416 |
| 35.4653 | 2.329053 | 2.113544 | 1.334423 | 2.650727 | 2.127747 | 2.348317 | 3.630058 | 0.491102 | 2.743766 | -0.291159 | 2.074983 | 0.712713 | 4.395438 | 2.951792 | 6.41857 | 2.239737 | 4.343661 | 0.243648 | 4.627044 | 2.522022 | 3.676254 | 1.054184 | 19.6845 |
| 35.4792 | 2.371186 | 2.028547 | 1.897389 | 2.903324 | 2.754693 | 1.954492 | 2.7181 | 0.169257 | 3.006991 | 0.798483 | 1.608709 | 2.506327 | 2.899376 | 3.552201 | 3.31668 | 1.677107 | 2.206321 | 0.926803 | 3.008758 | 2.083962 | 2.250964 | 1.035198 | 9.211278 |
| 35.4931 | 1.984343 | 2.201154 | 1.567262 | 2.52737 | 2.422138 | 2.046972 | 2.116979 | 0.331278 | 3.348266 | 1.369818 | 2.294162 | 1.791542 | 1.391678 | 2.300195 | 0.890777 | 2.219916 | 0.619053 | 0.750839 | 1.47711 | 1.847083 | 1.201716 | 2.947523 | 2.077131 |
| 35.5 | 1.266411 | 1.836771 | 1.317732 | 2.153315 | 1.916946 | 2.06422 | 1.758989 | 0.538601 | 3.314355 | 0.680214 | 2.257466 | 1.1091 | 0.645863 | 1.338061 | 0.674232 | 2.069932 | 0.312143 | 0.511649 | 1.004667 | 1.72483 | 0.754008 | 4.432464 | 0.855167 |
| 35.5139 | 0.508948 | 0.956093 | 1.06809 | 1.554708 | 1.129635 | 1.818086 | 1.261053 | 0.545328 | 3.162897 | -0.213213 | 1.742918 | 0.742802 | 0.389514 | 0.966497 | 0.438355 | 1.334067 | 0.172588 | 0.155873 | 0.711407 | 1.17265 | 0.440733 | 3.06311 | 0.604013 |
| 35.5278 | -0.014207 | 0.239573 | 0.914988 | 1.147985 | 0.581716 | 1.326942 | 0.900923 | 0.29215 | 2.822876 | -0.92003 | 1.362087 | 0.423614 | 0.248389 | 0.799843 | 0.211603 | 0.174832 | -0.048552 | -0.353164 | 0.437854 | 0.355023 | 0.096357 | 0.501828 | 0.165509 |
| 35.5417 | -0.271241 | -0.090048 | 0.696243 | 0.998467 | 0.306895 | 0.992163 | 0.88138 | 0.055565 | 1.825058 | -0.772595 | 0.953908 | 0.213212 | 0.241032 | 0.68637 | -0.036539 | -1.016731 | -0.269551 | -0.741577 | 0.277157 | -0.395313 | -0.273575 | -4.553674 | -0.349918 |
| 35.5556 | -0.762093 | -0.159044 | 0.230154 | 0.848172 | 0.201046 | 0.832418 | 1.071847 | -0.33544 | 1.538191 | -0.314489 | 0.604401 | 0.338048 | 0.299274 | 0.682044 | -0.140453 | -1.7146 | -0.22115 | -0.784956 | 0.504259 | -1.118531 | -0.441751 | -3.72948 | -0.443736 |
| 35.5694 | -0.985011 | -0.4849 | 0.547824 | 0.406438 | -0.164177 | 0.55686 | 0.970677 | -0.537834 | 1.028701 | -0.494712 | 0.152219 | 0.545999 | 0.242583 | 0.588966 | -0.082237 | -1.261394 | -0.125327 | -0.427571 | 0.355578 | -1.367635 | -0.49026 | 0.445282 | -0.330202 |
| 35.5833 | -1.044665 | -0.930414 | 0.665928 | -0.571491 | -0.63382 | 0.217467 | 0.92279 | -0.637597 | 0.490586 | -1.402586 | -0.42801 | 0.596951 | 0.096644 | 0.325335 | 0.037561 | -0.493898 | 0.064256 | -0.062508 | 0.56575 | -1.37539 | -0.208598 | 5.302679 | -0.157661 |
| 35.5972 | -1.034991 | -1.308661 | 0.572999 | -1.061655 | -1.406546 | -0.084148 | 0.723817 | -0.50725 | 0.344583 | -2.108728 | -1.354933 | 0.557858 | -0.013272 | 0.178092 | 0.155363 | -0.069833 | 0.016089 | 0.116348 | 0.717354 | -1.490735 | 0.09886 | 8.291204 | -0.107524 |
| 35.6111 | -1.061758 | -1.672486 | 0.282723 | -1.105055 | -2.226366 | -0.66919 | 0.685933 | -0.276181 | 0.058314 | -2.917998 | -1.958295 | 0.415364 | -0.009659 | 0.102133 | 0.184636 | -0.002053 | -0.006604 | 0.049162 | 0.695216 | -1.622523 | 0.249969 | 10.879312 | -0.22295 |
| 35.625 | -1.092505 | -1.973869 | 0.120839 | -1.000664 | -3.115067 | -1.108575 | 0.346914 | -0.489828 | -0.513062 | -3.640289 | -2.252185 | 0.134783 | 0.034092 | 0.000332 | 0.104428 | -0.155705 | -0.012649 | -0.132021 | 0.486023 | -1.929875 | 0.179944 | 7.928488 | -0.422448 |
| 35.6389 | -1.04198 | -2.079198 | -0.015676 | -0.674791 | -3.43892 | -1.844534 | 0.486402 | -0.728313 | -0.80665 | -3.330318 | -2.393954 | -0.192683 | 0.142848 | -0.133026 | -0.164787 | -0.453899 | -0.093908 | -0.242182 | 0.033429 | -1.980107 | -0.080042 | 4.944797 | -0.707026 |
| 35.6528 | -0.802463 | -2.038527 | -0.053192 | -0.243138 | -2.70155 | -1.706678 | 0.746798 | -0.457313 | -1.148506 | -2.927299 | -2.30512 | -0.480908 | 0.198781 | -0.25991 | -0.394862 | -0.606135 | -0.213113 | -0.249192 | -0.392658 | -1.838326 | -0.355157 | 2.309655 | -1.078546 |
| 35.6667 | -0.519561 | -1.848429 | -0.138569 | 0.172072 | -1.75742 | -1.469733 | 0.999144 | 0.011294 | -1.597929 | -2.191888 | -2.249439 | -0.589366 | 0.200264 | -0.517451 | -0.478185 | -0.491958 | -0.318989 | -0.110478 | -0.583878 | -1.506078 | -0.480639 | -1.94985 | -1.126432 |
| 35.6806 | -0.344746 | -1.59227 | -0.266603 | 0.533787 | -0.821954 | -1.15256 | 1.155209 | 0.31754 | -1.856293 | -1.668744 | -2.097065 | -0.519552 | 0.155018 | -0.818953 | -0.466969 | -0.14938 | -0.396438 | 0.097283 | -0.285232 | -0.799195 | -0.484633 | -4.977985 | -0.990629 |
| 35.6944 | -0.271944 | -1.374547 | -0.435223 | 0.855913 | -0.038372 | -0.832599 | 1.224795 | 0.480727 | -1.910067 | -1.24546 | -1.780924 | -0.344314 | 0.11875 | -0.966919 | -0.249799 | 0.307374 | -0.475976 | 0.311802 | 0.093039 | -0.036719 | -0.327479 | -6.434217 | -0.705714 |
| 35.7083 | -0.275098 | -1.254388 | 0.036867 | 1.069957 | 0.581338 | -0.609493 | 1.195764 | 0.515266 | -1.798605 | -1.27647 | -1.36908 | -0.172764 | 0.065235 | -0.803203 | 0.02599 | 0.664757 | -0.48718 | 0.378882 | 0.259411 | 0.652693 | -0.130835 | -3.222643 | -0.395838 |
| 35.7222 | -0.414527 | -1.386233 | 0.771772 | 1.077139 | 0.682805 | -0.553966 | 1.04812 | 0.541497 | -1.805534 | -1.567363 | -1.207989 | -0.182874 | 0.029475 | -0.566341 | 0.272965 | 1.050385 | -0.43658 | 0.377772 | 0.167015 | 1.273735 | -0.04832 | 2.786717 | -0.256837 |
| 35.7361 | -0.573285 | -1.539075 | 1.046423 | 0.925745 | 0.050186 | -0.962157 | 0.643104 | 0.763116 | -1.704342 | -1.929882 | -1.222866 | -0.289193 | 0.043813 | -0.494467 | 0.451468 | 1.278733 | -0.437448 | 0.237357 | -0.048244 | 1.503323 | -0.053305 | 8.595325 | -0.279238 |
| 35.75 | -0.774597 | -1.621526 | 0.614336 | 0.643633 | -1.030458 | -1.808504 | -0.013784 | 0.254661 | -1.962775 | -2.298689 | -1.48133 | -0.439156 | 0.044549 | -0.404525 | 0.555503 | 1.328152 | -0.436626 | 0.101275 | -0.120961 | 1.40035 | -0.065941 | 10.10127 | -0.351094 |
| 35.7639 | -0.886697 | -1.567244 | -0.233487 | 0.342522 | -1.901107 | -2.550978 | -0.51043 | -0.626421 | -2.545518 | -2.168349 | -1.630948 | -0.533932 | 0.064936 | -0.289925 | 0.437829 | 1.082013 | -0.472801 | 0.027091 | -0.168126 | 0.916826 | -0.066737 | 9.595707 | -0.441607 |
| 35.7778 | -1.019101 | -1.240537 | -1.346906 | 0.058391 | -2.529781 | -2.585244 | -0.770923 | -1.458104 | -3.32105 | -1.520524 | -1.741084 | -0.51757 | 0.073823 | -0.163654 | 0.192192 | 0.636046 | -0.37303 | 0.068151 | -0.273916 | 0.096302 | -0.01192 | 3.266538 | -0.451501 |
| 35.7917 | -1.172284 | -0.939065 | -1.925155 | 0.012844 | -3.079815 | -2.362294 | -1.014666 | -1.683951 | -4.373763 | -0.857924 | -2.033388 | -0.410729 | 0.093769 | -0.093114 | 0.068915 | 0.211424 | -0.259697 | 0.107647 | -0.432651 | -0.670875 | 0.05394 | -1.643585 | -0.500307 |
| 35.8056 | -1.272396 | -0.44943 | -2.10291 | -0.144694 | -3.140116 | -2.504893 | -1.214167 | -1.791967 | -4.981229 | -0.501777 | -2.617754 | -0.313625 | 0.04754 | -0.086239 | 0.051623 | -0.091401 | -0.150185 | 0.176028 | -0.495772 | -0.816816 | 0.116166 | -4.095661 | -0.573262 |
| 35.8194 | -1.270978 | 0.141263 | -1.820181 | -0.359896 | -2.869884 | -2.20343 | -1.326779 | -1.611591 | -4.305671 | -0.335442 | -2.725859 | -0.357429 | -0.070594 | -0.14535 | 0.082354 | -0.140974 | -0.051267 | 0.200317 | -0.478961 | -0.604251 | 0.175943 | -5.602434 | -0.663568 |
| 35.8333 | -1.404292 | 0.664989 | -1.901437 | -0.493667 | -3.272227 | -1.958663 | -1.223234 | -1.580564 | -3.2411 | -0.295044 | -2.785872 | -0.521393 | -0.190498 | -0.177911 | 0.133865 | -0.212941 | -0.021839 | 0.15933 | -0.473185 | -0.354957 | 0.303741 | -6.422074 | -0.673228 |
| 35.8472 | -1.222378 | 1.110246 | -1.709256 | -0.36302 | -3.515272 | -2.189921 | -0.957154 | -1.545294 | -2.284918 | -0.265059 | -2.437595 | -0.679024 | -0.344951 | -0.220937 | 0.155942 | -0.258476 | -0.044526 | 0.01147 | -0.42177 | -0.10879 | 0.464472 | 2.450719 | -0.590363 |
| 35.8611 | -0.74211 | 1.362622 | -1.354914 | -0.090491 | -2.829686 | -1.941676 | -0.611555 | -1.388103 | -1.581833 | -0.194023 | -1.817087 | -0.712459 | -0.514588 | -0.235055 | 0.136487 | -0.186127 | -0.085512 | -0.082095 | -0.380762 | 0.228359 | 0.552913 | 8.419223 | -0.541487 |
| 35.875 | -0.024462 | 1.495451 | -0.762803 | 0.110085 | -1.731577 | -1.222553 | -0.237799 | -0.611087 | -1.196996 | -0.07373 | -1.153709 | -0.684115 | -0.694034 | -0.202829 | 0.071262 | -0.145817 | -0.079749 | -0.018562 | -0.36237 | 0.59752 | 0.465093 | 6.892516 | -0.502382 |
| 35.8889 | 0.604135 | 1.565301 | -0.085799 | 0.145301 | -0.412879 | -0.520635 | -0.173947 | 0.505714 | -0.930775 | 0.061731 | -2.051208 | -0.577847 | -0.812972 | -0.150529 | 0.022687 | 0.005557 | -0.009321 | 0.144871 | -0.302459 | 0.865043 | 0.28501 | 12.287592 | -0.459575 |
| 35.9028 | -0.292551 | 0.353842 | 0.272919 | -0.368102 | 0.189692 | -0.435954 | -1.730181 | 1.546551 | -1.648261 | -0.446448 | -11.130126 | -0.427823 | -0.798465 | -0.076693 | 0.048264 | 0.213877 | 0.106901 | 0.319862 | -0.184295 | 1.113926 | 0.200023 | 9.393119 | -0.311837 |
| 35.9167 | -3.005717 | -3.984617 | -0.636065 | -1.641497 | -1.579812 | -1.900109 | -4.716266 | 1.421214 | -5.182344 | -0.890115 | -25.654492 | -0.323992 | -0.949374 | -0.052902 | 0.054263 | 0.376208 | 0.087431 | 0.411084 | -0.145706 | 1.381234 | 0.275317 | 1.112805 | -1.090147 |
| 35.9306 | -5.318892 | -8.143591 | -2.668817 | -2.05456 | -4.373431 | -3.865042 | -5.440134 | -1.095564 | -8.296648 | -1.113751 | -25.508981 | -0.335473 | -2.298765 | -0.147617 | -1.015368 | 0.336546 | -0.277515 | 0.468433 | -0.905447 | 0.777088 | 0.136116 | -6.368315 | -7.149447 |
| 35.9444 | -5.112944 | -5.63039 | -2.407674 | -1.084656 | -5.469473 | -4.267728 | -3.334625 | -4.708707 | -7.175763 | -0.927515 | -12.68933 | -1.322732 | -5.668834 | -1.764292 | -4.741727 | -1.186305 | -1.46735 | -0.399323 | -3.894473 | -2.463734 | -1.522011 | -8.215284 | -17.312724 |
| 35.9583 | -2.615951 | -2.114639 | -1.032701 | -0.114865 | -3.95224 | -2.954807 | -1.431295 | -6.281677 | -4.899505 | -0.338803 | -4.565597 | -2.861267 | -7.246353 | -3.542637 | -6.899104 | -2.924478 | -1.916439 | -1.173405 | -4.885363 | -6.446543 | -2.722828 | -5.157104 | -17.139496 |
| 35.9722 | -1.065832 | -1.101405 | -0.427873 | 0.470707 | -3.020937 | -1.9807 | -0.525011 | -5.429275 | -3.542412 | 0.044885 | -2.453558 | -2.497764 | -4.87109 | -1.56222 | -4.700097 | -1.774633 | -1.49775 | -1.193968 | -2.413253 | -5.194769 | -2.321336 | -8.175881 | -9.282055 |
| 35.9861 | -0.466143 | -0.987185 | -0.173077 | 0.498787 | -2.774601 | -1.379674 | -0.128314 | -3.0103 | -2.830941 | -0.032339 | -1.656049 | 0.226658 | -0.261475 | 1.032258 | -1.051352 | 0.673071 | 0.133353 | 0.466541 | 1.361333 | -1.623202 | -0.031213 | -8.786605 | -0.871627 |
| 36.0069 | -0.171364 | -1.049756 | 0.01766 | 0.349954 | -2.118152 | -0.990512 | 0.028513 | -0.777851 | -1.529555 | -0.245072 | -0.982308 | 1.309952 | 1.231001 | 2.097905 | 1.041832 | 1.829494 | 1.463252 | 1.448592 | 2.763023 | 0.683561 | 1.572981 | -6.737871 | 2.08976 |
| 36.0208 | 0.063796 | -1.036352 | 0.106387 | 0.30404 | -1.643148 | -0.834556 | 0.07071 | 0.959109 | 0.009448 | -0.100134 | -0.536283 | 0.644796 | 0.626635 | 1.363214 | 1.954958 | 1.84913 | 1.331381 | 1.221332 | 1.795252 | 1.415384 | 1.505695 | -5.064295 | 2.188711 |
| 36.0347 | 0.227554 | -0.979567 | 0.209214 | 0.340879 | -1.479001 | -0.71013 | 0.114182 | 2.480444 | 1.33919 | 0.081111 | -0.262965 | 0.302077 | 0.246131 | 0.849706 | 2.713878 | 1.466689 | 1.173172 | 0.685592 | 0.950856 | 1.770937 | 1.171304 | -2.240617 | 1.938638 |
| 36.0486 | 0.383624 | -1.066835 | 0.351665 | 0.389367 | -0.839836 | -0.487235 | 0.230374 | 3.869347 | 1.967152 | -0.405837 | 0.25832 | 0.535434 | 0.286408 | 0.856441 | 2.813551 | 1.050051 | 1.151847 | 0.547539 | 0.51108 | 2.46637 | 1.174401 | 3.722616 | 1.827269 |
| 36.0625 | 0.60075 | -1.273804 | 0.529757 | 0.497021 | 0.056743 | -0.236891 | 0.360123 | 5.118152 | 1.940827 | -1.155134 | 0.612453 | 0.570557 | 0.443512 | 0.733385 | 2.206143 | 0.493438 | 0.836805 | 0.370355 | 0.248485 | 2.557956 | 0.828093 | 7.129816 | 1.483889 |
| 36.0764 | 0.949104 | -1.240868 | 0.707789 | 0.619872 | 1.026945 | 0.033904 | 0.475315 | 6.267906 | 1.877532 | -1.537544 | 1.129362 | 0.224575 | 0.412638 | 0.483454 | 1.498547 | 0.132022 | 0.435568 | 0.167032 | 0.093415 | 2.242526 | 0.365172 | 10.236296 | 1.149965 |
| 36.0903 | 1.490381 | -0.856053 | 0.901488 | 0.729833 | 1.929286 | 0.413264 | 0.597389 | 5.97577 | 1.952023 | -1.227096 | 2.076801 | -0.156141 | 0.107314 | 0.27323 | 0.969832 | 0.019312 | 0.146864 | -0.000484 | 0.018184 | 2.590969 | -0.084377 | 11.078927 | 0.703934 |
| 36.1042 | 1.964037 | -0.291603 | 1.100087 | 0.884678 | 2.762142 | 0.962852 | 0.640117 | 5.537392 | 1.71053 | -0.35646 | 2.611033 | 0.001896 | 0.011431 | -0.019749 | 0.495086 | -0.016384 | -0.039485 | -0.050467 | -0.218495 | 2.740327 | -0.429318 | 8.383986 | 0.215181 |
| 36.1181 | 2.263434 | 0.165218 | 1.39003 | 1.013991 | 3.752224 | 1.427295 | 0.533889 | 5.251575 | 1.601536 | 0.019413 | 2.835117 | 0.297098 | 0.177728 | -0.413421 | 0.218872 | 0.054111 | -0.100185 | -0.010919 | -0.543545 | 1.994397 | -0.470515 | 4.315008 | -0.2293 |
| 36.1319 | 2.323279 | 0.600882 | 1.671601 | 1.090773 | 4.98498 | 2.232282 | 0.46694 | 4.204025 | 1.452553 | 0.078402 | 3.237173 | 0.514285 | 0.324641 | -0.334771 | 0.264424 | 0.280186 | -0.028468 | 0.075049 | -0.442409 | 1.603877 | -0.151928 | 1.742653 | -0.314071 |
| 36.1458 | 2.215358 | 0.912754 | 1.768666 | 1.070222 | 5.304168 | 2.214227 | 0.507969 | 3.552166 | 1.554494 | 0.124109 | 3.942988 | 0.556106 | 0.396557 | 0.072648 | 0.336082 | 0.521521 | 0.103178 | 0.133471 | -0.027798 | 1.348634 | 0.217899 | -0.142066 | -0.008054 |
| 36.1597 | 2.158642 | 1.069743 | 1.687158 | 1.035384 | 5.014682 | 1.800337 | 0.544934 | 3.61049 | 1.996188 | 0.101102 | 3.789894 | 0.575483 | 0.500749 | 0.324446 | 0.334553 | 0.602197 | 0.195467 | 0.138308 | 0.265791 | 0.751107 | 0.337408 | -0.447139 | 0.439186 |
| 36.1736 | 2.112734 | 1.110016 | 1.457465 | 1.003617 | 4.592551 | 0.928005 | 0.580578 | 3.214235 | 2.086983 | 0.171681 | 3.788038 | 0.576513 | 0.528447 | 0.36335 | 0.19461 | 0.484615 | 0.172951 | 0.049438 | 0.327288 | -0.078528 | 0.174489 | 1.053585 | 0.546567 |
| 36.1875 | 2.061857 | 1.050088 | 1.261839 | 1.046849 | 4.539167 | 0.788784 | 0.670453 | 2.678734 | 1.841045 | 0.094452 | 3.833471 | 0.497242 | 0.416503 | 0.284604 | -0.152851 | 0.288306 | 0.092494 | 0.002861 | 0.200116 | -0.953629 | -0.064939 | 3.802719 | 0.069786 |
| 36.2014 | 1.997056 | 0.946822 | 1.088705 | 1.112907 | 4.384943 | 0.972101 | 0.724788 | 2.420356 | 1.58178 | 0.070792 | 3.291779 | 0.500716 | 0.428393 | 0.245441 | -0.406419 | 0.277692 | 0.101486 | 0.141178 | 0.14918 | -1.513662 | -0.172847 | 6.198671 | -0.191537 |
| 36.2153 | 2.080199 | 0.913138 | 0.974728 | 1.207542 | 3.725311 | 0.868384 | 0.756864 | 2.105662 | 1.347659 | 0.066353 | 2.607885 | 0.644009 | 0.616437 | 0.19628 | -0.530093 | 0.493368 | 0.151628 | 0.346089 | 0.224392 | -1.582436 | -0.097519 | 4.708348 | 0.05899 |
| 36.2292 | 1.890004 | 0.961382 | 0.940858 | 1.297788 | 2.767391 | 0.410138 | 0.775888 | 1.230597 | 0.655113 | 0.050889 | 1.95198 | 0.751958 | 0.630484 | 0.044313 | -0.691008 | 0.710241 | 0.153034 | 0.408947 | 0.201019 | -1.585989 | 0.084886 | 2.777306 | 0.14595 |
| 36.2431 | 1.693563 | 1.098379 | 0.980194 | 1.338667 | 1.702831 | -0.054147 | 0.829085 | 0.236237 | 0.129397 | 0.017877 | 1.310832 | 0.702431 | 0.518521 | -0.049877 | -0.610972 | 0.738526 | 0.16787 | 0.376316 | 0.090296 | -1.666154 | 0.302543 | 1.53951 | 0.078751 |
| 36.2569 | 1.565402 | 1.292663 | 1.037548 | 1.296585 | 0.735805 | -0.253387 | 0.928763 | -0.491551 | -0.089185 | 0.003644 | 0.872684 | 0.550567 | 0.460782 | -0.001718 | -0.184008 | 0.638503 | 0.254941 | 0.34926 | 0.09468 | -1.491021 | 0.504987 | 1.680525 | 0.285253 |
| 36.2708 | 1.36574 | 1.369494 | 1.040674 | 1.21399 | -0.057157 | -0.327258 | 1.01014 | -1.189327 | -0.296922 | -0.056202 | -0.078622 | 0.47274 | 0.430779 | 0.164935 | 0.248453 | 0.514602 | 0.335326 | 0.332029 | 0.197492 | -1.019131 | 0.543488 | 2.494526 | 0.473959 |
| 36.2847 | 1.131532 | 1.395787 | 0.99779 | 1.181457 | -0.612817 | -0.425312 | 1.071966 | -1.82511 | -0.457083 | -0.117897 | -0.835385 | 0.489873 | 0.394827 | 0.316438 | 0.40691 | 0.408409 | 0.380486 | 0.315602 | 0.316027 | -0.49101 | 0.452093 | 2.676899 | 0.470262 |
| 36.2986 | 0.996911 | 1.385377 | 1.0539 | 1.148522 | -1.000932 | -0.681188 | 1.159623 | -2.162483 | -0.429195 | -0.093378 | -0.473114 | 0.516287 | 0.371832 | 0.42219 | 0.377909 | 0.365407 | 0.360107 | 0.266362 | 0.401877 | -0.069443 | 0.146648 | 1.539969 | 0.427782 |
| 36.3125 | 0.931328 | 1.215503 | 1.122012 | 1.079559 | -1.005539 | -0.915429 | 1.150471 | -1.953324 | -0.396252 | -0.044633 | 0.117867 | 0.475132 | 0.276493 | 0.437467 | 0.219717 | 0.336149 | 0.238548 | 0.156521 | 0.369322 | 0.181378 | -0.205274 | 1.25141 | 0.206792 |
| 36.3264 | 0.90873 | 0.960505 | 1.128625 | 1.016304 | -0.944904 | -0.951983 | 1.070822 | -1.56759 | -0.346213 | -0.065811 | 0.024816 | 0.445049 | 0.142479 | 0.393886 | 0.074372 | 0.326912 | 0.1552 | 0.046155 | 0.270913 | 0.326623 | -0.324829 | 2.589044 | -0.058858 |
| 36.3403 | 0.980579 | 0.933482 | 1.110545 | 1.058467 | -1.125197 | -0.856743 | 1.064712 | -1.491608 | -0.500897 | -0.109848 | -0.399111 | 0.50524 | 0.158035 | 0.3842 | 0.114226 | 0.392951 | 0.172266 | 0.042179 | 0.212134 | 0.51845 | -0.126096 | 5.789741 | -0.087738 |
| 36.3542 | 1.093104 | 0.90755 | 1.037226 | 1.112368 | -1.528403 | -0.853719 | 1.104452 | -1.77254 | -0.810255 | -0.235311 | -0.767193 | 0.607172 | 0.282765 | 0.433364 | 0.314773 | 0.495599 | 0.207313 | 0.102155 | 0.25239 | 0.811641 | 0.163823 | 4.608581 | 0.141235 |
| 36.3681 | 0.953494 | 0.644696 | 0.949768 | 1.067228 | -1.845667 | -0.997935 | 1.110046 | -1.988654 | -1.117796 | -0.359898 | -0.929248 | 0.681912 | 0.436305 | 0.537242 | 0.531056 | 0.53086 | 0.175539 | 0.103667 | 0.321265 | 1.114362 | 0.268149 | 2.040825 | 0.40817 |
| 36.3819 | 0.500219 | 0.249266 | 0.89824 | 1.013786 | -1.913882 | -1.133871 | 1.027668 | -2.024763 | -1.284848 | -0.437044 | -1.220522 | 0.687554 | 0.481667 | 0.600259 | 0.595137 | 0.441138 | 0.074416 | 0.034235 | 0.35976 | 1.326168 | 0.106131 | -0.890775 | 0.524482 |
| 36.3958 | 0.553801 | 0.386378 | 1.005546 | 1.023175 | -1.749534 | -1.174356 | 1.091773 | -1.941829 | -1.070828 | -0.374195 | -0.174707 | 0.49645 | 0.302348 | 0.498239 | 0.426511 | 0.280496 | -0.046951 | -0.031302 | 0.29684 | 1.249199 | -0.144043 | -2.598229 | 0.461873 |
| 36.4097 | 1.403025 | 2.167789 | 1.907553 | 1.184379 | -0.721547 | -1.126659 | 1.987459 | -1.575588 | -0.160522 | 0.027571 | 9.073669 | 0.260944 | 0.090139 | 0.369073 | 0.158355 | 0.199243 | -0.11112 | -0.010183 | 0.236057 | 0.993929 | -0.293592 | -2.901615 | 0.287602 |
| 36.4236 | 3.143656 | 5.739313 | 3.655559 | 1.728462 | 0.437915 | -0.831547 | 3.667536 | -0.995684 | 1.080509 | 0.82119 | 27.851782 | 0.348704 | 0.305592 | 0.335914 | 0.087392 | 0.314706 | -0.11163 | 0.081722 | 0.384586 | 0.96921 | -0.126535 | -2.539647 | 0.91587 |
| 36.4375 | 4.576068 | 8.216363 | 3.260317 | 2.904949 | 2.612438 | 0.474499 | 5.013772 | -1.336987 | 2.178671 | 2.022715 | 31.207182 | 1.018607 | 2.171982 | 0.237479 | 1.052121 | 0.769659 | -0.39471 | 0.101977 | 1.433771 | 1.818046 | 0.522429 | -2.256008 | 5.78935 |
| 36.4514 | 4.880778 | 6.368798 | 2.186514 | 2.744871 | 3.381463 | 1.391277 | 4.260299 | -0.115506 | 2.251761 | 1.995517 | 13.759442 | 2.189306 | 5.437957 | 0.166952 | 3.181517 | 1.005525 | -0.937063 | 0.218564 | 3.838409 | 3.592494 | 1.58926 | -1.855659 | 14.647488 |
| 36.4653 | 3.084446 | 3.970214 | 1.456707 | 1.401721 | 2.956423 | 1.114074 | 2.320814 | 1.242989 | 2.263923 | 0.68723 | 3.352107 | 2.838508 | 5.276768 | 1.532126 | 3.687386 | 0.997248 | -0.82717 | 0.92934 | 5.74549 | 4.538574 | 1.593761 | -0.490666 | 17.75193 |
| 36.4792 | 1.619255 | 2.169008 | 0.859707 | 0.771951 | 3.714753 | 0.773039 | 1.622227 | 1.09343 | 1.308256 | 0.24549 | 2.632545 | 3.460628 | 2.987354 | 2.789728 | 2.142512 | 1.387252 | 0.156804 | 1.180859 | 4.178478 | 3.491718 | 1.786012 | 2.731664 | 9.766939 |
| 36.4931 | 1.508816 | 1.507085 | 0.578345 | 0.75542 | 3.712789 | 0.81043 | 1.33029 | 1.0474 | 1.241774 | 0.112828 | 3.247707 | 2.231269 | 1.223028 | 1.822534 | 1.316217 | 1.779521 | 0.892735 | 0.764665 | 1.75807 | 2.168424 | 2.052539 | 5.554884 | 2.659524 |
| 36.5 | 1.874857 | 1.565283 | 0.487072 | 1.225283 | 3.805011 | 0.923302 | 1.259745 | 1.326943 | 1.65213 | 0.043702 | 3.591701 | 1.162529 | 0.606984 | 1.020574 | 1.309633 | 1.565477 | 0.901802 | 0.603425 | 0.835344 | 1.733814 | 1.703216 | 4.407546 | 1.159379 |
| 36.5139 | 2.234266 | 1.747949 | 0.538022 | 1.999187 | 4.285488 | 1.35442 | 1.692679 | 0.899489 | 1.498207 | 0.017641 | 4.01695 | 0.584886 | 0.719831 | 0.691862 | 1.119765 | 1.155501 | 0.706492 | 0.398026 | 0.547904 | 1.563925 | 0.942766 | 9.808185 | 0.836383 |
| 36.5278 | 2.30306 | 1.762622 | 0.596855 | 2.543786 | 3.494524 | 0.998845 | 2.037208 | 0.041655 | 0.704827 | -0.0744 | 3.466039 | 0.245621 | 0.555979 | 0.568745 | 0.525441 | 0.683394 | 0.412196 | -0.032863 | 0.463835 | 0.947292 | 0.144336 | 3.283171 | 0.304511 |
| 36.5417 | 2.105551 | 1.647022 | 0.719558 | 3.129033 | 3.260328 | 0.923087 | 1.88383 | -0.838111 | 0.012567 | -0.258223 | 2.612639 | 0.038802 | -0.126482 | 0.440178 | -0.046566 | 0.187965 | 0.162342 | -0.472717 | 0.386518 | -0.096839 | -0.811202 | -1.576941 | -0.270231 |
| 36.5556 | 1.843052 | 1.313859 | 0.964995 | 3.483381 | 2.714869 | 0.974576 | 2.229892 | -1.281311 | -0.803043 | -0.413667 | 2.001472 | 0.050785 | -0.057647 | 0.339852 | -0.346429 | -0.225443 | 0.084764 | -0.656866 | 0.319955 | -0.969132 | -1.57523 | -9.110714 | -0.769638 |
| 36.5694 | 1.474764 | 0.805673 | 1.302451 | 3.133095 | 1.789991 | 0.957196 | 2.154 | -0.972885 | -1.228481 | -0.53119 | 1.449551 | 0.306427 | 0.104906 | 0.258417 | -0.195929 | -0.386006 | 0.239105 | -0.569857 | 0.256379 | -1.228348 | -1.660857 | -9.669109 | -0.911262 |
| 36.5833 | 1.106113 | 0.28923 | 1.418309 | 1.605653 | 0.851874 | 0.254203 | 0.879024 | -0.724135 | -0.813367 | -0.678993 | 0.345295 | 0.571163 | 0.169635 | 0.306654 | 0.347013 | -0.554566 | 0.489788 | -0.273604 | 0.273068 | -1.511922 | -1.286269 | -11.356137 | -0.577838 |
| 36.5972 | 0.449986 | -0.195623 | 0.934637 | -0.078201 | -0.147872 | -0.698824 | -0.478482 | -0.859708 | -0.091427 | -0.749632 | -1.036983 | 0.560545 | 0.195482 | 0.392725 | 0.711309 | -0.373436 | 0.742288 | 0.044772 | 0.238704 | -1.770451 | -0.514198 | -5.24257 | -0.411686 |
| 36.6111 | 0.111561 | -0.67902 | 0.604203 | -1.442153 | -1.131609 | -1.273505 | -1.484046 | -1.826285 | 0.113057 | -0.787635 | -2.607352 | 0.317344 | 0.093441 | 0.38391 | 0.656887 | -0.54584 | 0.663897 | 0.157682 | 0.218049 | -2.228895 | -0.390611 | 1.038995 | -0.627796 |
| 36.625 | -0.192235 | -1.071785 | 0.457796 | -1.795924 | -2.251499 | -1.759212 | -1.68004 | -3.122314 | -0.097028 | -0.866283 | -3.635396 | -0.033104 | -0.053822 | 0.247752 | 0.095724 | -0.831318 | 0.387542 | 0.081737 | 0.146135 | -2.406174 | -0.833002 | 7.066887 | -0.707716 |
| 36.6389 | -0.495465 | -1.40578 | 0.522219 | -1.929188 | -3.699763 | -1.153681 | -0.751166 | -4.189514 | -0.624109 | -1.026096 | -4.083797 | -0.384246 | -0.196186 | 0.082264 | -0.594414 | -1.081565 | 0.188399 | -0.100761 | 0.141374 | -2.689556 | -1.108124 | 15.795937 | -0.660652 |
| 36.6528 | -0.497152 | -1.487054 | 0.74733 | -1.482529 | -3.740975 | -1.43232 | -0.249166 | -3.528417 | -0.314406 | -1.08445 | -4.486759 | -0.694043 | 0.16401 | -0.079211 | -0.818672 | -1.155725 | 0.043432 | -0.33337 | 0.108436 | -2.198934 | -1.008833 | 13.322347 | -0.705898 |
| 36.6667 | -0.206629 | -1.359778 | 1.123471 | -0.96797 | -4.253216 | -1.203328 | 0.290178 | -1.594516 | -0.045422 | -0.91984 | -4.354478 | -0.856271 | 1.978341 | -0.162227 | -0.588549 | -1.186557 | 0.188812 | -0.3925 | 0.084963 | -1.475304 | -0.675741 | 10.244423 | -0.83656 |
| 36.6806 | 0.060778 | -1.052719 | 1.367128 | -0.531363 | -4.056092 | -1.849321 | 0.511471 | 0.939466 | 0.377729 | -0.671152 | -3.84517 | -0.810867 | 0.690823 | -0.090332 | -0.202434 | -0.92752 | 0.32299 | -0.268129 | 0.135658 | -1.231378 | 0.0333 | 2.202401 | -0.826147 |
| 36.6944 | 0.134628 | -0.775937 | 1.225017 | -0.162303 | -4.310247 | -1.848565 | 0.654118 | 2.883279 | 0.321352 | -0.519311 | -2.793006 | -0.687875 | -1.153978 | 0.01924 | 0.32011 | -0.585584 | 0.563227 | -0.01277 | 0.116751 | -0.93823 | 0.536798 | -2.983587 | -0.64699 |
| 36.7083 | 0.030556 | -0.679876 | 0.789911 | 0.361353 | -3.61241 | -1.746342 | 0.640631 | 3.910125 | 0.518976 | -0.511467 | -1.71829 | -0.507235 | -1.551467 | -0.063227 | 0.677198 | -0.255023 | 0.630202 | 0.224044 | -0.073139 | -0.484713 | 0.462726 | -9.452938 | -0.809499 |
| 36.7222 | -0.151925 | -0.667622 | 0.525265 | 0.976014 | -2.155669 | -1.513379 | 0.509581 | 3.557846 | 0.232447 | -0.506986 | -0.670551 | -0.394899 | -0.261062 | -0.211345 | 0.71398 | -0.118243 | 0.455353 | 0.191008 | -0.275145 | 0.13838 | -0.283214 | -10.761543 | -0.552848 |
| 36.7361 | -0.204608 | -0.65711 | 0.350175 | 1.541277 | -1.324671 | -1.399617 | 0.203563 | 3.06472 | -0.475744 | -0.542655 | -0.167897 | -0.303024 | -0.223394 | -0.302679 | 0.347396 | -0.117968 | 0.190343 | 0.097064 | -0.318652 | 0.855541 | -0.788895 | -4.681241 | -0.285563 |
| 36.75 | 0.087254 | -0.659446 | -0.03086 | 1.94721 | -0.980775 | -1.507148 | -0.207097 | 2.61298 | -0.718075 | -0.616175 | -0.054715 | -0.264401 | -0.259227 | -0.322841 | -0.176569 | 0.163859 | -0.071999 | 0.041916 | -0.281891 | 1.472358 | -0.785449 | 3.446627 | -0.397169 |
| 36.7639 | 0.326746 | -0.807111 | -0.840684 | 2.119653 | -0.517698 | -1.401678 | -0.501084 | 1.508887 | -1.741052 | -0.721113 | 0.008583 | -0.339443 | -0.330302 | -0.278455 | -0.433446 | 0.621966 | -0.441784 | -0.011746 | -0.193425 | 1.818184 | -0.365943 | 9.160737 | -0.313834 |
| 36.7778 | 0.232263 | -1.080768 | -0.958607 | 2.04076 | -0.108216 | -1.127382 | -0.8346 | -0.231522 | -3.790713 | -0.898549 | -0.959071 | -0.595465 | -0.415698 | -0.27936 | -0.109796 | 0.91304 | -0.792083 | -0.094428 | -0.130862 | 2.011502 | -0.388463 | 10.494968 | 0.062156 |
| 36.7917 | -0.223256 | -1.258826 | -0.913471 | 1.724325 | -0.241147 | -0.965354 | -1.338604 | -1.982453 | -3.567794 | -1.186476 | -1.822758 | -0.824157 | 1.196895 | -0.289576 | 0.441408 | 1.020049 | -0.809911 | 0.032204 | -0.133056 | 1.737961 | -0.323724 | 13.974339 | -0.083052 |
| 36.8056 | -0.779102 | -1.2413 | -1.115172 | 1.235164 | -0.743108 | -1.040169 | -1.745301 | -3.505749 | -2.966583 | -1.46994 | -2.254418 | -0.898817 | 2.034466 | -0.300239 | 0.949864 | 0.935888 | -0.62329 | 0.313501 | -0.126287 | 1.367861 | -0.253009 | 13.879346 | 1.39207 |
| 36.8194 | -1.009059 | -0.926378 | -1.491757 | 0.867627 | -1.660152 | -1.753251 | -1.864233 | -4.10492 | -2.379636 | -1.700928 | -2.697909 | -0.821711 | 1.026473 | -0.284727 | 1.212197 | 0.829095 | -0.289334 | 0.610278 | -0.200048 | 0.816749 | 0.093051 | 2.730006 | 0.173681 |
| 36.8333 | -0.923701 | -0.561174 | -2.037805 | 0.758881 | -2.419318 | -2.27055 | -1.913868 | -5.10661 | -1.949242 | -1.682331 | -3.018 | -0.69771 | -1.667737 | -0.271536 | 1.198286 | 0.648643 | -0.127002 | 0.794421 | -0.2574 | 0.324364 | 0.247063 | -2.640653 | -1.176069 |
| 36.8472 | -0.688071 | -0.134301 | -1.951043 | 0.792676 | -2.45771 | -1.888011 | -1.821275 | -6.030853 | -1.427254 | -2.198334 | -2.806434 | -0.604212 | -1.544815 | -0.228083 | 1.146749 | 0.358172 | -0.08101 | 0.707768 | -0.306019 | -0.019075 | 0.37446 | -8.299484 | -1.329464 |
| 36.8611 | -0.222112 | 0.304202 | -1.475499 | 0.723673 | -2.068871 | -1.338928 | -1.445116 | -6.05328 | -1.239437 | -2.349512 | -2.100968 | -0.530422 | -0.366845 | -0.178332 | 0.968091 | 0.156982 | -0.173574 | 0.452049 | -0.317291 | -0.230734 | 0.229282 | -6.717532 | -0.265638 |
| 36.875 | 0.378597 | 0.735193 | -0.766783 | 0.652425 | -1.688344 | -0.84799 | -1.01074 | -4.502979 | -0.642693 | -1.740884 | -1.338538 | -0.472033 | -0.167946 | -0.122044 | 0.582943 | 0.009239 | -0.189583 | 0.243851 | -0.288674 | -0.34827 | 0.263073 | -5.129284 | -0.227814 |
| 36.8889 | 0.740046 | 1.138828 | -0.138311 | 0.193286 | -1.268045 | -0.415582 | -0.912845 | -1.463017 | 0.451068 | -1.679903 | -1.3262 | -0.359208 | -0.196027 | -0.337296 | 0.262424 | -0.035016 | -0.197018 | 0.145398 | -0.274413 | -0.479659 | 1.023534 | -2.349981 | -0.346113 |
| 36.9028 | -0.348145 | -0.093145 | -0.652642 | -2.213731 | -1.462266 | -0.103199 | -3.238156 | 0.80634 | 1.514028 | -2.983203 | -7.560918 | -0.255162 | -0.192213 | -0.681649 | 0.156421 | 0.042725 | -0.08084 | 0.251016 | -0.272905 | -0.623534 | 1.255689 | 2.381658 | -0.547774 |
| 36.9167 | -3.264351 | -4.002823 | -2.77743 | -5.295673 | -2.481664 | -0.360298 | -7.127692 | -1.396803 | 0.891438 | -6.080416 | -20.132351 | -0.173702 | -0.415327 | -0.574046 | 0.202125 | 0.229003 | 0.120952 | 0.458005 | -0.317432 | -0.489272 | 1.337913 | 5.932434 | -1.550603 |
| 36.9306 | -4.980178 | -7.805779 | -3.478115 | -3.998001 | -3.877878 | -1.014135 | -6.957634 | -6.67687 | -2.272995 | -6.598045 | -20.79099 | -0.370779 | -1.720706 | -0.377082 | -0.2802 | 0.058671 | 0.728669 | 0.391802 | -1.28265 | -1.187681 | 1.266982 | 4.474374 | -6.742885 |
| 36.9444 | -3.485882 | -4.926038 | -2.15355 | -1.275838 | -3.132353 | -1.287304 | -3.62216 | -9.36238 | -2.587268 | -3.001347 | -10.602246 | -1.627788 | -4.193065 | -1.133561 | -1.25673 | -0.724485 | 1.057093 | -0.462149 | -4.451124 | -3.961192 | -0.855039 | 4.684337 | -13.982802 |
| 36.9583 | -1.077538 | -0.863193 | -0.500756 | 0.48847 | -2.141498 | -0.945758 | -1.145652 | -7.220344 | -0.869297 | -0.626042 | -3.108013 | -2.873518 | -1.96515 | -1.92386 | -3.580298 | -2.425986 | 0.262682 | -1.710576 | -5.668464 | -6.468343 | -2.877057 | -0.903996 | -10.645021 |
| 36.9722 | -0.184449 | 0.320399 | -0.083942 | 1.210433 | -2.881714 | -0.648788 | -0.628764 | -5.100363 | -0.496802 | -0.237568 | -1.555045 | -1.919618 | -2.482349 | -1.012534 | -3.049408 | -1.763016 | -0.594469 | -1.553536 | -3.083445 | -5.011247 | -2.190179 | -3.589075 | -6.562854 |
| 36.9861 | -0.245629 | -0.116574 | -0.2753 | 1.238693 | -3.277455 | -0.370458 | -0.726442 | -3.971161 | -1.079273 | -0.08597 | -1.657622 | 0.137024 | -2.618332 | 0.525473 | -1.238092 | 0.639653 | 0.828753 | 0.029606 | 0.60399 | -0.687352 | -0.33533 | -6.824333 | -1.791997 |
| 37.0069 | -0.49061 | -0.811441 | -0.46868 | 1.044348 | -2.50589 | -0.277837 | -0.796136 | -3.109641 | -1.460515 | 0.004748 | -1.317432 | 1.046883 | -1.413082 | 1.308529 | 0.16613 | 1.412005 | 1.466101 | 0.840416 | 1.812748 | 1.527178 | 1.153309 | -8.822436 | -0.025535 |
| 37.0208 | -0.806762 | -1.368163 | -0.553499 | 0.826882 | -2.224366 | -0.25118 | -0.847108 | -2.819036 | -1.647494 | -0.348733 | -1.086014 | 0.579612 | -0.972414 | 0.86738 | 0.649025 | 1.15848 | 0.796341 | 0.589044 | 1.068554 | 1.569862 | 1.466341 | -11.763208 | 0.992786 |
| 37.0347 | -1.060395 | -1.612451 | -0.54761 | 0.61735 | -2.355196 | -0.232959 | -0.789809 | -2.074543 | -1.654986 | -0.673124 | -1.053353 | 0.214224 | 0.083033 | 0.400616 | 1.364669 | 1.047627 | 0.370164 | 0.583342 | 0.439855 | 1.326331 | 1.201584 | -10.638728 | 0.774524 |
| 37.0486 | -1.130647 | -1.440753 | -0.483383 | 0.470655 | -2.404498 | -0.150403 | -0.714285 | -0.794884 | -1.244008 | -0.688094 | -0.869685 | 0.189551 | 0.422679 | 0.384793 | 2.121162 | 0.809187 | 0.307022 | 0.755874 | 0.186827 | 1.565052 | 1.916096 | -8.637896 | 0.965675 |
| 37.0625 | -1.002283 | -1.076503 | -0.456225 | 0.408382 | -2.146905 | -0.100103 | -0.567794 | 0.820422 | -0.432527 | -0.62842 | -0.388 | 0.216448 | 0.627274 | 0.438839 | 2.238349 | 0.581438 | 0.385548 | 0.823143 | 0.146862 | 1.82807 | 2.181247 | -7.008919 | 1.257728 |
| 37.0764 | -0.531914 | -0.71515 | -0.295062 | 0.448817 | -1.569046 | -0.084279 | -0.340612 | 2.276865 | 0.545322 | -0.63693 | 0.312006 | 0.218938 | 0.448783 | 0.413188 | 1.75359 | 0.407434 | 0.42177 | 0.652774 | 0.140959 | 1.656254 | 1.575614 | -6.400224 | 1.137574 |
| 37.0903 | -0.059341 | -0.539816 | -0.033394 | 0.542655 | -0.813334 | -0.021063 | -0.102581 | 3.335678 | 1.477873 | -0.552228 | 1.095758 | 0.077015 | 0.141076 | 0.368645 | 1.090025 | 0.071894 | 0.273801 | 0.255874 | 0.029781 | 1.247285 | 0.633218 | -8.308547 | 0.563236 |

| | | | |
|---|---|---|---|
| 34.4792 | 2.64924 | 34.4722 | 8.225337 |
| 34.4931 | 3.007158 | 34.4861 | 1.854185 |
| 34.5 | 3.075496 | 34.5 | 0.949355 |
| 34.5139 | 2.656514 | 34.5139 | 0.653242 |
| 34.5278 | 2.100327 | 34.5278 | 0.109922 |
| 34.5417 | 1.552476 | 34.5417 | -0.400336 |
| 34.5556 | 1.056215 | 34.5556 | -0.64666 |
| 34.5694 | 0.112288 | 34.5694 | -0.638341 |
| 34.5833 | -1.100043 | 34.5833 | -0.680568 |
| 34.5972 | -2.151501 | 34.5972 | -0.7029 |
| 34.6111 | -2.911751 | 34.6111 | -0.906638 |
| 34.625 | -3.124337 | 34.625 | -0.965825 |
| 34.6389 | -2.787518 | 34.6389 | -0.732936 |
| 34.6528 | -2.592479 | 34.6528 | -0.346541 |
| 34.6667 | -2.108469 | 34.6667 | 0.080443 |
| 34.6806 | -1.572614 | 34.6806 | 0.47359 |
| 34.6944 | -0.980691 | 34.6944 | 0.828116 |
| 34.7083 | -0.647378 | 34.7083 | 0.876795 |
| 34.7222 | -0.556335 | 34.7222 | 0.624789 |
| 34.7361 | -0.621668 | 34.7361 | 0.147135 |
| 34.75 | -0.677233 | 34.75 | -0.290198 |
| 34.7639 | -0.892824 | 34.7639 | -0.54711 |
| 34.7778 | -1.314763 | 34.7778 | -0.647206 |
| 34.7917 | -1.859646 | 34.7917 | -0.649315 |
| 34.8056 | -2.410037 | 34.8056 | -0.618593 |
| 34.8194 | -2.829004 | 34.8194 | -0.562321 |
| 34.8333 | -2.867256 | 34.8333 | -0.476819 |
| 34.8472 | -2.979838 | 34.8472 | -0.286863 |
| 34.8611 | -2.273506 | 34.8611 | -0.116479 |
| 34.875 | -0.929198 | 34.875 | -0.104235 |
| 34.8889 | -1.153821 | 34.8889 | -0.075794 |
| 34.9028 | -11.794476 | 34.9028 | 0.033593 |
| 34.9167 | -28.284357 | 34.9167 | -1.074569 |
| 34.9306 | -24.647908 | 34.9306 | -9.344092 |
| 34.9444 | -10.611492 | 34.9444 | -20.09496 |
| 34.9583 | -2.747789 | 34.9583 | -18.822915 |
| 34.9722 | -1.8744 | 34.9722 | -8.652979 |
| 34.9861 | -2.240505 | 34.9861 | -2.690254 |
| 35.0069 | -2.131781 | 35 | -0.574871 |
| 35.0208 | -1.891135 | 35.0139 | 0.645469 |
| 35.0347 | -1.888718 | 35.0278 | 1.558157 |
| 35.0486 | -1.786363 | 35.0417 | 1.95174 |
| 35.0625 | -1.430535 | 35.0556 | 2.001013 |
| 35.0764 | -0.58866 | 35.0694 | 1.868549 |
| 35.0903 | 0.903519 | 35.0833 | 1.387982 |
| 35.1042 | 1.669049 | 35.0972 | 0.776888 |
| 35.1181 | 1.41135 | 35.1111 | 0.587575 |
| 35.1319 | 1.390897 | 35.125 | 0.465885 |
| 35.1458 | 1.56059 | 35.1389 | 0.407857 |
| 35.1597 | 1.352013 | 35.1528 | 0.358975 |
| 35.1736 | 1.095802 | 35.1667 | 0.395818 |
| 35.1875 | 0.700999 | 35.1806 | 0.382015 |
| 35.2014 | 0.046692 | 35.1944 | 0.227951 |
| 35.2153 | -0.362331 | 35.2083 | 0.062697 |
| 35.2292 | -0.602017 | 35.2222 | 0.333722 |
| 35.2431 | -0.409645 | 35.2361 | 0.516638 |
| 35.2569 | -0.127827 | 35.25 | 0.285805 |
| 35.2708 | -0.380713 | 35.2639 | 0.032527 |
| 35.2847 | -1.17077 | 35.2778 | 0.010482 |
| 35.2986 | -1.046222 | 35.2917 | 0.020264 |
| 35.3125 | -0.866945 | 35.3056 | 0.085165 |
| 35.3264 | -1.361167 | 35.3194 | 0.397888 |
| 35.3403 | -1.393386 | 35.3333 | 0.606228 |
| 35.3542 | -1.17586 | 35.3472 | 0.397539 |
| 35.3681 | -1.336504 | 35.3611 | 0.171813 |
| 35.3819 | -1.378513 | 35.375 | 0.22862 |
| 35.3958 | -0.500408 | 35.3889 | 0.33657 |
| 35.4097 | 6.697633 | 35.4028 | 0.310475 |
| 35.4236 | 19.941489 | 35.4167 | 0.765873 |
| 35.4375 | 23.785753 | 35.4306 | 6.148671 |
| 35.4514 | 10.120124 | 35.4444 | 16.232416 |
| 35.4653 | 2.074983 | 35.4583 | 19.6845 |
| 35.4792 | 1.608709 | 35.4722 | 9.211278 |
| 35.4931 | 2.294162 | 35.4861 | 2.077131 |
| 35.5 | 2.257466 | 35.5 | 0.855167 |
| 35.5139 | 1.742918 | 35.5139 | 0.604013 |
| 35.5278 | 1.362087 | 35.5278 | 0.165509 |
| 35.5417 | 0.953908 | 35.5417 | -0.349918 |
| 35.5556 | 0.604401 | 35.5556 | -0.443736 |
| 35.5694 | 0.152219 | 35.5694 | -0.330202 |
| 35.5833 | -0.42801 | 35.5833 | -0.157661 |
| 35.5972 | -1.354933 | 35.5972 | -0.107524 |
| 35.6111 | -1.958295 | 35.6111 | -0.22295 |
| 35.625 | -2.252185 | 35.625 | -0.422448 |
| 35.6389 | -2.393954 | 35.6389 | -0.707026 |
| 35.6528 | -2.30512 | 35.6528 | -1.078546 |
| 35.6667 | -2.249439 | 35.6667 | -1.126432 |
| 35.6806 | -2.097065 | 35.6806 | -0.990629 |
| 35.6944 | -1.780924 | 35.6944 | -0.705714 |
| 35.7083 | -1.36908 | 35.7083 | -0.395838 |
| 35.7222 | -1.207989 | 35.7222 | -0.256837 |
| 35.7361 | -1.222866 | 35.7361 | -0.279238 |
| 35.75 | -1.48133 | 35.75 | -0.351094 |
| 35.7639 | -1.630948 | 35.7639 | -0.441607 |
| 35.7778 | -1.741084 | 35.7778 | -0.451501 |
| 35.7917 | -2.033388 | 35.7917 | -0.500307 |
| 35.8056 | -2.617754 | 35.8056 | -0.573262 |
| 35.8194 | -2.725859 | 35.8194 | -0.663568 |
| 35.8333 | -2.785872 | 35.8333 | -0.673228 |
| 35.8472 | -2.437595 | 35.8472 | -0.590363 |
| 35.8611 | -1.817087 | 35.8611 | -0.541487 |
| 35.875 | -1.153709 | 35.875 | -0.502382 |
| 35.8889 | -2.051208 | 35.8889 | -0.459575 |
| 35.9028 | -11.130126 | 35.9028 | -0.311837 |
| 35.9167 | -25.654492 | 35.9167 | -1.090147 |
| 35.9306 | -25.508981 | 35.9306 | -7.149447 |
| 35.9444 | -12.68933 | 35.9444 | -17.312724 |
| 35.9583 | -4.565597 | 35.9583 | -17.139496 |
| 35.9722 | -2.453558 | 35.9722 | -9.282055 |
| 35.9861 | -1.656049 | 35.9861 | -0.871627 |
| 36.0069 | -0.982308 | 36 | 2.08976 |
| 36.0208 | -0.536283 | 36.0139 | 2.188711 |
| 36.0347 | -0.262965 | 36.0278 | 1.938638 |
| 36.0486 | 0.25832 | 36.0417 | 1.827269 |
| 36.0625 | 0.612453 | 36.0556 | 1.483889 |
| 36.0764 | 1.129362 | 36.0694 | 1.149965 |
| 36.0903 | 2.076801 | 36.0833 | 0.703934 |
| 36.1042 | 2.611033 | 36.0972 | 0.215181 |
| 36.1181 | 2.835117 | 36.1111 | -0.2293 |
| 36.1319 | 3.237173 | 36.125 | -0.314071 |
| 36.1458 | 3.942988 | 36.1389 | -0.008054 |
| 36.1597 | 3.789894 | 36.1528 | 0.439186 |
| 36.1736 | 3.788038 | 36.1667 | 0.546567 |
| 36.1875 | 3.833471 | 36.1806 | 0.069786 |
| 36.2014 | 3.291779 | 36.1944 | -0.191537 |
| 36.2153 | 2.607885 | 36.2083 | 0.05899 |
| 36.2292 | 1.95198 | 36.2222 | 0.14595 |
| 36.2431 | 1.310832 | 36.2361 | 0.078751 |
| 36.2569 | 0.872684 | 36.25 | 0.285253 |
| 36.2708 | -0.078622 | 36.2639 | 0.473959 |
| 36.2847 | -0.835385 | 36.2778 | 0.470262 |
| 36.2986 | -0.473114 | 36.2917 | 0.427782 |
| 36.3125 | 0.117867 | 36.3056 | 0.206792 |
| 36.3264 | 0.024816 | 36.3194 | -0.058858 |
| 36.3403 | -0.399111 | 36.3333 | -0.087738 |
| 36.3542 | -0.767193 | 36.3472 | 0.141235 |
| 36.3681 | -0.929248 | 36.3611 | 0.40817 |
| 36.3819 | -1.220522 | 36.375 | 0.524482 |
| 36.3958 | -0.174707 | 36.3889 | 0.461873 |
| 36.4097 | 9.073669 | 36.4028 | 0.287602 |
| 36.4236 | 27.851782 | 36.4167 | 0.91587 |
| 36.4375 | 31.207182 | 36.4306 | 5.78935 |
| 36.4514 | 13.759442 | 36.4444 | 14.647488 |
| 36.4653 | 3.352107 | 36.4583 | 17.75193 |
| 36.4792 | 2.632545 | 36.4722 | 9.766939 |
| 36.4931 | 3.247707 | 36.4861 | 2.659524 |
| 36.5 | 3.591701 | 36.5 | 1.159379 |
| 36.5139 | 4.01695 | 36.5139 | 0.836383 |
| 36.5278 | 3.466039 | 36.5278 | 0.304511 |
| 36.5417 | 2.612639 | 36.5417 | -0.270231 |
| 36.5556 | 2.001472 | 36.5556 | -0.769638 |
| 36.5694 | 1.449551 | 36.5694 | -0.911262 |
| 36.5833 | 0.345295 | 36.5833 | -0.577838 |
| 36.5972 | -1.036983 | 36.5972 | -0.411686 |
| 36.6111 | -2.607352 | 36.6111 | -0.627796 |
| 36.625 | -3.635396 | 36.625 | -0.707716 |
| 36.6389 | -4.083797 | 36.6389 | -0.660652 |
| 36.6528 | -4.486759 | 36.6528 | -0.705898 |
| 36.6667 | -4.354478 | 36.6667 | -0.83656 |
| 36.6806 | -3.84517 | 36.6806 | -0.826147 |
| 36.6944 | -2.793006 | 36.6944 | -0.64699 |
| 36.7083 | -1.71829 | 36.7083 | -0.809499 |
| 36.7222 | -0.670551 | 36.7222 | -0.552848 |
| 36.7361 | -0.167897 | 36.7361 | -0.285563 |
| 36.75 | -0.054715 | 36.75 | -0.397169 |
| 36.7639 | 0.008583 | 36.7639 | -0.313834 |
| 36.7778 | -0.959071 | 36.7778 | 0.062156 |
| 36.7917 | -1.822758 | 36.7917 | -0.083052 |
| 36.8056 | -2.254418 | 36.8056 | 1.39207 |
| 36.8194 | -2.697909 | 36.8194 | 0.173681 |
| 36.8333 | -3.018 | 36.8333 | -1.176069 |
| 36.8472 | -2.806434 | 36.8472 | -1.329464 |
| 36.8611 | -2.100968 | 36.8611 | -0.265638 |
| 36.875 | -1.338538 | 36.875 | -0.227814 |
| 36.8889 | -1.3262 | 36.8889 | -0.346113 |
| 36.9028 | -7.560918 | 36.9028 | -0.547774 |
| 36.9167 | -20.132351 | 36.9167 | -1.550603 |
| 36.9306 | -20.79099 | 36.9306 | -6.742885 |
| 36.9444 | -10.602246 | 36.9444 | -13.982802 |
| 36.9583 | -3.108013 | 36.9583 | -10.645021 |
| 36.9722 | -1.555045 | 36.9722 | -6.562854 |
| 36.9861 | -1.657622 | 36.9861 | -1.791997 |
| 37.0069 | -1.317432 | 37 | -0.025535 |
| 37.0208 | -1.086014 | 37.0139 | 0.992786 |
| 37.0347 | -1.053353 | 37.0278 | 0.774524 |
| 37.0486 | -0.869685 | 37.0417 | 0.965675 |
| 37.0625 | -0.388 | 37.0556 | 1.257728 |
| 37.0764 | 0.312006 | 37.0694 | 1.137574 |
| 37.0903 | 1.095758 | 37.0833 | 0.563236 |

| | | | | | | | | | | | | | | | | | | | | | | | |
|---|---|---|---|---|---|---|---|---|---|---|---|---|---|---|---|---|---|---|---|---|---|---|---|
| 37.1042 | 0.250403 | -0.559333 | 0.146496 | 0.641412 | -0.172103 | 0.100958 | 0.018099 | 4.361855 | 2.488415 | -0.51193 | 1.511558 | -0.013257 | -0.176771 | 0.363794 | 0.63532 | -0.326706 | 0.134019 | -0.105792 | -0.135137 | 1.025869 | -0.109481 | -10.56965 | -0.008774 |
| 37.1181 | 0.572155 | -0.403639 | 0.312907 | 0.686524 | 0.314223 | 0.210052 | 0.07535 | 5.582313 | 3.374386 | -0.501001 | 1.925578 | 0.036634 | -0.216674 | 0.459523 | 0.464955 | -0.502617 | 0.134704 | -0.163342 | -0.192682 | 0.70225 | -0.495684 | -6.974692 | -0.272534 |
| 37.1319 | 1.02846 | -0.045463 | 0.482876 | 0.665681 | 0.663642 | 0.225053 | 0.132753 | 6.334311 | 4.145245 | -0.486502 | 2.661933 | 0.262719 | -0.064954 | 0.542254 | 0.422773 | -0.32094 | 0.259545 | 0.099516 | -0.049407 | 0.332133 | -0.772824 | -0.002084 | -0.268671 |
| 37.1458 | 1.342797 | 0.158415 | 0.544796 | 0.561467 | 0.825386 | 0.193809 | 0.088348 | 6.966481 | 4.407538 | -0.612215 | 3.41151 | 0.473566 | 0.111876 | 0.585169 | 0.437108 | 0.070361 | 0.377595 | 0.400497 | 0.211256 | 0.240531 | -0.967484 | 0.03905 | -0.107529 |
| 37.1597 | 1.198743 | 0.110486 | 0.473531 | 0.411651 | 0.865644 | 0.172736 | 0.069554 | 6.748059 | 4.384829 | -0.694732 | 4.015856 | 0.55444 | 0.171956 | 0.555626 | 0.375409 | 0.410787 | 0.31078 | 0.448132 | 0.415645 | 0.267966 | -0.910551 | -0.00059 | 0.085615 |
| 37.1736 | 1.030202 | -0.072069 | 0.399729 | 0.273153 | 0.813271 | 0.117529 | 0.127883 | 4.959957 | 4.192877 | -0.563485 | 4.223404 | 0.573678 | 0.214082 | 0.489283 | 0.295283 | 0.670692 | 0.09893 | 0.269068 | 0.546954 | 0.012757 | -0.761238 | -0.085458 | 0.219951 |
| 37.1875 | 0.831183 | -0.31269 | 0.382701 | 0.251815 | 0.78124 | 0.089826 | 0.126851 | 4.577573 | 3.79573 | -0.48381 | 4.446053 | 0.502238 | 0.181672 | 0.399492 | 0.307456 | 0.835416 | -0.0187 | 0.165378 | 0.504584 | -0.454642 | -0.520837 | -0.087998 | 0.191029 |
| 37.2014 | 0.571398 | -0.517076 | 0.343859 | 0.341779 | 0.745461 | 0.093772 | 0.008716 | 5.204805 | 2.963429 | -0.549986 | 3.71623 | 0.328034 | 0.062124 | 0.28575 | 0.200063 | 0.783635 | -0.022298 | 0.174007 | 0.329375 | -0.951216 | -0.262934 | -0.057177 | -0.046683 |
| 37.2153 | 0.283099 | -0.577837 | 0.262672 | 0.464815 | 0.60444 | 0.078255 | -0.133496 | 4.84115 | 2.596913 | -0.512197 | 2.770993 | 0.172021 | -0.004675 | 0.208123 | -0.155457 | 0.661098 | -0.029884 | 0.257483 | 0.140238 | -1.26367 | -0.060356 | -0.02748 | -0.156667 |
| 37.2292 | 0.112562 | -0.589842 | 0.202673 | 0.619386 | 0.404255 | 0.02156 | -0.256411 | 3.859129 | 1.506287 | -0.456034 | 1.849508 | 0.136286 | 0.001506 | 0.175666 | -0.560263 | 0.521399 | -0.058079 | 0.428462 | 0.020875 | -1.274216 | -0.021052 | 0.025774 | -0.113892 |
| 37.2431 | 0.202543 | -0.565075 | 0.218776 | 0.84538 | 0.29739 | -0.088139 | -0.327293 | 2.681549 | 0.534465 | -0.408422 | 1.09992 | 0.195811 | 0.057449 | 0.14122 | -0.923675 | 0.363872 | -0.145475 | 0.529097 | 0.012463 | -1.049016 | -0.037928 | 0.035481 | -0.122379 |
| 37.2569 | 0.510969 | -0.299432 | 0.362927 | 1.101712 | 0.342576 | -0.185204 | -0.268463 | 1.860238 | 0.397255 | -0.294752 | 0.901616 | 0.317522 | 0.192576 | 0.092096 | -0.863839 | 0.373608 | -0.289678 | 0.447174 | 0.094812 | -0.645038 | 0.176273 | -0.028055 | -0.027098 |
| 37.2708 | 0.832751 | 0.167604 | 0.510291 | 1.203422 | 0.413832 | -0.15078 | -0.083472 | 1.380781 | 0.876914 | -0.157829 | 0.977081 | 0.489118 | 0.377607 | 0.082068 | -0.607741 | 0.587683 | -0.316354 | 0.158046 | 0.185803 | -0.241735 | 0.517036 | -0.064362 | 0.052359 |
| 37.2847 | 1.036584 | 0.503401 | 0.529245 | 1.13866 | 0.272306 | -0.018579 | 0.119276 | 1.308374 | 1.679485 | -0.180162 | 1.056995 | 0.614436 | 0.489316 | 0.077843 | -0.393621 | 0.831538 | -0.276531 | -0.075784 | 0.185878 | 0.070393 | 0.537657 | -0.004171 | -0.022402 |
| 37.2986 | 0.956351 | 0.514779 | 0.505639 | 1.058588 | -0.192794 | 0.026345 | 0.215486 | 1.401968 | 2.379619 | -0.196258 | 1.158017 | 0.620245 | 0.446871 | 0.04995 | -0.190044 | 0.912718 | -0.238995 | -0.145658 | 0.161744 | 0.361631 | 0.210264 | 8.1925 | -0.011248 |
| 37.3125 | 0.726847 | 0.352464 | 0.542844 | 1.023298 | -0.613348 | 0.00307 | 0.291395 | 1.558022 | 2.720037 | -0.086835 | 1.1502 | 0.522316 | 0.277536 | 0.088462 | -0.093115 | 0.777053 | -0.257161 | -0.102687 | 0.134771 | 0.538922 | -0.234241 | 12.160247 | -0.034138 |
| 37.3264 | 0.630916 | 0.341481 | 0.670321 | 1.079785 | -0.731668 | 0.041537 | 0.467722 | 1.369626 | 2.159759 | -0.025834 | 0.930207 | 0.391143 | 0.104655 | 0.078192 | -0.093067 | 0.492654 | -0.289427 | -0.014285 | 0.03484 | 0.40372 | -0.497914 | 11.452486 | -0.136696 |
| 37.3403 | 0.638259 | 0.423375 | 0.759902 | 1.152642 | -0.738656 | 0.107731 | 0.625418 | 0.400861 | 1.070044 | -0.118296 | 0.326254 | 0.348442 | 0.070712 | 0.057668 | -0.015742 | 0.326139 | -0.243358 | 0.121707 | -0.040645 | 0.169615 | -0.346659 | 8.474336 | -0.140449 |
| 37.3542 | 0.547568 | 0.3404 | 0.691106 | 1.156536 | -0.951103 | 0.093922 | 0.572271 | -0.920784 | -0.300576 | -0.313334 | -0.361935 | 0.417236 | 0.201496 | 0.1358 | 0.063601 | 0.298157 | -0.11112 | 0.195658 | 0.046492 | 0.221487 | 0.148417 | 8.062345 | 0.029102 |
| 37.3681 | 0.373409 | 0.155712 | 0.513553 | 1.064163 | -1.272576 | 0.001981 | 0.371298 | -1.828183 | -1.37636 | -0.48785 | -0.747219 | 0.479455 | 0.357758 | 0.267548 | 0.016561 | 0.335324 | -0.045021 | 0.073065 | 0.209686 | 0.594205 | 0.613428 | 7.389426 | 0.270006 |
| 37.3819 | 0.234415 | 0.076824 | 0.394992 | 0.983006 | -1.372952 | -0.034924 | 0.250362 | -2.130531 | -1.926599 | -0.584983 | -0.904591 | 0.381374 | 0.343412 | 0.27546 | -0.119209 | 0.300229 | -0.160195 | -0.205519 | 0.236985 | 0.842248 | 0.803573 | 6.831247 | 0.299432 |
| 37.3958 | 0.344169 | 0.356625 | 0.554181 | 0.995238 | -1.129857 | -0.035641 | 0.520101 | -2.490322 | -2.410838 | -0.363583 | -0.181694 | 0.217803 | 0.143134 | 0.18347 | -0.089788 | 0.180384 | -0.288583 | -0.383682 | 0.131101 | 0.776021 | 0.706014 | 5.00752 | 0.141632 |
| 37.4097 | 1.519505 | 1.915979 | 1.55908 | 1.382907 | -0.24482 | -0.507124 | 2.100864 | -1.748047 | -2.122311 | 1.173936 | 5.525443 | 0.196233 | 0.015592 | 0.154705 | 0.158859 | 0.136021 | -0.24646 | -0.293364 | 0.094854 | 0.654641 | 0.344422 | 3.082433 | 0.10806 |
| 37.4236 | 3.78593 | 4.753197 | 3.143579 | 3.167036 | 1.901743 | -2.013527 | 5.127872 | -0.241323 | -0.735554 | 3.811873 | 15.936924 | 0.333012 | 0.181004 | 0.24077 | 0.459764 | 0.197491 | -0.104258 | -0.05387 | 0.307125 | 0.774814 | 0.003684 | 1.893406 | 0.741828 |
| 37.4375 | 4.284682 | 8.498719 | 4.172278 | 6.129139 | 2.901224 | -2.291714 | 7.242361 | 0.794714 | 0.109072 | 5.536658 | 17.763444 | 0.883966 | 1.593552 | 0.50218 | 1.227505 | 0.483726 | -0.042784 | 0.311363 | 1.564818 | 1.507772 | 0.35149 | 1.048784 | 5.121255 |
| 37.4514 | 4.767511 | 5.301027 | 3.227737 | 4.400778 | 3.541285 | -0.252553 | 6.510642 | 1.541082 | 1.877454 | 3.110337 | 9.067596 | 1.870955 | 4.220654 | 0.658268 | 3.056674 | 0.937984 | -0.175106 | 0.481054 | 4.224498 | 3.034413 | 1.270143 | 0.5259 | 13.912867 |
| 37.4653 | 3.650203 | 4.322904 | 2.427817 | 1.867617 | 3.119632 | 0.583406 | 3.757825 | 3.023488 | 4.316927 | 1.432091 | 4.207603 | 1.945611 | 4.453464 | 0.000243 | 3.69098 | 0.792405 | -0.824317 | 0.481657 | 5.834616 | 3.787928 | 1.761619 | -0.006809 | 15.21083 |
| 37.4792 | 2.964335 | 3.294144 | 2.102916 | 0.878657 | 4.322257 | 0.393118 | 2.706727 | 2.984367 | 5.545267 | 1.256066 | 4.165049 | 2.150611 | 2.421129 | 0.873872 | 2.513363 | 1.125932 | 0.384982 | 1.217465 | 3.915245 | 3.235293 | 2.92408 | 3.612687 | 8.950058 |
| 37.4931 | 2.945872 | 2.847873 | 1.92132 | 0.560323 | 5.883757 | 0.311361 | 2.341501 | 2.376726 | 3.534031 | 1.500709 | 4.325684 | 1.888149 | 1.270741 | 1.080676 | 1.709022 | 1.384165 | 0.696518 | 0.991378 | 1.451083 | 2.165815 | 4.027865 | 4.467261 | 2.990042 |
| 37.5 | 2.870122 | 2.71977 | 1.573406 | 0.784321 | 7.205378 | 0.514634 | 2.289656 | 1.222093 | 1.633988 | 1.524155 | 3.486437 | 0.911428 | 1.480845 | 0.620948 | 1.419848 | 0.852845 | 0.604175 | 0.837773 | 0.527156 | 1.994918 | 3.20939 | 8.060835 | 1.129525 |
| 37.5139 | 2.610145 | 2.636372 | 1.243285 | 1.147251 | 6.500893 | 1.197416 | 2.240893 | 0.291105 | 0.974833 | 1.323287 | 2.934965 | 0.369585 | 2.85869 | 0.469822 | 0.992126 | 0.657744 | 0.425619 | 0.690941 | 0.318781 | 2.780201 | 1.713615 | 6.625849 | 0.838155 |
| 37.5278 | 2.206359 | 2.282795 | 0.958595 | 1.514339 | 5.056666 | 1.786618 | 2.101442 | -0.936965 | -0.754547 | 1.029418 | 2.791991 | 0.04305 | 2.697882 | 0.624488 | 0.48956 | 0.813835 | 0.294711 | 0.235472 | 0.249128 | 2.587513 | 0.929148 | 5.965125 | 1.668377 |
| 37.5417 | 1.709867 | 1.883058 | 0.809282 | 1.527239 | 4.200567 | 2.151898 | 1.92055 | -1.730577 | -2.609588 | 0.786894 | 1.970058 | -0.057454 | 0.82018 | 0.781905 | 0.037803 | 0.712025 | 0.256138 | -0.420046 | 0.201647 | 1.53297 | -0.528533 | 3.057099 | 0.861743 |
| 37.5556 | 1.244374 | 1.580451 | 0.762643 | 1.355737 | 3.261911 | 2.215502 | 1.74644 | -1.904065 | -4.036934 | 0.516507 | 0.809468 | -0.044843 | -1.620329 | 0.816488 | -0.125109 | 0.524716 | 0.233121 | -0.697922 | 0.252367 | 0.85312 | -1.428125 | 0.791587 | -0.309839 |
| 37.5694 | 1.232268 | 1.360861 | 0.632806 | 1.143185 | 2.751481 | 1.82188 | 1.569966 | -2.164068 | -4.174998 | 0.292954 | -0.617613 | 0.145293 | -3.541157 | 0.573041 | -0.022608 | 0.104332 | 0.296626 | -0.722982 | 0.334037 | 0.53599 | -2.183176 | -0.941191 | -1.282524 |
| 37.5833 | 0.900791 | 1.289209 | 0.422282 | 0.903871 | 2.457528 | 1.186523 | 1.314676 | -2.346831 | -3.905741 | 0.109825 | -1.605488 | 0.375818 | -3.372252 | 0.450712 | 0.209843 | 0.059302 | 0.478875 | -0.519743 | 0.33669 | 0.441517 | -3.126858 | -4.337453 | -0.565388 |
| 37.5972 | 0.776839 | 1.319603 | 0.416633 | 0.613946 | 1.825971 | 0.176002 | 1.0766 | -2.735106 | -3.274641 | 0.022308 | -2.392714 | 0.414184 | -2.002078 | 0.688688 | 0.31065 | 0.380683 | 0.634429 | 0.007486 | 0.412711 | 0.365019 | -2.526368 | -6.788988 | 0.087781 |
| 37.6111 | 0.528116 | 1.32139 | 0.498697 | 0.306398 | 0.609463 | -0.856019 | 0.928708 | -3.322777 | -2.911195 | 0.067105 | -2.550647 | 0.290288 | -0.581813 | 0.894434 | 0.332543 | 0.625136 | 0.728197 | 0.280887 | 0.431384 | -0.055468 | -3.576253 | -13.786211 | -0.215498 |
| 37.625 | 0.148077 | 1.106166 | 0.538392 | -0.05548 | -0.937436 | -1.736628 | 0.855523 | -4.426211 | -2.868554 | 0.077734 | -2.871432 | 0.062471 | -0.486663 | 0.91662 | 0.253994 | 0.681222 | 0.656405 | 0.059171 | 0.347204 | -0.819999 | -4.307523 | -13.361356 | -0.719946 |
| 37.6389 | 0.036477 | 0.869814 | 0.39766 | -0.554401 | -2.243018 | -2.131447 | 0.62459 | -5.57298 | -2.295348 | 0.061113 | -3.412781 | -0.314807 | -0.758175 | 0.642385 | -0.045652 | 0.61447 | 0.396241 | -0.474366 | 0.22599 | -1.702702 | -3.606661 | -8.634287 | -0.762007 |
| 37.6528 | 0.128688 | 0.721925 | -0.099292 | -0.742702 | -3.264916 | -0.981581 | 0.089761 | -6.249658 | -1.919181 | 0.020074 | -3.671118 | -0.788056 | -1.069097 | 0.317535 | -0.391904 | 0.402914 | 0.049139 | -0.950406 | 0.150064 | -2.481572 | -0.498036 | -1.051997 | -0.762819 |
| 37.6667 | 0.144963 | 0.688501 | -0.874283 | -0.612714 | -3.68504 | -0.958464 | -0.681858 | -5.885144 | -0.957447 | -0.22095 | -2.941724 | -0.907509 | -1.109151 | 0.097405 | -0.482821 | 0.094328 | -0.09925 | -1.142062 | 0.188231 | -2.823513 | 1.574526 | 3.477885 | -0.49095 |
| 37.6806 | 0.242213 | 0.613942 | -1.131451 | -0.466023 | -4.566375 | -0.874979 | -1.12211 | -4.446719 | -0.441208 | -0.642881 | -2.444572 | -0.743098 | 0.521763 | 0.075379 | -0.236483 | -0.054338 | -0.040836 | -0.780555 | 0.283968 | -2.858777 | 1.974393 | 14.502017 | 0.075237 |
| 37.6944 | 0.085588 | 0.44483 | -0.965265 | -0.523267 | -5.567403 | -0.269762 | -0.955711 | -2.903585 | 0.138274 | -0.904255 | -1.95362 | -0.562264 | 1.870322 | 0.12736 | 0.131709 | -0.102948 | 0.033015 | -0.273424 | 0.359937 | -2.657153 | 0.31983 | 13.304895 | 1.522092 |
| 37.7083 | -0.344081 | 0.263988 | -0.812964 | -0.502032 | -5.556013 | -0.175965 | -0.910733 | -1.620677 | 0.112223 | -1.309043 | -1.544275 | -0.457252 | 3.615348 | 0.167354 | 0.326607 | -0.301613 | 0.045276 | 0.076284 | 0.343581 | -2.269324 | 0.682122 | 4.379894 | 2.233942 |
| 37.7222 | -0.644584 | 0.103265 | -0.806948 | 0.043716 | -5.101997 | -0.087912 | -0.691262 | -1.095127 | -0.528789 | -1.617409 | -1.206554 | -0.516954 | 1.575453 | 0.142308 | 0.126941 | -0.439722 | -0.037342 | -0.002911 | 0.240331 | -1.823591 | 0.937434 | 1.205736 | 0.555988 |
| 37.7361 | -0.565223 | -0.146649 | -0.857423 | 1.106788 | -4.610043 | -0.032248 | -0.066349 | -1.211811 | -1.694903 | -1.893047 | -1.002723 | -0.808158 | -2.046603 | 0.059375 | -0.237364 | -0.61819 | -0.161383 | -0.369657 | 0.067934 | -1.835431 | 1.217605 | -1.33263 | -1.909321 |
| 37.75 | -0.71522 | -0.465254 | -0.800788 | 1.786566 | -3.681761 | -0.019247 | 0.326074 | -1.846707 | -3.100012 | -2.197059 | -1.22819 | -1.08886 | -3.187943 | -0.067358 | -0.820721 | -0.942921 | -0.326943 | -0.66335 | -0.117479 | -1.901616 | 0.538362 | -8.090059 | -3.071335 |
| 37.7639 | -0.839321 | -0.793204 | -0.820791 | 2.015351 | -2.701349 | -0.049759 | 0.284993 | -2.59014 | -4.114302 | -2.082212 | -1.562709 | -1.336522 | -1.980828 | -0.178921 | -1.2971 | -0.907028 | -0.460669 | -0.660902 | -0.191959 | -1.711869 | -1.688983 | -6.055394 | -1.985486 |
| 37.7778 | -1.080297 | -1.088144 | -0.933001 | 2.07521 | -2.222553 | -0.122569 | -0.082935 | -3.58725 | -5.018392 | -2.145574 | -1.865547 | -1.288173 | -0.486398 | -0.198923 | -1.358348 | -0.581317 | -0.540983 | -0.372181 | -0.170111 | -1.395924 | -1.364572 | -6.665941 | -1.293438 |
| 37.7917 | -1.655958 | -1.455619 | -0.951624 | 1.997437 | -1.894973 | -0.198085 | -0.607412 | -4.472581 | -5.257247 | -1.914925 | -2.263218 | -1.120889 | -0.38658 | -0.145948 | -1.177406 | -0.187711 | -0.606807 | -0.027807 | -0.135617 | -1.068819 | 0.146246 | -1.025806 | -0.630928 |
| 37.8056 | -1.861337 | -1.653183 | -0.809775 | 1.915954 | -1.671364 | -0.225251 | -1.040379 | -5.378015 | -5.017504 | -2.233347 | -2.486178 | -0.88712 | -0.100918 | -0.158162 | -0.686816 | -0.103437 | -0.604389 | 0.154697 | -0.114551 | -0.676287 | 1.725845 | 3.087049 | -0.130208 |
| 37.8194 | -1.269313 | -1.740027 | -0.568849 | 1.932886 | -1.433693 | -0.239337 | -1.368148 | -6.14924 | -5.079197 | -2.443648 | -2.705805 | -0.791932 | -0.049323 | -0.244428 | -0.066895 | 0.007477 | -0.59249 | 0.122066 | -0.201792 | -0.455015 | 2.73619 | 6.458185 | 0.402105 |
| 37.8333 | -1.082154 | -1.627886 | -0.337723 | 1.96947 | -1.208826 | -0.30184 | -1.570636 | -5.584512 | -4.537404 | -2.301445 | -2.518718 | -0.749056 | 0.391933 | -0.339877 | 0.503953 | 0.108627 | -0.440917 | 0.148579 | -0.333723 | -0.241879 | 2.548562 | 4.6848 | 0.480188 |
| 37.8472 | -0.275403 | -0.916937 | -0.165133 | 1.887389 | -0.994366 | -0.40626 | -1.593837 | -4.995036 | -4.115053 | -2.187951 | -2.109672 | -0.930934 | 1.148662 | -0.457944 | 0.844428 | 0.052706 | -0.214643 | 0.27674 | -0.458973 | -0.000438 | 4.565692 | 4.033706 | 1.038057 |
| 37.8611 | 0.19949 | -0.229191 | -0.012935 | 1.714016 | -0.602821 | -0.477929 | -1.487988 | -4.426326 | -4.06456 | -1.632986 | -1.760882 | -1.056477 | 2.388162 | -0.439703 | 0.967483 | -0.012142 | -0.04699 | 0.360786 | -0.484697 | 0.269251 | 6.735769 | 4.536237 | 1.565654 |
| | | | | | | | | | | | | -1.037941 | 0.727918 | -0.429261 | 1.068268 | -0.082415 | 0.101858 | 0.377054 | -0.382325 | 0.510372 | 2.989907 | 1.784239 | 1.004138 |
| | | | | | | | | | | | | -0.893965 | -0.46633 | -0.366748 | 1.279629 | -0.068309 | 0.179957 | 0.41189 | -0.296922 | 0.719509 | -0.579243 | 2.474829 | -0.015876 |

| | | | | |
|---|---|---|---|---|
| 37.1042 | 1.511558 | | 37.0972 | -0.008774 |
| 37.1181 | 1.925578 | | 37.1111 | -0.272534 |
| 37.1319 | 2.661933 | | 37.125 | -0.268671 |
| 37.1458 | 3.41151 | | 37.1389 | -0.107529 |
| 37.1597 | 4.015856 | | 37.1528 | 0.085615 |
| 37.1736 | 4.223404 | | 37.1667 | 0.219951 |
| 37.1875 | 4.446053 | | 37.1806 | 0.191029 |
| 37.2014 | 3.71623 | | 37.1944 | -0.046683 |
| 37.2153 | 2.770993 | | 37.2083 | -0.156667 |
| 37.2292 | 1.849508 | | 37.2222 | -0.113892 |
| 37.2431 | 1.09992 | | 37.2361 | -0.122379 |
| 37.2569 | 0.901616 | | 37.25 | -0.027098 |
| 37.2708 | 0.977081 | | 37.2639 | 0.052359 |
| 37.2847 | 1.056995 | | 37.2778 | -0.022402 |
| 37.2986 | 1.158017 | | 37.2917 | -0.011248 |
| 37.3125 | 1.1502 | | 37.3056 | -0.034138 |
| 37.3264 | 0.930207 | | 37.3194 | -0.136696 |
| 37.3403 | 0.326254 | | 37.3333 | -0.140449 |
| 37.3542 | -0.361935 | | 37.3472 | 0.029102 |
| 37.3681 | -0.747219 | | 37.3611 | 0.270006 |
| 37.3819 | -0.904591 | | 37.375 | 0.299432 |
| 37.3958 | -0.181694 | | 37.3889 | 0.141632 |
| 37.4097 | 5.525443 | | 37.4028 | 0.10806 |
| 37.4236 | 15.936924 | | 37.4167 | 0.741828 |
| 37.4375 | 17.763444 | | 37.4306 | 5.121255 |
| 37.4514 | 9.067596 | | 37.4444 | 13.912867 |
| 37.4653 | 4.207603 | | 37.4583 | 15.21083 |
| 37.4792 | 4.165049 | | 37.4722 | 8.950058 |
| 37.4931 | 4.325684 | | 37.4861 | 2.990042 |
| 37.5 | 3.486437 | | 37.5 | 1.129525 |
| 37.5139 | 2.934965 | | 37.5139 | 0.838155 |
| 37.5278 | 2.791991 | | 37.5278 | 1.668377 |
| 37.5417 | 1.970058 | | 37.5417 | 0.861743 |
| 37.5556 | 0.809468 | | 37.5556 | -0.309839 |
| 37.5694 | -0.617613 | | 37.5694 | -1.282524 |
| 37.5833 | -1.605488 | | 37.5833 | -0.565388 |
| 37.5972 | -2.392714 | | 37.5972 | 0.087781 |
| 37.6111 | -2.550647 | | 37.6111 | -0.215498 |
| 37.625 | -2.871432 | | 37.625 | -0.719946 |
| 37.6389 | -3.412781 | | 37.6389 | -0.762007 |
| 37.6528 | -3.671118 | | 37.6528 | -0.762819 |
| 37.6667 | -2.941724 | | 37.6667 | -0.49095 |
| 37.6806 | -2.444572 | | 37.6806 | 0.075237 |
| 37.6944 | -1.95362 | | 37.6944 | 1.522092 |
| 37.7083 | -1.544275 | | 37.7083 | 2.233942 |
| 37.7222 | -1.206554 | | 37.7222 | 0.555988 |
| 37.7361 | -1.002723 | | 37.7361 | -1.909321 |
| 37.75 | -1.22819 | | 37.75 | -3.071335 |
| 37.7639 | -1.562709 | | 37.7639 | -1.985486 |
| 37.7778 | -1.865547 | | 37.7778 | -1.293438 |
| 37.7917 | -2.263218 | | 37.7917 | -0.630928 |
| 37.8056 | -2.486178 | | 37.8056 | -0.130208 |
| 37.8194 | -2.705805 | | 37.8194 | 0.402105 |
| 37.8333 | -2.518718 | | 37.8333 | 0.480188 |
| 37.8472 | -2.109672 | | 37.8472 | 1.038057 |
| 37.8611 | -1.760882 | | 37.8611 | 1.565654 |
| | | | 37.875 | 1.004138 |
| | | | 37.8889 | -0.015876 |

**Supplementary Table S7 (b)** Direct motion in 100 mM NaCl stressed lettuce and their respective controls under dimG day/night lighting conditions

| Time | dimGcontrol1 | dimGcontrol2 | dimGcontrol3 | dimGcontrol4 | dimGcontrol5 | dimGcontrol6 | dimGcontrol7 | dimGcontrol8 | dimGcontrol9 | dimGcontrolnoplar | dimGcontroltank | dimGStress1 | dimGStress2 | dimGStress3 | dimGStress4 | dimGStress5 | dimGStress6 | dimGStress7 | dimGStress8 | dimGStress9 | dimGStress10 | dimGStressnoplan | dimGstresstank | | Time | Mean dimGL_control | | Time | Mean dimGL_stress |
|---|---|---|---|---|---|---|---|---|---|---|---|---|---|---|---|---|---|---|---|---|---|---|---|---|---|---|---|---|---|
| 24 | -0.521689 | -1.889034 | -1.392723 | -1.091552 | -0.342768 | -1.576081 | -0.374496 | -0.096819 | -0.741038 | 0.025927 | -1.266965 | -3.289443 | -1.319695 | -0.759037 | -0.320991 | -0.26295 | -0.276093 | -0.795027 | -0.197411 | 0.151016 | -1.279097 | 0.066069 | -2.055841 | | 24.01 | -1.266965 | | 24 | -2.055841 |
| 24.013889 | -0.238862 | -1.350883 | -1.076383 | -0.633803 | -0.224004 | -0.940478 | -0.273078 | 0.101338 | -0.393197 | 0.024712 | -0.529204 | -1.982778 | -0.845516 | -0.323452 | -0.149662 | -0.170274 | -0.290734 | -0.428551 | -0.114674 | 0.004824 | -1.283453 | 0.023812 | -1.461978 | | 24.02 | -0.529204 | | 24.014 | -1.461978 |
| 24.027778 | -0.070358 | -0.8412 | -0.5981 | -0.231807 | -0.105076 | -0.237654 | -0.061933 | 0.065978 | -0.101413 | 0.149218 | -0.162022 | -0.493348 | -0.23695 | 0.051291 | -0.069163 | -0.031882 | -0.282483 | -0.163006 | 0.066746 | -0.155596 | -0.821581 | 0.106662 | -0.92674 | | 24.03 | -0.162022 | | 24.028 | -0.92674 |
| 24.041667 | 0.27139 | -0.468568 | -0.174662 | 0.119035 | -0.038562 | 0.136617 | 0.153373 | -0.085343 | 0.08013 | 0.182894 | -0.129837 | 0.020227 | -0.086233 | 0.370545 | 0.002025 | 0.029053 | -0.380421 | -0.181739 | 0.056393 | -0.34931 | -0.432068 | 0.019082 | -0.620468 | | 24.05 | -0.129837 | | 24.042 | -0.620468 |
| 24.055556 | 0.839268 | 0.017055 | -0.145389 | 0.440502 | -0.034951 | 0.413334 | 0.170784 | -0.392866 | -0.039085 | 0.096721 | -0.034039 | -0.669323 | -1.01 | -0.180684 | -0.64276 | -0.168823 | -1.603289 | -1.036819 | -0.808912 | -0.886768 | -0.937422 | -0.350312 | -2.31686 | | 24.06 | -0.034039 | | 24.056 | -2.31686 |
| 24.069444 | 1.78478 | 1.267036 | -0.298294 | 0.952348 | 0.407546 | 1.047033 | 0.158933 | -0.617251 | -0.357237 | 2.340926 | 0.841507 | -3.130448 | -3.164918 | -2.331743 | -2.856621 | -0.860338 | -5.12048 | -3.278867 | -2.806457 | -2.399389 | -2.701773 | -1.010206 | -6.787287 | | 24.08 | 0.841507 | | 24.069 | -6.787287 |
| 24.083333 | 3.009239 | 3.074012 | -0.122316 | 1.649915 | 1.44551 | 2.009328 | 0.41883 | -0.281363 | -0.159963 | 10.742011 | 2.267131 | -3.26364 | -2.940436 | -3.485135 | -4.236913 | -1.655769 | -4.145098 | -2.922388 | -3.039524 | -2.286402 | -3.082708 | -1.310687 | -8.240116 | | 24.09 | 2.267131 | | 24.083 | -8.240116 |
| 24.097222 | 2.78679 | 3.0579 | 0.25126 | 1.905341 | 1.763352 | 2.123693 | 0.531716 | 0.413213 | 0.675821 | NaN | 2.699345 | -0.454755 | -0.571441 | -1.125795 | -2.08442 | -0.906495 | -1.704291 | -1.130388 | -1.194429 | -1.047482 | -0.665866 | -0.887571 | -3.752076 | | 24.1 | 2.699345 | | 24.097 | -3.752076 |
| 24.111111 | 1.387258 | 1.350063 | 0.491065 | 1.506838 | 1.081911 | 2.218813 | 0.437322 | 0.807487 | 1.618184 | -0.145824 | 1.908073 | 1.176376 | 0.677241 | 0.016611 | 0.688188 | 0.119089 | 1.78854 | 1.102097 | 0.747808 | 0.561928 | 1.030888 | -0.144991 | 0.883943 | | 24.12 | 1.908073 | | 24.111 | 0.883943 |
| 24.125 | 0.590878 | 0.480252 | 0.524411 | 0.929142 | 0.518562 | 1.603648 | 0.418497 | 0.638054 | 1.655934 | 0.224697 | 1.381518 | 0.900659 | 0.424404 | -0.05041 | 1.197227 | 0.451618 | 2.131942 | 1.170038 | 0.781214 | 0.590803 | 0.938333 | 0.403527 | 1.437682 | | 24.13 | 1.381518 | | 24.125 | 1.437682 |
| 24.138889 | 0.379665 | 0.346715 | 0.391261 | 0.324001 | 0.158618 | 0.970586 | 0.287381 | 0.262651 | 0.778298 | 0.058424 | 1.043397 | -0.137983 | -0.323091 | -0.701254 | 0.068703 | 0.073802 | -0.878974 | -0.565921 | -0.336447 | -0.542705 | 0.088782 | 0.200794 | -0.324064 | | 24.15 | 1.043397 | | 24.139 | -0.324064 |
| 24.152778 | 0.157645 | 0.093264 | 0.068146 | -0.130151 | 0.035675 | 0.549508 | 0.029214 | 0.018447 | 0.253908 | 0.090598 | 0.462607 | -0.65464 | -0.610809 | -0.709636 | -0.636383 | -0.167461 | -2.44341 | -1.502995 | -0.695233 | -1.046084 | -0.481318 | -0.038323 | -1.281107 | | 24.16 | 0.462607 | | 24.153 | -1.281107 |
| 24.166667 | 0.06533 | -0.05507 | -0.162467 | -0.34433 | 0.11879 | 0.424241 | -0.128712 | 0.022892 | 0.031546 | 0.148496 | 0.023351 | -0.533154 | -0.433451 | -0.503425 | -0.449234 | -0.09931 | -1.58035 | -0.93479 | -0.435599 | -0.760042 | -0.485338 | -0.039025 | -1.022531 | | 24.17 | 0.023351 | | 24.167 | -1.022531 |
| 24.180556 | 0.349325 | 0.041217 | -0.058781 | -0.174086 | 0.218824 | 0.328955 | -0.026199 | 0.10235 | 0.07497 | 0.20882 | 0.016478 | -0.265807 | -0.210879 | -0.433822 | 0.21101 | -0.056286 | -0.285265 | -0.326061 | -0.26958 | -0.465528 | -0.347839 | -0.010458 | -0.414175 | | 24.19 | 0.016478 | | 24.181 | -0.414175 |
| 24.194444 | 0.736299 | 0.210027 | 0.173969 | 0.216591 | 0.210062 | 0.192857 | 0.168742 | 0.114195 | 0.243271 | 0.08552 | -0.008039 | -0.108387 | -0.113853 | -0.48529 | 0.677855 | -0.100905 | 0.187372 | -0.195804 | -0.266894 | -0.359446 | -0.324409 | -0.069366 | -0.144062 | | 24.2 | -0.008039 | | 24.194 | -0.144062 |
| 24.208333 | 0.743418 | 0.222018 | 0.255856 | 0.369319 | 0.089152 | -0.130377 | 0.182989 | 0.022838 | 0.328037 | -0.096524 | -0.239751 | -0.17443 | -0.186344 | -0.611782 | 0.638116 | -0.181487 | -0.030129 | -0.349439 | -0.374872 | -0.148647 | -0.422242 | -0.138159 | -0.330112 | | 24.22 | -0.239751 | | 24.208 | -0.330112 |
| 24.222222 | 0.321633 | 0.082409 | 0.050102 | 0.168013 | 0.013977 | -0.41883 | 0.015934 | -0.025781 | 0.254087 | -0.212619 | -0.385027 | -0.258716 | -0.269155 | -0.610933 | 0.522317 | -0.221207 | -0.376051 | -0.424768 | -0.433404 | 0.294418 | -0.500907 | -0.190004 | -0.278724 | | 24.23 | -0.385027 | | 24.222 | -0.278724 |
| 24.236111 | -0.205164 | -0.069442 | -0.215477 | -0.154689 | -0.002263 | -0.655818 | -0.134465 | 0.00834 | 0.188967 | -0.161923 | -0.324965 | -0.184706 | -0.218239 | -0.489343 | 0.447367 | -0.127698 | -0.498807 | -0.35184 | -0.344923 | 0.750024 | -0.457399 | -0.173316 | 0.001102 | | 24.24 | -0.324965 | | 24.236 | 0.001102 |
| 24.25 | -0.423627 | -0.061474 | -0.334032 | -0.375688 | -0.10873 | -0.604059 | -0.247956 | 0.061312 | 0.190902 | -0.168034 | -0.13341 | -0.062435 | -0.144993 | -0.166873 | 0.327591 | -0.055924 | -0.506293 | -0.28069 | -0.248698 | 0.673466 | -0.428929 | -0.100505 | 0.001332 | | 24.26 | -0.13341 | | 24.25 | 0.001332 |
| [illegible] | | | | | | | | | | | | | | | | | | | | | | | | | | | | | |

| | | | | | | | | | | | | | | | | | | | | | | | | | | | | |
|---|---|---|---|---|---|---|---|---|---|---|---|---|---|---|---|---|---|---|---|---|---|---|---|---|---|---|---|---|
| 27.027778 | -0.250259 | 0.061027 | -0.194442 | -0.308309 | 0.342635 | 0.160026 | -0.188668 | -0.234484 | -1.340163 | 0.021846 | -0.415828 | 0.983945 | 0.726659 | 0.213976 | 0.806224 | 0.062497 | 0.925386 | 0.965786 | 0.38126 | 0.328828 | 0.184748 | -0.045109 | 0.276141 | | 27.03 | -0.415828 | 27.028 | 0.276141 |
| 27.041667 | 0.298263 | 1.134499 | 0.315788 | 0.01736 | 0.652572 | 0.676509 | 0.126118 | 0.095788 | -1.738151 | 0.455425 | 1.649814 | -0.01676 | 0.233765 | -0.090309 | 0.964345 | -0.140655 | 0.95661 | 0.754899 | 0.312001 | 0.982422 | 0.669091 | -0.975695 | -1.996178 | | 27.05 | 1.649814 | 27.042 | -1.996178 |
| 27.055556 | 0.926587 | 1.931628 | 0.578995 | 0.251218 | 1.327597 | 1.344705 | 0.582999 | 0.75556 | -0.860684 | 3.218203 | 5.482765 | 0.868952 | 0.824772 | -0.037063 | 0.902538 | -0.076068 | 0.764873 | 0.521802 | 0.266428 | 0.793559 | 0.287298 | 4.528555 | -1.425845 | | 27.06 | 5.482765 | 27.056 | -1.425845 |
| 27.069444 | 1.551851 | 2.589643 | 0.735731 | 0.247171 | 1.482757 | 0.474731 | 0.722897 | 1.472265 | -0.406556 | 2.200572 | 8.989161 | 0.554261 | 0.195349 | -0.065792 | 0.746267 | -0.042206 | 0.533696 | 0.261279 | 0.459342 | 0.166039 | -0.240681 | -0.390904 | -0.314328 | | 27.08 | 8.989161 | 27.069 | -0.314328 |
| 27.083333 | -0.067279 | -0.256364 | 2.406926 | 0.938483 | -0.810073 | -3.518572 | 0.47653 | -1.231079 | -0.001953 | NaN | 1.65021 | 0.375406 | -0.181082 | -0.04943 | 0.758872 | 0.081789 | 0.442289 | 0.154916 | 1.024066 | -0.169149 | -0.510553 | -1.199327 | 0.284698 | | 27.09 | 1.65021 | 27.083 | 0.284698 |
| 27.097222 | -1.825208 | -2.507622 | 0.029512 | 0.942081 | -0.522668 | -1.998154 | -0.189122 | -2.190439 | 0.895438 | -0.542562 | -5.45241 | 0.118579 | 0.258188 | 0.18824 | 0.99458 | 0.312782 | 0.710914 | 0.339427 | 1.410165 | 0.210703 | -0.068152 | -0.291513 | 0.274324 | | 27.1 | -5.45241 | 27.097 | 0.274324 |
| 27.111111 | -0.828322 | -0.962811 | 0.076711 | 1.081985 | 0.442486 | 1.270206 | 0.467853 | -0.613908 | 2.02873 | -0.419866 | -3.007591 | 0.557095 | 0.81159 | 0.403512 | 1.070588 | 0.498447 | 0.848072 | 0.540581 | 1.399742 | 0.53522 | 0.517681 | 0.96227 | 0.15611 | | 27.12 | -3.007591 | 27.111 | 0.15611 |
| 27.125 | 0.30058 | 0.09067 | -0.587214 | 1.086531 | 1.070639 | 2.057738 | 1.056344 | 0.558146 | 2.279224 | 0.307904 | 2.661337 | 0.346918 | 0.286871 | 0.291668 | 0.528308 | 0.395565 | 0.377393 | 0.348145 | 0.917247 | 0.000863 | 0.335251 | 0.920969 | 0.109969 | | 27.13 | 2.661337 | 27.125 | 0.109969 |
| 27.138889 | 0.435564 | 0.249954 | 0.260334 | 0.745531 | 0.938428 | 0.672257 | 0.743803 | 0.447845 | 2.034983 | 0.274679 | 1.564095 | -0.736886 | -0.645156 | 0.027014 | -0.361266 | 0.136168 | -0.207937 | -0.061329 | 0.149122 | -0.607089 | -0.208597 | 0.033288 | 0.020946 | | 27.15 | 1.564095 | 27.139 | 0.020946 |
| 27.152778 | 0.088886 | -0.051198 | 0.612875 | 0.198593 | 0.494948 | 0.303085 | 0.656259 | 0.676987 | 2.018496 | -0.045816 | 1.0881 | -0.704791 | -0.778695 | -0.062254 | -0.851923 | -0.005404 | -0.428878 | -0.272766 | -0.256871 | -0.455607 | -0.385081 | -0.479172 | -0.060807 | | 27.16 | 1.0881 | 27.153 | -0.060807 |
| 27.166667 | -0.66755 | -1.001216 | 1.323161 | -0.219117 | -0.38916 | 0.27502 | 0.818254 | 1.534068 | 3.4677 | -0.161074 | 1.186689 | 0.161634 | 0.101575 | 0.196149 | -0.599886 | 0.138448 | -0.18658 | -0.078515 | -0.077739 | 0.168508 | -0.049155 | -0.371542 | 0.257996 | | 27.17 | 1.186689 | 27.167 | 0.257996 |
| 27.180556 | -0.883858 | -1.395165 | 1.74406 | 0.054756 | -1.153328 | 0.459748 | 0.969138 | 1.407148 | 3.591431 | NaN | 1.546052 | 1.387462 | 1.550456 | 0.678711 | 0.085844 | 0.551659 | 0.537866 | 0.51186 | 0.487176 | 0.923189 | 0.822876 | 0.831549 | 0.566245 | | 27.19 | 1.546052 | 27.181 | 0.566245 |
| 27.194444 | -0.026557 | -0.329694 | 1.486814 | 0.96525 | -0.030387 | 0.4073 | 0.942107 | 0.75996 | 2.14154 | -3.036991 | 1.709433 | 1.486327 | 1.921098 | 0.734758 | 0.456729 | 0.559052 | 0.90123 | 0.80774 | 0.743765 | 0.916102 | 1.144853 | 1.800495 | 0.282415 | | 27.2 | 1.709433 | 27.194 | 0.282415 |
| 27.208333 | 0.659447 | 0.526908 | 1.348972 | 1.439311 | 0.157215 | 0.259773 | 0.843286 | 0.316124 | 1.121003 | -0.483986 | 1.578883 | 0.305681 | 0.816959 | 0.357088 | 0.147616 | 0.214861 | 0.539451 | 0.506929 | 0.29488 | 0.065353 | 0.456602 | 1.05726 | -0.057393 | | 27.22 | 1.578883 | 27.208 | -0.057393 |
| 27.222222 | 0.845438 | 0.798643 | 0.877769 | 0.96167 | -0.220306 | 0.091979 | 0.41117 | 0.059776 | 0.49619 | 0.01932 | 0.87557 | -0.614306 | -0.092195 | 0.112812 | -0.256709 | -0.085038 | 0.088099 | 0.07295 | -0.369732 | -0.369357 | -0.07104 | 0.233923 | -0.201042 | | 27.23 | 0.87557 | 27.222 | -0.201042 |
| 27.236111 | 0.570583 | 0.574262 | 0.140903 | 0.014548 | -0.582605 | -0.074128 | -0.190088 | -0.036259 | -0.086163 | 0.077417 | -0.301154 | -0.690676 | -0.241822 | 0.068639 | -0.489678 | -0.262443 | -0.198169 | -0.23318 | -0.798018 | -0.276966 | -0.061674 | -0.013471 | -0.336655 | | 27.24 | -0.301154 | 27.236 | -0.336655 |
| 27.25 | 0.100665 | 0.02345 | -0.214924 | -0.55194 | -0.66549 | -0.166036 | -0.547984 | -0.047649 | -0.507262 | 0.0336 | -1.118434 | -0.083858 | 0.067706 | 0.074468 | -0.560544 | -0.293163 | -0.341516 | -0.401317 | -0.786623 | -0.074549 | 0.017197 | -0.080235 | -0.391848 | | 27.26 | -1.118434 | 27.25 | -0.391848 |
| 27.263889 | -0.161952 | -0.344215 | -0.111406 | -0.328262 | -0.458945 | -0.127812 | -0.324294 | 0.032539 | -0.368907 | -0.241684 | -1.080055 | 0.793605 | 0.58973 | 0.102601 | -0.461191 | -0.129918 | -0.309795 | -0.256015 | -0.335348 | 0.073177 | -0.009219 | -0.049568 | -0.202047 | | 27.27 | -1.080055 | 27.264 | -0.202047 |
| 27.277778 | -0.006568 | -0.293169 | 0.29411 | 0.375833 | -0.372379 | 0.017109 | 0.25789 | 0.188673 | 0.115505 | -0.558369 | -0.218114 | 1.200657 | 0.828833 | 0.103426 | -0.335371 | 0.044752 | -0.194334 | 0.000108 | 0.20839 | 0.123209 | -0.09047 | -0.091485 | 0.141971 | | 27.28 | -0.218114 | 27.278 | 0.141971 |
| 27.291667 | 0.408725 | 0.189054 | 0.696631 | 1.048651 | -0.049864 | 0.317625 | 0.773525 | 0.334878 | 0.705698 | -0.079524 | 1.243358 | 0.744684 | 0.648825 | 0.07426 | -0.244473 | 0.147437 | -0.062659 | 0.246992 | 0.358215 | -0.027717 | -0.128645 | -0.169036 | 0.37325 | | 27.3 | 1.243358 | 27.292 | 0.37325 |
| 27.305556 | 0.696441 | 0.583129 | 0.766528 | 1.185372 | 0.293353 | 0.454369 | 0.867815 | 0.3459 | 1.121613 | 0.492519 | 2.011096 | 0.233687 | 0.409125 | 0.08117 | -0.024191 | 0.190728 | 0.069387 | 0.175644 | 0.199858 | -0.131053 | -0.0494 | -0.093191 | 0.351769 | | 27.31 | 2.011096 | 27.306 | 0.351769 |
| [illegible] | [illegible] | [illegible] | [illegible] | [illegible] | [illegible] | [illegible] | [illegible] | [illegible] | [illegible] | [illegible] | [illegible] | [illegible] | [illegible] | [illegible] | [illegible] | [illegible] | [illegible] | [illegible] | [illegible] | [illegible] | [illegible] | [illegible] | [illegible] | | [illegible] | [illegible] | [illegible] | [illegible] |

| | | | | | | | | | | | | | | | | | | | | | | | | | | | | |
|---|---|---|---|---|---|---|---|---|---|---|---|---|---|---|---|---|---|---|---|---|---|---|---|---|---|---|---|---|
| 30.083333 | 2.22207 | 1.828268 | 0.508948 | 1.818965 | 0.769755 | 0.682379 | 2.233018 | 0.109317 | 0.692229 | 0.306477 | 1.728451 | 0.140599 | 0.254716 | 2.139221 | 0.638478 | 0.870344 | 0.450777 | 0.444845 | 0.676198 | 0.117414 | 0.109254 | 0.006625 | 0.241514 | | 30.09 | 1.728451 | | 30.083 | 0.241514 |
| 30.097222 | 1.989351 | 1.794003 | 0.657163 | 1.507571 | 0.595532 | 0.600411 | 1.970211 | 0.07247 | 0.996667 | 0.196636 | 1.329324 | 0.319238 | 0.443278 | 2.627791 | 0.634831 | 0.913741 | 0.30424 | 0.257204 | 0.669781 | 0.115319 | 0.088214 | 0.039031 | 0.254571 | | 30.1 | 1.329324 | | 30.097 | 0.254571 |
| 30.111111 | 1.855956 | 1.676969 | 0.84205 | 0.981277 | 0.833862 | 0.464613 | 1.362054 | -0.049519 | 0.709605 | -0.037307 | 1.094319 | 0.143438 | 0.557773 | 2.475316 | 0.621971 | 0.900159 | 0.092873 | 0.092425 | 0.519414 | 0.087841 | 0.135599 | -0.006076 | 0.226196 | | 30.12 | 1.094319 | | 30.111 | 0.226196 |
| 30.125 | 2.41273 | 1.378586 | 1.066193 | 0.451986 | 0.951397 | 0.733107 | 0.776675 | 0.102751 | 0.784873 | 0.138528 | 1.753815 | -0.138359 | 0.527415 | 1.999744 | 0.511166 | 0.828549 | -0.087925 | 0.124865 | 0.361969 | 0.101376 | 0.231481 | -0.019298 | 0.234098 | | 30.13 | 1.753815 | | 30.125 | 0.234098 |
| 30.138889 | 2.254738 | 0.396615 | 1.422982 | -0.500294 | 0.975056 | 1.085511 | 0.08221 | 0.545382 | 1.617136 | 0.496434 | 2.425936 | -0.190683 | 0.566592 | 1.170753 | 0.33564 | 0.706868 | -0.088932 | 0.289798 | 0.292287 | 0.189481 | 0.246821 | 0.030913 | 0.275529 | | 30.15 | 2.425936 | | 30.139 | 0.275529 |
| 30.152778 | 2.091375 | 0.039262 | 1.941568 | -0.486996 | 0.857242 | 1.138308 | -0.089174 | 0.882316 | 1.869804 | -0.342295 | 2.326145 | 0.043544 | 0.708398 | 0.40186 | 0.187197 | 0.469235 | -0.026588 | 0.3676 | 0.148992 | 0.288132 | 0.130009 | 0.021883 | 0.31044 | | 30.16 | 2.326145 | | 30.153 | 0.31044 |
| 30.166667 | 2.319783 | 0.45738 | 2.665435 | 0.490121 | 0.643661 | 0.820091 | 0.18955 | 1.110518 | 1.899858 | 0.873136 | 1.979494 | 0.436257 | 1.104662 | -0.15262 | 0.096727 | 0.202977 | -0.070996 | 0.235781 | -0.052094 | 0.354504 | -0.039825 | -0.043589 | 0.344297 | | 30.17 | 1.979494 | | 30.167 | 0.344297 |
| 30.180556 | 2.041709 | 1.156042 | 3.159423 | 1.182381 | 0.544243 | 0.794201 | 0.459602 | 1.333636 | 1.563888 | 0.18711 | 1.916398 | 0.755481 | 1.393716 | -0.164004 | 0.114344 | 0.006514 | -0.067703 | 0.110895 | -0.155256 | 0.382038 | -0.063599 | 0.019587 | 0.379067 | | 30.19 | 1.916398 | | 30.181 | 0.379067 |
| 30.194444 | 1.795788 | 1.620494 | 3.214287 | 1.409302 | 0.411315 | 0.97342 | 0.880023 | 1.363061 | 1.333601 | -0.026155 | 1.839937 | 0.780518 | 1.38618 | 0.16332 | 0.199177 | -0.019745 | 0.062121 | 0.113938 | -0.029645 | 0.303981 | -0.019341 | 0.144638 | 0.400814 | | 30.2 | 1.839937 | | 30.194 | 0.400814 |
| 30.208333 | 1.377258 | 1.900395 | 2.845568 | 1.227012 | 0.308154 | 0.88283 | 1.191352 | 1.280051 | 1.343577 | NaN | 1.8639 | 0.555878 | 0.887212 | 0.562465 | 0.26282 | 0.060869 | 0.215219 | 0.265004 | 0.214235 | 0.167932 | -0.068812 | 0.133674 | 0.340097 | | 30.22 | 1.8639 | | 30.208 | 0.340097 |
| 30.222222 | 0.554736 | 1.901184 | 2.26019 | 0.762976 | 0.30323 | 0.763044 | 1.106939 | 1.090285 | 1.721339 | NaN | 1.665233 | 0.448893 | 0.491594 | 0.789225 | 0.31089 | 0.167811 | 0.246225 | 0.471951 | 0.484916 | 0.098557 | -0.095741 | 0.08746 | 0.379576 | | 30.23 | 1.665233 | | 30.222 | 0.379576 |
| 30.236111 | -0.187548 | 1.842422 | 1.858614 | 0.258504 | 0.376677 | 0.702483 | 0.833453 | 0.805162 | 2.618805 | NaN | 1.45302 | 0.888725 | 0.602664 | 0.760878 | 0.437781 | 0.237361 | 0.247401 | 0.69276 | 0.878382 | 0.173117 | 0.065529 | 0.192159 | 0.584734 | | 30.24 | 1.45302 | | 30.236 | 0.584734 |
| 30.25 | -0.19299 | 1.440255 | 1.965897 | 0.25002 | 0.585349 | 0.678589 | 0.970463 | 0.655893 | 3.092043 | NaN | 1.163791 | 1.449094 | 0.830646 | 0.357171 | 0.409265 | 0.149395 | 0.313101 | 0.748182 | 0.990518 | 0.271915 | 0.305245 | 0.347871 | 0.612689 | | 30.26 | 1.163791 | | 30.25 | 0.612689 |
| 30.263889 | 0.451132 | 1.013894 | 1.700633 | 0.638058 | 0.798048 | 0.633538 | 1.308338 | 0.933669 | 2.626886 | NaN | 1.123069 | 1.354284 | 0.772948 | 0.046583 | 0.460024 | 0.43273 | 0.373492 | 0.603377 | 0.656339 | 0.288689 | 0.470818 | 0.341829 | 0.443967 | | 30.27 | 1.123069 | | 30.264 | 0.443967 |
| 30.277778 | 0.962266 | 0.860461 | 1.537675 | 0.919989 | 0.962328 | 0.643432 | 1.498462 | 1.325162 | 2.752783 | NaN | 1.250451 | 1.155281 | 0.402395 | 0.297164 | 0.586804 | 0.672366 | 0.298743 | 0.45481 | 0.455597 | 0.265118 | 0.554113 | 0.204684 | 0.271868 | | 30.28 | 1.250451 | | 30.278 | 0.271868 |
| 30.291667 | 1.077052 | 0.852981 | 0.835485 | 0.845517 | 1.066899 | 0.537615 | 1.038575 | 1.31979 | 2.163701 | NaN | 1.233576 | 0.57404 | -0.35954 | 0.687621 | 0.696221 | 0.640431 | 0.088359 | 0.226121 | 0.38162 | 0.219511 | 0.427601 | 0.119841 | 0.170915 | | 30.3 | 1.233576 | | 30.292 | 0.170915 |
| 30.305556 | 0.790903 | 0.759901 | -0.203154 | 0.621382 | 0.943688 | 0.180524 | 0.332895 | 0.632889 | 0.553615 | NaN | 0.774232 | 0.225701 | -0.721011 | 0.970815 | 0.571319 | 0.447765 | -0.069045 | 0.012458 | 0.279184 | 0.155382 | 0.190993 | 0.089246 | 0.116086 | | 30.31 | 0.774232 | | 30.306 | 0.116086 |
| 30.319444 | 0.36617 | 0.411164 | -0.751117 | 0.394458 | 0.673375 | -0.040671 | -0.092593 | -0.147449 | -0.496978 | NaN | 0.156537 | 0.39926 | -0.415456 | 1.046744 | 0.313044 | 0.344257 | -0.002187 | -0.018176 | 0.167604 | 0.090221 | 0.074647 | 0.075786 | 0.163207 | | 30.33 | 0.156537 | | 30.319 | 0.163207 |
| 30.333333 | 0.380171 | 0.12296 | -0.525148 | 0.387061 | 0.352481 | 0.071379 | -0.035782 | -0.411175 | -0.289678 | NaN | 0.107987 | 0.80866 | 0.385546 | 0.58439 | 0.061122 | 0.364438 | 0.145576 | 0.045248 | 0.115446 | 0.038455 | 0.074415 | 0.132153 | 0.288671 | | 30.34 | 0.107987 | | 30.333 | 0.288671 |
| 30.347222 | 1.023584 | 0.057434 | -0.1636 | 0.679111 | 0.142792 | 0.210272 | 0.331949 | -0.132796 | 0.41658 | NaN | 0.531783 | 1.037164 | 1.185636 | 0.16781 | -0.052466 | 0.402144 | 0.254405 | 0.188815 | 0.192998 | 0.030615 | 0.197599 | 0.211684 | 0.383349 | | 30.35 | 0.531783 | | 30.347 | 0.383349 |
| 30.361111 | 1.773742 | 0.243778 | 0.116844 | 1.115963 | 0.1343 | 0.356728 | 0.641734 | 0.095942 | 1.069406 | NaN | 1.108315 | 0.738911 | 1.499589 | -0.184505 | 0.079259 | 0.408215 | 0.301974 | 0.335864 | 0.337152 | 0.074803 | 0.515135 | 0.23756 | 0.433748 | | 30.37 | 1.108315 | | 30.361 | 0.433748 |
| 30.375 | 2.234164 | 0.499216 | 0.304679 | 1.492219 | 0.184148 | 0.397225 | 0.727291 | 0.121636 | 1.593826 | NaN | 1.380656 | 0.422484 | 1.408404 | -0.212799 | 0.274871 | 0.365137 | 0.187651 | 0.297189 | 0.380025 | 0.094809 | 0.856255 | 0.14457 | 0.396911 | | 30.38 | 1.380656 | | 30.375 | 0.396911 |
| 30.388889 | 2.059713 | 0.60695 | 0.235465 | 1.507456 | 0.168367 | 0.255676 | 0.596647 | -0.011093 | 1.277317 | NaN | 1.15203 | 0.183512 | 1.064741 | 0.161775 | 0.4543 | 0.34364 | 0.022756 | 0.124193 | 0.303765 | 0.04338 | 0.928687 | 0.028266 | 0.321966 | | 30.4 | 1.15203 | | 30.389 | 0.321966 |
| 30.402778 | 1.575255 | 0.609082 | 0.094565 | 1.265986 | 0.04566 | 0.071732 | 0.39662 | -0.2845 | 0.236062 | NaN | 0.634068 | 0.24834 | 0.780765 | 0.721897 | 0.505587 | 0.383978 | 0.015067 | 0.03449 | 0.159882 | -0.009954 | 0.70164 | -0.001366 | 0.288626 | | 30.41 | 0.634068 | | 30.403 | 0.288626 |
| 30.416667 | 1.082831 | 0.71364 | 0.095874 | 1.022655 | -0.198427 | 0.048931 | 0.268698 | -0.416821 | -0.77747 | NaN | 0.005736 | 0.524682 | 0.551215 | 1.163619 | 0.415782 | 0.505276 | 0.161175 | 0.107843 | 0.08007 | -0.004599 | 0.301967 | 0.056012 | 0.254079 | | 30.42 | 0.005736 | | 30.417 | 0.254079 |
| 30.430556 | 0.893681 | 0.979219 | 0.207924 | 0.957427 | -0.288959 | 0.041782 | 0.27126 | -0.350195 | -1.357011 | NaN | -0.415426 | 0.626771 | 0.463821 | 1.080063 | 0.227386 | 0.550224 | 0.303534 | 0.2369 | 0.18399 | 0.020634 | -0.013212 | 0.122895 | 0.201919 | | 30.44 | -0.415426 | | 30.431 | 0.201919 |
| 30.444444 | 0.867941 | 1.100717 | 0.46228 | 1.038251 | 0.119807 | -0.044685 | 0.429473 | -0.067608 | -0.855752 | NaN | -0.531152 | 0.639641 | 0.724813 | 0.755354 | 0.033158 | 0.476942 | 0.220087 | 0.287524 | 0.446672 | -0.0295 | -0.084598 | 0.061314 | 0.326637 | | 30.45 | -0.531152 | | 30.444 | 0.326637 |
| 30.458333 | 1.514239 | 1.212737 | 1.155202 | 1.721516 | 1.219365 | -0.022619 | 0.88131 | 0.409155 | 0.47926 | NaN | -0.143468 | 0.861838 | 1.464017 | 0.579247 | -0.027017 | 0.422305 | 0.072262 | 0.296286 | 0.756952 | -0.159391 | 0.198223 | -0.114081 | 0.680118 | | 30.47 | -0.143468 | | 30.458 | 0.680118 |
| 30.472222 | 2.508045 | 1.285934 | 2.317647 | 2.643606 | 2.52922 | 0.252192 | 1.712915 | 1.012763 | 1.572271 | NaN | 0.586341 | 1.023591 | 1.852934 | 0.86673 | 0.245119 | 0.578926 | 0.186814 | 0.444246 | 1.20029 | -0.515119 | 0.870139 | -0.175628 | 1.043424 | | 30.48 | 0.586341 | | 30.472 | 1.043424 |
| 30.486111 | 3.206381 | 1.517444 | 3.884331 | 3.061644 | 2.701469 | 0.477314 | 2.496759 | 1.643401 | 2.052809 | NaN | 1.511446 | 1.172773 | 1.616923 | 1.323571 | 0.753404 | 0.91364 | 0.510512 | 0.786664 | 1.338591 | -0.92769 | 1.591526 | -0.097439 | 1.519577 | | 30.49 | 1.511446 | | 30.486 | 1.519577 |
| 30.5 | 2.893307 | 2.021494 | 4.301991 | 2.67587 | 2.20816 | 0.295726 | 2.354091 | 2.333832 | 2.275819 | NaN | 1.605812 | 1.372556 | 1.191121 | 1.68636 | 1.102015 | 1.146104 | 0.656188 | 1.095959 | 1.277842 | -0.698959 | 1.841748 | 0.03781 | 1.830771 | | 30.5 | 1.605812 | | 30.5 | 1.830771 |
| 30.513889 | 2.445659 | 1.838943 | 3.212917 | 1.673411 | 2.005351 | 0.594871 | 1.605754 | 1.842627 | 2.017377 | NaN | 0.998268 | 1.28453 | 0.550251 | 1.382365 | 1.016907 | 1.010961 | 0.527961 | 0.898313 | 0.990655 | -0.341443 | 1.25724 | 0.145577 | 1.64594 | | 30.51 | 0.998268 | | 30.514 | 1.64594 |
| 30.527778 | 1.030323 | 1.397214 | 1.918875 | 0.995985 | 1.756557 | 0.653319 | 0.83351 | 1.47532 | 1.616886 | 0.071819 | 0.781523 | 0.841306 | -0.114634 | 0.624544 | 0.619132 | 0.59486 | 0.058713 | 0.187029 | 0.533891 | -0.126239 | 0.193318 | 0.134194 | 1.267043 | | 30.53 | 0.781523 | | 30.528 | 1.267043 |
| 30.541667 | -0.383901 | 1.088952 | 0.750314 | 0.382603 | 1.860512 | 0.645132 | 0.058995 | 1.311309 | 1.632887 | 0.128451 | 0.981829 | 0.662297 | -0.740805 | -0.26212 | 0.256984 | 0.112278 | -0.428036 | -0.595235 | 0.081938 | -0.105911 | -0.8437 | 0.032504 | 0.935123 | | 30.54 | 0.981829 | | 30.542 | 0.935123 |
| 30.555556 | -1.143796 | 0.885343 | -0.082312 | 0.045814 | 1.868294 | 0.628043 | -0.486178 | 0.890712 | 0.785192 | 0.109578 | 1.258323 | 0.394502 | -1.335379 | -0.985802 | 0.06196 | -0.178326 | -0.554868 | -0.971165 | -0.297666 | -0.146934 | -1.362026 | -0.011962 | 0.41117 | | 30.56 | 1.258323 | | 30.556 | 0.41117 |
| 30.569444 | -0.993983 | 0.771558 | -0.79672 | -0.112431 | 1.627098 | 0.549123 | -0.516178 | 0.362948 | -0.496262 | NaN | 1.585401 | -0.01162 | -1.95451 | -1.469884 | -0.017761 | -0.310247 | -0.501279 | -0.950985 | -0.384814 | -0.044455 | -1.684215 | -0.016974 | -0.018755 | | 30.57 | 1.585401 | | 30.569 | -0.018755 |
| 30.583333 | -1.220727 | 0.505127 | -1.524013 | -0.06081 | 1.185898 | 0.30724 | -0.187337 | -0.181147 | -1.063763 | NaN | 1.083849 | 0.081459 | -2.055844 | -1.111788 | -0.085073 | -0.261313 | -0.225663 | -0.604533 | -0.265481 | 0.061546 | -1.667668 | -0.034512 | -0.236287 | | 30.58 | 1.083849 | | 30.583 | -0.236287 |
| 30.597222 | -1.282349 | -0.11186 | -1.993062 | -0.185201 | 0.520261 | 0.014561 | -0.020545 | -0.849341 | -1.177266 | NaN | -0.93042 | 0.577564 | -1.689083 | -0.280345 | -0.186011 | -0.375714 | 0.045515 | -0.146936 | -0.139214 | 0.23044 | -0.826069 | -0.005448 | -0.180425 | | 30.6 | -0.93042 | | 30.597 | -0.180425 |
| 30.611111 | -0.966271 | -0.843539 | -1.76406 | -0.372418 | -0.192653 | -0.404826 | 0.064214 | -1.155807 | -1.160122 | NaN | -2.013025 | 0.808312 | -1.144407 | 0.442799 | -0.294142 | -0.507229 | 0.075979 | -0.004018 | -0.107414 | 0.4283 | 0.106231 | 0.00892 | -0.091527 | | 30.61 | -2.013025 | | 30.611 | -0.091527 |
| 30.625 | 0.673893 | -1.11258 | -2.32458 | -0.388573 | -0.862433 | -0.611703 | -0.036296 | -1.181752 | -1.453365 | NaN | -1.347135 | 0.521127 | -0.447465 | 0.958553 | -0.298147 | -0.486844 | -0.052361 | -0.18822 | -0.101726 | 0.557604 | 0.612054 | -0.027943 | -0.104069 | | 30.63 | -1.347135 | | 30.625 | -0.104069 |
| 30.638889 | 1.359589 | -1.0503 | -2.047877 | -0.313423 | -1.398572 | -0.568153 | -0.173976 | -1.136532 | -0.976656 | NaN | -0.992642 | 0.001322 | 0.1396 | 1.026179 | -0.301775 | -0.310066 | -0.183831 | -0.403747 | -0.081246 | 0.50588 | 0.496189 | -0.062626 | -0.246051 | | 30.64 | -0.992642 | | 30.639 | -0.246051 |
| 30.652778 | 1.081754 | -0.693451 | -0.102407 | -0.012961 | -1.390943 | -0.542205 | -0.215845 | -1.029083 | 0.198437 | NaN | -0.467919 | -0.300545 | 0.297784 | 0.790526 | -0.228448 | -0.071341 | -0.222566 | -0.367191 | -0.149533 | 0.395973 | 0.027925 | -0.08118 | -0.346417 | | 30.65 | -0.467919 | | 30.653 | -0.346417 |
| 30.666667 | 1.025975 | -0.242228 | 1.116792 | 0.402948 | -1.501361 | -0.741907 | -0.16163 | -1.228351 | 1.082715 | NaN | -0.35894 | -0.336798 | 0.0958 | 0.521936 | -0.057923 | 0.18173 | -0.152695 | -0.21101 | -0.234618 | 0.256848 | -0.343827 | -0.068555 | -0.31036 | | 30.67 | -0.35894 | | 30.667 | -0.31036 |
| 30.680556 | 0.867142 | -0.137672 | 1.643589 | 0.766201 | -1.862491 | -0.476097 | 0.010342 | -1.550126 | 1.572241 | NaN | -1.205049 | -0.078638 | 0.140996 | 0.166648 | 0.13228 | 0.335995 | -0.112247 | -0.123956 | -0.265098 | 0.149855 | -0.407861 | -0.019012 | -0.109957 | | 30.68 | -1.205049 | | 30.681 | -0.109957 |
| 30.694444 | 0.409464 | -0.404671 | 1.242355 | 1.075551 | -1.881129 | -0.03015 | 0.178982 | -1.571249 | 2.115943 | NaN | -2.074724 | 0.417524 | 0.222873 | -0.329029 | 0.239886 | 0.313716 | -0.112001 | -0.097398 | -0.219243 | 0.090906 | -0.304701 | 0.063697 | 0.100576 | | 30.69 | -2.074724 | | 30.694 | 0.100576 |
| 30.708333 | -0.154365 | -0.70418 | 0.20293 | 1.382212 | -1.313184 | -0.053784 | 0.306772 | -1.014481 | 1.19082 | NaN | -2.215155 | 0.69439 | 0.068595 | -0.816342 | 0.194898 | 0.182397 | -0.100202 | -0.034617 | -0.103571 | 0.09405 | -0.178009 | 0.146012 | 0.220571 | | 30.71 | -2.215155 | | 30.708 | 0.220571 |
| 30.722222 | -0.341664 | -0.825825 | -0.143772 | 1.209171 | -0.688384 | 0.041269 | 0.286942 | -0.744348 | 0.446189 | NaN | -1.561717 | 0.712399 | -0.174559 | -0.922939 | 0.094255 | 0.01791 | -0.067628 | 0.048785 | 0.018314 | 0.142768 | -0.102139 | 0.137726 | 0.273807 | | 30.72 | -1.561717 | | 30.722 | 0.273807 |
| 30.736111 | -0.223336 | -0.806782 | -0.539653 | 0.771005 | -0.111465 | 0.069513 | 0.147881 | -0.597986 | -0.091702 | NaN | -0.567869 | 0.241533 | -0.516763 | -0.916704 | -0.073857 | -0.193163 | -0.062127 | 0.01029 | 0.139124 | 0.167328 | -0.233714 | 0.038519 | 0.285835 | | 30.74 | -0.567869 | | 30.736 | 0.285835 |
| 30.75 | 0.188454 | -0.688535 | -0.598089 | 0.105116 | 0.407455 | 0.066123 | -0.186837 | -0.440074 | -0.618775 | NaN | 0.251398 | -0.284654 | -1.140358 | -0.969914 | -0.348793 | -0.435626 | -0.086501 | -0.161145 | 0.056393 | 0.172464 | -0.440323 | -0.067128 | 0.316475 | | 30.75 | 0.251398 | | 30.75 | 0.316475 |
| 30.763889 | 0.201074 | -0.494158 | -0.642174 | -0.592208 | 0.726612 | 0.229129 | -0.524451 | -0.252637 | -1.356489 | NaN | 0.902502 | -0.659485 | -1.724488 | -1.01638 | -0.65377 | -0.714296 | -0.155429 | -0.382988 | -0.10875 | 0.173284 | -0.391318 | -0.123615 | 0.190261 | | 30.76 | 0.902502 | | 30.764 | 0.190261 |
| 30.777778 | -0.464491 | -0.429282 | -0.707328 | -0.938359 | 0.71488 | 0.662273 | -0.845351 | 0.135464 | -1.679817 | NaN | 1.38843 | -0.589128 | -1.972724 | -0.854081 | -0.848217 | -0.753426 | -0.134864 | -0.430189 | -0.312033 | 0.225438 | -0.069595 | -0.106003 | -0.05128 | | 30.78 | 1.38843 | | 30.778 | -0.05128 |
| 30.791667 | -0.795331 | -0.768803 | -1.813194 | -1.314115 | 0.355645 | 0.756309 | -0.769443 | 0.166484 | -1.423036 | NaN | 1.604467 | -0.420397 | -1.900283 | -0.756618 | -0.841443 | -0.60655 | -0.181993 | -0.409267 | -0.428058 | 0.275061 | 0.206219 | -0.008003 | -0.427674 | | 30.79 | 1.604467 | | 30.792 | -0.427674 |
| 30.805556 | -0.827792 | -1.355603 | -3.387755 | -1.588904 | -0.253523 | 0.28156 | -1.054718 | 0.060363 | -1.091741 | NaN | -0.865609 | -0.512046 | -1.933621 | -0.892486 | -0.72043 | -0.445171 | -0.207784 | -0.360012 | -0.579225 | 0.363931 | 0.244068 | 0.090917 | -0.808081 | | 30.81 | -0.865609 | | 30.806 | -0.808081 |
| 30.819444 | -0.912161 | -1.84699 | -4.047065 | -1.61835 | -0.968169 | 0.008663 | -1.295748 | -0.112933 | -0.900152 | NaN | -2.330573 | -0.728805 | -1.808477 | -0.853734 | -0.475533 | -0.323042 | -0.164044 | -0.210479 | -0.77522 | 0.443471 | 0.206806 | 0.083778 | -1.015278 | | 30.82 | -2.330573 | | 30.819 | -1.015278 |
| 30.833333 | -0.727102 | -1.655534 | -3.792266 | -1.52778 | -1.651224 | -0.02678 | -1.359089 | -0.195289 | -0.976974 | -1.099725 | -1.933311 | -1.117134 | -1.363868 | -0.571979 | -0.226508 | -0.242542 | 0.061349 | 0.091715 | -0.801747 | 0.439818 | 0.209405 | -0.031618 | -0.981001 | | 30.83 | -1.933311 | | 30.833 | -0.981001 |
| 30.847222 | -0.348796 | -1.209402 | -3.272382 | -1.364303 | -1.904049 | -0.206694 | -1.275746 | -0.458549 | -1.055954 | -0.579912 | -1.97327 | -1.352569 | -0.966836 | -0.288451 | -0.070023 | -0.15552 | 0.290544 | 0.409965 | -0.79255 | 0.328967 | 0.283989 | -0.122588 | -0.787515 | | 30.85 | -1.97327 | | 30.847 | -0.787515 |
| 30.861111 | 0.286297 | -0.661695 | -2.54589 | -1.076786 | -1.848236 | -0.453871 | -0.787948 | -0.821103 | -0.768715 | -0.261752 | -2.35617 | -1.174821 | -0.709671 | -0.165929 | 0.001348 | -0.023342 | 0.448888 | 0.508732 | -0.727054 | 0.242379 | 0.350863 | -0.103964 | -0.487253 | | 30.86 | -2.35617 | | 30.861 | -0.487253 |
| 30.875 | 1.038441 | 0.168626 | -1.558515 | -0.246425 | -1.427258 | -0.454118 | 0.003793 | -0.890964 | -0.405111 | 0.016017 | -1.727736 | -0.691483 | -0.42877 | -0.037682 | 0.034269 | 0.146741 | 0.432274 | 0.434076 | -0.563561 | 0.218882 | 0.299924 | -0.071335 | -0.209296 | | 30.88 | -1.727736 | | 30.875 | -0.209296 |
| 30.888889 | 1.994213 | 0.264544 | -0.315518 | 0.451095 | -0.833163 | -0.353409 | 0.980509 | -0.637749 | 0.201499 | 0.143091 | -1.068412 | -0.22186 | -0.257057 | 0.016618 | 0.024648 | 0.269212 | 0.268886 | 0.139361 | -0.358996 | 0.211471 | 0.210402 | -0.096567 | -0.017283 | | 30.89 | -1.068412 | | 30.889 | -0.017283 |
| 30.902778 | 2.706017 | 0.195878 | 0.971204 | 0.899709 | -0.232531 | -0.278921 | 1.825737 | -0.388872 | 0.790779 | 0.56459 | -0.645935 | 0.018636 | -0.112342 | -0.067139 | 0.000427 | 0.273605 | 0.047906 | -0.140297 | -0.312581 | 0.184294 | 0.127542 | -0.085954 | 0.057879 | | 30.9 | -0.645935 | | 30.903 | 0.057879 |
| 30.916667 | 2.838253 | 0.357329 | 1.954419 | 0.881362 | 0.318197 | -0.230049 | 2.200109 | -0.460804 | 1.024024 | 1.035196 | -0.19833 | -0.00153 | 0.022678 | -0.221279 | -0.034711 | 0.118566 | -0.059165 | -0.187801 | -0.342361 | 0.156655 | 0.079301 | -0.050475 | 0.074991 | | 30.92 | -0.19833 | | 30.917 | 0.074991 |
| 30.930556 | 2.404517 | 0.494662 | 2.484465 | 0.83912 | 0.6667 | -0.215841 | 1.925898 | -0.562421 | 1.090984 | 1.216735 | -0.122414 | -0.19632 | 0.026408 | -0.338385 | -0.117798 | -0.098083 | -0.02187 | -0.079914 | -0.278808 | 0.0943 | 0.027087 | -0.016623 | 0.032679 | | 30.93 | -0.122414 | | 30.931 | 0.032679 |
| 30.944444 | 1.078559 | 0.228627 | 2.069025 | 0.711596 | 0.2235 | -0.558798 | 1.217976 | -0.579882 | 0.502032 | 0.459105 | -1.026861 | -0.404599 | -0.333937 | -0.588514 | -0.475219 | -0.633831 | -0.139468 | -0.437722 | -0.47412 | -0.270638 | -0.211764 | 0.022993 | -0.306901 | | 30.94 | -1.026861 | | 30.944 | -0.306901 |
| 30.958333 | -1.600776 | -0.645366 | 0.122079 | -0.444106 | -1.1674 | -1.107231 | -0.266776 | -1.13793 | -0.774406 | NaN | -1.977814 | -0.741925 | -1.196202 | -1.300217 | -1.303569 | -1.662617 | -0.556898 | -1.57547 | -1.211494 | -1.014464 | -0.82579 | 0.122137 | -1.061799 | | 30.96 | -1.977814 | | 30.958 | -1.061799 |
| 30.972222 | -3.713823 | -1.128337 | -2.439712 | -1.760968 | -2.211743 | -1.069234 | -1.875076 | -1.006862 | -1.865537 | NaN | -1.373192 | -0.983428 | -1.836234 | -1.825413 | -2.054406 | -1.932303 | -0.851922 | -2.173964 | -1.742215 | -1.094769 | -1.219049 | 0.170381 | -1.664925 | | 30.97 | -1.373192 | | 30.972 | -1.664925 |
| 30.986111 | -3.423251 | -0.897623 | -3.519606 | -2.074417 | -2.064095 | -0.317575 | -3.264517 | -0.219349 | -1.987367 | NaN | -0.259454 | -0.531649 | -1.07382 | -1.149643 | -1.28351 | -1.28187 | -0.439071 | -1.216089 | -1.334483 | -0.304182 | -0.428856 | 0.05617 | -0.857553 | | 30.99 | -0.259454 | | 30.986 | -0.857553 |
| 31 | -2.547764 | -1.048206 | -3.304091 | -1.71635 | -1.364252 | 0.136014 | -4.186978 | 0.47461 | -1.899298 | NaN | 0.047136 | 0.242443 | -0.240925 | -0.35816 | -0.279049 | -0.220665 | 0.004188 | -0.129223 | -0.775323 | 0.457226 | 0.484962 | -0.038935 | 0.421297 | | 31.01 | 0.047136 | | 31 | 0.421297 |
| 31.013889 | -2.004047 | -0.850631 | -2.545223 | -1.472384 | -0.706679 | 0.362557 | -3.526186 | 0.449894 | -1.864476 | -0.849157 | -0.045219 | 0.70492 | 0.003338 | 0.122876 | 0.353562 | 0.253959 | 0.112584 | 0.187951 | -0.339467 | 0.628732 | 0.67687 | -0.090321 | 0.904367 | | 31.02 | -0.045219 | | 31.014 | 0.904367 |
| 31.027778 | -1.564428 | -1.133056 | -1.865892 | -0.913054 | -0.371976 | 0.310028 | -1.713941 | 0.411871 | -1.411309 | -0.913764 | -0.47723 | 0.988383 | -0.048632 | 0.449501 | 0.485716 | 0.315541 | 0.125942 | 0.181875 | -0.030335 | 0.430017 | 0.403327 | -0.220968 | 0.837695 | | 31.03 | -0.47723 | | 31.028 | 0.837695 |
| 31.041667 | -1.794952 | -1.555916 | -1.316473 | -0.424121 | -0.256774 | 0.371088 | 0.15298 | 0.363788 | -0.861323 | -0.381484 | -0.317791 | 1.041023 | 0.133735 | 0.916815 | 0.58856 | 0.397631 | 0.176737 | 0.412544 | 0.416372 | 0.428583 | 0.217509 | -0.395673 | 0.751312 | | 31.05 | -0.317791 | | 31.042 | 0.751312 |
| 31.055556 | -1.211861 | -1.053215 | -0.471184 | 0.074093 | -0.001036 | 0.575945 | 2.048875 | 0.438309 | -0.289049 | -0.042157 | -0.053687 | 0.899235 | 0.490257 | 1.40873 | 0.749335 | 0.538794 | 0.346468 | 0.850182 | 1.01885 | 0.732317 | 0.302748 | -0.258998 | 0.831766 | | 31.06 | -0.053687 | | 31.056 | 0.831766 |
| 31.069444 | -0.038589 | -0.027605 | 0.86632 | 0.916544 | 0.283462 | 1.005857 | 3.963056 | 0.59551 | 0.404122 | -0.1043 | 0.264431 | 0.885012 | 0.762085 | 1.539339 | 0.765224 | 0.781072 | 0.462561 | 1.125988 | 1.311308 | 0.886795 | 0.415613 | 0.103842 | 0.888444 | | 31.08 | 0.264431 | | 31.069 | 0.888444 |
| 31.083333 | 1.195094 | 0.972756 | 1.350445 | 1.521197 | 0.387501 | 1.126302 | 4.64499 | 0.421275 | 1.004535 | 0.104137 | 0.924761 | 0.709469 | 0.64333 | 1.138335 | 0.625873 | 0.805251 | 0.619648 | 1.200032 | 1.09166 | 0.594438 | 0.243477 | 0.264591 | 0.55512 | | 31.09 | 0.924761 | | 31.083 | 0.55512 |
| 31.097222 | 1.717079 | 1.464085 | 1.240068 | 1.368769 | 0.374921 | 0.831793 | 4.067246 | 0.101993 | 1.066664 | 0.525619 | 1.955665 | 0.262544 | 0.29134 | 0.643694 | 0.624762 | 0.749203 | 0.663433 | 1.023826 | 0.746735 | 0.325655 | -0.123342 | 5.416506 | 0.112683 | | 31.1 | 1.955665 | | 31.097 | 0.112683 |
| 31.111111 | 1.422581 | 1.355521 | 0.371374 | 0.879778 | 0.312829 | 0.395469 | 2.76187 | -0.02465 | 0.631384 | 0.203545 | 1.262755 | -0.250752 | -0.032828 | 0.200797 | 0.591438 | 0.608148 | 0.351271 | 0.308411 | 0.380267 | 0.191451 | -0.372857 | 1.647102 | -0.216242 | | 31.12 | 1.262755 | | 31.111 | -0.216242 |
| 31.125 | 0.91419 | 1.080211 | -0.226854 | 0.656622 | 0.196179 | -0.026974 | 1.489358 | -0.028345 | 0.050017 | -0.205404 | -0.425544 | -0.45068 | -0.183379 | 0.002702 | 0.416896 | 0.271959 | -0.08822 | -0.438447 | 0.189947 | 0.083495 | -0.343237 | -1.881998 | -0.306133 | | 31.13 | -0.425544 | | 31.125 | -0.306133 |
| 31.138889 | 0.818004 | 1.474597 | -0.208554 | 0.44851 | 0.019973 | -0.15802 | 0.688713 | 0.156718 | -0.320567 | -0.022563 | 0.285868 | -0.156292 | -0.177231 | 0.30338 | 0.188483 | 0.064299 | -0.396166 | -0.846525 | 0.100172 | -0.029412 | -0.1733 | -5.038144 | -0.137062 | | 31.15 | 0.285868 | | 31.139 | -0.137062 |
| 31.152778 | 1.081465 | 1.887086 | 0.218928 | 0.4642 | -0.020049 | 0.026519 | 0.375426 | 0.316441 | -0.411587 | 0.156191 | 1.685491 | 0.202845 | -0.07689 | 0.670917 | 0.12873 | -0.07 | -0.246833 | -0.718763 | -0.020836 | -0.07795 | -0.017099 | 0.048171 | 0.108296 | | 31.16 | 1.685491 | | 31.153 | 0.108296 |
| 31.166667 | 1.400567 | 1.918654 | 0.828233 | 0.523564 | 0.077944 | 0.17384 | 0.372071 | 0.213599 | -0.159686 | 1.390258 | 1.258749 | 0.223485 | -0.045859 | 0.804411 | 0.083245 | -0.206502 | -0.121234 | -0.470558 | -0.172764 | -0.09691 | 0.028234 | 0.024603 | 0.162275 | | 31.17 | 1.258749 | | 31.167 | 0.162275 |
| 31.180556 | 1.912332 | 2.098978 | 1.434302 | 0.606583 | 0.208219 | 0.228194 | 0.546464 | 0.062013 | 0.214302 | 0.98838 | 0.777034 | -0.028567 | -0.096528 | 0.653395 | -0.056483 | -0.132715 | 0.059648 | -0.141884 | -0.208415 | -0.126224 | 0.001974 | -0.079554 | 0.090217 | | 31.19 | 0.777034 | | 31.181 | 0.090217 |
| 31.194444 | 2.540056 | 1.931454 | 1.650701 | 0.675518 | 0.344908 | 0.284388 | 0.845277 | -0.057007 | 0.68155 | NaN | 0.992728 | -0.14742 | -0.10798 | 0.336307 | -0.210423 | 0.06909 | 0.137238 | 0.087235 | -0.110532 | -0.153029 | -0.001485 | -0.159079 | -0.034601 | | 31.2 | 0.992728 | | 31.194 | -0.034601 |
| 31.208333 | 2.733412 | 1.529762 | 1.493953 | 0.621128 | 0.361958 | 0.279007 | 0.963825 | -0.171927 | 0.8956 | NaN | 1.188694 | -0.217083 | -0.162062 | -0.055471 | -0.319734 | 0.073724 | 0.015366 | -0.152531 | -0.074078 | -0.156363 | 0.009262 | -0.183401 | -0.191745 | | 31.22 | 1.188694 | | 31.208 | -0.191745 |
| 31.222222 | 2.573783 | 1.032388 | 0.997631 | 0.44487 | 0.290388 | 0.250866 | 0.694713 | -0.226707 | 0.626894 | NaN | 0.963508 | -0.233535 | -0.255974 | -0.366229 | -0.360162 | -0.179545 | -0.205856 | -0.589189 | -0.157959 | -0.154977 | -0.003761 | -0.181866 | -0.342236 | | 31.23 | 0.963508 | | 31.222 | -0.342236 |
| 31.236111 | 2.086786 | 0.266586 | 0.527027 | 0.283305 | 0.2249 | 0.198446 | 0.325923 | -0.159448 | 0.164004 | NaN | -0.096147 | -0.078459 | -0.201094 | -0.263432 | -0.301564 | -0.421519 | -0.353813 | -0.783104 | -0.186932 | -0.131048 | 0.01153 | -0.146127 | -0.321413 | | 31.24 | -0.096147 | | 31.236 | -0.321413 |
| 31.25 | 1.229656 | -0.422323 | 0.427741 | 0.147721 | 0.195759 | 0.096199 | 0.033463 | -0.03792 | -0.092984 | NaN | -1.566009 | 0.035547 | -0.042589 | -0.053217 | -0.160434 | -0.379936 | -0.348189 | -0.659067 | -0.10853 | -0.093407 | 0.066446 | 1.368083 | -0.068719 | | 31.26 | -1.566009 | | 31.25 | -0.068719 |
| 31.263889 | 0.336387 | -0.627212 | 0.447495 | 0.014985 | 0.156354 | 0.100388 | -0.091619 | 0.095489 | -0.148334 | NaN | -1.844905 | 0.060759 | 0.163711 | 0.221394 | -0.091089 | -0.162534 | -0.225198 | -0.379921 | 0.082078 | -0.045543 | 0.093493 | 0.124716 | 0.193488 | | 31.27 | -1.844905 | | 31.264 | 0.193488 |
| 31.277778 | -0.199547 | -0.281751 | 0.651251 | -0.096817 | 0.070049 | 0.309961 | -0.048153 | 0.190806 | -0.098148 | NaN | -0.60593 | 0.037698 | 0.342208 | 0.348498 | -0.022309 | 0.103686 | -0.082193 | -0.084494 | 0.247437 | 0.021206 | 0.079884 | -1.056602 | 0.269672 | | 31.28 | -0.60593 | | 31.278 | 0.269672 |
| 31.291667 | -0.382234 | 0.207855 | 0.89434 | -0.047193 | 0.066751 | 0.489632 | 0.138591 | 0.299893 | 0.032939 | NaN | 0.998384 | 0.199607 | 0.437918 | 0.372226 | 0.098892 | 0.263076 | 0.107992 | 0.254761 | 0.288628 | 0.107937 | 0.077782 | -0.024844 | 0.282187 | | 31.3 | 0.998384 | | 31.292 | 0.282187 |
| 31.305556 | -0.271739 | 0.720608 | 1.043964 | 0.07487 | 0.052268 | 0.59826 | 0.424752 | 0.353009 | 0.200674 | NaN | 2.468074 | 0.348267 | 0.41982 | 0.301384 | 0.136705 | 0.328995 | 0.199438 | 0.423133 | 0.230749 | 0.148377 | 0.06346 | 0.0449 | 0.202141 | | 31.31 | 2.468074 | | 31.306 | 0.202141 |
| 31.319444 | -0.281727 | 0.750472 | 1.136524 | 0.024937 | 0.001274 | 0.543385 | 0.395007 | 0.30542 | 0.311645 | NaN | 2.703176 | 0.403667 | 0.346395 | 0.180251 | 0.036578 | 0.288598 | 0.151793 | 0.331312 | 0.200605 | 0.114427 | 0.045782 | 1.377191 | 0.09907 | | 31.33 | 2.703176 | | 31.319 | 0.09907 |
| 31.333333 | -0.521547 | 0.1587 | 1.088161 | -0.038628 | -0.075057 | 0.269331 | -0.074472 | 0.214039 | 0.292097 | NaN | 1.299513 | 0.35696 | 0.360097 | 0.128969 | -0.055894 | 0.241403 | 0.02051 | 0.113076 | 0.249755 | 0.058174 | 0.058257 | 1.143212 | 0.026753 | | 31.34 | 1.299513 | | 31.333 | 0.026753 |
| 31.347222 | -0.94369 | -0.459637 | 0.974087 | -0.195918 | -0.142021 | 0.050325 | -0.535625 | 0.195157 | 0.21874 | NaN | -0.255656 | 0.248989 | 0.466979 | 0.222513 | -0.070922 | 0.246788 | -0.115472 | -0.121618 | 0.297723 | 0.016662 | 0.085551 | -0.483891 | 0.078556 | | 31.35 | -0.255656 | | 31.347 | 0.078556 |
| 31.361111 | -1.270216 | -0.675249 | 0.828817 | -0.335127 | -0.178954 | -0.018347 | -0.669786 | 0.26185 | 0.174329 | NaN | -0.927273 | 0.132952 | 0.543252 | 0.393959 | -0.022861 | 0.297403 | -0.135769 | -0.194351 | 0.299177 | 0.004177 | 0.100305 | -1.519163 | 0.228208 | | 31.37 | -0.927273 | | 31.361 | 0.228208 |
| 31.375 | -1.010085 | -0.273319 | 0.621769 | -0.269166 | -0.152622 | 0.207396 | -0.338521 | 0.360204 | 0.144047 | NaN | 0.054914 | 0.150879 | 0.567658 | 0.523176 | 0.054422 | 0.404275 | -0.056207 | -0.094788 | 0.371405 | 0.026871 | 0.124072 | -0.744103 | 0.363372 | | 31.38 | 0.054914 | | 31.375 | 0.363372 |
| 31.388889 | -0.592021 | -0.043287 | 0.553373 | 0.002798 | -0.094165 | 0.329558 | -0.032019 | 0.392344 | 0.102946 | NaN | 0.451588 | 0.291869 | 0.540862 | 0.519034 | 0.151067 | 0.557076 | 0.040587 | 0.108121 | 0.544574 | 0.0248 | 0.14333 | 0.119069 | 0.394754 | | 31.4 | 0.451588 | | 31.389 | 0.394754 |
| 31.402778 | -0.583405 | -0.598155 | 0.647764 | 0.310018 | -0.20567 | 0.038813 | -0.159767 | 0.278797 | 0.013442 | NaN | -1.442425 | 0.404995 | 0.501407 | 0.344435 | 0.190205 | 0.648322 | 0.058039 | 0.295831 | 0.694156 | -0.03899 | 0.104827 | 1.447205 | 0.26929 | | 31.41 | -1.442425 | | 31.403 | 0.26929 |
| 31.416667 | -0.611671 | -0.981075 | 0.7462 | 0.439313 | -0.42764 | -0.237216 | -0.33764 | 0.13948 | -0.00331 | NaN | -2.429202 | 0.340063 | 0.44996 | 0.14432 | 0.200806 | 0.747431 | 0.018883 | 0.39079 | 0.697075 | -0.085938 | 0.022796 | 4.626684 | 0.070715 | | 31.42 | -2.429202 | | 31.417 | 0.070715 |
| 31.430556 | 0.062767 | -0.236903 | 0.706171 | 0.47128 | -0.603926 | -0.007834 | -0.066204 | 0.09966 | 0.118904 | NaN | -0.14915 | 0.185152 | 0.441 | 0.058857 | 0.263331 | 0.849593 | 0.008183 | 0.404781 | 0.640739 | -0.057704 | 0.012654 | 7.412917 | -0.055708 | | 31.44 | -0.14915 | | 31.431 | -0.055708 |
| 31.444444 | 1.092318 | 0.884535 | 1.154295 | 0.725978 | -0.594499 | 0.527604 | 0.501013 | 0.23245 | 0.500413 | NaN | 2.853961 | 0.331092 | 0.772104 | 0.360944 | 0.564192 | 1.144364 | 0.088206 | 0.596326 | 0.779938 | 0.082922 | 0.12421 | 6.292636 | -0.012179 | | 31.45 | 2.853961 | | 31.444 | -0.012179 |
| 31.458333 | 1.912593 | 0.986447 | 2.710592 | 1.194621 | -0.42348 | 0.744437 | 1.038319 | 0.440406 | 1.179426 | NaN | 2.552529 | 0.872425 | 1.784367 | 0.86005 | 1.086579 | 1.763244 | 0.244571 | 1.027881 | 1.193769 | 0.311519 | 0.336913 | 5.413469 | 0.14352 | | 31.47 | 2.552529 | | 31.458 | 0.14352 |
| 31.472222 | 2.537371 | 0.489285 | 3.710699 | 1.834001 | 0.186365 | 0.75537 | 1.980654 | 0.345846 | 1.523775 | NaN | 0.915627 | 1.1119 | 2.535268 | 1.437948 | 1.229712 | 2.16372 | 0.315767 | 1.23635 | 1.356342 | 0.366171 | 0.547123 | 4.727393 | 0.288586 | | 31.48 | 0.915627 | | 31.472 | 0.288586 |
| 31.486111 | 3.252521 | 0.680736 | 3.308564 | 2.668562 | 1.064273 | 0.766436 | 3.279172 | -0.023785 | 1.334992 | NaN | 1.292962 | 1.039803 | 1.754742 | 2.198972 | 0.939452 | 2.026213 | 0.386104 | 1.158573 | 1.207135 | 0.123242 | 0.802187 | 1.626424 | 0.451032 | | 31.49 | 1.292962 | | 31.486 | 0.451032 |
| 31.5 | 3.662005 | 0.961074 | 2.78643 | 2.819789 | 1.425544 | 0.393447 | 3.209204 | -0.192265 | 1.480611 | NaN | 1.898928 | 1.003731 | 0.887972 | 2.727781 | 0.776565 | 1.691755 | 0.192777 | 1.098888 | 1.092541 | -0.19627 | 1.174374 | -0.124394 | 0.475126 | | 31.5 | 1.898928 | | 31.5 | 0.475126 |
| 31.513889 | 3.411817 | 0.598663 | 2.043255 | 2.373578 | 1.176499 | 0.057788 | 1.803038 | -0.076666 | 0.981085 | NaN | 0.869776 | 0.951933 | 0.139124 | 1.675313 | 0.675514 | 1.020037 | 0.059215 | 0.775871 | 0.679858 | -0.422631 | 0.885378 | 1.681575 | 0.215945 | | 31.51 | 0.869776 | | 31.514 | 0.215945 |
| 31.527778 | 2.454556 | 0.134493 | 1.510446 | 1.379017 | 0.961834 | -0.078983 | 0.570653 | 0.120785 | 0.716253 | NaN | 0.077889 | 0.327586 | -0.525581 | 0.267633 | 0.417592 | 0.162498 | -0.040805 | 0.222926 | 0.130709 | -0.509312 | 0.265843 | 0.210279 | -0.148799 | | 31.53 | 0.077889 | | 31.528 | -0.148799 |
| 31.541667 | 1.515367 | 0.201343 | 1.036792 | 0.594438 | 1.360408 | -0.282372 | -0.232434 | 0.381083 | 1.199345 | NaN | 0.483269 | -0.362177 | -1.098145 | -1.168958 | 0.221588 | -0.566304 | -0.284351 | -0.316953 | -0.374399 | -0.49356 | -0.320073 | -1.131658 | -0.426414 | | 31.54 | 0.483269 | | 31.542 | -0.426414 |
| 31.555556 | 0.636262 | 0.334722 | 0.86763 | 0.033789 | 2.116308 | -0.416136 | -0.954145 | 0.639727 | 1.257909 | NaN | 0.481307 | -0.610816 | -1.478069 | -1.706863 | 0.120805 | -1.040359 | -0.425803 | -0.642918 | -0.50891 | -0.366771 | -0.64613 | 0.906963 | -0.556128 | | 31.56 | 0.481307 | | 31.556 | -0.556128 |
| 31.569444 | -0.357786 | 0.311691 | 0.169199 | -0.123847 | 2.01314 | -0.354134 | -1.490788 | 0.755361 | 0.63276 | NaN | -0.317373 | -0.380552 | -1.652878 | -1.374365 | 0.091058 | -1.011928 | -0.575908 | -0.953009 | -0.353737 | -0.217099 | -0.727766 | -0.16232 | -0.575194 | | 31.57 | -0.317373 | | 31.569 | -0.575194 |
| 31.583333 | -0.950366 | 0.2963 | -0.104174 | -0.13002 | 1.491985 | -0.170644 | -1.363245 | 0.670607 | 0.007753 | NaN | -0.357637 | -0.036347 | -1.83275 | -1.008346 | 0.020826 | -0.754309 | -0.481644 | -0.915548 | -0.536517 | -0.099661 | -0.733437 | -1.667312 | -0.485582 | | 31.58 | -0.357637 | | 31.583 | -0.485582 |
| 31.597222 | -1.605599 | 0.16699 | -0.168672 | -0.285318 | 1.401243 | 0.156583 | -1.171638 | 0.264641 | -1.045162 | NaN | -0.393389 | 0.113937 | -1.801877 | -0.715953 | -0.122403 | -0.805068 | -0.242864 | -0.576694 | -0.635551 | -0.022965 | -0.699279 | 0.994994 | -0.332894 | | 31.6 | -0.393389 | | 31.597 | -0.332894 |
| 31.611111 | -1.491722 | -0.410493 | -0.209164 | -0.53395 | 1.165697 | -0.059666 | -1.180034 | -0.310133 | -2.054197 | NaN | -1.324479 | 0.131941 | -1.45358 | -0.486822 | -0.289447 | -0.929214 | 0.035803 | -0.230583 | -0.712232 | 0.043973 | -0.593787 | 6.299546 | -0.196977 | | 31.61 | -1.324479 | | 31.611 | -0.196977 |
| 31.625 | -1.331293 | -0.726251 | -0.360245 | -0.34146 | 0.727604 | -0.444169 | -1.3318 | -0.750947 | -2.004585 | NaN | -1.484402 | 0.037725 | -0.920759 | -0.383772 | -0.385278 | -0.961647 | 0.232838 | -0.062927 | -0.637348 | 0.037325 | -0.443307 | 6.369974 | -0.136739 | | 31.63 | -1.484402 | | 31.625 | -0.136739 |
| 31.638889 | -0.409642 | -0.613514 | -0.381303 | 0.021465 | 0.172688 | -0.529618 | -1.28601 | -1.037505 | -1.53384 | NaN | -0.516215 | -0.073181 | -0.243472 | -0.264685 | -0.362143 | -0.884358 | 0.084908 | -0.098008 | -0.34171 | -0.01523 | -0.255907 | 5.399834 | -0.171056 | | 31.64 | -0.516215 | | 31.639 | -0.171056 |
| 31.652778 | 0.279076 | -0.456812 | -0.177057 | 0.358041 | -0.583576 | -0.385054 | -0.95139 | -0.995101 | -0.967892 | NaN | 0.635136 | -0.165884 | 0.558969 | -0.036946 | -0.224718 | -0.615498 | -0.178994 | -0.233535 | 0.136509 | -0.038107 | -0.121496 | 5.471953 | -0.142693 | | 31.65 | 0.635136 | | 31.653 | -0.142693 |
| 31.666667 | 0.994417 | -0.519321 | 0.006333 | 0.558642 | -1.590846 | -0.324518 | -0.576028 | -0.603801 | -0.329561 | NaN | 1.165322 | -0.206905 | 1.28271 | 0.279903 | -0.114509 | -0.098181 | -0.48034 | -0.35653 | 0.495516 | -0.022547 | 0.073347 | 3.089149 | 0.004331 | | 31.67 | 1.165322 | | 31.667 | 0.004331 |
| 31.680556 | 1.683945 | -0.676422 | 0.091147 | 0.79726 | -2.217388 | -0.399244 | -0.167597 | -0.193181 | 0.723182 | NaN | 0.874007 | -0.349092 | 1.59274 | 0.367382 | -0.016361 | 0.434865 | -0.543206 | -0.281508 | 0.570691 | 0.034648 | 0.291152 | -1.224133 | 0.114972 | | 31.68 | 0.874007 | | 31.681 | 0.114972 |
| 31.694444 | 1.745972 | -0.801703 | 0.165967 | 0.636257 | -2.267104 | -0.547081 | 0.227432 | -0.088564 | 1.575083 | NaN | 0.043296 | -0.540615 | 1.33508 | 0.410198 | 0.080008 | 0.780502 | -0.323984 | -0.07014 | 0.332213 | 0.117706 | 0.394934 | -4.70046 | 0.064216 | | 31.69 | 0.043296 | | 31.694 | 0.064216 |
| 31.708333 | 2.247867 | -0.739699 | 0.409129 | 0.672117 | -1.889979 | -1.056586 | 0.418554 | -0.439389 | 2.34012 | NaN | -0.934179 | -0.572121 | 0.783031 | 0.399265 | 0.125871 | 0.822847 | -0.189223 | 0.091209 | -0.155718 | 0.172841 | 0.436877 | -7.752094 | -0.081493 | | 31.71 | -0.934179 | | 31.708 | -0.081493 |
| 31.722222 | 2.30544 | -0.379577 | 0.611995 | 0.149433 | -1.599833 | -1.34588 | 0.365336 | -1.076538 | 2.060138 | NaN | -1.942465 | -0.38809 | 0.128078 | 0.356669 | 0.186313 | 0.684317 | -0.069293 | 0.163886 | -0.581145 | 0.16329 | 0.159124 | -6.854621 | -0.281414 | | 31.72 | -1.942465 | | 31.722 | -0.281414 |
| 31.736111 | 1.41479 | 0.279252 | 0.680078 | 0.012014 | -1.540359 | -1.1376 | 0.676277 | -0.92758 | 1.62439 | NaN | -1.631063 | -0.185699 | -0.376539 | 0.219475 | 0.259397 | 0.314005 | 0.009478 | 0.095016 | -0.739306 | 0.135549 | -0.253161 | -8.500046 | -0.53659 | | 31.74 | -1.631063 | | 31.736 | -0.53659 |
| 31.75 | 0.217723 | 0.689136 | 0.075923 | -0.365046 | -1.482241 | -1.207992 | 0.288525 | -0.835175 | 0.537427 | NaN | -1.248653 | 0.033106 | -0.683292 | -0.076548 | 0.285056 | -0.255255 | 0.04469 | 0.035059 | -0.60437 | 0.106293 | -0.617248 | -8.428834 | -0.578628 | | 31.75 | -1.248653 | | 31.75 | -0.578628 |
| 31.763889 | -0.519949 | 0.446662 | -0.20186 | -0.690625 | -1.573753 | -1.62542 | -0.450836 | -0.970389 | -0.605101 | NaN | -1.962756 | 0.027197 | -0.96885 | -0.470514 | 0.245316 | -0.815404 | 0.082268 | 0.047855 | -0.352531 | 0.102663 | -0.808698 | -7.994129 | -0.579173 | | 31.76 | -1.962756 | | 31.764 | -0.579173 |
| 31.777778 | -1.448289 | -0.562434 | -0.472906 | -1.054004 | -1.504998 | -2.03764 | -1.144947 | -0.93069 | -1.303491 | NaN | -3.152301 | -0.265211 | -1.32769 | -0.837 | 0.062391 | -1.230282 | 0.087848 | 0.083565 | -0.241919 | 0.084651 | -0.522321 | -5.556495 | -0.55585 | | 31.78 | -3.152301 | | 31.778 | -0.55585 |
| 31.791667 | -2.065071 | -1.356583 | -0.593297 | -0.875039 | -1.287282 | -1.997481 | -1.459745 | -0.660694 | -1.546139 | NaN | -3.043534 | -0.624584 | -1.497215 | -1.031308 | -0.142729 | -1.501211 | 0.118999 | 0.105684 | -0.35852 | 0.057411 | -0.274328 | -0.179114 | -0.491274 | | 31.79 | -3.043534 | | 31.792 | -0.491274 |
| 31.805556 | -2.268898 | -1.825241 | -0.857502 | -0.854078 | -1.195827 | -1.587315 | -1.611379 | -0.611149 | -1.929942 | NaN | -2.068242 | -0.777293 | -1.428429 | -0.718316 | -0.288383 | -1.320582 | 0.123157 | 0.11507 | -0.479277 | 0.016589 | 0.013607 | -0.159755 | -0.303622 | | 31.81 | -2.068242 | | 31.806 | -0.303622 |
| 31.819444 | -2.442461 | -1.852074 | -1.349913 | -0.902806 | -1.170644 | -1.490733 | -1.993819 | -0.669652 | -2.147785 | NaN | -1.621107 | -0.679408 | -1.082584 | -0.452116 | -0.400444 | -1.04331 | 0.117894 | 0.046005 | -0.413771 | 0.021948 | 0.179513 | -0.072913 | -0.149639 | | 31.82 | -1.621107 | | 31.819 | -0.149639 |
| 31.833333 | -2.39515 | -1.851645 | -1.88043 | -0.986202 | -1.031314 | -1.643804 | -2.596809 | -0.851325 | -2.244717 | NaN | -1.518484 | -0.410963 | -0.763771 | -0.368809 | -0.461814 | -0.665379 | 0.152587 | 0.038029 | -0.255369 | 0.04553 | 0.175273 | 7.99828 | 0.00285 | | 31.83 | -1.518484 | | 31.833 | 0.00285 |
| 31.847222 | -2.207768 | -1.798938 | -2.262277 | -0.959925 | -0.852517 | -1.835494 | -2.708459 | -1.056243 | -1.977189 | NaN | -1.166771 | -0.095749 | -0.472722 | -0.229225 | -0.523421 | -0.337484 | 0.279202 | 0.092725 | -0.08823 | 0.07769 | 0.074844 | 4.20321 | 0.095609 | | 31.85 | -1.166771 | | 31.847 | 0.095609 |
| 31.861111 | -1.962007 | -2.518483 | -1.969417 | -0.89307 | -0.593411 | -1.474211 | -2.014933 | -1.258584 | -1.508769 | NaN | -1.276094 | 0.135467 | -0.49952 | -0.085719 | -0.537205 | -0.007054 | 0.347793 | 0.123631 | 0.103372 | 0.1382 | -0.007223 | 0.683195 | 0.22315 | | 31.86 | -1.276094 | | 31.861 | 0.22315 |
| 31.875 | -1.106744 | -1.7144 | -1.198572 | -0.628938 | -0.343793 | -1.006606 | -1.392622 | -1.210544 | -1.461783 | NaN | -1.29084 | 0.141275 | -0.514712 | 0.009208 | -0.520332 | 0.097533 | 0.39138 | 0.124199 | 0.298785 | 0.207879 | -0.067338 | -1.572015 | 0.279519 | | 31.88 | -1.29084 | | 31.875 | 0.279519 |
| 31.888889 | -0.097505 | -0.889404 | -0.527328 | -0.060573 | -0.085025 | -0.553742 | -0.765039 | -0.747772 | -0.934211 | NaN | -0.49339 | -0.147844 | -0.525689 | 0.004757 | -0.413885 | 0.000539 | 0.308752 | 0.013019 | 0.425478 | 0.278223 | -0.033019 | -3.394735 | 0.214515 | | 31.89 | -0.49339 | | 31.889 | 0.214515 |
| 31.902778 | 0.648721 | -0.233384 | 0.112249 | 0.53866 | 0.135399 | -0.113589 | -0.113049 | -0.404165 | -0.105814 | NaN | -0.190527 | -0.370186 | -0.392956 | 0.005545 | -0.225049 | 0.083933 | 0.174745 | -0.010812 | 0.497129 | 0.322683 | 0.09381 | -0.078119 | 0.163372 | | 31.9 | -0.190527 | | 31.903 | 0.163372 |
| 31.916667 | 1.075597 | 0.610747 | 0.666171 | 1.018273 | 0.297367 | 0.093649 | 0.565248 | -0.236959 | 0.735631 | NaN | -0.280416 | -0.33822 | -0.23415 | 0.065924 | -0.060292 | 0.329647 | 0.069199 | 0.040063 | 0.434052 | 0.31508 | 0.25982 | -0.095485 | 0.21566 | | 31.92 | -0.280416 | | 31.917 | 0.21566 |
| 31.930556 | 1.221614 | 1.15912 | 1.195453 | 1.241454 | 0.388497 | 0.151064 | 1.147015 | -0.205046 | 1.476814 | NaN | -0.85807 | -0.121246 | -0.168027 | 0.03943 | -0.019916 | 0.577877 | 0.020115 | 0.080191 | 0.271751 | 0.160997 | 0.304609 | -0.099807 | 0.186391 | | 31.93 | -0.85807 | | 31.931 | 0.186391 |
| 31.944444 | 0.618379 | 0.897872 | 0.91745 | 0.590372 | 0.042908 | -0.266274 | 1.002835 | -0.362551 | 1.315099 | NaN | -1.66776 | -0.153822 | -0.489238 | -0.414897 | -0.457453 | 0.020937 | -0.11742 | -0.488271 | -0.296518 | -0.472112 | -0.018613 | -0.100321 | -0.397176 | | 31.94 | -1.66776 | | 31.944 | -0.397176 |
| 31.958333 | -0.874282 | -0.665871 | -1.006334 | -1.464642 | -0.73639 | -1.081467 | -0.37005 | -0.575607 | -0.627426 | NaN | -0.725083 | -0.982031 | -1.376348 | -1.389775 | -1.418812 | -1.649335 | -0.486434 | -1.834469 | -1.414344 | -1.855335 | -0.79323 | -0.117929 | -1.690519 | | 31.96 | -0.725083 | | 31.958 | -1.690519 |
| 31.972222 | -2.476581 | -2.608063 | -3.083684 | -3.614297 | -1.468946 | -1.560037 | -2.337576 | -0.705042 | -2.865126 | NaN | 0.281888 | -1.928093 | -1.786461 | -1.540205 | -1.479224 | -3.044241 | -0.621028 | -2.014831 | -2.055754 | -2.326132 | -1.158104 | -0.015262 | -2.069609 | | 31.97 | 0.281888 | | 31.972 | -2.069609 |
| 31.986111 | -3.248408 | -3.308253 | -2.946296 | -3.592755 | -1.364 | -1.261273 | -2.74619 | -0.731637 | -3.289976 | NaN | -0.964203 | -1.725199 | -0.679125 | -0.598079 | -0.540373 | -2.231024 | -0.150026 | -0.711235 | -1.381171 | -0.689284 | -0.486523 | 0.257311 | -0.800018 | | 31.99 | -0.964203 | | 31.986 | -0.800018 |
| 32 | -3.10062 | -3.31527 | -2.449875 | -2.906548 | -1.149771 | -1.611479 | -2.226079 | -0.604347 | -2.743265 | NaN | -1.650558 | -0.711094 | 0.28036 | -0.101738 | 0.341124 | -0.620729 | 0.310973 | 0.539709 | -0.460431 | 0.511028 | 0.289689 | 0.379555 | 0.41974 | | 32.01 | -1.650558 | | 32 | 0.41974 |
| 32.013889 | -2.427111 | -3.482816 | -2.689881 | -2.183364 | -0.965697 | -1.824247 | -2.239382 | -0.344966 | -2.310607 | NaN | -0.655103 | 0.008121 | 0.285509 | -0.105384 | 0.520609 | 0.281503 | 0.321852 | 0.863912 | 0.138223 | 0.833722 | 0.596589 | 0.171672 | 0.911274 | | 32.02 | -0.655103 | | 32.014 | 0.911274 |
| 32.027778 | -1.565746 | -3.170529 | -2.355594 | -1.091054 | -0.852092 | -1.478853 | -1.635647 | -0.221454 | -0.992693 | NaN | -0.523223 | 0.566769 | 0.000612 | -0.018507 | 0.432778 | 0.719369 | 0.240986 | 0.715582 | 0.40997 | 0.661462 | 0.529693 | -0.008813 | 1.080067 | | 32.03 | -0.523223 | | 32.028 | 1.080067 |
| 32.041667 | -0.505206 | -2.200008 | -1.316463 | -0.142597 | -0.46548 | -0.623837 | -0.519403 | -0.078827 | 0.018891 | NaN | -0.101657 | 1.209557 | 0.020209 | 0.311195 | 0.419483 | 1.024108 | 0.224604 | 0.704086 | 0.58947 | 0.500135 | 0.636486 | -0.048433 | 1.41802 | | 32.05 | -0.101657 | | 32.042 | 1.41802 |
| 32.055556 | 0.508571 | -0.944198 | -0.473662 | 0.467682 | 0.095069 | 0.470492 | 0.064461 | 0.194856 | 0.682492 | NaN | 1.242717 | 1.405292 | 0.038982 | 0.7856 | 0.360827 | 1.047166 | 0.202973 | 0.54165 | 0.683792 | 0.408273 | 0.649585 | -0.030216 | 1.431919 | | 32.06 | 1.242717 | | 32.056 | 1.431919 |
| 32.069444 | 1.114929 | -0.243968 | 0.056566 | 0.787066 | 0.477819 | 1.375559 | 0.376294 | 0.37413 | 1.12169 | NaN | 2.064629 | 1.323898 | 0.085945 | 1.14797 | 0.253287 | 0.847951 | 0.170465 | 0.358263 | 0.574437 | 0.355104 | 0.586155 | -0.006412 | 1.314465 | | 32.08 | 2.064629 | | 32.069 | 1.314465 |
| 32.083333 | 1.406104 | 0.074953 | 0.368361 | 0.982631 | 0.690062 | 1.728056 | 0.709373 | 0.430955 | 1.294144 | NaN | 1.733879 | 1.30146 | 0.273028 | 1.20671 | 0.082518 | 0.650142 | 0.141862 | 0.083698 | 0.368813 | 0.319172 | 0.574753 | 0.028191 | 1.12221 | | 32.09 | 1.733879 | | 32.083 | 1.12221 |
| 32.097222 | 1.50972 | 0.173519 | 0.578493 | 1.185448 | 0.84503 | 1.728922 | 1.005773 | 0.428695 | 1.402193 | NaN | 0.837888 | 1.08941 | 0.419433 | 1.184725 | 0.029343 | 0.562037 | 0.190174 | 0.03614 | 0.3001 | 0.243479 | 0.49101 | 0.057692 | 0.814879 | | 32.1 | 0.837888 | | 32.097 | 0.814879 |
| 32.111111 | 1.734263 | 0.3962 | 0.616721 | 1.237271 | 0.9326 | 1.828237 | 1.090503 | 0.552625 | 1.464992 | NaN | 0.850775 | 0.710676 | 0.342497 | 1.262208 | 0.113855 | 0.514521 | 0.117566 | 0.085816 | 0.542171 | 0.22204 | 0.282356 | 0.040631 | 0.489737 | | 32.12 | 0.850775 | | 32.111 | 0.489737 |
| 32.125 | 2.05354 | 0.932578 | 0.626745 | 1.070934 | 0.868263 | 1.733265 | 0.953326 | 0.738371 | 1.635161 | NaN | 1.746715 | 0.271523 | 0.153698 | 1.022529 | 0.303914 | 0.64669 | 0.040832 | 0.144337 | 1.018878 | 0.18081 | 0.056352 | -0.01172 | 0.188269 | | 32.13 | 1.746715 | | 32.125 | 0.188269 |
| 32.138889 | 2.023842 | 1.206203 | 0.688352 | 0.974678 | 0.76671 | 1.448257 | 0.922755 | 0.772576 | 2.097002 | NaN | 1.606206 | -0.052142 | -0.025951 | 0.546621 | 0.479611 | 0.951576 | -0.063463 | 0.25769 | 1.186021 | 0.100609 | -0.041682 | -0.048196 | 0.007264 | | 32.15 | 1.606206 | | 32.139 | 0.007264 |
| 32.152778 | 1.827123 | 1.279406 | 0.733126 | 1.013099 | 0.603053 | 1.044159 | 1.035929 | 0.87468 | 2.547258 | NaN | 1.426015 | -0.155164 | -0.093098 | 0.084976 | 0.603864 | 1.048203 | -0.037824 | 0.258306 | 0.907608 | 0.025558 | 0.033962 | -0.017756 | -0.082245 | | 32.16 | 1.426015 | | 32.153 | -0.082245 |
| 32.166667 | 1.760273 | 1.465675 | 0.695162 | 0.967734 | 0.541827 | 0.787082 | 1.179305 | 0.927673 | 2.513323 | NaN | 2.31592 | -0.205214 | -0.051375 | -0.168383 | 0.607376 | 0.88238 | 0.004777 | 0.146599 | 0.45531 | 0.001522 | 0.062627 | 0.054294 | -0.113762 | | 32.17 | 2.31592 | | 32.167 | -0.113762 |
| 32.180556 | 1.523438 | 1.453677 | 0.67969 | 0.868655 | 0.429289 | 0.644545 | 1.687096 | 0.6494 | 2.509327 | NaN | 1.989834 | -0.281639 | -0.008726 | -0.161534 | 0.491038 | 0.514226 | 0.0161 | 0.053973 | 0.057968 | -0.022789 | -0.069156 | 0.077615 | -0.182317 | | 32.19 | 1.989834 | | 32.181 | -0.182317 |
| 32.194444 | 1.268294 | 1.405586 | 0.601662 | 0.614702 | 0.385435 | 0.575003 | 2.136371 | 0.438377 | 2.2426 | NaN | 1.246406 | -0.398632 | -0.066272 | -0.148492 | 0.248 | 0.021114 | -0.04115 | -0.068262 | -0.217034 | -0.057763 | -0.255952 | 0.025806 | -0.274542 | | 32.2 | 1.246406 | | 32.194 | -0.274542 |
| 32.208333 | 1.149327 | 1.678199 | 0.500682 | 0.252492 | 0.401193 | 0.536832 | 2.312487 | 0.412927 | 2.09586 | NaN | 1.885529 | -0.394593 | -0.130243 | -0.161822 | 0.094814 | -0.304193 | -0.037771 | -0.073258 | -0.251434 | -0.02142 | -0.264198 | -0.017184 | -0.253388 | | 32.22 | 1.885529 | | 32.208 | -0.253388 |
| 32.222222 | 0.918925 | 1.853409 | 0.717352 | -0.106764 | 0.461806 | 0.380771 | 2.547019 | 0.411779 | 1.92116 | NaN | 2.028298 | -0.207938 | -0.055501 | -0.098162 | 0.121232 | -0.303478 | 0.030591 | 0.067104 | -0.008826 | 0.095082 | -0.081322 | 0.0102 | -0.172399 | | 32.23 | 2.028298 | | 32.222 | -0.172399 |
| 32.236111 | 0.944308 | 1.925166 | 1.092215 | -0.118919 | 0.494547 | 0.194612 | 2.614276 | 0.379158 | 1.633756 | NaN | 1.722086 | 0.08058 | 0.097122 | 0.080175 | 0.21556 | 0.094835 | 0.078625 | 0.214994 | 0.316048 | 0.17298 | 0.131843 | 0.014543 | -0.069245 | | 32.24 | 1.722086 | | 32.236 | -0.069245 |
| 32.25 | 1.119192 | 1.782955 | 1.213453 | 0.124723 | 0.489242 | 0.214966 | 2.429473 | 0.301788 | 1.237695 | NaN | 1.388008 | 0.293475 | 0.169338 | 0.282844 | 0.230481 | 0.55062 | 0.063753 | 0.25722 | 0.573764 | 0.155162 | 0.27938 | -0.019287 | 0.040383 | | 32.26 | 1.388008 | | 32.25 | 0.040383 |
| 32.263889 | 1.241807 | 1.544451 | 1.200194 | 0.322862 | 0.477859 | 0.545234 | 2.437708 | 0.219878 | 0.776796 | NaN | 1.14134 | 0.336759 | 0.175233 | 0.409948 | 0.144155 | 0.743464 | 0.03895 | 0.241851 | 0.623693 | 0.112031 | 0.356144 | 0.01148 | 0.159409 | | 32.27 | 1.14134 | | 32.264 | 0.159409 |
| 32.277778 | 1.493524 | 1.644943 | 1.088752 | 0.361541 | 0.566907 | 0.874077 | 2.418913 | 0.316578 | 0.580718 | NaN | 2.059538 | 0.364132 | 0.211499 | 0.48964 | 0.067719 | 0.699505 | 0.038432 | 0.235801 | 0.41854 | 0.127589 | 0.285678 | 0.071725 | 0.235086 | | 32.28 | 2.059538 | | 32.278 | 0.235086 |
| 32.291667 | 1.563756 | 1.788516 | 0.943617 | 0.243939 | 0.631983 | 0.880578 | 2.148538 | 0.386886 | 0.548389 | NaN | 2.428877 | 0.46061 | 0.319144 | 0.610847 | 0.10914 | 0.51605 | 0.082226 | 0.286858 | 0.178046 | 0.206366 | 0.159849 | 0.064762 | 0.265851 | | 32.3 | 2.428877 | | 32.292 | 0.265851 |
| 32.305556 | 1.221607 | 1.776267 | 0.956046 | 0.094502 | 0.447685 | 0.613573 | 1.62753 | 0.264757 | 0.385642 | NaN | 0.534116 | 0.511186 | 0.412406 | 0.785446 | 0.20678 | 0.330793 | 0.107629 | 0.345062 | 0.039469 | 0.252409 | 0.006929 | -0.006257 | 0.247868 | | 32.31 | 0.534116 | | 32.306 | 0.247868 |
| 32.319444 | 1.025263 | 1.60298 | 0.969574 | 0.091756 | 0.225357 | 0.332673 | 1.003647 | 0.065049 | 0.091906 | NaN | -1.376985 | 0.456231 | 0.394858 | 0.983764 | 0.226419 | 0.23422 | 0.068198 | 0.321472 | 0.098755 | 0.203006 | -0.103385 | -0.056879 | 0.092883 | | 32.33 | -1.376985 | | 32.319 | 0.092883 |
| 32.333333 | 1.417994 | 1.638098 | 1.004817 | 0.315921 | 0.290689 | 0.375513 | 0.364885 | 0.132839 | 0.050265 | NaN | -0.378849 | 0.310162 | 0.387624 | 0.981457 | 0.181787 | 0.230569 | 0.061757 | 0.278384 | 0.290294 | 0.14762 | -0.081624 | -0.018679 | -0.02039 | | 32.34 | -0.378849 | | 32.333 | -0.02039 |
| 32.347222 | 2.043661 | 1.817279 | 1.03796 | 0.570203 | 0.519564 | 0.666341 | 0.058462 | 0.411998 | 0.36552 | NaN | 1.909953 | 0.107932 | 0.426146 | 0.726271 | 0.158445 | 0.266027 | 0.111398 | 0.248942 | 0.425023 | 0.105923 | 0.009573 | 0.047201 | 0.022529 | | 32.35 | 1.909953 | | 32.347 | 0.022529 |
| 32.361111 | 2.060826 | 1.745533 | 1.186707 | 0.641298 | 0.538178 | 0.735639 | 0.036193 | 0.39064 | 0.58927 | NaN | 1.399125 | -0.007689 | 0.470962 | 0.308624 | 0.174364 | 0.318798 | 0.115922 | 0.183387 | 0.413138 | 0.045426 | 0.095429 | 0.064461 | 0.166342 | | 32.37 | 1.399125 | | 32.361 | 0.166342 |
| 32.375 | 1.574095 | 1.474799 | 0.967434 | 0.415109 | 0.389437 | 0.411036 | -0.204498 | 0.140778 | 0.498467 | NaN | -0.433052 | 0.062676 | 0.514836 | -0.096925 | 0.168048 | 0.35868 | 0.02972 | 0.028772 | 0.307392 | -0.003523 | 0.1141 | 0.054382 | 0.302754 | | 32.38 | -0.433052 | | 32.375 | 0.302754 |
| 32.388889 | 1.079627 | 1.112243 | 0.266225 | 0.008254 | 0.265144 | -0.129811 | -0.789496 | -0.020914 | 0.270176 | NaN | -0.66574 | 0.286072 | 0.517159 | -0.277628 | 0.151407 | 0.315821 | -0.014901 | -0.0694 | 0.155687 | -0.018076 | 0.098258 | 0.061592 | 0.378056 | | 32.4 | -0.66574 | | 32.389 | 0.378056 |
| 32.402778 | 0.685645 | 0.690048 | -0.254465 | -0.346105 | 0.154522 | -0.683729 | -1.290184 | -0.088984 | 0.074226 | NaN | -0.879945 | 0.504415 | 0.459113 | -0.063661 | 0.191785 | 0.221628 | 0.034024 | 0.000125 | 0.035899 | 0.018638 | 0.105457 | 0.072793 | 0.377219 | | 32.41 | -0.879945 | | 32.403 | 0.377219 |
| 32.416667 | 0.636967 | 0.631398 | -0.354483 | -0.359323 | 0.08426 | -1.019514 | -1.552793 | -0.091481 | -0.072979 | NaN | -0.763266 | 0.579168 | 0.340613 | 0.332298 | 0.207585 | 0.284924 | 0.050003 | 0.126533 | 0.075869 | 0.029207 | 0.084494 | 0.04359 | 0.301094 | | 32.42 | -0.763266 | | 32.417 | 0.301094 |
| 32.430556 | 0.987631 | 1.104962 | -0.266997 | -0.242635 | 0.124827 | -1.104454 | -1.497799 | 0.012868 | -0.228618 | NaN | 1.237539 | 0.585171 | 0.240376 | 0.639306 | 0.174983 | 0.474077 | -0.035521 | 0.085057 | 0.2424 | 0.002185 | 0.001569 | -0.010943 | 0.115558 | | 32.44 | 1.237539 | | 32.431 | 0.115558 |
| 32.444444 | 1.273399 | 1.559976 | 0.403579 | 0.130676 | 0.254176 | -0.693722 | -1.000742 | 0.139252 | -0.019727 | NaN | 1.717408 | 0.773326 | 0.532071 | 0.962748 | 0.281726 | 1.04753 | -0.083916 | 0.287219 | 0.633331 | 0.183907 | 0.049348 | -0.043659 | 0.062737 | | 32.45 | 1.717408 | | 32.444 | 0.062737 |
| 32.458333 | 1.852898 | 2.02594 | 2.015691 | 1.090667 | 0.583177 | 0.253688 | 0.294205 | 0.309174 | 0.771169 | NaN | 0.276345 | 1.317245 | 1.338942 | 1.73794 | 0.701425 | 2.05769 | 0.042146 | 0.82903 | 1.317968 | 0.618628 | 0.177373 | -0.008641 | 0.280879 | | 32.47 | 0.276345 | | 32.458 | 0.280879 |
| 32.472222 | 2.719033 | 1.899782 | 2.6062 | 1.926992 | 0.859052 | 1.344493 | 2.034777 | 0.494171 | 1.414304 | NaN | 0.185418 | 1.51551 | 2.333096 | 2.153143 | 0.954173 | 2.178574 | 0.180001 | 1.068253 | 1.663572 | 0.684883 | 0.083873 | -0.11972 | 0.571263 | | 32.48 | 0.185418 | | 32.472 | 0.571263 |
| 32.486111 | 3.13328 | 1.485665 | 2.308598 | 2.254129 | 0.69902 | 1.779064 | 3.649319 | 0.79484 | 1.853037 | NaN | 1.867285 | 1.493448 | 2.319595 | 2.20003 | 0.757565 | 1.777659 | 0.232276 | 0.837497 | 1.444157 | 0.321931 | 0.10026 | -0.108117 | 0.9129 | | 32.49 | 1.867285 | | 32.486 | 0.9129 |
| 32.5 | 3.136932 | 1.236581 | 2.455046 | 3.370978 | 0.680896 | 1.752973 | 4.375818 | 1.158101 | 2.118336 | NaN | 2.618506 | 1.652728 | 2.199924 | 2.647393 | 0.453406 | 1.520972 | 0.267914 | 0.801209 | 1.160977 | 0.063765 | 0.461043 | 11.651033 | 1.165032 | | 32.5 | 2.618506 | | 32.5 | 1.165032 |
| 32.513889 | 2.702285 | 0.954201 | 2.580285 | 3.749931 | 0.896608 | 1.537357 | 4.04504 | 1.204458 | 1.980531 | NaN | 1.874775 | 1.207561 | 1.618076 | 2.234248 | 0.202586 | 1.10407 | 0.32921 | 0.571769 | 0.874596 | -0.06661 | 0.81944 | 9.742876 | 0.925109 | | 32.51 | 1.874775 | | 32.514 | 0.925109 |
| 32.527778 | 2.103603 | 0.813351 | 2.322117 | 3.120216 | 1.076733 | 1.173132 | 3.085385 | 1.011772 | 1.24362 | NaN | 1.478812 | 0.177006 | 0.857127 | 0.77156 | -0.05386 | 0.310603 | 0.372692 | 0.101721 | 0.018045 | -0.233578 | 1.09086 | 3.916042 | 0.47975 | | 32.53 | 1.478812 | | 32.528 | 0.47975 |
| 32.541667 | 1.331536 | 0.605184 | 2.048359 | 1.74499 | 1.377138 | 0.332715 | 2.295281 | 0.886496 | 0.899906 | NaN | 0.965158 | -0.663666 | 0.212728 | -0.911915 | -0.242037 | -0.355747 | -0.039505 | -0.383086 | -0.875348 | -0.308528 | 1.175391 | -3.284198 | -0.032001 | | 32.54 | 0.965158 | | 32.542 | -0.032001 |
| 32.555556 | 0.679105 | 0.299695 | 1.720206 | 0.815119 | 1.817332 | -0.159195 | 1.894048 | 0.628423 | 0.569937 | NaN | 0.465104 | -0.893092 | -0.198511 | -1.992435 | -0.436054 | -0.60409 | -0.168663 | -0.448492 | -1.044911 | -0.25406 | 1.311586 | -7.718916 | -0.266146 | | 32.56 | 0.465104 | | 32.556 | -0.266146 |
| 32.569444 | 0.562096 | 0.240215 | 1.326358 | 0.086468 | 1.842403 | -0.391433 | 1.482954 | 0.498831 | 0.225302 | NaN | 0.629965 | -0.669725 | -0.431133 | -2.022456 | -0.243252 | -0.594924 | -0.263927 | -0.317597 | -0.567197 | -0.244016 | 0.867735 | -9.993035 | -0.276412 | | 32.57 | 0.629965 | | 32.569 | -0.276412 |
| 32.583333 | 0.473099 | 0.205769 | 1.136409 | -0.556052 | 1.653945 | -0.024803 | 1.226108 | 0.449065 | -0.06311 | NaN | 0.533632 | -0.53071 | -0.629913 | -1.752348 | 0.034859 | -0.572335 | -0.212314 | -0.246388 | -0.279123 | -0.230271 | 0.194178 | -12.001064 | -0.239895 | | 32.58 | 0.533632 | | 32.583 | -0.239895 |
| 32.597222 | 0.276686 | 0.092099 | 0.737944 | -0.669709 | 1.209203 | 0.399766 | 0.92278 | 0.242558 | -0.64509 | NaN | 0.233638 | -0.239862 | -0.86534 | -1.448035 | 0.171868 | -0.689556 | -0.188451 | -0.27831 | -0.375851 | -0.055823 | -0.498918 | -13.723273 | -0.203674 | | 32.6 | 0.233638 | | 32.597 | -0.203674 |
| 32.611111 | -0.057945 | -0.126194 | 0.53041 | -0.741482 | 0.803068 | 0.293887 | -0.022811 | 0.116889 | -1.129679 | NaN | -0.422836 | 0.061117 | -1.079176 | -1.082995 | 0.081415 | -0.968237 | -0.169986 | -0.298361 | -0.851618 | 0.117951 | -1.072355 | -9.504077 | -0.270356 | | 32.61 | -0.422836 | | 32.611 | -0.270356 |
| 32.625 | -0.323252 | -0.261396 | 0.269944 | -0.575171 | 0.518012 | -0.056911 | -1.414123 | 0.062122 | -1.599328 | NaN | -1.176723 | 0.152942 | -1.259933 | -0.88825 | -0.11335 | -1.252844 | -0.063477 | -0.220823 | -1.185901 | 0.16728 | -1.5112 | -3.654003 | -0.492893 | | 32.63 | -1.176723 | | 32.625 | -0.492893 |
| 32.638889 | -0.550059 | -0.272395 | 0.102356 | -0.57754 | 0.247769 | -0.863201 | -2.977438 | -0.014639 | -2.059155 | NaN | -1.663783 | -0.016969 | -1.42049 | -0.881361 | -0.336293 | -1.253368 | -0.021707 | -0.081544 | -1.153099 | 0.065592 | -1.449414 | 10.693792 | -0.789196 | | 32.64 | -1.663783 | | 32.639 | -0.789196 |
| 32.652778 | -0.656177 | -0.291285 | 0.232128 | -0.766294 | 0.129604 | -1.979841 | -3.765336 | -0.052299 | -2.212279 | NaN | -2.001562 | -0.245676 | -1.111212 | -0.739999 | -0.471647 | -1.001091 | 0.054228 | 0.171021 | -0.87635 | -0.066186 | -1.257574 | 13.15474 | -0.74636 | | 32.65 | -2.001562 | | 32.653 | -0.74636 |
| 32.666667 | -0.633882 | -0.276104 | 0.413648 | -0.741791 | 0.181206 | -2.737684 | -3.769226 | 0.197385 | -1.625667 | NaN | -1.958053 | -0.406192 | -0.658435 | -0.301436 | -0.276731 | -0.656356 | 0.219686 | 0.396788 | -0.245595 | -0.078129 | -0.964295 | 10.346995 | -0.46933 | | 32.67 | -1.958053 | | 32.667 | -0.46933 |
| 32.680556 | -0.399135 | -0.325296 | 0.581443 | -0.304691 | 0.318108 | -2.245879 | -2.163759 | 0.511429 | -0.378024 | NaN | -1.052435 | -0.369513 | -0.210398 | 0.290556 | 0.104537 | -0.259374 | 0.365736 | 0.499143 | 0.26395 | -0.049524 | -0.418322 | 11.660673 | 0.045155 | | 32.68 | -1.052435 | | 32.681 | 0.045155 |
| 32.694444 | -0.08337 | -0.469809 | 0.727713 | 0.384147 | 0.330804 | -1.405578 | -0.799194 | 0.565931 | 0.711673 | NaN | -0.30923 | -0.310102 | 0.079836 | 0.648738 | 0.326416 | 0.079189 | 0.364401 | 0.371167 | 0.615725 | -0.013857 | -0.057182 | 7.562526 | 0.336026 | | 32.69 | -0.30923 | | 32.694 | 0.336026 |
| 32.708333 | 0.229923 | -0.535777 | 0.682376 | 1.028858 | 0.176045 | -0.647951 | -0.049105 | 0.33269 | 1.453877 | NaN | 0.578242 | -0.412248 | 0.167826 | 0.713527 | 0.290329 | 0.312792 | 0.303249 | 0.16417 | 0.714439 | 0.047962 | 0.045075 | 2.361341 | 0.385891 | | 32.71 | 0.578242 | | 32.708 | 0.385891 |
| 32.722222 | 0.48372 | -0.432915 | 0.588383 | 1.661772 | -0.124649 | 0.068892 | 0.291861 | -0.057914 | 1.51522 | NaN | 0.970324 | -0.617461 | 0.123535 | 0.604965 | 0.170947 | 0.543184 | 0.153679 | 0.029933 | 0.566777 | 0.097816 | 0.110623 | -5.123753 | 0.293611 | | 32.72 | 0.970324 | | 32.722 | 0.293611 |
| 32.736111 | 0.336773 | -0.483219 | 0.482657 | 1.702174 | -0.639487 | 0.656859 | 0.334896 | -0.361485 | 2.000007 | NaN | 0.578846 | -0.88519 | 0.047417 | 0.117029 | 0.116278 | 0.704684 | -0.025872 | -0.012205 | 0.147323 | 0.133104 | 0.143923 | -10.534586 | -0.045928 | | 32.74 | 0.578846 | | 32.736 | -0.045928 |
| 32.75 | -0.251041 | -0.541385 | 0.385899 | 1.431746 | -1.173751 | 0.715541 | 0.320359 | -0.391579 | 2.149329 | NaN | -0.225945 | -1.216886 | -0.156135 | -0.48082 | 0.053602 | 0.677506 | -0.139978 | -0.030155 | -0.395996 | 0.073314 | 0.063644 | -11.70298 | -0.57765 | | 32.75 | -0.225945 | | 32.75 | -0.57765 |
| 32.763889 | -0.748543 | -0.397986 | 0.253343 | 0.312814 | -1.572903 | 0.797633 | 0.360117 | -0.218908 | 1.626426 | NaN | -0.2593 | -1.369886 | -0.362857 | -0.855926 | -0.009112 | 0.525529 | -0.116784 | 0.149557 | -0.730692 | -0.040994 | -0.0637 | -12.864205 | -0.956457 | | 32.76 | -0.2593 | | 32.764 | -0.956457 |
| 32.777778 | -1.169434 | -0.287251 | 0.014634 | -1.125475 | -1.891106 | 0.525388 | 0.296513 | 0.022371 | 0.699331 | NaN | -0.396394 | -1.141436 | -0.344808 | -0.759028 | 0.088626 | 0.262881 | 0.09406 | 0.275731 | -0.656228 | -0.021857 | -0.022028 | 0.014409 | -0.940009 | | 32.78 | -0.396394 | | 32.778 | -0.940009 |
| 32.791667 | -1.417964 | -0.282118 | -0.33729 | -2.075204 | -2.187091 | 0.013927 | 0.291306 | 0.207559 | -0.27736 | NaN | -1.027186 | -0.51454 | -0.379828 | -0.545258 | 0.261533 | -0.062728 | 0.279192 | 0.343643 | -0.410172 | 0.124731 | -0.001696 | 0.111875 | -0.605154 | | 32.79 | -1.027186 | | 32.792 | -0.605154 |
| 32.805556 | -1.685656 | -0.366762 | -0.659828 | -2.401705 | -2.193184 | -0.565835 | 0.024413 | 0.290127 | -1.243472 | NaN | -1.959822 | 0.13267 | -0.454328 | -0.318136 | 0.327799 | -0.473688 | 0.346831 | 0.287139 | -0.209005 | 0.269976 | -0.045487 | 0.123429 | -0.369975 | | 32.81 | -1.959822 | | 32.806 | -0.369975 |
| 32.819444 | -1.999719 | -0.537668 | -0.96362 | -2.322837 | -1.973821 | -1.051448 | -0.650649 | 0.15861 | -1.440436 | NaN | -2.857657 | 0.511415 | -0.515248 | -0.124048 | 0.202655 | -0.893538 | 0.191742 | 0.181795 | -0.445676 | 0.308714 | -0.09462 | -0.027967 | -0.235169 | | 32.82 | -2.857657 | | 32.819 | -0.235169 |
| 32.833333 | -2.164066 | -0.762138 | -1.025153 | -2.037393 | -1.509102 | -1.285147 | -1.345319 | -0.318504 | -2.040112 | NaN | -3.184347 | 0.656121 | -0.467021 | 0.071227 | 0.070271 | -1.005177 | 0.123968 | 0.155496 | -0.576723 | 0.306885 | -0.026151 | -0.091976 | -0.109431 | | 32.83 | -3.184347 | | 32.833 | -0.109431 |
| 32.847222 | -2.136881 | -0.96178 | -0.890656 | -1.694037 | -0.85756 | -1.539806 | -1.836987 | -0.783433 | -2.852829 | NaN | -3.052971 | 0.673096 | -0.286271 | 0.266917 | -0.013317 | -0.854848 | 0.180323 | 0.213556 | -0.445691 | 0.293663 | 0.052644 | -0.076435 | 0.095303 | | 32.85 | -3.052971 | | 32.847 | 0.095303 |
| 32.861111 | -1.747622 | -0.970658 | -0.680325 | -1.122043 | -0.150443 | -1.889328 | -1.93308 | -0.781211 | -2.804278 | NaN | -2.466881 | 0.547461 | -0.140234 | 0.475733 | -0.079752 | -0.581926 | 0.215908 | 0.200842 | -0.133634 | 0.232357 | 0.059088 | 5.135954 | 0.282936 | | 32.86 | -2.466881 | | 32.861 | 0.282936 |
| 32.875 | -1.106672 | -0.724873 | -0.500288 | -0.706544 | 0.408212 | -1.776147 | -1.416653 | -0.772415 | -3.005306 | NaN | -1.700157 | 0.258766 | -0.072676 | 0.723889 | -0.091599 | -0.314228 | 0.234599 | 0.14743 | 0.110053 | 0.163309 | 0.040069 | 9.886133 | 0.368334 | | 32.88 | -1.700157 | | 32.875 | 0.368334 |
| 32.888889 | -0.490082 | -0.474344 | -0.414958 | -0.287793 | 0.872113 | -1.237407 | -1.02977 | -0.69802 | -2.74708 | NaN | -0.870311 | 0.004212 | -0.05782 | 0.898838 | 0.036617 | -0.061978 | 0.318832 | 0.177344 | 0.244221 | 0.188325 | 0.078529 | 6.848729 | 0.380428 | | 32.89 | -0.870311 | | 32.889 | 0.380428 |
| 32.902778 | 0.044889 | -0.271048 | -0.080262 | 0.310283 | 1.005949 | -0.351727 | -0.603336 | -0.409169 | -1.305582 | NaN | -0.427628 | -0.082427 | 0.033713 | 0.85171 | 0.223892 | 0.088677 | 0.513622 | 0.319565 | 0.33591 | 0.284534 | 0.162162 | -0.32313 | 0.440692 | | 32.9 | -0.427628 | | 32.903 | 0.440692 |
| 32.916667 | 0.474746 | -0.056837 | 0.558627 | 1.157601 | 1.197392 | 0.527537 | 0.039398 | -0.122651 | 0.624616 | NaN | -0.366055 | -0.062065 | 0.099841 | 0.664017 | 0.350956 | 0.153409 | 0.694111 | 0.484777 | 0.36063 | 0.336218 | 0.218129 | -8.335789 | 0.485651 | | 32.92 | -0.366055 | | 32.917 | 0.485651 |
| 32.930556 | 0.790839 | 0.065686 | 0.971373 | 2.006359 | 1.319215 | 1.118568 | 0.912422 | 0.102917 | 2.169722 | NaN | -0.401381 | -0.020915 | 0.06164 | 0.363661 | 0.240743 | 0.104357 | 0.661756 | 0.476646 | 0.298531 | 0.229021 | 0.179356 | -10.552107 | 0.442576 | | 32.93 | -0.401381 | | 32.931 | 0.442576 |
| 32.944444 | 0.803541 | -0.204238 | 0.64073 | 1.871451 | 1.143337 | 0.810757 | 0.950279 | 0.142846 | 2.401631 | NaN | -0.122697 | -0.266407 | -0.321328 | -0.333707 | -0.214254 | -0.374527 | 0.365689 | 0.081186 | -0.387607 | -0.301651 | -0.023395 | -9.681577 | 0.216724 | | 32.94 | -0.122697 | | 32.944 | 0.216724 |
| 32.958333 | -0.19 | -0.866864 | -0.663728 | 0.150408 | 0.202414 | -0.895917 | -0.703266 | -0.113856 | 0.592463 | NaN | 1.277887 | -1.313474 | -1.275895 | -1.789299 | -0.97653 | -1.627158 | -0.165692 | -0.762849 | -2.205072 | -1.340224 | -0.408137 | -6.814324 | -0.5996 | | 32.96 | 1.277887 | | 32.958 | -0.5996 |
| 32.972222 | -2.105242 | -2.525381 | -1.914152 | -2.140652 | -1.022373 | -2.892516 | -2.694702 | -0.668658 | -2.240514 | NaN | 1.338186 | -2.510785 | -2.116972 | -2.667886 | -1.712715 | -2.676846 | -0.306608 | -0.965824 | -3.617429 | -1.867107 | -0.531403 | 0.235939 | -1.767079 | | 32.97 | 1.338186 | | 32.972 | -1.767079 |
| 32.986111 | -3.743397 | -4.276566 | -1.978958 | -3.512474 | -1.726781 | -3.283274 | -3.010683 | -1.005556 | -3.613133 | NaN | -1.047515 | -2.116012 | -1.155835 | -1.507863 | -1.096943 | -1.668266 | -0.124373 | -0.2108 | -2.884858 | -0.566263 | -0.068381 | 0.156342 | -1.77548 | | 32.99 | -1.047515 | | 32.986 | -1.77548 |
| 33 | -4.914501 | -4.551943 | -1.630095 | -3.333432 | -2.207363 | -2.918799 | -1.94246 | -0.983283 | -3.595062 | -1.368476 | -2.860192 | -0.850869 | -0.278793 | -0.191808 | -0.100028 | -0.621354 | 0.179193 | 0.267619 | -1.460954 | 0.405925 | 0.341061 | 0.055712 | -0.97302 | | 33.01 | -2.860192 | | 33 | -0.97302 |
| 33.013889 | -3.894193 | -4.369875 | -1.374683 | -2.445703 | -2.173551 | -2.746 | -1.53111 | -0.848036 | -4.330022 | -2.200235 | -3.501508 | 0.299297 | 0.304308 | 0.433103 | 0.519724 | 0.051872 | 0.272738 | 0.143777 | -0.466983 | 0.429111 | 0.492808 | 0.039592 | 0.106728 | | 33.02 | -3.501508 | | 33.014 | 0.106728 |
| 33.027778 | -1.858063 | -3.807908 | -1.074326 | -1.50288 | -1.47616 | -2.426143 | -1.088929 | -0.712746 | -3.662266 | -2.623912 | -3.685334 | 1.047003 | 0.539045 | 0.604412 | 0.751542 | 0.489041 | 0.297518 | -0.14482 | 0.365369 | 0.252873 | 0.48086 | 0.07794 | 0.878527 | | 33.03 | -3.685334 | | 33.028 | 0.878527 |
| 33.041667 | -0.655987 | -2.664754 | -0.911558 | -0.698184 | -0.806489 | -2.491329 | -0.699828 | -0.641401 | -2.747579 | -2.32231 | -3.097137 | 1.337002 | 0.533983 | 0.650989 | 0.731564 | 0.720361 | 0.242982 | -0.247328 | 0.875793 | 0.282788 | 0.364053 | 0.050267 | 1.254679 | | 33.05 | -3.097137 | | 33.042 | 1.254679 |
| 33.055556 | 0.14288 | -1.473766 | -0.696352 | -0.15752 | -0.081376 | -2.255034 | -0.343505 | -0.372672 | -1.861572 | -0.400402 | -0.971703 | 1.083785 | 0.448622 | 0.736074 | 0.58767 | 0.833371 | 0.216682 | -0.174405 | 1.117884 | 0.319563 | 0.280747 | 0.011851 | 1.492047 | | 33.06 | -0.971703 | | 33.056 | 1.492047 |
| 33.069444 | 0.692422 | -0.091594 | -0.428566 | 0.434842 | 0.479291 | -1.874756 | -0.0126 | -0.009347 | -1.154506 | 1.465287 | 1.217814 | 0.725671 | 0.422632 | 1.009947 | 0.474534 | 0.842901 | 0.281366 | -0.099655 | 1.224441 | 0.229298 | 0.378057 | 0.057745 | 1.766111 | | 33.08 | 1.217814 | | 33.069 | 1.766111 |
| 33.083333 | 0.829974 | 0.505532 | -0.132054 | 0.920704 | 0.777633 | -1.268085 | 0.235926 | 0.062623 | -0.737743 | 1.494047 | 1.221433 | 0.664647 | 0.343356 | 1.245571 | 0.197952 | 0.717177 | 0.281737 | -0.160299 | 1.218923 | 0.105779 | 0.571687 | 0.07387 | 1.919796 | | 33.09 | 1.221433 | | 33.083 | 1.919796 |
| 33.097222 | 0.814555 | 0.485856 | 0.095884 | 1.202214 | 0.962173 | -0.652322 | 0.497419 | 0.008545 | -0.594818 | 0.685225 | 0.322679 | 0.870332 | 0.167506 | 1.184856 | -0.001193 | 0.547592 | 0.185052 | -0.304649 | 1.082785 | 0.114587 | 0.325045 | 0.023052 | 1.50162 | | 33.1 | 0.322679 | | 33.097 | 1.50162 |
| 33.111111 | 1.072757 | 0.646563 | 0.416095 | 1.540607 | 1.153054 | 0.454796 | 0.821979 | 0.120185 | -0.216094 | 0.261624 | 0.163937 | 1.282421 | 0.293312 | 1.193044 | -0.009809 | 0.565084 | 0.146071 | -0.186573 | 1.054447 | 0.073922 | 0.114997 | 0.041725 | 0.872117 | | 33.12 | 0.163937 | | 33.111 | 0.872117 |
| 33.125 | 1.497898 | 0.856844 | 0.875465 | 2.040552 | 1.125727 | 1.593475 | 0.991123 | 0.328984 | 0.517987 | 0.092423 | 0.690932 | 1.075829 | 0.110464 | 1.230943 | 0.100672 | 0.586391 | 0.167075 | -0.165926 | 1.146891 | -0.088442 | 0.045456 | 0.034337 | 0.392944 | | 33.13 | 0.690932 | | 33.125 | 0.392944 |

| | | | | | | | | | | | | | | | | | | | | | | | | | | | | |
|---|---|---|---|---|---|---|---|---|---|---|---|---|---|---|---|---|---|---|---|---|---|---|---|---|---|---|---|---|
| 33.138889 | 1.921559 | 0.761165 | 1.075493 | 1.738072 | 1.241598 | 2.358361 | 1.064097 | 0.486303 | 1.418678 | -0.16017 | 1.057476 | 0.599507 | -0.180632 | 1.373142 | 0.180645 | 0.443311 | 0.074919 | -0.084584 | 1.249559 | -0.208523 | 0.049962 | -0.041439 | 0.23008 | | 33.15 | 1.057476 | 33.139 | 0.23008 |
| 33.152778 | 2.20875 | 0.680866 | 1.002052 | 1.276034 | 1.263901 | 2.728285 | 1.101216 | 0.642041 | 2.359369 | -0.229908 | 1.560227 | 0.256195 | -0.301512 | 1.053353 | 0.274764 | 0.25445 | 0.05276 | 0.14859 | 1.323052 | -0.162804 | 0.13041 | -0.042804 | 0.443065 | | 33.16 | 1.560227 | 33.153 | 0.443065 |
| 33.166667 | 2.222959 | 0.428052 | 0.934735 | 0.96383 | 1.170588 | 2.753598 | 1.070535 | 0.760651 | 3.093558 | -0.132219 | 2.077804 | 0.130106 | -0.303402 | 0.32238 | 0.329296 | 0.103088 | 0.038158 | 0.348139 | 1.120472 | -0.069216 | 0.220768 | 0.00685 | 0.752701 | | 33.17 | 2.077804 | 33.167 | 0.752701 |
| 33.180556 | 1.847826 | 0.033651 | 0.802522 | 0.763948 | 0.859099 | 2.500848 | 0.887121 | 0.831152 | 3.533142 | -0.547633 | 2.009822 | 0.006334 | -0.308785 | -0.502127 | 0.240787 | -0.160773 | -0.033491 | 0.311623 | 0.439786 | -0.006329 | 0.210188 | 0.015809 | 0.754627 | | 33.19 | 2.009822 | 33.181 | 0.754627 |
| 33.194444 | 1.500434 | -0.270396 | 0.639664 | 0.506616 | 0.515695 | 2.003459 | 0.724817 | 0.930527 | 3.287387 | -1.072067 | 1.818554 | -0.227475 | -0.345934 | -0.979084 | 0.127777 | -0.42099 | -0.119818 | 0.153975 | -0.175191 | -0.038062 | 0.193695 | 0.007183 | 0.347894 | | 33.2 | 1.818554 | 33.194 | 0.347894 |
| 33.208333 | 1.143586 | -0.162551 | 0.417853 | 0.187822 | 0.261788 | 1.731035 | 0.563463 | 1.109032 | 2.711771 | -0.522895 | 2.285113 | -0.499155 | -0.305955 | -0.92812 | 0.102696 | -0.66917 | -0.068681 | 0.010511 | -0.442515 | -0.068004 | 0.229696 | -0.005012 | -0.163489 | | 33.22 | 2.285113 | 33.208 | -0.163489 |
| 33.222222 | 0.611833 | -0.149465 | 0.136722 | -0.148324 | 0.001714 | 1.062423 | 0.279109 | 1.087587 | 2.547802 | -0.154062 | 2.335983 | -0.570309 | -0.320421 | -0.855116 | 0.135382 | -0.740548 | -0.024037 | -0.086748 | -0.58098 | -0.10335 | 0.194002 | -0.042982 | -0.460631 | | 33.23 | 2.335983 | 33.222 | -0.460631 |
| 33.236111 | 0.027275 | -0.497891 | 0.005039 | -0.321236 | -0.127913 | 0.242606 | 0.036482 | 0.759217 | 2.732695 | -0.787058 | 1.046639 | -0.498064 | -0.308258 | -0.594435 | 0.163497 | -0.548271 | 0.008779 | -0.059628 | -0.337805 | -0.184314 | 0.175525 | -0.057918 | -0.605996 | | 33.24 | 1.046639 | 33.236 | -0.605996 |
| 33.25 | -0.152783 | -0.148967 | 0.058256 | -0.263762 | -0.098287 | -0.333906 | -0.181517 | 0.660754 | 2.685031 | -0.412199 | 0.981669 | -0.208674 | -0.09044 | -0.239181 | 0.250676 | -0.097828 | 0.073974 | 0.199598 | 0.253949 | -0.087941 | 0.204486 | -0.015343 | -0.106414 | | 33.26 | 0.981669 | 33.25 | -0.106414 |
| 33.263889 | 0.092777 | 0.821187 | 0.270927 | -0.041511 | 0.081286 | -0.342575 | -0.142394 | 0.631763 | 2.313206 | 0.641034 | 2.022409 | 0.128721 | 0.210164 | 0.06252 | 0.306084 | 0.384065 | 0.097548 | 0.367862 | 0.806487 | 0.088161 | 0.223382 | 0.018915 | 0.392719 | | 33.27 | 2.022409 | 33.264 | 0.392719 |
| 33.277778 | 0.287389 | 1.048121 | 0.381404 | 0.112118 | 0.23622 | -0.244943 | -0.083095 | 0.420077 | 1.321012 | 0.274937 | 1.586896 | 0.331857 | 0.377956 | 0.212432 | 0.319884 | 0.803629 | 0.04034 | 0.339353 | 1.062702 | 0.169716 | 0.209326 | 0.025179 | 0.579025 | | 33.28 | 1.586896 | 33.278 | 0.579025 |
| 33.291667 | 0.518767 | 0.885372 | 0.433421 | 0.258591 | 0.426819 | -0.153892 | 0.175213 | 0.353818 | 1.095274 | -0.179194 | 1.260849 | 0.388018 | 0.466174 | 0.280126 | 0.325895 | 1.027821 | -0.053326 | 0.314017 | 0.936545 | 0.230045 | 0.243117 | 0.031947 | 0.601179 | | 33.3 | 1.260849 | 33.292 | 0.601179 |
| 33.305556 | 0.941372 | 1.466648 | 0.455666 | 0.408759 | 0.652703 | -0.144712 | 0.293544 | 0.361442 | 1.150794 | 0.622223 | 2.275676 | 0.349806 | 0.438636 | 0.33165 | 0.293092 | 0.965684 | -0.175012 | 0.253651 | 0.512169 | 0.360997 | 0.209631 | 0.006196 | 0.494402 | | 33.31 | 2.275676 | 33.306 | 0.494402 |
| 33.319444 | 1.01657 | 1.700539 | 0.404963 | 0.375565 | 0.692848 | -0.479847 | 0.131137 | 0.260479 | 0.848736 | 0.936344 | 2.125035 | 0.316157 | 0.147546 | 0.386154 | 0.193089 | 0.683932 | -0.23123 | 0.128508 | 0.083497 | 0.410663 | 0.150285 | -0.026358 | 0.363674 | | 33.33 | 2.125035 | 33.319 | 0.363674 |
| 33.333333 | 0.822908 | 1.312223 | 0.5032 | 0.253216 | 0.524589 | -1.069023 | -0.098367 | 0.182514 | 0.226533 | 0.344645 | 0.845225 | 0.421444 | -0.125879 | 0.434686 | 0.155546 | 0.379365 | -0.161485 | 0.095479 | -0.196166 | 0.39396 | 0.138751 | 0.015123 | 0.270174 | | 33.34 | 0.845225 | 33.333 | 0.270174 |
| 33.347222 | 0.780929 | 1.087222 | 0.728378 | 0.279859 | 0.381025 | -1.304094 | -0.082824 | 0.146571 | -0.261388 | 0.035107 | 0.048823 | 0.570589 | -0.144764 | 0.512406 | 0.13549 | 0.219986 | -0.090181 | 0.147027 | -0.224008 | 0.337344 | 0.134506 | 0.050507 | 0.266368 | | 33.35 | 0.048823 | 33.347 | 0.266368 |
| 33.361111 | 0.905528 | 1.09953 | 0.90027 | 0.320005 | 0.301764 | -1.408676 | 0.059034 | 0.10336 | -0.455421 | 0.000935 | -0.16596 | 0.619499 | -0.002316 | 0.48616 | 0.10103 | 0.218141 | -0.092859 | 0.148273 | -0.05241 | 0.21769 | 0.091991 | 0.019894 | 0.244256 | | 33.37 | -0.16596 | 33.361 | 0.244256 |
| 33.375 | 0.926822 | 1.019394 | 0.968123 | 0.27133 | 0.191497 | -1.333211 | 0.2117 | 0.078037 | -0.412333 | -0.171984 | 0.0311 | 0.546608 | 0.185441 | 0.467853 | 0.144123 | 0.373683 | -0.051569 | 0.139082 | 0.323674 | 0.122221 | 0.103217 | 0.023229 | 0.282566 | | 33.38 | 0.0311 | 33.375 | 0.282566 |
| 33.388889 | 0.99992 | 0.955273 | 0.974012 | 0.310089 | 0.173435 | -1.026262 | 0.346955 | 0.101971 | -0.310366 | -0.258344 | 0.284931 | 0.434252 | 0.327056 | 0.378193 | 0.214065 | 0.559479 | 0.016488 | 0.145882 | 0.651764 | 0.106154 | 0.116532 | 0.038341 | 0.237931 | | 33.4 | 0.284931 | 33.389 | 0.237931 |
| 33.402778 | 1.029146 | 0.902236 | 0.818399 | 0.234088 | 0.134264 | -1.120986 | 0.253781 | 0.007555 | -0.274123 | -0.207546 | 0.297027 | 0.311007 | 0.386443 | 0.256418 | 0.233713 | 0.580907 | -0.017323 | 0.143026 | 0.719472 | 0.089252 | 0.058355 | -0.003797 | 0.183759 | | 33.41 | 0.297027 | 33.403 | 0.183759 |
| 33.416667 | 0.82683 | 0.541677 | 0.576171 | 0.081022 | -0.026903 | -1.59304 | 0.011607 | -0.101487 | -0.385733 | -0.415101 | -0.411432 | 0.266105 | 0.388142 | 0.290049 | 0.266086 | 0.508654 | -0.011946 | 0.170503 | 0.641907 | 0.097531 | 0.032547 | 0.009709 | 0.217186 | | 33.42 | -0.411432 | 33.417 | 0.217186 |
| 33.430556 | 0.506725 | 0.296415 | 0.511706 | 0.016345 | -0.136292 | -1.836489 | -0.16804 | -0.079908 | -0.512268 | -0.539459 | -1.090878 | 0.349461 | 0.493241 | 0.437612 | 0.288931 | 0.443511 | 0.073181 | 0.237985 | 0.611616 | 0.141759 | 0.084841 | 0.099387 | 0.306724 | | 33.44 | -1.090878 | 33.431 | 0.306724 |
| 33.444444 | 0.399707 | 0.958991 | 0.763751 | 0.429395 | -0.074141 | -1.540821 | -0.001047 | 0.101389 | -0.19872 | -0.10791 | -0.350014 | 0.527244 | 0.830486 | 0.726085 | 0.301583 | 0.745722 | 0.150352 | 0.406053 | 1.039029 | 0.282579 | 0.115338 | 0.117789 | 0.24707 | | 33.45 | -0.350014 | 33.444 | 0.24707 |
| 33.458333 | 0.651487 | 2.424832 | 1.463057 | 1.243415 | 0.311875 | -0.293156 | 0.667346 | 0.449999 | 0.517337 | 0.431303 | 0.682016 | 0.981447 | 1.572687 | 1.267568 | 0.476762 | 1.656932 | 0.288597 | 0.725576 | 2.349395 | 0.662321 | 0.234588 | 0.083823 | 0.204783 | | 33.47 | 0.682016 | 33.458 | 0.204783 |
| 33.472222 | 1.458478 | 3.564491 | 2.203797 | 2.122619 | 0.731209 | 0.588339 | 1.418797 | 0.76899 | 1.478971 | 0.707794 | 1.302233 | 1.348512 | 1.850093 | 1.785655 | 0.76568 | 2.251846 | 0.31232 | 0.807282 | 2.646429 | 0.936042 | 0.339754 | 0.027834 | 0.432428 | | 33.48 | 1.302233 | 33.472 | 0.432428 |
| 33.486111 | 2.181524 | 2.745991 | 2.338331 | 2.615995 | 1.002854 | 1.200624 | 1.97421 | 0.874241 | 2.143286 | 0.918346 | 1.802129 | 1.432054 | 1.441906 | 2.03507 | 0.932798 | 2.104326 | 0.195341 | 0.565206 | 2.594157 | 0.721111 | 0.206047 | 0.052095 | 0.830256 | | 33.49 | 1.802129 | 33.486 | 0.830256 |
| 33.5 | 2.7069 | 2.268517 | 2.163577 | 2.800639 | 0.876984 | 1.537639 | 1.609849 | 0.855036 | 2.875505 | 0.890409 | 2.142249 | 1.226198 | 1.181227 | 2.04017 | 0.688804 | 1.953994 | -0.029775 | 0.36157 | 2.814725 | 0.511348 | -0.096323 | 5.59464 | 0.94822 | | 33.5 | 2.142249 | 33.5 | 0.94822 |
| 33.513889 | 2.757392 | 2.38924 | 1.671492 | 2.276725 | 1.161302 | 1.472441 | 1.44966 | 0.831981 | 3.054784 | 1.275738 | 1.835785 | 0.683819 | 1.059104 | 1.713129 | 0.424016 | 1.693267 | -0.139345 | -0.055039 | 2.401165 | 0.337478 | -0.190417 | 9.104715 | 0.44987 | | 33.51 | 1.835785 | 33.514 | 0.44987 |
| 33.527778 | 2.679938 | 2.623429 | 1.297065 | 1.949599 | 1.196041 | 1.724263 | 0.9402 | 0.763711 | 2.394697 | 2.215121 | 1.776361 | -0.02209 | 1.007957 | 0.805391 | 0.097632 | 1.254141 | -0.20268 | -0.425684 | 1.462477 | -0.118437 | 0.067676 | 9.299542 | -0.145592 | | 33.53 | 1.776361 | 33.528 | -0.145592 |
| 33.541667 | 2.344456 | 2.546292 | 1.009988 | 1.586851 | 1.04755 | 1.501119 | 0.222239 | 0.65982 | 1.641607 | 2.127852 | 1.694998 | -0.484434 | 1.033591 | -0.001858 | -0.147227 | 0.817219 | -0.277179 | -0.680812 | 0.665601 | -0.580508 | 0.589195 | 7.58983 | -0.547138 | | 33.54 | 1.694998 | 33.542 | -0.547138 |
| 33.555556 | 1.705866 | 2.549412 | 0.745991 | 1.183251 | 1.101846 | 1.206909 | -0.206573 | 0.646882 | 1.063344 | 0.49462 | 1.634736 | -0.410698 | 1.016846 | -0.436169 | -0.440646 | 0.464682 | -0.214067 | -0.858662 | 0.199328 | -0.561142 | 0.8762 | 4.68606 | -0.26867 | | 33.56 | 1.634736 | 33.556 | -0.26867 |
| 33.569444 | 1.044556 | 2.251278 | 0.608802 | 0.765718 | 1.049198 | 1.135946 | -0.293047 | 0.705788 | 0.585109 | NaN | 1.169246 | -0.132817 | 0.89422 | -0.539127 | -0.287631 | 0.140951 | -0.014144 | -0.454808 | -0.137036 | -0.525425 | 1.139132 | 1.553918 | 0.004966 | | 33.57 | 1.169246 | 33.569 | 0.004966 |
| 33.583333 | 0.372264 | 1.451283 | 0.435475 | 0.577186 | 0.786963 | 1.070773 | -0.029418 | 0.5117 | -0.06304 | NaN | 0.494764 | 0.023591 | 0.736592 | -0.689438 | -0.050312 | -0.245504 | 0.127673 | -0.090471 | -0.420436 | -0.50837 | 0.969211 | -2.112816 | 0.023133 | | 33.58 | 0.494764 | 33.583 | 0.023133 |
| 33.597222 | -0.188574 | 0.317246 | 0.411267 | 0.232495 | 0.302157 | 0.818529 | 0.391038 | -0.046905 | -0.794916 | NaN | -0.223336 | -0.085755 | 0.551452 | -0.713551 | 0.060364 | -0.595651 | 0.122449 | -0.021504 | -0.639815 | -0.377034 | 0.552406 | -6.23204 | -0.020219 | | 33.6 | -0.223336 | 33.597 | -0.020219 |
| 33.611111 | -0.905781 | -0.516512 | 0.417561 | -0.262743 | -0.359692 | 0.266512 | 0.563068 | -0.611912 | -1.499299 | NaN | -0.721828 | -0.305224 | 0.355342 | -0.572556 | 0.013011 | -0.802154 | 0.021049 | -0.015958 | -0.745463 | -0.143396 | 0.112996 | -5.636398 | -0.208984 | | 33.61 | -0.721828 | 33.611 | -0.208984 |
| 33.625 | -1.220519 | -1.148214 | 0.377017 | -0.635435 | -0.934492 | -0.246862 | 0.534961 | -1.070855 | -2.00622 | NaN | -1.005273 | -0.512109 | 0.176734 | -0.458444 | -0.141232 | -0.958277 | -0.081307 | -0.070629 | -0.906794 | 0.012332 | -0.320089 | -4.529257 | -0.514591 | | 33.63 | -1.005273 | 33.625 | -0.514591 |
| 33.638889 | -1.211855 | -1.637831 | 0.271366 | -0.71724 | -1.176901 | -0.648455 | 0.456384 | -1.663942 | -2.699146 | NaN | -1.622008 | -0.569336 | 0.009549 | -0.38312 | -0.379737 | -1.144418 | -0.248537 | -0.183655 | -1.046651 | 0.000704 | -0.720307 | -2.456402 | -0.759882 | | 33.64 | -1.622008 | 33.639 | -0.759882 |
| 33.652778 | -0.989571 | -1.721749 | 0.260323 | -0.577956 | -1.092738 | -0.877271 | 0.314909 | -2.057141 | -3.200421 | NaN | -2.607427 | -0.281084 | -0.078393 | -0.159054 | -0.452446 | -1.20055 | -0.2045 | -0.179671 | -1.069896 | -0.028428 | -0.82937 | 0.743041 | -0.728726 | | 33.65 | -2.607427 | 33.653 | -0.728726 |
| 33.666667 | -0.916646 | -1.640885 | 0.318786 | -0.188178 | -0.885427 | -0.746649 | 0.230372 | -1.923494 | -3.130205 | NaN | -2.792959 | 0.138036 | -0.043139 | 0.234192 | -0.229542 | -0.883923 | 0.011242 | -0.027229 | -0.88653 | -0.004554 | -0.635906 | 1.59932 | -0.386806 | | 33.67 | -2.792959 | 33.667 | -0.386806 |
| 33.680556 | -0.777096 | -1.759559 | 0.532805 | 0.26678 | -0.579682 | -0.602263 | 0.391066 | -1.237941 | -2.598982 | NaN | -2.246539 | 0.421013 | -0.017169 | 0.639998 | 0.132747 | -0.531243 | 0.230715 | 0.09369 | -0.487423 | 0.015871 | -0.443223 | 0.503252 | 0.014074 | | 33.68 | -2.246539 | 33.681 | 0.014074 |
| 33.694444 | -0.425222 | -1.511888 | 0.859765 | 0.919807 | -0.280326 | -0.538036 | 0.586427 | -0.490496 | -1.430798 | NaN | -1.470396 | 0.503292 | -0.050195 | 0.897434 | 0.257136 | -0.228832 | 0.304788 | 0.140638 | -0.043173 | 0.01383 | -0.234854 | 1.723315 | 0.223008 | | 33.69 | -1.470396 | 33.694 | 0.223008 |
| 33.708333 | -0.505022 | -1.186433 | 1.076116 | 1.304414 | -0.069911 | -0.355033 | 0.659241 | -0.082859 | 0.192793 | NaN | -0.861922 | 0.403413 | -0.054382 | 0.938135 | 0.218265 | 0.011534 | 0.31869 | 0.222818 | 0.32335 | 0.072887 | 0.033765 | 2.889218 | 0.237103 | | 33.71 | -0.861922 | 33.708 | 0.237103 |
| 33.722222 | -0.600084 | -0.801848 | 1.169193 | 1.30721 | -0.065955 | -0.093566 | 0.669082 | 0.025044 | 1.671384 | -0.21154 | -0.593624 | 0.290473 | -0.003247 | 0.820418 | 0.13549 | 0.204599 | 0.384156 | 0.362532 | 0.493137 | 0.132325 | 0.184444 | 0.004738 | 0.127304 | | 33.72 | -0.593624 | 33.722 | 0.127304 |
| 33.736111 | -0.724027 | -0.654524 | 1.105764 | 1.0548 | -0.2545 | 0.282224 | 0.57917 | 0.075076 | 2.50818 | -0.822997 | -0.222092 | 0.157842 | -0.008791 | 0.561017 | 0.065214 | 0.209755 | 0.367341 | 0.346872 | 0.306573 | 0.101394 | 0.149517 | -4.731934 | -0.044308 | | 33.74 | -0.222092 | 33.736 | -0.044308 |
| 33.75 | -0.841927 | -0.313832 | 0.818015 | 0.627123 | -0.731231 | 0.649824 | 0.36587 | 0.180998 | 2.507263 | 0.012815 | 0.42465 | -0.05227 | -0.127442 | 0.2791 | 0.02799 | 0.04299 | 0.341162 | 0.222558 | -0.032808 | -0.014139 | -0.053399 | -10.540323 | -0.200631 | | 33.75 | 0.42465 | 33.75 | -0.200631 |
| 33.763889 | -1.56781 | -0.60025 | 0.404818 | -0.094339 | -1.395762 | 0.842942 | -0.091558 | 0.293228 | 2.083381 | 0.358042 | 0.311133 | -0.334277 | -0.233225 | -0.050527 | 0.041774 | -0.077036 | 0.298867 | 0.101084 | -0.331024 | -0.145899 | -0.180957 | 0.015814 | -0.273247 | | 33.76 | 0.311133 | 33.764 | -0.273247 |
| 33.777778 | -2.739018 | -1.722564 | -0.206791 | -1.053983 | -2.182486 | 0.872488 | -0.526652 | 0.244384 | 1.320492 | 0.227095 | -0.871777 | -0.603849 | -0.308023 | -0.451129 | 0.124376 | -0.029258 | 0.11582 | 0.062885 | -0.580739 | -0.205339 | -0.137425 | 0.101864 | -0.264634 | | 33.78 | -0.871777 | 33.778 | -0.264634 |
| 33.791667 | -3.686317 | -3.087549 | -0.814053 | -1.756849 | -2.875422 | 0.752795 | -0.63147 | -0.013182 | 0.066866 | -0.173698 | -2.122704 | -0.917568 | -0.508152 | -0.871668 | 0.118282 | 0.13033 | 0.041901 | -0.024749 | -0.780517 | -0.234612 | -0.035659 | 0.116411 | -0.336505 | | 33.79 | -2.122704 | 33.792 | -0.336505 |
| 33.805556 | -4.095457 | -3.41866 | -1.167753 | -2.180365 | -3.710799 | 0.650126 | -0.419645 | -0.347947 | -1.201912 | -0.632142 | -2.739297 | -1.113413 | -0.883967 | -1.204191 | -0.039351 | 0.139921 | -0.000994 | -0.094187 | -0.80219 | -0.281449 | 0.093019 | -0.018557 | -0.453043 | | 33.81 | -2.739297 | 33.806 | -0.453043 |
| 33.819444 | -3.625499 | -3.305207 | -1.262275 | -2.507548 | -4.290124 | 0.47137 | -0.162277 | -0.616108 | -2.544174 | -0.528472 | -3.096457 | -1.10425 | -1.117788 | -1.536147 | -0.228455 | -0.022206 | -0.01145 | -0.058107 | -0.612442 | -0.247034 | 0.194933 | -0.100936 | -0.465689 | | 33.82 | -3.096457 | 33.819 | -0.465689 |
| 33.833333 | -3.470982 | -3.799037 | -1.319227 | -2.697441 | -4.097785 | 0.272725 | -0.042276 | -0.73545 | -3.863717 | -0.502921 | -3.232306 | -0.806668 | -1.071583 | -1.103521 | -0.295959 | -0.278354 | 0.103435 | 0.08646 | -0.286582 | -0.073326 | 0.259138 | -0.029819 | -0.296124 | | 33.83 | -3.232306 | 33.833 | -0.296124 |
| 33.847222 | -2.990414 | -4.201798 | -1.390557 | -2.476399 | -3.446172 | 0.014067 | -0.063994 | -0.710758 | -4.634564 | -0.636635 | -3.214586 | -0.425802 | -0.899098 | -0.585116 | -0.244086 | -0.501758 | -0.069202 | 0.059906 | 0.013586 | 0.14566 | 0.217447 | -0.014592 | -0.081626 | | 33.85 | -3.214586 | 33.847 | -0.081626 |
| 33.861111 | -2.08405 | -4.159824 | -1.114422 | -2.066809 | -2.379764 | -0.241696 | -0.117777 | -0.5812 | -4.645121 | -0.441353 | -3.022662 | -0.101436 | -0.762666 | -0.136826 | -0.171804 | -0.703087 | -0.24066 | 0.019385 | 0.190661 | 0.251072 | 0.100003 | -0.112258 | 0.086875 | | 33.86 | -3.022662 | 33.861 | 0.086875 |
| 33.875 | -1.397983 | -3.462212 | -0.728 | -1.444679 | -0.996755 | -0.458723 | -0.181601 | -0.53295 | -4.282653 | -0.377457 | -2.699663 | 0.173814 | -0.623226 | 0.333015 | -0.068032 | -0.653861 | -0.267017 | 0.015369 | 0.335896 | 0.295703 | 0.01229 | -0.3629 | 0.214669 | | 33.88 | -2.699663 | 33.875 | 0.214669 |
| 33.888889 | -0.700566 | -2.861657 | -0.209737 | -0.637033 | 0.451682 | -0.32976 | -0.129428 | -0.427537 | -4.072814 | -0.399094 | -2.408106 | 0.391071 | -0.443469 | 0.845908 | 0.137012 | -0.503951 | -0.04389 | 0.109946 | 0.431682 | 0.321639 | 0.036969 | -0.364143 | 0.297885 | | 33.89 | -2.408106 | 33.889 | 0.297885 |
| 33.902778 | 0.037537 | -2.359582 | 0.321057 | 0.324031 | 1.957594 | -0.019764 | 0.224659 | -0.213335 | -3.160793 | -0.37271 | -1.723896 | 0.527604 | -0.360652 | 1.117493 | 0.264413 | -0.331567 | 0.123049 | 0.086196 | 0.490772 | 0.322752 | 0.174415 | -0.087728 | 0.372796 | | 33.9 | -1.723896 | 33.903 | 0.372796 |
| 33.916667 | 0.695221 | -1.669767 | 0.776042 | 1.267224 | 2.826437 | 0.626218 | 0.645816 | 0.069642 | -2.078063 | -0.205253 | -0.499833 | 0.554439 | -0.213146 | 1.159644 | 0.209479 | -0.261617 | 0.164795 | -0.04612 | 0.440515 | 0.305713 | 0.314479 | 0.04484 | 0.354991 | | 33.92 | -0.499833 | 33.917 | 0.354991 |
| 33.930556 | 0.896783 | -1.179748 | 0.986965 | 1.89009 | 1.79378 | 1.246099 | 0.983374 | 0.3142 | -1.114113 | -0.100541 | 0.157509 | 0.415897 | -0.197822 | 0.934994 | 0.081053 | -0.285071 | 0.038125 | -0.270685 | 0.239573 | 0.175968 | 0.298774 | 0.084849 | 0.198223 | | 33.93 | 0.157509 | 33.931 | 0.198223 |
| 33.944444 | 0.32007 | -1.361187 | 0.424526 | 1.708429 | 0.698739 | 0.650205 | 0.765451 | 0.013076 | -1.369155 | -0.165027 | -0.172106 | -0.221093 | -0.653308 | 0.064038 | -0.086628 | -0.713552 | -0.350302 | -0.899558 | -0.573086 | -0.499766 | -0.080117 | 0.117403 | -0.213055 | | 33.94 | -0.172106 | 33.944 | -0.213055 |
| 33.958333 | -1.480071 | -3.083865 | -1.219874 | -0.446812 | -0.397104 | -1.409879 | -0.287304 | -0.792201 | -3.686259 | -0.591861 | -1.759192 | -1.734698 | -1.963164 | -1.79107 | -0.907374 | -1.838945 | -0.939462 | -2.082796 | -2.248233 | -2.129584 | -0.900326 | 0.191196 | -1.35136 | | 33.96 | -1.759192 | 33.958 | -1.35136 |
| 33.972222 | -3.926768 | -5.210303 | -2.287295 | -3.05727 | -2.044503 | -1.950739 | -1.138696 | -1.343752 | -4.830693 | -1.063716 | -3.574208 | -3.1251 | -3.255064 | -3.201962 | -1.655404 | -2.852561 | -1.150658 | -2.599447 | -3.40974 | -3.329913 | -1.417856 | 0.186264 | -3.171449 | | 33.97 | -3.574208 | 33.972 | -3.171449 |
| 33.986111 | -4.274013 | -6.141394 | -2.463041 | -3.476463 | -2.873664 | -1.792827 | -0.994801 | -1.34511 | -4.792632 | -1.012617 | -3.377561 | -2.590836 | -1.641435 | -1.927442 | -0.822612 | -1.70852 | -0.493137 | -1.179171 | -1.982869 | -1.7754 | -0.686076 | 0.158962 | -2.906631 | | 33.99 | -3.377561 | 33.986 | -2.906631 |
| 34 | -3.852819 | -4.471153 | -2.444304 | -2.126838 | -2.665841 | -1.770383 | -0.773373 | -1.061649 | -4.884678 | -0.352234 | -2.118755 | -1.067705 | -0.249301 | 0.092708 | 0.302293 | -0.101057 | 0.277526 | 0.258299 | 0.271854 | 0.162069 | 0.310011 | 0.11235 | -0.696278 | | 34.01 | -2.118755 | 34 | -0.696278 |
| 34.013889 | -3.73633 | -3.582321 | -1.845874 | -1.502237 | -2.357156 | -1.436038 | -0.461919 | -1.002345 | -4.149059 | -0.302858 | -1.76111 | 0.055335 | 0.509102 | 0.795299 | 0.764211 | 0.623487 | 0.431577 | 0.73531 | 1.275433 | 1.116193 | 0.72598 | 0.006751 | 0.867556 | | 34.02 | -1.76111 | 34.014 | 0.867556 |
| 34.027778 | -2.951631 | -2.771062 | -1.249045 | -0.943636 | -1.485209 | -1.134943 | -0.15965 | -0.943063 | -3.34049 | -0.426183 | -1.724497 | 1.007367 | 0.822983 | 1.168976 | 0.773544 | 0.798244 | 0.522835 | 0.649378 | 1.445981 | 1.353411 | 0.732252 | -0.014305 | 1.51837 | | 34.03 | -1.724497 | 34.028 | 1.51837 |
| 34.041667 | -1.69243 | -2.308492 | -0.692349 | -0.401766 | -0.642348 | -1.023573 | 0.085908 | -0.736632 | -2.265422 | -0.500408 | -1.503018 | 1.580123 | 0.869407 | 1.472206 | 1.159507 | 0.790336 | 0.724364 | 0.58212 | 1.456992 | 1.417583 | 0.667499 | 0.066695 | 1.893522 | | 34.05 | -1.503018 | 34.042 | 1.893522 |
| 34.055556 | -0.684188 | -1.683471 | -0.381026 | 0.004623 | 0.027997 | -1.083387 | 0.472314 | -0.500251 | -1.455713 | -0.192381 | -0.339105 | 1.357628 | 0.715724 | 1.511588 | 1.748749 | 0.710397 | 0.80108 | 0.509311 | 1.293652 | 1.421728 | 0.437712 | 0.090174 | 1.960783 | | 34.06 | -0.339105 | 34.056 | 1.960783 |
| 34.069444 | -0.235091 | -1.146248 | -0.171611 | 0.171081 | 0.454248 | -1.007775 | 0.651457 | -0.260278 | -0.822934 | 0.29656 | 0.872032 | 1.043194 | 0.591381 | 1.009113 | 1.17593 | 0.6813 | 0.623767 | 0.485556 | 1.079272 | 1.202362 | 0.255022 | 0.072542 | 1.777713 | | 34.08 | 0.872032 | 34.069 | 1.777713 |
| 34.083333 | 0.111659 | -0.742979 | 0.274611 | 0.418281 | 0.837559 | -0.766595 | 0.828741 | -0.117995 | -0.419141 | 0.552049 | 1.076748 | 0.789721 | 0.73082 | 0.610517 | 0.683357 | 0.849712 | 0.635655 | 0.538323 | 1.089991 | 1.003336 | 0.23701 | 0.091848 | 1.774896 | | 34.09 | 1.076748 | 34.083 | 1.774896 |
| 34.097222 | 0.69631 | -0.310188 | 0.639517 | 0.746439 | 1.031718 | -0.412545 | 0.872555 | -0.056225 | -0.001024 | 0.688802 | 0.739357 | 0.78159 | 0.7902 | 0.359765 | 0.585179 | 0.945402 | 0.53517 | 0.552644 | 1.157362 | 0.912651 | 0.213299 | 0.085915 | 1.744566 | | 34.1 | 0.739357 | 34.097 | 1.744566 |
| 34.111111 | 1.407953 | -0.065744 | 0.793965 | 1.139652 | 0.498169 | 0.017109 | 0.705161 | -0.087709 | 0.261588 | 0.357318 | 0.212542 | 0.948169 | 0.604208 | 0.125883 | 0.439666 | 1.070528 | 0.314676 | 0.366902 | 1.092062 | 0.731389 | 0.116559 | 0.027325 | 1.526101 | | 34.12 | 0.212542 | 34.111 | 1.526101 |
| 34.125 | 1.880326 | 0.078542 | 1.019644 | 1.541047 | 0.55329 | 0.537935 | 0.701117 | -0.022737 | 0.524264 | -0.012054 | 0.02403 | 1.290415 | 0.395097 | 0.230852 | 0.180953 | 1.171283 | 0.182578 | 0.246584 | 0.857605 | 0.389524 | 0.059301 | -0.000278 | 1.64521 | | 34.13 | 0.02403 | 34.125 | 1.64521 |
| 34.138889 | 1.805218 | 0.522518 | 1.270021 | 1.72953 | 0.905734 | 1.233563 | 1.00333 | 0.23359 | 0.963678 | 0.094386 | 1.338288 | 1.72197 | 0.41323 | 0.880554 | 0.018541 | 1.023022 | 0.150439 | 0.292477 | 0.820662 | 0.056478 | 0.205919 | 0.030156 | 1.799037 | | 34.15 | 1.338288 | 34.139 | 1.799037 |
| 34.152778 | 1.62099 | 0.818901 | 1.308234 | 2.006021 | 1.292799 | 2.033253 | 1.364849 | 0.385483 | 1.37552 | 0.388413 | 2.527754 | 1.841247 | 0.288196 | 1.125166 | -0.040019 | 0.733618 | 0.104588 | 0.295198 | 0.721367 | -0.260796 | 0.226598 | 0.019222 | 1.710948 | | 34.16 | 2.527754 | 34.153 | 1.710948 |
| 34.166667 | 1.207374 | 0.680942 | 1.054219 | 2.219623 | 1.351547 | 3.227984 | 1.338608 | 0.292929 | 1.333037 | 0.277458 | 2.215337 | 1.365716 | -0.063997 | 0.991022 | -0.016263 | 0.223028 | 0.008465 | 0.153137 | 0.509239 | -0.418605 | 0.165852 | -0.041572 | 1.241958 | | 34.17 | 2.215337 | 34.167 | 1.241958 |
| 34.180556 | 0.779935 | 0.208274 | 0.783391 | 2.157902 | 1.030163 | 3.588087 | 1.116232 | 0.150001 | 1.002293 | -0.087564 | 1.561802 | 0.617561 | -0.199985 | 0.375713 | -0.011654 | -0.203777 | 0.027061 | -0.014818 | 0.261243 | -0.63532 | 0.097305 | -0.110075 | 0.532347 | | 34.19 | 1.561802 | 34.181 | 0.532347 |
| 34.194444 | 0.236663 | -0.956168 | 0.62285 | 1.379716 | 0.2009 | 3.21992 | 0.816503 | 0.082734 | 1.065882 | -0.720405 | 1.175672 | 0.067412 | -0.231317 | 0.045469 | 0.098715 | -0.427093 | 0.077821 | -0.148045 | 0.061138 | -0.743167 | 0.035172 | -0.18175 | -0.040174 | | 34.2 | 1.175672 | 34.194 | -0.040174 |
| 34.208333 | -0.590456 | -3.159487 | 0.529375 | 0.857532 | -1.03103 | 2.146155 | 0.405541 | 0.125567 | 1.4406 | -1.026044 | 0.584068 | -0.1515 | -0.172258 | 0.181079 | 0.236125 | -0.312189 | 0.093872 | -0.103322 | -0.03492 | -0.547875 | 0.15896 | -0.025942 | -0.310964 | | 34.22 | 0.584068 | 34.208 | -0.310964 |
| 34.222222 | -0.75074 | -3.794431 | 0.25257 | 0.212605 | -1.86377 | 1.507278 | 0.156558 | 0.237393 | 1.908014 | -0.849524 | 0.234639 | -0.141362 | -0.179427 | 0.269811 | 0.31081 | -0.169504 | 0.155612 | 0.062965 | -0.044494 | -0.298947 | 0.313894 | 0.222232 | -0.357938 | | 34.23 | 0.234639 | 34.222 | -0.357938 |
| 34.236111 | -0.24646 | -2.27164 | -0.114703 | 0.279051 | -1.639591 | 1.031369 | 0.22656 | 0.334994 | 2.077788 | -0.165944 | 0.349714 | -0.087359 | -0.223499 | 0.121796 | 0.25461 | -0.155546 | 0.139036 | 0.12945 | -0.09497 | -0.10011 | 0.291986 | 0.198822 | -0.4016 | | 34.24 | 0.349714 | 34.236 | -0.4016 |
| 34.25 | 0.14416 | -1.598067 | -0.267289 | 0.159616 | -1.379393 | 0.597167 | 0.288194 | 0.361327 | 1.890145 | -0.283031 | 0.43666 | -0.18641 | -0.382666 | 0.01902 | 0.281251 | -0.073484 | 0.094393 | 0.033934 | -0.110934 | 0.03132 | 0.218227 | 0.028118 | -0.36508 | | 34.26 | 0.43666 | 34.25 | -0.36508 |
| 34.263889 | 0.208639 | -1.621595 | -0.190298 | -0.144765 | -1.390405 | 0.044969 | 0.33532 | 0.336577 | 1.805903 | -0.328447 | 0.471438 | -0.316223 | -0.41995 | -0.054047 | 0.34086 | 0.086137 | 0.108096 | 0.043565 | -0.004701 | 0.155405 | 0.236025 | -0.008678 | -0.22482 | | 34.27 | 0.471438 | 34.264 | -0.22482 |
| 34.277778 | 0.072986 | -1.629243 | -0.188405 | -0.404709 | -1.243043 | -0.764384 | 0.361414 | 0.298669 | 1.73888 | NaN | 0.670422 | -0.3959 | -0.355088 | -0.178101 | 0.556903 | 0.303262 | 0.212501 | 0.201449 | 0.164552 | 0.238414 | 0.266195 | 0.064912 | -0.072534 | | 34.28 | 0.670422 | 34.278 | -0.072534 |
| 34.291667 | 0.004094 | -1.444678 | -0.158781 | -0.33891 | -1.085198 | -1.79475 | 0.282152 | 0.196912 | 1.822771 | NaN | 0.79939 | -0.327718 | -0.175656 | -0.071665 | 0.59935 | 0.485582 | 0.217012 | 0.294364 | 0.263391 | 0.295249 | 0.246802 | 0.073876 | 0.06121 | | 34.3 | 0.79939 | 34.292 | 0.06121 |
| 34.305556 | -0.045756 | -1.255599 | -0.146521 | -0.150614 | -1.002981 | -2.506743 | 0.149675 | 0.058067 | 2.039084 | NaN | 0.965902 | -0.085631 | 0.054168 | 0.2669 | 0.629012 | 0.567918 | 0.104056 | 0.27844 | 0.346059 | 0.403653 | 0.207893 | 0.01073 | 0.186668 | | 34.31 | 0.965902 | 34.306 | 0.186668 |
| 34.319444 | -0.421065 | -1.562128 | -0.071397 | -0.209259 | -1.057773 | -2.570532 | -0.133509 | -0.056199 | 2.297877 | NaN | 0.9915 | 0.307192 | 0.33744 | 0.661621 | 0.669737 | 0.592666 | 0.080036 | 0.276399 | 0.405729 | 0.564488 | 0.270254 | 0.046511 | 0.392703 | | 34.33 | 0.9915 | 34.319 | 0.392703 |
| 34.333333 | -0.954865 | -2.402187 | 0.17907 | -0.635533 | -1.622595 | -2.110009 | -0.230921 | -0.155388 | 2.693628 | NaN | 0.475144 | 0.674111 | 0.521058 | 0.88561 | 0.585539 | 0.514839 | 0.115017 | 0.267977 | 0.378001 | 0.659038 | 0.333171 | 0.141021 | 0.59379 | | 34.34 | 0.475144 | 34.333 | 0.59379 |
| 34.347222 | -0.838687 | -2.262739 | 0.408277 | -0.671219 | -1.600527 | -1.149091 | -0.133261 | -0.21379 | 2.445923 | -1.949198 | 0.020439 | 0.727471 | 0.444643 | 0.776833 | 0.176235 | 0.213898 | 0.028651 | 0.14267 | 0.242234 | 0.561195 | 0.257875 | 0.099016 | 0.583681 | | 34.35 | 0.020439 | 34.347 | 0.583681 |
| 34.361111 | -0.190678 | -1.224907 | 0.410711 | -0.158503 | -0.888392 | -0.349871 | 0.313451 | -0.073148 | 1.825676 | -0.587962 | -0.159403 | 0.587825 | 0.310032 | 0.530442 | -0.285883 | -0.126499 | -0.046145 | 0.039603 | 0.148256 | 0.307531 | 0.170714 | 0.007748 | 0.44376 | | 34.37 | -0.159403 | 34.361 | 0.44376 |
| 34.375 | -0.027279 | -0.688296 | 0.301123 | 0.05562 | -0.935764 | -0.149595 | 0.621806 | 0.012919 | 1.68554 | 0.016609 | -0.734446 | 0.594832 | 0.309924 | 0.428274 | -0.368395 | -0.249276 | 0.027355 | 0.109445 | 0.16054 | 0.167255 | 0.179458 | 0.045269 | 0.281659 | | 34.38 | -0.734446 | 34.375 | 0.281659 |
| 34.388889 | -0.314284 | -0.670881 | 0.042903 | 0.105726 | -1.670289 | -0.450794 | 0.578586 | -0.080598 | 1.527927 | -0.021347 | -1.783525 | 0.644303 | 0.367534 | 0.468069 | -0.179925 | -0.193537 | 0.092203 | 0.183588 | 0.158813 | 0.039782 | 0.133813 | 0.116025 | 0.139873 | | 34.4 | -1.783525 | 34.389 | 0.139873 |
| 34.402778 | -0.537024 | -0.512977 | -0.213911 | 0.288901 | -1.774103 | -0.828466 | 0.394191 | -0.257728 | 1.011985 | 0.113545 | -1.886416 | 0.715438 | 0.341898 | 0.393368 | -0.019691 | -0.049026 | 0.092361 | 0.182641 | 0.235912 | -0.079078 | 0.048902 | 0.107416 | 0.12193 | | 34.41 | -1.886416 | 34.403 | 0.12193 |
| 34.416667 | -0.20612 | -0.476247 | -0.18935 | 0.219508 | -1.135486 | -0.900188 | 0.301432 | -0.27301 | 0.678225 | 0.070041 | -0.95189 | 0.87303 | 0.308481 | 0.514936 | 0.140692 | 0.179158 | 0.133734 | 0.203141 | 0.362751 | -0.033526 | -0.000528 | 0.068957 | 0.272394 | | 34.42 | -0.95189 | 34.417 | 0.272394 |
| 34.430556 | 0.080905 | -0.098007 | 0.155783 | 0.276811 | -0.312699 | -0.531187 | 0.473563 | -0.175557 | 0.652877 | 0.026322 | -0.354436 | 0.99131 | 0.289556 | 0.691382 | 0.21462 | 0.425155 | 0.257945 | 0.25469 | 0.550965 | 0.095527 | -0.06962 | 0.047809 | 0.457882 | | 34.44 | -0.354436 | 34.431 | 0.457882 |
| 34.444444 | 0.573324 | 0.618718 | 0.708319 | 0.592708 | 0.09297 | -0.125913 | 0.7854 | -0.094088 | 0.898793 | 0.030579 | -0.692152 | 1.088627 | 0.503391 | 0.982136 | 0.1658 | 0.756852 | 0.270011 | 0.368288 | 0.703459 | 0.30906 | -0.022472 | -0.014243 | 0.474788 | | 34.45 | -0.692152 | 34.444 | 0.474788 |
| 34.458333 | 1.256995 | 1.731054 | 1.603568 | 1.310777 | 0.545149 | 0.615703 | 1.254802 | 0.046628 | 1.857688 | 0.016935 | -1.306115 | 1.383384 | 1.251304 | 1.575676 | 0.15888 | 1.462183 | 0.270162 | 0.574111 | 1.290855 | 0.792962 | 0.238093 | -0.105143 | 0.29159 | | 34.47 | -1.306115 | 34.458 | 0.29159 |
| 34.472222 | 2.007974 | 2.88103 | 2.533312 | 2.233331 | 1.319804 | 2.0657 | 1.79769 | 0.35015 | 3.094038 | 0.037482 | -0.518528 | 1.560222 | 1.630055 | 1.812829 | 0.400301 | 2.233384 | 0.322194 | 0.732476 | 2.317463 | 1.234537 | 0.521899 | -0.088281 | 0.391137 | | 34.48 | -0.518528 | 34.472 | 0.391137 |
| 34.486111 | 2.831293 | 3.67295 | 3.387066 | 2.972525 | 2.000585 | 2.632243 | 2.248429 | 0.668236 | 3.675761 | 0.020302 | 1.149834 | 1.389892 | 1.408492 | 1.574769 | 0.633489 | 2.237034 | 0.404818 | 0.627733 | 2.039772 | 1.288888 | 0.529269 | 0.023185 | 0.843839 | | 34.49 | 1.149834 | 34.486 | 0.843839 |
| 34.5 | 3.329201 | 4.414964 | 3.412571 | 3.367244 | 2.232727 | 2.776269 | 2.364773 | 0.600496 | 4.138336 | 0.096331 | 2.64924 | 1.193346 | 1.126557 | 1.393652 | 0.622984 | 1.957092 | 0.426068 | 0.372695 | 1.458712 | 0.950259 | 0.35191 | 0.095964 | 0.949355 | | 34.5 | 2.64924 | 34.5 | 0.949355 |
| 34.513889 | 3.078135 | 5.347975 | 2.601438 | 3.084925 | 2.065163 | 2.610835 | 1.96112 | 0.400751 | 4.213203 | 0.172831 | 3.007158 | 0.865996 | 1.120026 | 1.220674 | 0.388417 | 1.701263 | 0.345155 | 0.198111 | 1.208857 | 0.685676 | 0.151954 | 0.221586 | 0.653242 | | 34.51 | 3.007158 | 34.514 | 0.653242 |
| 34.527778 | 2.266453 | 5.79653 | 1.625306 | 1.956765 | 1.831685 | 2.868307 | 1.307858 | 0.514119 | 4.508533 | 0.234649 | 3.075496 | 0.408929 | 1.36848 | 0.961737 | 0.218045 | 1.308862 | -0.00331 | 0.011737 | 0.77998 | 0.185146 | -0.093494 | 0.330246 | 0.109922 | | 34.53 | 3.075496 | 34.528 | 0.109922 |
| 34.541667 | 1.429498 | 5.68836 | 0.699126 | 0.781481 | 1.287587 | 2.585379 | 0.66518 | 0.732643 | 4.413524 | 0.223424 | 2.656514 | -0.101225 | 1.387853 | 0.504432 | 0.034067 | 1.080837 | -0.359683 | -0.217161 | 0.352381 | -0.54676 | -0.303854 | 0.231709 | -0.400336 | | 34.54 | 2.656514 | 34.542 | -0.400336 |

| | | | | | | | | | | | | | | | | | | | | | | | | | | | | |
|---|---|---|---|---|---|---|---|---|---|---|---|---|---|---|---|---|---|---|---|---|---|---|---|---|---|---|---|---|
| 36.194444 | 2.298072 | 1.296381 | 2.284543 | 1.097706 | 2.767229 | 0.958818 | 0.776739 | 1.354251 | 1.157085 | 0.157397 | 3.667733 | 0.483747 | 0.494058 | 0.41466 | -0.141706 | 0.293907 | 0.081721 | 0.142521 | 0.290503 | -1.033317 | -0.095467 | 4.883104 | 0.177681 | | 36.2 | 3.667733 | 36.194 | 0.177681 |
| 36.208333 | 2.254519 | 1.296755 | 1.725681 | 1.072479 | 3.395824 | 0.394627 | 0.743052 | 1.188833 | 1.406533 | 0.122728 | 3.827554 | 0.418402 | 0.435274 | 0.237426 | -0.384375 | 0.30824 | -0.037873 | 0.176589 | 0.272925 | -1.404144 | -0.241397 | 6.026801 | 0.180977 | | 36.22 | 3.827554 | 36.208 | 0.180977 |
| 36.222222 | 2.193917 | 0.989699 | 1.292191 | 1.061743 | 3.743155 | 0.389259 | 0.705317 | 1.418493 | 1.361743 | 0.078704 | 3.042607 | 0.427809 | 0.408878 | 0.023559 | -0.523718 | 0.469461 | -0.025283 | 0.245008 | 0.153166 | -1.459388 | -0.178821 | 5.061691 | 0.097826 | | 36.23 | 3.042607 | 36.222 | 0.097826 |
| 36.236111 | 2.136007 | 0.761982 | 0.999288 | 1.114252 | 4.052017 | 0.528849 | 0.693893 | 1.004625 | 0.854246 | 0.042503 | 2.495677 | 0.45259 | 0.41249 | -0.119087 | -0.51324 | 0.700939 | 0.094396 | 0.305569 | 0.010989 | -1.455266 | 0.094247 | 2.7611 | 0.117394 | | 36.24 | 2.495677 | 36.236 | 0.117394 |
| 36.25 | 2.329864 | 0.858147 | 0.908494 | 1.245797 | 3.646914 | 0.76045 | 0.782713 | 1.106354 | 0.493636 | 0.069295 | 1.710819 | 0.508494 | 0.464696 | -0.074884 | -0.275735 | 0.796185 | 0.251234 | 0.363078 | 0.031178 | -1.402538 | 0.359276 | 1.656649 | 0.123841 | | 36.26 | 1.710819 | 36.25 | 0.123841 |
| 36.263889 | 2.147967 | 1.166838 | 0.915818 | 1.372821 | 2.569708 | 0.249291 | 0.854777 | 0.813147 | 0.211957 | 0.078998 | 1.130057 | 0.50966 | 0.468876 | 0.101663 | 0.010523 | 0.693367 | 0.388491 | 0.424642 | 0.154897 | -0.993733 | 0.447775 | 1.82915 | 0.253352 | | 36.27 | 1.130057 | 36.264 | 0.253352 |
| 36.277778 | 1.914614 | 1.274796 | 0.816012 | 1.402124 | 1.293632 | -0.125275 | 0.892314 | -0.138116 | 0.050749 | 0.008871 | 0.615118 | 0.520339 | 0.398516 | 0.186969 | 0.160354 | 0.50299 | 0.400061 | 0.354962 | 0.261844 | -0.610577 | 0.356819 | 2.33573 | 0.406925 | | 36.28 | 0.615118 | 36.278 | 0.406925 |
| 36.291667 | 1.59335 | 1.255138 | 0.705503 | 1.369933 | 0.157881 | -0.335596 | 0.886892 | -0.921712 | -0.153259 | -0.019942 | 0.161835 | 0.441589 | 0.239926 | 0.22505 | 0.160824 | 0.282283 | 0.268175 | 0.234987 | 0.307931 | -0.233025 | 0.151448 | 2.177107 | 0.365565 | | 36.3 | 0.161835 | 36.292 | 0.365565 |
| 36.305556 | 1.302232 | 1.248909 | 0.69758 | 1.322456 | -0.625677 | -0.477369 | 0.966852 | -1.505351 | -0.221556 | -0.035825 | -0.104164 | 0.389748 | 0.152166 | 0.297722 | 0.116588 | 0.181538 | 0.192147 | 0.15382 | 0.323759 | 0.048496 | -0.059853 | 1.915723 | 0.277457 | | 36.31 | -0.104164 | 36.306 | 0.277457 |
| 36.319444 | 1.073864 | 1.293413 | 0.740062 | 1.290336 | -1.091887 | -0.528936 | 1.118809 | -1.99284 | -0.237042 | -0.027238 | -0.156377 | 0.43448 | 0.193955 | 0.391155 | 0.118958 | 0.2553 | 0.231236 | 0.161529 | 0.3061 | 0.251955 | -0.137242 | 2.112646 | 0.260284 | | 36.33 | -0.156377 | 36.319 | 0.260284 |
| 36.333333 | 1.002347 | 1.263354 | 0.825905 | 1.234811 | -1.296243 | -0.713693 | 1.23842 | -2.133544 | -0.23399 | -0.051997 | -0.207676 | 0.564175 | 0.317889 | 0.427691 | 0.168752 | 0.352934 | 0.268673 | 0.159429 | 0.312887 | 0.430361 | -0.067782 | 3.686191 | 0.317208 | | 36.34 | -0.207676 | 36.333 | 0.317208 |
| 36.347222 | 0.948067 | 1.05839 | 0.826798 | 1.144666 | -1.272989 | -0.796296 | 1.293436 | -1.878327 | -0.304011 | -0.043579 | -0.278074 | 0.644799 | 0.399514 | 0.431915 | 0.24625 | 0.418487 | 0.214085 | 0.094209 | 0.297621 | 0.661125 | 0.053651 | 5.546234 | 0.364714 | | 36.35 | -0.278074 | 36.347 | 0.364714 |
| 36.361111 | 0.923398 | 0.808404 | 0.7979 | 1.095528 | -1.276026 | -0.718865 | 1.234553 | -1.543409 | -0.391186 | -0.106739 | -0.226682 | 0.643233 | 0.371645 | 0.470956 | 0.277577 | 0.469617 | 0.077847 | 0.012004 | 0.248049 | 0.849778 | 0.137767 | 4.2833 | 0.358115 | | 36.37 | -0.226682 | 36.361 | 0.358115 |
| 36.375 | 0.940075 | 0.646099 | 0.790872 | 1.112301 | -1.519874 | -0.901054 | 1.13515 | -1.454615 | -0.469251 | -0.259383 | -0.290117 | 0.567815 | 0.278925 | 0.490211 | 0.31798 | 0.412026 | -0.013453 | -0.020675 | 0.210834 | 1.027271 | 0.102262 | 1.508017 | 0.375832 | | 36.38 | -0.290117 | 36.375 | 0.375832 |
| 36.388889 | 1.159461 | 0.570474 | 0.799955 | 1.137035 | -2.027843 | -1.159512 | 1.128394 | -1.514088 | -0.66661 | -0.378512 | -0.419877 | 0.510836 | 0.231988 | 0.498826 | 0.343562 | 0.262691 | -0.020179 | 0.009691 | 0.219342 | 1.049538 | -0.024057 | -1.35456 | 0.379236 | | 36.4 | -0.419877 | 36.389 | 0.379236 |
| 36.402778 | 1.025888 | 0.472109 | 0.777397 | 1.11542 | -2.119893 | -1.393523 | 1.106767 | -1.735874 | -1.002143 | -0.466567 | -0.560973 | 0.41903 | 0.19561 | 0.504269 | 0.262069 | 0.129273 | -0.025076 | 0.056274 | 0.246334 | 1.032981 | -0.146513 | -3.822783 | 0.345937 | | 36.41 | -0.560973 | 36.403 | 0.345937 |
| 36.416667 | 0.605573 | 0.334621 | 0.71017 | 1.072074 | -2.112486 | -1.557348 | 1.031655 | -1.835962 | -1.149982 | -0.488847 | -0.787012 | 0.339422 | 0.193971 | 0.455101 | 0.025898 | 0.057371 | -0.141442 | 0.011544 | 0.212734 | 0.854251 | -0.247121 | -4.078444 | 0.283685 | | 36.42 | -0.787012 | 36.417 | 0.283685 |
| 36.430556 | 0.616806 | 0.395362 | 0.654701 | 1.160851 | -1.899171 | -1.496382 | 0.99399 | -1.936419 | -1.095361 | -0.439753 | -0.832252 | 0.504538 | 0.238311 | 0.479666 | -0.174682 | 0.087632 | -0.2337 | -0.042381 | 0.190316 | 0.912436 | -0.167473 | -2.845659 | 0.27966 | | 36.44 | -0.832252 | 36.431 | 0.27966 |
| 36.444444 | 1.310942 | 1.315021 | 0.993533 | 1.408642 | -0.87679 | -1.009186 | 1.262569 | -2.012274 | -0.688813 | -0.064728 | -1.099565 | 1.385352 | 0.776121 | 1.049265 | -0.139768 | 0.377414 | -0.147666 | 0.117062 | 0.575075 | 1.640322 | 0.070023 | -0.821615 | 0.86628 | | 36.45 | -1.099565 | 36.444 | 0.86628 |
| 36.458333 | 2.896782 | 3.398552 | 1.737621 | 1.872909 | 0.942775 | 0.251919 | 1.897978 | -1.58076 | 0.368665 | 0.68577 | -1.110195 | 3.225653 | 2.252691 | 2.182894 | 0.165209 | 1.044488 | 0.243844 | 0.500704 | 1.506292 | 3.039192 | 0.630862 | 1.733359 | 1.594575 | | 36.47 | -1.110195 | 36.458 | 1.594575 |
| 36.472222 | 3.901688 | 5.12186 | 1.932326 | 1.901729 | 2.206905 | 1.154379 | 2.274495 | 0.003234 | 1.583664 | 0.873518 | 0.025062 | 3.868769 | 2.129676 | 2.753543 | 0.688339 | 1.640202 | 0.726183 | 0.787714 | 2.029644 | 3.223121 | 1.388637 | 4.393831 | 1.916208 | | 36.48 | 0.025062 | 36.472 | 1.916208 |
| 36.486111 | 2.908767 | 3.799983 | 1.387372 | 1.257387 | 2.734542 | 1.063013 | 2.020731 | 1.178815 | 2.111645 | 0.537417 | 1.538939 | 2.219997 | 1.104609 | 1.788655 | 1.116275 | 1.82715 | 0.944982 | 0.701628 | 1.466115 | 2.080403 | 1.961795 | 5.671357 | 1.475247 | | 36.49 | 1.538939 | 36.486 | 1.475247 |
| 36.5 | 1.619255 | 2.169008 | 0.859707 | 0.771951 | 3.714753 | 0.773039 | 1.622227 | 1.09343 | 1.308256 | 0.24549 | 2.632545 | 1.162529 | 0.606984 | 1.020574 | 1.309633 | 1.565477 | 0.901802 | 0.603425 | 0.835344 | 1.733814 | 1.703216 | 4.407546 | 1.159379 | | 36.5 | 2.632545 | 36.5 | 1.159379 |
| 36.513889 | 1.508816 | 1.507085 | 0.578345 | 0.75542 | 3.712789 | 0.81043 | 1.33029 | 1.0474 | 1.241774 | 0.112828 | 3.247707 | 0.584886 | 0.719831 | 0.691862 | 1.119765 | 1.155501 | 0.706492 | 0.398026 | 0.547904 | 1.563925 | 0.942766 | 9.808185 | 0.836383 | | 36.51 | 3.247707 | 36.514 | 0.836383 |
| 36.527778 | 1.874857 | 1.565283 | 0.487072 | 1.225283 | 3.805011 | 0.923302 | 1.259745 | 1.326943 | 1.65213 | 0.043702 | 3.591701 | 0.245621 | 0.555979 | 0.568745 | 0.525441 | 0.683394 | 0.412196 | -0.032863 | 0.463835 | 0.947292 | 0.144336 | 3.283171 | 0.304511 | | 36.53 | 3.591701 | 36.528 | 0.304511 |
| 36.541667 | 2.234266 | 1.747949 | 0.538022 | 1.999187 | 4.285488 | 1.35442 | 1.692679 | 0.899489 | 1.498207 | 0.017641 | 4.01695 | 0.038802 | -0.126482 | 0.440178 | -0.046566 | 0.187965 | 0.162342 | -0.472717 | 0.386518 | -0.096839 | -0.811202 | -1.576941 | -0.270231 | | 36.54 | 4.01695 | 36.542 | -0.270231 |
| 36.555556 | 2.30306 | 1.762622 | 0.596855 | 2.543786 | 3.494524 | 0.998845 | 2.037208 | 0.041655 | 0.704827 | -0.0744 | 3.466039 | 0.050785 | -0.057647 | 0.339852 | -0.346429 | -0.225443 | 0.084764 | -0.656866 | 0.319955 | -0.969132 | -1.57523 | -9.110714 | -0.769638 | | 36.56 | 3.466039 | 36.556 | -0.769638 |
| 36.569444 | 2.105551 | 1.647022 | 0.719558 | 3.129033 | 3.260328 | 0.923087 | 1.88383 | -0.838111 | 0.012567 | -0.258223 | 2.612639 | 0.306427 | 0.104906 | 0.258417 | -0.195929 | -0.386006 | 0.239105 | -0.569857 | 0.256379 | -1.228348 | -1.660857 | -9.669109 | -0.911262 | | 36.57 | 2.612639 | 36.569 | -0.911262 |
| 36.583333 | 1.843052 | 1.313859 | 0.964995 | 3.483381 | 2.714869 | 0.974576 | 2.229892 | -1.281311 | -0.803043 | -0.413667 | 2.001472 | 0.571163 | 0.169635 | 0.306654 | 0.347013 | -0.554566 | 0.489788 | -0.273604 | 0.273068 | -1.511922 | -1.286269 | -11.356137 | -0.577838 | | 36.58 | 2.001472 | 36.583 | -0.577838 |
| 36.597222 | 1.474764 | 0.805673 | 1.302451 | 3.133095 | 1.789991 | 0.957196 | 2.154 | -0.972885 | -1.228481 | -0.53119 | 1.449551 | 0.560545 | 0.195482 | 0.392725 | 0.711309 | -0.373436 | 0.742288 | 0.044772 | 0.238704 | -1.770451 | -0.514198 | -5.24257 | -0.411686 | | 36.6 | 1.449551 | 36.597 | -0.411686 |
| 36.611111 | 1.106113 | 0.28923 | 1.418309 | 1.605653 | 0.851874 | 0.254203 | 0.879024 | -0.724135 | -0.813367 | -0.678993 | 0.345295 | 0.317344 | 0.093441 | 0.38391 | 0.656887 | -0.54584 | 0.663897 | 0.157682 | 0.218049 | -2.228895 | -0.390611 | 1.038995 | -0.627796 | | 36.61 | 0.345295 | 36.611 | -0.627796 |
| 36.625 | 0.449986 | -0.195623 | 0.934637 | -0.078201 | -0.147872 | -0.698824 | -0.478482 | -0.859708 | -0.091427 | -0.749632 | -1.036983 | -0.033104 | -0.053822 | 0.247752 | 0.095724 | -0.831318 | 0.387542 | 0.081737 | 0.146135 | -2.406174 | -0.833002 | 7.066887 | -0.707716 | | 36.63 | -1.036983 | 36.625 | -0.707716 |
| 36.638889 | 0.111561 | -0.67902 | 0.604203 | -1.442153 | -1.131609 | -1.273505 | -1.484046 | -1.826285 | 0.113057 | -0.787635 | -2.607352 | -0.384246 | -0.196186 | 0.082264 | -0.594414 | -1.081565 | 0.188399 | -0.100761 | 0.141374 | -2.689556 | -1.108124 | 15.795937 | -0.660652 | | 36.64 | -2.607352 | 36.639 | -0.660652 |
| 36.652778 | -0.192235 | -1.071785 | 0.457796 | -1.795924 | -2.251499 | -1.759212 | -1.68004 | -3.122314 | -0.097028 | -0.866283 | -3.635396 | -0.694043 | 0.16401 | -0.079211 | -0.818672 | -1.155725 | 0.043432 | -0.33337 | 0.108436 | -2.198934 | -1.008833 | 13.322347 | -0.705898 | | 36.65 | -3.635396 | 36.653 | -0.705898 |
| 36.666667 | -0.495465 | -1.40578 | 0.522219 | -1.929188 | -3.699763 | -1.153681 | -0.751166 | -4.189514 | -0.624109 | -1.026096 | -4.083797 | -0.856271 | 1.978341 | -0.162227 | -0.588549 | -1.186557 | 0.188812 | -0.3925 | 0.084963 | -1.475304 | -0.675741 | 10.244423 | -0.83656 | | 36.67 | -4.083797 | 36.667 | -0.83656 |
| 36.680556 | -0.497152 | -1.487054 | 0.74733 | -1.482529 | -3.740975 | -1.43232 | -0.249166 | -3.528417 | -0.314406 | -1.08445 | -4.486759 | -0.810867 | 0.690823 | -0.090332 | -0.202434 | -0.92752 | 0.32299 | -0.268129 | 0.135658 | -1.231378 | 0.0333 | 2.202401 | -0.826147 | | 36.68 | -4.486759 | 36.681 | -0.826147 |
| 36.694444 | -0.206629 | -1.359778 | 1.123471 | -0.96797 | -4.253216 | -1.203328 | 0.290178 | -1.594516 | -0.045422 | -0.91984 | -4.354478 | -0.687875 | -1.153978 | 0.01924 | 0.32011 | -0.585584 | 0.563227 | -0.01277 | 0.116751 | -0.93823 | 0.536798 | -2.983587 | -0.64699 | | 36.69 | -4.354478 | 36.694 | -0.64699 |
| 36.708333 | 0.060778 | -1.052719 | 1.367128 | -0.531363 | -4.056092 | -1.849321 | 0.511471 | 0.939466 | 0.377729 | -0.671152 | -3.84517 | -0.507235 | -1.551467 | -0.063227 | 0.677198 | -0.255023 | 0.630202 | 0.224044 | -0.073139 | -0.484713 | 0.462726 | -9.452938 | -0.809499 | | 36.71 | -3.84517 | 36.708 | -0.809499 |
| 36.722222 | 0.134628 | -0.775937 | 1.225017 | -0.162303 | -4.310247 | -1.848565 | 0.654118 | 2.883279 | 0.321352 | -0.519311 | -2.793006 | -0.394899 | -0.261062 | -0.211345 | 0.71398 | -0.118243 | 0.455353 | 0.191008 | -0.275145 | 0.13838 | -0.283214 | -10.761543 | -0.552848 | | 36.72 | -2.793006 | 36.722 | -0.552848 |
| 36.736111 | 0.030556 | -0.679876 | 0.789911 | 0.361353 | -3.61241 | -1.746342 | 0.640631 | 3.910125 | 0.518976 | -0.511467 | -1.71829 | -0.303024 | -0.223394 | -0.302679 | 0.347396 | -0.117968 | 0.190343 | 0.097064 | -0.318652 | 0.855541 | -0.788895 | -4.681241 | -0.285563 | | 36.74 | -1.71829 | 36.736 | -0.285563 |
| 36.75 | -0.151925 | -0.667622 | 0.525265 | 0.976014 | -2.155669 | -1.513379 | 0.509581 | 3.557846 | 0.232447 | -0.506986 | -0.670551 | -0.264401 | -0.259227 | -0.322841 | -0.176569 | 0.163859 | -0.071999 | 0.041916 | -0.281891 | 1.472358 | -0.785449 | 3.446627 | -0.397169 | | 36.75 | -0.670551 | 36.75 | -0.397169 |
| 36.763889 | -0.204608 | -0.65711 | 0.350175 | 1.541277 | -1.324671 | -1.399617 | 0.203563 | 3.06472 | -0.475744 | -0.542655 | -0.167897 | -0.339443 | -0.330302 | -0.278455 | -0.433446 | 0.621966 | -0.441784 | -0.011746 | -0.193425 | 1.818184 | -0.365943 | 9.160737 | -0.313834 | | 36.76 | -0.167897 | 36.764 | -0.313834 |
| 36.777778 | 0.087254 | -0.659446 | -0.03086 | 1.94721 | -0.980775 | -1.507148 | -0.207097 | 2.61298 | -0.718075 | -0.616175 | -0.054715 | -0.595465 | -0.415698 | -0.27936 | -0.109796 | 0.91304 | -0.792083 | -0.094428 | -0.130862 | 2.011502 | -0.388463 | 10.494968 | 0.062156 | | 36.78 | -0.054715 | 36.778 | 0.062156 |
| 36.791667 | 0.326746 | -0.807111 | -0.840684 | 2.119653 | -0.517698 | -1.401678 | -0.501084 | 1.508887 | -1.741052 | -0.721113 | 0.008583 | -0.824157 | 1.196895 | -0.289576 | 0.441408 | 1.020049 | -0.809911 | 0.032204 | -0.133056 | 1.737961 | -0.323724 | 13.974339 | -0.083052 | | 36.79 | 0.008583 | 36.792 | -0.083052 |
| 36.805556 | 0.232263 | -1.080768 | -0.958607 | 2.04076 | -0.108216 | -1.127382 | -0.8346 | -0.231522 | -3.790713 | -0.898549 | -0.959071 | -0.898817 | 2.034466 | -0.300239 | 0.949864 | 0.935888 | -0.62329 | 0.313501 | -0.126287 | 1.367861 | -0.253009 | 13.879346 | 1.39207 | | 36.81 | -0.959071 | 36.806 | 1.39207 |
| 36.819444 | -0.223256 | -1.258826 | -0.913471 | 1.724325 | -0.241147 | -0.965354 | -1.338604 | -1.982453 | -3.567794 | -1.186476 | -1.822758 | -0.821711 | 1.026473 | -0.284727 | 1.212197 | 0.829095 | -0.289334 | 0.610278 | -0.200048 | 0.816749 | 0.093051 | 2.730006 | 0.173681 | | 36.82 | -1.822758 | 36.819 | 0.173681 |
| 36.833333 | -0.779102 | -1.2413 | -1.115172 | 1.235164 | -0.743108 | -1.040169 | -1.745301 | -3.505749 | -2.966583 | -1.46994 | -2.254418 | -0.69771 | -1.667737 | -0.271536 | 1.198286 | 0.648643 | -0.127002 | 0.794421 | -0.2574 | 0.324364 | 0.247063 | -2.640653 | -1.176069 | | 36.83 | -2.254418 | 36.833 | -1.176069 |
| 36.847222 | -1.009059 | -0.926378 | -1.491757 | 0.867627 | -1.660152 | -1.753251 | -1.864233 | -4.10492 | -2.379636 | -1.700928 | -2.697909 | -0.604212 | -1.544815 | -0.228083 | 1.146749 | 0.358172 | -0.08101 | 0.707768 | -0.306019 | -0.019075 | 0.37446 | -8.299484 | -1.329464 | | 36.85 | -2.697909 | 36.847 | -1.329464 |
| 36.861111 | -0.923701 | -0.561174 | -2.037805 | 0.758881 | -2.419318 | -2.27055 | -1.913868 | -5.10661 | -1.949242 | -1.682331 | -3.018 | -0.530422 | -0.366845 | -0.178332 | 0.968091 | 0.156982 | -0.173574 | 0.452049 | -0.317291 | -0.230734 | 0.229282 | -6.717532 | -0.265638 | | 36.86 | -3.018 | 36.861 | -0.265638 |
| 36.875 | -0.688071 | -0.134301 | -1.951043 | 0.792676 | -2.45771 | -1.888011 | -1.821275 | -6.030853 | -1.427254 | -2.198334 | -2.806434 | -0.472033 | -0.167946 | -0.122044 | 0.582943 | 0.009239 | -0.189583 | 0.243851 | -0.288674 | -0.34827 | 0.263073 | -5.129284 | -0.227814 | | 36.88 | -2.806434 | 36.875 | -0.227814 |
| 36.888889 | -0.222112 | 0.304202 | -1.475499 | 0.723673 | -2.068871 | -1.338928 | -1.445116 | -6.05328 | -1.239437 | -2.349512 | -2.100968 | -0.359208 | -0.196027 | -0.337296 | 0.262424 | -0.035016 | -0.197018 | 0.145398 | -0.274413 | -0.479659 | 1.023534 | -2.349981 | -0.346113 | | 36.89 | -2.100968 | 36.889 | -0.346113 |
| 36.902778 | 0.378597 | 0.735193 | -0.766783 | 0.652425 | -1.688344 | -0.84799 | -1.01074 | -4.502979 | -0.642693 | -1.740884 | -1.338538 | -0.255162 | -0.192213 | -0.681649 | 0.156421 | 0.042725 | -0.08084 | 0.251016 | -0.272905 | -0.623534 | 1.255689 | 2.381658 | -0.547774 | | 36.9 | -1.338538 | 36.903 | -0.547774 |
| 36.916667 | 0.908105 | 1.260584 | -0.0011 | 0.526792 | -1.181393 | -0.472885 | -0.482681 | -1.443527 | 0.471752 | -1.373254 | -0.292516 | -0.153733 | -0.097206 | -0.582091 | 0.258064 | 0.204948 | 0.03881 | 0.439914 | -0.176676 | -0.411325 | 1.251602 | 5.760928 | -0.493185 | | 36.92 | -0.292516 | 36.917 | -0.493185 |
| 36.930556 | 1.200148 | 1.474464 | 0.384996 | 0.347663 | -0.790039 | -0.278927 | -0.132329 | 1.679901 | 1.650765 | -0.901982 | 0.723715 | -0.128729 | -0.113125 | -0.28551 | 0.274239 | 0.343356 | 0.081274 | 0.583241 | -0.155263 | -0.199726 | 1.415987 | 4.948019 | -0.283436 | | 36.93 | 0.723715 | 36.931 | -0.283436 |
| 36.944444 | 0.479115 | 0.1287 | 0.050896 | 0.12781 | -1.029083 | -0.574484 | -0.230912 | 2.810922 | 1.758696 | -1.089374 | 1.030613 | -0.435506 | 0.228402 | -0.570904 | 0.030166 | -0.064822 | -0.319124 | 0.342118 | -0.665254 | -0.869708 | 0.803859 | 6.704677 | -0.956764 | | 36.94 | 1.030613 | 36.944 | -0.956764 |
| 36.958333 | -1.645333 | -2.894137 | -1.299282 | -0.145387 | -2.274658 | -1.59832 | -0.88359 | -0.271708 | -0.586558 | -2.120515 | 0.072467 | -1.34685 | 0.599138 | -1.795177 | -1.022422 | -1.86633 | -1.410378 | -0.696824 | -1.880637 | -2.885426 | -0.594052 | 7.015288 | 0.039646 | | 36.96 | 0.072467 | 36.958 | 0.039646 |
| 36.972222 | -3.034511 | -3.667526 | -1.851948 | -0.028605 | -3.065424 | -2.18604 | -1.237282 | -4.263038 | -2.562149 | -2.116479 | -1.683056 | -2.27395 | 0.141559 | -2.457523 | -2.588492 | -2.974183 | -2.659402 | -1.956266 | -2.758034 | -4.273174 | -1.360764 | 0.895226 | -2.302024 | | 36.97 | -1.683056 | 36.972 | -2.302024 |
| 36.986111 | -1.765918 | -1.427879 | -0.981338 | 0.662675 | -2.398208 | -1.607715 | -0.917717 | -5.405575 | -1.211851 | -1.012376 | -1.328471 | -1.215931 | -2.614828 | -0.728651 | -2.644604 | -1.059432 | -1.066912 | -1.314924 | -1.158724 | -2.425132 | -1.216947 | -4.940138 | -2.91873 | | 36.99 | -1.328471 | 36.986 | -2.91873 |
| 37 | -0.521785 | -0.025007 | -0.373252 | 1.346946 | -2.181229 | -0.968672 | -0.763172 | -4.21413 | 0.088772 | -0.660202 | -1.057456 | 0.563084 | -3.569375 | 1.000457 | -0.798004 | 1.171217 | 0.996342 | 0.111621 | 1.117061 | 0.377772 | -0.242663 | -7.119302 | -1.633833 | | 37.01 | -1.057456 | 37 | -1.633833 |
| 37.013889 | -0.085748 | 0.059661 | -0.580199 | 1.620495 | -3.012496 | -0.370023 | -0.805114 | -3.099885 | -0.6551 | -0.601664 | -0.93231 | 0.893152 | -2.470706 | 1.240794 | 0.308832 | 1.746137 | 1.426784 | 0.747746 | 1.526671 | 1.401923 | 1.108473 | -9.040982 | -0.072578 | | 37.02 | -0.93231 | 37.014 | -0.072578 |
| 37.027778 | -0.308775 | -0.334416 | -0.89242 | 1.185083 | -3.090968 | -0.10351 | -0.850543 | -2.525655 | -1.363594 | -0.293871 | -1.006725 | 0.452519 | -0.574193 | 0.60802 | 0.836708 | 1.526801 | 0.957646 | 0.848434 | 0.766601 | 1.117476 | 1.129909 | -11.793536 | 0.898995 | | 37.03 | -1.006725 | 37.028 | 0.898995 |
| 37.041667 | -0.729972 | -0.731143 | -0.965693 | 1.012265 | -2.231451 | -0.080349 | -0.811895 | -1.841486 | -1.522272 | -0.325203 | -0.947421 | 0.292137 | 0.413145 | 0.3175 | 1.488804 | 1.691763 | 0.504521 | 0.962128 | 0.230022 | 1.146081 | 1.337547 | -9.935533 | 0.59779 | | 37.05 | -0.947421 | 37.042 | 0.59779 |
| 37.055556 | -0.929185 | -1.098556 | -1.001773 | 0.868913 | -2.20122 | -0.092221 | -0.816479 | -1.295154 | -1.646906 | -0.649343 | -1.057354 | 0.14595 | 0.455822 | 0.238992 | 1.943292 | 1.212665 | 0.475142 | 0.990884 | 0.050751 | 1.353388 | 1.633247 | -7.511596 | 0.948267 | | 37.06 | -1.057354 | 37.056 | 0.948267 |
| 37.069444 | -1.097042 | -1.300388 | -1.041165 | 0.777393 | -2.412044 | -0.155026 | -0.851924 | -0.763398 | -1.493981 | -0.775375 | -1.055869 | 0.032461 | 0.351105 | 0.109452 | 1.66617 | 0.759278 | 0.342645 | 0.759975 | -0.022987 | 1.388107 | 1.682744 | -8.019689 | 0.897631 | | 37.08 | -1.055869 | 37.069 | 0.897631 |
| 37.083333 | -0.857337 | -1.308693 | -1.186734 | 0.690286 | -2.400539 | -0.171451 | -0.711336 | -0.084067 | -1.068655 | -0.730752 | -0.755329 | -0.114751 | 0.091655 | 0.160162 | 1.120692 | 0.370898 | 0.169863 | 0.408962 | -0.076589 | 1.164361 | 1.15717 | -6.713814 | 0.594897 | | 37.09 | -0.755329 | 37.083 | 0.594897 |
| 37.097222 | -0.550311 | -1.048502 | -0.939067 | 0.567581 | -1.94341 | -0.118233 | -0.522565 | 0.57646 | -0.233502 | -0.65379 | -0.235561 | -0.121568 | -0.190144 | 0.283365 | 0.617188 | -0.123464 | 0.126686 | 0.100891 | -0.075371 | 0.900329 | 0.355249 | -12.262341 | 0.313421 | | 37.1 | -0.235561 | 37.097 | 0.313421 |
| 37.111111 | -0.149163 | -0.636239 | -0.530039 | 0.567523 | -1.110551 | -0.03271 | -0.340605 | 0.836415 | 0.8925 | -0.607445 | 0.396669 | 0.003728 | -0.281456 | 0.421806 | 0.237197 | -0.336737 | 0.191662 | -0.047344 | -0.080778 | 0.813502 | -0.1495 | -9.167821 | 0.167586 | | 37.12 | 0.396669 | 37.111 | 0.167586 |
| 37.125 | 0.229334 | -0.335535 | -0.181593 | 0.662678 | -0.362813 | 0.014089 | -0.119559 | 1.074549 | 1.811594 | -0.568186 | 1.047918 | 0.149614 | -0.189804 | 0.49792 | 0.071671 | -0.361407 | 0.229818 | -0.033211 | -0.076884 | 0.552509 | -0.584594 | 0.030571 | 0.019446 | | 37.13 | 1.047918 | 37.125 | 0.019446 |
| 37.138889 | 0.419618 | -0.358301 | -0.011983 | 0.745838 | 0.042259 | -0.001019 | -0.063389 | 1.656241 | 2.495964 | -0.549402 | 1.449587 | 0.276327 | -0.113501 | 0.471756 | 0.027847 | -0.39251 | 0.213654 | 0.070152 | -0.029722 | 0.292245 | -1.027144 | -0.005487 | -0.142266 | | 37.15 | 1.449587 | 37.139 | -0.142266 |
| 37.152778 | 0.634015 | -0.361837 | 0.163459 | 0.75132 | 0.352264 | 0.007129 | -0.088296 | 2.083616 | 3.120425 | -0.563393 | 1.997054 | 0.372589 | -0.100747 | 0.424534 | 0.072195 | -0.127715 | 0.174797 | 0.228238 | 0.10898 | 0.25126 | -1.092167 | -0.080902 | -0.144028 | | 37.16 | 1.997054 | 37.153 | -0.144028 |
| 37.166667 | 0.951907 | -0.141229 | 0.33905 | 0.734538 | 0.65619 | 0.058184 | -0.148814 | 2.455087 | 3.383016 | -0.580301 | 2.637995 | 0.442066 | -0.13238 | 0.455059 | 0.174228 | 0.22438 | 0.142461 | 0.229368 | 0.246101 | 0.152923 | -0.839508 | -0.077521 | -0.063903 | | 37.17 | 2.637995 | 37.167 | -0.063903 |
| 37.180556 | 1.081324 | 0.005097 | 0.433152 | 0.631778 | 0.83237 | 0.104442 | -0.104583 | 2.320398 | 3.871397 | -0.597545 | 3.482049 | 0.497364 | -0.01444 | 0.475636 | 0.27184 | 0.588999 | 0.116481 | 0.206037 | 0.319256 | -0.197885 | -0.561812 | 0.005562 | 0.016478 | | 37.19 | 3.482049 | 37.181 | 0.016478 |
| 37.194444 | 1.015187 | 0.083049 | 0.457663 | 0.462147 | 0.886299 | 0.110091 | 0.055228 | 1.660957 | 3.661258 | -0.597219 | 4.035851 | 0.473927 | 0.05188 | 0.484469 | 0.300041 | 0.888228 | 0.111179 | 0.251855 | 0.336941 | -0.522974 | -0.310963 | 0.050137 | 0.034292 | | 37.2 | 4.035851 | 37.194 | 0.034292 |
| 37.208333 | 0.92194 | -0.012783 | 0.387699 | 0.252547 | 0.876806 | 0.068367 | 0.080348 | 2.022772 | 2.419148 | -0.602362 | 4.376232 | 0.318867 | 0.05452 | 0.352954 | 0.052026 | 0.871115 | 0.007524 | 0.271351 | 0.216694 | -0.812307 | -0.093362 | 0.016118 | 0.023963 | | 37.22 | 4.376232 | 37.208 | 0.023963 |
| 37.222222 | 0.612448 | -0.347133 | 0.19871 | 0.156659 | 0.715955 | 0.005236 | -0.053635 | 2.361935 | 1.697127 | -0.633369 | 4.382592 | 0.065966 | -0.071798 | 0.190914 | -0.389757 | 0.607207 | -0.152108 | 0.261131 | 0.023378 | -1.044814 | -0.039385 | -0.026324 | -0.061761 | | 37.23 | 4.382592 | 37.222 | -0.061761 |
| 37.236111 | 0.201237 | -0.694033 | 0.052585 | 0.374615 | 0.609981 | -0.020389 | -0.205615 | 3.260484 | 1.396098 | -0.634882 | 3.756967 | -0.023233 | -0.169957 | 0.108575 | -0.795494 | 0.260151 | -0.24396 | 0.375694 | -0.140402 | -1.143551 | -0.052598 | -0.021673 | -0.134237 | | 37.24 | 3.756967 | 37.236 | -0.134237 |
| 37.25 | -0.061383 | -0.699869 | 0.083971 | 0.728014 | 0.529111 | -0.007805 | -0.257825 | 3.51037 | 0.525095 | -0.493088 | 2.904205 | 0.153621 | -0.057059 | 0.156114 | -0.930859 | 0.197703 | -0.169525 | 0.466802 | -0.075487 | -0.872111 | 0.05668 | 0.03267 | -0.05879 | | 37.26 | 2.904205 | 37.25 | -0.05879 |
| 37.263889 | 0.036464 | -0.385352 | 0.205479 | 0.992472 | 0.473752 | -0.0099 | -0.228138 | 3.180529 | 0.044398 | -0.373695 | 1.701351 | 0.42142 | 0.170937 | 0.214515 | -0.762708 | 0.425625 | -0.113526 | 0.292715 | 0.116631 | -0.378869 | 0.307886 | 0.106186 | 0.082245 | | 37.27 | 1.701351 | 37.264 | 0.082245 |
| 37.277778 | 0.334403 | -0.091015 | 0.290467 | 1.149957 | 0.4173 | -0.080341 | -0.223522 | 2.408275 | -0.130728 | -0.344717 | 0.969344 | 0.569835 | 0.291887 | 0.161974 | -0.495372 | 0.824743 | -0.228866 | 0.004442 | 0.227946 | 0.027027 | 0.455648 | 0.072524 | 0.10684 | | 37.28 | 0.969344 | 37.278 | 0.10684 |
| 37.291667 | 0.685324 | 0.164124 | 0.32541 | 1.279052 | 0.339454 | -0.163465 | -0.193163 | 1.631678 | 0.32468 | -0.33428 | 0.524257 | 0.542221 | 0.234826 | 0.147213 | -0.2439 | 1.140287 | -0.373898 | -0.254948 | 0.205158 | 0.307549 | 0.284335 | -0.066784 | 0.038495 | | 37.3 | 0.524257 | 37.292 | 0.038495 |
| 37.305556 | 0.898161 | 0.296512 | 0.364049 | 1.396042 | 0.235043 | -0.160873 | -0.055404 | 1.441839 | 1.070308 | -0.26763 | 0.723553 | 0.415201 | 0.089778 | 0.16997 | -0.157583 | 1.056182 | -0.391302 | -0.274905 | 0.124135 | 0.462188 | -0.0792 | 9.183975 | 0.029244 | | 37.31 | 0.723553 | 37.306 | 0.029244 |
| 37.319444 | 0.889115 | 0.353157 | 0.405112 | 1.425904 | 0.00436 | -0.058869 | 0.073499 | 1.628523 | 2.114842 | -0.172537 | 1.088406 | 0.338275 | 0.040146 | 0.178101 | -0.0422 | 0.835824 | -0.29627 | -0.088153 | 0.056167 | 0.424339 | -0.346833 | 13.312794 | 0.031937 | | 37.33 | 1.088406 | 37.319 | 0.031937 |
| 37.333333 | 0.821369 | 0.433642 | 0.419634 | 1.332153 | -0.430344 | -0.001411 | 0.208329 | 1.715263 | 2.597868 | -0.134044 | 1.205108 | 0.372224 | 0.063301 | 0.165822 | 0.016913 | 0.634521 | -0.181601 | 0.115686 | 0.01326 | 0.282831 | -0.337953 | 10.902455 | -0.004067 | | 37.34 | 1.205108 | 37.333 | -0.004067 |
| 37.347222 | 0.778712 | 0.540652 | 0.470751 | 1.26579 | -0.784223 | -0.023668 | 0.389934 | 1.591171 | 2.562133 | -0.134362 | 1.053386 | 0.444395 | 0.127626 | 0.173263 | 0.003545 | 0.446356 | -0.129157 | 0.158064 | 0.019963 | 0.229808 | -0.156071 | 9.528097 | -0.092764 | | 37.35 | 1.053386 | 37.347 | -0.092764 |
| 37.361111 | 0.714285 | 0.505992 | 0.572182 | 1.369666 | -0.944722 | -0.039799 | 0.616166 | 1.014378 | 1.731359 | -0.191654 | 0.7063 | 0.455452 | 0.120485 | 0.189929 | -0.085457 | 0.344215 | -0.148813 | 0.011019 | 0.032997 | 0.405048 | 0.301267 | 8.796513 | -0.156633 | | 37.37 | 0.7063 | 37.361 | -0.156633 |
| 37.375 | 0.510917 | 0.300218 | 0.641342 | 1.514348 | -1.036495 | -0.024723 | 0.664349 | -0.04689 | 0.470746 | -0.207987 | 0.229237 | 0.345766 | 0.089102 | 0.217032 | -0.246238 | 0.359474 | -0.168833 | -0.179356 | 0.04782 | 0.619845 | 0.690342 | 7.467506 | -0.086484 | | 37.38 | 0.229237 | 37.375 | -0.086484 |
| 37.388889 | 0.302216 | 0.151649 | 0.587645 | 1.492041 | -1.231827 | 0.002991 | 0.469738 | -1.147754 | -0.847646 | -0.257728 | -0.321285 | 0.272859 | 0.11254 | 0.285332 | -0.305489 | 0.464532 | -0.204326 | -0.32571 | 0.090923 | 0.693422 | 0.920355 | 6.136484 | 0.035543 | | 37.4 | -0.321285 | 37.389 | 0.035543 |
| 37.402778 | 0.19516 | 0.187685 | 0.481349 | 1.242333 | -1.375028 | -0.031144 | 0.188783 | -2.000437 | -1.761744 | -0.368241 | -0.759829 | 0.257206 | 0.064488 | 0.299208 | -0.122093 | 0.516323 | -0.217199 | -0.327172 | 0.089701 | 0.660968 | 0.715112 | 4.124966 | 0.063255 | | 37.41 | -0.759829 | 37.403 | 0.063255 |
| 37.416667 | 0.095154 | 0.235869 | 0.381363 | 1.065893 | -1.333208 | -0.077515 | 0.13383 | -2.564583 | -2.337753 | -0.446798 | -0.976373 | 0.218299 | -0.065368 | 0.181526 | 0.104948 | 0.379463 | -0.255962 | -0.307853 | 0.042363 | 0.640934 | 0.196332 | 2.548482 | -0.012799 | | 37.42 | -0.976373 | 37.417 | -0.012799 |
| 37.430556 | 0.093322 | 0.30167 | 0.406722 | 1.02036 | -1.205726 | -0.021713 | 0.363733 | -3.094705 | -2.83068 | -0.43395 | -1.282358 | 0.271388 | -0.090384 | 0.143778 | 0.256185 | 0.38314 | -0.221309 | -0.230122 | 0.081904 | 0.798269 | -0.272072 | 1.821513 | 0.069717 | | 37.44 | -1.282358 | 37.431 | 0.069717 |
| 37.444444 | 0.672646 | 1.058479 | 0.868157 | 1.133489 | -0.537754 | 0.365526 | 1.210723 | -3.241386 | -2.995664 | -0.280529 | -1.720435 | 0.786519 | 0.311043 | 0.414291 | 0.582336 | 0.773736 | 0.183837 | 0.011436 | 0.523455 | 1.661849 | -0.282284 | 1.844711 | 0.789797 | | 37.45 | -1.720435 | 37.444 | 0.789797 |
| 37.458333 | 2.338724 | 3.021942 | 1.802187 | 1.556376 | 0.946011 | 1.050822 | 2.968888 | -1.826261 | -1.416299 | 0.487306 | -1.213815 | 1.989074 | 1.290845 | 1.04887 | 1.195679 | 1.704359 | 0.856293 | 0.433868 | 1.279337 | 2.980338 | 0.775783 | 2.867071 | 2.074961 | | 37.47 | -1.213815 | 37.458 | 2.074961 |
| 37.472222 | 3.780563 | 4.525818 | 2.411731 | 1.835649 | 2.893224 | 1.276374 | 3.992285 | 1.032613 | 1.501813 | 0.825062 | 0.700425 | 2.544826 | 1.563913 | 1.464106 | 1.482895 | 1.991068 | 1.047335 | 0.779323 | 1.605559 | 3.504247 | 2.954658 | 4.988119 | 2.576425 | | 37.48 | 0.700425 | 37.472 | 2.576425 |
| 37.486111 | 3.504687 | 4.174815 | 2.273274 | 1.506229 | 3.0094 | 0.797492 | 3.387257 | 2.881119 | 4.238117 | 1.096913 | 3.20367 | 1.904751 | 1.240884 | 1.155404 | 1.495431 | 1.566472 | 0.845611 | 0.938837 | 1.151303 | 2.149202 | 4.016303 | 4.587914 | 2.129212 | | 37.49 | 3.20367 | 37.486 | 2.129212 |
| 37.5 | 2.964335 | 3.294144 | 2.102916 | 0.878657 | 4.322257 | 0.393118 | 2.706727 | 2.984367 | 5.545267 | 1.256066 | 4.165049 | 0.911428 | 1.480845 | 0.620948 | 1.419848 | 0.852845 | 0.604175 | 0.837773 | 0.527156 | 1.994918 | 3.20939 | 8.060835 | 1.129525 | | 37.5 | 4.165049 | 37.5 | 1.129525 |
| 37.513889 | 2.945872 | 2.847873 | 1.92132 | 0.560323 | 5.883757 | 0.311361 | 2.341501 | 2.376726 | 3.534031 | 1.500709 | 4.325684 | 0.369585 | 2.85869 | 0.469822 | 0.992126 | 0.657744 | 0.425619 | 0.690941 | 0.318781 | 2.780201 | 1.713615 | 6.625849 | 0.838155 | | 37.51 | 4.325684 | 37.514 | 0.838155 |
| 37.527778 | 2.870122 | 2.71977 | 1.573406 | 0.784321 | 7.205378 | 0.514634 | 2.289656 | 1.222093 | 1.633988 | 1.524155 | 3.486437 | 0.04305 | 2.697882 | 0.624488 | 0.48956 | 0.813835 | 0.294711 | 0.235472 | 0.249128 | 2.587513 | 0.929148 | 5.965125 | 1.668377 | | 37.53 | 3.486437 | 37.528 | 1.668377 |
| 37.541667 | 2.610145 | 2.636372 | 1.243285 | 1.147251 | 6.500893 | 1.197416 | 2.240893 | 0.291105 | 0.974833 | 1.323287 | 2.934965 | -0.057454 | 0.82018 | 0.781905 | 0.037803 | 0.712025 | 0.256138 | -0.420046 | 0.201647 | 1.53297 | -0.528533 | 3.057099 | 0.861743 | | 37.54 | 2.934965 | 37.542 | 0.861743 |
| 37.555556 | 2.206359 | 2.282795 | 0.958595 | 1.514339 | 5.056666 | 1.786618 | 2.101442 | -0.936965 | -0.754547 | 1.029418 | 2.791991 | -0.044843 | -1.620329 | 0.816488 | -0.125109 | 0.524716 | 0.233121 | -0.697922 | 0.252367 | 0.85312 | -1.428125 | 0.791587 | -0.309839 | | 37.56 | 2.791991 | 37.556 | -0.309839 |
| 37.569444 | 1.709867 | 1.883058 | 0.809282 | 1.527239 | 4.200567 | 2.151898 | 1.92055 | -1.730577 | -2.609588 | 0.786894 | 1.970058 | 0.145293 | -3.541157 | 0.573041 | -0.022608 | 0.104332 | 0.296626 | -0.722982 | 0.334037 | 0.53599 | -2.183176 | -0.941191 | -1.282524 | | 37.57 | 1.970058 | 37.569 | -1.282524 |
| 37.583333 | 1.244374 | 1.580451 | 0.762643 | 1.355737 | 3.261911 | 2.215502 | 1.74644 | -1.904065 | -4.036934 | 0.516507 | 0.809468 | 0.375818 | -3.372252 | 0.450712 | 0.209843 | 0.059302 | 0.478875 | -0.519743 | 0.33669 | 0.441517 | -3.126858 | -4.337453 | -0.565388 | | 37.58 | 0.809468 | 37.583 | -0.565388 |
| 37.597222 | 1.232268 | 1.360861 | 0.632806 | 1.143185 | 2.751481 | 1.82188 | 1.569966 | -2.164068 | -4.174998 | 0.292954 | -0.617613 | 0.414184 | -2.002078 | 0.688688 | 0.31065 | 0.380683 | 0.634429 | 0.007486 | 0.412711 | 0.365019 | -2.526368 | -6.788988 | 0.087781 | | 37.6 | -0.617613 | 37.597 | 0.087781 |
| 37.611111 | 0.900791 | 1.289209 | 0.422282 | 0.903871 | 2.457528 | 1.186523 | 1.314676 | -2.346831 | -3.905741 | 0.109825 | -1.605488 | 0.290288 | -0.581813 | 0.894434 | 0.332543 | 0.625136 | 0.728197 | 0.280887 | 0.431384 | -0.055468 | -3.576253 | -13.786211 | -0.215498 | | 37.61 | -1.605488 | 37.611 | -0.215498 |
| 37.625 | 0.776839 | 1.319603 | 0.416633 | 0.613946 | 1.825971 | 0.176002 | 1.0766 | -2.735106 | -3.274641 | 0.022308 | -2.392714 | 0.062471 | -0.486663 | 0.91662 | 0.253994 | 0.681222 | 0.656405 | 0.059171 | 0.347204 | -0.819999 | -4.307523 | -13.361356 | -0.719946 | | 37.63 | -2.392714 | 37.625 | -0.719946 |
| 37.638889 | 0.528116 | 1.32139 | 0.498697 | 0.306398 | 0.609463 | -0.856019 | 0.928708 | -3.322777 | -2.911195 | 0.067105 | -2.550647 | -0.314807 | -0.758175 | 0.642385 | -0.045652 | 0.61447 | 0.396241 | -0.474366 | 0.22599 | -1.702702 | -3.606661 | -8.634287 | -0.762007 | | 37.64 | -2.550647 | 37.639 | -0.762007 |
| 37.652778 | 0.148077 | 1.106166 | 0.538392 | -0.05548 | -0.937436 | -1.736628 | 0.855523 | -4.426211 | -2.868554 | 0.077734 | -2.871432 | -0.788056 | -1.069097 | 0.317535 | -0.391904 | 0.402914 | 0.049139 | -0.950406 | 0.150064 | -2.481572 | -0.498036 | -1.051997 | -0.762819 | | 37.65 | -2.871432 | 37.653 | -0.762819 |
| 37.666667 | 0.036477 | 0.869814 | 0.39766 | -0.554401 | -2.243018 | -2.131447 | 0.62459 | -5.57298 | -2.295348 | 0.061113 | -3.412781 | -0.907509 | -1.109151 | 0.097405 | -0.482821 | 0.094328 | -0.09925 | -1.142062 | 0.188231 | -2.823513 | 1.574526 | 3.477885 | -0.49095 | | 37.67 | -3.412781 | 37.667 | -0.49095 |
| 37.680556 | 0.128688 | 0.721925 | -0.099292 | -0.742702 | -3.264916 | -0.981581 | 0.089761 | -6.249658 | -1.919181 | 0.020074 | -3.671118 | -0.743098 | 0.521763 | 0.075379 | -0.236483 | -0.054338 | -0.040836 | -0.780555 | 0.283968 | -2.858777 | 1.974393 | 14.502017 | 0.075237 | | 37.68 | -3.671118 | 37.681 | 0.075237 |
| 37.694444 | 0.144963 | 0.688501 | -0.874283 | -0.612714 | -3.68504 | -0.958464 | -0.681858 | -5.885144 | -0.957447 | -0.22095 | -2.941724 | -0.562264 | 1.870322 | 0.12736 | 0.131709 | -0.102948 | 0.033015 | -0.273424 | 0.359937 | -2.657153 | 0.31983 | 13.304895 | 1.522092 | | 37.69 | -2.941724 | 37.694 | 1.522092 |
| 37.708333 | 0.242213 | 0.613942 | -1.131451 | -0.466023 | -4.566375 | -0.874979 | -1.12211 | -4.446719 | -0.441208 | -0.642881 | -2.444572 | -0.457252 | 3.615348 | 0.167354 | 0.326607 | -0.301613 | 0.045276 | 0.076284 | 0.343581 | -2.269324 | 0.682122 | 4.379894 | 2.233942 | | 37.71 | -2.444572 | 37.708 | 2.233942 |
| 37.722222 | 0.085588 | 0.44483 | -0.965265 | -0.523267 | -5.567403 | -0.269762 | -0.955711 | -2.903585 | 0.138274 | -0.904255 | -1.95362 | -0.516954 | 1.575453 | 0.142308 | 0.126941 | -0.439722 | -0.037342 | -0.002911 | 0.240331 | -1.823591 | 0.937434 | 1.205736 | 0.555988 | | 37.72 | -1.95362 | 37.722 | 0.555988 |
| 37.736111 | -0.344081 | 0.263988 | -0.812964 | -0.502032 | -5.556013 | -0.175965 | -0.910733 | -1.620677 | 0.112223 | -1.309043 | -1.544275 | -0.808158 | -2.046603 | 0.059375 | -0.237364 | -0.61819 | -0.161383 | -0.369657 | 0.067934 | -1.835431 | 1.217605 | -1.33263 | -1.909321 | | 37.74 | -1.544275 | 37.736 | -1.909321 |
| 37.75 | -0.644584 | 0.103265 | -0.806948 | 0.043716 | -5.101997 | -0.087912 | -0.691262 | -1.095127 | -0.528789 | -1.617409 | -1.206554 | -1.08886 | -3.187943 | -0.067358 | -0.820721 | -0.942921 | -0.326943 | -0.66335 | -0.117479 | -1.901616 | 0.538362 | -8.090059 | -3.071335 | | 37.75 | -1.206554 | 37.75 | -3.071335 |
| 37.763889 | -0.565223 | -0.146649 | -0.857423 | 1.106788 | -4.610043 | -0.032248 | -0.066349 | -1.211811 | -1.694903 | -1.893047 | -1.002723 | -1.336522 | -1.980828 | -0.178921 | -1.2971 | -0.907028 | -0.460669 | -0.660902 | -0.191959 | -1.711869 | -1.688983 | -6.055394 | -1.985486 | | 37.76 | -1.002723 | 37.764 | -1.985486 |
| 37.777778 | -0.71522 | -0.465254 | -0.800788 | 1.786566 | -3.681761 | -0.019247 | 0.326074 | -1.846707 | -3.100012 | -2.197059 | -1.22819 | -1.288173 | -0.486398 | -0.198923 | -1.358348 | -0.581317 | -0.540983 | -0.372181 | -0.170111 | -1.395924 | -1.364572 | -6.665941 | -1.293438 | | 37.78 | -1.22819 | 37.778 | -1.293438 |
| 37.791667 | -0.839321 | -0.793204 | -0.820791 | 2.015351 | -2.701349 | -0.049759 | 0.284993 | -2.59014 | -4.114302 | -2.082212 | -1.562709 | -1.120889 | -0.38658 | -0.145948 | -1.177406 | -0.187711 | -0.606807 | -0.027807 | -0.135617 | -1.068819 | 0.146246 | -1.025806 | -0.630928 | | 37.79 | -1.562709 | 37.792 | -0.630928 |
| 37.805556 | -1.080297 | -1.088144 | -0.933001 | 2.07521 | -2.222553 | -0.122569 | -0.082935 | -3.58725 | -5.018392 | -2.145574 | -1.865547 | -0.88712 | -0.100918 | -0.158162 | -0.686816 | -0.103437 | -0.604389 | 0.154697 | -0.114551 | -0.676287 | 1.725845 | 3.087049 | -0.130208 | | 37.81 | -1.865547 | 37.806 | -0.130208 |
| 37.819444 | -1.655958 | -1.455619 | -0.951624 | 1.997437 | -1.894973 | -0.198085 | -0.607412 | -4.472581 | -5.257247 | -1.914925 | -2.263218 | -0.791932 | -0.049323 | -0.244428 | -0.066895 | 0.007477 | -0.59249 | 0.122066 | -0.201792 | -0.455015 | 2.73619 | 6.458185 | 0.402105 | | 37.82 | -2.263218 | 37.819 | 0.402105 |
| 37.833333 | -1.861337 | -1.653183 | -0.809775 | 1.915954 | -1.671364 | -0.225251 | -1.040379 | -5.378015 | -5.017504 | -2.233347 | -2.486178 | -0.749056 | 0.391933 | -0.339877 | 0.503953 | 0.108627 | -0.440917 | 0.148579 | -0.333723 | -0.241879 | 2.548562 | 4.6848 | 0.480188 | | 37.83 | -2.486178 | 37.833 | 0.480188 |
| 37.847222 | -1.269313 | -1.740027 | -0.568849 | 1.932886 | -1.433693 | -0.239337 | -1.368148 | -6.14924 | -5.079197 | -2.443648 | -2.705805 | -0.930934 | 1.148662 | -0.457944 | 0.844428 | 0.052706 | -0.214643 | 0.27674 | -0.458973 | -0.000438 | 4.565692 | 4.033706 | 1.038057 | | 37.85 | -2.705805 | 37.847 | 1.038057 |
| 37.861111 | -1.082154 | -1.627886 | -0.337723 | 1.96947 | -1.208826 | -0.30184 | -1.570636 | -5.584512 | -4.537404 | -2.301445 | -2.518718 | -1.056477 | 2.388162 | -0.439703 | 0.967483 | -0.012142 | -0.04699 | 0.360786 | -0.484697 | 0.269251 | 6.735769 | 4.536237 | 1.565654 | | 37.86 | -2.518718 | 37.861 | 1.565654 |
| 37.875 | | | | | | | | | | | | -1.037941 | 0.727918 | -0.429261 | 1.068268 | -0.082415 | 0.101858 | 0.377054 | -0.382325 | 0.510372 | 2.989907 | 1.784239 | 1.004138 | | | | 37.875 | 1.004138 |
| 37.888889 | | | | | | | | | | | | -0.893965 | -0.46633 | -0.366748 | 1.279629 | -0.068309 | 0.179957 | 0.41189 | -0.296922 | 0.719509 | -0.579243 | 2.474829 | -0.015876 | | | | 37.889 | -0.015876 |
| 37.902778 | | | | | | | | | | | | -0.822574 | -2.375736 | -0.312778 | 1.52376 | -0.008634 | 0.259216 | 0.503748 | -0.310071 | 0.888746 | -4.113241 | 0.852719 | -0.980529 | | | | 37.903 | -0.980529 |

**Supplementary Table S7 (c)** Mean day-to-day motion rate in Control and Stress tanks of lettuce measured under dimGday/night and RGBday dimGnight, calculated using either 24 h motion integration or 22 h motion integration excluding day/night transitions.

| Row | RGBStress | RGBControl | RGB22Stress | RGB22Control | dimGControl | dimGStress | dimG22Control | dimG22Stress |
|---|---|---|---|---|---|---|---|---|
| 24.5 | 0.874032479 | 1.276410009 | 0.635233733 | 1.075548645 | 0.86564808 | 0.867007372 | 0.788130894 | 0.823122542 |
| 25.5 | 0.853581252 | 1.267851317 | 0.726873026 | 1.114506035 | 0.817140347 | 0.925751985 | 0.769657607 | 0.891628281 |
| 26.5 | 0.591694295 | 1.761032322 | 0.475238034 | 1.194461185 | 1.342212902 | 0.659423408 | 1.288793183 | 0.62380653 |
| 27.5 | 0.942779315 | 2.457090912 | 0.608523781 | 1.198013725 | 1.169052987 | 0.570076847 | 1.096720871 | 0.540705254 |
| 28.5 | 0.618618419 | 2.845517689 | 0.382259411 | 1.460562328 | 1.423451044 | 0.608115683 | 1.366512254 | 0.560851659 |
| 29.5 | 0.695363295 | 2.873060971 | 0.439558561 | 1.474349509 | 1.43440757 | 0.495387634 | 1.383916948 | 0.472083758 |
| 30.5 | 0.968247624 | 2.49811591 | 0.417309332 | 1.235126115 | 1.071393841 | 0.320163552 | 0.976523091 | 0.307813003 |
| 31.5 | 1.135477107 | 2.916188433 | 0.491065471 | 1.385050459 | 1.225282643 | 0.451852758 | 1.140005504 | 0.428288972 |
| 32.5 | 1.639371177 | 3.27972569 | 0.652998135 | 1.630475414 | 1.420985498 | 0.48831974 | 1.377104828 | 0.457665509 |
| 33.5 | 1.966711989 | 3.469672021 | 0.703396748 | 1.523690611 | 1.381393306 | 0.715035736 | 1.297561028 | 0.687179123 |
| 34.5 | 1.951614498 | 3.085146396 | 0.656587509 | 1.445272953 | 1.420401457 | 0.559090285 | 1.275009626 | 0.542799151 |
| 35.5 | 1.868607009 | 3.839794339 | 0.694692508 | 1.952838455 | 1.668011787 | 0.714588343 | 1.598037566 | 0.681872534 |
| 36.5 | 1.613949131 | 3.072520525 | 0.656923836 | 1.788789954 | 1.777969118 | 0.695456373 | 1.691299299 | 0.666465662 |

**Supplementary Table S7 (d)** Mean day-to-day motion rate of individual lettuce plants under dimGday/night and RGBday dimGnight. Data represent means of 9–10 plants

| Interval | RGBControlMean | RGBControlSEM | dimGControlMea | dimGControlSEM | RGBStressMean | RGBStressSEM | dimGStressMean | dimGStressSEM |
|---|---|---|---|---|---|---|---|---|
| 24.5 | 0.654917872 | 0.064486232 | 0.590628281 | 0.063216553 | 0.528158681 | 0.042107669 | 0.483526073 | 0.034450747 |
| 25.5 | 0.647833272 | 0.062138374 | 0.568713779 | 0.069723857 | 0.568091893 | 0.045200865 | 0.499048916 | 0.036431339 |
| 26.5 | 0.795115828 | 0.05289016 | 0.729958421 | 0.062948219 | 0.577157154 | 0.05666755 | 0.511623683 | 0.055551974 |
| 27.5 | 0.91981825 | 0.093174238 | 0.784749274 | 0.098499289 | 0.554235414 | 0.058667439 | 0.468655613 | 0.056578997 |
| 28.5 | 0.975598339 | 0.104355277 | 0.816860093 | 0.112349661 | 0.545124178 | 0.058918994 | 0.45699458 | 0.063816985 |
| 29.5 | 1.140782556 | 0.101111099 | 0.973025519 | 0.108014447 | 0.578273329 | 0.05512029 | 0.473513022 | 0.060404902 |
| 30.5 | 1.028587644 | 0.105480572 | 0.877985603 | 0.100255753 | 0.542320987 | 0.050410114 | 0.426750935 | 0.051921115 |
| 31.5 | 1.19381405 | 0.117681441 | 1.007036462 | 0.102428314 | 0.538688684 | 0.051013042 | 0.438295099 | 0.064845666 |
| 32.5 | 1.349668039 | 0.219131836 | 1.114574216 | 0.163954339 | 0.61220115 | 0.063391554 | 0.465546379 | 0.072065404 |
| 33.5 | 1.285325872 | 0.198299665 | 1.148667758 | 0.17454792 | 0.704950848 | 0.070719612 | 0.503882067 | 0.066101943 |
| 34.5 | 1.258751117 | 0.134882971 | 1.079664351 | 0.137532691 | 0.65272048 | 0.091486593 | 0.475873795 | 0.077254853 |
| 35.5 | 1.49981685 | 0.190376295 | 1.296906678 | 0.154061127 | 0.70044182 | 0.10422227 | 0.572236876 | 0.075709423 |

**Supplementary Table S7 (e)** Direct motion in 100 mM NaCl stressed Amaranth and their respective controls under RGBday dimGnight lighting condition

| Time | RGBControl1 | RGBControl2 | RGBControl3 | RGBControl4 | RGBControl5 | RGBControl6 | RGBControl7 | RGBControl8 | RGBControl9 | RGBControl10 | RGBControl11 | RGBControl12 | RGBControltank | RGBStress1 | RGBStress2 | RGBStress3 | RGBStress4 | RGBStress5 | RGBStress6 | RGBStress7 | RGBStress8 | RGBStress9 | RGBStress10 | RGBStresstank | | Time | Mean ControlRGB | Mean StressRGB |
|---|---|---|---|---|---|---|---|---|---|---|---|---|---|---|---|---|---|---|---|---|---|---|---|---|---|---|---|---|
| 18.1041667 | 0.070784 | -0.009059 | 0.050035 | 0.076298 | 0.048139 | 0.054986 | 0.039462 | 0.052778 | 0.069895 | -0.615068 | -0.024734 | -0.323009 | 0.032835 | 0.56602 | 0.43218 | 0.554137 | 0.459779 | 0.292788 | 0.558321 | 0.619378 | 0.311394 | 0.563685 | 0.463269 | 0.538536 | | 18.097 | 0.032835 | 0.541189 |
| 18.1180556 | -0.001883 | -0.049514 | 0.014201 | 0.017248 | -0.014395 | -0.002345 | -0.040228 | -0.0026 | -0.005879 | -0.062464 | -0.107654 | -0.206411 | -0.060605 | 0.470302 | 0.420299 | 0.485745 | 0.387468 | 0.385459 | 0.45114 | 0.539163 | 0.345571 | 0.4847 | 0.428447 | 0.507787 | | 18.111 | -0.060605 | 0.510046 |
| 18.1319444 | 0.040049 | -0.031024 | 0.014217 | -0.028214 | -0.067434 | -0.014656 | -0.095207 | -0.019354 | -0.035399 | -0.058958 | -0.100264 | -0.139774 | -0.123781 | 0.373731 | 0.347042 | 0.39414 | 0.387466 | 0.403462 | 0.433924 | 0.460176 | 0.321011 | 0.426755 | 0.328087 | 0.40596 | | 18.125 | -0.123781 | 0.431817 |
| 18.1458333 | 0.053023 | -0.006999 | -0.024167 | -0.112494 | -0.083386 | -0.027695 | -0.085528 | -0.031966 | -0.04461 | -0.049094 | -0.106538 | -0.098242 | -0.038694 | 0.250487 | 0.229155 | 0.272731 | 0.278651 | 0.356023 | 0.373066 | 0.344525 | 0.252339 | 0.310972 | 0.186846 | 0.2822 | | 18.139 | -0.038694 | 0.327325 |
| 18.1597222 | 0.033157 | -0.02122 | -0.066381 | -0.18856 | -0.081257 | -0.033315 | -0.077626 | -0.042574 | -0.055263 | -0.020521 | -0.169568 | -0.082284 | 0.045093 | 0.091804 | 0.086054 | 0.127579 | 0.134774 | 0.227485 | 0.234028 | 0.189304 | 0.12572 | 0.116905 | 0.004637 | 0.156166 | | 18.153 | 0.045093 | 0.201772 |
| 18.1736111 | 0.007283 | -0.052296 | -0.078945 | -0.241611 | -0.096234 | -0.0338 | -0.109591 | -0.045779 | -0.068798 | -0.02055 | -0.212871 | -0.06909 | -0.017362 | 0.045251 | 0.05378 | 0.096118 | 0.075056 | 0.128602 | 0.194568 | 0.150779 | 0.0717 | 0.038095 | -0.087139 | 0.110761 | | 18.167 | -0.017362 | 0.14295 |
| 18.1875 | -0.055915 | -0.078145 | -0.071135 | -0.280487 | -0.125507 | -0.044306 | -0.159138 | -0.053004 | -0.099332 | -0.059723 | -0.200823 | -0.070253 | -0.145697 | 0.128568 | 0.130164 | 0.162411 | 0.130163 | 0.127351 | 0.247246 | 0.225094 | 0.13537 | 0.096128 | -0.068889 | 0.130567 | | 18.181 | -0.145697 | 0.162676 |
| 18.2013889 | -0.108674 | -0.097976 | -0.058221 | -0.298913 | -0.139465 | -0.066802 | -0.187065 | -0.089767 | -0.13831 | -0.105349 | -0.19365 | -0.100517 | -0.18884 | 0.205671 | 0.204527 | 0.210974 | 0.197725 | 0.157288 | 0.272303 | 0.285011 | 0.216058 | 0.161521 | -0.041796 | 0.170983 | | 18.194 | -0.18884 | 0.190261 |
| 18.2152778 | -0.10987 | -0.089444 | -0.057665 | -0.25861 | -0.11614 | -0.080057 | -0.162337 | -0.119786 | -0.143167 | -0.104864 | -0.169969 | -0.115739 | -0.14042 | 0.206696 | 0.226076 | 0.213218 | 0.212104 | 0.162925 | 0.253372 | 0.27984 | 0.237064 | 0.170557 | -0.051235 | 0.195302 | | 18.208 | -0.14042 | 0.206799 |
| 18.2291667 | -0.094889 | -0.061929 | -0.040071 | -0.178839 | -0.088496 | -0.077355 | -0.138324 | -0.110559 | -0.12762 | -0.07736 | -0.133268 | -0.089165 | -0.132523 | 0.061879 | 0.089138 | 0.059563 | 0.07558 | 0.018129 | 0.049548 | 0.084518 | 0.083156 | 0.002878 | -0.180971 | 0.016569 | | 18.222 | -0.132523 | 0.036406 |
| 18.2430556 | -0.072242 | -0.051496 | -0.050877 | -0.121818 | -0.087865 | -0.089511 | -0.14462 | -0.095616 | -0.124013 | -0.07271 | -0.103077 | -0.072303 | -0.17057 | -0.158337 | -0.134289 | -0.171455 | -0.149668 | -0.21592 | -0.24759 | -0.196015 | -0.167269 | -0.248948 | -0.419866 | -0.259055 | | 18.236 | -0.17057 | -0.227509 |
| 18.2569444 | -0.045367 | -0.078063 | -0.094752 | -0.112565 | -0.104002 | -0.102115 | -0.148385 | -0.098424 | -0.124778 | -0.095813 | -0.120306 | -0.09931 | -0.169985 | -0.186338 | -0.168737 | -0.177823 | -0.177847 | -0.225732 | -0.260961 | -0.211083 | -0.190743 | -0.24857 | -0.390269 | -0.223099 | | 18.25 | -0.169985 | -0.217401 |
| 18.2708333 | -0.044968 | -0.095181 | -0.08339 | -0.116707 | -0.10631 | -0.083056 | -0.129257 | -0.088295 | -0.109062 | -0.104955 | -0.134139 | -0.109521 | -0.127274 | -0.042055 | -0.033693 | -0.015237 | -0.027944 | -0.05396 | -0.062324 | -0.040705 | -0.020329 | -0.053677 | -0.172767 | 0.026209 | | 18.264 | -0.127274 | -0.014568 |
| 18.2847222 | -0.060761 | -0.074503 | -0.083386 | -0.109668 | -0.078342 | -0.074627 | -0.088416 | -0.07629 | -0.089212 | -0.073799 | -0.120005 | -0.085015 | -0.08223 | -0.004764 | 0.0053 | 0.010982 | 0.006523 | -0.01732 | -0.027314 | -0.018182 | 0.024779 | -0.010363 | -0.095062 | 0.061056 | | 18.278 | -0.08223 | 0.031271 |
| 18.2986111 | -0.075928 | -0.083385 | -0.158704 | -0.118261 | -0.068658 | -0.104904 | -0.063753 | -0.088873 | -0.094713 | -0.06106 | -0.141703 | -0.09246 | -0.06275 | -0.111663 | -0.098105 | -0.105769 | -0.098636 | -0.11947 | -0.145663 | -0.139913 | -0.086998 | -0.122764 | -0.17968 | -0.11727 | | 18.292 | -0.06275 | -0.099111 |
| 18.3125 | -0.081993 | -0.105802 | -0.160659 | -0.054031 | -0.08473 | -0.115069 | -0.082215 | -0.097419 | -0.104487 | -0.07801 | -0.152821 | -0.106248 | -0.071298 | -0.212242 | -0.203692 | -0.21218 | -0.205404 | -0.208361 | -0.241657 | -0.248785 | -0.203386 | -0.221827 | -0.261251 | -0.254389 | | 18.306 | -0.071298 | -0.228389 |
| 18.3263889 | -0.100177 | -0.088297 | -0.062998 | 0.047719 | -0.085545 | -0.081533 | -0.110852 | -0.080751 | -0.098473 | -0.083405 | -0.095042 | -0.083895 | -0.104079 | -0.227971 | -0.223423 | -0.245802 | -0.236286 | -0.224882 | -0.26538 | -0.277762 | -0.230028 | -0.22955 | -0.245683 | -0.292184 | | 18.319 | -0.104079 | -0.286728 |
| 18.3402778 | -0.114922 | -0.068188 | 0.000591 | 0.065853 | -0.07288 | -0.045034 | -0.114613 | -0.057744 | -0.081782 | -0.08419 | -0.022586 | -0.048841 | -0.14358 | -0.139371 | -0.151633 | -0.17834 | -0.153983 | -0.126406 | -0.182048 | -0.195902 | -0.138142 | -0.121229 | -0.127467 | -0.235619 | | 18.333 | -0.14358 | -0.236168 |
| 18.3541667 | -0.06341 | -0.048733 | -0.012403 | -0.002782 | -0.049291 | -0.038846 | -0.077455 | -0.041555 | -0.052296 | -0.067155 | 0.005935 | -0.02621 | -0.112742 | -0.008329 | -0.053101 | -0.053685 | -0.03693 | 0.012874 | -0.035081 | -0.065584 | -0.036282 | 0.023517 | 0.002991 | -0.089811 | | 18.347 | -0.112742 | -0.082623 |
| 18.3680556 | 0.011899 | -0.01572 | -0.046887 | -0.091497 | -0.018273 | -0.046504 | -0.018329 | -0.028976 | -0.025865 | -0.029503 | -0.009722 | -0.026679 | -0.020642 | 0.079906 | 0.023035 | 0.049686 | 0.033154 | 0.099923 | 0.079546 | 0.026742 | -0.006612 | 0.11725 | 0.087819 | 0.068229 | | 18.361 | -0.020642 | 0.074842 |
| 18.3819444 | 0.034201 | 0.008411 | -0.045439 | -0.116028 | 0.000891 | -0.034912 | 0.020009 | -0.00607 | -0.009716 | -0.001871 | -0.012792 | -0.019817 | 0.038488 | 0.122316 | 0.098465 | 0.123455 | 0.098544 | 0.159608 | 0.153846 | 0.098147 | 0.037612 | 0.188281 | 0.16718 | 0.175668 | | 18.375 | 0.038488 | 0.174432 |
| 18.3958333 | 0.013153 | -0.004336 | -0.03077 | -0.083767 | -0.006924 | -0.026631 | 0.00719 | -0.005661 | -0.013505 | -0.010821 | -0.00463 | -0.012432 | 0.015364 | 0.11442 | 0.123117 | 0.1308 | 0.128265 | 0.180977 | 0.162844 | 0.119603 | 0.085135 | 0.21882 | 0.196253 | 0.183759 | | 18.389 | 0.015364 | 0.169982 |
| [illegible] | [illegible] | [illegible] | [illegible] | [illegible] | [illegible] | [illegible] | [illegible] | [illegible] | [illegible] | [illegible] | [illegible] | [illegible] | [illegible] | [illegible] | [illegible] | [illegible] | [illegible] | [illegible] | [illegible] | [illegible] | [illegible] | [illegible] | [illegible] | [illegible] | | [illegible] | [illegible] | [illegible] |

| | | | | | | | | | |
|---|---|---|---|---|---|---|---|---|---|
| 21.5625 | -0.501354 | -1.911992 | 4.063564 | 3.42038 | -1.252795 | 1.010278 | -3.228059 | 0.381867 | -0.608295 |
| 21.5763889 | 0.190344 | -1.442168 | 1.242512 | 1.336558 | -0.796957 | 0.284509 | -1.483714 | 0.289617 | -0.327773 |
| 21.5902778 | 0.124095 | -0.302885 | 0.073306 | 0.180232 | -0.157671 | 0.088578 | 0.194423 | 0.270727 | 0.087672 |
| 21.6041667 | 0.256334 | 0.073074 | -0.050303 | 0.071825 | 0.087994 | 0.15508 | 0.527277 | 0.37214 | 0.227087 |
| 21.6180556 | 0.244197 | 0.036301 | -0.11585 | 0.025169 | 0.052362 | 0.108585 | 0.401717 | 0.227324 | 0.159974 |
| 21.6319444 | 0.114978 | -0.030339 | -0.137403 | 0.017584 | -0.006955 | 0.038572 | 0.511035 | 0.085806 | 0.073403 |
| 21.6458333 | -0.022954 | -0.0429 | -0.077901 | 0.081614 | -0.022071 | 0.019725 | 0.555227 | -0.00385 | 0.031393 |
| 21.6597222 | -0.118348 | -0.036755 | 0.034155 | 0.144077 | -0.016749 | 0.028103 | 0.326259 | -0.060734 | 0.065418 |
| 21.6736111 | -0.105119 | -0.018691 | 0.105345 | 0.155519 | -0.001037 | 0.050984 | 0.186684 | -0.078579 | 0.164223 |
| 21.6875 | -0.001587 | -0.007808 | 0.070968 | 0.107876 | 0.004455 | 0.052884 | 0.093818 | -0.067017 | 0.214879 |
| 21.7013889 | 0.106367 | -0.020794 | 0.012932 | 0.039153 | -0.007281 | 0.030445 | 0.013505 | -0.046592 | 0.146085 |
| 21.7152778 | 0.165985 | -0.039942 | -0.046227 | 0.0131 | 0.000494 | 0.015701 | -0.03175 | -0.0053 | 0.034215 |
| 21.7291667 | 0.161329 | -0.031344 | -0.112995 | 0.006194 | 0.022722 | 0.017645 | -0.03495 | 0.022711 | -0.072987 |
| 21.7430556 | 0.135335 | -0.006335 | -0.06496 | -0.045134 | 0.036585 | 0.008279 | -0.028423 | 0.029785 | -0.157494 |
| 21.7569444 | 0.160875 | 0.009912 | 0.061978 | -0.03944 | 0.050891 | 0.010955 | 0.014092 | 0.040547 | -0.053272 |
| 21.7708333 | 0.222465 | 0.018586 | 0.172905 | 0.01542 | 0.079503 | 0.025011 | 0.047948 | 0.02307 | 0.21543 |
| 21.7847222 | 0.266092 | 0.022251 | 0.217247 | 0.065331 | 0.099829 | 0.052388 | 0.102282 | 0.01885 | 0.251814 |
| 21.7986111 | 0.277173 | 0.017259 | 0.170211 | 0.100858 | 0.092406 | 0.060148 | 0.109244 | -0.007722 | 0.176781 |
| 21.8125 | 0.205995 | 0.004245 | 0.072991 | 0.100291 | 0.057786 | 0.047872 | 0.04007 | -0.056393 | 0.049478 |
| 21.8263889 | 0.048133 | -0.015083 | 0.008071 | 0.062159 | 0.023834 | 0.025447 | -0.11615 | -0.081315 | -0.146991 |
| 21.8402778 | -0.125309 | -0.019811 | -0.005802 | -0.004613 | 0.00472 | -0.013772 | -0.249658 | -0.154048 | -0.278774 |
| 21.8541667 | -0.281535 | -0.017392 | 0.003259 | -0.090423 | -0.018937 | -0.059498 | -0.378733 | -0.181409 | -0.262876 |
| 21.8680556 | -0.391975 | -0.015773 | 0.010045 | -0.152234 | -0.041719 | -0.083774 | -0.9518 | -0.178478 | -0.128756 |
| 21.8819444 | -0.456434 | -0.033935 | -0.005081 | -0.188998 | -0.085493 | -0.112003 | -1.268583 | -0.189329 | -0.041215 |
| [illegible] | [illegible] | [illegible] | [illegible] | [illegible] | [illegible] | [illegible] | [illegible] | [illegible] | [illegible] |

| | | | | | | | | | | | | | | | | | | | | | | | | | | | | |
|---|---|---|---|---|---|---|---|---|---|---|---|---|---|---|---|---|---|---|---|---|---|---|---|---|---|---|---|---|
| 25.0486111 | -1.029236 | 2.096274 | -4.464197 | -2.533581 | 0.731167 | -1.226267 | 1.151992 | -1.122082 | -0.422757 | 0.444661 | -0.708118 | -0.682191 | 7.52081 | -0.699622 | 0.77144 | 0.215175 | -1.307773 | -0.932132 | -0.040162 | -1.260669 | -1.746274 | -1.454069 | -0.529964 | 3.23058 | | 25.042 | 7.52081 | 3.719095 |
| 25.0625 | -0.568451 | 1.340348 | -3.632157 | -2.392387 | 0.582116 | -1.701122 | 0.882847 | -2.080322 | -0.833635 | 0.00274 | -1.085176 | -1.129276 | 5.613678 | -0.219764 | 0.415431 | -0.522748 | -1.580084 | -1.167964 | 0.412179 | -1.056637 | -1.618287 | -1.372983 | -1.022674 | 1.916525 | | 25.056 | 5.613678 | 2.683657 |
| 25.0763889 | -0.492583 | 0.326409 | -1.128601 | -1.297271 | 0.177428 | -0.866393 | -0.300639 | -2.191791 | -1.217454 | -0.534347 | -0.547532 | -1.49778 | 1.805577 | -0.194157 | 0.528336 | -0.043934 | -0.667968 | -0.508673 | 0.369817 | -0.33896 | -0.540861 | -1.524864 | -0.422695 | 0.542247 | | 25.069 | 1.805577 | 0.889118 |
| 25.0902778 | -0.379322 | -0.053954 | -0.057274 | -0.683346 | -0.080466 | -0.259373 | -1.338056 | -1.340618 | -1.140262 | -0.626978 | -0.493785 | -1.416666 | -0.29655 | -0.268943 | 0.523853 | 0.218185 | -0.107988 | 0.005425 | 0.020831 | -0.436074 | -0.206019 | -2.230953 | -0.182984 | -0.067916 | | 25.083 | -0.29655 | 0.047814 |
| 25.1041667 | -0.214935 | -0.155981 | 0.065711 | -0.438591 | -0.144422 | -0.054072 | -1.827357 | -0.593885 | -0.678624 | -0.30844 | -0.728317 | -0.814714 | -0.440813 | -0.43391 | 0.33848 | 0.008072 | -0.080757 | -0.083469 | -0.362575 | -0.516521 | -0.257809 | -1.444099 | -0.14196 | -0.258908 | | 25.097 | -0.440813 | -0.226257 |
| 25.1180556 | 0.094551 | -0.112336 | 0.18696 | -0.192244 | -0.133858 | 0.117764 | -1.368144 | -0.039581 | -0.116226 | 0.060495 | -0.670592 | -0.294555 | -0.207832 | -0.421175 | 0.106463 | -0.221612 | -0.097047 | -0.235156 | -0.586979 | -0.515928 | -0.364787 | -0.475538 | 0.114682 | -0.236228 | | 25.111 | -0.207832 | -0.287056 |
| 25.1319444 | 0.529468 | 0.006509 | 0.280698 | 0.042605 | -0.06931 | 0.292265 | -0.686082 | 0.408694 | 0.359045 | 0.277782 | -0.273871 | 0.10143 | -0.007948 | -0.366055 | -0.116134 | -0.320457 | -0.10325 | -0.224175 | -0.501704 | -0.463708 | -0.471377 | -0.318775 | 0.17305 | -0.276258 | | 25.125 | -0.007948 | -0.304634 |
| 25.1458333 | 0.635879 | 0.179933 | 0.181706 | 0.181284 | 0.049942 | 0.361219 | -0.00418 | 0.733207 | 0.603629 | 0.353962 | 0.22249 | 0.531303 | 0.244837 | -0.262078 | -0.209701 | -0.251536 | -0.076938 | 0.002102 | -0.163359 | -0.372959 | -0.484887 | -0.2333 | 0.149732 | -0.283775 | | 25.139 | 0.244837 | -0.28625 |
| 25.1597222 | 0.341973 | 0.392132 | -0.183188 | 0.250139 | 0.199683 | 0.254505 | 0.616227 | 0.773396 | 0.650763 | 0.281752 | 0.85281 | 0.956946 | 0.479272 | -0.11333 | -0.161599 | -0.14732 | -0.077544 | 0.202318 | -0.035129 | -0.247453 | -0.38761 | -0.067138 | 0.120994 | -0.203802 | | 25.153 | 0.479272 | -0.216965 |
| 25.1736111 | 0.139524 | 0.366201 | -0.362201 | 0.233259 | 0.187096 | 0.073872 | 0.882163 | 0.459034 | 0.565779 | 0.177158 | 1.627081 | 1.08008 | 0.502475 | -0.061798 | -0.132569 | -0.151299 | -0.175297 | 0.222064 | -0.209669 | -0.062381 | -0.277668 | 0.156682 | 0.039375 | -0.139555 | | 25.167 | 0.502475 | -0.179017 |
| 25.1875 | 0.142055 | -0.076773 | -0.292519 | 0.116401 | 0.068491 | -0.015405 | 0.472476 | 0.03378 | 0.379836 | 0.113137 | 0.671714 | 0.630343 | 0.368407 | -0.102969 | -0.171453 | -0.256888 | -0.258401 | 0.095577 | -0.578439 | -0.116409 | -0.108257 | 0.425091 | -0.075129 | -0.13524 | | 25.181 | 0.368407 | -0.165729 |
| 25.2013889 | -0.046905 | -0.315237 | 0.031177 | 0.05215 | 0.060555 | 0.021785 | -0.118494 | 0.166343 | 0.537315 | 0.176855 | -0.322416 | 0.033065 | 0.301035 | -0.174257 | -0.227569 | -0.366113 | -0.245598 | -0.065486 | -1.200915 | -0.492535 | 0.038135 | 0.194545 | -0.191707 | -0.20729 | | 25.194 | 0.301035 | -0.185465 |
| 25.2152778 | -0.392022 | 0.037124 | 0.334603 | 0.111111 | 0.185478 | 0.118872 | -0.326437 | 0.645609 | 0.854387 | 0.25637 | -0.402494 | -0.384294 | 0.3539 | -0.138236 | -0.202778 | -0.298151 | -0.123136 | -0.158222 | -1.232025 | -0.705317 | 0.029221 | -0.083703 | -0.19548 | -0.235379 | | 25.208 | 0.3539 | -0.165822 |
| 25.2291667 | -0.368838 | 0.288158 | 0.553487 | 0.249727 | 0.253187 | 0.156508 | 0.208821 | 0.755706 | 0.51079 | 0.262281 | 0.034539 | -0.484464 | 0.296565 | 0.067115 | -0.100373 | -0.183963 | 0.026412 | -0.17868 | -0.469971 | -0.263728 | -0.057788 | -0.027135 | -0.069002 | -0.051891 | | 25.222 | 0.296565 | 0.001508 |
| 25.2430556 | -0.006107 | 0.094711 | 0.405952 | 0.109055 | 0.21732 | 0.135949 | 1.06686 | 0.442152 | -0.096466 | 0.161491 | 0.090744 | -0.254922 | 0.092792 | 0.294563 | -0.024617 | -0.134128 | 0.048795 | -0.202663 | 0.194612 | 0.545093 | -0.091949 | 0.230716 | 0.07927 | 0.176476 | | 25.236 | 0.092792 | 0.202809 |
| 25.2569444 | 0.164178 | -0.033341 | -0.08673 | 0.104654 | 0.214801 | 0.144191 | 1.766042 | 0.412721 | -0.231378 | 0.072311 | -0.525801 | 0.358339 | 0.141287 | 0.222072 | -0.073112 | -0.198638 | -0.118523 | -0.265339 | 0.704448 | 0.823827 | -0.051307 | 0.324251 | 0.269573 | 0.22871 | | 25.25 | 0.141287 | 0.244822 |
| 25.2708333 | -0.082125 | 0.014543 | -0.502063 | 0.127181 | 0.224802 | 0.174072 | 1.865882 | 0.646587 | 0.192034 | 0.055055 | -0.539674 | 1.186318 | 0.441002 | 0.08063 | -0.091937 | -0.272425 | -0.220188 | -0.244811 | 0.460321 | 0.339944 | 0.06045 | 0.22217 | 0.461184 | 0.190719 | | 25.264 | 0.441002 | 0.201503 |
| 25.2847222 | -0.253036 | 0.100027 | -0.222701 | 0.195729 | 0.101661 | 0.131515 | 1.320807 | 0.793556 | 0.813422 | 0.08234 | -0.150467 | 1.338661 | 0.562991 | 0.032953 | 0.094656 | -0.232623 | -0.098264 | -0.143167 | 0.021926 | -0.06885 | 0.23594 | 0.122313 | 0.158251 | 0.116889 | | 25.278 | 0.562991 | 0.1715 |
| 25.2986111 | -0.164078 | 0.048944 | 0.149507 | 0.100328 | -0.06821 | 0.013493 | 0.527122 | 0.485241 | 0.8002 | 0.096816 | -0.120906 | 0.084874 | 0.365686 | -0.191212 | 0.227147 | -0.237822 | -0.003364 | -0.141669 | -0.370117 | 0.360517 | 0.212845 | -0.026071 | -0.308642 | -0.000657 | | 25.292 | 0.365686 | 0.07484 |
| 25.3125 | 0.036351 | -0.054411 | -0.257086 | -0.067568 | -0.105347 | -0.067184 | 0.149263 | 0.221848 | 0.054713 | 0.053453 | -0.079383 | -0.451066 | 0.166527 | -0.205014 | 0.074311 | -0.291467 | -0.049421 | -0.189771 | -0.591351 | 0.723516 | 0.083533 | -0.171278 | -0.505878 | -0.098665 | | 25.306 | 0.166527 | -0.058802 |
| 25.3263889 | 0.192578 | -0.058272 | -0.309245 | 0.079055 | -0.024369 | 0.030764 | 0.245886 | 0.274745 | -0.311002 | -0.004201 | 0.131359 | -0.821836 | 0.064591 | 0.191144 | -0.170311 | -0.243817 | -0.097878 | -0.085464 | -0.400661 | 0.663732 | 0.078455 | -0.173797 | -0.361521 | -0.018023 | | 25.319 | 0.064591 | -0.02789 |
| 25.3402778 | 0.256235 | 0.002887 | 0.371165 | 0.211475 | -0.020159 | 0.136772 | 0.753145 | 0.677643 | 0.008253 | 0.003014 | 0.239838 | -0.615289 | 0.04654 | 0.460676 | -0.283405 | -0.119866 | -0.02467 | 0.164957 | -0.021701 | 0.601047 | 0.286039 | -0.083871 | -0.111991 | 0.170067 | | 25.333 | 0.04654 | 0.174195 |
| 25.3541667 | 0.170487 | 0.065618 | 0.87711 | 0.156081 | -0.099355 | 0.071023 | 1.085084 | 0.576225 | 0.314404 | 0.045061 | 0.213993 | 0.18719 | 0.102602 | 0.362766 | -0.156056 | -0.065598 | 0.063805 | 0.332612 | 0.188511 | 0.511826 | 0.48814 | 0.096528 | 0.029708 | 0.278962 | | 25.347 | 0.102602 | 0.325042 |
| 25.3680556 | -0.097219 | 0.063291 | 0.658275 | 0.145458 | 0.000371 | -0.002643 | 0.864268 | 0.292191 | 0.296746 | 0.026951 | 0.267841 | 0.421073 | 0.169456 | 0.20682 | -0.064385 | -0.157397 | -0.014428 | 0.219963 | 0.080208 | 0.376683 | 0.521788 | 0.205895 | -0.034127 | 0.192377 | | 25.361 | 0.169456 | 0.243882 |
| 25.3819444 | -0.24456 | 0.052945 | -0.012402 | 0.157375 | 0.205401 | 0.03091 | 0.587358 | 0.400854 | 0.026384 | -0.022859 | 0.329445 | 0.292384 | 0.162142 | 0.20883 | -0.054527 | -0.260393 | -0.078697 | 0.077418 | -0.036414 | 0.144658 | 0.451931 | 0.225605 | -0.133986 | 0.122563 | | 25.375 | 0.162142 | 0.158649 |
| 25.3958333 | -0.058077 | 0.058698 | -0.414669 | 0.192446 | 0.251921 | 0.084913 | 0.781081 | 0.43089 | -0.087049 | 0.010751 | 0.408118 | 0.210381 | 0.163513 | 0.325977 | 0.015681 | -0.231728 | 0.040957 | 0.051482 | 0.050302 | 0.142127 | 0.426661 | 0.193354 | -0.102177 | 0.190187 | | 25.389 | 0.163513 | 0.25332 |
| 25.4097222 | 0.289947 | 0.068376 | 0.038139 | 0.205535 | 0.052544 | 0.090266 | 1.085932 | 0.558441 | 0.148553 | 0.075234 | 0.61288 | 0.178715 | 0.172465 | 0.416483 | -0.001599 | -0.186757 | 0.173002 | 0.071165 | 0.120318 | 0.255087 | 0.295052 | 0.284122 | -0.030581 | 0.206871 | | 25.403 | 0.172465 | 0.285257 |
| 25.4236111 | 0.565457 | 0.098796 | 0.621298 | 0.335654 | -0.044926 | 0.142812 | 1.107842 | 0.135882 | 0.494433 | 0.086974 | 0.84084 | 0.187881 | 0.24332 | 0.586916 | -0.048813 | -0.107546 | 0.23891 | 0.251944 | 0.191061 | 0.331255 | 0.154012 | 0.497164 | 0.089784 | 0.190266 | | 25.417 | 0.24332 | 0.224473 |
| 25.4375 | 0.566564 | 0.181644 | 0.636744 | 0.461428 | 0.049969 | 0.205292 | 1.057366 | 0.114487 | 0.449109 | 0.087827 | 0.901069 | 0.301062 | 0.314835 | 0.771715 | -0.040757 | -0.082316 | 0.21879 | 0.586186 | 0.193827 | 0.259663 | 0.100902 | 0.485667 | 0.22329 | 0.107426 | | 25.431 | 0.314835 | 0.087635 |
| 25.4513889 | 0.27415 | 0.124887 | 0.37768 | 0.229289 | 0.116302 | 0.191136 | 0.865404 | 0.521005 | 0.29829 | 0.034389 | 0.894954 | 0.353266 | 0.300437 | 0.640352 | -0.126697 | -0.178446 | 0.059132 | 0.755088 | -0.01679 | -0.061947 | -0.039927 | 0.05384 | -0.062628 | -0.189052 | | 25.444 | 0.300437 | -0.210775 |
| 25.4652778 | 0.027098 | -0.114739 | 0.156642 | -0.108865 | 0.001846 | 0.038956 | 0.654368 | 0.499391 | 0.237655 | -0.111477 | 1.081274 | 0.208789 | 0.049566 | 0.352529 | -0.2562 | -0.392515 | -0.118925 | 0.70161 | -0.238752 | -0.395255 | -0.262609 | -0.342434 | -0.365194 | -0.495417 | | 25.458 | 0.049566 | -0.489404 |
| 25.4791667 | 0.083633 | -0.258342 | 0.045821 | -0.228109 | -0.147693 | -0.066724 | 0.805925 | 0.062424 | 0.475617 | -0.189154 | 0.766751 | 0.02872 | -0.115686 | 0.124306 | -0.342383 | -0.462106 | -0.194107 | 0.640346 | -0.348775 | -0.678332 | -0.232524 | -0.462338 | -0.655248 | -0.556428 | | 25.472 | -0.115686 | -0.514877 |
| 25.4930556 | 0.52876 | -0.158687 | -0.359902 | -0.060271 | -0.090187 | 0.01441 | 1.231077 | -0.340083 | 0.806667 | -0.073433 | 0.988274 | 0.220772 | -0.087878 | 0.055579 | -0.303311 | -0.261166 | -0.221024 | 0.523827 | -0.193119 | -0.571646 | -0.111401 | -0.304496 | -0.550715 | -0.409199 | | 25.486 | -0.087878 | -0.384146 |
| 25.5069444 | 1.100115 | -0.048223 | -0.621487 | 0.097238 | 0.091729 | 0.104498 | 1.508384 | -0.43792 | 0.783237 | 0.078532 | 0.917759 | 0.632942 | 0.060508 | 0.02996 | -0.23511 | -0.091635 | -0.223849 | 0.450504 | -0.081332 | -0.325797 | -0.02965 | -0.240635 | -0.399964 | -0.269021 | | 25.5 | 0.060508 | -0.238222 |
| 25.5208333 | 1.266423 | -0.027539 | -0.837084 | 0.398721 | 0.210359 | 0.179966 | 1.304069 | 0.218611 | 0.520859 | 0.085728 | 0.933644 | 0.98779 | -0.186047 | -0.047966 | -0.273369 | -0.05994 | -0.216574 | 0.378125 | -0.187157 | -0.173848 | -0.117464 | -0.252588 | -0.307969 | -0.245128 | | 25.514 | -0.186047 | -0.260384 |
| 25.5347222 | 0.84337 | -0.487606 | -0.11075 | 1.734732 | 0.154366 | 0.538281 | 0.520584 | 0.841154 | 0.061627 | -0.011624 | 1.353213 | 1.247372 | -2.5804 | -0.048586 | -0.442351 | -0.194608 | -0.010618 | 0.499304 | -0.748827 | 0.194566 | -0.17424 | -0.117005 | 0.277126 | -0.343401 | | 25.528 | -2.5804 | -0.776549 |
| 25.5486111 | 1.128483 | -1.478477 | 1.851025 | 3.838349 | -0.244386 | 1.007921 | -0.401597 | 1.574869 | -0.73843 | 0.149428 | 2.041934 | 1.490994 | -8.198129 | -0.140318 | -0.930139 | -0.506199 | 0.29857 | 0.52191 | -1.864225 | 1.24624 | -0.004535 | -0.035222 | 1.102797 | -2.194453 | | 25.542 | -8.198129 | -3.211197 |
| 25.5625 | 0.764639 | -2.092795 | 2.031667 | 3.996319 | -0.51832 | 1.038533 | -0.01673 | 1.127852 | -1.172729 | -0.049307 | 1.356766 | 1.447263 | -9.602041 | 0.417521 | -1.361185 | -1.168776 | -0.454965 | -0.193489 | -1.915214 | 1.785314 | -0.591718 | -0.483281 | 0.729348 | -5.368836 | | 25.556 | -9.602041 | -4.910626 |
| 25.5763889 | 0.571168 | -1.218893 | 0.901109 | 1.838715 | -0.25494 | 0.439458 | 0.89594 | 0.085063 | -0.481314 | -0.200753 | 1.189299 | 0.871863 | -4.420104 | 0.979244 | -0.995456 | -0.686761 | -0.745151 | -0.547191 | -1.047711 | 0.973028 | -0.845827 | -0.479696 | -0.172944 | -5.092175 | | 25.569 | -4.420104 | -4.515995 |
| 25.5902778 | 0.276407 | -0.100431 | 0.575531 | 0.510115 | -0.1013 | 0.180386 | 1.343612 | -0.499367 | -0.133001 | -0.095593 | 0.601824 | 0.340944 | -0.893811 | 0.118964 | -0.506217 | 0.115826 | -0.314707 | -0.162797 | -0.201222 | 0.341837 | -0.38425 | -0.206059 | -0.133111 | -0.936658 | | 25.583 | -0.893811 | -0.893515 |
| 25.6041667 | 0.339962 | 0.134874 | 0.537647 | 0.300519 | -0.047868 | 0.183633 | 0.937199 | -0.620275 | 0.178437 | 0.041266 | 0.301083 | 0.419218 | -0.392947 | -0.008157 | -0.403888 | 0.369857 | -0.000634 | -0.022233 | 0.103888 | 0.461646 | -0.085233 | 0.098219 | 0.085441 | 0.162611 | | 25.597 | -0.392947 | 0.246918 |
| 25.6180556 | 0.286102 | 0.143762 | 0.149963 | 0.160245 | -0.016034 | 0.133851 | 0.465412 | -0.119649 | 0.145107 | 0.111148 | -0.05778 | 0.63207 | -0.252431 | 0.050578 | -0.302053 | 0.328716 | 0.10826 | -0.053382 | 0.144181 | 0.459257 | 0.235752 | 0.268128 | 0.157221 | 0.307734 | | 25.611 | -0.252431 | 0.37364 |
| 25.6319444 | 0.043733 | 0.077426 | -0.265719 | 0.051559 | 0.048818 | 0.096978 | 0.214902 | 0.403956 | 0.077061 | 0.045817 | -0.396182 | 0.84549 | -0.060564 | 0.066372 | -0.063294 | 0.200866 | 0.09264 | -0.207913 | 0.085192 | 0.302698 | 0.165148 | 0.277466 | 0.112478 | 0.273255 | | 25.625 | -0.060564 | 0.301962 |
| 25.6458333 | -0.265694 | -0.060007 | -0.781505 | -0.028253 | 0.045945 | 0.16304 | 0.077945 | 0.622811 | 0.021229 | -0.061849 | -0.646555 | 1.165767 | 0.114614 | -0.122143 | -0.037547 | 0.031093 | 0.008253 | -0.257381 | -0.036027 | 0.035501 | 0.026676 | 0.124032 | -0.037439 | 0.14936 | | 25.639 | 0.114614 | 0.166434 |
| 25.6597222 | -0.525394 | -0.133724 | -0.809716 | -0.094227 | 0.115858 | 0.150768 | -0.042098 | 0.642435 | 0.24091 | -0.059308 | -0.39403 | 1.257261 | 0.215562 | -0.363517 | -0.074886 | -0.10382 | -0.043976 | -0.262609 | -0.080705 | -0.21643 | 0.034526 | 0.00231 | -0.185496 | 0.071177 | | 25.653 | 0.215562 | 0.079733 |
| 25.6736111 | -0.565137 | -0.127624 | -0.303814 | 0.046371 | 0.042269 | 0.080904 | -0.223358 | -0.056298 | 0.071793 | -0.016853 | -0.284304 | 0.787165 | 0.054635 | -0.534362 | -0.072621 | -0.196568 | -0.079898 | -0.214254 | -0.077976 | -0.288291 | -0.035471 | -0.07301 | -0.286397 | 0.005163 | | 25.667 | 0.054635 | -0.002878 |
| 25.6875 | -0.412707 | -0.018378 | 0.019205 | 0.2244 | -0.162251 | 0.020046 | -0.47979 | -0.454544 | -0.094751 | -0.033854 | -0.179509 | 0.143498 | -0.232237 | -0.435756 | -0.079823 | -0.268222 | -0.112595 | -0.138004 | -0.048708 | -0.219656 | -0.201414 | -0.106517 | -0.275312 | -0.083995 | | 25.681 | -0.232237 | -0.111245 |

| | | | | | | | | | | | | | | | | | | | | | | | | | | | | |
|---|---|---|---|---|---|---|---|---|---|---|---|---|---|---|---|---|---|---|---|---|---|---|---|---|---|---|---|---|
| 28.5347222 | 1.737547 | -0.22818 | 0.846624 | 0.685312 | 0.936488 | 1.684731 | 0.257775 | 0.904384 | 0.87006 | 1.11415 | 3.061225 | 1.176896 | -0.005185 | 0.372556 | -0.046146 | 0.102537 | 0.12946 | 0.510772 | -1.009129 | -0.610869 | 0.220949 | -0.429489 | -1.125033 | 2.655077 | | 28.528 | -0.005185 | 1.385615 |
| 28.5486111 | 2.791694 | -1.711493 | 1.522388 | 1.983735 | 0.383881 | 1.49794 | -1.586513 | 0.022497 | 0.159195 | 0.228727 | 2.623365 | 1.307174 | -0.183329 | 0.39167 | -1.141602 | -0.098549 | 0.326413 | -0.099771 | -2.761371 | -1.370654 | -0.151485 | -1.189519 | -3.55414 | 4.23595 | | 28.542 | -0.183329 | 3.512657 |
| 28.5625 | 2.834762 | -2.041083 | 1.416438 | 2.102135 | 0.128374 | 1.510994 | -1.056505 | -0.068467 | 0.663613 | -0.359622 | 2.700961 | 1.846525 | -1.408622 | 1.529774 | -2.087929 | -0.786871 | -0.536101 | -1.079962 | -3.159718 | -1.508853 | -0.561881 | -1.877004 | -4.980249 | 3.795514 | | 28.556 | -1.408622 | 0.81087 |
| 28.5763889 | 1.700006 | -1.630885 | 0.384022 | 0.751889 | -0.053688 | 0.646125 | 0.17218 | 0.34475 | 0.717141 | -0.583168 | 1.985664 | 1.583832 | -0.362387 | 1.41921 | -1.178091 | -0.608369 | -0.480445 | -1.123736 | -1.765373 | -1.220667 | 0.171648 | -1.395825 | -1.706509 | 0.254071 | | 28.569 | -0.362387 | -0.419732 |
| 28.5902778 | 1.129541 | -0.287124 | -0.140373 | 0.152224 | -0.139748 | -0.02161 | 1.418668 | 0.08864 | 0.406001 | -0.12086 | 1.50782 | 1.252745 | -1.197965 | 0.16952 | -0.292794 | -0.609561 | -0.01116 | -0.437146 | -0.718511 | -0.603157 | 1.313656 | -0.652112 | -0.113074 | -0.215277 | | 28.583 | -1.197965 | -0.326187 |
| 28.6041667 | 0.721841 | 0.049303 | -0.403052 | 0.38377 | -0.365763 | -0.220521 | 0.832058 | -0.079006 | -0.229087 | 0.257437 | 1.245006 | 1.347643 | -1.440581 | -0.001567 | 0.087057 | -0.805238 | 0.212874 | -0.113417 | -0.207621 | -0.195383 | 1.061579 | -0.191367 | 1.039234 | -0.044428 | | 28.597 | -1.440581 | 0.030642 |
| 28.6180556 | 0.302509 | 0.042848 | 0.150848 | 0.262043 | -0.385243 | -0.104757 | 0.513415 | -0.069533 | 0.103221 | -0.175347 | 0.875416 | 1.227043 | -0.476564 | 0.115673 | 0.131187 | -0.461132 | 0.116663 | 0.013234 | -0.050009 | -0.064798 | 0.299881 | -0.020815 | 1.916335 | -0.028986 | | 28.611 | -0.476564 | -0.021816 |
| 28.6319444 | 0.073632 | -0.014706 | 0.913055 | 0.136405 | -0.57404 | 0.148852 | 0.411486 | 0.42331 | 0.286857 | -0.154535 | 0.201933 | 0.710083 | 1.572377 | 0.535971 | -0.099644 | 0.003072 | -0.23596 | 0.029382 | -0.066529 | -0.082741 | -0.841885 | -0.033462 | 0.526013 | -0.252033 | | 28.625 | 1.572377 | -0.250147 |
| 28.6458333 | -0.163482 | -0.040012 | 0.394114 | 0.097273 | -0.406901 | 0.355406 | 0.53918 | 0.326877 | -0.383038 | -0.105586 | -0.955281 | 0.191329 | 2.179347 | 0.230491 | -0.22296 | 0.066228 | -0.566741 | -0.073541 | -0.133194 | -0.136142 | -1.80331 | -0.116436 | -0.140221 | -0.365722 | | 28.639 | 2.179347 | -0.418945 |
| 28.6597222 | -0.34746 | -0.042512 | -0.542194 | 0.021813 | -0.561801 | 0.745967 | 0.507972 | 0.068267 | -0.507412 | 0.064917 | -2.016465 | 0.103178 | 1.252839 | -0.007661 | -0.221759 | -0.021371 | -0.717128 | -0.143377 | -0.169961 | -0.169416 | -0.792561 | -0.170358 | -0.669095 | -0.222063 | | 28.653 | 1.252839 | -0.287244 |
| 28.6736111 | -0.362078 | -0.016056 | -1.026376 | 0.030582 | -1.363446 | 0.761199 | 0.072921 | -0.196714 | 0.181817 | -0.145302 | -1.658252 | 0.097629 | 0.547856 | 0.036619 | -0.361136 | -0.09116 | -0.532344 | -0.10867 | -0.1701 | -0.165174 | 0.613126 | -0.170581 | -1.053475 | 0.077492 | | 28.667 | 0.547856 | -0.036156 |
| 28.6875 | -0.200167 | 0.006848 | -0.977531 | 0.236038 | -0.53043 | 0.53449 | -0.414156 | -0.551708 | 0.532118 | -0.027607 | -1.225844 | 0.045836 | -0.160801 | -0.408002 | -0.302455 | -0.108129 | -0.175969 | 0.03503 | -0.109206 | -0.109811 | 1.186319 | -0.099595 | -1.292354 | 0.28153 | | 28.681 | -0.160801 | 0.173558 |
| 28.7013889 | -0.111943 | 0.012136 | -0.379791 | 0.333902 | -0.161601 | 0.144987 | -0.401564 | -0.107631 | 0.407763 | 1.019108 | -0.945966 | -0.098645 | -0.409414 | -1.226564 | 0.025536 | -0.099618 | 0.117725 | 0.207124 | 0.023295 | 0.007278 | 0.810229 | 0.035712 | -1.307857 | 0.29797 | | 28.694 | -0.409414 | 0.125665 |
| 28.7152778 | -0.235372 | 0.027814 | 0.278413 | 0.319673 | -0.190699 | -0.07349 | 0.312929 | -0.043576 | -0.201556 | 0.726204 | 0.350473 | -0.266286 | -0.244382 | -1.082677 | 0.25792 | -0.097473 | 0.310426 | 0.309789 | 0.144895 | 0.113877 | -0.152178 | 0.158223 | -0.567822 | 0.134066 | | 28.708 | -0.244382 | 0.011168 |
| 28.7291667 | -0.116141 | 0.061874 | 0.53357 | 0.241774 | 0.016162 | -0.135669 | 0.97946 | 0.252165 | -0.200479 | -0.126012 | 1.317693 | -0.305028 | 0.292621 | 0.664778 | 0.203425 | -0.140295 | 0.353146 | 0.314835 | 0.176082 | 0.139878 | -0.921395 | 0.191093 | 0.341576 | -0.0704 | | 28.722 | 0.292621 | -0.06938 |
| 28.7430556 | 0.1888 | 0.06008 | 0.365487 | 0.231755 | -0.515587 | -0.140272 | 1.128503 | 0.246449 | 0.099517 | -0.154709 | 1.969586 | -0.234691 | 0.697986 | 1.347572 | 0.123191 | -0.186686 | 0.337107 | 0.248885 | 0.122576 | 0.087101 | -1.578556 | 0.130474 | 1.019381 | -0.236968 | | 28.736 | 0.697986 | -0.19795 |
| 28.7569444 | 0.358806 | 0.040659 | 0.817376 | 0.04598 | 0.030148 | -0.10689 | 0.694056 | -0.08789 | -0.021148 | 0.180847 | 2.249394 | -0.089391 | 0.50999 | 0.746217 | 0.150067 | -0.19491 | 0.259202 | 0.15119 | 0.034307 | 0.005293 | -1.012232 | 0.032469 | 1.10266 | -0.237804 | | 28.75 | 0.50999 | -0.220381 |
| 28.7708333 | 0.262254 | 0.016605 | 0.654596 | -0.034579 | 0.56363 | -0.064064 | -0.062008 | -0.278453 | 0.000923 | -0.28849 | 1.357518 | 0.063492 | 0.169129 | -0.125925 | -0.208811 | -0.105521 | -0.127259 | 0.066378 | -0.05319 | -0.071343 | 0.168191 | -0.056099 | 1.049206 | -0.159556 | | 28.764 | 0.169129 | -0.082375 |
| 28.7847222 | -0.107853 | -0.017153 | 0.524219 | -0.152229 | 0.256745 | -0.125765 | -0.950716 | -0.56893 | -0.243741 | -0.787803 | 0.373685 | 0.219246 | -0.930016 | -0.284515 | -0.695321 | -0.027486 | -0.293345 | -0.017247 | -0.1154 | -0.127029 | 1.074487 | -0.118847 | 1.161658 | -0.078222 | | 28.778 | -0.930016 | 0.081708 |
| 28.7986111 | -0.490292 | -0.023643 | 0.592058 | -0.200683 | 0.314164 | -0.172316 | -1.841459 | -0.683437 | -0.296534 | -0.661531 | -0.478374 | 0.395745 | -2.333166 | -1.041359 | -0.85905 | -0.021565 | -0.281438 | -0.074426 | -0.122772 | -0.140351 | 1.083042 | -0.131206 | 0.38323 | 0.023619 | | 28.792 | -2.333166 | 0.124221 |
| 28.8125 | -0.601349 | -0.035388 | -0.021007 | -0.046198 | 0.017352 | -0.290578 | -2.125768 | -0.525655 | -0.602728 | -0.268558 | -1.034056 | 0.573332 | -1.860979 | -2.324882 | -0.764896 | -0.044162 | -0.157116 | -0.109808 | -0.084373 | -0.114699 | 0.489947 | -0.095855 | -0.207252 | 0.01803 | | 28.806 | -1.860979 | 0.022509 |
| 28.8263889 | -0.667639 | -0.09039 | -0.623959 | -0.051178 | -0.013467 | -0.475217 | -1.642837 | -0.38188 | -0.560193 | 0.184772 | -1.161685 | 0.727413 | 0.010394 | -2.639363 | -0.649793 | -0.146826 | -0.120595 | -0.172888 | -0.060413 | -0.092229 | -0.351665 | -0.067911 | -0.515453 | -0.084867 | | 28.819 | 0.010394 | -0.141545 |
| 28.8402778 | -0.923956 | -0.169529 | -1.304302 | -0.084807 | 0.525789 | -0.677499 | -1.04859 | -0.262311 | -0.090441 | 0.117409 | -0.883226 | 0.651233 | 1.221217 | -1.655494 | -0.361959 | -0.293769 | -0.334576 | -0.298964 | -0.086883 | -0.092748 | -1.036115 | -0.078701 | -0.706407 | -0.2453 | | 28.833 | 1.221217 | -0.285856 |
| 28.8541667 | -0.966447 | -0.195946 | -1.222798 | 0.127908 | 0.022651 | -0.772038 | 0.086221 | -0.092398 | 0.415558 | -0.083778 | -1.471701 | 0.226377 | 0.723936 | -0.951188 | -0.428703 | -0.482357 | -0.666289 | -0.438545 | -0.101295 | -0.094371 | -1.575945 | -0.090324 | -0.848788 | -0.362387 | | 28.847 | 0.723936 | -0.408471 |
| [illegible] | [illegible] | [illegible] | [illegible] | [illegible] | [illegible] | [illegible] | [illegible] | [illegible] | [illegible] | [illegible] | [illegible] | [illegible] | [illegible] | [illegible] | [illegible] | [illegible] | [illegible] | [illegible] | [illegible] | [illegible] | [illegible] | [illegible] | [illegible] | [illegible] | | [illegible] | [illegible] | [illegible] |

| | | | | | | | | | | | | | | | |
|---|---|---|---|---|---|---|---|---|---|---|---|---|---|---|---|
| 32.0208333 | -1.925757 | -0.515483 | -3.038532 | -0.137654 | -1.07654 | -0.852057 | -1.187244 | -2.314912 | -2.098381 | -0.73152 | -0.596129 | -0.414681 | -0.931884 | -1.345344 | [illegible] |
| 32.0347222 | -1.675059 | 0.676626 | -3.681405 | -0.449608 | -0.985697 | -0.829056 | -0.765679 | -2.47647 | -1.553749 | 0.113302 | 0.002037 | -0.458385 | -0.220767 | -1.303974 | [illegible] |
| 32.0486111 | -2.076118 | 2.077258 | -4.005801 | -1.275745 | -0.944595 | -2.22782 | 0.916086 | -2.445289 | -0.856309 | -0.503891 | -0.994822 | -0.893341 | -1.323549 | -1.950979 | [illegible] |
| 32.0625 | -1.805395 | 1.483488 | -2.390378 | -1.666061 | -2.033358 | -3.338598 | 2.165128 | -3.096761 | -1.507692 | -1.293004 | -1.806301 | -1.258525 | -2.921757 | -1.73874 | [illegible] |
| 32.0763889 | -2.358558 | 0.100267 | -0.292272 | -0.880105 | -2.325326 | -2.851963 | -0.22895 | -3.590782 | -2.889423 | -2.857001 | -2.412483 | -1.137728 | -4.210901 | -1.667161 | [illegible] |
| 32.0902778 | -1.211959 | -0.144287 | 0.88714 | -0.580721 | -2.046114 | -2.745558 | -1.096779 | -3.956119 | -2.225997 | -2.632342 | -2.399016 | -0.792216 | -4.506052 | -1.15485 | [illegible] |
| 32.1041667 | -0.409285 | -0.261854 | 0.405562 | -0.742493 | -1.334544 | -2.017772 | -0.270229 | -3.682145 | -1.554442 | -1.492906 | -0.824157 | -0.919833 | -3.922076 | -0.453365 | [illegible] |
| 32.1180556 | -0.441935 | -0.3104 | -0.310622 | -0.545156 | -0.83377 | -0.664398 | -0.186155 | -2.606112 | -0.804257 | -0.567887 | 0.021959 | -0.871205 | -2.161934 | 0.322557 | [illegible] |
| 32.1319444 | -0.27328 | -0.330051 | -0.37892 | -0.317741 | -0.442281 | 0.276068 | -0.508565 | -1.99077 | -0.249113 | -0.046036 | 0.481221 | -0.330671 | -0.540216 | 0.944102 | [illegible] |
| 32.1458333 | 0.083797 | -0.288733 | -0.199998 | -0.196489 | 0.210096 | 0.79682 | -0.638532 | -1.137186 | 0.105996 | 0.27205 | 0.682015 | -0.19047 | 0.331648 | 1.326815 | [illegible] |
| 32.1597222 | 0.482226 | -0.150705 | -0.10521 | -0.006304 | 0.613752 | 1.08185 | -0.203913 | 0.020295 | 0.499031 | 0.71792 | 0.643939 | -0.523503 | 0.802156 | 1.20504 | [illegible] |
| 32.1736111 | 0.987131 | 0.007192 | -0.188569 | 0.157848 | 0.877904 | 1.271082 | 0.376615 | 1.212307 | 0.885965 | 1.077575 | 0.365538 | -1.089709 | 1.155621 | 0.736979 | [illegible] |
| 32.1875 | 1.817625 | 0.057365 | -0.298763 | 0.676472 | 1.56073 | 1.131372 | 0.821387 | 2.13918 | 1.174901 | 1.229512 | 0.476187 | -1.692165 | 1.238787 | 0.296084 | [illegible] |
| 32.2013889 | 2.230887 | -0.040518 | -0.206835 | 0.448873 | 2.162855 | 0.830274 | 0.863807 | 2.644519 | 1.200456 | 1.499422 | 0.827262 | -1.753798 | 1.098394 | 0.06452 | [illegible] |
| 32.2152778 | 1.301306 | -0.18718 | 0.027943 | 0.219281 | 1.335387 | 0.388294 | 0.451481 | 2.264564 | 0.94005 | 1.134997 | 1.157817 | -1.173956 | 0.858902 | 0.033209 | [illegible] |
| 32.2291667 | 0.624246 | -0.242544 | 0.416692 | 0.060815 | 0.572086 | 0.06493 | 0.136909 | 1.392335 | 0.745084 | 0.628268 | 1.384325 | -0.610339 | 0.418817 | 0.181404 | [illegible] |
| 32.2430556 | -0.293837 | -0.101525 | 0.896297 | -0.113576 | 0.060999 | 0.205795 | -0.021568 | 1.094438 | 0.765738 | 0.376894 | 0.666397 | -0.196245 | 0.307967 | 0.355994 | [illegible] |
| 32.2569444 | -0.699609 | 0.130157 | 1.34508 | 0.022314 | 0.975206 | 0.69349 | -0.576489 | 1.557707 | 0.675285 | 0.26396 | -0.42605 | 0.069979 | 1.107817 | 0.442001 | [illegible] |
| 32.2708333 | -0.418965 | 0.230305 | 1.399934 | 0.494876 | 1.552029 | 1.073853 | -0.668876 | 2.338626 | 0.4992 | 0.567732 | -0.821555 | 0.193848 | 2.221581 | 0.319919 | [illegible] |
| 32.2847222 | 0.187161 | 0.270092 | 1.136961 | 0.322838 | 2.230615 | 0.891083 | -0.553881 | 2.266675 | 0.382236 | 1.357463 | -0.307108 | 0.225529 | 2.820025 | 0.126188 | [illegible] |
| 32.2986111 | 0.987722 | 0.440096 | 0.803395 | 0.094392 | 2.577989 | 0.610512 | -0.141135 | 1.024418 | 0.480959 | 1.549335 | 1.140118 | 0.265756 | 2.559206 | 0.168649 | [illegible] |
| 32.3125 | 1.811062 | 0.603736 | 0.544086 | -0.017268 | 1.976722 | 0.541871 | 0.602415 | -0.543294 | 1.130238 | 1.013073 | 1.694254 | 0.293163 | 1.591557 | 0.45 | [illegible] |
| 32.3263889 | 1.255882 | 0.661342 | 0.596841 | -0.097532 | 1.038993 | 0.483851 | 1.234961 | -1.017538 | 1.865644 | 0.114599 | 1.007393 | 0.290724 | 0.582887 | 0.672189 | [illegible] |
| 32.3402778 | 0.564599 | 0.706454 | 0.976943 | -0.118881 | -0.35121 | 0.591706 | 0.944597 | -0.403918 | 1.122361 | -0.210293 | 0.235245 | 0.527872 | 0.254215 | 0.7313 | [illegible] |
| 32.3541667 | 0.128058 | 0.764041 | 1.751044 | 0.062101 | -0.875173 | 1.023936 | 0.519124 | 0.642061 | 1.173612 | 0.769276 | 0.202653 | 0.977266 | 1.141156 | 0.792274 | [illegible] |
| 32.3680556 | -0.115679 | 0.863829 | 0.98665 | 0.306004 | 0.183321 | 1.48881 | 0.434735 | 2.028582 | 1.44242 | 1.981138 | 1.198042 | 1.404683 | 2.390335 | 0.755822 | [illegible] |
| 32.3819444 | 0.354297 | 0.97624 | 0.387702 | 0.32815 | 1.293654 | 1.468066 | 0.365516 | 2.888187 | 0.915425 | 2.539245 | 1.836253 | 1.641705 | 3.157506 | 0.726654 | [illegible] |
| 32.3958333 | 1.383775 | 0.974493 | 0.095523 | 0.008367 | 1.726073 | 1.250292 | 0.323654 | 2.2777 | 1.226048 | 1.555562 | 2.108285 | 1.723905 | 3.099497 | 0.865218 | [illegible] |
| 32.4097222 | 2.365885 | 0.919654 | 0.22059 | -0.307378 | 1.394725 | 1.045075 | 0.606565 | 1.590748 | 1.768316 | 0.640998 | 1.797666 | 1.575417 | 2.570888 | 1.039646 | [illegible] |
| 32.4236111 | 2.711398 | 0.773711 | 0.809312 | 0.300175 | 0.702636 | 1.089317 | 0.784338 | 2.236954 | 0.160703 | -0.504687 | 1.260548 | 1.289749 | 2.150692 | 1.291039 | [illegible] |
| 32.4375 | 2.104064 | 0.472376 | 1.673098 | 0.639196 | 0.37964 | 1.283191 | 0.803283 | 3.378725 | 0.928439 | -0.312285 | 1.193051 | 1.11144 | 1.773216 | 1.66384 | [illegible] |
| 32.4513889 | 1.441152 | 0.235615 | 1.752012 | 0.273941 | -0.128871 | 1.675032 | 0.675227 | 2.502436 | 2.027679 | 0.485666 | 1.314979 | 0.987691 | 1.332072 | 1.944275 | [illegible] |
| 32.4652778 | 1.215357 | 0.124637 | 0.521728 | -0.197033 | 0.489885 | 1.463243 | 0.400921 | 1.812032 | 1.486411 | 0.909642 | 1.480369 | 1.014565 | 1.132653 | 1.996172 | [illegible] |
| 32.4791667 | 1.540134 | 0.520688 | -0.67966 | -0.381599 | 1.049566 | 1.291901 | 0.36858 | 2.247232 | 0.75331 | 1.020038 | 1.552439 | 1.307352 | 1.646773 | 1.856791 | [illegible] |
| 32.4930556 | 2.346972 | 1.795255 | -0.925077 | -0.184609 | 2.918302 | 1.738276 | 0.587122 | 1.890878 | 1.1525 | 1.231621 | 2.164254 | 1.790625 | 2.404243 | 1.850737 | [illegible] |
| 32.5069444 | 2.654224 | 1.387429 | 0.608634 | 0.097729 | 3.135359 | 1.727975 | 0.98442 | 1.582044 | 1.620531 | 1.590789 | 2.666232 | 2.115481 | 2.303436 | 2.177289 | [illegible] |
| 32.5208333 | 2.376583 | 0.592235 | 1.443284 | 0.367267 | 2.092938 | 1.14956 | 0.956152 | 1.01699 | 0.463936 | 1.793778 | 2.665629 | 2.088835 | 1.056006 | 2.143584 | [illegible] |
| 32.5347222 | 2.263566 | 0.154379 | 1.643444 | 1.242131 | 0.371728 | 1.285124 | -0.172905 | 0.481625 | 0.900583 | 1.365913 | 2.714342 | 2.287756 | 1.320335 | 1.990558 | [illegible] |
| 32.5486111 | 2.77545 | -0.14157 | 2.639164 | 1.905313 | -0.913778 | 1.946687 | -2.74551 | 0.976362 | 0.167942 | 1.48587 | 3.536514 | 1.814974 | 5.805415 | 2.169933 | [illegible] |
| 32.5625 | 3.533358 | -0.265089 | 2.404475 | 1.146468 | 0.458035 | 2.08454 | -3.5786 | 0.171434 | -0.08428 | 2.581155 | 2.930658 | 1.954187 | 9.883758 | 1.619428 | [illegible] |
| 32.5763889 | 2.119749 | -0.272692 | 0.310907 | 0.218104 | 3.36061 | 1.133479 | -1.28722 | -0.206829 | 0.15022 | 2.811257 | 2.368706 | 1.234683 | 6.422284 | 0.812031 | [illegible] |
| 32.5902778 | 1.502137 | 0.134936 | -0.73581 | 0.269053 | 4.398615 | 0.223259 | 0.255796 | -0.346513 | -0.011906 | 3.7098 | 3.361402 | 1.038426 | 3.636707 | -0.357484 | [illegible] |
| 32.6041667 | 0.925816 | 0.37297 | -1.029312 | 0.523322 | 2.545959 | -0.522607 | 0.580414 | 0.165931 | 0.38294 | 1.93858 | 2.976367 | 1.183895 | 3.565569 | -0.674214 | [illegible] |
| 32.6180556 | 0.27962 | 0.135097 | -0.903734 | 0.226888 | 0.770737 | -0.947151 | 0.399809 | 0.660676 | 1.327381 | 2.378132 | 1.651712 | 1.333599 | 2.995596 | 0.075822 | [illegible] |
| 32.6319444 | 0.998146 | 0.659746 | -0.575236 | -0.146607 | -0.614099 | -1.006181 | 0.185486 | 0.396064 | 1.370748 | 4.417337 | 0.213291 | 1.361724 | 1.227429 | 1.891004 | [illegible] |
| 32.6458333 | 0.797235 | 0.606834 | -0.841767 | -0.816825 | 1.177334 | 0.303365 | -0.04202 | -0.954367 | 0.719657 | 3.126252 | -1.843925 | 1.280376 | -1.793425 | 3.456399 | [illegible] |
| 32.6597222 | 0.028681 | -0.345335 | -2.066231 | 0.345398 | 0.457869 | 1.376225 | -0.273984 | -1.150426 | 0.221604 | 1.763287 | -1.939728 | 0.841762 | -1.614167 | 2.395334 | [illegible] |
| 32.6736111 | 0.934258 | 0.00458 | -1.983729 | 1.235053 | -2.328538 | 1.109494 | -0.412547 | -1.270224 | -0.469146 | -0.734373 | -2.197769 | 0.053431 | -0.945125 | -0.728321 | [illegible] |
| 32.6875 | 0.549264 | 0.053868 | -0.490745 | 0.246413 | -3.300691 | 0.452657 | -0.278095 | -0.336812 | -0.484862 | -2.531672 | -2.670035 | -0.315102 | -0.689659 | -3.09573 | [illegible] |
| 32.7013889 | 1.920692 | -0.024262 | 0.480863 | -0.552477 | -0.261287 | 0.09114 | 0.052656 | 0.132158 | -0.405598 | -1.285816 | -1.689176 | 0.013808 | 0.491315 | -3.425885 | [illegible] |
| 32.7152778 | 1.453917 | 0.091548 | 0.997831 | 0.117339 | 2.665065 | 0.137197 | -0.201075 | -0.354797 | 0.775195 | 1.133907 | -0.494706 | 0.435906 | 2.497647 | -2.98616 | [illegible] |
| 32.7291667 | -0.671516 | 0.34109 | 1.139149 | 0.642172 | 2.867524 | 0.099574 | -0.106204 | -0.640305 | 0.465164 | 3.582082 | 1.469465 | 0.412533 | 1.3896 | -0.46141 | [illegible] |
| 32.7430556 | -0.93189 | 0.118389 | 1.649708 | 0.740411 | 1.392349 | -0.216193 | 0.43664 | 0.20372 | 0.171064 | 1.69468 | 3.483066 | 0.342166 | 2.545063 | 2.383459 | [illegible] |
| 32.7569444 | -1.114036 | 0.34857 | 1.786054 | 0.620581 | 0.981446 | -0.438129 | 1.119408 | 0.850733 | -0.053697 | -0.382244 | 3.264898 | -0.051274 | 3.201241 | 2.221381 | [illegible] |
| 32.7708333 | 1.295862 | 0.165851 | 0.839432 | -0.327548 | 1.968697 | -0.635477 | 0.936542 | 0.805414 | -0.384837 | -0.076488 | 2.176552 | -0.142051 | 3.257345 | 0.442475 | [illegible] |
| 32.7847222 | 2.822544 | -0.260805 | -0.797633 | -0.480236 | -0.223814 | -0.364164 | 0.390873 | -0.438712 | -0.906002 | 0.224918 | 0.502051 | -0.632227 | -1.828358 | -1.608545 | [illegible] |
| 32.7986111 | 2.030759 | -0.669079 | -2.165802 | 0.013718 | -0.515483 | -0.030752 | 0.273383 | -0.576971 | -0.612285 | -0.554497 | -1.262862 | -1.250104 | -4.159059 | -2.727255 | [illegible] |
| 32.8125 | 0.172702 | -0.764722 | -2.218871 | -0.19149 | -0.787106 | 0.150524 | 0.208332 | -0.097934 | 0.234422 | -1.83629 | -1.761048 | -1.641305 | -2.662597 | -3.234115 | [illegible] |
| 32.8263889 | -1.056225 | -0.258581 | -1.220793 | -0.143228 | 0.502242 | 0.252969 | -0.012333 | 0.17096 | 1.047745 | -0.918669 | -1.065588 | -2.080693 | -0.058315 | -3.326033 | [illegible] |
| [illegible] | [illegible] | [illegible] | [illegible] | [illegible] | [illegible] | [illegible] | [illegible] | [illegible] | [illegible] | [illegible] | [illegible] | [illegible] | [illegible] | [illegible] | [illegible] |

| | | | | | | | | | | | | | | | | | | | | | | | | | | | | |
|---|---|---|---|---|---|---|---|---|---|---|---|---|---|---|---|---|---|---|---|---|---|---|---|---|---|---|---|---|
| 35.5069444 | 2.258244 | 0.990458 | 0.968188 | -0.102711 | 2.525782 | 2.108626 | 1.128785 | 4.55032 | 3.42734 | 2.08891 | 3.301075 | 4.646646 | 4.826302 | 2.704498 | 1.947951 | 0.07515 | 1.620746 | 0.281428 | 0.18569 | 0.135672 | 0.840749 | 0.14832 | 1.650998 | 1.236606 | | 35.5 | 4.826302 | 1.243146 |
| 35.5208333 | 2.370347 | 2.462579 | 0.998307 | 0.157684 | 2.565206 | 2.125879 | 2.21841 | 4.048084 | 2.436487 | 2.347947 | 3.789969 | 3.407031 | 4.929279 | 1.779473 | 1.63084 | -0.004656 | 1.301349 | 0.248271 | 0.033811 | 0.092129 | 0.950794 | 0.103886 | 1.578587 | 1.520867 | | 35.514 | 4.929279 | 1.373818 |
| 35.5347222 | 3.892529 | 2.298155 | 1.532327 | -0.25768 | 2.381925 | 2.483615 | 0.989184 | 2.39486 | 2.647419 | 2.0527 | 4.979937 | 2.503375 | 6.43394 | 2.011737 | 0.995212 | -0.406336 | 1.189179 | 0.098686 | -0.786912 | -0.132203 | 1.3222 | -0.064861 | 1.675021 | 3.719466 | | 35.528 | 6.43394 | 2.540737 |
| 35.5486111 | 8.701207 | 1.25114 | 0.395934 | -1.045381 | 2.468325 | 3.319581 | -2.65161 | 1.390218 | 5.819176 | 0.978619 | 6.783326 | 2.741797 | 11.443132 | 3.670602 | 0.472514 | -1.10476 | 1.638659 | -0.361129 | -2.058202 | -0.403029 | 1.789085 | -0.486905 | 1.75185 | 6.275689 | | 35.542 | 11.443132 | 4.508514 |
| 35.5625 | 7.958773 | 0.911075 | 1.539073 | -1.260386 | 2.030525 | 3.513925 | -4.034068 | 0.97897 | 6.535424 | 1.524664 | 6.970685 | 2.21749 | 13.046573 | 5.456069 | -0.076272 | -1.666757 | 1.08627 | -1.399653 | -2.229701 | -0.687455 | 1.120751 | -1.37132 | 1.276743 | 3.75143 | | 35.556 | 13.046573 | 3.932885 |
| 35.5763889 | 2.848503 | -0.034758 | 2.2337 | -0.55641 | 2.073952 | 1.973835 | -3.330206 | 1.486296 | 4.607119 | 3.250184 | 6.486802 | 1.27256 | 7.646757 | 4.761298 | -0.651287 | -1.367039 | -0.213271 | -1.548673 | -1.583374 | -1.133283 | 0.264337 | -1.406918 | 0.825311 | 0.770047 | | 35.569 | 7.646757 | 0.836547 |
| 35.5902778 | 0.994357 | -0.75186 | 3.16314 | -0.087588 | 1.414867 | 0.04241 | -1.626371 | 0.529963 | 2.178525 | 4.458789 | 0.612453 | 1.017114 | 5.566485 | 3.723715 | -0.454078 | -0.794841 | -0.397706 | -0.932406 | -0.871769 | -0.78493 | 0.43819 | -0.847874 | 0.75273 | -0.093067 | | 35.583 | 5.566485 | -0.03677 |
| 35.6041667 | 1.510998 | -0.404412 | 3.303758 | 0.221427 | 0.932469 | -0.323849 | -0.485274 | -0.008216 | 0.489833 | 1.033126 | 1.379994 | 2.87607 | 5.363144 | 2.602397 | -1.389379 | -0.379623 | -0.047194 | -0.619359 | -0.395706 | -0.372425 | 1.237915 | -0.392613 | 1.195119 | -0.099993 | | 35.597 | 5.363144 | -0.041303 |
| 35.6180556 | 1.178131 | 0.499824 | 1.590311 | 0.52318 | 1.184897 | -0.016545 | 0.376814 | -0.232072 | -0.810352 | -0.600928 | 2.182655 | 2.798104 | 3.437134 | 1.302035 | -0.390008 | -0.198692 | 0.278469 | -0.550631 | -0.185595 | -0.167151 | 2.373344 | -0.186253 | 1.759462 | 0.08131 | | 35.611 | 3.437134 | -0.009535 |
| 35.6319444 | -0.750923 | 0.050897 | 0.273068 | 0.440438 | 1.411453 | 0.630197 | 0.539167 | -0.14598 | -0.481187 | -0.774326 | -0.535799 | 0.237558 | -0.026798 | 1.393835 | 0.85424 | -0.09977 | 0.489609 | -0.65256 | -0.075766 | -0.076467 | 2.318392 | -0.082095 | 1.481491 | -0.259657 | | 35.625 | -0.026798 | -0.499682 |
| 35.6458333 | -1.021052 | -0.33514 | -0.878416 | 0.000845 | -0.088131 | 0.594319 | 0.554368 | -0.597923 | -0.16244 | 1.468201 | -1.871547 | -1.661312 | -2.445635 | 0.03583 | 0.048357 | -0.084481 | 0.521376 | -0.674227 | -0.07123 | -0.070607 | 1.118705 | -0.069346 | 0.066015 | -0.946412 | | 35.639 | -2.445635 | -1.133227 |
| 35.6597222 | -1.958801 | -0.066171 | -1.886258 | 0.179715 | 0.260692 | 0.734729 | -0.242636 | -0.554424 | 0.831377 | 2.108101 | -1.133873 | -1.004074 | -2.0801 | -2.697625 | 0.276778 | -0.104119 | 0.444847 | -0.514359 | -0.108403 | -0.089843 | -1.24488 | -0.104623 | -0.385487 | -0.92054 | | 35.653 | -2.0801 | -0.959919 |
| 35.6736111 | -3.699313 | -1.286897 | -2.917646 | 0.470103 | -0.034002 | 0.444646 | -0.127773 | 1.534898 | 2.089909 | 1.616311 | 0.929742 | 0.010922 | 1.316809 | -3.89981 | 0.363564 | -0.060114 | 0.249993 | -0.254996 | -0.057404 | -0.046823 | -3.624905 | -0.067247 | 0.115536 | -1.121017 | | 35.667 | 1.316809 | -0.529395 |
| 35.6875 | -6.230034 | -1.338899 | -2.911169 | 0.532788 | -0.915771 | -0.073752 | -0.166661 | 0.908681 | 1.258645 | -0.348538 | 2.883713 | 0.955503 | 3.840005 | -3.154814 | 0.262452 | 0.018661 | -0.121563 | -0.117579 | 0.043343 | 0.029069 | -1.803705 | 0.030605 | -1.757528 | -2.134902 | | 35.681 | 3.840005 | -1.928892 |
| 35.7013889 | -3.038637 | -2.536308 | -1.130206 | 0.34173 | 0.709796 | -0.564421 | -0.392055 | 0.146804 | -0.290775 | -2.116236 | 3.047348 | 1.74806 | 1.836974 | -3.448368 | 1.093735 | 0.044979 | -0.183952 | -0.025265 | 0.083243 | 0.056065 | 0.346187 | 0.080911 | 0.302425 | -1.906585 | | 35.694 | 1.836974 | -2.223441 |
| 35.7152778 | -1.332795 | -1.075523 | 1.678959 | 0.303699 | 1.918094 | -1.325593 | -0.290954 | -0.189286 | -1.188883 | -3.492408 | 2.9118 | 0.115131 | -2.870607 | -2.902397 | 0.274209 | 0.038396 | -0.197025 | 0.171614 | 0.077948 | 0.056893 | 2.000633 | 0.074249 | -0.080346 | -0.329812 | | 35.708 | -2.870607 | -0.894603 |
| 35.7291667 | 1.549914 | -0.836405 | 3.444772 | 0.265265 | 1.298258 | -0.677155 | -0.878568 | -0.918875 | -0.61214 | -2.766436 | 2.732432 | -2.406034 | -7.710964 | -0.123176 | 0.401023 | 0.07136 | -0.172228 | 0.427324 | 0.098154 | 0.083219 | 2.160756 | 0.08098 | -0.650498 | 0.178008 | | 35.722 | -7.710964 | 0.018433 |
| 35.7430556 | 2.04338 | -0.393176 | 1.992357 | -0.712896 | 0.438266 | -0.818141 | -0.595442 | -1.796331 | -0.231016 | -1.71097 | -1.923513 | -2.027779 | -4.21768 | -0.212625 | 1.089416 | 0.137235 | -0.141059 | 0.747046 | 0.151754 | 0.139492 | 1.809647 | 0.12999 | -0.579312 | 1.062588 | | 35.736 | -4.21768 | 1.022957 |
| 35.7569444 | 1.715822 | -0.082771 | 2.001215 | -0.033759 | 0.42071 | -0.732046 | 1.012629 | -0.299624 | 0.768189 | 1.130579 | -2.909931 | -0.796463 | 1.854515 | -2.445187 | 1.387199 | 0.168182 | -0.128624 | 1.01083 | 0.186054 | 0.167321 | 0.655776 | 0.16764 | -1.593936 | 1.115027 | | 35.75 | 1.854515 | 0.22334 |
| 35.7708333 | 0.356229 | -0.475302 | 0.485429 | 0.188917 | 0.029386 | 0.450981 | 0.592534 | 0.940571 | 0.951215 | 3.326235 | -3.666583 | 0.019133 | 6.454473 | -3.319592 | 0.736736 | 0.145027 | -0.101605 | 0.959913 | 0.160585 | 0.144808 | -0.471395 | 0.149515 | -1.676266 | -0.247115 | | 35.764 | 6.454473 | -0.413383 |
| 35.7847222 | 0.276181 | 0.876771 | -0.702291 | -0.12448 | 0.694289 | 0.946242 | 0.642045 | 1.410276 | 0.974354 | 6.054391 | -1.867784 | -0.577486 | 6.892743 | -2.027505 | 0.410645 | 0.113098 | -0.194785 | 0.655446 | 0.113778 | 0.110173 | -1.574727 | 0.109149 | -1.9227 | -0.739404 | | 35.778 | 6.892743 | -0.588445 |
| 35.7986111 | -0.187964 | 2.26508 | -1.613621 | 0.104067 | 1.034546 | 0.743562 | 1.539305 | 0.731202 | 0.378037 | 1.358687 | 0.066631 | -0.191419 | 6.308315 | -0.460373 | 0.045147 | 0.089864 | -0.140974 | 0.382318 | 0.075892 | 0.08483 | -2.25856 | 0.073429 | -1.13738 | -1.148902 | | 35.792 | 6.308315 | -1.142516 |
| 35.8125 | 0.083349 | 1.189445 | -1.343359 | 0.037623 | -0.214956 | 0.572219 | 3.219188 | 0.298461 | -0.060652 | 2.289873 | 0.660579 | 0.5734 | 4.890927 | 0.514722 | 0.795413 | 0.06371 | -0.241959 | 0.199555 | 0.052019 | 0.060786 | -2.159576 | 0.055452 | 0.506459 | -2.114724 | | 35.806 | 4.890927 | -2.191648 |
| 35.8263889 | 0.793128 | 0.287293 | -2.628067 | -0.135067 | -0.333886 | 0.409608 | 3.449039 | 0.238392 | -0.771118 | 2.065041 | 0.70702 | 0.842729 | 1.26284 | -0.329094 | 1.703441 | 0.053307 | -0.600539 | 0.092242 | 0.055436 | 0.052204 | -0.83903 | 0.062957 | -0.896208 | -1.668269 | | 35.819 | 1.26284 | -1.527648 |
| 35.8402778 | -2.871549 | -0.491626 | -3.118114 | -0.158816 | 0.504306 | -0.148805 | 2.711647 | 0.264634 | -1.55658 | -0.004357 | 0.073209 | 1.400517 | -2.890283 | -1.962731 | 2.353921 | 0.059939 | -1.520447 | 0.022392 | 0.076793 | 0.064115 | 0.185611 | 0.077146 | -1.542225 | -0.69953 | | 35.833 | -2.890283 | -0.477286 |
| 35.8541667 | -0.450416 | -0.57204 | -1.837912 | -0.219594 | -0.144995 | -0.923347 | 1.338267 | -0.104143 | -1.67156 | -0.608354 | -1.367134 | 0.629182 | -1.616636 | -4.093609 | 2.163703 | 0.080333 | -1.289465 | -0.027644 | 0.109981 | 0.096405 | 1.085276 | 0.108112 | -0.340346 | -0.027872 | | 35.847 | -1.616636 | 0.087574 |
| 35.8680556 | 3.355775 | -0.001253 | -0.872656 | 0.030989 | -0.351602 | -1.697491 | 0.241806 | -0.364933 | -0.535448 | 0.957678 | -3.102862 | 0.491113 | -0.093768 | -5.506312 | 1.83345 | 0.112562 | -0.755598 | -0.0975 | 0.129675 | 0.114242 | 0.949174 | 0.13759 | 0.469824 | 0.023719 | | 35.861 | -0.093768 | 0.329964 |
| 35.8819444 | 5.69437 | -0.462587 | -0.665364 | 0.182359 | -1.2106 | -2.552788 | -0.511108 | -0.507624 | 0.570298 | 0.753319 | -4.705851 | -0.140394 | -1.522275 | -6.204101 | 1.42735 | 0.098296 | -1.075159 | -0.197231 | 0.099118 | 0.079702 | -0.290507 | 0.108305 | 0.998506 | -0.602104 | | 35.875 | -1.522275 | -0.068795 |
| 35.8958333 | 3.442432 | -1.267533 | -0.912055 | 0.555706 | -1.690624 | -1.944438 | -2.184626 | -0.052585 | 1.587241 | -1.044079 | -4.815057 | -1.788976 | -2.60122 | -5.676192 | 0.903793 | 0.085452 | -1.431683 | -0.277112 | 0.08214 | 0.065669 | -1.870508 | 0.075172 | 1.098342 | -1.211702 | | 35.889 | -2.60122 | -0.846419 |
| 35.9097222 | 0.36597 | -1.518212 | -1.433569 | 0.01436 | -2.767987 | -1.74079 | -1.806533 | 0.349435 | 0.721629 | -2.081456 | -2.71296 | -0.483454 | -2.214183 | -4.869448 | 0.438457 | 0.134043 | -1.751407 | -0.339543 | 0.139544 | 0.129175 | -3.171443 | 0.123239 | 1.129697 | -1.445903 | | 35.903 | -2.214183 | -1.056187 |
| 35.9236111 | -1.01825 | -0.950119 | -2.218035 | -0.051181 | -3.314244 | -2.92622 | -0.563234 | 0.527601 | 0.069726 | -3.372822 | -0.873297 | -1.641031 | -1.965803 | -4.132441 | 0.445662 | 0.175888 | -2.167141 | -0.358856 | 0.198961 | 0.183986 | -3.504067 | 0.17985 | 1.240141 | -1.528681 | | 35.917 | -1.965803 | -1.376154 |
| 35.9375 | -1.143245 | 0.320022 | -2.822036 | 0.096676 | -1.781922 | -3.031026 | 0.565558 | 0.16488 | -1.122395 | -1.70921 | -0.691193 | -2.152492 | -4.371786 | -3.362361 | -0.49621 | 0.184649 | -2.506661 | -0.422659 | 0.212111 | 0.191039 | -3.242663 | 0.198378 | 1.675753 | -1.402263 | | 35.931 | -4.371786 | -1.284743 |
| 35.9513889 | -1.865507 | 0.305385 | -1.868423 | 0.057281 | -0.33398 | -1.926148 | 0.616412 | -0.575468 | -1.359322 | -0.945037 | -1.512052 | -1.899069 | -4.752978 | -2.317233 | -0.415152 | 0.19991 | -2.83293 | -0.414761 | 0.219839 | 0.183882 | -2.568514 | 0.214108 | 2.23916 | -0.936645 | | 35.944 | -4.752978 | -0.955413 |
| 35.9652778 | -4.415525 | -0.75195 | -0.374681 | -0.013794 | -0.838631 | -1.08499 | -0.096946 | -1.032748 | -1.74578 | 0.077641 | -2.074856 | -1.792013 | -3.684453 | -0.941394 | -0.954983 | 0.203847 | -3.206573 | -0.429045 | 0.230328 | 0.184037 | -1.20977 | 0.231137 | 1.627464 | -0.627967 | | 35.958 | -3.684453 | -0.735628 |
| 35.9791667 | -6.84875 | -4.213804 | 0.607223 | -0.336518 | -0.55101 | -0.775727 | -1.109651 | -1.780028 | -1.459091 | -0.774234 | -3.212768 | -2.234879 | -3.730793 | 0.31875 | -1.394298 | 0.205216 | -3.323879 | -0.456138 | 0.242209 | 0.20089 | -0.497587 | 0.243662 | 0.125206 | -0.29738 | | 35.972 | -3.730793 | -0.419387 |
| 35.9930556 | -5.203176 | -6.520686 | 1.020056 | -0.650738 | -0.912762 | -0.649127 | 0.782921 | -2.691565 | -2.293717 | -0.635385 | -4.28584 | -1.902968 | -5.306993 | 1.229821 | -1.690058 | 0.129442 | -3.469345 | -0.493686 | 0.249883 | 0.212349 | -0.065855 | 0.244748 | -0.953534 | -0.432996 | | 35.986 | -5.306993 | -0.626209 |
| 36.0069444 | -2.673653 | -3.224098 | 1.135935 | -0.590981 | -1.555729 | -1.544705 | 1.68262 | -3.640281 | -1.459526 | 0.711299 | -3.814415 | -1.812701 | -6.139585 | 1.77042 | -2.005569 | -0.022102 | -3.190312 | -0.499745 | 0.243576 | 0.202365 | 0.009452 | 0.23297 | -1.008385 | -0.646837 | | 36 | -6.139585 | -0.940312 |
| 36.0208333 | 1.918001 | -0.636031 | 0.600607 | 0.03655 | -2.67399 | -2.919414 | 2.366374 | -2.833641 | -1.056576 | 0.557896 | -2.852269 | -2.328855 | -5.079582 | 1.799238 | -1.711845 | -0.090562 | -2.581229 | -0.342063 | 0.370534 | 0.268062 | -0.167255 | 0.340598 | -1.014089 | -0.786764 | | 36.014 | -5.079582 | -1.134698 |
| 36.0347222 | 6.044896 | 0.277603 | -0.793043 | 1.212222 | -3.343037 | -3.97514 | 2.572495 | -2.064806 | -0.057405 | 0.653897 | -2.804948 | -5.261507 | -4.688929 | 0.273022 | -1.172035 | 0.049026 | -2.0869 | 0.006319 | 0.690584 | 0.333077 | -0.633901 | 0.614533 | -1.186618 | -1.46421 | | 36.028 | -4.688929 | -2.208781 |
| 36.0486111 | 4.346631 | -1.725253 | -2.42677 | 2.033482 | -3.214857 | -4.616022 | 1.679488 | -0.43417 | -0.098548 | -1.651605 | -3.575874 | -6.411415 | -8.463999 | -2.484457 | -0.690979 | 0.482708 | -2.153208 | 0.225885 | 1.40587 | 0.19231 | -1.544949 | 0.600712 | -2.469237 | -3.344501 | | 36.042 | -8.463999 | -4.202068 |
| 36.0625 | 1.339201 | -3.015353 | -2.595935 | 1.659679 | -4.009518 | -4.080717 | 1.3718 | -2.70149 | -1.690745 | -2.157117 | -3.62743 | -5.741199 | -9.758442 | -1.614821 | -0.383838 | 0.686604 | -2.318071 | 0.068377 | 1.859285 | 0.190733 | -1.80127 | 0.266253 | -2.361724 | -3.575318 | | 36.056 | -9.758442 | -3.933377 |
| 36.0763889 | -1.22246 | -4.53518 | -1.040559 | 0.184582 | -5.093181 | -2.921439 | 0.368194 | -4.145097 | -3.978917 | -4.680685 | -4.925101 | -5.665355 | -10.326193 | 0.131232 | -0.389261 | 0.507875 | -2.018349 | 0.398107 | 1.276851 | 0.457994 | -0.721993 | 0.422757 | -2.375239 | -1.647096 | | 36.069 | -10.326193 | -1.542322 |
| 36.0902778 | -3.564097 | -3.590453 | -1.486625 | -0.378272 | -4.44721 | -1.286377 | -0.018098 | -6.301149 | -6.641843 | -4.058735 | -5.408063 | -3.508179 | -10.778841 | 1.186595 | -0.59185 | 0.169895 | -1.371513 | 0.715355 | 0.727122 | 0.626191 | 0.216156 | 0.574443 | -2.423982 | -0.495825 | | 36.083 | -10.778841 | -0.669519 |
| 36.1041667 | -4.362352 | -2.843429 | -2.166582 | -0.232796 | -4.064418 | -0.687807 | -0.083822 | -5.18201 | -4.734672 | -3.350559 | -4.786332 | -1.916842 | -8.266878 | 1.398999 | -0.374669 | 0.47875 | -0.244104 | 0.643598 | 0.559761 | 0.532616 | 0.496861 | 0.525043 | -1.416944 | -0.10694 | | 36.097 | -8.266878 | -0.341208 |
| 36.1180556 | -2.862641 | -1.485487 | -1.468148 | 0.08599 | -2.693047 | 0.258366 | 0.242263 | -3.719917 | -2.701337 | -1.875345 | -2.544032 | -1.848622 | -4.842443 | 1.469485 | -0.443483 | 0.439024 | 0.288468 | 0.536393 | 0.406621 | 0.363783 | 0.40108 | 0.37762 | -0.485346 | 0.191008 | | 36.111 | -4.842443 | 0.039081 |
| 36.1319444 | -1.098792 | -1.353059 | -0.488369 | 0.151763 | -1.599068 | 0.540283 | 0.397372 | -3.330938 | -1.989089 | -0.819837 | -0.792536 | -2.116267 | -2.497679 | 1.43206 | -0.509459 | 0.194666 | 0.320439 | 0.525448 | 0.248315 | 0.20831 | 0.24843 | 0.226877 | 0.174093 | 0.413911 | | 36.125 | -2.497679 | 0.361356 |
| 36.1458333 | 0.55647 | -1.246542 | 0.006809 | 0.050615 | -1.002441 | 0.467943 | 0.327013 | -2.133736 | -1.624444 | -0.021151 | 0.51206 | -1.506208 | -0.764224 | 1.062612 | -0.176406 | 0.012737 | 0.335181 | 0.46277 | 0.148745 | 0.107687 | 0.18707 | 0.123932 | -0.232346 | 0.460202 | | 36.139 | -0.764224 | 0.468858 |
| 36.1597222 | 1.881523 | 0.157999 | 0.265851 | 0.034505 | -0.230645 | 0.373827 | 0.289012 | -1.613103 | -1.169725 | 0.659833 | 1.850214 | -0.274881 | 0.890206 | 0.094822 | -0.00901 | -0.046052 | 0.706426 | 0.258714 | 0.116294 | 0.077618 | 0.186688 | 0.089318 | -0.531757 | 0.097683 | | 36.153 | 0.890206 | 0.120903 |
| 36.1736111 | 2.6334 | 2.843568 | 0.543445 | 0.152182 | 0.800069 | -0.125108 | 0.33484 | -1.022726 | -0.469121 | 1.302327 | 2.815797 | 1.446651 | 1.986201 | -1.922694 | -0.789761 | -0.057528 | 0.986229 | 0.002464 | 0.033765 | 0.00901 | 0.065066 | 0.0172 | -0.150623 | -0.536864 | | 36.167 | 1.986201 | -0.515023 |
| 36.1875 | 2.793345 | 4.484748 | 0.963494 | 0.204052 | 2.5054 | -0.916247 | 0.608542 | -0.531509 | 0.224196 | 2.136386 | 2.812162 | 2.851405 | 2.313151 | -3.190105 | -1.068586 | -0.035134 | 0.824502 | -0.198207 | -0.077128 | -0.081665 | -0.062124 | -0.078427 | 0.103092 | -0.67022 | | 36.181 | 2.313151 | -0.654198 |
| 36.2013889 | 1.870142 | 3.214335 | 1.108426 | 0.275388 | 3.823429 | -1.648698 | 0.234971 | -0.645236 | 0.602008 | 2.777648 | 1.597945 | 4.107607 | 1.948438 | -3.222462 | -0.911465 | 0.051971 | 0.506937 | -0.26457 | -0.090704 | -0.078754 | 0.126594 | -0.077202 | -0.362619 | -0.229337 | | 36.194 | 1.948438 | -0.084689 |
| 36.2152778 | 0.524025 | 1.199731 | 0.794467 | 0.337935 | 4.045936 | -1.103383 | -0.204421 | -0.421339 | 0.592444 | 2.707122 | 0.351572 | 3.674882 | 0.837332 | -1.949845 | -0.991893 | 0.092883 | 0.551671 | -0.270789 | -0.043863 | -0.022605 | 0.721198 | -0.021224 | -0.858659 | 0.465331 | | 36.208 | 0.837332 | 0.584125 |
| 36.2291667 | -0.58298 | -1.435423 | 0.372612 | 0.197025 | 2.936817 | -0.66573 | -0.142285 | 0.421387 | 0.53823 | 1.948682 | -0.415048 | 1.391575 | -0.486896 | -0.684359 | -0.756579 | 0.117202 | 0.953219 | -0.320994 | -0.029563 | -0.015943 | 1.178214 | -0.015885 | -0.611077 | 0.926123 | | 36.222 | -0.486896 | 0.963379 |
| 36.2430556 | 0.141841 | -1.972194 | 0.168114 | 0.290029 | 0.936914 | -0.208822 | 0.739688 | 0.618786 | 0.523202 | 0.958479 | -0.179033 | -0.802296 | -0.68733 | 0.113748 | -0.357187 | 0.093524 | 1.518353 | -0.37316 | -0.029781 | -0.032106 | 1.318477 | -0.03496 | -0.4173 | 0.974796 | | 36.236 | -0.68733 | 1.054926 |
| 36.2569444 | 1.871177 | -2.638904 | 0.035485 | -0.218287 | -0.895713 | 0.114122 | 0.928492 | 0.369539 | 0.592402 | -0.015537 | 0.222271 | -2.495916 | -0.092505 | 0.092488 | -0.018809 | 0.058442 | 1.717333 | -0.322047 | -0.00586 | -0.016765 | 1.169254 | -0.018542 | -0.251163 | 0.665003 | | 36.25 | -0.092505 | 0.757539 |
| 36.2708333 | 2.713319 | -1.589646 | -0.098249 | -0.586872 | -0.574965 | 0.229541 | -1.700406 | 0.275436 | 0.720839 | -0.256644 | 0.731476 | -1.672664 | 0.124283 | -0.620625 | -0.350329 | -0.023199 | 1.146536 | -0.225607 | -0.001591 | -0.010558 | 0.962458 | -0.01048 | -0.03281 | 0.206753 | | 36.264 | 0.124283 | 0.316761 |
| 36.2847222 | 2.857906 | 0.940588 | -0.089995 | -0.695995 | -0.098095 | 0.984871 | -2.09977 | 0.196307 | 1.090023 | 0.216446 | 1.070482 | -0.094472 | 0.659587 | -1.301804 | -0.693934 | -0.101658 | 0.378963 | -0.18863 | -0.035199 | -0.047598 | 0.780118 | -0.054554 | 0.148896 | -0.23512 | | 36.278 | 0.659587 | -0.150972 |
| 36.2986111 | 2.408978 | 3.278771 | 0.03365 | -0.64113 | 0.810407 | 2.483586 | -0.431931 | 0.288883 | 1.864277 | 1.217776 | 1.260293 | 1.479672 | 1.529674 | -0.824598 | -2.83219 | -0.115256 | -0.065568 | -0.297034 | -0.060673 | -0.075462 | 0.60974 | -0.09662 | 0.263686 | -0.45039 | | 36.292 | 1.529674 | -0.428213 |
| 36.3125 | 0.972908 | 4.148722 | 0.213359 | -0.098006 | 1.593885 | 1.543443 | 0.149496 | 0.6326 | 2.578868 | 1.801535 | 0.971196 | 2.934038 | 2.114003 | 0.657782 | -2.876305 | -0.1228 | 0.167726 | -0.425892 | -0.093286 | -0.076957 | 0.620157 | -0.103805 | 0.456651 | -0.285473 | | 36.306 | 2.114003 | -0.395026 |
| 36.3263889 | -0.008368 | 2.443941 | 0.480047 | 0.520114 | 2.037215 | 2.067879 | 0.913971 | 1.102297 | 3.323703 | 1.952959 | 1.215047 | 3.988028 | 2.538129 | 2.046324 | -0.260233 | -0.121012 | 1.153331 | -0.457533 | -0.136464 | -0.076786 | 0.769188 | -0.10102 | 0.714339 | 0.084669 | | 36.319 | 2.538129 | -0.137608 |
| 36.3402778 | -0.430219 | 1.244944 | 0.881631 | 0.677428 | 2.539687 | 2.794502 | 1.685195 | 1.829035 | 3.517368 | 1.759263 | 1.634106 | 4.402496 | 3.500055 | 2.322229 | 1.698772 | -0.133905 | 1.679452 | -0.41832 | -0.135675 | -0.085451 | 0.852552 | -0.102926 | 0.905149 | 0.45006 | | 36.333 | 3.500055 | 0.256409 |
| 36.3541667 | -0.736855 | 0.853062 | 0.925207 | 0.600842 | 2.654108 | 3.253136 | 2.147818 | 1.670027 | 2.987181 | 1.447719 | 1.943418 | 3.687758 | 4.343904 | 1.891748 | 1.950775 | -0.170984 | 1.312722 | -0.360121 | -0.095079 | -0.08172 | 0.770509 | -0.09458 | 0.924477 | 0.75579 | | 36.347 | 4.343904 | 0.674902 |
| 36.3680556 | -0.777741 | 0.846008 | 0.765127 | 0.253349 | 2.425137 | 2.57848 | 2.151338 | 1.493186 | 2.267682 | 1.511255 | 1.877459 | 2.549835 | 4.281429 | 1.386264 | 1.726135 | -0.178997 | 1.374427 | -0.287321 | -0.06807 | -0.073515 | 0.60721 | -0.081768 | 0.866583 | 0.909073 | | 36.361 | 4.281429 | 0.98749 |
| 36.3819444 | -0.300354 | 0.666242 | 0.671175 | 0.077493 | 1.907386 | 1.947004 | 1.357805 | 1.526655 | 2.057037 | 1.861293 | 1.756795 | 1.239001 | 3.443537 | 0.682204 | 0.63772 | -0.169887 | 0.80761 | -0.159548 | -0.08162 | -0.068604 | 0.459893 | -0.078744 | 0.814759 | 0.896271 | | 36.375 | 3.443537 | 1.085863 |
| 36.3958333 | 0.703601 | 0.547854 | 0.568983 | 0.013006 | 1.775233 | 1.569111 | 0.855364 | 1.547357 | 2.265558 | 2.514625 | 1.645484 | 0.459095 | 2.385643 | 0.166128 | 1.270066 | -0.083587 | 0.548287 | 0.074551 | -0.065706 | -0.03029 | 0.415522 | -0.036811 | 0.924263 | 0.818534 | | 36.389 | 2.385643 | 1.063332 |
| 36.4097222 | 1.361255 | 0.392001 | 0.451449 | 0.244006 | 2.25473 | 1.320487 | 0.55902 | 1.441624 | 2.134481 | 2.983015 | 1.9057 | 1.26782 | 1.763148 | 0.576643 | 1.844146 | 0.063855 | 1.003297 | 0.379869 | 0.027306 | 0.043765 | 0.557826 | 0.052497 | 1.07595 | 0.958271 | | 36.403 | 1.763148 | 1.177617 |
| 36.4236111 | 0.847666 | 0.316208 | 0.3754 | 0.559124 | 2.098285 | 1.478162 | 0.03653 | 1.139022 | 2.304438 | 2.45745 | 2.518886 | 2.847606 | 1.770558 | 1.640778 | 1.645555 | 0.142672 | 1.974297 | 0.67675 | 0.090813 | 0.08699 | 0.769404 | 0.105596 | 1.20633 | 1.263355 | | 36.417 | 1.770558 | 1.416767 |
| 36.4375 | -0.327052 | 0.259303 | 0.417894 | 0.515109 | 1.524915 | 2.098129 | -0.325617 | 1.091318 | 2.385018 | 1.873301 | 3.195117 | 4.986876 | 2.067103 | 2.624547 | 1.405641 | 0.114786 | 2.682298 | 0.890703 | 0.031524 | 0.048709 | 0.954827 | 0.065782 | 1.254018 | 1.58082 | | 36.431 | 2.067103 | 1.599723 |
| 36.4513889 | -1.130023 | -0.131279 | 0.223237 | 0.007074 | 1.276283 | 2.648029 | -0.046413 | 1.29153 | 2.486026 | 1.549439 | 2.935776 | 5.413006 | 2.255932 | 3.273311 | 1.309655 | -0.042166 | 2.723149 | 0.875616 | -0.151537 | -0.109786 | 0.941347 | -0.128143 | 1.331007 | 1.578824 | | 36.444 | 2.255932 | 1.568825 |
| 36.4652778 | -1.122654 | -0.574732 | -0.072399 | -0.68335 | 1.330041 | 2.4787 | 0.195335 | 1.636063 | 2.285204 | 1.608582 | 1.951284 | 4.600138 | 1.881638 | 3.639167 | -0.350249 | -0.298938 | 1.989572 | 0.620629 | -0.391338 | -0.355332 | 0.756892 | -0.415247 | 1.531929 | 1.300563 | | 36.458 | 1.881638 | 1.355857 |
| 36.4791667 | -0.735734 | -0.532422 | -0.157228 | -0.509542 | 1.521589 | 1.457011 | 0.650589 | 1.87127 | 2.353686 | 2.066713 | 1.236902 | 3.502338 | 1.406071 | 4.170631 | 0.176124 | -0.391412 | 2.330311 | 0.399069 | -0.453181 | -0.409408 | 0.462612 | -0.442492 | 1.831677 | 1.254936 | | 36.472 | 1.406071 | 1.342617 |
| 36.4930556 | -0.472707 | 0.025246 | -0.005909 | 0.354778 | 2.236776 | 0.650318 | 1.77022 | 2.456544 | 2.543983 | 2.314477 | 2.484997 | 2.470719 | 1.804598 | 5.047111 | 2.319666 | -0.319906 | 1.964997 | 0.347415 | -0.275188 | -0.261579 | 0.364386 | -0.267584 | 2.149209 | 1.351951 | | 36.486 | 1.804598 | 1.51748 |
| 36.5069444 | 0.328929 | 0.640627 | -0.058762 | 0.903861 | 3.125655 | 0.380568 | 1.742616 | 3.734277 | 2.860577 | 2.356501 | 4.416978 | 1.785745 | 2.981176 | 5.838975 | 4.22895 | -0.285539 | 0.515324 | 0.357965 | -0.166659 | -0.176621 | 0.32477 | -0.182321 | 2.198777 | 1.319286 | | 36.5 | 2.981176 | 1.496446 |
| 36.5208333 | 1.622331 | 0.99699 | -0.182514 | 0.941989 | 3.972344 | 0.956863 | 0.654669 | 4.853097 | 3.828789 | 1.89584 | 6.098673 | 2.724487 | 4.788264 | 5.442857 | 4.308383 | -0.367077 | 0.880463 | 0.307064 | -0.295963 | -0.232026 | 0.516116 | -0.209876 | 1.976051 | 1.490135 | | 36.514 | 4.788264 | 1.40754 |
| 36.5347222 | 4.0884 | 0.576377 | -0.287519 | 0.766184 | 4.531397 | 2.643875 | -0.438406 | 4.207623 | 5.772462 | 2.112177 | 6.854574 | 3.895695 | 8.807119 | 5.223964 | 2.958742 | -0.701071 | 1.102213 | 0.254749 | -1.01378 | -0.441326 | 1.092717 | -0.28308 | 1.831622 | 3.65084 | | 36.528 | 8.807119 | 1.96525 |
| 36.5486111 | 8.443255 | -0.337787 | -0.430776 | 0.846842 | 5.106546 | 4.125652 | -1.450611 | 3.128081 | 6.327023 | 2.791175 | 7.649341 | 3.01344 | 16.01241 | 5.413056 | 1.028005 | -1.39212 | 1.190708 | -0.018057 | -2.075734 | -0.712733 | 1.565605 | -0.643466 | 1.768171 | 5.9328 | | 36.542 | 16.01241 | 3.030816 |
| 36.5625 | 8.828716 | -0.784152 | 0.259063 | 0.97597 | 3.886994 | 4.232139 | -2.320912 | 1.18629 | 4.438485 | 2.314293 | 4.868664 | 2.600838 | 14.794838 | 5.536805 | -0.805354 | -2.114639 | -0.208931 | -1.372017 | -2.567908 | -1.354354 | 1.091134 | -1.747544 | 0.78753 | 2.453954 | | 36.556 | 14.794838 | 0.865361 |
| 36.5763889 | 2.113127 | -0.606181 | 1.988201 | 1.133326 | 2.264482 | 2.154744 | -2.600195 | -0.164183 | 3.098991 | 1.932335 | 2.816691 | 0.136243 | 5.814441 | 2.632411 | -0.748991 | -1.880709 | -1.450021 | -1.706479 | -2.082724 | -1.964095 | 0.192153 | -1.982874 | 0.023254 | -1.015261 | | 36.569 | 5.814441 | -0.955441 |
| 36.5902778 | -2.695634 | -0.456763 | 2.615767 | 1.238757 | 1.195408 | 0.179365 | -1.518429 | 0.303292 | 3.532919 | 2.248321 | 2.234296 | -0.539184 | 2.587168 | 1.231398 | -0.363342 | -1.081603 | -0.999557 | -0.866698 | -1.224324 | -1.123522 | 0.469874 | -1.218802 | 0.357663 | -0.853187 | | 36.583 | 2.587168 | -0.9555 |
| 36.6041667 | -3.020518 | -0.154488 | 3.589935 | 1.671296 | 0.821881 | 0.398063 | 0.754346 | 3.111064 | 1.096284 | 2.021781 | 1.67017 | 1.073307 | 5.409573 | 1.844987 | 0.659602 | -0.413085 | -0.100662 | -0.286996 | -0.462447 | -0.427803 | 1.34706 | -0.483918 | 1.102268 | -0.151716 | | 36.597 | 5.409573 | -0.262856 |
| 36.6180556 | 0.411032 | 0.100781 | 2.539137 | 1.685588 | 1.707514 | 1.717832 | 1.935737 | 3.224251 | 1.115338 | 1.479946 | 2.69558 | 3.859148 | 8.579923 | 2.029281 | 1.400805 | -0.115419 | 0.670483 | -0.051391 | -0.106464 | -0.108547 | 2.804745 | -0.123823 | 1.500296 | 0.576796 | | 36.611 | 8.579923 | 0.420022 |
| 36.6319444 | 5.126213 | 0.84377 | 2.069077 | 1.780461 | 2.420668 | 2.107223 | 3.014455 | 2.300482 | 0.2835 | 0.078847 | 2.045221 | 3.263225 | 8.458663 | 1.52858 | 1.05043 | 0.01763 | 0.848699 | 0.015987 | 0.056344 | 0.042678 | 3.699027 | 0.055611 | 0.803481 | 0.636358 | | 36.625 | 8.458663 | 0.373508 |
| 36.6458333 | 6.974912 | 2.229238 | 1.612584 | 1.47441 | 1.746331 | 0.53307 | 3.264367 | 1.1955 | -0.857573 | -3.015952 | -0.573213 | -0.052774 | 3.424766 | 0.308854 | -2.050367 | 0.021064 | 0.93657 | -0.010184 | 0.081881 | 0.074923 | 1.370686 | 0.087512 | 0.059637 | -0.531024 | | 36.639 | 3.424766 | -0.814407 |
| 36.6597222 | 4.137007 | 2.238276 | 1.069549 | 1.57099 | 0.322862 | 0.27305 | -2.683275 | -2.056321 | -2.799326 | -1.116834 | -3.333692 | -2.400058 | -4.487224 | -1.723137 | -3.523362 | 0.037615 | 1.267531 | 0.0042 | 0.106209 | 0.113201 | -1.061218 | 0.105502 | -0.011476 | -1.203784 | | 36.653 | -4.487224 | -1.45005 |
| 36.6736111 | 0.475828 | 2.880368 | 0.907418 | 1.908631 | -0.427954 | 0.552809 | -3.038673 | -1.832865 | -0.822591 | 1.0557 | -3.129785 | -1.073547 | -4.533556 | -3.608281 | -1.250435 | 0.19342 | 0.200213 | 0.143947 | 0.260502 | 0.277245 | -1.670007 | 0.248352 | 0.435719 | -1.121953 | | 36.667 | -4.533556 | -1.102536 |
| 36.6875 | -2.774233 | 0.192415 | 0.734873 | 1.682007 | 0.510464 | -0.382443 | -2.96966 | 1.840053 | -0.022204 | 1.580132 | -0.587267 | -1.101181 | -0.499633 | -4.246714 | 0.720331 | 0.353075 | -0.417011 | 0.218113 | 0.371165 | 0.392237 | -0.979664 | 0.349014 | 1.266422 | -0.65216 | | 36.681 | -0.499633 | -0.417948 |
| 36.7013889 | -1.062483 | -2.412556 | 0.444358 | 0.582671 | 0.219784 | -0.003913 | -0.27766 | 2.527 | 1.358057 | -2.654627 | 0.542028 | -0.187707 | 5.683373 | -2.210468 | 1.269836 | 0.329838 | -0.202231 | 0.146442 | 0.332369 | 0.340653 | 0.949364 | 0.308389 | 1.326215 | 0.138506 | | 36.694 | 5.683373 | 0.21387 |
| 36.7152778 | -0.378799 | -2.338066 | 0.535943 | 0.294417 | 1.233089 | 0.665008 | 3.023824 | 0.309626 | 0.423316 | -1.870029 | 0.936205 | 2.125273 | 8.273493 | 1.524725 | 1.853767 | 0.283269 | -0.00593 | 0.067224 | 0.283541 | 0.263319 | 2.186347 | 0.261747 | 0.972495 | 0.626989 | | 36.708 | 8.273493 | 0.651554 |
| 36.7291667 | -4.735009 | -0.494513 | 0.433745 | -0.039124 | 2.559559 | 0.88818 | 2.441149 | 0.047648 | -0.248209 | 0.086284 | 3.703516 | 1.721482 | 5.996124 | 4.402421 | 1.930307 | 0.272142 | -0.684798 | 0.016195 | 0.287807 | 0.241918 | 2.582778 | 0.261771 | 1.350557 | 0.541483 | | 36.722 | 5.996124 | 0.667264 |
| 36.7430556 | -1.141869 | 1.816162 | 0.149427 | -0.482758 | 1.187755 | 0.599178 | 1.488051 | -0.238241 | -0.664169 | -0.641064 | 2.305525 | 0.719541 | 3.682553 | 5.224017 | 0.700446 | 0.257085 | -0.279049 | -0.029989 | 0.297187 | 0.232543 | 2.015749 | 0.263036 | -0.208067 | 0.322898 | | 36.736 | 3.682553 | 0.075196 |
| 36.7569444 | 4.050419 | 1.767544 | -0.43986 | -0.014154 | -0.305302 | 0.668859 | 0.219715 | 0.240976 | -0.994078 | 0.376307 | 1.216673 | -1.580302 | 3.588489 | 3.188644 | -0.682076 | 0.236117 | 0.28562 | -0.054335 | 0.289494 | 0.203132 | 0.566644 | 0.256265 | -2.701386 | -0.214211 | | 36.75 | 3.588489 | -0.62673 |
| 36.7708333 | 4.452696 | 1.490031 | -1.155073 | -0.288953 | -0.239838 | 0.373279 | 0.872033 | 0.500129 | -0.688752 | -0.250078 | -0.553999 | -1.046852 | 3.613195 | -4.201263 | -0.816038 | 0.226704 | 0.613917 | -0.059741 | 0.274954 | 0.193131 | -1.373122 | 0.252702 | -3.257683 | -0.458815 | | 36.764 | 3.613195 | -0.929594 |
| 36.7847222 | 1.452507 | 1.511711 | -0.254717 | -0.459082 | -0.49416 | -0.129204 | 1.137673 | 0.228488 | -0.077071 | -0.287401 | -1.554171 | -0.24862 | 2.475748 | -5.725864 | -0.292583 | 0.239103 | 0.607014 | -0.035177 | 0.262479 | 0.207294 | -1.706747 | 0.250632 | -2.354726 | -0.441451 | | 36.778 | 2.475748 | -0.75601 |
| 36.7986111 | 2.366449 | 1.785308 | 0.682363 | -0.174582 | -0.435976 | -0.247605 | 0.565426 | -0.120823 | 0.555028 | -0.223666 | -2.310896 | 0.861683 | 2.252968 | -4.479742 | -0.055069 | 0.210573 | -0.244846 | -0.060864 | 0.248132 | 0.241246 | -1.233032 | 0.240227 | -1.275608 | -0.903549 | | 36.792 | 2.252968 | -0.321108 |
| 36.8125 | 1.458832 | 2.13631 | 0.386539 | 0.099261 | -0.754912 | -0.073495 | 0.986306 | -0.518424 | 0.667867 | -0.344385 | -3.195535 | -0.020696 | -0.142511 | -3.653945 | 0.122276 | 0.191474 | -0.377434 | -0.098478 | 0.238254 | 0.264474 | -0.566146 | 0.226357 | -0.261965 | -0.587467 | | 36.806 | -0.142511 | -0.072468 |
| 36.8263889 | 1.007299 | 1.937229 | -0.576527 | 0.128061 | -1.141572 | -0.206216 | 3.118073 | -0.832173 | 0.593872 | 0.408138 | -3.098754 | -1.508436 | -0.688433 | -5.16929 | 0.586453 | 0.19869 | -0.673818 | -0.110506 | 0.247033 | 0.272563 | 0.626573 | 0.228869 | 0.112347 | -0.052683 | | 36.819 | -0.688433 | 0.270869 |
| 36.8402778 | -4.349314 | 1.055922 | -0.919159 | 0.249735 | -1.456023 | -0.537653 | 4.787723 | -0.955485 | 0.257778 | 0.563271 | -1.554728 | -1.76398 | 0.928441 | -4.76618 | 0.172746 | 0.214059 | -0.944033 | -0.094567 | 0.264809 | 0.273619 | 2.870863 | 0.247361 | -0.059397 | 0.232685 | | 36.833 | 0.928441 | 0.304825 |
| 36.8541667 | -5.229839 | 0.32636 | -1.241572 | 0.312836 | -0.873573 | -0.864741 | 4.398651 | -0.625662 | -0.253998 | -0.73363 | -0.166667 | -0.490776 | 3.043175 | -1.283073 | -0.572191 | 0.210185 | -1.513013 | -0.126969 | 0.244484 | 0.229991 | 3.573667 | 0.243423 | -0.497598 | 0.187208 | | 36.847 | 3.043175 | 0.057142 |
| 36.8680556 | -0.440632 | 1.139496 | -2.146769 | 0.595714 | -0.575928 | -1.679324 | 3.893783 | -0.189157 | -0.495579 | -1.317006 | -0.187778 | -1.14392 | 1.489607 | 1.573376 | -1.336613 | 0.181675 | -1.207509 | -0.170575 | 0.178646 | 0.151933 | 1.620059 | 0.202244 | -0.860392 | -0.121843 | | 36.861 | 1.489607 | -0.330673 |
| 36.8819444 | -2.786511 | -0.966854 | -2.607778 | -0.112362 | -0.68135 | -1.745427 | 6.208353 | 0.724354 | -0.983345 | 0.205408 | -1.85687 | -1.355799 | -4.14774 | 3.00101 | -1.918791 | 0.165643 | -1.127621 | -0.210473 | 0.137473 | 0.100884 | -0.300652 | 0.159192 | -1.23857 | -0.594431 | | 36.875 | -4.14774 | -0.89057 |
| 36.8958333 | -1.760048 | -2.08888 | -2.161715 | -0.62202 | -0.381766 | -1.653288 | 2.890212 | 1.183451 | -1.073551 | -1.20909 | -4.495578 | -1.352864 | -6.995267 | 2.74147 | -2.0514 | 0.178761 | -1.903157 | -0.226376 | 0.156996 | 0.105267 | -2.549091 | 0.15452 | -1.40754 | -1.193725 | | 36.889 | -6.995267 | -1.449344 |
| 36.9097222 | 0.83982 | -1.320964 | -1.501966 | -0.528627 | -1.358509 | -0.281876 | -0.874502 | 0.539254 | -1.148762 | -0.81874 | -5.361686 | -0.660206 | -4.957502 | -0.772191 | -1.802834 | 0.201519 | -2.848112 | -0.213261 | 0.187131 | 0.129573 | -4.655975 | 0.176635 | -1.297066 | -1.872268 | | 36.903 | -4.957502 | -1.873241 |
| 36.9236111 | 1.62568 | -0.775925 | -1.002641 | -0.595548 | -2.009056 | 0.489399 | -5.456175 | -0.753033 | 0.372002 | 0.004807 | -7.082277 | -1.035124 | -3.093373 | -3.388517 | -1.267375 | 0.177087 | -3.67215 | -0.2335 | 0.163007 | 0.127184 | -4.751424 | 0.171686 | -1.19327 | -2.549572 | | 36.917 | -3.093373 | -2.491982 |
| 36.9375 | 2.855335 | 0.892354 | -0.85127 | -0.411696 | -0.665485 | -0.145614 | -5.292403 | -1.333367 | 1.218207 | 0.617877 | -7.147328 | -1.694253 | -3.961119 | -4.771344 | -0.939026 | 0.104227 | -4.423695 | -0.294713 | 0.086026 | 0.087267 | -3.979492 | 0.118939 | -1.109064 | -2.667491 | | 36.931 | -3.961119 | -2.623018 |
| 36.9513889 | 6.496119 | 1.680384 | -1.109631 | -1.014996 | 1.176664 | -1.49133 | -3.059963 | -1.685812 | 1.262646 | 0.337903 | -5.162642 | -5.401754 | -2.481964 | -5.555902 | -0.750154 | 0.034972 | -4.839013 | -0.345845 | 0.008625 | 0.028452 | -2.558236 | 0.048893 | -1.013033 | -2.526079 | | 36.944 | -2.481964 | -2.467855 |
| 36.9652778 | 5.516269 | 0.825313 | -1.639719 | -0.807937 | 1.285009 | -3.513747 | -2.404382 | -1.991583 | 0.711049 | -1.270724 | -2.47064 | -2.683136 | -0.982585 | -5.531765 | -0.665562 | -0.021952 | -4.756182 | -0.395838 | -0.046139 | -0.032821 | -1.426046 | -0.022758 | -0.758654 | -2.287749 | | 36.958 | -0.982585 | -2.273836 |
| 36.9791667 | 3.617943 | -0.389175 | -2.191567 | -0.855181 | 2.6989 | -5.333248 | -1.08831 | -2.49322 | -0.249276 | -2.112719 | 0.222598 | -0.119519 | -0.559784 | -4.739969 | -0.807919 | -0.080197 | -4.229211 | -0.471965 | -0.094655 | -0.096412 | -0.903268 | -0.090214 | -0.630667 | -2.066269 | | 36.972 | -0.559784 | -2.250727 |
| 36.9930556 | 1.590108 | -1.2686 | -1.738425 | -0.246989 | 1.385692 | -5.057131 | 0.250599 | -2.789769 | -1.476504 | -2.50998 | 1.209852 | 1.143876 | -2.468835 | -3.370778 | -1.129671 | -0.147523 | -3.466485 | -0.553611 | -0.126561 | -0.136663 | -0.700662 | -0.13166 | -0.651398 | -1.984465 | | 36.986 | -2.468835 | -2.03788 |
| 37.0069444 | -0.413366 | -3.054396 | -1.480972 | 1.245681 | -1.126346 | -5.108521 | 0.489952 | -3.305817 | -2.466027 | -3.30609 | 0.940351 | 2.607121 | -5.481475 | -2.177358 | -1.494625 | -0.159049 | -2.363296 | -0.551859 | -0.110706 | -0.120785 | -0.65402 | -0.11607 | -0.554411 | -1.674548 | | 37 | -5.481475 | -1.722173 |
| 37.0208333 | -4.097036 | -4.621411 | -2.155147 | 1.814787 | -2.400259 | -4.298788 | 1.078511 | -2.494616 | -2.670262 | -3.356977 | 0.555423 | 2.095298 | -6.97662 | -1.642806 | -1.450635 | -0.014912 | -1.328802 | -0.335964 | 0.079621 | 0.036975 | -0.631247 | 0.066269 | -0.300478 | -1.392816 | | 37.014 | -6.97662 | -1.554356 |
| 37.0347222 | -9.3127 | -5.025397 | -3.12282 | 1.233517 | -3.79546 | -2.384051 | 2.366319 | -1.755962 | -4.331387 | -3.09346 | 0.011527 | -0.054495 | -8.524974 | -2.691371 | -1.16842 | 0.305136 | -0.793242 | -0.104715 | 0.415749 | 0.266844 | -0.918193 | 0.389146 | -0.443354 | -2.031605 | | 37.028 | -8.524974 | -3.060826 |
| 37.0486111 | -10.030538 | -0.130319 | -3.914855 | -0.005009 | -5.014766 | -1.372999 | 1.370484 | -2.173348 | -6.562519 | -2.631632 | -3.001557 | -2.668723 | -10.037674 | -4.63153 | -1.048902 | 0.705236 | -1.258336 | -0.536568 | 0.907658 | 0.16282 | -1.696095 | 0.313922 | -1.470373 | -4.770127 | | 37.042 | -10.037674 | -5.906149 |
| 37.0625 | -8.73329 | -0.545138 | -3.623712 | -1.095587 | -5.52504 | -0.961358 | -1.26422 | -2.829973 | -7.166093 | -2.889401 | -5.97476 | -5.107819 | -11.145586 | -3.6109 | -0.867779 | 0.959364 | -1.692786 | -1.199797 | 1.322488 | 0.0999 | -1.997273 | -0.058113 | -1.534444 | -4.962767 | | 37.056 | -11.145586 | -4.454178 |
| 37.0763889 | -6.460167 | -1.49275 | -2.432335 | -1.024446 | -6.656264 | -2.326149 | -4.044133 | -3.428096 | -5.064442 | -4.299313 | -7.484797 | -6.203459 | -11.714143 | -2.063167 | -0.820286 | 0.391601 | -1.081313 | -0.69173 | 0.465416 | 0.054812 | -1.750054 | -0.210989 | -1.151628 | -2.775796 | | 37.069 | -11.714143 | -2.249232 |
| 37.0902778 | -5.772643 | -2.445344 | -0.713065 | -2.549267 | -6.938138 | -2.721798 | -4.647682 | -3.84266 | -5.420749 | -3.68713 | -6.598011 | -4.578658 | -11.31421 | -1.091837 | -0.813238 | 0.182277 | -0.516136 | -0.35537 | 0.078084 | 0.058662 | -1.001143 | -0.039335 | -0.889476 | -1.057833 | | 37.083 | -11.31421 | -1.053586 |
| 37.1041667 | -5.741535 | -2.891063 | -0.39918 | -2.730566 | -5.675131 | -2.190804 | -4.469854 | -3.300821 | -4.808595 | -3.542519 | -3.974976 | -3.830788 | -9.51377 | -0.577911 | -1.429606 | 0.203668 | -0.386017 | -0.25258 | 0.108936 | 0.131096 | -0.23091 | 0.077997 | -0.56061 | -0.484961 | | 37.097 | -9.51377 | -0.57101 |
| 37.1180556 | -3.241312 | -1.612758 | -0.555834 | -1.963781 | -3.198568 | -1.184246 | -1.565003 | -2.553248 | -3.657867 | -3.445691 | -1.44359 | -3.838156 | -6.158228 | -0.067497 | -1.91788 | 0.182087 | -0.199101 | -0.142232 | 0.130505 | 0.149445 | 0.137582 | 0.11712 | -0.535489 | -0.096985 | | 37.111 | -6.158228 | -0.216189 |
| 37.1319444 | -0.280424 | -0.416229 | -0.401977 | -0.679258 | -1.972374 | -0.898399 | 0.010076 | -2.327018 | -2.470704 | -2.613087 | -0.046456 | -2.818664 | -2.916397 | 0.548414 | -1.5794 | 0.119354 | 0.243433 | -0.047766 | 0.114979 | 0.142545 | 0.172311 | 0.121139 | -0.546407 | 0.178091 | | 37.125 | -2.916397 | 0.069504 |
| 37.1458333 | 2.22817 | 0.482448 | -0.104296 | -0.026703 | -1.20926 | -0.226427 | 0.709723 | -2.04555 | -1.333025 | -1.099007 | 1.010673 | -1.527684 | -0.385701 | 1.094521 | -1.002504 | 0.026895 | 0.718573 | -0.016154 | 0.089558 | 0.105426 | 0.092119 | 0.086759 | -0.273734 | 0.319473 | | 37.139 | -0.385701 | 0.243766 |
| 37.1597222 | 4.279176 | 2.027194 | 0.264275 | 0.448422 | -0.276137 | 0.732857 | 2.178735 | -1.613129 | -0.861994 | -0.018742 | 2.01158 | 0.251355 | 2.015182 | 1.606271 | -0.468383 | -0.045713 | 1.146724 | -0.018583 | 0.074723 | 0.063048 | 0.067838 | 0.055995 | 0.018629 | 0.448469 | | 37.153 | 2.015182 | 0.414114 |
| 37.1736111 | 6.202062 | 2.930678 | 0.465127 | 0.749774 | 0.59956 | 1.488987 | 3.163827 | -0.644302 | -0.502255 | 0.781779 | 2.505513 | 1.033976 | 4.105218 | 1.675834 | 0.069269 | -0.089717 | 1.28963 | -0.022571 | 0.025361 | 0.009495 | 0.129275 | 0.022348 | 0.263193 | 0.579636 | | 37.167 | 4.105218 | 0.560272 |
| 37.1875 | 6.450953 | 1.279338 | 0.364024 | 0.781492 | 1.643925 | 2.006464 | 2.632487 | 0.218323 | -0.329344 | 1.744708 | 1.891464 | 1.746336 | 4.581709 | 1.045978 | 0.507235 | -0.085755 | 1.087033 | -0.046134 | -0.041011 | -0.047295 | 0.230264 | -0.020491 | 0.176251 | 0.543626 | | 37.181 | 4.581709 | 0.545859 |
| 37.2013889 | 3.691049 | -0.261995 | 0.339967 | 0.810653 | 3.703032 | 1.635983 | 1.810979 | 0.491016 | 0.594897 | 2.589772 | 0.452655 | 2.671755 | 3.103633 | 0.239238 | 0.903119 | -0.029664 | 0.697772 | -0.066001 | -0.040634 | -0.048376 | 0.458367 | -0.032038 | -0.170597 | 0.335132 | | 37.194 | 3.103633 | 0.379787 |
| 37.2152778 | -0.382792 | 0.426795 | 0.461559 | 1.150047 | 4.900284 | 0.774413 | 0.709361 | 0.666033 | 2.319588 | 1.988762 | -0.940341 | 3.240623 | 1.007483 | -0.250812 | 1.288389 | 0.022066 | 0.56837 | -0.036834 | 0.005352 | 0.002239 | 0.848769 | -0.00415 | -0.390433 | 0.286602 | | 37.208 | 1.007483 | 0.375331 |
| 37.2291667 | -3.993394 | 0.415329 | 0.742057 | 1.066728 | 3.850946 | 0.22312 | 0.150277 | 0.739194 | 3.513249 | 0.328905 | -1.875258 | 1.072896 | -0.792677 | -0.14755 | 1.489243 | 0.028062 | 1.01354 | 0.017406 | 0.053023 | 0.049187 | 1.231598 | 0.037968 | -0.293493 | 0.574273 | | 37.222 | -0.792677 | 0.662956 |
| 37.2430556 | -4.427144 | 0.677017 | 0.760255 | 0.314492 | 2.122294 | 0.262848 | -0.340982 | 0.598745 | 3.411005 | -1.399692 | -1.500862 | -1.335627 | -1.215301 | 0.225645 | 1.270582 | 0.004166 | 1.739004 | 0.044876 | 0.087476 | 0.070497 | 1.516093 | 0.069224 | -0.383013 | 0.887176 | | 37.236 | -1.215301 | 0.918497 |
| 37.2569444 | -2.054513 | -0.099411 | 0.445646 | -0.28704 | 0.25573 | 0.753763 | -0.788126 | 0.195719 | 2.649004 | -1.689948 | -0.768606 | -2.475376 | -0.910188 | 0.409965 | 0.839308 | -0.011903 | 2.129254 | 0.040307 | 0.078567 | 0.066498 | 1.601643 | 0.075565 | -0.621038 | 0.92945 | | 37.25 | -0.910188 | 0.924575 |
| 37.2708333 | 0.105085 | -2.071571 | 0.114238 | -0.202063 | -1.559597 | 1.599421 | -1.018711 | 0.045827 | 1.501354 | -0.532797 | -0.20721 | -1.293821 | -0.361585 | 0.194878 | 0.645306 | -0.024518 | 1.808507 | 0.030804 | 0.075635 | 0.069156 | 1.537109 | 0.081601 | -0.662672 | 0.727211 | | 37.264 | -0.361585 | 0.668122 |
| 37.2847222 | 0.18676 | -2.498063 | 0.02293 | 0.483011 | -1.218378 | 2.731193 | -0.364726 | -0.041844 | 0.937082 | 0.676233 | -0.7639 | 0.59859 | 0.602325 | -0.315711 | 0.551005 | -0.004628 | 1.07723 | 0.055695 | 0.126781 | 0.107065 | 1.380993 | 0.120012 | -0.334834 | 0.458286 | | 37.278 | 0.602325 | 0.401675 |
| 37.2986111 | 1.05703 | -0.601587 | 0.16696 | 0.615457 | 0.699479 | 3.625001 | 0.607567 | 0.230639 | 1.273953 | 1.196418 | -0.895724 | 1.529077 | 1.529043 | -0.785348 | 0.374301 | 0.024414 | 0.516331 | 0.098553 | 0.170318 | 0.1488 | 1.122056 | 0.162693 | 0.023738 | 0.217675 | | 37.292 | 1.529043 | 0.200623 |
| 37.3125 | 2.727311 | 1.003299 | 0.415891 | 0.058264 | 2.348993 | 3.078119 | 1.306456 | 0.434751 | 1.855214 | 1.684111 | 1.522633 | 2.376241 | 2.268584 | -0.833427 | 0.206395 | 0.060546 | 0.238986 | 0.124685 | 0.183476 | 0.178244 | 0.860258 | 0.189006 | 0.302442 | 0.120847 | | 37.306 | 2.268584 | 0.114672 |
| 37.3263889 | 2.630182 | 1.645848 | 0.604558 | 0.131445 | 2.950832 | 2.185206 | 1.114338 | 0.808127 | 1.892191 | 1.877988 | 3.843607 | 3.858519 | 2.490896 | -0.35934 | 0.037021 | 0.077876 | 0.399699 | 0.113171 | 0.164103 | 0.183141 | 0.686905 | 0.186176 | 0.500697 | 0.192047 | | 37.319 | 2.490896 | 0.204781 |
| 37.3402778 | 2.038409 | 0.711064 | 0.690852 | 1.115272 | 2.522464 | 1.596674 | 0.700828 | 1.121297 | 2.085736 | 1.558304 | 4.793407 | 6.101448 | 2.719808 | 0.512519 | -0.167978 | 0.046858 | 0.814092 | 0.043821 | 0.093346 | 0.125603 | 0.554317 | 0.121304 | 0.565766 | 0.3706 | | 37.333 | 2.719808 | 0.335457 |
| 37.3541667 | 1.246224 | 0.511138 | 0.685027 | 2.174833 | 2.052276 | 1.324616 | 0.264691 | 1.287641 | 2.305206 | 1.206173 | 4.297314 | 6.050336 | 3.720792 | 1.535213 | -0.22851 | 0.004188 | 1.20405 | -0.045094 | 0.033675 | 0.051754 | 0.524104 | 0.043802 | 0.555115 | 0.516555 | | 37.347 | 3.720792 | 0.423358 |
| 37.3680556 | 2.493207 | 1.086808 | 0.522645 | 1.912228 | 1.77346 | 1.058378 | -0.017492 | 1.166752 | 2.432384 | 1.264135 | 2.698287 | 4.244015 | 4.887443 | 2.400272 | -0.09003 | 0.02211 | 1.440663 | -0.079457 | 0.039731 | 0.046548 | 0.563044 | 0.035811 | 0.578582 | 0.577817 | | 37.361 | 4.887443 | 0.461035 |
| 37.3819444 | 3.241879 | 1.540269 | 0.249224 | 1.25547 | 1.820584 | 1.548755 | 0.075641 | 0.813611 | 2.15454 | 1.489886 | 0.927863 | 3.349635 | 4.888132 | 2.597197 | 0.243368 | 0.053445 | 1.231104 | -0.048077 | 0.066232 | 0.083158 | 0.587918 | 0.069205 | 0.679159 | 0.569513 | | 37.375 | 4.888132 | 0.459016 |
| 37.3958333 | 1.090799 | 1.964791 | 0.155119 | 0.665315 | 2.110068 | 2.253137 | 0.461759 | 0.669859 | 2.040434 | 1.838905 | 0.643762 | 2.393873 | 3.707386 | 1.901673 | 0.744383 | 0.064721 | 0.906592 | -0.028515 | 0.062404 | 0.085408 | 0.599868 | 0.067783 | 0.752048 | 0.505007 | | 37.389 | 3.707386 | 0.457649 |
| 37.4097222 | -1.259056 | 1.721917 | 0.286181 | 0.519731 | 2.190167 | 2.220565 | 0.845512 | 0.891672 | 1.802982 | 2.057158 | 1.565656 | 1.994691 | 2.275432 | 1.057161 | 1.215278 | 0.033049 | 0.896376 | -0.036493 | 0.036123 | 0.0556 | 0.520856 | 0.040616 | 0.761649 | 0.570234 | | 37.403 | 2.275432 | 0.653059 |
| 37.4236111 | -3.481221 | 0.827813 | 0.544514 | 0.849598 | 1.833139 | 1.54866 | 1.188501 | 1.114666 | 1.556689 | 2.010456 | 3.068208 | 2.787423 | 1.058387 | 0.244531 | 1.722284 | 0.002705 | 1.146124 | -0.033578 | 0.013527 | 0.020312 | 0.439202 | 0.017776 | 0.780191 | 0.766565 | | 37.417 | 1.058387 | 0.949881 |
| 37.4375 | -3.116462 | -0.226523 | 0.781487 | 0.95184 | 1.445972 | 0.477075 | 1.235686 | 1.056847 | 1.58812 | 1.991384 | 4.25189 | 4.088793 | 1.089243 | -0.025892 | 1.857745 | -0.021822 | 1.347701 | -0.037725 | -0.003935 | -0.01363 | 0.362992 | -0.014813 | 0.796003 | 0.919617 | | 37.431 | 1.089243 | 1.119294 |
| 37.4513889 | 0.288343 | -0.379331 | 0.575074 | 1.100273 | 1.420688 | -0.644231 | 0.593571 | 1.127533 | 1.976411 | 2.343663 | 4.522269 | 4.109861 | 2.163768 | 0.658685 | 1.342143 | -0.100465 | 1.458703 | -0.103813 | -0.080134 | -0.079612 | 0.325766 | -0.111519 | 0.952375 | 0.887376 | | 37.444 | 2.163768 | 1.130854 |
| 37.4652778 | 4.306056 | -2.121204 | -0.027396 | 1.00614 | 1.162071 | -1.185422 | -0.374597 | 1.343923 | 2.599876 | 2.598767 | 3.688868 | 3.515916 | 2.668299 | 0.760639 | 1.045977 | -0.207862 | 1.641046 | -0.190463 | -0.206607 | -0.172546 | 0.214988 | -0.232762 | 1.390834 | 0.576978 | | 37.458 | 2.668299 | 0.985255 |
| 37.4791667 | 6.550611 | -1.966906 | -0.340253 | 0.658122 | 1.103072 | -0.467318 | -1.190038 | 1.260564 | 3.144183 | 3.054961 | 2.607247 | 3.616179 | 2.618283 | 0.41839 | 1.248453 | -0.190752 | 2.226525 | -0.172125 | -0.190646 | -0.142762 | 0.273876 | -0.162877 | 2.058081 | -0.319045 | | 37.472 | 2.618283 | 0.700368 |
| 37.4930556 | 6.564719 | 0.07204 | -0.237218 | 0.519773 | 1.577149 | 1.439884 | -0.963132 | 1.545834 | 3.825865 | 3.968058 | 1.922852 | 4.260283 | 3.122034 | -0.790657 | 1.190257 | -0.085595 | 2.792388 | -0.070039 | -0.048392 | -0.029406 | 0.574519 | -0.022892 | 2.797632 | -0.37215 | | 37.486 | 3.122034 | 1.062346 |
| 37.5069444 | 5.26502 | 2.050047 | -0.097965 | 0.700257 | 1.972836 | 3.716773 | 0.532757 | 2.137248 | 3.926038 | 4.773715 | 2.082903 | 4.982293 | 3.975231 | 2.874528 | 1.958672 | 0.002293 | 2.9811 | -0.001267 | 0.042926 | 0.049213 | 0.94113 | 0.063891 | 3.121897 | 1.274004 | | 37.5 | 3.975231 | 1.582373 |
| 37.5208333 | 3.413255 | 2.457556 | -0.172653 | 1.276015 | 2.357202 | 4.578941 | 3.781398 | 2.609508 | 3.612215 | 5.47755 | 3.085634 | 4.898108 | 5.550476 | 5.456564 | 2.035106 | -0.022489 | 2.777793 | 0.01537 | -0.031173 | 0.026408 | 1.091699 | 0.079318 | 3.031806 | 2.243381 | | 37.514 | 5.550476 | 2.051178 |
| 37.5347222 | 3.875382 | 1.376851 | 0.061977 | 1.657197 | 2.800842 | 4.351072 | 5.962796 | 3.755504 | 4.261098 | 4.959937 | 4.738376 | 4.821238 | 9.959868 | 3.742282 | 1.921853 | -0.455211 | 2.955874 | -0.013069 | -0.825107 | -0.361005 | 1.105691 | -0.096573 | 3.006331 | 3.138266 | | 37.528 | 9.959868 | 2.013941 |
| 37.5486111 | 7.018985 | -0.43274 | 0.97818 | 1.998044 | 3.37288 | 7.039487 | 4.249737 | 0.498157 | 6.569347 | 2.832696 | 7.641104 | 4.537116 | 17.801825 | 1.881184 | 1.10521 | -1.393439 | 2.914948 | -0.46187 | -2.239877 | -1.091943 | 0.928362 | -0.653022 | 2.566649 | 3.64168 | | 37.542 | 17.801825 | 1.098253 |
| 37.5625 | 10.616281 | -0.880855 | 1.592299 | 2.134442 | 4.215864 | 7.261305 | 2.496409 | -0.60544 | 4.038671 | 1.707474 | 9.361085 | 4.126065 | 19.056264 | 1.176164 | 0.655954 | -1.921038 | 1.730149 | -1.936956 | -2.549217 | -1.548643 | 0.579854 | -1.681191 | 1.871773 | 1.047354 | | 37.556 | 19.056264 | -0.333199 |
| 37.5763889 | 4.20838 | -0.192921 | 0.785002 | 2.153675 | 2.89725 | 4.41662 | -0.451472 | 0.65684 | 3.120073 | 2.123331 | 6.556027 | 1.166085 | 10.278165 | 1.405971 | -0.127121 | -1.725061 | 0.632832 | -1.918178 | -1.817444 | -1.67147 | 0.505175 | -1.728704 | 1.322731 | -0.219447 | | 37.569 | 10.278165 | -0.547542 |
| 37.5902778 | 0.574691 | -0.349386 | 0.29755 | 2.928695 | 2.640706 | 4.186531 | -2.931389 | 1.688332 | 2.69267 | 3.476184 | 1.679959 | -0.543819 | 3.134663 | 0.908926 | -0.459212 | -0.862567 | 0.81814 | -0.963334 | -0.954047 | -0.879843 | 1.2847 | -0.978492 | 1.258349 | -0.205127 | | 37.583 | 3.134663 | -0.357764 |
| 37.6041667 | -1.309173 | 0.180317 | 0.656255 | 2.914552 | 2.946837 | 4.692528 | -1.371549 | 2.499272 | 1.754442 | 3.554042 | 0.212131 | 0.036498 | 1.753439 | -1.159798 | -0.267424 | -0.331382 | 1.235353 | -0.39434 | -0.36925 | -0.330486 | 2.323213 | -0.399913 | 1.823385 | 0.252512 | | 37.597 | 1.753439 | 0.136553 |
| 37.6180556 | 0.920905 | 0.306838 | 0.512897 | 1.430949 | 2.835005 | 2.679135 | 2.27293 | 1.433016 | 0.787323 | 2.420746 | -1.103033 | 0.78411 | 2.013749 | -4.622314 | -0.123764 | -0.146432 | 1.046845 | -0.207626 | -0.15323 | -0.12959 | 3.658903 | -0.168653 | 2.181156 | 0.459549 | | 37.611 | 2.013749 | 0.371569 |
| 37.6319444 | 1.97599 | 1.420868 | -0.264855 | -0.514477 | 1.770427 | 0.554676 | 4.847732 | -0.651127 | -0.542433 | 0.816452 | -3.16108 | -0.382247 | 1.697471 | -4.024209 | 0.128254 | -0.083918 | 0.773364 | -0.140785 | -0.070311 | -0.052926 | 3.977078 | -0.075646 | 1.64039 | 0.310747 | | 37.625 | 1.697471 | 0.219243 |
| 37.6458333 | 1.002065 | 3.284359 | -0.320437 | -1.495752 | -0.427084 | -0.694526 | 4.787354 | -1.194597 | -0.975001 | -0.079648 | -2.734991 | -0.461616 | 3.261955 | -0.799887 | 0.422617 | -0.050926 | 0.594552 | -0.095162 | -0.0144 | 0.000986 | 0.980146 | -0.016557 | 0.690701 | 0.237939 | | 37.639 | 3.261955 | 0.160279 |
| 37.6597222 | -1.57928 | 2.181579 | -0.19756 | -0.408246 | -0.714178 | -0.340663 | 5.247097 | 0.434641 | 0.179598 | 0.682296 | 1.05978 | 1.666624 | 6.910704 | 1.204944 | 0.332576 | -0.043654 | 0.035203 | -0.077743 | 0.007059 | 0.035583 | -1.256912 | 0.007561 | -0.207787 | 0.011749 | | 37.653 | 6.910704 | -0.037689 |
| 37.6736111 | -2.588347 | 2.50939 | 0.390275 | 1.567741 | 0.043021 | 1.115126 | 5.920434 | 1.478093 | 0.668297 | 3.592208 | 3.172372 | 0.475592 | 10.127716 | 2.131455 | 0.122032 | -0.058772 | -0.880114 | -0.099918 | -0.01822 | 0.022551 | -2.842494 | -0.017466 | -1.174312 | -0.449795 | | 37.667 | 10.127716 | -0.433691 |
| 37.6875 | -3.4014 | 0.506714 | 1.207647 | 1.550512 | 2.504387 | 1.397211 | -2.672415 | 1.28127 | 0.395926 | 3.591089 | 0.970255 | 0.757067 | 8.430097 | -0.753369 | -0.325267 | -0.070299 | -1.023042 | -0.131226 | -0.054219 | -0.020456 | -1.521123 | -0.061565 | -1.978543 | -0.643462 | | 37.681 | 8.430097 | -0.774845 |
| 37.7013889 | -1.428896 | -3.360911 | 2.401121 | 0.643107 | 2.384643 | 0.700867 | 0.518552 | -0.060418 | 0.375807 | 1.37148 | -1.082707 | 1.563019 | 1.146353 | -2.234482 | -0.453363 | -0.033585 | -0.496834 | -0.112099 | -0.045443 | -0.0306 | 0.955844 | -0.053557 | -1.798621 | -0.354093 | | 37.694 | 1.146353 | -0.668672 |
| 37.7152778 | 1.92291 | -6.350968 | 4.477114 | 0.298543 | 1.691514 | 0.911622 | 4.110618 | 0.055444 | 0.254732 | 0.370214 | 1.49484 | -1.205152 | -5.419043 | -2.481938 | -0.485487 | 0.051015 | 0.234722 | -0.029951 | 0.029123 | 0.02752 | 1.810776 | 0.032159 | -0.955931 | -0.224453 | | 37.708 | -5.419043 | -0.402006 |
| 37.7291667 | 2.140449 | -2.271835 | 4.900639 | -0.536948 | 0.10211 | 1.348068 | 4.596254 | -0.183641 | 0.204831 | -0.369277 | 4.237629 | -2.841179 | -5.923719 | -1.615742 | -0.553032 | 0.18609 | 0.712681 | 0.095767 | 0.155587 | 0.136887 | 2.006006 | 0.16442 | -0.370393 | -0.15701 | | 37.722 | -5.923719 | -0.132351 |
| 37.7430556 | 2.893164 | 1.836825 | 2.54646 | 0.074591 | -0.61692 | 0.727425 | -0.27487 | -1.448938 | 0.429285 | -1.365086 | 4.230092 | -1.924719 | -2.979641 | -0.656119 | -0.642742 | 0.364873 | 0.681004 | 0.266916 | 0.341426 | 0.292225 | 1.260893 | 0.348711 | -0.225676 | 0.082288 | | 37.736 | -2.979641 | 0.07571 |
| 37.7569444 | 1.840683 | 3.48473 | 0.669006 | 0.632347 | -1.277344 | 0.327942 | -2.270433 | -0.543852 | 0.350401 | -0.77624 | 0.067122 | -0.655016 | 0.498368 | 0.202959 | -0.641413 | 0.542226 | 0.770131 | 0.43844 | 0.544975 | 0.469289 | 0.415512 | 0.548452 | -0.217216 | 0.240857 | | 37.75 | 0.498368 | 0.175279 |
| 37.7708333 | 1.117201 | 1.860213 | 0.199735 | -0.189468 | -0.82519 | 0.297749 | -1.359726 | 0.31651 | -0.693664 | -0.156477 | -2.370011 | 0.150651 | 3.846559 | 0.712155 | -0.610541 | 0.618961 | 0.541012 | 0.526574 | 0.65523 | 0.590909 | -1.087566 | 0.652265 | -0.264519 | 0.24429 | | 37.764 | 3.846559 | 0.131188 |
| 37.7847222 | 1.19242 | -0.027297 | -0.219072 | -0.256185 | -0.30482 | 0.391193 | -0.183975 | 0.215488 | -1.539782 | -2.059307 | -3.359591 | 0.507907 | 4.565585 | 0.956613 | -0.444043 | 0.590316 | 0.08241 | 0.516566 | 0.639936 | 0.608291 | -1.457827 | 0.631432 | -0.197764 | 0.101405 | | 37.778 | 4.565585 | 0.056141 |
| 37.7986111 | 0.756771 | -2.946091 | -0.739531 | 0.573728 | -0.157452 | 0.072479 | 1.536963 | 0.440541 | -0.14711 | -0.005714 | -3.804223 | 0.809645 | 2.097376 | 0.820639 | -0.027898 | 0.489063 | -0.72558 | 0.440997 | 0.543611 | 0.544298 | -1.067467 | 0.541342 | -0.246958 | -0.114179 | | 37.792 | 2.097376 | 0.00599 |
| 37.8125 | 0.000359 | -2.727372 | -1.046896 | 0.274779 | 0.22032 | -0.60171 | -0.099623 | 0.311378 | 0.553154 | 1.860312 | -3.746705 | 1.058147 | -0.602292 | -0.229585 | 0.421529 | 0.36725 | -2.070353 | 0.340279 | 0.420455 | 0.430831 | -0.540044 | 0.424327 | -0.552965 | -0.310445 | | 37.806 | -0.602292 | -0.07639 |
| 37.8263889 | -0.816788 | -2.092522 | -1.237466 | 1.362705 | 0.488496 | -1.126061 | -2.538948 | 0.179736 | 1.380129 | 2.117731 | -2.778614 | 1.208943 | -2.314257 | -1.951795 | 0.163351 | 0.25477 | -3.167055 | 0.240466 | 0.300053 | 0.305591 | 0.838226 | 0.305309 | -1.081807 | -0.429231 | | 37.819 | -2.314257 | -0.302833 |
| 37.8402778 | -4.740982 | -1.134384 | -1.242421 | 0.905251 | 0.143408 | -1.134438 | -2.882074 | -0.211963 | 1.100415 | -1.005144 | -1.230134 | 2.133467 | -2.676911 | -4.118482 | -0.349567 | 0.18022 | -3.370176 | 0.15684 | 0.204136 | 0.200902 | 2.584803 | 0.213005 | -1.231063 | -0.687761 | | 37.833 | -2.676911 | -0.650114 |
| 37.8541667 | -5.504971 | -1.185834 | -1.53612 | 0.787987 | 0.089125 | -1.235512 | -1.798784 | -0.464879 | 0.538105 | -1.705554 | -0.8367 | 1.693343 | -4.698314 | -5.107357 | -0.946134 | 0.139875 | -2.516631 | 0.087838 | 0.141732 | 0.11971 | 3.043718 | 0.15685 | -1.036756 | -1.140735 | | 37.847 | -4.698314 | -1.043686 |
| 37.8680556 | -4.820319 | -1.06712 | -1.714917 | -0.23117 | 0.218708 | -0.774246 | -1.257773 | -0.93795 | 0.089629 | 0.059269 | -1.039011 | -0.078139 | -7.36439 | -4.8221 | -1.595001 | 0.104836 | -1.649478 | 0.029097 | 0.090037 | 0.05457 | 1.068037 | 0.110326 | -0.944561 | -1.270557 | | 37.861 | -7.36439 | -1.172215 |
| 37.8819444 | -1.943869 | -1.123209 | -1.734792 | -0.743659 | -0.409411 | -0.253546 | -0.78142 | -0.625938 | -0.091898 | 0.176178 | -1.27587 | -1.998299 | -11.029248 | -3.659836 | -1.664721 | 0.045049 | -1.293434 | -0.024179 | 0.024812 | -0.004702 | -0.986253 | 0.047345 | -1.104041 | -1.137136 | | 37.875 | -11.029248 | -1.077439 |
| 37.8958333 | 0.59737 | -0.65886 | -1.691767 | -0.898905 | -0.695551 | -0.530357 | 0.30038 | -0.842776 | -0.47024 | -1.221033 | -0.556097 | -2.403584 | -10.927532 | -2.072978 | -1.307947 | -0.013221 | -2.313997 | -0.076753 | -0.036119 | -0.055472 | -3.732262 | -0.016404 | -1.531815 | -1.236575 | | 37.889 | -10.927532 | -1.134317 |
| 37.9097222 | 4.895663 | 0.579053 | -1.662688 | 0.254417 | -0.972967 | -1.268809 | 0.780759 | -0.838383 | -0.903547 | -1.573589 | -2.914727 | 0.733787 | -3.75798 | -1.022202 | -0.96286 | -0.032446 | -3.867204 | -0.1017 | -0.051848 | -0.076883 | -5.046008 | -0.03738 | -2.036558 | -1.660547 | | 37.903 | -3.75798 | -1.551038 |
| 37.9236111 | 3.128706 | 0.749716 | -0.871832 | 0.535113 | -1.389286 | -2.370869 | 1.617662 | 0.253856 | -1.257145 | -0.972944 | -5.195427 | -0.139423 | 2.841483 | -0.255801 | -0.856087 | 0.003999 | -4.639053 | -0.082316 | -0.02073 | -0.051958 | -5.316405 | -0.011432 | -2.385162 | -2.145557 | | 37.917 | 2.841483 | -1.996225 |
| 37.9375 | 0.444681 | 0.11302 | -0.23134 | -1.638864 | -0.863361 | -4.842603 | 4.771427 | 0.82439 | -0.557288 | 0.124288 | -4.582634 | 0.056212 | 10.018195 | -0.00614 | -0.935887 | 0.032413 | -4.200874 | -0.050947 | 0.010945 | -0.010235 | -4.499802 | 0.015841 | -2.25853 | -2.323449 | | 37.931 | 10.018195 | -2.353143 |
| 37.9513889 | -0.023222 | -0.253779 | -0.383909 | -2.46293 | 0.640067 | -6.53879 | 4.798189 | 0.496758 | 0.175763 | -0.531266 | -2.259164 | 1.573864 | 9.973233 | -0.162195 | -0.982019 | 0.036925 | -3.445982 | -0.030139 | 0.020634 | 0.017841 | -2.97314 | 0.029501 | -1.843929 | -2.005054 | | 37.944 | 9.973233 | -2.111741 |
| 37.9652778 | -0.946871 | -0.860461 | -0.546468 | -0.694488 | 0.528535 | -8.275183 | 0.529154 | 0.163004 | 0.772546 | -0.310286 | 0.405984 | 1.493449 | 3.534035 | -0.039495 | -0.99979 | 0.038092 | -2.769472 | -0.021007 | 0.033353 | 0.033794 | -1.34106 | 0.043013 | -1.441642 | -1.594197 | | 37.958 | 3.534035 | -1.710228 |
| 37.9791667 | -1.794634 | -1.089482 | -0.280679 | 0.0382 | -0.64618 | -4.951364 | -2.17797 | -0.062669 | 1.172852 | 0.306254 | -0.348889 | -0.071292 | -2.126614 | -0.070672 | -0.989404 | 0.020331 | -2.389433 | -0.034979 | 0.028906 | 0.029962 | -0.682009 | 0.036346 | -1.243136 | -1.349351 | | 37.972 | -2.126614 | -1.457583 |
| 37.9930556 | -1.878371 | -1.354406 | -0.471442 | 1.419654 | -1.953696 | -0.003117 | 0.816221 | -0.238437 | 1.109393 | -0.331638 | -3.323501 | -1.746255 | -6.391837 | -0.339292 | -1.071051 | -0.01122 | -2.191684 | -0.061746 | 0.00245 | 0.010861 | -0.501784 | 0.012529 | -1.106495 | -1.18861 | | 37.986 | -6.391837 | -1.356336 |
| 38.0069445 | -2.214275 | -1.756215 | -1.2876 | 2.206624 | -3.447068 | 3.398236 | 2.689343 | -0.518872 | 0.664083 | -0.382173 | -6.075548 | -2.531454 | -8.357868 | -0.652788 | -1.05947 | -0.004326 | -1.676677 | -0.0476 | 0.024937 | 0.021166 | -0.474951 | 0.028193 | -0.817277 | -1.084331 | | 38 | -8.357868 | -1.198299 |
| 38.0208333 | -3.061794 | -2.275121 | -1.835276 | 0.468312 | -4.166893 | 5.515719 | 1.747594 | -0.248638 | -0.812662 | 2.323817 | -6.673034 | -4.467417 | -10.341673 | -0.89027 | -0.956592 | 0.126056 | -1.178592 | 0.081986 | 0.194413 | 0.14729 | -0.436917 | 0.171647 | -0.458589 | -0.994708 | | 38.014 | -10.341673 | -1.121232 |

**Supplementary Table S7 (f)** Direct motion in 100 mM NaCl stressed Amaranth and their respective controls under dimG day/night lighting conditions

| Time | dimGControl1 | dimGControl2 | dimGControl3 | dimGControl4 | dimGControl5 | dimGControl6 | dimGControl7 | dimGControl8 | dimGControl9 | dimGControl10 | dimGControl11 | dimGControl12 | dimGControltank | dimG Stress1 | dimG Stress2 | dimG Stress3 | dimG Stress4 | dimG Stress5 | dimG Stress6 | dimG Stress7 | dimG Stress8 | dimG Stress9 | dimG Stress10 | dimGStresstank | | Time | Mean ControldimG | Mean StressdimG |
|---|---|---|---|---|---|---|---|---|---|---|---|---|---|---|---|---|---|---|---|---|---|---|---|---|---|---|---|---|
| 18 | 0.257636 | 0.033696 | 0.101469 | 0.033852 | 0.095972 | 0.095759 | 0.129426 | 0.091965 | 0.176731 | 0.073003 | 0.050673 | -0.279135 | 0.258951 | 0.447187 | 0.309397 | 0.480633 | 0.362183 | -0.071754 | 0.435376 | 0.576147 | 0.173775 | 0.50304 | 0.44958 | 0.580823 | | 18.09722 | 0.258951 | 0.49336 |
| 18.014 | 0.091484 | -0.015292 | 0.077028 | 0.042138 | 0.015714 | 0.034376 | 0.015456 | 0.019582 | 0.087541 | 0.034707 | -0.01693 | -0.283633 | 0.063093 | 0.550543 | 0.488158 | 0.553745 | 0.534068 | 0.252538 | 0.466632 | 0.648778 | 0.3306 | 0.567292 | 0.488074 | 0.358855 | | 18.11111 | 0.063093 | 0.370954 |
| 18.028 | 0.161061 | -0.013051 | 0.008584 | -0.018234 | -0.048836 | -0.037042 | -0.054148 | -0.032838 | 0.008561 | -0.033126 | -0.097606 | -0.267522 | -0.04064 | 0.507672 | 0.477342 | 0.529817 | 0.524514 | 0.431616 | 0.528177 | 0.623467 | 0.374177 | 0.545222 | 0.43105 | 0.464297 | | 18.125 | -0.04064 | 0.444188 |
| 18.042 | 0.178665 | -0.018682 | -0.009506 | -0.082086 | -0.091325 | -0.066211 | -0.08768 | -0.050639 | -0.03204 | -0.060714 | -0.153388 | -0.187368 | -0.075982 | 0.400006 | 0.363442 | 0.411232 | 0.424085 | 0.458519 | 0.51925 | 0.502128 | 0.329429 | 0.414225 | 0.266931 | 0.241727 | | 18.13889 | -0.075982 | 0.278917 |
| 18.056 | 0.084869 | -0.039773 | -0.041196 | -0.161774 | -0.119662 | -0.078188 | -0.112098 | -0.066123 | -0.079994 | -0.075351 | -0.205238 | -0.127997 | -0.093308 | 0.214555 | 0.192081 | 0.225818 | 0.255708 | 0.378708 | 0.390213 | 0.326564 | 0.196002 | 0.206343 | 0.027401 | -0.064316 | | 18.15278 | -0.093308 | 0.019811 |
| 18.069 | 0.077528 | -0.029552 | -0.081655 | -0.230659 | -0.115223 | -0.06863 | -0.121774 | -0.074643 | -0.110741 | -0.066595 | -0.231178 | -0.100316 | -0.076685 | 0.001229 | 0.017361 | 0.049783 | 0.059098 | 0.188242 | 0.192876 | 0.140737 | 0.025477 | -0.012526 | -0.231832 | -0.076131 | | 18.16667 | -0.076685 | -0.050764 |
| 18.083 | 0.092575 | -0.022389 | -0.088018 | -0.267025 | -0.112269 | -0.054523 | -0.139083 | -0.077742 | -0.121754 | -0.058517 | -0.232106 | -0.103914 | -0.063397 | -0.055057 | -0.008608 | 0.0289 | 0.00321 | 0.054112 | 0.120256 | 0.086065 | -0.014177 | -0.093951 | -0.344528 | 0.126441 | | 18.18056 | -0.063397 | 0.058439 |
| 18.097 | 0.005711 | -0.057048 | -0.046729 | -0.264947 | -0.122824 | -0.056371 | -0.176685 | -0.089946 | -0.13399 | -0.077484 | -0.235988 | -0.126632 | -0.103236 | 0.076576 | 0.123662 | 0.156894 | 0.127143 | 0.083286 | 0.227967 | 0.20802 | 0.121983 | 0.035522 | -0.273747 | 0.183439 | | 18.19444 | -0.103236 | 0.13249 |
| 18.111 | -0.081936 | -0.081016 | -0.018042 | -0.208596 | -0.123564 | -0.059965 | -0.183416 | -0.101847 | -0.144012 | -0.089058 | -0.232833 | -0.132946 | -0.133982 | 0.178898 | 0.220676 | 0.239551 | 0.224306 | 0.125401 | 0.299274 | 0.298717 | 0.224409 | 0.166419 | -0.211362 | 0.14897 | | 18.20833 | -0.133982 | 0.153171 |
| 18.125 | -0.105163 | -0.066304 | -0.017427 | -0.146672 | -0.097665 | -0.059831 | -0.158846 | -0.103455 | -0.14043 | -0.067786 | -0.199381 | -0.1038 | -0.13025 | 0.109457 | 0.154778 | 0.147599 | 0.13724 | 0.040819 | 0.152672 | 0.182915 | 0.127587 | 0.067193 | -0.35113 | 0.119332 | | 18.22222 | -0.13025 | 0.10139 |
| 18.139 | -0.081676 | -0.040176 | -0.027106 | -0.118937 | -0.073799 | -0.071092 | -0.149023 | -0.109252 | -0.129604 | -0.051768 | -0.148025 | -0.08022 | -0.143075 | -0.077255 | -0.038695 | -0.068039 | -0.071143 | -0.135682 | -0.152581 | -0.091156 | -0.090181 | -0.178765 | -0.572733 | -0.042129 | | 18.23611 | -0.143075 | -0.096466 |
| 18.153 | -0.028786 | -0.036559 | -0.085909 | -0.134776 | -0.086823 | -0.099809 | -0.157643 | -0.119669 | -0.12203 | -0.073705 | -0.171916 | -0.09739 | -0.160497 | -0.179309 | -0.158742 | -0.187054 | -0.16743 | -0.21128 | -0.311245 | -0.21925 | -0.165928 | -0.24573 | -0.563751 | -0.212457 | | 18.25 | -0.160497 | -0.226049 |
| 18.167 | -0.020405 | -0.0797 | -0.125174 | -0.145825 | -0.117356 | -0.110629 | -0.166759 | -0.10963 | -0.1256 | -0.109401 | -0.213536 | -0.123306 | -0.16437 | -0.110014 | -0.103841 | -0.106004 | -0.080527 | -0.116404 | -0.184326 | -0.108446 | -0.052671 | -0.105068 | -0.34654 | -0.205839 | | 18.26389 | -0.16437 | -0.14036 |
| 18.181 | -0.088582 | -0.119518 | -0.108995 | -0.137084 | -0.129835 | -0.099808 | -0.175324 | -0.103698 | -0.12738 | -0.120578 | -0.187632 | -0.130059 | -0.164828 | -0.032939 | -0.017372 | -0.01026 | 0.00017 | -0.047241 | -0.052009 | -0.024774 | 0.03297 | -0.017919 | -0.23791 | -0.110632 | | 18.27778 | -0.164828 | -0.045195 |
| 18.194 | -0.103641 | -0.099354 | -0.102149 | -0.117702 | -0.105138 | -0.092337 | -0.148969 | -0.10926 | -0.114196 | -0.100685 | -0.118001 | -0.11352 | -0.121251 | -0.077084 | -0.041047 | -0.042578 | -0.044317 | -0.086457 | -0.106387 | -0.095107 | -0.013155 | -0.068425 | -0.239152 | -0.01724 | | 18.29167 | -0.121251 | -0.037484 |
| 18.208 | -0.013365 | -0.045245 | -0.093218 | -0.058832 | -0.05626 | -0.073722 | -0.086189 | -0.075157 | -0.078339 | -0.060879 | -0.046576 | -0.065832 | -0.035904 | -0.150827 | -0.12037 | -0.128883 | -0.13329 | -0.129505 | -0.199118 | -0.190093 | -0.104478 | -0.14071 | -0.24353 | -0.008312 | | 18.30556 | -0.035904 | -0.064816 |
| 18.222 | 0.034169 | -0.009676 | -0.069764 | 0.025045 | -0.028158 | -0.0459 | -0.043899 | -0.027133 | -0.050036 | -0.037012 | -0.00955 | -0.029734 | -0.006961 | -0.188186 | -0.180014 | -0.198686 | -0.196078 | -0.149524 | -0.248664 | -0.241881 | -0.173811 | -0.189827 | -0.239124 | -0.118403 | | 18.31944 | -0.006961 | -0.126301 |
| 18.236 | -0.046179 | -0.051306 | -0.076136 | 0.040936 | -0.061177 | -0.046663 | -0.061801 | -0.032254 | -0.063894 | -0.071234 | -0.059828 | -0.062809 | -0.078423 | -0.226841 | -0.240122 | -0.244959 | -0.239277 | -0.167026 | -0.281806 | -0.285029 | -0.220983 | -0.228105 | -0.22112 | -0.239803 | | 18.33333 | -0.078423 | -0.214082 |
| 18.25 | -0.112749 | -0.102364 | -0.076589 | -0.005579 | -0.090835 | -0.057652 | -0.085763 | -0.057445 | -0.079063 | -0.101153 | -0.099095 | -0.09245 | -0.140056 | -0.165617 | -0.195221 | -0.170896 | -0.171611 | -0.072764 | -0.180956 | -0.216726 | -0.154199 | -0.147295 | -0.102448 | -0.210888 | | 18.34722 | -0.140056 | -0.185336 |
| 18.264 | -0.053652 | -0.051551 | -0.054129 | -0.048898 | -0.04535 | -0.046751 | -0.048839 | -0.042295 | -0.048472 | -0.058767 | -0.048571 | -0.044158 | -0.082072 | 0.016978 | -0.022814 | 0.001484 | -0.005291 | 0.113668 | 0.019953 | -0.039505 | -0.018352 | 0.040108 | 0.100023 | -0.014121 | | 18.36111 | -0.082072 | -0.015583 |
| 18.278 | 0.019893 | 0.010926 | -0.037684 | -0.072075 | 0.004465 | -0.027773 | -0.008083 | -0.016719 | -0.01416 | -0.012852 | -0.007683 | -0.004049 | -0.000731 | 0.139494 | 0.100632 | 0.113009 | 0.102564 | 0.231058 | 0.134911 | 0.083504 | 0.019176 | 0.158669 | 0.23074 | 0.165204 | | 18.375 | -0.000731 | 0.120726 |
| 18.292 | 0.02425 | 0.001861 | -0.033395 | -0.079146 | -0.000085 | -0.028624 | -0.01896 | -0.021498 | -0.01912 | -0.01827 | -0.019452 | -0.01413 | -0.005837 | 0.166488 | 0.136458 | 0.149342 | 0.140182 | 0.268274 | 0.170124 | 0.134307 | 0.031306 | 0.214115 | 0.278513 | 0.222307 | | 18.38889 | -0.005837 | 0.17907 |
| 18.306 | -0.008669 | -0.036712 | -0.031039 | -0.074746 | -0.028872 | -0.041985 | -0.04914 | -0.038339 | -0.044468 | -0.043904 | -0.036518 | -0.032439 | -0.055133 | 0.162427 | 0.136369 | 0.151351 | 0.147332 | 0.262236 | 0.186837 | 0.147317 | 0.077971 | 0.255856 | 0.27481 | 0.180357 | | 18.40278 | -0.055133 | 0.183937 |
| 18.319 | -0.040282 | -0.060678 | -0.036546 | -0.081593 | -0.050849 | -0.054041 | -0.069084 | -0.052917 | -0.061712 | -0.063902 | -0.034621 | -0.050096 | -0.09778 | 0.112246 | 0.083422 | 0.091883 | 0.091261 | 0.202067 | 0.132057 | 0.082073 | 0.067969 | 0.212577 | 0.196288 | 0.081938 | | 18.41667 | -0.09778 | 0.106592 |
| 18.333 | -0.028134 | -0.062054 | -0.042763 | -0.096853 | -0.05533 | -0.062257 | -0.065475 | -0.05767 | -0.059076 | -0.067815 | -0.023571 | -0.051841 | -0.118544 | 0.062761 | 0.022451 | 0.026111 | 0.031167 | 0.144978 | 0.057514 | 0.002726 | 0.03073 | 0.140611 | 0.121683 | 0.0051 | | 18.43056 | -0.118544 | 0.010911 |
| 18.347 | 0.030608 | -0.059764 | -0.050456 | -0.106973 | -0.056272 | -0.071015 | -0.062545 | -0.082384 | -0.05709 | -0.073545 | -0.035821 | -0.048695 | -0.127871 | 0.034171 | -0.005668 | -0.008615 | 0.001979 | 0.124629 | 0.00716 | -0.049165 | 0.005816 | 0.085138 | 0.075682 | -0.042803 | | 18.44444 | -0.127871 | -0.049199 |
| 18.361 | 0.058384 | -0.138004 | -0.118212 | -0.234294 | -0.15174 | -0.170305 | -0.177627 | -0.313711 | -0.130167 | -0.174377 | -0.154228 | -0.155115 | -0.185379 | -0.096947 | -0.121861 | -0.120433 | -0.118661 | -0.037809 | -0.159075 | -0.26426 | -0.118985 | -0.134143 | -0.122126 | -0.212786 | | 18.45833 | -0.185379 | -0.207719 |
| 18.375 | 0.026104 | -0.315622 | -0.14301 | -0.377634 | -0.340242 | -0.359291 | -0.433932 | -0.715942 | -0.250974 | -0.334313 | -0.456488 | -0.372438 | -0.402793 | -0.291312 | -0.320308 | -0.286997 | -0.254138 | -0.249532 | -0.40301 | -0.562888 | -0.277567 | -0.393605 | -0.37107 | -0.719596 | | 18.47222 | -0.402793 | -0.604676 |
| 18.389 | -0.001825 | -0.476211 | 0.431454 | 0.037718 | -0.477562 | -0.272878 | -0.777998 | -0.670833 | -0.04643 | -0.347697 | -0.681148 | -0.346771 | -1.029221 | -0.183746 | -0.400143 | -0.3711 | -0.220449 | -0.286281 | -0.520468 | -0.55304 | -0.270472 | -0.290454 | -0.329082 | -3.32342 | | 18.48611 | -1.029221 | -2.392149 |
| 18.403 | 0.709815 | -0.84791 | 1.432844 | 0.879604 | -0.570401 | 0.140186 | -1.248166 | 0.261346 | 0.272019 | -0.310043 | -0.38985 | -0.081732 | -2.093219 | 0.245875 | -0.422545 | -0.70603 | -0.122942 | -0.260317 | -0.148178 | -0.463095 | -0.187513 | 0.14642 | -0.145431 | -9.108863 | | 18.5 | -2.093219 | -7.271394 |
| 18.417 | 0.995832 | -1.017845 | 1.541297 | 0.984965 | -0.575032 | 0.194446 | -1.242423 | -0.168155 | -0.09726 | -0.769606 | 0.117672 | -0.054503 | -3.767556 | -0.018566 | -0.531312 | -1.012725 | -0.224972 | -0.19925 | -0.114145 | -0.560903 | -0.247617 | 0.555532 | -0.443636 | -11.783637 | | 18.51389 | -3.767556 | -11.909655 |
| 18.431 | 0.456532 | -0.387519 | 0.009084 | -0.007976 | -0.142438 | -0.06211 | -0.133726 | -0.448193 | -0.540378 | -0.626743 | 0.430067 | 0.090111 | -1.86802 | -0.362745 | -0.411042 | -0.311602 | -0.218246 | 0.220173 | -0.088216 | -0.801048 | -0.148864 | 0.00198 | -0.631865 | -2.160136 | | 18.52778 | -1.86802 | -3.016824 |
| 18.444 | -0.609437 | 0.852227 | -1.75168 | -1.176698 | 0.39856 | -0.450317 | 1.095195 | -0.489943 | -0.220225 | 0.31416 | -0.056189 | -0.069331 | 2.750943 | -1.210604 | -0.603788 | -0.072329 | -0.86549 | -0.277372 | -0.491101 | -0.784166 | -0.847277 | -1.272286 | -0.49491 | 10.556035 | | 18.54167 | 2.750943 | 8.862467 |
| 18.458 | -0.567193 | 0.983415 | -1.920381 | -1.242675 | 0.429261 | -0.623034 | 1.140257 | -0.719607 | -0.001593 | 0.499624 | -0.190614 | 0.058119 | 2.858122 | -2.753771 | -2.129464 | -1.438218 | -2.552367 | -1.984559 | -2.02364 | -2.101481 | -2.27194 | -2.955358 | -1.566726 | 8.983255 | | 18.55556 | 2.858122 | 6.381064 |
| 18.472 | -0.258433 | 0.412398 | -0.924521 | -0.527338 | 0.10831 | -0.385337 | 0.39336 | -0.406662 | 0.034936 | 0.128786 | -0.098883 | 0.201807 | 1.003003 | -3.02595 | -2.881185 | -2.545564 | -3.02781 | -2.695209 | -3.055914 | -2.647335 | -2.587836 | -3.420203 | -2.060236 | 1.356701 | | 18.56944 | 1.003003 | 0.165567 |
| 18.486 | -0.029057 | 0.044946 | -0.225517 | -0.010601 | -0.04929 | -0.094311 | 0.061238 | -0.005542 | 0.103352 | -0.067056 | 0.011027 | 0.34173 | 0.12875 | -1.833543 | -1.689252 | -1.44545 | -1.69223 | -1.491782 | -1.751405 | -1.409369 | -1.377563 | -1.796407 | -1.077424 | -1.150189 | | 18.58333 | 0.12875 | -1.266633 |
| 18.5 | 0.094737 | 0.11911 | -0.029517 | 0.164056 | 0.017815 | 0.056537 | 0.152859 | 0.139105 | 0.164569 | -0.009964 | 0.123831 | 0.353102 | 0.129212 | -0.652222 | -0.648757 | -0.449558 | -0.594868 | -0.247715 | -0.55711 | -0.321425 | -0.279001 | -0.537073 | -0.251684 | -0.50293 | | 18.59722 | 0.129212 | -0.39405 |
| 18.514 | 0.171175 | 0.214615 | -0.046389 | 0.152568 | 0.06894 | 0.05387 | 0.241778 | 0.099562 | 0.093191 | 0.031776 | 0.204469 | 0.228288 | 0.142903 | -0.126986 | -0.13026 | 0.017457 | -0.114681 | 0.474488 | -0.018818 | 0.168353 | 0.167966 | 0.033952 | 0.06533 | -0.016419 | | 18.61111 | 0.142903 | 0.090378 |
| 18.528 | 0.183144 | 0.195338 | -0.087217 | 0.091771 | 0.070272 | 0.021609 | 0.278119 | 0.060744 | 0.038875 | 0.019932 | 0.183417 | 0.19819 | 0.10924 | 0.095491 | 0.086624 | 0.205644 | 0.090271 | 0.540436 | 0.186826 | 0.370486 | 0.296126 | 0.293422 | 0.215366 | 0.145874 | | 18.625 | 0.10924 | 0.262236 |
| 18.542 | 0.175551 | 0.162786 | -0.082285 | 0.05696 | 0.092294 | 0.025168 | 0.252724 | 0.081649 | 0.048748 | 0.026447 | 0.172145 | 0.264586 | 0.088364 | 0.301943 | 0.259271 | 0.346393 | 0.271463 | 0.497069 | 0.39336 | 0.57531 | 0.389311 | 0.484971 | 0.414186 | 0.320121 | | 18.63889 | 0.088364 | 0.421193 |
| 18.556 | 0.151864 | 0.111656 | -0.028811 | 0.040468 | 0.118387 | 0.036628 | 0.187981 | 0.106501 | 0.078719 | 0.036843 | 0.147702 | 0.321012 | 0.06215 | 0.496564 | 0.430698 | 0.504599 | 0.45394 | 0.522668 | 0.619025 | 0.784126 | 0.5226 | 0.669854 | 0.621877 | 0.530098 | | 18.65278 | 0.06215 | 0.586478 |
| 18.569 | 0.0528 | 0.028212 | -0.024317 | -0.000426 | 0.090862 | -0.000788 | 0.078544 | 0.073589 | 0.061254 | -0.00076 | 0.093274 | 0.277666 | 0.002931 | 0.607476 | 0.546425 | 0.603908 | 0.570002 | 0.514858 | 0.775403 | 0.891229 | 0.64307 | 0.787196 | 0.77744 | 0.648349 | | 18.66667 | 0.002931 | 0.658139 |
| 18.583 | -0.033907 | -0.033004 | -0.037024 | -0.039154 | 0.050114 | -0.042599 | 0.001137 | 0.036565 | 0.038342 | -0.041467 | 0.029039 | 0.169621 | -0.033653 | 0.639173 | 0.601682 | 0.62657 | 0.613436 | 0.482246 | 0.817732 | 0.873443 | 0.708326 | 0.822333 | 0.827903 | 0.652252 | | 18.68056 | -0.033653 | 0.647881 |
| 18.597 | -0.037879 | -0.015999 | -0.013011 | -0.030018 | 0.042413 | -0.042057 | -0.001904 | 0.026036 | 0.039611 | -0.026286 | -0.016344 | 0.113967 | -0.012165 | 0.665549 | 0.644496 | 0.642749 | 0.643679 | 0.51479 | 0.837383 | 0.7964 | 0.756815 | 0.84558 | 0.85154 | 0.684903 | | 18.69444 | -0.012165 | 0.688558 |
| 18.611 | -0.02066 | -0.008011 | 0.021771 | 0.002113 | 0.023864 | -0.02489 | 0.008282 | 0.021004 | 0.03284 | -0.013141 | -0.106876 | 0.061945 | 0.001214 | 0.762348 | 0.733047 | 0.755173 | 0.738788 | 0.677608 | 0.959267 | 0.835337 | 0.863222 | 0.972025 | 0.966514 | 0.832535 | | 18.70833 | 0.001214 | 0.818061 |
| 18.625 | -0.005995 | -0.005166 | 0.060806 | 0.032538 | 0.003493 | -0.002484 | 0.0102 | 0.026033 | 0.026542 | -0.005788 | -0.12412 | 0.03373 | 0.000344 | 0.926378 | 0.887222 | 0.93833 | 0.896557 | 0.88367 | 1.155847 | 0.979914 | 0.996664 | 1.187383 | 1.135992 | 1.012294 | | 18.72222 | 0.000344 | 0.979269 |
| 18.639 | 0.019012 | 0.040464 | 0.076342 | 0.052711 | 0.01721 | 0.006073 | 0.025102 | 0.026837 | 0.030482 | 0.023672 | -0.046085 | 0.050546 | 0.021082 | 0.981195 | 0.954836 | 0.987188 | 0.94571 | 0.936189 | 1.170677 | 1.02986 | 0.997075 | 1.28188 | 1.174633 | 1.037634 | | 18.73611 | 0.021082 | 1.020883 |
| 18.653 | -0.0101 | 0.019189 | 0.042391 | 0.037503 | -0.004153 | -0.022679 | 0.009275 | 0.012168 | 0.004826 | 0.003571 | -0.040565 | 0.015089 | 0.015962 | 0.771553 | 0.741941 | 0.766375 | 0.734238 | 0.725558 | 0.891529 | 0.81349 | 0.726332 | 1.039465 | 0.937437 | 0.801002 | | 18.75 | 0.015962 | 0.802715 |
| 18.667 | -0.073338 | -0.083623 | -0.002559 | -0.005292 | -0.064735 | -0.058529 | -0.04738 | -0.010127 | -0.045767 | -0.064303 | -0.12334 | -0.050419 | -0.03391 | 0.418025 | 0.363623 | 0.403736 | 0.374696 | 0.36041 | 0.489036 | 0.432248 | 0.336356 | 0.575309 | 0.499156 | 0.426294 | | 18.76389 | -0.03391 | 0.420537 |
| 18.681 | -0.054438 | -0.109758 | -0.004916 | -0.01243 | -0.065717 | -0.05311 | -0.077617 | -0.017391 | -0.058579 | -0.068699 | -0.157306 | -0.035914 | -0.052554 | 0.180402 | 0.129474 | 0.155592 | 0.135046 | 0.098832 | 0.214831 | 0.163991 | 0.112656 | 0.237372 | 0.187252 | 0.179356 | | 18.77778 | -0.052554 | 0.147758 |
| 18.694 | -0.015316 | -0.056381 | -0.006303 | 0.005604 | -0.02384 | -0.040772 | -0.090775 | -0.022785 | -0.046026 | -0.027591 | -0.148805 | 0.007076 | -0.039104 | 0.065998 | 0.033611 | 0.025048 | 0.023205 | -0.030727 | 0.062257 | 0.026724 | 0.040042 | 0.059261 | 0.044408 | 0.039031 | | 18.79167 | -0.039104 | -0.011066 |
| 18.708 | -0.069624 | -0.067461 | -0.062213 | -0.020102 | -0.048339 | -0.074448 | -0.161921 | -0.073068 | -0.071676 | -0.057735 | -0.188261 | -0.04505 | -0.065218 | -0.044988 | -0.069215 | -0.075474 | -0.076693 | -0.12155 | -0.076957 | -0.088195 | -0.023026 | -0.097525 | -0.045164 | -0.099483 | | 18.80556 | -0.065218 | -0.148238 |
| 18.722 | -0.117486 | -0.11765 | -0.09159 | -0.052447 | -0.095589 | -0.089586 | -0.211902 | -0.115125 | -0.093541 | -0.111077 | -0.205897 | -0.113778 | -0.087874 | -0.07469 | -0.09615 | -0.066729 | -0.08765 | -0.093168 | -0.089367 | -0.09143 | -0.013717 | -0.128008 | -0.047806 | -0.132853 | | 18.81944 | -0.087874 | -0.130509 |
| 18.736 | -0.089527 | -0.117822 | -0.055325 | -0.034913 | -0.080896 | -0.053318 | -0.171204 | -0.094221 | -0.066526 | -0.096241 | -0.152667 | -0.0935 | -0.056013 | -0.012717 | -0.038205 | 0.016387 | -0.017429 | -0.01164 | 0.018755 | 0.001288 | 0.041446 | -0.043325 | 0.017032 | -0.053332 | | 18.83333 | -0.056013 | 0.011823 |
| 18.75 | -0.058059 | -0.099552 | -0.047709 | -0.024643 | -0.049771 | -0.043928 | -0.147359 | -0.073507 | -0.050161 | -0.06012 | -0.074969 | -0.049516 | -0.028452 | -0.020575 | -0.044147 | -0.010574 | -0.03526 | -0.066367 | 0.010091 | -0.017148 | 0.002057 | -0.051338 | 0.00799 | -0.044011 | | 18.84722 | -0.028452 | 0.005231 |
| 18.764 | -0.066402 | -0.085888 | -0.081985 | -0.037973 | -0.039889 | -0.063154 | -0.122366 | -0.072056 | -0.065997 | -0.045134 | -0.027578 | -0.042091 | -0.026558 | -0.152046 | -0.150861 | -0.16344 | -0.171598 | -0.26043 | -0.150473 | -0.191164 | -0.154773 | -0.211792 | -0.113845 | -0.174895 | | 18.86111 | -0.026558 | -0.216761 |
| 18.778 | -0.091405 | -0.086694 | -0.082019 | -0.036178 | -0.041622 | -0.052874 | -0.090411 | -0.042692 | -0.072105 | -0.045531 | -0.015171 | -0.067341 | -0.004741 | -0.258372 | -0.237597 | -0.262631 | -0.273231 | -0.401817 | -0.239569 | -0.310139 | -0.274018 | -0.343864 | -0.222377 | -0.289149 | | 18.875 | -0.004741 | -0.398142 |
| 18.792 | -0.108574 | -0.106476 | -0.045843 | -0.025504 | -0.049866 | -0.025309 | -0.074345 | -0.002233 | -0.062226 | -0.053191 | -0.027412 | -0.101664 | 0.029858 | -0.240098 | -0.2186 | -0.223871 | -0.254826 | -0.396842 | -0.15345 | -0.279973 | -0.267697 | -0.34089 | -0.223227 | -0.280767 | | 18.88889 | 0.029858 | -0.362656 |
| 18.806 | -0.11569 | -0.128869 | -0.01921 | -0.026475 | -0.052375 | -0.01931 | -0.09228 | -0.007434 | -0.056722 | -0.054045 | -0.035439 | -0.117281 | 0.031054 | -0.154249 | -0.132333 | -0.142412 | -0.17027 | -0.284677 | -0.030864 | -0.220852 | -0.20488 | -0.260716 | -0.171521 | -0.193162 | | 18.90278 | 0.031054 | -0.207667 |
| 18.819 | -0.123432 | -0.140766 | -0.011938 | -0.0186 | -0.0346 | -0.017968 | -0.091575 | -0.017141 | -0.052175 | -0.033623 | -0.02408 | -0.110392 | 0.0177 | -0.06626 | -0.046501 | -0.078605 | -0.087515 | -0.143658 | 0.016676 | -0.167897 | -0.12265 | -0.150117 | -0.119745 | -0.093933 | | 18.91667 | 0.0177 | -0.061554 |
| 18.833 | -0.13539 | -0.131652 | -0.025421 | -0.006169 | -0.009692 | -0.018748 | -0.061414 | -0.02198 | -0.047949 | -0.001885 | -0.005262 | -0.094263 | 0.018066 | -0.038065 | -0.001589 | -0.060017 | -0.050984 | -0.068862 | -0.070051 | -0.145456 | -0.065853 | -0.084234 | -0.097663 | -0.030637 | | 18.93056 | 0.018066 | -0.007477 |
| 18.847 | -0.15451 | -0.114746 | -0.050106 | -0.017012 | -0.011859 | -0.040179 | -0.047972 | -0.063445 | -0.065196 | 0.00433 | -0.009355 | -0.093317 | -0.002126 | -0.102405 | -0.032658 | -0.121697 | -0.098714 | -0.124076 | -0.293757 | -0.207234 | -0.10349 | -0.140383 | -0.158451 | -0.063496 | | 18.94444 | -0.002126 | -0.067422 |
| 18.861 | -0.18481 | -0.117082 | -0.04694 | -0.032147 | -0.040942 | -0.045797 | -0.060275 | -0.096744 | -0.091291 | -0.01991 | -0.044164 | -0.129592 | -0.029607 | -0.166434 | -0.090392 | -0.171799 | -0.152667 | -0.19425 | -0.450851 | -0.261248 | -0.180227 | -0.220276 | -0.24119 | -0.121978 | | 18.95833 | -0.029607 | -0.130687 |
| 18.875 | -0.218985 | -0.127041 | -0.012474 | -0.026076 | -0.059034 | -0.01683 | -0.081426 | -0.079242 | -0.078093 | -0.037371 | -0.058274 | -0.166712 | -0.019299 | -0.12303 | -0.071682 | -0.107105 | -0.105983 | -0.165422 | -0.422643 | -0.203551 | -0.16636 | -0.185441 | -0.210559 | -0.060432 | | 18.97222 | -0.019299 | -0.051505 |
| 18.889 | -0.255483 | -0.103479 | 0.009807 | -0.014535 | -0.057121 | 0.003378 | -0.084915 | -0.056393 | -0.063366 | -0.024332 | -0.03119 | -0.164751 | -0.003062 | -0.006033 | 0.021674 | 0.023897 | 0.012009 | -0.073451 | -0.260164 | -0.082803 | -0.066538 | -0.061674 | -0.086617 | 0.116497 | | 18.98611 | -0.003062 | 0.130599 |
| 18.903 | -0.23922 | -0.064812 | 0.011863 | -0.000261 | -0.044599 | 0.011146 | -0.057792 | -0.044075 | -0.041762 | -0.000401 | 0.003238 | -0.138366 | 0.00287 | 0.104561 | 0.117211 | 0.131917 | 0.114771 | 0.004889 | -0.114328 | 0.044403 | 0.034657 | 0.056617 | 0.037286 | 0.240652 | | 19 | 0.00287 | 0.269533 |
| 18.917 | -0.115482 | -0.032779 | 0.103934 | 0.131195 | 0.007941 | 0.1075 | -0.006238 | 0.133847 | 0.01662 | 0.054881 | 0.174516 | 0.002049 | -0.06067 | 0.243958 | 0.194396 | 0.211139 | 0.223812 | 0.114975 | 0.065395 | 0.204889 | 0.167253 | 0.2431 | 0.125008 | -0.070259 | | 19.01389 | -0.06067 | 0.083328 |
| 18.931 | 0.212793 | -0.202596 | 0.542407 | 0.642214 | 0.012908 | 0.404105 | -0.165626 | 0.640396 | 0.060293 | 0.026971 | 0.493295 | 0.315544 | -0.844132 | 0.347018 | 0.113143 | 0.0826 | 0.381393 | -0.007687 | 0.232515 | 0.246192 | 0.354659 | 0.527051 | -0.051407 | -2.91552 | | 19.02778 | -0.844132 | -2.033311 |
| 18.944 | 0.793033 | -0.56311 | 1.26012 | 1.327223 | -0.138136 | 0.747778 | -0.58612 | 1.086243 | 0.056936 | -0.141198 | 0.741333 | 0.608838 | -2.364306 | 0.481194 | -0.218569 | -0.441304 | 0.303615 | -0.845164 | 0.106493 | -0.132319 | 0.386314 | 0.517892 | -0.611948 | -10.113572 | | 19.04167 | -2.364306 | -6.89362 |
| 18.958 | 0.253494 | -0.381092 | 0.936708 | 0.963994 | -0.093077 | 0.574991 | -0.337862 | 0.847983 | 0.251754 | -0.030733 | 0.359908 | 0.558583 | -1.610217 | 0.103874 | -0.369355 | -0.805701 | -0.248826 | -1.438492 | -0.510671 | -0.347025 | -0.280597 | 0.010538 | -0.97399 | -9.12125 | | 19.05556 | -1.610217 | -7.715458 |
| 18.972 | -0.506941 | 0.509933 | -0.583175 | -0.481216 | 0.316328 | -0.04916 | 0.83936 | -0.135883 | 0.353347 | 0.417545 | -0.05438 | 0.050026 | 1.826071 | -0.417245 | -0.045891 | 0.151859 | -0.32809 | -0.088353 | -0.169647 | 0.172365 | -0.460122 | -0.121943 | 0.23449 | 7.752947 | | 19.06944 | 1.826071 | 7.318583 |
| 18.986 | -0.64025 | 0.782063 | -1.153767 | -1.02893 | 0.441315 | -0.345109 | 1.182497 | -0.317885 | 0.158091 | 0.465576 | -0.195268 | -0.283817 | 2.702155 | -0.232827 | 0.393008 | 0.731701 | -0.034846 | 0.63956 | 0.422577 | 0.514463 | -0.097149 | 0.081384 | 0.743708 | 9.561139 | | 19.08333 | 2.702155 | 6.865038 |
| 19 | -0.172553 | 0.294101 | -0.503386 | -0.541448 | 0.191397 | -0.12325 | 0.567693 | -0.127402 | -0.080515 | 0.098083 | -0.031291 | -0.15196 | 1.04342 | 0.211427 | 0.337189 | 0.433596 | 0.20439 | 0.054246 | 0.395931 | 0.241065 | 0.099138 | 0.20655 | 0.451307 | 3.639051 | | 19.09722 | 1.04342 | 2.537986 |
| 19.014 | -0.000553 | -0.004472 | -0.079262 | -0.177309 | 0.013207 | -0.001523 | 0.131537 | 0.008962 | -0.078953 | -0.05297 | 0.004238 | -0.057112 | 0.094233 | 0.330275 | 0.305079 | 0.309659 | 0.308782 | -0.067701 | 0.362863 | 0.022302 | 0.173931 | 0.317169 | 0.332416 | 0.566131 | | 19.11111 | 0.094233 | 0.432359 |
| 19.028 | 0.068592 | -0.033331 | -0.01888 | -0.102691 | -0.029388 | -0.00717 | 0.049018 | -0.047137 | -0.048089 | -0.041429 | -0.096824 | -0.069821 | -0.045058 | 0.293883 | 0.304002 | 0.281729 | 0.241475 | 0.2066 | 0.315649 | -0.016304 | 0.055915 | 0.266038 | 0.34882 | 0.451958 | | 19.125 | -0.045058 | 0.503715 |
| 19.042 | 0.087556 | -0.030384 | -0.032395 | -0.1145 | -0.032941 | -0.0411 | 0.048367 | -0.10625 | -0.091449 | -0.038294 | -0.189882 | -0.099093 | -0.0466 | 0.114044 | 0.198117 | 0.165938 | 0.082045 | 0.330045 | 0.141373 | -0.185663 | -0.143153 | 0.098357 | 0.238802 | 0.59079 | | 19.13889 | -0.0466 | 0.590848 |
| 19.056 | 0.054385 | -0.038561 | -0.054641 | -0.11142 | -0.031414 | -0.066185 | -0.000576 | -0.130139 | -0.17685 | -0.038589 | -0.277814 | -0.109062 | -0.071082 | -0.02653 | -0.008352 | -0.006973 | -0.02535 | 0.123883 | -0.034935 | -0.354775 | -0.304955 | -0.064721 | 0.018452 | -0.076983 | | 19.15278 | -0.071082 | -0.147644 |
| 19.069 | 0.016097 | -0.037838 | -0.079647 | -0.086024 | -0.029413 | -0.067102 | -0.054868 | -0.110451 | -0.234397 | -0.037895 | -0.27664 | -0.104665 | -0.101019 | -0.042894 | -0.103856 | -0.067902 | -0.02999 | -0.055666 | -0.091177 | -0.341737 | -0.329308 | -0.074474 | -0.088294 | -0.626179 | | 19.16667 | -0.101019 | -0.70625 |
| 19.083 | 0.006994 | -0.020514 | -0.076499 | -0.054435 | -0.024034 | -0.049334 | -0.080498 | -0.077049 | -0.235793 | -0.033564 | -0.215189 | -0.100605 | -0.107032 | -0.025932 | -0.03291 | 0.005714 | -0.014391 | 0.089068 | -0.093116 | -0.140559 | -0.224132 | -0.02455 | 0.027045 | -0.075497 | | 19.18056 | -0.107032 | -0.092023 |
| 19.097 | 0.026784 | -0.015918 | -0.057002 | -0.041651 | -0.028758 | -0.042337 | -0.11884 | -0.056886 | -0.188916 | -0.035212 | -0.193552 | -0.109958 | -0.083365 | -0.057737 | 0.000045 | 0.02472 | -0.04961 | 0.196321 | -0.157613 | -0.011525 | -0.137365 | -0.068798 | 0.096822 | 0.480869 | | 19.19444 | -0.083365 | 0.505208 |
| 19.111 | 0.035398 | -0.026092 | -0.037557 | -0.04513 | -0.041409 | -0.043781 | -0.149719 | -0.04654 | -0.145576 | -0.041754 | -0.151988 | -0.125966 | -0.071631 | -0.084111 | -0.104134 | -0.063731 | -0.082577 | 0.047811 | -0.262649 | -0.057605 | -0.080926 | -0.17474 | -0.047708 | -0.005138 | | 19.20833 | -0.071631 | -0.026901 |
| 19.125 | 0.010532 | -0.032016 | -0.015536 | -0.044088 | -0.047038 | -0.044693 | -0.175186 | -0.035397 | -0.114737 | -0.042769 | -0.125156 | -0.144635 | -0.079069 | -0.07728 | -0.186186 | -0.114182 | -0.081795 | -0.021903 | -0.299138 | -0.111842 | -0.05297 | -0.233352 | -0.17261 | -0.552726 | | 19.22222 | -0.079069 | -0.63061 |
| 19.139 | -0.013685 | -0.033679 | -0.007156 | -0.042094 | -0.035759 | -0.04606 | -0.169921 | -0.02626 | -0.073781 | -0.03448 | -0.10698 | -0.165145 | -0.041719 | -0.094461 | -0.17608 | -0.102002 | -0.108574 | 0.032768 | -0.296004 | -0.09651 | -0.071235 | -0.279755 | -0.149011 | -0.379662 | | 19.23611 | -0.041719 | -0.409815 |
| 19.153 | 0.006164 | -0.032798 | -0.01278 | -0.035664 | -0.014545 | -0.037972 | -0.103588 | -0.022478 | -0.037639 | -0.023437 | -0.087676 | -0.16536 | -0.00049 | -0.136752 | -0.184779 | -0.118287 | -0.152143 | -0.033619 | -0.297935 | -0.107562 | -0.108177 | -0.349985 | -0.101499 | -0.176574 | | 19.25 | -0.00049 | -0.135101 |
| 19.167 | 0.024425 | -0.016978 | -0.031409 | -0.03214 | 0.000849 | -0.034206 | -0.03361 | -0.021281 | -0.022938 | -0.014917 | -0.055078 | -0.155804 | -0.000444 | -0.105183 | -0.191428 | -0.101216 | -0.12474 | -0.121962 | -0.202123 | -0.081168 | -0.081469 | -0.332305 | -0.021243 | -0.132455 | | 19.26389 | -0.000444 | -0.102718 |
| 19.181 | -0.004832 | -0.007263 | -0.06187 | -0.051176 | -0.00467 | -0.056149 | -0.031772 | -0.041509 | -0.03894 | -0.021688 | -0.032751 | -0.180407 | -0.009474 | -0.026227 | -0.272144 | -0.039091 | -0.058437 | -0.126395 | -0.080355 | -0.010843 | -0.021645 | -0.242989 | 0.07584 | -0.046868 | | 19.27778 | -0.009474 | -0.039601 |
| 19.194 | -0.051814 | -0.039333 | -0.061311 | -0.074335 | -0.034958 | -0.077101 | -0.079667 | -0.070009 | -0.059334 | -0.053511 | -0.078 | -0.23686 | -0.039828 | -0.061317 | -0.016105 | -0.065796 | -0.089341 | -0.261467 | -0.121128 | -0.05538 | -0.057125 | -0.265775 | 0.015995 | -0.04445 | | 19.29167 | -0.039828 | -0.037488 |
| 19.208 | -0.059379 | -0.048367 | -0.025166 | -0.069682 | -0.050471 | -0.067411 | -0.078378 | -0.079376 | -0.047477 | -0.064589 | -0.112536 | -0.256065 | -0.053566 | -0.186771 | -0.157233 | -0.187862 | -0.203947 | -0.491261 | -0.305623 | -0.207848 | -0.161821 | -0.397659 | -0.158093 | -0.171543 | | 19.30556 | -0.053566 | -0.164435 |
| 19.222 | -0.048966 | -0.016416 | -0.008303 | -0.055784 | -0.051071 | -0.057478 | -0.02769 | -0.082856 | -0.023992 | -0.0464 | -0.082858 | -0.218462 | -0.033281 | -0.334741 | -0.310343 | -0.327149 | -0.328254 | -0.624621 | -0.484969 | -0.348589 | -0.251509 | -0.507619 | -0.345678 | -0.311163 | | 19.31944 | -0.033281 | -0.338006 |
| 19.236 | -0.055585 | -0.018299 | -0.007224 | -0.04703 | -0.065379 | -0.051835 | -0.015519 | -0.077505 | -0.003237 | -0.040507 | -0.059715 | -0.175632 | -0.015487 | -0.472672 | -0.473254 | -0.441039 | -0.434047 | -0.59876 | -0.528556 | -0.435559 | -0.333213 | -0.597072 | -0.531019 | -0.403983 | | 19.33333 | -0.015487 | -0.457252 |
| 19.25 | -0.026766 | -0.011082 | 0.015113 | -0.015381 | -0.057816 | -0.025646 | -0.053704 | -0.046111 | 0.031901 | -0.025447 | -0.034521 | -0.102611 | 0.023681 | -0.505281 | -0.497805 | -0.455839 | -0.435937 | -0.397894 | -0.401322 | -0.440511 | -0.338409 | -0.568591 | -0.590475 | -0.416584 | | 19.34722 | 0.023681 | -0.440724 |
| 19.264 | 0.000828 | 0.028332 | -0.004566 | 0.007025 | -0.031393 | -0.014732 | -0.137554 | -0.008981 | 0.060976 | -0.007691 | -0.00112 | -0.033131 | 0.040502 | -0.3656 | -0.33036 | -0.316194 | -0.264267 | -0.12882 | -0.171718 | -0.289979 | -0.155016 | -0.326242 | -0.412769 | -0.298639 | | 19.36111 | 0.040502 | -0.270382 |
| 19.278 | -0.001264 | 0.063453 | -0.038094 | 0.001459 | -0.004125 | -0.012686 | -0.165794 | 0.028441 | 0.079069 | 0.001547 | 0.028081 | 0.026715 | 0.011503 | -0.148139 | -0.095537 | -0.093485 | -0.010985 | 0.129675 | 0.027268 | -0.050119 | 0.08452 | 0.008987 | -0.124904 | -0.056223 | | 19.375 | 0.011503 | -0.011135 |
| 19.292 | 0.005056 | 0.088766 | -0.022997 | -0.004806 | 0.034829 | 0.010079 | -0.017256 | 0.066224 | 0.080483 | 0.017492 | 0.066131 | 0.103121 | 0.007196 | 0.009833 | 0.074244 | 0.07621 | 0.179978 | 0.293883 | 0.125858 | 0.171471 | 0.220921 | 0.254367 | 0.079136 | 0.177311 | | 19.38889 | 0.007196 | 0.209356 |
| 19.306 | -0.000122 | 0.061568 | -0.00961 | -0.024231 | 0.047956 | 0.014626 | 0.197259 | 0.048776 | 0.056628 | 0.000619 | 0.042246 | 0.152494 | 0.024159 | 0.024607 | 0.088092 | 0.094209 | 0.224193 | 0.282127 | 0.102169 | 0.287733 | 0.199487 | 0.31209 | 0.120402 | 0.197242 | | 19.40278 | 0.024159 | 0.236409 |
| 19.319 | -0.013985 | 0.015914 | -0.010751 | -0.044817 | 0.03712 | -0.003716 | 0.248329 | -0.0001 | 0.031252 | -0.029589 | -0.008829 | 0.168203 | 0.048862 | -0.037441 | -0.006305 | 0.009574 | 0.160749 | 0.191418 | 0.027433 | 0.304155 | 0.096222 | 0.255837 | 0.044054 | 0.041269 | | 19.41667 | 0.048862 | 0.086557 |
| 19.333 | -0.011296 | 0.023697 | -0.0064 | -0.029542 | 0.044698 | -0.012107 | 0.240304 | -0.009335 | 0.028081 | -0.029867 | 0.011237 | 0.20671 | 0.04771 | -0.075691 | -0.099213 | -0.074804 | 0.097836 | 0.097521 | -0.03365 | 0.288819 | 0.031633 | 0.220217 | -0.020617 | -0.074966 | | 19.43056 | 0.04771 | -0.060144 |
| 19.347 | 0.00375 | 0.046789 | 0.02886 | -0.00912 | 0.038744 | -0.013861 | 0.281859 | -0.033957 | 0.024211 | -0.031875 | 0.055515 | 0.242964 | 0.005731 | -0.058852 | -0.155558 | -0.101683 | 0.117114 | 0.070094 | -0.002941 | 0.255085 | 0.025951 | 0.235698 | -0.062902 | -0.0762 | | 19.44444 | 0.005731 | -0.10612 |
| 19.361 | 0.018596 | 0.00625 | 0.026574 | -0.115348 | -0.067703 | -0.092474 | 0.211562 | -0.23099 | -0.030723 | -0.108311 | 0.002788 | 0.186405 | -0.076408 | 0.010624 | -0.144953 | -0.079151 | 0.161086 | 0.084967 | 0.104214 | 0.094804 | 0.063339 | 0.248142 | -0.166992 | -0.058899 | | 19.45833 | -0.076408 | -0.129306 |
| 19.375 | -0.054625 | -0.109739 | -0.092876 | -0.326556 | -0.272905 | -0.312608 | -0.02092 | -0.619913 | -0.170868 | -0.276022 | -0.246699 | -0.000325 | -0.211747 | 0.032541 | -0.138291 | -0.096371 | 0.175081 | 0.021089 | 0.066412 | -0.109988 | 0.095669 | 0.108881 | -0.274052 | -0.152537 | | 19.47222 | -0.211747 | -0.202895 |
| 19.389 | -0.246491 | -0.098615 | -0.198658 | -0.346962 | -0.331557 | -0.403269 | -0.118796 | -0.698916 | -0.214927 | -0.340749 | -0.465095 | -0.167528 | -0.291959 | -0.084965 | -0.296723 | -0.228391 | 0.08356 | -0.020057 | -0.205567 | -0.302189 | -0.007242 | -0.162941 | -0.393022 | -0.341124 | | 19.48611 | -0.291959 | -0.359274 |
| 19.403 | -0.145808 | -0.048491 | -0.179047 | -0.177155 | -0.185666 | -0.212355 | -0.020489 | -0.345823 | -0.130603 | -0.206525 | -0.19104 | -0.005122 | -0.217398 | -0.328807 | -0.634036 | -0.456811 | -0.19599 | -0.112646 | -0.367366 | -0.545321 | -0.27092 | -0.343996 | -0.653108 | -0.635164 | | 19.5 | -0.217398 | -0.657126 |
| 19.417 | 0.019805 | -0.005512 | -0.112737 | -0.047089 | -0.044428 | -0.049169 | 0.109669 | -0.061851 | -0.025799 | -0.066222 | 0.083485 | 0.159428 | -0.075669 | -0.56737 | -0.852794 | -0.562015 | -0.414492 | -0.160112 | -0.461933 | -0.681529 | -0.458325 | -0.388889 | -0.894274 | -0.847048 | | 19.51389 | -0.075669 | -0.898918 |
| 19.431 | 0.092223 | 0.04922 | -0.072678 | 0.014722 | 0.028531 | 0.017969 | 0.169489 | 0.031408 | 0.038946 | 0.009377 | 0.188628 | 0.239871 | -0.007376 | -0.463791 | -0.652534 | -0.407723 | -0.331327 | 0.090304 | -0.362846 | -0.506995 | -0.30143 | -0.241469 | -0.726441 | -0.684903 | | 19.52778 | -0.007376 | -0.74033 |
| 19.444 | 0.149445 | 0.136363 | -0.116227 | 0.042592 | 0.072087 | -0.009696 | 0.198864 | 0.014245 | 0.042178 | 0.063037 | 0.292211 | 0.27397 | -0.06437 | -0.335401 | -0.312296 | -0.268421 | -0.226161 | 0.33795 | -0.328652 | -0.287862 | -0.157012 | -0.087653 | -0.464574 | -0.265531 | | 19.54167 | -0.06437 | -0.375654 |
| 19.458 | 0.198963 | 0.178757 | -0.233312 | 0.015988 | 0.044405 | -0.128983 | 0.16676 | -0.088386 | -0.034365 | 0.054781 | 0.26646 | 0.256226 | -0.211603 | -0.634142 | -0.419794 | -0.566765 | -0.516836 | 0.015302 | -0.699552 | -0.470411 | -0.442578 | -0.386368 | -0.593449 | -0.144654 | | 19.55556 | -0.211603 | -0.297376 |
| 19.472 | 0.201389 | 0.07235 | -0.199983 | 0.000306 | -0.044625 | -0.134793 | 0.066445 | -0.065748 | -0.061551 | -0.015837 | 0.092311 | 0.224128 | -0.25827 | -0.963301 | -0.666731 | -0.798951 | -0.774988 | -0.241566 | -0.852057 | -0.632371 | -0.711287 | -0.702448 | -0.8064 | -0.348464 | | 19.56944 | -0.25827 | -0.530399 |
| 19.486 | 0.197919 | -0.033891 | -0.009412 | 0.070031 | -0.061866 | 0.022076 | 0.000297 | 0.098639 | 0.047712 | -0.030491 | 0.001634 | 0.194841 | -0.134017 | -0.713127 | -0.390794 | -0.391696 | -0.408175 | 0.11251 | -0.31436 | -0.283779 | -0.375484 | -0.409444 | -0.488423 | -0.346114 | | 19.58333 | -0.134017 | -0.497301 |
| 19.5 | 0.199995 | 0.001296 | 0.099801 | 0.130792 | 0.007582 | 0.116969 | 0.020758 | 0.175458 | 0.153232 | 0.033839 | 0.053326 | 0.197537 | -0.007757 | -0.147104 | 0.175959 | 0.19812 | 0.190737 | 0.680291 | 0.344692 | 0.212159 | 0.212881 | 0.188243 | 0.113962 | 0.018225 | | 19.59722 | -0.007757 | -0.068521 |
| 19.514 | 0.171019 | 0.036937 | 0.098129 | 0.126386 | 0.048221 | 0.091764 | 0.037469 | 0.136488 | 0.160613 | 0.05405 | 0.076287 | 0.159328 | 0.022227 | 0.222196 | 0.492918 | 0.493675 | 0.498176 | 0.830295 | 0.635767 | 0.37721 | 0.524521 | 0.535802 | 0.446715 | 0.2706 | | 19.61111 | 0.022227 | 0.247722 |
| 19.528 | 0.126872 | 0.017788 | 0.102448 | 0.117344 | 0.043719 | 0.043427 | 0.011908 | 0.089209 | 0.131526 | 0.021452 | 0.037237 | 0.063215 | 0.010261 | 0.337769 | 0.508936 | 0.506126 | 0.510722 | 0.671792 | 0.667752 | 0.26483 | 0.536421 | 0.590383 | 0.480989 | 0.316674 | | 19.625 | 0.010261 | 0.300491 |
| 19.542 | 0.094949 | 0.008042 | 0.145462 | 0.139494 | 0.051567 | 0.025328 | 0.018111 | 0.073978 | 0.112094 | 0.006583 | 0.021983 | -0.01378 | 0.001845 | 0.367589 | 0.417882 | 0.423058 | 0.414181 | 0.431437 | 0.59142 | 0.139929 | 0.42561 | 0.518216 | 0.41822 | 0.265537 | | 19.63889 | 0.001845 | 0.238475 |
| 19.556 | 0.084678 | 0.015745 | 0.135263 | 0.169601 | 0.066714 | 0.024498 | 0.076054 | 0.091462 | 0.093724 | 0.008867 | 0.02071 | -0.03465 | -0.008945 | 0.344827 | 0.310559 | 0.32749 | 0.298239 | 0.192483 | 0.493623 | 0.08876 | 0.317726 | 0.417489 | 0.340096 | 0.190575 | | 19.65278 | -0.008945 | 0.195747 |
| 19.569 | 0.081423 | 0.012953 | 0.04039 | 0.174037 | 0.056556 | 0.017093 | 0.065863 | 0.109634 | 0.071738 | 0.01193 | 0.017537 | -0.028461 | -0.017448 | 0.302143 | 0.233746 | 0.261087 | 0.215875 | 0.01182 | 0.389461 | 0.054436 | 0.238084 | 0.343199 | 0.280467 | 0.122501 | | 19.66667 | -0.017448 | 0.204411 |
| 19.583 | 0.058851 | 0.003939 | -0.004284 | 0.157963 | 0.034659 | 0.015545 | -0.009004 | 0.119486 | 0.042297 | 0.0159 | 0.0192 | -0.018946 | -0.007369 | 0.30896 | 0.22697 | 0.253899 | 0.207758 | 0.012129 | 0.35667 | 0.119157 | 0.226853 | 0.360678 | 0.281124 | 0.1171 | | 19.68056 | -0.007369 | 0.258195 |
| 19.597 | 0.027736 | -0.012881 | 0.013181 | 0.119258 | 0.008925 | 0.019192 | -0.068336 | 0.106377 | 0.021071 | 0.0007 | 0.010356 | -0.02044 | -0.001096 | 0.322217 | 0.215422 | 0.257547 | 0.214725 | 0.100667 | 0.367823 | 0.222875 | 0.236184 | 0.359274 | 0.313478 | 0.175777 | | 19.69444 | -0.001096 | 0.327535 |
| 19.611 | 0.010131 | -0.017174 | 0.040177 | 0.057238 | -0.012721 | 0.026325 | -0.082519 | 0.07527 | 0.030179 | -0.014648 | 0.006583 | -0.012381 | 0.013747 | 0.265438 | 0.146759 | 0.229826 | 0.186761 | 0.169896 | 0.34129 | 0.25404 | 0.222292 | 0.288447 | 0.301612 | 0.209737 | | 19.70833 | 0.013747 | 0.333664 |
| 19.625 | -0.015991 | 0.002026 | 0.041647 | -0.013655 | -0.022548 | 0.008608 | -0.062714 | 0.000353 | 0.04441 | -0.020141 | 0.005435 | -0.010512 | 0.033701 | 0.17493 | 0.078589 | 0.167298 | 0.139825 | 0.136347 | 0.251458 | 0.18569 | 0.182424 | 0.201642 | 0.220357 | 0.159147 | | 19.72222 | 0.033701 | 0.273696 |
| 19.639 | -0.018433 | 0.017787 | 0.045389 | -0.045839 | -0.024721 | 0.003903 | -0.018382 | -0.023972 | 0.052231 | -0.005265 | 0.011372 | -0.013327 | 0.036988 | 0.152704 | 0.086735 | 0.142225 | 0.131529 | 0.128009 | 0.222702 | 0.155592 | 0.178659 | 0.180332 | 0.159128 | 0.132307 | | 19.73611 | 0.036988 | 0.213378 |
| 19.653 | 0.016245 | 0.033135 | 0.047935 | -0.029809 | -0.015841 | 0.032633 | 0.027755 | 0.054222 | 0.058209 | 0.032585 | 0.050439 | -0.009538 | 0.017826 | 0.201424 | 0.141584 | 0.1714 | 0.166082 | 0.176362 | 0.266955 | 0.204101 | 0.217724 | 0.236976 | 0.172652 | 0.172879 | | 19.75 | 0.017826 | 0.234585 |
| 19.667 | 0.000533 | -0.005173 | -0.012563 | -0.046708 | -0.046604 | 0.004306 | -0.01362 | 0.091251 | -0.004847 | -0.004699 | 0.017155 | -0.076852 | -0.048496 | 0.170226 | 0.113432 | 0.137506 | 0.134586 | 0.146143 | 0.202596 | 0.186745 | 0.194757 | 0.208527 | 0.149036 | 0.173695 | | 19.76389 | -0.048496 | 0.234937 |
| 19.681 | -0.021474 | -0.049656 | -0.067104 | -0.0614 | -0.063805 | -0.037605 | -0.063494 | 0.062491 | -0.082223 | -0.06104 | -0.040409 | -0.13817 | -0.069116 | 0.064083 | -0.00173 | 0.03449 | 0.027665 | 0.048454 | 0.060939 | 0.06946 | 0.116 | 0.076894 | 0.041933 | 0.082308 | | 19.77778 | -0.069116 | 0.142701 |
| 19.694 | -0.012674 | -0.019237 | -0.056959 | -0.019169 | -0.006714 | -0.024578 | -0.044042 | 0.012331 | -0.035867 | -0.032992 | -0.001966 | -0.112318 | -0.003939 | 0.014189 | -0.069806 | -0.031448 | -0.049248 | -0.01815 | -0.023124 | -0.009137 | -0.017563 | -0.005956 | -0.033669 | -0.000913 | | 19.79167 | -0.003939 | 0.040962 |
| 19.708 | -0.047671 | -0.006001 | -0.056927 | 0.009424 | 0.02621 | -0.012849 | -0.049443 | -0.064891 | 0.040771 | -0.01288 | 0.011502 | -0.078935 | 0.038959 | 0.040792 | -0.075062 | -0.024009 | -0.061427 | -0.020409 | -0.035 | 0.012362 | -0.047249 | 0.012887 | -0.017488 | 0.006562 | | 19.80556 | 0.038959 | 0.014961 |
| 19.722 | -0.074739 | -0.02512 | -0.050209 | 0.012926 | 0.011375 | -0.004198 | -0.092687 | -0.123856 | 0.050687 | -0.027966 | -0.018759 | -0.077924 | 0.056925 | 0.120786 | -0.024054 | 0.06304 | 0.000767 | 0.068344 | 0.036288 | 0.131155 | -0.015221 | 0.092821 | 0.081054 | 0.109298 | | 19.81944 | 0.056925 | 0.101766 |
| 19.736 | -0.039545 | -0.01245 | -0.029579 | 0.003323 | 0.005538 | 0.004652 | -0.135678 | -0.120655 | 0.034806 | -0.016684 | -0.004785 | -0.066835 | 0.061619 | 0.177225 | 0.126473 | 0.194307 | 0.121757 | 0.206076 | 0.169394 | 0.300347 | 0.082726 | 0.179472 | 0.229587 | 0.274472 | | 19.83333 | 0.061619 | 0.268796 |
| 19.75 | -0.019529 | 0.005692 | -0.042452 | -0.042338 | 0.008358 | -0.015527 | -0.227471 | -0.058102 | 0.004942 | 0.005744 | 0.01736 | -0.097464 | 0.039536 | 0.069167 | 0.242369 | 0.25046 | 0.179787 | 0.270152 | 0.223809 | 0.404161 | 0.138759 | 0.140879 | 0.314002 | 0.371291 | | 19.84722 | 0.039536 | 0.387841 |
| 19.764 | -0.052633 | -0.032957 | -0.05851 | -0.104202 | -0.015124 | -0.034569 | -0.251671 | 0.028864 | -0.057402 | -0.003403 | 0.004467 | -0.210036 | 0.036828 | -0.155552 | 0.19499 | 0.200493 | 0.117569 | 0.223248 | 0.1588 | 0.388921 | 0.079434 | -0.076252 | 0.250966 | 0.354331 | | 19.86111 | 0.036828 | 0.377337 |
| 19.778 | -0.096559 | -0.111299 | -0.042243 | -0.129693 | -0.052692 | -0.037531 | 0.070225 | 0.021893 | -0.10108 | -0.030863 | -0.035028 | -0.400824 | 0.071064 | -0.281502 | 0.06314 | 0.095431 | -0.000254 | 0.117441 | 0.000904 | 0.303844 | -0.048414 | -0.331141 | 0.099276 | 0.256703 | | 19.875 | 0.071064 | 0.278442 |
| 19.792 | -0.112917 | -0.142129 | -0.017681 | -0.100121 | -0.061354 | -0.042465 | -0.151198 | -0.073749 | -0.101206 | -0.02701 | -0.060006 | -0.617473 | 0.108867 | -0.283049 | -0.012889 | 0.001839 | -0.088921 | 0.020687 | -0.146111 | 0.225519 | -0.123281 | -0.492368 | -0.016174 | 0.155804 | | 19.88889 | 0.108867 | 0.206421 |
| 19.806 | -0.101639 | -0.138264 | 0.004629 | -0.034645 | -0.029717 | -0.051367 | -0.174853 | -0.195853 | -0.037359 | 0.006043 | -0.049042 | -0.667665 | 0.137998 | -0.243128 | 0.013595 | -0.012033 | -0.10271 | 0.009337 | -0.118045 | 0.211358 | -0.159868 | -0.506535 | -0.022971 | 0.135621 | | 19.90278 | 0.137998 | 0.194216 |
| 19.819 | -0.141777 | -0.126365 | 0.006669 | -0.000226 | -0.027522 | -0.050839 | -0.212784 | -0.268742 | -0.029732 | 0.008386 | -0.047174 | -0.569883 | 0.140732 | -0.245819 | 0.038553 | -0.002563 | -0.125396 | 0.028015 | -0.119427 | 0.153178 | -0.160183 | -0.415599 | -0.027886 | 0.157948 | | 19.91667 | 0.140732 | 0.156751 |
| 19.833 | -0.222937 | -0.143971 | 0.01177 | -0.03303 | -0.057696 | -0.048997 | -0.34304 | -0.218342 | -0.065823 | -0.022287 | -0.079311 | -0.455129 | 0.132291 | -0.306636 | -0.005433 | -0.065192 | -0.233028 | 0.011429 | -0.284952 | -0.094018 | -0.164321 | -0.32035 | -0.11121 | 0.070287 | | 19.93056 | 0.132291 | 0.03788 |
| 19.847 | -0.273613 | -0.122276 | 0.036258 | -0.095766 | -0.06116 | -0.044948 | -0.483231 | -0.185288 | -0.091143 | -0.032448 | -0.10312 | -0.333882 | 0.114943 | -0.362469 | -0.057314 | -0.127815 | -0.313613 | 0.02281 | -0.473644 | -0.415809 | -0.176624 | -0.260081 | -0.181755 | -0.060444 | | 19.94444 | 0.114943 | -0.085873 |
| 19.861 | -0.256834 | -0.077084 | 0.06556 | -0.114397 | -0.021519 | -0.037689 | -0.528101 | -0.209205 | -0.102133 | 0.002334 | -0.067692 | -0.299103 | 0.084304 | -0.376024 | -0.05791 | -0.133765 | -0.262381 | 0.090263 | -0.587693 | -0.625303 | -0.241492 | -0.225522 | -0.158944 | -0.122833 | | 19.95833 | 0.084304 | -0.184312 |
| 19.875 | -0.237521 | -0.036779 | 0.061593 | -0.103377 | 0.014103 | -0.039675 | -0.482478 | -0.197243 | -0.101078 | 0.037613 | -0.007261 | -0.39139 | 0.043261 | -0.375903 | -0.043158 | -0.129285 | -0.216136 | 0.179773 | -0.633291 | -0.579694 | -0.481359 | -0.193505 | -0.111903 | -0.136815 | | 19.97222 | 0.043261 | -0.232629 |
| 19.889 | -0.244571 | -0.048444 | 0.035447 | -0.106236 | -0.015146 | -0.068038 | -0.362582 | -0.18709 | -0.102459 | 0.011054 | -0.025668 | -0.486247 | -0.008206 | -0.384232 | -0.079906 | -0.140035 | -0.246754 | 0.176798 | -0.652709 | -0.486212 | -0.567358 | -0.216494 | -0.104259 | -0.163626 | | 19.98611 | -0.008206 | -0.260596 |
| 19.903 | -0.229531 | -0.077834 | 0.031104 | -0.124799 | -0.052904 | -0.066398 | -0.273478 | -0.158018 | -0.085196 | -0.023686 | -0.061801 | -0.441055 | -0.015433 | -0.363729 | -0.105843 | -0.110242 | -0.282315 | 0.084035 | -0.553436 | -0.356392 | -0.49064 | -0.246549 | -0.092851 | -0.161655 | | 20 | -0.015433 | -0.244651 |
| 19.917 | -0.129952 | -0.030308 | 0.075082 | -0.060529 | 0.020497 | 0.031438 | -0.096924 | 0.038986 | -0.007164 | 0.054021 | 0.09523 | -0.147841 | 0.065395 | -0.238419 | 0.000876 | 0.000012 | -0.216748 | 0.09772 | -0.31783 | -0.140115 | -0.236972 | -0.126851 | 0.006343 | -0.072935 | | 20.01389 | 0.065395 | -0.100514 |
| 19.931 | 0.056921 | 0.105769 | 0.151745 | 0.12253 | 0.208185 | 0.241631 | 0.198948 | 0.474446 | 0.105099 | 0.245724 | 0.480849 | 0.288045 | 0.203405 | -0.056246 | 0.164725 | 0.143632 | -0.064096 | 0.140549 | -0.011713 | 0.08301 | 0.051107 | 0.094833 | 0.138068 | -0.010941 | | 20.02778 | 0.203405 | 0.021031 |
| 19.944 | 0.133532 | 0.245589 | 0.202143 | 0.247467 | 0.313612 | 0.373011 | 0.416586 | 0.792814 | 0.190641 | 0.35278 | 0.603163 | 0.517342 | 0.274551 | -0.244782 | -0.077675 | -0.099002 | -0.20785 | -0.150154 | -0.17888 | -0.140263 | -0.133681 | -0.180846 | -0.133504 | -0.473464 | | 20.04167 | 0.274551 | -0.311562 |
| 19.958 | 0.047133 | 0.170629 | 0.118996 | 0.166563 | 0.201759 | 0.267814 | 0.368067 | 0.631939 | 0.14713 | 0.225294 | 0.236363 | 0.321741 | 0.231268 | -0.769319 | -0.77308 | -0.77246 | -0.667745 | -0.837189 | -0.947643 | -0.884537 | -0.863613 | -0.916506 | -0.85976 | -1.348762 | | 20.05556 | 0.231268 | -1.070254 |
| 19.972 | -0.001016 | -0.006425 | 0.070935 | 0.034777 | 0.037035 | 0.148497 | 0.125892 | 0.267784 | 0.066083 | 0.05249 | -0.060519 | 0.055851 | 0.175895 | -0.796519 | -0.859431 | -0.844084 | -0.643672 | -0.982394 | -1.01122 | -0.904935 | -1.070536 | -1.004919 | -0.969517 | -1.40661 | | 20.06944 | 0.175895 | -1.16481 |
| 19.986 | -0.016472 | -0.079277 | 0.090759 | -0.003614 | -0.02711 | 0.071765 | -0.126128 | -0.045299 | 0.018902 | -0.007204 | -0.161503 | -0.095555 | 0.119487 | -0.211054 | -0.243585 | -0.207071 | -0.209997 | -0.395842 | -0.404625 | -0.227049 | -0.504451 | -0.30668 | -0.286662 | -0.482525 | | 20.08333 | 0.119487 | -0.385942 |
| 20 | -0.017502 | -0.075309 | 0.104211 | 0.012095 | -0.028952 | -0.008931 | -0.307648 | -0.143949 | -0.002896 | -0.002543 | -0.150167 | -0.156677 | 0.062887 | 0.099699 | 0.155113 | 0.14917 | -0.107354 | 0.080163 | -0.196271 | -0.02169 | -0.138891 | 0.027376 | 0.158086 | 0.094639 | | 20.09722 | 0.062887 | 0.114886 |
| 20.014 | 0.004935 | -0.047295 | 0.117315 | 0.021489 | -0.016854 | -0.018064 | -0.170944 | -0.009349 | 0.007356 | 0.008676 | -0.094181 | -0.08916 | 0.068447 | 0.055027 | 0.174584 | 0.124318 | 0.004407 | -0.470725 | -0.157874 | -0.237671 | -0.097507 | -0.041519 | 0.119469 | 0.247634 | | 20.11111 | 0.068447 | 0.271135 |
| 20.028 | 0.019786 | -0.009609 | 0.127225 | 0.005321 | -0.015044 | 0.039148 | 0.094084 | 0.114088 | 0.011899 | 0.011863 | -0.067674 | 0.029074 | 0.091832 | 0.01334 | 0.157244 | 0.117972 | 0.265172 | -0.412906 | 0.11739 | -0.167957 | -0.025928 | -0.031846 | 0.064125 | 0.365615 | | 20.125 | 0.091832 | 0.287885 |
| 20.042 | -0.001479 | -0.001299 | 0.086514 | -0.023264 | -0.033051 | 0.047967 | 0.129039 | 0.058078 | -0.020659 | -0.004818 | -0.088878 | 0.094471 | 0.062506 | -0.016075 | 0.144536 | 0.134198 | 0.282969 | -0.405271 | 0.206954 | 0.082862 | 0.019245 | -0.047517 | 0.053869 | 0.202781 | | 20.13889 | 0.062506 | 0.098726 |
| 20.056 | -0.006589 | -0.019144 | 0.021598 | -0.036615 | -0.047243 | 0.001718 | 0.010536 | -0.074901 | -0.048338 | -0.018511 | -0.095827 | 0.078182 | 0.00719 | -0.059318 | 0.095379 | 0.093665 | 0.070032 | -0.658268 | 0.038699 | 0.244684 | -0.048732 | -0.195372 | 0.014988 | -0.023968 | | 20.15278 | 0.00719 | -0.043871 |
| 20.069 | 0.030487 | -0.031077 | 0.000923 | -0.0287 | -0.049076 | -0.022242 | 0.005512 | -0.065409 | -0.056184 | -0.012465 | -0.06839 | -0.004554 | -0.025999 | -0.056538 | 0.079062 | 0.067541 | 0.010116 | -0.492154 | 0.006467 | 0.211499 | -0.107875 | -0.292384 | 0.020637 | -0.132604 | | 20.16667 | -0.025999 | -0.08579 |
| 20.083 | 0.064966 | -0.049926 | -0.004855 | -0.02134 | -0.055367 | -0.029267 | 0.058451 | 0.019935 | -0.059458 | -0.013264 | -0.05432 | -0.131974 | -0.045945 | -0.043753 | 0.091768 | 0.073173 | 0.046049 | -0.1618 | 0.078852 | 0.005756 | -0.097902 | -0.308459 | 0.067347 | -0.10791 | | 20.18056 | -0.045945 | -0.085867 |
| 20.097 | 0.044912 | -0.08417 | -0.01532 | -0.035967 | -0.068543 | -0.042476 | 0.109533 | 0.025148 | -0.065267 | -0.037006 | -0.070253 | -0.276928 | -0.045866 | -0.068869 | 0.084523 | 0.062192 | 0.038835 | -0.044669 | 0.034778 | -0.146778 | -0.051591 | -0.334319 | 0.109509 | -0.088042 | | 20.19444 | -0.045866 | -0.104822 |
| 20.111 | 0.02419 | -0.094468 | -0.003972 | -0.043695 | -0.049195 | -0.033824 | 0.209221 | -0.007415 | -0.049998 | -0.031346 | -0.080501 | -0.376664 | 0.010279 | -0.091008 | 0.065895 | 0.038465 | 0.063174 | -0.06762 | 0.00428 | -0.094556 | 0.021963 | -0.32642 | 0.154534 | -0.032913 | | 20.20833 | 0.010279 | -0.103702 |
| 20.125 | -0.000887 | -0.05936 | 0.008897 | -0.034428 | -0.019768 | -0.027607 | 0.216359 | -0.02358 | -0.047012 | -0.001769 | -0.067217 | -0.327928 | 0.057513 | -0.09757 | 0.017356 | -0.01512 | 0.051549 | -0.140135 | 0.048091 | -0.025473 | 0.074459 | -0.318754 | 0.184231 | -0.032155 | | 20.22222 | 0.057513 | -0.125776 |
| 20.139 | -0.073985 | -0.025061 | -0.036929 | -0.034751 | -0.033818 | -0.056333 | 0.068193 | -0.05063 | -0.092984 | -0.019765 | -0.075808 | -0.185481 | 0.015434 | -0.074056 | -0.014746 | -0.047255 | 0.024051 | -0.160587 | 0.068962 | 0.007058 | 0.101282 | -0.273237 | 0.195203 | -0.052053 | | 20.23611 | 0.015434 | -0.108844 |
| 20.153 | -0.119663 | -0.000423 | -0.056691 | -0.012717 | -0.051499 | -0.047145 | 0.0079 | -0.044265 | -0.101841 | -0.04389 | -0.084343 | -0.013219 | -0.041461 | 0.096363 | 0.108126 | 0.099343 | 0.176324 | 0.009138 | 0.145571 | 0.133404 | 0.215065 | -0.009889 | 0.30172 | 0.13217 | | 20.25 | -0.041461 | 0.105052 |
| 20.167 | -0.101639 | 0.041989 | -0.032318 | 0.021922 | -0.032289 | -0.007128 | -0.00701 | -0.002303 | -0.04642 | -0.031308 | -0.066786 | 0.06779 | -0.034321 | 0.270171 | 0.249606 | 0.260837 | 0.292677 | 0.174929 | 0.182794 | 0.208283 | 0.295339 | 0.255123 | 0.347504 | 0.334714 | | 20.26389 | -0.034321 | 0.290877 |
| 20.181 | -0.099141 | 0.03849 | -0.058822 | 0.013835 | -0.026273 | -0.026483 | -0.070635 | -0.000653 | -0.019523 | -0.033591 | -0.084414 | 0.015041 | -0.038331 | 0.103799 | 0.098864 | 0.088273 | 0.040684 | 0.032 | -0.065987 | -0.028924 | 0.086116 | 0.127807 | 0.102698 | 0.205217 | | 20.27778 | -0.038331 | 0.127406 |
| 20.194 | -0.125331 | -0.021551 | -0.09723 | -0.004044 | -0.040937 | -0.067852 | -0.080849 | -0.025171 | -0.038269 | -0.054894 | -0.112145 | -0.072002 | -0.077243 | -0.250743 | -0.200748 | -0.253978 | -0.355241 | -0.257733 | -0.415085 | -0.391811 | -0.246486 | -0.243279 | -0.198133 | 0.120342 | | 20.29167 | -0.077243 | 0.046432 |
| 20.208 | -0.132119 | -0.05093 | -0.083714 | 0.004696 | -0.03497 | -0.059154 | -0.002968 | -0.0322 | -0.064857 | -0.041339 | -0.102338 | -0.111997 | -0.070444 | -0.362917 | -0.28815 | -0.363699 | -0.501288 | -0.342838 | -0.480048 | -0.504346 | -0.331405 | -0.4582 | -0.25981 | 0.170605 | | 20.30556 | -0.070444 | 0.139112 |
| 20.222 | -0.099062 | 0.005321 | -0.037444 | 0.019701 | -0.009568 | -0.027245 | 0.08331 | -0.00703 | -0.066684 | -0.007549 | -0.072333 | -0.105277 | -0.023923 | -0.147668 | -0.166316 | -0.199943 | -0.328188 | -0.189478 | -0.198524 | -0.252375 | -0.080529 | -0.276054 | -0.113674 | -0.197985 | | 20.31944 | -0.023923 | -0.204378 |
| 20.236 | -0.050409 | 0.090904 | 0.02001 | 0.034201 | 0.016414 | 0.004134 | 0.162151 | 0.055072 | -0.026633 | 0.012283 | -0.005117 | -0.06252 | 0.000783 | 0.207689 | 0.087812 | 0.10985 | 0.051599 | 0.115779 | 0.251574 | 0.219768 | 0.346298 | 0.238331 | 0.132193 | -0.357598 | | 20.33333 | 0.000783 | -0.392725 |
| 20.25 | 0.005346 | 0.123504 | 0.051087 | 0.049151 | 0.026913 | 0.030193 | 0.250518 | 0.096762 | 0.02521 | 0.016711 | 0.070232 | -0.00428 | 0.01562 | 0.402536 | 0.289756 | 0.334181 | 0.301144 | 0.314733 | 0.46965 | 0.513227 | 0.562187 | 0.58973 | 0.230925 | 0.111652 | | 20.34722 | 0.01562 | 0.089824 |
| 20.264 | 0.018131 | 0.081472 | 0.025121 | 0.036882 | 0.014233 | 0.029504 | 0.373322 | 0.065006 | 0.04375 | 0.00996 | 0.084198 | 0.041167 | 0.041918 | 0.21521 | 0.169683 | 0.184295 | 0.1611 | 0.151164 | 0.247213 | 0.334564 | 0.351306 | 0.482056 | 0.047023 | 0.254179 | | 20.36111 | 0.041918 | 0.240753 |
| 20.278 | -0.016919 | 0.00488 | -0.032142 | -0.001088 | -0.017017 | -0.004119 | 0.396324 | 0.03334 | 0.014488 | -0.017308 | 0.044264 | 0.060463 | 0.021955 | -0.100867 | -0.112921 | -0.130921 | -0.102781 | -0.148959 | -0.08548 | 0.004702 | 0.071528 | 0.268447 | -0.193343 | 0.004581 | | 20.375 | 0.021955 | -0.004642 |
| 20.292 | -0.029586 | -0.044978 | -0.045787 | -0.010734 | -0.030566 | -0.020901 | 0.349712 | 0.079116 | -0.008842 | -0.032177 | 0.066416 | 0.084487 | -0.029555 | -0.137995 | -0.200521 | -0.225628 | -0.118204 | -0.220095 | -0.107157 | -0.036529 | 0.065887 | 0.335021 | -0.234447 | -0.051185 | | 20.38889 | -0.029555 | -0.013982 |
| 20.306 | 0.000467 | -0.019375 | -0.00692 | 0.017804 | -0.004514 | 0.009699 | 0.398189 | 0.161938 | 0.012445 | -0.002669 | 0.192323 | 0.12056 | -0.044956 | 0.114922 | -0.125308 | -0.13895 | 0.022204 | -0.137315 | 0.047437 | 0.165848 | 0.205337 | 0.484272 | -0.196925 | 0.047579 | | 20.40278 | -0.044956 | 0.138822 |
| 20.319 | 0.039069 | 0.052364 | 0.011226 | 0.023393 | 0.012816 | 0.021807 | 0.401137 | 0.15204 | 0.029035 | 0.018938 | 0.264998 | 0.133919 | -0.02818 | 0.373961 | -0.143052 | -0.15023 | 0.031781 | -0.179332 | 0.017151 | 0.26299 | 0.22535 | 0.426859 | -0.239778 | -0.004568 | | 20.41667 | -0.02818 | 0.142881 |
| 20.333 | 0.073441 | 0.103627 | 0.007873 | 0.006325 | -0.002538 | -0.014951 | 0.436737 | 0.095738 | 0.008317 | -0.00748 | 0.207264 | 0.154133 | -0.035536 | 0.426967 | -0.362517 | -0.357909 | -0.160293 | -0.428053 | -0.242221 | 0.0871 | 0.024015 | 0.217534 | -0.445505 | -0.286142 | | 20.43056 | -0.035536 | -0.106352 |
| 20.347 | 0.113574 | 0.09001 | 0.026807 | -0.008959 | -0.031397 | -0.02727 | 0.596008 | 0.052126 | -0.02116 | -0.066185 | 0.067497 | 0.181099 | -0.109276 | 0.319391 | -0.579574 | -0.530397 | -0.338415 | -0.6534 | -0.42883 | -0.087954 | -0.170866 | 0.023831 | -0.67906 | -0.511537 | | 20.44444 | -0.109276 | -0.341073 |
| 20.361 | 0.170782 | 0.029732 | 0.018455 | -0.082218 | -0.094804 | -0.077353 | 0.77928 | -0.121517 | -0.05849 | -0.175303 | -0.071211 | 0.149567 | -0.20346 | 0.230918 | -0.567924 | -0.460081 | -0.275971 | -0.66522 | -0.314304 | -0.158701 | -0.099294 | 0.02393 | -0.761416 | -0.458228 | | 20.45833 | -0.20346 | -0.338654 |
| 20.375 | 0.125756 | -0.114012 | -0.054787 | -0.18917 | -0.19259 | -0.252453 | 0.756353 | -0.432455 | -0.123217 | -0.307764 | -0.278111 | -0.029528 | -0.285915 | 0.100165 | -0.453329 | -0.314643 | -0.109976 | -0.573566 | -0.167226 | -0.232516 | 0.04489 | 0.04739 | -0.695215 | -0.401032 | | 20.47222 | -0.285915 | -0.278085 |
| 20.389 | -0.013672 | -0.067085 | -0.117064 | -0.116039 | -0.16702 | -0.314832 | 0.665833 | -0.473073 | -0.104794 | -0.287913 | -0.411148 | -0.157258 | -0.259974 | 0.201997 | -0.1814 | -0.022103 | 0.16057 | -0.268839 | 0.087037 | -0.046027 | 0.19798 | 0.176655 | -0.246102 | -0.215384 | | 20.48611 | -0.259974 | -0.144005 |
| 20.403 | 0.08857 | 0.041962 | -0.119406 | 0.068755 | -0.018154 | -0.124399 | 0.660024 | -0.12252 | -0.01326 | -0.10764 | -0.124228 | 0.002384 | -0.121467 | 0.545747 | 0.322821 | 0.415068 | 0.508992 | 0.218278 | 0.729369 | 0.499797 | 0.473787 | 0.715176 | 0.294794 | 0.217107 | | 20.5 | -0.121467 | 0.212274 |
| 20.417 | 0.163792 | 0.071616 | -0.085939 | 0.161851 | 0.055292 | -0.009464 | 0.620886 | 0.147441 | 0.052436 | 0.002317 | 0.042243 | 0.080028 | -0.040973 | 0.645694 | 0.589687 | 0.661761 | 0.655211 | 0.457756 | 1.037911 | 0.77232 | 0.605298 | 1.048645 | 0.475914 | 0.429489 | | 20.51389 | -0.040973 | 0.424422 |
| 20.431 | 0.13234 | 0.051583 | -0.094013 | 0.154503 | 0.038028 | -0.008614 | 0.481763 | 0.144065 | 0.033475 | 0.005185 | 0.036639 | 0.049353 | -0.049471 | 0.304051 | 0.388546 | 0.429217 | 0.391944 | 0.222998 | 0.735636 | 0.558455 | 0.321939 | 0.804095 | 0.229475 | 0.175529 | | 20.52778 | -0.049471 | 0.194779 |
| 20.444 | 0.165709 | 0.084707 | -0.153456 | 0.11388 | 0.03168 | -0.083 | 0.314308 | 0.03145 | -0.013383 | 0.006425 | 0.051336 | 0.039502 | -0.117773 | -0.361624 | -0.192517 | -0.262372 | -0.281824 | -0.429626 | -0.005339 | -0.048482 | -0.343171 | 0.121822 | -0.352626 | -0.232178 | | 20.54167 | -0.117773 | -0.327261 |
| 20.458 | 0.266909 | 0.147663 | -0.255552 | 0.084937 | 0.047675 | -0.18027 | 0.200787 | -0.075449 | -0.043946 | 0.023596 | 0.103084 | 0.107637 | -0.22497 | -1.037148 | -0.88272 | -1.115505 | -1.075917 | -1.182812 | -0.837041 | -0.753274 | -1.083568 | -0.63673 | -0.919849 | -0.471953 | | 20.55556 | -0.22497 | -0.793478 |
| 20.472 | 0.30396 | 0.103321 | -0.203972 | 0.085186 | 0.03733 | -0.121106 | 0.151465 | 0.025348 | 0.003738 | 0.013141 | 0.066335 | 0.214529 | -0.216804 | -0.840266 | -0.76545 | -0.898337 | -0.901242 | -0.885755 | -0.642322 | -0.567905 | -0.841864 | -0.575769 | -0.622159 | -0.291455 | | 20.56944 | -0.216804 | -0.535567 |
| 20.486 | 0.326224 | 0.021676 | -0.043763 | 0.119205 | 0.021541 | 0.047956 | 0.19937 | 0.192053 | 0.087154 | 0.00191 | 0.007501 | 0.288603 | -0.084302 | 0.155527 | 0.095996 | 0.146482 | 0.112395 | 0.223941 | 0.481132 | 0.46922 | 0.197526 | 0.381209 | 0.310138 | 0.380539 | | 20.58333 | -0.084302 | 0.24716 |
| 20.5 | 0.357631 | 0.022469 | 0.007434 | 0.135014 | 0.015498 | 0.091719 | 0.150942 | 0.210529 | 0.096033 | 0.014837 | 0.040089 | 0.311146 | -0.047436 | 0.745816 | 0.561405 | 0.639118 | 0.621316 | 0.737548 | 0.977212 | 0.984825 | 0.746573 | 0.899088 | 0.73666 | 0.67152 | | 20.59722 | -0.047436 | 0.618067 |
| 20.514 | 0.373172 | 0.000248 | -0.042923 | 0.105201 | -0.030146 | 0.013583 | -0.059784 | 0.099129 | 0.032278 | -0.018129 | 0.051708 | 0.272799 | -0.144556 | 0.546855 | 0.340051 | 0.343509 | 0.363222 | 0.433794 | 0.613925 | 0.745676 | 0.482501 | 0.656869 | 0.428803 | 0.332119 | | 20.61111 | -0.144556 | 0.293745 |
| 20.528 | 0.285653 | -0.084493 | -0.060054 | 0.068085 | -0.095246 | -0.035663 | -0.081303 | 0.021957 | -0.018458 | -0.090114 | -0.001249 | 0.209497 | -0.205641 | 0.084462 | -0.067456 | -0.093324 | -0.07762 | 0.001346 | 0.091427 | 0.313129 | 0.006658 | 0.211699 | -0.052726 | -0.135174 | | 20.625 | -0.205641 | -0.139072 |
| 20.542 | 0.142417 | -0.146242 | -0.061486 | 0.026316 | -0.125949 | -0.053945 | 0.06074 | -0.031528 | -0.040884 | -0.127756 | -0.048282 | 0.175828 | -0.169258 | -0.247487 | -0.288658 | -0.289784 | -0.337296 | -0.213491 | -0.158093 | 0.000836 | -0.266508 | -0.094721 | -0.295 | -0.340057 | | 20.63889 | -0.169258 | -0.294983 |
| 20.556 | 0.039398 | -0.143466 | -0.076578 | -0.030407 | -0.122926 | -0.065596 | 0.020643 | -0.05421 | -0.04491 | -0.122503 | -0.0539 | 0.170588 | -0.106502 | -0.34523 | -0.281669 | -0.265949 | -0.375033 | -0.207373 | -0.185487 | -0.166431 | -0.265894 | -0.214606 | -0.316746 | -0.294798 | | 20.65278 | -0.106502 | -0.211366 |
| 20.569 | -0.015807 | -0.104436 | -0.081272 | -0.060097 | -0.100221 | -0.053939 | 0.004842 | -0.026096 | -0.032574 | -0.086688 | -0.033229 | 0.223576 | -0.062215 | -0.293488 | -0.16967 | -0.170256 | -0.278137 | -0.112442 | -0.161056 | -0.245782 | -0.146819 | -0.204363 | -0.268846 | -0.151242 | | 20.66667 | -0.062215 | -0.068578 |
| 20.583 | -0.024154 | -0.074825 | -0.072962 | -0.047543 | -0.074759 | -0.031631 | 0.026911 | 0.016289 | -0.012273 | -0.055634 | -0.01085 | 0.213835 | -0.041867 | -0.222355 | -0.1087 | -0.125893 | -0.20858 | -0.061285 | -0.198971 | -0.319026 | -0.1064 | -0.172353 | -0.273806 | -0.066343 | | 20.68056 | -0.041867 | -0.007297 |
| 20.597 | 0.000498 | -0.046467 | -0.0522 | -0.016032 | -0.053422 | -0.016047 | 0.056322 | 0.040356 | 0.005625 | -0.036375 | -0.000939 | 0.152707 | -0.041696 | -0.192675 | -0.146832 | -0.17472 | -0.218686 | -0.099982 | -0.275812 | -0.398153 | -0.158556 | -0.198641 | -0.282378 | -0.085092 | | 20.69444 | -0.041696 | -0.051947 |
| 20.611 | 0.003561 | -0.009954 | -0.030714 | 0.01589 | -0.0319 | -0.006141 | 0.076521 | 0.0298 | 0.016199 | -0.018759 | 0.009008 | 0.092768 | -0.026668 | -0.183744 | -0.193109 | -0.230451 | -0.245921 | -0.147953 | -0.300713 | -0.430995 | -0.20012 | -0.239323 | -0.180069 | -0.128535 | | 20.70833 | -0.026668 | -0.112771 |
| 20.625 | -0.023565 | -0.000105 | -0.022779 | 0.037219 | -0.028472 | -0.012561 | 0.061801 | -0.026477 | 0.000331 | -0.024591 | -0.001667 | 0.021932 | -0.004237 | -0.167384 | -0.175941 | -0.204517 | -0.206515 | -0.129479 | -0.23702 | -0.321054 | -0.196493 | -0.191649 | -0.072641 | -0.131019 | | 20.72222 | -0.004237 | -0.08651 |
| 20.639 | -0.044801 | -0.018169 | -0.018675 | 0.024985 | -0.043382 | -0.034239 | 0.015963 | -0.087292 | -0.033727 | -0.050568 | -0.032706 | -0.039987 | -0.005182 | -0.177325 | -0.099232 | -0.106562 | -0.116699 | -0.033966 | -0.102802 | -0.188768 | -0.103636 | -0.091644 | -0.041549 | -0.058522 | | 20.73611 | -0.005182 | 0.028356 |
| 20.653 | -0.043747 | -0.033395 | 0.001265 | -0.009129 | -0.050218 | -0.042407 | -0.012484 | -0.097314 | -0.053723 | -0.060007 | -0.05175 | -0.076535 | -0.016658 | -0.173031 | -0.009616 | -0.017612 | -0.035769 | 0.096852 | 0.033577 | -0.101484 | -0.001203 | -0.069288 | -0.048532 | 0.060858 | | 20.75 | -0.016658 | 0.125003 |
| 20.667 | -0.026674 | -0.026718 | 0.006988 | -0.037255 | -0.028168 | -0.023832 | 0.008375 | -0.045294 | -0.045726 | -0.035385 | -0.037805 | -0.09188 | -0.017764 | -0.136638 | 0.045634 | 0.013538 | -0.007267 | 0.182037 | 0.097871 | -0.055123 | 0.062317 | -0.155262 | -0.050317 | 0.162293 | | 20.76389 | -0.017764 | 0.194747 |
| 20.681 | -0.000339 | -0.002869 | -0.016088 | -0.043046 | 0.002552 | -0.001778 | 0.030548 | -0.000803 | -0.020141 | -0.003715 | -0.000264 | -0.09138 | 0.00006 | -0.116171 | 0.09295 | 0.02791 | 0.000693 | 0.207216 | 0.114044 | -0.017669 | 0.023497 | -0.197725 | -0.025225 | 0.194826 | | 20.77778 | 0.00006 | 0.212983 |
| 20.694 | 0.017588 | 0.011698 | -0.04474 | -0.036231 | 0.020442 | 0.015226 | 0.018266 | -0.008345 | 0.006264 | 0.012998 | 0.036971 | -0.09428 | 0.036866 | -0.141554 | 0.1571 | 0.093621 | 0.055685 | 0.259381 | 0.166383 | 0.060275 | 0.042732 | -0.032983 | 0.062794 | 0.226579 | | 20.79167 | 0.036866 | 0.225254 |
| 20.708 | -0.002333 | 0.005134 | -0.025287 | -0.020636 | 0.028934 | 0.030732 | -0.042202 | -0.083298 | 0.028457 | 0.022063 | 0.040425 | -0.10193 | 0.077426 | -0.190299 | 0.185079 | 0.183708 | 0.118103 | 0.321948 | 0.217687 | 0.143011 | 0.146191 | 0.165931 | 0.170935 | 0.268205 | | 20.80556 | 0.077426 | 0.269389 |
| 20.722 | -0.034898 | -0.045045 | 0.028731 | -0.054025 | 0.024706 | 0.022721 | -0.147162 | -0.221168 | 0.028574 | 0.016998 | 0.005774 | -0.170021 | 0.11423 | -0.22572 | 0.160851 | 0.206624 | 0.071939 | 0.315534 | 0.149365 | 0.132884 | 0.147566 | 0.193731 | 0.19114 | 0.302259 | | 20.81944 | 0.11423 | 0.325807 |
| 20.736 | -0.054481 | -0.12368 | 0.101045 | -0.083229 | 0.009621 | -0.00708 | -0.234108 | -0.311226 | 0.000724 | -0.002298 | -0.031391 | -0.28272 | 0.143594 | -0.169332 | 0.117549 | 0.16071 | -0.045027 | 0.24308 | -0.025318 | 0.021465 | 0.116787 | 0.081329 | 0.137906 | 0.322885 | | 20.83333 | 0.143594 | 0.34818 |
| 20.75 | -0.124317 | -0.194337 | 0.119318 | -0.115901 | -0.013075 | -0.021538 | -0.243623 | -0.266453 | -0.0411 | -0.028438 | -0.071819 | -0.429472 | 0.11994 | -0.121961 | 0.046138 | 0.064063 | -0.156661 | 0.146448 | -0.269983 | -0.184647 | 0.05544 | -0.092565 | 0.037623 | 0.2464 | | 20.84722 | 0.11994 | 0.280725 |
| 20.764 | -0.222815 | -0.226438 | 0.038782 | -0.095196 | -0.025526 | -0.015471 | -0.397519 | -0.191994 | -0.072186 | -0.043389 | -0.102002 | -0.560839 | 0.074174 | -0.160224 | -0.075642 | -0.076017 | -0.20458 | 0.056342 | -0.579133 | -0.558419 | -0.08015 | -0.371645 | -0.112311 | 0.062578 | | 20.86111 | 0.074174 | 0.09251 |
| 20.778 | -0.311733 | -0.228276 | -0.115307 | -0.153571 | -0.031569 | -0.026596 | -1.327366 | -0.156813 | -0.100275 | -0.039425 | -0.108161 | -0.621575 | 0.054218 | -0.302021 | -0.191891 | -0.200879 | -0.220606 | 0.044728 | -0.781608 | -1.0314 | -0.370798 | -0.63834 | -0.239694 | -0.120216 | | 20.875 | 0.054218 | -0.121253 |
| 20.792 | -0.37522 | -0.203883 | -0.238888 | -0.306872 | -0.045611 | -0.053601 | -0.625985 | -0.19581 | -0.108771 | -0.025479 | -0.162598 | -0.596403 | 0.046649 | -0.497381 | -0.25814 | -0.260613 | -0.236803 | 0.147313 | -0.756097 | -1.000029 | -0.632198 | -0.690444 | -0.306507 | -0.238972 | | 20.88889 | 0.046649 | -0.23753 |
| 20.806 | -0.40302 | -0.146966 | -0.221727 | -0.322388 | -0.049451 | -0.061848 | 0.004295 | -0.184402 | -0.105357 | -0.00891 | -0.250291 | -0.524943 | 0.073217 | -0.704538 | -0.335 | -0.312927 | -0.37376 | 0.295227 | -0.646855 | -0.815785 | -0.691605 | -0.739944 | -0.375145 | -0.320774 | | 20.90278 | 0.073217 | -0.338695 |
| 20.819 | -0.450624 | -0.130631 | -0.115465 | -0.217747 | -0.071753 | -0.068733 | 0.157661 | -0.17226 | -0.14819 | -0.033894 | -0.339461 | -0.485725 | 0.08657 | -0.87868 | -0.473797 | -0.412368 | -0.628427 | 0.3036 | -0.605996 | -0.892319 | -0.688064 | -0.830575 | -0.486953 | -0.452808 | | 20.91667 | 0.08657 | -0.486623 |
| 20.833 | -0.483973 | -0.118092 | 0.02832 | -0.149278 | -0.088671 | -0.068067 | -0.096698 | -0.169467 | -0.138163 | -0.055543 | -0.359236 | -0.456916 | 0.069989 | -1.012952 | -0.584607 | -0.473968 | -0.830445 | 0.0417 | -0.574323 | -1.056858 | -0.695392 | -0.90859 | -0.565555 | -0.57941 | | 20.93056 | 0.069989 | -0.636794 |
| 20.847 | -0.443949 | -0.064653 | 0.123176 | -0.139831 | -0.06845 | -0.056219 | -0.287623 | -0.130118 | -0.088389 | -0.026462 | -0.246389 | -0.352512 | 0.048809 | -1.162932 | -0.573792 | -0.443129 | -0.795766 | -0.151862 | -0.559476 | -1.081906 | -0.731022 | -0.894971 | -0.533361 | -0.599142 | | 20.94444 | 0.048809 | -0.707898 |
| 20.861 | -0.421067 | -0.060827 | 0.130557 | -0.213745 | -0.069457 | -0.053418 | -0.541922 | -0.088235 | -0.068251 | -0.016091 | -0.118207 | -0.229381 | 0.011883 | -1.278574 | -0.517346 | -0.368748 | -0.619572 | -0.201042 | -0.579417 | -0.949788 | -0.863964 | -0.749999 | -0.441981 | -0.539763 | | 20.95833 | 0.011883 | -0.678655 |
| 20.875 | -0.373814 | -0.099223 | 0.137789 | -0.221692 | -0.072692 | -0.03554 | -0.996947 | -0.065282 | -0.068688 | -0.008042 | -0.06715 | -0.102323 | 0.011567 | -1.088795 | -0.409523 | -0.27025 | -0.430063 | -0.163751 | -0.569728 | -0.747258 | -1.197913 | -0.491629 | -0.324584 | -0.409132 | | 20.97222 | 0.011567 | -0.577518 |
| 20.889 | -0.326985 | -0.114162 | 0.143789 | -0.176845 | -0.036616 | -0.024581 | -1.147318 | -0.062325 | -0.088536 | 0.023334 | -0.081141 | 0.009736 | 0.060151 | -0.646632 | -0.255681 | -0.231563 | -0.248908 | -0.064605 | -0.546786 | -0.602433 | -0.813185 | -0.253558 | -0.205479 | -0.256333 | | 20.98611 | 0.060151 | -0.404612 |
| 20.903 | -0.272956 | -0.129816 | 0.120471 | -0.139983 | -0.033385 | -0.024481 | -0.815375 | -0.085914 | -0.13991 | 0.023939 | -0.056008 | 0.036538 | 0.097002 | -0.29888 | -0.158655 | -0.195977 | -0.146388 | 0.055005 | -0.594169 | -0.481359 | -0.334932 | -0.222863 | -0.162228 | -0.148151 | | 21 | 0.097002 | -0.249493 |
| 20.917 | -0.131712 | -0.085029 | 0.146511 | -0.052389 | 0.023506 | 0.064912 | -0.43471 | 0.022465 | -0.111791 | 0.082714 | 0.149297 | 0.103874 | 0.13989 | -0.104011 | -0.056311 | -0.154679 | -0.079826 | 0.127733 | -0.557927 | -0.276745 | -0.018601 | -0.23275 | -0.123192 | -0.060743 | | 21.01389 | 0.13989 | -0.10365 |
| 20.931 | 0.092224 | 0.070617 | 0.186663 | 0.150314 | 0.19822 | 0.292852 | -0.071098 | 0.411899 | 0.072132 | 0.2673 | 0.414663 | 0.35126 | 0.23897 | 0.098606 | 0.128311 | -0.071057 | 0.027786 | 0.243358 | -0.161601 | 0.10515 | 0.022471 | -0.054077 | 0.016446 | -0.002649 | | 21.02778 | 0.23897 | 0.014763 |
| 20.944 | 0.15308 | 0.194168 | 0.222369 | 0.280978 | 0.29627 | 0.448598 | 0.122362 | 0.691654 | 0.228684 | 0.353901 | 0.173461 | 0.484846 | 0.314076 | 0.179878 | 0.130826 | -0.060548 | 0.055455 | 0.262168 | 0.248863 | 0.403576 | -0.280087 | -0.091684 | 0.073976 | -0.219544 | | 21.04167 | 0.314076 | -0.098288 |
| 20.958 | 0.070222 | 0.11249 | 0.197412 | 0.213154 | 0.208142 | 0.329555 | 0.022802 | 0.468736 | 0.214513 | 0.227305 | -0.231147 | 0.318473 | 0.276559 | 0.030315 | -0.099064 | -0.234902 | -0.05168 | 0.108463 | 0.322095 | 0.314859 | -0.568899 | -0.286863 | -0.101814 | -0.657774 | | 21.05556 | 0.276559 | -0.440554 |
| 20.972 | 0.003086 | -0.042237 | 0.167971 | 0.088562 | 0.08785 | 0.186927 | -0.242395 | -0.012814 | 0.09673 | 0.088979 | -0.309265 | 0.070789 | 0.223088 | -0.022349 | -0.09471 | -0.271355 | -0.04828 | 0.070742 | 0.396136 | 0.206169 | -0.636265 | -0.234091 | -0.230376 | -0.653179 | | 21.06944 | 0.223088 | -0.457649 |
| 20.986 | 0.025815 | -0.115125 | 0.205232 | 0.030584 | 0.03093 | 0.062328 | -0.704469 | -0.427082 | -0.0132 | 0.033927 | -0.532609 | -0.125427 | 0.157398 | 0.081234 | 0.076394 | -0.130639 | 0.082353 | 0.228078 | 0.156665 | 0.193503 | -0.318475 | -0.078693 | -0.078632 | -0.175691 | | 21.08333 | 0.157398 | -0.090353 |
| 21 | 0.031813 | -0.116931 | 0.094256 | -0.005441 | 0.01181 | -0.026167 | -1.212447 | -0.305331 | -0.057317 | 0.020912 | -0.56136 | -0.26012 | 0.068333 | 0.089661 | 0.103002 | -0.064104 | 0.096361 | 0.294718 | -0.443311 | 0.025854 | -0.002564 | -0.186788 | 0.138671 | 0.148371 | | 21.09722 | 0.068333 | 0.111149 |
| 21.014 | 0.032667 | -0.053753 | -0.007685 | 0.008956 | 0.011928 | -0.008711 | -0.982257 | 0.04124 | -0.008953 | 0.02088 | -0.011134 | -0.259978 | 0.039317 | 0.071839 | 0.102443 | -0.009239 | -0.035972 | 0.235348 | -0.583588 | -0.042716 | 0.177832 | -0.163572 | 0.243789 | 0.188216 | | 21.11111 | 0.039317 | 0.139987 |
| 21.028 | 0.026907 | 0.066628 | -0.031378 | 0.016106 | 0.004216 | 0.020892 | -0.37451 | 0.223426 | 0.024929 | 0.005582 | 0.646182 | -0.189601 | 0.044424 | 0.113326 | 0.17256 | 0.065794 | -0.060504 | 0.180326 | -0.329868 | 0.17093 | 0.169627 | 0.16576 | 0.264169 | 0.217203 | | 21.125 | 0.044424 | 0.228201 |
| 21.042 | 0.03218 | 0.206781 | -0.008758 | -0.03591 | 0.004096 | 0.010772 | 0.15113 | 0.135883 | -0.015291 | -0.001172 | 0.92244 | -0.105866 | 0.04635 | 0.092244 | 0.166541 | 0.141726 | -0.032789 | 0.131421 | -0.260983 | 0.271805 | 0.077674 | 0.432288 | 0.178414 | 0.246882 | | 21.13889 | 0.04635 | 0.282808 |
| 21.056 | 0.04689 | 0.25868 | 0.081213 | -0.087763 | 0.011339 | -0.025853 | 0.485155 | 0.06181 | -0.094813 | 0.006745 | 0.610789 | -0.026513 | 0.03684 | 0.026695 | 0.041681 | 0.131125 | -0.051712 | 0.053544 | -0.32554 | 0.031447 | -0.037711 | 0.4008 | 0.063548 | 0.160547 | | 21.15278 | 0.03684 | 0.185608 |
| 21.069 | 0.002101 | 0.106797 | -0.00224 | -0.067476 | -0.011798 | -0.069388 | 0.42062 | 0.086618 | -0.122508 | -0.014969 | 0.064258 | 0.006466 | 0.020488 | 0.060193 | -0.059429 | 0.015197 | -0.099393 | -0.021111 | -0.23518 | -0.244398 | -0.139297 | 0.094162 | -0.024172 | -0.072779 | | 21.16667 | 0.020488 | -0.049122 |
| 21.083 | -0.025712 | -0.085731 | -0.038936 | 0.012613 | -0.033178 | -0.116057 | 0.267797 | 0.172193 | -0.0725 | -0.038049 | -0.46499 | 0.024151 | 0.020056 | 0.214061 | -0.040718 | -0.076363 | -0.142681 | -0.050116 | 0.014031 | -0.199826 | -0.13928 | -0.431371 | -0.073648 | -0.224135 | | 21.18056 | 0.020056 | -0.22068 |
| 21.097 | 0.038447 | -0.071282 | -0.020487 | 0.058976 | -0.019493 | -0.105857 | 0.388511 | 0.209272 | -0.009319 | -0.020936 | -0.5305 | 0.081734 | 0.003878 | 0.278388 | 0.008354 | -0.084063 | -0.202987 | -0.093963 | 0.069954 | -0.130445 | -0.006923 | -0.710385 | -0.107471 | -0.207274 | | 21.19444 | 0.003878 | -0.215061 |
| 21.111 | 0.141854 | 0.087654 | -0.004771 | 0.077372 | 0.004422 | -0.00212 | 0.768086 | 0.086932 | 0.019852 | 0.006866 | -0.007988 | 0.165545 | -0.009936 | 0.194711 | -0.048247 | -0.053389 | -0.285032 | -0.155249 | -0.045677 | -0.23424 | 0.131864 | -0.540699 | -0.134618 | -0.157361 | | 21.20833 | -0.009936 | -0.162155 |
| 21.125 | 0.197918 | 0.183823 | 0.00196 | 0.102663 | 0.033234 | 0.109045 | 0.494238 | -0.002848 | 0.027378 | 0.029372 | 0.381586 | 0.251468 | 0.055616 | 0.191138 | -0.104926 | 0.005961 | -0.247221 | -0.135169 | 0.023349 | -0.224568 | 0.186281 | -0.274629 | -0.084354 | -0.079488 | | 21.22222 | 0.055616 | -0.037289 |
| 21.139 | 0.204981 | 0.124981 | -0.001506 | 0.132172 | 0.045603 | 0.071916 | -0.235119 | 0.086272 | 0.018722 | 0.032624 | 0.174487 | 0.248017 | 0.144851 | 0.21598 | -0.051598 | 0.038784 | -0.13868 | -0.085871 | 0.168096 | 0.003911 | 0.209196 | -0.020151 | -0.009195 | 0.050117 | | 21.23611 | 0.144851 | 0.120677 |
| 21.153 | 0.192918 | 0.033349 | -0.039537 | 0.119867 | 0.027311 | -0.017457 | -0.353115 | 0.130529 | 0.004067 | 0.010905 | -0.146648 | 0.165486 | 0.176514 | 0.083226 | 0.013013 | 0.002238 | -0.010153 | -0.084606 | 0.093329 | 0.218139 | 0.226575 | 0.126148 | 0.006424 | 0.144408 | | 21.25 | 0.176514 | 0.179746 |
| 21.167 | 0.209183 | 0.027805 | -0.045715 | 0.103183 | 0.023239 | -0.048486 | 0.123691 | 0.128409 | 0.008267 | 0.007586 | -0.123392 | 0.12737 | 0.12686 | -0.025847 | 0.008096 | -0.048917 | 0.152811 | -0.076784 | -0.062125 | 0.19507 | 0.214693 | 0.233719 | -0.012492 | 0.135687 | | 21.26389 | 0.12686 | 0.119929 |
| 21.181 | 0.229324 | 0.036429 | 0.012116 | 0.14048 | 0.036057 | -0.026583 | 0.57171 | 0.23364 | 0.030962 | 0.022282 | -0.008636 | 0.146064 | 0.041134 | -0.034537 | -0.065202 | -0.080799 | 0.291965 | -0.060574 | -0.077311 | 0.029184 | 0.154631 | 0.25172 | -0.042621 | 0.066269 | | 21.27778 | 0.041134 | 0.054559 |
| 21.194 | 0.199759 | 0.004419 | 0.028938 | 0.168932 | 0.042961 | -0.012787 | 0.6015 | 0.289554 | 0.045439 | 0.029795 | -0.122312 | 0.148481 | -0.032133 | -0.013031 | -0.139942 | -0.083911 | 0.339263 | -0.07535 | -0.03788 | 0.025606 | 0.165367 | 0.120586 | -0.044769 | 0.057999 | | 21.29167 | -0.032133 | 0.078177 |
| 21.208 | 0.089897 | -0.015333 | -0.024235 | 0.125554 | 0.033376 | -0.019022 | 0.161552 | 0.186548 | 0.034656 | 0.022349 | -0.240962 | 0.113933 | -0.039648 | 0.045441 | -0.101689 | -0.059757 | 0.310553 | -0.09901 | 0.020916 | 0.114835 | 0.163868 | -0.033527 | -0.029198 | 0.056579 | | 21.30556 | -0.039648 | 0.089587 |
| 21.222 | -0.037934 | -0.016798 | -0.049805 | 0.081094 | -0.012128 | -0.036202 | -0.210525 | 0.050179 | 0.004022 | -0.019879 | -0.131307 | 0.060494 | -0.01822 | 0.076872 | -0.009337 | -0.059749 | 0.187252 | -0.150751 | 0.093793 | 0.028105 | 0.005277 | -0.145694 | -0.070337 | -0.052026 | | 21.31944 | -0.01822 | -0.009632 |
| 21.236 | -0.058893 | 0.046289 | 0.00703 | 0.092511 | -0.02103 | 0.013035 | -0.236061 | 0.030191 | 0.0168 | -0.017338 | 0.085534 | 0.063279 | 0.003517 | 0.11236 | 0.03914 | -0.017964 | 0.134958 | -0.096543 | 0.169668 | -0.030378 | -0.017565 | -0.041399 | -0.054208 | -0.033549 | | 21.33333 | 0.003517 | 0.009072 |
| 21.25 | 0.018939 | 0.113384 | 0.07026 | 0.103403 | 0.029402 | 0.125151 | 0.08308 | 0.131931 | 0.079254 | 0.044695 | 0.270127 | 0.094322 | 0.06572 | 0.196286 | 0.032116 | 0.116562 | 0.185991 | 0.0415 | 0.322193 | 0.203332 | 0.257017 | 0.242587 | 0.075406 | 0.196761 | | 21.34722 | 0.06572 | 0.241482 |
| 21.264 | 0.058718 | 0.080591 | 0.066231 | 0.064416 | 0.033482 | 0.115815 | 0.514862 | 0.179453 | 0.114604 | 0.047076 | 0.258522 | 0.066032 | 0.118693 | 0.234856 | 0.020314 | 0.146148 | 0.171508 | 0.076362 | 0.430124 | 0.388249 | 0.497822 | 0.353439 | 0.093505 | 0.323067 | | 21.36111 | 0.118693 | 0.380137 |
| 21.278 | 0.079689 | 0.084114 | 0.050053 | 0.05066 | 0.016883 | 0.025115 | 0.908489 | 0.161734 | 0.133082 | 0.025046 | 0.15303 | 0.084856 | 0.124168 | 0.266957 | 0.033609 | 0.059998 | 0.118587 | -0.007065 | 0.366006 | 0.341144 | 0.463201 | 0.224035 | 0.007989 | 0.279773 | | 21.375 | 0.124168 | 0.330665 |

| | | | | | | | | | | | | | | | | | | | | | | | | | | | | |
|---|---|---|---|---|---|---|---|---|---|---|---|---|---|---|---|---|---|---|---|---|---|---|---|---|---|---|---|---|
| 21.292 | 0.115876 | 0.171859 | 0.027695 | 0.069975 | 0.028603 | 0.012993 | 1.023868 | 0.21336 | 0.153446 | 0.03717 | 0.23221 | 0.18992 | 0.128881 | 0.385092 | 0.05293 | 0.044628 | 0.169726 | -0.094841 | 0.23594 | 0.31926 | 0.433263 | 0.117937 | 0.00626 | 0.230625 | | 21.38889 | 0.128881 | 0.291646 |
| 21.306 | 0.116818 | 0.224174 | -0.053602 | 0.075322 | 0.019295 | 0.046264 | 0.777266 | 0.311755 | 0.149754 | 0.023973 | 0.469216 | 0.285456 | 0.103733 | 0.521256 | 0.048534 | 0.094393 | 0.265913 | -0.120605 | 0.233776 | 0.477324 | 0.540648 | 0.180264 | 0.057461 | 0.273427 | | 21.40278 | 0.103733 | 0.379435 |
| 21.319 | 0.162734 | 0.246842 | -0.052142 | 0.103664 | 0.032982 | 0.095308 | 0.480216 | 0.382578 | 0.151915 | 0.024384 | 0.666437 | 0.40742 | 0.034741 | 0.6758 | 0.143354 | 0.119378 | 0.333432 | 0.219872 | 0.30637 | 0.735213 | 0.713979 | 0.375292 | 0.08237 | 0.39913 | | 21.41667 | 0.034741 | 0.539839 |
| 21.333 | 0.289276 | 0.258812 | 0.044252 | 0.16902 | 0.083907 | 0.141338 | 0.310695 | 0.408808 | 0.192713 | 0.061484 | 0.715443 | 0.589228 | 0.012485 | 0.714859 | 0.253581 | 0.081928 | 0.351019 | 0.05048 | 0.330638 | 0.985688 | 0.747106 | 0.554514 | 0.015591 | 0.40683 | | 21.43056 | 0.012485 | 0.51875 |
| 21.347 | 0.398876 | 0.15964 | 0.061386 | 0.19806 | 0.087534 | 0.129958 | 0.318151 | 0.377769 | 0.220618 | 0.039834 | 0.508481 | 0.720362 | 0.010914 | 0.59287 | 0.238648 | 0.067435 | 0.327495 | -0.049992 | 0.281397 | 1.003337 | 0.637435 | 0.642594 | -0.076377 | 0.256744 | | 21.44444 | 0.010914 | 0.339649 |
| 21.361 | 0.495886 | 0.021635 | -0.022345 | 0.069521 | 0.009484 | 0.017777 | 0.439775 | 0.148266 | 0.137268 | -0.057198 | 0.105668 | 0.635995 | -0.061704 | 0.558889 | 0.241403 | 0.204956 | 0.395842 | 0.042187 | 0.399057 | 0.675948 | 0.677939 | 0.685644 | -0.044257 | 0.244296 | | 21.45833 | -0.061704 | 0.315767 |
| 21.375 | 0.496505 | -0.061488 | -0.141188 | -0.134839 | -0.097102 | -0.184554 | 0.671688 | -0.241343 | 0.018218 | -0.176727 | -0.287625 | 0.383281 | -0.102458 | 0.676399 | 0.363662 | 0.437114 | 0.545876 | 0.272168 | 0.701166 | 0.335349 | 0.807065 | 0.829213 | 0.140398 | 0.332477 | | 21.47222 | -0.102458 | 0.395928 |
| 21.389 | 0.217478 | -0.004802 | -0.243468 | -0.155365 | -0.106214 | -0.28546 | 1.133854 | -0.314951 | 0.016865 | -0.185762 | -0.332544 | 0.138 | 0.163037 | 0.798326 | 0.366339 | 0.525548 | 0.597871 | 0.504711 | 1.0045 | 0.189432 | 0.854998 | 0.704662 | 0.361937 | 0.325837 | | 21.48611 | 0.163037 | 0.358095 |
| 21.403 | 0.201944 | 0.068501 | -0.327172 | -0.028981 | -0.018598 | -0.122489 | 1.678857 | 0.041396 | 0.141912 | -0.07256 | -0.153136 | 0.242739 | 0.403105 | 0.809149 | 0.338662 | 0.443336 | 0.499601 | 0.59419 | 1.115568 | 0.389976 | 0.591456 | 0.728981 | 0.341503 | 0.205336 | | 21.5 | 0.403105 | 0.225431 |
| 21.417 | 0.227286 | -0.145711 | -0.222216 | 0.120463 | -0.013211 | 0.024705 | 1.512442 | 0.297914 | 0.26971 | -0.046324 | 0.084424 | 0.436446 | -0.115692 | 0.639754 | 0.336877 | 0.319944 | 0.383525 | 0.560784 | 0.892821 | 0.611429 | 0.370776 | 0.739298 | 0.243749 | 0.088807 | | 21.51389 | -0.115692 | 0.109079 |
| 21.431 | 0.235198 | -0.311693 | -0.07384 | 0.178573 | -0.053283 | 0.090305 | 0.74613 | 0.262929 | 0.223334 | -0.065827 | 0.146882 | 0.524408 | -0.688948 | 0.341137 | 0.220171 | 0.151713 | 0.288662 | 0.543767 | 0.611299 | 0.588576 | 0.265269 | 0.55313 | 0.179099 | 0.064131 | | 21.52778 | -0.688948 | 0.092459 |
| 21.444 | 0.250735 | -0.056645 | -0.013437 | 0.13708 | -0.002709 | 0.043396 | 0.258225 | 0.13105 | 0.146301 | 0.039753 | 0.185187 | 0.523429 | -0.508551 | -0.21462 | -0.096418 | -0.265598 | -0.151749 | 0.12397 | 0.148026 | 0.112492 | -0.133882 | -0.047985 | -0.130197 | -0.030988 | | 21.54167 | -0.508551 | -0.103795 |
| 21.458 | 0.191443 | 0.158211 | -0.096865 | 0.094946 | 0.002276 | -0.098426 | 0.036181 | -0.071146 | 0.131862 | 0.075655 | 0.185456 | 0.462329 | -0.340114 | -0.92673 | -0.768357 | -1.013609 | -0.935921 | -0.666723 | -0.577908 | -0.763185 | -0.919474 | -0.873248 | -0.739227 | -0.404481 | | 21.55556 | -0.340114 | -0.609009 |
| 21.472 | 0.083585 | 0.065834 | -0.134045 | 0.081988 | -0.069366 | -0.117018 | 0.003921 | -0.071769 | 0.109872 | 0.002297 | 0.10837 | 0.304946 | -0.33041 | -1.182263 | -1.215713 | -1.382415 | -1.295603 | -1.035453 | -0.902595 | -1.348625 | -1.247233 | -1.294854 | -1.063791 | -0.818616 | | 21.56944 | -0.33041 | -0.974687 |
| 21.486 | 0.08026 | -0.019608 | -0.071462 | 0.058123 | -0.057006 | 0.017725 | 0.23247 | 0.18263 | 0.155573 | -0.006028 | 0.022237 | 0.276868 | -0.146583 | -0.774683 | -0.994479 | -1.025538 | -0.962144 | -0.60709 | -0.576563 | -1.166966 | -0.942905 | -1.018373 | -0.602885 | -0.788907 | | 21.58333 | -0.146583 | -0.835807 |
| 21.5 | 0.16086 | -0.013432 | -0.079489 | 0.041979 | 0.004379 | 0.107197 | 0.482953 | 0.302766 | 0.202219 | 0.04208 | 0.039313 | 0.367677 | -0.030088 | -0.274943 | -0.524613 | -0.565598 | -0.498285 | -0.092363 | -0.147036 | -0.681906 | -0.582001 | -0.544103 | -0.160276 | -0.469291 | | 21.59722 | -0.030088 | -0.449478 |
| 21.514 | 0.21695 | 0.013882 | -0.139223 | 0.011508 | 0.031711 | 0.107016 | 0.452602 | 0.265046 | 0.167953 | 0.071003 | 0.067685 | 0.498001 | -0.040711 | 0.077194 | -0.161028 | -0.232041 | -0.165003 | 0.129005 | 0.198104 | -0.235627 | -0.290535 | -0.132967 | 0.050812 | -0.222825 | | 21.61111 | -0.040711 | -0.183986 |
| 21.528 | 0.206519 | 0.016735 | -0.114107 | 0.025353 | 0.031693 | 0.104086 | 0.45899 | 0.170654 | 0.124347 | 0.074662 | 0.077812 | 0.495015 | -0.031811 | 0.204071 | -0.011576 | -0.058924 | -0.023728 | 0.172696 | 0.369159 | 0.05507 | -0.09458 | 0.080795 | 0.070815 | -0.14594 | | 21.625 | -0.031811 | -0.103726 |
| 21.542 | 0.102208 | -0.007399 | -0.069785 | 0.073925 | 0.014301 | 0.071658 | 0.555699 | 0.062181 | 0.075087 | 0.054563 | 0.060276 | 0.339781 | -0.007809 | 0.098433 | -0.048408 | -0.065735 | -0.056529 | 0.00621 | 0.205035 | 0.177611 | -0.086556 | 0.078679 | -0.145042 | -0.203357 | | 21.63889 | -0.007809 | -0.173388 |
| 21.556 | -0.018395 | -0.009598 | -0.002439 | 0.135886 | 0.012254 | 0.046454 | 0.436291 | -0.020193 | 0.064003 | 0.044304 | 0.051389 | 0.158693 | 0.02541 | -0.017207 | -0.09864 | -0.128468 | -0.122931 | -0.154632 | 0.017169 | 0.254746 | -0.057109 | 0.017171 | -0.419898 | -0.214464 | | 21.65278 | 0.02541 | -0.225502 |
| 21.569 | -0.083814 | -0.000923 | 0.089552 | 0.162084 | 0.011931 | 0.039184 | 0.251441 | -0.065582 | 0.11957 | 0.033559 | 0.03617 | 0.054568 | 0.040907 | -0.022214 | -0.123632 | -0.174333 | -0.168405 | -0.19193 | -0.047241 | 0.208886 | -0.056474 | -0.032695 | -0.422432 | -0.131919 | | 21.66667 | 0.040907 | -0.203494 |
| 21.583 | -0.061098 | -0.008051 | 0.084371 | 0.128055 | -0.000111 | 0.028666 | 0.143276 | -0.091864 | 0.178258 | 0.016904 | 0.006818 | -0.030951 | 0.012197 | -0.037102 | -0.180563 | -0.240322 | -0.244434 | -0.101448 | -0.126003 | -0.002278 | -0.169992 | -0.12473 | -0.464895 | -0.116818 | | 21.68056 | 0.012197 | -0.205471 |
| 21.597 | 0.018491 | -0.033692 | 0.015164 | 0.039849 | -0.020549 | 0.005013 | 0.04391 | -0.097979 | 0.166896 | -0.005898 | -0.02383 | -0.06238 | -0.049064 | -0.214403 | -0.281289 | -0.342445 | -0.360667 | -0.029076 | -0.290776 | -0.268359 | -0.393679 | -0.28711 | -0.53696 | -0.25471 | | 21.69444 | -0.049064 | -0.283444 |
| 21.611 | 0.118035 | -0.035216 | -0.053542 | 0.009018 | -0.020457 | 0.002694 | -0.026832 | -0.054855 | 0.070909 | -0.017922 | -0.007053 | 0.002052 | -0.073473 | -0.347936 | -0.364147 | -0.415758 | -0.43266 | -0.012435 | -0.445813 | -0.476231 | -0.571409 | -0.397818 | -0.603558 | -0.378321 | | 21.70833 | -0.073473 | -0.353271 |
| 21.625 | 0.176581 | -0.013892 | -0.083717 | 0.028493 | 0.020525 | 0.030039 | -0.026112 | 0.022496 | -0.011293 | -0.007053 | 0.051336 | 0.116948 | -0.022304 | -0.266171 | -0.38764 | -0.398739 | -0.385536 | -0.139759 | -0.46565 | -0.579176 | -0.573444 | -0.369644 | -0.585733 | -0.3745 | | 21.72222 | -0.022304 | -0.326476 |
| 21.639 | 0.148367 | -0.018572 | -0.105534 | -0.003338 | 0.040822 | 0.036739 | -0.010414 | 0.054246 | -0.110398 | -0.00028 | 0.072768 | 0.206807 | 0.032713 | -0.215203 | -0.354383 | -0.321086 | -0.290535 | -0.321717 | -0.400233 | -0.568175 | -0.444664 | -0.234705 | -0.465765 | -0.327537 | | 21.73611 | 0.032713 | -0.2341 |
| 21.653 | 0.140177 | -0.00249 | 0.027161 | -0.025731 | 0.062169 | 0.045027 | 0.005905 | 0.07213 | -0.118047 | 0.047223 | 0.081426 | 0.202601 | 0.081987 | -0.16963 | -0.23967 | -0.178668 | -0.14943 | -0.34797 | -0.244611 | -0.314973 | -0.075543 | -0.025443 | -0.238737 | -0.185842 | | 21.75 | 0.081987 | -0.077996 |
| 21.667 | 0.204125 | 0.044995 | 0.155272 | 0.019334 | 0.094055 | 0.061573 | 0.054692 | 0.071747 | 0.128457 | 0.122022 | 0.107753 | 0.192051 | 0.169356 | -0.035098 | 0.066487 | 0.136046 | 0.137919 | -0.11005 | 0.157115 | 0.261134 | 0.331385 | 0.325939 | 0.156073 | 0.205513 | | 21.76389 | 0.169356 | 0.221332 |
| 21.681 | 0.235996 | 0.019041 | 0.187503 | 0.049183 | 0.07891 | 0.043493 | 0.077338 | 0.034275 | 0.245715 | 0.135099 | 0.081192 | 0.120147 | 0.196291 | 0.281228 | 0.566991 | 0.624125 | 0.597186 | 0.243159 | 0.783982 | 1.01455 | 0.773323 | 0.802377 | 0.734006 | 0.747825 | | 21.77778 | 0.196291 | 0.7111 |
| 21.694 | 0.223767 | -0.047427 | 0.146214 | 0.055678 | 0.041275 | 0.015066 | 0.08253 | -0.020887 | 0.191517 | 0.089762 | 0.033676 | 0.014028 | 0.13794 | 0.691745 | 0.994905 | 1.064999 | 0.996418 | 0.38662 | 1.302166 | 1.490737 | 0.958578 | 1.206977 | 1.231466 | 1.127878 | | 21.79167 | 0.13794 | 1.108359 |
| 21.708 | 0.229072 | -0.03174 | 0.094245 | 0.079709 | 0.051702 | 0.02115 | 0.079501 | -0.066185 | 0.088959 | 0.056039 | 0.071354 | -0.053428 | 0.115265 | 0.903634 | 1.136181 | 1.251002 | 1.116363 | 0.220622 | 1.429788 | 1.400019 | 0.88727 | 1.283865 | 1.367783 | 1.231301 | | 21.80556 | 0.115265 | 1.211048 |
| 21.722 | 0.162252 | 0.034104 | 0.043165 | 0.086067 | 0.070621 | 0.03479 | -0.0044 | -0.078469 | -0.031364 | -0.001079 | 0.153102 | -0.118917 | 0.136529 | 0.973825 | 1.135349 | 1.229889 | 1.061456 | 0.4247 | 1.250894 | 1.118028 | 0.79546 | 1.158784 | 1.307173 | 1.239042 | | 21.81944 | 0.136529 | 1.209194 |
| 21.736 | -0.000158 | 0.027525 | 0.011082 | 0.042612 | 0.045414 | 0.015367 | -0.173104 | -0.143693 | -0.213146 | -0.084641 | 0.243479 | -0.175743 | 0.105698 | 0.804803 | 1.007313 | 1.027008 | 0.84901 | 0.819818 | 0.866228 | 0.744469 | 0.755526 | 0.959485 | 1.086713 | 1.138468 | | 21.83333 | 0.105698 | 1.149082 |
| 21.75 | -0.193312 | -0.021771 | 0.018788 | -0.039935 | -0.006246 | -0.025052 | -0.303063 | -0.176872 | -0.280109 | -0.102967 | 0.24877 | -0.286697 | 0.059358 | 0.275255 | 0.631851 | 0.584804 | 0.416332 | 0.885116 | 0.274764 | 0.167038 | 0.657196 | 0.495353 | 0.586177 | 0.80972 | | 21.84722 | 0.059358 | 0.864394 |
| 21.764 | -0.365936 | -0.049721 | 0.040277 | -0.11674 | -0.050154 | -0.059492 | -0.994759 | -0.16789 | -0.19025 | -0.077504 | 0.054047 | -0.467875 | 0.043072 | -0.276651 | 0.195251 | 0.096098 | -0.077706 | 0.369955 | -0.283963 | -0.439211 | 0.211896 | -0.005337 | -0.048754 | 0.365784 | | 21.86111 | 0.043072 | 0.443202 |
| 21.778 | -0.463455 | -0.050255 | 0.007611 | -0.175434 | -0.079974 | -0.108489 | -1.753837 | -0.148656 | -0.081607 | -0.048919 | -0.039278 | -0.695838 | -0.00972 | -0.623984 | -0.22255 | -0.309185 | -0.4659 | -0.425684 | -0.649559 | -1.147543 | -0.352884 | -0.280058 | -0.61593 | -0.104337 | | 21.875 | -0.00972 | -0.057782 |
| 21.792 | -0.458703 | -0.037711 | -0.067251 | -0.217348 | -0.101889 | -0.136463 | 0.207548 | -0.251818 | -0.062172 | -0.052117 | -0.080546 | -0.522078 | -0.013421 | -0.768569 | -0.603463 | -0.601318 | -0.780318 | -1.053755 | -0.861683 | -1.439062 | -0.832101 | -0.427138 | -1.143713 | -0.571009 | | 21.88889 | -0.013421 | -0.54754 |
| 21.806 | -0.423255 | 0.003042 | -0.102281 | -0.229302 | -0.107601 | -0.114446 | 0.938434 | -0.388752 | -0.149864 | -0.082684 | -0.100726 | -0.086179 | 0.066009 | -1.049281 | -0.957508 | -0.881577 | -1.100214 | -1.422361 | -1.041636 | -1.34643 | -1.209409 | -0.725993 | -1.662328 | -0.989585 | | 21.90278 | 0.066009 | -1.02834 |
| 21.819 | -0.487234 | 0.033214 | -0.098249 | -0.239612 | -0.125185 | -0.084035 | 0.697714 | -0.339007 | -0.228868 | -0.142465 | -0.18328 | 0.081541 | 0.111096 | -1.259621 | -1.321446 | -1.151471 | -1.494645 | -1.639432 | -1.360114 | -1.529336 | -1.382542 | -1.176883 | -2.020277 | -1.307639 | | 21.91667 | 0.111096 | -1.382479 |
| 21.833 | -0.59627 | 0.05771 | -0.03421 | -0.218315 | -0.117375 | -0.049959 | 0.339394 | -0.118128 | -0.18754 | -0.159827 | -0.292723 | -0.174474 | 0.128303 | -1.335793 | -1.406019 | -1.186265 | -1.478128 | -1.659458 | -1.520462 | -1.591082 | -1.297426 | -1.385618 | -1.974695 | -1.274369 | | 21.93056 | 0.128303 | -1.350264 |
| 21.847 | -0.576371 | 0.047996 | 0.020176 | -0.217315 | -0.049942 | -0.013103 | 0.161038 | -0.089891 | -0.112786 | -0.052657 | -0.356607 | -0.605554 | 0.155642 | -1.2085 | -1.133192 | -0.954412 | -1.141013 | -1.521691 | -1.411909 | -1.343441 | -1.107759 | -1.242932 | -1.526366 | -0.934802 | | 21.94444 | 0.155642 | -0.990311 |
| 21.861 | -0.432121 | -0.046627 | 0.097166 | -0.243041 | 0.004556 | -0.004534 | -0.062874 | -0.212637 | -0.10622 | 0.018599 | -0.273229 | -0.968944 | 0.175541 | -1.076012 | -0.846061 | -0.668189 | -0.851248 | -1.168286 | -1.30199 | -1.060532 | -0.984238 | -1.089962 | -1.029499 | -0.636522 | | 21.95833 | 0.175541 | -0.677283 |
| 21.875 | -0.230663 | -0.12374 | 0.217205 | -0.235243 | 0.045432 | -0.010816 | -0.393565 | -0.319967 | -0.232003 | 0.043716 | -0.319224 | -0.870977 | 0.209201 | -0.971427 | -0.625121 | -0.440269 | -0.671613 | -0.912241 | -1.209146 | -0.979043 | -0.935112 | -1.084964 | -0.687573 | -0.506622 | | 21.97222 | 0.209201 | -0.551463 |
| 21.889 | -0.108061 | -0.090258 | 0.18102 | -0.219717 | 0.094301 | -0.014533 | -0.682856 | -0.25747 | -0.208487 | 0.074359 | -0.790032 | -0.689281 | 0.23529 | -0.664254 | -0.445367 | -0.229345 | -0.445713 | -0.979524 | -1.066405 | -1.006183 | -0.822737 | -0.998715 | -0.435169 | -0.487753 | | 21.98611 | 0.23529 | -0.562543 |
| 21.903 | -0.079199 | -0.03971 | 0.099228 | -0.198914 | 0.108184 | 0.014244 | -0.710917 | -0.179551 | -0.000071 | 0.083461 | -0.942192 | -0.472784 | 0.235896 | -0.400309 | -0.424413 | -0.113098 | -0.217459 | -1.271503 | -0.823398 | -0.84793 | -0.653102 | -0.943134 | -0.323029 | -0.476482 | | 22 | 0.235896 | -0.58303 |
| 21.917 | -0.000372 | 0.078339 | 0.117393 | -0.096595 | 0.14493 | 0.118137 | -0.619748 | -0.008827 | 0.124413 | 0.147785 | -0.662916 | 0.275049 | 0.275208 | -0.215737 | -0.431089 | -0.168957 | -0.061928 | -1.2979 | -0.586263 | -0.507627 | -0.522023 | -0.92315 | -0.357597 | -0.432242 | | 22.01389 | 0.275208 | -0.551717 |
| 21.931 | 0.135207 | 0.189757 | 0.2114 | 0.069788 | 0.266308 | 0.300651 | -0.327861 | 0.243315 | 0.182254 | 0.303473 | -0.023297 | 1.299388 | 0.38203 | -0.179731 | -0.375108 | -0.231891 | -0.035413 | -1.057321 | -0.267155 | -0.214576 | -0.352198 | -0.785943 | -0.4898 | -0.433074 | | 22.02778 | 0.38203 | -0.538902 |
| 21.944 | 0.174501 | 0.100463 | 0.275669 | 0.189663 | 0.286053 | 0.413123 | -0.225818 | 0.38716 | 0.133645 | 0.345689 | 0.305956 | 1.45974 | 0.437273 | -0.314451 | -0.439747 | -0.362174 | -0.114492 | -0.763075 | -0.270348 | -0.233334 | -0.346305 | -0.786196 | -0.564144 | -0.569487 | | 22.04167 | 0.437273 | -0.625847 |
| 21.958 | 0.150919 | -0.144245 | 0.261421 | 0.145222 | 0.13398 | 0.314402 | -0.625284 | 0.007895 | 0.146088 | 0.192142 | -0.001054 | 0.437142 | 0.366559 | 0.035948 | -0.291556 | -0.22681 | 0.086706 | -0.593342 | -0.406653 | -0.034434 | -0.166141 | -0.339774 | -0.188099 | -0.417474 | | 22.05556 | 0.366559 | -0.41321 |
| 21.972 | 0.264575 | -0.338814 | 0.34815 | -0.048207 | -0.005203 | 0.186607 | -1.343882 | -0.599574 | 0.116743 | -0.038697 | -0.318746 | -0.304947 | 0.254078 | 0.965361 | 0.254142 | 0.231178 | 0.585311 | -0.425872 | -0.122604 | 0.698584 | 0.291648 | 0.72487 | 0.66747 | 0.336554 | | 22.06944 | 0.254078 | 0.397663 |
| 21.986 | 0.252466 | -0.551265 | 0.390636 | -0.146231 | -0.066828 | 0.076199 | -2.207673 | -0.833029 | -0.038393 | -0.083494 | -0.95322 | -0.811661 | 0.124563 | 0.918274 | 0.646553 | 0.41301 | 0.658255 | -0.320288 | -0.148299 | 1.098452 | 0.378926 | 1.501244 | 1.187497 | 0.862205 | | 22.08333 | 0.124563 | 0.844473 |
| 22 | -0.03769 | -0.476377 | 0.270396 | -0.130139 | -0.028545 | -0.065042 | -2.278052 | -0.493517 | -0.10958 | 0.044021 | -1.57047 | -0.611325 | 0.022904 | -0.04613 | 0.594741 | 0.320502 | 0.450672 | -0.040958 | -0.110998 | 0.723956 | 0.094461 | 0.66124 | 0.886773 | 0.549943 | | 22.09722 | 0.022904 | 0.402659 |
| 22.014 | -0.163415 | -0.042896 | 0.11477 | -0.048181 | 0.043214 | -0.080829 | -1.123428 | 0.079786 | 0.011745 | 0.194244 | -0.940432 | 0.161346 | 0.008607 | -0.369736 | 0.494021 | 0.481439 | 0.591316 | 0.294807 | 0.388517 | 0.435372 | 0.34205 | -0.197061 | 0.533872 | 0.212966 | | 22.11111 | 0.008607 | 0.070966 |
| 22.028 | 0.179268 | 0.338038 | 0.068791 | 0.026263 | 0.065972 | -0.019521 | -0.308556 | 0.444108 | 0.120336 | 0.200378 | 0.109745 | 0.641263 | 0.097084 | -0.122736 | 0.514456 | 0.629 | 0.795001 | 0.599232 | 0.923422 | 0.461788 | 0.732029 | -0.360141 | 0.611551 | 0.307878 | | 22.125 | 0.097084 | 0.217551 |
| 22.042 | 0.477365 | 0.345684 | 0.053818 | 0.060195 | 0.063202 | 0.083 | 0.254474 | 0.419767 | 0.129764 | 0.043849 | 0.696817 | 0.243959 | 0.147749 | -0.022502 | 0.619211 | 0.831619 | 0.694101 | 0.679343 | 1.067566 | 0.464321 | 1.007445 | -0.05219 | 0.829948 | 0.429743 | | 22.13889 | 0.147749 | 0.437227 |
| 22.056 | 0.205828 | -0.039141 | 0.03733 | 0.055984 | 0.084686 | 0.12388 | 0.531737 | 0.222734 | -0.003726 | -0.060858 | 0.514123 | -0.69378 | 0.135834 | 0.008127 | 0.77271 | 0.995859 | 0.638962 | 0.677203 | 0.766041 | 0.335004 | 1.123756 | 0.493195 | 0.83691 | 0.565727 | | 22.15278 | 0.135834 | 0.662203 |
| 22.069 | -0.189629 | -0.288677 | 0.001381 | 0.069244 | 0.082277 | 0.046788 | 0.491012 | 0.134058 | -0.139045 | 0.016484 | 0.151256 | -0.851035 | 0.107617 | 0.387 | 0.888455 | 0.995341 | 0.867969 | 1.138781 | 0.772804 | 0.279015 | 1.15408 | 1.03628 | 0.810604 | 0.811012 | | 22.16667 | 0.107617 | 0.953879 |
| 22.083 | -0.150732 | -0.18361 | -0.022338 | 0.086784 | 0.025384 | -0.009235 | 0.275543 | 0.116012 | -0.010385 | 0.121458 | 0.038712 | -0.052629 | 0.113431 | 0.895494 | 0.84462 | 0.862852 | 0.907052 | 2.120393 | 1.087621 | 0.424369 | 0.964099 | 1.151946 | 0.847745 | 0.922838 | | 22.18056 | 0.113431 | 1.054192 |
| 22.097 | 0.087835 | -0.08383 | -0.037739 | 0.174676 | -0.004996 | 0.054794 | 0.150956 | 0.079499 | 0.232106 | 0.136345 | 0.241591 | 0.686952 | 0.17689 | 1.111971 | 0.72776 | 0.708714 | 0.599839 | 2.082538 | 1.078529 | 0.492584 | 0.601559 | 0.742833 | 0.824964 | 0.777341 | | 22.19444 | 0.17689 | 0.709256 |
| 22.111 | 0.130398 | 0.100089 | -0.015333 | 0.250875 | 0.019601 | 0.0974 | 0.233009 | 0.039437 | 0.201274 | 0.072472 | 0.48973 | 0.543219 | 0.226245 | 0.9935 | 0.712063 | 0.654393 | 0.594077 | 1.376876 | 0.839586 | 0.435289 | -0.07902 | 0.374384 | 0.731508 | 0.631598 | | 22.20833 | 0.226245 | 0.461398 |
| 22.125 | 0.100564 | 0.381928 | -0.01895 | 0.230572 | 0.022835 | 0.040134 | 0.523601 | 0.073735 | -0.019694 | 0.024928 | 0.456614 | 0.241863 | 0.192457 | 0.482093 | 0.755623 | 0.586221 | 0.664498 | 1.112533 | 0.706799 | 0.460338 | 0.215948 | 0.281382 | 0.596516 | 0.652148 | | 22.22222 | 0.192457 | 0.603809 |
| 22.139 | 0.182195 | 0.322991 | -0.091153 | 0.193323 | -0.013544 | 0.028297 | 0.942238 | 0.181651 | -0.012665 | 0.039078 | 0.416475 | 0.07379 | 0.126904 | -0.048653 | 0.597906 | 0.37192 | 0.310294 | 1.207037 | 0.522611 | 0.781168 | 1.008796 | 0.346976 | 0.362282 | 0.608002 | | 22.23611 | 0.126904 | 0.78041 |
| 22.153 | 0.235976 | -0.02453 | -0.182678 | 0.183968 | -0.027426 | 0.101072 | 0.918482 | 0.37834 | 0.274582 | 0.06812 | 0.587308 | -0.090412 | 0.101927 | 0.060862 | 0.449647 | 0.217365 | 0.08201 | 1.145547 | 0.249217 | 1.099893 | 1.686519 | 0.582438 | 0.164176 | 0.583197 | | 22.25 | 0.101927 | 0.79923 |
| 22.167 | 0.159872 | -0.103025 | -0.197174 | 0.14842 | -0.005823 | 0.143856 | 0.311872 | 0.552552 | 0.495659 | 0.035641 | 0.449569 | -0.149813 | 0.111759 | 0.499384 | 0.496563 | 0.279247 | 0.306935 | 0.947882 | 0.271428 | 0.737435 | 0.710647 | 0.681335 | 0.107008 | 0.61099 | | 22.26389 | 0.111759 | 0.748417 |
| 22.181 | 0.149228 | 0.199626 | -0.12963 | 0.144773 | 0.017558 | 0.114769 | 0.014046 | 0.522777 | 0.402803 | 0.009335 | -0.151742 | 0.252358 | 0.115359 | 0.451206 | 0.467649 | 0.3276 | 0.494585 | 0.828429 | 0.442473 | 0.118482 | -0.100386 | 0.507778 | 0.059321 | 0.464227 | | 22.27778 | 0.115359 | 0.483117 |
| 22.194 | 0.250625 | 0.354971 | -0.019395 | 0.166141 | 0.035242 | 0.05707 | 0.243218 | 0.413035 | 0.03508 | 0.039697 | -0.185446 | 0.456819 | 0.113799 | 0.308249 | 0.321858 | 0.283659 | 0.409708 | 0.806952 | 0.504976 | 0.104464 | -0.548192 | 0.348769 | 0.057317 | 0.333251 | | 22.29167 | 0.113799 | 0.371076 |
| 22.208 | 0.257431 | 0.176682 | -0.026956 | 0.121474 | 0.014147 | -0.005242 | 0.568337 | 0.307619 | -0.159732 | 0.053727 | 0.276736 | 0.278733 | 0.111075 | 0.674646 | 0.305962 | 0.300353 | 0.359738 | 0.859959 | 0.566443 | 0.799521 | 0.301869 | 0.386411 | 0.198629 | 0.565227 | | 22.30556 | 0.111075 | 0.626748 |
| 22.222 | 0.124165 | -0.108403 | -0.092813 | 0.031147 | -0.051993 | -0.05625 | 0.625403 | 0.224664 | -0.059063 | -0.002955 | 0.429238 | 0.023796 | 0.084326 | 0.851313 | 0.435315 | 0.346925 | 0.303451 | 0.609957 | 0.583573 | 1.295266 | 1.056292 | 0.334042 | 0.272672 | 0.732774 | | 22.31944 | 0.084326 | 0.757972 |
| 22.236 | 0.043954 | -0.3194 | -0.066671 | 0.010877 | -0.09222 | -0.057672 | 0.56246 | 0.169042 | 0.158153 | -0.049414 | 0.052159 | -0.066006 | 0.038684 | 0.36739 | 0.526871 | 0.327006 | 0.209308 | 0.208315 | 0.471779 | 0.995042 | 0.976784 | 0.195122 | 0.234852 | 0.482859 | | 22.33333 | 0.038684 | 0.470031 |
| 22.25 | 0.069894 | -0.220712 | -0.023173 | 0.023971 | -0.075489 | -0.013416 | 0.572003 | 0.123525 | 0.218838 | -0.03503 | -0.112368 | 0.222783 | 0.01639 | -0.10707 | 0.441975 | 0.203121 | 0.164289 | 0.066809 | 0.304068 | 0.301733 | 0.386444 | 0.076069 | 0.207081 | 0.167283 | | 22.34722 | 0.01639 | 0.142084 |
| 22.264 | 0.107376 | 0.104058 | -0.076458 | 0.000139 | -0.046116 | 0.023295 | 0.449506 | 0.021547 | 0.056116 | 0.009157 | -0.070782 | 0.774666 | 0.045338 | 0.033406 | 0.168148 | -0.017208 | 0.108783 | 0.167338 | 0.133004 | -0.138735 | 0.145351 | -0.005198 | 0.146884 | 0.030158 | | 22.36111 | 0.045338 | 0.057054 |
| 22.278 | 0.071632 | 0.3372 | -0.150268 | -0.058307 | -0.043295 | -0.020947 | 0.266834 | -0.02288 | -0.123359 | 0.002901 | -0.228554 | 0.78249 | 0.043496 | 0.323126 | -0.09324 | -0.22264 | 0.032508 | 0.277356 | 0.039994 | -0.197088 | 0.06961 | -0.018425 | 0.15766 | 0.030555 | | 22.375 | 0.043496 | 0.146812 |
| 22.292 | 0.017843 | 0.269212 | -0.100755 | -0.077776 | -0.059582 | -0.08154 | 0.283693 | 0.179063 | -0.129505 | -0.042556 | -0.305237 | 0.210463 | 0.000243 | 0.10107 | -0.183403 | -0.259579 | -0.046692 | 0.321732 | 0.113822 | 0.138319 | 0.258704 | 0.061963 | 0.328424 | 0.094681 | | 22.38889 | 0.000243 | 0.292007 |
| 22.306 | 0.048452 | 0.072185 | 0.052181 | -0.012226 | -0.060717 | -0.059995 | 0.439323 | 0.465651 | 0.04498 | -0.041511 | 0.025129 | -0.181427 | -0.015364 | -0.289557 | -0.06208 | -0.14306 | -0.047577 | 0.418121 | 0.248765 | 0.6934 | 0.443468 | 0.138931 | 0.443287 | 0.200949 | | 22.40278 | -0.015364 | 0.432709 |
| 22.319 | 0.151492 | -0.022135 | 0.130435 | 0.045291 | -0.026885 | 0.015613 | 0.4997 | 0.490332 | 0.241558 | 0.008013 | 0.504096 | -0.111774 | 0.01037 | -0.01448 | 0.202096 | -0.035505 | 0.081486 | 0.64277 | 0.25192 | 0.86035 | 0.447285 | 0.175871 | 0.261004 | 0.267432 | | 22.41667 | 0.01037 | 0.442753 |
| 22.333 | 0.26013 | 0.055276 | 0.102272 | 0.035511 | 0.031506 | 0.062381 | 0.595955 | 0.264493 | 0.289675 | 0.052062 | 0.904638 | 0.335601 | 0.067846 | 0.737639 | 0.375852 | -0.071838 | 0.280485 | 1.18917 | 0.236134 | 0.432904 | 0.424642 | 0.279171 | -0.107516 | 0.327281 | | 22.43056 | 0.067846 | 0.435039 |
| 22.347 | 0.318739 | 0.164845 | 0.00932 | 0.012214 | 0.053007 | 0.043336 | 0.866571 | 0.234025 | 0.259347 | 0.033965 | 1.133349 | 1.118392 | 0.059233 | 1.094098 | -0.038565 | -0.163778 | 0.272696 | 1.828584 | 0.355905 | -0.007511 | 0.556648 | 0.441409 | -0.214549 | 0.324376 | | 22.44444 | 0.059233 | 0.444006 |
| 22.361 | 0.337333 | 0.091426 | -0.130183 | -0.044902 | -0.045964 | -0.093893 | 0.844724 | 0.239737 | 0.188966 | -0.096596 | 0.97807 | 1.742164 | -0.096883 | 0.839999 | -0.461119 | -0.237871 | -0.10314 | 1.42203 | 0.446709 | -0.160158 | 0.592879 | 0.468974 | -0.218311 | 0.177337 | | 22.45833 | -0.096883 | 0.274075 |
| 22.375 | 0.275171 | -0.218376 | -0.267333 | -0.141735 | -0.198599 | -0.334524 | 0.396945 | 0.057298 | 0.035271 | -0.287605 | 0.363395 | 0.973232 | -0.345473 | 0.251553 | -0.385575 | -0.290802 | -0.324625 | 1.159384 | 0.346479 | -0.121315 | 0.599343 | 0.31189 | -0.212045 | -0.037504 | | 22.47222 | -0.345473 | 0.010967 |
| 22.389 | 0.196068 | -0.409309 | -0.391601 | -0.180475 | -0.229298 | -0.400984 | 0.118935 | -0.098044 | -0.016441 | -0.291987 | 0.023247 | 0.137513 | -0.513372 | 0.240202 | -0.18585 | -0.320182 | -0.351785 | 1.117594 | 0.136883 | 0.00058 | 0.403636 | 0.217182 | -0.134943 | -0.1424 | | 22.48611 | -0.513372 | -0.140148 |
| 22.403 | 0.365409 | -0.245295 | -0.441105 | -0.059715 | -0.12109 | -0.231259 | 0.310259 | 0.099423 | 0.041335 | -0.094415 | 0.282728 | 0.183182 | -0.551433 | 0.862393 | 0.516596 | -0.43771 | -0.3433 | 0.881107 | 0.08522 | 0.192453 | -0.102454 | 0.078395 | -0.179727 | -0.257795 | | 22.5 | -0.551433 | -0.23728 |
| 22.417 | 0.384938 | -0.05197 | -0.310161 | 0.137511 | -0.008328 | -0.106196 | 0.732694 | 0.170156 | 0.079448 | 0.040608 | 0.381741 | -0.185981 | -0.48412 | 1.308469 | 0.019373 | -0.616105 | -0.526721 | 0.504557 | 0.126212 | 0.241442 | -0.479682 | -0.063885 | -0.233183 | -0.49456 | | 22.51389 | -0.48412 | -0.41412 |
| 22.431 | 0.326948 | 0.065348 | -0.174679 | 0.183487 | 0.038368 | -0.061747 | 1.215038 | 0.1447 | 0.101906 | 0.080348 | 0.100646 | -0.329533 | -0.381735 | 1.302718 | -0.496281 | -0.706817 | -0.873255 | 0.110226 | 0.259798 | 0.077005 | -0.626882 | -0.153985 | -0.25356 | -0.714991 | | 22.52778 | -0.381735 | -0.619997 |
| 22.444 | 0.236139 | 0.146156 | -0.282294 | 0.1253 | 0.040265 | -0.094092 | 1.460277 | 0.111959 | 0.139885 | 0.088025 | 0.00963 | 0.587502 | -0.317093 | -0.29962 | -0.88599 | -0.992561 | -1.412898 | -0.535433 | 0.075627 | -0.463393 | -0.775753 | -0.478281 | -0.717169 | -1.042206 | | 22.54167 | -0.317093 | -1.051791 |
| 22.458 | 0.135147 | 0.152967 | -0.37915 | 0.056888 | 0.003896 | -0.168392 | 1.427317 | 0.069969 | 0.24664 | 0.082269 | 0.037376 | 1.269483 | -0.31255 | -1.365165 | -1.621961 | -1.847465 | -2.091377 | -1.569408 | -1.0332 | -1.410452 | -1.507669 | -1.312783 | -1.617094 | -1.541945 | | 22.55556 | -0.31255 | -1.69819 |
| 22.472 | 0.023452 | 0.019967 | -0.385407 | -0.015632 | -0.064754 | -0.14464 | 1.007323 | 0.05568 | 0.230495 | 0.028998 | 0.05861 | 0.577452 | -0.25241 | -1.933027 | -1.706035 | -1.924231 | -1.975281 | -1.82574 | -1.597819 | -1.588065 | -1.743626 | -1.479133 | -1.749044 | -1.587549 | | 22.56944 | -0.25241 | -1.760163 |
| 22.486 | -0.013545 | -0.130594 | -0.152956 | -0.016601 | -0.086801 | 0.020123 | 0.481865 | 0.093829 | 0.252301 | -0.008041 | -0.043947 | 0.115086 | -0.078728 | -1.255712 | -0.973972 | -0.955888 | -0.985503 | -0.988012 | -0.87353 | -0.785199 | -1.016848 | -0.529173 | -0.990618 | -0.930746 | | 22.58333 | -0.078728 | -0.942402 |
| 22.5 | 0.017874 | -0.143159 | -0.121585 | -0.012465 | -0.053477 | 0.102741 | 0.277363 | 0.066783 | 0.21069 | 0.014239 | -0.092564 | -0.039521 | -0.020852 | -0.559558 | -0.452415 | -0.275129 | -0.308719 | -0.33808 | -0.233367 | -0.174801 | -0.451869 | 0.188383 | -0.405698 | -0.254767 | | 22.59722 | -0.020852 | -0.178897 |
| 22.514 | -0.0079 | -0.125514 | -0.213413 | -0.065552 | -0.056586 | 0.040631 | 0.258924 | -0.039953 | 0.07276 | -0.002542 | -0.096417 | -0.177953 | -0.154803 | -0.29678 | -0.348392 | -0.016357 | -0.193958 | -0.197117 | -0.151705 | -0.033672 | -0.358815 | 0.316972 | -0.284594 | -0.083315 | | 22.61111 | -0.154803 | -0.025304 |
| 22.528 | -0.079497 | -0.097655 | -0.216198 | -0.091599 | -0.056211 | -0.002398 | 0.296718 | -0.013576 | -0.041804 | -0.0259 | -0.055309 | -0.145389 | -0.255159 | -0.14283 | -0.3916 | -0.029375 | -0.291419 | -0.170287 | -0.144622 | -0.018751 | -0.331364 | 0.237101 | -0.36535 | -0.188102 | | 22.625 | -0.255159 | -0.180676 |
| 22.542 | -0.151227 | -0.045807 | -0.088554 | -0.049055 | -0.012317 | -0.00107 | 0.270179 | 0.062134 | -0.086725 | -0.014408 | -0.010412 | 0.01091 | -0.274751 | -0.003704 | -0.278504 | -0.100834 | -0.333607 | -0.10326 | 0.01286 | -0.011257 | -0.277506 | 0.206225 | -0.432009 | -0.291618 | | 22.63889 | -0.274751 | -0.306584 |
| 22.556 | -0.250634 | -0.040908 | 0.015878 | -0.0161 | 0.003675 | -0.020173 | 0.169042 | 0.060394 | -0.096156 | -0.028485 | -0.013241 | 0.070974 | -0.271274 | -0.058363 | -0.206089 | -0.124639 | -0.310451 | -0.014958 | 0.12653 | -0.016994 | -0.220745 | 0.104491 | -0.424838 | -0.3534 | | 22.65278 | -0.271274 | -0.360661 |
| 22.569 | -0.329862 | -0.063631 | 0.048352 | 0.003481 | -0.012118 | -0.030443 | -0.003367 | 0.027778 | -0.050259 | -0.051765 | -0.039876 | 0.065963 | -0.183416 | -0.17741 | -0.162727 | -0.10273 | -0.212139 | 0.060836 | 0.193933 | 0.047147 | -0.174376 | 0.051398 | -0.238731 | -0.321779 | | 22.66667 | -0.183416 | -0.32901 |
| 22.583 | -0.336694 | -0.032328 | 0.050238 | 0.031971 | 0.004121 | -0.016487 | -0.151085 | -0.017603 | 0.082094 | -0.01805 | -0.008549 | 0.094697 | -0.007776 | 0.011466 | -0.001205 | -0.00983 | -0.031093 | 0.191654 | 0.224409 | 0.142599 | -0.15237 | 0.093495 | 0.010979 | -0.170896 | | 22.68056 | -0.007776 | -0.186235 |
| 22.597 | -0.215317 | 0.016992 | 0.009131 | 0.033174 | 0.028638 | -0.014698 | -0.305484 | -0.109823 | 0.248128 | 0.015034 | 0.03858 | 0.10884 | 0.127977 | 0.12894 | 0.131772 | 0.044116 | 0.08805 | 0.350598 | 0.142048 | 0.152307 | -0.087347 | 0.10186 | 0.137029 | 0.002546 | | 22.69444 | 0.127977 | -0.056859 |
| 22.611 | 0.013264 | -0.002216 | -0.063121 | -0.013324 | 0.002637 | -0.023922 | -0.534326 | -0.18003 | 0.250399 | -0.01266 | 0.005645 | 0.058283 | 0.134685 | -0.018426 | 0.114867 | 0.008171 | 0.08618 | 0.374363 | 0.025558 | 0.082124 | -0.123272 | 0.015602 | 0.188613 | 0.114437 | | 22.70833 | 0.134685 | 0.029137 |
| 22.625 | 0.358305 | -0.048901 | -0.091678 | -0.064366 | -0.029072 | -0.013666 | -0.688488 | -0.159408 | 0.104697 | -0.047583 | -0.055213 | 0.005983 | 0.068696 | -0.173805 | 0.053937 | -0.026316 | 0.063593 | 0.351966 | -0.035685 | 0.055978 | -0.206232 | -0.070765 | 0.22469 | 0.192834 | | 22.72222 | 0.068696 | 0.144676 |
| 22.639 | 0.667807 | -0.054801 | -0.057632 | -0.084383 | -0.029292 | 0.001896 | -0.448131 | -0.06724 | -0.062183 | -0.045423 | -0.04712 | 0.041354 | 0.006094 | -0.174484 | 0.073343 | 0.000087 | 0.075829 | 0.293449 | 0.005321 | 0.09956 | -0.212635 | -0.099148 | 0.249722 | 0.231604 | | 22.73611 | 0.006094 | 0.247402 |
| 22.653 | 0.687699 | -0.035523 | 0.050465 | -0.071165 | -0.018048 | -0.001945 | -0.100586 | 0.026915 | -0.177599 | -0.039335 | -0.008459 | 0.116179 | -0.028692 | -0.055154 | 0.117453 | 0.019984 | 0.079642 | 0.072567 | 0.09262 | 0.150233 | -0.106035 | -0.161377 | 0.158929 | 0.168466 | | 22.75 | -0.028692 | 0.223202 |
| 22.667 | 0.481996 | -0.017855 | 0.175414 | 0.006449 | -0.016943 | -0.026676 | 0.236275 | 0.071227 | -0.285907 | -0.052911 | 0.03974 | 0.141607 | -0.052642 | 0.084883 | 0.113624 | 0.004896 | 0.043901 | -0.346769 | 0.183233 | 0.16675 | -0.018821 | -0.241882 | 0.053256 | 0.068406 | | 22.76389 | -0.052642 | 0.096607 |
| 22.681 | 0.325492 | 0.006045 | 0.221142 | 0.026772 | -0.020221 | -0.05808 | 0.572996 | 0.0638 | -0.288512 | -0.071887 | 0.098408 | 0.100316 | -0.091275 | 0.151512 | 0.08296 | -0.038918 | 0.008003 | -0.810751 | 0.240022 | 0.161648 | 0.016242 | -0.20115 | 0.06446 | -0.015909 | | 22.77778 | -0.091275 | 0.007926 |
| 22.694 | 0.192304 | 0.009455 | 0.195781 | 0.00013 | -0.012769 | -0.078861 | 0.516588 | 0.034056 | -0.26424 | -0.069897 | 0.216416 | 0.050583 | -0.094391 | 0.044675 | 0.062678 | -0.043892 | 0.012646 | -0.499946 | 0.264593 | 0.11891 | 0.022596 | 0.045161 | 0.165987 | -0.001876 | | 22.79167 | -0.094391 | 0.074474 |
| 22.708 | 0.047315 | 0.01029 | 0.155115 | -0.040905 | -0.00018 | -0.05185 | 0.134609 | -0.019476 | -0.090701 | -0.040881 | 0.211823 | 0.020264 | -0.047682 | -0.105332 | 0.051991 | -0.013893 | 0.053156 | -0.15865 | 0.231781 | -0.022042 | -0.007177 | 0.365509 | 0.269007 | 0.143501 | | 22.80556 | -0.047682 | 0.233432 |
| 22.722 | -0.157999 | -0.041415 | 0.048673 | -0.097746 | 0.004807 | -0.052958 | -0.263074 | -0.102623 | 0.106986 | -0.018726 | 0.159294 | 0.007683 | 0.003288 | -0.015491 | 0.039344 | 0.040855 | 0.086612 | 0.255304 | 0.166241 | -0.28918 | -0.072861 | 0.533951 | 0.363184 | 0.250132 | | 22.81944 | 0.003288 | 0.371016 |
| 22.736 | -0.436765 | -0.06561 | -0.103035 | -0.143252 | 0.00789 | -0.040903 | -0.594463 | -0.172898 | 0.205763 | -0.005602 | 0.117459 | -0.0384 | 0.051615 | 0.097253 | 0.005875 | 0.092318 | 0.127629 | 0.783023 | 0.064149 | -0.389489 | -0.143006 | 0.341138 | 0.345528 | 0.261169 | | 22.83333 | 0.051615 | 0.378551 |
| 22.75 | -0.709683 | -0.044715 | -0.167662 | -0.128135 | 0.004695 | -0.013935 | -0.874612 | -0.161971 | 0.169857 | -0.002597 | 0.137099 | -0.140694 | 0.061714 | -0.022657 | -0.033602 | 0.079893 | 0.161159 | 0.567032 | 0.01568 | -0.274191 | -0.158285 | 0.010449 | 0.121719 | 0.217607 | | 22.84722 | 0.061714 | 0.277312 |
| 22.764 | -0.748072 | -0.02047 | -0.058734 | -0.106298 | -0.008011 | -0.004624 | -1.104295 | -0.069807 | 0.025953 | -0.029982 | 0.089731 | -0.284401 | 0.016056 | -0.126606 | -0.112142 | -0.02235 | 0.086194 | -0.116203 | -0.053873 | -0.28838 | -0.247437 | -0.383266 | -0.297072 | 0.163305 | | 22.86111 | 0.016056 | 0.199352 |
| 22.778 | -0.576075 | -0.036441 | 0.16703 | -0.144931 | -0.027533 | -0.025502 | -0.872302 | 0.00327 | -0.194143 | -0.084393 | 0.149214 | -0.475514 | -0.027043 | -0.206246 | -0.228789 | -0.163539 | -0.135944 | -0.750855 | -0.747706 | -0.402818 | -0.324174 | -0.60027 | -0.607344 | 0.081571 | | 22.875 | -0.027043 | 0.087543 |
| 22.792 | -0.48984 | -0.101711 | 0.316289 | -0.178031 | -0.052635 | -0.059158 | -0.410035 | -0.016231 | -0.350444 | -0.15423 | 0.270164 | -0.458718 | -0.044819 | -0.481557 | -0.315492 | -0.263627 | -0.37481 | -0.687395 | -0.606245 | -0.521536 | -0.415286 | -0.690824 | -0.597094 | -0.040405 | | 22.88889 | -0.044819 | -0.053953 |
| 22.806 | -0.342401 | -0.190043 | 0.215077 | -0.185551 | -0.079398 | -0.078106 | -0.117774 | -0.12926 | -0.403739 | -0.227782 | 0.146735 | -0.217183 | 0.010993 | -0.76486 | -0.352971 | -0.275886 | -0.449589 | -0.4794 | -0.472403 | -0.654725 | -0.499024 | -0.651324 | -0.442783 | -0.112105 | | 22.90278 | 0.010993 | -0.097071 |
| 22.819 | -0.141804 | -0.204246 | -0.002747 | -0.178971 | -0.109681 | -0.094056 | -0.21648 | -0.284629 | -0.398306 | -0.282471 | -0.04712 | -0.020393 | 0.118685 | -0.866391 | -0.36227 | -0.20797 | -0.406809 | -0.882342 | -0.469665 | -0.780242 | -0.592453 | -0.592171 | -0.386243 | -0.120745 | | 22.91667 | 0.118685 | -0.07673 |
| 22.833 | -0.121979 | -0.131068 | -0.105053 | -0.159272 | -0.135904 | -0.07874 | -0.694252 | -0.510635 | -0.429587 | -0.269872 | -0.229834 | -0.032637 | 0.235184 | -0.799728 | -0.331577 | -0.14012 | -0.364089 | -0.775539 | -0.415041 | -0.901796 | -0.570092 | -0.52669 | -0.474214 | -0.107422 | | 22.93056 | 0.235184 | -0.119379 |
| 22.847 | -0.268793 | 0.019901 | 0.099681 | -0.171423 | -0.122943 | -0.007769 | -1.03731 | -0.819375 | -0.265771 | -0.144991 | -0.61127 | -0.186927 | 0.361712 | -0.678598 | -0.272821 | -0.047866 | -0.317599 | -0.500864 | -0.362974 | -0.975845 | -0.390022 | -0.468569 | -0.560453 | -0.16067 | | 22.94444 | 0.361712 | -0.228841 |
| 22.861 | -0.333991 | 0.118833 | 0.096247 | -0.243099 | -0.082167 | 0.035604 | -1.055084 | -0.988948 | -0.216767 | -0.011586 | -1.175775 | -0.462358 | 0.447092 | -0.653529 | -0.252979 | -0.013976 | -0.285631 | -0.430799 | -0.499731 | -0.925833 | -0.369065 | -0.400372 | -0.578045 | -0.250052 | | 22.95833 | 0.447092 | -0.359435 |
| 22.875 | -0.310923 | 0.069852 | -0.053964 | -0.291566 | -0.08515 | 0.007119 | -0.804889 | -0.99944 | -0.242691 | -0.000368 | -1.047673 | -0.759112 | 0.387316 | -0.900926 | -0.34873 | -0.114798 | -0.331416 | -0.621146 | -0.622744 | -0.734557 | -0.563133 | -0.528588 | -0.579306 | -0.319961 | | 22.97222 | 0.387316 | -0.450278 |
| 22.889 | -0.289658 | -0.067568 | -0.144348 | -0.285278 | -0.113165 | -0.038645 | -0.586598 | -1.134793 | -0.280974 | -0.077707 | -0.868266 | -0.892069 | 0.246759 | -1.321196 | -0.550136 | -0.122961 | -0.475916 | -0.958563 | -0.702026 | -0.624715 | -0.534865 | -0.697089 | -0.574488 | -0.403183 | | 22.98611 | 0.246759 | -0.515009 |
| 22.903 | -0.242854 | -0.141106 | -0.123906 | -0.216 | -0.090565 | -0.054063 | -0.48346 | -0.714994 | -0.259967 | -0.091891 | -0.584752 | -0.725808 | 0.137625 | -1.517286 | -0.691173 | -0.126191 | -0.441568 | -1.186831 | -0.65774 | -0.593254 | -0.458197 | -0.703359 | -0.432275 | -0.432719 | | 23 | 0.137625 | -0.519853 |
| 22.917 | -0.032222 | -0.064385 | -0.058735 | -0.075016 | 0.029806 | 0.042078 | -0.389545 | -0.194699 | -0.168992 | 0.021192 | -0.163508 | -0.453654 | 0.135017 | -1.088011 | -0.60321 | -0.102005 | -0.21416 | -1.629428 | -0.174607 | -0.423588 | -0.299208 | -0.425869 | -0.191931 | -0.314525 | | 23.01389 | 0.135017 | -0.422705 |
| 22.931 | 0.263796 | 0.062048 | 0.122804 | 0.11243 | 0.217948 | 0.252931 | -0.233583 | 0.327094 | 0.009371 | 0.217251 | 0.338614 | -0.078813 | 0.225286 | -0.221215 | -0.317734 | 0.064369 | 0.080563 | -1.467757 | 0.430055 | 0.002335 | 0.028079 | 0.079686 | 0.170084 | -0.028299 | | 23.02778 | 0.225286 | -0.185473 |
| 22.944 | 0.250621 | 0.045749 | 0.255231 | 0.20876 | 0.252725 | 0.372974 | -0.329046 | 0.504084 | 0.077489 | 0.287641 | 0.340774 | 0.060623 | 0.282881 | 0.435954 | -0.127195 | 0.149926 | 0.244206 | -0.405742 | 0.537405 | 0.318069 | 0.247117 | 0.166885 | 0.53479 | 0.077883 | | 23.04167 | 0.282881 | -0.048635 |
| 22.958 | -0.036274 | -0.227602 | 0.197317 | 0.107216 | 0.082507 | 0.212963 | -0.560895 | -0.058191 | -0.125402 | 0.096061 | -0.060117 | -0.153375 | 0.276871 | 0.704664 | 0.079121 | 0.164598 | 0.21933 | -0.187052 | 0.289206 | 0.302547 | 0.034049 | -0.160284 | 0.738879 | -0.106172 | | 23.05556 | 0.276871 | -0.233648 |
| 22.972 | -0.023719 | -0.552072 | 0.194497 | -0.023294 | -0.038828 | 0.055505 | -0.926375 | -0.979447 | -0.60794 | -0.182785 | 0.062895 | -0.340259 | 0.232652 | 0.994598 | 0.627776 | 0.242011 | 0.229183 | -1.27285 | -0.106138 | 0.426144 | -0.620125 | -0.45445 | 1.094021 | -0.058107 | | 23.06944 | 0.232652 | -0.291253 |
| 22.986 | 0.420612 | -0.335088 | 0.109424 | -0.016861 | 0.053554 | 0.182565 | -1.246635 | -1.22657 | -0.618456 | -0.172248 | 0.606607 | -0.19378 | 0.235313 | 1.567869 | 1.094001 | 0.345045 | 0.430387 | -1.865398 | -0.528146 | 1.252507 | -0.565251 | 0.27446 | 1.608728 | 0.484498 | | 23.08333 | 0.235313 | 0.226505 |
| 23 | 0.807311 | 0.391014 | -0.0512 | 0.055817 | 0.256496 | 0.91411 | -1.415085 | -0.532931 | -0.221857 | 0.041631 | 1.104981 | -0.070147 | 0.37894 | 2.653605 | 0.729052 | 0.622618 | 0.428326 | -2.115003 | -0.747261 | 3.752741 | -0.180521 | 3.35162 | 2.760185 | 0.828597 | | 23.09722 | 0.37894 | 0.75339 |
| 23.014 | 0.248957 | 0.386366 | 0.103592 | 0.06535 | -0.026398 | 0.814698 | -1.195784 | 0.133801 | 0.032072 | 0.202675 | -0.215473 | -0.141306 | 0.426973 | 1.758179 | -0.38034 | 0.431231 | 0.305171 | -1.841777 | -0.450964 | 2.225199 | -0.055756 | 2.724553 | 2.465103 | 0.557523 | | 23.11111 | 0.426973 | 0.545837 |
| 23.028 | -0.318129 | 0.049581 | 0.043827 | -0.009178 | 0.018148 | 0.268486 | -0.503592 | 0.505816 | 0.187406 | 0.227533 | -0.275358 | -0.039547 | 0.297508 | -0.023134 | 0.251478 | 0.222139 | 0.084368 | 0.048414 | 0.029651 | -0.009207 | 0.048727 | -0.378368 | 0.533829 | 0.004846 | | 23.125 | 0.297508 | -0.164501 |
| 23.042 | -0.471154 | 0.133405 | -0.021304 | -0.052142 | 0.094799 | 0.079033 | 0.14713 | 0.48946 | 0.242226 | 0.139309 | -0.289021 | 0.131747 | 0.133247 | -0.761766 | 0.453247 | 0.698763 | 0.228446 | 2.355512 | 0.123656 | 0.220448 | 0.108962 | -0.881949 | -0.502653 | 0.164658 | | 23.13889 | 0.133247 | 0.10347 |
| 23.056 | -0.447529 | 0.060743 | -0.241156 | -0.091568 | -0.027088 | 0.050112 | 0.535251 | 0.170015 | 0.155257 | -0.007696 | -0.565412 | -0.05303 | 0.008965 | -0.713261 | 0.785895 | 0.718354 | 0.121153 | 3.716932 | -0.080222 | 0.843481 | 0.122006 | 0.121977 | -0.750798 | 0.749457 | | 23.15278 | 0.008965 | 1.108672 |
| 23.069 | -0.255068 | -0.414268 | -0.310995 | -0.101709 | -0.300676 | 0.016363 | 0.561492 | -0.06863 | 0.063286 | -0.143486 | -1.041496 | -0.457751 | -0.127174 | -0.834289 | 1.1806 | 0.25233 | -0.15208 | 4.016266 | -0.345879 | 0.817926 | 0.130771 | 0.060931 | -0.361936 | 0.909783 | | 23.16667 | -0.127174 | 1.686765 |
| 23.083 | 0.092074 | -0.604166 | -0.063557 | 0.010486 | -0.437216 | 0.021095 | 0.375034 | 0.03084 | 0.030079 | -0.123926 | -1.12894 | -0.630455 | -0.137015 | -0.714499 | 1.528082 | -0.088643 | -0.246201 | 2.812751 | -0.566749 | 0.475664 | 0.382762 | -0.609734 | 0.240938 | 0.938053 | | 23.18056 | -0.137015 | 1.474074 |
| 23.097 | 0.475479 | -0.162764 | 0.285228 | 0.148471 | -0.369523 | 0.07368 | 0.195442 | 0.458965 | 0.045668 | 0.039952 | -0.543059 | -0.508877 | 0.017602 | 0.067647 | 1.242393 | 0.130592 | 0.119198 | 0.837021 | -0.063472 | 0.215468 | 0.85203 | -0.400396 | 0.63309 | 0.597324 | | 23.19444 | 0.017602 | 1.05727 |
| 23.111 | 0.648742 | 0.283063 | 0.3527 | 0.190507 | -0.083188 | 0.142196 | 0.251848 | 0.909352 | 0.101685 | 0.119382 | 0.283098 | -0.137268 | 0.24109 | 1.192687 | 0.267267 | 0.737586 | 0.413396 | -0.546385 | 0.133639 | 0.172252 | 0.739369 | -0.488537 | 0.584029 | 0.251413 | | 23.20833 | 0.24109 | 0.301552 |
| 23.125 | 0.490991 | 0.508431 | 0.233599 | 0.179299 | 0.147955 | 0.212153 | 0.595008 | 0.962346 | 0.279487 | 0.038701 | 1.157357 | 0.227602 | 0.389521 | 0.705914 | -0.534981 | 0.563018 | 0.035074 | -0.753015 | -0.140482 | -0.11775 | 0.006953 | -1.085248 | -0.065704 | -0.288064 | | 23.22222 | 0.389521 | -0.4957 |
| 23.139 | 0.203107 | 0.381135 | 0.263371 | 0.131088 | 0.010987 | 0.224566 | 1.219466 | 0.58693 | 0.463992 | -0.017684 | 1.621096 | 0.316503 | 0.33646 | -0.760681 | -0.021525 | -0.43806 | -0.635573 | 0.055092 | -0.444739 | -0.689479 | -1.012041 | -1.438033 | -1.235216 | -0.820429 | | 23.23611 | 0.33646 | -0.726842 |
| 23.153 | 0.000783 | -0.014799 | 0.327296 | 0.038636 | -0.139476 | 0.152449 | 1.960484 | 0.261419 | 0.471687 | 0.01967 | 1.374611 | 0.199582 | 0.161131 | -0.819517 | 1.251648 | -1.214563 | -0.561719 | 1.176391 | -0.217929 | -0.895608 | -0.773631 | -1.045625 | -2.996555 | -0.511022 | | 23.25 | 0.161131 | -0.378633 |
| 23.167 | 0.090197 | -0.106577 | 0.249801 | 0.013835 | -0.131975 | 0.073262 | 1.818987 | 0.189995 | 0.236764 | 0.040363 | 0.804488 | 0.176144 | 0.089147 | 0.685359 | 1.115239 | -1.700298 | 0.17803 | 2.677825 | 0.133427 | -0.887446 | 0.205569 | -0.165415 | -3.206611 | 0.241838 | | 23.26389 | 0.089147 | 0.281645 |
| 23.181 | 0.325837 | -0.001646 | 0.06408 | 0.099877 | 0.030781 | 0.04308 | 1.03624 | 0.240687 | -0.006712 | 0.022634 | 0.151339 | 0.260127 | 0.144212 | 1.443745 | 0.235274 | -1.221755 | 0.510847 | 1.938387 | 0.171043 | -1.240415 | 0.517502 | 0.277384 | -1.903044 | 0.465969 | | 23.27778 | 0.144212 | 0.433181 |
| 23.194 | 0.278915 | -0.014126 | -0.009875 | 0.136391 | 0.006779 | -0.006608 | 0.485571 | 0.233443 | -0.074999 | 0.035124 | -0.439597 | 0.255486 | 0.144522 | 0.526928 | -0.34005 | -0.586897 | 0.259607 | 0.004419 | 0.239493 | -1.760916 | 0.144208 | 0.375525 | -1.003099 | 0.250555 | | 23.29167 | 0.144522 | 0.255281 |
| 23.208 | 0.049606 | -0.124063 | 0.037796 | 0.044309 | -0.131374 | -0.127189 | -0.155703 | 0.168486 | 0.010632 | 0.022082 | -0.532022 | 0.065596 | 0.058429 | -0.124678 | -0.315966 | -0.376449 | -0.124071 | -1.007416 | 0.306238 | -1.54232 | 0.046657 | 0.326367 | 0.157404 | -0.160671 | | 23.30556 | 0.058429 | -0.100955 |
| 23.222 | 0.000358 | -0.148285 | 0.066567 | -0.02203 | -0.182373 | -0.240894 | -0.844194 | 0.132731 | 0.071122 | -0.126608 | -0.387014 | -0.347659 | -0.066978 | -0.534578 | -0.490218 | -0.80472 | -0.744354 | -1.483097 | -0.443203 | -1.567902 | -0.384697 | -0.430445 | 0.036455 | -0.980314 | | 23.31944 | -0.066978 | -0.897943 |
| 23.236 | 0.096663 | -0.05732 | 0.11927 | 0.125691 | -0.048986 | -0.152267 | -0.609847 | 0.313243 | 0.099356 | -0.216831 | -0.220568 | -0.42202 | -0.082751 | -1.178962 | -1.086819 | -1.656756 | -1.521616 | -1.525102 | -1.853593 | -2.468298 | -1.642139 | -1.653078 | -1.304501 | -1.888622 | | 23.33333 | -0.082751 | -1.829217 |
| 23.25 | 0.165817 | 0.056453 | 0.137274 | 0.282769 | 0.133467 | 0.187487 | 0.533404 | 0.636886 | 0.193923 | 0.014519 | 0.239245 | 0.030278 | 0.135273 | -1.37039 | -0.760502 | -1.427686 | -1.187902 | -0.321481 | -1.619238 | -2.414538 | -1.799934 | -1.091785 | -1.308443 | -1.364724 | | 23.34722 | 0.135273 | -1.343722 |
| 23.264 | 0.166822 | -0.019644 | 0.081892 | 0.196546 | 0.09541 | 0.46365 | 1.570778 | 0.709056 | 0.254742 | 0.236213 | 0.607866 | 0.417647 | 0.27623 | -0.738624 | 0.164913 | -0.545966 | -0.137253 | 1.220307 | -0.144742 | -0.972301 | -0.038359 | 0.544518 | -0.058275 | 0.068651 | | 23.36111 | 0.27623 | 0.112229 |
| 23.278 | 0.085597 | -0.185211 | 0.085643 | 0.031205 | -0.076723 | 0.366626 | 2.401394 | 0.62123 | 0.161237 | 0.173453 | 0.497253 | 0.367474 | 0.160547 | 0.082908 | 0.335569 | -0.062253 | 0.13841 | 1.018196 | 0.457344 | -0.112409 | 1.174964 | 1.05146 | 0.237524 | 0.736349 | | 23.375 | 0.160547 | 0.822522 |
| 23.292 | 0.024747 | -0.119448 | 0.192873 | 0.023469 | -0.132134 | 0.17554 | 2.198864 | 0.603118 | 0.22057 | 0.081491 | 0.331317 | 0.256894 | 0.126181 | 0.875465 | 0.140157 | 0.172442 | 0.057178 | 0.456761 | 0.469573 | 0.248106 | 1.247065 | 0.882112 | -0.249213 | 0.817045 | | 23.38889 | 0.126181 | 0.947266 |
| 23.306 | 0.098757 | 0.077983 | 0.486953 | 0.107021 | -0.023973 | 0.083014 | 1.232461 | 0.406348 | 0.269836 | 0.069189 | 0.204021 | 0.303835 | 0.18095 | 1.187706 | 0.103311 | 0.203342 | 0.125004 | 0.250042 | 0.490881 | 0.329464 | 0.817503 | 0.56672 | -0.221849 | 0.690605 | | 23.40278 | 0.18095 | 0.855385 |
| 23.319 | 0.273164 | 0.12852 | 0.407815 | 0.100219 | 0.041269 | 0.10344 | 0.656298 | 0.203942 | 0.150716 | 0.07399 | 0.062123 | 0.457079 | 0.129713 | 1.16754 | 0.227979 | 0.052648 | 0.250641 | 0.236273 | 0.505629 | 0.058926 | 0.24252 | 0.326799 | 0.325772 | 0.526201 | | 23.41667 | 0.129713 | 0.664277 |
| 23.333 | 0.41243 | 0.051237 | 0.203113 | 0.056335 | 0.007787 | 0.118266 | 0.341372 | 0.290092 | 0.053404 | 0.079983 | -0.031151 | 0.587739 | 0.035986 | 1.00065 | 0.38481 | 0.036931 | 0.2706 | 0.529416 | 0.60423 | -0.168923 | 0.141494 | 0.221313 | 0.464568 | 0.449778 | | 23.43056 | 0.035986 | 0.572759 |
| 23.347 | 0.446985 | 0.072461 | -0.057148 | -0.008924 | -0.023219 | 0.072677 | 0.162119 | 0.429499 | 0.049955 | 0.049945 | -0.079274 | 0.575931 | -0.01239 | 0.718941 | 0.396474 | 0.114202 | 0.241479 | 1.12732 | 0.55273 | -0.082023 | 0.53709 | 0.348946 | 0.319632 | 0.418201 | | 23.44444 | -0.01239 | 0.567771 |
| 23.361 | 0.437164 | 0.096874 | -0.234918 | -0.038304 | -0.009118 | -0.048517 | 0.079366 | 0.367797 | 0.057974 | -0.062952 | -0.172231 | 0.428961 | -0.053885 | -0.048587 | 0.367064 | 0.074242 | 0.191302 | 1.967873 | 0.276875 | 0.068927 | 0.991777 | 0.481405 | 0.319994 | 0.280029 | | 23.45833 | -0.053885 | 0.541312 |
| 23.375 | 0.464241 | -0.049221 | -0.331297 | -0.110187 | -0.022684 | -0.247938 | 0.069451 | 0.042727 | 0.054438 | -0.231691 | -0.373575 | 0.157887 | -0.143088 | -0.069629 | 0.334535 | -0.137774 | 0.047101 | 2.629593 | -0.099342 | 0.045372 | 1.110554 | 0.364667 | 0.39445 | 0.093717 | | 23.47222 | -0.143088 | 0.45969 |
| 23.389 | 0.465912 | -0.195692 | -0.221766 | 0.02775 | -0.031344 | -0.266228 | 0.127649 | -0.088324 | 0.188111 | -0.230403 | -0.517244 | 0.171458 | -0.211908 | 1.269351 | 0.354678 | -0.276924 | -0.13003 | 2.071631 | -0.126887 | -0.06823 | 0.580448 | 0.186813 | 0.459886 | 0.056811 | | 23.48611 | -0.211908 | 0.291102 |
| 23.403 | 0.514249 | -0.149651 | -0.080243 | 0.209163 | 0.171278 | -0.048631 | 0.389023 | 0.063537 | 0.371088 | -0.036136 | -0.214113 | 0.790882 | -0.206509 | 2.464645 | 0.503914 | -0.25507 | -0.03893 | 1.68036 | 0.295852 | -0.075008 | 0.503464 | 0.366318 | 0.479475 | 0.04876 | | 23.5 | -0.206509 | 0.141993 |
| 23.417 | 0.593539 | -0.010912 | -0.144576 | 0.312338 | 0.249146 | 0.098369 | 0.757548 | 0.279168 | 0.40231 | 0.160557 | 0.072004 | 1.405361 | -0.200101 | 2.695549 | 0.469119 | -0.29744 | 0.048949 | 1.303112 | 0.688298 | 0.021876 | 0.620237 | 0.568587 | 0.315965 | -0.15294 | | 23.51389 | -0.200101 | -0.154138 |
| 23.431 | 0.628043 | 0.136842 | -0.402677 | 0.269766 | 0.202823 | 0.118116 | 1.110529 | 0.386729 | 0.28865 | 0.231238 | 0.216134 | 1.443253 | -0.194051 | 1.16799 | 0.331701 | -0.326803 | -0.002733 | 0.869572 | 0.56153 | 0.162749 | 0.668243 | 0.524407 | 0.268991 | -0.382613 | | 23.52778 | -0.194051 | -0.429238 |
| 23.444 | 0.635582 | 0.355051 | -0.589378 | 0.220808 | 0.132601 | 0.067029 | 1.487317 | 0.309692 | 0.280704 | 0.255246 | 0.339172 | 1.095103 | -0.164302 | 0.751057 | 0.187972 | -0.568072 | -0.368 | 0.369552 | -0.126789 | 0.205738 | 0.553511 | 0.218363 | 0.290684 | -0.70792 | | 23.54167 | -0.164302 | -0.923717 |
| 23.458 | 0.542402 | 0.309386 | -0.601382 | 0.134698 | -0.036785 | -0.016419 | 1.474075 | 0.101857 | 0.32445 | 0.240055 | 0.472394 | 0.841695 | -0.124347 | -0.243211 | -0.267986 | -1.147058 | -1.001381 | -0.781157 | -0.756548 | -0.352936 | -0.117164 | -0.255117 | 0.165779 | -1.319188 | | 23.55556 | -0.124347 | -1.609461 |
| 23.472 | 0.359666 | 0.399754 | -0.345175 | 0.097793 | -0.085911 | 0.083744 | 1.160735 | -0.044836 | 0.303228 | 0.152121 | 0.592259 | 0.53715 | -0.087455 | -2.558473 | -0.590259 | -1.349741 | -1.081509 | -1.88852 | -1.146849 | -1.073294 | -0.84407 | -0.457208 | -0.44669 | -1.698788 | | 23.56944 | -0.087455 | -1.903538 |
| 23.486 | 0.155279 | 0.121548 | 0.1163 | 0.153068 | -0.105181 | 0.312773 | 0.878662 | -0.013441 | 0.302947 | 0.082889 | 0.338185 | 0.279891 | -0.099232 | -1.625589 | -0.634469 | -0.663404 | -0.887487 | -1.128756 | -0.886071 | -0.955125 | -0.967048 | 0.058184 | -0.781422 | -1.307761 | | 23.58333 | -0.099232 | -1.394161 |
| 23.5 | 0.056545 | 0.087123 | 0.447253 | 0.125959 | 0.001533 | 0.458399 | 0.671125 | 0.018498 | 0.148954 | 0.088505 | 0.057301 | 0.23319 | -0.157333 | 0.024389 | -0.292294 | 0.257255 | -0.52885 | -0.602385 | -0.36459 | -0.364042 | -0.609075 | 0.812138 | -0.548825 | -0.564525 | | 23.59722 | -0.157333 | -0.544272 |
| 23.514 | -0.032436 | 0.140708 | 0.462473 | 0.073255 | 0.074248 | 0.414889 | 0.509097 | -0.072089 | 0.055634 | 0.110166 | 0.05765 | 0.326482 | -0.208491 | 0.992851 | -0.031755 | 0.670073 | -0.227899 | 0.340402 | -0.021621 | 0.073103 | -0.127884 | 0.85468 | -0.258602 | -0.124495 | | 23.61111 | -0.208491 | -0.020742 |
| 23.528 | -0.200493 | 0.136582 | 0.22707 | -0.015819 | 0.050853 | 0.2812 | 0.324757 | 0.04894 | 0.041027 | 0.058964 | -0.117604 | 0.375574 | -0.220395 | 0.89467 | -0.071456 | 0.275238 | -0.1469 | 0.499781 | 0.021068 | 0.133 | -0.248355 | 0.597439 | -0.227499 | -0.265301 | | 23.625 | -0.220395 | -0.196198 |
| 23.542 | -0.402796 | -0.016274 | -0.009568 | -0.103612 | 0.005339 | 0.14287 | 0.095504 | 0.241113 | -0.013716 | 0.006846 | -0.232447 | 0.346333 | -0.120764 | 0.135317 | -0.145581 | -0.087084 | -0.154047 | 0.705341 | -0.107798 | 0.059785 | -0.405063 | 0.38676 | -0.266669 | -0.46108 | | 23.63889 | -0.120764 | -0.519798 |
| 23.556 | -0.383518 | -0.175141 | -0.118225 | -0.125355 | 0.012269 | 0.071672 | -0.209032 | 0.386068 | 0.100944 | 0.014379 | -0.188158 | 0.305754 | 0.127054 | -0.375598 | -0.032799 | -0.102507 | -0.050118 | 0.169657 | -0.160806 | 0.135288 | -0.120213 | 0.355996 | -0.135604 | -0.338825 | | 23.65278 | 0.127054 | -0.400112 |
| 23.569 | -0.151039 | -0.191429 | -0.162145 | -0.027737 | 0.037189 | 0.066817 | -0.392315 | 0.293135 | 0.218701 | 0.035223 | -0.089712 | 0.221359 | 0.363364 | -0.131616 | 0.093034 | -0.050739 | -0.012889 | 0.248925 | -0.07039 | 0.16412 | 0.107471 | 0.283355 | 0.047813 | -0.104145 | | 23.66667 | 0.363364 | -0.187317 |
| 23.583 | 0.154056 | 0.013478 | -0.186667 | 0.015249 | 0.028155 | 0.075356 | -0.326944 | 0.182384 | 0.213692 | 0.017641 | -0.027748 | 0.100908 | 0.297546 | 0.193963 | 0.04473 | -0.131968 | -0.100643 | -0.443676 | -0.099237 | 0.017857 | 0.011572 | 0.088145 | 0.098917 | -0.027116 | | 23.68056 | 0.297546 | -0.078395 |
| 23.597 | 0.408926 | 0.203783 | -0.147431 | 0.051449 | 0.006817 | 0.083044 | -0.121471 | -0.013749 | 0.149514 | -0.005999 | -0.014209 | -0.001139 | 0.113345 | -0.12493 | -0.07937 | -0.244468 | -0.188942 | -1.940882 | -0.084288 | -0.116344 | -0.129536 | -0.103366 | 0.085848 | -0.015337 | | 23.69444 | 0.113345 | -0.00803 |
| 23.611 | 0.570291 | 0.32323 | -0.061914 | 0.131417 | 0.013268 | 0.099249 | 0.023236 | -0.081028 | 0.166511 | 0.008552 | 0.028232 | -0.015769 | 0.047946 | -0.607905 | -0.115015 | -0.234733 | -0.175366 | -1.84736 | 0.014286 | -0.045246 | -0.130512 | -0.152044 | 0.141877 | 0.065546 | | 23.70833 | 0.047946 | 0.077954 |
| 23.625 | 0.571499 | 0.282263 | -0.003289 | 0.208509 | 0.025104 | 0.110542 | -0.029393 | -0.133546 | 0.089237 | 0.034901 | 0.129071 | 0.022242 | 0.103053 | -0.30012 | -0.089165 | -0.135673 | -0.085985 | -1.217267 | 0.109844 | 0.174578 | -0.104801 | -0.155999 | 0.190112 | 0.109321 | | 23.72222 | 0.103053 | 0.086345 |
| 23.639 | 0.412902 | 0.155777 | 0.005391 | 0.185021 | 0.019978 | 0.106924 | -0.173922 | -0.141174 | 0.009714 | 0.047241 | 0.212521 | 0.102991 | 0.141604 | 0.250047 | -0.111357 | -0.110727 | -0.003515 | -0.307311 | 0.10596 | 0.333654 | -0.163327 | -0.282624 | 0.143849 | 0.080978 | | 23.73611 | 0.141604 | 0.002122 |
| 23.653 | 0.223813 | -0.015212 | -0.020833 | 0.113114 | 0.014124 | 0.094008 | -0.313648 | -0.064848 | -0.073105 | 0.044164 | 0.150269 | 0.198376 | 0.123416 | 0.418534 | -0.18541 | -0.225685 | 0.022498 | 0.638626 | 0.00919 | 0.370541 | -0.332084 | -0.496955 | 0.03712 | -0.029506 | | 23.75 | 0.123416 | -0.102514 |
| 23.667 | 0.089361 | -0.186313 | -0.009389 | 0.046399 | 0.017583 | 0.073257 | -0.442956 | 0.001519 | -0.11216 | 0.028636 | 0.059065 | 0.25578 | 0.076499 | 0.469533 | -0.203178 | -0.321653 | 0.047644 | 1.583159 | 0.028828 | 0.338095 | -0.26203 | -0.648443 | 0.093623 | -0.06871 | | 23.76389 | 0.076499 | -0.067844 |
| 23.681 | -0.015819 | -0.278196 | 0.058233 | 0.007543 | 0.037227 | 0.054639 | -0.499388 | 0.053951 | -0.122267 | 0.020916 | 0.016758 | 0.283615 | 0.017081 | 0.476068 | -0.135647 | -0.264778 | 0.064067 | 1.919965 | 0.141983 | 0.32085 | 0.052192 | -0.577065 | 0.402181 | 0.046327 | | 23.77778 | 0.017081 | 0.129245 |
| 23.694 | -0.134496 | -0.206294 | 0.136126 | -0.005876 | 0.058997 | 0.021579 | -0.419499 | 0.126573 | -0.136482 | 0.015729 | 0.008436 | 0.260721 | -0.057204 | 0.27996 | -0.070419 | -0.147834 | 0.120656 | 1.090337 | 0.309017 | 0.292753 | 0.259082 | -0.496678 | 0.649138 | 0.142683 | | 23.79167 | -0.057204 | 0.248613 |
| 23.708 | -0.3095 | -0.177642 | 0.101389 | 0.010236 | 0.066461 | -0.041125 | -0.317264 | 0.162317 | -0.170675 | 0.007148 | 0.033693 | 0.172465 | -0.086803 | -0.148597 | -0.054914 | -0.050587 | 0.206389 | 0.400965 | 0.408068 | 0.188698 | 0.275139 | -0.328004 | 0.606993 | 0.12222 | | 23.80556 | -0.086803 | 0.223627 |
| 23.722 | -0.449117 | -0.125208 | -0.047691 | 0.027276 | 0.0525 | -0.097546 | -0.361693 | 0.119643 | -0.152828 | 0.000669 | 0.113038 | 0.039457 | -0.042656 | -0.696605 | 0.017665 | 0.090344 | 0.285195 | 0.091975 | 0.394132 | 0.243228 | 0.246554 | -0.106893 | 0.505693 | 0.177471 | | 23.81944 | -0.042656 | 0.271692 |
| 23.736 | -0.556913 | -0.134882 | -0.211433 | 0.026799 | 0.036861 | -0.113635 | -0.656223 | 0.055592 | -0.028698 | -0.00847 | 0.080992 | -0.091844 | 0.026366 | -1.233731 | 0.1667 | 0.253316 | 0.396693 | -0.089999 | 0.419492 | 0.135761 | 0.206381 | 0.04617 | 0.516696 | 0.325263 | | 23.83333 | 0.026366 | 0.397593 |
| 23.75 | -0.649499 | 0.005457 | -0.293028 | -0.021651 | 0.019267 | -0.078032 | -0.859216 | -0.062941 | 0.101757 | -0.013548 | 0.100109 | -0.081625 | 0.082841 | -0.621252 | 0.267868 | 0.309352 | 0.429332 | -0.320606 | 0.53403 | -0.015063 | 0.154178 | 0.049938 | 0.535548 | 0.459305 | | 23.84722 | 0.082841 | 0.535642 |
| 23.764 | -0.289556 | 0.138603 | -0.282683 | -0.107949 | 0.000175 | -0.055852 | -0.798185 | -0.169623 | 0.116666 | -0.038145 | 0.103946 | 0.033909 | 0.156005 | 0.431691 | 0.200491 | 0.22408 | 0.374808 | -0.643777 | 0.399612 | -0.012328 | 0.060693 | -0.045867 | 0.47954 | 0.498379 | | 23.86111 | 0.156005 | 0.596583 |
| 23.778 | 0.265893 | 0.184575 | -0.264721 | -0.159353 | 0.00073 | -0.025965 | -0.581963 | -0.19847 | 0.085025 | -0.058333 | 0.106334 | 0.068685 | 0.193144 | 1.305765 | -0.039997 | 0.142163 | 0.350258 | -0.509533 | -0.088641 | 0.124977 | -0.00721 | -0.218985 | 0.3852 | 0.470546 | | 23.875 | 0.193144 | 0.582135 |
| 23.792 | 0.47436 | 0.204286 | -0.243934 | -0.155113 | 0.022715 | 0.001644 | -0.374038 | -0.182174 | -0.035924 | -0.080503 | 0.09779 | -0.090562 | 0.149288 | 1.382852 | -0.389169 | 0.1415 | 0.284143 | -0.452578 | -0.039319 | 0.201913 | -0.055428 | -0.408806 | -0.682514 | 0.363864 | | 23.88889 | 0.149288 | 0.416487 |
| 23.806 | 0.118858 | 0.063327 | -0.253772 | -0.149932 | 0.016828 | -0.040635 | -0.304114 | -0.196504 | -0.245065 | -0.125369 | -0.039184 | -0.364787 | -0.00634 | 0.518522 | -0.671607 | 0.176411 | 0.16694 | -0.522001 | 0.247049 | -0.078023 | -0.067736 | -0.47669 | -0.408422 | 0.314013 | | 23.90278 | -0.00634 | 0.304187 |
| 23.819 | -0.375243 | -0.122686 | -0.274835 | -0.153095 | -0.043921 | -0.159499 | -0.51513 | -0.352833 | -0.456961 | -0.18285 | -0.140043 | -0.543496 | -0.205915 | -0.455932 | -0.804746 | 0.191508 | 0.091947 | -0.573971 | 0.173932 | -0.31714 | -0.112213 | -0.479937 | 0.603938 | 0.316951 | | 23.91667 | -0.205915 | 0.288718 |
| 23.833 | -0.978504 | -0.243978 | -0.267853 | -0.198203 | -0.126678 | -0.29822 | -0.917808 | -0.490435 | -0.53125 | -0.202331 | -0.110109 | -0.559358 | -0.339292 | -1.225913 | -0.71773 | 0.280493 | 0.211062 | -0.772677 | -0.060208 | -0.396236 | -0.220482 | -0.577068 | 1.536813 | 0.228742 | | 23.93056 | -0.339292 | 0.190956 |
| 23.847 | -1.468414 | -0.302257 | -0.38773 | -0.288437 | -0.191811 | -0.381061 | -1.030972 | -0.681434 | -0.532915 | -0.171673 | -0.286598 | -0.373377 | -0.340469 | -2.167535 | -0.588837 | 0.300515 | 0.270664 | -0.996497 | -0.238262 | -0.392099 | -0.302881 | -0.631253 | 1.550016 | 0.050895 | | 23.94444 | -0.340469 | -0.08536 |
| 23.861 | -1.358921 | -0.203325 | -0.409772 | -0.385123 | -0.217145 | -0.349051 | -0.621107 | -0.809688 | -0.425109 | -0.107416 | -0.602479 | -0.218575 | -0.18005 | -2.997868 | -0.560968 | 0.275386 | 0.184198 | -1.38532 | -0.165774 | -0.46185 | -0.431644 | -0.792768 | 0.124798 | -0.136493 | | 23.95833 | -0.18005 | -0.402572 |
| 23.875 | -0.938193 | -0.158514 | -0.222565 | -0.488648 | -0.216555 | -0.220109 | -0.32845 | -0.842837 | -0.341135 | -0.066721 | -0.489461 | -0.386185 | 0.001705 | -3.138735 | -0.593976 | 0.177888 | 0.115818 | -2.212923 | 0.029964 | -0.471847 | -0.500398 | -1.093732 | -0.695139 | -0.358568 | | 23.97222 | 0.001705 | -0.526291 |
| 23.889 | -0.536441 | -0.164679 | -0.066227 | -0.486658 | -0.206965 | -0.146361 | -0.528543 | -1.010056 | -0.29165 | -0.085858 | -0.588819 | -0.720338 | 0.035498 | -2.363998 | -0.623443 | -0.013672 | 0.098229 | -2.308943 | 0.003811 | -0.435629 | -0.514618 | -1.383105 | -1.60478 | -0.6271 | | 23.98611 | 0.035498 | -0.764076 |
| 23.903 | -0.184508 | -0.108198 | -0.296683 | -0.302441 | -0.195055 | -0.097979 | -0.736265 | -1.232028 | -0.252427 | -0.116757 | -0.629628 | -0.759296 | -0.016294 | -1.295272 | -0.605632 | -0.206313 | 0.129861 | -1.865008 | -0.024067 | -0.50687 | -0.623676 | -1.478162 | -1.347726 | -0.762026 | | 24 | -0.016294 | -1.020102 |
| 23.917 | 0.144083 | 0.077988 | -0.472123 | -0.029775 | -0.117601 | -0.00791 | -0.574739 | -0.869008 | -0.124492 | -0.068436 | -0.398867 | -0.411766 | 0.061173 | -0.311474 | -0.479117 | -0.174466 | 0.209263 | -1.119201 | 0.236527 | -0.468343 | -0.546349 | -1.005706 | -0.706082 | -0.669506 | | 24.01389 | 0.061173 | -0.923705 |
| 23.931 | 0.37117 | 0.293273 | -0.292198 | 0.104763 | 0.052327 | 0.161515 | -0.171932 | -0.211253 | 0.080622 | 0.100794 | 0.086171 | 0.024119 | 0.215383 | 0.421774 | -0.207112 | 0.037738 | 0.334482 | -0.097853 | 0.670853 | -0.204846 | -0.332887 | -0.296056 | -0.219771 | -0.299023 | | 24.02778 | 0.215383 | -0.606046 |
| 23.944 | 0.3569 | 0.329089 | -0.051823 | 0.224162 | 0.11927 | 0.247795 | 0.105871 | -0.05764 | 0.089222 | 0.172253 | 0.343945 | -0.033937 | 0.240925 | 0.577578 | -0.169363 | 0.004941 | 0.269896 | 0.546894 | 0.633638 | -0.173826 | -0.293779 | -0.406139 | 0.030039 | -0.422825 | | 24.04167 | 0.240925 | -0.710957 |
| 23.958 | 0.028762 | 0.076556 | -0.015552 | 0.068106 | -0.051795 | -0.013397 | -0.256518 | -0.789991 | -0.229503 | -0.068186 | 0.32713 | -0.590808 | 0.033252 | 0.04511 | -0.507423 | -0.317681 | -0.15114 | 0.509979 | -0.191447 | -0.491249 | -0.838044 | -0.969767 | -0.014058 | -0.971472 | | 24.05556 | 0.033252 | -1.128312 |
| 23.972 | -0.379073 | -0.197555 | 0.046072 | -0.170081 | -0.239455 | -0.772949 | -1.014595 | -1.706136 | -0.840712 | -0.515164 | 0.587139 | -1.581969 | -0.173764 | -0.276095 | -0.528041 | -0.422887 | -0.461671 | 0.308476 | -1.251149 | -0.498956 | -1.873259 | -1.123617 | 0.358481 | -1.094178 | | 24.06944 | -0.173764 | -1.387515 |
| 23.986 | -0.525979 | -0.378262 | -0.079111 | -0.297424 | -0.317074 | -1.168322 | -2.551015 | -1.978048 | -0.98969 | -0.597077 | 0.497285 | -1.20588 | -0.328285 | -0.431179 | -0.328317 | -0.276878 | -0.298159 | 0.454903 | -1.917454 | -0.284625 | -2.298239 | -1.327645 | 0.609501 | -1.018295 | | 24.08333 | -0.328285 | -1.357173 |
| 24 | -0.319364 | -0.526881 | -0.274994 | -0.326298 | -0.344303 | -0.562328 | -3.190758 | -1.311533 | -0.689404 | -0.363689 | -1.073761 | -1.788442 | -0.495812 | -0.988801 | -0.591228 | -0.407529 | -0.283954 | 0.982678 | -2.890735 | -0.354139 | -1.583051 | -1.985309 | 0.263325 | -1.315362 | | 24.09722 | -0.495812 | -1.781278 |
| 24.014 | 0.058007 | -0.426808 | -0.241465 | -0.218659 | -0.290361 | -0.015518 | -1.940565 | -0.510657 | -0.434491 | -0.108535 | -1.489909 | -0.920791 | -0.422628 | -1.333799 | -1.014664 | -0.502649 | -0.419072 | -0.002785 | -2.636286 | -0.460471 | -1.145208 | -1.943467 | 0.231866 | -1.431261 | | 24.11111 | -0.422628 | -1.768501 |
| 24.028 | 0.478014 | -0.066468 | 0.112677 | -0.097156 | -0.120748 | 0.40171 | -0.554788 | 0.199321 | -0.062724 | 0.167564 | -0.533804 | -0.187478 | -0.045255 | -0.876698 | -0.301518 | -0.324087 | -0.222585 | -0.528845 | -1.110847 | -0.407706 | -0.637453 | -0.8483 | 0.008707 | -0.957125 | | 24.125 | -0.045255 | -1.177367 |
| 24.042 | 0.786507 | 0.258884 | 0.481819 | -0.032304 | 0.06165 | 0.560208 | 0.363372 | 0.821028 | 0.293682 | 0.374179 | 0.166427 | 0.320105 | 0.357524 | -0.474765 | -0.244947 | -0.144586 | -0.044834 | -0.320856 | -0.71261 | -0.362548 | -0.206973 | -0.108111 | -0.081122 | -0.49083 | | 24.13889 | 0.357524 | -0.627275 |
| 24.056 | 0.685535 | 0.335257 | 0.371689 | 0.00073 | 0.198273 | 0.474245 | 1.019741 | 0.935199 | 0.582248 | 0.335888 | 0.548092 | 0.468732 | 0.498018 | -0.356712 | -0.216439 | -0.077655 | -0.0208 | -0.195614 | -0.419345 | -0.395129 | 0.003386 | 0.228131 | 0.019529 | -0.296811 | | 24.15278 | 0.498018 | -0.408066 |
| 24.069 | 0.110935 | 0.14528 | -0.063792 | 0.057014 | 0.196747 | 0.305093 | 1.296066 | 0.706122 | 0.632579 | 0.151642 | 0.830169 | 0.445441 | 0.404604 | -0.276665 | -0.028887 | -0.012572 | -0.119508 | -0.25016 | -0.177904 | -0.580484 | 0.034314 | 0.333072 | 0.103288 | -0.266506 | | 24.16667 | 0.404604 | -0.331147 |
| 24.083 | -0.417764 | -0.180395 | -0.27713 | 0.154286 | 0.055711 | 0.121178 | 0.97941 | 0.542848 | 0.587225 | 0.026518 | 1.020669 | 0.497256 | 0.294782 | -0.204655 | 0.184096 | -0.006867 | -0.178403 | -0.477811 | -0.143997 | -0.751849 | 0.001433 | 0.402483 | 0.140716 | -0.250402 | | 24.18056 | 0.294782 | -0.234092 |
| 24.097 | -0.403806 | -0.325023 | 0.145132 | 0.176334 | -0.077249 | 0.000757 | 0.20849 | 0.616323 | 0.183473 | -0.024998 | 0.855132 | 0.507735 | 0.282796 | -0.122632 | 0.33433 | 0.008439 | -0.126722 | -0.803946 | -0.099709 | -0.706003 | 0.008742 | 0.592362 | 0.082059 | -0.037462 | | 24.19444 | 0.282796 | 0.000283 |
| 24.111 | -0.023693 | -0.118138 | 0.399846 | 0.057995 | -0.099095 | 0.014782 | -0.471182 | 0.8099 | 0.075136 | 0.027478 | -0.01786 | 0.340613 | 0.321335 | 0.147506 | 0.531059 | 0.032222 | -0.01718 | -0.859672 | 0.08546 | -0.409181 | 0.154998 | 0.886931 | -0.048872 | 0.267041 | | 24.20833 | 0.321335 | 0.36684 |
| 24.125 | 0.186366 | 0.343678 | 0.042075 | -0.020381 | -0.011613 | 0.119696 | -0.217213 | 0.901561 | 0.269305 | 0.128826 | -0.005934 | 0.028491 | 0.414824 | 0.521015 | 0.748689 | 0.012869 | 0.018325 | -0.681427 | 0.300172 | -0.012763 | 0.472805 | 1.168697 | -0.08196 | 0.567864 | | 24.22222 | 0.414824 | 0.901652 |
| 24.139 | 0.200581 | 0.584805 | -0.355916 | 0.001672 | 0.069199 | 0.205057 | 0.608786 | 0.733846 | 0.405213 | 0.125888 | 0.251005 | -0.191242 | 0.431401 | 1.017141 | 1.016485 | 0.075731 | 0.013629 | -0.413134 | 0.406873 | 0.497438 | 0.884595 | 0.945887 | 0.036947 | 0.967585 | | 24.23611 | 0.431401 | 1.404833 |
| 24.153 | 0.141448 | 0.128487 | -0.287 | 0.081196 | 0.123705 | 0.25181 | 1.487436 | 0.559767 | 0.061935 | 0.055297 | 0.072656 | -0.01638 | 0.330327 | 1.350404 | 0.961617 | 0.326996 | 0.060892 | -0.199631 | 0.580099 | 1.143329 | 1.278656 | 0.301045 | 0.200465 | 1.287425 | | 24.25 | 0.330327 | 1.84863 |
| 24.167 | 0.11718 | -0.238404 | 0.060362 | 0.164043 | 0.170855 | 0.23068 | 0.965708 | 0.437087 | -0.270926 | 0.031974 | -0.341307 | 0.438709 | 0.113266 | 1.562728 | 0.420663 | 0.57738 | 0.222806 | 0.019437 | 0.860016 | 1.62428 | 1.461819 | -0.002087 | 0.362267 | 1.402096 | | 24.26389 | 0.113266 | 1.883964 |
| 24.181 | 0.214466 | -0.101331 | 0.105688 | 0.268102 | 0.15814 | 0.139507 | 0.168456 | 0.305542 | -0.212559 | 0.065808 | -0.490562 | 0.772646 | 0.007005 | 1.383573 | -0.239943 | 0.572082 | 0.447155 | 0.172314 | 0.635977 | 1.732993 | 1.474694 | -0.263674 | 0.615107 | 1.274858 | | 24.27778 | 0.007005 | 1.571137 |
| 24.194 | 0.23699 | 0.288247 | -0.066632 | 0.257499 | 0.083779 | -0.016611 | 0.256193 | 0.395141 | 0.054778 | 0.078297 | -0.307827 | 0.59736 | 0.193524 | 1.086703 | -0.953546 | 0.286796 | 0.531119 | 0.205608 | 0.009892 | 1.398043 | 1.291097 | -0.473854 | 0.697271 | 0.918641 | | 24.29167 | 0.193524 | 1.188096 |
| 24.208 | 0.139906 | 0.215933 | -0.122369 | 0.264373 | 0.05275 | -0.118994 | 0.78617 | 0.575407 | 0.164457 | -0.009668 | -0.129612 | 0.285539 | 0.379821 | 0.636418 | -1.052076 | 0.101023 | 0.446043 | 0.269029 | -0.343455 | 1.040093 | 1.068574 | -0.199804 | 0.631225 | 0.717973 | | 24.30556 | 0.379821 | 0.902106 |
| 24.222 | 0.117032 | -0.141202 | 0.135253 | 0.262114 | 0.099565 | -0.036829 | 0.82959 | 0.713539 | 0.376229 | -0.056649 | -0.0616 | 0.29494 | 0.411295 | 0.284246 | -0.622288 | 0.138169 | 0.213938 | 0.372801 | -0.389597 | 0.639348 | 1.066763 | 0.670193 | 0.463224 | 0.737898 | | 24.31944 | 0.411295 | 0.917588 |
| 24.236 | 0.238744 | -0.181627 | 0.374644 | 0.185646 | 0.150642 | 0.109674 | 0.765251 | 0.747753 | 0.431237 | 0.019763 | -0.062463 | 0.371953 | 0.33385 | 0.129069 | -0.145053 | 0.174442 | 0.016055 | 0.396719 | -0.247816 | 0.205918 | 0.882589 | 1.302914 | 0.116763 | 0.768101 | | 24.33333 | 0.33385 | 0.960596 |
| 24.25 | 0.352879 | 0.067972 | 0.354785 | 0.137665 | 0.112866 | 0.192702 | 0.897007 | 0.54755 | 0.171316 | 0.110651 | -0.00654 | 0.312118 | 0.219509 | 0.219305 | 0.222559 | 0.259254 | -0.017079 | 0.441056 | -0.006213 | 0.044934 | 0.893621 | 1.358159 | -0.151662 | 0.681318 | | 24.34722 | 0.219509 | 0.999788 |
| 24.264 | 0.299388 | 0.213899 | 0.336798 | 0.058827 | 0.056245 | 0.181087 | 1.515655 | 0.267416 | 0.069475 | 0.090599 | 0.192404 | 0.085804 | 0.169024 | 0.512759 | 0.431802 | 0.305397 | 0.087524 | 0.45905 | 0.238242 | 0.318458 | 0.893618 | 1.033135 | -0.123499 | 0.532888 | | 24.36111 | 0.169024 | 0.753739 |
| 24.278 | 0.10493 | 0.119476 | 0.301972 | -0.035719 | 0.045254 | 0.145001 | 1.890044 | 0.305509 | 0.136505 | 0.049947 | 0.33219 | -0.067314 | 0.243478 | 0.663043 | 0.43305 | 0.273471 | 0.224039 | 0.37305 | 0.078888 | 0.591892 | 0.757784 | 0.125451 | 0.171622 | 0.525956 | | 24.375 | 0.243478 | 0.673555 |
| 24.292 | -0.138443 | 0.085736 | 0.422395 | -0.028096 | 0.055593 | 0.13843 | 1.478913 | 0.541045 | 0.156632 | 0.056624 | 0.233058 | 0.044086 | 0.289379 | 0.586422 | 0.314996 | 0.182693 | 0.361499 | 0.237942 | -0.335886 | 0.818753 | 0.493132 | -0.50672 | 0.549439 | 0.442439 | | 24.38889 | 0.289379 | 0.557677 |
| 24.306 | -0.151679 | 0.244291 | 0.395133 | 0.02799 | 0.104167 | 0.168093 | 1.090169 | 0.53816 | 0.152427 | 0.094854 | 0.065336 | 0.216557 | 0.275051 | 0.568131 | 0.163515 | 0.165573 | 0.385606 | 0.33048 | -0.155293 | 0.993326 | 0.181512 | -0.381839 | 0.675444 | 0.376996 | | 24.40278 | 0.275051 | 0.514483 |
| 24.319 | 0.232381 | 0.415094 | 0.29511 | 0.124205 | 0.181161 | 0.21303 | 0.948922 | 0.423111 | 0.273527 | 0.134373 | -0.03503 | 0.284964 | 0.329043 | 0.616567 | -0.075125 | 0.201996 | 0.267779 | 0.526675 | 0.761839 | 0.980165 | 0.009138 | 0.305269 | 0.536313 | 0.362209 | | 24.41667 | 0.329043 | 0.506415 |
| 24.333 | 0.694577 | 0.305783 | 0.460522 | 0.217912 | 0.164875 | 0.21205 | 0.763993 | 0.334129 | 0.486049 | 0.151012 | -0.023703 | 0.334373 | 0.413118 | 0.57513 | -0.414402 | 0.197585 | 0.128511 | 0.68282 | 1.390553 | 0.761306 | -0.057489 | 0.542636 | 0.449243 | 0.250143 | | 24.43056 | 0.413118 | 0.388596 |
| 24.347 | 0.936213 | 0.020674 | 0.491481 | 0.217445 | 0.075774 | 0.179624 | 0.553648 | 0.414864 | 0.488091 | 0.10652 | 0.072375 | 0.37682 | 0.339468 | 0.583089 | -0.569612 | 0.136349 | 0.056143 | 0.766202 | 0.761683 | 0.390987 | -0.052771 | 0.291674 | 0.361173 | 0.158162 | | 24.44444 | 0.339468 | 0.254776 |
| 24.361 | 0.879959 | -0.20922 | -0.012624 | 0.20537 | -0.029556 | 0.101123 | 0.593759 | 0.446967 | 0.3332 | -0.037013 | 0.286927 | 0.373561 | 0.120335 | 0.828141 | -0.400291 | 0.036618 | 0.062202 | 0.834906 | -0.002903 | 0.126721 | 0.099712 | 0.034525 | 0.298032 | 0.123693 | | 24.45833 | 0.120335 | 0.176355 |
| 24.375 | 0.422878 | -0.187563 | -0.233748 | 0.090595 | -0.183335 | -0.047021 | 0.932375 | 0.191434 | 0.221611 | -0.200287 | 0.432342 | 0.280251 | -0.080958 | 1.015942 | -0.114599 | -0.082129 | 0.140877 | 0.828421 | -0.6465 | 0.074543 | 0.313665 | -0.088615 | 0.232535 | 0.107684 | | 24.47222 | -0.080958 | 0.172303 |
| 24.389 | 0.096732 | 0.090378 | -0.193578 | 0.064799 | -0.184787 | -0.076431 | 1.49096 | 0.093853 | 0.283563 | -0.158033 | 0.605041 | 0.260213 | -0.092182 | 0.568701 | 0.124817 | -0.047412 | 0.278992 | 0.8021 | -0.815196 | 0.124494 | 0.470428 | -0.05428 | -0.088171 | 0.179952 | | 24.48611 | -0.092182 | 0.233753 |
| 24.403 | 0.536257 | 0.361748 | 0.326015 | 0.206891 | 0.0876 | 0.097758 | 1.642086 | 0.273494 | 0.404454 | 0.025393 | 1.107876 | 0.526492 | 0.001035 | 0.39764 | 0.3209 | 0.095729 | 0.358595 | 0.943693 | -0.309274 | 0.37427 | 0.648528 | 0.525141 | 0.285804 | 0.26126 | | 24.5 | 0.001035 | 0.316523 |
| 24.417 | 1.350061 | 0.358826 | 0.219583 | 0.289533 | 0.380467 | 0.29257 | 1.168546 | 0.561928 | 0.40474 | 0.09802 | 0.579508 | 0.817149 | 0.22359 | 0.418013 | 0.344209 | 0.204362 | 0.303864 | 1.042267 | 0.242201 | 0.4782 | 0.742741 | 0.731902 | 0.618265 | 0.168641 | | 24.51389 | 0.22359 | 0.198132 |
| 24.431 | 2.033882 | 0.137342 | 0.105169 | 0.260556 | 0.471093 | 0.38176 | 0.709557 | 0.63476 | 0.349027 | 0.079063 | 0.131052 | 0.87209 | 0.234735 | 0.275167 | 0.178346 | 0.134802 | 0.145565 | 1.032831 | 0.628959 | 0.293904 | 0.634077 | 0.585974 | 0.485871 | -0.073326 | | 24.52778 | 0.234735 | -0.040792 |
| 24.444 | 1.862094 | 0.228895 | -0.447965 | 0.267768 | 0.370502 | 0.38145 | 0.375313 | 0.557146 | 0.322977 | 0.113636 | 0.006692 | 0.951141 | -0.037744 | -0.050901 | -0.118656 | -0.139486 | -0.187365 | 0.695846 | 0.650093 | -0.081465 | 0.384716 | 0.474219 | -0.20591 | -0.342459 | | 24.54167 | -0.037744 | -0.364892 |
| 24.458 | 1.288225 | 0.411324 | -1.411458 | 0.228383 | 0.241165 | 0.286574 | 0.352052 | 0.30721 | 0.29475 | 0.145815 | 0.106613 | 1.00264 | -0.215307 | -0.308765 | -0.503365 | -0.607712 | -0.699054 | 0.227347 | 0.44344 | -0.588145 | 0.228144 | 0.175205 | -0.708054 | -0.66171 | | 24.55556 | -0.215307 | -0.739843 |
| 24.472 | 0.593576 | 0.363622 | -1.329723 | 0.083769 | 0.150997 | 0.267589 | 0.910273 | 0.122556 | 0.231734 | 0.079387 | -0.047154 | 0.867182 | -0.174444 | -0.199692 | -0.594502 | -0.703979 | -0.841746 | 0.261252 | 0.073371 | -0.612167 | 0.176168 | -0.044323 | -0.657922 | -0.778757 | | 24.56944 | -0.174444 | -0.831507 |
| 24.486 | 0.033263 | 0.393916 | -0.776058 | 0.113182 | 0.103742 | 0.32373 | 1.521966 | 0.102066 | 0.322861 | 0.023345 | 0.321024 | 0.714225 | -0.029019 | 0.259864 | -0.226509 | -0.137093 | -0.385483 | 0.369242 | -0.202543 | -0.001821 | 0.183835 | 0.404663 | -0.227829 | -0.455898 | | 24.58333 | -0.029019 | -0.481908 |
| 24.5 | -0.160908 | 0.321072 | -0.135414 | 0.145895 | 0.151412 | 0.322937 | 1.286406 | -0.018304 | 0.395806 | 0.050482 | 0.705595 | 0.689013 | 0.030133 | 0.568996 | 0.178843 | 0.576854 | 0.053413 | 0.475661 | 0.17989 | 0.472434 | 0.324828 | 0.904701 | 0.114697 | -0.027006 | | 24.59722 | 0.030133 | -0.092528 |
| 24.514 | -0.153704 | 0.327315 | 0.265366 | 0.022895 | 0.150558 | 0.281588 | 0.835333 | -0.24502 | 0.313782 | 0.048807 | 0.5097 | 0.583371 | -0.024048 | 0.493775 | 0.021295 | 0.600088 | 0.070702 | 0.274702 | 0.317199 | 0.369758 | 0.195831 | 0.812837 | 0.058768 | -0.064746 | | 24.61111 | -0.024048 | -0.123411 |
| 24.528 | -0.057049 | 0.271661 | 0.376899 | -0.102674 | -0.007999 | 0.231316 | 0.527154 | -0.163935 | 0.107258 | -0.010165 | 0.150109 | 0.410358 | -0.080536 | 0.302268 | -0.552411 | 0.258329 | -0.12508 | -0.011427 | 0.104289 | 0.06152 | -0.081903 | 0.50975 | -0.193764 | -0.345439 | | 24.625 | -0.080536 | -0.344153 |
| 24.542 | 0.044237 | 0.120181 | 0.419652 | -0.126244 | -0.084446 | 0.220721 | 0.227675 | 0.198591 | 0.074223 | -0.030837 | -0.054265 | 0.26784 | -0.072443 | 0.1346 | -0.753621 | 0.13683 | -0.205058 | -0.086341 | -0.088271 | -0.041537 | -0.195189 | 0.51603 | -0.268209 | -0.427735 | | 24.63889 | -0.072443 | -0.379901 |
| 24.556 | -0.007733 | -0.003286 | 0.386779 | -0.111418 | 0.105045 | 0.191808 | -0.153583 | 0.536764 | 0.03697 | -0.006923 | -0.148873 | 0.260876 | -0.062858 | -0.809561 | -0.474764 | 0.200053 | -0.116893 | -0.018024 | -0.042019 | 0.020746 | -0.133069 | 0.642999 | -0.155816 | -0.246925 | | 24.65278 | -0.062858 | -0.200382 |
| 24.569 | -0.09068 | -0.077634 | 0.157556 | -0.056421 | 0.033636 | 0.127118 | -0.463681 | 0.494128 | 0.144561 | 0.017348 | -0.205672 | 0.290483 | -0.027672 | -0.723234 | -0.204446 | 0.158021 | -0.039483 | 0.031727 | 0.004949 | 0.032554 | -0.099355 | 0.57234 | -0.063864 | -0.110103 | | 24.66667 | -0.027672 | -0.034737 |
| 24.583 | -0.118659 | -0.072968 | -0.09634 | 0.064784 | -0.01107 | 0.078497 | -0.53543 | 0.070177 | 0.086132 | 0.01444 | -0.278174 | 0.215827 | -0.008707 | -0.096148 | -0.08815 | 0.026097 | -0.0021 | 0.020527 | 0.006646 | 0.019405 | -0.09172 | 0.362749 | -0.013072 | -0.026714 | | 24.68056 | -0.008707 | 0.068694 |
| 24.597 | 0.053072 | 0.020772 | -0.301159 | 0.190507 | -0.01267 | 0.062863 | -0.446139 | -0.336044 | -0.012334 | 0.000789 | -0.337298 | 0.020242 | -0.059241 | 0.326921 | 0.013376 | -0.03341 | 0.06885 | 0.054395 | 0.091132 | 0.086096 | 0.070656 | 0.311144 | 0.069516 | 0.114909 | | 24.69444 | -0.059241 | 0.205621 |

| | | | | | | | | | | | | | | | | | | | | | | | | |
|---|---|---|---|---|---|---|---|---|---|---|---|---|---|---|---|---|---|---|---|---|---|---|---|---|
| 24.611 | 0.208249 | 0.138966 | -0.353535 | 0.162621 | 0.003024 | 0.054635 | -0.213269 | -0.458313 | 0.074641 | 0.000109 | -0.21467 | -0.126438 | -0.094399 | 0.430316 | 0.149708 | 0.006652 | 0.171636 | 0.121457 | 0.181502 | 0.196217 | 0.258463 | 0.33728 | 0.143564 | 0.305444 |
| 24.625 | 0.223277 | 0.175656 | -0.209708 | 0.103645 | 0.032729 | 0.039819 | 0.136029 | -0.309553 | -0.078256 | 0.018134 | -0.220726 | -0.108642 | -0.047513 | 0.371798 | 0.222949 | 0.061173 | 0.198648 | 0.16132 | 0.19182 | 0.228865 | 0.320281 | 0.374538 | 0.136708 | 0.388652 |
| 24.639 | 0.149283 | 0.123778 | -0.055433 | 0.040472 | 0.03007 | 0.044317 | 0.341605 | -0.103426 | -0.031398 | 0.035174 | -0.186201 | 0.099373 | 0.059355 | 0.258762 | 0.226888 | 0.098946 | 0.146378 | 0.166497 | 0.136493 | 0.216753 | 0.29867 | 0.569421 | 0.135551 | 0.108338 |
| 24.653 | 0.088457 | 0.04237 | 0.083895 | -0.009672 | 0.027423 | 0.074303 | 0.250752 | 0.062749 | 0.01463 | 0.029096 | -0.191273 | 0.338068 | 0.125215 | 0.18494 | 0.196415 | 0.108642 | 0.058768 | 0.15131 | 0.05956 | 0.176036 | 0.252753 | 0.570602 | 0.16211 | -0.487959 |
| 24.667 | 0.042552 | -0.049492 | 0.152318 | -0.068412 | 0.018796 | 0.102405 | 0.000493 | 0.143522 | 0.073284 | 0.018556 | -0.152548 | 0.45375 | 0.179074 | 0.127587 | 0.152763 | 0.060933 | -0.006695 | 0.115447 | 0.013756 | 0.119452 | 0.187096 | -1.227154 | 0.144792 | -0.92564 |
| 24.681 | -0.07174 | -0.159416 | 0.140338 | -0.121447 | -0.000775 | 0.076496 | -0.311607 | 0.220282 | 0.122868 | 0.003099 | -0.093003 | 0.415578 | 0.194058 | 0.057514 | 0.091315 | 0.020822 | -0.027054 | 0.071746 | 0.016426 | 0.056526 | 0.112203 | -1.225705 | 0.076037 | -0.37433 |
| 24.694 | -0.242134 | -0.278297 | 0.015581 | -0.174702 | -0.018354 | -0.03119 | -0.710303 | 0.219467 | -0.033112 | -0.017478 | -0.073524 | 0.292719 | 0.132707 | -0.019339 | 0.04864 | 0.001112 | -0.024436 | 0.021129 | 0.035392 | -0.03687 | 0.059566 | -0.788514 | 0.032891 | -0.105671 |
| 24.708 | -0.491043 | -0.413993 | -0.170237 | -0.169967 | -0.038584 | -0.248593 | -1.090329 | 0.08012 | -0.206187 | -0.045166 | 0.007734 | 0.161252 | 0.035518 | -0.13066 | 0.028179 | -0.02284 | -0.042514 | -0.041228 | 0.016326 | -0.159918 | 0.010407 | -0.515016 | -0.03306 | 0.127609 |
| 24.722 | -0.739047 | -0.566859 | -0.257997 | -0.138818 | -0.036757 | -0.41955 | -1.258626 | -0.077856 | -0.260827 | -0.081567 | 0.092462 | 0.024465 | -0.063563 | -0.249383 | -0.017199 | -0.005327 | -0.00924 | -0.104264 | -0.01747 | -0.271548 | -0.031709 | -0.456436 | -0.158178 | 0.135301 |
| 24.736 | -1.315388 | -0.364556 | -0.152194 | -0.077974 | 0.005821 | -0.305849 | -1.062532 | -0.129132 | -0.281798 | -0.08024 | 0.098811 | -0.197478 | -0.15305 | -0.277591 | -0.058958 | 0.048862 | 0.029544 | -0.138024 | -0.025259 | -0.29663 | -0.032114 | -0.247124 | -0.19604 | 0.128854 |
| 24.75 | -1.48942 | -0.233522 | -0.065291 | -0.03017 | 0.056721 | -0.191673 | -0.674679 | -0.115296 | -0.273596 | -0.068145 | -0.079724 | -0.59583 | -0.148913 | -0.255345 | -0.072877 | 0.043147 | 0.076681 | -0.166721 | -0.006728 | -0.302071 | -0.011008 | -0.054711 | -0.180489 | 0.141183 |
| 24.764 | -0.612064 | 0.057295 | -0.392616 | -0.040363 | 0.032579 | -0.026984 | -0.259683 | -0.02277 | -0.240882 | -0.048115 | -0.343812 | -1.020414 | -0.04155 | -0.240931 | -0.097964 | -0.048761 | 0.090584 | -0.240861 | -0.066591 | -0.331777 | 0.000623 | 0.084822 | -0.161233 | 0.139559 |
| 24.778 | 0.093061 | 0.189386 | -0.741156 | -0.06497 | -0.028541 | 0.086032 | -0.091191 | 0.037559 | -0.204591 | -0.035301 | -0.630744 | -0.928306 | 0.037467 | -0.266798 | -0.153832 | -0.19583 | 0.067608 | -0.38484 | -0.122258 | -0.386712 | -0.023222 | 0.132826 | -0.109896 | 0.119275 |
| 24.792 | 0.456423 | 0.101576 | -0.564342 | -0.06605 | -0.054039 | -0.003677 | -0.227189 | -0.170341 | -0.179906 | -0.025261 | -0.923403 | -0.257905 | 0.048631 | -0.303347 | -0.226257 | -0.238936 | 0.013549 | -0.535766 | -0.132765 | -0.419889 | -0.071241 | 0.119068 | -0.12456 | 0.089995 |
| 24.806 | 0.262008 | -0.085737 | -0.234909 | -0.096539 | -0.065282 | -0.206685 | -0.516751 | -0.38757 | -0.159409 | -0.032762 | -0.946426 | 0.62584 | 0.039405 | -0.341398 | -0.295582 | -0.066513 | -0.033098 | -0.74393 | -0.159342 | -0.3955 | -0.112873 | 0.028724 | -0.216351 | 0.016985 |
| 24.819 | -0.42183 | -0.318252 | -0.071806 | -0.15594 | -0.096152 | -0.43698 | -0.897013 | -0.581947 | -0.227066 | -0.06042 | -0.711384 | 1.172728 | -0.007621 | -0.438091 | -0.380778 | 0.074004 | -0.086165 | -1.001236 | -0.260956 | -0.345144 | -0.188379 | -0.127864 | -0.423437 | -0.094878 |
| 24.833 | -0.856105 | -0.439929 | -0.162033 | -0.183511 | -0.134515 | -0.786928 | -1.277402 | -0.732042 | -0.296548 | -0.091512 | -0.535292 | 0.763 | -0.091198 | -0.572522 | -0.515048 | 0.107202 | -0.137443 | -1.351856 | -0.344509 | -0.377029 | -0.310713 | -0.263343 | -0.756215 | -0.214632 |
| 24.847 | -0.981792 | -0.490913 | -0.227085 | -0.17266 | -0.206348 | -0.811142 | -1.390372 | -0.821092 | -0.312539 | -0.145114 | -0.316796 | -0.066957 | -0.127018 | -0.726 | -0.696783 | 0.056436 | -0.233485 | -1.6764 | -0.434766 | -0.445038 | -0.436034 | -0.286236 | -0.937153 | -0.279234 |
| 24.861 | -1.460394 | -0.322926 | -0.251097 | -0.217684 | -0.301367 | -0.521149 | -1.257445 | -0.852264 | -0.183846 | -0.24763 | -0.240658 | -0.680628 | -0.03471 | -0.818264 | -0.795524 | -0.234976 | -0.327499 | -1.858345 | -0.465231 | -0.473362 | -0.594476 | -0.31939 | -0.812749 | -0.356459 |
| 24.875 | -2.113702 | -0.113105 | -0.341896 | -0.321214 | -0.35934 | -0.316183 | -1.049466 | -0.870135 | -0.24929 | -0.330383 | -0.373136 | -1.688048 | 0.042196 | -0.828218 | -0.770183 | -0.329296 | -0.367145 | -1.759752 | -0.464697 | -0.404052 | -0.623036 | -0.343896 | -0.472909 | -0.398232 |
| 24.889 | -2.190045 | -0.005998 | -0.490156 | -0.317491 | -0.357748 | -0.250269 | -0.536836 | -1.199984 | -0.475428 | -0.331586 | -0.443947 | -2.752122 | -0.080527 | -0.798318 | -0.64686 | -0.285365 | -0.460114 | -1.446258 | -0.41464 | -0.326322 | -0.605653 | -0.39111 | -0.202785 | -0.388823 |
| 24.903 | -1.463636 | -0.018934 | -0.647828 | -0.240846 | -0.325998 | -0.238248 | -0.131724 | -1.241702 | -0.555916 | -0.279323 | -0.378366 | -2.819694 | -0.263062 | -0.714024 | -0.379629 | -0.212353 | -0.441365 | -1.138498 | -0.247051 | -0.176334 | -0.63459 | -0.412054 | 0.050305 | -0.317103 |
| 24.917 | -0.564784 | -0.008995 | -0.710935 | -0.17587 | -0.233648 | -0.143266 | 0.05002 | -1.013657 | -0.440622 | -0.119788 | -0.094337 | -1.871967 | -0.403973 | -0.46795 | 0.018353 | -0.032819 | -0.208347 | -0.680031 | 0.090581 | 0.138414 | -0.550945 | -0.369417 | 0.256764 | -0.164912 |
| 24.931 | 0.112164 | 0.091693 | -0.504949 | 0.005849 | 0.003054 | 0.084843 | -0.006079 | -0.488923 | -0.198176 | 0.128715 | 0.370813 | -0.777675 | -0.345603 | -0.114889 | 0.510492 | 0.25578 | 0.040559 | -0.161719 | 0.52156 | 0.520315 | -0.300061 | -0.023168 | 0.457436 | 0.0233 |
| 24.944 | 0.201301 | 0.127588 | -0.297769 | 0.150721 | 0.113612 | 0.079055 | -0.333905 | -0.482325 | -0.089937 | 0.179313 | 0.540622 | -0.357129 | -0.360666 | -0.108176 | 0.515078 | 0.181074 | -0.056106 | -0.141061 | 0.446806 | 0.317699 | -0.464528 | -0.067521 | 0.23579 | -0.165827 |
| 24.958 | -0.233438 | -0.072958 | -0.202862 | -0.003826 | -0.07807 | -0.316825 | -0.845114 | -1.348601 | -0.36851 | -0.15706 | 0.280473 | -0.66671 | -0.792997 | -0.504441 | -0.11138 | -0.29648 | -0.486093 | -0.627964 | -0.353361 | -0.454434 | -0.970541 | -0.972435 | -0.50112 | -0.828833 |
| 24.972 | -0.611251 | -0.289536 | -0.050676 | -0.367229 | -0.264628 | -0.710231 | -1.229828 | -2.170196 | -0.921154 | -0.69385 | 0.017752 | -1.557562 | -1.252824 | -0.666883 | -0.151848 | -0.351777 | -0.601396 | -0.707852 | -0.574796 | -0.663436 | -1.012053 | -1.480078 | -0.773198 | -1.060804 |
| 24.986 | -0.632824 | -0.335425 | -0.077962 | -0.484255 | -0.218236 | -0.609411 | -1.740614 | -2.254974 | -1.39104 | -0.946415 | -0.332489 | -2.520062 | -1.256287 | -0.68536 | 0.325529 | -0.011874 | -0.368354 | -0.264812 | -0.20093 | -0.220931 | -0.550146 | -1.812766 | -0.462302 | -0.749805 |
| 25 | -0.592219 | -0.337076 | 0.023942 | -0.490214 | -0.135067 | -0.286752 | -2.469942 | -1.293387 | -1.040063 | -0.576973 | -0.792818 | -1.707217 | -0.920557 | -0.801351 | 0.547299 | -0.034836 | -0.225943 | -0.153313 | -0.263351 | -0.328495 | -0.398966 | -2.026343 | -0.328077 | -0.517459 |
| 25.014 | -0.100724 | -0.353573 | 0.202994 | -0.322917 | -0.150425 | 0.019404 | -2.387117 | -0.391427 | -0.444454 | -0.116897 | -0.899184 | -0.915214 | -0.446811 | -0.718307 | 0.352855 | -0.298987 | -0.184206 | -0.349789 | -0.677845 | -0.500339 | -0.432326 | -1.067613 | 0.041674 | -0.415699 |
| 25.028 | 0.551255 | -0.159762 | 0.450398 | -0.0633 | -0.119206 | 0.319319 | -1.550474 | 0.230828 | 0.173005 | 0.196854 | -0.635425 | -0.199136 | -0.075532 | -0.506718 | 0.088346 | -0.404778 | -0.086242 | -0.441229 | -0.847776 | -0.584108 | -0.523933 | -0.390701 | 0.307669 | -0.357503 |
| 25.042 | 0.990657 | 0.219465 | 0.405349 | 0.130426 | -0.004355 | 0.56724 | -0.602601 | 0.717484 | 0.569143 | 0.373579 | -0.017558 | 0.440954 | 0.238436 | -0.32554 | -0.102021 | -0.257649 | 0.038385 | -0.168249 | -0.458337 | -0.527253 | -0.548305 | -0.189618 | 0.448093 | -0.329575 |
| 25.056 | 0.658444 | 0.550968 | 0.05454 | 0.247793 | 0.134273 | 0.583202 | 0.346056 | 0.917344 | 0.706678 | 0.369091 | 0.503641 | 1.119178 | 0.5252 | -0.122909 | -0.115701 | -0.080052 | 0.097355 | 0.297195 | 0.055531 | -0.394179 | -0.435326 | 0.018044 | 0.423256 | -0.251787 |
| 25.069 | 0.251277 | 0.726885 | -0.346712 | 0.270717 | 0.16324 | 0.301915 | 0.944619 | 0.721552 | 0.656338 | 0.256182 | 1.259965 | 1.58044 | 0.653913 | 0.013027 | -0.097551 | -0.082825 | -0.04387 | 0.442061 | -0.032722 | -0.240072 | -0.386279 | 0.125839 | 0.235231 | -0.20716 |
| 25.083 | 0.186408 | 0.06824 | -0.419762 | 0.197415 | 0.05429 | 0.03615 | 0.889011 | 0.14578 | 0.496559 | 0.146064 | 1.541917 | 1.31556 | 0.535164 | 0.032036 | -0.141663 | -0.154261 | -0.258306 | 0.311035 | -0.526104 | -0.101593 | -0.274884 | 0.192689 | -0.005941 | -0.247129 |
| 25.097 | 0.111972 | -0.663311 | -0.075341 | -0.011241 | -0.0034 | -0.011464 | 0.339238 | 0.000534 | 0.502345 | 0.148898 | 0.180807 | 0.577064 | 0.494493 | -0.037777 | -0.208198 | -0.369671 | -0.33179 | 0.14498 | -1.528563 | -0.257965 | -0.033415 | 0.260949 | -0.140941 | -0.337303 |
| 25.111 | -0.401072 | -0.179638 | 0.329217 | -0.0169 | 0.129977 | 0.126396 | -0.091519 | 0.56485 | 0.912281 | 0.267004 | -0.448655 | -0.282277 | 0.726812 | -0.132362 | -0.283273 | -0.452812 | -0.242232 | -0.088007 | -2.686394 | -0.669667 | 0.117191 | 0.064451 | -0.235939 | -0.407796 |
| 25.125 | -0.646573 | 0.536997 | 0.60424 | 0.159754 | 0.260093 | 0.236866 | 0.151467 | 0.985552 | 0.726742 | 0.328173 | -0.223907 | -0.616363 | 0.80557 | -0.00124 | -0.220056 | -0.343245 | -0.001468 | -0.221927 | -1.668345 | -0.630689 | 0.051558 | -0.054443 | -0.181462 | -0.241537 |
| 25.139 | -0.318097 | 0.448398 | 0.563396 | 0.188916 | 0.243437 | 0.240255 | 0.875565 | 0.769977 | 0.022529 | 0.232966 | 0.121952 | -0.686179 | 0.506925 | 0.352573 | -0.04945 | -0.105211 | 0.152538 | -0.236642 | -0.450294 | 0.186959 | -0.091079 | 0.110105 | 0.025267 | 0.128212 |
| 25.153 | 0.101783 | 0.106867 | 0.15938 | 0.043361 | 0.216236 | 0.27215 | 1.672669 | 0.687781 | -0.36061 | 0.120898 | -0.284599 | 0.316728 | 0.388786 | 0.610339 | 0.080463 | 0.014241 | 0.045404 | -0.216089 | 0.663844 | 1.123872 | -0.067891 | 0.325182 | 0.372524 | 0.449967 |
| 25.167 | 0.030643 | 0.013573 | -0.333039 | 0.099158 | 0.248387 | 0.323592 | 1.98959 | 0.816568 | 0.054941 | 0.097427 | -0.690001 | 1.290572 | 0.681713 | 0.503415 | 0.078533 | -0.092121 | -0.128851 | -0.162184 | 1.826331 | 0.976717 | 0.138635 | 0.365774 | 0.85891 | 0.501463 |
| 25.181 | -0.214484 | 0.181357 | -0.23355 | 0.183248 | 0.162101 | 0.251983 | 1.705076 | 0.940856 | 0.816086 | 0.119292 | -0.290733 | 2.160374 | 0.812229 | 0.270735 | 0.114728 | -0.198068 | -0.121068 | -0.062152 | 1.019988 | 0.283976 | 0.437586 | 0.255792 | 0.783045 | 0.387439 |
| 25.194 | -0.274317 | 0.234133 | -0.110967 | 0.088351 | -0.0145 | 0.071164 | 1.056615 | 0.722524 | 0.935516 | 0.153828 | -0.146241 | 1.03602 | 0.600965 | 0.035002 | 0.358241 | -0.125756 | 0.038304 | 0.043928 | 0.067837 | 0.240302 | 0.623731 | 0.18705 | 0.171954 | 0.264615 |

| | | |
|---|---|---|
| 24.70833 | -0.094399 | [illegible] |
| 24.72222 | -0.047513 | [illegible] |
| 24.73611 | 0.059355 | [illegible] |
| 24.75 | 0.125215 | [illegible] |
| 24.76389 | 0.179074 | [illegible] |
| 24.77778 | 0.194058 | [illegible] |
| 24.79167 | 0.132707 | [illegible] |
| 24.80556 | 0.035518 | [illegible] |
| 24.81944 | -0.063563 | [illegible] |
| 24.83333 | -0.15305 | [illegible] |
| 24.84722 | -0.148913 | [illegible] |
| 24.86111 | -0.04155 | [illegible] |
| 24.875 | 0.037467 | [illegible] |
| 24.88889 | 0.048631 | [illegible] |
| 24.90278 | 0.039405 | [illegible] |
| 24.91667 | -0.007621 | [illegible] |
| 24.93056 | -0.091198 | [illegible] |
| 24.94444 | -0.127018 | [illegible] |
| 24.95833 | -0.03471 | [illegible] |
| 24.97222 | 0.042196 | [illegible] |
| 24.98611 | -0.080527 | [illegible] |
| 25 | -0.263062 | [illegible] |
| 25.01389 | -0.403973 | [illegible] |
| 25.02778 | -0.345603 | [illegible] |
| 25.04167 | -0.360666 | [illegible] |
| 25.05556 | -0.792997 | [illegible] |
| 25.06944 | -1.252824 | [illegible] |
| 25.08333 | -1.256287 | [illegible] |
| 25.09722 | -0.920557 | [illegible] |
| 25.11111 | -0.446811 | [illegible] |
| 25.125 | -0.075532 | [illegible] |
| 25.13889 | 0.238436 | [illegible] |
| 25.15278 | 0.5252 | [illegible] |
| 25.16667 | 0.653913 | [illegible] |
| 25.18056 | 0.535164 | [illegible] |
| 25.19444 | 0.494493 | [illegible] |
| 25.20833 | 0.726812 | [illegible] |
| 25.22222 | 0.80557 | [illegible] |
| 25.23611 | 0.506925 | [illegible] |
| 25.25 | 0.388786 | [illegible] |
| 25.26389 | 0.681713 | [illegible] |
| 25.27778 | 0.812229 | [illegible] |
| 25.29167 | 0.600965 | [illegible] |

| | | | | | | | | | | | | | | | | | | | | | | | | | | | | |
|---|---|---|---|---|---|---|---|---|---|---|---|---|---|---|---|---|---|---|---|---|---|---|---|---|---|---|---|---|
| 27.931 | -0.181448 | -0.535452 | -0.847974 | 0.034858 | 0.514439 | -0.091991 | -1.241745 | -0.598457 | -0.316158 | -0.394854 | -0.188373 | -2.173055 | -0.297588 | -0.51099 | 0.418768 | -0.249811 | -0.11229 | -0.071308 | 0.418639 | 0.330401 | 0.148534 | 0.368501 | -0.104887 | -0.151771 | | 28.02778 | -0.297588 | -0.142915 |
| 27.944 | -0.102401 | -0.412418 | -0.048636 | 0.158886 | 0.450955 | -0.038271 | -0.842593 | -0.411493 | -0.100075 | -0.235555 | 0.157114 | -2.056116 | 0.050326 | -0.466166 | 0.233887 | -0.422539 | -0.223787 | -0.095923 | 0.354773 | 0.209629 | 0.060134 | 0.288977 | -0.073799 | -0.24575 | | 28.04167 | 0.050326 | -0.137452 |
| 27.958 | -0.694682 | -0.787464 | 0.260313 | -0.164387 | -0.068251 | -0.773622 | -0.07 | -1.258516 | -0.341213 | -0.539667 | -0.181362 | -1.41514 | -0.229447 | -0.776602 | -0.271573 | -0.99573 | -0.622272 | -0.502108 | -0.126426 | -0.286178 | -0.416981 | -0.200808 | -0.379737 | -0.684293 | | 28.05556 | -0.229447 | -0.543359 |
| 27.972 | -1.866228 | -2.041186 | -0.050833 | -0.852082 | -1.10091 | -2.09741 | 0.127706 | -2.650356 | -0.931384 | -1.02782 | -0.774356 | -0.748521 | -0.763991 | -1.216168 | -0.518905 | -1.761548 | -0.974799 | -0.82369 | -0.260319 | -0.376226 | -0.783946 | -0.344758 | -0.475434 | -1.042547 | | 28.06944 | -0.763991 | -1.041115 |
| 27.986 | -2.573747 | -1.896372 | -0.625267 | -1.441273 | -0.902607 | -2.753886 | -0.714085 | -3.508235 | -1.278285 | -0.983035 | -1.035794 | 0.069663 | -1.350685 | -1.494145 | -0.509453 | -2.64985 | -1.119245 | -1.183025 | 0.064009 | -0.080095 | -1.02622 | -0.00467 | -0.333286 | -1.129905 | | 28.08333 | -1.350685 | -1.340735 |
| 28 | -2.028727 | -0.661067 | -0.694474 | -1.778879 | -0.243776 | -1.680973 | -2.99747 | -2.785916 | -0.836717 | -0.398142 | -1.219855 | 1.24194 | -1.583185 | -1.142917 | -0.42096 | -2.733321 | -0.781383 | -1.214689 | 0.269612 | 0.073796 | -1.20485 | 0.238122 | -0.387094 | -0.951993 | | 28.09722 | -1.583185 | -1.305884 |
| 28.014 | -0.865146 | -0.040902 | -0.204907 | -1.321859 | -0.005334 | -0.252195 | -4.630833 | -1.274332 | -0.573914 | 0.020678 | -1.335445 | 0.930052 | -1.202363 | -0.458697 | -0.290355 | -1.674583 | -0.136214 | -0.977606 | 0.213188 | 0.01923 | -0.978355 | 0.206977 | -0.198432 | -0.643865 | | 28.11111 | -1.202363 | -0.942487 |
| 28.028 | 0.131853 | 0.193924 | 0.287724 | -0.627188 | 0.146023 | 0.581984 | -2.676701 | 0.136854 | -0.280742 | 0.166682 | -0.913998 | -0.220886 | -0.44398 | -0.069974 | -0.281193 | -0.683739 | 0.287321 | -0.775599 | 0.057569 | -0.107227 | -0.616188 | 0.08767 | -0.191761 | -0.358257 | | 28.125 | -0.44398 | -0.554301 |
| 28.042 | 0.841615 | 0.184907 | 0.488971 | -0.059347 | 0.594107 | 0.93441 | -1.198824 | 0.860119 | -0.013187 | 0.23818 | -0.488062 | -1.129594 | 0.211322 | -0.086377 | -0.154237 | -0.15335 | 0.34636 | -0.490902 | -0.078228 | -0.225941 | -0.393196 | 0.004475 | -0.142169 | -0.195835 | | 28.13889 | 0.211322 | -0.2256 |
| 28.056 | 1.379584 | 0.168759 | 0.505617 | 0.362847 | 0.446546 | 1.246869 | -0.378581 | 1.347329 | 0.318258 | 0.277307 | 0.042827 | -1.837622 | 0.695713 | -0.110499 | 0.121185 | 0.239025 | 0.245463 | -0.144074 | -0.14334 | -0.270252 | -0.369397 | -0.008838 | 0.026482 | -0.06669 | | 28.15278 | 0.695713 | -0.007759 |
| 28.069 | 1.521346 | 0.220351 | 0.391628 | 0.602544 | 0.177553 | 1.459281 | 0.170478 | 1.799016 | 0.752617 | 0.288219 | 0.380875 | -1.87113 | 1.050678 | 0.2235 | 0.458632 | 0.603043 | 0.107937 | -0.006818 | -0.124641 | -0.251889 | -0.518046 | 0.001425 | 0.228985 | 0.059198 | | 28.16667 | 1.050678 | 0.096281 |
| 28.083 | 1.384789 | 0.195172 | 0.254825 | 0.550013 | -0.205809 | 1.075406 | 0.696766 | 1.458363 | 0.947208 | 0.251197 | 0.42085 | -1.719934 | 1.098841 | 0.712004 | 0.842062 | 0.717782 | -0.045336 | -0.103887 | -0.079694 | -0.248701 | -0.558558 | 0.000647 | 0.391122 | 0.191053 | | 28.18056 | 1.098841 | 0.200425 |
| 28.097 | 1.418306 | 0.043464 | 0.130554 | 0.258781 | 0.062568 | 0.089724 | 1.180985 | 0.447641 | 0.664077 | 0.343522 | 0.418087 | -1.416398 | 0.596868 | 1.123081 | 1.038499 | 0.720223 | -0.019195 | -0.201807 | -0.015247 | -0.237956 | -0.178197 | 0.029798 | 0.328279 | 0.359187 | | 28.19444 | 0.596868 | 0.330684 |
| 28.111 | 1.914096 | -0.064971 | -0.216636 | 0.01306 | 0.676413 | -0.688326 | 1.51509 | -0.31531 | 0.220138 | 0.449403 | 0.63644 | -0.988855 | 0.017171 | 1.356321 | 0.779393 | 0.701613 | 0.345806 | -0.176332 | 0.064025 | -0.193762 | 0.523076 | 0.083329 | 0.08147 | 0.473077 | | 28.20833 | 0.017171 | 0.468825 |
| 28.125 | 2.125709 | 0.093985 | -0.458046 | -0.053112 | 0.695297 | -0.661931 | 1.455535 | -0.510941 | 0.111451 | 0.408964 | 0.981355 | -0.288555 | -0.169249 | 1.231074 | 0.341127 | 0.635797 | 0.689159 | -0.150446 | 0.088663 | -0.172019 | 1.18428 | 0.095714 | -0.162029 | 0.530667 | | 28.22222 | -0.169249 | 0.543746 |
| 28.139 | 1.22603 | 0.377727 | -0.277143 | 0.154338 | 0.301409 | 0.072635 | 1.08873 | -0.317429 | 0.203301 | 0.19253 | 1.907416 | 0.731837 | 0.00559 | 0.760016 | 0.009505 | 0.602353 | 0.672886 | -0.096643 | 0.05724 | -0.184259 | 1.530982 | 0.076207 | -0.23997 | 0.459343 | | 28.23611 | 0.00559 | 0.444544 |
| 28.153 | -0.663532 | 0.343786 | -0.068014 | 0.182248 | -0.143785 | 0.416549 | 0.496386 | -0.086377 | 0.311016 | 0.025672 | 2.468803 | 1.437697 | 0.261445 | 0.14035 | 0.07658 | 0.648105 | 0.499204 | 0.036031 | 0.008869 | -0.218591 | 1.429918 | 0.039866 | -0.201184 | 0.285934 | | 28.25 | 0.261445 | 0.224971 |
| 28.167 | -0.585659 | 0.172876 | -0.078419 | 0.144015 | -0.445657 | 0.329813 | -0.278308 | 0.10687 | 0.188986 | 0.731185 | 1.020335 | 1.412272 | 0.391346 | -0.218236 | 0.534376 | 0.744636 | 0.522987 | 0.182876 | -0.055818 | -0.267081 | 1.146013 | -0.028062 | 0.075675 | 0.175722 | | 28.26389 | 0.391346 | 0.219063 |
| 28.181 | 0.860977 | 0.366335 | 0.224744 | 0.218327 | -0.160922 | 0.336663 | -0.856186 | 0.442208 | -0.097022 | 0.845537 | -1.044742 | 0.758331 | 0.2951 | 0.420344 | 0.974327 | 0.942454 | 0.699552 | 0.28433 | -0.136354 | -0.346827 | 1.176414 | -0.107855 | 0.192492 | 0.34201 | | 28.27778 | 0.2951 | 0.430471 |
| 28.194 | 0.975134 | 0.608929 | 0.921884 | 0.391446 | 0.284219 | 0.510863 | -0.751484 | 0.802783 | -0.056783 | 0.209127 | -2.125972 | -0.047358 | 0.160455 | 1.381321 | 1.01447 | 1.211375 | 0.903335 | 0.282957 | -0.197932 | -0.398504 | 1.434351 | -0.158423 | -0.211865 | 0.661869 | | 28.29167 | 0.160455 | 0.80935 |
| 28.208 | -0.119275 | 0.257179 | 1.721947 | 0.354079 | 0.599973 | 0.481306 | -0.042426 | 0.773461 | 0.545267 | -0.409014 | -1.051611 | -0.562908 | 0.179505 | 1.837397 | 0.856594 | 1.33019 | 1.087871 | 0.348112 | -0.211846 | -0.394791 | 1.659319 | -0.163456 | -0.357005 | 0.963651 | | 28.30556 | 0.179505 | 1.242234 |
| 28.222 | -0.342284 | -0.222873 | 1.746173 | 0.159726 | 0.393822 | 0.36492 | 0.877327 | 0.66013 | 0.970795 | -0.382305 | -0.01713 | -0.135868 | 0.601524 | 1.638929 | 0.6436 | 0.977573 | 1.157629 | 0.470245 | -0.188505 | -0.354138 | 2.161443 | -0.139688 | -0.124579 | 1.11818 | | 28.31944 | 0.601524 | 1.385158 |
| 28.236 | 0.800037 | -0.404923 | 0.658445 | 0.230834 | 0.13823 | 0.180062 | 1.632461 | 0.860844 | 1.083114 | 0.724973 | 0.499123 | 0.839573 | 1.216091 | 1.406848 | 0.594053 | 0.712085 | 0.846078 | 0.602554 | -0.131614 | -0.293392 | 2.160961 | -0.09367 | 0.532902 | 1.043928 | | 28.33333 | 1.216091 | 1.193433 |
| 28.25 | 1.406445 | -0.147252 | -0.928876 | 0.608192 | 0.261735 | 0.155007 | 2.030476 | 1.091137 | 0.852973 | 1.624475 | 0.496136 | 1.451698 | 1.125784 | 1.254096 | 0.594454 | 0.762733 | 0.636673 | 0.652415 | -0.043382 | -0.219981 | 1.812806 | -0.017113 | 1.152649 | 0.897622 | | 28.34722 | 1.125784 | 0.967995 |
| 28.264 | 0.493349 | 0.256496 | -1.807633 | 0.801137 | 0.390325 | 0.385952 | 1.863289 | 1.333525 | 0.422793 | 1.642782 | 0.281371 | 1.13362 | 0.350157 | 1.017645 | 0.624785 | 0.945304 | 0.699486 | 0.590931 | 0.026424 | -0.168715 | 1.695526 | 0.051131 | 1.435773 | 0.759647 | | 28.36111 | 0.350157 | 0.799095 |
| 28.278 | -0.682339 | 0.453638 | -1.134415 | 0.399261 | 0.315721 | 0.807328 | 1.262728 | 1.032708 | 0.415316 | 0.331432 | 0.082434 | 0.536959 | -0.321969 | 0.814231 | 0.656172 | 0.932059 | 1.008344 | 0.484182 | 0.019344 | -0.203064 | 1.501294 | 0.047136 | 1.477386 | 0.620032 | | 28.375 | -0.321969 | 0.693498 |
| 28.292 | -0.293162 | 0.376769 | 0.421885 | 0.110964 | 0.363571 | 1.069464 | 0.94342 | 0.879205 | 0.945024 | -0.326905 | 0.136411 | 0.343997 | 0.144299 | 0.969336 | 0.717887 | 0.819381 | 1.047276 | 0.508288 | -0.011314 | -0.24607 | 1.376992 | 0.016723 | 1.40132 | 0.580175 | | 28.38889 | 0.144299 | 0.741994 |
| 28.306 | 1.219709 | 0.330677 | 1.449524 | 0.282721 | 0.408115 | 0.89442 | 1.381125 | 1.168194 | 1.418067 | 0.074634 | 0.534317 | 0.675377 | 1.341777 | 1.430498 | 0.859513 | 0.753455 | 0.667193 | 0.697172 | 0.035474 | -0.19778 | 1.331425 | 0.06289 | 1.164356 | 0.775537 | | 28.40278 | 1.341777 | 0.859504 |
| 28.319 | 1.997683 | 0.45655 | 1.644384 | 0.74878 | 0.61637 | 0.775346 | 1.997694 | 1.422649 | 1.545316 | 0.552808 | 1.138026 | 1.004362 | 1.850037 | 2.092276 | 1.135457 | 0.883076 | 0.300779 | 1.004375 | 0.194406 | -0.037807 | 1.350432 | 0.227714 | 0.685435 | 0.995837 | | 28.41667 | 1.850037 | 1.039757 |
| 28.333 | 1.216007 | 0.633749 | 0.541245 | 1.029128 | 0.978605 | 0.919059 | 2.546606 | 1.802468 | 1.175779 | 1.031626 | 1.707531 | 1.119738 | 1.64004 | 2.554601 | 1.407109 | 1.212949 | 0.173939 | 1.235027 | 0.395213 | 0.161139 | 1.436815 | 0.441103 | 0.982785 | 1.196554 | | 28.43056 | 1.64004 | 1.305584 |
| 28.347 | 0.796861 | 0.667335 | -0.097001 | 0.70405 | 0.978353 | 1.059286 | 2.909144 | 2.121669 | 0.993964 | 1.058513 | 2.009172 | 0.969438 | 1.139957 | 2.513556 | 1.438103 | 1.471674 | 0.50174 | 1.313392 | 0.450197 | 0.290345 | 1.385403 | 0.496052 | 2.024346 | 1.225447 | | 28.44444 | 1.139957 | 1.324327 |
| 28.361 | 1.043693 | 0.578323 | -0.624853 | 0.296766 | 0.74945 | 0.910334 | 3.051749 | 1.720021 | 1.034896 | 0.542877 | 2.239903 | 0.852085 | -0.049041 | 2.308444 | 1.164331 | 1.368853 | 0.917778 | 1.226016 | 0.209545 | 0.238342 | 1.050416 | 0.212761 | 2.048717 | 0.95522 | | 28.45833 | -0.049041 | 1.020797 |
| 28.375 | 1.539923 | 0.489576 | -0.839101 | 0.057649 | 0.480646 | 0.728368 | 2.784594 | 0.898621 | 1.049614 | -0.093375 | 2.64156 | 0.889921 | -0.675307 | 2.234886 | 0.854892 | 1.171458 | 1.066075 | 0.886146 | -0.256357 | -0.054745 | 0.659569 | -0.218494 | 1.417737 | 0.463047 | | 28.47222 | -0.675307 | 0.507703 |
| 28.389 | 2.269658 | 0.553741 | -0.500421 | 0.141897 | 0.296818 | 0.762569 | 2.116418 | 0.637869 | 0.840857 | -0.435448 | 2.931979 | 1.209863 | 0.020104 | 2.425777 | 0.876074 | 0.707704 | 1.078098 | 0.54973 | -0.3276 | -0.212263 | 0.682623 | -0.239933 | 1.273583 | 0.149159 | | 28.48611 | 0.020104 | 0.163091 |
| 28.403 | 2.666935 | 0.930672 | 1.072882 | 0.420889 | 0.531185 | 1.265816 | 1.917385 | 0.667872 | 0.98073 | 0.142726 | 3.258233 | 1.529671 | 1.803533 | 2.256996 | 1.140628 | 0.719929 | 0.750117 | 0.755487 | -0.050646 | -0.132041 | 1.250725 | -0.004219 | 1.197394 | 0.193195 | | 28.5 | 1.803533 | 0.07409 |
| 28.417 | 2.274975 | 1.361395 | 1.411761 | 0.242281 | 0.846426 | 1.854136 | 1.800955 | 1.223261 | 1.354888 | 0.78019 | 3.684284 | 1.598652 | 2.54656 | 1.461313 | 0.948115 | 0.464086 | 0.356609 | 0.866201 | 0.033543 | -0.149644 | 1.661473 | 0.075858 | 0.79236 | 0.322202 | | 28.51389 | 2.54656 | 0.235622 |
| 28.431 | 2.155725 | 1.551178 | 1.067623 | -0.090746 | 1.146913 | 1.913298 | 1.756021 | 1.593626 | 1.605001 | 1.442333 | 3.347216 | 1.663685 | 2.315746 | 1.030387 | 0.496584 | 0.522497 | -0.042434 | 0.775253 | -0.111025 | -0.285562 | 0.848029 | -0.055628 | -0.373553 | 0.359759 | | 28.52778 | 2.315746 | 0.450625 |
| 28.444 | 1.802779 | 0.579806 | 0.521993 | -0.094612 | 1.200568 | 1.253918 | 1.451917 | 1.034049 | 1.23254 | 1.757804 | 1.978909 | 1.724704 | 1.792141 | 0.249989 | 0.026053 | 0.528061 | -0.404844 | 0.289487 | -0.538252 | -0.637785 | -0.209809 | -0.437292 | -1.942276 | 0.104255 | | 28.54167 | 1.792141 | 0.257414 |
| 28.458 | 1.516174 | 0.284498 | 0.160158 | 0.088808 | 0.921616 | 0.552486 | 1.361772 | 0.454014 | 0.791775 | 1.391176 | 1.065068 | 2.135936 | 1.19746 | -0.00069 | -0.585149 | -0.032598 | -0.819929 | -0.523148 | -1.226049 | -1.280537 | -0.923216 | -1.038467 | -3.146005 | -0.454374 | | 28.55556 | 1.19746 | -0.545967 |
| 28.472 | 1.440414 | 0.183678 | 0.221199 | -0.013255 | 0.611033 | 0.274155 | 1.954301 | -0.026596 | 0.603137 | 0.489061 | 0.92812 | 2.05949 | 0.210376 | -0.17951 | -0.751347 | -0.484522 | -0.793106 | -0.813357 | -1.420281 | -1.430875 | -0.464053 | -1.23544 | -1.205556 | -0.785134 | | 28.56944 | 0.210376 | -0.814536 |
| 28.486 | 1.08083 | 0.051745 | -0.086786 | -0.021321 | 0.140103 | 0.020123 | 1.509419 | 0.058993 | 0.445861 | -0.142013 | 1.330867 | 1.390842 | -0.792867 | -0.134206 | -0.36047 | -0.51624 | -0.222712 | -0.558041 | -0.909475 | -0.88178 | 0.670947 | -0.835472 | -0.272858 | -0.469531 | | 28.58333 | -0.792867 | -0.388356 |
| 28.5 | 0.864798 | 0.087785 | 0.024347 | 0.334199 | -0.16619 | -0.173136 | 1.155721 | 0.467858 | -0.01655 | 0.384956 | 1.47857 | 1.329481 | -1.35819 | 0.041133 | -0.029131 | -0.599586 | 0.178968 | -0.21443 | -0.360084 | -0.339917 | 1.167497 | -0.345344 | 0.382015 | -0.105123 | | 28.59722 | -1.35819 | -0.044748 |
| 28.514 | 0.469012 | 0.104369 | 0.124024 | 0.266003 | -0.30103 | -0.16852 | 0.688529 | 0.033837 | -0.095996 | 0.172178 | 1.05824 | 1.319709 | -1.137276 | 0.024951 | 0.175653 | -0.697286 | 0.248795 | -0.005756 | -0.083104 | -0.093578 | 0.519333 | -0.061706 | 1.78998 | -0.017228 | | 28.61111 | -1.137276 | -0.001979 |
| 28.528 | 0.132599 | 0.006609 | 0.643206 | 0.180875 | -0.374977 | 0.025254 | 0.38353 | 0.08658 | 0.379115 | -0.073773 | 0.574268 | 0.896866 | 0.510727 | 0.33704 | 0.001071 | -0.214092 | -0.049573 | 0.041061 | -0.062693 | -0.081693 | -0.199564 | -0.029989 | 1.135973 | -0.175148 | | 28.625 | 0.510727 | -0.200736 |
| 28.542 | -0.058981 | -0.075961 | 0.828541 | 0.173097 | -0.58355 | 0.263181 | 0.414702 | 0.410833 | 0.188295 | -0.445329 | -0.356356 | 0.380095 | 2.1313 | 0.424457 | -0.214512 | 0.050989 | -0.441962 | -0.033577 | -0.12443 | -0.136143 | -1.139555 | -0.10079 | 0.145974 | -0.382503 | | 28.63889 | 2.1313 | -0.435552 |
| 28.556 | -0.275617 | -0.057186 | 0.022477 | 0.140697 | -0.414423 | 0.545882 | 0.563041 | 0.165074 | -0.235934 | -0.295499 | -1.569024 | 0.13416 | 1.925534 | 0.049108 | -0.167351 | 0.05913 | -0.656625 | -0.125029 | -0.153924 | -0.162757 | -1.142817 | -0.146488 | -0.405312 | -0.319383 | | 28.65278 | 1.925534 | -0.392189 |
| 28.569 | -0.374069 | -0.012263 | -0.947034 | -0.051465 | -0.997849 | 0.86081 | 0.323169 | -0.273509 | -0.209994 | -0.011415 | -1.951507 | 0.110061 | 0.95548 | 0.080956 | -0.276544 | -0.055691 | -0.689928 | -0.146461 | -0.172339 | -0.171238 | 0.038286 | -0.167108 | -0.875886 | -0.052 | | 28.66667 | 0.95548 | -0.126347 |
| 28.583 | -0.304633 | 0.002602 | -1.148796 | 0.023689 | -1.128093 | 0.688852 | -0.167 | -0.617582 | 0.470724 | -0.014822 | -1.301986 | 0.077309 | 0.201146 | -0.105335 | -0.377089 | -0.133626 | -0.365459 | -0.045096 | -0.139494 | -0.131332 | 1.295823 | -0.127731 | -1.268641 | 0.171262 | | 28.68056 | 0.201146 | 0.138086 |
| 28.597 | -0.158141 | 0.021268 | -0.716242 | 0.25223 | -0.173343 | 0.289339 | -0.505217 | -0.359482 | 0.492465 | 0.458602 | -1.227495 | -0.034025 | -0.300741 | -0.855835 | -0.161351 | -0.111361 | -0.012015 | 0.149838 | -0.028026 | -0.029436 | 0.992552 | -0.013529 | -1.506602 | 0.326701 | | 28.69444 | -0.300741 | 0.227491 |
| 28.611 | -0.172637 | 0.000452 | 0.136815 | 0.253784 | -0.424965 | -0.023156 | -0.143255 | -0.001678 | 0.045529 | 0.609853 | -0.234173 | -0.169778 | -0.257368 | -1.522949 | 0.159334 | -0.068 | 0.234053 | 0.275201 | 0.104642 | 0.080923 | 0.332425 | 0.115974 | -0.738039 | 0.234265 | | 28.70833 | -0.257368 | 0.079635 |
| 28.625 | -0.161497 | 0.031056 | 0.515026 | 0.197927 | -0.17634 | -0.112792 | 0.6965 | 0.12542 | -0.023591 | 0.520207 | 0.885299 | -0.241795 | 0.030258 | -0.247791 | 0.285387 | -0.083946 | 0.366488 | 0.32435 | 0.17997 | 0.146556 | -0.554422 | 0.189384 | 0.118099 | 0.039113 | | 28.72222 | 0.030258 | 0.005119 |
| 28.639 | 0.015187 | 0.048019 | 0.515895 | 0.282032 | 0.497087 | -0.144276 | 1.180469 | 0.494069 | 0.040338 | -0.08102 | 1.70988 | -0.24356 | 0.554953 | 1.289812 | 0.234949 | -0.153004 | 0.347626 | 0.284345 | 0.144792 | 0.116617 | -1.311794 | 0.156061 | 0.887601 | -0.161059 | | 28.73611 | 0.554953 | -0.126739 |
| 28.653 | 0.289889 | 0.022953 | 0.703625 | 0.160143 | -0.034011 | -0.15225 | 0.95765 | 0.077433 | 0.090107 | 0.116245 | 2.252944 | -0.197316 | 0.562257 | 1.039284 | 0.139683 | -0.212462 | 0.322209 | 0.184965 | 0.058721 | 0.029163 | -1.455647 | 0.058643 | 1.130934 | -0.260471 | | 28.75 | 0.562257 | -0.294928 |
| 28.667 | 0.368988 | 0.012928 | 0.719938 | -0.011491 | 0.291178 | -0.09696 | 0.381164 | -0.227305 | 0.183257 | -0.034546 | 2.056858 | -0.019781 | 0.299075 | 0.281852 | 0.014182 | -0.168789 | 0.070368 | 0.105345 | -0.012916 | -0.040116 | -0.378141 | -0.023009 | 1.14253 | -0.180915 | | 28.76389 | 0.299075 | -0.200771 |
| 28.681 | 0.099452 | 0.017395 | 0.604969 | -0.093016 | 0.262641 | -0.076986 | -0.510274 | -0.453484 | 0.010602 | -0.68225 | 0.754622 | 0.140182 | -0.404341 | -0.196745 | -0.445951 | -0.057351 | -0.245071 | 0.033918 | -0.070458 | -0.086318 | 0.798706 | -0.07329 | 1.231088 | -0.07829 | | 28.77778 | -0.404341 | 0.026444 |
| 28.694 | -0.293461 | 0.027274 | 0.492684 | -0.175485 | 0.201611 | -0.10912 | -1.478482 | -0.583522 | -0.303669 | -0.671802 | -0.036268 | 0.332666 | -1.847983 | -0.461854 | -0.797844 | 0.008182 | -0.274149 | -0.032015 | -0.091842 | -0.109694 | 1.270979 | -0.091533 | 0.586986 | 0.014355 | | 28.79167 | -1.847983 | 0.125865 |
| 28.708 | -0.597221 | 0.001377 | -0.124342 | -0.166292 | 0.218497 | -0.213729 | -2.072166 | -0.601824 | -0.444814 | -0.612533 | -0.788281 | 0.513323 | -2.325181 | -1.714916 | -0.826171 | -0.009293 | -0.200282 | -0.079379 | -0.081143 | -0.107874 | 0.811804 | -0.087585 | -0.036131 | 0.031565 | | 28.80556 | -2.325181 | 0.075888 |
| 28.722 | -0.627364 | -0.070928 | -0.477563 | -0.071037 | -0.291621 | -0.389066 | -1.787686 | -0.429086 | -0.558552 | 0.017449 | -1.081725 | 0.657224 | -0.806877 | -2.650221 | -0.715757 | -0.088219 | -0.117616 | -0.145781 | -0.09056 | -0.117662 | 0.062522 | -0.094389 | -0.410763 | -0.061523 | | 28.81944 | -0.806877 | -0.080242 |
| 28.736 | -0.775271 | -0.152699 | -1.139378 | -0.073282 | 0.196316 | -0.593705 | -1.380537 | -0.384014 | -0.343111 | 0.195589 | -1.159157 | 0.693453 | 0.864073 | -2.23111 | -0.484228 | -0.232137 | -0.236274 | -0.249368 | -0.12087 | -0.137493 | -0.827678 | -0.116139 | -0.666243 | -0.21447 | | 28.83333 | 0.864073 | -0.23087 |
| 28.75 | -0.963999 | -0.179421 | -1.183665 | 0.058604 | 0.081127 | -0.791512 | -0.520258 | -0.219256 | 0.197773 | 0.014991 | -1.38781 | 0.470529 | 1.034803 | -1.243338 | -0.388615 | -0.414008 | -0.517905 | -0.3801 | -0.129112 | -0.128735 | -1.220984 | -0.122302 | -0.834943 | -0.316391 | | 28.84722 | 1.034803 | -0.344212 |
| 28.764 | -1.060988 | -0.181667 | -0.857892 | 0.595117 | -0.146146 | -0.679905 | 0.810377 | -0.004212 | 0.396 | -0.18307 | -2.063437 | -0.100278 | 0.492336 | -0.846415 | -0.47571 | -0.633995 | -0.854434 | -0.514625 | -0.088054 | -0.092936 | -1.975578 | -0.09566 | -0.812226 | -0.34289 | | 28.86111 | 0.492336 | -0.425137 |
| 28.778 | -1.129265 | -0.27477 | -0.869822 | 0.540621 | -0.34957 | -0.406481 | 0.810107 | -0.104854 | -0.010342 | -0.354359 | -1.795614 | -0.853251 | -0.076145 | -0.950363 | -0.6003 | -0.839454 | -1.130101 | -0.634212 | -0.033003 | -0.058614 | -1.690929 | -0.050262 | -0.780357 | -0.380136 | | 28.875 | -0.076145 | -0.42441 |
| 28.792 | -1.083701 | -0.472641 | -0.428074 | 0.160381 | -0.559169 | -0.281348 | -0.296097 | -0.199917 | -0.396476 | -0.535866 | -0.209436 | -1.530071 | -0.410242 | -1.228058 | -0.757162 | -0.694413 | -1.465495 | -0.618338 | 0.0166 | -0.030275 | -0.705225 | -0.005204 | -1.010648 | -0.370397 | | 28.88889 | -0.410242 | -0.357074 |
| 28.806 | -0.921276 | -0.578824 | 0.386675 | -0.261698 | -0.045249 | -0.366255 | -1.234942 | -0.295549 | -0.945378 | -0.651415 | 1.310823 | -2.229644 | -0.440379 | -1.538311 | -0.858841 | -0.675707 | -1.72348 | -0.593936 | 0.083494 | -0.006294 | -0.133617 | 0.02442 | -1.249998 | -0.365597 | | 28.90278 | -0.440379 | -0.307542 |
| 28.819 | -1.073454 | -0.605272 | 0.632801 | -0.345701 | -0.453016 | -0.706226 | -1.765559 | -0.852162 | -1.363382 | -0.575025 | 1.638922 | -3.011838 | -0.406638 | -1.593613 | -1.070857 | -0.654495 | -1.702549 | -0.526529 | 0.073434 | 0.018805 | 0.007078 | 0.048967 | -1.265258 | -0.46419 | | 28.91667 | -0.406638 | -0.374298 |
| 28.833 | -1.50291 | -0.640431 | 0.273882 | -0.057068 | -0.941539 | -1.188564 | -1.716258 | -1.593875 | -1.243402 | -0.724334 | -1.021338 | -3.32149 | -0.74437 | -1.442532 | -1.392839 | -0.518342 | -1.507683 | -0.405563 | 0.089327 | 0.045541 | -0.427394 | 0.079597 | -1.079953 | -0.633747 | | 28.93056 | -0.74437 | -0.612568 |
| 28.847 | -2.013658 | -0.52445 | -0.338173 | 0.102191 | -0.855136 | -1.329564 | -1.157492 | -2.139447 | -1.158355 | -0.935143 | -1.763553 | -2.551516 | -1.403562 | -1.315568 | -1.735231 | -0.483197 | -1.018883 | -0.295387 | 0.112957 | 0.077202 | -0.999321 | 0.111555 | -0.919651 | -0.834264 | | 28.94444 | -1.403562 | -0.899801 |
| 28.861 | -2.174766 | -0.429451 | -0.991566 | -0.054215 | -0.61113 | -1.383996 | -1.744748 | -2.003535 | -1.023666 | -1.172132 | -2.564781 | -1.641584 | -1.829601 | -1.336802 | -1.435165 | -0.336836 | -0.554559 | -0.213785 | 0.160536 | 0.120166 | -1.336589 | 0.15708 | -0.698916 | -0.873369 | | 28.95833 | -1.829601 | -1.000395 |
| 28.875 | -2.142383 | -0.434773 | -2.079591 | -0.2895 | -0.3792 | -1.332645 | -2.384217 | -1.773561 | -1.190483 | -1.230183 | -3.841689 | -0.570556 | -2.065783 | -1.304843 | -0.951822 | -0.247448 | -0.215086 | -0.12901 | 0.19709 | 0.147397 | -1.381301 | 0.198989 | -0.505635 | -0.756662 | | 28.97222 | -2.065783 | -0.886644 |
| 28.889 | -2.020074 | -0.546151 | -2.246987 | -0.517364 | -0.144979 | -1.002103 | -2.39619 | -2.037662 | -1.576941 | -1.587894 | -3.381403 | 0.283471 | -2.214418 | -1.100543 | -0.392994 | -0.390105 | -0.105974 | -0.024881 | 0.155464 | 0.112539 | -1.30836 | 0.169792 | -0.409306 | -0.611701 | | 28.98611 | -2.214418 | -0.702049 |
| 28.903 | -1.591211 | -0.733935 | -1.623432 | -0.699856 | -0.001532 | -0.761426 | -1.130422 | -2.335328 | -1.868488 | -1.686667 | -2.278462 | 0.7906 | -2.044736 | -0.805842 | 0.200633 | -0.332571 | -0.08659 | 0.046047 | 0.095586 | 0.056862 | -1.072757 | 0.105675 | -0.349475 | -0.408466 | | 29 | -2.044736 | -0.464672 |
| 28.917 | -0.922028 | -0.805374 | -0.709052 | -0.579695 | 0.042036 | -0.549955 | -0.254436 | -2.204247 | -1.453497 | -1.288566 | -0.833142 | 0.657978 | -1.30931 | -0.490008 | 0.765057 | 0.067909 | -0.004786 | 0.169599 | 0.211613 | 0.14057 | -0.664437 | 0.197706 | -0.18195 | -0.120424 | | 29.01389 | -1.30931 | -0.135546 |
| 28.931 | -0.267588 | -0.724652 | 0.17592 | -0.284219 | 0.166786 | -0.29765 | -0.16923 | -1.144002 | -0.837081 | -0.882962 | 0.522332 | 0.769386 | -0.305225 | -0.152669 | 0.966466 | 0.398396 | 0.195912 | 0.369731 | 0.508831 | 0.369624 | -0.199041 | 0.466568 | 0.121994 | 0.178509 | | 29.02778 | -0.305225 | 0.220962 |
| 28.944 | -0.110342 | -0.710989 | 0.555179 | -0.218906 | 0.124587 | -0.27321 | -1.028975 | -0.786384 | -0.460456 | -0.699362 | 1.178711 | 0.759493 | 0.351784 | -0.145937 | 0.527115 | 0.086717 | 0.206201 | 0.26483 | 0.554614 | 0.375475 | -0.104876 | 0.512723 | 0.202372 | 0.203914 | | 29.04167 | 0.351784 | 0.263475 |
| 28.958 | -0.978304 | -1.007245 | 0.188841 | -0.569905 | -0.538663 | -0.970012 | -2.185462 | -1.485886 | -0.871987 | -0.918563 | 0.258505 | 0.698645 | 0.031528 | -0.443436 | -0.287515 | -0.906358 | -0.188345 | -0.189598 | 0.101374 | 0.035355 | -0.444596 | 0.106033 | -0.136439 | -0.175379 | | 29.05556 | 0.031528 | -0.127198 |
| 28.972 | -2.886448 | -2.138394 | -0.51248 | -1.369014 | -1.279636 | -2.230037 | -1.872339 | -2.758177 | -1.995664 | -1.282178 | -1.582734 | -0.68718 | -1.167997 | -0.786921 | -0.957079 | -1.677677 | -0.648258 | -0.463347 | -0.096018 | -0.103861 | -0.938851 | -0.122443 | -0.383586 | -0.664798 | | 29.06944 | -1.167997 | -0.678245 |
| 28.986 | -4.503311 | -2.547603 | -1.301936 | -1.670757 | -1.572106 | -2.807587 | -1.178022 | -3.67528 | -2.382352 | -1.167832 | -2.630733 | -3.010436 | -2.251033 | -1.241219 | -1.454327 | -1.782838 | -1.2394 | -0.63449 | 0.051169 | 0.037702 | -1.549514 | 0.022333 | -0.505704 | -1.005058 | | 29.08333 | -2.251033 | -1.163693 |
| 29 | -2.837935 | -1.869228 | -1.477759 | -1.843912 | -1.058644 | -2.561048 | -1.869781 | -3.466882 | -2.054081 | -0.610521 | -1.98822 | -2.499817 | -2.49489 | -1.423527 | -1.273478 | -1.697122 | -1.321945 | -0.933806 | 0.111527 | 0.093752 | -1.872013 | 0.105643 | -0.707136 | -1.018608 | | 29.09722 | -2.49489 | -1.288893 |
| 29.014 | -1.074585 | -1.479369 | -0.752476 | -1.632283 | -0.416171 | -2.076353 | -3.86845 | -1.997519 | -1.576849 | -0.00852 | -1.288337 | -2.502818 | -2.183011 | -0.954183 | -0.202844 | -1.625848 | -0.408864 | -1.214212 | 0.064561 | 0.061775 | -1.297438 | 0.063324 | -0.829463 | -0.738926 | | 29.11111 | -2.183011 | -1.00594 |
| 29.028 | 0.084754 | -1.038931 | 0.029755 | -0.792915 | 0.045501 | -0.896539 | -4.027705 | -0.633398 | -0.96059 | 0.382515 | -0.214718 | -1.624138 | -1.424774 | -0.225838 | 0.463702 | -1.398888 | 0.116457 | -1.21551 | 0.036025 | 0.02957 | -0.468172 | 0.03473 | -0.918395 | -0.364892 | | 29.125 | -1.424774 | -0.558579 |
| 29.042 | 0.759876 | -0.659159 | 0.69086 | -0.434232 | 0.219082 | 0.182804 | -2.868286 | 0.341803 | -0.433325 | 0.520297 | 0.709855 | -0.412729 | -0.548853 | 0.228369 | 0.783978 | -1.033361 | 0.295531 | -1.124478 | 0.001528 | -0.02144 | 0.101282 | 0.007808 | -1.054552 | -0.125601 | | 29.13889 | -0.548853 | -0.249212 |
| 29.056 | 1.446026 | -0.338782 | 0.986556 | 0.018781 | 0.168984 | 1.063352 | -1.758893 | 0.807924 | 0.073484 | 0.528093 | 1.301092 | 0.241134 | 0.282009 | 0.335629 | 0.738648 | -0.679943 | 0.249131 | -0.837013 | -0.023523 | -0.070331 | 0.360468 | -0.011825 | -0.794097 | -0.04352 | | 29.15278 | 0.282009 | -0.075969 |
| 29.069 | 2.665633 | -0.220766 | 0.790326 | 0.987631 | 0.092549 | 1.552551 | -0.399928 | 1.087649 | 0.800774 | 0.511331 | 1.841509 | 0.660313 | 1.110255 | 0.205567 | 0.512212 | -0.133638 | 0.170248 | -0.627928 | -0.005175 | -0.068513 | 0.255095 | 0.000738 | -0.298572 | -0.049951 | | 29.16667 | 1.110255 | 0.018843 |
| 29.083 | 3.212441 | -0.632227 | 0.337197 | 2.104075 | 0.023956 | 1.617535 | 0.953686 | 1.246002 | 1.666361 | 0.467178 | 1.987235 | 0.821127 | 1.82304 | 0.055782 | 0.467501 | 0.40751 | 0.210583 | -0.615649 | 0.057436 | -0.01009 | 0.043728 | 0.051102 | 0.231598 | 0.010689 | | 29.18056 | 1.82304 | 0.102331 |
| 29.097 | 1.913374 | -2.152505 | -0.163241 | 1.642493 | 0.012887 | 0.958734 | 2.183468 | 1.309222 | 1.967744 | 0.98085 | 1.162562 | 0.703708 | 1.91652 | 0.324793 | 0.774866 | 0.640127 | 0.48038 | -0.749858 | 0.145384 | 0.083859 | 0.190143 | 0.134467 | 0.828853 | 0.215958 | | 29.19444 | 1.91652 | 0.291666 |
| 29.111 | 1.238272 | -3.331745 | -0.687311 | 0.274378 | 0.960936 | 0.186325 | 3.161501 | 1.251187 | 1.023923 | 0.654926 | 0.111223 | 0.430889 | 1.026513 | 1.064764 | 0.934841 | 0.314396 | 0.970923 | -0.82352 | 0.201126 | 0.150231 | 0.856214 | 0.194652 | 1.004254 | 0.508464 | | 29.20833 | 1.026513 | 0.593459 |
| 29.125 | 0.617419 | -0.697277 | -1.421306 | -0.620098 | 1.728882 | -0.508116 | 3.217826 | 1.160895 | -0.124839 | 0.322422 | 0.052533 | 0.056182 | -0.094814 | 2.028445 | 0.589122 | 0.058633 | 1.218719 | -0.826161 | 0.222641 | 0.176414 | 1.532937 | 0.218542 | 1.164951 | 0.778438 | | 29.22222 | -0.094814 | 0.860701 |
| 29.139 | 1.408032 | 0.555054 | -1.095156 | -0.305531 | 1.084671 | -0.72004 | 1.648685 | 0.990683 | -0.210554 | 0.110206 | -0.348478 | -0.244903 | -0.167985 | 2.317932 | 0.318795 | 0.142296 | 0.929729 | -0.59063 | 0.2805 | 0.221541 | 1.607129 | 0.272724 | 1.532859 | 0.884447 | | 29.23611 | -0.167985 | 0.960758 |
| 29.153 | 2.585926 | 1.277573 | -0.276042 | 1.12128 | 0.485147 | -0.38285 | 0.376274 | 0.56488 | 0.432036 | 0.20594 | -0.613809 | -0.20932 | 0.613803 | 1.500768 | 0.44652 | 0.46613 | 0.682381 | -0.277051 | 0.322155 | 0.260677 | 1.503859 | 0.318046 | 1.699905 | 0.817146 | | 29.25 | 0.613803 | 0.808945 |
| 29.167 | 0.58663 | 1.366096 | 0.261831 | 1.783572 | 0.299506 | -0.176937 | -0.939832 | -0.09197 | 0.751194 | 0.452125 | -1.011952 | 0.333833 | 1.191435 | 0.44769 | 0.702971 | 0.676111 | 0.704434 | -0.102527 | 0.215424 | 0.172306 | 1.58861 | 0.218866 | 1.378251 | 0.587824 | | 29.26389 | 1.191435 | 0.534618 |
| 29.181 | -0.637922 | 0.993001 | 0.504319 | 0.759968 | 0.016465 | 0.008411 | -1.610098 | -0.234247 | 0.105299 | 1.186904 | -0.81164 | 1.243688 | 1.18861 | -0.327459 | 0.928612 | 0.452847 | 0.893635 | -0.141771 | 0.021541 | 0.01143 | 1.708272 | 0.025986 | 0.777505 | 0.473006 | | 29.27778 | 1.18861 | 0.463745 |
| 29.194 | -0.845415 | 0.342351 | 0.80477 | -0.122075 | -0.499015 | 0.35563 | -1.102959 | 0.100313 | -0.423052 | 2.401301 | 0.07199 | 1.745601 | 0.611136 | -0.547643 | 0.995973 | 0.199242 | 1.119095 | -0.223077 | -0.099767 | -0.09062 | 1.893057 | -0.107674 | 0.929765 | 0.504829 | | 29.29167 | 0.611136 | 0.615759 |
| 29.208 | 0.768338 | -0.282417 | 1.297689 | -0.870688 | -0.414761 | 0.688119 | 0.146242 | 0.645509 | 0.070416 | 1.237394 | 0.796012 | 1.554333 | 0.094227 | 0.004783 | 0.896229 | 0.270904 | 1.215048 | -0.227866 | -0.09818 | -0.100265 | 1.635115 | -0.122607 | 0.114361 | 0.592401 | | 29.30556 | 0.094227 | 0.701885 |
| 29.222 | 2.019187 | -0.719078 | 1.351158 | 0.447653 | 0.163191 | 0.37248 | 1.402342 | 0.988355 | 1.242223 | -0.09837 | 0.717818 | 1.551409 | -0.011283 | 1.246958 | 0.721945 | 0.48909 | 1.39734 | -0.063125 | 0.00377 | -0.010643 | 1.217949 | -0.02401 | -0.488088 | 0.729347 | | 29.31944 | -0.011283 | 0.828876 |
| 29.236 | 1.614977 | -0.923719 | 0.827402 | 1.515121 | 0.569186 | -0.038517 | 1.730654 | 0.243235 | 2.129447 | -1.236966 | 0.076605 | 1.546592 | 0.072519 | 2.343355 | 0.658796 | 0.606751 | 1.441654 | 0.260159 | 0.12305 | 0.109269 | 0.924403 | 0.100121 | -0.678457 | 0.795136 | | 29.33333 | 0.072519 | 0.942595 |
| 29.25 | 1.106661 | -0.918567 | 0.255977 | 0.671588 | 0.699378 | 0.131914 | 0.96128 | 0.325692 | 1.275442 | 0.00313 | -0.199451 | 1.199101 | 0.518368 | 2.394193 | 0.591641 | 0.374101 | 1.384535 | 0.536883 | 0.149722 | 0.131921 | 1.031712 | 0.12395 | -0.364735 | 0.799523 | | 29.34722 | 0.518368 | 0.836648 |
| 29.264 | 0.665618 | -0.684529 | 0.130591 | 0.022179 | 0.882152 | 1.001542 | 0.203628 | 1.037008 | -0.163205 | 0.714258 | 0.349937 | 1.299026 | 1.339019 | 2.007007 | 0.689641 | 0.20873 | 1.095576 | 0.559598 | 0.104902 | 0.062011 | 1.512207 | 0.069211 | 0.201879 | 0.75009 | | 29.36111 | 1.339019 | 0.690694 |
| 29.278 | 0.877132 | -0.401597 | 0.494108 | -0.345596 | 1.260066 | 1.50145 | -0.162373 | 1.968255 | -0.494465 | 1.342789 | 0.875629 | 1.524961 | 2.034437 | 1.441969 | 1.146848 | 0.293609 | 0.779966 | 0.468527 | 0.075237 | 0.010693 | 1.791883 | 0.037354 | 0.433217 | 0.709489 | | 29.375 | 2.034437 | 0.660013 |
| 29.292 | 0.748359 | -0.05133 | 0.719636 | -0.15125 | 1.554269 | 1.068443 | -0.084457 | 2.684846 | 0.205326 | 1.318829 | 0.64326 | 1.368025 | 2.066033 | 1.100658 | 1.683376 | 0.339338 | 0.552922 | 0.404819 | 0.028918 | -0.024893 | 1.691172 | -0.004565 | 1.065193 | 0.634069 | | 29.38889 | 2.066033 | 0.607131 |
| 29.306 | 0.128572 | 0.274442 | -0.17916 | 0.350673 | 1.342927 | 0.50187 | 0.278326 | 1.312693 | 1.337949 | 0.561419 | 0.13888 | 1.046318 | 1.253305 | 0.638384 | 1.886441 | 0.273183 | 0.470107 | 0.454651 | -0.010798 | -0.038818 | 1.48218 | -0.046091 | 1.790645 | 0.544676 | | 29.40278 | 1.253305 | 0.499226 |
| 29.319 | 0.154715 | 0.50703 | -2.305468 | 0.609001 | 0.901346 | 0.546402 | 0.797811 | 0.549335 | 1.663476 | 0.090258 | -0.079564 | 0.742272 | 0.024163 | 0.605208 | 1.73976 | 0.435156 | 0.568508 | 0.676799 | 0.054666 | 0.021253 | 1.381 | 0.013225 | 1.937362 | 0.56272 | | 29.41667 | 0.024163 | 0.601691 |
| 29.333 | 0.980944 | 0.621205 | -2.431983 | 0.456379 | 0.494394 | 1.149385 | 1.32385 | 1.410333 | 1.174758 | -0.108545 | 0.212762 | 0.838694 | -0.67873 | 1.19758 | 1.465152 | 0.835204 | 0.799979 | 1.010021 | 0.112447 | 0.051944 | 1.440524 | 0.076407 | 1.966148 | 0.713794 | | 29.43056 | -0.67873 | 0.811052 |
| 29.347 | 2.05835 | 0.668121 | -0.989254 | 0.336432 | 0.043374 | 1.607938 | 1.747287 | 2.640036 | 0.788069 | 0.310786 | 0.851143 | 1.116575 | 0.399916 | 2.207521 | 1.115274 | 1.190797 | 1.001712 | 1.113242 | -0.02647 | -0.082976 | 1.376802 | -0.046345 | 1.878613 | 0.820969 | | 29.44444 | 0.399916 | 0.937698 |
| 29.361 | 2.756915 | 0.553802 | 1.906392 | 0.161535 | -0.100995 | 1.472355 | 2.37482 | 2.401208 | 1.075073 | 0.632726 | 1.697954 | 1.222747 | 1.173723 | 3.339307 | 1.265818 | 1.350261 | 1.041899 | 0.732092 | -0.378732 | -0.386719 | 1.002294 | -0.426915 | 1.951938 | 0.786056 | | 29.45833 | 1.173723 | 0.881911 |
| 29.375 | 1.944096 | 0.258616 | 2.482487 | -0.104382 | 0.248072 | 1.24514 | 2.782107 | 2.456922 | 1.583068 | 0.994722 | 2.485093 | 1.087489 | 1.211838 | 2.666598 | 1.188174 | 1.417348 | 0.87113 | 0.47402 | -0.812952 | -0.686231 | 0.475997 | -0.823549 | 1.745086 | 0.414446 | | 29.47222 | 1.211838 | 0.622169 |
| 29.389 | 0.78444 | 0.077057 | 0.356741 | 0.005827 | 0.790586 | 1.406687 | 2.929577 | 2.106676 | 1.946397 | 1.373428 | 2.94972 | 1.076835 | 0.996729 | 1.288684 | 0.890648 | 1.045593 | 0.496177 | 0.589007 | -0.701259 | -0.582033 | 0.146822 | -0.615232 | 0.908929 | 0.222664 | | 29.48611 | 0.996729 | 0.32613 |
| 29.403 | 0.713341 | 0.09506 | -0.255546 | 0.436782 | 1.661498 | 1.779077 | 3.217598 | 1.580236 | 1.799189 | 1.132221 | 3.465694 | 1.364177 | 1.592466 | 1.256057 | 0.912352 | 0.93231 | 0.354558 | 1.00739 | -0.228782 | -0.225189 | 0.521903 | -0.181299 | 0.892388 | 0.32094 | | 29.5 | 1.592466 | 0.387266 |
| 29.417 | 1.515673 | 0.138148 | 1.355249 | 0.545461 | 2.549434 | 1.888898 | 3.20044 | 0.871552 | 1.148272 | 1.541995 | 3.083364 | 1.593048 | 1.988534 | 1.492069 | 1.066106 | 1.053412 | 0.349817 | 1.155172 | 0.065868 | 0.026913 | 1.17336 | 0.087771 | 0.735034 | 0.61722 | | 29.51389 | 1.988534 | 0.572379 |
| 29.431 | 2.292687 | 0.289725 | 1.511701 | 0.631929 | 2.966688 | 1.530861 | 3.111595 | 0.58571 | 0.435444 | 1.523739 | 2.159188 | 1.722074 | 2.125159 | 1.709964 | 0.9682 | 1.120566 | 0.351763 | 1.164907 | 0.197762 | 0.140044 | 1.576737 | 0.199008 | 1.551077 | 0.780522 | | 29.52778 | 2.125159 | 0.648839 |
| 29.444 | 2.914028 | 0.66533 | 0.81974 | 0.547393 | 2.172238 | 1.24407 | 2.611654 | 0.113327 | 0.606315 | 1.299592 | 2.364668 | 1.936474 | 1.93148 | 1.343259 | 0.526501 | 0.927761 | 0.112891 | 0.832379 | -0.079836 | -0.185719 | 1.207377 | -0.03061 | 1.019813 | 0.562544 | | 29.54167 | 1.93148 | 0.346085 |
| 29.458 | 1.766596 | 1.106891 | 0.084941 | 0.284583 | 1.408967 | 1.357834 | 1.953074 | 0.402554 | 1.197211 | 0.544858 | 2.839348 | 2.278231 | 1.310039 | 0.481404 | -0.151313 | 0.266123 | -0.503289 | 0.223927 | -0.857982 | -1.014303 | 0.313729 | -0.716085 | 0.011195 | -0.166116 | | 29.55556 | 1.310039 | -0.343174 |
| 29.472 | 0.976883 | 1.063417 | -0.199286 | 0.038499 | 0.985387 | 1.055635 | 1.069813 | 0.387221 | 0.829377 | 0.829187 | 2.38623 | 1.810276 | 0.688787 | 0.009672 | -0.459316 | -0.063065 | -0.724468 | -0.210884 | -1.249218 | -1.373756 | 0.090812 | -1.115775 | -1.117896 | -0.676284 | | 29.56944 | 0.688787 | -0.708062 |
| 29.486 | 0.327979 | 0.617412 | -0.140396 | -0.012366 | 0.969676 | 0.324454 | 0.773258 | 0.439081 | 1.132759 | 0.636068 | 1.292513 | 1.256919 | -0.083009 | 0.716964 | -0.069793 | 0.010583 | -0.117826 | -0.10913 | -0.674835 | -0.728941 | 0.553464 | -0.665848 | -1.616443 | -0.280968 | | 29.58333 | -0.083009 | -0.219169 |
| 29.5 | 0.081918 | 0.396058 | -0.580472 | 0.025666 | 0.961468 | -0.022435 | 0.42919 | 0.107782 | 0.546591 | 0.597016 | 0.794624 | 1.365643 | -0.668677 | 1.125775 | 0.633708 | 0.079592 | 0.585146 | 0.180567 | 0.098178 | 0.049589 | 1.656224 | 0.050903 | 0.203483 | 0.378966 | | 29.59722 | -0.668677 | 0.29515 |
| 29.514 | 0.431311 | 0.001737 | -0.856103 | 0.050307 | 0.979075 | -0.345899 | 0.310393 | -0.460181 | 0.343535 | 0.5976 | -0.067203 | 1.524335 | -0.641728 | 0.917227 | 0.930383 | 0.191393 | 0.722549 | 0.449516 | 0.468965 | 0.403081 | 1.431012 | 0.459267 | 0.627535 | 0.611276 | | 29.61111 | -0.641728 | 0.447425 |
| 29.528 | 0.024318 | -0.524315 | -0.181208 | -0.080747 | 0.176054 | -0.40059 | 0.335559 | 0.235223 | -0.305625 | 0.490348 | -0.729623 | 1.548887 | 0.108576 | 0.550129 | 0.391271 | 0.346532 | 0.404809 | 0.639433 | 0.416004 | 0.355294 | 0.795003 | 0.444579 | 0.920294 | 0.520997 | | 29.625 | 0.108576 | 0.412282 |
| 29.542 | -0.321913 | -1.024011 | 0.419886 | 0.08174 | -0.151479 | 0.420799 | 0.429104 | 0.573478 | -0.831257 | 0.087577 | -1.987198 | 1.496336 | 0.913668 | 0.005172 | -0.24006 | 0.431796 | 0.004143 | 0.574593 | 0.145874 | 0.113957 | -0.000813 | 0.169093 | 0.768427 | 0.259706 | | 29.63889 | 0.913668 | 0.275838 |
| 29.556 | -0.734041 | -1.274435 | 0.289715 | 0.165996 | -0.354832 | 1.458282 | 0.4768 | 0.731746 | -0.060477 | 0.304645 | -2.450916 | 1.306894 | 1.791056 | -0.552047 | -0.455187 | 0.369223 | -0.398667 | 0.316542 | -0.104041 | -0.103966 | -1.522722 | -0.103078 | 0.532892 | -0.043724 | | 29.65278 | 1.791056 | 0.080965 |
| 29.569 | -0.949921 | -1.136612 | 0.047457 | -0.016157 | -0.239402 | 0.773199 | 0.389335 | 0.439271 | 0.744619 | 0.907949 | -1.387022 | 0.853648 | 2.36779 | 0.03775 | -0.402797 | 0.18195 | -0.567731 | 0.082773 | -0.214173 | -0.198874 | -0.705904 | -0.216193 | 0.384381 | -0.109407 | | 29.66667 | 2.36779 | 0.00248 |
| 29.583 | -0.47266 | -0.710804 | -0.06704 | 0.248249 | -1.114793 | 0.281275 | 0.113086 | -0.101108 | 0.312577 | 0.795347 | 0.058258 | 0.149971 | 1.516634 | 0.020433 | -0.178685 | -0.030614 | -0.582511 | 0.043148 | -0.18332 | -0.166184 | 0.51494 | -0.183917 | 0.276818 | -0.017603 | | 29.68056 | 1.516634 | 0.0545 |
| 29.597 | 0.095967 | -0.218425 | -0.6192 | 0.227734 | -2.076473 | 0.065831 | -0.186518 | -0.08177 | -0.097901 | 0.083827 | 1.335504 | -0.621591 | -0.354049 | -0.461267 | 0.059567 | -0.203602 | -0.462105 | 0.147376 | -0.087299 | -0.077987 | 1.592786 | -0.090843 | 0.35167 | 0.042887 | | 29.69444 | -0.354049 | -0.055987 |
| 29.611 | 0.302042 | 0.320326 | 0.071018 | -0.084244 | -0.467871 | 0.141747 | -0.327302 | -0.136951 | -0.253149 | -0.695337 | 1.987135 | -1.495311 | -1.553473 | -0.611698 | -0.064376 | -0.378726 | -0.414706 | 0.199715 | -0.064695 | -0.057845 | 0.842427 | -0.057071 | -0.359206 | -0.215574 | | 29.70833 | -1.553473 | -0.434439 |
| 29.625 | 0.323948 | 0.923592 | 0.250505 | -0.028588 | 0.755773 | 0.281475 | -0.30345 | 0.167241 | -0.121986 | -0.293738 | 1.759452 | -2.144766 | -1.032442 | -1.078514 | -0.232029 | -0.555468 | -0.528816 | 0.126244 | -0.158535 | -0.150023 | 0.097263 | -0.135522 | -0.266727 | -0.479747 | | 29.72222 | -1.032442 | -0.797446 |
| 29.639 | 0.472726 | 1.235873 | 0.426123 | -0.005334 | 0.73072 | 0.267207 | -0.14495 | 0.199331 | 0.017134 | 0.564605 | 1.471885 | -2.176211 | 0.024573 | -1.873661 | -0.424085 | -0.472594 | -0.603837 | 0.037188 | -0.278281 | -0.265796 | -0.756639 | -0.258686 | -0.299725 | -0.737711 | | 29.73611 | 0.024573 | -0.849551 |
| 29.653 | 0.811705 | 1.325767 | 0.762176 | 0.008224 | -0.768074 | 0.11929 | 0.114549 | -0.163137 | -0.634375 | 0.701714 | 0.813437 | -1.916124 | 0.894503 | -1.968712 | -0.722504 | -0.132814 | -0.460219 | 0.032285 | -0.34999 | -0.321785 | -1.758742 | -0.342226 | -0.359439 | -0.81388 | | 29.75 | 0.894503 | -0.774948 |
| 29.667 | 1.098167 | 1.447381 | 0.723003 | 0.092013 | -0.644579 | -0.026735 | 0.192826 | -0.18669 | -0.24006 | 0.098203 | -0.046169 | -0.932976 | 1.216518 | -0.751581 | -0.777398 | 0.162217 | -0.095587 | 0.114705 | -0.340927 | -0.294994 | -1.949453 | -0.330623 | -0.383375 | -0.699168 | | 29.76389 | 1.216518 | -0.571949 |
| 29.681 | 0.893272 | 1.521905 | 0.655468 | 0.198677 | -0.271953 | -0.180837 | -0.115383 | -0.17671 | 0.371856 | -0.86459 | -0.749894 | -0.077173 | 0.90737 | 0.218796 | -0.743027 | 0.13628 | 0.282892 | 0.223953 | -0.276671 | -0.222461 | -1.321421 | -0.264066 | -0.576036 | -0.476368 | | 29.77778 | 0.90737 | -0.284823 |
| 29.694 | 0.29175 | 1.199879 | 0.671052 | 0.206024 | -0.280757 | -0.272159 | 0.585489 | -0.287202 | 0.531484 | 0.447666 | -1.369956 | 0.793591 | 0.085598 | 0.295379 | -0.31878 | -0.017259 | 0.371865 | 0.240484 | -0.252751 | -0.199725 | -0.402933 | -0.247371 | -0.75439 | -0.252507 | | 29.79167 | 0.085598 | -0.047269 |
| 29.708 | -0.435288 | 0.930777 | 0.446612 | 0.029954 | -0.558674 | -0.269798 | 0.558964 | -0.567178 | 0.896364 | 0.228811 | -1.902274 | 2.025575 | -0.453282 | 0.052582 | -0.180568 | -0.16723 | 0.250181 | 0.14541 | -0.261548 | -0.209227 | -0.01736 | -0.255303 | -0.833695 | -0.126071 | | 29.80556 | -0.453282 | 0.017069 |
| 29.722 | -1.286873 | 0.702746 | 0.214583 | -0.116326 | -0.750719 | -0.241856 | 0.113239 | -0.422643 | 0.285859 | -0.793516 | -2.155416 | 2.515946 | -0.514647 | -0.164075 | -0.428525 | -0.271865 | 0.051584 | 0.112267 | -0.233685 | -0.175557 | -0.195818 | -0.216372 | -0.67673 | -0.090232 | | 29.81944 | -0.514647 | -0.036237 |
| 29.736 | -1.718635 | 0.53516 | -0.343939 | -0.030056 | -0.173282 | -0.212993 | -1.106558 | -0.02034 | 0.364409 | 0.044214 | -1.579126 | 1.742626 | -0.760063 | -0.493153 | -0.717513 | -0.41533 | -0.335501 | 0.058044 | -0.212202 | -0.160344 | 0.127381 | -0.190076 | -0.387942 | -0.216711 | | 29.83333 | -0.760063 | -0.218653 |
| 29.75 | -1.927004 | 0.398873 | -0.137761 | -0.199001 | 0.352402 | -0.210653 | -1.825425 | 0.318579 | -0.481494 | -0.230039 | -0.286697 | 0.331263 | -0.634845 | -1.369831 | -1.003191 | -0.835258 | -0.803577 | -0.05683 | -0.24879 | -0.222817 | 0.281277 | -0.226501 | -0.353551 | -0.538458 | | 29.84722 | -0.634845 | -0.505242 |
| 29.764 | -2.029018 | 0.279866 | 0.02738 | -0.186022 | -0.07867 | -0.239132 | -1.747075 | 0.583927 | -1.086341 | -0.363282 | 1.121054 | -1.286877 | -0.47359 | -1.784962 | -1.265443 | -1.271153 | -1.162806 | -0.212616 | -0.277046 | -0.279204 | -1.09839 | -0.25306 | -0.595293 | -0.734874 | | 29.86111 | -0.47359 | -0.666345 |
| 29.778 | -1.529667 | 0.131977 | -0.117628 | -0.019894 | -0.360965 | -0.277574 | -1.421389 | 0.109605 | -0.933868 | -0.368535 | 1.444628 | -2.940172 | -0.649535 | -1.367394 | -1.300945 | -0.892734 | -1.44976 | -0.369165 | -0.219511 | -0.247872 | -1.754943 | -0.214239 | -0.648264 | -0.686793 | | 29.875 | -0.649535 | -0.655982 |
| 29.792 | -1.186842 | 0.052476 | -0.466182 | 0.030656 | -0.848703 | -0.261918 | -0.038187 | -0.647199 | -0.87845 | -0.468036 | 0.312282 | -3.158741 | -1.191966 | -1.761504 | -1.54767 | -0.720896 | -1.73592 | -0.452507 | -0.177354 | -0.197911 | -2.0633 | -0.16239 | -0.663483 | -0.675517 | | 29.88889 | -1.191966 | -0.633935 |
| 29.806 | -1.188507 | -0.005437 | -1.34326 | 0.041735 | -1.424849 | -0.36726 | 0.171364 | -1.096813 | -0.965895 | -0.821181 | -1.159589 | -2.954413 | -1.626077 | -2.512978 | -1.961829 | -0.713356 | -1.92017 | -0.505926 | -0.168174 | -0.170716 | -2.415039 | -0.142288 | -0.737166 | -0.867687 | | 29.90278 | -1.626077 | -0.783319 |
| 29.819 | -1.518732 | -0.138792 | -1.777325 | 0.112533 | -1.909752 | -0.609434 | -0.391983 | -1.66378 | -1.054585 | -1.223249 | -2.614775 | -2.697125 | -2.126368 | -3.220255 | -2.368477 | -0.897615 | -1.633562 | -0.55905 | -0.158676 | -0.154331 | -1.939362 | -0.141052 | -0.958107 | -1.064919 | | 29.91667 | -2.126368 | -1.026156 |
| 29.833 | -1.373486 | -0.342698 | -1.459921 | 0.232087 | -1.387785 | -1.091947 | -0.066654 | -1.839111 | -1.16267 | -1.416305 | -4.650075 | -2.084289 | -2.792356 | -3.727491 | -1.635148 | -1.039247 | -1.522055 | -0.660917 | -0.141033 | -0.139582 | -1.123945 | -0.136394 | -1.396019 | -1.224773 | | 29.93056 | -2.792356 | -1.251793 |
| 29.847 | -1.551565 | -0.650839 | -0.798959 | 0.098552 | -1.221603 | -1.687317 | -0.244298 | -2.045777 | -1.548958 | -1.215266 | -5.298571 | -1.256618 | -3.501494 | -3.109295 | -0.785782 | -1.399849 | -1.362874 | -0.767868 | -0.101364 | -0.112895 | -0.514312 | -0.103203 | -1.731902 | -1.189219 | | 29.94444 | -3.501494 | -1.384505 |
| 29.861 | -2.090026 | -1.060791 | -0.578737 | -0.272855 | -0.896216 | -1.874937 | -1.471713 | -2.597267 | -1.34281 | -1.342693 | -3.67187 | -0.819473 | -2.779515 | -2.262277 | -0.352451 | -2.227375 | -1.176717 | -0.777761 | -0.032438 | -0.039768 | -0.190702 | -0.035083 | -1.832631 | -0.995828 | | 29.95833 | -2.779515 | -1.310211 |
| 29.875 | -2.470669 | -1.695252 | 1.083339 | -0.379079 | -0.971783 | -1.615131 | -2.82335 | -3.410301 | -0.980199 | -1.33591 | -1.550141 | -0.706911 | -1.408555 | -1.640191 | -0.388141 | -2.196651 | -0.965446 | -0.768195 | 0.041588 | 0.062665 | -0.27082 | 0.033297 | -1.662774 | -0.770581 | | 29.97222 | -1.408555 | -0.996454 |
| 29.889 | -2.899117 | -2.158126 | 2.276316 | -0.197793 | -0.927341 | -1.368593 | -2.189021 | -3.697506 | -0.803245 | -1.386808 | -0.406592 | -0.479573 | -0.55841 | -0.986102 | -0.54406 | -1.596596 | -0.432374 | -0.640482 | 0.119344 | 0.147104 | -0.505855 | 0.104913 | -1.315891 | -0.510764 | | 29.98611 | -0.55841 | -0.644276 |
| 29.903 | -2.630724 | -1.798901 | 2.044891 | -0.049593 | -0.569247 | -1.212167 | -1.731559 | -3.052146 | -1.047283 | -1.425086 | -0.069893 | 0.084336 | -0.176382 | -0.431706 | -0.50367 | -0.706687 | 0.107241 | -0.420738 | 0.225453 | 0.222962 | -0.553048 | 0.207709 | -0.850251 | -0.181079 | | 30 | -0.176382 | -0.231414 |
| 29.917 | -1.657858 | -1.297797 | 1.883322 | 0.180958 | -0.736305 | -0.994431 | -0.605534 | -2.063803 | -1.197137 | -1.076495 | 0.133677 | 0.760383 | -0.727885 | 0.042499 | -0.321308 | 0.110288 | 0.445875 | -0.195703 | 0.406249 | 0.354595 | -0.314923 | 0.373003 | -0.371926 | 0.152755 | | 30.01389 | -0.727885 | 0.142435 |
| 29.931 | -0.784124 | -0.924271 | 1.643116 | 0.354891 | -0.588416 | -0.498211 | 0.381519 | -1.512911 | -0.79288 | -0.559126 | 0.658332 | 2.202756 | -1.744905 | 0.355391 | -0.086688 | 0.605477 | 0.620799 | 0.019871 | 0.701862 | 0.578865 | 0.021944 | 0.639232 | 0.147187 | 0.418466 | | 30.02778 | -1.744905 | 0.427202 |
| 29.944 | -0.485211 | -0.657446 | 0.247058 | 0.178946 | -0.549229 | -0.309502 | 1.44828 | -1.418423 | -0.683283 | -0.365282 | 0.739889 | 1.885732 | -2.661674 | 0.545605 | 0.124618 | 0.709526 | 0.794872 | 0.266368 | 1.075706 | 0.886964 | 0.401839 | 0.994306 | 0.639302 | 0.684682 | | 30.04167 | -2.661674 | 0.737719 |
| 29.958 | -1.196297 | -0.352524 | 0.141792 | -0.674925 | -0.933422 | -1.107115 | 0.980317 | -2.741131 | -1.4887 | -0.880476 | -0.198179 | 0.051309 | -2.901552 | 0.581991 | 0.273958 | 0.709335 | 0.79771 | 0.487888 | 1.233652 | 1.122855 | 0.684688 | 1.193135 | 0.904531 | 0.795073 | | 30.05556 | -2.901552 | 0.907636 |
| 29.972 | -2.624206 | -0.073512 | 0.618749 | -1.261973 | -1.727723 | -2.620688 | -2.227221 | -3.77364 | -2.845843 | -1.903038 | -1.576543 | -1.263488 | -3.230306 | 0.116477 | -0.006419 | 0.496129 | 0.298558 | 0.239215 | 1.082151 | 1.06986 | 0.498936 | 1.058175 | 0.632007 | 0.400162 | | 30.06944 | -3.230306 | 0.444907 |
| 29.986 | -2.992482 | 0.035729 | 1.877986 | -2.141127 | -2.173784 | -2.779695 | -2.415663 | -4.006056 | -3.410587 | -1.669602 | -2.575523 | -0.162994 | -3.252013 | -0.576368 | -0.749089 | 0.223778 | -0.756264 | -0.384875 | 0.809015 | 0.813856 | -0.204272 | 0.781685 | -0.058996 | -0.194576 | | 30.08333 | -3.252013 | -0.24631 |
| 30 | -1.793651 | -0.185597 | 0.415191 | -2.060707 | -1.412481 | -2.535484 | -2.152451 | -3.349134 | -3.766645 | -1.287374 | -2.3183 | -2.015518 | -3.003162 | -0.995318 | -1.132028 | 0.575553 | -1.261001 | -0.70691 | 0.645912 | 0.640217 | -0.983022 | 0.64825 | -0.299058 | -0.423944 | | 30.09722 | -3.003162 | -0.554425 |
| 30.014 | -0.8462 | -0.489876 | -0.925156 | -1.083574 | -0.610911 | -2.367946 | -3.917212 | -2.330514 | -2.649905 | -0.650743 | -1.809431 | -2.196901 | -1.678009 | -0.796956 | -0.741404 | 0.937485 | -0.436631 | -0.913074 | 0.500479 | 0.493742 | -1.191951 | 0.524883 | -0.293944 | -0.362211 | | 30.11111 | -1.678009 | -0.500119 |
| 30.028 | -0.147479 | -0.624512 | -1.128113 | -0.341624 | 0.180199 | -1.722649 | -4.097383 | -1.078633 | -0.936826 | 0.017417 | -1.034791 | -0.239004 | -0.499985 | -0.313687 | -0.125379 | 0.608247 | 0.232036 | -1.046615 | 0.308317 | 0.305226 | -0.708846 | 0.328897 | -0.356835 | -0.170964 | | 30.125 | -0.499985 | -0.238758 |
| 30.042 | 0.522817 | -0.564349 | -0.549135 | -0.170672 | 0.303444 | -0.671317 | -2.641681 | 0.228202 | -0.00118 | 0.490274 | -0.211469 | 0.595358 | 0.081277 | 0.504008 | 0.371556 | -0.075553 | 0.676402 | -0.88347 | 0.194295 | 0.187876 | 0.028521 | 0.198563 | -0.136354 | 0.110785 | | 30.13889 | 0.081277 | 0.13928 |
| 30.056 | 1.434554 | -0.34162 | 0.145001 | 0.130975 | 0.685907 | 0.468938 | -0.632032 | 1.012598 | 0.492956 | 0.77336 | 0.602245 | 0.694663 | 0.873941 | 1.034237 | 0.75713 | -0.651299 | 0.767938 | -0.426738 | 0.147263 | 0.147149 | 0.627023 | 0.149358 | 0.08389 | 0.364904 | | 30.15278 | 0.873941 | 0.434891 |
| 30.069 | 2.641916 | 0.02324 | 0.258151 | 0.955049 | 1.053833 | 1.658641 | 1.242096 | 1.302018 | 1.209653 | 0.838136 | 1.55577 | 0.532852 | 1.744852 | 0.994304 | 1.105202 | -0.835534 | 0.573022 | -0.02214 | 0.107044 | 0.118742 | 0.955175 | 0.115159 | 0.13698 | 0.47103 | | 30.16667 | 1.744852 | 0.563306 |
| 30.083 | 3.910379 | 0.363738 | -0.085681 | 1.303936 | 1.191791 | 1.838312 | 2.567679 | 1.118355 | 2.004228 | 1.088167 | 2.610442 | 0.14663 | 2.109054 | 0.600577 | 1.581887 | -0.746999 | 0.549518 | 0.185573 | 0.086009 | 0.09665 | 0.926628 | 0.092937 | 0.187249 | 0.432947 | | 30.18056 | 2.109054 | 0.496232 |
| 30.097 | 2.347344 | 0.606672 | -0.993668 | 0.494861 | 0.73018 | 1.122783 | 3.154646 | 0.881307 | 2.240645 | 1.745759 | 2.513398 | -0.239995 | 1.458282 | 0.397832 | 2.036724 | -0.578191 | 0.899863 | 0.074265 | 0.095411 | 0.10511 | 0.795132 | 0.098318 | 0.309623 | 0.384615 | | 30.19444 | 1.458282 | 0.444664 |
| 30.111 | 0.513535 | 0.766218 | -2.117751 | -0.266915 | 0.18588 | 0.285714 | 2.574422 | 0.69263 | 1.491289 | 0.731756 | 1.644181 | -0.677766 | 0.451644 | 0.718754 | 1.981798 | -0.403707 | 1.346753 | -0.268031 | 0.131987 | 0.144546 | 0.830096 | 0.132354 | 0.543998 | 0.508933 | | 30.20833 | 0.451644 | 0.56101 |
| 30.125 | -0.610701 | 0.804342 | -1.489721 | -0.481449 | 0.141482 | -0.211763 | 0.735682 | 0.582223 | 0.389048 | -0.08004 | 0.133384 | -1.040196 | 0.103875 | 1.261741 | 1.372394 | -0.232065 | 1.259089 | -0.526482 | 0.170173 | 0.179582 | 1.067398 | 0.169186 | 0.935551 | 0.724866 | | 30.22222 | 0.103875 | 0.765098 |
| 30.139 | 0.867864 | 0.634199 | -0.107768 | 0.175985 | 0.248874 | -0.29911 | -0.537575 | 0.345162 | -0.265242 | -0.485158 | -0.329594 | -0.887378 | 0.588799 | 1.360717 | 0.55214 | 0.0744 | 0.738421 | -0.618866 | 0.091414 | 0.105685 | 1.416334 | 0.082961 | 1.208978 | 0.780668 | | 30.23611 | 0.588799 | 0.838612 |
| 30.153 | 2.175818 | 0.236629 | 0.732737 | 1.571379 | 0.296567 | -0.191573 | -0.955854 | 0.082995 | -0.139824 | 0.133837 | -0.820168 | -0.088709 | 1.262618 | 1.090827 | -0.130895 | 0.288049 | 0.44546 | -0.771842 | -0.078479 | -0.048314 | 1.27413 | -0.095983 | 1.002801 | 0.595651 | | 30.25 | 1.262618 | 0.627788 |
| 30.167 | 1.003193 | 0.034623 | 0.692948 | 1.852202 | 0.534183 | -0.353485 | -1.363062 | 0.150651 | 0.428615 | 0.449778 | -1.061085 | 1.039294 | 1.443287 | 0.771749 | -0.146459 | 0.293602 | 0.826948 | -0.783278 | -0.060182 | -0.027086 | 1.058594 | -0.061912 | 0.867189 | 0.407447 | | 30.26389 | 1.443287 | 0.373941 |
| 30.181 | -0.612074 | 0.166414 | 0.501271 | 0.5355 | 0.627337 | -0.190371 | -1.100725 | 0.48794 | 0.524769 | 0.769455 | -0.597921 | 1.865573 | 1.132432 | 0.928157 | 0.355333 | 0.320244 | 1.409404 | -0.478987 | 0.148445 | 0.174266 | 1.164481 | 0.169775 | 1.050912 | 0.47599 | | 30.27778 | 1.132432 | 0.442877 |
| 30.194 | -0.891653 | 0.160106 | 0.772275 | -0.441762 | 0.481858 | 0.434875 | -0.529074 | 0.822409 | -0.054212 | 1.34554 | -0.105114 | 1.359245 | 0.6175 | 1.275383 | 0.987829 | 0.330047 | 1.372432 | -0.170456 | 0.273809 | 0.292394 | 1.301789 | 0.295033 | 1.013557 | 0.679982 | | 30.29167 | 0.6175 | 0.640829 |
| 30.208 | 0.31187 | -0.238108 | 1.366225 | -0.843685 | -0.483725 | 0.91406 | 0.263542 | 0.8704 | 0.377877 | 1.461523 | 0.93934 | 0.357732 | 0.547853 | 1.072411 | 1.649003 | 0.291639 | 1.150896 | -0.099866 | 0.212194 | 0.220922 | 1.184509 | 0.221192 | 0.550988 | 0.688818 | | 30.30556 | 0.547853 | 0.645472 |
| 30.222 | 1.727612 | -0.467873 | 1.978116 | 0.181478 | -0.883283 | 0.82517 | 1.116667 | 0.451015 | 0.803595 | 0.883703 | 1.591435 | 0.071994 | 1.444848 | 0.381826 | 1.631749 | 0.224327 | 0.982117 | -0.101495 | 0.063692 | 0.073443 | 0.701171 | 0.073353 | -0.007895 | 0.353523 | | 30.31944 | 1.444848 | 0.3695 |
| 30.236 | 1.401114 | -0.318198 | 1.298391 | 1.126646 | -0.471597 | 0.562201 | 1.475645 | 0.225753 | 1.30083 | -0.011489 | 1.182351 | 0.193657 | 1.899279 | -0.057538 | 1.139873 | 0.246065 | 1.139633 | 0.073001 | -0.003668 | 0.023633 | 0.324337 | 0.013996 | -0.609441 | 0.027307 | | 30.33333 | 1.899279 | 0.094072 |
| 30.25 | 0.371138 | -0.121702 | 0.023565 | 0.756759 | 0.622386 | 0.489102 | 1.500995 | 0.775898 | 1.353413 | -0.603209 | 0.388699 | 0.587921 | 1.187639 | -0.003634 | 0.884734 | 0.425768 | 1.339075 | 0.302839 | 0.020172 | 0.057492 | 0.402714 | 0.037796 | -0.751222 | 0.045433 | | 30.34722 | 1.187639 | 0.154446 |
| 30.264 | -0.253475 | -0.050935 | -1.339468 | -0.007852 | 1.058592 | 0.794492 | 1.348807 | 1.724255 | 1.324601 | -0.336416 | -0.070969 | 1.047928 | 0.663973 | 0.700307 | 0.85714 | 0.579954 | 1.091345 | 0.367086 | 0.009126 | 0.038658 | 0.630005 | 0.024384 | -0.418842 | 0.314349 | | 30.36111 | 0.663973 | 0.421329 |
| 30.278 | 0.633488 | -0.027445 | -0.957034 | -0.204066 | 0.930289 | 1.080748 | 0.655278 | 2.340391 | 0.904564 | 0.528964 | 0.197926 | 1.280963 | 0.829216 | 1.374836 | 0.808764 | 0.704918 | 0.806961 | 0.33735 | -0.075642 | -0.048816 | 0.875204 | -0.063779 | 0.184635 | 0.606151 | | 30.375 | 0.829216 | 0.66619 |
| 30.292 | 1.963532 | 0.124541 | 0.037194 | -0.228929 | 0.652893 | 1.168714 | 0.260884 | 2.140721 | 0.646588 | 0.956567 | 1.131392 | 1.43382 | 1.380636 | 1.525645 | 0.784841 | 0.866895 | 0.683463 | 0.363493 | -0.150906 | -0.119562 | 1.099735 | -0.153146 | 0.980627 | 0.75469 | | 30.38889 | 1.380636 | 0.75842 |
| 30.306 | 2.216261 | 0.412488 | 1.419538 | -0.183147 | 0.397835 | 1.222964 | 0.6064 | 1.348013 | 0.806493 | 1.37747 | 1.910376 | 1.68769 | 2.294272 | 1.30573 | 0.833304 | 1.034248 | 0.794619 | 0.539359 | -0.117452 | -0.10395 | 1.289617 | -0.12616 | 1.632191 | 0.806772 | | 30.40278 | 2.294272 | 0.859854 |
| 30.319 | 1.495893 | 0.624835 | 2.458911 | 0.106109 | 0.950381 | 1.207604 | 1.093914 | 1.002644 | 1.304234 | 1.707223 | 1.722987 | 1.992873 | 2.685786 | 1.387999 | 0.860089 | 1.078599 | 0.978543 | 0.824293 | -0.080155 | -0.092458 | 1.374765 | -0.080981 | 2.102348 | 0.871204 | | 30.41667 | 2.685786 | 0.94137 |
| 30.333 | 1.27856 | 0.619392 | 2.258099 | 0.236068 | 1.193772 | 1.115084 | 1.384374 | 1.292342 | 1.624561 | 1.499795 | 1.268318 | 2.13912 | 2.150589 | 1.884031 | 0.710796 | 0.907845 | 1.020994 | 1.066052 | -0.16313 | -0.16989 | 1.412531 | -0.165086 | 2.096197 | 0.915486 | | 30.43056 | 2.150589 | 1.064776 |
| 30.347 | 1.374285 | 0.602612 | 1.643078 | -0.090937 | 0.732103 | 1.085324 | 1.198621 | 1.853933 | 1.729332 | 1.088661 | 1.140963 | 2.058484 | 1.416835 | 2.713352 | 1.036007 | 0.810056 | 1.003832 | 1.227495 | -0.237585 | -0.222718 | 1.672105 | -0.255654 | 1.712701 | 1.029287 | | 30.44444 | 1.416835 | 1.208513 |
| 30.361 | 1.762594 | 0.59728 | 0.90947 | -0.414634 | 0.286371 | 1.105268 | 0.888165 | 2.640787 | 1.303665 | 0.876957 | 1.563503 | 1.725212 | 0.953981 | 3.245733 | 1.80626 | 0.748135 | 0.854569 | 1.245235 | -0.325384 | -0.283735 | 2.047642 | -0.373066 | 1.485892 | 1.024919 | | 30.45833 | 0.953981 | 1.197258 |
| 30.375 | 2.292171 | 0.641422 | 0.258867 | -1.181749 | 0.363264 | 1.397488 | 0.767004 | 4.210899 | 0.561402 | 0.906352 | 2.29943 | 1.613409 | 0.941566 | 2.915 | 2.168986 | 0.748033 | 0.554624 | 1.173397 | -0.579816 | -0.429325 | 1.678786 | -0.557661 | 1.244968 | 0.786232 | | 30.47222 | 0.941566 | 0.907604 |
| 30.389 | 2.583603 | 0.839829 | 0.261895 | -1.090644 | 1.119205 | 1.93829 | 1.069974 | 4.687418 | 0.204627 | 0.794212 | 2.810207 | 1.63277 | 1.864508 | 2.057803 | 1.906798 | 0.880584 | 0.387821 | 1.138974 | -0.585431 | -0.432462 | 1.094607 | -0.491966 | 1.050533 | 0.744053 | | 30.48611 | 1.864508 | 0.765985 |
| 30.403 | 2.143134 | 1.153022 | 0.890474 | -0.338652 | 1.680038 | 2.709147 | 1.648927 | 5.524789 | 0.677334 | 1.386366 | 3.187161 | 1.746394 | 2.810166 | 1.801283 | 1.117104 | 1.213612 | 0.38209 | 1.1895 | -0.392493 | -0.324207 | 0.743757 | -0.354415 | 0.57104 | 0.809868 | | 30.5 | 2.810166 | 0.724217 |
| 30.417 | 1.64555 | 1.286251 | 1.367351 | 0.342746 | 0.844433 | 2.819737 | 2.173872 | 3.467159 | 1.594275 | 2.096486 | 2.966985 | 1.956081 | 3.163658 | 1.595918 | 0.92018 | 1.24479 | 0.510946 | 1.163926 | -0.268445 | -0.234965 | 0.704815 | -0.260698 | -0.524393 | 0.67491 | | 30.51389 | 3.163658 | 0.608551 |
| 30.431 | 1.634384 | 1.313154 | 1.084311 | 0.605776 | 0.949089 | 2.042292 | 2.432811 | 2.053335 | 2.066601 | 2.236295 | 2.143494 | 2.248435 | 2.596781 | 1.306487 | 1.316929 | 1.179723 | 0.493348 | 0.935774 | -0.22582 | -0.22658 | 0.859454 | -0.217822 | -0.592674 | 0.600171 | | 30.52778 | 2.596781 | 0.658236 |
| 30.444 | 1.320465 | 1.015438 | -0.400241 | 0.184045 | 2.130526 | 1.638593 | 2.150178 | 0.921328 | 1.487991 | 2.027449 | 1.972086 | 2.432918 | 1.878404 | 1.464313 | 1.390777 | 0.949397 | 0.190587 | 0.552863 | -0.475942 | -0.520149 | 0.854143 | -0.417795 | 0.727998 | 0.69545 | | 30.54167 | 1.878404 | 0.549053 |
| 30.458 | 0.806615 | 0.870502 | -1.26603 | -0.238382 | 1.889163 | 1.335888 | 1.498355 | 0.633027 | 0.952748 | 1.793954 | 2.371295 | 2.273141 | 1.407383 | 1.266384 | 0.057454 | -0.060974 | -0.734543 | -0.240505 | -1.138831 | -1.217338 | 0.474857 | -0.999131 | 1.301635 | 0.252574 | | 30.55556 | 1.407383 | -0.137478 |
| 30.472 | 0.247433 | 0.671515 | -0.776395 | 0.007444 | 1.269936 | 1.012919 | 0.698924 | 0.214576 | 0.795794 | 1.455832 | 2.002611 | 1.215296 | 1.052668 | 0.639 | -0.330877 | -0.863687 | -1.534819 | -0.919584 | -1.65465 | -1.673616 | -0.046999 | -1.499038 | 0.0472 | -0.501719 | | 30.56944 | 1.052668 | -0.722767 |
| 30.486 | 1.261689 | 0.52908 | 0.18282 | 0.598205 | 1.440469 | 0.629978 | 0.36471 | -0.242476 | 0.796097 | 1.49831 | 1.369288 | 0.44303 | 1.917974 | 0.624136 | -0.784035 | -0.889752 | -1.009058 | -0.833756 | -1.334475 | -1.292792 | -0.041349 | -1.293645 | -0.538746 | -0.673468 | | 30.58333 | 1.917974 | -0.687544 |
| 30.5 | 1.542357 | 0.775629 | 1.086053 | 0.659354 | 1.554143 | 0.194276 | 0.318337 | -0.444796 | 1.068687 | 1.734469 | 1.452749 | 0.605837 | 2.315778 | 0.881604 | -1.16912 | -0.652616 | -0.39174 | -0.480648 | -0.656374 | -0.619456 | 0.756235 | -0.672251 | -0.066354 | -0.318735 | | 30.59722 | 2.315778 | -0.342344 |
| 30.514 | 1.277049 | 0.966245 | 1.305533 | -0.006315 | 0.646682 | -0.126689 | 0.310929 | -0.449381 | 0.254231 | 0.807467 | 0.606592 | 0.854274 | 1.683381 | 0.632857 | -0.400769 | -0.66407 | -0.256713 | -0.183935 | -0.161064 | -0.154485 | 0.928124 | -0.1706 | 0.385787 | -0.07838 | | 30.61111 | 1.683381 | -0.210631 |
| 30.528 | 0.516281 | 0.743097 | 0.773166 | -0.284256 | -0.63478 | -0.244051 | 0.342585 | -0.035142 | -0.164961 | -0.087045 | -0.106144 | 0.856705 | 0.372673 | 1.549785 | 0.301188 | -0.557557 | -0.174231 | 0.047475 | 0.046396 | 0.026323 | 0.535497 | 0.047542 | 0.477582 | -0.016361 | | 30.625 | 0.372673 | -0.10606 |
| 30.542 | 0.776135 | 0.494254 | 0.312546 | -0.344059 | -0.936367 | 0.132911 | 0.268564 | 0.317157 | -0.693421 | -0.454348 | -1.149243 | 0.997401 | -0.100043 | 1.202002 | 0.961327 | -0.496304 | -0.048667 | 0.100878 | -0.006201 | -0.025881 | -0.290514 | -0.001471 | 0.108205 | -0.134133 | | 30.63889 | -0.100043 | -0.228248 |
| 30.556 | 0.282376 | 0.306986 | -0.455878 | -0.041216 | -0.330748 | 1.188832 | 0.210235 | 0.100373 | -0.362344 | -0.461153 | -2.320006 | 0.733464 | 0.296522 | -0.783138 | 1.143672 | -0.484009 | -0.057887 | 0.004475 | -0.123499 | -0.11493 | -1.527361 | -0.118342 | -0.191014 | -0.374474 | | 30.65278 | 0.296522 | -0.452633 |
| 30.569 | -1.230726 | 0.325567 | -0.548957 | 0.147562 | -0.004853 | 0.91171 | 0.264744 | 0.164748 | 0.497204 | -0.746489 | -1.520315 | 0.254369 | 0.704038 | 0.412581 | 0.262777 | -0.38864 | -0.348095 | -0.047039 | -0.123127 | -0.095435 | -1.726072 | -0.118101 | -0.263128 | -0.460153 | | 30.66667 | 0.704038 | -0.51216 |
| 30.583 | -2.207352 | 0.252166 | -0.338822 | -0.063015 | 0.122405 | 0.534397 | 0.378042 | 0.022528 | 0.946004 | -1.795506 | -0.306829 | -0.135276 | 0.190076 | -0.65248 | 0.022849 | -0.268671 | -0.71423 | -0.014955 | -0.043269 | -0.025351 | -0.046835 | -0.04146 | -0.222383 | -0.234958 | | 30.68056 | 0.190076 | -0.33713 |
| 30.597 | -2.451862 | 0.088826 | -0.157383 | -0.052272 | 0.867795 | 0.252624 | 0.470684 | -0.107702 | 0.448637 | -0.775211 | 1.198486 | -0.114432 | 0.269324 | -3.384025 | -0.272784 | -0.419042 | -0.927729 | 0.018334 | 0.010493 | 0.009135 | 1.846342 | 0.01062 | -0.57996 | -0.096186 | | 30.69444 | 0.269324 | -0.392635 |
| 30.611 | -1.448418 | -0.014564 | 0.370907 | 0.168056 | 0.307046 | -0.015329 | 0.253234 | -0.116837 | -0.076263 | 0.594002 | 2.287917 | -0.022198 | -0.023838 | -1.780676 | -0.331406 | -0.938719 | -0.835264 | -0.006685 | -0.003023 | -0.007936 | 1.971435 | 0.004755 | -1.544976 | -0.317676 | | 30.70833 | -0.023838 | -0.515726 |
| 30.625 | 0.071104 | -0.015957 | 1.052963 | 0.511487 | 1.133523 | -0.183178 | 0.107422 | 0.204187 | -0.112412 | 0.417693 | 1.627997 | -0.638745 | 0.381247 | 0.549829 | -0.490533 | -1.304734 | -0.328914 | -0.027496 | -0.014951 | -0.007202 | 0.702367 | -0.000821 | -0.791688 | -0.369362 | | 30.72222 | 0.381247 | -0.428586 |
| 30.639 | 1.269968 | 0.029937 | 0.330807 | 0.778904 | 2.11228 | -0.19914 | 0.069804 | 0.63354 | -0.77401 | 1.117742 | 0.726809 | -0.180164 | 1.250061 | 0.059534 | 0.206342 | -1.010974 | 0.281067 | 0.012514 | 0.022687 | 0.029803 | -0.179208 | 0.028417 | -0.057575 | -0.264987 | | 30.73611 | 1.250061 | -0.102462 |
| 30.653 | 2.043609 | 0.036432 | 0.200271 | 0.811141 | 1.561908 | -0.086736 | -0.123513 | 0.738254 | -0.439221 | 0.162617 | 0.270473 | 0.114512 | 2.032765 | -1.993682 | 0.223996 | -0.53576 | 0.754851 | 0.068479 | 0.061736 | 0.056703 | -1.164996 | 0.056226 | 0.375817 | -0.237317 | | 30.75 | 2.032765 | 0.005398 |
| 30.667 | 2.748458 | -0.00839 | 0.53337 | 0.627364 | 0.223206 | 0.02664 | -0.278514 | 0.131855 | -0.010893 | 0.435067 | -0.164531 | -0.064054 | 2.074975 | -1.369825 | -0.029238 | -0.314553 | 1.04381 | 0.106494 | 0.079643 | 0.070882 | -2.734802 | 0.075562 | 0.582782 | -0.408451 | | 30.76389 | 2.074975 | -0.139142 |
| 30.681 | 1.729883 | -0.080726 | 1.451694 | 0.39138 | -0.102566 | 0.113175 | -0.366172 | -0.496089 | 0.574644 | 0.51934 | -0.881847 | -0.557864 | 0.740492 | 0.011861 | -0.344639 | -0.285183 | 0.853421 | 0.14939 | 0.095273 | 0.099264 | -3.105396 | 0.103765 | 0.450664 | -0.829933 | | 30.77778 | 0.740492 | -0.69965 |
| 30.694 | 0.432225 | -0.184289 | 1.088934 | -0.135877 | 0.588345 | 0.095038 | -0.247241 | -0.562053 | 0.659944 | -1.199203 | -1.466825 | -1.107574 | -1.146641 | -0.227327 | -0.588367 | -0.3241 | 0.326451 | 0.185739 | 0.114589 | 0.12716 | -1.475388 | 0.13155 | 0.207112 | -1.019495 | | 30.79167 | -1.146641 | -1.128681 |
| 30.708 | -0.665763 | -0.30571 | 0.058875 | -0.214442 | 1.073461 | 0.026673 | 0.08646 | -0.439344 | 1.088801 | -0.719331 | -1.943845 | -1.454212 | -2.526973 | -1.590485 | -0.601571 | -0.404272 | -0.247841 | 0.172249 | 0.102707 | 0.111507 | 0.377043 | 0.118595 | -0.159009 | -0.742305 | | 30.80556 | -2.526973 | -0.908149 |
| 30.722 | -2.042244 | -0.397982 | -0.663296 | -0.1159 | 1.509792 | -0.031692 | 0.459648 | -0.333931 | -0.018714 | -0.171217 | -2.170092 | -1.438772 | -2.79243 | -0.756967 | -0.773548 | -0.442942 | -0.777026 | 0.135319 | 0.066526 | 0.07212 | 1.899386 | 0.074179 | -0.662931 | -0.205977 | | 30.81944 | -2.79243 | -0.572073 |
| 30.736 | -1.835871 | -0.423081 | -1.394624 | -0.111107 | 1.726336 | -0.126703 | 0.267845 | -0.176356 | -0.648481 | -0.331 | -1.658006 | -1.207946 | -2.17163 | -0.353227 | -0.448269 | -0.427686 | -0.967076 | 0.138756 | 0.064644 | 0.068694 | 2.610312 | 0.067769 | -0.981877 | 0.140514 | | 30.83333 | -2.17163 | -0.203109 |
| 30.75 | -0.968382 | -0.367431 | -3.174472 | -0.047951 | 1.01227 | -0.352634 | -0.375434 | -0.415331 | -0.529684 | -0.685462 | -0.929873 | -0.848331 | -1.745615 | 0.323229 | -0.221854 | -0.312295 | -0.813838 | 0.191775 | 0.114117 | 0.114456 | 1.66312 | 0.12182 | -1.300744 | 0.410803 | | 30.84722 | -1.745615 | 0.246832 |
| 30.764 | -0.374401 | -0.393971 | -2.597464 | 0.046048 | -0.230281 | -0.637617 | -0.491309 | 0.160225 | -0.534886 | -0.662626 | -0.224262 | -0.404577 | -1.466924 | 0.503475 | 0.070899 | -0.068919 | -0.67571 | 0.242312 | 0.186884 | 0.175055 | 0.336024 | 0.200391 | -1.327784 | 0.339462 | | 30.86111 | -1.466924 | 0.440627 |
| 30.778 | -0.010061 | -0.456655 | -1.462732 | 0.085418 | -1.442176 | -1.037099 | 0.191889 | 0.123332 | -0.823832 | -0.499778 | 0.043223 | 0.371286 | -0.96503 | -0.616652 | -0.236327 | 0.049995 | -0.715159 | 0.223734 | 0.246985 | 0.225081 | -0.972729 | 0.262908 | -1.23107 | -0.038102 | | 30.875 | -0.96503 | 0.218488 |
| 30.792 | -0.183909 | -0.543541 | -0.967558 | 0.052896 | -1.439692 | -1.282762 | 0.699388 | -0.19277 | -0.651708 | -0.448871 | -0.482975 | 1.108591 | -0.790698 | -1.100348 | -0.927738 | 0.19311 | -0.855134 | 0.158061 | 0.294638 | 0.272219 | -1.841296 | 0.312316 | -1.177055 | -0.532145 | | 30.88889 | -0.790698 | -0.297291 |
| 30.806 | -0.50608 | -0.721353 | 0.833773 | 0.11763 | -1.597579 | -1.29202 | 0.112276 | -0.61291 | -0.109543 | -0.591042 | -0.403705 | 0.673348 | -0.822134 | -1.397522 | -1.589613 | 0.210387 | -1.009184 | 0.097017 | 0.326267 | 0.303318 | -2.7078 | 0.342652 | -1.127864 | -0.774887 | | 30.90278 | -0.822134 | -0.74364 |
| 30.819 | -0.885506 | -0.968266 | 0.974557 | 0.043198 | -0.963184 | -0.973313 | -0.305515 | -1.069514 | 0.013335 | -0.802926 | -0.879697 | -0.31851 | -0.372931 | -1.953195 | -2.184637 | -0.050458 | -1.346785 | 0.045092 | 0.318726 | 0.305871 | -1.527638 | 0.331324 | -1.01072 | -0.648318 | | 30.91667 | -0.372931 | -0.754824 |
| 30.833 | -1.770707 | -1.025529 | 0.811267 | -0.069661 | -0.815787 | -0.965821 | -0.706614 | -1.413887 | -0.373393 | -0.944028 | -1.961958 | -0.852586 | -0.992448 | -2.026243 | -2.21614 | -0.381593 | -1.632219 | -0.021027 | 0.28571 | 0.289167 | 0.036179 | 0.299935 | -1.010656 | -0.327644 | | 30.93056 | -0.992448 | -0.476718 |
| 30.847 | -2.998784 | -1.086801 | 2.011025 | -0.074412 | -0.335643 | -1.161416 | -0.652525 | -2.127971 | -0.584098 | -1.150171 | -2.047697 | -1.532608 | -1.007083 | -1.814902 | -1.653801 | -0.562303 | -1.568936 | -0.108934 | 0.25431 | 0.260865 | 0.969036 | 0.275279 | -0.945711 | -0.09472 | | 30.94444 | -1.007083 | -0.234603 |
| 30.861 | -4.238359 | -1.15926 | 2.77642 | -0.152369 | -0.25045 | -1.530449 | -0.024006 | -3.507511 | -0.542202 | -1.873683 | -2.300444 | -2.710901 | -0.712149 | -1.694215 | -1.13816 | -0.624176 | -1.23067 | -0.219979 | 0.220555 | 0.21936 | 0.876084 | 0.239126 | -0.720511 | -0.144403 | | 30.95833 | -0.712149 | -0.15609 |
| 30.875 | -4.238004 | -1.238428 | 2.390541 | -0.087094 | -0.224665 | -1.677454 | -0.725884 | -4.248403 | -1.216449 | -2.175407 | -3.546545 | -4.104078 | -0.817309 | -1.616846 | -0.668624 | -0.706552 | -1.077343 | -0.378638 | 0.171031 | 0.167668 | 0.249944 | 0.177541 | -0.536771 | -0.326038 | | 30.97222 | -0.817309 | -0.32974 |
| 30.889 | -3.391503 | -1.231081 | 0.893605 | 0.030395 | -0.777602 | -1.499673 | -1.532373 | -2.733509 | -1.318253 | -2.287476 | -3.456218 | -4.042461 | -0.525061 | -1.572862 | -0.344609 | -0.811354 | -0.768086 | -0.518444 | 0.10602 | 0.107151 | -0.290383 | 0.108153 | -0.423428 | -0.422615 | | 30.98611 | -0.525061 | -0.458215 |
| 30.903 | -2.466981 | -1.160053 | -1.235734 | 0.019264 | -1.745107 | -1.34033 | -2.069031 | -1.591486 | -1.387541 | -2.235278 | -2.573706 | -3.612251 | 0.051325 | -1.464308 | -0.169006 | -0.818644 | -0.340605 | -0.603404 | 0.063286 | 0.061228 | -0.654017 | 0.065582 | -0.443874 | -0.41662 | | 31 | 0.051325 | -0.495703 |
| 30.917 | -1.301464 | -1.063177 | -1.327839 | 0.02644 | -1.392005 | -1.073283 | -1.881295 | -1.568641 | -1.278447 | -1.638635 | -1.328638 | -3.049164 | 0.729752 | -1.172394 | -0.08015 | -0.525752 | -0.037718 | -0.558864 | 0.163212 | 0.131984 | -0.748025 | 0.147245 | -0.366322 | -0.309717 | | 31.01389 | 0.729752 | -0.414002 |
| 30.931 | -0.348628 | -0.803471 | 0.374083 | 0.262528 | -0.883071 | -0.634487 | -1.489896 | -1.938288 | -1.021165 | -0.742833 | -0.216139 | -2.214926 | 0.429959 | -0.808168 | 0.025001 | -0.263515 | 0.123163 | -0.360655 | 0.40282 | 0.320547 | -0.660905 | 0.357747 | -0.145203 | -0.145716 | | 31.02778 | 0.429959 | -0.240634 |
| 30.944 | -0.374514 | -0.7619 | 1.217011 | 0.352344 | -1.106466 | -0.5068 | -2.253574 | -1.907429 | -1.045359 | -0.500617 | 0.135471 | -1.319046 | -1.453493 | -0.772979 | -0.042847 | -0.379765 | 0.007758 | -0.304408 | 0.502734 | 0.364185 | -0.620489 | 0.445899 | 0.05235 | -0.176657 | | 31.04167 | -1.453493 | -0.197294 |
| 30.958 | -1.525245 | -0.815739 | 1.762 | 0.054822 | -2.800937 | -1.231159 | -2.036553 | -2.777873 | -1.812398 | -1.110622 | -0.496563 | -0.216698 | -4.001652 | -0.897303 | -0.421128 | -0.688556 | -0.359245 | -0.469577 | 0.312087 | 0.232158 | -0.840947 | 0.277293 | -0.0169 | -0.523898 | | 31.05556 | -4.001652 | -0.448862 |
| 30.972 | -2.620065 | -0.986071 | 0.968418 | -0.531689 | -2.495711 | -3.107096 | -2.21442 | -4.773818 | -2.860442 | -2.303313 | -1.576529 | 0.119305 | -5.999712 | -1.313457 | -0.791395 | -0.944465 | -0.718791 | -0.702223 | 0.23895 | 0.240413 | -1.34894 | 0.1972 | -0.372597 | -1.170032 | | 31.06944 | -5.999712 | -1.085925 |
| 30.986 | -2.348374 | -1.436779 | -0.273877 | -1.723606 | -1.969402 | -4.954494 | -1.565376 | -6.428463 | -3.622719 | -2.572744 | -2.448391 | 0.486767 | -6.054165 | -2.99149 | -0.908814 | -1.286478 | -1.071317 | -1.274893 | 0.306359 | 0.331428 | -2.599285 | 0.273682 | -0.860116 | -1.785767 | | 31.08333 | -6.054165 | -1.728078 |
| 31 | -0.832198 | -2.344862 | -0.867636 | -1.759461 | -1.511704 | -3.372222 | -0.294377 | -5.74373 | -3.753185 | -2.070088 | -1.879239 | 0.701147 | -4.847575 | -2.158581 | -0.787196 | -1.310734 | -1.006919 | -2.021746 | 0.205811 | 0.239403 | -2.531027 | 0.19689 | -1.139696 | -1.796467 | | 31.09722 | -4.847575 | -1.802936 |
| 31.014 | -0.324939 | -2.427969 | -1.991439 | -1.212507 | -0.939118 | -2.042421 | -0.586641 | -3.352588 | -2.642556 | -1.050217 | -0.986003 | -0.615385 | -2.922408 | -0.226526 | -0.361294 | -0.741187 | -0.6141 | -1.796757 | -0.008741 | 0.016177 | -1.465854 | -0.020616 | -0.906712 | -0.941765 | | 31.11111 | -2.922408 | -1.066333 |
| 31.028 | -0.380891 | -2.019937 | -2.20433 | -0.490945 | -0.69304 | -0.713662 | -1.520849 | -1.619607 | -1.259131 | -0.201228 | -0.125921 | -0.772698 | -1.180921 | 1.331137 | -0.123531 | -0.223216 | -0.176541 | -1.166992 | -0.136473 | -0.130174 | -0.404831 | -0.152631 | -0.506028 | -0.062526 | | 31.125 | -1.180921 | -0.172724 |
| 31.042 | -0.174516 | -1.614044 | -1.269575 | -0.214474 | -0.235231 | 0.316023 | -1.634346 | -0.663615 | -0.374256 | 0.381508 | 0.556625 | -0.482752 | -0.310984 | 1.856091 | -0.054189 | 0.120399 | 0.262526 | -0.300149 | -0.100048 | -0.106303 | 0.377635 | -0.10413 | -0.166354 | 0.49471 | | 31.13889 | -0.310984 | 0.491997 |
| 31.056 | 0.197833 | -1.272382 | -0.262587 | 0.034433 | 0.443281 | 1.199055 | -0.740269 | 0.040656 | 0.11944 | 0.743727 | 0.723694 | -0.330752 | 0.243831 | 1.536424 | -0.012873 | 0.464946 | 0.479692 | 0.52463 | 0.003285 | -0.006371 | 0.650365 | 0.006313 | -0.094156 | 0.776323 | | 31.15278 | 0.243831 | 0.84477 |
| 31.069 | 0.811348 | -0.985922 | 0.211393 | 0.73621 | 0.902232 | 1.902559 | 0.528855 | 0.781287 | 0.730506 | 0.917382 | 0.731677 | -0.256746 | 0.743955 | 0.994666 | 0.166141 | 0.767212 | 0.478667 | 1.000456 | 0.061185 | 0.057298 | 0.838281 | 0.06512 | -0.035693 | 0.837782 | | 31.16667 | 0.743955 | 0.903654 |
| 31.083 | 1.624665 | -0.676277 | 0.414259 | 1.704038 | 0.963978 | 2.291064 | 1.586176 | 1.365443 | 1.42134 | 0.908688 | 1.275741 | -0.549216 | 1.104334 | 0.63878 | 0.617027 | 0.883564 | 0.54547 | 0.893046 | 0.153859 | 0.154119 | 1.036601 | 0.16659 | 0.467428 | 0.835963 | | 31.18056 | 1.104334 | 0.911088 |
| 31.097 | 2.035887 | -0.393074 | 0.650886 | 1.727049 | 1.451814 | 2.118256 | 2.539912 | 1.859531 | 1.861155 | 0.875391 | 2.747544 | -0.929762 | 1.527174 | 0.426001 | 1.225844 | 0.823161 | 0.875336 | 0.531712 | 0.35034 | 0.341189 | 1.232929 | 0.371167 | 1.146052 | 0.817078 | | 31.19444 | 1.527174 | 0.906955 |
| 31.111 | 1.365384 | -0.005265 | 0.38729 | 0.375764 | 2.118605 | 1.276013 | 3.47387 | 1.897014 | 1.721881 | 1.092677 | 2.733651 | -1.007172 | 1.689672 | 0.115181 | 1.473585 | 0.578929 | 1.321314 | 0.107462 | 0.498151 | 0.479029 | 1.169761 | 0.513132 | 1.523327 | 0.600296 | | 31.20833 | 1.689672 | 0.673485 |
| 31.125 | 0.512111 | 0.699036 | -0.171519 | -0.794079 | 1.260081 | 0.579014 | 1.403685 | 1.314863 | 0.985922 | 0.875628 | 2.58789 | -0.618188 | 1.352584 | -0.295907 | 1.099784 | 0.192131 | 1.467835 | -0.255404 | 0.470337 | 0.448067 | 0.851875 | 0.470356 | 1.785451 | 0.249643 | | 31.22222 | 1.352584 | 0.324572 |
| 31.139 | 0.506709 | 1.536052 | -0.424973 | -1.133893 | 0.884852 | 0.027096 | 0.132643 | 1.084071 | 0.146979 | 0.47488 | 0.748731 | -0.043072 | 1.286327 | -0.507025 | 0.505258 | -0.258791 | 1.410217 | -0.364105 | 0.322117 | 0.296622 | 0.636466 | 0.308242 | 1.889194 | 0.083619 | | 31.23611 | 1.286327 | 0.134155 |
| 31.153 | 1.393105 | 2.225531 | 0.458142 | 0.087462 | 0.842124 | -0.268468 | -0.611177 | 1.211841 | -0.002786 | 0.020894 | -0.885386 | 0.2557 | 1.637587 | -0.276253 | -0.187622 | -0.651405 | 1.259144 | -0.335015 | 0.122034 | 0.102677 | 0.816206 | 0.10229 | 1.340393 | -0.00592 | | 31.25 | 1.637587 | -0.021586 |
| 31.167 | 1.756412 | 2.776461 | 2.575279 | 1.607087 | 0.863451 | -0.249576 | -1.168581 | 1.480218 | 0.320782 | -0.00859 | -0.818359 | 0.202111 | 1.89836 | 0.040166 | -0.472053 | -0.857583 | 1.08144 | -0.331109 | -0.062871 | -0.059747 | 1.005661 | -0.087599 | -0.048091 | -0.093749 | | 31.26389 | 1.89836 | -0.19986 |
| 31.181 | 0.762593 | 3.35519 | 3.342871 | 1.014827 | 1.298619 | 0.139393 | -1.405202 | 1.421063 | 0.715612 | 0.451674 | -0.719185 | 0.116157 | 1.503624 | 0.408956 | 0.06356 | -0.729071 | 1.035974 | -0.411719 | -0.176839 | -0.138493 | 1.125324 | -0.210904 | 0.007849 | 0.008396 | | 31.27778 | 1.503624 | -0.069388 |
| 31.194 | -0.511118 | 2.719521 | 2.650676 | 0.108159 | 1.079365 | 0.950241 | -0.441173 | 0.736503 | 0.978617 | 0.78012 | 0.537048 | 0.049821 | 1.110589 | 0.882837 | 0.818842 | -0.445311 | 1.078382 | -0.398658 | -0.158526 | -0.108575 | 1.089482 | -0.186445 | 1.657385 | 0.392487 | | 31.29167 | 1.110589 | 0.458662 |
| 31.208 | -0.507161 | 2.016688 | 0.810789 | -0.304629 | 1.345404 | 1.596106 | 0.569106 | 0.200871 | 0.837826 | 1.108591 | 1.554957 | 0.317885 | 0.903205 | 1.287105 | 1.340044 | -0.009446 | 1.311821 | -0.15407 | 0.052 | 0.072609 | 1.435728 | 0.050244 | 3.565197 | 0.815653 | | 31.30556 | 0.903205 | 0.978281 |
| 31.222 | 0.097357 | 1.491986 | -1.238823 | -0.283301 | 0.487542 | 0.995833 | 1.081496 | -0.508378 | 0.754173 | 1.128363 | 1.809825 | 1.166549 | 1.179293 | 1.353369 | 1.656179 | 0.479052 | 1.616096 | 0.30482 | 0.36301 | 0.348818 | 1.771799 | 0.374787 | 2.771356 | 1.168784 | | 31.31944 | 1.179293 | 1.313541 |
| 31.236 | 0.335716 | 1.046494 | -1.666008 | 0.622923 | -0.51773 | 0.509641 | 0.801017 | -0.076895 | 0.999025 | 0.748314 | 0.650226 | 2.049367 | 2.8108 | 1.056513 | 1.880053 | 0.783932 | 1.655399 | 0.648361 | 0.529777 | 0.503757 | 1.725521 | 0.545668 | 0.919734 | 1.139553 | | 31.33333 | 2.8108 | 1.191508 |

| | | | | | | | | | | | | | | | | | | | | | | | | | | | | |
|---|---|---|---|---|---|---|---|---|---|---|---|---|---|---|---|---|---|---|---|---|---|---|---|---|---|---|---|---|
| 34.569 | -1.442123 | -2.621983 | 1.266824 | -0.065062 | 2.071359 | 0.68585 | -0.137597 | 1.084309 | 1.100331 | -0.54342 | -0.767971 | 0.101523 | -1.10393 | 0.163428 | -0.430145 | 0.313413 | -0.46212 | 0.463375 | 0.395226 | 0.371188 | -3.116103 | 0.39331 | -1.558756 | 0.490185 | | 34.66667 | -1.10393 | 0.802522 |
| 34.583 | 0.109893 | -1.397612 | -0.078331 | 0.077017 | 0.827143 | 0.527749 | -0.048588 | 1.002837 | 0.209006 | -1.016248 | -0.044725 | -0.472746 | 3.01991 | 1.048567 | 2.467236 | 0.370347 | 0.162911 | 0.576221 | 0.433436 | 0.41915 | -1.895262 | 0.446385 | -0.198948 | 0.588222 | | 34.68056 | 3.01991 | 0.721836 |
| 34.597 | 2.406926 | 1.159127 | -0.899596 | -0.001247 | -0.794175 | 0.554117 | 1.000551 | 0.35343 | -0.020871 | 1.216923 | -0.170477 | -0.598223 | 5.941992 | 3.112261 | 3.312837 | 0.408482 | 0.720934 | 0.624294 | 0.452639 | 0.431955 | -0.105241 | 0.477593 | 0.643032 | 1.172262 | | 34.69444 | 5.941992 | 0.841704 |
| 34.611 | 3.514185 | 3.506817 | -2.97026 | -0.192083 | -0.153114 | 0.847383 | 2.725387 | -0.845029 | -0.660832 | 1.60492 | 0.045846 | -0.439237 | 4.492715 | -0.424633 | 1.864505 | 0.383899 | 0.891359 | 0.564904 | 0.407873 | 0.385194 | 1.684373 | 0.426509 | 1.24199 | 0.976253 | | 34.70833 | 4.492715 | 0.397296 |
| 34.625 | 4.35273 | 2.963098 | -2.786724 | -0.325191 | -1.27267 | 0.969695 | 2.110869 | -1.462633 | -2.00474 | 0.948748 | -0.387721 | -0.145992 | -0.221142 | -3.832692 | 1.481632 | 0.355663 | 0.868073 | 0.489866 | 0.363243 | 0.342061 | 3.316282 | 0.355977 | 1.233164 | 0.040763 | | 34.72222 | -0.221142 | -0.280004 |
| 34.639 | 3.760035 | 1.456491 | -1.205162 | -0.230981 | -1.376989 | 0.580895 | 1.963255 | -2.852848 | -0.838113 | 1.659831 | -1.869666 | 0.344598 | -3.084296 | -2.754016 | 1.37056 | 0.329945 | 0.585697 | 0.401129 | 0.339887 | 0.314682 | 2.594452 | 0.323761 | 0.699149 | -0.408508 | | 34.73611 | -3.084296 | -0.702204 |
| 34.653 | 1.387542 | 1.202642 | -0.049016 | -0.198602 | -0.669073 | -0.016335 | 1.338747 | 0.005109 | 1.207606 | 2.687666 | -2.327214 | 0.817257 | -0.741768 | -0.417379 | 1.241349 | 0.247975 | 0.329964 | 0.275196 | 0.259917 | 0.246352 | 0.686684 | 0.264187 | 0.469432 | -0.605934 | | 34.75 | -0.741768 | -0.67246 |
| 34.667 | -0.961021 | 3.42592 | 0.707853 | 0.070537 | 0.252925 | -0.329043 | 0.688567 | 2.517676 | 1.264389 | 0.652269 | -1.175709 | 1.116964 | 3.574236 | 0.481873 | 0.968953 | 0.159732 | 0.004049 | 0.185 | 0.175674 | 0.178354 | -0.647769 | 0.188519 | 0.099493 | -0.522267 | | 34.76389 | 3.574236 | -0.437479 |
| 34.681 | -3.020576 | 2.794403 | 2.323093 | 0.009376 | 0.880789 | -0.253164 | 1.447027 | 1.668124 | 0.830587 | -0.485903 | -0.27236 | 1.3625 | 6.467291 | 1.222398 | 0.302195 | 0.119166 | -0.567488 | 0.131096 | 0.148351 | 0.14822 | -2.803966 | 0.150682 | -0.221491 | -0.407414 | | 34.77778 | 6.467291 | -0.208458 |
| 34.694 | -1.656767 | 1.158412 | 1.983236 | 0.310018 | 0.576706 | -0.40467 | 1.657022 | 0.678206 | 0.339324 | -0.84017 | 1.476123 | 0.971373 | 8.326622 | 1.489421 | -0.023336 | 0.042054 | -1.161602 | 0.01334 | 0.074055 | 0.063425 | -2.604458 | 0.075026 | -0.447035 | -0.430914 | | 34.79167 | 8.326622 | -0.11456 |
| 34.708 | 0.63261 | 0.275329 | 1.657695 | 0.332857 | -0.165429 | -0.5242 | -0.982647 | 0.041002 | 0.198903 | -1.830094 | 2.303867 | 1.193583 | 5.166945 | 2.57167 | -1.466527 | -0.048801 | -1.011986 | -0.124864 | -0.050909 | -0.057007 | -2.034039 | -0.03899 | -0.48928 | -0.764169 | | 34.80556 | 5.166945 | -0.260912 |
| 34.722 | 1.662708 | 0.070854 | 1.110232 | -0.294171 | -0.694803 | -0.635937 | 0.62649 | -0.417256 | -0.314554 | 1.390714 | 2.204673 | 0.158833 | 1.5563 | 2.497289 | -2.141783 | -0.064566 | -0.975058 | -0.196445 | -0.070934 | -0.064757 | -0.50736 | -0.059953 | -0.42448 | -0.679002 | | 34.81944 | 1.5563 | -0.262925 |
| 34.736 | 2.297845 | -0.73281 | -0.162408 | -0.128785 | -0.873766 | -0.558158 | 1.330671 | -1.029341 | -0.620714 | 0.974381 | 0.250872 | -0.597513 | -1.129979 | 1.605483 | -1.510615 | 0.005168 | -0.862006 | -0.209857 | 0.013797 | 0.023027 | 0.793077 | 0.009185 | -0.558884 | -0.370691 | | 34.83333 | -1.129979 | -0.321207 |
| 34.75 | 1.585291 | -1.999081 | -0.721608 | -1.306211 | -0.874961 | -0.279373 | 0.35679 | -1.205157 | -1.356964 | 0.956772 | -1.476838 | -1.781916 | -3.84707 | -0.147683 | -1.328117 | 0.046754 | -0.975372 | -0.288408 | 0.071302 | 0.080437 | 1.370432 | 0.064509 | -0.746983 | -0.354792 | | 34.84722 | -3.84707 | -0.531301 |
| 34.764 | 0.801156 | -1.590588 | -0.690099 | -0.573507 | -1.298175 | -0.097258 | -1.012449 | -1.551427 | -1.040282 | 1.723297 | -3.073378 | -1.788661 | -4.999066 | -2.822885 | -1.200775 | 0.025991 | -1.705233 | -0.444611 | 0.065821 | 0.070312 | 1.113094 | 0.075878 | -1.044217 | -0.722919 | | 34.86111 | -4.999066 | -0.922753 |
| 34.778 | 0.341751 | -0.411914 | -0.362852 | -0.132029 | -2.358382 | -0.549316 | -1.791506 | -1.016448 | 0.333625 | -0.727416 | -4.848579 | -1.909079 | -3.40161 | -4.745193 | -1.051179 | 0.003909 | -1.608695 | -0.663961 | 0.021698 | 0.017419 | -0.189898 | 0.046142 | -1.310345 | -1.271697 | | 34.875 | -3.40161 | -1.374753 |
| 34.792 | -1.416198 | 1.373747 | 0.004743 | 0.504738 | -1.340079 | -1.360578 | -0.601868 | -0.328583 | 1.276295 | -0.424166 | -5.340486 | -1.773882 | -0.76362 | -4.645051 | -1.219811 | 0.022805 | -1.544565 | -1.007871 | -0.020969 | -0.026376 | -2.063597 | 0.000713 | -1.499238 | -1.824456 | | 34.88889 | -0.76362 | -1.793687 |
| 34.806 | -3.701444 | 1.396624 | 0.098795 | 0.273895 | -1.279877 | -1.864296 | 1.15751 | 0.599 | 1.734002 | 2.546695 | -1.32224 | -2.436577 | 0.37405 | -3.01796 | -1.54112 | 0.049855 | -1.805329 | -1.290223 | -0.027481 | -0.026467 | -3.243006 | -0.013827 | -1.64122 | -2.570164 | | 34.90278 | 0.37405 | -2.544179 |
| 34.819 | -3.830487 | -0.07435 | -0.064318 | 0.092983 | -1.88988 | -2.019024 | 1.577508 | 1.013391 | 1.464528 | 2.689072 | 0.608559 | -2.394038 | 0.595648 | -1.782683 | -2.274243 | 0.043187 | -1.917733 | -1.561948 | -0.030699 | -0.016043 | -4.138851 | -0.019019 | -1.536688 | -2.799972 | | 34.91667 | 0.595648 | -2.965316 |
| 34.833 | -2.619959 | -0.879334 | -0.319066 | 0.133484 | -1.23794 | -2.073601 | -0.091748 | 0.386538 | 0.392244 | 1.762333 | 2.377214 | -1.552329 | -0.512538 | -1.660502 | -2.755519 | -0.033228 | -2.879246 | -1.744551 | -0.067523 | -0.049671 | -3.45162 | -0.064588 | -1.276213 | -2.684161 | | 34.93056 | -0.512538 | -2.97511 |
| 34.847 | -0.474301 | -2.162423 | -0.621428 | -0.780068 | -1.448528 | -2.537026 | -1.633578 | -0.690137 | -0.702132 | -1.01891 | 0.816112 | 1.707562 | -3.112764 | -3.427129 | -2.729491 | -0.082106 | -3.665764 | -1.806249 | -0.102017 | -0.097816 | -1.315397 | -0.110098 | -1.080212 | -2.438074 | | 34.94444 | -3.112764 | -2.906785 |
| 34.861 | 1.0399 | -3.121099 | -0.804722 | -0.912359 | -2.025271 | -2.509991 | -2.901093 | -2.105982 | -3.125889 | -2.767328 | -0.399857 | 1.323902 | -6.489951 | -4.748099 | -2.52669 | -0.103348 | -4.028852 | -1.973242 | -0.120135 | -0.130978 | 0.370203 | -0.125348 | -1.109487 | -2.18523 | | 34.95833 | -6.489951 | -2.682148 |
| 34.875 | 0.283036 | -2.780511 | -0.865689 | -0.535063 | -2.755 | -1.909247 | -3.649838 | -3.874443 | -3.365597 | -2.348826 | -1.707122 | -1.308249 | -8.762678 | -4.170224 | -2.503084 | -0.191062 | -3.721276 | -1.917214 | -0.178128 | -0.18929 | 0.954406 | -0.171842 | -1.150448 | -1.97793 | | 34.97222 | -8.762678 | -2.430736 |
| 34.889 | -1.500599 | -0.522692 | -1.008578 | -0.097532 | -2.134975 | -1.320011 | -4.662277 | -5.624253 | -2.750239 | -2.945074 | -2.485284 | -4.145055 | -8.169484 | -3.358787 | -2.298972 | -0.254398 | -3.278812 | -1.787013 | -0.266255 | -0.260248 | 0.749188 | -0.254137 | -1.172285 | -1.762732 | | 34.98611 | -8.169484 | -2.148624 |
| 34.903 | -3.775587 | 1.313705 | -0.754869 | 0.015187 | -0.580642 | -1.254853 | -5.035884 | -6.463834 | -1.334707 | -1.938896 | -2.787053 | -4.701043 | -5.760156 | -2.578117 | -1.863236 | -0.286901 | -2.831292 | -1.417991 | -0.258264 | -0.24498 | 0.083591 | -0.252861 | -1.103275 | -1.611178 | | 35 | -5.760156 | -1.871811 |
| 34.917 | -4.158492 | 2.706634 | 0.253088 | 0.302612 | 0.133238 | -1.577475 | -5.855016 | -4.036769 | 1.756623 | -0.700929 | -1.733655 | -5.34238 | -2.984709 | -1.8337 | -1.43496 | -0.117202 | -2.20142 | -1.031258 | -0.039097 | -0.063519 | -0.206278 | -0.056914 | -0.84551 | -1.332455 | | 35.01389 | -2.984709 | -1.49039 |
| 34.931 | -4.01833 | 2.194575 | 0.971587 | 0.594506 | 0.56557 | -1.237814 | -3.429903 | -0.764642 | 3.460846 | 0.602028 | -1.470307 | -4.97162 | -1.57073 | -1.328893 | -1.145251 | 0.141082 | -1.152033 | -0.662598 | 0.294622 | 0.207241 | -0.204076 | 0.242949 | -0.47631 | -0.938847 | | 35.02778 | -1.57073 | -0.990457 |
| 34.944 | -2.855101 | -0.415677 | 1.259506 | 0.413159 | 0.19015 | -0.762695 | -0.772724 | 0.754837 | 1.608088 | 1.278428 | -1.185763 | -4.257172 | -2.131693 | -1.081615 | -0.99299 | 0.156848 | -0.484831 | -0.543509 | 0.307709 | 0.207594 | -0.407252 | 0.263536 | -0.391219 | -0.826531 | | 35.04167 | -2.131693 | -0.754244 |
| 34.958 | -2.505095 | -2.978015 | 0.367469 | 0.136863 | -1.508807 | -1.04462 | 0.749947 | -1.676403 | -0.431098 | -0.665194 | -1.956929 | -4.176289 | -5.148354 | -1.116841 | -1.010521 | -0.134227 | -0.839537 | -0.850663 | -0.129428 | -0.139307 | -1.066126 | -0.152351 | -0.784666 | -1.291877 | | 35.05556 | -5.148354 | -1.095231 |
| 34.972 | -2.968513 | -4.854137 | -0.492357 | -0.330614 | -4.383901 | -2.327158 | 0.814254 | -3.524422 | -2.828224 | -3.326406 | -3.62223 | -4.226959 | -8.672078 | -0.828543 | -0.836509 | -0.34094 | -1.515828 | -0.816011 | -0.29839 | -0.248182 | -1.511089 | -0.354085 | -0.997423 | -1.845772 | | 35.06944 | -8.672078 | -1.659599 |
| 34.986 | -3.795428 | -4.87187 | -1.471412 | -0.144631 | -5.661294 | -3.644578 | -0.090397 | -3.799693 | -5.075819 | -5.169585 | -5.911147 | -3.568022 | -10.09945 | -0.301458 | 0.436517 | -0.193616 | -1.406265 | -0.122961 | -0.088126 | -0.049951 | -1.511578 | -0.145336 | -0.88187 | -1.572063 | | 35.08333 | -10.09945 | -1.440897 |
| 35 | -4.020255 | -5.074142 | -1.410256 | -0.43156 | -5.086747 | -3.685791 | -1.448309 | -4.123583 | -5.387484 | -4.871431 | -6.095187 | -0.188828 | -8.483471 | -0.160844 | -0.196705 | 0.051181 | -0.494855 | 0.716653 | 0.049639 | 0.059805 | -1.257233 | 0.015108 | -0.463002 | -0.516005 | | 35.09722 | -8.483471 | -0.456763 |
| 35.014 | -2.497711 | -3.140244 | -0.424803 | -1.044823 | -4.180173 | -2.804722 | -1.262128 | -4.128128 | -3.764317 | -2.854031 | -3.376347 | 0.066642 | -5.315838 | -0.202908 | -0.529482 | 0.168489 | 0.405728 | 0.781913 | 0.034398 | 0.033658 | -0.534997 | 0.021586 | 0.111852 | 0.130803 | | 35.11111 | -5.315838 | 0.152679 |
| 35.028 | -1.319775 | -1.454463 | 0.053802 | -0.906285 | -2.077928 | -1.167077 | -1.164536 | -3.791217 | -1.959108 | -1.137184 | -1.141884 | -1.189348 | -2.844371 | 0.682134 | -0.841441 | 0.07928 | 0.731694 | 0.407219 | -0.023214 | -0.032117 | -0.195831 | -0.028897 | 0.134402 | 0.386882 | | 35.125 | -2.844371 | 0.286124 |
| 35.042 | -0.302131 | -0.321451 | -0.046617 | -0.573265 | -0.582751 | 0.105008 | -0.837897 | -3.06245 | -0.994352 | 0.143682 | 0.109818 | -1.896419 | -1.148574 | 1.473565 | -0.2441 | -0.14264 | 0.64596 | -0.111414 | -0.100726 | -0.10456 | -0.050534 | -0.109447 | -0.051942 | 0.256824 | | 35.13889 | -1.148574 | 0.099091 |
| 35.056 | 1.179938 | 1.170975 | -0.214095 | -0.537197 | 0.585414 | 0.82083 | -0.489662 | -1.932313 | -0.372151 | 1.301479 | 1.276823 | -2.228299 | 0.505255 | 1.588153 | 0.858497 | -0.408433 | 0.426638 | -0.572244 | -0.149795 | -0.142461 | 0.013619 | -0.159634 | -0.537773 | -0.232929 | | 35.15278 | 0.505255 | -0.494151 |
| 35.069 | 3.33327 | 3.708315 | -0.067226 | -0.134457 | 1.209727 | 0.955468 | 0.337121 | -0.794925 | 0.292781 | 2.08763 | 2.221531 | -0.549402 | 2.152693 | 1.15587 | 0.097139 | -0.675037 | 0.172966 | -0.642996 | -0.128265 | -0.129597 | 0.068804 | -0.132973 | -1.189249 | -0.744266 | | 35.16667 | 2.152693 | -0.97102 |
| 35.083 | 4.700507 | 4.37505 | 0.526797 | 0.193675 | 1.775344 | 0.778967 | 0.970242 | 0.440714 | 1.047682 | 2.182733 | 2.474127 | 2.618828 | 3.047182 | 1.164259 | -0.3588 | -0.81817 | -0.176493 | -0.423092 | -0.071399 | -0.09351 | 0.278358 | -0.070236 | -1.385632 | -0.575434 | | 35.18056 | 3.047182 | -0.678421 |
| 35.097 | 3.752774 | 2.390769 | 1.422018 | 0.624016 | 2.838277 | 0.528852 | 0.9401 | 1.722799 | 1.361462 | 2.5163 | 1.904789 | 3.418912 | 3.070732 | 1.013318 | 0.714133 | -0.640155 | -0.313195 | -0.252206 | -0.02314 | -0.050728 | 0.779685 | -0.018672 | -0.952666 | 0.209281 | | 35.19444 | 3.070732 | 0.238691 |
| 35.111 | 2.225052 | 0.038076 | 1.36606 | 0.526698 | 3.00063 | 0.250987 | 0.684815 | 2.467162 | 1.265171 | 2.044078 | 1.802177 | 2.959798 | 2.324708 | 1.027748 | 1.049544 | -0.038741 | 0.057018 | -0.110356 | 0.014468 | 0.003275 | 1.144328 | 0.018624 | -0.583119 | 1.113568 | | 35.20833 | 2.324708 | 1.158361 |
| 35.125 | 0.859778 | -1.912318 | 0.614819 | 0.255531 | 1.81615 | 0.096325 | 0.64945 | 2.811628 | 0.739615 | 1.718022 | 2.100504 | 0.7971 | 1.122725 | 1.180596 | 2.174317 | 0.619046 | 0.942973 | 0.089354 | 0.061435 | 0.061781 | 1.136916 | 0.05794 | -0.725702 | 1.571235 | | 35.22222 | 1.122725 | 1.804279 |
| 35.139 | 0.592887 | -2.255254 | -0.27321 | -0.313824 | 0.565189 | 0.043322 | 0.604759 | 3.344747 | -0.122987 | 1.645294 | 2.143888 | -2.000197 | -0.083495 | 0.624146 | 1.428944 | 0.814584 | 2.176295 | 0.386702 | 0.109795 | 0.10577 | 0.892929 | 0.100802 | -0.635399 | 1.263267 | | 35.23611 | -0.083495 | 1.658759 |
| 35.153 | 1.356689 | -2.976222 | -0.971113 | -0.743011 | -1.010814 | -0.01086 | 0.598359 | 2.338261 | -0.869049 | 1.533993 | 1.669086 | -3.876671 | -1.166463 | -0.821818 | 1.107903 | 0.703969 | 3.219698 | 0.712932 | 0.139851 | 0.131043 | 0.364234 | 0.135427 | -0.013385 | 0.436131 | | 35.25 | -1.166463 | 0.704156 |
| 35.167 | 1.931831 | 0.613323 | -1.283483 | -0.143039 | -1.493379 | 0.506129 | 0.663637 | 0.626552 | -0.501208 | 1.034841 | 0.907358 | -1.746676 | -0.470458 | -1.488734 | -0.391901 | 0.512889 | 2.84835 | 0.882926 | 0.152962 | 0.143239 | 0.039921 | 0.151154 | 0.56849 | -0.444007 | | 35.26389 | -0.470458 | -0.513237 |
| 35.181 | 1.203773 | 3.191403 | 2.187751 | 0.144134 | -0.111349 | 1.568207 | 1.527128 | 0.154199 | 0.613714 | -0.073015 | 0.186947 | 0.502272 | 2.230508 | -0.757332 | 0.028711 | 0.221041 | 1.673617 | 0.752463 | 0.128223 | 0.119033 | 0.154381 | 0.117513 | 0.670851 | -0.544745 | | 35.27778 | 2.230508 | -0.705332 |
| 35.194 | 0.645357 | 4.961821 | 4.290555 | 0.833081 | 0.839391 | 2.51259 | 1.141186 | 0.734056 | 1.715696 | -0.122813 | 0.651263 | 3.405833 | 3.994666 | 0.405658 | 1.203636 | 0.087275 | 0.780616 | 0.459785 | 0.079182 | 0.075051 | 0.628533 | 0.065815 | 0.451633 | 0.311894 | | 35.29167 | 3.994666 | 0.179076 |
| 35.208 | 0.803737 | 3.680515 | 2.983879 | 0.57367 | 1.666196 | 2.532471 | 0.246944 | 1.687667 | 2.811456 | 0.598449 | 1.84481 | 4.660936 | 3.680862 | 1.584963 | 1.169311 | 0.020113 | 0.586192 | 0.131778 | 0.033194 | 0.022012 | 1.245138 | 0.038148 | 0.125453 | 1.171932 | | 35.30556 | 3.680862 | 1.145415 |
| 35.222 | 0.991739 | 2.706675 | 0.844229 | -0.166971 | 2.390879 | 1.830685 | -0.522418 | 2.418961 | 3.396325 | 0.99213 | 1.885467 | 3.186008 | 1.878912 | 1.524322 | 1.963245 | -0.044894 | 0.801548 | -0.053731 | -0.015231 | -0.038125 | 1.723169 | 0.012322 | 0.041232 | 1.381207 | | 35.31944 | 1.878912 | 1.421689 |
| 35.236 | 0.987817 | 1.841749 | -2.34113 | 0.157179 | 2.685256 | 1.662738 | -0.775803 | 2.581748 | 3.095765 | 1.337698 | 0.325587 | 1.514173 | 0.309014 | 1.134093 | 0.410623 | -0.067836 | 0.950212 | -0.110617 | -0.030236 | -0.056857 | 1.787309 | -0.004625 | -0.108105 | 0.670644 | | 35.33333 | 0.309014 | 0.800764 |
| 35.25 | 1.092527 | 0.85441 | -2.209926 | 0.625068 | 2.820071 | 2.199097 | -0.211983 | 2.767155 | 2.77503 | 1.789853 | -0.465063 | 0.369671 | 0.203285 | 0.451756 | -1.383251 | -0.015456 | 1.034909 | -0.070391 | 0.012902 | -0.009389 | 1.580179 | 0.011855 | -0.784142 | 0.110371 | | 35.34722 | 0.203285 | 0.038414 |
| 35.264 | 1.730377 | -0.439801 | 0.31377 | 0.118136 | 2.606392 | 2.521979 | 0.54739 | 3.249291 | 2.50767 | 2.289859 | -0.324315 | 1.89629 | 1.450582 | -0.103678 | 0.032388 | 0.045064 | 1.237185 | 0.060823 | 0.058158 | 0.041569 | 1.106362 | 0.039638 | -0.503374 | 0.495941 | | 35.36111 | 1.450582 | 0.588577 |
| 35.278 | 2.640115 | 0.74625 | 2.29408 | -0.245215 | 2.014558 | 2.886121 | 0.889349 | 3.617691 | 2.248498 | 2.524575 | 0.971267 | 3.665912 | 3.363388 | 0.524877 | 1.353264 | 0.035512 | 1.582409 | 0.221005 | 0.058752 | 0.039157 | 0.657413 | 0.04667 | 0.72007 | 1.227174 | | 35.375 | 3.363388 | 1.449178 |
| 35.292 | 2.118082 | 2.076006 | 1.925082 | 0.102954 | 1.566708 | 2.567076 | 0.806304 | 3.00511 | 1.881999 | 2.360236 | 2.447255 | 5.266467 | 4.483599 | 1.552989 | 2.26049 | -0.005469 | 1.893603 | 0.323359 | 0.007082 | -0.010373 | 0.431399 | 0.006618 | 1.984987 | 1.674943 | | 35.38889 | 4.483599 | 1.862721 |
| 35.306 | 0.756513 | 2.433187 | 1.10649 | 0.492537 | 1.098168 | 1.317126 | 0.73125 | 1.943712 | 1.525694 | 2.045983 | 2.461994 | 4.983766 | 3.843499 | 2.514684 | 2.458011 | -0.012739 | 1.969751 | 0.435871 | -0.026953 | -0.040072 | 0.417084 | -0.025864 | 2.012254 | 1.698118 | | 35.40278 | 3.843499 | 1.754822 |
| 35.319 | 0.111277 | 1.691165 | 0.745042 | 0.277165 | 1.09246 | 0.601037 | 0.654007 | 0.998989 | 1.437869 | 1.901174 | 1.988241 | 3.391728 | 2.606055 | 3.040864 | 2.5099 | 0.014911 | 2.094327 | 0.603531 | -0.015227 | -0.034133 | 0.62959 | -0.007914 | 2.237197 | 1.720967 | | 35.41667 | 2.606055 | 1.611115 |
| 35.333 | -0.193212 | 1.355072 | 0.354844 | 0.364992 | 1.520404 | 0.908376 | 0.573342 | 0.834807 | 1.739597 | 1.614839 | 1.449429 | 2.278287 | 1.824489 | 2.888316 | 1.628951 | -0.002344 | 2.463126 | 0.70574 | -0.036829 | -0.04773 | 0.945619 | -0.013818 | 2.482131 | 1.741027 | | 35.43056 | 1.824489 | 1.663126 |
| 35.347 | 0.801366 | 0.649205 | 0.691298 | 0.21802 | 1.953933 | 1.74033 | 0.553643 | 1.299229 | 1.98589 | 1.176366 | 1.466059 | 1.555692 | 1.729589 | 2.723455 | 2.12661 | -0.025978 | 2.643228 | 0.715159 | -0.049151 | -0.053628 | 1.073924 | -0.043159 | 2.338734 | 1.761988 | | 35.44444 | 1.729589 | 1.865615 |
| 35.361 | 2.805747 | -1.131182 | 1.167181 | -0.21494 | 2.029818 | 2.666909 | 0.438913 | 2.614653 | 2.309766 | 1.224106 | 1.839342 | 1.184647 | 2.064968 | 4.072727 | 3.103245 | -0.044111 | 2.661562 | 0.606689 | -0.055146 | -0.061157 | 0.932958 | -0.1047 | 1.835132 | 1.708238 | | 35.45833 | 2.064968 | 1.936034 |
| 35.375 | 4.568299 | -3.029268 | 1.314672 | -0.490919 | 2.024434 | 2.856303 | 0.270775 | 3.50867 | 3.403005 | 1.373495 | 2.30019 | 1.593064 | 2.520507 | 5.757798 | 3.815152 | -0.095543 | 2.740335 | 0.424841 | -0.149917 | -0.120542 | 0.860976 | -0.182405 | 1.54607 | 1.679151 | | 35.47222 | 2.520507 | 1.907639 |
| 35.389 | 4.852614 | -2.906707 | 1.542267 | -0.121515 | 1.897579 | 2.282523 | 0.097076 | 4.532724 | 3.526958 | 1.947793 | 2.645205 | 2.712218 | 3.370289 | 6.294905 | 3.415164 | -0.057894 | 2.866458 | 0.246146 | -0.108778 | -0.04738 | 0.961057 | -0.049709 | 1.790727 | 1.694605 | | 35.48611 | 3.370289 | 1.770728 |
| 35.403 | 3.842497 | -0.826832 | 1.724516 | -0.037096 | 2.164101 | 2.234974 | 0.378779 | 6.47812 | 3.370494 | 2.197158 | 3.340238 | 4.591749 | 4.733073 | 4.922714 | 2.339894 | 0.060545 | 2.856358 | 0.263993 | 0.081986 | 0.104479 | 1.074682 | 0.116448 | 2.235139 | 1.595606 | | 35.5 | 4.733073 | 1.635207 |
| 35.417 | 3.029968 | 2.068973 | 1.657823 | 0.271348 | 2.818033 | 2.786706 | 1.770513 | 5.828177 | 3.163981 | 2.356498 | 3.854777 | 4.479249 | 5.475755 | 2.99042 | 1.785816 | 0.110245 | 2.351449 | 0.319373 | 0.163109 | 0.165897 | 1.098018 | 0.171176 | 1.975893 | 1.563101 | | 35.51389 | 5.475755 | 1.525748 |
| 35.431 | 3.049557 | 3.518254 | 1.987587 | 0.37572 | 2.755224 | 2.10136 | 2.754642 | 4.227468 | 2.101086 | 2.799446 | 4.620587 | 3.77977 | 4.627926 | 1.324418 | 1.860322 | 0.053679 | 1.581132 | 0.28914 | 0.09959 | 0.089977 | 1.308705 | 0.113159 | 1.845333 | 1.516675 | | 35.52778 | 4.627926 | 1.512042 |
| 35.444 | 3.563863 | 3.016681 | 2.798931 | 0.141202 | 2.010572 | 1.442156 | 1.762419 | 3.067954 | 1.931078 | 2.084924 | 5.036225 | 4.388161 | 3.521561 | 0.44345 | 0.991782 | -0.250497 | 0.805739 | -0.017977 | -0.209906 | -0.219362 | 1.474163 | -0.130505 | 1.657104 | 1.526239 | | 35.54167 | 3.521561 | 1.356152 |
| 35.458 | 4.05658 | 1.809143 | 2.27533 | -0.126585 | 1.844139 | 0.876098 | 0.152647 | 2.339042 | 2.618264 | 1.759111 | 4.382545 | 4.621821 | 3.463082 | 0.916203 | 0.352204 | -0.809016 | 0.074191 | -0.666328 | -0.806757 | -0.794965 | 1.560069 | -0.640887 | 1.227254 | 1.278248 | | 35.55556 | 3.463082 | 0.835379 |
| 35.472 | 2.430912 | 0.790037 | 1.551791 | -0.226899 | 1.647059 | 0.758132 | -0.629376 | 1.227175 | 3.478061 | 3.040296 | 5.297613 | 2.743691 | 4.104981 | 1.957315 | -0.267299 | -1.151923 | -0.509986 | -1.094606 | -1.235407 | -1.172252 | 0.672619 | -1.07548 | 0.745108 | 0.366676 | | 35.56944 | 4.104981 | 0.075679 |
| 35.486 | 1.236619 | -0.193758 | 2.357005 | -0.111537 | 1.844195 | 0.392958 | -1.539261 | 0.733886 | 2.227466 | 4.318676 | 3.708992 | 1.071302 | 4.549235 | 2.776306 | -0.598152 | -0.928794 | -0.537825 | -0.968179 | -1.011199 | -0.963595 | 0.244514 | -0.945517 | 0.639329 | -0.162776 | | 35.58333 | 4.549235 | -0.192684 |
| 35.5 | 1.263016 | -0.736096 | 3.782087 | 0.093886 | 0.850955 | -0.415957 | -1.276828 | 0.715528 | 1.345556 | 2.494919 | 0.177032 | 2.227497 | 5.514387 | 3.073615 | -1.836584 | -0.50565 | -0.201399 | -0.663902 | -0.535525 | -0.510776 | 0.773321 | -0.526291 | 0.949992 | -0.111139 | | 35.59722 | 5.514387 | -0.123066 |
| 35.514 | 1.986723 | 0.189153 | 2.445221 | 0.341011 | 0.871693 | -0.345424 | 0.06307 | -0.031373 | 0.017234 | 0.291622 | 3.008906 | 3.569282 | 4.377458 | 1.564586 | -0.979332 | -0.274829 | 0.144115 | -0.556009 | -0.271008 | -0.258146 | 1.732903 | -0.27148 | 1.716245 | -0.012209 | | 35.61111 | 4.377458 | -0.003066 |
| 35.528 | 0.458594 | 0.182021 | 0.788521 | 0.668828 | 1.303511 | 0.805557 | 0.476273 | -0.215056 | -1.34225 | -1.212967 | 0.925492 | 0.889071 | 1.501094 | 1.386716 | 0.756664 | -0.171867 | 0.364516 | -0.613543 | -0.162728 | -0.147316 | 2.647216 | -0.162502 | 1.620727 | 0.107729 | | 35.625 | 1.501094 | -0.169502 |
| 35.542 | -2.000192 | -0.61802 | -0.20572 | 0.392222 | 1.388865 | 0.887242 | 0.465651 | -1.069553 | -1.398599 | 0.162694 | -1.396475 | -0.522731 | -1.950507 | 0.859025 | 1.590776 | -0.094333 | 0.491843 | -0.713477 | -0.079238 | -0.077554 | 1.818783 | -0.07725 | 0.315688 | -0.469699 | | 35.63889 | -1.950507 | -0.689355 |
| 35.556 | -2.268205 | -0.091044 | -1.377265 | 0.057972 | 0.061093 | 0.679794 | -0.056681 | -0.234584 | -0.204687 | 1.793102 | -1.662206 | -1.504751 | -2.861492 | -1.634203 | 1.320299 | -0.072868 | 0.514137 | -0.602558 | -0.060802 | -0.066751 | -0.325718 | -0.064747 | -0.535671 | -0.954394 | | 35.65278 | -2.861492 | -1.052965 |
| 35.569 | -2.492966 | -0.640747 | -2.426657 | -0.15147 | 0.874371 | 0.509868 | -0.632154 | 0.57379 | 1.163839 | 1.412911 | -0.935876 | -0.360315 | -0.510139 | -3.601589 | 1.018985 | -0.0726 | 0.354984 | -0.387251 | -0.066483 | -0.067076 | -3.114459 | -0.080761 | 0.090167 | -1.158539 | | 35.66667 | -0.510139 | -0.894533 |
| 35.583 | -5.674047 | -2.401296 | -3.143405 | 0.140853 | -0.730677 | 0.201901 | -0.426364 | 1.823826 | 0.440851 | 1.058478 | 2.087636 | 0.163692 | 2.709794 | -3.886414 | 0.674098 | -0.02011 | 0.034158 | -0.17215 | -0.002302 | -0.005924 | -2.739069 | -0.014823 | 0.017893 | -1.429765 | | 35.68056 | 2.709794 | -0.724249 |
| 35.597 | -2.748481 | -2.19072 | -2.129339 | 0.15629 | 0.525465 | -0.26182 | -0.886612 | 0.727161 | -0.038989 | -1.209203 | 3.327936 | 1.164626 | 3.298267 | -2.808343 | 0.738767 | 0.029079 | -0.168523 | -0.07193 | 0.057741 | 0.039463 | -0.857668 | 0.056922 | -0.678213 | -1.940605 | | 35.69444 | 3.298267 | -2.07233 |
| 35.611 | -1.188971 | -1.846713 | -0.005708 | 0.841872 | 1.290002 | -0.794078 | -0.370753 | -0.042139 | -0.519862 | -3.359114 | 3.065426 | 0.48782 | 0.188829 | -2.795153 | 0.206367 | 0.024017 | -0.243021 | 0.029338 | 0.051697 | 0.031515 | 0.576844 | 0.053974 | -0.045298 | -1.055142 | | 35.70833 | 0.188829 | -1.597683 |
| 35.625 | 0.063818 | -0.96423 | 2.297118 | 0.763618 | 1.724576 | -1.468722 | -0.647813 | -1.047027 | -0.466866 | -4.231494 | 2.032522 | -1.089525 | -5.22793 | -0.949521 | -0.549046 | 0.055445 | -0.215064 | 0.281661 | 0.083874 | 0.068093 | 2.663096 | 0.075167 | -0.38791 | 0.32545 | | 35.72222 | -5.22793 | -0.235958 |
| 35.639 | 0.981685 | -1.283387 | 1.875062 | 0.096287 | 0.369956 | -0.535847 | -0.623294 | -1.546869 | -0.287693 | -3.506123 | -0.029946 | -2.211179 | -6.261386 | 0.39639 | 0.817892 | 0.137298 | -0.131215 | 0.630378 | 0.167268 | 0.152888 | 2.618666 | 0.147805 | -0.633269 | 0.896013 | | 35.73611 | -6.261386 | 0.717409 |
| 35.653 | 0.99388 | 0.030238 | 2.041132 | -0.20518 | -1.025442 | -0.680071 | 0.236067 | -1.942329 | -0.001613 | -0.913812 | -2.531344 | -1.31336 | -0.981012 | -0.233334 | 0.608551 | 0.172944 | -0.1097 | 0.907946 | 0.197761 | 0.183782 | 1.170633 | 0.175786 | -0.986441 | 1.512546 | | 35.75 | -0.981012 | 0.697606 |
| 35.667 | 0.192176 | 0.504078 | 1.19862 | -0.070865 | -0.30433 | 0.115112 | 0.665134 | 0.505794 | 0.620808 | 2.059487 | -2.834649 | -0.353549 | 4.151777 | -4.073984 | -0.468872 | 0.147121 | -0.130127 | 1.042937 | 0.160863 | 0.14568 | 0.006324 | 0.141101 | -1.518553 | 0.13689 | | 35.76389 | 4.151777 | -0.377362 |
| 35.681 | 0.361765 | 0.283589 | -0.026102 | 0.19776 | 0.570138 | 1.000455 | 0.420436 | 1.457434 | 0.527746 | 3.621161 | -2.077896 | 0.922159 | 8.702984 | -2.425266 | 1.089825 | 0.106649 | -0.211166 | 0.796087 | 0.107827 | 0.098073 | -0.9961 | 0.091472 | -1.919098 | -0.75357 | | 35.77778 | 8.702984 | -0.780728 |
| 35.694 | 0.632282 | 1.523468 | -1.210188 | 0.33934 | 1.145036 | 1.027442 | 1.159255 | 0.547074 | 0.382966 | 2.154781 | -1.300804 | -0.740256 | 6.912714 | -1.171418 | 0.070341 | 0.088485 | -0.259742 | 0.489004 | 0.081393 | 0.087231 | -2.149481 | 0.072476 | -2.171185 | -0.951214 | | 35.79167 | 6.912714 | -0.607961 |
| 35.708 | -0.376105 | 2.407575 | -1.499284 | 0.204143 | 0.354732 | 0.833021 | 2.542814 | 0.644097 | -0.08776 | 2.462496 | -0.169458 | -1.072897 | 5.130351 | 0.367562 | 0.154215 | 0.082741 | -0.210866 | 0.280864 | 0.080752 | 0.088848 | -2.343902 | 0.083244 | -0.761239 | -1.193267 | | 35.80556 | 5.130351 | -0.995536 |
| 35.722 | -0.191808 | 1.084169 | -1.787924 | -0.150364 | -1.328823 | 0.554067 | 3.29879 | 0.391684 | -1.011245 | 1.457194 | 0.573493 | 0.614908 | 3.066139 | 0.051969 | 1.435934 | 0.069498 | -0.567828 | 0.148091 | 0.075628 | 0.071801 | -1.235259 | 0.083219 | 0.433666 | -2.077424 | | 35.81944 | 3.066139 | -2.047933 |
| 35.736 | -0.398325 | -0.329451 | -3.0446 | -0.389179 | 0.156142 | 0.185716 | 3.417535 | 0.351356 | -1.057282 | -0.215959 | 0.52188 | 1.100151 | -1.335535 | -1.041102 | 2.414884 | 0.051099 | -1.47967 | 0.057051 | 0.06227 | 0.054252 | -0.285983 | 0.064801 | -0.982252 | -1.461786 | | 35.83333 | -1.335535 | -1.442383 |
| 35.75 | -0.162737 | -0.621473 | -2.403001 | 0.49299 | -0.1891 | -0.500328 | 1.705111 | 0.07795 | -1.53446 | -0.633472 | -0.646822 | 0.81515 | -3.026438 | -2.741917 | 2.599412 | 0.05938 | -1.313646 | -0.010942 | 0.07028 | 0.068227 | 0.637466 | 0.06775 | -1.071482 | -0.462541 | | 35.84722 | -3.026438 | -0.276741 |
| 35.764 | 2.021121 | -0.51666 | -1.129797 | -0.304324 | -0.236763 | -1.259725 | 0.334508 | -0.297516 | -1.495211 | -0.060216 | -2.21757 | 0.391234 | -0.899108 | -4.447394 | 1.915737 | 0.089118 | -0.785845 | -0.071312 | 0.097595 | 0.097639 | 1.077815 | 0.103538 | 0.054362 | 0.06519 | | 35.86111 | -0.899108 | 0.337158 |
| 35.778 | 4.193788 | -0.734724 | -0.803602 | -0.358951 | -0.643913 | -2.190363 | -0.111521 | -0.441159 | 0.365396 | 0.402459 | -4.190593 | 0.11795 | -0.999049 | -6.064892 | 1.522099 | 0.089732 | -0.909983 | -0.166239 | 0.087684 | 0.074748 | 0.272274 | 0.101978 | 0.7167 | -0.126112 | | 35.875 | -0.999049 | 0.142657 |
| 35.792 | 4.42461 | -0.910093 | -0.7286 | 0.337275 | -1.175092 | -2.60697 | -2.640329 | -0.30202 | 1.215086 | -0.100677 | -4.838122 | 0.198255 | -2.811194 | -6.008213 | 1.111851 | 0.072408 | -1.27426 | -0.260391 | 0.066199 | 0.040587 | -1.182636 | 0.068063 | 1.138519 | -0.828793 | | 35.88889 | -2.811194 | -0.378108 |
| 35.806 | 0.813016 | -1.582236 | -1.005727 | 0.232645 | -1.919795 | -2.553835 | -2.218693 | 0.13915 | 1.136457 | 0.0048 | -3.729137 | -0.989409 | -2.388204 | -5.292952 | 0.560066 | 0.108277 | -1.556247 | -0.322509 | 0.110725 | 0.086424 | -2.284797 | 0.092643 | 1.186596 | -1.482375 | | 35.90278 | -2.388204 | -1.02265 |
| 35.819 | -0.40316 | -1.223772 | -1.829268 | -0.049545 | -3.525341 | -1.795711 | -1.097181 | 0.616998 | 0.437544 | -1.02882 | -1.534971 | -1.429152 | -1.745911 | -4.413725 | 0.517335 | 0.172652 | -1.89977 | -0.337503 | 0.18986 | 0.167671 | -3.493786 | 0.165704 | 1.283835 | -1.482392 | | 35.91667 | -1.745911 | -1.320743 |
| 35.833 | -0.94276 | 0.087076 | -2.602441 | 0.033157 | -2.374863 | -2.687731 | 0.313992 | 0.528001 | -0.586822 | -1.683785 | -0.494059 | -2.135279 | -3.201376 | -3.71779 | -0.169262 | 0.203777 | -2.322271 | -0.375984 | 0.230134 | 0.201396 | -3.320789 | 0.212758 | 1.314679 | -1.47894 | | 35.93056 | -3.201376 | -1.367417 |
| 35.847 | -1.654707 | 0.083548 | -2.621616 | 0.057092 | -0.90573 | -2.441501 | 1.209873 | -0.170788 | -0.963391 | 0.179067 | -1.350407 | -2.179714 | -5.107654 | -2.909516 | -0.079923 | 0.212922 | -2.673946 | -0.391522 | 0.237224 | 0.201513 | -3.070573 | 0.23265 | 1.717699 | -1.166327 | | 35.94444 | -5.107654 | -1.065911 |
| 35.861 | -3.088038 | 0.867997 | -1.041231 | -0.004155 | -0.459363 | -1.395992 | 0.822398 | -0.653077 | -1.56306 | -0.068242 | -1.91673 | -1.321732 | -4.407056 | -1.670515 | -0.369336 | 0.217099 | -2.955481 | -0.394583 | 0.244342 | 0.201649 | -1.754409 | 0.245166 | 1.759041 | -0.615653 | | 35.95833 | -4.407056 | -0.610862 |
| 35.875 | -4.479647 | -2.837665 | 0.192359 | -0.217551 | -0.314491 | -0.835639 | -0.538884 | -1.083318 | -1.28671 | 0.01558 | -2.467043 | -0.307287 | -3.427227 | -0.30165 | -1.10623 | 0.196333 | -3.238871 | -0.449923 | 0.239654 | 0.198223 | -0.840889 | 0.236518 | 0.650569 | -0.315366 | | 35.97222 | -3.427227 | -0.337682 |
| 35.889 | -5.435213 | -5.119394 | 0.825843 | -0.571388 | -0.539175 | -0.60051 | -0.159089 | -2.405004 | -1.985326 | -0.716409 | -3.593617 | -2.286201 | -4.435693 | 0.758412 | -1.521696 | 0.172361 | -3.495484 | -0.49149 | 0.23273 | 0.197631 | -0.286628 | 0.225152 | -0.779224 | -0.43536 | | 35.98611 | -4.435693 | -0.580524 |
| 35.903 | -4.372957 | -4.833813 | 1.109979 | -0.662609 | -1.111854 | -0.95322 | 1.257689 | -3.647345 | -1.96998 | 0.193841 | -4.294112 | -3.82482 | -6.413153 | 1.545805 | -1.881052 | 0.069809 | -3.448335 | -0.509459 | 0.23928 | 0.207249 | 0.025823 | 0.233793 | -0.909398 | -0.704843 | | 36 | -6.413153 | -0.986125 |
| 35.917 | -1.046125 | -1.95238 | 0.908471 | -0.29233 | -2.235775 | -2.18871 | 2.050074 | -3.338958 | -1.106058 | 0.916467 | -3.361496 | -5.422165 | -5.947344 | 1.97655 | -1.888584 | -0.049102 | -2.871436 | -0.424249 | 0.330464 | 0.272962 | -0.007531 | 0.313983 | -0.924288 | -0.70344 | | 36.01389 | -5.947344 | -0.944608 |
| 35.931 | 3.406457 | 0.210609 | 0.457043 | 0.367332 | -2.839567 | -3.204259 | 2.525114 | -2.640665 | -1.308173 | 0.509861 | -1.997118 | -4.741815 | -3.363916 | 1.921331 | -1.505453 | -0.008631 | -2.018017 | -0.237692 | 0.546315 | 0.425993 | -0.08422 | 0.496564 | -0.883389 | -0.577997 | | 36.02778 | -3.363916 | -0.722977 |
| 35.944 | 4.454807 | -0.943293 | -0.090026 | 0.734187 | -2.835887 | -3.926069 | 1.821407 | 0.103029 | 0.739238 | -1.144367 | -1.077631 | -6.300673 | -2.00956 | 1.120782 | -1.316868 | -0.027408 | -1.429026 | -0.172246 | 0.52888 | 0.389556 | -0.32888 | 0.485029 | -1.205244 | -0.639075 | | 36.04167 | -2.00956 | -0.83719 |
| 35.958 | 3.554837 | -4.651425 | -0.346194 | 0.391579 | -3.221554 | -3.970897 | 0.2187 | -0.770968 | 0.018042 | -1.530689 | -1.404529 | -6.905151 | -4.07534 | 0.295239 | -1.317316 | -0.227088 | -1.640537 | -0.253044 | 0.139099 | 0.1036 | -0.860265 | 0.104567 | -1.918082 | -1.250555 | | 36.05556 | -4.07534 | -1.30308 |
| 35.972 | 2.922773 | -6.030845 | -0.381331 | -0.387824 | -5.905182 | -4.106409 | -0.175365 | -3.883994 | -2.602786 | -4.304964 | -2.932602 | -6.776661 | -8.226958 | 0.359885 | -1.284794 | -0.398254 | -2.667765 | 0.026289 | 0.109133 | 0.136659 | -0.981824 | 0.045359 | -2.658744 | -1.517653 | | 36.06944 | -8.226958 | -1.510513 |
| 35.986 | -1.693387 | -4.792558 | -0.808532 | -1.052152 | -6.168336 | -2.748648 | -0.257002 | -7.016095 | -5.751789 | -5.52018 | -5.353574 | -5.175716 | -11.334502 | 1.083388 | -0.880855 | -0.390088 | -2.171069 | 0.582761 | 0.461585 | 0.466078 | -0.337565 | 0.406625 | -3.497636 | -1.008443 | | 36.08333 | -11.334502 | -1.079203 |
| 36 | -4.417354 | -3.565542 | -3.027586 | -0.752256 | -4.900791 | -1.100588 | -0.288875 | -7.411139 | -5.841656 | -5.216777 | -6.451445 | -2.570449 | -10.536048 | 1.52858 | -0.325916 | 0.382705 | -1.024687 | 0.783667 | 0.597838 | 0.576766 | 0.397506 | 0.570854 | -2.251906 | -0.381601 | | 36.09722 | -10.536048 | -0.582266 |
| 36.014 | -4.543342 | -2.281024 | -2.719856 | -0.348055 | -3.410642 | -0.387211 | 0.052051 | -6.20869 | -3.915051 | -2.944889 | -3.876007 | -2.086262 | -7.077455 | 1.534807 | -0.77409 | 0.408134 | -0.065216 | 0.628289 | 0.469419 | 0.442493 | 0.493657 | 0.451712 | -1.133489 | -0.09218 | | 36.11111 | -7.077455 | -0.349716 |
| 36.028 | -2.489814 | -1.622326 | -1.413956 | -0.055491 | -2.126636 | 0.512642 | 0.601461 | -4.793731 | -2.417531 | -1.367591 | -1.583163 | -2.492643 | -3.887991 | 1.668306 | -1.026365 | 0.254924 | 0.180287 | 0.571342 | 0.294873 | 0.27049 | 0.32597 | 0.279196 | -0.267838 | 0.202077 | | 36.125 | -3.887991 | 0.057758 |
| 36.042 | -0.41221 | -1.553465 | -0.454149 | 0.040039 | -1.321059 | 0.967706 | 0.497845 | -3.053456 | -1.866087 | -0.442337 | -0.098282 | -2.058267 | -1.794773 | 1.611326 | -0.714436 | 0.075563 | 0.318145 | 0.595343 | 0.144424 | 0.135954 | 0.206795 | 0.133374 | -0.348207 | 0.404653 | | 36.13889 | -1.794773 | 0.42215 |
| 36.056 | 1.357617 | -0.856789 | 0.235057 | 0.110331 | -0.725009 | 0.767994 | 0.370739 | -2.451348 | -1.474713 | 0.282416 | 1.337837 | -1.11546 | 0.069549 | 1.039556 | -0.040968 | -0.01809 | 0.615855 | 0.402243 | 0.049394 | 0.051876 | 0.186695 | 0.045201 | -1.120928 | 0.352396 | | 36.15278 | 0.069549 | 0.406788 |
| 36.069 | 2.587892 | 1.145895 | 0.584117 | 0.200811 | 0.179183 | 0.305748 | 0.217898 | -1.401099 | -0.83306 | 1.014067 | 2.553178 | 0.641435 | 1.903588 | -0.269685 | -0.032701 | -0.058132 | 1.014693 | 0.09641 | 0.012878 | 0.003908 | 0.150821 | 0.009185 | -0.157406 | -0.047681 | | 36.16667 | 1.903588 | -0.032363 |
| 36.083 | 3.325575 | 3.829778 | 0.972255 | 0.262205 | 1.452848 | -0.503582 | 0.526616 | -0.713162 | -0.086255 | 1.642559 | 3.105348 | 2.54177 | 2.937123 | -2.26759 | -0.807623 | -0.03007 | 1.161346 | -0.13868 | -0.017785 | -0.041864 | 0.038951 | -0.028845 | 0.301557 | -0.594161 | | 36.18056 | 2.937123 | -0.573954 |
| 36.097 | 2.676962 | 3.819088 | 1.344368 | 0.096928 | 3.296575 | -1.209854 | 0.343636 | -0.408652 | 0.45238 | 2.712635 | 2.293823 | 4.193634 | 2.890027 | -3.823448 | -0.63771 | 0.054145 | 0.901568 | -0.292303 | -0.055538 | -0.080718 | 0.034756 | -0.071036 | 0.208136 | -0.604385 | | 36.19444 | 2.890027 | -0.508626 |
| 36.111 | 1.384955 | 2.779694 | 1.406146 | 0.355432 | 4.596481 | -1.498428 | -0.069561 | 0.034202 | 0.784226 | 3.193638 | 1.057585 | 4.551527 | 1.884684 | -3.384192 | -0.93959 | 0.104288 | 0.6871 | -0.390081 | -0.075554 | -0.08863 | 0.426331 | -0.079875 | -0.857724 | -0.100683 | | 36.20833 | 1.884684 | 0.14214 |
| 36.125 | -0.175522 | 0.289161 | 0.946564 | 0.573735 | 3.859411 | -0.765457 | -0.009175 | 0.998132 | 0.858504 | 2.724527 | -0.11864 | 2.613487 | 0.188244 | -1.829312 | -1.107335 | 0.200648 | 1.056192 | -0.44247 | -0.069034 | -0.070085 | 1.149946 | -0.059565 | -0.926188 | 0.627684 | | 36.22222 | 0.188244 | 0.809671 |
| 36.139 | -1.389149 | -1.23369 | 0.450908 | 0.767067 | 1.877653 | 0.074557 | 0.590342 | 0.597771 | 0.755366 | 1.586528 | -0.178676 | 0.300138 | -0.73447 | -0.475568 | -0.836942 | 0.266109 | 1.642479 | -0.46656 | -0.050676 | -0.050407 | 1.466369 | -0.036945 | -0.517924 | 1.109834 | | 36.23611 | -0.73447 | 1.16033 |
| 36.153 | 0.434103 | -1.945713 | 0.176848 | 0.165513 | -0.254799 | 0.169767 | 0.391425 | 0.498296 | 0.568444 | 0.385566 | 0.129194 | -2.040561 | -0.23504 | 0.273843 | -0.441619 | 0.157457 | 2.078531 | -0.444873 | -0.036249 | -0.038618 | 1.435167 | -0.025377 | -0.300512 | 0.944029 | | 36.25 | -0.23504 | 0.994157 |
| 36.167 | 2.269295 | -2.864672 | -0.0307 | -0.458809 | -0.905143 | 0.421497 | 0.041943 | 0.364206 | 0.41198 | -0.230054 | 0.572317 | -2.195102 | -0.225467 | -0.04013 | -0.371315 | 0.058301 | 1.675751 | -0.390458 | -0.034858 | -0.038843 | 1.201311 | -0.031569 | -0.341143 | 0.440314 | | 36.26389 | -0.225467 | 0.506611 |
| 36.181 | 3.038868 | -0.470212 | -0.037487 | -0.591637 | -0.038865 | 0.829913 | -1.269771 | 0.360115 | 0.605413 | -0.0737 | 0.929846 | -0.544502 | 0.205851 | -1.14475 | -0.290996 | -0.069645 | 0.780675 | -0.290169 | -0.046224 | -0.048782 | 0.983029 | -0.045438 | -0.034403 | -0.040388 | | 36.27778 | 0.205851 | 0.029119 |
| 36.194 | 3.447607 | 2.10166 | 0.057605 | -0.904476 | 0.482422 | 2.300399 | -1.369764 | 0.339472 | 1.188656 | 0.668487 | 1.215662 | 0.906952 | 1.226739 | -1.423579 | -0.649394 | -0.096696 | 0.155846 | -0.23682 | -0.048409 | -0.043305 | 0.868928 | -0.041806 | 0.259168 | -0.380942 | | 36.29167 | 1.226739 | -0.3585 |
| 36.208 | 2.546799 | 3.984357 | 0.17036 | -0.367768 | 1.00698 | 2.906639 | -0.365148 | 0.619547 | 2.302986 | 1.601907 | 1.124757 | 2.35032 | 1.85633 | -0.413251 | -3.221761 | -0.108365 | -0.02156 | -0.370356 | -0.040974 | -0.035312 | 0.818129 | -0.040618 | 0.468709 | -0.447772 | | 36.30556 | 1.85633 | -0.51668 |
| 36.222 | 0.820017 | 3.746738 | 0.428499 | 0.441611 | 1.452561 | 2.077419 | 0.355613 | 1.125381 | 3.035473 | 1.961134 | 1.127098 | 3.555676 | 2.291362 | 1.404677 | -3.125765 | -0.141225 | 0.588338 | -0.544243 | -0.062442 | -0.052541 | 0.889039 | -0.068745 | 0.731463 | -0.180192 | | 36.31944 | 2.291362 | -0.374971 |
| 36.236 | -0.262568 | 2.106143 | 0.959581 | 0.637659 | 2.126716 | 2.444641 | 1.035214 | 2.032876 | 3.727824 | 1.93079 | 1.536109 | 4.039404 | 3.017121 | 2.589046 | 0.220519 | -0.122478 | 1.80948 | -0.583957 | -0.090097 | -0.069843 | 0.98271 | -0.086756 | 0.939509 | 0.261137 | | 36.33333 | 3.017121 | 0.063658 |
| 36.25 | -0.720044 | 1.418293 | 1.476962 | 0.84895 | 2.527509 | 3.776237 | 1.765893 | 2.085621 | 3.073767 | 1.637662 | 1.981576 | 4.077506 | 3.928246 | 2.789676 | 2.049564 | -0.080635 | 2.197649 | -0.485239 | -0.093905 | -0.070546 | 1.066678 | -0.078538 | 1.089051 | 0.776797 | | 36.34722 | 3.928246 | 0.647788 |
| 36.264 | -0.80419 | 1.422816 | 1.333867 | 0.860521 | 2.474334 | 3.458157 | 2.317798 | 1.780501 | 2.127106 | 1.483409 | 2.129547 | 3.132564 | 4.338884 | 2.163765 | 2.444724 | -0.10338 | 2.154795 | -0.39767 | -0.095331 | -0.075215 | 0.95491 | -0.077343 | 1.090874 | 1.160825 | | 36.36111 | 4.338884 | 1.112045 |
| 36.278 | -0.555403 | 1.251432 | 0.971371 | 0.526956 | 2.186841 | 2.027913 | 1.877146 | 1.680468 | 1.737364 | 1.681611 | 1.86055 | 1.742559 | 3.804109 | 1.471438 | 2.086443 | -0.172704 | 2.005777 | -0.288235 | -0.09795 | -0.075884 | 0.70591 | -0.08466 | 1.076945 | 1.259368 | | 36.375 | 3.804109 | 1.377886 |
| 36.292 | 0.217089 | 0.862974 | 0.756322 | 0.408705 | 1.743712 | 1.652105 | 1.008801 | 1.526331 | 1.773969 | 2.203712 | 1.780532 | 0.576064 | 2.848176 | 0.672161 | 1.023925 | -0.14045 | 1.109113 | -0.060951 | -0.082377 | -0.046878 | 0.523926 | -0.059541 | 1.097567 | 1.12094 | | 36.38889 | 2.848176 | 1.44566 |
| 36.306 | 1.218067 | 0.698736 | 0.630607 | 0.422581 | 1.924476 | 1.539559 | 0.893954 | 1.594335 | 2.015562 | 2.942668 | 1.903928 | 1.097574 | 2.0495 | 0.466803 | 2.223378 | -0.041207 | 0.85193 | 0.287238 | -0.051383 | 0.008033 | 0.524292 | 0.004254 | 1.170534 | 1.045946 | | 36.40278 | 2.0495 | 1.437467 |
| 36.319 | 1.379541 | 0.323848 | 0.632719 | 0.799301 | 2.448104 | 1.585203 | 0.583708 | 1.585198 | 2.061099 | 3.007004 | 2.415451 | 2.251189 | 1.792751 | 1.364451 | 2.396335 | 0.050021 | 1.278353 | 0.653214 | -0.01143 | 0.04644 | 0.734722 | 0.056944 | 1.235262 | 1.24177 | | 36.41667 | 1.792751 | 1.532715 |
| 36.333 | 0.476252 | 0.291921 | 0.698196 | 0.889298 | 2.154949 | 2.146496 | -0.056599 | 1.393013 | 2.098455 | 2.368517 | 3.162842 | 3.97997 | 1.985246 | 2.591555 | 1.811293 | 0.097129 | 2.652253 | 0.994205 | 0.024516 | 0.048967 | 1.013793 | 0.069073 | 1.296455 | 1.672669 | | 36.43056 | 1.985246 | 1.798097 |
| 36.347 | -0.61743 | 0.252263 | 0.851718 | 0.806404 | 1.65363 | 3.176696 | -0.316135 | 1.54287 | 2.172476 | 1.962603 | 3.759263 | 5.858541 | 2.330947 | 3.747516 | 1.358565 | 0.061343 | 3.020523 | 1.222595 | -0.005985 | 0.007662 | 1.148801 | 0.017666 | 1.434003 | 1.931021 | | 36.44444 | 2.330947 | 1.960498 |
| 36.361 | -1.144393 | -0.163347 | 0.598588 | 0.097881 | 1.656779 | 3.577441 | -0.009668 | 1.912035 | 2.297331 | 1.670823 | 3.187767 | 5.811358 | 2.446674 | 4.556028 | -0.196223 | -0.111949 | 2.669898 | 1.089381 | -0.200245 | -0.1675 | 1.14855 | -0.214793 | 1.704342 | 1.77199 | | 36.45833 | 2.446674 | 1.81655 |
| 36.375 | -0.847914 | -0.469422 | -0.002484 | -0.559032 | 1.457862 | 2.920541 | 0.481087 | 2.493293 | 2.248023 | 1.734642 | 2.230748 | 4.315751 | 1.986655 | 5.158828 | -0.597571 | -0.391922 | 2.461242 | 0.78266 | -0.56142 | -0.440443 | 0.872905 | -0.544087 | 2.000862 | 1.555666 | | 36.47222 | 1.986655 | 1.634212 |
| 36.389 | -0.414793 | -0.24008 | -0.048573 | 0.044739 | 1.695545 | 1.654938 | 1.193977 | 3.197983 | 2.320748 | 2.184183 | 1.552635 | 3.145386 | 1.943755 | 6.118854 | 1.543273 | -0.464407 | 3.037859 | 0.490808 | -0.573329 | -0.439759 | 0.481766 | -0.500828 | 2.255394 | 1.664723 | | 36.48611 | 1.943755 | 1.840284 |
| 36.403 | -0.082817 | 0.480117 | 0.087557 | 0.934874 | 2.729564 | 0.840132 | 2.154749 | 4.37426 | 2.639089 | 2.616775 | 3.437389 | 2.738825 | 2.664885 | 7.273056 | 3.807735 | -0.308461 | 2.299611 | 0.473549 | -0.292366 | -0.253603 | 0.405956 | -0.26689 | 2.53196 | 1.815788 | | 36.5 | 2.664885 | 2.026464 |
| 36.417 | 1.023087 | 1.161852 | -0.101215 | 1.27703 | 3.528826 | 0.609672 | 1.550581 | 6.297872 | 3.182655 | 2.719464 | 5.721598 | 2.668232 | 3.882676 | 7.558265 | 4.93302 | -0.207767 | 1.288535 | 0.511077 | -0.145177 | -0.147207 | 0.493044 | -0.144945 | 2.425109 | 1.700404 | | 36.51389 | 3.882676 | 1.991004 |
| 36.431 | 2.425914 | 1.508082 | -0.078961 | 1.366972 | 4.131529 | 1.920143 | 0.670146 | 6.210681 | 4.271854 | 2.420712 | 6.882807 | 4.489141 | 5.282666 | 6.114597 | 4.448371 | -0.245335 | 1.689647 | 0.44348 | -0.181499 | -0.198695 | 0.891199 | -0.176265 | 1.999971 | 1.462752 | | 36.52778 | 5.282666 | 1.548615 |
| 36.444 | 4.075883 | 1.127311 | 0.014455 | 1.0798 | 4.497125 | 4.257776 | 0.163598 | 4.707508 | 4.671965 | 3.028116 | 6.327293 | 5.691837 | 6.693816 | 3.305308 | 2.708599 | -0.568098 | 1.393679 | 0.01301 | -0.499987 | -0.549028 | 1.385515 | -0.452055 | 1.383008 | 1.130434 | | 36.54167 | 6.693816 | 0.977865 |
| 36.458 | 5.474317 | 0.892943 | 0.103192 | 0.832817 | 3.546336 | 4.382624 | -0.361995 | 0.055623 | 4.21459 | 3.739717 | 5.658903 | 4.718726 | 7.141082 | 1.093172 | 0.54085 | -1.218428 | -0.073751 | -0.791848 | -1.138867 | -1.216833 | 1.313575 | -1.023986 | 0.526249 | 0.588342 | | 36.55556 | 7.141082 | 0.155448 |
| 36.472 | 3.221268 | 0.680291 | 0.8734 | 0.807207 | 1.738088 | 3.393417 | -1.072551 | 0.007737 | 4.143513 | 2.61044 | 2.911185 | 3.136682 | 4.285964 | -0.054477 | -0.592497 | -1.577356 | -1.538014 | -1.264739 | -1.608391 | -1.644153 | 0.477649 | -1.478101 | -0.102024 | -0.648375 | | 36.56944 | 4.285964 | -1.072776 |
| 36.486 | -0.89503 | -0.143165 | 1.875602 | 1.11098 | 1.139437 | 0.512399 | -2.017512 | -0.458568 | 2.513381 | 3.019182 | 1.60935 | 0.211778 | 1.852671 | 0.59016 | -0.525569 | -1.236479 | -1.263566 | -0.983811 | -1.367829 | -1.292138 | 0.215517 | -1.327503 | 0.135131 | -1.07749 | | 36.58333 | 1.852671 | -1.176523 |
| 36.5 | -3.84008 | -0.367555 | 2.772677 | 1.598845 | 0.509579 | -0.1245 | -0.537216 | 1.932467 | 2.492714 | 2.587656 | 1.555309 | 0.063119 | 3.030126 | 1.603431 | 0.525723 | -0.672729 | -0.539623 | -0.488491 | -0.758712 | -0.69213 | 0.952362 | -0.776592 | 0.837597 | -0.462648 | | 36.59722 | 3.030126 | -0.589132 |
| 36.514 | -1.838014 | -0.063943 | 2.51349 | 2.011401 | 1.241242 | 1.030171 | 1.450432 | 4.553534 | 2.522322 | 1.434323 | 1.728919 | 1.780901 | 7.205159 | 1.993916 | 1.267765 | -0.288974 | 0.210246 | -0.194731 | -0.314868 | -0.291046 | 1.87996 | -0.341171 | 1.349787 | 0.315803 | | 36.61111 | 7.205159 | 0.144331 |
| 36.528 | 3.063573 | 0.374324 | 2.015323 | 1.593969 | 2.08047 | 2.119536 | 2.310041 | 3.125416 | 0.913531 | 1.476229 | 2.508715 | 3.34642 | 9.376131 | 1.805869 | 0.911442 | -0.08229 | 0.839202 | -0.058608 | -0.071272 | -0.068295 | 3.076948 | -0.084627 | 1.122613 | 0.541087 | | 36.625 | 9.376131 | 0.250264 |
| 36.542 | 6.253875 | 1.407982 | 1.938559 | 1.272518 | 2.319875 | 1.42714 | 4.654534 | 1.746741 | -0.348452 | -0.079959 | 0.393483 | 1.324464 | 6.871989 | 0.941465 | -1.474817 | 0.041734 | 0.896288 | 0.017585 | 0.080952 | 0.073735 | 2.233688 | 0.088764 | 0.302607 | -0.024568 | | 36.63889 | 6.871989 | -0.403223 |
| 36.556 | 3.954245 | 1.430489 | 1.284223 | 1.589341 | 1.150007 | 0.748566 | 0.438295 | -0.263301 | -1.005759 | -0.159835 | -2.4521 | -0.934611 | -0.892081 | -0.628181 | -2.734033 | 0.098347 | 0.578889 | 0.051299 | 0.157517 | 0.145916 | 0.114154 | 0.163676 | 0.001372 | -0.630008 | | 36.65278 | -0.892081 | -0.924148 |
| 36.569 | 2.515146 | 2.460474 | 0.954017 | 1.253503 | -0.258676 | 0.658196 | -2.351989 | -2.896269 | -1.220168 | 0.894379 | -2.913932 | -1.710299 | -5.572529 | -2.598182 | -1.923445 | 0.153543 | 0.369269 | 0.104366 | 0.228966 | 0.225589 | -1.193649 | 0.212217 | 0.266777 | -0.886443 | | 36.66667 | -5.572529 | -0.97093 |
| 36.583 | -0.858699 | 2.131989 | 0.747346 | 1.815085 | 0.310744 | 0.008417 | -4.59457 | -0.12922 | 0.274423 | 0.449745 | -1.569253 | -0.956856 | -2.94257 | -3.919177 | -0.243009 | 0.280819 | 0.222015 | 0.193598 | 0.330884 | 0.347215 | -1.345368 | 0.308674 | 0.87777 | -0.934873 | | 36.68056 | -2.94257 | -0.660999 |
| 36.597 | -0.11404 | -0.062793 | 0.520001 | 1.289865 | 0.240243 | -0.296624 | -2.748124 | 3.185207 | 1.548186 | -0.779424 | -0.021952 | -0.333131 | 3.563379 | -3.276291 | 1.175857 | 0.359051 | -0.686641 | 0.212176 | 0.371153 | 0.393296 | -0.13263 | 0.357064 | 1.299931 | -0.28509 | | 36.69444 | 3.563379 | -0.09206 |
| 36.611 | -4.901746 | -3.265366 | 0.386062 | 0.551012 | 1.365718 | 0.173201 | 1.284496 | 1.923694 | 1.391403 | -1.837189 | 1.776673 | 1.751989 | 8.525373 | -0.664925 | 1.671042 | 0.326222 | 0.122184 | 0.133385 | 0.324381 | 0.32631 | 1.263554 | 0.309799 | 1.117109 | 0.437748 | | 36.70833 | 8.525373 | 0.467319 |
| 36.625 | -5.849791 | -2.060075 | 0.545236 | 0.599116 | 2.107849 | 0.99702 | 3.766405 | 0.494229 | 0.626731 | -0.855066 | 1.776383 | 3.095557 | 8.472821 | 3.098101 | 2.107151 | 0.255913 | -0.318745 | 0.030918 | 0.26786 | 0.240924 | 1.935948 | 0.246062 | 1.079098 | 0.505944 | | 36.72222 | 8.472821 | 0.730508 |
| 36.639 | -2.767716 | 0.312398 | 0.335098 | -0.411462 | 2.910764 | 0.848813 | 1.636825 | -0.279834 | -0.336942 | -0.838173 | 3.149154 | 0.649213 | 3.815954 | 5.03997 | 1.336433 | 0.225268 | -0.569922 | -0.047143 | 0.25565 | 0.205312 | 2.135728 | 0.225173 | 0.383119 | 0.426643 | | 36.73611 | 3.815954 | 0.487044 |
| 36.653 | 1.432138 | 2.555484 | -0.029122 | 0.032643 | 0.686786 | 0.77295 | 0.840179 | -0.192978 | -0.617422 | -1.979246 | 2.113366 | -0.668559 | 4.151907 | 4.830341 | -0.0566 | 0.239471 | 0.059061 | -0.07374 | 0.290777 | 0.214622 | 1.15567 | 0.254452 | -2.103438 | -0.020507 | | 36.75 | 4.151907 | -0.291087 |
| 36.667 | 4.048485 | 1.766804 | -1.571028 | -0.08835 | -0.458955 | 0.854998 | 1.315944 | 0.22373 | -0.45606 | -1.313459 | 0.431741 | -0.957585 | 4.314929 | -0.120184 | -1.072724 | 0.261803 | 0.371135 | -0.020015 | 0.320062 | 0.227817 | -0.10623 | 0.289459 | -2.954434 | -0.353563 | | 36.76389 | 4.314929 | -0.837717 |
| 36.681 | 1.326327 | 2.019529 | -1.128754 | -0.259511 | -0.359972 | 0.09626 | 2.486448 | 0.382277 | -0.515853 | 0.232919 | -1.003031 | -0.56089 | 3.375996 | -5.695938 | -0.386139 | 0.255252 | 0.650534 | -0.03149 | 0.290592 | 0.209758 | -1.608752 | 0.269716 | -2.877517 | -0.453982 | | 36.77778 | 3.375996 | -0.904434 |
| 36.694 | 1.115509 | 1.994396 | 0.23024 | 0.373048 | -0.473751 | -0.167871 | 0.819078 | 0.216515 | 0.03303 | 0.958878 | -1.903431 | 1.061657 | 2.480268 | -4.950632 | -0.130017 | 0.213373 | 0.625998 | -0.05568 | 0.247152 | 0.216532 | -1.629253 | 0.233308 | -1.853851 | -0.638594 | | 36.79167 | 2.480268 | -0.548165 |
| 36.708 | 1.979724 | 2.018583 | 0.62482 | -0.267406 | -0.737356 | -0.265403 | -0.53407 | -0.254276 | 0.264735 | 0.777263 | -2.486482 | 1.831223 | 0.802426 | -3.866198 | 0.016501 | 0.182531 | -0.293618 | -0.086077 | 0.229554 | 0.251225 | -0.998192 | 0.222717 | -0.712708 | -0.755989 | | 36.80556 | 0.802426 | -0.231267 |
| 36.722 | 0.733943 | 1.800809 | 0.092795 | -0.422326 | -0.819521 | -0.185771 | 1.273105 | -0.763429 | 0.392588 | 1.837205 | -3.321234 | -0.783491 | -1.00944 | -4.570332 | 0.394222 | 0.175858 | -0.405955 | -0.109843 | 0.227674 | 0.261918 | -0.31687 | 0.221551 | 0.019073 | -0.306289 | | 36.81944 | -1.00944 | 0.147069 |
| 36.736 | -2.372267 | 1.170042 | -0.706449 | 0.027759 | -1.340196 | -0.383892 | 3.68444 | -1.02913 | 0.43935 | 0.169497 | -2.609231 | -1.381478 | -0.340783 | -5.50669 | 0.557745 | 0.195949 | -0.77991 | -0.11121 | 0.246235 | 0.264048 | 1.803949 | 0.233528 | -0.093647 | 0.091321 | | 36.83333 | -0.340783 | 0.295202 |
| 36.75 | -5.599361 | 0.561701 | -0.9388 | 0.113405 | -1.203761 | -0.752204 | 5.865202 | -0.630159 | -0.184255 | -0.278574 | -0.206925 | -1.310424 | 2.29289 | -3.28171 | -0.199072 | 0.215814 | -1.447664 | -0.094202 | 0.252495 | 0.244183 | 3.322679 | 0.24145 | -0.394914 | 0.225259 | | 36.84722 | 2.29289 | 0.183571 |
| 36.764 | -1.002014 | 0.362302 | -1.781517 | 0.163671 | -0.656881 | -0.99765 | 7.514595 | -0.566773 | -0.591354 | -0.848739 | 0.326722 | -1.696671 | 2.614221 | 0.259155 | -0.895154 | 0.218822 | -1.327699 | -0.141442 | 0.220152 | 0.191744 | 2.645097 | 0.225504 | -0.586971 | 0.055148 | | 36.86111 | 2.614221 | -0.112379 |
| 36.778 | -0.077337 | 0.445095 | -2.462288 | 0.431278 | -0.564715 | -1.427821 | 6.24728 | 0.214407 | -0.701153 | 0.944166 | -0.877322 | -1.146061 | -0.114191 | 2.599988 | -1.58285 | 0.202429 | -1.036818 | -0.172793 | 0.179367 | 0.144824 | 0.741661 | 0.194563 | -1.013312 | -0.31328 | | 36.875 | -0.114191 | -0.558894 |
| 36.792 | -0.864334 | -1.897204 | -2.370054 | -0.209618 | -0.221933 | -1.617602 | 3.455257 | 1.025943 | -1.448074 | 0.319658 | -3.173653 | -0.117471 | -5.812615 | 3.207861 | -2.085527 | 0.188218 | -1.349529 | -0.205595 | 0.167618 | 0.122849 | -1.23372 | 0.172306 | -1.314309 | -0.860567 | | 36.88889 | -5.812615 | -1.137107 |
| 36.806 | 1.126501 | -1.171101 | -1.777289 | -0.701988 | -1.014886 | -0.948173 | 0.26595 | 1.013354 | -0.859251 | -1.610294 | -4.909066 | 0.305011 | -5.985511 | 1.485074 | -2.006576 | 0.193498 | -2.513911 | -0.214097 | 0.189196 | 0.135115 | -3.594826 | 0.181743 | -1.366245 | -1.505826 | | 36.90278 | -5.985511 | -1.666616 |
| 36.819 | 1.797502 | -1.917976 | -1.187266 | -0.912054 | -1.707006 | 0.371301 | -1.477514 | -0.177647 | -0.509126 | -0.318267 | -5.547438 | 0.707043 | -3.332518 | -2.707442 | -1.538255 | 0.193662 | -3.453894 | -0.218653 | 0.181846 | 0.139103 | -4.550216 | 0.185708 | -1.213888 | -2.286289 | | 36.91667 | -3.332518 | -2.246171 |
| 36.833 | 1.83908 | -0.004938 | -0.775546 | -1.023715 | -0.308173 | 0.2344 | -4.592938 | -1.208638 | 0.725764 | 1.245583 | -7.596959 | -1.391859 | -3.113816 | -3.985013 | -1.043483 | 0.137604 | -4.095263 | -0.268685 | 0.110652 | 0.100094 | -4.170962 | 0.140609 | -1.183801 | -2.630297 | | 36.93056 | -3.113816 | -2.581219 |
| 36.847 | 4.433318 | 1.455004 | -0.881387 | -1.405324 | 0.951175 | -0.864728 | -4.81337 | -1.773393 | 1.132707 | 0.339033 | -6.290577 | -4.022305 | -3.636824 | -5.455968 | -0.832596 | 0.047217 | -4.618592 | -0.332251 | 0.025631 | 0.036255 | -3.303997 | 0.06389 | -1.146535 | -2.604235 | | 36.94444 | -3.636824 | -2.590191 |
| 36.861 | 5.817947 | 0.858905 | -1.27275 | -1.146701 | 1.71841 | -2.689853 | -3.058243 | -1.897633 | 1.096651 | 0.424181 | -3.437334 | -4.140361 | -1.704272 | -5.757984 | -0.683295 | -0.016775 | -4.855142 | -0.386352 | -0.042624 | -0.031678 | -1.841823 | -0.015835 | -0.908057 | -2.41006 | | 36.95833 | -1.704272 | -2.396249 |
| 36.875 | 3.530832 | -0.210853 | -1.821193 | -0.871931 | 2.251324 | -4.752587 | -1.367302 | -2.297499 | 0.384552 | -0.973694 | -0.585239 | -1.388821 | -0.516153 | -5.216486 | -0.794598 | -0.070965 | -4.556433 | -0.436843 | -0.089525 | -0.086074 | -1.127182 | -0.075704 | -0.705066 | -2.182469 | | 36.97222 | -0.516153 | -2.258375 |
| 36.889 | 2.322033 | -0.757743 | -2.129681 | -0.643869 | 2.131043 | -5.800867 | -1.003143 | -3.018067 | -0.879603 | -1.342354 | 1.071461 | 0.164243 | -1.336628 | -3.999397 | -0.973343 | -0.120562 | -3.826047 | -0.504884 | -0.109269 | -0.109116 | -0.76923 | -0.105165 | -0.662769 | -2.06909 | | 36.98611 | -1.336628 | -2.093684 |
| 36.903 | 0.24307 | -3.556025 | -1.352495 | 0.489206 | 0.038871 | -5.179919 | -0.864349 | -3.135481 | -2.154124 | -2.861622 | 0.658143 | 2.21265 | -4.147973 | -2.66667 | -1.303819 | -0.147554 | -2.924158 | -0.546246 | -0.104045 | -0.106405 | -0.653213 | -0.110027 | -0.566932 | -1.826903 | | 37 | -4.147973 | -1.836222 |
| 36.917 | -1.876687 | -4.889387 | -1.809062 | 1.848042 | -2.061599 | -4.97797 | 0.711926 | -2.679787 | -1.751589 | -3.961157 | 0.157516 | 2.390999 | -6.379034 | -1.688923 | -1.464505 | -0.057994 | -1.74927 | -0.436187 | 0.019093 | -0.015729 | -0.593603 | -0.003901 | -0.347922 | -1.418336 | | 37.01389 | -6.379034 | -1.451422 |
| 36.931 | -4.753925 | -6.225662 | -2.073811 | 1.727567 | -2.743011 | -3.65152 | 2.279902 | -1.830999 | -3.799066 | -3.072676 | 0.683701 | 1.074659 | -6.440447 | -1.248444 | -1.286775 | 0.137231 | -0.760927 | -0.172525 | 0.284872 | 0.178216 | -0.541448 | 0.234489 | -0.056206 | -0.965213 | | 37.02778 | -6.440447 | -0.938878 |
| 36.944 | -5.901399 | -4.184972 | -1.924291 | 0.88706 | -3.06403 | -2.824575 | 1.372241 | -1.426305 | -3.447639 | -2.520408 | -0.547166 | -1.246357 | -6.50247 | -1.611219 | -1.277177 | 0.138909 | -0.249697 | -0.183197 | 0.294587 | 0.171577 | -0.758318 | 0.260865 | -0.146073 | -0.832818 | | 37.04167 | -6.50247 | -0.858773 |
| 36.958 | -6.328708 | -1.947014 | -1.68022 | 0.098196 | -4.625534 | -2.413143 | -0.488962 | -1.778663 | -3.554346 | -2.554819 | -3.612783 | -3.308526 | -8.029363 | -2.195529 | -1.511066 | -0.194081 | -0.538582 | -0.538322 | -0.170429 | -0.146013 | -1.337165 | -0.161746 | -0.794028 | -1.485748 | | 37.05556 | -8.029363 | -1.479683 |
| 36.972 | -6.559134 | -1.943078 | -2.085949 | -0.7323 | -5.303761 | -3.10243 | -3.774884 | -3.040763 | -4.480171 | -4.095869 | -6.279774 | -6.472271 | -11.255969 | -2.233389 | -1.544739 | -0.303422 | -0.910096 | -0.715081 | -0.398568 | -0.300309 | -1.818507 | -0.433371 | -1.140316 | -2.084927 | | 37.06944 | -11.255969 | -2.04689 |
| 36.986 | -6.080441 | -2.740451 | -2.131261 | -2.099946 | -6.74628 | -3.80808 | -5.00792 | -4.924205 | -5.682136 | -4.951688 | -7.059055 | -5.406157 | -13.362284 | -1.687601 | -1.161016 | -0.087208 | -0.716041 | -0.602112 | -0.179602 | -0.09677 | -1.660257 | -0.208842 | -1.163677 | -1.693431 | | 37.08333 | -13.362284 | -1.633162 |
| 37 | -6.631829 | -3.130588 | -1.443679 | -3.597387 | -6.288359 | -3.314386 | -5.364825 | -5.295282 | -5.140589 | -3.890881 | -5.214596 | -3.185632 | -12.000011 | -1.140034 | -1.420713 | 0.113454 | -0.447775 | -0.440869 | 0.043804 | 0.104028 | -0.698978 | 0.04118 | -0.945233 | -0.84338 | | 37.09722 | -12.000011 | -0.968019 |
| 37.014 | -5.023853 | -1.656863 | -1.100602 | -3.273994 | -4.523402 | -2.153184 | -2.703881 | -3.553177 | -4.051294 | -4.48089 | -2.84229 | -3.002294 | -8.685762 | -0.723897 | -2.401634 | 0.1452 | -0.4013 | -0.352775 | 0.083708 | 0.121705 | -0.045377 | 0.086479 | -0.639305 | -0.440021 | | 37.11111 | -8.685762 | -0.607877 |
| 37.028 | -1.964893 | -0.665974 | -0.759443 | -1.8175 | -2.69937 | -0.956227 | -0.917425 | -3.298635 | -2.812003 | -3.686872 | -0.900868 | -2.221347 | -4.919008 | 0.054548 | -2.251118 | 0.097096 | -0.117528 | -0.250502 | 0.065286 | 0.088739 | 0.055508 | 0.067018 | -0.939218 | -0.208959 | | 37.125 | -4.919008 | -0.338157 |
| 37.042 | 0.375 | 0.251589 | -0.342602 | -0.452489 | -1.654615 | -0.653439 | 0.277491 | -3.331937 | -1.834689 | -2.278278 | 0.417814 | -1.766709 | -1.93422 | 0.83654 | -1.483195 | 0.050843 | 0.395803 | -0.125816 | 0.06652 | 0.083923 | 0.04873 | 0.068183 | -0.715857 | 0.011733 | | 37.13889 | -1.93422 | -0.058737 |
| 37.056 | 3.242533 | 1.271089 | 0.249016 | 0.281552 | -0.881461 | 0.253974 | 1.457279 | -2.760939 | -1.072915 | -0.715908 | 1.507593 | -0.558015 | 0.638145 | 1.559791 | -0.716667 | 0.001619 | 0.934947 | -0.065345 | 0.052677 | 0.069476 | 0.036086 | 0.056128 | -0.175296 | 0.291733 | | 37.15278 | 0.638145 | 0.22096 |
| 37.069 | 5.814605 | 2.50174 | 0.566876 | 0.775147 | -0.009056 | 1.341474 | 3.014409 | -1.761622 | -0.755027 | 0.440492 | 2.479152 | 0.783047 | 3.140421 | 2.008882 | -0.146304 | -0.044142 | 1.389238 | -0.074627 | 0.023708 | 0.028143 | 0.065283 | 0.022838 | 0.238497 | 0.558395 | | 37.16667 | 3.140421 | 0.478856 |
| 37.083 | 7.835168 | 2.390532 | 0.5416 | 1.360333 | 1.03497 | 1.994422 | 3.295095 | -0.13778 | -0.201651 | 1.203778 | 2.7231 | 1.67794 | 4.629333 | 1.752089 | 0.359274 | -0.036838 | 1.51489 | -0.100117 | -0.001637 | -0.018478 | 0.155039 | -0.014157 | 0.374818 | 0.661868 | | 37.18056 | 4.629333 | 0.625297 |
| 37.097 | 5.668826 | 0.936898 | 0.385673 | 1.461383 | 2.673345 | 2.222513 | 1.984392 | 1.237987 | 0.111172 | 1.517697 | 1.378481 | 2.743261 | 4.552917 | 0.8636 | 0.714563 | -0.017735 | 1.153923 | -0.110246 | -0.019817 | -0.047738 | 0.361577 | -0.039744 | 0.114728 | 0.55103 | | 37.19444 | 4.552917 | 0.58907 |
| 37.111 | 2.261138 | 0.101626 | 0.530285 | 1.32455 | 4.717897 | 1.377096 | 1.361763 | 1.939668 | 1.363182 | 2.096515 | -0.192881 | 2.658859 | 2.563896 | 0.03898 | 1.120558 | -0.015159 | 0.85825 | -0.089668 | -0.014245 | -0.039422 | 0.772712 | -0.028765 | -0.283693 | 0.445853 | | 37.20833 | 2.563896 | 0.539706 |
| 37.125 | -2.315638 | 0.205056 | 0.787605 | 1.112596 | 4.822139 | 0.615073 | 0.694036 | 1.396239 | 2.990336 | 1.086675 | -1.192228 | 1.903591 | 0.628231 | -0.28596 | 1.49432 | -0.003421 | 1.046369 | -0.052376 | 0.010878 | -0.005308 | 1.408843 | 0.00819 | -0.47067 | 0.609375 | | 37.22222 | 0.628231 | 0.695497 |
| 37.139 | -4.498743 | 0.705516 | 0.91487 | 0.533947 | 2.8707 | 0.493199 | 0.030031 | 0.914224 | 3.458913 | -0.61891 | -1.743231 | -0.451073 | -0.551242 | 0.088666 | 1.673466 | 0.016225 | 1.73291 | -0.003418 | 0.043664 | 0.036536 | 1.668974 | 0.047471 | -0.500849 | 0.999164 | | 37.23611 | -0.551242 | 1.019506 |
| 37.153 | -3.271515 | 1.747905 | 0.662792 | -0.341786 | 0.792292 | 0.931985 | -0.41654 | 0.276787 | 2.879379 | -2.256037 | -0.549076 | -2.592885 | -0.566332 | 0.420005 | 1.279294 | 0.037446 | 2.432938 | 0.03765 | 0.082706 | 0.086064 | 1.85307 | 0.090175 | -0.630758 | 1.176514 | | 37.25 | -0.566332 | 1.170275 |
| 37.167 | -1.025934 | -0.383938 | 0.299191 | -0.384333 | -0.904249 | 1.802589 | -0.731649 | 0.09598 | 2.049827 | -1.010466 | -0.111102 | -1.909971 | -0.479818 | 0.446249 | 0.909627 | 0.049158 | 2.521428 | 0.047138 | 0.114845 | 0.118835 | 1.858499 | 0.126584 | -0.711467 | 1.021785 | | 37.26389 | -0.479818 | 0.947055 |
| 37.181 | -0.121678 | -2.818305 | 0.066272 | 0.181055 | -1.420903 | 2.838895 | -0.669469 | 0.066392 | 1.24817 | 0.124836 | -0.50732 | -0.171651 | 0.276687 | -0.031675 | 0.741545 | 0.070063 | 1.894899 | 0.058687 | 0.136414 | 0.130826 | 1.73619 | 0.14514 | -0.69025 | 0.692286 | | 37.27778 | 0.276687 | 0.610627 |
| 37.194 | 0.065384 | -2.049727 | 0.114037 | 0.771607 | -0.105881 | 3.630659 | 0.240473 | 0.140488 | 0.930109 | 0.959076 | -1.223658 | 1.238909 | 1.080123 | -0.769211 | 0.525901 | 0.066219 | 1.017079 | 0.081178 | 0.151405 | 0.141178 | 1.461957 | 0.152491 | -0.244181 | 0.402201 | | 37.29167 | 1.080123 | 0.364254 |
| 37.208 | 1.5763 | -0.312337 | 0.387868 | 0.393895 | 1.554233 | 3.967762 | 0.927018 | 0.330015 | 1.403945 | 1.501398 | -0.552784 | 1.971217 | 1.585616 | -1.228544 | 0.278107 | 0.081325 | 0.442492 | 0.089201 | 0.158 | 0.150409 | 1.151291 | 0.158408 | 0.134313 | 0.191842 | | 37.30556 | 1.585616 | 0.147216 |
| 37.222 | 2.609354 | 1.68563 | 0.63969 | 0.079863 | 2.448098 | 2.945888 | 1.280895 | 0.445045 | 1.895147 | 1.975942 | 2.192078 | 2.898043 | 2.070427 | -0.989791 | 0.087956 | 0.08358 | 0.299649 | 0.072022 | 0.146814 | 0.146222 | 0.899637 | 0.150529 | 0.391941 | 0.155225 | | 37.31944 | 2.070427 | 0.147291 |
| 37.236 | 3.248274 | 0.975478 | 0.757722 | 0.809385 | 2.683018 | 2.106655 | 1.144793 | 0.965963 | 1.843552 | 1.997214 | 4.796989 | 4.66203 | 2.598739 | -0.136862 | -0.057214 | 0.04392 | 0.634704 | -0.004286 | 0.09962 | 0.107807 | 0.724721 | 0.103905 | 0.572316 | 0.301936 | | 37.33333 | 2.598739 | 0.279824 |
| 37.25 | 1.531274 | 0.145766 | 0.752411 | 1.829426 | 2.440505 | 1.867638 | 0.7702 | 1.66546 | 2.266988 | 1.627408 | 5.118295 | 6.217344 | 3.571263 | 1.044315 | -0.280944 | -0.001662 | 1.170297 | -0.088424 | 0.062053 | 0.070343 | 0.642821 | 0.062352 | 0.615381 | 0.457403 | | 37.34722 | 3.571263 | 0.382585 |
| 37.264 | 2.014226 | 0.834868 | 0.644449 | 2.445285 | 2.043938 | 1.715516 | 0.168021 | 1.714433 | 2.526941 | 1.344416 | 4.281543 | 5.308796 | 4.666751 | 2.312024 | -0.285481 | 0.025398 | 1.601064 | -0.113509 | 0.0673 | 0.074861 | 0.652768 | 0.068425 | 0.649149 | 0.587675 | | 37.36111 | 4.666751 | 0.45463 |
| 37.278 | 4.372333 | 1.535394 | 0.524365 | 1.774973 | 1.906141 | 1.582365 | -0.004198 | 1.40143 | 2.343393 | 1.528851 | 2.222225 | 4.243615 | 5.286347 | 3.075636 | -0.022034 | 0.062108 | 1.688456 | -0.095129 | 0.077581 | 0.089164 | 0.714582 | 0.087969 | 0.771491 | 0.65082 | | 37.375 | 5.286347 | 0.488292 |
| 37.292 | 3.352842 | 2.122771 | 0.407629 | 1.149614 | 1.948218 | 2.096794 | 0.246513 | 0.918482 | 2.033693 | 1.772193 | 0.597217 | 3.623584 | 4.51171 | 3.057386 | 0.565106 | 0.071407 | 1.373667 | -0.063958 | 0.073576 | 0.087695 | 0.724062 | 0.087569 | 0.886266 | 0.599364 | | 37.38889 | 4.51171 | 0.457176 |
| 37.306 | -0.801877 | 2.403415 | 0.42001 | 0.937714 | 2.290561 | 2.506459 | 0.703365 | 0.837281 | 1.779925 | 2.059485 | 0.954509 | 2.76706 | 2.971899 | 2.013246 | 1.117693 | 0.055184 | 1.075217 | -0.064033 | 0.079354 | 0.045607 | 0.589403 | 0.081861 | 0.925157 | 0.387385 | | 37.40278 | 2.971899 | 0.444368 |
| 37.319 | -3.148259 | 2.196665 | 0.566116 | 0.995715 | 2.180197 | 2.149918 | 0.995051 | 1.107856 | 1.12405 | 2.104458 | 2.363277 | 2.685876 | 1.256077 | 0.860666 | 1.602803 | 0.027346 | 1.238082 | -0.099029 | 0.225597 | -0.065045 | 0.313949 | 0.099445 | 0.893342 | -0.47267 | | 37.41667 | 1.256077 | 0.332447 |
| 37.333 | -3.920625 | 1.385957 | 0.715098 | 1.080678 | 1.906515 | 1.380455 | 1.278695 | 1.389161 | 1.235774 | 1.947763 | 3.666727 | 3.827533 | 0.102953 | -0.084541 | 2.048986 | 0.009525 | 1.621253 | -0.128827 | 0.381594 | 0.200622 | 0.154325 | 0.058598 | 0.825688 | -1.099944 | | 37.43056 | 0.102953 | 0.180058 |
| 37.347 | -1.776247 | -0.302491 | 0.849023 | 1.384648 | 1.408726 | 0.223671 | 1.242922 | 0.979789 | 1.58494 | 2.175941 | 5.125063 | 5.589232 | 1.514143 | 0.163856 | 1.789882 | -0.001931 | 1.960808 | -0.028771 | 0.01189 | 0.015508 | 0.52329 | 0.00108 | 1.084183 | 1.105584 | | 37.44444 | 1.514143 | 1.444706 |
| 37.361 | 2.496198 | -1.084695 | 0.452589 | 1.320299 | 1.25137 | -0.803565 | 0.458599 | 1.07111 | 2.620408 | 2.779382 | 5.179689 | 4.612916 | 3.5488 | 1.20984 | 1.265716 | -0.069487 | 2.04143 | 0.001881 | -0.465935 | -0.227754 | 0.661061 | -0.139557 | 1.533164 | 3.255071 | | 37.45833 | 3.5488 | 2.508915 |
| 37.375 | 6.504624 | -0.866966 | -0.169351 | 0.842175 | 1.086852 | -0.928549 | -0.644897 | 2.120472 | 3.335013 | 2.954878 | 3.979368 | 3.57085 | 3.699991 | 0.671936 | 1.251025 | -0.199068 | 2.402702 | -0.181589 | -0.498011 | -0.063926 | 0.351326 | -0.364688 | 1.975369 | 2.080404 | | 37.47222 | 3.699991 | 1.925163 |
| 37.389 | 7.734165 | -0.675084 | -0.240471 | 0.56013 | 1.336233 | 0.362832 | -1.320383 | 2.931695 | 3.871915 | 3.39158 | 2.208042 | 4.383357 | 3.268557 | 0.678479 | 1.593375 | -0.200348 | 3.222234 | -0.19969 | -0.262949 | -0.095429 | 0.422905 | -0.160414 | 2.750315 | 0.616166 | | 37.48611 | 3.268557 | 1.383065 |
| 37.403 | 7.238076 | 1.199661 | -0.018489 | 0.708886 | 1.935581 | 3.079185 | -0.471041 | 2.817279 | 4.018206 | 4.593759 | 2.097659 | 4.913512 | 3.76003 | -0.426 | 1.814262 | -0.071638 | 3.908879 | -0.111091 | -0.054312 | -0.023661 | 0.872947 | -0.004928 | 3.407647 | 0.395039 | | 37.5 | 3.76003 | 1.53484 |
| 37.417 | 4.381169 | 2.945264 | -0.036746 | 1.226714 | 2.299856 | 5.265959 | 1.684491 | 3.841752 | 4.439816 | 5.534691 | 2.683008 | 5.577976 | 4.943053 | 3.051107 | 2.435393 | -0.02083 | 3.844567 | -0.106218 | -0.009305 | -0.013472 | 1.174212 | 0.01465 | 3.593303 | 1.895324 | | 37.51389 | 4.943053 | 2.111019 |
| 37.431 | 2.628076 | 3.132901 | -0.183974 | 1.797581 | 2.405837 | 5.008848 | 5.761982 | 5.50602 | 4.397727 | 5.530058 | 4.104137 | 5.561173 | 6.534151 | 4.149689 | 2.235083 | -0.058802 | 3.732976 | -0.15461 | -0.036907 | -0.041324 | 1.242514 | -0.009684 | 3.219056 | 2.591598 | | 37.52778 | 6.534151 | 2.705191 |
| 37.444 | 2.373672 | 2.740889 | -0.188806 | 2.182853 | 2.318189 | 4.903497 | 6.70299 | 4.172624 | 4.64809 | 5.161995 | 5.760602 | 6.201041 | 8.829627 | 1.311912 | 2.109303 | -0.461161 | 3.207644 | -0.526109 | -0.403824 | -0.413176 | 1.009745 | -0.319937 | 2.803776 | 2.299333 | | 37.54167 | 8.829627 | 2.122356 |
| 37.458 | 3.620245 | 2.149631 | 0.397262 | 2.567345 | 2.618213 | 6.409821 | 4.158916 | 0.026736 | 4.80428 | 3.882207 | 7.883663 | 6.597549 | 10.934021 | 0.581523 | 0.858049 | -1.287868 | 1.963183 | -1.388515 | -1.245273 | -1.264648 | 0.497614 | -1.082618 | 2.150025 | 1.118359 | | 37.55556 | 10.934021 | 0.812629 |
| 37.472 | 2.747883 | 1.288922 | 0.479471 | 2.143028 | 2.675258 | 5.053876 | 1.04557 | -1.285647 | 3.590104 | 2.922033 | 6.960952 | 3.934761 | 8.762747 | 1.011065 | -0.268898 | -1.621755 | 0.662428 | -1.776563 | -1.651176 | -1.689929 | 0.073817 | -1.535019 | 1.054989 | -0.134714 | | 37.56944 | 8.762747 | -0.35055 |
| 37.486 | 0.725863 | 0.927472 | 0.116231 | 2.399987 | 2.582495 | 4.757271 | -1.733556 | 0.282298 | 2.69149 | 3.016968 | 2.530569 | 1.004337 | 4.212454 | 0.55431 | -0.549437 | -1.080488 | 0.472172 | -1.21489 | -1.175917 | -1.152584 | 0.704442 | -1.174772 | 0.879421 | -0.401896 | | 37.58333 | 4.212454 | -0.644418 |
| 37.5 | -0.984 | -0.139078 | 0.471837 | 3.071234 | 2.814842 | 5.162657 | -2.4005 | 1.971616 | 1.947907 | 3.504725 | 0.803918 | -0.012269 | 1.550914 | -0.361445 | -0.428457 | -0.524915 | 1.103382 | -0.594725 | -0.57142 | -0.53381 | 1.754898 | -0.61474 | 1.486755 | 0.030478 | | 37.59722 | 1.550914 | -0.139909 |
| 37.514 | -0.808192 | -0.056635 | 0.785779 | 2.355476 | 2.860788 | 4.494636 | 0.648631 | 2.133763 | 1.165052 | 2.873805 | -0.796371 | 0.153362 | 1.65476 | -2.187329 | -0.315507 | -0.229703 | 1.201827 | -0.281753 | -0.234067 | -0.21236 | 3.005099 | -0.262368 | 2.064071 | 0.390848 | | 37.61111 | 1.65476 | 0.324591 |
| 37.528 | 2.479067 | 0.537667 | 0.131525 | 0.351829 | 2.625549 | 1.558637 | 3.796777 | 0.061639 | 0.095786 | 1.490363 | -2.198633 | -0.727793 | 1.793662 | -3.914655 | -0.198446 | -0.096106 | 0.910162 | -0.150962 | -0.083183 | -0.065992 | 4.305184 | -0.094857 | 2.010537 | 0.381955 | | 37.625 | 1.793662 | 0.30083 |
| 37.542 | 2.005237 | 2.252397 | -0.418105 | -1.112636 | 0.500317 | -0.217386 | 6.130951 | -0.982792 | -0.485396 | 0.864669 | -3.388528 | -0.014732 | 2.637516 | -1.485377 | 0.383334 | -0.047061 | 0.668969 | -0.095619 | -0.021121 | 0.001583 | 2.298647 | -0.024152 | 1.221582 | 0.306396 | | 37.63889 | 2.637516 | 0.183071 |
| 37.556 | -0.449986 | 3.203685 | -0.254946 | -0.843215 | -1.25299 | -0.768868 | 3.840775 | -0.060877 | 0.057644 | -0.703933 | -0.074326 | 4.016207 | 5.468266 | 0.368613 | 0.352493 | -0.054117 | 0.381887 | -0.087262 | -0.019044 | 0.014915 | -0.231152 | -0.017336 | 0.168973 | 0.1305 | | 37.65278 | 5.468266 | 0.019215 |
| 37.569 | -2.737048 | 3.885677 | 0.02598 | 1.239669 | -0.589207 | 0.219532 | 6.805292 | 1.311588 | 0.888478 | 0.62516 | 2.334717 | 4.521224 | 8.538323 | 1.946198 | 0.259517 | -0.082847 | -0.554566 | -0.120419 | -0.052585 | -0.008952 | -2.173111 | -0.050287 | -0.779459 | -0.278906 | | 37.66667 | 8.538323 | -0.299658 |
| 37.583 | -1.4633 | 1.785449 | 0.710065 | 1.613663 | 1.541376 | 0.478803 | -1.977199 | 1.86078 | 0.584775 | 2.343133 | 1.689088 | 3.070557 | 11.746475 | 0.84861 | -0.199089 | -0.085334 | -1.108487 | -0.143737 | -0.062046 | -0.02197 | -2.570497 | -0.06436 | -1.716213 | -0.661424 | | 37.68056 | 11.746475 | -0.6501 |
| 37.597 | -0.849082 | -2.017965 | 1.828478 | 0.899898 | 2.939808 | 0.037024 | -2.481235 | 0.058401 | 0.467148 | -0.695829 | 0.381133 | -0.018061 | 4.12977 | -1.633739 | -0.400704 | -0.041977 | -0.807549 | -0.123959 | -0.042173 | -0.024409 | -0.485758 | -0.048704 | -2.002238 | -0.545868 | | 37.69444 | 4.12977 | -0.754301 |
| 37.611 | 1.230466 | -5.115418 | 2.958726 | 0.427303 | 1.884988 | 1.02617 | 3.352964 | -0.140406 | 0.299049 | -0.845774 | 1.694949 | -1.022492 | -2.914909 | -2.39913 | -0.468319 | 0.010592 | -0.162625 | -0.069525 | -0.00638 | -0.011285 | 1.175181 | -0.01008 | -1.373227 | -0.267532 | | 37.70833 | -2.914909 | -0.551726 |
| 37.625 | 0.973434 | -3.226504 | 4.778589 | -0.310629 | 0.743546 | 1.894536 | 4.942504 | -0.165682 | -0.151611 | -0.162096 | 2.400624 | -2.567857 | -6.696324 | -2.103023 | -0.511458 | 0.121541 | 0.551174 | 0.036283 | 0.096707 | 0.080155 | 1.864388 | 0.098214 | -0.565386 | -0.248949 | | 37.72222 | -6.696324 | -0.257088 |
| 37.639 | 1.042656 | 0.11352 | 3.905006 | -0.263583 | 0.006347 | 1.115609 | 1.593358 | -0.626139 | 0.234375 | -0.775296 | 4.783061 | -2.177354 | -4.988129 | -1.097315 | -0.575179 | 0.296002 | 0.761359 | 0.196871 | 0.275828 | 0.23698 | 2.399079 | 0.280729 | -0.064934 | 0.03656 | | 37.73611 | -4.988129 | -0.001837 |
| 37.653 | 2.063576 | 2.734425 | 1.877037 | 0.30615 | -1.025009 | 0.358239 | -1.184975 | -1.306929 | 0.994877 | -2.209252 | 1.786259 | -1.058177 | -0.759526 | -0.189505 | -0.629057 | 0.465182 | 0.816581 | 0.356559 | 0.455185 | 0.394264 | 1.174391 | 0.462282 | -0.196828 | 0.228153 | | 37.75 | -0.759526 | 0.152967 |
| 37.667 | 0.990426 | 2.693836 | 0.459995 | 0.378975 | -1.392323 | 0.141782 | -5.887327 | 1.720458 | 1.024881 | -1.269706 | -1.550843 | -0.784822 | 2.069445 | 0.485309 | -0.676873 | 0.564349 | 0.639971 | 0.468633 | 0.577084 | 0.507396 | -0.685959 | 0.580158 | -0.309269 | 0.181712 | | 37.76389 | 2.069445 | 0.099754 |
| 37.681 | 0.99353 | 0.422311 | -0.006561 | 0.06856 | -0.882079 | 0.422571 | -1.467724 | 1.189059 | -0.360323 | -2.560001 | -2.934117 | 0.300072 | 4.869529 | 0.826491 | -0.600532 | 0.588267 | 0.194307 | 0.506622 | 0.620707 | 0.568979 | -1.568162 | 0.61313 | -0.224726 | 0.126836 | | 37.77778 | 4.869529 | 0.061789 |
| 37.694 | 0.919558 | -0.834359 | -0.557067 | 0.30772 | -0.136132 | 0.213548 | 0.28668 | 0.586031 | 0.142696 | -1.480367 | -3.710461 | 0.726544 | 3.801009 | 0.912871 | -0.258721 | 0.531613 | -0.319083 | 0.473563 | 0.58293 | 0.571828 | -1.764437 | 0.573139 | -0.105392 | 0.03113 | | 37.79167 | 3.801009 | 0.057434 |
| 37.708 | 0.242946 | -1.909475 | -0.920787 | 0.80049 | -0.097448 | -0.165823 | 1.029433 | 0.006719 | 0.930041 | 0.564847 | -3.907249 | 0.945356 | 0.971656 | 0.431437 | 0.232916 | 0.445799 | -1.33415 | 0.402309 | 0.493989 | 0.508291 | -0.77084 | 0.498312 | -0.289967 | -0.151451 | | 37.80556 | 0.971656 | 0.022567 |
| 37.722 | 0.127386 | -2.589852 | -1.186871 | 0.883035 | 0.739488 | -0.750587 | -0.518615 | 0.541394 | 0.929713 | 1.669963 | -3.283011 | 0.810489 | -1.779171 | -0.974983 | 0.490692 | 0.345093 | -2.700188 | 0.318975 | 0.391459 | 0.404826 | 0.137391 | 0.403756 | -0.754381 | -0.336691 | | 37.81944 | -1.779171 | -0.169302 |
| 37.736 | -1.891807 | -1.649796 | -1.189941 | 0.967037 | 0.105573 | -1.046744 | -2.478423 | 0.289206 | 0.974756 | 1.304041 | -1.820657 | 0.826996 | -2.650704 | -3.134621 | -0.070402 | 0.249226 | -3.29694 | 0.228193 | 0.288772 | 0.289835 | 1.801504 | 0.297991 | -1.288751 | -0.519219 | | 37.83333 | -2.650704 | -0.457541 |
| 37.75 | -4.849998 | -1.07005 | -1.375362 | -0.036309 | 0.004961 | -1.387666 | -2.745116 | -0.333356 | 0.697825 | -0.662872 | -0.704599 | 1.274379 | -3.187244 | -4.861531 | -0.62231 | 0.162878 | -3.000427 | 0.128358 | 0.181888 | 0.1644 | 2.933222 | 0.191896 | -1.175912 | -0.908525 | | 37.84722 | -3.187244 | -0.843538 |
| 37.764 | -3.460866 | -0.697365 | -1.597465 | 0.478744 | 0.176207 | -1.113569 | -1.992767 | -0.618008 | 0.538332 | -0.690819 | -0.554373 | 1.215343 | -5.225927 | -5.151671 | -1.329609 | 0.09096 | -1.754867 | 0.033575 | 0.081966 | 0.051858 | 1.951184 | 0.101137 | -1.033773 | -1.355341 | | 37.86111 | -5.225927 | -1.161966 |
| 37.778 | -3.414733 | -0.815442 | -1.731275 | -0.808801 | -0.095657 | -0.465812 | -1.048939 | -0.682694 | 0.149671 | -1.15095 | -1.509989 | -1.667368 | -9.854222 | -4.39912 | -1.748017 | 0.051284 | -1.444859 | -0.020646 | 0.023491 | -0.004631 | 0.022115 | 0.049674 | -1.039715 | -1.235333 | | 37.875 | -9.854222 | -1.14644 |
| 37.792 | -2.30181 | -1.036441 | -1.728378 | -1.475851 | -0.561819 | -0.215999 | -0.141476 | -0.614181 | -0.412835 | 0.835614 | -2.39144 | -3.752263 | -12.997507 | -2.886888 | -1.499115 | 0.023783 | -1.697258 | -0.051092 | -0.005613 | -0.027493 | -2.453783 | 0.014777 | -1.335941 | -1.108043 | | 37.88889 | -12.997507 | -1.096169 |
| 37.806 | 3.140619 | 0.422203 | -1.722781 | -1.03278 | -0.791571 | -0.986785 | 0.616235 | -0.860192 | -0.980196 | -1.615188 | -2.651173 | -2.325273 | -8.013034 | -1.444584 | -1.086033 | -0.01439 | -3.164121 | -0.080961 | -0.028073 | -0.051078 | -4.672053 | -0.014538 | -1.74594 | -1.49831 | | 37.90278 | -8.013034 | -1.348744 |
| 37.819 | 6.17773 | -0.324716 | -1.348969 | 0.27198 | -1.425152 | -2.643033 | 2.078602 | -0.683231 | -0.924516 | -2.116774 | -3.552274 | 1.296367 | -0.805023 | -0.61875 | -0.857648 | -0.012619 | -4.352493 | -0.078039 | -0.015661 | -0.04319 | -5.336194 | -0.003377 | -2.163469 | -1.846114 | | 37.91667 | -0.805023 | -1.753975 |
| 37.833 | 3.827654 | 0.498994 | -0.435952 | -0.111996 | -1.12384 | -4.520751 | 3.890492 | 0.157059 | -0.730193 | -1.137155 | -6.367742 | 0.219861 | 7.000437 | -0.114909 | -0.873243 | 0.029399 | -4.514836 | -0.048593 | 0.017597 | -0.00985 | -5.326816 | 0.025114 | -2.332627 | -2.369561 | | 37.93056 | 7.000437 | -2.286794 |
| 37.847 | -2.056824 | 0.954925 | -0.20036 | -2.56625 | 0.181154 | -6.82772 | 3.683124 | 0.884093 | 0.189278 | -0.166961 | -2.198346 | 1.903685 | 11.07322 | -0.01851 | -0.940799 | 0.046717 | -3.870286 | -0.031011 | 0.035512 | 0.015338 | -3.782265 | 0.037494 | -2.087885 | -2.141871 | | 37.94444 | 11.07322 | -2.243503 |
| 37.861 | -2.15592 | -0.450281 | -0.361128 | -0.857658 | 0.271426 | -7.130914 | 3.23921 | 0.568715 | 0.410713 | -0.657209 | -0.154187 | 2.390212 | 6.755429 | -0.098232 | -0.965766 | 0.04033 | -3.039471 | -0.026951 | 0.036145 | 0.028752 | -1.882166 | 0.044925 | -1.612515 | -1.742464 | | 37.95833 | 6.755429 | -1.877499 |
| 37.875 | -1.849973 | -0.78997 | -0.407617 | -0.080111 | -0.195153 | -6.257952 | 1.021996 | 0.026265 | 1.079577 | 0.07784 | 0.668848 | 0.729136 | 0.634768 | -0.04664 | -0.999852 | 0.012493 | -2.491252 | -0.039324 | 0.027551 | 0.021077 | -0.923752 | 0.033539 | -1.344106 | -1.467778 | | 37.97222 | 0.634768 | -1.579473 |

| | | | | | | | | | | | | | | | | | | | | | | | | | | | | |
|---|---|---|---|---|---|---|---|---|---|---|---|---|---|---|---|---|---|---|---|---|---|---|---|---|---|---|---|---|
| 37.889 | -1.742277 | -0.516656 | -0.260198 | 0.472136 | -1.562526 | -2.435867 | -0.564996 | -0.186926 | 1.170983 | 0.492466 | -2.059809 | -1.777172 | -4.631411 | -0.200795 | -1.067624 | -0.021011 | -2.333107 | -0.073922 | -0.02164 | -0.014861 | -0.597216 | -0.013793 | -1.240736 | -1.266994 | | 37.98611 | -4.631411 | -1.455759 |
| 37.903 | -2.338495 | -2.088464 | -0.761814 | 1.642229 | -3.112508 | 1.809589 | 2.518749 | -0.420077 | 0.845996 | -0.137173 | -5.68132 | -4.277988 | -7.589501 | -0.497506 | -1.099502 | -0.025262 | -2.029574 | -0.075346 | -0.033006 | -0.015695 | -0.530107 | -0.019162 | -1.022624 | -1.145467 | | 38 | -7.589501 | -1.336154 |
| 37.917 | -2.232413 | -1.620448 | -1.50569 | 0.65014 | -3.210123 | 4.804574 | 2.53593 | -0.342366 | 0.213572 | 2.054226 | -6.147206 | -4.030802 | -8.444878 | -0.732142 | -0.969664 | 0.079979 | -1.325049 | 0.040469 | 0.134944 | 0.111629 | -0.419665 | 0.131407 | -0.602351 | -0.97973 | | 38.01389 | -8.444878 | -1.07209 |

**Supplementary Table S7 (g)** Mean day-to-day motion rate in Control and Stress tanks of Amaranth measured under dimGday/night and RGBday dimGnight, calculated using either 24 h motion integration or 22 h motion integration excluding day/night transitions.

| Row | RGB22Control | RGB22Stress | RGBControl | RGBStress | dimG22Control | dimG22Stress | dimGControl | dimGStress |
|---|---|---|---|---|---|---|---|---|
| 18.5 | 0.54479302 | 0.690216507 | 0.755063598 | 0.908435061 | 0.165404692 | 0.661371742 | 0.077433799 | 0.285763979 |
| 19.5 | 0.48691375 | 0.599609597 | 0.699635065 | 0.813158757 | 0.067563852 | 0.24307877 | 0.082894063 | 0.235884477 |
| 20.5 | 0.50961553 | 0.694267977 | 0.728555724 | 0.884525072 | 0.084127178 | 0.354009276 | 0.123380787 | 0.384653414 |
| 21.5 | 0.51901935 | 0.648268357 | 0.744161996 | 0.831940682 | 0.117570906 | 0.324208151 | 0.150745909 | 0.342892146 |
| 22.5 | 0.541142709 | 0.576727374 | 0.748117543 | 0.670777596 | 0.137292243 | 0.3893113 | 0.146121438 | 0.4574556 |
| 23.5 | 0.546375919 | 0.475757308 | 0.774329853 | 0.559711611 | 0.159249844 | 0.409248271 | 0.210752425 | 0.408435598 |
| 24.5 | 0.615799055 | 0.445763878 | 0.821371508 | 0.541372139 | 0.339271828 | 0.327900474 | 0.437671361 | 0.382014978 |
| 25.5 | 0.668498911 | 0.537617449 | 0.869141917 | 0.635394462 | 0.556629527 | 0.511713647 | 0.548277729 | 0.476758793 |
| 26.5 | 0.705862699 | 0.316183684 | 0.867924089 | 0.354792493 | 0.574395335 | 0.372139142 | 0.599903071 | 0.416755278 |
| 27.5 | 0.752019322 | 0.373710361 | 0.893210111 | 0.462585561 | 0.705260177 | 0.425469574 | 0.869904461 | 0.426943433 |
| 28.5 | 0.865967475 | 0.56264795 | 1.011919719 | 0.698620092 | 0.944385427 | 0.479649237 | 1.161667143 | 0.516311421 |
| 29.5 | 1.059872863 | 0.565491133 | 1.237705227 | 0.75180018 | 1.158848819 | 0.404111092 | 1.355811058 | 0.469491748 |
| 30.5 | 1.663373393 | 0.805096486 | 1.876148405 | 1.047767738 | 1.864878111 | 0.518882073 | 1.949923372 | 0.596499546 |
| 31.5 | 2.127596189 | 0.696268285 | 2.289374734 | 0.864707267 | 2.109968085 | 0.539787957 | 2.291192124 | 0.566269718 |
| 32.5 | 2.223259283 | 0.801374841 | 2.465729357 | 0.987275421 | 2.290304651 | 0.664989014 | 2.635189208 | 0.818798016 |
| 33.5 | 2.933985348 | 0.92346411 | 3.296282316 | 1.140909039 | 2.847592848 | 0.863388054 | 3.11438738 | 1.024033804 |
| 34.5 | 3.176626496 | 0.846411054 | 3.733184435 | 1.06097033 | 2.824745736 | 0.820955815 | 3.199420572 | 0.918459853 |
| 35.5 | 3.257550073 | 0.841319236 | 3.848596847 | 1.040614787 | 3.045115513 | 0.732446502 | 3.549895001 | 0.907234573 |
| 36.5 | 4.077928816 | 0.699844154 | 4.816938187 | 0.91247367 | 3.982925396 | 0.697420817 | 3.892857621 | 0.732231014 |

**Supplementary Table S7 (h)** Mean day-to-day motion rate of individual Amaranth plants under dimGday/night and RGBday dimGnight. Data represent means of 9–10 plants

| Interval | RGBControlMean | RGBControlSEM | RGBStressMean | RGBStressSEM | dimGControlMean | dimGControlSEM | dimGStressMean | dimGStressSEM |
|---|---|---|---|---|---|---|---|---|
| 18.5 | 0.216394114 | 0.030361512 | 0.264790928 | 0.015478555 | 0.087153741 | 0.011554016 | 0.235967022 | 0.010454294 |
| 19.5 | 0.21122412 | 0.035069737 | 0.306810224 | 0.011456207 | 0.102183661 | 0.017855493 | 0.272900772 | 0.012819876 |
| 20.5 | 0.243565235 | 0.037191571 | 0.41151498 | 0.017486954 | 0.159567751 | 0.03556145 | 0.432601714 | 0.019497727 |
| 21.5 | 0.255259958 | 0.032408387 | 0.39138105 | 0.020923852 | 0.194696629 | 0.039687716 | 0.429371861 | 0.034714891 |
| 22.5 | 0.281049763 | 0.035083846 | 0.476613414 | 0.060714627 | 0.256222933 | 0.046219521 | 0.541577105 | 0.078776185 |
| 23.5 | 0.344668413 | 0.048554031 | 0.325650563 | 0.038919162 | 0.332742975 | 0.058859749 | 0.363818559 | 0.03757468 |
| 24.5 | 0.429095721 | 0.063567579 | 0.355059231 | 0.02342799 | 0.433008437 | 0.082918451 | 0.378493094 | 0.033653181 |
| 25.5 | 0.56702043 | 0.08250643 | 0.4536194 | 0.052682656 | 0.600789702 | 0.094743466 | 0.51765048 | 0.073342231 |
| 26.5 | 0.655378776 | 0.101764679 | 0.421121707 | 0.04784345 | 0.700159974 | 0.113903346 | 0.488037517 | 0.0693524 |
| 27.5 | 0.694064255 | 0.083492114 | 0.493085668 | 0.07696444 | 0.764177182 | 0.097257803 | 0.533740897 | 0.094375556 |
| 28.5 | 0.830219522 | 0.069265355 | 0.555006053 | 0.086450784 | 0.933757001 | 0.080945529 | 0.61040802 | 0.095229174 |
| 29.5 | 0.877336905 | 0.079624421 | 0.557120656 | 0.078240149 | 0.984851603 | 0.086168735 | 0.60669702 | 0.105378513 |
| 30.5 | 0.986831868 | 0.096669949 | 0.644820425 | 0.105423391 | 1.057757734 | 0.092808816 | 0.686366711 | 0.123953282 |
| 31.5 | 1.033053746 | 0.099880951 | 0.669654857 | 0.132002511 | 1.092365833 | 0.104610347 | 0.710474947 | 0.146844956 |
| 32.5 | 1.170215758 | 0.115093161 | 0.748488432 | 0.150057579 | 1.238648225 | 0.13007652 | 0.775101871 | 0.174829931 |
| 33.5 | 1.348796547 | 0.129105496 | 0.791898241 | 0.181777737 | 1.4831063 | 0.155109291 | 0.809789951 | 0.199855802 |
| 34.5 | 1.513969945 | 0.152495046 | 0.734873194 | 0.192635105 | 1.549254305 | 0.15122363 | 0.761032328 | 0.20055943 |
| 35.5 | 1.627858352 | 0.158018178 | 0.862374903 | 0.234012329 | 1.689042309 | 0.138884334 | 0.86048133 | 0.246006152 |
| 36.5 | 1.818862628 | 0.183349425 | 0.712633054 | 0.175125167 | 1.634404768 | 0.158431508 | 0.661155158 | 0.170271477 |

**Supplementary Table S7 (i)** Direct motion in 100 mM NaCl stressed Tomato and their respective controls under RGBday dimGnight lighting condition

| Time | RGBStress1 | RGBStress2 | RGBStress3 | RGBStress4 | RGBStress5 | RGBStress6 | RGBStress7 | RGBStress8 | RGBStress9 | RGBStress10 | RGBStressTank | RGBControl1 | RGBControl2 | RGBControl3 |
|---|---|---|---|---|---|---|---|---|---|---|---|---|---|---|
| 18.0069444 | -0.177844 | -0.260765 | -0.524374 | -0.170428 | -0.152694 | -0.349393 | -0.046511 | -0.066308 | -0.196936 | -0.16369 | -0.223129 | -0.658897 | 0.153506 | 0.07075 |
| 18.0208333 | -0.252416 | -0.373509 | -0.494709 | -0.209322 | -0.195617 | -0.41389 | -0.078669 | -0.063329 | -0.328695 | -0.284156 | -0.120787 | -0.729586 | 0.119192 | -0.1428 |
| 18.0347222 | -0.323715 | -0.430014 | -0.488433 | -0.294146 | -0.271845 | -0.546793 | -0.134452 | -0.110637 | -0.46417 | -0.346241 | -0.18748 | -0.716039 | 0.058524 | -0.311773 |
| 18.0486111 | -0.33739 | -0.446274 | -0.441181 | -0.351736 | -0.317763 | -0.613954 | -0.173718 | -0.169759 | -0.528589 | -0.309337 | -0.249861 | -0.545501 | 0.024901 | -0.367482 |
| 18.0625 | -0.275326 | -0.39558 | -0.323776 | -0.29783 | -0.316724 | -0.544526 | -0.159854 | -0.172888 | -0.50535 | -0.287759 | -0.151397 | -0.361892 | 0.018943 | -0.28434 |
| 18.0763889 | -0.216148 | -0.332547 | -0.200471 | -0.206952 | -0.279447 | -0.43035 | -0.160726 | -0.158076 | -0.435486 | -0.299579 | 0.030329 | -0.242486 | 0.004736 | -0.170237 |
| 18.0902778 | -0.205752 | -0.277339 | -0.14878 | -0.159245 | -0.251537 | -0.347623 | -0.193341 | -0.163081 | -0.3624 | -0.298989 | 0.15912 | -0.127458 | -0.035416 | -0.121378 |
| 18.1041667 | -0.164723 | -0.226234 | -0.169233 | -0.166 | -0.245446 | -0.344344 | -0.181949 | -0.159707 | -0.278848 | -0.222375 | 0.088696 | -0.04631 | -0.067153 | -0.098436 |
| 18.1180556 | -0.100675 | -0.156794 | -0.202624 | -0.194637 | -0.214195 | -0.357434 | -0.148781 | -0.148338 | -0.178663 | -0.062005 | -0.172532 | -0.093727 | -0.081641 | -0.083278 |
| 18.1319444 | -0.067922 | -0.109011 | -0.188505 | -0.179733 | -0.166482 | -0.325253 | -0.147271 | -0.128869 | -0.093386 | 0.052618 | -0.298637 | -0.265267 | -0.135209 | -0.140116 |
| 18.1458333 | -0.048318 | -0.092285 | -0.146956 | -0.121311 | -0.143526 | -0.255919 | -0.130831 | -0.08681 | -0.06447 | 0.036693 | -0.133643 | -0.42409 | -0.187601 | -0.223166 |
| 18.1597222 | -0.04943 | -0.10651 | -0.140794 | -0.102718 | -0.147002 | -0.225072 | -0.094823 | -0.071492 | -0.08378 | -0.0077 | -0.078714 | -0.476318 | -0.14559 | -0.207943 |
| 18.1736111 | -0.062515 | -0.126097 | -0.148739 | -0.116038 | -0.13942 | -0.212426 | -0.078589 | -0.078754 | -0.097823 | -0.034636 | -0.119141 | -0.518263 | -0.087389 | -0.169501 |
| 18.1875 | -0.066918 | -0.117845 | -0.144005 | -0.112959 | -0.121041 | -0.192068 | -0.077702 | -0.06391 | -0.09551 | -0.079705 | -0.047555 | -0.585832 | -0.10142 | -0.231157 |
| 18.2013889 | -0.087603 | -0.140841 | -0.140053 | -0.107822 | -0.126908 | -0.166993 | -0.067483 | -0.058184 | -0.134047 | -0.152312 | 0.027404 | -0.612755 | -0.151132 | -0.295348 |
| 18.2152778 | -0.090925 | -0.161936 | -0.145924 | -0.111326 | -0.137999 | -0.15418 | -0.055523 | -0.069283 | -0.15617 | -0.15513 | 0.026734 | -0.584063 | -0.168015 | -0.258214 |
| 18.2291667 | -0.061693 | -0.123866 | -0.148261 | -0.117886 | -0.134217 | -0.152896 | -0.055039 | -0.069289 | -0.111696 | -0.0796 | 0.000465 | -0.538669 | -0.155579 | -0.210085 |
| 18.2430556 | -0.030118 | -0.078739 | -0.125432 | -0.08815 | -0.116048 | -0.12108 | -0.034643 | -0.035073 | -0.087837 | -0.089376 | 0.057952 | -0.474311 | -0.125741 | -0.169337 |
| 18.2569444 | -0.000615 | -0.029422 | -0.093374 | -0.055719 | -0.065763 | -0.079819 | -0.003593 | -0.001892 | -0.051897 | -0.053656 | 0.00649 | -0.366744 | -0.060588 | -0.080144 |
| 18.2708333 | 0.00005 | -0.004739 | -0.097416 | -0.077822 | -0.022153 | -0.080978 | -0.022335 | -0.023647 | 0.00691 | 0.077785 | -0.199631 | -0.2867 | 0.000268 | -0.022373 |
| 18.2847222 | -0.029188 | -0.037056 | -0.103031 | -0.092315 | -0.033755 | -0.08793 | -0.051023 | -0.05389 | -0.00047 | 0.092862 | -0.23424 | -0.234965 | 0.03135 | -0.036543 |
| 18.2986111 | -0.042166 | -0.066503 | -0.101243 | -0.079478 | -0.068834 | -0.085659 | -0.048786 | -0.058215 | -0.044106 | 0.01148 | -0.153112 | -0.221106 | 0.022218 | -0.061745 |
| 18.3125 | -0.043265 | -0.051824 | -0.095572 | -0.072733 | -0.074422 | -0.085003 | -0.041667 | -0.055122 | -0.063306 | -0.035001 | -0.098068 | -0.223958 | -0.031753 | -0.096323 |
| 18.3263889 | -0.042164 | -0.020621 | -0.085578 | -0.068878 | -0.045623 | -0.05627 | -0.046651 | -0.045697 | -0.049437 | -0.035304 | -0.072321 | -0.207384 | -0.049334 | -0.142378 |
| 18.3402778 | -0.040549 | -0.005964 | -0.045308 | -0.032221 | 0.002937 | 0.010581 | -0.053678 | -0.028139 | -0.028526 | -0.031189 | -0.017132 | -0.172185 | 0.005187 | -0.103699 |
| 18.3541667 | -0.033388 | -0.003381 | 0.022075 | 0.026782 | 0.04206 | 0.068313 | -0.041255 | -0.007608 | -0.029215 | -0.048545 | 0.111759 | -0.191269 | 0.051491 | -0.007663 |
| 18.3680556 | -0.019269 | -0.002399 | 0.045826 | 0.052531 | 0.039019 | 0.066885 | -0.014804 | 0.003618 | -0.047315 | -0.061571 | 0.176793 | -0.334352 | 0.006725 | -0.036108 |
| 18.3819444 | -0.042906 | -0.024212 | 0.006705 | 0.013756 | 0.021639 | 0.022645 | -0.030615 | -0.025455 | -0.068291 | -0.064414 | 0.097549 | -0.578166 | -0.101588 | -0.17186 |
| 18.3958333 | -0.095275 | -0.062242 | -0.050619 | -0.055004 | 0.004341 | -0.025258 | -0.105943 | -0.097499 | -0.096437 | -0.079427 | -0.052147 | -0.70225 | -0.128574 | -0.203349 |
| 18.4097222 | -0.111672 | -0.073199 | -0.048103 | -0.047833 | -0.003904 | -0.02419 | -0.13292 | -0.117281 | -0.137288 | -0.144701 | -0.026586 | -0.577539 | -0.034377 | -0.084395 |
| 18.4236111 | -0.048261 | 0.035226 | -0.074739 | -0.098149 | 0.001364 | -0.129177 | -0.103241 | -0.13153 | -0.01463 | 0.152983 | -0.497794 | -0.196578 | 0.031142 | -0.145779 |
| 18.4375 | 0.261592 | 0.669739 | -0.572587 | -0.904176 | -0.154105 | -1.038885 | -0.334198 | -0.63214 | 0.844363 | 2.638124 | -4.972013 | 0.290796 | -0.139719 | -1.34152 |
| 18.4513889 | 0.774006 | 1.754867 | -1.335338 | -2.407393 | -0.604901 | -2.704392 | -0.734302 | -1.445976 | 1.864969 | 7.079355 | -14.187483 | 0.123663 | -0.78047 | -3.848044 |
| 18.4652778 | 1.216296 | 1.305097 | -1.460006 | -2.473311 | -1.205037 | -2.975433 | -0.679215 | -1.468268 | 1.710243 | 8.106751 | -19.928118 | -1.509558 | -2.717994 | -5.41693 |
| 18.4791667 | 0.329758 | -0.285417 | -0.952722 | -1.107017 | -0.992398 | -1.609077 | -0.257309 | -0.829708 | 0.524803 | 3.132343 | -10.796403 | -2.077855 | -2.618002 | -3.599579 |
| 18.4930556 | -0.294157 | -0.430864 | -0.498085 | -0.262516 | -0.462012 | -0.909859 | -0.055201 | -0.313825 | -0.170296 | 0.310357 | -1.521903 | -0.908244 | -1.290178 | -1.526274 |
| 18.5069444 | -0.254343 | -0.186602 | -0.243932 | -0.067953 | -0.302958 | -0.707908 | 0.018417 | -0.065953 | -0.122541 | -0.041866 | 0.167232 | 0.088899 | -0.380428 | -0.622722 |
| 18.5208333 | -0.178393 | -0.041702 | -0.145434 | -0.038647 | -0.202628 | -0.478432 | -0.024714 | -0.008463 | -0.06155 | -0.000606 | 0.123761 | 0.920244 | -0.077873 | -0.286248 |
| 18.5347222 | -0.15097 | -0.000247 | -0.078608 | -0.04305 | -0.126547 | -0.258228 | -0.08178 | -0.021398 | -0.060123 | -0.020548 | 0.023313 | 1.47306 | 0.061812 | -0.066288 |
| 18.5486111 | -0.100311 | 0.011516 | 0.01713 | -0.047691 | -0.092923 | -0.131635 | -0.08514 | -0.022781 | -0.040041 | -0.039108 | 0.035956 | 1.223627 | 0.174112 | 0.06444 |
| 18.5625 | -0.029734 | 0.071306 | 0.153077 | -0.033824 | -0.075059 | -0.087081 | -0.048682 | 0.007903 | -0.002992 | -0.022411 | 0.121643 | 0.963063 | 0.226766 | 0.10472 |
| 18.5763889 | -0.021247 | 0.104803 | 0.329408 | -0.031074 | -0.062292 | -0.072358 | -0.020838 | 0.017063 | -0.020081 | -0.019298 | 0.205889 | 0.73287 | 0.20167 | 0.104082 |
| 18.5902778 | -0.038556 | 0.073022 | 0.451293 | -0.032585 | -0.046168 | -0.031552 | -0.001503 | 0.029352 | -0.041781 | -0.018066 | 0.239101 | 0.569897 | 0.197412 | 0.102499 |
| 18.6041667 | -0.027291 | 0.003159 | 0.50935 | -0.029703 | -0.050636 | 0.003484 | 0.027585 | 0.101119 | -0.015621 | 0.00771 | 0.198833 | 0.515675 | 0.298137 | 0.120836 |
| 18.6180556 | -0.038577 | -0.090981 | 0.486854 | -0.043318 | -0.117381 | -0.09822 | 0.067066 | 0.098194 | 0.000853 | 0.032438 | 0.114247 | 0.486046 | 0.425384 | 0.258047 |
| 18.6319444 | -0.058732 | -0.161188 | 0.437528 | -0.051073 | -0.236389 | -0.374797 | 0.104835 | 0.004374 | 0.018081 | 0.061009 | 0.077287 | 0.361951 | 0.500715 | 0.401139 |
| 18.6458333 | -0.074527 | -0.202776 | 0.360283 | -0.008169 | -0.38051 | -0.731286 | 0.098864 | -0.114201 | 0.026102 | 0.04139 | 0.133969 | 0.173414 | 0.491223 | 0.396621 |
| 18.6597222 | -0.115644 | -0.287434 | 0.318024 | 0.11335 | -0.507387 | -1.07622 | 0.074224 | -0.144441 | -0.022262 | -0.038157 | 0.275293 | 0.061636 | 0.416717 | 0.34379 |
| 18.6736111 | -0.176408 | -0.375008 | 0.333229 | 0.259865 | -0.595862 | -1.290213 | 0.051644 | -0.13641 | -0.104527 | -0.099032 | 0.347272 | 0.098577 | 0.338412 | 0.345062 |
| 18.6875 | -0.217013 | -0.360306 | 0.252454 | 0.230268 | -0.501053 | -1.002747 | 0.000829 | -0.089437 | -0.163469 | -0.08197 | 0.272029 | 0.319586 | 0.273103 | 0.280577 |
| 18.7013889 | -0.299863 | -0.256698 | 0.100114 | 0.080787 | -0.369887 | -0.638311 | -0.052051 | -0.051377 | -0.234049 | -0.053029 | 0.119757 | 0.51526 | 0.16487 | 0.106259 |
| 18.7152778 | -0.29734 | -0.218814 | -0.040388 | -0.095447 | -0.27863 | -0.332939 | -0.044562 | -0.041137 | -0.383051 | -0.093533 | -0.052995 | 0.597651 | 0.061719 | 0.058379 |
| 18.7291667 | -0.257254 | -0.205352 | -0.124658 | -0.239188 | -0.278359 | -0.204183 | -0.02154 | -0.04905 | -0.514193 | -0.156195 | -0.186281 | 0.675444 | 0.021042 | 0.185503 |
| 18.7430556 | -0.216264 | -0.154682 | -0.143323 | -0.285427 | -0.303731 | -0.129447 | 0.004724 | 0.024785 | -0.489965 | -0.144204 | -0.222396 | 0.777942 | -0.011469 | 0.225789 |
| 18.7569444 | -0.172787 | -0.078067 | -0.071594 | -0.235219 | -0.303623 | 0.025034 | 0.045563 | 0.116925 | -0.357455 | -0.098529 | -0.142964 | 0.900243 | -0.091434 | 0.094395 |
| 18.7708333 | -0.165584 | -0.065666 | 0.025345 | -0.121324 | -0.285591 | 0.243277 | 0.060742 | 0.123332 | -0.272126 | -0.100315 | -0.010455 | 0.846346 | -0.194234 | -0.050887 |
| 18.7847222 | -0.139733 | -0.073451 | 0.129712 | -0.024746 | -0.151425 | 0.430497 | 0.074758 | 0.094557 | -0.234752 | -0.125615 | 0.116279 | 0.472155 | -0.207534 | -0.025971 |
| 18.7986111 | -0.069541 | -0.011787 | 0.220707 | 0.054876 | 0.055478 | 0.493232 | 0.093235 | 0.089247 | -0.152237 | -0.106705 | 0.219411 | 0.282157 | -0.103421 | 0.17113 |
| 18.8125 | 0.004359 | 0.087464 | 0.292404 | 0.141618 | 0.184022 | 0.531488 | 0.106408 | 0.09234 | -0.020716 | -0.053921 | 0.291922 | 0.539147 | 0.014869 | 0.377061 |
| 18.8263889 | 0.049395 | 0.159607 | 0.307574 | 0.18851 | 0.195398 | 0.601108 | 0.078939 | 0.075675 | 0.072617 | -0.031369 | 0.291451 | 0.837526 | 0.089131 | 0.458319 |
| 18.8402778 | 0.092307 | 0.200879 | 0.282399 | 0.192448 | 0.190017 | 0.705515 | 0.034195 | 0.04944 | 0.119411 | -0.03813 | 0.211864 | 0.356623 | 0.158817 | 0.386338 |
| 18.8541667 | 0.15437 | 0.232853 | 0.26476 | 0.185475 | 0.253953 | 0.807103 | 0.003973 | 0.023211 | 0.181613 | -0.04955 | 0.121485 | -0.406631 | 0.245835 | 0.227846 |
| 18.8680556 | 0.249166 | 0.219892 | 0.236195 | 0.176579 | 0.37813 | 0.936949 | -0.004943 | -0.01482 | 0.247935 | -0.065307 | 0.069506 | -0.308121 | 0.383384 | 0.123103 |
| 18.8819444 | 0.377759 | 0.23793 | 0.220364 | 0.166324 | 0.474624 | 1.065169 | 0.012248 | -0.026403 | 0.264663 | -0.055285 | 0.070708 | 0.500672 | 0.481981 | 0.156441 |
| 18.8958333 | 0.431686 | 0.302611 | 0.196461 | 0.141285 | 0.463942 | 0.982991 | 0.026302 | -0.01991 | 0.193177 | -0.031342 | 0.086156 | 1.425891 | 0.51471 | 0.277246 |
| 18.9097222 | 0.49165 | 0.308954 | 0.156285 | 0.083397 | 0.369641 | 0.777358 | 0.006141 | -0.03554 | 0.106939 | -0.0415 | 0.068088 | 1.385038 | 0.456019 | 0.365337 |
| 18.9236111 | 0.357289 | 0.227992 | 0.166482 | 0.121407 | 0.312963 | 0.713309 | 0.027763 | 0.036275 | -0.045929 | -0.500957 | 1.275327 | 0.252411 | 0.41444 | 0.409839 |
| 18.9375 | -0.373396 | 0.018497 | 0.657619 | 0.762531 | 0.631124 | 1.254644 | 0.381976 | 0.695177 | -0.768157 | -3.351529 | 8.783574 | -0.377637 | 0.868494 | 0.834554 |
| 18.9513889 | -1.226931 | -0.448269 | 1.716157 | 2.014949 | 0.942117 | 2.917171 | 1.223682 | 1.981035 | -1.876959 | -8.037728 | 18.452403 | -0.570623 | 1.680743 | 1.888235 |
| 18.9652778 | -1.147324 | -0.578107 | 1.85915 | 2.037157 | 0.713433 | 2.76593 | 1.395722 | 2.155876 | -2.065661 | -7.5585 | 16.347115 | -0.558164 | 0.846186 | 1.96744 |
| 18.9791667 | -0.41816 | -0.125995 | 0.851126 | 0.888358 | 0.210474 | 1.042524 | 0.502082 | 0.801157 | -1.096277 | -2.932391 | 6.080496 | -0.312959 | 0.302937 | 0.924594 |
| 18.9930556 | -0.177092 | -0.43178 | -0.169686 | -0.075488 | -0.07917 | -0.082017 | -0.06169 | -0.04417 | -0.186927 | -0.316296 | 0.259783 | -0.157203 | 0.055401 | 0.287838 |
| 19.0069444 | -0.168812 | -0.340773 | -0.438658 | -0.215974 | -0.073246 | -0.182405 | -0.107935 | -0.122753 | -0.045201 | 0.084926 | -0.695411 | -0.173368 | 0.177302 | 0.162152 |
| 19.0208333 | -0.160919 | -0.155898 | -0.377651 | -0.128968 | 0.040781 | -0.062798 | -0.051557 | -0.050411 | -0.109373 | -0.083039 | -0.294313 | -0.209924 | 0.187353 | 0.04543 |
| 19.0347222 | -0.161112 | -0.201022 | -0.310701 | -0.101327 | 0.112766 | -0.033952 | -0.054843 | -0.056513 | -0.149422 | -0.193599 | -0.103436 | -0.236288 | 0.07268 | -0.070341 |
| 19.0486111 | -0.163028 | -0.29191 | -0.272149 | -0.123709 | 0.066067 | -0.04266 | -0.104826 | -0.10504 | -0.170246 | -0.222084 | -0.056075 | -0.279546 | -0.06906 | -0.143758 |
| 19.0625 | -0.144415 | -0.323973 | -0.243928 | -0.13073 | -0.015658 | -0.043014 | -0.124605 | -0.132195 | -0.174687 | -0.178748 | -0.062562 | -0.335389 | -0.270718 | -0.228941 |
| 19.0763889 | -0.111534 | -0.317557 | -0.182179 | -0.092966 | -0.039229 | -0.038596 | -0.094982 | -0.112095 | -0.132023 | -0.09782 | -0.044477 | -0.347317 | -0.408903 | -0.268484 |
| 19.0902778 | -0.095589 | -0.22927 | -0.103546 | -0.041271 | -0.027314 | -0.025368 | -0.057392 | -0.073053 | -0.07398 | -0.047761 | 0.011172 | -0.303782 | -0.326187 | -0.16453 |
| 19.1041667 | -0.096477 | -0.154869 | -0.08753 | -0.029017 | -0.021784 | -0.028744 | -0.043355 | -0.046463 | -0.041916 | -0.039712 | -0.007202 | -0.268489 | -0.220515 | -0.059138 |
| 19.1180556 | -0.09272 | -0.111959 | -0.119261 | -0.059377 | -0.042708 | -0.072036 | -0.051891 | -0.055298 | -0.038084 | -0.036855 | -0.113585 | -0.237423 | -0.233074 | -0.113826 |
| 19.1319444 | -0.073568 | -0.098222 | -0.114091 | -0.070401 | -0.060738 | -0.093756 | -0.070502 | -0.06829 | -0.035281 | -0.015581 | -0.125073 | -0.251733 | -0.300803 | -0.21112 |
| 19.1458333 | -0.054737 | -0.065561 | -0.083219 | -0.049049 | -0.057336 | -0.070254 | -0.069557 | -0.051478 | -0.036396 | -0.017135 | -0.039905 | -0.32967 | -0.323395 | -0.205422 |
| 19.1597222 | -0.040763 | 0.082852 | -0.060655 | -0.030192 | -0.045696 | -0.03884 | -0.050107 | -0.009648 | -0.051334 | -0.062688 | 0.03548 | -0.361785 | -0.314997 | -0.18294 |
| 19.1736111 | -0.035509 | 0.295824 | -0.018006 | -0.025947 | -0.038761 | -0.017646 | -0.041392 | 0.018792 | -0.059729 | -0.101094 | 0.13331 | -0.309956 | -0.271864 | -0.223256 |
| 19.1875 | -0.070885 | 0.245965 | -0.029858 | -0.05687 | -0.062084 | -0.053624 | -0.076903 | -0.027157 | -0.075982 | -0.096226 | 0.096691 | -0.254606 | -0.242728 | -0.216771 |
| 19.2013889 | -0.077358 | 0.079096 | -0.074341 | -0.062749 | -0.057309 | -0.070807 | -0.072538 | -0.054713 | -0.059357 | -0.043776 | -0.046952 | -0.243999 | -0.243353 | -0.169869 |
| 19.2152778 | -0.029913 | 0.025765 | -0.051608 | -0.010542 | 0.00012 | 0.005014 | -0.003578 | 0.015915 | -0.023474 | -0.041276 | 0.018377 | -0.302763 | -0.309116 | -0.224325 |
| 19.2291667 | -0.037968 | -0.024086 | -0.054501 | -0.027239 | 0.002547 | 0.019459 | 0.001759 | 0.02586 | -0.04032 | -0.080755 | 0.042031 | -0.404769 | -0.395356 | -0.365787 |
| 19.2430556 | -0.050257 | -0.036085 | -0.124001 | -0.126373 | -0.048485 | -0.078479 | -0.045177 | -0.048505 | -0.038541 | -0.016174 | -0.275804 | -0.417385 | -0.395016 | -0.422707 |
| 19.2569444 | 0.003769 | 0.016639 | -0.122956 | -0.115199 | -0.044089 | -0.096545 | -0.028734 | -0.034723 | 0.005295 | 0.040947 | -0.347702 | -0.300961 | -0.28091 | -0.318959 |
| 19.2708333 | 0.008572 | -0.004472 | 0.012092 | 0.021008 | 0.014448 | 0.015911 | 0.023281 | 0.041162 | -0.00147 | -0.024356 | 0.093152 | -0.182754 | -0.16615 | -0.137168 |
| 19.2847222 | -0.037619 | -0.040643 | 0.073724 | 0.052178 | 0.015468 | 0.037654 | 0.014202 | 0.029507 | -0.018552 | -0.033127 | 0.249334 | -0.071544 | -0.054685 | -0.032666 |
| 19.2986111 | -0.060052 | -0.036773 | -0.012822 | -0.041049 | -0.042911 | -0.053203 | -0.055916 | -0.043066 | 0.00626 | 0.071986 | -0.052418 | 0.000208 | 0.022892 | -0.053051 |
| 19.3125 | -0.046919 | -0.037087 | -0.041263 | -0.050871 | -0.049284 | -0.06369 | -0.065583 | -0.043604 | 0.022385 | 0.103785 | -0.103521 | -0.057386 | -0.007728 | -0.030827 |
| 19.3263889 | -0.013321 | -0.013375 | 0.015232 | 0.024258 | -0.006304 | -0.001513 | 0.001187 | 0.026723 | -0.004295 | -0.004628 | 0.128639 | -0.131651 | -0.070134 | 0.017936 |
| 19.3402778 | -0.001995 | 0.00953 | 0.061001 | 0.072793 | 0.021942 | 0.040343 | 0.042832 | 0.06046 | -0.029861 | -0.081969 | 0.211251 | -0.10812 | -0.064344 | 0.000489 |
| 19.3541667 | 0.000621 | 0.010737 | 0.066448 | 0.070444 | 0.029905 | 0.04617 | 0.035221 | 0.042708 | -0.016322 | -0.054582 | 0.101956 | -0.040049 | 0.001321 | -0.024491 |
| 19.3680556 | 0.000723 | 0.024399 | 0.032301 | 0.031134 | 0.015257 | 0.016949 | 0.014502 | 0.014257 | -0.007113 | -0.011886 | -0.018832 | 0.004486 | 0.07105 | -0.042674 |
| 19.3819444 | -0.015323 | 0.034705 | 0.000643 | -0.006786 | -0.015488 | -0.019749 | -0.001979 | -0.002336 | -0.041426 | -0.033027 | 0.015438 | -0.020999 | 0.094909 | -0.099145 |
| 19.3958333 | -0.038951 | 0.03832 | -0.023603 | -0.038084 | -0.036663 | -0.040623 | -0.017809 | -0.018755 | -0.063787 | -0.062312 | 0.06434 | -0.146336 | 0.042182 | -0.171862 |
| 19.4097222 | -0.039162 | 0.046336 | -0.047858 | -0.061626 | -0.028237 | -0.033679 | -0.036755 | -0.037525 | -0.023862 | -0.001201 | -0.033478 | -0.266488 | -0.030316 | -0.202356 |
| 19.4236111 | 0.038684 | 0.143769 | -0.106954 | -0.143822 | -0.040557 | -0.116049 | -0.088313 | -0.124828 | 0.167446 | 0.417505 | -0.81532 | -0.209248 | -0.013858 | -0.267618 |
| 19.4375 | 0.294948 | 0.584199 | -0.602042 | -0.894196 | -0.374601 | -1.016593 | -0.540109 | -0.85469 | 1.06911 | 2.707095 | -5.817596 | -0.017536 | 0.082127 | -1.036704 |
| 19.4513889 | 0.793735 | 1.437618 | -1.335187 | -2.201285 | -0.894063 | -2.609508 | -1.370287 | -2.234765 | 2.196241 | 6.761564 | -15.807102 | -0.06987 | -0.3082 | -2.612554 |
| 19.4652778 | 1.153202 | 1.779734 | -1.722394 | -2.165628 | -1.341012 | -2.615359 | -1.430269 | -2.400072 | 1.904404 | 7.706843 | -19.867264 | -1.043984 | -1.27128 | -3.051432 |
| 19.4791667 | 0.561809 | 0.405189 | -0.977711 | -0.990439 | -0.756901 | -0.902064 | -0.619762 | -1.153808 | 0.717983 | 3.398283 | -10.565356 | -1.096539 | -1.100235 | -1.500126 |
| 19.4930556 | -0.029045 | 0.226957 | -0.36024 | -0.166088 | 0.031616 | -0.036899 | -0.067305 | -0.285368 | 0.081022 | 0.436282 | -1.501075 | -0.46078 | 0.009185 | -0.207686 |
| 19.5069444 | -0.045154 | 0.216647 | -0.097559 | 0.005864 | 0.193637 | 0.089512 | 0.082123 | -0.040997 | 0.060219 | 0.030613 | 0.045968 | -0.074157 | 0.533309 | 0.231428 |
| 19.5208333 | -0.067069 | 0.101324 | -0.002458 | -0.072909 | 0.114894 | 0.017346 | 0.063017 | -0.035526 | 0.045573 | 0.004031 | 0.025907 | 0.078894 | 0.492801 | 0.286146 |
| 19.5347222 | -0.040295 | 0.020014 | 0.200924 | -0.131179 | 0.061649 | 0.033962 | 0.002915 | -0.048547 | 0.014659 | 0.00111 | 0.033787 | 0.161387 | 0.353638 | 0.279412 |
| 19.5486111 | 0.102794 | 0.004236 | 0.562846 | -0.107806 | 0.140293 | 0.273634 | 0.019967 | -0.013951 | 0.011328 | 0.027751 | 0.113345 | 0.195376 | 0.256274 | 0.284575 |
| 19.5625 | 0.27021 | 0.015975 | 1.007614 | -0.045363 | 0.325751 | 0.719388 | 0.068605 | 0.026281 | 0.045081 | 0.090081 | 0.150222 | 0.18776 | 0.173269 | 0.287469 |
| 19.5763889 | 0.375372 | 0.034294 | 1.40263 | 0.051452 | 0.534015 | 1.300016 | 0.090835 | 0.043562 | 0.093737 | 0.12535 | 0.153982 | 0.146762 | 0.068226 | 0.267548 |
| 19.5902778 | 0.598165 | 0.173193 | 1.881981 | 0.200438 | 0.777726 | 2.008982 | 0.050419 | 0.07561 | 0.268848 | 0.209374 | 0.271577 | 0.121057 | 0.007628 | 0.231166 |
| 19.6041667 | 0.940624 | 0.314955 | 2.102438 | 0.356469 | 0.9374 | 2.605026 | 0.032282 | 0.086642 | 0.570222 | 0.386247 | 0.393067 | 0.136845 | 0.033312 | 0.231531 |
| 19.6180556 | 1.31888 | 0.341779 | 1.814653 | 0.415892 | 0.843695 | 2.277968 | 0.003279 | 0.068516 | 0.883897 | 0.611911 | 0.387622 | 0.172078 | 0.073285 | 0.283393 |
| 19.6319444 | 1.401484 | 0.252452 | 1.462625 | 0.544599 | 0.543948 | 1.341499 | 0.07979 | 0.0974 | 1.003328 | 0.79895 | 0.40615 | 0.211771 | 0.083084 | 0.359044 |
| 19.6458333 | 1.235782 | 0.192608 | 1.174788 | 0.73109 | 0.32754 | 0.750802 | 0.162073 | 0.142826 | 0.893319 | 0.791771 | 0.406095 | 0.280947 | 0.151085 | 0.451163 |
| 19.6597222 | 0.947833 | 0.182052 | 0.95891 | 0.966076 | 0.130053 | 0.220661 | 0.174511 | 0.159659 | 0.691501 | 0.660071 | 0.280376 | 0.329617 | 0.302544 | 0.563369 |
| 19.6736111 | 0.554531 | 0.172908 | 0.708573 | 1.007931 | -0.0168 | -0.211233 | 0.129103 | 0.105406 | 0.464265 | 0.438769 | 0.121745 | 0.29787 | 0.424864 | 0.815758 |
| 19.6875 | 0.191943 | 0.233559 | 0.429255 | 0.669147 | -0.064732 | -0.381968 | 0.098184 | 0.034687 | 0.280313 | 0.247575 | 0.116744 | 0.218128 | 0.414816 | 0.639836 |
| 19.7013889 | -0.039396 | 0.299378 | 0.243009 | 0.323235 | 0.048361 | -0.173237 | 0.098659 | 0.033532 | 0.18858 | 0.15901 | 0.165391 | 0.164502 | 0.312456 | 0.319179 |
| 19.7152778 | -0.102799 | 0.33973 | 0.115549 | 0.036324 | 0.215216 | 0.250251 | 0.073133 | 0.035892 | 0.087443 | 0.093064 | 0.0679 | 0.142237 | 0.111013 | 0.248595 |
| 19.7291667 | -0.009905 | 0.338826 | -0.083245 | -0.182982 | 0.329996 | 0.712525 | 0.020565 | 0.032512 | -0.012899 | 0.031805 | -0.105293 | 0.152388 | -0.045258 | 0.304404 |
| 19.7430556 | 0.147012 | 0.333225 | -0.323464 | -0.29696 | 0.269864 | 0.605968 | -0.002271 | 0.063619 | -0.016938 | 0.023726 | -0.146193 | 0.175439 | -0.023587 | 0.27374 |
| 19.7569444 | 0.131877 | 0.311773 | -0.481112 | -0.324934 | 0.049308 | 0.125322 | 0.014076 | 0.072888 | 0.030704 | 0.01127 | -0.036874 | 0.20651 | 0.096465 | 0.198775 |
| 19.7708333 | -0.006567 | 0.222386 | -0.358375 | -0.186471 | -0.135024 | -0.309902 | 0.021954 | 0.012116 | 0.047221 | -0.044885 | 0.119611 | 0.240411 | 0.173094 | 0.171123 |
| 19.7847222 | -0.082755 | 0.147327 | 0.013331 | 0.035774 | -0.16477 | -0.368599 | 0.013373 | -0.023907 | 0.047933 | -0.071173 | 0.213885 | 0.246626 | 0.120439 | 0.155726 |
| 19.7986111 | -0.04395 | 0.184344 | 0.345225 | 0.143873 | -0.129993 | -0.290189 | 0.000042 | -0.008292 | 0.048438 | -0.022195 | 0.196986 | 0.222332 | 0.03101 | 0.111529 |
| 19.8125 | 0.006561 | 0.255305 | 0.315191 | 0.03227 | -0.018559 | -0.056072 | -0.016635 | 0.018266 | 0.014548 | 0.030315 | 0.069384 | 0.190153 | 0.008521 | 0.044474 |
| 19.8263889 | -0.038408 | 0.297479 | -0.017518 | -0.139543 | 0.09427 | 0.252575 | -0.033754 | 0.001509 | -0.092711 | -0.009928 | -0.151024 | 0.16153 | 0.014996 | 0.010517 |
| 19.8402778 | -0.130186 | 0.267772 | -0.398494 | -0.263432 | 0.111296 | 0.315242 | -0.039045 | -0.028604 | -0.216341 | -0.124935 | -0.272002 | 0.150864 | -0.022876 | 0.036385 |
| 19.8541667 | -0.187759 | 0.223302 | -0.652927 | -0.402267 | 0.010099 | 0.003349 | -0.044471 | -0.033071 | -0.25556 | -0.235127 | -0.149002 | 0.138131 | -0.165986 | 0.059342 |
| 19.8680556 | -0.33035 | 0.168669 | -0.759422 | -0.52804 | -0.148361 | -0.39205 | -0.071623 | -0.03406 | -0.243331 | -0.331573 | 0.066156 | 0.094149 | -0.375356 | 0.010971 |
| 19.8819444 | -0.617668 | 0.054682 | -0.742932 | -0.552211 | -0.292071 | -0.75903 | -0.053948 | -0.040322 | -0.291165 | -0.465211 | 0.178806 | 0.101785 | -0.341526 | -0.091514 |
| 19.8958333 | -0.930452 | -0.075905 | -0.824554 | -0.507649 | -0.366643 | -0.947761 | -0.006889 | -0.056518 | -0.491463 | -0.648654 | 0.071143 | 0.191608 | -0.148453 | -0.144582 |
| 19.9097222 | -0.993725 | -0.093445 | -1.022604 | -0.479688 | -0.334191 | -0.838596 | -0.036715 | -0.10182 | -0.821513 | -0.788284 | -0.278908 | 0.275618 | -0.002019 | -0.14828 |
| 19.9236111 | -0.923375 | -0.050699 | -1.182629 | -0.433146 | -0.256225 | -0.704664 | -0.070898 | -0.07572 | -1.294343 | -1.26968 | 0.511783 | 0.269599 | -0.053516 | -0.127971 |
| 19.9375 | -1.438399 | -0.307208 | -1.037062 | -0.15864 | -0.041949 | -0.005202 | 0.204833 | 0.490664 | -1.931696 | -4.019758 | 7.785267 | 0.378901 | -0.058111 | 0.167394 |
| 19.9513889 | -2.057585 | -1.77598 | -0.367211 | 0.575962 | 0.235362 | 2.140898 | 1.089957 | 1.75226 | -2.247004 | -8.037673 | 15.305208 | 0.469533 | 0.165233 | 0.803103 |
| 19.9652778 | -1.290113 | -1.264986 | 0.078996 | 1.060073 | 0.316749 | 2.371103 | 1.21651 | 1.894141 | -2.247377 | -7.10248 | 12.22301 | 0.209396 | 0.099081 | 1.136361 |
| 19.9791667 | -0.635713 | -0.58369 | -0.309481 | 0.535357 | -0.08295 | 0.450851 | 0.502392 | 0.812606 | -1.040747 | -2.926155 | 4.715642 | -0.006058 | -0.111497 | 0.5673 |
| 19.9930556 | -0.430819 | -0.171397 | -0.614183 | 0.085667 | -0.37126 | -0.643207 | 0.090164 | 0.09833 | -0.24491 | -0.53976 | 0.891728 | 0.009998 | -0.001291 | 0.050535 |
| 20.0069444 | -0.24287 | -0.146218 | -0.454213 | 0.044631 | -0.364233 | -0.65598 | 0.046091 | 0.015438 | -0.114212 | -0.165149 | 0.327208 | 0.012834 | -0.016136 | -0.109371 |
| 20.0208333 | -0.10727 | -0.137059 | -0.340283 | 0.042687 | -0.262258 | -0.45009 | 0.060239 | 0.028345 | -0.118474 | -0.164633 | 0.198206 | 0.012049 | -0.032637 | -0.153516 |
| 20.0347222 | -0.037577 | -0.071595 | -0.386611 | -0.016846 | -0.176549 | -0.252925 | 0.033913 | 0.003918 | -0.127539 | -0.151501 | 0.013398 | 0.006259 | -0.019557 | -0.17496 |
| 20.0486111 | 0.013095 | 0.007428 | -0.397998 | -0.096082 | -0.108971 | -0.159254 | -0.029329 | -0.031055 | -0.059207 | -0.03822 | -0.118101 | -0.043463 | -0.024664 | -0.138407 |
| 20.0625 | 0.0417 | 0.03973 | -0.277732 | -0.133423 | -0.115924 | -0.222806 | -0.073456 | -0.070867 | 0.017266 | 0.080533 | -0.149169 | -0.109484 | -0.039449 | -0.089706 |
| 20.0763889 | 0.048149 | 0.032551 | -0.126659 | -0.069992 | -0.147586 | -0.281795 | -0.050094 | -0.053478 | 0.007316 | 0.061443 | -0.026768 | -0.163863 | -0.030664 | -0.060004 |
| 20.0902778 | 0.02104 | -0.0167 | -0.008509 | 0.076875 | -0.122314 | -0.183771 | 0.005617 | -0.002796 | -0.048603 | -0.019953 | 0.097202 | -0.170872 | -0.03685 | -0.051362 |
| 20.1041667 | -0.034348 | -0.066939 | 0.064675 | 0.187713 | -0.065536 | -0.067454 | 0.003305 | 0.01444 | -0.096342 | -0.037486 | 0.092705 | -0.165958 | -0.06969 | -0.062573 |
| 20.1180556 | -0.062398 | -0.036563 | 0.051587 | 0.185701 | -0.049802 | -0.066581 | -0.053348 | -0.013402 | -0.101229 | 0.006773 | 0.007881 | -0.175555 | -0.095136 | -0.121105 |
| 20.1319444 | -0.043985 | 0.019441 | -0.005514 | 0.116415 | -0.095258 | -0.147571 | -0.094632 | -0.03915 | -0.082469 | 0.02768 | -0.008058 | -0.191625 | -0.102533 | -0.186011 |
| 20.1458333 | -0.029771 | -0.001269 | -0.013606 | 0.075529 | -0.133692 | -0.197364 | -0.09687 | -0.034584 | -0.129055 | -0.072685 | 0.090776 | -0.17949 | -0.092987 | -0.154996 |
| 20.1597222 | -0.012574 | -0.028981 | 0.035066 | 0.081075 | -0.125982 | -0.177939 | -0.077929 | -0.007468 | -0.176511 | -0.142285 | 0.125646 | -0.169391 | -0.098065 | -0.087811 |
| 20.1736111 | 0.023164 | -0.025225 | 0.073984 | 0.061569 | -0.121184 | -0.151746 | -0.07742 | -0.005687 | -0.16803 | -0.086404 | -0.029139 | -0.189667 | -0.131547 | -0.084407 |
| 20.1875 | -0.000313 | -0.058525 | 0.0902 | -0.010983 | -0.15774 | -0.157632 | -0.120176 | -0.085617 | -0.200112 | -0.055248 | -0.235382 | -0.238436 | -0.191528 | -0.114092 |
| 20.2013889 | -0.031564 | -0.08127 | 0.141989 | -0.071438 | -0.156351 | -0.123316 | -0.141417 | -0.152846 | -0.198463 | -0.032486 | -0.296519 | -0.31967 | -0.271394 | -0.167167 |
| 20.2152778 | 0.019251 | -0.044553 | 0.271159 | -0.013113 | -0.070566 | -0.020375 | -0.079517 | -0.095183 | -0.091572 | 0.038091 | -0.146749 | -0.347774 | -0.270303 | -0.198 |
| 20.2291667 | 0.077281 | -0.007669 | 0.387243 | 0.094758 | 0.009173 | 0.05534 | 0.003377 | -0.000426 | 0.014946 | 0.080126 | 0.0548 | -0.281833 | -0.18284 | -0.140104 |
| 20.2430556 | 0.077047 | -0.012341 | 0.371885 | 0.115118 | 0.01822 | 0.07458 | 0.02121 | 0.016102 | 0.054954 | 0.062705 | 0.077908 | -0.255564 | -0.161896 | -0.122833 |
| 20.2569444 | 0.061083 | -0.040589 | 0.290462 | 0.073972 | 0.010159 | 0.085335 | -0.002135 | -0.016304 | 0.072665 | 0.048335 | -0.04849 | -0.320336 | -0.259801 | -0.213734 |
| 20.2708333 | 0.081826 | -0.023684 | 0.254065 | 0.05953 | 0.021478 | 0.116139 | 0.003421 | -0.018661 | 0.136033 | 0.079164 | -0.058774 | -0.318778 | -0.316259 | -0.257076 |
| 20.2847222 | 0.098308 | 0.010226 | 0.276098 | 0.048913 | 0.024565 | 0.131159 | 0.023317 | -0.007417 | 0.196344 | 0.115934 | 0.000925 | -0.217733 | -0.235176 | -0.173645 |
| 20.2986111 | 0.103367 | 0.002177 | 0.258434 | 0.012901 | 0.031866 | 0.156868 | 0.026777 | -0.012842 | 0.211583 | 0.114131 | -0.014616 | -0.132122 | -0.119595 | -0.064965 |
| 20.3125 | 0.155478 | -0.019128 | 0.222394 | -0.011414 | 0.069991 | 0.228041 | 0.0366 | -0.00781 | 0.223518 | 0.143843 | -0.047785 | -0.10297 | -0.065895 | -0.037147 |
| 20.3263889 | 0.197769 | -0.052928 | 0.286153 | -0.011657 | 0.101068 | 0.281635 | 0.048023 | 0.008476 | 0.233664 | 0.182139 | -0.039638 | -0.225136 | -0.186853 | -0.206143 |
| 20.3402778 | 0.199202 | -0.081601 | 0.493456 | 0.068221 | 0.133727 | 0.348145 | 0.065677 | 0.05842 | 0.261144 | 0.197502 | 0.053638 | -0.452788 | -0.428602 | -0.480777 |
| 20.3541667 | 0.207538 | -0.057818 | 0.730081 | 0.243143 | 0.228858 | 0.483416 | 0.091421 | 0.105356 | 0.377447 | 0.298106 | 0.137768 | -0.429888 | -0.427027 | -0.464536 |
| 20.3680556 | 0.251692 | 0.002356 | 0.922686 | 0.414369 | 0.312707 | 0.60548 | 0.109545 | 0.102274 | 0.518333 | 0.416345 | 0.106145 | -0.125432 | -0.121225 | -0.155854 |
| 20.3819444 | 0.361138 | 0.070667 | 1.123224 | 0.561331 | 0.3443 | 0.665322 | 0.123766 | 0.101169 | 0.600238 | 0.489106 | 0.048577 | 0.167816 | 0.169335 | 0.105398 |
| 20.3958333 | 0.480456 | 0.111196 | 1.329622 | 0.628796 | 0.38567 | 0.651007 | 0.142061 | 0.098645 | 0.645458 | 0.555898 | 0.075032 | 0.334526 | 0.292031 | 0.20333 |
| 20.4097222 | 0.576611 | 0.147588 | 1.373926 | 0.626089 | 0.421451 | 0.719198 | 0.139408 | 0.111892 | 0.810488 | 0.703479 | 0.176713 | 0.410427 | 0.293022 | 0.189806 |
| 20.4236111 | 0.732881 | 0.248394 | 1.258321 | 0.4863 | 0.461514 | 0.768932 | 0.076515 | 0.047488 | 1.17802 | 1.18995 | -0.210132 | 0.479396 | 0.284264 | 0.031849 |
| 20.4375 | 1.273328 | 0.661875 | 0.473582 | -0.557611 | 0.55496 | 0.070629 | -0.237919 | -0.571101 | 2.12423 | 3.697694 | -3.858669 | 0.484615 | 0.189245 | -0.761347 |
| 20.4513889 | 1.913869 | 1.32779 | -0.653883 | -2.185047 | 0.468665 | -1.343309 | -0.759141 | -1.605996 | 3.26921 | 8.162852 | -11.771305 | -0.191837 | 0.051176 | -2.68524 |
| 20.4652778 | 3.003198 | 1.415973 | -0.341811 | -2.211561 | 0.213378 | -1.432906 | -0.566902 | -1.655244 | 2.563928 | 8.776207 | -15.172125 | -0.879164 | -1.336313 | -3.168209 |
| 20.4791667 | 1.957328 | 0.309403 | 1.245292 | -0.594582 | -0.003624 | -0.128934 | -0.089202 | -0.701262 | 1.46923 | 3.887231 | -7.336182 | -0.103287 | -0.374298 | -1.186111 |
| 20.4930556 | 1.672087 | 0.639918 | 2.209112 | 0.329934 | 0.660015 | 0.881022 | 0.152286 | -0.018608 | 1.241732 | 1.400751 | -0.590853 | 1.392868 | 0.940974 | 0.262475 |
| 20.5069444 | 1.198792 | 0.580493 | 2.249135 | 0.352452 | 0.583596 | 0.897409 | 0.182592 | 0.097887 | 1.686607 | 1.370934 | 0.941082 | 2.078393 | 1.077214 | 0.531538 |
| 20.5208333 | 0.738582 | 0.494625 | 1.298215 | 0.193336 | 0.092451 | -0.088478 | 0.059654 | 0.158198 | 1.174934 | 1.38926 | 0.451625 | 1.771709 | 0.888404 | 0.381281 |
| 20.5347222 | 0.435975 | 0.325555 | 0.579517 | 0.057572 | 0.138936 | 0.347907 | 0.144656 | 0.11346 | 0.260718 | 0.577385 | -0.41264 | 0.990923 | 0.730005 | 0.424533 |
| 20.5486111 | 0.484487 | 0.409531 | 0.091177 | 0.015826 | 0.08847 | 0.045912 | 0.204072 | 0.09653 | -0.477761 | 0.129766 | -0.741333 | 0.365296 | 0.372137 | 0.641617 |
| 20.5625 | 0.58326 | 0.593572 | 0.538537 | 0.261301 | 0.414317 | 0.938197 | 0.179383 | 0.104357 | -0.111327 | 0.087018 | -0.172855 | 0.408352 | 0.103189 | 1.074687 |
| 20.5763889 | 0.531665 | 0.575499 | 1.10245 | 0.456759 | 0.237232 | 0.415306 | 0.083274 | 0.09238 | 0.464407 | 0.444039 | 0.394511 | 0.475838 | 0.199751 | 1.320533 |
| 20.5902778 | 0.343055 | 0.49432 | 1.005719 | 0.34072 | 0.17683 | 0.217423 | 0.040436 | 0.077821 | 0.282495 | 0.41656 | 0.293079 | 0.297455 | 0.56253 | 1.008237 |
| 20.6041667 | 0.207083 | 0.430284 | 0.586961 | 0.120337 | -0.020688 | -0.079213 | 0.136524 | 0.067742 | 0.068476 | 0.134809 | 0.220723 | 0.230179 | 0.628227 | 0.611745 |
| 20.6180556 | 0.425329 | 0.860142 | 0.740148 | 0.131641 | 0.201057 | 0.489232 | 0.136594 | 0.083583 | 0.522293 | 0.103373 | 0.56623 | 0.374782 | 0.320679 | 0.513266 |
| 20.6319444 | 0.361926 | 0.82487 | 0.943924 | 0.347219 | 0.158006 | 0.319302 | 0.029533 | 0.088772 | 0.758827 | 0.186571 | 0.637644 | 0.425866 | 0.073886 | 0.61322 |
| 20.6458333 | -0.145126 | 0.20757 | 0.414629 | 0.21214 | 0.137828 | 0.30972 | -0.026058 | -0.014449 | 0.088518 | -0.041791 | 0.105976 | 0.007758 | 0.220468 | 0.542702 |
| 20.6597222 | -0.290402 | -0.024635 | -0.569828 | -0.13635 | 0.224656 | 0.475326 | -0.036959 | -0.047983 | -0.417056 | -0.326577 | -0.479201 | -0.258487 | 0.409055 | 0.20879 |
| 20.6736111 | -0.081856 | 0.206899 | -1.118505 | -0.352256 | 0.027881 | 0.041379 | -0.029062 | -0.042491 | -0.186518 | -0.34246 | -0.607527 | 0.22399 | 0.300085 | 0.196832 |
| 20.6875 | 0.28803 | 0.680658 | -0.643157 | 0.053131 | 0.036532 | 0.133244 | -0.078162 | 0.00826 | 0.071383 | -0.189375 | -0.303417 | 1.347012 | 0.06001 | 0.57002 |
| 20.7013889 | 0.200935 | 0.575056 | 0.306316 | 0.443611 | 0.123971 | 0.18657 | -0.05213 | 0.010557 | -0.31048 | -0.322871 | 0.022815 | 1.360639 | 0.090164 | 0.73415 |
| 20.7152778 | -0.218787 | 0.022529 | 0.450054 | 0.240553 | 0.428702 | 0.839675 | -0.039865 | -0.044482 | -0.584594 | -0.564063 | 0.118758 | -0.33821 | 0.189075 | 0.368863 |
| 20.7291667 | -0.438764 | -0.327458 | -0.177964 | -0.066082 | 0.247057 | 0.525718 | 0.012967 | -0.054213 | -0.163689 | -0.455072 | -0.065476 | -1.69923 | 0.208654 | 0.074254 |
| 20.7430556 | -0.146476 | 0.109546 | -0.661811 | -0.128075 | -0.251178 | -0.5223 | -0.105376 | -0.052605 | 0.388535 | 0.087135 | -0.401212 | -0.708391 | 0.009186 | -0.015766 |
| 20.7569444 | 0.33031 | 0.830654 | -0.5401 | -0.06349 | -0.597784 | -1.224287 | -0.223768 | -0.061159 | 0.103375 | 0.275324 | -0.434147 | 1.00105 | -0.086009 | -0.016601 |
| 20.7708333 | 0.25258 | 0.647922 | 0.340517 | 0.106051 | -0.190778 | -0.50886 | -0.215746 | -0.149385 | -0.22472 | 0.097689 | 0.210305 | 1.941762 | 0.142354 | 0.03219 |
| 20.7847222 | -0.183012 | -0.168564 | 0.781943 | 0.290467 | 0.12685 | 0.330354 | -0.14883 | -0.184764 | -0.044734 | -0.10921 | 0.714022 | 0.347125 | -0.053191 | 0.059781 |
| 20.7986111 | -0.408081 | -0.352285 | 0.22607 | 0.014641 | 0.085901 | 0.253072 | -0.16732 | -0.143789 | 0.184636 | 0.055466 | 0.204388 | -1.488755 | -0.078409 | -0.093117 |
| 20.8125 | -0.20408 | 0.209436 | -0.711765 | -0.198652 | -0.505781 | -1.203573 | -0.205285 | -0.106804 | -0.040364 | 0.312911 | -0.814572 | -1.625893 | -0.820236 | -0.360518 |
| 20.8263889 | -0.035658 | 0.521606 | -1.050103 | -0.378899 | -0.281935 | -0.744241 | -0.175052 | -0.093316 | -0.481121 | 0.088112 | -1.28981 | -0.079578 | -0.369764 | -0.505476 |
| 20.8402778 | -0.306995 | -0.082797 | -1.050299 | -0.202257 | 0.107615 | 0.201316 | -0.114475 | -0.064241 | -0.207559 | -0.357669 | -0.629474 | 0.77518 | 0.05957 | -0.415639 |
| 20.8541667 | -0.462011 | -0.455183 | -0.651544 | -0.202468 | 0.255678 | 0.756423 | -0.06694 | -0.056675 | -0.283173 | -0.812634 | 0.237282 | -0.051719 | 0.028732 | -0.282178 |
| 20.8680556 | -0.47841 | -0.441334 | -0.924337 | -0.374788 | -0.15763 | -0.378652 | -0.067989 | -0.140033 | -0.863762 | -0.850257 | 0.214787 | -1.310537 | -0.273193 | -0.230426 |
| 20.8819444 | -0.539171 | -0.339347 | -2.072191 | -0.537314 | -0.525461 | -1.383565 | -0.096583 | -0.228416 | -1.096352 | -0.457465 | -0.551939 | -1.345189 | -1.018452 | -0.367137 |
| 20.8958333 | -0.816265 | -0.481678 | -3.120579 | -0.595485 | -0.190412 | -0.689073 | -0.046275 | -0.193061 | -0.611419 | -0.497113 | -1.031577 | -0.541095 | -0.54009 | -0.596806 |

| Time | RGBControl4 | RGBControl5 | RGBControl6 | RGBControl7 | RGBControl8 | RGBControl9 | RGBControl10 | RGBControl11 | RGBControl12 | RGBControlTank | Time | Mean ControlRGB | Mean StressRGB |
|---|---|---|---|---|---|---|---|---|---|---|---|---|---|
| 18.0069444 | 0.11226 | 0.085068 | -0.153998 | -0.330776 | -0.272133 | -0.314186 | 0.121874 | -0.036751 | -1.025192 | -0.041588 | 18.0972222 | 0.032835 | 0.541189 |
| 18.0208333 | 0.023343 | 0.02557 | -0.24697 | -0.327674 | -0.592209 | -0.4347 | 0.110339 | -0.13748 | -0.838439 | -0.126377 | 18.1111111 | -0.060605 | 0.510046 |
| 18.0347222 | -0.093619 | -0.079693 | -0.340821 | -0.387763 | -0.905577 | -0.48519 | 0.06861 | -0.210116 | -0.322787 | -0.227248 | 18.125 | -0.123781 | 0.431817 |
| 18.0486111 | -0.189339 | -0.139233 | -0.363919 | -0.400506 | -1.022275 | -0.475329 | 0.03512 | -0.222814 | -0.113434 | -0.319116 | 18.1388889 | -0.038694 | 0.327325 |
| 18.0625 | -0.213639 | -0.139289 | -0.304894 | -0.320314 | -0.815494 | -0.380894 | 0.034993 | -0.160195 | -0.051887 | -0.367293 | 18.1527778 | 0.045093 | 0.201772 |
| 18.0763889 | -0.171747 | -0.076712 | -0.216408 | -0.20045 | -0.564529 | -0.277707 | 0.045689 | -0.097295 | -0.022042 | -0.330959 | 18.1666667 | -0.017362 | 0.14295 |
| 18.0902778 | -0.083608 | -0.008084 | -0.155395 | -0.110434 | -0.367792 | -0.184826 | 0.041406 | -0.069661 | 0.006234 | -0.079058 | 18.1805556 | -0.145697 | 0.162676 |
| 18.1041667 | -0.019908 | 0.022356 | -0.134268 | -0.073771 | -0.24941 | -0.129684 | 0.021016 | -0.084593 | 0.028757 | 0.170244 | 18.1944444 | -0.18884 | 0.190261 |
| 18.1180556 | -0.000637 | 0.01931 | -0.152686 | -0.092028 | -0.18483 | -0.097277 | -0.004687 | -0.091386 | 0.040125 | 0.266804 | 18.2083333 | -0.14042 | 0.206799 |
| 18.1319444 | -0.048738 | -0.013408 | -0.223766 | -0.160794 | -0.176171 | -0.099617 | -0.02372 | -0.084164 | 0.012362 | 0.097906 | 18.2222222 | -0.132523 | 0.036406 |
| 18.1458333 | -0.113291 | -0.048606 | -0.284624 | -0.215759 | -0.173282 | -0.098216 | -0.010718 | -0.062612 | -0.039657 | -0.26454 | 18.2361111 | -0.17057 | -0.227509 |
| 18.1597222 | -0.081385 | -0.013708 | -0.243354 | -0.182275 | -0.125662 | -0.039683 | 0.033973 | -0.025488 | -0.018628 | -0.232278 | 18.25 | -0.169985 | -0.217401 |
| 18.1736111 | -0.018154 | 0.039745 | -0.170215 | -0.133962 | -0.065123 | 0.021776 | 0.065465 | 0.023058 | 0.040478 | 0.021369 | 18.2638889 | -0.127274 | -0.014568 |
| 18.1875 | -0.052792 | 0.016482 | -0.189659 | -0.194596 | -0.06779 | 0.001775 | 0.041779 | 0.028922 | 0.041327 | -0.015242 | 18.2777778 | -0.08223 | 0.031271 |
| 18.2013889 | -0.108515 | -0.041844 | -0.256139 | -0.309898 | -0.121609 | -0.081764 | -0.018197 | -0.014657 | 0.009596 | -0.180816 | 18.2916667 | -0.06275 | -0.099111 |
| 18.2152778 | -0.117542 | -0.068147 | -0.294099 | -0.372712 | -0.167859 | -0.142853 | -0.057748 | -0.030203 | -0.027731 | -0.35781 | 18.3055556 | -0.071298 | -0.228389 |
| 18.2291667 | -0.101795 | -0.062774 | -0.311485 | -0.333605 | -0.201347 | -0.149501 | -0.071956 | -0.042376 | -0.023833 | -0.275912 | 18.3194444 | -0.104079 | -0.286728 |
| 18.2430556 | -0.045596 | -0.025406 | -0.276993 | -0.22888 | -0.200468 | -0.109423 | -0.066855 | -0.078931 | 0.057458 | 0.23285 | 18.3333333 | -0.14358 | -0.236168 |
| 18.2569444 | 0.025815 | 0.032123 | -0.178707 | -0.09414 | -0.13611 | -0.049087 | -0.027833 | -0.066254 | 0.108963 | 0.451229 | 18.3472222 | -0.112742 | -0.082623 |
| 18.2708333 | 0.057191 | 0.060273 | -0.090861 | -0.009826 | -0.069854 | -0.010807 | 0.006897 | -0.018546 | 0.069571 | 0.20694 | 18.3611111 | -0.020642 | 0.074842 |
| 18.2847222 | 0.042076 | 0.048475 | -0.056012 | -0.017961 | -0.046883 | -0.001455 | 0.008443 | -0.008441 | 0.019556 | 0.025387 | 18.375 | 0.038488 | 0.174432 |
| 18.2986111 | -0.027831 | -0.007321 | -0.076106 | -0.089466 | -0.040211 | -0.027469 | -0.019276 | -0.039131 | -0.045208 | -0.19706 | 18.3888889 | 0.015364 | 0.169982 |
| 18.3125 | -0.096675 | -0.064309 | -0.109251 | -0.144665 | -0.0561 | -0.074452 | -0.052404 | -0.081201 | -0.093894 | -0.384624 | 18.4027778 | -0.067526 | 0.102823 |
| 18.3263889 | -0.076143 | -0.045391 | -0.100687 | -0.121545 | -0.074491 | -0.076562 | -0.045008 | -0.081791 | -0.0564 | -0.25323 | 18.4166667 | -0.10287 | 0.08597 |
| 18.3402778 | 0.002651 | 0.026036 | -0.051508 | -0.047969 | -0.036932 | -0.029062 | -0.003702 | -0.049737 | -0.0057 | -0.118423 | 18.4305556 | -0.051869 | 0.104383 |
| 18.3541667 | 0.042425 | 0.056896 | -0.016163 | 0.001821 | 0.003946 | -0.016155 | 0.005606 | -0.054871 | -0.020839 | -0.158426 | 18.4444444 | -0.101245 | -0.061784 |
| 18.3680556 | -0.01457 | -0.009815 | -0.066177 | -0.037915 | -0.055608 | -0.097592 | -0.069856 | -0.143452 | -0.113521 | -0.224054 | 18.4583333 | -0.333986 | -0.388153 |
| 18.3819444 | -0.123932 | -0.131737 | -0.182075 | -0.150268 | -0.20049 | -0.214298 | -0.191184 | -0.26695 | -0.219914 | -0.18706 | 18.4722222 | -0.449664 | -0.447809 |
| 18.3958333 | -0.136969 | -0.159817 | -0.213302 | -0.192537 | -0.237661 | -0.216973 | -0.230417 | -0.263262 | -0.187437 | -0.050566 | 18.4861111 | -0.244265 | -0.213022 |
| 18.4097222 | -0.038023 | -0.066173 | -0.119913 | -0.100072 | -0.111346 | -0.099592 | -0.157154 | -0.137269 | -0.068322 | -0.023778 | 18.5 | 0.009763 | -0.076703 |
| 18.4236111 | 0.123831 | 0.047818 | 0.018049 | -0.002428 | -0.131057 | 0.019334 | -0.077601 | -0.063362 | 0.256901 | 1.548743 | 18.5138889 | -0.298052 | -0.283427 |
| 18.4375 | 0.580867 | 0.272049 | 0.188879 | 0.040948 | -1.076114 | 0.122526 | -0.164966 | -0.069187 | 2.024573 | 11.168965 | 18.5277778 | -2.830116 | -1.881362 |
| 18.4513889 | 0.667719 | 0.219114 | 0.185466 | -0.122785 | -2.807591 | -0.170582 | -0.662722 | -0.039268 | 5.259112 | 27.867575 | 18.5416667 | -7.741367 | -6.547624 |
| 18.4652778 | -0.950328 | -0.893771 | -1.123383 | -0.79684 | -4.213231 | -1.387415 | -1.769262 | -0.659846 | 5.674774 | 28.172131 | 18.5555556 | -9.566361 | -10.48582 |
| 18.4791667 | -1.979645 | -1.795514 | -2.420998 | -2.105356 | -3.356824 | -1.853659 | -2.224375 | -0.864239 | 2.384088 | 10.01934 | 18.5694444 | -4.710957 | -7.668576 |
| 18.4930556 | -1.343296 | -0.963743 | -2.163151 | -1.739128 | -1.262134 | -0.980595 | -1.353845 | -0.745025 | -0.082345 | 0.1136 | 18.5833333 | -0.612239 | -1.985507 |
| 18.5069444 | -0.656047 | -0.2919 | -1.407859 | -1.253886 | -0.365835 | -0.253414 | -0.51869 | -0.268751 | -0.054604 | -0.589622 | 18.5972222 | 0.164606 | -0.10695 |
| 18.5208333 | -0.291384 | 0.017041 | -0.781766 | -0.786497 | -0.197764 | -0.090307 | -0.164998 | -0.032217 | 0.161144 | -0.205364 | 18.6111111 | 0.146081 | 0.218548 |
| 18.5347222 | -0.076074 | 0.127662 | -0.322313 | -0.413022 | -0.016673 | -0.023871 | -0.074761 | 0.069901 | 0.233719 | -0.026755 | 18.625 | 0.067256 | 0.36408 |
| 18.5486111 | 0.030194 | 0.187137 | -0.169189 | -0.365041 | 0.151419 | 0.056099 | -0.021532 | 0.157787 | 0.283362 | 0.061928 | 18.6388889 | -0.006084 | 0.513103 |
| 18.5625 | 0.037218 | 0.214087 | -0.258605 | -0.533586 | 0.224869 | 0.168057 | 0.085883 | 0.215489 | 0.315921 | 0.066216 | 18.6527778 | -0.0023 | 0.605626 |
| 18.5763889 | 0.019271 | 0.234298 | -0.336833 | -0.644752 | 0.241942 | 0.327137 | 0.218327 | 0.242054 | 0.344206 | 0.063955 | 18.6666667 | 0.037879 | 0.638109 |
| 18.5902778 | 0.016713 | 0.263693 | -0.195913 | -0.373234 | 0.231528 | 0.518584 | 0.360161 | 0.266356 | 0.400587 | 0.150476 | 18.6805556 | 0.040817 | 0.686475 |
| 18.6041667 | 0.07019 | 0.300844 | 0.119189 | 0.178137 | 0.217515 | 0.722589 | 0.493414 | 0.306498 | 0.465404 | 0.367114 | 18.6944444 | -0.007237 | 0.758858 |
| 18.6180556 | 0.200654 | 0.360394 | 0.516226 | 0.995151 | 0.337223 | 0.909991 | 0.559834 | 0.356171 | 0.508107 | 0.589807 | 18.7083333 | -0.021236 | 0.898201 |
| 18.6319444 | 0.332131 | 0.43948 | 0.946583 | 2.216242 | 0.491081 | 0.987486 | 0.604522 | 0.411071 | 0.519412 | 0.672442 | 18.7222222 | 0.021859 | 1.020164 |
| 18.6458333 | 0.379113 | 0.477094 | 1.173882 | 2.983112 | 0.51219 | 0.886133 | 0.571647 | 0.433544 | 0.509591 | 0.593733 | 18.7361111 | 0.016448 | 0.883507 |
| 18.6597222 | 0.345946 | 0.427375 | 0.922347 | 2.002649 | 0.420461 | 0.676222 | 0.431803 | 0.386861 | 0.441555 | 0.455096 | 18.75 | -0.035306 | 0.548847 |
| 18.6736111 | 0.333122 | 0.356336 | 0.605214 | 1.076133 | 0.342651 | 0.439246 | 0.177227 | 0.326618 | 0.36083 | 0.359152 | 18.7638889 | -0.062031 | 0.304192 |
| 18.6875 | 0.295592 | 0.285083 | 0.267487 | 0.372884 | 0.212561 | 0.157311 | -0.117824 | 0.27535 | 0.288016 | 0.315258 | 18.7777778 | -0.061474 | 0.154165 |
| 18.7013889 | 0.156254 | 0.172383 | -0.082903 | -0.264672 | -0.022764 | -0.103639 | -0.330897 | 0.164804 | 0.187158 | 0.182147 | 18.7916667 | -0.05083 | -0.058411 |
| 18.7152778 | 0.050592 | 0.112155 | -0.335518 | -0.86736 | -0.170294 | -0.117213 | -0.263867 | 0.074961 | 0.123317 | 0.069679 | 18.8055556 | -0.056074 | -0.153158 |
| 18.7291667 | 0.07563 | 0.14762 | -0.400877 | -1.581543 | -0.123963 | 0.05038 | -0.031692 | 0.074233 | 0.157557 | 0.158507 | 18.8194444 | -0.059146 | -0.050399 |
| 18.7430556 | 0.115164 | 0.156105 | -0.298499 | -1.73906 | -0.100088 | 0.216976 | 0.19199 | 0.0556 | 0.207407 | 0.318913 | 18.8333333 | -0.015078 | 0.018197 |
| 18.7569444 | 0.081133 | 0.070348 | -0.110454 | -0.674407 | -0.140239 | 0.240618 | 0.307318 | -0.03599 | 0.260756 | 0.411514 | 18.8472222 | 0.009862 | -0.094702 |
| 18.7708333 | 0.013043 | -0.064326 | -0.132107 | -0.29482 | -0.134321 | 0.228702 | 0.289808 | -0.12004 | 0.242007 | 0.323777 | 18.8611111 | -0.047578 | -0.313766 |
| 18.7847222 | -0.000468 | -0.132827 | -0.070619 | 0.022822 | -0.001833 | 0.196926 | 0.304067 | -0.133363 | 0.129953 | 0.109252 | 18.875 | -0.071093 | -0.40522 |
| 18.7986111 | 0.108574 | -0.080422 | 0.192857 | 0.674997 | 0.175042 | 0.136885 | 0.172481 | -0.068863 | 0.044712 | 0.031522 | 18.8888889 | -0.010877 | -0.299832 |
| 18.8125 | 0.27934 | -0.002004 | 0.573556 | 1.84342 | 0.335828 | 0.071738 | 0.052029 | 0.004067 | 0.028207 | 0.208676 | 18.9027778 | 0.042403 | -0.142702 |
| 18.8263889 | 0.400782 | 0.012756 | 0.692405 | 1.657109 | 0.38099 | -0.010345 | -0.039977 | 0.01036 | -0.076733 | 0.360022 | 18.9166667 | 0.074662 | -0.037549 |
| 18.8402778 | 0.432428 | -0.035768 | 0.697374 | 1.234723 | 0.334356 | -0.123671 | -0.132842 | -0.012876 | -0.197707 | 0.162312 | 18.9305556 | 0.063323 | -0.027999 |
| 18.8541667 | 0.316244 | -0.073912 | 0.743839 | 0.99579 | 0.299549 | -0.181211 | -0.150884 | -0.02143 | -0.20779 | -0.138748 | 18.9444444 | -0.013882 | -0.075883 |
| 18.8680556 | 0.204338 | -0.028106 | 0.770143 | 0.796355 | 0.413259 | -0.150797 | -0.104223 | -0.038815 | -0.156365 | -0.175578 | 18.9583333 | -0.053196 | -0.061474 |
| 18.8819444 | 0.179742 | 0.029453 | 0.722852 | 0.696973 | 0.574897 | -0.067549 | -0.087592 | -0.03145 | -0.123639 | 0.092119 | 18.9722222 | -0.031356 | 0.066425 |
| 18.8958333 | 0.240477 | 0.149877 | 0.617695 | 0.570094 | 0.636622 | 0.053987 | -0.110431 | 0.00593 | -0.107614 | 0.406678 | 18.9861111 | -0.018343 | 0.182716 |
| 18.9097222 | 0.249559 | 0.185213 | 0.566223 | 0.388965 | 0.541812 | 0.097182 | 0.032439 | 0.026198 | -0.180036 | 0.472719 | 19 | 0.013309 | 0.240324 |
| 18.9236111 | 0.161564 | 0.187139 | 0.586293 | 0.154889 | 0.486407 | 0.002438 | 0.041346 | 0.028834 | -0.816592 | -1.50746 | 19.0138889 | 0.747779 | 0.8889 |
| 18.9375 | 0.025116 | 0.512204 | 0.391514 | -0.624392 | 0.885622 | -0.218371 | -0.197635 | -0.13729 | -3.642726 | -14.18706 | 19.0277778 | 4.245023 | 3.306638 |
| 18.9513889 | -0.736688 | 0.196807 | -0.449475 | -1.316084 | 1.950403 | -0.738524 | -1.486482 | -0.044998 | -6.071648 | -32.770718 | 19.0416667 | 7.395805 | 2.658779 |
| 18.9652778 | -1.342848 | -0.332967 | -0.434981 | -1.113531 | 1.436197 | -0.752599 | -1.37633 | -0.337862 | -4.917143 | -28.042621 | 19.0555556 | 6.361797 | 1.533515 |
| 18.9791667 | -0.510415 | -0.11274 | -0.112525 | -0.526247 | 0.537656 | -0.412449 | -0.72218 | -0.082713 | -1.779932 | -10.55478 | 19.0694444 | 3.524762 | 3.128312 |
| 18.9930556 | 0.066405 | 0.173606 | 0.001234 | -0.331504 | 0.065709 | -0.199074 | -0.359194 | 0.071643 | -0.521388 | -3.966654 | 19.0833333 | 3.19438 | 3.468069 |
| 19.0069444 | 0.024365 | 0.174446 | -0.119953 | -0.374495 | -0.093272 | -0.252375 | -0.199382 | 0.036538 | -0.581421 | -3.014255 | 19.0972222 | 1.89839 | 2.260119 |
| 19.0208333 | -0.058694 | 0.073393 | -0.282109 | -0.494164 | -0.284692 | -0.302676 | -0.083941 | -0.147299 | -0.682598 | -1.463629 | 19.1111111 | 0.261913 | 0.596758 |
| 19.0347222 | -0.113884 | -0.03639 | -0.406173 | -0.621756 | -0.468974 | -0.291733 | -0.02661 | -0.200161 | -0.536316 | -0.51286 | 19.125 | -0.006062 | 0.020452 |
| 19.0486111 | -0.153418 | -0.146363 | -0.452749 | -0.652855 | -0.552631 | -0.239298 | -0.007141 | -0.1861 | -0.27677 | 0.038046 | 19.1388889 | -0.048359 | -0.201187 |
| 19.0625 | -0.185337 | -0.227931 | -0.474126 | -0.664211 | -0.583047 | -0.22237 | -0.027803 | -0.241859 | -0.052543 | 0.931082 | 19.1527778 | -0.102334 | -0.114953 |
| 19.0763889 | -0.157288 | -0.205974 | -0.464248 | -0.63874 | -0.552161 | -0.208287 | -0.047719 | -0.409914 | 0.060612 | 1.44042 | 19.1666667 | -0.07327 | 0.030352 |
| 19.0902778 | -0.191917 | -0.153761 | -0.41604 | -0.507209 | -0.354996 | -0.18913 | -0.100859 | -0.312833 | -0.124334 | -0.179641 | 19.1805556 | -0.030692 | -0.081543 |
| 19.1041667 | -0.207639 | -0.105135 | -0.332465 | -0.335627 | -0.168691 | -0.15498 | -0.128885 | -0.201231 | -0.292477 | -1.53031 | 19.1944444 | -0.096258 | -0.192692 |
| 19.1180556 | -0.094489 | -0.052424 | -0.218325 | -0.230225 | -0.197005 | -0.121151 | -0.064112 | -0.162827 | -0.059609 | -0.045531 | 19.2083333 | -0.121904 | -0.020952 |
| 19.1319444 | -0.066956 | -0.080749 | -0.197585 | -0.235847 | -0.31259 | -0.18028 | -0.085743 | -0.234196 | 0.049037 | 1.133798 | 19.2222222 | -0.031702 | 0.143857 |
| 19.1458333 | -0.210682 | -0.167258 | -0.294918 | -0.261594 | -0.3052 | -0.286201 | -0.228251 | -0.307427 | -0.272733 | -0.370499 | 19.2361111 | 0.037726 | 0.00941 |
| 19.1597222 | -0.294448 | -0.18769 | -0.340574 | -0.245631 | -0.238479 | -0.294173 | -0.281835 | -0.288896 | -0.474489 | -1.694695 | 19.25 | 0.03682 | -0.088702 |
| 19.1736111 | -0.227789 | -0.134188 | -0.292188 | -0.191544 | -0.225352 | -0.226008 | -0.199604 | -0.244524 | -0.298392 | -1.001053 | 19.2638889 | -0.036229 | -0.025714 |
| 19.1875 | -0.151557 | -0.082599 | -0.222789 | -0.134571 | -0.212798 | -0.182493 | -0.128083 | -0.223536 | -0.131079 | -0.259394 | 19.2777778 | -0.082974 | -0.073932 |
| 19.2013889 | -0.1366 | -0.07598 | -0.19844 | -0.114396 | -0.193528 | -0.181509 | -0.1253 | -0.207865 | -0.118413 | -0.242363 | 19.2916667 | -0.027783 | -0.210126 |
| 19.2152778 | -0.179073 | -0.120693 | -0.241613 | -0.150836 | -0.233728 | -0.205238 | -0.158473 | -0.215204 | -0.109514 | -0.208917 | 19.3055556 | 0.008372 | -0.270921 |
| 19.2291667 | -0.239088 | -0.189937 | -0.314373 | -0.213787 | -0.312619 | -0.237032 | -0.205874 | -0.270358 | -0.05387 | 0.118192 | 19.3194444 | -0.021624 | -0.350586 |
| 19.2430556 | -0.212442 | -0.187001 | -0.310035 | -0.24239 | -0.345733 | -0.217701 | -0.203798 | -0.293791 | 0.077613 | 0.774584 | 19.3333333 | -0.027482 | -0.420538 |
| 19.2569444 | -0.133844 | -0.107451 | -0.256993 | -0.213865 | -0.280544 | -0.161049 | -0.146583 | -0.223904 | 0.122205 | 0.842352 | 19.3472222 | -0.015565 | -0.343539 |
| 19.2708333 | -0.102849 | -0.041529 | -0.206394 | -0.131874 | -0.126879 | -0.117649 | -0.109609 | -0.112086 | -0.050243 | -0.253193 | 19.3611111 | -0.006189 | -0.136933 |
| 19.2847222 | -0.013653 | 0.055099 | -0.089968 | -0.036032 | -0.034041 | -0.034671 | -0.019007 | -0.037143 | -0.031779 | -0.212732 | 19.375 | 0.016732 | 0.093667 |
| 19.2986111 | 0.083443 | 0.121191 | 0.034868 | 0.000402 | -0.059853 | 0.038384 | 0.071991 | -0.028342 | 0.178891 | 0.896502 | 19.3888889 | 0.016888 | 0.207897 |
| 19.3125 | -0.022969 | 0.022419 | -0.010322 | -0.027675 | -0.040007 | -0.024545 | -0.008596 | -0.044276 | 0.038205 | -0.230876 | 19.4027778 | -0.009989 | 0.175442 |
| 19.3263889 | -0.165623 | -0.097567 | -0.100494 | -0.060554 | 0.000464 | -0.106599 | -0.123759 | -0.047624 | -0.216646 | -1.920541 | 19.4166667 | -0.032508 | 0.023663 |
| 19.3402778 | -0.12576 | -0.068978 | -0.068809 | -0.067603 | -0.015259 | -0.077038 | -0.104137 | -0.037997 | -0.130605 | -1.407286 | 19.4305556 | -0.053756 | -0.15822 |
| 19.3541667 | -0.014125 | 0.016744 | 0.009014 | -0.048989 | -0.04211 | -0.021329 | -0.032701 | -0.032343 | 0.029691 | -0.136562 | 19.4444444 | -0.113346 | -0.172879 |
| 19.3680556 | 0.047897 | 0.04773 | 0.042364 | -0.056633 | -0.102505 | -0.013512 | -0.001062 | -0.068586 | 0.124086 | 0.841559 | 19.4583333 | -0.256015 | -0.07839 |
| 19.3819444 | 0.02249 | -0.000447 | 0.008006 | -0.106388 | -0.17527 | -0.037126 | -0.015026 | -0.122788 | 0.147425 | 1.249701 | 19.4722222 | -0.298905 | -0.105226 |
| 19.3958333 | -0.12544 | -0.122642 | -0.102489 | -0.145821 | -0.168279 | -0.093425 | -0.099307 | -0.128115 | -0.006254 | 0.213848 | 19.4861111 | -0.142405 | -0.351919 |
| 19.4097222 | -0.278084 | -0.235368 | -0.161972 | -0.141468 | -0.133271 | -0.154971 | -0.203364 | -0.103474 | -0.199113 | -1.096039 | 19.5 | -0.030256 | -0.629551 |
| 19.4236111 | -0.131487 | -0.111246 | 0.017157 | -0.009996 | -0.171011 | -0.064379 | -0.090768 | -0.034553 | 0.077907 | 0.419025 | 19.5138889 | -0.354204 | -0.941744 |
| 19.4375 | 0.664691 | 0.566144 | 0.497855 | 0.285449 | -0.632361 | 0.327599 | 0.684323 | 0.143126 | 1.919636 | 11.221666 | 19.5277778 | -2.571221 | -3.077864 |
| 19.4513889 | 1.416391 | 1.178057 | 0.939469 | 0.597089 | -1.753037 | 0.769226 | 2.027995 | 0.861269 | 4.997996 | 31.780385 | 19.5416667 | -6.830476 | -6.65721 |
| 19.4652778 | 0.218033 | 0.108222 | 0.53511 | 0.372676 | -1.951646 | 0.393404 | 1.302441 | 1.997902 | 6.414182 | 33.855133 | 19.5555556 | -8.665727 | -8.008611 |
| 19.4791667 | -0.611491 | -0.642356 | -0.248501 | -0.195154 | -1.355683 | -0.176993 | -0.531059 | 1.070518 | 3.219808 | 13.129472 | 19.5694444 | -4.444392 | -5.346015 |
| 19.4930556 | -0.369235 | -0.28402 | 0.035159 | -0.145535 | -0.418225 | 0.018672 | -0.402166 | 0.516433 | 0.204736 | 1.536255 | 19.5833333 | -0.675556 | -1.541204 |
| 19.5069444 | 0.034408 | 0.09161 | 0.294656 | 0.085029 | 0.080373 | 0.219108 | -0.04814 | 0.567621 | 0.184708 | 0.093863 | 19.5972222 | 0.021965 | 0.132387 |
| 19.5208333 | 0.194341 | 0.237941 | 0.326105 | 0.201376 | 0.242028 | 0.266938 | 0.128359 | 0.551772 | 0.581089 | 0.105256 | 19.6111111 | 0.000923 | 0.259348 |
| 19.5347222 | 0.212343 | 0.240576 | 0.296062 | 0.252954 | 0.222817 | 0.236569 | 0.167779 | 0.476444 | 0.786882 | 0.059486 | 19.625 | -0.020922 | 0.267185 |
| 19.5486111 | 0.218301 | 0.229545 | 0.26055 | 0.325311 | 0.185993 | 0.207417 | 0.190237 | 0.431935 | 0.933577 | 0.110341 | 19.6388889 | -0.022065 | 0.230758 |
| 19.5625 | 0.225001 | 0.222467 | 0.20711 | 0.318719 | 0.196119 | 0.192754 | 0.194798 | 0.347395 | 1.082619 | 0.158679 | 19.6527778 | -0.007926 | 0.177582 |
| 19.5763889 | 0.220414 | 0.205807 | 0.109395 | 0.166142 | 0.239681 | 0.177303 | 0.162281 | 0.261281 | 1.069624 | 0.129636 | 19.6666667 | 0.014822 | 0.198856 |
| 19.5902778 | 0.226666 | 0.213748 | 0.027312 | -0.095343 | 0.351768 | 0.180644 | 0.14374 | 0.191343 | 0.975321 | 0.084842 | 19.6805556 | 0.002541 | 0.274817 |
| 19.6041667 | 0.244859 | 0.231395 | 0.054407 | -0.354748 | 0.531553 | 0.198483 | 0.141004 | 0.14992 | 0.890467 | 0.083341 | 19.6944444 | -0.012519 | 0.336646 |
| 19.6180556 | 0.253035 | 0.230504 | 0.153228 | -0.496875 | 0.715019 | 0.221806 | 0.123383 | 0.116904 | 0.745248 | 0.055796 | 19.7083333 | 0.013116 | 0.330301 |
| 19.6319444 | 0.281162 | 0.239505 | 0.454546 | -0.251572 | 0.801215 | 0.269066 | 0.113993 | 0.145104 | 0.630687 | -0.016118 | 19.7222222 | 0.004187 | 0.24698 |
| 19.6458333 | 0.37325 | 0.315781 | 0.931051 | 0.59342 | 0.771035 | 0.369256 | 0.146158 | 0.231259 | 0.497336 | 0.030516 | 19.7361111 | -0.025439 | 0.186143 |
| 19.6597222 | 0.440327 | 0.402355 | 1.427371 | 1.968479 | 0.651394 | 0.436727 | 0.172536 | 0.29246 | 0.316528 | 0.158188 | 19.75 | -0.033461 | 0.19342 |
| 19.6736111 | 0.410688 | 0.377894 | 2.134935 | 4.79327 | 0.489397 | 0.429979 | 0.130899 | 0.292787 | 0.137776 | 0.129701 | 19.7638889 | -0.040743 | 0.18586 |
| 19.6875 | 0.327058 | 0.258627 | 1.717094 | 3.633527 | 0.354691 | 0.366782 | 0.068249 | 0.249879 | 0.074806 | 0.120077 | 19.7777778 | -0.02224 | 0.117569 |
| 19.7013889 | 0.271027 | 0.180508 | 0.85367 | 1.61167 | 0.250213 | 0.267793 | 0.036951 | 0.204943 | 0.0832 | 0.171064 | 19.7916667 | 0.014448 | 0.037481 |
| 19.7152778 | 0.28425 | 0.171578 | 0.330682 | 0.577248 | 0.195632 | 0.176973 | 0.03183 | 0.169886 | 0.099729 | 0.229044 | 19.8055556 | 0.048089 | 0.080894 |
| 19.7291667 | 0.335352 | 0.235781 | 0.004122 | -0.160041 | 0.168127 | 0.120631 | 0.027294 | 0.157276 | 0.077495 | 0.309698 | 19.8194444 | 0.08332 | 0.257249 |
| 19.7430556 | 0.288914 | 0.241407 | -0.287237 | -1.018124 | 0.11692 | 0.062432 | 0.009999 | 0.14341 | -0.003902 | 0.242098 | 19.8333333 | 0.088434 | 0.370424 |
| 19.7569444 | 0.186652 | 0.161354 | -0.690749 | -2.626218 | 0.099273 | 0.02583 | -0.001749 | 0.118199 | -0.045965 | 0.129157 | 19.8472222 | 0.059546 | 0.326382 |
| 19.7708333 | 0.133206 | 0.115302 | -0.722953 | -3.344817 | 0.148555 | 0.052682 | 0.010235 | 0.119505 | -0.030248 | 0.113235 | 19.8611111 | 0.055886 | 0.271131 |
| 19.7847222 | 0.123159 | 0.101112 | -0.16187 | -1.197495 | 0.189799 | 0.095679 | 0.039729 | 0.15481 | 0.006772 | 0.123175 | 19.875 | 0.083309 | 0.238706 |
| 19.7986111 | 0.107469 | 0.088531 | 0.270038 | 0.733131 | 0.181054 | 0.107871 | 0.06888 | 0.202799 | 0.034847 | 0.070094 | 19.8888889 | 0.107105 | 0.19753 |
| 19.8125 | 0.061768 | 0.058432 | 0.48451 | 1.839779 | 0.125453 | 0.07849 | 0.064859 | 0.205697 | 0.017559 | 0.01676 | 19.9027778 | 0.114955 | 0.170771 |
| 19.8263889 | 0.013103 | 0.008444 | 0.344636 | 1.328468 | 0.064837 | 0.042743 | 0.033286 | 0.157603 | -0.062844 | 0.034759 | 19.9166667 | 0.114046 | 0.139421 |
| 19.8402778 | 0.0045 | -0.014838 | 0.172353 | 0.464712 | 0.052282 | 0.04991 | 0.016541 | 0.145823 | -0.125893 | 0.122675 | 19.9305556 | 0.116615 | 0.033325 |
| 19.8541667 | 0.015479 | -0.009494 | 0.007185 | -0.000151 | 0.045912 | 0.059611 | 0.022009 | 0.153795 | -0.167083 | 0.168659 | 19.9444444 | 0.136544 | -0.122864 |
| 19.8680556 | -0.006709 | -0.029004 | -0.168525 | -0.227026 | -0.013035 | 0.015173 | 0.013697 | 0.10923 | -0.269098 | 0.143965 | 19.9583333 | 0.10198 | -0.254577 |
| 19.8819444 | -0.058007 | -0.07751 | -0.33556 | -0.366006 | -0.061408 | -0.046867 | -0.012508 | 0.059014 | -0.419611 | 0.098018 | 19.9722222 | -0.005233 | -0.32404 |
| 19.8958333 | -0.094737 | -0.120484 | -0.420423 | -0.46772 | 0.074895 | -0.035052 | -0.032652 | 0.013624 | -0.583184 | 0.108017 | 19.9861111 | -0.063943 | -0.292326 |
| 19.9097222 | -0.124 | -0.151573 | -0.444805 | -0.575898 | 0.447345 | 0.061343 | -0.042527 | -0.043442 | -0.747661 | 0.164482 | 20 | -0.008294 | -0.173405 |
| 19.9236111 | -0.202694 | -0.112139 | -0.517047 | -0.82309 | 0.414 | 0.003372 | -0.073884 | -0.219793 | -1.361788 | -1.699134 | 20.0138889 | 0.678315 | 0.283512 |
| 19.9375 | -0.429164 | 0.273143 | -0.766815 | -1.796237 | 0.421393 | -0.432206 | -0.365966 | -0.922976 | -4.000693 | -14.399054 | 20.0277778 | 4.159415 | 2.828576 |
| 19.9513889 | -0.993292 | 0.257363 | -1.353198 | -1.898583 | 0.927111 | -1.055822 | -1.452556 | -1.616613 | -6.265471 | -35.792869 | 20.0416667 | 8.449172 | 6.177772 |
| 19.9652778 | -1.50364 | -0.430937 | -0.936776 | -1.748978 | 1.139668 | -1.402866 | -2.025636 | -0.809642 | -4.722978 | -32.095243 | 20.0555556 | 6.752139 | 6.22299 |
| 19.9791667 | -0.78039 | -0.343934 | -0.447377 | -0.719151 | 0.073754 | -0.921076 | -0.873127 | -0.237512 | -2.02805 | -12.677817 | 20.0694444 | 2.683416 | 2.4745 |
| 19.9930556 | -0.150114 | -0.055297 | -0.214561 | -0.270499 | -0.45261 | -0.569057 | -0.118013 | -0.229476 | -0.741812 | -1.954489 | 20.0833333 | 0.412326 | 0.450968 |
| 20.0069444 | -0.063512 | -0.004366 | -0.199957 | -0.144312 | -0.415016 | -0.476619 | 0.004779 | -0.282261 | -0.46894 | -0.329593 | 20.0972222 | 0.050896 | 0.086426 |
| 20.0208333 | -0.050325 | 0.004733 | -0.233299 | -0.107178 | -0.312454 | -0.412806 | 0.032702 | -0.228071 | -0.309466 | 0.005274 | 20.1111111 | 0.003616 | 0.106954 |
| 20.0347222 | -0.027411 | 0.022036 | -0.245391 | -0.053043 | -0.271629 | -0.313851 | 0.071875 | -0.13322 | -0.155645 | 0.326385 | 20.125 | -0.016798 | 0.134755 |
| 20.0486111 | -0.045335 | 0.001361 | -0.232497 | -0.053365 | -0.252969 | -0.229152 | 0.053066 | -0.126971 | -0.117891 | 0.04177 | 20.1388889 | -0.001232 | 0.039842 |
| 20.0625 | -0.072584 | -0.016411 | -0.196226 | -0.059442 | -0.2184 | -0.167072 | 0.012888 | -0.140824 | -0.104545 | -0.374653 | 20.1527778 | -0.005992 | -0.02606 |
| 20.0763889 | -0.057208 | 0.008754 | -0.142077 | -0.03667 | -0.170454 | -0.115288 | 0.015706 | -0.096541 | -0.006851 | -0.346542 | 20.1666667 | -0.021307 | -0.017773 |
| 20.0902778 | -0.033479 | 0.025259 | -0.134065 | -0.033369 | -0.133917 | -0.0877 | 0.020792 | -0.068711 | 0.067653 | -0.057196 | 20.1805556 | -0.019109 | -0.013849 |
| 20.1041667 | -0.044137 | 0.001777 | -0.187064 | -0.042666 | -0.094385 | -0.079804 | 0.001101 | -0.080694 | 0.042327 | 0.113462 | 20.1944444 | -0.027627 | -0.02551 |
| 20.1180556 | -0.083221 | -0.029235 | -0.255944 | -0.076756 | -0.085538 | -0.07087 | -0.008743 | -0.091941 | -0.022941 | 0.028474 | 20.2083333 | -0.120491 | -0.035676 |
| 20.1319444 | -0.103215 | -0.049725 | -0.293524 | -0.100486 | -0.148998 | -0.073262 | 0.004015 | -0.109301 | -0.0554 | 0.040448 | 20.2222222 | -0.190105 | -0.076221 |
| 20.1458333 | -0.062476 | -0.037969 | -0.246686 | -0.081957 | -0.229945 | -0.091696 | 0.017709 | -0.129115 | -0.036981 | 0.347485 | 20.2361111 | -0.052195 | -0.047707 |
| 20.1597222 | -0.04086 | -0.033552 | -0.177718 | -0.077709 | -0.303224 | -0.155016 | -0.004025 | -0.15695 | -0.049917 | 0.333158 | 20.25 | 0.072868 | 0.117381 |
| 20.1736111 | -0.073959 | -0.048201 | -0.161508 | -0.108619 | -0.400513 | -0.234904 | -0.035891 | -0.173106 | -0.085153 | -0.086745 | 20.2638889 | 0.038292 | 0.164456 |
| 20.1875 | -0.116754 | -0.07785 | -0.168578 | -0.143148 | -0.413606 | -0.252284 | -0.066207 | -0.17034 | -0.055797 | -0.302302 | 20.2777778 | 0.041936 | -0.099869 |
| 20.2013889 | -0.179224 | -0.145243 | -0.212537 | -0.203811 | -0.324944 | -0.212809 | -0.136075 | -0.197944 | -0.039317 | -0.32087 | 20.2916667 | 0.06728 | -0.469414 |
| 20.2152778 | -0.186997 | -0.15544 | -0.225966 | -0.214561 | -0.235272 | -0.147321 | -0.152909 | -0.218948 | -0.043112 | -0.225703 | 20.3055556 | 0.031681 | -0.555918 |
| 20.2291667 | -0.100511 | -0.062488 | -0.163708 | -0.141209 | -0.120918 | -0.055681 | -0.079012 | -0.194978 | 0.012163 | -0.03074 | 20.3194444 | -0.00166 | -0.105978 |
| 20.2430556 | -0.081908 | -0.03194 | -0.165528 | -0.135528 | -0.108683 | -0.065021 | -0.078444 | -0.204357 | 0.026084 | -0.083701 | 20.3333333 | -0.043427 | 0.467837 |
| 20.2569444 | -0.194967 | -0.140632 | -0.272805 | -0.250912 | -0.238109 | -0.193242 | -0.203582 | -0.259385 | -0.069327 | -0.335823 | 20.3472222 | -0.083037 | 0.497544 |
| 20.2708333 | -0.251045 | -0.208085 | -0.311981 | -0.313479 | -0.30273 | -0.230744 | -0.260218 | -0.234014 | -0.117813 | -0.232173 | 20.3611111 | -0.006098 | 0.069125 |
| 20.2847222 | -0.169563 | -0.145876 | -0.21521 | -0.244264 | -0.184973 | -0.116619 | -0.186185 | -0.117137 | -0.051093 | 0.000144 | 20.375 | 0.073023 | -0.173471 |
| 20.2986111 | -0.070966 | -0.060928 | -0.094687 | -0.137579 | 0.002901 | -0.005111 | -0.114416 | -0.047423 | 0.003369 | -0.172738 | 20.3888889 | -0.002229 | 0.036769 |
| 20.3125 | -0.032383 | -0.028381 | -0.037437 | -0.068587 | 0.122147 | 0.043941 | -0.095464 | -0.052814 | 0.03826 | -0.276078 | 20.4027778 | -0.062988 | 0.224563 |
| 20.3263889 | -0.1653 | -0.164604 | -0.167922 | -0.172651 | -0.003097 | -0.081332 | -0.219809 | -0.191879 | -0.051153 | -0.171337 | 20.4166667 | -0.048223 | -0.054931 |
| 20.3402778 | -0.406746 | -0.414527 | -0.408135 | -0.382509 | -0.265563 | -0.296783 | -0.447449 | -0.41624 | -0.254141 | -0.118568 | 20.4305556 | -0.077299 | -0.432153 |
| 20.3541667 | -0.395691 | -0.41786 | -0.375368 | -0.340725 | -0.198251 | -0.225593 | -0.455365 | -0.384281 | -0.23842 | 0.059373 | 20.4444444 | -0.171492 | -0.46925 |
| 20.3680556 | -0.1233 | -0.152518 | -0.0847 | -0.046784 | 0.223135 | 0.123595 | -0.228885 | -0.052891 | -0.037267 | 0.014128 | 20.4583333 | -0.297665 | -0.388974 |
| 20.3819444 | 0.138154 | 0.101525 | 0.159327 | 0.192053 | 0.556332 | 0.403288 | -0.019847 | 0.235977 | 0.167101 | 0.051284 | 20.4722222 | -0.33802 | -0.269728 |
| 20.3958333 | 0.263629 | 0.189237 | 0.274094 | 0.284508 | 0.654652 | 0.502566 | 0.042791 | 0.286903 | 0.373491 | 0.690688 | 20.4861111 | -0.216724 | 0.022112 |
| 20.4097222 | 0.244826 | 0.160299 | 0.28742 | 0.311629 | 0.726613 | 0.505918 | -0.027062 | 0.185897 | 0.506653 | 0.863331 | 20.5 | -0.04581 | 0.363005 |
| 20.4236111 | 0.231979 | 0.162047 | 0.27949 | 0.416332 | 0.909343 | 0.633424 | -0.005934 | 0.18265 | 0.776434 | 1.627809 | 20.5138889 | -0.294907 | 0.126869 |
| 20.4375 | 0.477955 | 0.063929 | 0.245222 | 0.617318 | 0.333776 | 1.021502 | 0.603461 | 0.24538 | 1.89124 | 11.812073 | 20.5277778 | -2.491272 | -2.446559 |
| 20.4513889 | 0.460026 | -0.820081 | -0.108726 | 1.194218 | -1.059198 | 1.206553 | 1.215672 | 0.359202 | 4.100882 | 31.87329 | 20.5416667 | -6.618239 | -7.621672 |
| 20.4652778 | -0.642796 | -2.398412 | -0.522684 | 0.89535 | -1.385449 | 0.562679 | 0.661895 | 0.112727 | 4.635132 | 34.412678 | 20.5555556 | -8.652126 | -6.356985 |
| 20.4791667 | -0.566141 | -1.282821 | -0.321547 | -0.320008 | -0.068949 | 0.381388 | -0.651593 | 0.311784 | 2.795376 | 14.850537 | 20.5694444 | -4.376163 | -4.424503 |
| 20.4930556 | 0.370873 | 0.112562 | 0.956461 | -0.000649 | 1.547183 | 0.876032 | -0.203916 | 0.986221 | 1.151186 | 2.271033 | 20.5833333 | -0.705096 | -0.283892 |
| 20.5069444 | 0.592267 | 0.37772 | 1.54554 | 0.330517 | 1.756305 | 1.173902 | -0.029388 | 1.275614 | 0.786348 | 0.261739 | 20.5972222 | -0.099553 | 0.454435 |
| 20.5208333 | 0.540157 | 0.30489 | 1.460797 | 0.752755 | 1.513252 | 1.384384 | -0.118921 | 1.243347 | 0.712363 | -0.195639 | 20.6111111 | -0.171316 | 0.004132 |
| 20.5347222 | 0.498853 | 0.34103 | 1.113163 | 1.447028 | 1.235797 | 1.597819 | -0.217277 | 0.712908 | 0.929653 | -0.368851 | 20.625 | -0.15627 | -0.248042 |
| 20.5486111 | 0.418748 | 0.386926 | 0.944949 | 1.936371 | 0.676389 | 0.552826 | -0.223175 | 0.080837 | 0.660514 | -0.206241 | 20.6388889 | -0.075571 | -0.226412 |
| 20.5625 | 0.177455 | 0.277527 | 0.765704 | 2.036307 | 0.336866 | -0.182781 | -0.16296 | -0.169445 | 0.208604 | 0.006463 | 20.6527778 | -0.031667 | -0.09744 |
| 20.5763889 | 0.125422 | -0.036769 | 0.414299 | 0.985986 | 0.15326 | -0.110793 | -0.113736 | 0.186422 | 0.062625 | 0.276686 | 20.6666667 | -0.052549 | -0.023082 |
| 20.5902778 | 0.13982 | -0.169994 | 0.193447 | -0.033743 | 0.448634 | 0.405148 | -0.106944 | 0.679217 | 0.187928 | 0.323702 | 20.6805556 | -0.085536 | -0.035182 |
| 20.6041667 | 0.146246 | -0.121877 | 0.200297 | 0.042796 | 0.656916 | 0.621021 | -0.117123 | 0.667966 | 0.443753 | 0.214596 | 20.6944444 | -0.068855 | -0.100736 |
| 20.6180556 | 0.162013 | -0.020235 | 0.389097 | 0.101761 | 0.411042 | 0.312969 | -0.122 | 0.265577 | 0.914174 | 0.176805 | 20.7083333 | -0.021496 | -0.114966 |
| 20.6319444 | 0.148373 | 0.08166 | 0.345213 | 0.174148 | 0.188015 | 0.208503 | -0.080633 | -0.0979 | 0.763746 | 0.104619 | 20.7222222 | 0.004674 | -0.022203 |
| 20.6458333 | 0.084179 | -0.075478 | 0.029638 | 0.524626 | 0.741528 | 0.386488 | 0.006568 | 0.007982 | 0.669202 | -0.048831 | 20.7361111 | -0.007532 | 0.066863 |
| 20.6597222 | 0.023992 | -0.112234 | 0.017026 | 0.61086 | 0.814325 | 0.275307 | 0.031436 | 0.259588 | 0.062911 | -0.159443 | 20.75 | -0.014887 | 0.146921 |
| 20.6736111 | 0.006584 | -0.035826 | 0.160651 | 0.251794 | 0.296858 | -0.166931 | -0.001174 | 0.285608 | 0.180226 | -0.056004 | 20.7638889 | 0.005585 | 0.193853 |
| 20.6875 | 0.150924 | -0.034055 | 0.33076 | 0.150137 | -0.117188 | -0.375864 | -0.013411 | 0.074018 | 0.201128 | 0.145479 | 20.7777778 | 0.032041 | 0.22369 |
| 20.7013889 | 0.298565 | 0.052639 | 0.589409 | 0.383185 | -0.053263 | 0.147032 | -0.009458 | -0.118417 | 0.290356 | 0.271681 | 20.7916667 | 0.060395 | 0.265064 |
| 20.7152778 | 0.320006 | 0.112059 | 0.332749 | 0.321362 | 0.312011 | 0.721732 | 0.098189 | -0.136173 | 0.305198 | 0.119673 | 20.8055556 | 0.068636 | 0.306318 |
| 20.7291667 | 0.244421 | 0.110937 | 0.131943 | 0.22316 | 0.23281 | 0.479661 | 0.049037 | -0.071442 | 0.12471 | -0.074867 | 20.8194444 | 0.06182 | 0.332233 |
| 20.7430556 | 0.163367 | 0.059009 | 0.146816 | 0.176561 | -0.13762 | -0.231981 | -0.074132 | -0.055075 | -0.001465 | -0.040276 | 20.8333333 | 0.085261 | 0.342147 |
| 20.7569444 | 0.058433 | 0.002966 | 0.147585 | 0.054602 | -0.506377 | -0.742134 | -0.04433 | -0.052851 | -0.180105 | 0.137429 | 20.8472222 | 0.122407 | 0.265686 |
| 20.7708333 | -0.061834 | -0.123263 | 0.140997 | -0.118045 | -0.309009 | -0.125979 | 0.002006 | 0.008641 | -0.052091 | 0.157592 | 20.8611111 | 0.133576 | 0.04166 |
| 20.7847222 | -0.146932 | -0.255196 | 0.053579 | -0.156323 | 0.513132 | 0.82486 | 0.037642 | 0.118925 | 0.248047 | -0.076872 | 20.875 | 0.081405 | -0.222723 |
| 20.7986111 | -0.234565 | -0.213894 | 0.015184 | -0.008683 | 1.255406 | 1.586125 | -0.046225 | 0.06309 | 0.134139 | -0.248994 | 20.8888889 | 0.02969 | -0.375 |
| 20.8125 | -0.32739 | -0.25005 | 0.018926 | 0.224306 | 0.581261 | 0.755889 | -0.193835 | -0.2386 | -0.450266 | -0.308973 | 20.9027778 | 0.055935 | -0.465834 |
| 20.8263889 | -0.36166 | -0.277306 | -0.152564 | 0.120339 | -0.748659 | -0.748567 | -0.272843 | -0.500949 | -0.739241 | -0.210786 | 20.9166667 | 0.1177 | -0.52944 |
| 20.8402778 | -0.376855 | -0.326159 | -0.533437 | -0.41086 | -0.808678 | -0.804203 | -0.34657 | -0.42447 | -0.457806 | -0.175631 | 20.9305556 | 0.112921 | -0.622583 |
| 20.8541667 | -0.471937 | -0.430525 | -0.712301 | -0.835315 | -0.2951 | -0.261096 | -0.447977 | -0.120697 | -0.187395 | -0.13845 | 20.9444444 | 0.024852 | -0.720417 |
| 20.8680556 | -0.549609 | -0.360345 | -0.47981 | -0.744164 | -0.415138 | -0.08992 | -0.232642 | -0.098383 | -0.404877 | -0.034034 | 20.9583333 | -0.012363 | -0.690097 |
| 20.8819444 | -0.50849 | -0.236166 | -0.245103 | -0.6115 | -1.153761 | -0.827037 | -0.158091 | -0.483422 | -0.616023 | 0.017866 | 20.9722222 | 0.043026 | -0.485818 |
| 20.8958333 | -0.473019 | -0.218874 | -0.33249 | -0.854203 | -1.513635 | -1.579272 | 0.256418 | -0.771043 | -0.470676 | -0.067849 | 20.9861111 | 0.060698 | -0.271145 |

| | | | | | | | | | | | | | | | | | | | | | | | | | | | | |
|---|---|---|---|---|---|---|---|---|---|---|---|---|---|---|---|---|---|---|---|---|---|---|---|---|---|---|---|---|
| 20.9097222 | -1.163759 | -0.777224 | -2.753793 | -0.602209 | -0.244496 | -0.765857 | -0.124424 | -0.155574 | -0.445322 | -1.084923 | -0.670835 | -0.275067 | -0.088614 | -0.695291 | -0.542941 | -0.293236 | -0.784626 | -1.20159 | -1.262285 | -1.566724 | -0.066065 | -0.827864 | -0.427534 | -0.27566 | | 21 | 0.076262 | -0.14253 |
| 20.9236111 | -1.362111 | -1.066355 | -1.106682 | -0.557499 | -0.526674 | -1.264116 | -0.12338 | -0.136182 | -1.169104 | -2.062087 | 0.923781 | -0.901836 | -0.074093 | -0.67614 | -0.642372 | -0.17774 | -1.001697 | -1.582373 | -1.055948 | -1.302093 | -0.423272 | -0.851391 | -1.055619 | -2.281844 | | 21.0138889 | 0.713086 | 0.490418 |
| 20.9375 | -1.609697 | -0.915749 | -0.874081 | -0.077821 | -0.744927 | -1.168865 | -0.008559 | 0.295071 | -2.509981 | -4.249661 | 7.361346 | -1.787717 | -0.520703 | -0.101619 | -0.410316 | 0.676837 | -0.528398 | -2.367255 | -1.145833 | -1.245451 | 0.095864 | -1.297952 | -4.009807 | -14.603905 | | 21.0277778 | 4.243004 | 2.811655 |
| 20.9513889 | -1.963595 | -1.586823 | -1.807567 | 1.072571 | -0.540786 | 0.341291 | 0.412823 | 1.461773 | -2.900773 | -8.17528 | 15.615426 | -1.528288 | -0.720318 | 1.407303 | -0.408559 | 1.172548 | -0.375032 | -3.266636 | -0.804462 | -1.852905 | -1.483592 | -1.661957 | -7.127744 | -33.903904 | | 21.0416667 | 8.89474 | 4.826703 |
| 20.9652778 | -1.056317 | -1.510408 | -1.019883 | 1.596478 | -0.3119 | 1.186458 | 0.740585 | 1.539248 | -2.164805 | -7.090876 | 11.600601 | -1.039464 | -0.908888 | 1.524388 | -0.759783 | 0.406256 | -0.317367 | -2.261742 | -0.175962 | -1.633456 | -1.763615 | -1.518553 | -5.958908 | -30.765092 | | 21.0555556 | 6.963785 | 5.737116 |
| 20.9791667 | -0.977497 | -1.329726 | -0.463073 | 0.733311 | -0.194802 | 0.208141 | 0.233489 | 0.635832 | -1.022001 | -3.057259 | 5.004704 | -0.753984 | 0.016199 | 0.47262 | -0.545233 | -0.003376 | -0.175004 | -0.730304 | -0.356674 | -1.351815 | -0.664813 | -0.829615 | -2.741914 | -12.446126 | | 21.0694444 | 2.6714 | 2.21514 |
| 20.9930556 | -0.534042 | -0.840223 | -0.219348 | 0.153515 | -0.29195 | -0.519214 | -0.052706 | 0.053457 | -0.18025 | -0.80579 | 0.977611 | -0.787569 | 0.38224 | -0.385777 | -0.312316 | -0.037749 | -0.296414 | -0.147477 | -0.562177 | -1.091307 | -0.09037 | -0.44113 | -1.33298 | -2.884137 | | 21.0833333 | 0.361889 | 0.427342 |
| 21.0069444 | -0.248808 | -0.291922 | -0.278566 | -0.098763 | -0.331853 | -0.672388 | -0.071745 | -0.074036 | -0.158005 | -0.392841 | 0.307479 | -0.987399 | -0.388504 | -0.723546 | -0.318647 | -0.056486 | -0.564644 | -0.27492 | -0.616073 | -0.876458 | -0.04514 | -0.316323 | -1.122362 | -1.451927 | | 21.0972222 | 0.027941 | 0.107966 |
| 21.0208333 | -0.222724 | -0.149109 | -0.69263 | -0.317936 | -0.376702 | -0.825282 | -0.113545 | -0.11393 | -0.370093 | -0.338264 | -0.15355 | -1.010892 | -0.877526 | -0.575885 | -0.300121 | -0.140878 | -0.595213 | -0.417509 | -0.637945 | -0.674423 | -0.041168 | -0.233145 | -0.772765 | -0.67492 | | 21.1111111 | 0.081164 | 0.151477 |
| 21.0347222 | -0.195834 | -0.203121 | -0.82639 | -0.423494 | -0.41387 | -1.033947 | -0.204727 | -0.151329 | -0.509274 | -0.365448 | -0.275871 | -0.678402 | -0.468842 | -0.212922 | -0.239706 | -0.184141 | -0.481912 | -0.479348 | -0.623168 | -0.489493 | 0.002189 | -0.224963 | -0.329672 | -0.082147 | | 21.125 | 0.097272 | 0.272243 |
| 21.0486111 | -0.064858 | -0.169528 | -0.682964 | -0.437215 | -0.408349 | -0.980666 | -0.248921 | -0.139794 | -0.399568 | -0.336026 | -0.011655 | -0.398927 | -0.176832 | -0.072951 | -0.206828 | -0.194379 | -0.454786 | -0.538195 | -0.511534 | -0.457226 | -0.007403 | -0.255644 | -0.133099 | 0.002388 | | 21.1388889 | 0.055657 | 0.246525 |
| 21.0625 | -0.026691 | -0.13068 | -0.72616 | -0.492378 | -0.302178 | -0.693376 | -0.218967 | -0.088748 | -0.193432 | -0.194964 | 0.127991 | -0.362172 | -0.21659 | -0.184754 | -0.234083 | -0.209346 | -0.496725 | -0.609334 | -0.424561 | -0.484832 | -0.048033 | -0.26216 | -0.12147 | -0.287586 | | 21.1527778 | -0.017741 | 0.040946 |
| 21.0763889 | -0.085519 | -0.169951 | -0.860351 | -0.5268 | -0.216155 | -0.493578 | -0.092099 | -0.045216 | -0.132282 | -0.056699 | 0.013551 | -0.413541 | -0.350745 | -0.269954 | -0.223884 | -0.185368 | -0.462997 | -0.513595 | -0.357042 | -0.413612 | -0.039713 | -0.244068 | -0.117596 | -0.352053 | | 21.1666667 | -0.054629 | -0.1009 |
| 21.0902778 | -0.043744 | -0.166851 | -0.695498 | -0.555236 | -0.22964 | -0.484192 | 0.021236 | -0.030766 | -0.158442 | -0.0651 | -0.057889 | -0.379015 | -0.28322 | -0.27837 | -0.19341 | -0.165709 | -0.387291 | -0.355304 | -0.294704 | -0.29209 | -0.022924 | -0.258063 | -0.105714 | -0.164376 | | 21.1805556 | -0.006872 | -0.074671 |
| 21.1041667 | 0.03839 | -0.106305 | -0.385774 | -0.438542 | -0.257985 | -0.508628 | 0.048593 | -0.036046 | -0.119432 | -0.146356 | 0.035127 | -0.287003 | -0.101131 | -0.275382 | -0.18571 | -0.163088 | -0.334698 | -0.261121 | -0.281732 | -0.251741 | -0.040498 | -0.261986 | -0.146732 | -0.230837 | | 21.1944444 | 0.030373 | -0.073743 |
| 21.1180556 | 0.013952 | -0.090702 | -0.143048 | -0.209418 | -0.233209 | -0.417983 | 0.022058 | -0.043813 | -0.05054 | -0.165353 | 0.087771 | -0.171279 | 0.018978 | -0.205937 | -0.123853 | -0.096828 | -0.258217 | -0.172083 | -0.248676 | -0.207058 | -0.000985 | -0.184577 | -0.199948 | -0.18729 | | 21.2083333 | 0.052522 | -0.136807 |
| 21.1319444 | -0.085178 | -0.11098 | 0.01493 | -0.047073 | -0.184564 | -0.316666 | 0.002962 | -0.048257 | -0.019297 | -0.078797 | -0.117062 | -0.103052 | -0.018237 | -0.105098 | -0.006389 | 0.021963 | -0.157456 | -0.060188 | -0.126595 | -0.071246 | 0.077747 | -0.071518 | -0.162405 | 0.149313 | | 21.2222222 | 0.084858 | -0.136426 |
| 21.1458333 | -0.078618 | -0.095386 | 0.079824 | -0.070612 | -0.142166 | -0.246285 | 0.015472 | -0.027409 | 0.021727 | 0.057491 | -0.36747 | -0.112914 | -0.104319 | -0.059831 | 0.031399 | 0.082535 | -0.101289 | -0.025324 | -0.011964 | 0.026799 | 0.067677 | -0.014844 | -0.06806 | 0.08049 | | 21.2361111 | 0.098611 | -0.042325 |
| 21.1597222 | 0.071188 | -0.009498 | -0.034906 | -0.510967 | -0.098218 | -0.114088 | 0.02119 | -0.005565 | 0.096555 | 0.105107 | -0.152537 | -0.089993 | -0.049271 | -0.027989 | 0.002707 | 0.056814 | -0.072348 | -0.079677 | 0.022739 | 0.030878 | -0.011747 | -0.008339 | -0.033258 | -0.383324 | | 21.25 | 0.105365 | 0.035091 |
| 21.1736111 | 0.190108 | 0.095684 | 0.177892 | -0.142567 | -0.044234 | 0.044837 | 0.015048 | 0.008867 | 0.162419 | 0.083662 | 0.343772 | 0.032562 | 0.161974 | 0.015409 | 0.011682 | 0.051855 | -0.019225 | -0.099451 | 0.053254 | 0.075921 | -0.010667 | 0.040554 | 0.008957 | -0.358239 | | 21.2638889 | 0.06977 | 0.035424 |
| 21.1875 | 0.177157 | 0.162751 | 0.279752 | -0.02555 | 0.023861 | 0.136758 | 0.044686 | 0.024411 | 0.177943 | 0.070335 | 0.467409 | 0.137103 | 0.31596 | 0.051606 | 0.081702 | 0.105976 | 0.034185 | -0.034033 | 0.140627 | 0.179432 | 0.075612 | 0.10249 | 0.121006 | 0.275839 | | 21.2777778 | -0.017262 | 0.013431 |
| 21.2013889 | 0.134142 | 0.171866 | 0.433086 | 0.220365 | 0.072081 | 0.129461 | 0.084629 | 0.037338 | 0.127387 | 0.033227 | 0.253889 | 0.110714 | 0.225499 | 0.049921 | 0.117234 | 0.145616 | 0.082224 | 0.050151 | 0.220964 | 0.199571 | 0.109344 | 0.104488 | 0.161656 | 0.5146 | | 21.2916667 | -0.06566 | 0.027375 |
| 21.2152778 | 0.231849 | 0.219363 | 0.576141 | 0.55187 | 0.108084 | 0.166155 | 0.123192 | 0.085178 | 0.158158 | 0.087779 | 0.100903 | 0.07014 | 0.111741 | 0.032511 | 0.083421 | 0.123512 | 0.13332 | 0.11929 | 0.226848 | 0.17875 | 0.074706 | 0.1731 | 0.144939 | 0.062922 | | 21.3055556 | -0.041473 | 0.057855 |
| 21.2291667 | 0.337543 | 0.250357 | 0.536096 | 0.571025 | 0.129003 | 0.275068 | 0.168363 | 0.149873 | 0.255036 | 0.153722 | 0.122548 | 0.133798 | 0.187012 | 0.060991 | 0.045919 | 0.091296 | 0.175844 | 0.186369 | 0.197033 | 0.196012 | 0.037061 | 0.287136 | 0.190326 | -0.301486 | | 21.3194444 | 0.011272 | 0.038221 |
| 21.2430556 | 0.235581 | 0.179275 | 0.464006 | 0.438823 | 0.099919 | 0.221748 | 0.121878 | 0.151439 | 0.21919 | 0.093789 | 0.139347 | 0.276796 | 0.340954 | 0.136961 | 0.077458 | 0.106772 | 0.206156 | 0.273098 | 0.229528 | 0.246003 | 0.059083 | 0.249863 | 0.205069 | -0.086154 | | 21.3333333 | 0.046738 | 0.00461 |
| 21.2569444 | 0.194352 | 0.263768 | 0.580155 | 0.455302 | 0.078762 | 0.126646 | 0.07833 | 0.135618 | 0.186842 | 0.136338 | 0.032297 | 0.400508 | 0.362529 | 0.161195 | 0.153482 | 0.155852 | 0.230038 | 0.38663 | 0.352721 | 0.368913 | 0.119762 | 0.207406 | 0.210889 | 0.408177 | | 21.3472222 | 0.048397 | 0.055349 |
| 21.2708333 | 0.284802 | 0.408583 | 0.618843 | 0.496032 | 0.093436 | 0.175519 | 0.124748 | 0.15106 | 0.236811 | 0.230853 | -0.012662 | 0.381744 | 0.283437 | 0.156046 | 0.178186 | 0.155191 | 0.261199 | 0.437513 | 0.404853 | 0.462447 | 0.102537 | 0.318488 | 0.242647 | 0.535358 | | 21.3611111 | 0.077699 | 0.139214 |
| 21.2847222 | 0.285238 | 0.309192 | 0.494175 | 0.433063 | 0.111445 | 0.268346 | 0.12766 | 0.114985 | 0.237901 | 0.17892 | 0.106739 | 0.300147 | 0.274378 | 0.12952 | 0.141831 | 0.098742 | 0.283885 | 0.397917 | 0.315931 | 0.397727 | 0.038649 | 0.339411 | 0.214302 | 0.276915 | | 21.375 | 0.132555 | 0.130078 |
| 21.2986111 | 0.303603 | 0.265187 | 0.586762 | 0.412465 | 0.174309 | 0.310123 | 0.051603 | 0.07569 | 0.288391 | 0.165338 | 0.256966 | 0.324833 | 0.314435 | 0.224398 | 0.1496 | 0.116165 | 0.331958 | 0.411817 | 0.330991 | 0.36786 | 0.030297 | 0.363424 | 0.226192 | -0.038682 | | 21.3888889 | 0.164447 | 0.148419 |
| 21.3125 | 0.359986 | 0.376362 | 0.876955 | 0.555743 | 0.239876 | 0.31534 | 0.028356 | 0.087085 | 0.351271 | 0.222587 | 0.245195 | 0.395576 | 0.26421 | 0.340566 | 0.180378 | 0.186415 | 0.386884 | 0.48034 | 0.450189 | 0.443587 | 0.028551 | 0.590988 | 0.279798 | -0.258803 | | 21.4027778 | 0.14776 | 0.262768 |
| 21.3263889 | 0.322268 | 0.38334 | 0.837305 | 0.580718 | 0.191098 | 0.28733 | 0.079135 | 0.092099 | 0.290194 | 0.197887 | 0.099382 | 0.390507 | 0.204382 | 0.27711 | 0.180369 | 0.202812 | 0.382406 | 0.407695 | 0.390272 | 0.44347 | 0.03414 | 0.60943 | 0.258726 | -0.002797 | | 21.4166667 | 0.022169 | 0.284804 |
| 21.3402778 | 0.347548 | 0.416052 | 0.749751 | 0.458323 | 0.174303 | 0.343212 | 0.143064 | 0.147736 | 0.277822 | 0.231736 | 0.136421 | 0.446342 | 0.254062 | 0.182339 | 0.216653 | 0.201362 | 0.374629 | 0.31594 | 0.390159 | 0.57979 | 0.10218 | 0.495872 | 0.358124 | 0.604351 | | 21.4305556 | -0.083988 | 0.236914 |
| 21.3541667 | 0.44735 | 0.530564 | 0.958053 | 0.414493 | 0.278839 | 0.455583 | 0.197901 | 0.227493 | 0.374754 | 0.329385 | 0.363284 | 0.586162 | 0.290391 | 0.290003 | 0.30907 | 0.261986 | 0.445927 | 0.348903 | 0.637088 | 0.86891 | 0.156772 | 0.701111 | 0.581828 | 0.926209 | | 21.4444444 | -0.108306 | 0.233678 |
| 21.3680556 | 0.532579 | 0.612904 | 1.04345 | 0.428499 | 0.328784 | 0.461975 | 0.189247 | 0.225907 | 0.415368 | 0.383756 | 0.417185 | 0.701035 | 0.330396 | 0.501251 | 0.430517 | 0.360667 | 0.599892 | 0.554136 | 0.776599 | 0.929658 | 0.17378 | 1.017601 | 0.739529 | 1.047336 | | 21.4583333 | -0.198236 | 0.234278 |
| 21.3819444 | 0.712751 | 0.733028 | 1.072029 | 0.510972 | 0.373522 | 0.569456 | 0.19782 | 0.240051 | 0.487176 | 0.515492 | 0.347773 | 0.867583 | 0.461827 | 0.659143 | 0.585051 | 0.477452 | 0.766391 | 0.780949 | 0.808409 | 0.94692 | 0.213941 | 1.116587 | 0.892726 | 0.989302 | | 21.4722222 | -0.253689 | 0.340613 |
| 21.3958333 | 0.977762 | 0.911577 | 1.422568 | 0.660801 | 0.499949 | 0.864225 | 0.286824 | 0.341379 | 0.695883 | 0.740168 | 0.357255 | 1.050856 | 0.540507 | 0.723748 | 0.673631 | 0.582977 | 0.851807 | 0.810011 | 1.083969 | 1.252916 | 0.213028 | 1.00273 | 1.098725 | 0.398615 | | 21.4861111 | -0.347105 | 0.465112 |
| 21.4097222 | 1.136822 | 0.994011 | 1.910534 | 0.649993 | 0.625908 | 1.153362 | 0.390527 | 0.435044 | 0.919132 | 0.918906 | 0.390585 | 1.197073 | 0.501943 | 0.769486 | 0.685807 | 0.614285 | 0.895128 | 0.864757 | 1.464429 | 1.585631 | 0.192829 | 0.876369 | 1.281133 | 0.025148 | | 21.5 | -0.461024 | 0.416011 |
| 21.4236111 | 1.315861 | 1.13644 | 2.096507 | 0.359387 | 0.672212 | 1.218706 | 0.469059 | 0.425121 | 1.17083 | 1.348715 | 0.154915 | 1.385422 | 0.530395 | 0.836977 | 0.796216 | 0.604378 | 1.011078 | 1.058534 | 1.531512 | 1.790435 | 0.291923 | 0.838271 | 1.703448 | 1.992846 | | 21.5138889 | -0.380375 | 0.004068 |
| 21.4375 | 1.698632 | 1.598298 | 1.395914 | -0.587903 | 0.581237 | 0.569508 | 0.414308 | -0.074681 | 1.960203 | 3.488277 | -1.294759 | 1.46222 | 0.718856 | 0.289937 | 1.262852 | 0.394358 | 0.904223 | 1.251659 | 1.131278 | 1.940692 | 0.907402 | 1.315494 | 3.336023 | 13.68222 | | 21.5277778 | -2.078631 | -2.228093 |
| 21.4513889 | 2.625092 | 2.318399 | 0.334272 | -2.182966 | 0.359521 | -0.763837 | 0.1534 | -0.813655 | 3.18716 | 7.699929 | -5.81807 | 1.404358 | 0.879576 | -1.160868 | 1.493945 | -0.484052 | 0.336403 | 1.463386 | 1.031893 | 1.950984 | 1.965311 | 2.989385 | 5.883815 | 34.854497 | | 21.5416667 | -6.457039 | -6.190531 |

| | | | | | | | | | | | | | | | | | | | | | | | | | | | | |
|---|---|---|---|---|---|---|---|---|---|---|---|---|---|---|---|---|---|---|---|---|---|---|---|---|---|---|---|---|
| 23.8402778 | -0.365406 | -0.588271 | -0.644216 | -0.327063 | -0.755236 | -2.010198 | 0.118422 | -0.887201 | 0.066421 | -1.757214 | 0.279374 | -0.412825 | 0.210585 | -0.312271 | -0.015472 | 0.246452 | 0.541296 | 0.268089 | 1.628375 | 0.518077 | 0.725892 | 1.088693 | 2.110544 | -0.250685 | | 23.9305556 | -0.361722 | 0.113209 |
| 23.8541667 | -0.670008 | -1.891144 | -2.22889 | -1.614512 | -0.659317 | -1.138505 | -0.284985 | -0.576475 | -0.462785 | -1.027428 | -0.910113 | -0.427405 | 0.692732 | -0.511716 | -0.102835 | -0.173138 | -0.487224 | -0.944911 | -0.887059 | -1.093645 | 0.469102 | 1.363433 | 0.970804 | -1.37426 | | 23.9444444 | -0.297414 | -0.228259 |
| 23.8680556 | -0.597547 | -1.520481 | -3.004908 | -2.178208 | -0.697314 | -0.953891 | -0.921287 | -0.279905 | -1.324333 | -0.18611 | -1.88256 | -1.24492 | -1.883662 | -0.092714 | -0.056098 | -0.472198 | -0.803429 | -1.439682 | -1.634372 | -1.856625 | -0.440503 | -1.087737 | -2.261715 | -1.381974 | | 23.9583333 | -0.075427 | -0.475566 |
| 23.8819444 | -0.588552 | -0.931557 | -1.801561 | -1.479584 | -0.9274 | -1.698679 | -0.800618 | -0.547067 | -1.424563 | -0.588767 | -1.429931 | -1.964539 | -2.454873 | -0.205954 | -0.441677 | -0.3474 | -0.641451 | -1.312439 | -0.384508 | -1.3626 | -0.914367 | -2.473007 | -2.293255 | -1.006752 | | 23.9722222 | 0.040612 | -0.600873 |
| 23.8958333 | -0.895533 | -1.592441 | -1.559409 | -1.767548 | -1.063303 | -1.949186 | -0.52999 | -0.99522 | -0.826177 | -1.43995 | -0.413335 | -2.074736 | -1.479357 | -1.015777 | -0.745209 | -0.604492 | -0.310677 | -1.411477 | -0.612092 | -1.493861 | -1.159026 | -0.359736 | -1.723249 | -1.070421 | | 23.9861111 | 0.03718 | -0.947533 |
| 23.9097222 | -1.647432 | -2.338548 | -2.729101 | -2.194303 | -1.312587 | -2.021851 | -0.500431 | -1.18365 | -0.919394 | -1.994304 | -1.217322 | -1.731862 | -0.999614 | -1.580742 | -0.954437 | -0.791605 | -0.669783 | -2.387191 | -1.90006 | -1.583203 | -1.117833 | 2.053737 | -0.473464 | -1.690671 | | 24 | 0.019923 | -1.001584 |
| 23.9236111 | -2.19116 | -2.734083 | -4.175255 | -2.053072 | -1.403372 | -2.034213 | -0.762763 | -0.855345 | -1.615602 | -2.020588 | -3.551359 | -2.230444 | -1.854689 | -1.01557 | -0.938499 | -0.937969 | -1.623298 | -2.728138 | -2.518307 | -2.028942 | -1.301854 | -0.858546 | -1.084475 | -3.722216 | | 24.0138889 | 0.711522 | -0.591529 |
| 23.9375 | -2.921267 | -2.346135 | -4.251115 | -1.870402 | -1.459349 | -1.724596 | -1.445897 | -0.714765 | -3.132597 | -3.620605 | -5.658674 | -3.211467 | -1.833965 | -0.620169 | -1.032728 | -1.095807 | -2.029573 | -1.905788 | -2.06881 | -2.662221 | -1.147966 | -4.570644 | -3.392912 | -15.015104 | | 24.0277778 | 4.632477 | 0.796546 |
| 23.9513889 | -3.784856 | -2.38884 | -3.271775 | -1.824069 | -1.019478 | -0.56445 | -1.939169 | -0.1237 | -4.782724 | -6.889016 | -5.619608 | -3.248411 | -1.077866 | -0.181726 | -1.254552 | -0.744662 | -1.476872 | -1.57132 | -0.486806 | -3.047637 | -0.006145 | -6.057189 | -5.18059 | -33.777039 | | 24.0416667 | 9.271693 | 4.177643 |
| 23.9652778 | -2.693463 | -2.066076 | -1.896181 | -0.619093 | -0.143054 | 1.285285 | -1.490833 | 0.226693 | -3.827481 | -6.310381 | -5.108811 | -2.594708 | -0.791395 | -0.490641 | -1.430495 | -0.802633 | -0.978846 | -1.66967 | -0.402872 | -2.299466 | -0.336851 | -3.606379 | -4.399131 | -31.671038 | | 24.0555556 | 7.297442 | 2.438708 |
| 23.9791667 | -1.233924 | -1.142173 | -0.933521 | -0.003685 | 0.257982 | 0.829402 | -0.244498 | 0.173064 | -1.44825 | -3.007643 | -1.908439 | -2.141425 | -1.161161 | -1.93003 | -1.880382 | -0.950516 | -1.008224 | -2.168295 | -1.094809 | -1.52547 | -0.319382 | -1.477338 | -2.268567 | -14.002405 | | 24.0694444 | 2.690354 | 1.416783 |
| 23.9930556 | -0.091997 | -0.402236 | -0.354868 | 0.219763 | 0.77347 | 0.905304 | 1.172943 | 0.318553 | 0.736575 | -0.41924 | -1.274909 | -1.689658 | -1.201553 | -2.076102 | -1.723424 | -0.670563 | -1.023054 | -2.267474 | -1.199979 | -0.819577 | -0.07821 | -0.443639 | -0.785369 | -2.879479 | | 24.0833333 | 0.175116 | -0.101641 |
| 24.0069444 | -0.092565 | -1.000756 | -1.735008 | -0.460736 | 0.905323 | 1.107335 | 1.826195 | 0.072816 | 1.158785 | 0.922175 | -3.361388 | -1.128786 | -0.44008 | -0.904905 | -0.635229 | 0.257114 | -0.886763 | -1.730729 | -0.836732 | -0.613602 | -0.163633 | -0.809917 | -0.273691 | -0.272059 | | 24.0972222 | -0.27595 | -1.292501 |
| 24.0208333 | -0.419945 | -1.936258 | -3.730569 | -1.80083 | -0.015679 | -0.125125 | 0.578581 | -1.065008 | 1.029532 | 0.778602 | -5.35862 | -1.166683 | -0.524354 | -0.500848 | -0.288139 | 0.101922 | -0.890337 | -1.44999 | -0.756931 | -1.021775 | -0.796948 | -1.502701 | -0.398514 | 0.166354 | | 24.1111111 | -0.18539 | -1.549685 |
| 24.0347222 | 0.212106 | 0.908961 | 0.475733 | -0.197813 | 0.362376 | 0.966203 | 0.293884 | -0.194583 | 1.313786 | 1.100748 | -0.92963 | -1.451064 | -1.102234 | -0.972338 | -0.721577 | -0.744143 | -1.143039 | -1.417192 | -0.982669 | -1.6734 | -1.474694 | -1.642279 | -0.934323 | -0.486354 | | 24.125 | 0.028486 | -0.861513 |
| 24.0486111 | 2.054994 | 3.22117 | 3.645404 | 1.094391 | 1.299287 | 2.801881 | 0.555962 | 0.450568 | 2.491571 | 1.837809 | 2.499494 | -1.863714 | -1.27233 | -1.01429 | -0.938695 | -1.153099 | -1.285864 | -1.435236 | -1.253544 | -1.606081 | -1.171773 | -1.253094 | -1.14289 | -1.051322 | | 24.1388889 | 0.247532 | -0.347726 |
| 24.0625 | 1.974424 | 2.584567 | 2.87364 | 0.989127 | 1.285162 | 2.671045 | 0.523493 | 0.582421 | 1.772891 | 1.539759 | 2.290095 | -1.468065 | -1.016271 | -0.613296 | -0.618937 | -0.698536 | -0.86576 | -1.109086 | -1.060275 | -0.969915 | -0.477564 | -0.578741 | -0.853876 | -0.690672 | | 24.1527778 | 0.337069 | -0.207937 |
| 24.0763889 | 1.418302 | 1.670627 | 1.875782 | 0.702881 | 0.705134 | 1.6578 | 0.630048 | 0.587211 | 1.087901 | 1.191937 | 1.18654 | -0.695882 | -0.506178 | -0.598143 | -0.295687 | -0.118148 | -0.3892 | -0.696407 | -0.598399 | -0.467429 | -0.096918 | -0.113093 | -0.452743 | -0.162249 | | 24.1666667 | 0.213638 | -0.129412 |
| 24.0902778 | 1.204852 | 1.245397 | 1.335359 | 0.703308 | 0.520728 | 1.300256 | 0.941548 | 0.74894 | 1.083676 | 1.319414 | 0.557857 | -0.204059 | -0.185 | -0.726128 | -0.223141 | -0.053152 | -0.33295 | -0.472324 | -0.330293 | -0.400283 | -0.121925 | -0.236987 | -0.356375 | -0.224209 | | 24.1805556 | 0.049583 | -0.026162 |
| 24.1041667 | 1.276744 | 1.177991 | 0.925966 | 0.751486 | 0.508037 | 1.174727 | 0.991626 | 0.871029 | 1.200428 | 1.434165 | 0.288256 | -0.164489 | -0.1834 | -0.263637 | -0.234596 | -0.163606 | -0.524959 | -0.390708 | -0.252398 | -0.435472 | -0.189876 | -0.370851 | -0.442126 | -0.495903 | | 24.1944444 | 0.009639 | 0.138011 |
| 24.1180556 | 1.285918 | 1.021449 | 0.948096 | 0.695117 | 0.468054 | 0.997603 | 0.876552 | 0.802264 | 1.239671 | 1.37016 | 0.308676 | -0.231062 | -0.257142 | 0.018047 | -0.219722 | -0.10657 | -0.504341 | -0.308124 | -0.230234 | -0.288899 | -0.087999 | -0.129701 | -0.421061 | -0.481333 | | 24.2083333 | 0.054203 | 0.384889 |
| 24.1319444 | 1.029674 | 0.876265 | 1.377177 | 0.573274 | 0.434144 | 0.830269 | 0.797381 | 0.675585 | 1.323902 | 1.365837 | 0.617755 | -0.208937 | -0.232186 | -0.306678 | -0.248167 | -0.01959 | -0.311647 | -0.290072 | -0.270069 | -0.161979 | -0.040455 | 0.031439 | -0.30519 | -0.221409 | | 24.2222222 | 0.145 | 0.725029 |
| 24.1458333 | 0.854406 | 0.871229 | 1.64548 | 0.529369 | 0.376956 | 0.719788 | 0.808136 | 0.615788 | 1.237423 | 1.321013 | 0.793831 | -0.188669 | -0.290925 | -0.559471 | -0.238054 | -0.039897 | -0.204537 | -0.266694 | -0.282131 | -0.187905 | -0.094789 | -0.162328 | -0.238338 | -0.006997 | | 24.2361111 | 0.219677 | 1.0424 |
| 24.1597222 | 0.756198 | 0.906035 | 1.550739 | 0.473588 | 0.331664 | 0.602178 | 0.811555 | 0.58047 | 0.983932 | 1.178842 | 0.791643 | -0.125052 | -0.356406 | -0.305102 | -0.163542 | -0.07065 | -0.208228 | -0.164872 | -0.231028 | -0.202305 | -0.090563 | -0.215838 | -0.249431 | -0.019775 | | 24.25 | 0.185715 | 1.217582 |
| 24.1736111 | 0.757581 | 0.859052 | 1.376574 | 0.374691 | 0.259218 | 0.449124 | 0.621794 | 0.475529 | 0.723163 | 1.024713 | 0.874893 | -0.019739 | -0.280141 | -0.069834 | -0.125127 | -0.06394 | -0.265031 | -0.095924 | -0.215124 | -0.159418 | -0.014188 | 0.089104 | -0.317744 | -0.184036 | | 24.2638889 | 0.036648 | 1.154665 |
| 24.1875 | 0.723451 | 0.757725 | 1.099382 | 0.24359 | 0.159133 | 0.243437 | 0.348521 | 0.296915 | 0.524013 | 0.80812 | 0.876658 | 0.009391 | -0.141087 | -0.089161 | -0.110561 | -0.036452 | -0.240758 | -0.102496 | -0.19191 | -0.126789 | 0.070551 | 0.320511 | -0.328836 | -0.389117 | | 24.2777778 | -0.013647 | 0.914433 |
| 24.2013889 | 0.614018 | 0.588113 | 0.846964 | 0.166712 | 0.064811 | 0.083687 | 0.104976 | 0.15717 | 0.32705 | 0.638671 | 0.743056 | -0.069239 | -0.103115 | -0.10993 | -0.049665 | -0.048374 | -0.099489 | -0.088239 | -0.059786 | -0.108796 | 0.106737 | 0.204067 | -0.243758 | -0.410038 | | 24.2916667 | 0.105119 | 0.598577 |
| 24.2152778 | 0.480863 | 0.377914 | 0.596135 | 0.147194 | -0.021291 | -0.012883 | -0.08723 | 0.060544 | 0.087766 | 0.384409 | 0.538631 | -0.191066 | -0.15855 | 0.041614 | 0.031366 | -0.056743 | 0.017718 | -0.034133 | 0.079696 | -0.081374 | 0.072598 | -0.020457 | -0.172647 | -0.25975 | | 24.3055556 | 0.219425 | 0.433059 |
| 24.2291667 | 0.383458 | 0.289889 | 0.347595 | 0.025786 | -0.03973 | -0.037208 | -0.194666 | -0.019187 | -0.04321 | 0.187445 | 0.354512 | -0.236282 | -0.119508 | 0.205313 | 0.0914 | 0.018465 | 0.064102 | 0.074264 | 0.119722 | -0.038013 | -0.008935 | 0.005464 | -0.171856 | -0.253914 | | 24.3194444 | 0.23806 | 0.502021 |
| 24.2430556 | 0.339517 | 0.293496 | 0.167047 | -0.10693 | -0.017516 | -0.036958 | -0.166776 | -0.042376 | -0.03292 | 0.097788 | 0.283849 | -0.10591 | 0.058957 | 0.137149 | 0.083558 | 0.057121 | 0.141773 | 0.211646 | 0.172765 | 0.038598 | -0.016301 | 0.289511 | -0.115759 | -0.143899 | | 24.3333333 | 0.180867 | 0.688123 |
| 24.2569444 | 0.300101 | 0.225082 | 0.070236 | -0.094089 | -0.057769 | -0.077999 | -0.102398 | -0.019443 | -0.119197 | -0.045538 | 0.237309 | 0.079474 | 0.148945 | -0.014187 | 0.035278 | 0.006377 | 0.246907 | 0.252544 | 0.266155 | 0.123588 | 0.061351 | 0.402711 | 0.007948 | 0.234656 | | 24.3472222 | 0.073096 | 0.593509 |
| 24.2708333 | 0.240967 | 0.143214 | -0.063565 | -0.078129 | -0.142765 | -0.147546 | -0.187056 | -0.083569 | -0.305165 | -0.196479 | 0.029489 | 0.141577 | 0.124573 | 0.08189 | 0.03681 | 0.020531 | 0.289241 | 0.21827 | 0.309495 | 0.210027 | 0.12486 | 0.225462 | 0.033819 | 0.405926 | | 24.3611111 | 0.044964 | 0.43814 |
| 24.2847222 | 0.188534 | 0.085658 | -0.235352 | -0.112216 | -0.212137 | -0.194672 | -0.348407 | -0.189294 | -0.508658 | -0.293856 | -0.176343 | 0.212834 | 0.250883 | 0.278063 | 0.111834 | 0.170477 | 0.317244 | 0.327852 | 0.322446 | 0.283204 | 0.157564 | 0.193785 | 0.022115 | 0.201409 | | 24.375 | 0.138697 | 0.335155 |
| 24.2986111 | 0.154518 | 0.033666 | -0.241643 | -0.07506 | -0.197888 | -0.169623 | -0.303259 | -0.145417 | -0.512829 | -0.280372 | -0.106931 | 0.434491 | 0.543282 | 0.225235 | 0.195261 | 0.268516 | 0.410721 | 0.616524 | 0.329287 | 0.336389 | 0.212098 | 0.482628 | 0.171482 | 0.147696 | | 24.3888889 | 0.178028 | 0.335073 |
| 24.3125 | 0.102224 | 0.013815 | -0.116858 | -0.058849 | -0.098146 | -0.092165 | -0.123715 | -0.023913 | -0.28816 | -0.164561 | 0.034477 | 0.61941 | 0.63717 | 0.174936 | 0.242893 | 0.23203 | 0.491989 | 0.739981 | 0.330137 | 0.363802 | 0.26878 | 0.662568 | 0.322288 | 0.24224 | | 24.4027778 | 0.146694 | 0.38697 |
| 24.3263889 | 0.055706 | 0.037433 | -0.031225 | -0.106381 | -0.015185 | -0.04246 | -0.027332 | 0.02099 | -0.097794 | -0.062154 | 0.083369 | 0.636453 | 0.549282 | 0.365462 | 0.273859 | 0.201384 | 0.433461 | 0.542532 | 0.277526 | 0.371846 | 0.307754 | 0.458372 | 0.254274 | 0.115096 | | 24.4166667 | 0.201331 | 0.337862 |
| 24.3402778 | 0.042788 | 0.071094 | 0.032645 | -0.076138 | 0.026312 | -0.010262 | 0.02613 | 0.038357 | -0.057644 | -0.024382 | 0.095494 | 0.646329 | 0.705472 | 0.477223 | 0.412061 | 0.333125 | 0.36011 | 0.372273 | 0.334807 | 0.425866 | 0.316426 | 0.357603 | 0.250949 | 0.165802 | | 24.4305556 | 0.230393 | 0.189845 |
| 24.3541667 | 0.052195 | 0.105883 | 0.131762 | 0.026404 | 0.060744 | 0.039857 | 0.11785 | 0.07616 | -0.037185 | 0.025708 | 0.093607 | 0.858534 | 1.110781 | 0.469549 | 0.719458 | 0.583695 | 0.501646 | 0.581614 | 0.677314 | 0.58234 | 0.334064 | 0.779315 | 0.566271 | 0.599356 | | 24.4444444 | 0.063527 | 0.055706 |
| 24.3680556 | 0.115379 | 0.228708 | 0.257809 | 0.10821 | 0.119207 | 0.100412 | 0.248994 | 0.12931 | 0.101168 | 0.154583 | 0.130785 | 1.17447 | 1.395356 | 0.760193 | 1.104372 | 0.824489 | 0.801773 | 1.05363 | 1.047409 | 0.83871 | 0.425805 | 1.276149 | 0.977868 | 0.976743 | | 24.4583333 | -0.2112 | 0.048735 |
| 24.3819444 | 0.230078 | 0.43493 | 0.39001 | 0.169273 | 0.210855 | 0.186423 | 0.382034 | 0.207873 | 0.329213 | 0.31478 | 0.262611 | 1.434537 | 1.38242 | 1.377037 | 1.409663 | 1.009487 | 0.963457 | 1.234251 | 1.212569 | 1.144161 | 0.556041 | 1.470328 | 1.229119 | 0.964945 | | 24.4722222 | -0.310282 | 0.219259 |
| 24.3958333 | 0.299654 | 0.573857 | 0.48544 | 0.244369 | 0.28673 | 0.287531 | 0.486975 | 0.298877 | 0.454792 | 0.386077 | 0.343898 | 1.779391 | 1.552585 | 1.726197 | 1.624016 | 1.261465 | 1.061485 | 1.296936 | 1.419386 | 1.512305 | 0.743702 | 1.623208 | 1.508294 | 0.977061 | | 24.4861111 | -0.234716 | 0.37552 |
| 24.4097222 | 0.299094 | 0.552216 | 0.543828 | 0.326712 | 0.285487 | 0.34039 | 0.587492 | 0.36997 | 0.431835 | 0.370418 | 0.367824 | 2.465097 | 2.160119 | 1.716353 | 1.761156 | 1.423696 | 1.520378 | 1.914805 | 1.823311 | 2.079075 | 1.08369 | 2.062376 | 1.987976 | 1.659355 | | 24.5 | -0.036264 | 0.30106 |
| 24.4236111 | 0.382278 | 0.534291 | 0.607142 | 0.29941 | 0.241606 | 0.230644 | 0.790439 | 0.393559 | 0.571511 | 0.628092 | 0.560129 | 3.210737 | 2.780942 | 1.757865 | 1.839263 | 1.470487 | 2.190794 | 3.451861 | 2.166057 | 2.70921 | 1.325535 | 2.698483 | 2.843524 | 3.954732 | | 24.5138889 | -0.192919 | 0.095254 |
| 24.4375 | 0.924057 | 0.831912 | 0.717056 | -0.254123 | -0.018356 | -0.59738 | 1.288104 | 0.12374 | 1.705356 | 2.46793 | 1.525235 | 3.509205 | 2.293482 | 1.674509 | 1.966255 | 1.143077 | 2.317276 | 4.333956 | 1.672338 | 3.346938 | 1.223982 | 3.461994 | 5.112032 | 15.025667 | | 24.5277778 | -2.38257 | -0.192008 |
| 24.4513889 | 2.157659 | 1.489797 | 1.332108 | -1.157543 | -0.557456 | -1.862871 | 2.044905 | -0.225559 | 3.14264 | 5.701298 | 4.113967 | 2.799459 | 1.032515 | 1.56636 | 1.933423 | 0.496695 | 1.304978 | 2.475404 | 0.444059 | 3.779266 | 0.556494 | 4.763259 | 7.95918 | 32.865366 | | 24.5416667 | -7.387528 | -2.219029 |
| 24.4652778 | 3.060904 | 1.822429 | 2.386923 | -1.045126 | -0.226499 | -0.890035 | 2.359333 | 0.115614 | 3.742699 | 6.237397 | 3.065633 | 1.85705 | 0.635902 | 1.071366 | 1.30953 | -0.235299 | 0.479082 | 0.563056 | 0.033748 | 2.691794 | -0.541237 | 4.281467 | 7.125695 | 35.156042 | | 24.5555556 | -8.962083 | -4.510012 |
| 24.4791667 | 2.134416 | 1.402735 | 2.693991 | 0.087263 | 0.907594 | 1.456311 | 1.865563 | 1.045315 | 2.841159 | 4.157748 | 2.363076 | 1.999567 | 1.34959 | 1.723284 | 1.096043 | 0.085695 | 0.717871 | 0.0137 | 1.319962 | 1.820866 | 0.109807 | 1.996129 | 3.331999 | 15.458124 | | 24.5694444 | -4.584004 | -3.004462 |
| 24.4930556 | 1.39718 | 1.292212 | 2.176576 | 0.892332 | 1.373544 | 2.297925 | 1.114574 | 1.147695 | 1.877724 | 2.215075 | 2.238052 | 2.423557 | 2.01555 | 2.053611 | 1.308546 | 0.687351 | 1.333905 | 1.260551 | 2.869024 | 2.293768 | 0.928553 | 0.294672 | 0.389335 | 4.44058 | | 24.5833333 | -0.612997 | -0.833699 |
| 24.5069444 | 1.132628 | 1.345407 | 1.83803 | 0.860474 | 1.066208 | 1.72722 | 0.885127 | 0.937723 | 1.505646 | 1.62544 | 1.936763 | 2.551973 | 3.183855 | 2.19022 | 1.876094 | 1.037522 | 2.112234 | 3.798609 | 2.990388 | 3.013073 | 1.853237 | 0.95412 | 0.868008 | 3.638655 | | 24.5972222 | 0.036603 | -0.019323 |
| 24.5208333 | 0.918563 | 1.27484 | 2.135751 | 0.651623 | 0.686119 | 1.212828 | 0.662785 | 0.722468 | 1.54627 | 1.650111 | 2.150051 | 2.58454 | 4.594207 | 1.509371 | 1.343396 | 0.708308 | 2.441764 | 4.431083 | 1.884509 | 3.784365 | 1.889104 | 1.697339 | 3.984861 | 3.835252 | | 24.6111111 | -0.09427 | -0.173352 |
| 24.5347222 | 0.807373 | 1.458955 | 3.162167 | 0.505103 | 0.613723 | 1.186493 | 0.641459 | 0.691928 | 1.755997 | 1.933829 | 2.572569 | 2.403715 | 3.622285 | 0.88975 | 0.699845 | 0.590698 | 0.563193 | 1.726918 | 0.982045 | 2.387595 | 0.54607 | 1.41481 | 4.201929 | 2.437146 | | 24.625 | -0.142196 | -0.337074 |
| 24.5486111 | 0.921754 | 2.285661 | 3.716922 | 0.469892 | 0.721698 | 1.459587 | 0.717776 | 0.765469 | 1.656403 | 1.888572 | 2.596966 | 1.586077 | 2.59879 | 0.528172 | 0.351309 | 0.490644 | -0.201041 | -0.635577 | 0.930528 | 2.011296 | -0.165845 | 0.610736 | 1.767048 | 0.555117 | | 24.6388889 | -0.061946 | -0.279434 |
| 24.5625 | 0.950968 | 2.368144 | 2.739128 | 0.555448 | 0.742174 | 1.556531 | 0.677978 | 0.69987 | 1.28551 | 1.448024 | 2.301555 | 0.954376 | 0.868175 | 0.339011 | -0.030352 | 0.013277 | 0.193524 | -0.406625 | 0.962091 | 1.129938 | 0.805167 | -0.466711 | -0.740172 | -0.121989 | | 24.6527778 | -0.004357 | -0.107983 |
| 24.5763889 | 0.860051 | 2.082465 | 1.67219 | 0.576779 | 0.574028 | 1.154675 | 0.494463 | 0.551041 | 0.970284 | 0.994707 | 1.8006 | 0.898519 | 1.152113 | 0.53696 | 0.472315 | 0.213556 | 0.813502 | 0.75769 | 1.345848 | 1.105032 | 1.506916 | 1.095038 | -0.123468 | 0.183765 | | 24.6666667 | -0.005904 | 0.019337 |
| 24.5902778 | 0.883996 | 2.027456 | 1.171228 | 0.684783 | 0.409898 | 0.791913 | 0.305894 | 0.366798 | 0.80885 | 0.801841 | 1.337178 | 1.1369 | 2.021129 | 0.826841 | 0.662811 | 0.02642 | 0.643855 | 1.289581 | 1.278611 | 1.423224 | 1.404456 | 1.494582 | 0.170004 | 0.555382 | | 24.6805556 | -0.037092 | 0.109447 |
| 24.6041667 | 0.998855 | 2.015936 | 0.983604 | 0.943122 | 0.350695 | 0.640883 | 0.226112 | 0.301482 | 0.757898 | 0.7214 | 1.162339 | 0.373227 | 1.01357 | 1.238443 | 1.356544 | 0.05699 | -0.351918 | 0.34209 | 0.204433 | 0.890975 | 0.270591 | 0.47134 | 0.53177 | 0.649867 | | 24.6944444 | -0.082869 | 0.261481 |
| 24.6180556 | 1.055649 | 1.880898 | 0.966218 | 1.155194 | 0.375546 | 0.643235 | 0.229258 | 0.308856 | 0.686537 | 0.626003 | 1.143805 | 0.068864 | 0.158327 | 0.787571 | 0.533838 | 0.04381 | -0.039089 | 0.582035 | -0.356819 | 0.334412 | -0.209538 | -0.005899 | 0.269926 | 0.582642 | | 24.7083333 | -0.094325 | 0.39366 |
| 24.6319444 | 0.895067 | 1.628199 | 1.004142 | 1.173104 | 0.417222 | 0.692314 | 0.279304 | 0.348012 | 0.625331 | 0.544362 | 1.06461 | -0.022619 | -1.308046 | 0.358736 | -0.154152 | -0.241152 | 0.453611 | 0.314658 | 0.147924 | -0.277006 | 0.059706 | 0.162405 | -0.552678 | 0.189577 | | 24.7222222 | -0.026119 | 0.296246 |
| 24.6458333 | 0.688503 | 1.465968 | 1.159262 | 1.12691 | 0.427771 | 0.712855 | 0.316483 | 0.397304 | 0.62777 | 0.573187 | 0.976509 | 0.57641 | 0.450105 | -0.162075 | -0.546945 | -0.035445 | 0.831458 | -0.153833 | 0.331788 | -0.816387 | 0.582454 | 0.57868 | -0.135352 | -0.367029 | | 24.7361111 | 0.07766 | -0.292481 |
| 24.6597222 | 0.476205 | 1.472606 | 1.34899 | 0.956959 | 0.400233 | 0.685966 | 0.295736 | 0.395099 | 0.648333 | 0.661934 | 0.898425 | 0.106896 | -0.505803 | -0.346318 | -0.106457 | 0.197088 | 0.184787 | -0.453534 | -0.094044 | -0.075748 | 0.657105 | 0.487784 | -0.130829 | -0.491149 | | 24.75 | 0.171061 | -1.170434 |
| 24.6736111 | 0.378992 | 1.534042 | 1.376765 | 0.691407 | 0.362443 | 0.677824 | 0.243075 | 0.344169 | 0.697354 | 0.742842 | 0.798129 | -0.418099 | -0.287515 | 0.160336 | 0.593634 | 0.386442 | -0.309678 | -0.239029 | -0.055175 | 0.58233 | 0.470944 | 0.298452 | -0.836653 | 0.045694 | | 24.7638889 | 0.21093 | -0.882943 |
| 24.6875 | 0.35994 | 1.247364 | 1.038186 | 0.449534 | 0.350435 | 0.700749 | 0.168826 | 0.264046 | 0.644056 | 0.700114 | 0.734965 | -0.205869 | 0.025313 | 0.43347 | 0.575455 | 0.683828 | 0.206147 | 0.696582 | 0.196282 | 1.534359 | 0.899367 | -0.404134 | 0.444751 | 0.698082 | | 24.7777778 | 0.168242 | -0.296684 |
| 24.7013889 | 0.356942 | 0.940645 | 0.609962 | 0.34459 | 0.36332 | 0.683911 | 0.096461 | 0.225271 | 0.401216 | 0.462108 | 0.604503 | 0.607 | 0.985015 | 0.775058 | 0.300129 | 0.216395 | 0.140945 | -0.119946 | -0.327009 | -0.08845 | 0.428211 | -1.779792 | 0.466453 | 0.183973 | | 24.7916667 | 0.08055 | 0.175661 |
| 24.7152778 | 0.349663 | 0.581485 | 0.169109 | 0.307464 | 0.363533 | 0.632665 | 0.128049 | 0.246774 | 0.040727 | 0.104617 | 0.37097 | 1.184078 | 1.259198 | 0.431903 | -0.199718 | -0.464672 | -0.048538 | -0.104752 | -0.828498 | -2.28454 | -0.473041 | -0.995123 | -0.385178 | -0.589487 | | 24.8055556 | -0.031344 | 0.144745 |
| 24.7291667 | 0.35688 | 0.252452 | -0.075875 | 0.388453 | 0.322165 | 0.536502 | 0.235712 | 0.280816 | -0.174565 | -0.10586 | 0.123224 | 0.021726 | -0.853631 | -0.104532 | 0.441308 | 0.742529 | -0.464565 | -1.003987 | -0.020786 | -2.405399 | -1.222125 | -0.149953 | -1.390973 | -0.962743 | | 24.8194444 | -0.132243 | 0.124582 |
| 24.7430556 | 0.365681 | 0.132764 | -0.021561 | 0.51494 | 0.2325 | 0.406877 | 0.370606 | 0.265722 | -0.018398 | 0.030417 | 0.030517 | -0.849451 | -1.132162 | 0.006364 | 0.396556 | 0.144109 | 0.040816 | -0.247758 | 1.307015 | -0.125116 | -1.532536 | 1.473039 | -0.229766 | -0.396917 | | 24.8333333 | -0.136726 | 0.172271 |
| 24.7569444 | 0.411474 | 0.344248 | 0.265349 | 0.616153 | 0.143235 | 0.255983 | 0.492527 | 0.259832 | 0.281115 | 0.179538 | 0.162992 | -0.631238 | -0.198379 | 0.309865 | 0.176738 | -0.285445 | 0.753963 | 1.082272 | 0.544673 | 1.652576 | -0.457858 | -0.769272 | 0.341652 | 0.210464 | | 24.8472222 | -0.055533 | 0.169133 |
| 24.7708333 | 0.443705 | 0.734874 | 0.663554 | 0.665346 | 0.115919 | 0.201097 | 0.388585 | 0.257355 | 0.350655 | 0.333782 | 0.409403 | 0.64745 | 1.833412 | 0.412544 | -0.21295 | -0.542206 | 0.226763 | 0.333373 | -1.528343 | -0.408196 | 0.592991 | -2.329918 | -0.840905 | 0.037281 | | 24.8611111 | -0.002297 | 0.0939 |
| 24.7847222 | 0.397129 | 0.998539 | 0.764539 | 0.599323 | 0.140699 | 0.282185 | 0.075093 | 0.157884 | 0.060255 | 0.024441 | 0.538774 | 0.726387 | -0.274962 | -0.209088 | -0.702848 | -0.822463 | -0.32306 | -0.254389 | -1.443407 | -1.680247 | 0.573838 | -0.880257 | 0.475516 | -0.681582 | | 24.875 | 0.00843 | 0.026509 |
| 24.7986111 | 0.277974 | 0.696552 | 0.388315 | 0.373333 | 0.142885 | 0.312329 | -0.326099 | 0.020816 | -0.198993 | -0.204026 | 0.458838 | -0.608427 | -2.025268 | -0.213356 | -0.337589 | -0.061553 | -0.379069 | -0.64658 | 0.389416 | -1.151076 | -0.19165 | 2.55276 | -1.220366 | -0.817825 | | 24.8888889 | 0.035795 | -0.010434 |
| 24.8125 | 0.139798 | 0.003463 | -0.220955 | 0.158106 | 0.049015 | 0.188361 | -0.380077 | -0.235144 | -0.060062 | 0.06103 | 0.169763 | -1.482914 | -1.295105 | 0.022361 | 0.086748 | -0.92052 | 0.696663 | 0.200113 | 1.319849 | 0.49783 | -1.668715 | 0.82705 | 0.416095 | -1.039825 | | 24.9027778 | 0.058528 | -0.057709 |
| 24.8263889 | -0.021108 | -0.738406 | -0.673054 | 0.024907 | -0.021286 | -0.017613 | -0.026376 | -0.083756 | 0.169658 | 0.433889 | -0.223142 | -0.463779 | 1.087228 | -0.138872 | -0.271311 | -0.557394 | 0.748767 | 0.160498 | 0.250918 | 0.810823 | -1.491765 | -0.486444 | 0.283828 | -0.635475 | | 24.9166667 | -0.017065 | -0.170037 |
| 24.8402778 | -0.183533 | -1.078948 | -0.709858 | -0.056013 | -0.103025 | -0.321559 | 0.243294 | 0.097039 | 0.035454 | 0.396451 | -0.494347 | -0.151389 | -0.311649 | 0.032875 | -0.391613 | -0.059674 | -0.18935 | -0.72988 | -1.189275 | -0.935606 | -0.34247 | 1.254786 | 0.047683 | -0.52147 | | 24.9305556 | -0.148693 | -0.345807 |
| 24.8541667 | -0.28172 | -1.074546 | -0.555375 | -0.097601 | -0.269838 | -0.665635 | 0.135104 | -0.025215 | -0.50629 | -0.301824 | -0.753857 | -1.267705 | -1.886924 | -1.029972 | -1.225477 | 0.127907 | -0.872816 | -0.310087 | -1.97393 | -2.042995 | 0.006137 | 1.566138 | 0.031722 | -1.66676 | | 24.9444444 | -0.129536 | -0.482752 |
| 24.8680556 | -0.312069 | -1.146243 | -0.828174 | -0.375571 | -0.453524 | -0.954821 | -0.426097 | -0.294979 | -0.812571 | -0.808912 | -1.163404 | -1.944459 | -2.013155 | -0.798756 | -0.743286 | -0.546312 | -1.190427 | -0.070433 | -1.145892 | -1.598337 | -0.519643 | -0.900438 | -1.996162 | -2.048946 | | 24.9583333 | 0.000974 | -0.539416 |
| 24.8819444 | -0.345372 | -1.463416 | -1.582445 | -0.850393 | -0.654147 | -1.115295 | -0.819568 | -0.685867 | -0.673859 | -0.909051 | -1.552555 | -1.433362 | -0.238727 | -0.437875 | -0.203298 | -0.66282 | -0.538654 | -1.550191 | 0.043237 | -0.1759 | -1.444738 | -2.536721 | -0.788604 | -1.527048 | | 24.9722222 | -0.079152 | -0.519994 |
| 24.8958333 | -0.51695 | -1.943474 | -2.080237 | -1.091 | -0.866919 | -1.365445 | -1.120628 | -1.095608 | -0.728555 | -0.887838 | -1.777054 | -1.545777 | -0.293441 | -0.072303 | -0.297614 | -0.784923 | -0.864686 | -3.552294 | -1.182874 | -0.423061 | -1.515386 | -1.327309 | 0.611663 | -1.778842 | | 24.9861111 | -0.191598 | -0.430453 |
| 24.9097222 | -0.840783 | -2.181995 | -2.065339 | -0.936762 | -1.020517 | -1.696656 | -0.91128 | -0.928735 | -0.69713 | -0.816675 | -1.620195 | -2.661055 | -1.77145 | -0.033921 | -0.551831 | -0.630057 | -1.430374 | -3.773732 | -2.813647 | -1.413495 | -1.201258 | -0.070397 | -1.553614 | -2.039237 | | 25 | -0.366558 | -0.296663 |
| 24.9236111 | -1.2238 | -2.184817 | -1.817942 | -0.950697 | -1.043387 | -1.787881 | -0.457289 | -0.603573 | -0.997538 | -1.199938 | -1.838184 | -2.168226 | -2.520476 | -2.129728 | -1.822794 | -1.338189 | -1.997759 | -2.188381 | -3.515701 | -1.812667 | -1.701334 | 0.237365 | -2.69259 | -4.055056 | | 25.0138889 | 0.201674 | 0.208762 |
| 24.9375 | -1.995393 | -2.071827 | -1.588265 | -0.733494 | -0.804952 | -1.015461 | -0.394364 | -0.156925 | -2.370871 | -3.428105 | -4.018093 | -1.628279 | -1.011155 | -4.351364 | -2.859717 | -1.355864 | -1.320676 | -0.376404 | -2.690485 | -2.147895 | -2.087118 | -3.341532 | -3.030647 | -13.980471 | | 25.0277778 | 3.685899 | 1.968199 |
| 24.9513889 | -2.894533 | -1.788445 | -0.876077 | 0.232201 | -0.329229 | 0.870403 | -0.603893 | 0.96225 | -3.727186 | -5.924635 | -5.8807 | -2.109289 | 0.525629 | -1.782215 | -2.554329 | -1.260778 | -0.79341 | -0.084331 | -0.810926 | -2.053836 | -1.216759 | -8.331344 | -2.751049 | -30.954139 | | 25.0416667 | 7.52081 | 3.719095 |
| 24.9652778 | -1.932811 | -1.204957 | -0.609907 | 0.794017 | -0.075419 | 1.799036 | -0.26711 | 1.347271 | -3.190449 | -4.84702 | -6.926632 | -3.043619 | 0.154622 | -0.230282 | -2.054274 | -0.679627 | -1.052877 | -1.118403 | -0.409086 | -2.050899 | -1.221981 | -4.798022 | -2.910779 | -30.816913 | | 25.0555556 | 5.613678 | 2.683657 |
| 24.9791667 | -0.867791 | -0.888105 | -0.728228 | 0.253697 | -0.224622 | 0.616938 | -0.254584 | 0.433901 | -1.687551 | -2.194353 | -3.33585 | -3.266409 | -0.933756 | -0.081031 | -1.329551 | -0.740323 | -1.38569 | -2.584251 | -0.888886 | -1.470226 | -0.893154 | -1.246571 | -1.892913 | -12.855654 | | 25.0694444 | 1.805577 | 0.889118 |
| 24.9930556 | -0.523011 | -0.938096 | -0.662217 | -0.245643 | -0.405258 | -0.474927 | -0.531421 | -0.479932 | -0.725649 | -0.842227 | -1.147939 | -2.192519 | -1.142192 | -0.449352 | -0.901899 | -0.610933 | -1.497396 | -3.286869 | -0.918912 | -0.89594 | -0.428182 | -0.046764 | -1.268227 | -1.952232 | | 25.0833333 | -0.29655 | 0.047814 |
| 25.0069444 | -0.352701 | -0.731188 | -0.555171 | -0.211101 | -0.270399 | -0.637345 | -0.691107 | -0.732078 | -0.41362 | -0.412795 | -0.600652 | -0.791582 | -0.698007 | -0.685722 | -0.678991 | -0.224721 | -1.480086 | -3.243197 | -0.724588 | -0.875356 | -0.234383 | -0.787791 | -1.130727 | -0.509136 | | 25.0972222 | -0.440813 | -0.226257 |
| 25.0208333 | -0.274903 | -0.653941 | -0.45892 | -0.074811 | -0.126901 | -0.519892 | -0.671589 | -0.685867 | -0.244085 | -0.228767 | -0.338338 | -1.27737 | -1.508569 | -0.918254 | -1.128378 | -0.444407 | -1.851783 | -3.490479 | -0.913445 | -1.437837 | -1.06864 | -2.028666 | -1.50329 | -1.209604 | | 25.1111111 | -0.207832 | -0.287056 |
| 25.0347222 | -0.373375 | -0.881734 | -0.519839 | -0.113394 | -0.200504 | -0.456862 | -0.638613 | -0.582944 | -0.340611 | -0.33809 | -0.416618 | -2.597394 | -2.435322 | -1.347264 | -1.734447 | -1.18989 | -2.270634 | -3.40006 | -1.504463 | -2.082562 | -2.11924 | -2.550045 | -1.730933 | -1.953571 | | 25.125 | -0.007948 | -0.304634 |
| 25.0486111 | -0.497131 | -1.057172 | -0.645888 | -0.27188 | -0.344916 | -0.515934 | -0.637776 | -0.549588 | -0.50095 | -0.466113 | -0.619702 | -3.158777 | -2.433766 | -1.764033 | -1.738971 | -1.433818 | -1.795142 | -2.355441 | -1.725651 | -1.941754 | -2.085294 | -1.988379 | -1.361989 | -1.82895 | | 25.1388889 | 0.244837 | -0.28625 |
| 25.0625 | -0.557603 | -1.049203 | -0.666883 | -0.384314 | -0.424086 | -0.56563 | -0.638621 | -0.575139 | -0.59764 | -0.531309 | -0.769072 | -2.177818 | -1.905234 | -1.869733 | -1.271249 | -1.081566 | -1.173698 | -1.326003 | -1.529616 | -1.437867 | -1.254724 | -1.452949 | -0.98155 | -1.12242 | | 25.1527778 | 0.479272 | -0.216965 |
| 25.0763889 | -0.58477 | -0.959252 | -0.565489 | -0.361358 | -0.396326 | -0.517596 | -0.598168 | -0.564665 | -0.698996 | -0.634337 | -0.931739 | -1.133063 | -1.29393 | -1.601548 | -0.965011 | -0.810747 | -0.928007 | -0.831002 | -1.267951 | -1.157257 | -0.753888 | -1.319471 | -0.841136 | -0.877218 | | 25.1666667 | 0.502475 | -0.179017 |
| 25.0902778 | -0.605721 | -0.9458 | -0.535473 | -0.313732 | -0.335174 | -0.473481 | -0.496703 | -0.480821 | -0.783187 | -0.703741 | -1.021141 | -0.660983 | -0.666842 | -1.149621 | -0.904518 | -0.798083 | -0.965201 | -0.754111 | -0.91399 | -1.045601 | -0.611762 | -1.314215 | -0.813512 | -1.061384 | | 25.1805556 | 0.368407 | -0.165729 |
| 25.1041667 | -0.605613 | -1.00336 | -0.6507 | -0.38671 | -0.279114 | -0.490152 | -0.399441 | -0.406726 | -0.773739 | -0.705509 | -1.029561 | -0.91481 | -0.254227 | -0.992711 | -0.925883 | -0.873697 | -1.055333 | -0.908674 | -0.646644 | -0.874455 | -0.576455 | -1.283789 | -0.754863 | -1.126351 | | 25.1944444 | 0.301035 | -0.185465 |
| 25.1180556 | -0.669264 | -1.103549 | -0.811091 | -0.552036 | -0.257553 | -0.526186 | -0.376511 | -0.366642 | -0.693779 | -0.659966 | -1.017196 | -1.348401 | -0.567427 | -0.951954 | -0.811025 | -0.756326 | -1.025406 | -1.010537 | -0.633829 | -0.677995 | -0.499531 | -1.171335 | -0.663035 | -0.938028 | | 25.2083333 | 0.3539 | -0.165822 |
| 25.1319444 | -0.78014 | -1.169139 | -0.885156 | -0.674235 | -0.284078 | -0.539373 | -0.426333 | -0.360224 | -0.720933 | -0.722426 | -1.086107 | -1.235738 | -0.989946 | -0.77626 | -0.508748 | -0.47788 | -0.909195 | -0.937491 | -0.713637 | -0.541371 | -0.392732 | -0.944063 | -0.584374 | -0.503563 | | 25.2222222 | 0.296565 | 0.001508 |
| 25.1458333 | -0.857373 | -1.184367 | -0.846439 | -0.625398 | -0.335963 | -0.600267 | -0.464269 | -0.395965 | -0.732973 | -0.704527 | -0.920796 | -0.808024 | -0.978391 | -0.539754 | -0.374183 | -0.397134 | -0.816601 | -0.827495 | -0.694403 | -0.493223 | -0.327906 | -0.725618 | -0.569659 | -0.229333 | | 25.2361111 | 0.092792 | 0.202809 |
| 25.1597222 | -0.80035 | -1.06924 | -0.670134 | -0.523536 | -0.351712 | -0.625632 | -0.433888 | -0.407657 | -0.608532 | -0.557573 | -0.585935 | -0.470445 | -0.526723 | -0.488934 | -0.553563 | -0.538015 | -0.769071 | -0.689957 | -0.630582 | -0.485803 | -0.317493 | -0.692402 | -0.606018 | -0.3755 | | 25.25 | 0.141287 | 0.244822 |
| 25.1736111 | -0.623063 | -0.787143 | -0.515322 | -0.469476 | -0.297758 | -0.517398 | -0.33584 | -0.340234 | -0.432945 | -0.411457 | -0.34026 | -0.45859 | -0.419479 | -0.701864 | -0.709742 | -0.54428 | -0.744873 | -0.589002 | -0.641707 | -0.488862 | -0.361608 | -0.796455 | -0.638742 | -0.670016 | | 25.2638889 | 0.441002 | 0.201503 |
| 25.1875 | -0.505933 | -0.521518 | -0.447831 | -0.450603 | -0.231036 | -0.369152 | -0.247331 | -0.245907 | -0.318459 | -0.344292 | -0.338015 | -0.589224 | -0.644272 | -0.85282 | -0.588595 | -0.364804 | -0.702091 | -0.559047 | -0.689963 | -0.492993 | -0.40666 | -0.859607 | -0.669422 | -0.859763 | | 25.2777778 | 0.562991 | 0.1715 |
| 25.2013889 | -0.451069 | -0.355079 | -0.374132 | -0.349796 | -0.212044 | -0.315281 | -0.203667 | -0.197917 | -0.210345 | -0.277702 | -0.285058 | -0.508324 | -0.562236 | -0.710732 | -0.419775 | -0.287612 | -0.623386 | -0.542737 | -0.615631 | -0.467028 | -0.384165 | -0.745272 | -0.658733 | -0.781795 | | 25.2916667 | 0.365686 | 0.07484 |
| 25.2152778 | -0.407909 | -0.28641 | -0.220103 | -0.205983 | -0.232929 | -0.313714 | -0.175148 | -0.199033 | -0.14798 | -0.243819 | -0.128026 | -0.247355 | -0.2285 | -0.506821 | -0.405214 | -0.32588 | -0.477596 | -0.439531 | -0.400702 | -0.429532 | -0.309394 | -0.433637 | -0.509063 | -0.520804 | | 25.3055556 | 0.166527 | -0.058802 |
| 25.2291667 | -0.353707 | -0.218057 | -0.066521 | -0.19631 | -0.227936 | -0.272324 | -0.090307 | -0.187112 | -0.172801 | -0.291572 | -0.021595 | -0.121199 | -0.159579 | -0.389781 | -0.357226 | -0.253681 | -0.250022 | -0.206922 | -0.209721 | -0.382285 | -0.265067 | -0.160301 | -0.314053 | -0.352791 | | 25.3194444 | 0.064591 | -0.02789 |
| 25.2430556 | -0.218513 | -0.000495 | 0.0448 | -0.217372 | -0.136072 | -0.150878 | 0.048363 | -0.0893 | -0.129023 | -0.281187 | 0.067795 | -0.115505 | -0.15257 | -0.250621 | -0.08541 | -0.020271 | -0.047824 | 0.061585 | -0.083517 | -0.260448 | -0.192183 | -0.04056 | -0.19346 | -0.241688 | | 25.3333333 | 0.04654 | 0.174195 |
| 25.2569444 | -0.065418 | 0.252631 | 0.117136 | -0.071994 | -0.024203 | -0.001101 | 0.121259 | 0.027981 | -0.018543 | -0.190856 | 0.133113 | -0.097749 | 0.096038 | -0.082959 | 0.124995 | 0.079986 | -0.008825 | 0.110395 | 0.016743 | -0.122723 | -0.097131 | 0.053607 | -0.132115 | -0.082549 | | 25.3472222 | 0.102602 | 0.325042 |
| 25.2708333 | -0.015137 | 0.293076 | 0.11804 | 0.070645 | 0.00717 | 0.085951 | 0.079952 | 0.042973 | 0.073607 | -0.068519 | 0.191375 | -0.194621 | 0.10057 | -0.022751 | -0.02717 | -0.122898 | -0.126587 | -0.105335 | -0.025778 | -0.09805 | -0.124401 | 0.131746 | -0.117999 | 0.001296 | | 25.3611111 | 0.169456 | 0.243882 |
| 25.2847222 | 0.10719 | 0.283027 | 0.177575 | 0.093994 | 0.031254 | 0.140616 | 0.09504 | 0.049266 | 0.194261 | 0.082933 | 0.431797 | -0.262412 | -0.154342 | -0.085711 | -0.172079 | -0.223979 | -0.146421 | -0.180201 | -0.138423 | -0.122252 | -0.179599 | 0.178181 | -0.091303 | 0.025429 | | 25.375 | 0.162142 | 0.158649 |
| 25.2986111 | 0.407001 | 0.489969 | 0.380996 | 0.102627 | 0.128657 | 0.256647 | 0.28892 | 0.216209 | 0.299738 | 0.177189 | 0.755295 | -0.028586 | -0.138268 | -0.152166 | -0.011591 | -0.033223 | 0.016426 | 0.046677 | -0.139652 | -0.093296 | -0.159271 | 0.207017 | -0.056086 | 0.054703 | | 25.3888889 | 0.163513 | 0.25332 |
| 25.3125 | 0.604359 | 0.752757 | 0.513024 | 0.19817 | 0.230054 | 0.38379 | 0.46906 | 0.393384 | 0.351229 | 0.18426 | 0.898297 | 0.289095 | 0.120678 | -0.095463 | 0.121775 | 0.027183 | 0.122968 | 0.270397 | -0.088131 | -0.121696 | -0.105892 | 0.238569 | -0.088972 | -0.067317 | | 25.4027778 | 0.172465 | 0.285257 |
| 25.3263889 | 0.613667 | 0.880507 | 0.58534 | 0.432621 | 0.322445 | 0.516258 | 0.541996 | 0.435089 | 0.44986 | 0.256666 | 0.924788 | 0.395347 | 0.279026 | 0.188112 | 0.117985 | -0.083055 | 0.213407 | 0.439418 | -0.011797 | -0.071391 | -0.039321 | 0.324449 | -0.080211 | -0.212848 | | 25.4166667 | 0.24332 | 0.224473 |
| 25.3402778 | 0.647935 | 0.950564 | 0.769276 | 0.681419 | 0.507339 | 0.75068 | 0.621395 | 0.550661 | 0.674733 | 0.541608 | 1.116553 | 0.357885 | 0.392342 | 0.43411 | 0.261472 | 0.0162 | 0.389894 | 0.677768 | 0.044215 | 0.135042 | 0.084636 | 0.375598 | 0.02904 | -0.11623 | | 25.4305556 | 0.314835 | 0.087635 |
| 25.3541667 | 0.951067 | 1.244304 | 1.263054 | 0.794279 | 0.650508 | 0.965784 | 0.823141 | 0.845771 | 0.961999 | 1.0015 | 1.637152 | 0.486288 | 0.721662 | 0.428574 | 0.534827 | 0.329971 | 0.537204 | 0.99049 | 0.11848 | 0.317923 | 0.217034 | 0.407616 | 0.150864 | 0.113538 | | 25.4444444 | 0.300437 | -0.210775 |
| 25.3680556 | 1.351762 | 1.806386 | 1.814668 | 0.964347 | 0.693605 | 1.144768 | 1.083213 | 1.233841 | 1.196524 | 1.298317 | 2.21187 | 0.960065 | 1.236965 | 0.551926 | 0.821239 | 0.555557 | 0.690756 | 1.329018 | 0.388318 | 0.507799 | 0.410299 | 0.613195 | 0.33663 | 0.309321 | | 25.4583333 | 0.049566 | -0.489404 |
| 25.3819444 | 1.506882 | 2.344171 | 2.134488 | 1.43067 | 0.815807 | 1.476897 | 1.216799 | 1.450909 | 1.465663 | 1.486575 | 2.453432 | 1.479186 | 1.48408 | 0.982849 | 1.057396 | 0.615684 | 0.915943 | 1.662896 | 0.761186 | 0.80624 | 0.653319 | 1.004672 | 0.70407 | 0.741845 | | 25.4722222 | -0.115686 | -0.514877 |
| 25.3958333 | 1.600076 | 2.611454 | 2.179844 | 1.994182 | 1.142245 | 2.134251 | 1.290593 | 1.419018 | 1.825869 | 1.82722 | 2.687461 | 1.871211 | 1.606778 | 1.278409 | 1.312398 | 0.834062 | 1.131211 | 1.849207 | 1.055239 | 1.34467 | 0.93672 | 1.480731 | 1.276073 | 1.35489 | | 25.4861111 | -0.087878 | -0.384146 |
| 25.4097222 | 1.559753 | 2.534417 | 2.192152 | 1.696337 | 1.418176 | 2.581142 | 1.411262 | 1.491258 | 1.939136 | 2.219881 | 3.027331 | 2.507356 | 2.180081 | 1.40713 | 1.65939 | 1.395245 | 1.40396 | 2.093663 | 1.429265 | 2.074193 | 1.398713 | 1.972954 | 1.926917 | 1.775099 | | 25.5 | 0.060508 | -0.238222 |
| 25.4236111 | 1.642057 | 3.155536 | 2.557337 | 1.506116 | 1.405891 | 2.549826 | 1.593309 | 1.683803 | 1.890996 | 2.647377 | 3.514581 | 3.4522 | 3.065941 | 1.74678 | 2.147013 | 1.847849 | 1.828455 | 2.759913 | 2.074163 | 2.892589 | 2.066787 | 2.52544 | 2.816534 | 3.562976 | | 25.5138889 | -0.186047 | -0.260384 |
| 25.4375 | 2.064381 | 4.217222 | 2.449104 | 0.893399 | 1.365712 | 1.728447 | 2.036993 | 1.210313 | 2.614059 | 3.779129 | 5.926646 | 4.052346 | 2.722207 | 2.19045 | 2.450407 | 1.037021 | 2.249985 | 3.62042 | 2.56447 | 3.333245 | 1.961183 | 3.508822 | 4.389208 | 14.222439 | | 25.5277778 | -2.5804 | -0.776549 |
| 25.4513889 | 2.730602 | 3.586532 | 1.784149 | -0.418994 | 1.492187 | 0.514085 | 1.683243 | -0.772539 | 4.268867 | 6.11734 | 9.674419 | 4.026549 | 1.049191 | 2.893617 | 2.644474 | 0.008121 | 2.076743 | 3.181469 | 2.207505 | 3.314089 | 0.975663 | 5.076759 | 6.02012 | 30.983541 | | 25.5416667 | -8.198129 | -3.211197 |
| 25.4652778 | 3.165924 | 1.99175 | -0.095487 | -0.927921 | 1.235048 | 0.092629 | 0.757859 | -1.67485 | 4.025271 | 6.455336 | 10.06282 | 3.538944 | 0.115806 | 3.962855 | 2.258942 | 0.043809 | 1.416795 | 1.924017 | 2.185524 | 3.43319 | 0.406639 | 5.549461 | 6.258239 | 29.702242 | | 25.5555556 | -9.602041 | -4.910626 |
| 25.4791667 | 1.995693 | 0.677201 | -0.827027 | -0.892315 | 1.121619 | 0.612468 | 0.144547 | -1.26571 | 1.691231 | 3.396382 | 3.093973 | 2.949421 | 1.207847 | 2.812203 | 1.822302 | 0.766505 | 1.35151 | 1.123627 | 2.748256 | 3.617514 | 0.535755 | 5.060119 | 4.920685 | 13.414623 | | 25.5694444 | -4.420104 | -4.515995 |
| 25.4930556 | 1.765721 | 1.614467 | -0.672055 | -0.664007 | 1.277278 | 1.307897 | -0.05431 | -0.484864 | 1.335416 | 1.722439 | -0.199149 | 3.042393 | 2.34731 | 2.154578 | 1.662036 | 1.246826 | 1.663876 | 1.040169 | 3.277984 | 3.228224 | 2.122938 | 4.428805 | 2.683641 | 4.734138 | | 25.5833333 | -0.893811 | -0.893515 |
| 25.5069444 | 1.988792 | 2.474753 | 1.187698 | -0.124584 | 1.321879 | 1.780583 | 0.191753 | 0.47581 | 1.941386 | 1.96203 | 0.76103 | 2.254133 | 2.714027 | 2.210557 | 2.048656 | 1.524382 | 1.818187 | 1.409244 | 3.152417 | 2.914256 | 3.440358 | 4.126264 | 2.062903 | 4.205662 | | 25.5972222 | -0.392947 | 0.246918 |
| 25.5208333 | 1.357941 | 2.46144 | 2.327516 | 1.310058 | 1.36058 | 1.847663 | 0.145134 | 1.073285 | 2.059646 | 1.915697 | 3.156043 | 2.088877 | 3.736095 | 2.373061 | 2.17288 | 1.265175 | 2.120837 | 3.101186 | 2.649275 | 3.003337 | 3.697407 | 3.659417 | 2.105286 | 4.399354 | | 25.6111111 | -0.252431 | 0.37364 |
| 25.5347222 | 0.619053 | 2.123532 | 3.115186 | 3.04284 | 1.133794 | 2.188211 | 0.509515 | 1.329578 | 1.163102 | 1.534136 | 4.122353 | 3.013017 | 5.43219 | 2.770526 | 2.359015 | 1.088886 | 2.865842 | 4.025444 | 1.097014 | 2.157823 | 2.182296 | 1.994329 | 2.549569 | 4.014042 | | 25.625 | -0.060564 | 0.301962 |
| 25.5486111 | -0.294726 | -0.198311 | 1.821222 | 2.368942 | 0.406491 | 1.20154 | 0.299176 | 0.563535 | 0.42969 | 1.219703 | 2.892464 | 3.041461 | 5.762658 | 1.436783 | 0.718094 | 0.887014 | 1.59381 | 2.249522 | 0.624985 | 2.005151 | -0.139564 | 0.440829 | 1.604608 | 2.282919 | | 25.6388889 | 0.114614 | 0.166434 |
| 25.5625 | -0.30892 | -1.053088 | 0.79307 | 0.981033 | -0.196398 | -0.75949 | 0.157096 | -0.242917 | 0.505203 | 1.290466 | 1.512312 | 1.581629 | 2.22415 | 0.561131 | -0.032409 | 0.429458 | -0.167175 | -0.510261 | 1.079008 | 1.902309 | -0.210546 | -0.309371 | -0.654059 | 0.051716 | | 25.6527778 | 0.215562 | 0.079733 |
| 25.5763889 | 0.626937 | 0.90774 | -0.270285 | -0.919202 | -0.083755 | -0.356853 | 0.305315 | -0.569153 | 0.623693 | 1.17614 | 0.35986 | 0.411175 | -0.558655 | -0.008022 | -0.106746 | 0.163565 | -0.512059 | -0.632345 | 0.863516 | 0.108235 | 0.450196 | 0.089808 | -2.22272 | -0.810779 | | 25.6666667 | 0.054635 | -0.002878 |
| 25.5902778 | 0.951916 | 2.026362 | -1.111583 | -1.408146 | 0.106495 | 0.103844 | 0.605729 | -0.325221 | 0.241176 | 0.522286 | -1.760845 | 0.433489 | 0.62912 | 0.521336 | 0.725625 | 0.189108 | 0.123918 | 0.561204 | 0.709381 | 0.810798 | 1.87778 | 0.931175 | -2.06579 | 0.579526 | | 25.6805556 | -0.232237 | -0.111245 |
| 25.6041667 | 0.578479 | 1.464639 | -0.845641 | -0.765644 | -0.001089 | 0.001412 | 0.087173 | 0.004494 | -0.527427 | -0.60314 | -1.837264 | 0.524606 | 1.464462 | 0.944578 | 1.188195 | 0.321164 | 1.411788 | 2.551384 | 0.056203 | 0.856956 | 1.835047 | 0.234299 | 0.268088 | 2.003907 | | 25.6944444 | -0.367569 | -0.018657 |
| 25.6180556 | -0.079317 | -0.169998 | -0.562526 | -0.23707 | 0.02849 | 0.290818 | -0.469331 | 0.390514 | -0.510069 | -0.716488 | -0.825722 | 0.34283 | 1.235371 | 0.777129 | 0.750579 | -0.10918 | 1.266464 | 1.457213 | 0.495152 | 0.590543 | -0.433486 | -0.411233 | 1.603776 | 1.347613 | | 25.7083333 | -0.27912 | -0.182576 |
| 25.6319444 | -0.204176 | -1.238637 | -0.165752 | 0.757364 | 0.352936 | 0.799093 | -0.337305 | 0.239271 | -0.035661 | -0.586625 | 1.462273 | 0.202095 | 0.400135 | 0.267772 | -0.042438 | -0.138005 | -0.75391 | -1.546247 | 0.863986 | 0.366882 | -1.576497 | -0.531335 | -0.699546 | -0.506474 | | 25.7222222 | -0.174211 | -0.089773 |
| 25.6458333 | -0.375943 | -1.352991 | -0.037047 | 0.155747 | 0.221125 | 0.095431 | 0.463144 | 0.118414 | 0.202885 | -0.535826 | 1.099873 | -0.155313 | -1.066705 | -0.057866 | -0.29125 | -0.104117 | -1.501756 | -2.388763 | -0.132462 | -1.319706 | -0.541395 | -0.361537 | -2.731646 | -2.195791 | | 25.7361111 | -0.02079 | 0.03132 |
| 25.6597222 | 0.669453 | 1.093319 | -0.478412 | -0.793955 | -0.252363 | -0.571829 | 0.729717 | -0.05647 | 0.564918 | -0.258535 | -1.412949 | 0.334804 | -0.130712 | -0.187128 | -0.111152 | 0.05887 | -0.377391 | -0.678489 | -0.686775 | -1.727274 | 0.459429 | 0.009897 | -2.615884 | -1.198285 | | 25.75 | 0.16019 | 0.171068 |
| 25.6736111 | 0.925411 | 0.899317 | -0.754793 | -0.880596 | -0.312921 | 0.031609 | 0.436997 | -0.033069 | 0.044718 | 0.054508 | -3.052237 | 0.77273 | 1.649697 | 0.321957 | 0.568143 | 0.104441 | 0.757871 | 0.336105 | -0.629772 | -0.67737 | 1.13369 | 1.244594 | -1.123942 | -0.235 | | 25.7638889 | 0.400751 | 0.293008 |
| 25.6875 | 0.782185 | 0.177966 | -1.275532 | -0.644476 | -0.169697 | 0.151022 | -0.443191 | -0.192135 | -0.440768 | 0.414183 | -2.850806 | -0.338572 | 0.295997 | -0.189818 | 0.013374 | 0.014838 | 0.927919 | 1.3141 | 0.758824 | 0.97548 | 0.928512 | 1.176184 | -1.325062 | 0.385489 | | 25.7777778 | 0.530823 | 0.250438 |
| 25.7013889 | 0.338768 | 0.436416 | 0.107596 | -0.361353 | -0.099442 | -0.106628 | -0.652287 | -0.484931 | -0.821941 | -0.323758 | -0.981828 | -0.237698 | -0.376712 | 0.240102 | 0.033525 | -0.047803 | -0.178874 | 0.102588 | 1.36532 | 2.067065 | 1.077787 | -0.215355 | 1.678167 | 1.022073 | | 25.7916667 | 0.466911 | 0.144942 |
| 25.7152778 | -0.833867 | -0.857195 | 0.667999 | -0.211525 | 0.166647 | 0.407315 | -0.178013 | -0.204939 | -0.533051 | -0.591192 | 1.694253 | 0.507362 | 0.524713 | -0.262607 | -0.696014 | 0.062283 | -0.938596 | -1.66465 | 0.719679 | 0.79378 | 0.912315 | -0.212512 | 1.403955 | 0.223577 | | 25.8055556 | 0.239277 | 0.107856 |
| 25.7291667 | -0.296279 | 0.049771 | 0.807683 | 0.091303 | 0.555551 | 0.773312 | 0.300352 | 0.317664 | -0.261435 | -1.074759 | 2.395544 | 0.103967 | -0.297327 | -0.543564 | -0.952755 | -0.527093 | -0.004521 | -0.294143 | -0.10559 | -0.928013 | -0.382287 | 0.722804 | -0.570411 | -0.731619 | | 25.8194444 | -0.055317 | -0.043215 |
| 25.7430556 | 0.351015 | 0.640454 | 1.498081 | 0.186293 | -0.075632 | -0.548857 | 0.617745 | 0.720466 | 0.222222 | -0.450986 | 0.495656 | -1.586931 | -2.471933 | -0.235095 | -0.33819 | -0.739588 | 0.719347 | 0.692286 | -1.022711 | -1.832992 | -1.324748 | 0.368365 | -1.444225 | -1.410324 | | 25.8333333 | -0.285078 | -0.38985 |
| 25.7569444 | 0.581507 | 0.46071 | -0.328774 | -0.034809 | -0.386673 | -0.554951 | 0.176356 | 0.405073 | 0.228844 | 0.419341 | -1.256977 | -1.300355 | -1.501864 | 0.244247 | 0.391338 | -0.824872 | 0.484161 | 1.789492 | -0.207462 | -0.895271 | -0.938245 | 0.4599 | -1.077059 | -1.445829 | | 25.8472222 | -0.344953 | -0.721402 |
| 25.7708333 | 0.521913 | 0.207426 | -0.077719 | 0.391398 | -0.189183 | 0.242452 | 0.072055 | 0.142577 | -0.097759 | 0.399879 | -1.193346 | 0.153479 | 0.623146 | -0.093239 | -0.295083 | -0.405771 | -0.31307 | 0.934019 | 0.423984 | -0.035241 | -0.2233 | -1.069831 | -0.083005 | -0.828323 | | 25.8611111 | -0.23864 | -0.961016 |
| 25.7847222 | 0.130018 | -0.810493 | -0.847407 | 0.424417 | 0.225015 | 0.958984 | -0.2808 | -0.403351 | -0.586163 | -0.4972 | -0.251884 | 1.172664 | 2.174164 | -0.714008 | -1.028436 | -0.007271 | -0.270982 | -0.63748 | -0.75606 | 0.053642 | 0.894674 | 0.116435 | 1.257838 | -0.214992 | | 25.875 | -0.136969 | -1.071846 |
| 25.7986111 | 0.020945 | 0.006167 | -0.695082 | 0.014723 | 0.133261 | 0.225435 | -0.585396 | -0.630226 | -0.938201 | -1.634447 | 0.276833 | -0.235041 | -0.697691 | -0.983581 | -0.96161 | 0.273877 | 0.110155 | -1.014113 | -1.234364 | -0.108623 | 0.971491 | 0.71974 | 1.626166 | -1.136162 | | 25.8888889 | -0.109674 | -1.067326 |
| 25.8125 | -0.392125 | 0.821927 | 0.488338 | -0.865602 | -0.407674 | -0.663025 | -0.81141 | -0.397411 | -0.794478 | -1.819223 | 0.228394 | -1.854455 | -4.039591 | -0.598092 | -0.463034 | -0.568634 | -0.488852 | -1.038665 | -0.564913 | 0.097151 | -0.665315 | 3.62262 | 1.052456 | -0.618966 | | 25.9027778 | -0.042045 | -0.95258 |
| 25.8263889 | -0.969814 | 0.835973 | 1.193639 | -1.107412 | -0.595069 | -1.686553 | -0.219461 | 0.331732 | -0.218853 | -0.71207 | -0.493107 | -1.044733 | -1.968292 | -0.510117 | -0.145449 | -0.9415 | -0.816605 | -0.368884 | 0.313426 | -0.198372 | -1.97731 | 2.77559 | 0.21644 | -0.175347 | | 25.9166667 | -0.056041 | -0.828176 |
| 25.8402778 | -1.022707 | -0.873137 | 0.436273 | -0.872121 | -0.886065 | -2.229478 | 0.237937 | 0.418182 | 0.066871 | 0.247676 | -1.238107 | -0.1883 | 0.075418 | -0.341008 | -0.651876 | -0.895764 | 0.002221 | 0.243271 | 0.331959 | -0.37533 | -0.945854 | -0.295087 | 0.092156 | -0.321409 | | 25.9305556 | -0.216356 | -0.744904 |
| 25.8541667 | -1.145605 | -3.121333 | -1.455624 | -0.96571 | -0.775911 | -1.647946 | -0.011279 | -0.228436 | -0.30901 | 0.114776 | -1.343554 | -0.269619 | 0.427938 | -0.935569 | -0.958586 | -0.272214 | 0.399392 | 0.098766 | -0.72783 | -0.457425 | -0.140083 | -0.794375 | -0.619015 | -1.213886 | | 25.9444444 | -0.41286 | -0.549839 |
| 25.8680556 | -1.314829 | -3.906226 | -2.768372 | -1.442488 | -0.456581 | -1.083247 | -0.318686 | -0.591282 | -1.290679 | -1.360127 | -1.366313 | -0.680659 | -1.298559 | -0.864533 | -0.676526 | 0.318004 | -0.900265 | -1.923893 | -1.654195 | -0.436218 | -0.247255 | 0.117837 | 0.149867 | -1.576327 | | 25.9583333 | -0.449114 | -0.266835 |
| 25.8819444 | -1.40704 | -3.733171 | -4.545433 | -2.104694 | -0.198032 | -0.730497 | -0.942833 | -0.960483 | -2.020433 | -2.395415 | -3.088847 | -0.635413 | -1.961169 | -0.372948 | 0.560703 | 0.853051 | -2.014319 | -3.298281 | -1.748678 | -1.230619 | -2.302033 | 2.47564 | -0.004124 | -1.292482 | | 25.9722222 | -0.414886 | -0.222906 |
| 25.8958333 | -2.085727 | -3.264104 | -5.366524 | -2.711109 | -0.775125 | -1.449386 | -1.689102 | -1.325214 | -2.115208 | -2.538849 | -4.465825 | -1.50803 | -1.637707 | -0.313203 | 0.046832 | -0.626314 | -1.550016 | -3.219271 | -1.604848 | -2.673664 | -3.955474 | 2.729849 | -0.056414 | -1.953042 | | 25.9861111 | -0.398545 | -0.457177 |
| 25.9097222 | -2.441165 | -2.876716 | -3.499584 | -2.457985 | -1.825997 | -3.239412 | -1.675989 | -1.548297 | -1.951725 | -2.482819 | -3.595301 | -2.90726 | -1.901709 | -0.370133 | -1.066675 | -1.638472 | -0.647237 | -1.995101 | -1.638779 | -3.334382 | -3.433253 | -1.743285 | -0.991496 | -3.234402 | | 26 | -0.325292 | -0.592664 |
| 25.9236111 | -2.333833 | -2.750254 | -1.478661 | -1.779399 | -2.353506 | -4.298867 | -1.628975 | -1.366468 | -2.112913 | -3.004506 | -3.034845 | -2.622623 | -2.256834 | -1.377779 | -2.006228 | -1.530415 | -1.170571 | -1.365798 | -2.609364 | -2.319482 | -1.764835 | -5.738839 | -1.609543 | -4.169834 | | 26.0138889 | 0.327624 | 0.025146 |
| 25.9375 | -3.333405 | -3.173722 | -1.69162 | -0.864375 | -2.310769 | -3.771483 | -1.907815 | -0.832935 | -4.040395 | -6.028458 | -7.578459 | -1.938532 | -1.804865 | -3.636945 | -2.513868 | -1.136268 | -2.40051 | -1.57243 | -3.52455 | -1.0423 | -1.708687 | -4.841124 | -2.288579 | -9.701076 | | 26.0277778 | 3.595065 | 2.712016 |
| 25.9513889 | -3.916211 | -2.492795 | -2.995411 | 0.536256 | -1.738271 | -1.698013 | -1.664456 | 0.207189 | -5.938546 | -8.59438 | -14.703719 | -1.679386 | 0.250621 | -3.334848 | -1.89769 | -0.425478 | -2.094314 | -2.539365 | -2.270604 | -0.390672 | -1.259116 | -3.427219 | -3.815495 | -20.538774 | | 26.0416667 | 7.95608 | 4.526659 |
| 25.9652778 | -2.709856 | -1.511252 | -2.52188 | 0.97949 | -1.258924 | 0.337435 | -0.654738 | 0.839618 | -4.585844 | -6.591823 | -12.722089 | -1.228507 | 0.957944 | -0.646634 | -0.648821 | 0.268527 | -0.667578 | -1.808077 | 0.156831 | -0.591111 | 0.056207 | -2.226717 | -3.363417 | -21.255941 | | 26.0555556 | 6.018461 | 1.033701 |
| 25.9791667 | -1.414321 | -1.253933 | -1.673611 | 0.097779 | -0.52758 | 0.30858 | -0.145927 | 0.382142 | -1.703612 | -2.361771 | -5.609321 | -0.610711 | 0.77701 | 1.000652 | 0.210896 | 0.477762 | 0.312262 | -1.233735 | 0.894012 | -0.512917 | 0.752855 | -0.723816 | -1.451045 | -9.428512 | | 26.0694444 | 1.406911 | 0.98928 |
| 25.9930556 | -0.98555 | -1.169211 | -1.153983 | -0.395461 | -0.192409 | -0.073481 | -0.082466 | -0.124615 | -0.098156 | -0.301764 | -1.400795 | -0.262559 | 1.164517 | 1.449522 | 0.647708 | 0.687928 | 0.389097 | -1.01251 | 0.841758 | -0.144377 | 0.817052 | -0.352977 | 0.087397 | -1.499708 | | 26.0833333 | -0.559982 | 0.254303 |
| 26.0069444 | -0.685888 | -0.833293 | -0.840404 | -0.225524 | -0.027114 | -0.161077 | -0.127786 | -0.337711 | 0.146775 | 0.045426 | -0.476774 | -0.698003 | 0.534702 | 0.674059 | 0.239566 | 0.654877 | -0.172561 | -0.940537 | 0.514423 | -0.104203 | 0.296834 | -0.134715 | 0.295102 | -1.055447 | | 26.0972222 | -0.476333 | 0.118022 |
| 26.0208333 | -0.449378 | -0.540562 | -0.42206 | 0.010747 | 0.102617 | -0.164731 | -0.163531 | -0.244913 | 0.015849 | -0.065552 | -0.26571 | -1.957431 | -1.390219 | -0.397783 | -0.773715 | 0.023313 | -0.747458 | -1.787101 | -0.472331 | -0.766002 | -0.282049 | -0.303659 | -0.060067 | -1.578326 | | 26.1111111 | -0.006372 | 0.17577 |
| 26.0347222 | -0.369179 | -0.49214 | -0.24621 | 0.058076 | 0.253564 | -0.233422 | -0.240329 | -0.338455 | -0.217963 | -0.255274 | -0.316038 | -2.493021 | -2.547297 | -1.296045 | -1.39937 | -0.744252 | -1.085073 | -2.574656 | -1.412846 | -1.349585 | -1.788566 | -1.090358 | -0.775808 | -1.617103 | | 26.125 | 0.419065 | 0.253195 |
| 26.0486111 | -0.293916 | -0.448283 | -0.38924 | -0.043271 | -0.175571 | -0.332975 | -0.336014 | -0.352505 | -0.385386 | -0.450836 | -0.468281 | -1.79903 | -1.863751 | -1.77767 | -1.475808 | -1.003695 | -1.208387 | -2.214356 | -1.456289 | -1.491774 | -1.814469 | -1.257257 | -0.92731 | -1.511288 | | 26.1388889 | 0.767931 | 0.326119 |
| 26.0625 | -0.252947 | -0.428773 | -0.560371 | -0.194313 | -0.409593 | -0.364971 | -0.387413 | -0.346944 | -0.45137 | -0.547733 | -0.562577 | -1.274103 | -1.140441 | -1.931211 | -1.494486 | -0.847778 | -1.108658 | -1.521424 | -0.982025 | -1.416433 | -1.383992 | -1.22963 | -0.935917 | -1.422619 | | 26.1527778 | 0.968674 | 0.344244 |
| 26.0763889 | -0.288296 | -0.515311 | -0.529515 | -0.209246 | -0.459678 | -0.349807 | -0.391007 | -0.316291 | -0.576984 | -0.64665 | -0.560185 | -0.999169 | -0.800925 | -1.611785 | -1.341263 | -0.663563 | -0.766188 | -0.989333 | -0.500671 | -1.197013 | -0.973287 | -1.019612 | -0.859797 | -0.963772 | | 26.1666667 | 0.811013 | 0.354411 |
| 26.0902778 | -0.271189 | -0.457594 | -0.355947 | -0.136479 | -0.375833 | -0.344029 | -0.35956 | -0.275634 | -0.646584 | -0.63778 | -0.363415 | -0.623795 | -0.533133 | -1.114268 | -0.855973 | -0.522695 | -0.464857 | -0.684716 | -0.153014 | -0.74599 | -0.626692 | -0.836177 | -0.479393 | -0.257458 | | 26.1805556 | 0.407612 | 0.493814 |
| 26.1041667 | -0.221634 | -0.306925 | -0.22598 | -0.122292 | -0.301117 | -0.37551 | -0.305471 | -0.258632 | -0.507104 | -0.467059 | -0.110729 | -0.263616 | 0.055823 | -0.742207 | -0.339533 | -0.244059 | -0.342481 | -0.480817 | -0.031122 | -0.416319 | -0.266303 | -0.690372 | -0.102243 | 0.093817 | | 26.1944444 | 0.131855 | 0.73926 |
| 26.1180556 | -0.205929 | -0.274675 | -0.224612 | -0.163475 | -0.286153 | -0.458076 | -0.254461 | -0.243663 | -0.285855 | -0.291098 | 0.027372 | -0.266847 | 0.554856 | -0.38721 | -0.132721 | 0.09632 | -0.254788 | -0.333889 | -0.123451 | -0.415721 | -0.074149 | -0.461749 | -0.041552 | 0.032071 | | 26.2083333 | 0.224895 | 0.823987 |
| 26.1319444 | -0.192151 | -0.32759 | -0.239252 | -0.163696 | -0.277979 | -0.411492 | -0.171832 | -0.163122 | -0.124786 | -0.173922 | 0.072305 | -0.359637 | 0.391722 | 0.027355 | -0.044859 | 0.318848 | -0.030616 | -0.025983 | 0.00879 | -0.255066 | 0.095815 | -0.14346 | -0.012187 | 0.029877 | | 26.2222222 | 0.473231 | 0.697955 |
| 26.1458333 | -0.099129 | -0.30501 | -0.086213 | -0.057904 | -0.220047 | -0.241095 | -0.019597 | 0.010511 | 0.012328 | -0.07317 | 0.159514 | -0.091219 | 0.139577 | 0.423402 | 0.248262 | 0.437439 | 0.265833 | 0.406611 | 0.368954 | 0.278453 | 0.392228 | 0.248159 | 0.288871 | 0.271392 | | 26.2361111 | 0.543877 | 0.570334 |
| 26.1597222 | 0.035731 | -0.165319 | 0.235073 | 0.143361 | -0.132697 | -0.09016 | 0.166719 | 0.176277 | 0.183047 | 0.103413 | 0.397966 | 0.292451 | 0.304552 | 0.422123 | 0.509672 | 0.390783 | 0.294491 | 0.509554 | 0.409354 | 0.534688 | 0.491532 | 0.389152 | 0.408539 | 0.30485 | | 26.25 | 0.421858 | 0.467567 |
| 26.1736111 | 0.178388 | -0.012938 | 0.452392 | 0.412982 | -0.022783 | 0.002971 | 0.281937 | 0.258472 | 0.329231 | 0.299998 | 0.661958 | 0.370771 | 0.625518 | 0.277908 | 0.565901 | 0.302444 | 0.164618 | 0.35174 | 0.134197 | 0.353555 | 0.355562 | 0.200825 | 0.247006 | 0.080047 | | 26.2638889 | 0.387354 | 0.39279 |
| 26.1875 | 0.259554 | 0.061232 | 0.475763 | 0.572353 | 0.136371 | 0.115869 | 0.270876 | 0.267791 | 0.348394 | 0.402649 | 0.783353 | 0.137475 | 0.350042 | 0.312832 | 0.495413 | 0.257945 | 0.143372 | 0.273488 | -0.003217 | 0.250989 | 0.302678 | -0.027193 | 0.036296 | -0.074595 | | 26.2777778 | 0.521512 | 0.448671 |
| 26.2013889 | 0.279355 | 0.110573 | 0.466151 | 0.590787 | 0.291293 | 0.20519 | 0.201241 | 0.254566 | 0.246967 | 0.354005 | 0.734244 | -0.039903 | -0.186411 | 0.383426 | 0.413535 | 0.196956 | 0.187296 | 0.373003 | 0.077318 | 0.337781 | 0.271143 | -0.079445 | -0.02386 | -0.104129 | | 26.2916667 | 0.769579 | 0.602628 |
| 26.2152778 | 0.20058 | 0.140037 | 0.484319 | 0.527107 | 0.425694 | 0.161378 | 0.126499 | 0.2006 | 0.096655 | 0.155404 | 0.571404 | 0.258576 | 0.062184 | 0.40912 | 0.472598 | 0.261334 | 0.309761 | 0.587184 | 0.265229 | 0.34838 | 0.173451 | 0.046592 | -0.012668 | -0.030658 | | 26.3055556 | 0.51152 | 0.64153 |
| 26.2291667 | 0.102578 | 0.15788 | 0.466491 | 0.395771 | 0.509164 | 0.07133 | 0.099124 | 0.107335 | 0.103107 | 0.071693 | 0.495274 | 0.72251 | 0.569261 | 0.575243 | 0.736907 | 0.46266 | 0.597432 | 0.703518 | 0.470243 | 0.43249 | 0.220886 | 0.23676 | -0.009545 | 0.242535 | | 26.3194444 | -0.135416 | 0.564785 |
| 26.2430556 | 0.152642 | 0.277681 | 0.33824 | 0.243817 | 0.218814 | 0.109905 | 0.145029 | 0.087163 | 0.275498 | 0.190019 | 0.562237 | 0.827924 | 0.413876 | 0.889663 | 0.907765 | 0.56239 | 0.784959 | 0.563683 | 0.634146 | 0.699738 | 0.483898 | 0.372164 | 0.027884 | 0.381613 | | 26.3333333 | -0.425812 | 0.559636 |
| 26.2569444 | 0.298377 | 0.420863 | 0.189213 | 0.159893 | -0.02896 | 0.190764 | 0.191638 | 0.149216 | 0.363073 | 0.278836 | 0.548645 | 0.773535 | 0.391075 | 1.006735 | 0.84329 | 0.547633 | 0.666077 | 0.491392 | 0.716526 | 0.75659 | 0.63525 | 0.578343 | 0.062929 | 0.241371 | | 26.3472222 | -0.095325 | 0.657237 |
| 26.2708333 | 0.339964 | 0.36846 | 0.11509 | 0.149036 | -0.197116 | 0.147908 | 0.196633 | 0.201403 | 0.303757 | 0.251744 | 0.423311 | 0.853134 | 0.854214 | 0.74356 | 0.724389 | 0.594936 | 0.561119 | 0.753789 | 0.664302 | 0.48253 | 0.637906 | 0.828741 | 0.460371 | 0.321061 | | 26.3611111 | 0.415898 | 0.641161 |
| 26.2847222 | 0.225318 | 0.16619 | 0.117124 | 0.189092 | -0.156941 | 0.069531 | 0.210268 | 0.188288 | 0.233927 | 0.214343 | 0.344064 | 0.761164 | 0.763183 | 0.412263 | 0.519287 | 0.455161 | 0.560117 | 0.758918 | 0.511304 | 0.261706 | 0.537722 | 0.817412 | 0.480811 | 0.51308 | | 26.375 | 0.564954 | 0.467894 |
| 26.2986111 | 0.223687 | 0.201672 | 0.185722 | 0.282547 | -0.032893 | 0.161858 | 0.28722 | 0.18936 | 0.298118 | 0.311554 | 0.456878 | 0.544929 | 0.357331 | 0.363556 | 0.278646 | 0.14677 | 0.470991 | 0.464836 | 0.394381 | 0.266874 | 0.302575 | 0.699689 | 0.263226 | 0.442336 | | 26.3888889 | 0.400814 | 0.366629 |
| 26.3125 | 0.482769 | 0.578713 | 0.39665 | 0.455408 | 0.140781 | 0.384698 | 0.467733 | 0.324268 | 0.514831 | 0.592134 | 0.795837 | 0.563463 | 0.652919 | 0.638415 | 0.337469 | 0.180599 | 0.427197 | 0.600912 | 0.354583 | 0.385731 | 0.213747 | 0.745431 | 0.453864 | 0.398288 | | 26.4027778 | 0.268092 | 0.421464 |
| 26.3263889 | 0.882332 | 0.996162 | 0.774541 | 0.694662 | 0.296626 | 0.587716 | 0.707349 | 0.590578 | 0.709249 | 0.877725 | 1.239196 | 0.662551 | 1.077953 | 0.717889 | 0.653965 | 0.515692 | 0.541819 | 1.065038 | 0.363424 | 0.451788 | 0.312804 | 0.774989 | 0.710378 | 0.339685 | | 26.4166667 | 0.316822 | 0.419503 |
| 26.3402778 | 1.27831 | 1.332168 | 1.193916 | 0.924141 | 0.455702 | 0.822668 | 0.908807 | 0.875065 | 0.827772 | 1.082595 | 1.717183 | 0.767708 | 0.983714 | 0.596029 | 0.834698 | 0.693421 | 0.659415 | 1.178633 | 0.470387 | 0.592709 | 0.506464 | 0.736678 | 0.782707 | 0.339396 | | 26.4305556 | 0.461796 | 0.089845 |
| 26.3541667 | 1.632451 | 1.842239 | 1.553323 | 1.094689 | 0.730471 | 1.209163 | 1.112409 | 1.103254 | 1.053418 | 1.343696 | 2.098698 | 1.089599 | 0.986779 | 0.591577 | 0.913262 | 0.808786 | 0.83186 | 1.264279 | 0.72912 | 0.887783 | 0.799643 | 0.89589 | 0.85501 | 0.851178 | | 26.4444444 | 0.448676 | -0.323397 |
| 26.3680556 | 1.914086 | 2.504264 | 1.891705 | 1.348703 | 1.022727 | 1.645856 | 1.283381 | 1.352615 | 1.412661 | 1.829923 | 2.265412 | 1.418868 | 1.413203 | 0.919485 | 1.178043 | 1.079952 | 1.146073 | 1.917995 | 1.074322 | 1.197949 | 1.25843 | 1.252738 | 1.084138 | 1.404028 | | 26.4583333 | 0.227369 | -0.351472 |
| 26.3819444 | 2.168955 | 3.289137 | 2.172159 | 1.469669 | 1.186816 | 2.018309 | 1.455808 | 1.54986 | 1.708077 | 2.3819 | 2.385025 | 1.635129 | 1.8829 | 1.38321 | 1.598963 | 1.372558 | 1.527504 | 2.924931 | 1.433398 | 1.528403 | 1.719335 | 1.72153 | 1.642154 | 1.518847 | | 26.4722222 | 0.025504 | -0.233061 |
| 26.3958333 | 2.336018 | 3.939142 | 2.542661 | 1.78372 | 1.302522 | 2.473766 | 1.518272 | 1.634347 | 1.706202 | 2.307296 | 2.780729 | 1.944445 | 2.247042 | 1.681522 | 2.098523 | 1.682151 | 1.834856 | 3.425727 | 1.894713 | 2.006223 | 2.091467 | 2.374154 | 2.531634 | 1.848458 | | 26.4861111 | 0.127585 | 0.043147 |
| 26.4097222 | 2.342521 | 4.693043 | 3.178316 | 2.200114 | 1.473904 | 3.168459 | 1.538677 | 1.831073 | 1.498975 | 2.264942 | 3.25002 | 2.911437 | 3.118762 | 2.217067 | 2.784784 | 2.169386 | 2.138268 | 3.618247 | 2.743646 | 2.965872 | 2.739555 | 3.272114 | 3.503456 | 2.654396 | | 26.5 | 0.583566 | 0.189864 |
| 26.4236111 | 2.38703 | 5.749252 | 3.701464 | 2.488364 | 1.415952 | 3.479999 | 1.648669 | 1.891987 | 1.683946 | 2.766237 | 3.810093 | 4.365902 | 4.637893 | 3.156743 | 3.73839 | 2.728852 | 2.436145 | 3.808836 | 3.698873 | 4.005799 | 3.479254 | 4.295116 | 4.679763 | 5.211649 | | 26.5138889 | 0.566471 | -0.298132 |
| 26.4375 | 2.505821 | 5.041707 | 3.265176 | 1.229656 | 1.018802 | 2.218905 | 1.715355 | 1.00887 | 3.498289 | 4.942576 | 7.136894 | 4.899179 | 4.632447 | 3.537761 | 4.479474 | 2.838799 | 2.680167 | 3.749776 | 3.569819 | 4.112416 | 3.46149 | 5.890984 | 5.192568 | 15.231482 | | 26.5277778 | -1.908933 | -1.882615 |
| 26.4513889 | 2.974694 | 3.859422 | 1.919647 | -0.301162 | 0.729968 | 0.917674 | 1.47847 | -0.496504 | 5.864903 | 7.626138 | 12.106627 | 4.725034 | 2.672689 | 3.253543 | 4.743532 | 2.720447 | 3.000863 | 3.504033 | 2.835744 | 3.737922 | 3.358589 | 7.295183 | 5.943254 | 28.950565 | | 26.5416667 | -6.520316 | -3.806983 |
| 26.4652778 | 3.315967 | 1.799302 | 0.043294 | -1.012592 | 0.583453 | 0.542437 | 0.817081 | -0.880981 | 5.544441 | 7.5886 | 11.068779 | 3.558832 | 0.596619 | 3.358743 | 3.944249 | 2.411743 | 2.457516 | 2.642905 | 1.246743 | 1.922042 | 1.143525 | 5.588213 | 4.830863 | 23.180331 | | 26.5555556 | -7.960544 | -6.732384 |
| 26.4791667 | 0.700519 | -1.280993 | -2.378128 | -0.778247 | 0.60452 | 0.621676 | -0.032569 | -0.305671 | 2.73737 | 3.607453 | 2.818564 | 1.295307 | -0.231191 | -0.253026 | 0.41912 | 0.746914 | 0.57588 | 0.373193 | -2.086328 | -1.172055 | -0.498241 | 1.563545 | 1.392434 | 7.729782 | | 26.5694444 | -3.048992 | -6.30872 |
| 26.4930556 | -0.836753 | -1.00666 | -1.977702 | 0.438767 | 0.744397 | 0.880402 | 0.10474 | 0.017478 | 1.048023 | 0.676479 | -2.553694 | 0.256699 | -0.467066 | -0.381642 | 0.021203 | -0.020207 | 0.206306 | -0.324422 | -1.266655 | -0.58013 | 0.696314 | 0.447858 | 0.415573 | 2.114486 | | 26.5833333 | -0.260178 | -2.470043 |
| 26.5069444 | -0.020693 | 0.685796 | 0.690374 | 1.974363 | 1.050754 | 1.658322 | 0.678504 | 0.531803 | 1.753537 | 1.196057 | 0.64645 | 1.528899 | 2.114365 | 1.962113 | 1.597063 | -0.204647 | 2.209557 | 4.174026 | 1.803651 | 2.844094 | 4.515511 | 1.604057 | 1.965599 | 4.752888 | | 26.5972222 | 0.141759 | -0.7195 |
| 26.5208333 | 1.564672 | 2.204974 | 2.888881 | 2.983275 | 0.777947 | 1.725753 | 0.896374 | 0.425318 | 1.500714 | 1.688141 | 4.821279 | 1.88669 | 4.825719 | 2.859181 | 2.214326 | 0.753724 | 2.533802 | 4.472004 | 4.391449 | 5.342729 | 6.585249 | 1.766417 | 2.49093 | 7.451844 | | 26.6111111 | 0.209796 | -0.003285 |
| 26.5347222 | 2.046267 | 3.455633 | 2.787109 | 3.006611 | 0.271442 | 0.355104 | 0.508233 | 0.548245 | 1.131139 | 1.856698 | 6.142492 | 2.106839 | 5.667946 | 4.012944 | 3.747831 | 2.859903 | 1.680903 | 2.694155 | 3.453439 | 4.687153 | 4.149423 | 2.01197 | 2.503006 | 6.05864 | | 26.625 | 0.215886 | 0.270729 |
| 26.5486111 | 1.87851 | 3.544948 | 1.401957 | 1.845613 | 0.449848 | 1.138141 | 0.225451 | 0.429643 | 0.769269 | 1.392576 | 4.854355 | 1.578156 | 4.261072 | 1.700131 | 1.196602 | 1.957001 | 0.903665 | 0.912873 | 1.002454 | 2.295191 | 1.265838 | 1.45121 | 1.615427 | 1.72082 | | 26.6388889 | -0.016648 | 0.312552 |
| 26.5625 | 0.714781 | 1.492715 | 0.498938 | -0.525454 | 0.036289 | 0.072766 | 0.233283 | 0.485593 | 0.250483 | 0.367219 | 2.491893 | 0.672115 | 1.176964 | 0.364731 | -0.372242 | 0.451633 | -0.084336 | -0.953959 | -1.327072 | -0.382153 | -1.367525 | -0.448378 | 0.34236 | -1.480578 | | 26.6527778 | -0.257873 | 0.198058 |
| 26.5763889 | -0.186796 | -0.448881 | -0.545277 | -1.828471 | -0.163392 | -0.363146 | 0.002837 | -0.335369 | 0.191253 | -0.376624 | -0.779578 | -0.038108 | -0.179544 | 0.249687 | -0.206502 | 0.041202 | -0.26455 | -0.539082 | 0.571234 | 0.205551 | -0.270462 | -0.198372 | -0.309941 | -0.904717 | | 26.6666667 | -0.229715 | -0.106457 |
| 26.5902778 | -0.487475 | -1.388513 | -1.048261 | -1.095808 | -0.212286 | -0.305024 | -0.01429 | -0.765966 | 0.335628 | -0.005827 | -3.404827 | 0.448952 | 0.854212 | 1.006276 | 1.4621 | 1.010467 | 0.937449 | 1.47492 | 3.012342 | 2.735746 | 2.321319 | 0.097875 | 0.1469 | 1.549071 | | 26.6805556 | 0.305705 | -0.11037 |
| 26.6041667 | -0.488088 | -0.297791 | -0.49808 | 0.032494 | -0.072818 | 0.130191 | -0.149876 | -0.699085 | 0.394353 | 0.383469 | -3.527354 | 1.07017 | 2.089012 | 2.490388 | 3.511852 | 1.476205 | 1.323075 | 2.135326 | 3.180671 | 3.490109 | 3.540942 | -0.133774 | 0.533762 | 3.359777 | | 26.6944444 | 0.791385 | 0.257569 |
| 26.6180556 | -0.268483 | 0.534702 | 0.580448 | 0.474017 | 0.103438 | 0.617749 | -0.222572 | -0.147485 | -0.46707 | -0.618011 | -1.02655 | 0.955489 | 1.987523 | 0.748404 | 0.675655 | 0.495474 | 0.587774 | 0.732647 | 0.935461 | 1.059648 | 1.390436 | 1.970469 | 0.125134 | 1.38062 | | 26.7083333 | 0.574749 | 0.705792 |
| 26.6319444 | 0.411125 | 3.019238 | 3.392646 | 0.969095 | 0.462375 | 0.975914 | 0.014639 | 0.241421 | -0.448005 | -0.280623 | 2.03503 | 0.428772 | 0.748355 | 0.141806 | -0.666776 | -0.68661 | -0.980877 | -2.529564 | -2.146779 | -1.953497 | -2.243453 | -0.158493 | -0.413326 | -1.864869 | | 26.7222222 | 0.048064 | 1.184955 |
| 26.6458333 | 1.099519 | 2.876084 | 2.267468 | 0.405643 | 0.37605 | 0.677963 | 0.291357 | -0.38691 | -0.264923 | -0.194316 | 2.13351 | -0.239778 | -0.642604 | -0.137932 | -0.769344 | -0.633887 | -0.990318 | -2.783529 | -1.824144 | -2.446621 | -1.502989 | -1.320369 | -0.943808 | -1.866907 | | 26.7361111 | -0.291698 | 1.287867 |
| 26.6597222 | 0.740045 | 1.317703 | 0.901885 | 0.389061 | -0.097281 | -0.271768 | 0.2496 | -0.241192 | 0.339553 | 0.578573 | 0.898791 | 0.050217 | -1.308247 | -0.22891 | -0.461155 | -0.440883 | 0.389042 | -0.301689 | 0.451929 | -0.955177 | 0.369141 | -0.561988 | -0.903176 | -0.111078 | | 26.75 | -0.317791 | 0.746246 |
| 26.6736111 | -0.132407 | 0.25429 | 0.263358 | -0.313037 | -0.623836 | -0.729009 | -0.133656 | -0.059739 | 0.371734 | 0.767051 | 0.248458 | 1.177477 | 0.684979 | 0.47047 | 0.831123 | 0.283983 | 0.153088 | 0.136175 | 1.20331 | 0.729286 | 1.246525 | 0.772479 | 0.277447 | 0.926669 | | 26.7638889 | -0.219601 | 0.542588 |
| 26.6875 | -0.624455 | -0.432676 | -0.383389 | -0.730548 | -0.823905 | -0.92091 | -0.452415 | -0.143968 | -0.534496 | -0.241677 | -0.612057 | 1.313169 | 2.59866 | 0.903861 | 1.522118 | 1.245227 | -0.241369 | -0.094915 | 1.098445 | 0.756364 | 0.748258 | 2.470878 | 1.089516 | 0.578303 | | 26.7777778 | -0.19466 | 0.421282 |
| 26.7013889 | -0.797395 | -1.279791 | -1.832149 | -0.771594 | -0.494697 | -0.620971 | -0.592271 | -0.430794 | -1.052402 | -1.138732 | -2.025414 | 0.195333 | 0.017685 | 1.053955 | 1.142159 | 0.709841 | 1.110331 | 1.93519 | -0.162569 | 0.896665 | 0.29574 | 1.374994 | 1.778443 | -0.281725 | | 26.7916667 | -0.223877 | 0.074115 |
| 26.7152778 | -0.673391 | -2.278369 | -2.180593 | -0.478962 | 0.390825 | 0.264509 | -0.255513 | -0.230126 | -1.230057 | -1.406764 | -3.086192 | 0.042101 | 0.125967 | 0.352314 | -0.164161 | -0.596475 | 0.86307 | 0.038947 | -0.781814 | 0.439313 | 0.514634 | -1.706292 | -1.26648 | 0.031212 | | 26.8055556 | -0.147671 | -0.274389 |
| 26.7291667 | -0.304879 | -2.593545 | -2.51134 | -0.949253 | 0.804706 | 0.693275 | 0.157294 | 0.122374 | -0.032645 | -0.343344 | -2.712646 | -0.199411 | 0.456392 | 0.559859 | 0.345623 | -0.414028 | -0.65601 | -2.285966 | -0.402746 | -0.33982 | -0.250419 | -1.533787 | -0.022768 | -0.018316 | | 26.8194444 | -0.115929 | -0.369612 |
| 26.7430556 | -0.166384 | -2.026632 | -1.625163 | -0.755827 | 0.374007 | 0.012904 | 0.227024 | 0.245669 | 0.860211 | 0.843392 | -0.385625 | 0.248444 | 0.756764 | 0.411642 | 0.709248 | -0.23053 | -0.693829 | -1.029863 | 0.155255 | -1.457091 | -0.881537 | 1.755092 | 0.675237 | -0.47597 | | 26.8333333 | -0.15245 | -0.292084 |
| 26.7569444 | -0.136941 | -0.800785 | -0.852097 | -0.840077 | -0.210512 | -0.415864 | 0.236042 | 0.213142 | 0.680608 | 1.423931 | -0.123342 | 0.38788 | 0.017278 | 0.606793 | 1.593789 | 1.177923 | -0.395644 | 0.510981 | 1.364724 | -0.039388 | 0.173742 | 1.984363 | 1.52494 | -0.378654 | | 26.8472222 | -0.104476 | -0.082653 |

| | | | | | | | | | | | | | | | | | | | | | | | | | | | |
|---|---|---|---|---|---|---|---|---|---|---|---|---|---|---|---|---|---|---|---|---|---|---|---|---|---|---|---|
| 26.7708333 | -0.35336 | 0.052968 | -1.132372 | -1.342641 | -0.536503 | -0.852173 | 0.052116 | 0.169922 | 0.032196 | 0.746117 | -0.453885 | 0.418995 | -0.475402 | -0.706242 | -0.380275 | 0.035669 | 1.17803 | 2.608188 | 1.100612 | 1.258572 | 1.009415 | -0.268401 | 0.001986 | 0.415408 | 26.8611111 | -0.040188 | 0.090322 |
| 26.7847222 | -0.47122 | 0.774602 | -1.160232 | -1.762717 | -0.497129 | -1.080773 | -0.324743 | 0.00361 | -1.611361 | -1.572538 | -0.094529 | 0.405711 | 0.16741 | -0.321043 | -0.633648 | -0.751287 | 2.270398 | 2.984564 | -1.598532 | 0.488464 | 0.156242 | -1.462294 | -1.586121 | 0.522451 | 26.875 | 0.032857 | 0.120767 |
| 26.7986111 | 0.327994 | 1.391293 | -0.25637 | -1.03666 | 0.105419 | -0.220796 | -0.406978 | 0.115668 | -1.846412 | -2.656528 | 0.744622 | 0.959013 | 2.282286 | 0.225982 | 0.222669 | 0.146619 | -0.318775 | -0.917556 | -3.063648 | -2.102751 | -1.787131 | 0.278935 | 1.403822 | -0.854919 | 26.8888889 | -0.02296 | 0.076212 |
| 26.8125 | 0.331186 | 1.712809 | 1.600912 | 0.442027 | 0.540645 | 0.485495 | -0.096792 | 0.156798 | -0.517466 | -1.364626 | 2.035425 | 0.014874 | 0.22497 | 0.090907 | 0.726555 | 0.828101 | -1.415019 | -2.312196 | -0.773937 | -2.38933 | -2.509987 | 2.080441 | -1.094809 | -1.780058 | 26.9027778 | -0.186968 | -0.071267 |
| 26.8263889 | 0.05288 | 0.881683 | 2.55587 | 1.815544 | 0.501713 | 0.196302 | -0.120006 | -0.081093 | 0.577205 | -0.07389 | 2.641155 | -0.890043 | -2.247831 | 0.995481 | 1.841109 | 1.814165 | -0.592086 | -1.038736 | 1.417564 | -0.581241 | -0.468174 | -0.241113 | -0.204896 | -0.822762 | 26.9166667 | -0.469922 | -0.335917 |
| 26.8402778 | -0.027614 | -0.89311 | 0.798394 | 1.393033 | -0.112225 | -0.48705 | 0.154769 | -0.238689 | 0.718057 | 0.606222 | 1.36605 | -0.682619 | -1.653841 | 0.519768 | 0.636962 | 0.274578 | 0.067498 | 0.007065 | 0.159298 | -0.24256 | 0.080673 | -2.228311 | -1.359216 | -0.87911 | 26.9305556 | -0.75412 | -0.623847 |
| 26.8541667 | -1.17679 | -2.267517 | -1.731866 | -0.95447 | -0.871724 | -1.132181 | -0.243383 | -0.614878 | -0.698698 | -0.44811 | -1.413471 | -0.943204 | -1.391666 | -1.514239 | -1.127204 | -1.049436 | 0.13011 | 0.507924 | -2.684757 | -1.267174 | -0.993483 | -0.950693 | -2.887594 | -3.176778 | 26.9444444 | -1.027026 | -0.637136 |
| 26.8680556 | -2.373129 | -3.486571 | -3.180585 | -2.531948 | -1.547343 | -2.219109 | -1.149047 | -0.768346 | -2.116221 | -1.851645 | -4.498561 | -1.938276 | -2.586698 | -1.753832 | -0.840214 | -0.667504 | -0.811178 | -1.286122 | -3.411929 | -2.102177 | -2.586207 | 0.446535 | -1.118676 | -4.338581 | 26.9583333 | -1.090997 | -0.385595 |
| 26.8819444 | -2.379524 | -4.372662 | -3.975264 | -3.845295 | -1.342372 | -2.001507 | -1.836629 | -0.536698 | -2.658452 | -2.520206 | -5.548463 | -2.285346 | -3.348894 | -1.4207 | -2.03811 | -1.134223 | -1.731083 | -3.116101 | -1.651407 | -1.728005 | -3.430175 | 1.654351 | -0.453943 | -2.813398 | 26.9722222 | -1.197966 | -0.223826 |
| 26.8958333 | -2.827699 | -5.894776 | -5.027067 | -4.061896 | -1.068223 | -1.901698 | -1.795649 | -1.097639 | -2.062816 | -2.234392 | -4.735514 | -1.099008 | -1.958519 | 0.83094 | -1.214371 | -1.452948 | -1.772799 | -3.082876 | -1.77979 | -2.139445 | -2.835822 | 1.088601 | -0.818487 | -1.7424 | 26.9861111 | -1.125929 | -0.135383 |
| 26.9097222 | -3.408768 | -5.159751 | -4.882168 | -3.125935 | -1.367005 | -2.726178 | -1.998132 | -1.746776 | -1.129791 | -1.413095 | -3.54175 | -0.983542 | -1.444669 | 0.616771 | -1.313429 | -1.883228 | -1.102415 | -1.97671 | -3.677754 | -3.425992 | -1.914997 | -4.758168 | -2.62204 | -3.778422 | 27 | -1.003533 | -0.025567 |
| 26.9236111 | -3.530148 | -4.328204 | -4.355143 | -2.01333 | -1.781586 | -3.554252 | -1.906002 | -1.694626 | -1.494014 | -2.261698 | -4.469343 | -2.578274 | -3.178131 | -3.033626 | -2.202519 | -1.901751 | -1.305305 | -1.501323 | -4.607476 | -3.643152 | -2.248287 | -6.968566 | -3.570728 | -7.069853 | 27.0138889 | -0.411441 | 0.417212 |
| 26.9375 | -3.921089 | -3.590584 | -4.329967 | -0.928374 | -1.759001 | -3.357688 | -2.353659 | -1.345201 | -3.95418 | -5.842299 | -11.079406 | -4.538405 | -4.239093 | -4.935244 | -3.04525 | -2.544942 | -2.922057 | -2.9101 | -3.851223 | -3.696494 | -2.13688 | -4.444737 | -1.944383 | -11.123501 | 27.0277778 | 2.318075 | 1.705863 |
| 26.9513889 | -5.296047 | -2.525864 | -3.749004 | 0.426875 | -0.888074 | -1.763887 | -2.718363 | -0.695584 | -6.973756 | -9.858206 | -19.752008 | -3.672414 | -1.560839 | -3.717037 | -3.891792 | -3.181618 | -2.935304 | -3.244615 | -1.893045 | -2.499244 | -0.627463 | -3.556415 | -1.887887 | -16.336122 | 27.0416667 | 6.903258 | 3.759907 |
| 26.9652778 | -3.547148 | -1.5184 | -1.831173 | 0.501285 | -0.093976 | -0.140506 | -1.721649 | -0.128637 | -5.93451 | -7.718858 | -16.749524 | -2.428028 | -0.324943 | -0.830471 | -2.799176 | -3.037658 | -2.007973 | -1.675477 | -1.242242 | -1.957403 | -0.381175 | -3.423509 | -2.668588 | -16.283348 | 27.0555556 | 7.059936 | 1.254277 |
| 26.9791667 | -1.423456 | -1.009575 | -0.822912 | 0.098326 | -0.178179 | -0.159665 | -0.695642 | -0.132066 | -2.50324 | -2.965828 | -6.534422 | -1.722381 | -0.631917 | 0.534074 | -1.609978 | -2.131955 | -0.975126 | -0.709698 | -1.467365 | -1.550962 | -0.900935 | -2.747187 | -1.917865 | -7.400905 | 27.0694444 | 2.165198 | 0.723619 |
| 26.9930556 | -0.311811 | -0.677282 | -0.2946 | -0.356374 | -0.309286 | -0.487687 | -0.245832 | -0.281215 | -0.206656 | -0.481957 | -1.018098 | -1.39521 | -0.244267 | 0.56341 | -0.740944 | -0.865376 | -0.478706 | -0.316269 | -1.260209 | -0.973642 | -0.688793 | -1.791237 | -0.8482 | -1.589561 | 27.0833333 | -0.501418 | 0.099627 |
| 27.0069444 | 0.294716 | -0.160538 | 0.175035 | -0.166014 | -0.296395 | -0.389093 | -0.096277 | -0.339047 | 0.245986 | 0.113644 | 0.388567 | -1.692842 | -0.446081 | -0.191329 | -0.665201 | -0.101554 | -0.629644 | -0.471863 | -0.693485 | -0.616915 | -0.340114 | -1.127748 | -0.425534 | -1.22065 | 27.0972222 | -0.710488 | -0.014346 |
| 27.0208333 | 0.496386 | 0.010281 | 0.340853 | 0.143301 | -0.241835 | -0.237478 | -0.163565 | -0.417967 | -0.009456 | 0.049143 | 0.711126 | -2.348416 | -1.950876 | -1.120149 | -1.467409 | -0.484966 | -1.189858 | -1.659462 | -0.627155 | -0.883253 | -0.992943 | -1.473581 | -0.820157 | -1.921289 | 27.1111111 | -0.293665 | 0.132869 |
| 27.0347222 | 0.000577 | -0.388938 | 0.115521 | 0.223219 | -0.237931 | -0.314813 | -0.31081 | -0.526885 | -0.333225 | -0.33215 | 0.563557 | -2.564614 | -2.927832 | -1.679408 | -2.242319 | -1.415728 | -1.696064 | -2.948279 | -1.16498 | -1.662039 | -2.323001 | -2.039784 | -1.222663 | -2.193536 | 27.125 | 0.168966 | 0.273964 |
| 27.0486111 | -0.331002 | -0.744389 | -0.17171 | -0.034275 | -0.3006 | -0.527162 | -0.444711 | -0.573791 | -0.602707 | -0.727494 | 0.067166 | -2.567826 | -2.726195 | -1.902836 | -2.213274 | -1.645525 | -1.666635 | -3.162259 | -1.6072 | -1.932592 | -2.579383 | -2.18431 | -1.430577 | -2.026986 | 27.1388889 | 0.565773 | 0.342163 |
| 27.0625 | -0.586619 | -0.94588 | -0.348818 | -0.24629 | -0.355857 | -0.626268 | -0.549925 | -0.585238 | -0.703511 | -0.894588 | -0.511578 | -2.155564 | -1.893079 | -1.860853 | -1.503001 | -1.081362 | -1.252322 | -2.266302 | -1.451705 | -1.563293 | -1.640235 | -1.781201 | -1.672654 | -1.740715 | 27.1527778 | 0.848545 | 0.343595 |
| 27.0763889 | -0.67698 | -0.996052 | -0.388769 | -0.33661 | -0.368162 | -0.594717 | -0.63561 | -0.58673 | -0.592575 | -0.810864 | -0.637315 | -1.655297 | -1.421278 | -1.555385 | -0.930704 | -0.745326 | -0.932044 | -1.318157 | -0.923451 | -1.173854 | -0.874916 | -1.310909 | -1.5259 | -1.294047 | 27.1666667 | 0.821673 | 0.34893 |
| 27.0902778 | -0.678682 | -0.993408 | -0.464001 | -0.371703 | -0.373421 | -0.56056 | -0.640798 | -0.599476 | -0.537471 | -0.751896 | -0.477798 | -1.258689 | -1.092885 | -1.154762 | -0.909188 | -0.705777 | -0.825871 | -0.951941 | -0.488686 | -0.93258 | -0.802168 | -0.91056 | -1.244566 | -0.866885 | 27.1805556 | 0.466403 | 0.434683 |
| 27.1041667 | -0.794155 | -1.064509 | -0.614115 | -0.313553 | -0.383128 | -0.602675 | -0.625868 | -0.629405 | -0.730665 | -0.893691 | -0.603744 | -0.859359 | -0.468024 | -1.054275 | -1.030894 | -0.555226 | -0.578145 | -0.850544 | -0.311106 | -0.773515 | -0.721342 | -0.578935 | -1.008646 | -0.593696 | 27.1944444 | -0.009148 | 0.660747 |
| 27.1180556 | -1.079 | -1.255671 | -0.79668 | -0.295928 | -0.381612 | -0.689108 | -0.674894 | -0.690806 | -0.974338 | -1.043019 | -0.969861 | -0.558681 | -0.105837 | -1.101949 | -0.87853 | -0.332446 | -0.305921 | -0.702879 | -0.359127 | -0.552741 | -0.322738 | -0.531502 | -0.603974 | -0.324356 | 27.2083333 | -0.146177 | 0.891808 |
| 27.1319444 | -1.287915 | -1.402634 | -0.934702 | -0.372639 | -0.384613 | -0.775528 | -0.695808 | -0.724976 | -0.975663 | -1.008271 | -1.054842 | -0.382703 | -0.238254 | -1.007618 | -0.570338 | -0.336478 | -0.307739 | -0.581634 | -0.378503 | -0.394394 | -0.181368 | -0.609063 | -0.362929 | -0.128491 | 27.2222222 | 0.21695 | 0.910864 |
| 27.1458333 | -1.02388 | -1.15427 | -0.841468 | -0.460953 | -0.404132 | -0.781578 | -0.600246 | -0.660809 | -0.715425 | -0.780532 | -0.789806 | -0.223899 | -0.216038 | -0.581475 | -0.319417 | -0.312488 | -0.42469 | -0.532313 | -0.243118 | -0.245479 | -0.262235 | -0.48255 | -0.30508 | -0.073879 | 27.2361111 | 0.590566 | 0.641735 |
| 27.1597222 | -0.550269 | -0.676255 | -0.612826 | -0.519103 | -0.421349 | -0.660067 | -0.43059 | -0.45788 | -0.423071 | -0.434541 | -0.437611 | -0.060138 | 0.130665 | -0.04655 | -0.134437 | -0.052567 | -0.201587 | -0.422875 | -0.075275 | -0.116026 | -0.155848 | -0.266228 | -0.197456 | -0.095182 | 27.25 | 0.330691 | 0.324881 |
| 27.1736111 | -0.033933 | -0.198763 | -0.305904 | -0.434112 | -0.36892 | -0.481621 | -0.266458 | -0.230068 | -0.202687 | -0.094259 | 0.041608 | 0.043074 | 0.2756 | 0.279695 | -0.050678 | 0.056881 | 0.082374 | -0.224268 | 0.023165 | 0.034449 | 0.189881 | -0.242238 | 0.034978 | -0.118822 | 27.2638889 | -0.263003 | 0.321664 |
| 27.1875 | 0.333598 | 0.161267 | -0.05216 | -0.233036 | -0.261411 | -0.32351 | -0.174098 | -0.078246 | -0.102643 | 0.023826 | 0.419852 | 0.055489 | 0.119804 | 0.298622 | -0.008533 | -0.062383 | 0.007038 | -0.085924 | -0.025535 | 0.064013 | 0.178356 | -0.196375 | 0.105249 | -0.024176 | 27.2777778 | -0.666233 | 0.593174 |
| 27.2013889 | 0.431464 | 0.317749 | -0.014842 | -0.138064 | -0.186923 | -0.209953 | -0.173327 | -0.028177 | -0.159263 | -0.110125 | 0.337876 | 0.024658 | 0.011458 | 0.19203 | 0.067254 | -0.059196 | -0.102526 | 0.013228 | -0.137168 | -0.011093 | -0.165438 | -0.027203 | 0.011894 | 0.117285 | 27.2916667 | -0.289559 | 0.697975 |
| 27.2152778 | 0.305182 | 0.168581 | -0.107809 | -0.216074 | -0.173948 | -0.157309 | -0.188291 | -0.059953 | -0.252776 | -0.230635 | 0.082871 | -0.013148 | -0.000379 | 0.187435 | 0.25137 | 0.196595 | 0.161791 | 0.158789 | -0.110273 | 0.033472 | -0.089864 | -0.007056 | 0.097641 | 0.180086 | 27.3055556 | 0.758208 | 0.572022 |
| 27.2291667 | 0.093262 | -0.069258 | -0.189433 | -0.318029 | -0.175577 | -0.132088 | -0.222783 | -0.164021 | -0.280859 | -0.231261 | 0.015513 | -0.032658 | -0.072444 | 0.193386 | 0.294213 | 0.257226 | 0.355264 | 0.313836 | -0.027472 | 0.064434 | 0.144211 | -0.162227 | 0.273514 | 0.181723 | 27.3194444 | 1.120381 | 0.416639 |
| 27.2430556 | -0.015496 | -0.043487 | -0.165565 | -0.266352 | -0.141251 | -0.094159 | -0.219619 | -0.157738 | -0.296833 | -0.245434 | 0.00582 | 0.027161 | -0.045487 | 0.127191 | 0.162886 | 0.025522 | 0.113272 | 0.310851 | -0.036865 | -0.03579 | -0.034726 | -0.165839 | 0.125974 | -0.002527 | 27.3333333 | 0.430244 | 0.454269 |
| 27.2569444 | -0.045635 | 0.100718 | -0.103729 | -0.112953 | -0.092152 | -0.084689 | -0.139296 | -0.041932 | -0.298272 | -0.320623 | -0.039703 | 0.168822 | 0.192001 | 0.103342 | 0.189125 | 0.063432 | -0.013607 | 0.194508 | 0.011234 | 0.093047 | -0.123548 | -0.008839 | -0.074492 | -0.135977 | 27.3472222 | -0.162004 | 0.516863 |
| 27.2708333 | -0.020897 | 0.068087 | -0.064187 | -0.005076 | -0.047014 | -0.065451 | 0.011106 | 0.046767 | -0.182798 | -0.314336 | 0.065368 | 0.293741 | 0.421046 | 0.152522 | 0.460259 | 0.431094 | 0.316038 | 0.217815 | 0.204165 | 0.326812 | 0.238353 | 0.09774 | 0.060319 | 0.041222 | 27.3611111 | -0.224552 | 0.31507 |
| 27.2847222 | 0.029039 | -0.127678 | -0.029028 | 0.087728 | 0.00151 | 0.018312 | 0.15739 | 0.06096 | -0.066261 | -0.204421 | 0.243946 | 0.42139 | 0.532508 | 0.227484 | 0.636455 | 0.586081 | 0.5724 | 0.525107 | 0.383029 | 0.410982 | 0.481712 | 0.157447 | 0.276013 | 0.251353 | 27.375 | 0.281722 | 0.060073 |
| 27.2986111 | 0.118802 | -0.159721 | 0.065109 | 0.245015 | 0.027563 | 0.099823 | 0.261169 | 0.108352 | -0.0414 | -0.080481 | 0.318619 | 0.574407 | 0.671193 | 0.353437 | 0.59168 | 0.466069 | 0.461845 | 0.763641 | 0.393501 | 0.447079 | 0.459214 | 0.344864 | 0.395314 | 0.340118 | 27.3888889 | 0.903784 | -0.01531 |
| 27.3125 | 0.311597 | 0.136121 | 0.211065 | 0.449851 | 0.064866 | 0.182816 | 0.335589 | 0.259016 | 0.04613 | 0.05237 | 0.382389 | 0.792105 | 0.918481 | 0.555283 | 0.612879 | 0.543903 | 0.399369 | 0.711984 | 0.378975 | 0.620733 | 0.650653 | 0.546031 | 0.616532 | 0.475719 | 27.4027778 | 0.822334 | -0.007688 |
| 27.3263889 | 0.561678 | 0.602464 | 0.337571 | 0.601079 | 0.197048 | 0.3743 | 0.445422 | 0.423416 | 0.336846 | 0.280024 | 0.597374 | 1.09193 | 1.195628 | 0.846739 | 0.822084 | 0.880724 | 0.608527 | 0.741699 | 0.531721 | 0.840456 | 1.019425 | 0.656694 | 0.983092 | 0.674308 | 27.4166667 | 0.25315 | 0.054486 |
| 27.3402778 | 0.847173 | 0.99219 | 0.54899 | 0.737559 | 0.366771 | 0.633628 | 0.623734 | 0.599407 | 0.662097 | 0.573127 | 1.038115 | 1.30282 | 1.331238 | 1.11432 | 1.129202 | 1.174849 | 0.802716 | 1.075227 | 0.768167 | 1.041669 | 1.221488 | 0.85635 | 1.104405 | 0.872631 | 27.4305556 | -0.391969 | 0.167448 |
| 27.3541667 | 1.291889 | 1.283323 | 1.008968 | 0.97563 | 0.484733 | 0.846059 | 0.918492 | 0.909798 | 0.853758 | 0.886017 | 1.628573 | 1.315077 | 1.408484 | 1.326593 | 1.338984 | 1.281473 | 0.903351 | 1.511151 | 1.014499 | 1.324338 | 1.479649 | 1.292304 | 1.128637 | 1.227748 | 27.4444444 | -0.560362 | 0.152242 |
| 27.3680556 | 2.081879 | 1.742978 | 1.743528 | 1.457614 | 0.622289 | 1.116904 | 1.317965 | 1.388555 | 1.10394 | 1.275426 | 2.404074 | 1.430592 | 1.707333 | 1.232205 | 1.491955 | 1.488809 | 1.155152 | 2.00553 | 1.292628 | 1.65542 | 2.03242 | 1.796251 | 1.644871 | 1.657226 | 27.4583333 | 0.114974 | -0.023545 |
| 27.3819444 | 2.843073 | 2.561982 | 2.558158 | 1.956332 | 0.836132 | 1.572609 | 1.749411 | 1.824225 | 1.448358 | 1.734397 | 3.236472 | 1.810876 | 2.288963 | 1.209227 | 1.686097 | 1.798121 | 1.5314 | 2.538975 | 1.639184 | 2.09336 | 2.544138 | 2.147405 | 2.407423 | 2.055573 | 27.4722222 | 0.753484 | -0.063705 |
| 27.3958333 | 3.023256 | 3.703734 | 3.148939 | 2.141795 | 1.174495 | 2.350941 | 2.177466 | 2.159736 | 1.676281 | 2.154232 | 4.05789 | 2.435033 | 2.976286 | 1.717433 | 2.233639 | 2.200033 | 1.762493 | 3.04396 | 2.262843 | 2.753226 | 2.75966 | 2.558977 | 2.996871 | 2.524336 | 27.4861111 | 0.701945 | 0.051126 |
| 27.4097222 | 3.093011 | 4.956887 | 3.603274 | 2.628187 | 1.562951 | 3.200821 | 2.507766 | 2.384621 | 1.734187 | 2.707578 | 4.635211 | 3.202638 | 3.634853 | 3.24899 | 3.133315 | 2.667978 | 1.94338 | 3.514998 | 3.049539 | 3.695257 | 3.160101 | 3.238774 | 3.249435 | 3.09906 | 27.5 | 0.249551 | 0.065719 |
| 27.4236111 | 2.969325 | 5.774215 | 4.013548 | 3.113501 | 1.576111 | 3.210288 | 2.775116 | 2.533523 | 2.001454 | 3.497337 | 5.44761 | 3.981831 | 4.309072 | 4.865467 | 3.948835 | 3.189648 | 2.375298 | 3.973407 | 3.610226 | 4.542671 | 4.450241 | 4.270014 | 3.588884 | 4.955548 | 27.5138889 | -0.610324 | -0.04037 |
| 27.4375 | 3.337711 | 5.701071 | 3.825606 | 3.022092 | 1.339709 | 2.445785 | 3.133573 | 2.032917 | 3.494015 | 5.124876 | 10.417644 | 4.896093 | 4.369185 | 4.598495 | 4.714576 | 3.483944 | 2.922834 | 4.094868 | 3.802928 | 4.751996 | 4.854424 | 5.823488 | 4.022032 | 13.207197 | 27.5277778 | -1.819769 | -0.459498 |
| 27.4513889 | 4.00902 | 5.761513 | 3.720417 | 1.755074 | 1.217674 | 1.617753 | 3.997372 | 1.04828 | 5.750199 | 8.161653 | 19.837258 | 5.107566 | 3.090021 | 2.193682 | 3.705413 | 3.000808 | 2.755975 | 3.129073 | 3.114569 | 4.153147 | 1.735718 | 7.530106 | 3.681407 | 22.089094 | 27.5416667 | -3.909783 | -1.191189 |
| 27.4652778 | 3.30609 | 3.347052 | 2.552537 | 0.36908 | 0.686944 | 1.051917 | 3.262132 | 0.392141 | 6.186285 | 8.046643 | 21.613749 | 2.610486 | 0.251804 | -0.287964 | 1.115411 | 1.313192 | 1.397689 | 1.191289 | 1.263052 | 1.673559 | -2.164796 | 6.363255 | 2.159123 | 17.353402 | 27.5555556 | -5.428989 | -1.431789 |
| 27.4791667 | 0.293047 | -2.077227 | -0.092433 | -0.026902 | 0.266399 | 0.640897 | 0.681131 | -0.180716 | 2.821711 | 3.602889 | 6.236649 | 0.006368 | -1.672745 | -2.086361 | -0.446305 | -0.056856 | -0.644867 | -1.248971 | -1.450596 | -1.273246 | -3.692998 | 2.195403 | 0.674112 | 5.060264 | 27.5694444 | -1.734708 | -2.074962 |
| 27.4930556 | -0.952193 | -1.993657 | -0.021555 | -0.149391 | 0.660814 | 0.500532 | -0.579003 | -0.188569 | 0.843304 | 0.625439 | -3.761195 | -0.240993 | -0.858703 | -0.829731 | 0.096286 | -0.171749 | -0.273052 | -0.389432 | -0.738386 | -0.955779 | -1.217816 | -0.166037 | 1.06215 | 2.044715 | 27.5833333 | 0.03121 | -0.612937 |
| 27.5069444 | -0.493509 | 1.44761 | 1.183352 | 1.497377 | 1.565187 | 1.900211 | 0.525032 | 0.75882 | 1.419704 | 1.546262 | -1.746605 | 1.593002 | 1.854894 | 2.870705 | 2.097484 | 1.718184 | 2.422783 | 4.266518 | 3.268544 | 3.38827 | 3.549609 | 1.796918 | 1.836605 | 6.523623 | 27.5972222 | -0.110099 | 0.366825 |
| 27.5208333 | 1.765847 | 3.458004 | 2.582647 | 2.792342 | 1.731159 | 2.821523 | 1.252179 | 1.198093 | 2.126071 | 2.648225 | 2.639638 | 3.396352 | 3.603365 | 4.720408 | 3.952317 | 2.098784 | 3.656199 | 5.595001 | 6.527402 | 5.769749 | 7.028818 | 4.515564 | 2.536014 | 9.874026 | 27.6111111 | -0.72153 | 0.648008 |
| 27.5347222 | 2.158873 | 4.173115 | 3.11707 | 3.642048 | 1.83059 | 2.843587 | 1.491841 | 0.702964 | 1.870833 | 2.745126 | 5.46207 | 2.766655 | 4.771167 | 4.084761 | 3.065691 | 1.586196 | 1.657425 | 2.698463 | 5.928611 | 5.864892 | 5.223907 | 4.652059 | 4.333458 | 8.246356 | 27.625 | -0.990018 | 0.457622 |
| 27.5486111 | 1.602786 | 3.023602 | 2.236402 | 2.605337 | 1.167973 | 2.40668 | 0.985112 | 0.309057 | 1.300735 | 1.988694 | 6.585565 | 1.779634 | 4.447179 | 1.123592 | 0.893121 | 1.57283 | 0.795029 | 1.144432 | 2.661799 | 3.145349 | 1.317588 | 3.356021 | 3.327852 | 3.510755 | 27.6388889 | -0.224882 | 0.170167 |
| 27.5625 | 1.119783 | 1.822034 | 2.987158 | 0.202803 | 0.193606 | 1.136772 | 0.410911 | 0.261055 | 0.426145 | 0.346727 | 4.572494 | 0.313252 | 1.782447 | 0.497003 | -0.454942 | -0.143646 | 0.515643 | -1.383022 | -0.933418 | -0.331696 | -1.395918 | 0.736777 | 0.665161 | -1.009975 | 27.6527778 | 0.527454 | -0.081327 |
| 27.5763889 | 0.463429 | 0.16392 | 0.983061 | -0.771449 | -0.150304 | 0.247021 | -0.036688 | 0.108039 | 0.361023 | 0.381499 | 0.961023 | -0.317252 | -0.199367 | 1.210872 | 0.675592 | -0.250821 | -0.898008 | -2.50377 | -1.143608 | -1.404876 | -0.10106 | -1.889936 | -1.474092 | -2.440233 | 27.6666667 | 0.697878 | -0.13931 |
| 27.5902778 | 0.084489 | -0.641634 | -0.620481 | -0.856281 | -0.398764 | -0.816967 | -0.37034 | -0.254422 | 0.082576 | -0.237271 | -1.912517 | -0.272718 | -1.179798 | 1.517114 | 1.151977 | 0.082921 | -0.372165 | -0.42051 | 1.47586 | 1.256772 | 1.835732 | -1.905096 | -1.668878 | -0.732676 | 27.6805556 | 0.331858 | -0.094189 |
| 27.6041667 | -0.83021 | -2.051286 | -0.97506 | -0.072403 | -0.384992 | -0.608068 | -0.28377 | -0.202271 | -0.054953 | -0.484556 | -3.829893 | 0.522155 | -0.905169 | 0.964516 | 1.008801 | 0.572124 | 0.551532 | 1.260669 | 2.780592 | 2.986705 | 2.708719 | 0.472876 | 0.454152 | 2.115337 | 27.6944444 | 0.044085 | 0.101732 |
| 27.6180556 | -1.226689 | -2.055807 | -0.514651 | 0.4841 | 0.048995 | 0.342852 | -0.340731 | -0.230397 | -0.007249 | -0.261833 | -3.352211 | 1.32245 | 2.053317 | 1.635083 | 1.582087 | 0.840265 | -0.281598 | 0.4932 | 2.10407 | 2.097437 | 0.865132 | 2.723118 | 1.458518 | 3.101537 | 27.7083333 | -0.025502 | 0.282208 |
| 27.6319444 | -0.104337 | 1.244509 | 1.19221 | 1.608856 | 0.3942 | 0.945553 | -0.171058 | 0.050658 | -0.261433 | -0.302784 | -0.700204 | 0.242992 | 2.103673 | -0.041753 | -0.503497 | 0.138717 | -0.225296 | -0.213908 | -0.446915 | 0.119515 | -1.520481 | 1.143232 | 0.665938 | 0.154221 | 27.7222222 | -0.043004 | 0.26693 |
| 27.6458333 | 1.258702 | 3.68893 | 2.284227 | 0.749435 | 0.524699 | 0.988845 | 0.359231 | 0.313289 | -0.276099 | -0.301969 | 2.269515 | -1.533655 | -0.354105 | -1.027579 | -1.708954 | -0.997694 | 0.113067 | -1.024232 | -3.035947 | -2.358734 | -1.459096 | -2.978681 | -2.109602 | -3.18765 | 27.7361111 | 0.126825 | 0.040686 |
| 27.6597222 | 1.965267 | 2.33266 | 0.21905 | -0.459754 | 0.356619 | 0.650909 | 0.54957 | 0.320411 | 0.034242 | -0.186759 | 3.346576 | -1.854764 | -1.404039 | -0.341211 | -0.729521 | -0.999014 | -0.637873 | -1.196704 | -1.96105 | -1.814084 | 0.911676 | -2.753251 | -3.257463 | -3.40231 | 27.75 | 0.349854 | -0.122753 |
| 27.6736111 | 1.270831 | 0.375104 | -0.847607 | -0.186117 | -0.038493 | 0.494921 | 0.470493 | 0.623495 | 0.238655 | 0.261281 | 2.667877 | -0.337948 | -1.986378 | 0.178089 | 0.230131 | -0.407481 | -0.07283 | -0.486907 | -0.230527 | -0.207953 | 1.201483 | 0.480861 | 0.233826 | -0.374951 | 27.7638889 | 0.565384 | -0.223807 |
| 27.6875 | 0.545682 | -0.061496 | 0.686747 | 0.832125 | -0.297065 | 0.149384 | 0.054999 | 0.124079 | -0.012399 | 0.23799 | 1.911978 | 1.0279 | 0.184741 | 0.558763 | 0.880107 | 0.211094 | 0.123973 | 0.930431 | 1.815654 | 1.568713 | 0.663343 | 1.434952 | 0.702922 | 2.380454 | 27.7777778 | 0.405614 | -0.219479 |
| 27.7013889 | 0.192653 | 0.54873 | 0.998808 | 0.989399 | -0.379689 | -0.36732 | -0.281257 | -0.007513 | -0.679203 | -0.318171 | 0.037223 | 0.973648 | 3.797009 | 0.560419 | 0.542257 | 0.539633 | 0.527718 | 1.434941 | 2.84694 | 2.663958 | -0.262078 | 2.251976 | -0.157267 | 1.93215 | 27.7916667 | 0.07137 | -0.120327 |
| 27.7152778 | -0.436733 | 0.039885 | 0.95679 | 0.038267 | -0.3253 | -0.490224 | -0.237999 | 0.048398 | -0.862152 | -0.653931 | -1.654993 | 0.103515 | 2.410735 | -0.083253 | -0.219685 | 0.617059 | 0.838908 | 0.931883 | 0.158339 | 0.449012 | -1.089381 | -0.361067 | -0.704161 | -0.574193 | 27.8055556 | -0.375902 | -0.086314 |
| 27.7291667 | -1.015373 | -1.124536 | 0.256719 | -1.197993 | -0.217861 | -0.675051 | -0.022999 | 0.03795 | -1.230342 | -1.571894 | -2.921272 | 0.244561 | -0.512396 | -0.769789 | -1.488492 | -0.878713 | 0.515999 | 0.40008 | -2.61729 | -2.063073 | -0.161949 | -0.531851 | -0.83926 | -1.883569 | 27.8194444 | -0.463389 | -0.092906 |
| 27.7430556 | -1.537297 | -2.58037 | -1.253362 | -1.279486 | -0.137477 | -0.535737 | -0.367038 | -0.36541 | -0.40942 | -1.246265 | -3.159365 | 0.401769 | -0.641389 | 0.033826 | -0.780591 | -1.500146 | -0.474505 | -0.681393 | -1.823877 | -2.18058 | 0.160824 | 0.508991 | 0.040504 | -0.772162 | 27.8333333 | -0.260122 | -0.056315 |
| 27.7569444 | -2.393782 | -3.206793 | -0.195971 | -0.413594 | -0.069348 | -0.324219 | -0.560372 | -0.447753 | 0.755251 | -0.123564 | -1.548562 | 0.025262 | 0.532776 | 0.15025 | 0.21839 | -0.33057 | -0.428551 | -0.879103 | 0.448922 | 0.12441 | 0.563885 | 0.310384 | 0.963986 | 0.197302 | 27.8472222 | -0.213854 | 0.012585 |
| 27.7708333 | -0.802057 | -1.084159 | -0.486605 | 0.043477 | -0.048223 | -0.148192 | -0.057955 | -0.26573 | 0.764936 | 0.477489 | 0.544567 | -1.186657 | -0.401128 | 0.288991 | -0.10746 | -0.801834 | 0.213915 | 0.504146 | 1.850139 | 2.221092 | 0.843364 | 0.547659 | 0.451799 | 0.292298 | 27.8611111 | -0.334897 | 0.041627 |
| 27.7847222 | 0.317309 | 0.697467 | 0.113395 | 0.069887 | 0.032017 | -0.131223 | 0.478467 | 0.095539 | 0.458141 | 0.771818 | 0.851475 | -0.882112 | -2.193362 | -0.483252 | -0.486728 | -0.670151 | 0.278099 | 1.054433 | -0.266674 | 0.846011 | 0.162682 | 1.123594 | -0.07444 | -0.073474 | 27.875 | -0.485916 | -0.032956 |
| 27.7986111 | 1.031459 | 1.683429 | 0.454658 | -0.471261 | 0.06007 | 0.454824 | 0.191262 | 0.240183 | -0.292799 | 0.472147 | 0.041581 | 0.431562 | -0.130756 | -1.424162 | -1.480823 | 0.163345 | 1.084271 | 1.542436 | -2.326288 | -2.072593 | -0.410204 | -0.412806 | -2.181449 | -1.080463 | 27.8888889 | -0.666681 | -0.169141 |
| 27.8125 | -0.060023 | 0.070289 | -0.965987 | -1.1728 | -0.10817 | 0.214342 | -0.424016 | 0.109492 | -1.300942 | -1.259352 | -1.20205 | 0.550793 | 2.505726 | -1.152119 | -0.844736 | 0.479568 | 0.984631 | 0.513282 | -1.98981 | -2.755511 | -0.897309 | -0.35026 | -4.064536 | -3.037704 | 27.9027778 | -1.01683 | -0.440109 |
| 27.8263889 | -1.578467 | -2.259952 | -1.592921 | -0.527828 | -0.407271 | -0.861337 | -1.265105 | -0.234062 | -1.898443 | -2.921452 | -1.009507 | -0.756115 | 0.534759 | -0.883052 | 0.052447 | 0.30888 | -0.296542 | -0.893694 | -1.379189 | -1.57497 | -0.532727 | -0.481014 | -2.152011 | -3.972012 | 27.9166667 | -1.13767 | -0.744605 |
| 27.8402778 | -2.286481 | -2.890894 | -0.734081 | -0.229703 | -0.761934 | -2.041736 | -1.5302 | -0.517089 | -0.505831 | -1.949571 | 0.371964 | -1.441031 | -2.23685 | -0.899771 | 0.448568 | -0.187358 | -1.953362 | -2.127647 | -0.634964 | -0.308211 | -0.647559 | 0.484549 | -0.968063 | -2.967608 | 27.9305556 | -0.859351 | -1.022973 |
| 27.8541667 | -1.595572 | -1.769774 | -0.233342 | -0.836707 | -1.05835 | -2.568948 | -1.098502 | -0.695106 | -0.359462 | -1.607322 | 0.634701 | -0.756977 | -3.181576 | -0.459188 | 0.585091 | -0.391405 | -1.104633 | -0.513266 | -0.875692 | -0.805668 | -1.182557 | 2.080522 | 0.127526 | -0.980148 | 27.9444444 | -0.773939 | -1.016184 |
| 27.8680556 | -1.038889 | -1.238285 | -0.958631 | -2.054521 | -0.887213 | -1.676273 | -0.482317 | -0.349479 | -0.49942 | -1.216602 | -1.53971 | -0.453267 | -0.373048 | 0.13394 | 0.526166 | 0.889112 | -0.16692 | -0.016844 | -3.0629 | -2.925925 | -1.826048 | 2.358674 | -1.832786 | -0.936494 | 27.9583333 | -1.174369 | -0.87411 |
| 27.8819444 | -1.323398 | -2.970943 | -3.444479 | -3.50308 | -0.679505 | -0.901272 | -0.604322 | -0.37373 | -1.206081 | -1.273745 | -4.74338 | -1.923383 | 0.004134 | -1.463147 | -1.905698 | -0.77196 | 0.076429 | -1.275803 | -5.105096 | -4.533091 | -3.690517 | 1.898557 | -2.565004 | -4.127031 | 27.9722222 | -1.410758 | -0.619384 |
| 27.8958333 | -2.027886 | -5.299709 | -5.376893 | -4.185249 | -0.832291 | -1.473395 | -1.669074 | -1.031645 | -2.415028 | -2.717606 | -7.294332 | -3.346343 | -2.11246 | -3.692673 | -3.631082 | -1.636136 | -1.274442 | -2.281005 | -4.899469 | -4.478811 | -3.662636 | -2.162019 | -2.282995 | -6.313006 | 27.9861111 | -1.138234 | -0.475535 |
| 27.9097222 | -3.209029 | -6.663159 | -5.44928 | -4.053035 | -1.291723 | -2.365627 | -2.958449 | -1.637199 | -3.105261 | -3.994373 | -7.276066 | -1.81494 | -3.176158 | -1.820926 | -2.469545 | -2.496317 | -2.581622 | -2.878965 | -2.422681 | -3.102329 | -3.23537 | -3.818278 | -2.035581 | -5.159818 | 28 | -0.833542 | -0.421818 |
| 27.9236111 | -4.114105 | -5.838968 | -3.936784 | -3.047651 | -1.882539 | -3.212905 | -3.258081 | -2.104396 | -2.974145 | -3.803072 | -6.203696 | -1.211233 | -2.588313 | 0.270613 | -0.317673 | -1.472502 | -2.516877 | -2.922778 | -1.819503 | -3.158009 | -2.580148 | -4.700744 | -1.463301 | -3.590799 | 28.0138889 | -0.196732 | -0.187964 |
| 27.9375 | -4.399803 | -3.088558 | -2.879302 | -2.498662 | -2.092835 | -3.669193 | -3.565762 | -2.039007 | -4.326907 | -5.726603 | -13.659913 | -2.275152 | -1.637135 | 0.058309 | -0.738191 | -1.944983 | -1.389782 | -0.998811 | -4.052295 | -5.247212 | -2.857957 | -3.69824 | -1.487439 | -4.828026 | 28.0277778 | 3.125445 | 0.723831 |
| 27.9513889 | -5.149132 | -1.552667 | -2.83654 | -1.836491 | -1.551395 | -2.173774 | -3.37918 | -1.078267 | -6.21164 | -8.058062 | -25.523251 | -3.457641 | -1.624182 | -1.936971 | -3.520788 | -2.784701 | -1.892303 | -0.610234 | -4.128149 | -5.862157 | -1.974325 | -4.927259 | -3.515696 | -9.776659 | 28.0416667 | 8.448738 | 1.16666 |
| 27.9652778 | -4.297529 | -1.189794 | -2.531607 | -1.219988 | -0.82618 | -0.385082 | -2.386169 | -0.265225 | -5.386466 | -6.8883 | -21.484845 | -2.587495 | -1.23612 | -2.465742 | -4.911603 | -3.567862 | -2.365474 | -0.459259 | -2.311067 | -3.669541 | -0.659595 | -4.430147 | -4.634072 | -10.295955 | 28.0555556 | 7.51164 | 0.186524 |
| 27.9791667 | -2.533103 | -1.36902 | -1.754413 | -0.858274 | -0.342625 | -0.457497 | -1.186562 | -0.296228 | -2.615232 | -3.089967 | -8.436158 | -1.614626 | -1.647489 | -2.781547 | -3.685637 | -3.439221 | -2.001382 | -0.752981 | -0.896452 | -1.553221 | -1.626401 | -2.19944 | -3.231518 | -7.010842 | 28.0694444 | 2.561201 | 0.251536 |
| 27.9930556 | -0.727151 | -0.876256 | -1.155551 | -0.54867 | -0.3499 | -0.794488 | -0.669824 | -0.425314 | -0.753515 | -0.84464 | -1.581439 | -0.839755 | -0.93751 | -2.618865 | -2.274368 | -1.437865 | -1.034712 | -0.656882 | -0.105653 | -0.238932 | -1.614001 | -0.739816 | -0.902468 | -2.352292 | 28.0833333 | -0.453099 | -0.498063 |
| 28.0069444 | 0.227871 | 0.003393 | -0.609809 | -0.13923 | -0.357422 | -0.633594 | -0.438489 | -0.272743 | -0.23067 | -0.339585 | -0.25791 | -1.093342 | -0.256063 | -1.736117 | -0.685292 | 0.807033 | -0.239221 | -0.386772 | 0.265811 | 0.316496 | -1.168565 | -0.558769 | 0.4633 | -0.380195 | 28.0972222 | -0.929779 | -0.569859 |
| 28.0208333 | 0.379718 | 0.337976 | -0.169722 | 0.226527 | -0.228939 | -0.230025 | -0.157357 | -0.074491 | -0.076858 | -0.21833 | -0.164061 | -1.36257 | -0.821491 | -1.79849 | -1.230414 | 0.463672 | -0.310714 | -0.703594 | -0.195817 | -0.225057 | -1.165775 | -1.442658 | -0.124217 | -0.70463 | 28.1111111 | -0.40196 | -0.352013 |
| 28.0347222 | 0.097469 | 0.010441 | 0.015112 | 0.199157 | -0.143747 | -0.01998 | -0.071618 | -0.083918 | -0.118337 | -0.269252 | -0.13277 | -1.909786 | -2.487914 | -2.553311 | -2.448303 | -1.433917 | -0.995725 | -1.60078 | -1.017039 | -1.424116 | -1.376242 | -2.436881 | -1.818062 | -2.230633 | 28.125 | 0.085367 | -0.134237 |
| 28.0486111 | -0.116991 | -0.291562 | -0.176542 | -0.152803 | -0.224508 | -0.184219 | -0.293968 | -0.345642 | -0.228852 | -0.464724 | -0.220256 | -2.324409 | -3.204079 | -3.067426 | -3.09724 | -2.798401 | -1.518499 | -2.227763 | -1.254729 | -1.962456 | -1.836358 | -2.518339 | -2.105414 | -2.895474 | 28.1388889 | 0.315732 | -0.015469 |
| 28.0625 | -0.294669 | -0.518774 | -0.53687 | -0.573505 | -0.351401 | -0.458669 | -0.527731 | -0.568785 | -0.299528 | -0.665784 | -0.283719 | -1.984598 | -2.435959 | -2.725611 | -2.39374 | -2.534458 | -1.53544 | -2.218936 | -1.115148 | -1.745542 | -2.054058 | -1.804891 | -1.153834 | -2.412989 | 28.1527778 | 0.464397 | 0.038074 |
| 28.0763889 | -0.463204 | -0.743733 | -0.710099 | -0.73086 | -0.418649 | -0.633086 | -0.647377 | -0.676584 | -0.334056 | -0.767934 | -0.277205 | -1.328267 | -1.514309 | -2.149717 | -1.441071 | -1.315187 | -1.279083 | -1.761913 | -0.987539 | -1.269922 | -1.479556 | -1.017513 | -0.596077 | -1.655721 | 28.1666667 | 0.566484 | 0.082865 |
| 28.0902778 | -0.607902 | -0.939025 | -0.737207 | -0.609573 | -0.400174 | -0.5098 | -0.644863 | -0.602112 | -0.406967 | -0.803754 | -0.504112 | -0.986277 | -1.137699 | -1.746619 | -1.145877 | -0.854282 | -0.902575 | -1.384729 | -0.955814 | -0.984044 | -0.772427 | -0.555484 | -0.605584 | -1.068048 | 28.1805556 | 0.416146 | 0.160685 |
| 28.1041667 | -0.79084 | -1.085149 | -0.837595 | -0.618147 | -0.387438 | -0.366486 | -0.653963 | -0.576471 | -0.780072 | -0.876053 | -0.884375 | -0.762434 | -0.479984 | -1.507715 | -1.353602 | -0.309866 | -0.577723 | -0.694538 | -0.790346 | -1.003731 | -0.815003 | -0.434992 | -0.838816 | -0.108772 | 28.1944444 | 0.17088 | 0.300771 |
| 28.1180556 | -1.096758 | -1.299524 | -1.109004 | -0.849228 | -0.381872 | -0.570631 | -0.759874 | -0.779421 | -0.902003 | -0.964615 | -1.126504 | -0.568726 | -0.692327 | -1.334595 | -1.192258 | -1.191833 | -0.531041 | -0.609925 | -0.58171 | -1.140435 | -1.070373 | -0.39242 | -0.967261 | -0.898468 | 28.2083333 | -0.055433 | 0.45132 |
| 28.1319444 | -1.454941 | -1.553292 | -1.475832 | -1.176884 | -0.512758 | -0.971431 | -0.893321 | -1.036875 | -0.93792 | -0.963102 | -1.191476 | -0.505421 | -0.382664 | -1.087022 | -0.777893 | -0.489775 | -0.617193 | -0.896506 | -0.600398 | -0.90082 | -0.658648 | -0.394665 | -0.720037 | -0.782066 | 28.2222222 | -0.101677 | 0.458458 |
| 28.1458333 | -1.593967 | -1.567002 | -1.593406 | -1.290278 | -0.609649 | -1.119693 | -0.888809 | -1.067528 | -0.849929 | -0.808372 | -1.173296 | -0.579791 | -0.308965 | -0.839217 | -0.649931 | -0.202119 | -0.591015 | -0.990443 | -0.765556 | -0.680738 | -0.27698 | -0.505358 | -0.33736 | -0.56873 | 28.2361111 | -0.000938 | 0.324477 |
| 28.1597222 | -1.250188 | -1.206618 | -1.217808 | -0.957077 | -0.501055 | -0.809584 | -0.612227 | -0.725751 | -0.605285 | -0.54556 | -0.974652 | -0.598811 | -0.333085 | -0.623122 | -0.830611 | -0.493683 | -0.433097 | -0.685552 | -0.694874 | -0.639947 | -0.446161 | -0.585221 | -0.317799 | -0.464677 | 28.25 | 0.195747 | 0.24066 |
| 28.1736111 | -0.661116 | -0.660277 | -0.748583 | -0.565348 | -0.37328 | -0.529517 | -0.306136 | -0.359584 | -0.261522 | -0.232651 | -0.426663 | -0.450996 | -0.263808 | -0.314462 | -0.661975 | -0.54331 | -0.30488 | -0.421599 | -0.362766 | -0.552197 | -0.554316 | -0.475047 | -0.456568 | -0.388043 | 28.2638889 | 0.205352 | 0.296859 |
| 28.1875 | -0.240276 | -0.285672 | -0.549007 | -0.471643 | -0.352168 | -0.46463 | -0.201444 | -0.201203 | -0.057352 | -0.042576 | 0.018102 | -0.373489 | -0.245474 | -0.079002 | -0.325102 | -0.273206 | -0.262005 | -0.394191 | -0.187986 | -0.396617 | -0.287772 | -0.390589 | -0.363747 | -0.314154 | 28.2777778 | 0.064421 | 0.447462 |
| 28.2013889 | -0.208718 | -0.243457 | -0.458841 | -0.460431 | -0.258468 | -0.381976 | -0.20487 | -0.175788 | -0.159333 | -0.156128 | -0.011764 | -0.534813 | -0.388865 | -0.068987 | -0.210755 | -0.142062 | -0.194972 | -0.365559 | -0.207847 | -0.339221 | -0.07844 | -0.505805 | -0.185135 | -0.294717 | 28.2916667 | 0.049232 | 0.59143 |
| 28.2152778 | -0.396802 | -0.362575 | -0.320325 | -0.326585 | -0.09033 | -0.205708 | -0.213561 | -0.147022 | -0.357653 | -0.309025 | -0.220263 | -0.707421 | -0.517646 | -0.077709 | -0.171164 | -0.139501 | -0.076324 | -0.124807 | -0.155883 | -0.24417 | -0.088623 | -0.57738 | -0.184818 | -0.283434 | 28.3055556 | 0.071468 | 0.692239 |
| 28.2291667 | -0.374913 | -0.323758 | -0.206664 | -0.214277 | -0.052291 | -0.167105 | -0.223571 | -0.085587 | -0.347968 | -0.207036 | -0.191673 | -0.648333 | -0.36726 | 0.035301 | 0.011025 | -0.058283 | -0.073315 | -0.0251 | 0.019787 | 0.012146 | -0.023045 | -0.508665 | -0.153419 | -0.235209 | 28.3194444 | 0.394155 | 0.680945 |
| 28.2430556 | -0.100212 | -0.119432 | -0.159737 | -0.209679 | -0.106716 | -0.209839 | -0.289553 | -0.097638 | -0.223439 | -0.101954 | -0.026021 | -0.435639 | -0.063396 | 0.074667 | 0.141564 | 0.073115 | -0.098383 | -0.138683 | 0.000733 | 0.171186 | 0.128325 | -0.4176 | 0.014492 | -0.123965 | 28.3333333 | 0.783533 | 0.570389 |
| 28.2569444 | 0.230711 | 0.043214 | -0.139733 | -0.224537 | -0.040703 | -0.060919 | -0.365175 | -0.128181 | -0.21752 | -0.128343 | 0.048366 | -0.243459 | 0.072624 | 0.011815 | 0.166559 | 0.17321 | 0.055445 | -0.05607 | -0.107552 | 0.161619 | 0.128606 | -0.228533 | 0.092487 | 0.068984 | 28.3472222 | 0.589245 | 0.475143 |
| 28.2708333 | 0.435303 | 0.061591 | -0.072596 | -0.177593 | 0.050055 | 0.134308 | -0.298098 | -0.085157 | -0.303526 | -0.167831 | 0.071409 | 0.0049 | 0.19115 | 0.175965 | 0.336619 | 0.230684 | 0.277743 | 0.297889 | 0.1037 | 0.340013 | 0.21941 | 0.190791 | 0.249719 | 0.330934 | 28.3611111 | 0.076637 | 0.420802 |
| 28.2847222 | 0.460903 | 0.047385 | 0.082517 | -0.011963 | 0.059178 | 0.108962 | -0.039271 | 0.067961 | -0.255238 | -0.117369 | 0.235484 | 0.37029 | 0.535208 | 0.580902 | 0.63109 | 0.352676 | 0.380581 | 0.498314 | 0.517914 | 0.813889 | 0.57787 | 0.494627 | 0.54496 | 0.626769 | 28.375 | -0.149323 | 0.349797 |
| 28.2986111 | 0.450353 | 0.176227 | 0.230984 | 0.152757 | 0.086644 | 0.087103 | 0.197043 | 0.180124 | -0.003621 | 0.035163 | 0.519494 | 0.68226 | 0.921082 | 0.798743 | 0.866388 | 0.683393 | 0.463479 | 0.521367 | 0.613652 | 1.183451 | 0.801472 | 0.382568 | 0.679537 | 0.913784 | 28.3888889 | 0.129032 | 0.310858 |
| 28.3125 | 0.533448 | 0.398945 | 0.306899 | 0.2592 | 0.275993 | 0.382519 | 0.326189 | 0.243446 | 0.293543 | 0.235257 | 0.661552 | 0.934417 | 1.174145 | 0.767335 | 1.019209 | 1.076722 | 0.691326 | 0.775678 | 0.568073 | 1.261646 | 0.985263 | 0.295836 | 0.712583 | 1.1629 | 28.4027778 | 0.639236 | 0.445254 |
| 28.3263889 | 0.761804 | 0.645322 | 0.448504 | 0.467924 | 0.51398 | 0.781684 | 0.48198 | 0.399381 | 0.526511 | 0.493586 | 0.762179 | 1.201175 | 1.405164 | 0.763896 | 1.089828 | 1.126891 | 0.840335 | 1.076171 | 0.755643 | 1.332297 | 1.201409 | 0.556994 | 0.93752 | 1.351712 | 28.4166667 | 0.993826 | 0.701171 |
| 28.3402778 | 1.077592 | 0.952846 | 0.774653 | 0.819864 | 0.619999 | 0.850346 | 0.749122 | 0.64199 | 0.773282 | 0.790849 | 1.155227 | 1.3628 | 1.6379 | 0.793601 | 1.17623 | 1.279703 | 0.796723 | 1.066619 | 0.858658 | 1.460165 | 1.165336 | 0.785658 | 1.214465 | 1.496904 | 28.4305556 | 0.968542 | 0.854615 |
| 28.3541667 | 1.478174 | 1.462141 | 1.255977 | 1.274313 | 0.743486 | 0.982769 | 1.026949 | 0.929768 | 1.083241 | 1.08737 | 1.714146 | 1.484475 | 1.965623 | 1.059837 | 1.609926 | 1.879023 | 0.859952 | 1.184793 | 0.880914 | 1.71814 | 1.327326 | 0.889188 | 1.443181 | 1.788864 | 28.4444444 | 0.542945 | 0.716149 |
| 28.3680556 | 2.175543 | 2.341523 | 1.869693 | 1.749828 | 1.061131 | 1.54611 | 1.340616 | 1.32698 | 1.496804 | 1.56002 | 2.341998 | 1.89917 | 2.509016 | 1.831629 | 2.366813 | 2.479507 | 1.279031 | 1.846341 | 1.156547 | 2.170257 | 2.24526 | 1.291386 | 1.875655 | 2.448014 | 28.4583333 | -0.082666 | 0.364514 |
| 28.3819444 | 3.159309 | 3.5548 | 2.6091 | 2.233475 | 1.484832 | 2.347374 | 1.809778 | 1.858985 | 1.826382 | 2.233797 | 3.227681 | 2.560486 | 3.132281 | 2.743749 | 2.971043 | 2.841143 | 1.829505 | 2.562293 | 1.707642 | 2.798061 | 3.499503 | 2.056661 | 2.590794 | 3.284096 | 28.4722222 | -0.143311 | 0.144639 |
| 28.3958333 | 4.023153 | 4.837069 | 3.261407 | 2.644489 | 1.705796 | 2.755519 | 2.284751 | 2.423001 | 2.047953 | 2.652634 | 4.492716 | 3.097659 | 3.748065 | 3.063064 | 3.38833 | 3.423364 | 2.149343 | 2.945547 | 2.129527 | 3.281295 | 3.998138 | 2.746758 | 3.059542 | 4.07974 | 28.4861111 | 0.389072 | 0.148909 |
| 28.4097222 | 4.451698 | 5.79901 | 3.915329 | 3.134571 | 1.811754 | 2.950142 | 2.951207 | 3.029205 | 2.299792 | 3.061255 | 5.941011 | 3.407443 | 4.413367 | 3.441327 | 3.972998 | 4.124477 | 2.390585 | 3.547559 | 2.656202 | 3.866835 | 4.524797 | 3.254804 | 3.18458 | 5.026021 | 28.5 | 1.034295 | 0.183702 |
| 28.4236111 | 4.258529 | 6.325139 | 4.782347 | 3.853483 | 1.607656 | 2.977632 | 2.952955 | 2.856895 | 2.597534 | 3.374422 | 8.764206 | 3.765136 | 5.049046 | 4.019756 | 4.573289 | 4.541785 | 2.754939 | 4.300683 | 3.324228 | 4.559514 | 5.292267 | 4.015301 | 3.656035 | 6.256024 | 28.5138889 | 0.876739 | 0.37917 |
| 28.4375 | 4.370008 | 7.182787 | 5.488602 | 3.138209 | 1.370354 | 2.555371 | 2.802195 | 2.147377 | 3.833182 | 4.49462 | 15.389797 | 4.323903 | 4.670251 | 4.738385 | 4.790669 | 3.723643 | 3.00533 | 4.006656 | 3.591425 | 5.057899 | 4.466416 | 5.77993 | 4.538354 | 10.231291 | 28.5277778 | -0.005185 | 1.385615 |
| 28.4513889 | 5.160317 | 8.076006 | 5.965646 | 2.000631 | 1.509854 | 2.193248 | 3.942815 | 1.480274 | 6.133516 | 6.881749 | 27.833662 | 5.036062 | 3.138993 | 5.205138 | 5.177662 | 2.522377 | 3.086399 | 2.465932 | 3.477672 | 5.894567 | 2.21994 | 9.16109 | 5.6167 | 16.961175 | 28.5416667 | -0.183329 | 3.512657 |
| 28.4652778 | 5.07342 | 6.592507 | 5.281669 | 1.759649 | 1.462884 | 2.673469 | 5.102133 | 1.179313 | 6.263511 | 7.790238 | 26.283615 | 3.810722 | 1.123418 | 3.247719 | 3.797546 | 1.295902 | 2.154259 | 0.745867 | 1.670551 | 3.346643 | -0.766781 | 9.277098 | 4.046886 | 14.017712 | 28.5555556 | -1.408622 | 0.81087 |
| 28.4791667 | 0.299482 | -0.017294 | 1.775663 | 1.335051 | 0.768372 | 1.78351 | 3.458232 | 0.734363 | 3.563983 | 4.543352 | 9.156427 | 1.661432 | -0.694762 | -1.528433 | -0.040676 | -0.16713 | -0.023781 | -1.233912 | -0.624683 | -0.161708 | -1.781311 | 3.784234 | 0.639832 | 3.877309 | 28.5694444 | -0.362387 | -0.419732 |
| 28.4930556 | -2.324051 | -2.432874 | -0.148004 | 1.452729 | 0.425655 | 0.743553 | 1.107755 | 0.19527 | 1.359439 | 1.482964 | -3.434747 | 1.052364 | 0.383656 | -1.192902 | -0.318471 | 0.553383 | -0.40739 | -0.637473 | -0.074256 | 1.167864 | -0.177849 | 0.451035 | 0.191787 | 1.468963 | 28.5833333 | -1.197965 | -0.326187 |
| 28.5069444 | -1.464373 | -1.060589 | 0.77462 | 2.169192 | 0.525841 | 1.304133 | 0.708181 | -0.045359 | 1.512192 | 1.006298 | -3.050901 | 2.698979 | 2.182906 | 1.700235 | 2.023909 | 1.860392 | 1.146722 | 2.071193 | 3.736473 | 5.024132 | 3.970011 | 1.578537 | 1.429487 | 7.004241 | 28.5972222 | -1.440581 | 0.030642 |
| 28.5208333 | 0.995438 | 0.765534 | 2.228899 | 2.914736 | 0.352754 | 1.508548 | 0.713034 | 0.257364 | 1.774297 | 1.74781 | 2.262591 | 4.182911 | 4.84978 | 4.918796 | 3.958475 | 1.794528 | 1.949573 | 3.154531 | 6.09429 | 6.38735 | 5.543808 | 4.083062 | 2.760988 | 11.189982 | 28.6111111 | -0.476564 | -0.021816 |
| 28.5347222 | 1.902265 | 2.990094 | 2.243559 | 2.487939 | 2.218652 | 1.525091 | 0.567164 | 0.789147 | 1.372005 | 1.487877 | 5.599399 | 3.44098 | 6.239916 | 5.909926 | 5.300328 | 2.225955 | 2.05734 | 3.19267 | 5.977553 | 5.01263 | 4.394393 | 5.077026 | 4.069252 | 9.641852 | 28.625 | 1.579377 | -0.250147 |
| 28.5486111 | 1.944835 | 4.584715 | 2.151557 | 2.047326 | 0.586859 | 1.250899 | 0.576383 | 0.66961 | 0.9092 | 1.588841 | 9.202328 | 2.229228 | 4.999271 | 2.72789 | 2.798449 | 1.716651 | 2.121853 | 3.145687 | 3.735893 | 2.54595 | 1.205616 | 5.369219 | 5.202018 | 4.003372 | 28.6388889 | 2.179347 | -0.418945 |
| 28.5625 | 1.72714 | 4.184555 | 1.884982 | 0.415783 | 0.351409 | 0.850256 | 0.303346 | -0.15635 | -0.482072 | 0.511865 | 6.448178 | 1.669534 | 3.118354 | 0.094121 | -0.311722 | 0.774406 | 1.337521 | 1.614096 | 0.14142 | -0.456057 | -1.062356 | 2.175096 | 2.180897 | -1.183936 | 28.6527778 | 1.252839 | -0.287244 |
| 28.5763889 | 1.46262 | 2.909219 | 1.69287 | -0.064212 | 0.048885 | 0.004584 | 0.105299 | 0.085588 | -1.168206 | -0.768706 | 1.692234 | 0.686259 | 1.190129 | -0.478792 | -0.766074 | 0.171126 | -1.369816 | -1.293297 | -2.217281 | -1.704476 | -0.641963 | -1.756603 | -1.115633 | -2.42893 | 28.6666667 | 0.547856 | -0.036156 |
| 28.5902778 | 1.003615 | 0.516443 | 0.822251 | -0.608354 | -0.126917 | -0.513664 | 0.231497 | -0.083043 | -0.306451 | 0.509641 | -1.080637 | 0.135981 | 0.570741 | -0.220345 | -0.732382 | -0.620831 | -1.028194 | -0.263239 | -0.178513 | 0.134897 | 1.511355 | -2.619751 | -1.613902 | 0.127944 | 28.6805556 | -0.160801 | 0.173558 |
| 28.6041667 | -0.165674 | -1.331545 | -0.168978 | -0.004819 | -0.029372 | -0.516883 | -0.142776 | -0.580068 | -0.110687 | 0.442468 | -1.986813 | 0.138957 | -0.285878 | 1.14168 | 0.738491 | -0.649433 | 1.015954 | 2.208319 | 2.745174 | 3.438556 | 2.968312 | -0.66266 | -0.660811 | 3.496348 | 28.6944444 | -0.409414 | 0.125665 |
| 28.6180556 | -0.902527 | -1.387952 | -0.05034 | 0.950006 | 0.397015 | 0.324017 | -0.744789 | -0.535207 | 0.543223 | 0.749613 | -1.354035 | 0.586061 | 0.99489 | 2.603762 | 2.582733 | 0.571211 | 0.896564 | 2.039395 | 2.487832 | 2.697626 | 1.041235 | 0.546624 | 1.041547 | 3.963473 | 28.7083333 | -0.244382 | 0.011168 |
| 28.6319444 | -0.725155 | 0.769083 | 0.648682 | 0.394456 | 0.597762 | 1.174723 | -0.644142 | -0.03041 | 0.683317 | 0.577991 | 0.570093 | 0.589054 | 1.066235 | 0.965413 | 0.702424 | 0.366499 | 0.308027 | 0.391543 | 0.719034 | -0.642939 | -3.018492 | 2.376134 | 1.113143 | -0.270758 | 28.7222222 | 0.292621 | -0.06938 |
| 28.6458333 | -0.347311 | 2.298477 | 1.564688 | -0.039131 | 0.280424 | 1.032554 | -0.302831 | 0.46629 | 0.393423 | -0.624606 | 2.095789 | 0.082583 | -0.027848 | -1.023826 | -1.622701 | -0.579312 | -0.909505 | -1.770513 | -1.660439 | -2.920954 | -2.975692 | 2.179824 | -0.015729 | -4.541598 | 28.7361111 | 0.697986 | -0.19795 |
| 28.6597222 | 0.431042 | 2.189346 | 1.881577 | 0.109923 | 0.114332 | 0.114272 | 0.333008 | 0.640883 | -0.338028 | -1.621939 | 2.184501 | -0.114889 | -0.532464 | -1.137352 | -0.832332 | 0.27209 | -1.395334 | -2.157437 | -2.212488 | -0.988185 | 0.664369 | -3.526957 | -0.538884 | -4.523983 | 28.75 | 0.50999 | -0.220381 |
| 28.6736111 | 1.41142 | 0.104215 | 0.481305 | -0.342565 | -0.011319 | -0.324387 | 0.785945 | 0.852961 | -0.494683 | -1.327395 | 1.311107 | -0.051103 | -1.475953 | -0.493217 | -0.373778 | -0.410884 | -0.32206 | -0.646263 | -1.41353 | 0.649378 | 2.692712 | -4.307042 | -0.871398 | -1.524106 | 28.7638889 | 0.169129 | -0.082375 |
| 28.6875 | 0.431255 | -2.649111 | -1.371675 | 0.029702 | -0.137172 | -0.394059 | 0.710842 | 0.676504 | -0.331413 | -0.09553 | -0.560959 | 0.288709 | -0.490423 | -0.033239 | 0.621706 | -0.514244 | 0.312029 | 0.891824 | 1.163173 | 2.196651 | 2.432079 | -0.329714 | -1.059445 | 1.449769 | 28.7777778 | -0.930016 | 0.081708 |
| 28.7013889 | -0.656889 | -2.465588 | -1.494139 | 0.116531 | -0.196436 | 0.129609 | 0.321172 | 0.434582 | -0.476832 | 0.132337 | -2.116993 | 0.542181 | 1.198534 | 0.688637 | 1.377387 | 0.167303 | 0.447886 | 1.478945 | 2.238844 | 1.476078 | 1.151627 | 2.293459 | -0.070932 | 3.027656 | 28.7916667 | -2.333166 | 0.124221 |
| 28.7152778 | -0.875849 | -0.994838 | -0.0189 | 0.335024 | -0.203544 | -0.068982 | -0.19507 | -0.1311 | -0.457556 | 0.529166 | -2.178495 | 0.03719 | 1.804722 | 1.457232 | 1.773273 | 0.975513 | 1.024426 | 0.905384 | 0.189761 | -1.8021 | -1.724281 | 4.384569 | 1.573206 | 1.035749 | 28.8055556 | -1.860979 | 0.022509 |
| 28.7291667 | -0.755167 | 0.867222 | 0.468655 | -0.221012 | 0.238458 | 0.230072 | -0.291809 | -0.511647 | -0.268711 | 0.31486 | -1.171894 | -0.025828 | -1.12291 | 1.510463 | 0.333132 | -0.286401 | 0.320994 | -0.371946 | -1.010814 | -2.970188 | -2.288468 | 0.775974 | 2.171622 | -2.544957 | 28.8194444 | 0.010394 | -0.145549 |
| 28.7430556 | -0.337799 | 1.792359 | 0.415545 | -0.299902 | 0.345654 | -0.207009 | -0.307347 | -0.020206 | -0.31899 | -0.143214 | 0.468881 | -0.076549 | -1.118681 | -0.199688 | -0.48572 | 0.194498 | -1.157042 | -1.915825 | -2.055713 | -1.573458 | -1.398991 | -1.841529 | -0.99107 | -3.507154 | 28.8333333 | 1.221217 | -0.285856 |
| 28.7569444 | -0.21734 | 1.622748 | 1.062767 | -1.198806 | 0.300782 | -0.129311 | -0.393307 | -0.093682 | 0.174873 | 0.101128 | 1.267645 | 0.579078 | 2.297007 | -0.775197 | -1.01483 | -1.278332 | -1.313785 | -1.383949 | 0.4884 | 1.798753 | 0.41222 | -1.144296 | -3.919374 | -1.158792 | 28.8472222 | 0.723936 | -0.408471 |
| 28.7708333 | 0.368309 | -0.366171 | -1.053535 | -1.675269 | 0.250891 | 0.037205 | 0.043644 | 0.392138 | 0.249609 | 0.039507 | 1.303077 | 0.405838 | 1.344806 | -0.524529 | -0.696106 | -1.170919 | 0.322802 | 0.501763 | 2.401883 | 3.152516 | 1.884345 | 0.548018 | -1.072169 | 2.470918 | 28.8611111 | 0.161713 | -0.423157 |
| 28.7847222 | 0.267844 | -1.68165 | -0.92269 | -0.510946 | 0.032259 | -0.107139 | 0.323221 | 0.455499 | -0.324954 | -0.631305 | 0.637009 | -0.023152 | -1.350315 | -0.077143 | 0.716834 | 0.064696 | 1.292353 | 1.664578 | 0.056322 | 0.117275 | 0.642027 | 3.846906 | 3.148863 | 3.702767 | 28.875 | -0.258449 | -0.336993 |
| 28.7986111 | -1.603643 | -3.883231 | -1.326193 | -0.352667 | -0.14554 | -0.237024 | 0.43101 | 0.670013 | -1.218102 | -1.452152 | -1.750498 | -0.357497 | -1.798202 | 0.27011 | 1.518516 | 1.378353 | 1.32988 | 2.022677 | -1.20825 | -2.181703 | -2.282128 | 1.767965 | 3.006533 | -0.492714 | 28.8888889 | -0.447364 | -0.285421 |
| 28.8125 | -2.399536 | -2.675114 | -0.391532 | 0.133731 | -0.258514 | -0.462788 | -0.084067 | -0.069128 | -1.505125 | -1.686493 | -3.437779 | -1.109443 | -1.775867 | -0.189945 | -0.75085 | -0.450318 | -1.133024 | -1.382583 | -1.372207 | -2.622737 | -2.248869 | -2.808882 | -1.374435 | -5.603937 | 28.9027778 | -0.382647 | -0.351798 |
| 28.8263889 | -1.877019 | -0.600076 | -0.400552 | -0.118936 | -0.365377 | -0.672411 | -0.718037 | -0.537901 | -1.211688 | -1.456435 | -3.444016 | -1.079879 | -0.26482 | -0.83547 | -2.414984 | -1.831586 | -2.685276 | -3.779469 | -1.414962 | -1.469616 | -1.017929 | -0.862975 | -3.014736 | -6.227593 | 28.9166667 | -0.4688 | -0.517677 |
| 28.8402778 | -1.660334 | -0.12398 | -0.666857 | -0.27324 | -0.307324 | -0.451954 | -1.104531 | -1.108555 | -0.291321 | -0.735854 | -1.600617 | -0.386749 | 0.154171 | -1.427633 | -1.587985 | -0.885393 | -0.67498 | -1.598427 | -1.282518 | -0.844737 | 0.530545 | 1.355493 | -1.297011 | -2.27281 | 28.9305556 | -1.089485 | -0.786996 |
| 28.8541667 | -1.180672 | -1.370429 | -0.75849 | -0.47109 | -0.388438 | -0.778605 | -1.532285 | -1.661911 | 0.25792 | -0.173751 | -0.318913 | 0.167854 | -0.427225 | -1.186352 | -0.824949 | -0.722637 | 1.302478 | 0.316401 | -1.212775 | -1.246025 | 0.212617 | 3.861661 | 1.090882 | 0.407228 | 28.9444444 | -1.625027 | -0.956767 |
| 28.8680556 | -1.758748 | -4.651826 | -1.075341 | -0.769209 | -0.569752 | -1.08526 | -1.693654 | -1.206712 | -0.570884 | -0.769605 | -1.389005 | -0.332952 | -1.117385 | -0.789415 | 0.589192 | -0.644569 | 0.548924 | 0.577817 | -1.807445 | -3.335121 | -2.772891 | 7.889717 | 1.454726 | -1.731227 | 28.9583333 | -1.934816 | -0.937022 |
| 28.8819444 | -3.320777 | -7.049083 | -1.525039 | -1.98282 | -0.889535 | -1.585678 | -1.320934 | -0.987847 | -2.148187 | -1.813677 | -3.530295 | -2.95931 | -2.771889 | -1.990073 | -0.674674 | -1.306238 | -0.973503 | -0.742309 | -2.020998 | -4.134081 | -4.5339 | 6.124407 | 2.457518 | -5.771649 | 28.9722222 | -2.099446 | -0.781687 |
| 28.8958333 | -3.884864 | -7.36277 | -4.182769 | -4.456663 | -1.669126 | -2.576772 | -1.617647 | -0.965182 | -3.157722 | -3.582303 | -6.277485 | -4.357044 | -4.130766 | -4.455228 | -3.661029 | -2.369272 | -2.216689 | -2.195317 | -2.816478 | -4.846873 | -3.476154 | 3.509332 | 0.514119 | -7.746636 | 28.9861111 | -2.226414 | -0.587663 |
| 28.9097222 | -3.210336 | -5.994213 | -5.336451 | -4.627338 | -2.627822 | -3.394844 | -1.553845 | -1.146523 | -2.82826 | -3.041335 | -6.807349 | -2.823539 | -3.843875 | -5.316957 | -4.395242 | -2.567637 | -1.724043 | -2.913668 | -4.187773 | -4.864001 | -0.864585 | 0.946025 | -2.518571 | -6.478462 | 29 | -1.804195 | -0.356531 |
| 28.9236111 | -1.676885 | -3.210875 | -4.935271 | -3.693194 | -2.910738 | -3.240652 | -1.956906 | -1.941046 | -2.51481 | -2.599775 | -6.306192 | -1.494145 | -2.384612 | -3.355449 | -3.672937 | -3.225403 | -1.003152 | -2.76723 | -3.377632 | -3.694385 | -1.49216 | -1.31811 | -3.384629 | -3.637358 | 29.0138889 | -0.675148 | 0.014416 |
| 28.9375 | -2.236618 | -2.02612 | -4.806421 | -3.334467 | -2.589018 | -2.95558 | -3.676762 | -3.169199 | -3.645312 | -3.772521 | -15.260398 | -2.886296 | -0.833412 | -2.579447 | -2.804001 | -2.881068 | -2.334714 | -2.535151 | -3.718162 | -3.965858 | -3.84517 | -1.488416 | -2.661086 | -3.66447 | 29.0277778 | 1.945917 | 0.644272 |
| 28.9513889 | -3.764426 | -2.006117 | -3.915563 | -2.573435 | -2.265082 | -2.49448 | -4.964751 | -2.909045 | -5.721597 | -6.307392 | -31.143158 | -4.742565 | -1.139481 | -4.750742 | -4.269067 | -3.33396 | -3.387609 | -1.980967 | -4.705514 | -4.822012 | -3.244335 | -1.949066 | -4.103638 | -7.498932 | 29.0416667 | 4.469564 | 0.516283 |
| 28.9652778 | -3.797758 | -2.151443 | -2.834359 | -1.563212 | -1.658702 | -1.502564 | -3.871543 | -1.882877 | -4.986271 | -5.693372 | -27.263435 | -4.028561 | -1.389365 | -3.795561 | -5.001643 | -3.403274 | -2.73725 | -0.961596 | -3.086779 | -4.671715 | -1.386125 | -1.754952 | -3.148109 | -9.793812 | 29.0555556 | 3.704758 | 0.564609 |
| 28.9791667 | -1.901906 | -2.212679 | -1.838483 | -1.200209 | -0.954977 | -1.074005 | -1.791194 | -1.096147 | -2.638762 | -2.80555 | -11.604296 | -2.594982 | -1.811496 | -2.288659 | -4.188949 | -3.68042 | -1.470388 | -0.375151 | -2.14064 | -3.532049 | -0.909099 | 0.003182 | -1.168674 | -5.376783 | 29.0694444 | 1.28388 | 0.290092 |
| 28.9930556 | -0.647394 | -1.122172 | -0.965443 | -0.808795 | -0.555178 | -0.832544 | -0.557378 | -0.459614 | -1.106083 | -1.20474 | -2.581673 | -1.561475 | -1.178203 | -1.038076 | -2.650107 | -1.794719 | -0.711142 | -0.016979 | -1.66069 | -1.741366 | -0.140272 | 1.287965 | 0.534426 | -2.19479 | 29.0833333 | -0.761774 | -0.539229 |
| 29.0069444 | -0.326177 | -0.454615 | -0.491125 | -0.351834 | -0.110905 | -0.304065 | -0.087786 | 0.035221 | -0.59011 | -0.680236 | -0.709889 | -1.373217 | -1.229174 | -0.64663 | -0.315374 | 1.200054 | 0.069784 | 0.430793 | 0.146956 | -0.340637 | 0.055433 | 0.798936 | 1.002432 | -0.997011 | 29.0972222 | -1.169651 | -0.599525 |
| 29.0208333 | -0.298317 | -0.261326 | -0.21397 | 0.21072 | 0.229108 | 0.1322 | 0.094611 | 0.173262 | -0.267306 | -0.315553 | -0.532847 | -2.000831 | -2.33912 | -1.316561 | 0.085183 | 0.71081 | 0.007444 | -0.195585 | 0.560141 | -0.465955 | -0.80373 | -0.792576 | 0.111299 | -1.506305 | 29.1111111 | -1.085496 | -0.332133 |
| 29.0347222 | -0.507679 | -0.542778 | -0.233091 | 0.18265 | 0.154658 | 0.014517 | -0.086301 | -0.025409 | -0.145681 | -0.198304 | -0.580213 | -2.357314 | -3.590833 | -3.16096 | -1.855324 | -1.420844 | -0.761536 | -1.366401 | -0.903064 | -1.454213 | -1.8319 | -2.117219 | -0.663652 | -2.793289 | 29.125 | -0.694356 | -0.105968 |
| 29.0486111 | -0.898035 | -1.122423 | -0.432434 | -0.206533 | -0.173298 | -0.350099 | -0.382814 | -0.284973 | -0.221036 | -0.391943 | -0.690635 | -2.667346 | -3.883558 | -4.369793 | -3.319804 | -2.451361 | -1.303146 | -2.165728 | -1.88032 | -1.941935 | -2.061046 | -2.240604 | -0.971817 | -3.415894 | 29.1388889 | -0.197015 | -0.024046 |
| 29.0625 | -1.164902 | -1.61011 | -0.675503 | -0.582822 | -0.377112 | -0.560928 | -0.482055 | -0.410124 | -0.323812 | -0.609504 | -0.761853 | -2.597273 | -3.004422 | -3.655603 | -2.976934 | -2.100934 | -1.244382 | -1.965092 | -2.04972 | -1.648207 | -1.575847 | -1.463988 | -1.177362 | -2.892982 | 29.1527778 | 0.290248 | -0.061531 |
| 29.0763889 | -1.17804 | -1.684013 | -0.770349 | -0.722754 | -0.387182 | -0.506797 | -0.424192 | -0.479138 | -0.478418 | -0.679926 | -0.958118 | -2.033878 | -2.064136 | -2.227921 | -1.40597 | -1.296726 | -1.023346 | -1.205319 | -1.201408 | -1.200489 | -1.256346 | -0.856765 | -1.13476 | -1.992081 | 29.1666667 | 0.746523 | -0.09586 |
| 29.0902778 | -1.155572 | -1.600172 | -0.72686 | -0.635621 | -0.362758 | -0.379682 | -0.550712 | -0.623832 | -0.690652 | -0.725214 | -1.325181 | -1.507093 | -1.515452 | -1.453336 | -0.485863 | -0.954533 | -0.874985 | -0.74041 | -0.810976 | -1.136645 | -1.361214 | -0.596257 | -0.765407 | -1.459883 | 29.1805556 | 1.071488 | 0.05282 |
| 29.1041667 | -1.243752 | -1.636047 | -0.701459 | -0.564496 | -0.446774 | -0.50441 | -0.863204 | -0.853515 | -0.781861 | -0.799119 | -1.552256 | -1.219821 | -1.214503 | -1.53294 | -0.662389 | -0.979675 | -0.673032 | -0.977947 | -0.681202 | -1.20764 | -1.22202 | -0.410465 | -0.920664 | -1.418018 | 29.1944444 | 0.772601 | 0.374155 |
| 29.1180556 | -1.54062 | -1.929131 | -0.879883 | -0.814725 | -0.628992 | -0.846537 | -1.093385 | -1.106595 | -0.758957 | -0.923913 | -1.725534 | -1.048769 | -0.946073 | -1.708532 | -1.08176 | -1.022394 | -0.588346 | -1.292611 | -0.530741 | -1.050527 | -0.670232 | -0.391704 | -1.402748 | -1.420865 | 29.2083333 | 0.061314 | 0.676979 |
| 29.1319444 | -2.014012 | -2.319899 | -1.17459 | -1.119637 | -0.764 | -1.149017 | -1.096453 | -1.197858 | -0.771102 | -1.003087 | -2.05856 | -0.750527 | -0.481722 | -1.220519 | -0.928588 | -0.779573 | -0.65853 | -1.041966 | -0.576038 | -0.820667 | -0.530692 | -0.526449 | -1.252013 | -0.928622 | 29.2222222 | -0.281304 | 0.835046 |
| 29.1458333 | -2.441222 | -2.543893 | -1.318852 | -1.054109 | -0.668928 | -0.89515 | -0.952592 | -1.092507 | -0.831773 | -0.928669 | -2.290491 | -0.489426 | -0.157597 | -0.555926 | -0.376099 | -0.300713 | -0.680789 | -0.596006 | -0.516859 | -0.777224 | -0.861618 | -0.603486 | -0.59753 | -0.361869 | 29.2361111 | 0.028746 | 0.759068 |
| 29.1597222 | -2.436502 | -2.309925 | -1.162048 | -0.71538 | -0.576565 | -0.787028 | -0.717139 | -0.914031 | -0.785587 | -0.686279 | -2.005023 | -0.4737 | -0.222499 | -0.181051 | -0.131235 | -0.225466 | -0.503389 | -0.589415 | -0.437994 | -0.668847 | -0.752057 | -0.513655 | -0.301325 | -0.245148 | 29.25 | 0.485101 | 0.505606 |
| 29.1736111 | -2.035662 | -1.845681 | -0.815394 | -0.354439 | -0.506303 | -0.744948 | -0.631719 | -0.650235 | -0.570489 | -0.338716 | -1.338132 | -0.416626 | -0.290139 | -0.071368 | -0.302166 | -0.414612 | -0.287013 | -0.636943 | -0.383606 | -0.389929 | -0.211907 | -0.400469 | -0.423734 | -0.318913 | 29.2638889 | 0.704268 | 0.353601 |
| 29.1875 | -1.452559 | -1.308876 | -0.543439 | -0.161577 | -0.360783 | -0.592671 | -0.381971 | -0.343049 | -0.303576 | -0.119267 | -0.72028 | -0.232433 | -0.186024 | -0.020386 | -0.394385 | -0.41795 | -0.249579 | -0.562166 | -0.536463 | -0.270869 | 0.051204 | -0.448886 | -0.256184 | -0.287061 | 29.2777778 | 0.64442 | 0.387449 |
| 29.2013889 | -0.95528 | -0.93411 | -0.412442 | -0.168807 | -0.148165 | -0.274842 | -0.185403 | -0.150182 | -0.199899 | -0.128978 | -0.297706 | -0.19527 | -0.091757 | 0.098729 | -0.093457 | -0.10839 | -0.305286 | -0.393894 | -0.411182 | -0.356388 | -0.191195 | -0.532737 | -0.152358 | -0.258289 | 29.2916667 | 0.366962 | 0.453641 |
| 29.2152778 | -0.716338 | -0.686251 | -0.38071 | -0.264219 | -0.136391 | -0.198766 | -0.229733 | -0.1539 | -0.24285 | -0.222205 | -0.242584 | -0.317821 | -0.068256 | 0.235472 | 0.174254 | 0.109999 | -0.191582 | -0.19285 | -0.268876 | -0.320916 | 0.095306 | -0.464954 | -0.12775 | -0.219076 | 29.3055556 | 0.085122 | 0.428193 |
| 29.2291667 | -0.633948 | -0.557628 | -0.338111 | -0.277206 | -0.265814 | -0.327862 | -0.357396 | -0.216621 | -0.268407 | -0.20945 | -0.365598 | -0.330343 | -0.032427 | 0.281905 | 0.124647 | 0.100194 | 0.017321 | -0.040143 | -0.235088 | -0.064446 | 0.562882 | -0.302902 | 0.184938 | -0.057674 | 29.3194444 | -0.043827 | 0.501556 |
| 29.2430556 | -0.517179 | -0.524104 | -0.29571 | -0.247817 | -0.298313 | -0.39876 | -0.356287 | -0.231628 | -0.220469 | -0.116262 | -0.199262 | -0.091249 | 0.174979 | 0.316741 | 0.208391 | 0.31443 | 0.128512 | 0.038624 | -0.144606 | 0.279583 | 0.406271 | -0.140369 | 0.410195 | 0.232288 | 29.3333333 | 0.045376 | 0.592177 |
| 29.2569444 | -0.297459 | -0.428417 | -0.264644 | -0.317031 | -0.079793 | -0.080913 | -0.354868 | -0.186509 | -0.148066 | -0.066876 | 0.122708 | 0.412426 | 0.596604 | 0.449122 | 0.486416 | 0.632663 | 0.209122 | 0.139924 | 0.172296 | 0.496028 | 0.249042 | 0.059756 | 0.188988 | 0.536562 | 29.3472222 | 0.568591 | 0.580723 |
| 29.2708333 | 0.173002 | -0.018189 | -0.070393 | -0.310201 | 0.087407 | 0.132155 | -0.273735 | -0.013748 | -0.106421 | -0.091144 | 0.237443 | 0.9565 | 0.962757 | 0.553509 | 0.635987 | 0.648771 | 0.316593 | 0.189318 | 0.276684 | 0.60158 | 0.672494 | 0.246061 | 0.103343 | 0.81832 | 29.3611111 | 1.242443 | 0.486335 |
| 29.2847222 | 0.6926 | 0.3711 | 0.204826 | -0.110478 | 0.03117 | 0.022313 | -0.095461 | 0.148993 | -0.095739 | -0.150581 | 0.150185 | 1.131676 | 1.142024 | 0.550289 | 0.626939 | 0.607939 | 0.370476 | 0.213334 | 0.175296 | 0.648644 | 0.830848 | 0.266103 | 0.523192 | 0.931524 | 29.375 | 1.613087 | 0.342872 |
| 29.2986111 | 0.736974 | 0.288951 | 0.335803 | 0.061443 | -0.039462 | -0.000648 | -0.023197 | 0.117318 | -0.066312 | -0.134723 | 0.188204 | 0.984694 | 1.2097 | 0.758144 | 0.792226 | 0.855954 | 0.436526 | 0.404076 | 0.201681 | 0.757432 | 0.632873 | 0.251573 | 1.028029 | 1.008192 | 29.3888889 | 1.386518 | 0.230652 |
| 29.3125 | 0.634635 | 0.146594 | 0.368751 | 0.164065 | 0.167477 | 0.295673 | 0.037272 | 0.092156 | 0.088707 | 0.108062 | 0.662119 | 0.947903 | 1.3907 | 1.25083 | 1.22126 | 1.344742 | 0.705612 | 0.796814 | 0.542448 | 1.06235 | 0.89066 | 0.439745 | 1.277134 | 1.273998 | 29.4027778 | 0.599679 | 0.210788 |
| 29.3263889 | 0.871148 | 0.471137 | 0.587542 | 0.420828 | 0.605707 | 0.853079 | 0.286336 | 0.330492 | 0.359555 | 0.426447 | 1.322186 | 1.183057 | 1.66592 | 1.570399 | 1.641096 | 1.827054 | 1.02151 | 1.12175 | 0.6908 | 1.297442 | 1.382699 | 0.643702 | 1.555491 | 1.475693 | 29.4166667 | -0.442962 | 0.2965 |
| 29.3402778 | 1.331294 | 1.001553 | 0.913092 | 0.814461 | 0.847817 | 1.07684 | 0.704495 | 0.708297 | 0.660936 | 0.651361 | 1.819738 | 1.362881 | 1.827119 | 1.535411 | 1.856985 | 2.089653 | 1.142025 | 1.243858 | 0.752341 | 1.410238 | 1.499214 | 0.762768 | 1.946244 | 1.635559 | 29.4305556 | -0.505065 | 0.317984 |
| 29.3541667 | 1.782737 | 1.505329 | 1.199003 | 1.183554 | 0.922392 | 1.213734 | 1.134698 | 1.098689 | 0.863043 | 0.843919 | 2.114141 | 1.582786 | 2.056678 | 1.619842 | 1.948608 | 2.254024 | 1.108894 | 1.377411 | 1.021611 | 1.688491 | 1.730286 | 1.029582 | 2.250225 | 2.108597 | 29.4444444 | 0.523612 | 0.183134 |
| 29.3680556 | 2.393915 | 2.289119 | 1.553351 | 1.698556 | 1.164516 | 1.609729 | 1.530966 | 1.582467 | 1.0504 | 1.14086 | 2.502252 | 2.018411 | 2.670956 | 2.181493 | 2.41588 | 2.737409 | 1.347102 | 1.768288 | 1.596019 | 2.233473 | 2.36867 | 1.617429 | 2.88859 | 2.911005 | 29.4583333 | 1.218263 | -0.072732 |
| 29.3819444 | 3.229204 | 3.564825 | 2.156524 | 2.669582 | 1.716079 | 2.416921 | 1.998158 | 2.264708 | 1.426899 | 1.663773 | 3.524014 | 2.49413 | 3.536733 | 3.051249 | 3.508483 | 3.680048 | 1.832455 | 2.360193 | 2.249631 | 2.91255 | 3.014943 | 2.494 | 3.813377 | 3.922106 | 29.4722222 | 1.12085 | -0.190301 |
| 29.3958333 | 4.62769 | 5.33039 | 2.798895 | 3.688402 | 2.023295 | 2.992921 | 2.71574 | 2.914446 | 1.943806 | 2.311806 | 5.509121 | 2.984259 | 4.295827 | 3.995267 | 4.763649 | 4.408429 | 2.319062 | 2.994345 | 2.805883 | 3.692892 | 3.600831 | 3.532359 | 4.22909 | 5.139045 | 29.4861111 | 1.019131 | -0.161331 |
| 29.4097222 | 5.343819 | 7.040125 | 3.628722 | 4.548824 | 1.971107 | 3.06988 | 3.083944 | 3.157044 | 2.280009 | 2.800569 | 8.193878 | 3.66437 | 5.64483 | 5.095475 | 5.488018 | 5.104199 | 2.745931 | 3.703298 | 3.253005 | 4.949241 | 4.524591 | 4.467144 | 4.261237 | 6.625778 | 29.5 | 1.188042 | -0.044353 |
| 29.4236111 | 4.812034 | 8.109117 | 4.48695 | 4.978274 | 1.873806 | 3.118228 | 3.072741 | 3.142755 | 2.669254 | 3.527126 | 10.688455 | 4.306019 | 6.367584 | 6.129877 | 6.137531 | 5.084587 | 3.127046 | 4.32155 | 3.991312 | 5.878754 | 5.846646 | 5.310211 | 4.333654 | 8.172394 | 29.5138889 | 1.223328 | 0.448302 |
| 29.4375 | 4.636992 | 8.093223 | 5.059765 | 4.473052 | 1.802385 | 2.915756 | 3.554276 | 2.969417 | 4.719481 | 5.739465 | 18.135765 | 5.413652 | 5.649663 | 6.612205 | 7.236794 | 3.849769 | 3.663819 | 4.172203 | 5.535634 | 6.980909 | 4.647222 | 6.432667 | 4.297105 | 11.53216 | 29.5277778 | 1.408195 | 2.156398 |
| 29.4513889 | 4.373878 | 6.780691 | 5.841303 | 3.185096 | 1.61443 | 2.639759 | 3.732409 | 2.892999 | 8.335706 | 8.031413 | 30.445328 | 6.13078 | 3.58032 | 6.259696 | 7.247899 | 1.611371 | 4.411934 | 3.244302 | 6.030635 | 7.108363 | 0.573552 | 7.839263 | 2.884495 | 15.189483 | 29.5416667 | 0.842973 | 5.28722 |
| 29.4652778 | 3.852435 | 4.490262 | 5.89674 | 2.544126 | 1.104178 | 3.316469 | 3.739949 | 2.373323 | 8.398656 | 7.814266 | 28.855076 | 4.775405 | 0.596023 | 2.47173 | 3.536879 | -0.306718 | 2.791884 | 1.053977 | 2.550181 | 2.617175 | -1.97861 | 6.289837 | 1.010654 | 9.558938 | 29.5555556 | 0.71833 | 3.374487 |
| 29.4791667 | -0.270485 | -1.890126 | 2.30725 | 1.069925 | 0.502166 | 1.776346 | 2.476425 | 1.226999 | 3.699909 | 3.317 | 8.900158 | -0.145182 | -2.080106 | -2.953174 | -1.902746 | -1.441814 | 0.157944 | -0.487003 | -0.366464 | -1.226447 | -2.531529 | 1.505202 | -0.165316 | -1.535939 | 29.5694444 | 0.235348 | 0.60825 |
| 29.4930556 | -4.636534 | -5.647045 | 0.207009 | 0.522227 | 0.008797 | 0.676025 | 1.127488 | 0.384408 | 0.524268 | 0.569644 | -5.57306 | -1.281579 | -0.680249 | -2.930479 | -1.620005 | -0.000861 | 0.150162 | 0.93629 | 0.557538 | 0.261052 | -0.381107 | -1.120547 | 0.220448 | -0.237929 | 29.5833333 | -0.54386 | 0.14215 |
| 29.5069444 | -2.979853 | -2.545364 | 0.890837 | 2.141321 | 0.242317 | 1.107583 | 1.145178 | 0.698128 | 0.472156 | 1.397735 | -4.04033 | 1.767019 | 1.758717 | 0.056561 | 0.553475 | 1.645192 | 2.522551 | 3.346942 | 3.872642 | 5.502359 | 3.996751 | 2.343707 | 2.561971 | 9.123602 | 29.5972222 | -0.802984 | 0.384253 |
| 29.5208333 | 0.103662 | 0.375973 | 1.597417 | 3.070637 | 0.35358 | 1.786338 | 1.046039 | 1.277926 | 1.50187 | 2.794255 | 3.43859 | 4.268429 | 4.89852 | 4.209124 | 3.220188 | 2.221312 | 3.57312 | 4.73969 | 6.587715 | 7.930192 | 6.434175 | 5.679483 | 5.47389 | 15.838636 | 29.6111111 | -0.269791 | 0.40473 |
| 29.5347222 | 2.630907 | 3.275912 | 2.000139 | 2.634982 | -0.06053 | 1.758971 | 0.884797 | 1.311351 | 2.41352 | 2.773569 | 8.511256 | 3.765611 | 4.003797 | 5.106551 | 3.529067 | 0.923647 | 2.644916 | 3.930991 | 7.337097 | 5.997589 | 4.636728 | 6.672075 | 5.626784 | 12.496807 | 29.625 | 0.487717 | 0.353144 |
| 29.5486111 | 2.579081 | 3.918512 | 2.082239 | 1.644861 | 0.386794 | 1.666714 | 0.874445 | 0.279415 | 2.410627 | 2.728101 | 9.824458 | 2.446022 | 4.176048 | 2.960653 | 1.929987 | -0.496351 | 2.222542 | 2.1717 | 3.501649 | 2.258306 | 0.898626 | 4.903355 | 3.795523 | 4.044413 | 29.6388889 | 1.32616 | 0.234135 |
| 29.5625 | 1.405951 | 1.902233 | 1.469606 | -0.584344 | 0.57456 | 1.611941 | 0.666628 | -0.158807 | 1.213797 | 1.409997 | 4.697691 | 1.38413 | 2.750983 | -0.330778 | -1.332932 | -0.823942 | 0.654948 | -0.608005 | 0.933739 | -0.888182 | -2.395518 | 1.930632 | 0.897605 | -1.917129 | 29.6527778 | 2.347202 | 0.078762 |
| 29.5763889 | 0.737621 | 2.152084 | 1.066164 | -1.228686 | 0.36825 | 1.182242 | 0.413072 | 0.160792 | -0.0752 | -0.105228 | 0.717014 | 0.371988 | 0.76295 | -0.262015 | -0.998855 | 0.44736 | -0.990915 | -0.840532 | 0.017756 | -0.329195 | -1.669456 | -1.718801 | -0.366638 | -2.47797 | 29.6666667 | 2.285043 | 0.017743 |
| 29.5902778 | 0.610592 | 1.220357 | 1.396697 | -0.653607 | 0.158835 | 0.01623 | -0.072148 | -0.08494 | -0.772462 | -1.052968 | -0.800526 | -0.428836 | 0.920767 | -0.024561 | -0.436899 | 0.241255 | -0.79204 | 0.223582 | 0.933279 | 1.679244 | 1.521183 | -2.729684 | -0.602339 | 0.860608 | 29.6805556 | 0.622651 | -0.034693 |
| 29.6041667 | -0.039195 | -0.144194 | 0.830205 | 0.053515 | -0.001302 | -0.827087 | -0.694055 | -0.322503 | 0.035723 | -0.033769 | -0.58829 | -0.480733 | 0.617523 | 1.245577 | 0.931236 | -0.149153 | -0.007582 | 1.713374 | 1.64809 | 3.529793 | 5.100385 | -0.922939 | 1.147909 | 4.527309 | 29.6944444 | -1.233645 | -0.243412 |
| 29.6180556 | -0.682223 | -0.930078 | -0.252286 | 1.217719 | 0.034458 | -0.248665 | -0.539827 | 0.156751 | 0.584433 | 1.055562 | 0.184054 | -0.145787 | -0.530709 | 1.229332 | 1.797337 | 0.524653 | 0.605897 | 1.658578 | 1.180139 | 3.058742 | 3.351889 | 1.559409 | 2.229886 | 4.427942 | 29.7083333 | -1.558636 | -0.584735 |
| 29.6319444 | -0.82382 | -0.703445 | -0.690734 | 1.54305 | 0.267848 | 0.639103 | -0.239616 | 0.239058 | 1.043467 | 1.888761 | 0.061999 | 0.007366 | 0.35197 | 0.637091 | 0.949062 | -0.091244 | 0.697864 | 0.537277 | -0.643614 | -0.464857 | -0.756238 | 3.192499 | 0.803946 | -1.167132 | 29.7222222 | -0.705032 | -0.786199 |
| 29.6458333 | -0.222882 | 1.618038 | -0.001814 | 1.231936 | 0.131014 | 0.886455 | 0.095703 | 0.074103 | 1.027416 | 1.637078 | -0.458848 | -0.120563 | -1.068951 | -0.442323 | -1.756475 | -1.046587 | 0.330552 | -0.50864 | -2.176147 | -3.553729 | -4.479114 | 0.711035 | -2.813194 | -6.348446 | 29.7361111 | 0.506541 | -0.855844 |
| 29.6597222 | 0.706358 | 3.033515 | 1.205203 | 0.458832 | -0.102992 | 0.522299 | 0.113879 | -0.142367 | 0.192672 | 0.12767 | -0.117756 | -0.140706 | -0.447712 | -0.661534 | -0.738699 | 1.082952 | -1.52691 | -2.634824 | -0.800677 | -2.991288 | -3.844867 | -2.971032 | -4.198027 | -5.916352 | 29.75 | 1.191926 | -0.690406 |
| 29.6736111 | 1.205574 | 0.677793 | 1.051929 | -0.360557 | -0.006286 | 0.489998 | 0.154191 | 0.66966 | -0.364307 | -1.488217 | -0.019593 | -0.14384 | -0.325919 | -0.162666 | 0.175233 | 1.050379 | -1.914203 | -2.659996 | -0.187812 | -0.694296 | 0.65816 | -3.713882 | -2.437166 | -1.106252 | 29.7638889 | 1.125619 | -0.431824 |
| 29.6875 | 0.364824 | -1.589761 | 0.14414 | -0.153401 | 0.318561 | 0.800143 | 0.064055 | 0.284199 | -0.686476 | -2.270124 | -0.330546 | -0.046045 | 0.116444 | -0.10893 | 0.79341 | 0.646295 | -0.662856 | -0.279812 | 0.351282 | 2.757429 | 3.878295 | -1.046536 | 0.986929 | 3.577442 | 29.7777778 | 0.56009 | -0.146816 |

| | | | | | | | |
|---|---|---|---|---|---|---|---|
| 29.7013889 | -0.966839 | -2.709305 | -0.044351 | 0.533182 | -0.084526 | 0.102397 | -0.089213 |
| 29.7152778 | -1.653912 | -2.053461 | -0.205054 | 0.572799 | -0.431716 | -0.732078 | -0.219409 |
| 29.7291667 | -1.084808 | -0.148167 | -0.192807 | 0.292047 | -0.241442 | -0.591431 | -0.183319 |
| 29.7430556 | -0.195528 | 1.798456 | 0.048999 | -0.391597 | -0.013172 | -0.075932 | -0.073272 |
| 29.7569444 | 0.464767 | 1.119258 | 0.371074 | -1.600796 | 0.264052 | 0.719936 | 0.332692 |
| 29.7708333 | 0.531141 | -1.022758 | -0.073815 | -1.93841 | 0.40994 | 0.681296 | -0.362631 |
| 29.7847222 | -0.855171 | -2.292981 | 0.15064 | -1.026573 | 0.1923 | 0.015323 | -0.299933 |
| 29.7986111 | -2.672257 | -4.904649 | -0.619162 | 0.206559 | 0.029219 | -0.942304 | -0.687658 |
| 29.8125 | -3.210572 | -4.289386 | -1.154153 | 0.714835 | -0.41078 | -1.14407 | -0.203999 |
| 29.8263889 | -1.823255 | -0.901642 | -1.393623 | -0.120175 | -0.667376 | -1.017908 | -0.262162 |
| 29.8402778 | -0.616358 | -0.076388 | -2.096482 | -1.002304 | -0.520718 | -0.66094 | 0.137223 |
| 29.8541667 | -0.487337 | -2.193117 | -3.208197 | -1.927625 | -0.957394 | -1.244869 | -0.168355 |
| 29.8680556 | -2.167475 | -4.818578 | -2.556599 | -3.021196 | -1.702739 | -2.204782 | -1.330724 |
| 29.8819444 | -2.790433 | -5.248425 | -0.883093 | -2.222303 | -2.024856 | -2.978995 | -2.099466 |
| 29.8958333 | -3.218631 | -4.854291 | -0.634955 | -2.285964 | -1.851901 | -3.107971 | -1.420515 |
| 29.9097222 | -2.243571 | -3.661268 | -1.68785 | -2.887171 | -1.94137 | -2.81056 | -0.936355 |
| 29.9236111 | -1.222797 | -3.463003 | -3.006326 | -3.068927 | -2.064304 | -3.1738 | -1.712296 |
| 29.9375 | -2.267981 | -4.081837 | -5.38971 | -3.487142 | -2.590249 | -3.721036 | -3.194975 |
| 29.9513889 | -3.643321 | -3.424738 | -7.746578 | -2.374323 | -3.305581 | -3.699543 | -4.647902 |
| 29.9652778 | -4.367517 | -3.477392 | -5.182999 | -1.636685 | -2.495112 | -2.26836 | -3.428403 |
| 29.9791667 | -3.230256 | -3.059932 | -2.773182 | -1.546805 | -1.238165 | -1.176422 | -1.684933 |
| 29.9930556 | -1.131275 | -1.247995 | -1.089854 | -1.198939 | -0.580558 | -0.63585 | -0.597031 |
| 30.0069444 | -0.422622 | -0.275179 | -0.320592 | -0.637115 | -0.29666 | -0.349111 | 0.006574 |
| 30.0208333 | -0.539201 | -0.187765 | -0.090007 | -0.088503 | -0.103478 | -0.190696 | 0.20642 |
| 30.0347222 | -0.896567 | -0.420066 | -0.011577 | 0.084911 | -0.016875 | -0.198758 | 0.040594 |

| | | | | | | | | | | | | | | | | | | | | | | | | | | | | |
|---|---|---|---|---|---|---|---|---|---|---|---|---|---|---|---|---|---|---|---|---|---|---|---|---|---|---|---|---|
| 32.6319444 | -1.047906 | -0.456016 | 1.188431 | 0.89752 | -0.694342 | -1.596277 | -0.502578 | -0.802675 | 0.347075 | 1.643485 | -1.709179 | -0.791342 | -1.782134 | -0.044151 | 1.785684 | 0.139387 | 0.107788 | 2.102508 | -2.657217 | -3.212653 | 1.999446 | 0.348998 | 2.068818 | 5.866097 | | 32.7222222 | 1.3896 | -0.264676 |
| 32.6458333 | 0.684361 | 1.229368 | 1.066454 | -0.062121 | -0.8576 | -1.393356 | -0.675973 | -0.373935 | -0.629483 | 0.45116 | 0.940999 | -0.647005 | -1.370542 | 0.262704 | 2.300613 | 2.931715 | 0.039022 | 0.37199 | -1.324476 | -3.027939 | 2.6594 | 3.062911 | 6.800087 | 3.342528 | | 32.7361111 | 2.545063 | 0.073972 |
| 32.6597222 | 1.291824 | 2.10546 | -0.285039 | -1.147756 | -0.723131 | -0.776672 | -0.721437 | 0.325781 | -1.247959 | -0.058003 | 2.55306 | -0.074542 | -0.576758 | 0.320917 | 0.060137 | 0.649569 | -0.174523 | -0.086573 | 1.018572 | -0.86721 | 0.991655 | 2.795891 | 1.333668 | -2.986076 | | 32.75 | 3.201241 | -0.267221 |
| 32.6736111 | 0.760748 | 1.186356 | -1.415886 | -1.357401 | -0.420219 | 0.071218 | -0.855003 | 0.472709 | -1.33553 | -0.414904 | 1.402735 | 0.037462 | -1.091198 | -1.166329 | -3.04078 | -3.046488 | 0.321521 | 0.18529 | 2.699378 | 2.306751 | -1.28663 | -2.071414 | -0.842104 | -7.110757 | | 32.7638889 | 3.257345 | -0.540502 |
| 32.6875 | -0.056794 | 0.91564 | -1.417379 | -0.652594 | 0.472901 | 1.47793 | -0.287966 | 0.482806 | -0.946061 | -0.398988 | -0.403897 | -0.549853 | -0.994531 | -2.925418 | -5.896185 | -3.919189 | 0.917385 | 0.037257 | 3.135882 | 3.404709 | -2.811963 | -8.532506 | -0.306089 | -5.824658 | | 32.7777778 | -1.828358 | -0.495167 |
| 32.7013889 | -2.339677 | -2.843225 | -1.039084 | 0.064736 | 0.801091 | 1.042313 | -0.320563 | 0.174072 | -0.060558 | 0.622408 | -1.11417 | -0.368227 | -0.648493 | -1.555563 | -2.028574 | -1.453545 | 0.054518 | -0.826415 | 0.218643 | 2.836178 | -2.165367 | -7.378462 | -0.210036 | -1.135585 | | 32.7916667 | -4.159059 | -0.23475 |
| 32.7152778 | -1.773611 | -3.955381 | 0.549607 | 0.786433 | 0.300943 | 0.34389 | -0.928772 | -1.066527 | 0.767185 | 0.121509 | -2.510854 | -0.591826 | 0.017445 | 0.512635 | 1.510761 | 1.733869 | -0.558966 | -1.655123 | -1.546938 | 1.678889 | -1.062281 | -1.802043 | -0.191773 | 3.836526 | | 32.8055556 | -2.662597 | -0.144474 |
| 32.7291667 | -1.903731 | -3.602172 | 0.871709 | 0.695991 | -0.15867 | -0.702936 | -0.684381 | -1.440126 | 1.110103 | -0.167448 | -3.776159 | -0.307612 | -0.526409 | 2.335691 | 3.981487 | 4.18962 | -0.741765 | -0.973853 | -0.644571 | 0.753969 | 0.048342 | 3.180666 | 0.137635 | 5.855098 | | 32.8194444 | -0.058315 | -0.369068 |
| 32.7430556 | -2.883483 | -4.518003 | 0.075988 | 0.127129 | -1.05932 | -1.406379 | -0.007394 | -0.028625 | 0.902007 | -1.05708 | -2.997424 | 0.17594 | 0.031396 | 1.276389 | 2.67384 | 2.858107 | -0.558927 | -0.187473 | 1.120182 | -1.87325 | 1.691255 | 4.854071 | 3.599571 | 4.170756 | | 32.8333333 | 1.347914 | -0.511787 |
| 32.7569444 | -2.455079 | -2.88685 | -0.877326 | -0.729643 | -1.041395 | -0.820807 | 0.600687 | -0.306632 | 0.415693 | -0.254516 | -0.355284 | 0.849461 | -0.296588 | 0.281782 | -0.752864 | -0.556561 | 0.316432 | 0.715106 | 1.667585 | -1.814029 | 1.479161 | 4.372681 | 1.535583 | -0.805276 | | 32.8472222 | 1.618492 | -0.534454 |
| 32.7708333 | -0.033003 | 1.268925 | -1.01237 | -0.620944 | -0.23789 | 0.564023 | 0.850745 | 0.294697 | -0.243405 | 1.190242 | 0.949982 | 0.85623 | 0.23544 | -0.325027 | -0.713258 | -1.378407 | 0.296076 | -0.394424 | 0.162521 | -1.348786 | -0.495592 | 0.392476 | -1.187288 | -3.441595 | | 32.8611111 | 0.842983 | -0.589331 |
| 32.7847222 | 1.730177 | 4.526517 | -0.566294 | -0.005787 | 0.337833 | 1.000551 | 0.008936 | -0.329189 | -0.47879 | 2.47538 | 1.461106 | 0.388561 | 0.236807 | -1.187789 | 0.661899 | -1.074873 | 0.163221 | -0.186965 | -2.538438 | -1.72436 | -1.91399 | -3.441581 | -2.042361 | -3.705719 | | 32.875 | -0.850717 | -1.078227 |
| 32.7986111 | 1.577834 | 4.211774 | 0.520105 | 0.701431 | 0.629279 | 0.832214 | -0.843334 | -1.114397 | -0.066016 | 2.152724 | 1.481948 | -0.249472 | 0.132121 | -0.99558 | -0.295194 | -0.387091 | 0.33184 | 0.656075 | -4.52653 | -1.932108 | -0.304993 | -6.905024 | -0.349892 | -3.690432 | | 32.8888889 | -2.707232 | -2.065214 |
| 32.8125 | 1.015172 | 1.063836 | 1.284762 | 0.562836 | 0.148034 | -0.231202 | -1.032606 | -1.114439 | 0.03237 | 1.162655 | 1.25727 | -1.625785 | -0.575598 | -1.028983 | -1.487087 | 1.37811 | 0.597841 | 1.272354 | -2.205923 | -0.682112 | 1.820019 | -3.898761 | -0.714749 | -0.911399 | | 32.9027778 | -4.409099 | -2.581292 |
| 32.8263889 | 0.093538 | -1.432337 | 1.389472 | -0.376773 | -0.67417 | -1.11602 | -0.683387 | -0.60997 | -0.406028 | 0.922508 | 0.395237 | -1.436694 | -1.686724 | -0.208173 | 0.597524 | 3.382281 | 0.404726 | 0.897065 | 0.440458 | 0.858914 | 0.500807 | -0.674761 | -1.121028 | 2.070531 | | 32.9166667 | -4.57241 | -2.119996 |
| 32.8402778 | -1.824522 | -2.546731 | 1.231263 | -0.44477 | -1.785324 | -2.375284 | -1.10221 | -0.13757 | -1.087647 | -0.222037 | -1.076085 | -0.460238 | -2.200585 | 1.606564 | 1.707578 | -0.471204 | -1.26074 | -1.446332 | 0.817504 | -0.556988 | -0.259698 | 2.356615 | -0.629777 | -0.241235 | | 32.9305556 | -4.698413 | -1.289526 |
| 32.8541667 | -4.428676 | -3.30689 | -0.211096 | -0.826301 | -0.770179 | -1.29375 | -1.417155 | -1.337752 | -1.8669 | -2.225712 | -2.906893 | -0.330772 | -1.80719 | -0.87887 | -0.752697 | -2.725211 | -2.990252 | -3.89104 | 0.493464 | -2.31388 | -2.813753 | 2.125667 | -3.4215 | -5.436723 | | 32.9444444 | -4.178013 | -0.5675 |
| 32.8680556 | -5.137275 | -3.465939 | -1.434287 | -0.944405 | 0.43476 | 0.262176 | -0.66606 | -0.869022 | -2.494935 | -3.157355 | -4.785791 | -1.095003 | -2.273211 | -4.465098 | -3.091056 | -2.808336 | -3.709085 | -5.26074 | 0.006437 | -2.020091 | -4.162473 | -0.79814 | -2.321904 | -8.578596 | | 32.9583333 | -4.204866 | -0.22109 |
| 32.8819444 | -4.938492 | -4.949598 | -3.494772 | -1.095844 | 0.157361 | -0.278853 | -0.641232 | -0.996899 | -1.866895 | -2.941094 | -6.890402 | -0.871036 | -2.804682 | -7.138831 | -5.945032 | -2.744155 | -1.955445 | -3.166332 | 0.247404 | -0.79769 | -4.01172 | -4.928294 | -1.55211 | -7.947777 | | 32.9722222 | -5.896807 | -0.222591 |
| 32.8958333 | -4.535735 | -5.570647 | -4.690155 | -2.088281 | -1.747643 | -2.956095 | -0.766474 | -2.031235 | -0.94664 | -2.434754 | -9.901986 | -2.027673 | -3.585123 | -5.639849 | -5.166005 | -1.711597 | -0.975262 | -2.228373 | -1.692848 | -4.567706 | -5.511724 | -9.018148 | -2.690034 | -6.881824 | | 32.9861111 | -4.443617 | -0.494611 |
| 32.9097222 | -5.815993 | -7.245983 | -4.8713 | -3.004171 | -4.178483 | -6.123218 | -1.993611 | -3.949698 | -1.323893 | -2.576167 | -11.589395 | -5.375623 | -5.568515 | -2.445418 | -3.111398 | -3.066589 | -2.146309 | -2.615234 | -5.402496 | -7.871287 | -4.646351 | -8.263459 | -3.720395 | -8.176985 | | 33 | -1.644088 | -0.63393 |
| 32.9236111 | -6.966253 | -7.802778 | -4.312388 | -3.33981 | -4.444256 | -6.317201 | -3.085905 | -4.021752 | -2.920485 | -3.794962 | -13.82564 | -5.360833 | -4.976791 | -0.234839 | -2.217463 | -3.693238 | -4.470354 | -3.60494 | -8.222679 | -7.912309 | -4.588605 | -8.154296 | -5.296652 | -11.031489 | | 33.0138889 | 0.686538 | -0.553048 |
| 32.9375 | -8.515883 | -7.76424 | -4.696247 | -2.668545 | -3.376322 | -5.238434 | -4.373773 | -3.049478 | -6.604085 | -6.117288 | -27.043464 | -7.770706 | -4.791189 | -0.56327 | -4.034775 | -4.285773 | -6.996276 | -6.206633 | -9.342885 | -6.921861 | -3.331272 | -10.542962 | -5.841213 | -18.768638 | | 33.0277778 | 1.107562 | -0.843972 |
| 32.9513889 | -8.810532 | -6.935503 | -5.533298 | -2.432628 | -3.652176 | -4.902884 | -5.857445 | -3.122331 | -8.864599 | -6.955318 | -47.104537 | -12.867944 | -6.959305 | -4.444964 | -4.676937 | -3.421584 | -7.324081 | -6.29023 | -10.126001 | -6.982083 | -1.529839 | -13.178947 | -6.037222 | -21.976137 | | 33.0416667 | -2.201615 | -2.258907 |
| 32.9652778 | -10.417499 | -6.043464 | -5.140671 | -2.133616 | -2.707621 | -3.568 | -4.63553 | -3.215626 | -7.477067 | -5.937946 | -40.15342 | -11.732197 | -8.263372 | -5.200552 | -4.904403 | -2.925397 | -5.172077 | -3.741079 | -6.26579 | -6.152284 | -1.27749 | -9.82576 | -4.87517 | -16.444337 | | 33.0555556 | -4.179654 | -1.398999 |
| 32.9791667 | -7.11045 | -4.661467 | -3.415437 | -1.593074 | -1.306232 | -2.420485 | -2.380527 | -2.588998 | -3.214308 | -2.890148 | -17.431263 | -9.321337 | -7.163149 | -6.018493 | -3.493452 | -1.390668 | -2.007958 | -0.904805 | -2.715127 | -4.512201 | -1.767203 | -4.835169 | -2.686858 | -8.20539 | | 33.0694444 | -4.858852 | -0.394774 |
| 32.9930556 | -2.71495 | -1.438382 | -1.491407 | -1.004194 | -0.761224 | -1.596236 | -0.865109 | -1.529358 | -0.485739 | -0.892248 | -3.74293 | -6.047176 | -3.264791 | -5.691774 | -3.493638 | -0.873898 | -1.604694 | -1.083041 | -0.948319 | -3.470922 | -1.588202 | -3.770655 | -0.955707 | -3.343688 | | 33.0833333 | -5.172218 | -0.062365 |
| 33.0069444 | -0.354682 | 0.670826 | -0.585955 | -0.550075 | -0.471649 | -1.079382 | -0.561594 | -1.153366 | 0.140668 | -0.35632 | -0.311589 | -2.770744 | 0.226008 | -4.403121 | -3.967183 | -0.906702 | -2.32553 | -2.126016 | -1.162869 | -2.668614 | -1.088093 | -3.295646 | -0.642124 | -2.517517 | | 33.0972222 | -4.963041 | 0.083248 |
| 33.0208333 | 0.321041 | 0.994234 | -0.440321 | -0.401673 | -0.242162 | -0.744133 | -0.566983 | -1.0486 | -0.000224 | -0.507721 | 0.205163 | -1.859999 | 0.394601 | -3.622681 | -3.55092 | -1.078952 | -2.648068 | -1.914526 | -1.81672 | -1.564713 | -0.511045 | -2.706705 | -1.237598 | -3.129999 | | 33.1111111 | -3.689803 | 0.330081 |
| 33.0347222 | -0.056602 | 0.330161 | -0.582781 | -0.339895 | -0.204566 | -0.646122 | -0.367465 | -0.760967 | -0.278497 | -0.640609 | -0.084053 | -1.839232 | -1.196375 | -3.12894 | -2.63509 | -1.642149 | -2.360707 | -1.647103 | -2.00819 | -1.153211 | -0.92611 | -2.517808 | -2.289267 | -3.778245 | | 33.125 | -2.023543 | 0.619291 |
| 33.0486111 | -0.618775 | -0.386836 | -0.662958 | -0.239766 | -0.258416 | -0.682178 | -0.279951 | -0.629445 | -0.395244 | -0.57608 | -0.540333 | -2.423535 | -2.42471 | -3.4385 | -2.457497 | -2.507314 | -2.163045 | -2.698112 | -2.068725 | -1.516699 | -2.065967 | -3.079437 | -3.082303 | -4.577894 | | 33.1388889 | -0.527134 | 0.745125 |
| 33.0625 | -1.111394 | -0.849053 | -0.650836 | -0.173065 | -0.306788 | -0.657981 | -0.350141 | -0.65391 | -0.350147 | -0.485185 | -0.817854 | -2.722967 | -2.513884 | -3.744746 | -2.673687 | -2.91587 | -2.552274 | -3.409097 | -2.753204 | -2.212164 | -2.457104 | -4.487222 | -3.1048 | -5.08271 | | 33.1527778 | 0.884758 | 0.609195 |
| 33.0763889 | -1.451099 | -0.979949 | -0.669635 | -0.230576 | -0.335895 | -0.583136 | -0.441138 | -0.705588 | -0.331858 | -0.500832 | -1.107527 | -2.692983 | -2.547381 | -3.603993 | -3.028692 | -2.563092 | -3.144581 | -2.673153 | -3.174586 | -2.463701 | -2.043755 | -5.266946 | -2.429095 | -5.085394 | | 33.1666667 | 1.820314 | 0.198581 |
| 33.0902778 | -1.690038 | -0.85224 | -0.745127 | -0.425742 | -0.357451 | -0.464734 | -0.478482 | -0.739924 | -0.342784 | -0.578434 | -1.324783 | -2.497712 | -2.542282 | -2.945909 | -2.774309 | -1.887593 | -3.421391 | -1.913744 | -2.772877 | -2.003687 | -1.4912 | -4.769393 | -2.015817 | -4.589383 | | 33.1805556 | 1.858823 | -0.403228 |
| 33.1041667 | -2.064544 | -0.885088 | -0.865083 | -0.533027 | -0.371886 | -0.347197 | -0.514526 | -0.845076 | -0.375047 | -0.683992 | -1.164406 | -2.393197 | -1.808822 | -1.9262 | -1.915783 | -1.454492 | -3.239039 | -1.533344 | -2.413187 | -1.738915 | -0.963699 | -3.706262 | -1.728109 | -3.8011 | | 33.1944444 | 1.254904 | -0.435659 |
| 33.1180556 | -2.680906 | -1.351446 | -1.062114 | -0.564037 | -0.448215 | -0.378871 | -0.663858 | -1.147322 | -0.467439 | -0.835808 | -1.069447 | -2.110183 | -1.221134 | -1.259521 | -1.3427 | -1.163634 | -2.433783 | -0.941568 | -2.302319 | -1.646729 | -0.461536 | -2.82474 | -1.520331 | -2.768912 | | 33.2083333 | 0.660605 | 0.421252 |
| 33.1319444 | -3.481133 | -2.028015 | -1.300362 | -0.70111 | -0.537095 | -0.559821 | -0.966143 | -1.53001 | -0.704697 | -1.039034 | -1.834627 | -1.75631 | -1.372509 | -1.000417 | -1.091856 | -1.003353 | -1.570803 | -0.187061 | -1.563429 | -1.231847 | -0.134085 | -2.480383 | -1.003577 | -1.796503 | | 33.2222222 | 0.766088 | 1.273368 |
| 33.1458333 | -4.15239 | -2.274417 | -1.454772 | -0.972465 | -0.556583 | -0.688272 | -1.25136 | -1.641697 | -0.931256 | -1.181041 | -2.773315 | -1.496499 | -1.26124 | -1.005745 | -0.739314 | -0.405123 | -1.18921 | 0.132203 | -0.659803 | -0.602602 | -0.066641 | -1.731664 | -0.443318 | -1.085139 | | 33.2361111 | 1.449569 | 1.446677 |
| 33.1597222 | -4.49401 | -1.988073 | -1.351241 | -0.957015 | -0.449865 | -0.552982 | -1.263643 | -1.585469 | -0.859078 | -0.936341 | -2.746827 | -1.026908 | -0.164855 | -0.152213 | 0.147032 | 0.27457 | -0.541734 | 0.203831 | -0.689747 | -0.299536 | 0.096026 | -0.761607 | -0.408811 | -0.538636 | | 33.25 | 2.164329 | 0.979339 |
| 33.1736111 | -4.444173 | -1.216729 | -1.140075 | -0.655811 | -0.295591 | -0.251094 | -1.072183 | -1.399924 | -0.583457 | -0.479066 | -2.292403 | -0.551968 | 0.482956 | -0.008533 | 0.000405 | 0.532843 | 0.090227 | 0.489816 | -0.886104 | -0.205166 | 0.616043 | -0.184064 | -0.370096 | 0.050083 | | 33.2638889 | 2.122497 | 0.335018 |
| 33.1875 | -4.145245 | -0.788956 | -1.029421 | -0.448858 | -0.168292 | -0.053358 | -0.909898 | -1.176477 | -0.404902 | -0.265735 | -2.264633 | -0.637675 | -0.06971 | 0.085233 | -0.292405 | 0.217661 | 0.129966 | 0.66244 | -0.279227 | -0.060386 | 0.839502 | -0.015775 | -0.200871 | 0.361321 | | 33.2777778 | 1.721297 | -0.046477 |
| 33.2013889 | -3.761949 | -0.688052 | -0.949367 | -0.468869 | -0.046339 | 0.028109 | -0.841042 | -1.000778 | -0.419767 | -0.332058 | -2.123391 | -1.380002 | -0.657028 | -0.012936 | -0.252091 | 0.077508 | -0.252853 | 0.301982 | 0.203442 | -0.117769 | 0.342228 | -0.093595 | -0.022645 | 0.126685 | | 33.2916667 | 1.660068 | -0.109819 |
| 33.2152778 | -3.238239 | -0.576936 | -0.834193 | -0.476569 | 0.028171 | 0.067555 | -0.76792 | -0.764459 | -0.498825 | -0.408113 | -1.656983 | -1.941891 | -0.574753 | -0.359106 | -0.107892 | 0.334206 | -0.169386 | -0.160163 | -0.066423 | -0.274567 | -0.080615 | -0.010413 | 0.042305 | -0.279596 | | 33.3055556 | 1.945503 | 0.026212 |
| 33.2291667 | -2.77825 | -0.229363 | -0.649628 | -0.302866 | 0.031811 | 0.05641 | -0.565342 | -0.462715 | -0.444903 | -0.318825 | -1.135731 | -1.471185 | -0.228779 | -0.499916 | -0.192184 | 0.447792 | 0.316753 | -0.127011 | -0.142491 | -0.090486 | 0.171049 | 0.250117 | 0.181988 | -0.135856 | | 33.3194444 | 1.982177 | 0.282326 |
| 33.2430556 | -2.01803 | 0.055014 | -0.533629 | -0.129679 | -0.070648 | -0.064354 | -0.304911 | -0.216343 | -0.277894 | -0.136015 | -0.801146 | -0.467944 | -0.303478 | -0.170221 | -0.062823 | 0.161754 | 0.545595 | 0.080228 | 0.333277 | 0.132676 | 0.397177 | 0.481275 | 0.264636 | 0.182332 | | 33.3333333 | 1.786667 | 0.519088 |
| 33.2569444 | -1.386718 | -0.003766 | -0.495162 | -0.151479 | -0.216871 | -0.269171 | -0.102957 | -0.058447 | -0.189863 | -0.057284 | -0.63769 | 0.04169 | -0.149438 | -0.184138 | -0.11295 | 0.002333 | 0.43356 | -0.101682 | 0.323409 | -0.028186 | 0.040829 | 0.344878 | -0.038493 | -0.111255 | | 33.3472222 | 1.42712 | 0.475546 |
| 33.2708333 | -0.960311 | -0.097343 | -0.420509 | -0.159424 | -0.251749 | -0.43543 | -0.030456 | 0.075928 | -0.241984 | -0.113633 | -0.555705 | 0.332524 | 0.570894 | -0.255993 | -0.011647 | 0.24082 | 0.313336 | -0.106898 | 0.344974 | 0.104284 | -0.139555 | 0.39206 | -0.312976 | -0.177552 | | 33.3611111 | 1.373337 | 0.119513 |
| 33.2847222 | -0.137302 | 0.224323 | -0.225345 | 0.049739 | -0.129621 | -0.299774 | 0.002857 | 0.287756 | -0.238167 | -0.047181 | -0.308785 | 1.232789 | 1.136474 | -0.059921 | 0.05676 | 0.598907 | 0.689352 | 0.543753 | 1.172939 | 0.906473 | 0.598059 | 1.12909 | -0.069598 | 0.853231 | | 33.375 | 1.714644 | -0.158274 |
| 33.2986111 | 1.017226 | 0.723415 | -0.007248 | 0.250685 | 0.062335 | 0.15166 | 0.13169 | 0.509111 | -0.036802 | 0.255008 | 0.397563 | 2.523595 | 1.441417 | 0.848823 | 0.682135 | 0.864479 | 1.257732 | 1.023589 | 1.418382 | 1.248817 | 1.581646 | 2.228554 | 0.485433 | 1.977547 | | 33.3888889 | 2.229738 | -0.41481 |
| 33.3125 | 2.147866 | 0.969783 | 0.097698 | 0.246498 | 0.233053 | 0.571087 | 0.269792 | 0.683432 | 0.218678 | 0.541336 | 1.06469 | 3.25371 | 1.885302 | 1.395752 | 1.05643 | 1.337983 | 1.890994 | 1.273792 | 1.099226 | 1.040047 | 1.861169 | 3.169456 | 1.171533 | 2.61494 | | 33.4027778 | 2.210882 | -0.91325 |
| 33.3263889 | 3.00855 | 1.124132 | 0.152107 | 0.243561 | 0.394435 | 0.877281 | 0.455648 | 0.847938 | 0.329576 | 0.687873 | 1.502859 | 3.350763 | 2.410457 | 1.760276 | 1.382135 | 1.859288 | 2.775729 | 1.653129 | 1.288887 | 1.220194 | 1.697327 | 3.912932 | 1.878923 | 3.325021 | | 33.4166667 | 2.0231 | -0.924995 |
| 33.3402778 | 4.042239 | 1.63067 | 0.428731 | 0.554719 | 0.602521 | 1.190049 | 0.707049 | 1.160101 | 0.466131 | 0.973019 | 2.111712 | 3.499303 | 2.520517 | 2.149097 | 1.771058 | 2.319827 | 3.925257 | 2.244885 | 2.112956 | 2.021678 | 1.935403 | 4.435817 | 2.344334 | 4.153102 | | 33.4305556 | 2.144822 | 0.176319 |
| 33.3541667 | 4.918878 | 2.414504 | 0.841215 | 1.078713 | 0.847582 | 1.685352 | 1.021378 | 1.578427 | 0.828046 | 1.606574 | 2.892074 | 3.576505 | 2.571255 | 2.6068 | 2.492228 | 2.754011 | 4.507992 | 2.883352 | 2.530743 | 2.576925 | 2.586054 | 5.158606 | 2.69372 | 4.909541 | | 33.4444444 | 2.304755 | 1.161816 |
| 33.3680556 | 5.555615 | 3.292837 | 1.211542 | 1.631662 | 1.292843 | 2.551715 | 1.393209 | 2.115884 | 1.421827 | 2.448545 | 3.731841 | 3.760129 | 3.004645 | 3.11255 | 3.498802 | 3.50686 | 5.122444 | 3.629675 | 3.088995 | 3.05974 | 3.057049 | 6.745975 | 3.309227 | 5.855775 | | 33.4583333 | 2.113964 | 1.413528 |
| 33.3819444 | 6.525956 | 4.155464 | 1.626723 | 2.269507 | 1.934036 | 3.816248 | 1.834387 | 2.830354 | 2.127693 | 3.330787 | 5.182986 | 4.857547 | 4.134115 | 4.291722 | 4.349496 | 4.467858 | 5.808742 | 4.509885 | 4.02347 | 3.980808 | 3.539873 | 9.036704 | 4.491529 | 7.773808 | | 33.4722222 | 1.989494 | 1.461647 |
| 33.3958333 | 7.137784 | 5.166389 | 2.104957 | 3.121312 | 2.455666 | 4.489914 | 2.13167 | 3.33951 | 2.783942 | 3.883084 | 6.288093 | 7.274811 | 6.070505 | 5.842678 | 5.562578 | 5.517454 | 5.626768 | 4.89508 | 4.952523 | 5.440925 | 4.499545 | 10.216452 | 6.712362 | 10.45443 | | 33.4861111 | 2.129248 | 1.277774 |
| 33.4097222 | 7.096587 | 5.968064 | 2.595418 | 3.800039 | 2.223493 | 4.2188 | 2.477731 | 3.704147 | 2.987974 | 4.006814 | 6.808759 | 9.603651 | 7.934543 | 6.944028 | 6.875963 | 6.567665 | 5.977864 | 6.663409 | 5.994153 | 6.280081 | 5.201034 | 11.786997 | 7.70287 | 13.079314 | | 33.5 | 2.086541 | 1.111621 |
| 33.4236111 | 7.69862 | 6.245033 | 2.986976 | 4.008206 | 1.933913 | 3.362656 | 2.847724 | 3.972822 | 3.041165 | 4.36991 | 8.39349 | 10.5801 | 9.395241 | 7.980931 | 8.499868 | 6.892837 | 7.054796 | 8.326727 | 7.27365 | 8.288878 | 5.952592 | 11.988978 | 7.178443 | 15.973013 | | 33.5138889 | 1.986898 | 1.271676 |
| 33.4375 | 9.905491 | 5.42826 | 3.527413 | 3.668807 | 1.88519 | 3.536025 | 3.095261 | 3.470658 | 4.183127 | 6.228087 | 19.517951 | 11.280412 | 10.93309 | 8.247774 | 8.424222 | 6.885321 | 6.47276 | 6.349986 | 8.570494 | 10.058558 | 6.921448 | 12.616573 | 6.617024 | 24.769742 | | 33.5277778 | 4.38548 | 2.702456 |
| 33.4513889 | 12.963092 | 4.485477 | 4.976251 | 2.862071 | 2.056095 | 4.921729 | 3.894108 | 3.337579 | 7.515867 | 9.121696 | 42.31289 | 10.748928 | 10.964243 | 8.570557 | 8.210702 | 6.901731 | 5.343189 | 4.558552 | 10.59212 | 11.968536 | 6.696103 | 15.933971 | 7.264678 | 40.857915 | | 33.5416667 | 8.434702 | 3.199461 |
| 33.4652778 | 14.291121 | 5.732749 | 6.920194 | 2.336212 | 1.559464 | 4.571195 | 4.931259 | 3.84221 | 8.772193 | 9.730565 | 49.745616 | 9.044309 | 9.515957 | 8.005528 | 6.81003 | 4.274208 | 4.117444 | 2.446347 | 9.600312 | 9.227007 | 5.077828 | 15.741758 | 5.868118 | 36.827755 | | 33.5555556 | 9.664679 | 1.804682 |
| 33.4791667 | 5.605863 | 3.36578 | 5.924535 | 2.264041 | 1.233702 | 3.809834 | 3.06892 | 2.640381 | 4.385932 | 4.791285 | 24.144752 | 1.029608 | 1.442235 | 2.200248 | 3.653525 | 0.035253 | 1.605659 | 1.906308 | 4.876821 | 3.169017 | 0.958621 | 1.573841 | 2.803933 | 11.093654 | | 33.5694444 | 6.866045 | -0.786671 |
| 33.4930556 | -1.568081 | 1.302866 | 4.706402 | 2.250064 | 0.612388 | 1.785945 | 0.839698 | 1.576438 | 1.039182 | 1.434265 | 1.371363 | -4.899265 | 0.943605 | -0.188219 | 2.804302 | 1.071406 | 2.068428 | 4.380223 | 3.172859 | 2.963247 | 0.687618 | -5.517263 | 2.535226 | 2.532684 | | 33.5833333 | 3.420885 | -0.267378 |
| 33.5069444 | -1.455207 | 2.432327 | 3.324625 | 0.766879 | 0.357211 | 1.095587 | -0.127705 | 1.041497 | 0.873895 | 1.516487 | -2.155206 | -0.537709 | 2.851388 | 0.237878 | 4.040635 | 4.374569 | 2.197517 | 5.488261 | 3.604388 | 4.680275 | 3.595567 | 1.389679 | 4.667942 | 10.44036 | | 33.5972222 | 2.229783 | 0.453714 |
| 33.5208333 | 2.142435 | 3.98726 | 2.036395 | -0.827932 | 1.115421 | 1.772937 | 0.180712 | 1.777964 | 1.33348 | 2.439102 | 3.724086 | 3.438884 | 4.273116 | 3.100499 | 4.239713 | 6.75304 | 2.227569 | 3.581423 | 2.863848 | 5.693058 | 5.308333 | 3.765642 | 5.736175 | 16.972745 | | 33.6111111 | 2.441615 | 0.256083 |
| 33.5347222 | 5.678169 | 5.073998 | 0.988773 | -1.497881 | 1.700437 | 2.13796 | 0.836326 | 2.218523 | 1.482906 | 1.905277 | 7.574389 | 3.667851 | 5.12338 | 4.449446 | 3.708877 | 5.75516 | 1.749252 | 1.708516 | 2.220367 | 6.186487 | 7.285166 | 5.90702 | 3.743084 | 17.581694 | | 33.625 | 3.755405 | 0.42518 |
| 33.5486111 | 5.122059 | 4.372876 | 0.267979 | 0.019264 | 1.211426 | 0.712836 | 1.279563 | 1.617078 | 0.972435 | 2.056002 | 7.481222 | 2.233907 | 4.871592 | 3.650905 | 3.646095 | 2.963086 | 2.368623 | 2.863737 | 3.589546 | 5.421837 | 7.734368 | 6.481418 | 3.959817 | 13.425452 | | 33.6388889 | 1.753989 | 1.281033 |
| 33.5625 | 4.484582 | 4.578366 | 0.772481 | 1.403434 | 0.409768 | 0.56464 | 0.811606 | 1.416297 | 0.619169 | 0.872402 | 6.217107 | 1.668773 | 2.93209 | 2.937126 | 3.572306 | 3.77179 | 3.6488 | 4.510025 | 3.830023 | 3.703226 | 3.717456 | 5.338318 | 1.283989 | 7.657056 | | 33.6527778 | 0.626122 | 1.757399 |
| 33.5763889 | 2.486446 | 4.359051 | 1.672743 | 1.331541 | 0.562446 | 1.320471 | 0.66766 | 0.721821 | 0.624789 | 0.170607 | 3.734857 | 1.504461 | 2.736846 | 2.03448 | 1.926166 | 1.352786 | 3.469986 | 3.535725 | 3.20389 | 0.717837 | -0.356629 | 4.521327 | -0.588576 | 1.725613 | | 33.6666667 | -1.860232 | 1.265356 |
| 33.5902778 | 0.16939 | -0.082102 | 1.792384 | 0.978047 | 0.666036 | 1.573431 | 0.353884 | -0.727097 | 0.395492 | -0.106798 | 0.21423 | 1.013221 | 1.934089 | 1.567106 | 1.268124 | -0.991374 | 1.020198 | 1.058063 | 1.388151 | -1.995444 | -4.085344 | 1.151582 | -3.06075 | -1.388949 | | 33.6805556 | -0.378737 | 0.29321 |
| 33.6041667 | -0.93856 | -2.606865 | 0.934736 | 1.227842 | 0.378731 | 0.225206 | -0.092095 | -1.984274 | -0.041361 | 0.305289 | -2.347973 | 0.557741 | -0.177039 | 0.633823 | 1.536665 | 0.069334 | -1.569945 | -0.66546 | -0.333727 | -2.666718 | -3.046329 | -5.714497 | -8.4644 | -4.752697 | | 33.6944444 | 2.203619 | -0.489622 |
| 33.6180556 | -2.81414 | -3.58741 | 0.398963 | 0.666487 | -0.032523 | -1.201929 | -0.756364 | -2.924714 | 0.037653 | 0.722271 | -4.101843 | 0.334437 | -1.323786 | -0.355486 | -0.713707 | -0.129676 | -2.699353 | -1.930939 | -1.605699 | -1.903595 | -1.55776 | -10.270618 | -14.082181 | -6.842625 | | 33.7083333 | 4.472593 | -0.292125 |
| 33.6319444 | -5.183061 | -3.471899 | 0.177604 | -0.307276 | -0.520144 | -1.387325 | -1.024871 | -2.385946 | 0.546843 | 1.554149 | -5.244609 | 0.15086 | 0.792428 | -1.356088 | -1.990278 | -0.72347 | -0.668453 | -0.449107 | -1.98912 | -1.118627 | 1.505264 | -4.483077 | -5.374419 | -2.035943 | | 33.7222222 | -1.50255 | 0.074663 |
| 33.6458333 | -4.071685 | -2.314128 | 0.52762 | -0.211646 | -0.602311 | -1.06065 | -1.251639 | -1.069618 | 0.44123 | 1.377518 | -3.991715 | -0.293215 | 2.426531 | -0.542632 | -0.654268 | 0.552215 | 0.650058 | 2.037602 | -0.46859 | -0.347726 | 2.705033 | 1.578743 | 0.769747 | 4.769869 | | 33.7361111 | -0.966888 | 0.641327 |
| 33.6597222 | 0.460359 | 0.091033 | 1.140866 | 0.617041 | -0.543221 | -0.899047 | 0.594132 | 0.965098 | -0.19777 | -0.454888 | 2.523173 | -0.639569 | 0.748939 | 0.9476 | 1.806733 | 2.234151 | 0.76053 | 3.130233 | 2.264047 | 2.780689 | 0.714798 | 2.446739 | 2.006358 | 3.922916 | | 33.75 | -0.729503 | 0.955074 |
| 33.6736111 | 4.667642 | 3.817092 | 1.642954 | 1.512029 | -0.048191 | 0.145222 | 0.073239 | 1.051519 | -0.602866 | -1.85893 | 6.597216 | -0.914239 | -1.197745 | 0.687312 | 1.428645 | 0.76776 | -0.457968 | -0.521259 | 1.006384 | 0.530037 | -3.747089 | 0.284128 | -1.400723 | -3.237033 | | 33.7638889 | 0.664381 | 0.723693 |
| 33.6875 | 6.549825 | 6.63364 | 0.500975 | 0.913629 | 0.382235 | 0.945528 | 0.063862 | 0.156289 | -0.360109 | -1.785499 | 7.823308 | -1.609473 | -2.499895 | -1.891475 | -1.760601 | -2.963447 | -2.039573 | -2.937215 | -1.625294 | -3.644757 | -6.002362 | -0.918888 | 2.87003 | -9.627714 | | 33.7777778 | 0.62716 | 0.347083 |
| 33.7013889 | 4.501327 | 6.493911 | 0.068559 | -0.447224 | 0.131085 | 0.598541 | 0.086421 | -0.683635 | 0.197521 | -0.583269 | 6.561287 | -0.727241 | -3.818104 | -5.561789 | -2.979535 | -3.812343 | -0.732692 | -2.155989 | -4.034901 | -8.880034 | -7.172083 | -2.714436 | 2.524059 | -13.84379 | | 33.7916667 | 0.148497 | 0.076394 |
| 33.7152778 | 2.923822 | 4.519152 | 0.128161 | -0.714973 | -0.091717 | -0.086119 | 0.220717 | -1.110633 | 0.739388 | -0.474855 | 3.27574 | 0.874073 | -1.696503 | -5.235247 | -3.636954 | -2.172529 | 0.395861 | -0.13594 | -4.471609 | -4.692534 | -2.456322 | -2.289116 | 0.95917 | -5.985666 | | 33.8055556 | -0.399277 | -0.121743 |
| 33.7291667 | 1.070346 | 1.259322 | 0.86069 | -0.220972 | -0.169192 | 0.190783 | -0.176327 | -1.158395 | 0.800159 | 0.519783 | -0.335053 | 0.380219 | -1.727625 | -0.789453 | -1.973313 | -0.263435 | 1.658426 | 1.152528 | -2.033936 | -0.677491 | -0.136421 | -2.757385 | 3.326474 | 3.814273 | | 33.8194444 | -1.912246 | -0.525766 |
| 33.7430556 | -1.05016 | -1.439749 | 1.466484 | 0.400356 | 0.059949 | 0.233212 | -0.878647 | -1.089246 | 0.257586 | 0.675143 | -2.183158 | -0.360765 | -0.224011 | 5.067531 | 2.473095 | 2.07547 | 1.058401 | -0.387376 | 2.022293 | 2.312798 | 1.835984 | -2.594408 | 0.924312 | 6.481816 | | 33.8333333 | -2.429956 | -1.898725 |
| 33.7569444 | -2.477896 | -3.743644 | 0.445417 | 0.095711 | 0.292663 | 0.166135 | -1.928048 | -0.389446 | -0.84107 | -0.236242 | -3.716539 | -1.502462 | -0.235523 | 5.3112 | 4.494692 | 3.294289 | -1.012557 | -2.643931 | 2.061171 | 2.988319 | 0.047831 | 0.569553 | -1.147085 | 2.906672 | | 33.8472222 | -3.079195 | -2.157799 |
| 33.7708333 | -2.190216 | -5.301241 | -1.224531 | -0.287794 | 0.50411 | 0.676038 | -1.883712 | 0.238731 | -1.553473 | -0.787457 | -5.114998 | -1.553376 | -1.085759 | 2.334207 | 3.706676 | 2.449982 | -1.183281 | -1.690785 | 0.684614 | -0.518927 | -3.411333 | 3.806636 | -0.81567 | -2.775397 | | 33.8611111 | -4.194574 | -1.80378 |
| 33.7847222 | -4.341719 | -8.036996 | -2.748403 | -1.047577 | 0.053912 | -0.020985 | -0.23962 | 0.952079 | -1.501377 | 0.206242 | -6.423562 | -3.029683 | -4.405051 | 0.082173 | 2.353382 | 0.715873 | 0.205281 | 1.330172 | 1.540838 | -2.469643 | -1.842823 | 5.786729 | 0.782084 | -2.874226 | | 33.875 | -4.55254 | -1.492284 |
| 33.7986111 | -7.101876 | -9.120001 | -4.230923 | -1.035623 | -0.346769 | -0.71827 | 0.471242 | 0.655948 | -0.547823 | 1.268467 | -6.954364 | -2.351533 | -4.09448 | -1.130729 | -0.551141 | -2.356839 | 1.104868 | 1.752317 | 1.046944 | -0.601677 | 0.101383 | 1.739874 | 1.500458 | -1.343793 | | 33.8888889 | -3.328021 | -1.395909 |
| 33.8125 | -6.847825 | -6.085034 | -3.607703 | 0.227348 | -0.665186 | -0.586231 | 0.40285 | 0.937451 | 0.147184 | 1.869484 | -7.162059 | 0.598915 | 0.951073 | -2.494762 | -2.321025 | -1.416325 | 1.283606 | 1.022728 | 1.357997 | 3.270057 | 0.621332 | -2.231104 | -0.844173 | -1.662896 | | 33.9027778 | -1.967473 | -1.517859 |
| 33.8263889 | -7.717634 | -4.330889 | -1.992638 | 0.669805 | -0.718721 | -0.894896 | -0.830197 | -1.023548 | -0.056966 | 0.341459 | -7.302438 | 2.259876 | 3.474877 | -3.862338 | -5.517258 | -0.881509 | 0.277592 | -1.091237 | 1.145222 | 5.927533 | 1.498817 | -5.293926 | -0.109417 | -2.521571 | | 33.9166667 | -1.635512 | -1.709473 |
| 33.8402778 | -7.823207 | -3.757938 | -1.692449 | -0.286494 | -0.775672 | -1.490659 | -2.080107 | -1.968526 | -1.158508 | -2.550272 | -7.278534 | 1.327581 | 1.131837 | -2.500833 | -3.254988 | 0.916187 | -0.230685 | -2.156365 | 0.019664 | 4.686324 | 0.841042 | -8.148932 | -1.538182 | -1.808442 | | 33.9305556 | -2.460759 | -1.939564 |
| 33.8541667 | -9.167665 | -4.675008 | -1.999482 | -1.220309 | -0.116543 | -1.222601 | -2.728324 | -2.137385 | -2.642841 | -4.088511 | -8.558039 | -2.212461 | -3.325519 | 0.154264 | 1.529732 | 0.897289 | -1.490715 | -2.353279 | -2.189346 | -1.019554 | 0.096843 | -7.665739 | -4.189474 | -4.344439 | | 33.9444444 | -3.44628 | -1.969249 |
| 33.8680556 | -10.912535 | -5.926009 | -2.852435 | -2.445014 | 0.538819 | -0.134939 | -2.870689 | -2.13091 | -2.140106 | -2.808991 | -8.254363 | -7.897394 | -7.998058 | -0.752173 | 2.883852 | 1.7303 | -2.157456 | -0.874513 | -7.314599 | -6.153184 | -0.859274 | -1.434373 | -2.110786 | -8.124272 | | 33.9583333 | -4.276492 | -1.814388 |
| 33.8819444 | -9.950004 | -5.737376 | -2.875286 | -2.164824 | -0.012939 | 0.095992 | -0.912669 | -1.421917 | -0.866452 | -1.094553 | -6.500634 | -7.052087 | -2.229868 | -4.254398 | -0.242113 | 1.493082 | -2.71942 | -1.515664 | -5.994389 | -6.953603 | -2.01203 | 1.684705 | -1.982686 | -8.325618 | | 33.9722222 | -5.956753 | -1.590727 |
| 33.8958333 | -6.95502 | -4.155006 | -3.12102 | -1.920805 | -0.374099 | 0.088468 | 0.361674 | -1.563718 | -0.586964 | -1.638276 | -4.766457 | -4.95592 | -0.361278 | -6.682004 | -3.812195 | 0.487188 | -3.327094 | -1.997718 | -0.524756 | -2.187051 | -2.457272 | 2.721105 | -3.524562 | -8.624732 | | 33.9861111 | -5.432845 | -1.481545 |
| 33.9097222 | -4.279492 | -3.156112 | -3.020171 | -1.732177 | -0.801056 | -1.353986 | -0.783361 | -2.676229 | -1.631833 | -3.523618 | -6.230183 | -4.272926 | -4.187711 | -8.044337 | -7.353564 | -2.188258 | -4.852273 | -4.876613 | 3.648281 | -0.038791 | -3.379486 | -0.414949 | -4.739889 | -9.773247 | | 34 | -3.967293 | -1.478287 |
| 33.9236111 | -1.61159 | -2.277342 | -3.054942 | -1.925831 | -2.033486 | -3.637573 | -2.927296 | -3.577167 | -3.111899 | -5.833429 | -8.807543 | -6.894526 | -6.801414 | -7.956097 | -8.755794 | -2.90463 | -6.399669 | -6.971476 | -1.260323 | -3.912642 | -4.381819 | -7.481736 | -3.747679 | -13.609809 | | 34.0138889 | -3.44758 | -1.277042 |
| 33.9375 | -0.280643 | -3.157507 | -4.920648 | -3.05706 | -3.354585 | -4.876806 | -5.185502 | -4.109038 | -6.168815 | -7.714013 | -19.374396 | -10.499606 | -7.228279 | -5.291406 | -7.071067 | -1.881017 | -7.498288 | -5.758479 | -9.050278 | -8.118172 | -4.112204 | -15.568552 | -5.426802 | -23.865266 | | 34.0277778 | -5.36681 | -0.479615 |
| 33.9513889 | -1.816165 | -3.664186 | -6.2652 | -3.188327 | -4.249377 | -5.420218 | -7.048961 | -4.516187 | -10.37346 | -9.514224 | -34.911849 | -10.937121 | -8.771883 | -3.955712 | -5.597371 | -3.880923 | -9.747516 | -6.134191 | -16.465777 | -8.752723 | -1.816746 | -16.389422 | -7.40986 | -34.517454 | | 34.0416667 | -8.564904 | -1.14101 |
| 33.9652778 | -4.573276 | -4.171577 | -5.404672 | -2.78692 | -3.613718 | -4.782756 | -6.485743 | -4.767384 | -7.948198 | -7.031663 | -33.205108 | -14.742631 | -10.681016 | -6.156013 | -6.137167 | -4.138236 | -10.95494 | -5.046249 | -7.867449 | -5.611855 | -1.887567 | -12.543553 | -6.927972 | -29.605346 | | 34.0555556 | -8.711798 | -1.167836 |
| 33.9791667 | -5.408353 | -3.287093 | -2.806894 | -2.439633 | -2.540227 | -4.095103 | -3.836901 | -3.760872 | -3.23663 | -3.456853 | -16.563394 | -14.856719 | -9.11628 | -8.571014 | -6.859982 | -2.297064 | -10.787643 | -2.171415 | -0.715106 | -3.815215 | -1.992665 | -5.497219 | -5.511073 | -16.32181 | | 34.0694444 | -6.602838 | -0.535881 |
| 33.9930556 | -2.799224 | -1.575453 | -1.33238 | -1.761978 | -2.099635 | -3.304876 | -1.904554 | -2.247724 | -0.767451 | -1.14891 | -4.764902 | -7.875536 | -4.155148 | -5.920753 | -4.194805 | -0.272118 | -9.298546 | -0.965017 | -0.36098 | -4.48621 | -1.507137 | -3.648338 | -2.833505 | -7.054214 | | 34.0833333 | -5.850898 | -0.744557 |
| 34.0069444 | -0.547196 | -0.459251 | -0.876599 | -1.027565 | -1.474334 | -1.783425 | -1.360842 | -1.436282 | -0.359381 | -0.671296 | -1.402135 | -1.693978 | -0.378302 | -2.832028 | -2.321363 | -0.051554 | -5.990312 | -1.163184 | -1.595138 | -3.647776 | 0.150243 | -3.892677 | -1.102542 | -3.263341 | | 34.0972222 | -5.469273 | -0.672611 |
| 34.0208333 | 0.670568 | -0.166061 | -0.665128 | -0.599253 | -0.700073 | -0.530143 | -1.140586 | -1.063031 | -0.592932 | -0.888457 | -0.84987 | -0.814285 | 0.30152 | -1.504519 | -1.119288 | -0.205858 | -3.090053 | -0.457788 | -0.884024 | -0.766105 | 0.468551 | -1.694034 | -0.768821 | -2.166073 | | 34.1111111 | -3.030397 | -0.15936 |
| 34.0347222 | 0.874753 | -0.272858 | -0.495475 | -0.279572 | -0.154384 | -0.034463 | -0.844521 | -0.833863 | -0.687242 | -1.005768 | -0.457351 | -1.386353 | -1.023375 | -1.685328 | -1.543691 | -0.907045 | -1.196824 | 0.554504 | -2.542527 | -1.426472 | -0.74946 | -0.964146 | -1.268654 | -3.027685 | | 34.125 | -0.688814 | 0.390214 |
| 34.0486111 | 0.143375 | -0.481702 | -0.433308 | -0.098106 | -0.034052 | -0.165743 | -0.679973 | -0.824191 | -0.588874 | -0.861232 | -0.459954 | -2.738563 | -2.077116 | -2.993705 | -3.283386 | -2.518577 | -1.857739 | -0.433622 | -3.860661 | -2.062512 | -1.640052 | -2.815187 | -1.870589 | -4.39705 | | 34.1388889 | 0.606118 | 0.678218 |
| 34.0625 | -0.966417 | -0.74949 | -0.558219 | -0.174174 | -0.138194 | -0.513505 | -0.686472 | -0.938237 | -0.474732 | -0.780513 | -0.853887 | -4.472938 | -1.983247 | -3.40385 | -3.34191 | -3.285759 | -3.650231 | -1.74454 | -4.028335 | -2.375188 | -2.145272 | -5.561604 | -2.353716 | -5.121195 | | 34.1527778 | 1.880277 | 0.671527 |
| 34.0763889 | -1.581411 | -0.950529 | -0.667445 | -0.328588 | -0.294992 | -0.768405 | -0.661527 | -0.973432 | -0.370039 | -0.749866 | -1.030218 | -5.352594 | -1.81257 | -3.459874 | -2.832859 | -3.307472 | -4.264624 | -2.009909 | -4.550664 | -2.474812 | -2.540036 | -6.730487 | -2.290492 | -4.669394 | | 34.1666667 | 2.877621 | 0.663924 |
| 34.0902778 | -1.694412 | -1.04387 | -0.651584 | -0.298455 | -0.38416 | -0.80212 | -0.625492 | -0.98374 | -0.317321 | -0.761843 | -1.13612 | -5.206105 | -1.936104 | -3.966453 | -2.616167 | -2.574532 | -2.799203 | -1.377292 | -3.41393 | -1.47782 | -2.314617 | -6.426926 | -2.07739 | -3.760224 | | 34.1805556 | 2.746763 | 0.744553 |
| 34.1041667 | -1.67716 | -1.181941 | -0.607426 | -0.193592 | -0.411054 | -0.689652 | -0.735604 | -1.094911 | -0.274334 | -0.799566 | -1.267922 | -3.89291 | -2.05324 | -2.926419 | -1.722745 | -2.063461 | -1.101371 | -1.065579 | -2.969126 | -0.816183 | -1.741823 | -5.364309 | -1.975973 | -2.756279 | | 34.1944444 | 1.737588 | 0.903514 |
| 34.1180556 | -1.833691 | -1.500024 | -0.687894 | -0.381044 | -0.467618 | -0.725825 | -0.966825 | -1.280304 | -0.344453 | -0.922662 | -1.470288 | -2.77693 | -1.727776 | -1.02591 | -0.235725 | -1.319882 | 0.203118 | -0.687581 | -2.591699 | -0.56592 | -1.00073 | -3.879488 | -1.822233 | -1.945251 | | 34.2083333 | 0.782694 | 0.659297 |
| 34.1319444 | -2.006462 | -1.76339 | -0.819976 | -0.682503 | -0.557963 | -1.016403 | -1.214493 | -1.523948 | -0.592947 | -1.119095 | -1.839931 | -1.984572 | -1.263207 | -0.184499 | 0.712107 | -0.239829 | 0.378891 | -0.192266 | -0.864885 | -0.803186 | -0.635162 | -2.091613 | -1.374919 | -1.232883 | | 34.2222222 | 0.540487 | 0.061554 |
| 34.1458333 | -2.14726 | -1.70991 | -0.749596 | -0.736752 | -0.512454 | -1.048686 | -1.362309 | -1.713682 | -0.748318 | -1.167886 | -2.120742 | -1.184474 | -0.803861 | -0.602015 | 0.493158 | 0.285855 | 0.347317 | 0.402808 | 0.180746 | -0.923562 | -0.3878 | -0.646671 | -0.537103 | -0.533538 | | 34.2361111 | 0.999782 | -0.191179 |
| 34.1597222 | -2.136772 | -1.453112 | -0.610682 | -0.513804 | -0.30518 | -0.6458 | -1.351952 | -1.799025 | -0.5963 | -0.733185 | -1.870398 | -0.442324 | -0.455727 | 0.00942 | 0.759147 | -0.001671 | 0.954167 | 0.744268 | -0.37572 | -0.667264 | 0.129885 | 0.085849 | 0.531766 | 0.214855 | | 34.25 | 1.746446 | -0.16073 |
| 34.1736111 | -2.196954 | -1.115507 | -0.579765 | -0.390287 | -0.06011 | -0.049688 | -1.142485 | -1.681479 | -0.430347 | -0.203919 | -1.517259 | -0.148953 | -0.420838 | 1.614803 | 2.056571 | -0.08703 | 1.905545 | 0.637723 | -0.867971 | -0.687459 | 0.306723 | 0.423091 | 1.069834 | 0.957226 | | 34.2638889 | 2.233692 | -0.17652 |
| 34.1875 | -2.325195 | -0.782512 | -0.597987 | -0.284938 | 0.143914 | 0.540066 | -0.951902 | -1.318972 | -0.409616 | 0.021951 | -1.416289 | 0.07956 | -0.483225 | 1.709823 | 2.160943 | 0.392942 | 2.008593 | 0.511491 | -0.463192 | -0.857376 | -0.009557 | 1.082722 | 1.061655 | 1.192853 | | 34.2777778 | 2.152891 | 0.191683 |
| 34.2013889 | -1.966022 | -0.374242 | -0.498166 | -0.103227 | 0.299802 | 0.892124 | -0.775587 | -0.838685 | -0.330594 | 0.119792 | -1.044335 | 0.19546 | -0.318894 | 0.16371 | 0.56528 | 0.675456 | 1.088259 | 0.568993 | 0.257798 | -1.058227 | -0.395804 | 1.821554 | 0.979976 | 0.919031 | | 34.2916667 | 1.838967 | 1.068486 |
| 34.2152778 | -1.626093 | -0.036095 | -0.39821 | -0.085811 | 0.346541 | 0.912081 | -0.63756 | -0.504192 | -0.212242 | 0.203787 | -0.611195 | -0.18482 | -0.195159 | -0.637857 | -0.21819 | 0.519011 | 0.238729 | 0.394032 | 0.31988 | -1.034015 | -0.374657 | 1.457812 | 1.068062 | 0.303869 | | 34.3055556 | 1.24727 | 1.494476 |
| 34.2291667 | -1.319812 | 0.146513 | -0.42235 | -0.202335 | 0.341473 | 0.856111 | -0.530032 | -0.364755 | -0.126193 | 0.272475 | -0.680404 | -0.297316 | -0.21769 | -0.015419 | 0.555898 | 0.343789 | 0.65617 | 0.118927 | 0.123816 | -0.637505 | 0.00389 | 0.799082 | 1.117899 | 0.295742 | | 34.3194444 | 0.832982 | 1.343852 |
| 34.2430556 | -0.694318 | 0.320552 | -0.369332 | 0.013055 | 0.533876 | 1.111031 | -0.324434 | -0.105887 | -0.04674 | 0.37064 | -0.571756 | 0.666215 | 0.02213 | 0.621168 | 1.562768 | 0.507956 | 2.508071 | 0.099723 | 0.677509 | -0.507264 | 0.344636 | 1.522745 | 0.982033 | 1.115736 | | 34.3333333 | 1.147596 | 1.030289 |
| 34.2569444 | 0.34055 | 0.720192 | -0.135627 | 0.487088 | 1.102085 | 1.81134 | 0.027189 | 0.305988 | 0.051473 | 0.471334 | 0.306372 | 2.086582 | 0.864743 | 0.368099 | 1.574737 | 0.936858 | 4.041324 | 0.438764 | 0.605863 | -0.453271 | 0.432366 | 2.821592 | 0.975955 | 1.76481 | | 34.3472222 | 2.066645 | 0.827529 |
| 34.2708333 | 1.212636 | 0.983189 | 0.032616 | 0.634332 | 1.451517 | 2.144708 | 0.41338 | 0.596693 | -0.002959 | 0.459518 | 1.044199 | 2.940508 | 1.766418 | 0.2471 | 1.927024 | 1.677973 | 4.232306 | 0.897318 | 0.355369 | -0.395161 | 0.678608 | 2.671144 | 1.010366 | 2.214979 | | 34.3611111 | 3.201641 | 0.577207 |
| 34.2847222 | 1.710093 | 0.920244 | 0.109689 | 0.419616 | 1.387227 | 1.776393 | 0.649028 | 0.786299 | -0.102312 | 0.43424 | 1.237137 | 3.568014 | 2.13534 | 1.369156 | 3.042503 | 2.172442 | 3.685413 | 0.973018 | 0.890457 | 0.55273 | 1.306846 | 2.365845 | 1.142011 | 2.831616 | | 34.375 | 3.543155 | 0.149657 |
| 34.2986111 | 2.821741 | 0.889435 | 0.289572 | 0.503547 | 1.571103 | 1.825003 | 0.776775 | 1.149469 | 0.003407 | 0.579144 | 1.712576 | 4.591103 | 2.432961 | 2.143233 | 3.179051 | 2.237448 | 3.751739 | 1.227747 | 1.947027 | 1.464151 | 1.936452 | 2.972659 | 1.649756 | 3.567894 | | 34.3888889 | 2.594577 | -0.405175 |
| 34.3125 | 3.757274 | 1.141117 | 0.556425 | 0.922042 | 2.003022 | 2.410712 | 1.052206 | 1.612732 | 0.2332 | 0.850613 | 2.698076 | 5.075769 | 3.149535 | 1.935287 | 2.700359 | 2.650299 | 4.787465 | 1.908815 | 1.604159 | 1.801209 | 2.18442 | 3.987371 | 2.163135 | 4.187225 | | 34.4027778 | 1.465736 | -0.5434 |
| 34.3263889 | 4.116554 | 1.407388 | 0.803468 | 1.246144 | 2.279388 | 3.147431 | 1.323733 | 1.969556 | 0.433826 | 1.142807 | 3.522564 | 5.563722 | 3.884487 | 2.081296 | 3.008888 | 3.606079 | 6.210522 | 2.680366 | 0.876664 | 1.761609 | 2.369688 | 4.112819 | 2.487531 | 4.718864 | | 34.4166667 | 0.884793 | 0.383217 |
| 34.3402778 | 4.950127 | 1.577959 | 1.094199 | 1.313 | 2.179488 | 3.573129 | 1.699526 | 2.342741 | 0.70309 | 1.566914 | 4.225253 | 6.934244 | 4.094154 | 3.34232 | 4.380988 | 4.313107 | 5.532082 | 3.044798 | 0.969346 | 2.037429 | 2.596376 | 4.497949 | 3.191714 | 5.569648 | | 34.4305556 | 0.902288 | 1.780551 |
| 34.3541667 | 5.395253 | 1.786036 | 1.396908 | 1.579357 | 2.181033 | 3.934998 | 2.080423 | 2.774043 | 1.098286 | 2.18297 | 4.950809 | 8.726053 | 4.336103 | 4.216 | 4.805809 | 4.700646 | 4.373621 | 2.857559 | 2.152426 | 2.986676 | 3.128266 | 5.8342 | 4.516647 | 6.598113 | | 34.4444444 | 1.192925 | 2.453004 |
| 34.3680556 | 5.65683 | 2.486094 | 1.734187 | 2.065537 | 2.348265 | 4.844068 | 2.477805 | 3.301984 | 1.762017 | 3.049351 | 5.889613 | 10.647522 | 5.033155 | 4.548005 | 4.491247 | 5.102436 | 4.789827 | 3.729184 | 3.510071 | 4.263375 | 3.768495 | 7.889633 | 6.154186 | 8.005426 | | 34.4583333 | 1.444812 | 2.263139 |
| 34.3819444 | 6.292429 | 3.658977 | 2.083366 | 2.441479 | 2.608704 | 4.910797 | 2.747497 | 3.760611 | 2.436539 | 4.13073 | 7.214427 | 11.564837 | 6.113095 | 4.481238 | 4.020181 | 4.567465 | 6.184872 | 5.269922 | 4.518182 | 5.701721 | 4.393312 | 10.299865 | 7.195353 | 10.389454 | | 34.4722222 | 1.676755 | 1.628715 |
| 34.3958333 | 6.706705 | 4.84067 | 2.621311 | 2.655843 | 2.988379 | 5.18139 | 3.057115 | 4.25147 | 2.890093 | 5.069258 | 8.244749 | 13.213445 | 7.006448 | 6.258875 | 4.375878 | 4.343561 | 7.266424 | 6.477797 | 4.455978 | 5.93853 | 5.538872 | 14.151248 | 7.407045 | 13.546896 | | 34.4861111 | 1.907168 | 1.340989 |
| 34.4097222 | 8.32243 | 6.1383 | 3.253358 | 2.901713 | 2.13793 | 4.013808 | 2.609618 | 3.88839 | 3.23395 | 5.539859 | 9.316592 | 13.940979 | 7.545446 | 7.691517 | 6.158677 | 5.455942 | 6.816641 | 6.74733 | 5.196225 | 6.126733 | 6.23635 | 17.068911 | 8.228564 | 17.017263 | | 34.5 | 1.965514 | 1.374321 |
| 34.4236111 | 9.641508 | 6.984325 | 3.788197 | 3.354728 | 1.592503 | 4.204334 | 2.547751 | 3.76774 | 3.578293 | 6.041487 | 11.433259 | 13.167496 | 8.150771 | 6.948262 | 6.118099 | 6.311918 | 5.818344 | 5.949276 | 6.83201 | 7.358748 | 5.886641 | 18.04958 | 8.1409 | 19.420612 | | 34.5138889 | 1.615746 | 1.740439 |
| 34.4375 | 9.943726 | 7.625579 | 4.711051 | 3.828846 | 1.609366 | 3.793051 | 2.927072 | 3.257492 | 4.792929 | 7.030488 | 21.008393 | 14.762026 | 9.839189 | 5.706447 | 4.816889 | 5.155667 | 5.898453 | 5.88973 | 8.780887 | 8.847337 | 6.022136 | 20.251036 | 10.402859 | 31.039314 | | 34.5277778 | 3.050717 | 3.321628 |
| 34.4513889 | 9.782895 | 6.739326 | 5.216068 | 3.173108 | 2.412103 | 3.25547 | 3.647563 | 3.090422 | 7.191131 | 7.967993 | 35.855823 | 14.397862 | 11.197697 | 5.991639 | 4.141662 | 4.390268 | 7.965443 | 6.4337 | 10.879541 | 11.286501 | 7.965032 | 19.363043 | 10.039859 | 51.745374 | | 34.5416667 | 6.838663 | 4.976589 |
| 34.4652778 | 9.072558 | 5.620805 | 4.631294 | 1.482874 | 1.997986 | 2.265787 | 3.784025 | 2.367357 | 7.682116 | 5.894417 | 40.02541 | 8.488078 | 7.193211 | 7.291533 | 5.224731 | 3.815519 | 7.258394 | 6.239331 | 9.801318 | 9.78125 | 5.832885 | 15.584318 | 7.428328 | 47.172935 | | 34.5555556 | 9.004116 | 4.004444 |
| 34.4791667 | 3.190666 | 1.810978 | 3.149394 | 0.592921 | 0.869063 | 1.295078 | 1.930057 | 0.887919 | 3.443553 | 1.569754 | 15.677871 | -3.500212 | -1.234859 | 4.16812 | 4.142772 | 2.492791 | 2.570509 | 4.035507 | 3.188438 | 3.388571 | 0.727597 | 2.072166 | 2.990721 | 11.108893 | | 34.5694444 | 7.614197 | 1.0904 |
| 34.4930556 | -4.532528 | -1.54208 | 1.452694 | 0.47258 | 0.264706 | 1.04197 | -0.348882 | -0.421736 | 0.084223 | -1.038801 | -6.278062 | -8.02171 | -2.214946 | 3.18422 | 3.390167 | 1.079407 | -0.289051 | 2.640742 | -1.364771 | -0.776137 | -0.243777 | -4.95864 | 1.740264 | -3.939209 | | 34.5833333 | 8.219038 | 0.04559 |
| 34.5069444 | -4.558533 | 0.433973 | 1.727801 | 1.134079 | 0.38118 | 1.504374 | -1.384001 | -0.070768 | -0.037197 | -1.638012 | -7.768217 | -1.343476 | 2.185202 | 4.586053 | 4.668093 | 2.727162 | 0.56771 | 4.060184 | -0.925576 | -0.158499 | 1.764966 | -2.615419 | 1.450172 | 5.468061 | | 34.5972222 | 6.318131 | -0.005149 |
| 34.5208333 | -0.019903 | 2.97526 | 1.991897 | 1.469443 | 0.29586 | 1.804192 | 0.239148 | 1.441667 | 1.374279 | 1.173292 | 2.115841 | 6.248267 | 6.239159 | 4.246388 | 4.547271 | 4.122826 | 0.822791 | 2.873073 | 3.343657 | 4.13141 | 3.855764 | 4.122241 | 3.357382 | 15.827809 | | 34.6111111 | 0.756305 | 0.118472 |
| 34.5347222 | 4.922304 | 5.873421 | 1.494155 | 1.279584 | -0.291748 | 1.862635 | 2.790323 | 2.633798 | 2.061759 | 3.737965 | 9.000858 | 9.618268 | 10.062598 | 2.54541 | 2.168252 | 2.211847 | 0.723774 | 1.839111 | 7.622201 | 9.728308 | 7.60128 | 8.969342 | 3.354528 | 20.289361 | | 34.625 | -5.159745 | 0.565937 |
| 34.5486111 | 6.664427 | 5.587566 | 0.183371 | 0.591057 | 0.228303 | 2.169194 | 1.560882 | 2.211858 | 1.221519 | 1.318071 | 11.660589 | 6.71564 | 9.195189 | 1.146892 | 1.804655 | 2.911018 | 1.244802 | 2.864126 | 6.740586 | 9.635478 | 4.610023 | 12.192552 | 2.392484 | 15.213682 | | 34.6388889 | -6.414012 | 1.121051 |
| 34.5625 | 6.801066 | 4.44956 | 0.014727 | 0.370745 | 0.378174 | 1.504238 | 0.859432 | 1.1306 | 1.166784 | 0.729103 | 10.230114 | 2.218341 | 5.003634 | 1.947329 | 2.233319 | 2.055621 | 1.479004 | 3.433065 | 4.651473 | 5.49582 | 2.926548 | 9.950576 | 2.922065 | 9.73517 | | 34.6527778 | -3.290693 | 1.056998 |
| 34.5763889 | 7.320794 | 5.875205 | -0.19934 | 0.349944 | 0.050171 | 0.579505 | 1.071052 | 1.184986 | 1.067945 | -0.211605 | 5.753798 | 0.908924 | 2.385494 | 2.38798 | 3.602628 | 2.83498 | 0.877319 | 1.884509 | 3.256642 | -0.58923 | -1.455934 | 7.89311 | 1.324609 | 2.51655 | | 34.6666667 | 1.079583 | 0.712428 |
| 34.5902778 | 3.009802 | 2.728227 | 0.693677 | 0.14999 | 0.176126 | 1.043596 | 0.511486 | 0.492418 | 0.510599 | -0.906334 | 1.53727 | 0.533055 | 0.05092 | 2.226442 | 3.037834 | 1.460703 | 0.218225 | -0.465496 | 1.120033 | -1.295229 | 0.354344 | 3.456113 | 1.92835 | 1.420067 | | 34.6805556 | 4.810565 | 1.004953 |
| 34.6041667 | -0.850328 | -3.07985 | 0.705821 | -0.26396 | 0.110626 | 0.165121 | -0.682592 | -0.833694 | 0.223662 | -1.505867 | -2.082106 | 0.380957 | -0.850662 | 1.393635 | 1.56228 | 0.8139 | 0.400127 | -1.232351 | -1.445169 | -0.608602 | 1.926459 | -1.626066 | -1.001556 | -0.616436 | | 34.6944444 | 5.268429 | 0.76815 |
| 34.6180556 | -2.961684 | -2.180489 | 0.054877 | -0.307856 | 0.157479 | -0.060785 | -1.697924 | -1.602904 | 0.455956 | 0.326783 | -5.489431 | 0.757109 | -0.173865 | 0.417177 | -0.784805 | -0.721305 | 0.547973 | -1.289154 | -3.574243 | -1.134649 | -1.133007 | -5.492324 | -1.967132 | -1.475332 | | 34.7083333 | 2.206585 | -0.05194 |
| 34.6319444 | -4.431442 | -2.023451 | 0.675392 | 0.17399 | 0.343549 | 0.117413 | -1.496334 | -1.006346 | 0.623322 | 1.549625 | -6.892566 | 0.927393 | 1.98131 | -0.809418 | -2.48627 | -2.394506 | 0.703408 | -0.280786 | -1.420388 | 0.451345 | -0.12572 | -4.444898 | -0.618329 | -1.286564 | | 34.7222222 | -1.677413 | -0.565199 |
| 34.6458333 | -4.905672 | -1.8358 | 1.717508 | 0.904625 | 0.354426 | 0.614859 | 1.494032 | 1.56334 | 0.375581 | 2.735315 | -1.091573 | 0.298113 | 1.339116 | -1.286099 | -2.205515 | -3.336135 | -0.42404 | 0.911183 | 1.48063 | 1.144994 | 1.932117 | -2.545937 | -0.021349 | 0.570425 | | 34.7361111 | -1.86928 | -0.655654 |
| 34.6597222 | -2.497611 | -1.102695 | 2.310586 | 1.741322 | 0.135475 | 0.619412 | 2.050537 | 2.724814 | -0.318337 | 0.615468 | 3.732325 | -0.831257 | -1.487526 | -0.142721 | -0.993582 | -2.817755 | -1.158781 | 1.698571 | 2.429246 | 0.022901 | 1.087087 | -0.511241 | 0.177394 | -0.596887 | | 34.75 | 1.340051 | -0.566681 |
| 34.6736111 | 1.007945 | 0.229161 | 1.89606 | 1.922457 | -0.474552 | 0.143303 | 0.784785 | 3.152226 | -1.098061 | -0.681724 | 3.250785 | -2.899695 | -4.302702 | 1.217623 | 1.399912 | 0.004711 | -1.251742 | 0.385122 | 0.75833 | -1.248111 | 0.370998 | 1.625513 | -2.21982 | -4.451504 | | 34.7638889 | 5.068426 | -0.372357 |
| 34.6875 | 4.247332 | 0.419133 | 0.491047 | 1.121656 | -0.505536 | -0.386896 | -0.498676 | 1.047972 | -1.317566 | -0.673389 | 4.071884 | -6.433155 | -9.321327 | 0.832144 | 1.632934 | 0.770734 | -0.031354 | -0.17818 | -0.712265 | 0.219305 | -0.759414 | 0.83077 | -4.534578 | -4.922887 | | 34.7777778 | 7.001926 | -0.151806 |
| 34.7013889 | 5.000239 | 0.980571 | -0.87105 | -0.223892 | -0.328484 | -0.152448 | -0.069647 | 0.826528 | -0.346066 | -0.134908 | 2.983243 | -5.069061 | -7.744675 | -0.282045 | 0.073604 | -1.008712 | 1.19811 | -1.497106 | -1.446816 | 0.463035 | -1.593924 | -1.999392 | -5.233728 | -5.352225 | | 34.7916667 | 7.851415 | -0.163434 |
| 34.7152778 | 1.16193 | 1.086076 | -2.459655 | -1.197409 | 0.057438 | 0.644996 | 0.481544 | 0.236379 | 0.961145 | 0.080735 | 0.283446 | -3.46805 | -5.157995 | -2.456283 | -0.860832 | -1.949216 | 0.993066 | 0.012084 | -1.88571 | -3.128467 | -2.529533 | -1.565122 | -1.18936 | -6.033002 | | 34.8055556 | 2.914991 | -0.308348 |
| 34.7291667 | -2.178257 | 0.050315 | -2.534565 | -1.067371 | 0.323725 | 1.074079 | 0.752761 | -0.18633 | 1.714711 | 0.469736 | -2.115891 | -0.232195 | -1.205565 | -4.895456 | -2.343765 | -1.98265 | 0.083722 | 1.635373 | -2.223361 | -4.969376 | -1.714505 | -1.002249 | 1.450179 | -5.554156 | | 34.8194444 | 0.07043 | -0.36313 |
| 34.7430556 | -1.117964 | 0.035834 | -0.706048 | -0.257928 | 0.668846 | 0.774797 | 0.448626 | -0.672335 | 1.183518 | 0.352461 | -2.921525 | 2.059371 | 2.197915 | -2.913742 | -1.58039 | -1.406295 | -1.215698 | 1.609281 | -2.940578 | -2.572527 | 0.009164 | -0.511077 | 2.931142 | -3.422838 | | 34.8333333 | -2.124583 | -0.370711 |
| 34.7569444 | 0.742704 | 0.730904 | 0.485443 | -0.368457 | 0.082511 | -0.640981 | -0.926989 | -2.254527 | 0.34261 | 0.181206 | -2.382073 | 0.824515 | 0.988583 | 0.695445 | 0.655955 | 1.710289 | -0.474813 | 0.474774 | -0.833194 | 1.144382 | 1.141778 | 0.785792 | -0.421831 | 1.605473 | | 34.8472222 | -5.56424 | -0.604349 |
| 34.7708333 | -0.358455 | 0.970262 | 0.475241 | -0.825062 | -0.351686 | -1.419841 | -1.674396 | -2.008888 | -0.522527 | -0.626241 | -3.725107 | -2.829708 | -2.814795 | 3.606315 | 2.867958 | -0.52306 | 0.952033 | -0.38075 | 2.259129 | 4.028314 | 0.015984 | -1.953148 | 0.657072 | 0.577083 | | 34.8611111 | -3.603394 | -1.053479 |
| 34.7847222 | -1.300672 | 0.603994 | 0.031166 | -1.273525 | -0.061274 | -0.341529 | -2.263814 | -1.526953 | -1.110759 | -1.322532 | -1.213546 | -4.555838 | -3.185179 | 3.505649 | 1.494315 | -3.256825 | 1.610032 | -0.184467 | 2.202761 | -0.948813 | -3.622113 | -4.513554 | 5.315027 | -2.830643 | | 34.875 | -2.737747 | -1.646948 |
| 34.7986111 | -1.713856 | -1.002047 | 0.319667 | -0.234114 | 0.354847 | 1.064295 | -1.760787 | -0.8424 | -1.135731 | -0.552475 | 0.460042 | -3.682245 | -3.260033 | 1.827763 | -0.806675 | -2.856019 | 2.045386 | 0.944344 | 1.868021 | -0.839862 | -2.356756 | 0.697536 | 0.205639 | -3.358362 | | 34.8888889 | -0.048416 | -2.262818 |
| 34.8125 | -3.356837 | -2.385766 | 1.537607 | 0.888572 | 0.193354 | 0.212207 | -0.690887 | -0.355354 | -0.597701 | 0.915946 | 1.123707 | -1.870283 | -2.232131 | -0.399834 | -0.78647 | -1.011035 | 1.428024 | 1.581613 | 0.470911 | -0.38526 | 0.101627 | 9.090633 | -1.364628 | 0.083538 | | 34.9027778 | 0.916448 | -2.778169 |
| 34.8263889 | -3.919619 | -3.287739 | 1.127753 | 0.544563 | 0.627582 | 0.26443 | 1.15089 | 0.282756 | -1.061598 | 1.741375 | -0.098933 | -0.644347 | -0.976964 | -2.54018 | -0.162492 | 1.944691 | -0.093048 | 0.635283 | -1.742181 | -2.622707 | -0.252087 | 10.281077 | 2.118734 | -0.347019 | | 34.9166667 | 0.148275 | -2.957674 |
| 34.8402778 | -4.97397 | -3.084149 | -0.604266 | -0.371041 | -0.057015 | -1.082996 | 1.293927 | 0.73257 | -1.442486 | -0.39076 | -4.256286 | -3.707617 | -0.253937 | -5.205231 | -0.808426 | 2.561323 | -2.174668 | -2.154926 | -4.644809 | -5.96061 | -0.006565 | -2.15935 | 1.908268 | -10.697643 | | 34.9305556 | -1.766996 | -2.90185 |
| 34.8541667 | -5.062237 | -4.068719 | -1.93133 | -0.896311 | -0.521457 | -2.435103 | -0.14432 | 0.533301 | -1.479314 | -1.193582 | -8.41004 | -4.58765 | -1.710899 | -3.436304 | -2.004647 | -0.922279 | -4.253672 | -4.616823 | -8.045899 | -9.294408 | -1.771519 | -13.879698 | 0.074908 | -16.519741 | | 34.9444444 | -4.938365 | -2.760242 |
| 34.8680556 | -5.837108 | -4.17607 | -2.399474 | -1.509042 | -0.381195 | -2.795828 | -1.909332 | 0.021952 | -1.111252 | -1.975075 | -10.390161 | -1.797529 | -2.149595 | 0.983179 | -1.388179 | -2.461605 | -3.916968 | -5.915629 | -2.70674 | -4.660621 | -0.394571 | -7.170586 | -0.905439 | -9.707285 | | 34.9583333 | -8.336095 | -2.611582 |
| 34.8819444 | -6.535732 | -3.937626 | -1.919205 | -1.776644 | -1.110029 | -1.700691 | -2.343177 | -0.809113 | -1.556357 | -2.530791 | -9.170382 | -1.499109 | -3.271687 | 4.160834 | -0.508973 | -4.224755 | -3.644795 | -5.104692 | 2.9895 | 1.535772 | -0.249012 | 2.338768 | -1.237751 | -3.837683 | | 34.9722222 | -8.332068 | -2.296585 |
| 34.8958333 | -6.708469 | -3.350951 | -2.196239 | -2.250121 | -0.885567 | -0.672047 | -2.304416 | -0.956208 | -1.733852 | -2.837093 | -5.172132 | -5.193081 | -4.464283 | 1.202922 | 0.109144 | -2.626641 | -3.081054 | -3.431971 | 1.656157 | 1.877871 | -0.294187 | 0.845953 | -2.248736 | -0.750518 | | 34.9861111 | -6.882188 | -2.00122 |
| 34.9097222 | -5.186502 | -2.309306 | -1.125152 | -0.610276 | 0.092622 | 0.526655 | -3.27667 | -2.649088 | -1.890984 | -3.238183 | -3.280749 | -11.016107 | -4.480997 | -2.793186 | -0.823625 | -0.833989 | -2.787148 | -3.030253 | -1.941565 | -1.059039 | -0.969491 | -4.300661 | -0.759462 | -5.576872 | | 35 | -4.192751 | -1.733299 |
| 34.9236111 | -2.734559 | -2.029102 | -0.614518 | -0.266646 | 0.074776 | -1.319469 | -3.291904 | -3.335121 | -2.174508 | -3.493544 | -5.003033 | -14.188225 | -6.341982 | -6.410333 | -4.040078 | -1.94287 | -4.046575 | -5.008287 | -5.876069 | -5.492973 | -2.517855 | -7.691975 | -1.522424 | -16.231951 | | 35.0138889 | -2.69633 | -1.400784 |
| 34.9375 | -2.306378 | -3.460955 | -1.311367 | -1.732141 | -1.6202 | -4.005867 | -2.363414 | -4.241289 | -4.121286 | -4.737851 | -21.526621 | -11.028418 | -7.64996 | -8.472233 | -7.704664 | -3.18051 | -9.912646 | -9.535481 | -7.464809 | -7.574305 | -5.508429 | -14.237601 | -6.437947 | -34.503116 | | 35.0277778 | -5.395422 | -1.874876 |
| 34.9513889 | -4.924817 | -6.175963 | -3.728438 | -2.956553 | -4.499968 | -7.820541 | -4.132272 | -5.675306 | -6.729547 | -5.732999 | -42.297465 | -6.449878 | -8.398655 | -10.472145 | -11.471451 | -7.625852 | -15.965137 | -13.28053 | -12.034332 | -9.435492 | -5.575997 | -21.202181 | -9.622074 | -44.680473 | | 35.0416667 | -12.146826 | -3.503056 |
| 34.9652778 | -5.487728 | -6.394525 | -4.582398 | -3.149817 | -5.21321 | -8.167345 | -4.078844 | -5.662815 | -6.276606 | -5.122671 | -35.934867 | -7.652219 | -6.985463 | -9.977204 | -9.687646 | -5.400358 | -12.991356 | -9.639778 | -7.410479 | -7.647044 | -3.286346 | -17.705976 | -8.888807 | -36.416975 | | 35.0555556 | -13.695383 | -2.013808 |
| 34.9791667 | -5.71876 | -4.597677 | -3.137371 | -2.451157 | -4.26615 | -6.134149 | -2.742469 | -3.554073 | -3.529788 | -3.429229 | -18.470194 | -9.755091 | -3.843681 | -7.600704 | -4.485503 | -1.847031 | -10.152796 | -4.086107 | -3.02052 | -4.801481 | -3.69907 | -9.725367 | -5.604315 | -17.885126 | | 35.0694444 | -10.957972 | -1.364962 |
| 34.9930556 | -4.287526 | -2.59212 | -2.079047 | -1.891977 | -2.649507 | -2.739699 | -1.735722 | -2.138842 | -1.772434 | -2.461244 | -7.042856 | -8.099467 | -2.115497 | -5.735646 | -5.202773 | -3.210877 | -7.790857 | -0.602862 | -1.211577 | -3.964513 | -2.652567 | -2.202588 | -3.500169 | -8.71097 | | 35.0833333 | -8.220285 | -0.586579 |
| 35.0069444 | -2.778006 | -1.28246 | -1.617922 | -1.221195 | -1.414391 | -0.342792 | -1.436924 | -1.527944 | -1.284251 | -1.771501 | -3.338898 | -6.462143 | -2.010906 | -6.717683 | -8.460377 | -4.555816 | -7.031663 | -1.372132 | -1.758654 | -2.92684 | -1.707777 | -1.654319 | -3.68755 | -8.157546 | | 35.0972222 | -5.824223 | -0.096764 |
| 35.0208333 | -1.690438 | -0.914061 | -1.302083 | -0.843327 | -0.931805 | 0.488881 | -1.451826 | -1.345001 | -0.90794 | -1.200748 | -1.971182 | -5.513883 | -2.214575 | -5.071655 | -5.180666 | -2.284459 | -4.404869 | -1.586788 | -3.082983 | -2.72662 | -0.885092 | -2.936736 | -3.92016 | -6.32941 | | 35.1111111 | -3.448547 | 0.276617 |
| 35.0347222 | -1.180646 | -0.988585 | -1.100663 | -0.695185 | -0.777524 | -0.016154 | -1.386992 | -1.456936 | -0.758514 | -1.032282 | -1.662855 | -5.956029 | -1.243478 | -1.758528 | 0.36965 | 0.561447 | -0.945512 | -0.554549 | -3.817973 | -2.465491 | -1.12525 | -2.203244 | -3.482563 | -2.528958 | | 35.125 | -1.664529 | 0.299421 |
| 35.0486111 | -1.158305 | -1.087838 | -0.867848 | -0.697653 | -0.461122 | -0.148048 | -1.338932 | -1.457152 | -0.691361 | -1.006197 | -1.415743 | -7.702919 | -2.077003 | -1.503072 | 0.60462 | 0.260123 | 0.307112 | -0.159975 | -2.530299 | -2.484384 | -1.665252 | -2.510353 | -3.689356 | -2.537096 | | 35.1388889 | -0.254802 | -0.125659 |
| 35.0625 | -1.274353 | -0.988151 | -0.610414 | -0.737718 | -0.160462 | -0.03745 | -1.371881 | -1.530758 | -0.558 | -0.882313 | -0.946529 | -8.572963 | -3.697417 | -2.551476 | -2.335797 | -2.204037 | -0.582399 | -0.876041 | -3.085249 | -3.496811 | -2.398986 | -4.011691 | -3.352845 | -4.209253 | | 35.1527778 | 1.313048 | -0.598586 |
| 35.0763889 | -1.338644 | -1.055115 | -0.51768 | -0.741818 | -0.07496 | 0.009814 | -1.282339 | -1.517569 | -0.502731 | -0.731453 | -0.874637 | -7.13828 | -3.692484 | -3.802036 | -5.229491 | -5.273235 | -2.460223 | -1.503169 | -3.489833 | -3.98979 | -3.098052 | -4.893873 | -2.590735 | -5.656263 | | 35.1666667 | 2.558192 | -0.741655 |
| 35.0902778 | -1.373347 | -1.310354 | -0.513534 | -0.660284 | -0.170545 | -0.162171 | -1.106523 | -1.434729 | -0.600273 | -0.72631 | -1.229087 | -4.598533 | -2.715058 | -3.689636 | -5.982625 | -5.326229 | -3.77925 | -1.808993 | -3.355933 | -2.805552 | -2.975768 | -4.815726 | -2.38406 | -5.677168 | | 35.1805556 | 2.724013 | -0.303368 |
| 35.1041667 | -1.387722 | -1.469489 | -0.472732 | -0.522229 | -0.407156 | -0.576259 | -1.128309 | -1.498707 | -0.817668 | -0.906616 | -1.77477 | -2.944504 | -1.759981 | -2.473043 | -5.855271 | -3.804193 | -5.824715 | -1.681947 | -0.497887 | -0.414901 | -2.167852 | -3.642193 | -2.10478 | -5.064165 | | 35.1944444 | 2.142739 | 0.494132 |
| 35.1180556 | -1.343311 | -1.324833 | -0.492108 | -0.764924 | -0.798824 | -1.151036 | -1.508183 | -1.835412 | -1.031894 | -1.195226 | -2.266513 | -1.373631 | -0.556899 | -1.824345 | -5.220944 | -2.749494 | -6.599425 | -0.891373 | 1.111204 | 0.693321 | -1.256619 | -2.313119 | -1.997862 | -3.690107 | | 35.2083333 | 1.120627 | 1.285951 |
| 35.1319444 | -1.329011 | -0.88611 | -0.569142 | -1.223782 | -1.171801 | -1.633932 | -1.572094 | -1.944783 | -1.069784 | -1.353792 | -2.309341 | 0.183126 | 0.342862 | -1.008438 | -2.584493 | -1.367751 | -4.538214 | -0.099445 | -0.836267 | 0.005592 | -0.471644 | -2.042146 | -1.954841 | -2.292502 | | 35.2222222 | -0.137821 | 1.444934 |
| 35.1458333 | -1.17137 | -0.222818 | -0.561592 | -1.407374 | -1.241032 | -1.693322 | -1.496553 | -1.882838 | -0.862149 | -1.159938 | -1.824466 | 2.020388 | 0.850964 | -0.928753 | -1.044002 | -0.480491 | -3.23538 | -0.433038 | -3.530191 | -0.952828 | 0.250044 | -2.255266 | -1.592757 | -1.748315 | | 35.2361111 | -1.188613 | 1.103264 |
| 35.1597222 | -0.8382 | 0.411124 | -0.451843 | -1.064258 | -0.90428 | -1.242212 | -1.426639 | -1.791425 | -0.573694 | -0.701975 | -1.267429 | 3.821623 | 1.162477 | -0.351476 | -0.241932 | 0.077486 | -2.126381 | -0.607121 | -2.131969 | -0.502581 | 0.490033 | -1.273718 | -0.516106 | -0.615138 | | 35.25 | -0.829249 | 0.445695 |
| 35.1736111 | -0.998525 | 0.434426 | -0.430424 | -0.710575 | -0.421983 | -0.60005 | -1.413959 | -1.568092 | -0.384297 | -0.168717 | -0.817734 | 4.593031 | 1.153599 | 0.726234 | 1.577989 | 1.812544 | -0.035475 | 0.702458 | 1.316601 | 0.069212 | 0.577411 | 0.419675 | 0.498502 | 1.528684 | | 35.2638889 | 1.571811 | -0.374755 |
| 35.1875 | -1.088208 | 0.522214 | -0.399023 | -0.607481 | 0.045362 | -0.060861 | -1.401892 | -1.259307 | -0.345692 | 0.190403 | -0.846097 | 4.432307 | 0.958664 | 0.733034 | 2.281659 | 2.670626 | 2.16526 | 1.48453 | 2.20511 | 1.335732 | 0.987448 | 1.603413 | 1.283582 | 2.679148 | | 35.2777778 | 3.864355 | -0.390526 |
| 35.2013889 | -0.918606 | 0.979642 | -0.419034 | -0.699213 | 0.34796 | 0.29988 | -1.192087 | -0.873281 | -0.355527 | 0.330919 | -0.973667 | 4.010397 | 0.754313 | 0.618971 | 1.623217 | 2.27744 | 3.284916 | 1.672548 | 2.045112 | 1.892683 | 1.414797 | 1.81609 | 1.480503 | 2.270405 | | 35.2916667 | 3.993179 | 0.448922 |
| 35.2152778 | -0.517764 | 1.680989 | -0.354832 | -0.602895 | 0.578867 | 0.551084 | -1.006937 | -0.639256 | -0.2147 | 0.466619 | -0.745784 | 3.546482 | 0.587728 | 0.427743 | 1.076293 | 2.05178 | 2.746468 | 1.368443 | 1.695631 | 1.72246 | 1.406148 | 1.711289 | 1.322542 | 1.506465 | | 35.3055556 | 2.28089 | 0.93867 |
| 35.2291667 | 0.081677 | 1.988513 | -0.265093 | -0.413202 | 0.741146 | 0.748927 | -0.826289 | -0.502532 | 0.071184 | 0.623379 | -0.21341 | 2.651103 | 0.26025 | 0.475172 | 1.132114 | 2.242759 | 2.077885 | 1.036137 | 1.838459 | 1.125626 | 1.305969 | 1.950551 | 1.322786 | 1.420251 | | 35.3194444 | 0.306861 | 0.838215 |
| 35.2430556 | -0.054136 | 1.433573 | -0.242071 | -0.245769 | 0.782937 | 0.820376 | -0.621819 | -0.307863 | 0.186536 | 0.597771 | 0.086974 | 0.906687 | 0.026303 | 0.193138 | 1.380164 | 2.338451 | 2.531017 | 0.956658 | 0.689599 | 0.814743 | 1.966025 | 2.341446 | 1.566367 | 1.604069 | | 35.3333333 | -0.496693 | 0.141425 |
| 35.2569444 | -0.454226 | 0.806908 | -0.281678 | 0.004366 | 0.873782 | 0.976462 | -0.370496 | 0.020913 | 0.071274 | 0.485593 | -0.025305 | -0.304077 | 0.670408 | 0.424577 | 1.554478 | 2.054581 | 4.373524 | 1.336342 | 0.928764 | 1.863328 | 3.051028 | 2.827521 | 1.794435 | 2.005693 | | 35.3472222 | 0.483296 | 0.090575 |
| 35.2708333 | -0.290915 | 0.824307 | -0.140685 | 0.309281 | 1.159671 | 1.41586 | -0.08589 | 0.412358 | -0.061187 | 0.417823 | -0.136405 | 0.137692 | 1.894271 | 1.277361 | 2.350112 | 2.179803 | 8.266438 | 2.077694 | 1.801634 | 3.092845 | 3.451442 | 3.754219 | 1.919019 | 3.052807 | | 35.3611111 | 2.44288 | 0.873721 |
| 35.2847222 | 0.421657 | 1.318586 | 0.121857 | 0.525249 | 1.652448 | 2.234043 | 0.257726 | 0.684815 | -0.019535 | 0.38662 | 0.209908 | 1.550233 | 2.081224 | 2.174879 | 3.718535 | 2.46163 | 10.635408 | 3.036636 | 3.364183 | 3.22955 | 3.428359 | 4.921729 | 1.974255 | 4.244204 | | 35.375 | 4.476346 | 1.556632 |
| 35.2986111 | 1.112754 | 1.517052 | 0.382647 | 0.66984 | 2.232693 | 3.5204 | 0.446477 | 0.782832 | 0.046262 | 0.331458 | 0.73564 | 2.830588 | 1.584371 | 2.448474 | 4.010045 | 2.365037 | 8.644426 | 3.166666 | 1.735463 | 2.257707 | 3.789642 | 5.431815 | 2.192249 | 4.510306 | | 35.3888889 | 4.749026 | 1.682988 |
| 35.3125 | 1.594113 | 1.250489 | 0.548285 | 0.844902 | 2.37484 | 3.572458 | 0.50091 | 0.919168 | 0.172691 | 0.43827 | 1.116274 | 4.229894 | 1.459685 | 2.530664 | 3.698912 | 1.957596 | 6.33999 | 3.12584 | 0.618732 | 2.559834 | 4.367287 | 5.349512 | 2.619337 | 4.447439 | | 35.4027778 | 3.291838 | 1.559854 |
| 35.3263889 | 2.106115 | 1.364012 | 0.675504 | 0.959911 | 2.489589 | 3.880597 | 0.762889 | 1.264495 | 0.431385 | 0.867524 | 1.634896 | 6.452255 | 2.441459 | 3.156169 | 4.437131 | 2.043734 | 6.642799 | 4.043431 | 2.597412 | 4.254553 | 4.911836 | 6.134576 | 3.150326 | 5.578141 | | 35.4166667 | 1.908557 | 1.416425 |
| 35.3402778 | 2.354266 | 2.077032 | 0.817497 | 1.015269 | 2.874869 | 4.590664 | 1.122935 | 1.656945 | 0.782345 | 1.421525 | 2.413842 | 7.09743 | 3.603672 | 3.563572 | 5.551732 | 3.035759 | 7.251859 | 4.390049 | 3.725037 | 4.276682 | 4.454404 | 7.387158 | 3.546193 | 6.965831 | | 35.4305556 | 1.427175 | 1.409701 |
| 35.3541667 | 2.457009 | 3.015128 | 0.985509 | 1.205771 | 3.295689 | 5.002265 | 1.501608 | 2.017102 | 1.185778 | 1.924345 | 3.292243 | 7.388041 | 4.951115 | 4.087456 | 5.996305 | 3.95044 | 7.338119 | 4.440002 | 4.077459 | 3.775001 | 4.011312 | 8.472512 | 4.409505 | 7.772869 | | 35.4444444 | 1.639864 | 1.458609 |
| 35.3680556 | 2.737656 | 3.725418 | 1.104024 | 1.736292 | 3.634464 | 5.840409 | 1.991656 | 2.469331 | 1.684031 | 2.456833 | 4.312198 | 9.732466 | 5.980758 | 4.528594 | 6.695028 | 5.085165 | 7.45625 | 5.768398 | 3.094354 | 3.349713 | 4.856575 | 10.269241 | 5.79784 | 8.57129 | | 35.4583333 | 2.161948 | 1.469877 |
| 35.3819444 | 3.241647 | 3.430365 | 1.202024 | 2.326059 | 3.59164 | 5.959767 | 2.629944 | 3.00738 | 2.117846 | 3.177187 | 4.997773 | 12.344307 | 6.385776 | 5.090872 | 6.551226 | 5.21704 | 7.135474 | 7.683327 | 2.186142 | 3.260345 | 5.241825 | 12.966725 | 6.780312 | 9.427634 | | 35.4722222 | 2.680661 | 1.392816 |
| 35.3958333 | 3.559516 | 2.549474 | 1.190983 | 2.709202 | 2.80211 | 4.982728 | 3.552287 | 3.660665 | 2.138078 | 3.795481 | 4.945007 | 14.20674 | 6.695067 | 5.141708 | 6.173647 | 4.791504 | 6.892843 | 8.696219 | 2.436974 | 3.82466 | 5.137171 | 14.643319 | 6.734552 | 10.808716 | | 35.4861111 | 3.799088 | 1.269821 |
| 35.4097222 | 3.79931 | 2.19433 | 1.271347 | 2.739051 | 1.939806 | 3.584766 | 4.063724 | 4.189196 | 2.0823 | 4.266769 | 5.070633 | 15.184189 | 7.890323 | 5.24672 | 6.450454 | 5.759051 | 8.24482 | 9.991085 | 3.617425 | 4.235971 | 5.228416 | 17.471086 | 6.971646 | 13.24341 | | 35.5 | 4.826302 | 1.243146 |
| 35.4236111 | 3.978067 | 3.092958 | 1.424506 | 2.504845 | 1.856908 | 3.666445 | 3.950283 | 4.694812 | 2.375362 | 4.885954 | 5.478747 | 16.482427 | 9.419816 | 5.249152 | 7.385414 | 7.944225 | 8.377746 | 10.149358 | 4.630286 | 5.846421 | 5.81308 | 20.449055 | 8.600905 | 17.888301 | | 35.5138889 | 4.929279 | 1.373818 |
| 35.4375 | 3.615366 | 4.789582 | 2.139539 | 2.513786 | 2.226089 | 4.399081 | 3.521214 | 4.551037 | 4.137856 | 6.370372 | 9.462236 | 17.915382 | 10.021642 | 5.746211 | 8.673467 | 9.481619 | 8.270881 | 10.435291 | 7.056479 | 7.898958 | 7.813885 | 20.899405 | 10.124249 | 31.04156 | | 35.5277778 | 6.43394 | 2.540737 |
| 35.4513889 | 3.513181 | 6.296776 | 3.356896 | 2.960562 | 2.80049 | 4.220298 | 3.559306 | 3.958136 | 7.083708 | 9.301152 | 20.139047 | 19.728855 | 10.420497 | 7.283482 | 8.138956 | 8.345877 | 9.074154 | 8.992496 | 10.034148 | 9.206965 | 8.725411 | 20.556579 | 8.780594 | 51.742303 | | 35.5416667 | 11.443132 | 4.508514 |
| 35.4652778 | 4.51532 | 6.379235 | 3.585397 | 3.000017 | 2.062289 | 3.900609 | 3.689524 | 3.392362 | 7.123901 | 8.903268 | 32.444184 | 14.359596 | 6.923728 | 8.260196 | 7.231906 | 8.282617 | 8.472429 | 6.237389 | 8.199859 | 6.054287 | 6.917695 | 17.537528 | 7.03574 | 48.351843 | | 35.5555556 | 13.046573 | 3.932885 |
| 35.4791667 | 3.697872 | 3.905529 | 2.723702 | 2.170931 | 1.342826 | 2.204791 | 4.140256 | 3.283435 | 3.287535 | 4.967348 | 19.691606 | 0.720358 | 1.035986 | 4.587205 | 4.552696 | 4.719775 | 2.28482 | 1.243484 | 4.369422 | 1.130517 | 2.313813 | 5.963236 | 4.465148 | 18.195096 | | 35.5694444 | 7.646757 | 0.836547 |
| 35.4930556 | 2.181187 | 0.374383 | 2.129078 | 1.071126 | -0.092868 | -0.421297 | 2.284806 | 0.304916 | 1.153055 | 1.282153 | 1.510293 | -3.969004 | -0.605727 | 3.401756 | 4.290771 | 4.488045 | -2.360723 | 0.990986 | 0.023826 | 1.451638 | 1.983812 | -1.347572 | 5.461964 | 2.231072 | | 35.5833333 | 5.566485 | -0.03677 |
| 35.5069444 | 0.87663 | 0.143954 | 2.310454 | 1.426517 | -0.712774 | -1.173984 | 0.368603 | -1.37055 | 0.289564 | -0.225544 | -5.276167 | -0.058502 | 3.864621 | 3.725093 | 4.58711 | 5.497653 | -0.870613 | 3.089666 | -0.020365 | 3.890816 | 4.248916 | 3.737362 | 8.708124 | 10.562135 | | 35.5972222 | 5.363144 | -0.041303 |
| 35.5208333 | 0.431819 | 1.641413 | 1.984744 | 2.378817 | -0.603192 | -0.200478 | 0.087032 | -0.49618 | 0.444153 | -0.402949 | 0.68972 | 5.584847 | 7.420824 | 3.687001 | 3.361536 | 5.255394 | 1.258914 | 3.302061 | 3.825379 | 6.229743 | 4.669624 | 13.551578 | 2.697682 | 21.700047 | | 35.6111111 | 3.437134 | -0.009535 |
| 35.5347222 | 1.17455 | 1.867754 | 2.031068 | 3.07733 | 0.698233 | 1.24042 | -0.09167 | 1.048232 | 0.467015 | -0.41293 | 7.16746 | 9.364393 | 7.650772 | 2.088677 | 1.949522 | 5.982944 | 2.645864 | 2.733822 | 4.273245 | 5.901421 | 6.555291 | 15.056271 | 1.363861 | 25.598997 | | 35.625 | -0.026798 | -0.499682 |
| 35.5486111 | 2.732499 | 1.537812 | 2.419739 | 2.309129 | 1.322719 | 1.711172 | 0.087203 | 1.337341 | 0.727322 | 0.390124 | 10.710777 | 6.75123 | 5.855574 | 2.651064 | 2.126079 | 5.22974 | 2.770782 | 2.33029 | 0.254114 | 0.886666 | 3.758019 | 9.556868 | 4.235837 | 17.538338 | | 35.6388889 | -2.445635 | -1.133227 |

| | | | | | | | | | | | | | | | | | | | | | | | | | | | |
|---|---|---|---|---|---|---|---|---|---|---|---|---|---|---|---|---|---|---|---|---|---|---|---|---|---|---|---|
| 35.5625 | 5.542659 | 3.100121 | 1.941671 | 1.377483 | 0.462042 | 2.308873 | 1.205598 | 1.437157 | 1.222443 | 0.936921 | 10.591386 | 2.339269 | 2.012499 | 4.459873 | 3.709968 | 2.188425 | 2.976785 | 3.060011 | 0.684733 | -2.724233 | -4.600486 | 6.736662 | 4.154977 | 7.961275 | 35.6527778 | -2.0801 | [illegible] |
| 35.5763889 | 6.236462 | 5.115557 | 0.622034 | 0.294671 | 0.462149 | 1.172335 | 1.904 | 1.233106 | 0.77483 | -0.831799 | 4.895034 | 1.191648 | 1.854472 | 2.520436 | 3.128337 | 0.4337 | 0.916744 | 2.108407 | 1.373181 | -2.814968 | -8.077522 | 6.533217 | 0.258011 | 3.33018 | 35.6666667 | 1.316809 | [illegible] |
| 35.5902778 | 4.449312 | 3.929083 | -0.265938 | 0.19055 | 0.516325 | 1.048027 | 1.063765 | -0.542459 | 0.287447 | -1.725898 | 0.20204 | 0.261479 | 0.975158 | 1.53244 | 2.107074 | 0.782814 | -0.515235 | 0.813328 | 1.059842 | -2.367215 | -2.134653 | 4.82768 | 0.712226 | 1.570769 | 35.6805556 | 3.840005 | [illegible] |
| 35.6041667 | 0.37198 | 0.284287 | -1.575018 | -0.456662 | 0.488713 | 0.556742 | 0.65443 | -0.60175 | 1.174594 | 0.447653 | -2.317109 | 0.356066 | 0.119944 | 0.321133 | 1.197273 | 2.300203 | -0.373767 | 1.129193 | 1.034646 | 1.656824 | 0.855133 | 0.997199 | 0.111796 | 1.177474 | 35.6944444 | 1.836974 | [illegible] |
| 35.6180556 | -2.105212 | -2.778123 | -0.882635 | -1.25204 | -0.216058 | -1.798589 | 0.29424 | -1.518476 | 1.874927 | 3.591866 | -4.749508 | 1.021049 | 1.486653 | -0.630333 | -0.595382 | 0.135252 | 0.166982 | 0.022984 | 1.067245 | 7.091202 | 5.231546 | -2.923245 | -3.304383 | -0.651293 | 35.7083333 | -2.870607 | [illegible] |
| 35.6319444 | -2.82199 | -2.107382 | -1.237257 | -1.494363 | -0.066581 | -0.868936 | 0.078636 | -1.237387 | 0.599175 | 2.490312 | -4.031069 | 0.384311 | 1.217378 | -2.2621 | -2.280465 | -0.346822 | 1.252223 | -0.71313 | -0.274351 | 5.618227 | 4.918898 | -5.766618 | -4.709066 | -4.146144 | 35.7222222 | -7.710964 | [illegible] |
| 35.6458333 | -0.870761 | 0.352532 | -1.716918 | -1.491461 | 0.252685 | 1.2322 | -0.527188 | -2.022263 | -0.398921 | 1.011339 | -3.135784 | -1.141468 | -1.849069 | -2.171958 | -2.459902 | -1.831771 | 1.143647 | 0.919818 | -1.972831 | -0.510864 | -2.494125 | -4.496197 | -2.843117 | -5.466319 | 35.7361111 | -4.21768 | [illegible] |
| 35.6597222 | -0.560559 | -0.336124 | -0.776718 | -0.736236 | 0.387311 | 2.459123 | -1.712741 | -2.243969 | -0.741845 | 1.042233 | -3.997553 | -2.703936 | -4.02618 | -0.389949 | -1.136198 | -2.454313 | 0.192501 | 0.96093 | -1.210964 | -5.321344 | -8.71236 | -2.342656 | -4.242846 | -8.502041 | 35.75 | 1.854515 | [illegible] |
| 35.6736111 | -0.371077 | -0.161455 | 0.584575 | 0.86436 | 0.160501 | 1.772591 | -1.355537 | -1.872014 | -0.782697 | 1.722061 | -5.244684 | -4.798266 | -6.781772 | 1.763596 | 0.973367 | -0.882995 | -1.215997 | -1.180139 | 0.033348 | -4.91812 | -4.619748 | -1.38183 | -3.744492 | -9.541687 | 35.7638889 | 6.454473 | [illegible] |
| 35.6875 | -0.783061 | 0.324304 | 0.613106 | 1.540445 | 0.199957 | 1.790454 | 0.242255 | 0.002022 | -0.708435 | 0.981028 | -1.424203 | -7.813191 | -7.709498 | 1.345575 | 0.746715 | 0.20364 | -1.45631 | -1.525614 | -0.116191 | -1.445198 | 3.493602 | -3.789547 | -4.634623 | -7.233188 | 35.7777778 | 6.892743 | [illegible] |
| 35.7013889 | -1.717831 | -0.021016 | 0.31321 | 1.408817 | 0.290479 | 1.342506 | 1.800755 | 3.062095 | -0.466344 | 0.300518 | 3.568504 | -8.454753 | -9.32789 | -0.51323 | -0.965158 | -0.552579 | -1.285613 | -2.040067 | 0.441178 | 2.525151 | -0.440846 | -8.653023 | -6.195202 | -6.907455 | 35.7916667 | 6.308315 | [illegible] |
| 35.7152778 | -2.724549 | -0.792693 | 0.166559 | 1.251346 | 0.24136 | 1.074261 | 2.275586 | 3.969517 | 0.194238 | 0.015136 | 4.124058 | -5.096465 | -5.922009 | -2.731152 | -2.242515 | 0.283742 | -1.008784 | -2.428871 | 1.580166 | 4.735264 | -0.159847 | -2.347909 | 1.070564 | -1.2322 | 35.8055556 | 4.890927 | [illegible] |
| 35.7291667 | -3.675349 | -1.674746 | 0.410683 | 0.86307 | 0.128483 | 0.104563 | 1.853022 | 3.096815 | 0.826525 | -0.494527 | 1.816135 | -1.778699 | -0.375668 | -4.622185 | -3.213762 | 0.739712 | -0.060993 | -2.444735 | 0.702841 | 3.773267 | -1.706756 | 1.225505 | 3.952798 | 3.542929 | 35.8194444 | 1.26284 | [illegible] |
| 35.7430556 | -1.34176 | -1.366471 | 1.176849 | 0.38448 | 0.05725 | -0.570295 | 0.76962 | 1.176729 | 0.928104 | -1.510924 | 2.023427 | -0.743461 | 2.755901 | -5.844643 | -2.596314 | 1.725177 | 1.810816 | 1.121174 | -0.700601 | -2.632351 | -4.3899 | -0.598212 | 0.042272 | 0.680865 | 35.8333333 | -2.890283 | [illegible] |
| 35.7569444 | 0.854768 | -0.24345 | 1.500596 | -0.059394 | -0.358155 | -0.132716 | 0.611809 | 0.860592 | 0.629855 | -1.553168 | 3.040833 | -2.817806 | -0.419201 | -1.719648 | -1.392822 | -0.102171 | 1.775715 | 3.440142 | -1.261617 | -7.785835 | -4.14556 | 0.465636 | -1.735721 | -4.544125 | 35.8472222 | -1.616636 | [illegible] |
| 35.7708333 | 1.62639 | 0.078213 | 1.517611 | -0.607216 | -0.104469 | 0.460671 | 1.025466 | 0.511669 | 0.66346 | 0.858294 | 3.867275 | -2.76253 | -5.87115 | 3.89556 | 1.83766 | 1.222835 | -0.45153 | 3.19517 | -2.044364 | -6.50853 | 2.496758 | -4.448996 | 0.243599 | -5.896604 | 35.8611111 | -0.093768 | [illegible] |
| 35.7847222 | 1.953132 | -0.384278 | 0.520201 | -1.417591 | -0.021081 | 0.306315 | 0.897356 | 0.579354 | -0.057281 | 1.502825 | 3.965786 | -3.531094 | -8.887608 | 4.632668 | 3.917531 | 2.145096 | -1.536299 | 2.331192 | -2.777171 | -1.595115 | 4.076046 | -16.93416 | 2.152783 | -5.693349 | 35.875 | -1.522275 | [illegible] |
| 35.7986111 | 1.414048 | -1.08885 | -0.924772 | -2.048218 | -0.090625 | -0.345849 | -0.28008 | -0.12795 | -0.76293 | 1.579276 | 2.770699 | -2.485996 | -6.636325 | 1.994903 | 1.095053 | -0.207726 | -1.670823 | 0.321271 | -0.226836 | 5.726226 | 3.597848 | -9.946263 | 2.919827 | -3.116947 | 35.8888889 | -2.60122 | [illegible] |
| 35.8125 | 0.661332 | -1.49223 | -1.31172 | -1.911402 | -0.527091 | -1.996388 | -1.461546 | -0.941108 | -1.473561 | -0.185308 | -1.0557 | -3.113818 | -2.402135 | -1.1511 | -2.883985 | -2.162173 | -2.014768 | -1.623899 | 3.611076 | 7.47731 | 3.252865 | 4.657316 | 4.576585 | -0.761576 | 35.9027778 | -2.214183 | [illegible] |
| 35.8263889 | 0.89986 | -0.72265 | -1.534022 | -1.836402 | -0.548166 | -2.31354 | -1.708293 | -0.962669 | -2.3237 | -2.127979 | -4.049161 | -3.298336 | 2.613894 | -3.330644 | -4.359383 | -2.003646 | -2.275532 | -2.685396 | 3.487441 | 2.567329 | -1.75662 | 15.194518 | 1.691992 | -2.638123 | 35.9166667 | -1.965803 | [illegible] |
| 35.8402778 | 2.3535 | 0.676037 | -0.545754 | -0.484518 | -0.544976 | -1.36228 | -2.301564 | -1.281112 | -2.555988 | -2.998706 | -2.538103 | -3.69302 | 4.051688 | -2.897033 | -2.344055 | -2.486709 | -1.919293 | -5.127646 | 0.759744 | -3.617967 | -5.48471 | -1.687736 | -4.399073 | -9.529819 | 35.9305556 | -4.371786 | [illegible] |
| 35.8541667 | 4.358457 | 1.104229 | 0.324078 | 1.006022 | -0.458403 | 0.018726 | -4.544649 | -2.530791 | -1.224392 | -0.925591 | -2.109628 | -3.47523 | 2.145624 | -3.856874 | 0.078477 | -2.596257 | 0.190526 | -0.287305 | -1.529384 | -7.168879 | -2.702507 | -13.316082 | -8.486219 | -9.166903 | 35.9444444 | -4.752978 | [illegible] |
| 35.8680556 | 4.737163 | 0.433143 | -0.003088 | 0.413744 | -0.988157 | -0.469531 | -5.489143 | -4.315091 | 0.157692 | 1.402888 | -2.326995 | -0.222881 | -1.469016 | 0.838912 | 1.139551 | -2.204537 | 1.862357 | 3.127222 | -3.777645 | -10.204009 | -1.645188 | -12.043407 | -5.287074 | -6.433514 | 35.9583333 | -3.684453 | [illegible] |
| 35.8819444 | 2.422381 | -0.788018 | -0.794703 | -1.587622 | -0.659875 | -2.192374 | -6.113611 | -5.773352 | -0.732876 | -0.36542 | -4.486868 | 1.018906 | -3.929924 | 6.582869 | -0.331017 | -3.051091 | -0.835064 | 2.323409 | -2.853637 | -4.99949 | -0.427934 | -6.022036 | -3.206786 | -3.676467 | 35.9722222 | -3.730793 | [illegible] |
| 35.8958333 | -1.200526 | -2.261299 | -1.571006 | -2.516908 | 0.047583 | -2.199761 | -5.022363 | -5.749314 | -1.573905 | -3.103021 | -7.875727 | -2.736593 | -4.153643 | 8.275729 | -2.915844 | -6.039429 | -3.086172 | -1.905282 | -2.321863 | -2.199797 | -3.713474 | 0.150771 | -2.506141 | -11.460544 | 35.9861111 | -5.306993 | [illegible] |
| 35.9097222 | -4.600759 | -3.723021 | -2.54983 | -2.300609 | -0.149051 | -2.621236 | -4.802785 | -4.948349 | -3.80993 | -5.048386 | -11.370145 | -6.857804 | -4.870085 | 1.976762 | -2.393999 | -5.650162 | -5.65104 | -6.62333 | -2.57503 | -5.031002 | -5.431554 | 2.637978 | -3.374234 | -16.249445 | 36 | -6.139585 | [illegible] |
| 35.9236111 | -6.182762 | -3.463214 | -2.189165 | -2.741924 | -0.687533 | -0.271205 | -4.757725 | -3.219776 | -5.305766 | -6.802881 | -13.571831 | -6.182965 | -4.541736 | -1.836188 | -0.156853 | -1.128383 | -8.582599 | -9.553451 | -4.837807 | -11.063883 | -4.977721 | 0.407525 | -6.524725 | -15.235992 | 36.0138889 | -5.079582 | [illegible] |
| 35.9375 | -7.136647 | -4.021723 | -1.560971 | -3.234741 | -2.015696 | -0.638904 | -6.442465 | -2.007389 | -5.355568 | -7.783263 | -21.697663 | -6.245252 | -3.686154 | -6.912289 | -3.059611 | -0.774017 | -10.486209 | -10.984617 | -14.624774 | -15.879927 | -4.295425 | -8.480218 | -12.472078 | -29.447804 | 36.0277778 | -4.688929 | [illegible] |
| 35.9513889 | -5.555003 | -7.041056 | -0.974144 | -2.715745 | -3.418229 | -2.014392 | -8.053518 | -0.59067 | -6.52683 | -7.766218 | -31.514222 | -9.607309 | -5.397839 | -11.631255 | -10.033917 | -6.10454 | -11.958374 | -8.201194 | -17.873227 | -12.372385 | -7.337173 | -12.830127 | -17.915518 | -53.938041 | 36.0416667 | -8.463999 | [illegible] |
| 35.9652778 | -3.817371 | -6.912667 | -1.368202 | -2.2354 | -3.68857 | -4.847107 | -7.481398 | -0.813913 | -5.998517 | -6.213992 | -28.475984 | -12.812377 | -6.521769 | -8.847929 | -11.368868 | -8.391691 | -14.543766 | -9.72121 | -17.487705 | -9.442729 | -10.00062 | -16.063054 | -9.810875 | -52.616789 | 36.0555556 | -9.758442 | [illegible] |
| 35.9791667 | -3.663387 | -4.931329 | -1.187531 | -1.693484 | -2.916606 | -5.336921 | -5.368397 | -2.266876 | -3.678783 | -4.314051 | -15.381362 | -11.540875 | -5.970586 | -7.799917 | -9.248075 | -8.213007 | -15.436628 | -6.538275 | -12.024356 | -8.064859 | -5.749369 | -15.322418 | -5.133359 | -31.37893 | 36.0694444 | -10.326193 | [illegible] |
| 35.9930556 | -2.805564 | -2.47264 | -1.295567 | -1.326275 | -1.705576 | -4.212215 | -3.309091 | -2.499213 | -2.38574 | -3.100039 | -6.083086 | -8.340144 | -2.437584 | -5.701447 | -6.55998 | -5.792868 | -12.359312 | -6.240985 | -8.512866 | -7.380248 | -2.889996 | -9.977183 | -3.135771 | -13.816302 | 36.0833333 | -10.778841 | [illegible] |
| 36.0069444 | -1.752571 | -0.989137 | -1.387114 | -1.263895 | -1.227435 | -2.679208 | -2.74146 | -2.247374 | -2.12313 | -2.368255 | -3.157377 | -8.908842 | -1.775029 | -5.93642 | -4.083044 | -3.053021 | -7.119219 | -3.06904 | -9.580917 | -5.821059 | -1.180874 | -8.086497 | -4.310756 | -7.652203 | 36.0972222 | -8.266878 | [illegible] |
| 36.0208333 | -1.238882 | -0.427645 | -1.076518 | -1.089165 | -1.093742 | -2.032718 | -2.168917 | -1.606312 | -1.9166 | -1.893861 | -2.421998 | -10.524257 | -1.177089 | -4.843817 | -0.638696 | -0.299048 | -3.836547 | -0.908956 | -6.188703 | -2.075533 | -0.276099 | -9.740533 | -5.989864 | -5.152652 | 36.1111111 | -4.842443 | [illegible] |
| 36.0347222 | -0.993909 | -0.319166 | -0.530038 | -0.991776 | -1.017797 | -1.932496 | -1.618516 | -0.96693 | -1.47815 | -1.436736 | -2.187368 | -12.049356 | -2.262199 | -4.974945 | -1.633972 | -2.900197 | -2.460503 | -1.283459 | -2.716571 | -0.32012 | -0.200402 | -5.962794 | -4.80459 | -4.25374 | 36.125 | -2.497679 | [illegible] |
| 36.0486111 | -0.955038 | -0.388821 | -0.196477 | -0.471377 | -0.988245 | -1.88472 | -1.331909 | -0.59014 | -0.889412 | -1.032537 | -1.71478 | -10.573221 | -2.057467 | -5.812501 | -5.25216 | -6.170072 | -3.412044 | -2.266757 | -2.170968 | -0.673445 | -1.245279 | -5.149018 | -4.130912 | -5.148585 | 36.1388889 | -0.764224 | [illegible] |
| 36.0625 | -0.99327 | -0.554585 | -0.288926 | -0.403852 | -0.991524 | -1.950601 | -1.299428 | -0.623168 | -0.459552 | -0.670163 | -1.262053 | -6.53506 | -0.732271 | -6.352517 | -6.043334 | -5.260333 | -4.494783 | -2.493879 | -2.724272 | -1.59715 | -2.505753 | -5.651088 | -3.95963 | -5.727728 | 36.1527778 | 0.890206 | [illegible] |
| 36.0763889 | -1.068152 | -0.738656 | -0.679944 | -0.865322 | -1.061308 | -2.043706 | -1.255963 | -0.745237 | -0.313379 | -0.449016 | -1.161966 | -4.590383 | -1.048757 | -4.752815 | -4.77115 | -2.665445 | -5.898033 | -3.056942 | -4.284847 | -2.671827 | -3.513493 | -5.935394 | -2.42428 | -5.110323 | 36.1666667 | 1.986201 | [illegible] |
| 36.0902778 | -1.200311 | -0.995097 | -0.795739 | -1.176349 | -1.168717 | -2.121491 | -1.075351 | -0.683649 | -0.322951 | -0.389074 | -1.318346 | -4.544513 | -2.478121 | -4.322639 | -5.419119 | -2.91348 | -7.504271 | -3.584753 | -5.876096 | -2.773139 | -3.343798 | -5.384516 | -0.609231 | -4.986663 | 36.1805556 | 2.313151 | [illegible] |
| 36.1041667 | -1.353767 | -1.247331 | -0.593307 | -0.96371 | -1.272359 | -2.200639 | -0.888717 | -0.562791 | -0.269282 | -0.378699 | -1.320113 | -5.53632 | -4.540268 | -4.216037 | -6.6207 | -4.442604 | -7.788179 | -2.795552 | -4.475369 | -2.134871 | -2.916341 | -4.542166 | -1.071051 | -5.221919 | 36.1944444 | 1.948438 | [illegible] |
| 36.1180556 | -1.502373 | -1.474007 | -0.460048 | -0.64791 | -1.446407 | -2.439676 | -0.941956 | -0.626317 | -0.178611 | -0.329115 | -1.23536 | -3.395963 | -3.217803 | -3.011519 | -5.490303 | -3.317408 | -4.835878 | 0.128905 | -1.159925 | -0.826264 | -2.54952 | -3.97314 | -2.153008 | -4.874447 | 36.2083333 | 0.837332 | [illegible] |
| 36.1319444 | -1.724498 | -1.785346 | -0.654322 | -0.602281 | -1.875138 | -3.034798 | -1.326199 | -0.744266 | -0.220726 | -0.463046 | -1.562793 | -0.654314 | -1.323726 | -0.856081 | -2.910677 | -1.646976 | -1.246567 | 2.486003 | 2.795231 | 0.591596 | -1.390252 | -3.293805 | -1.889451 | -3.468952 | 36.2222222 | -0.486896 | [illegible] |
| 36.1458333 | -1.945512 | -2.165835 | -0.643392 | -0.751489 | -2.42191 | -4.080336 | -1.796077 | -0.754919 | -0.371288 | -0.625002 | -1.77925 | 1.511268 | -0.677626 | 1.029601 | -0.835466 | -0.66111 | 0.95851 | 2.351014 | 4.410433 | 1.087977 | -0.628819 | -1.620595 | -0.231239 | -1.495692 | 36.2361111 | -0.68733 | [illegible] |
| 36.1597222 | -2.024636 | -2.107848 | -0.662635 | -0.883707 | -2.479403 | -4.534107 | -1.850838 | -0.588214 | -0.543785 | -0.634194 | -1.241642 | 2.605409 | -0.975324 | 1.253464 | -0.245153 | 0.197722 | 1.843919 | 1.345369 | 2.565754 | 0.869967 | -0.258489 | 0.453132 | 0.96824 | 0.260793 | 36.25 | -0.092505 | [illegible] |
| 36.1736111 | -2.077444 | -1.710044 | -0.695297 | -0.80922 | -1.698628 | -3.778975 | -1.463995 | -0.454195 | -0.654815 | -0.467945 | -0.120437 | 3.403671 | -0.837139 | 1.202916 | 0.46899 | 1.178428 | 2.312422 | 0.593182 | 1.461731 | 0.786244 | 0.083378 | 2.318574 | 1.243962 | 1.581799 | 36.2638889 | 0.124283 | [illegible] |
| 36.1875 | -1.922711 | -1.2454 | -0.595163 | -0.393435 | -1.120887 | -2.461343 | -0.980681 | -0.480712 | -0.716011 | -0.312389 | 0.200764 | 4.338461 | -0.022122 | 1.664709 | 1.665454 | 2.084172 | 3.538429 | 1.627317 | 1.320261 | 1.133244 | 0.769061 | 3.638515 | 1.060861 | 2.43809 | 36.2777778 | 0.659587 | [illegible] |
| 36.2013889 | -1.488396 | -0.677784 | -0.457425 | -0.012771 | -0.898122 | -1.128177 | -0.630722 | -0.500959 | -0.860636 | -0.371889 | -0.464722 | 5.043144 | 0.789446 | 2.095325 | 2.528248 | 2.733422 | 5.275736 | 3.698332 | 3.078047 | 1.967179 | 1.768795 | 4.141123 | 1.329058 | 2.957575 | 36.2916667 | 1.529674 | [illegible] |
| 36.2152778 | -1.187754 | -0.129864 | -0.424612 | -0.058999 | -1.020667 | -0.37857 | -0.48619 | -0.43418 | -0.998834 | -0.500209 | -1.202816 | 5.670388 | 1.149661 | 2.537569 | 2.86063 | 2.989522 | 7.942388 | 5.26572 | 5.20705 | 2.927098 | 2.286856 | 4.182254 | 2.021585 | 3.49076 | 36.3055556 | 2.114003 | [illegible] |
| 36.2291667 | -0.821488 | 0.376803 | -0.345137 | -0.300003 | -1.036082 | -0.062162 | -0.34537 | -0.241842 | -0.945178 | -0.340524 | -1.490189 | 6.009521 | 1.302717 | 3.051773 | 3.119533 | 3.089047 | 10.233805 | 6.404504 | 6.398562 | 3.280137 | 2.402528 | 3.670756 | 2.067726 | 3.924726 | 36.3194444 | 2.538129 | [illegible] |
| 36.2430556 | -0.546609 | 0.733549 | -0.276868 | -0.416226 | -0.73989 | 0.201803 | 0.075424 | -0.07778 | -0.693456 | -0.027357 | -1.156689 | 6.021843 | 1.286109 | 2.996114 | 3.162872 | 2.96084 | 9.352936 | 6.883202 | 5.555803 | 3.422313 | 2.690476 | 2.829965 | 1.524554 | 3.945123 | 36.3333333 | 3.500055 | [illegible] |
| 36.2569444 | -0.391604 | 0.882405 | -0.221925 | -0.209329 | 0.049681 | 0.745317 | 0.714659 | 0.003635 | -0.431512 | 0.210375 | -0.487364 | 5.118895 | 1.039594 | 2.231762 | 2.554298 | 2.459875 | 8.015675 | 6.679038 | 3.534761 | 3.995927 | 2.612409 | 2.16718 | 1.161704 | 3.57821 | 36.3472222 | 4.343904 | [illegible] |
| 36.2708333 | -0.261635 | 0.891631 | -0.162622 | 0.21044 | 0.983872 | 1.549981 | 1.224257 | 0.046057 | -0.261037 | 0.304735 | 0.137856 | 4.438347 | 1.053407 | 1.316279 | 1.613673 | 1.608382 | 7.146189 | 5.843255 | 6.391592 | 4.839715 | 2.32544 | 2.664074 | 1.635542 | 3.41668 | 36.3611111 | 4.281429 | [illegible] |
| 36.2847222 | -0.107093 | 0.867026 | -0.090302 | 0.578356 | 1.9591 | 2.50149 | 1.410683 | 0.18054 | -0.09024 | 0.394654 | 0.575004 | 4.639751 | 1.794794 | 1.302591 | 1.846059 | 1.560952 | 7.558587 | 6.05256 | 7.703748 | 5.208281 | 2.247228 | 3.744924 | 2.172417 | 3.841921 | 36.375 | 3.443537 | [illegible] |
| 36.2986111 | 0.09027 | 0.875578 | -0.002247 | 0.676946 | 2.709771 | 3.217825 | 1.503564 | 0.419403 | 0.100529 | 0.577836 | 1.096162 | 5.241205 | 2.803262 | 2.599164 | 3.659046 | 2.859406 | 8.847487 | 7.251568 | 5.15811 | 3.972891 | 2.668553 | 4.42748 | 2.116706 | 4.897404 | 36.3888889 | 2.385643 | [illegible] |
| 36.3125 | 0.260799 | 0.922791 | 0.116432 | 0.712817 | 3.301852 | 3.534052 | 1.759631 | 0.690675 | 0.286512 | 0.794317 | 1.740013 | 5.906687 | 3.405405 | 4.392121 | 5.486647 | 4.57409 | 9.488244 | 8.510587 | 4.990043 | 4.108481 | 3.242308 | 4.889256 | 2.131769 | 6.260274 | 36.4027778 | 1.763148 | [illegible] |
| 36.3263889 | 0.381403 | 0.981783 | 0.212291 | 0.828866 | 3.552869 | 3.796557 | 2.240832 | 0.871429 | 0.393015 | 1.01883 | 2.277936 | 5.672611 | 3.744208 | 3.723677 | 5.538449 | 4.720593 | 9.470646 | 10.841977 | 5.967098 | 4.924777 | 3.82211 | 5.43231 | 2.659731 | 6.800788 | 36.4166667 | 1.770558 | [illegible] |
| 36.3402778 | 0.54044 | 1.021088 | 0.24734 | 1.104932 | 3.673085 | 4.140874 | 2.625191 | 0.931601 | 0.479028 | 1.348594 | 2.506791 | 4.258733 | 3.558553 | 2.279201 | 4.350466 | 3.565978 | 9.735954 | 11.987473 | 6.119016 | 5.406214 | 3.695133 | 6.799965 | 3.589538 | 6.914601 | 36.4305556 | 2.067103 | [illegible] |
| 36.3541667 | 0.819149 | 1.148852 | 0.238388 | 1.35676 | 3.568613 | 4.77089 | 2.896686 | 1.080047 | 0.748965 | 1.955141 | 2.452806 | 4.772737 | 3.846861 | 2.177964 | 3.621001 | 3.193094 | 10.049067 | 10.771488 | 7.483734 | 5.431145 | 3.177684 | 8.317362 | 4.023009 | 6.730658 | 36.4444444 | 2.255932 | [illegible] |
| 36.3680556 | 1.166188 | 1.477038 | 0.267256 | 1.278456 | 3.302278 | 5.110594 | 2.997103 | 1.507121 | 1.342176 | 2.926216 | 2.97044 | 6.688407 | 4.771511 | 2.659243 | 4.598311 | 4.383294 | 10.274444 | 10.423769 | 5.728024 | 5.377415 | 3.038485 | 9.688862 | 3.928133 | 7.132966 | 36.4583333 | 1.881638 | [illegible] |
| 36.3819444 | 1.507478 | 1.917961 | 0.437062 | 1.170419 | 3.18353 | 5.161295 | 3.317508 | 2.212057 | 2.014687 | 4.048196 | 4.084389 | 9.839113 | 6.373394 | 3.829994 | 6.407662 | 6.289698 | 9.471294 | 10.653262 | 3.523145 | 4.360623 | 3.40753 | 11.0003 | 3.805547 | 8.633477 | 36.4722222 | 1.406071 | [illegible] |
| 36.3958333 | 2.108422 | 2.550403 | 0.757805 | 1.487834 | 3.187445 | 5.990636 | 4.389088 | 3.302044 | 2.57596 | 5.097747 | 5.654111 | 13.193279 | 8.651303 | 5.479369 | 8.219313 | 9.081458 | 9.022898 | 11.487467 | 3.19075 | 4.409437 | 4.091865 | 12.289269 | 4.458366 | 11.11787 | 36.4861111 | 1.804598 | [illegible] |
| 36.4097222 | 2.809297 | 2.995586 | 1.11957 | 2.137224 | 3.329528 | 7.384686 | 5.68892 | 4.316479 | 3.115167 | 6.093336 | 7.063116 | 15.785085 | 9.862973 | 5.139811 | 7.954933 | 9.206791 | 8.636535 | 11.978936 | 5.958422 | 5.167592 | 4.858923 | 17.022655 | 6.388394 | 13.883508 | 36.5 | 2.981176 | [illegible] |
| 36.4236111 | 3.051098 | 2.794073 | 1.304338 | 2.660252 | 4.293457 | 8.481421 | 5.929284 | 5.044913 | 4.01398 | 6.805854 | 9.103303 | 16.212417 | 9.983535 | 4.110026 | 6.004852 | 6.948501 | 7.924182 | 11.576918 | 9.293417 | 8.465637 | 5.867258 | 19.684787 | 8.775196 | 16.701984 | 36.5138889 | 4.788264 | [illegible] |
| 36.4375 | 2.714591 | 3.48058 | 1.574605 | 2.933915 | 4.329258 | 6.484362 | 6.14582 | 5.340573 | 5.467516 | 7.968342 | 16.710631 | 16.119184 | 9.399126 | 3.09728 | 4.024618 | 4.328717 | 7.621791 | 11.195862 | 10.55961 | 10.941678 | 7.743532 | 21.830313 | 10.457234 | 24.532966 | 36.5277778 | 8.807119 | [illegible] |
| 36.4513889 | 3.113087 | 5.465171 | 1.857926 | 3.347282 | 4.483197 | 5.444776 | 7.279514 | 6.430534 | 8.15358 | 9.69561 | 32.23341 | 16.390887 | 8.40218 | 4.10311 | 5.489068 | 4.206862 | 7.454581 | 10.702885 | 10.892198 | 11.567224 | 9.720378 | 24.945275 | 10.159477 | 38.989152 | 36.5416667 | 16.01241 | [illegible] |
| 36.4652778 | 5.500555 | 5.85991 | 2.737874 | 2.328368 | 3.280956 | 4.241446 | 6.937293 | 6.92153 | 8.248863 | 11.128904 | 37.204155 | 15.749342 | 8.849429 | 5.47771 | 6.03371 | 5.35994 | 8.529903 | 10.55845 | 8.373037 | 8.485508 | 8.421243 | 21.637188 | 7.650473 | 39.238233 | 36.5555556 | 14.794838 | [illegible] |
| 36.4791667 | 7.557497 | 3.608642 | 1.909795 | 1.672859 | 1.534578 | 1.879729 | 2.515831 | 3.531508 | 4.391623 | 5.936677 | 15.224777 | 1.100957 | 2.671649 | 3.940715 | 4.929473 | 5.452383 | 3.455726 | 4.425969 | 2.12259 | 4.539008 | 3.742118 | 5.155169 | 3.600651 | 16.885617 | 36.5694444 | 5.814441 | [illegible] |
| 36.4930556 | 5.455391 | 1.550877 | 1.621762 | 1.516768 | 0.784922 | 1.412569 | -2.557827 | 0.008359 | 1.667457 | 1.803154 | -4.67512 | -7.410129 | -1.149735 | 4.733706 | 5.281249 | 7.508319 | 0.476826 | 1.12796 | 1.460706 | 1.423421 | 1.815391 | -2.453424 | 3.036236 | 4.903432 | 36.5833333 | 2.587168 | [illegible] |
| 36.5069444 | 5.079324 | 1.582343 | 1.425585 | 1.675405 | 0.474881 | 2.973416 | -1.077702 | 1.355962 | 0.655233 | -0.000999 | -3.087914 | -2.789439 | -0.280805 | 4.575986 | 8.400434 | 10.932374 | 1.15333 | 3.950553 | 2.983212 | 2.426319 | 3.965639 | 5.24225 | 8.178681 | 15.760289 | 36.5972222 | 5.409573 | [illegible] |
| 36.5208333 | 5.893213 | 1.569383 | 1.75221 | 1.596133 | 1.630649 | 3.567133 | 2.641491 | 2.792263 | 0.339862 | -0.35986 | 3.215475 | 3.275364 | 5.656063 | 5.352627 | 8.022759 | 12.570141 | 2.602966 | 3.16308 | 4.096547 | 4.965374 | 5.399504 | 12.25477 | 10.480681 | 28.304752 | 36.6111111 | 8.579923 | [illegible] |
| 36.5347222 | 6.472126 | 1.660158 | 2.004738 | 1.836297 | 1.845745 | 2.781126 | 3.608616 | 4.082972 | 0.227294 | 0.361591 | 8.282361 | 8.133484 | 8.591363 | 6.925129 | 7.03424 | 10.166302 | 3.217004 | 3.402194 | 3.513443 | 9.762891 | 4.875592 | 15.209148 | 10.680373 | 30.71347 | 36.625 | 8.458663 | [illegible] |
| 36.5486111 | 6.319744 | 1.721853 | 1.569097 | 0.949993 | 0.889118 | 2.222781 | 3.18973 | 2.238714 | 0.68809 | 0.764904 | 9.583635 | 9.634458 | 10.779669 | 4.662954 | 3.521116 | 3.860192 | 3.02588 | 2.793746 | 0.960241 | 5.884918 | 3.104247 | 12.589495 | 8.859269 | 20.409029 | 36.6388889 | 3.424766 | [illegible] |
| 36.5625 | 5.813858 | 1.287828 | 0.817419 | -0.008556 | 0.132873 | 0.617477 | 1.113785 | -0.177867 | 0.853248 | 1.178543 | 6.850877 | 8.09246 | 8.464117 | 2.864853 | 1.371228 | 0.548338 | 2.393154 | 2.283585 | 1.191789 | 2.336283 | 2.221962 | 5.959813 | 3.523295 | 6.930962 | 36.6527778 | -4.487224 | [illegible] |
| 36.5763889 | 5.395851 | 1.062923 | 0.227123 | -0.558885 | 0.118153 | 0.02129 | 1.110331 | 0.039871 | 1.069125 | 1.508781 | 3.769972 | 5.066828 | 5.747286 | 3.087276 | 1.276551 | 0.674413 | 0.920518 | 0.357306 | 1.626828 | -0.046547 | 3.803157 | 1.757938 | -1.765506 | -0.135156 | 36.6666667 | -4.533556 | [illegible] |
| 36.5902778 | 6.141053 | 2.105708 | 0.514369 | -0.280479 | 0.20877 | 0.130095 | 1.390217 | 0.642161 | 1.240298 | 1.416452 | 4.731536 | 2.149345 | 3.67425 | 4.210321 | 2.772162 | 1.940604 | 0.396903 | -0.234997 | 1.15596 | 0.522988 | 4.453908 | -0.943805 | -3.543131 | 0.325621 | 36.6805556 | -0.499633 | [illegible] |
| 36.6041667 | 5.400879 | 3.420966 | 0.740354 | 0.519218 | 0.138886 | -0.595051 | 1.007567 | -0.33241 | 1.234892 | 0.693316 | 5.210208 | 0.617803 | 1.155571 | 4.179025 | 4.167783 | 3.552554 | 0.080187 | 0.614886 | 0.40413 | 1.412042 | 1.739348 | -3.020754 | 1.03705 | 3.636237 | 36.6944444 | 5.683373 | [illegible] |
| 36.6180556 | 4.82802 | 3.86555 | 1.099214 | 1.600157 | 0.326943 | -1.202401 | -0.066577 | -1.01201 | 1.02816 | -0.165269 | 2.516737 | -0.274214 | -0.91041 | 2.649551 | 3.65544 | 4.559532 | 0.041587 | 1.188604 | 0.355811 | 1.594799 | 2.719825 | -3.898309 | 1.862996 | 3.007216 | 36.7083333 | 8.273493 | [illegible] |
| 36.6319444 | 2.31443 | 2.592748 | 1.523823 | 1.889474 | -0.246143 | -1.910215 | -1.552715 | -1.667393 | 0.504119 | -1.529047 | -0.452417 | -0.598189 | -1.092944 | 0.450736 | 1.943881 | 3.5227 | 0.157656 | 1.248482 | 1.150686 | 1.002215 | 4.399481 | -0.395137 | 4.127371 | 4.093908 | 36.7222222 | 5.996124 | [illegible] |
| 36.6458333 | -0.597433 | 0.87338 | 0.986492 | 1.076334 | -0.085718 | -2.724444 | -3.370384 | -3.937025 | -0.163514 | -2.173041 | -3.170904 | -2.604957 | -2.01669 | -1.773842 | -0.293768 | -0.417413 | 0.441696 | 0.757668 | 1.356452 | 1.678941 | 4.657476 | 3.759734 | 5.789517 | 2.511103 | 36.7361111 | 3.682553 | [illegible] |
| 36.6597222 | -2.666907 | -0.930563 | 0.110228 | -0.182605 | -0.832733 | -4.170566 | -3.709295 | -5.117016 | -0.529381 | -1.939431 | -5.812008 | -4.776889 | -4.805946 | -3.852648 | -3.611907 | -3.469646 | 1.078007 | 1.368228 | 0.800178 | 1.42565 | 0.695545 | 2.020527 | 2.879226 | -4.137079 | 36.75 | 3.588489 | [illegible] |
| 36.6736111 | -3.458073 | -2.773947 | -0.220769 | -1.350177 | -0.888595 | -3.93331 | -1.571006 | -2.868148 | -0.623858 | -0.610318 | -4.84012 | -5.216193 | -4.095019 | -3.648993 | -3.706529 | -3.68049 | 1.364535 | 1.228199 | 0.49415 | -0.097013 | -3.407106 | -5.236262 | -3.820819 | -12.364436 | 36.7638889 | 3.613195 | [illegible] |
| 36.6875 | -1.801132 | -2.116309 | 0.001147 | -1.69292 | -0.05503 | -1.504304 | 1.709482 | 0.384184 | -0.466725 | 0.332779 | 0.654346 | -7.65999 | -2.730553 | -3.342819 | -1.655853 | -1.607443 | 0.768549 | -0.350094 | -1.157874 | -0.512673 | -4.70748 | -9.060548 | -6.551897 | -16.793134 | 36.7777778 | 2.475748 | [illegible] |
| 36.7013889 | 0.296034 | -0.395771 | 0.768809 | -0.504742 | 0.6016 | 2.904899 | 2.62658 | 2.877164 | 0.052051 | 0.520916 | 6.745078 | -3.721614 | -1.802205 | 0.794723 | 1.080158 | 0.099211 | -0.048819 | -1.705405 | -3.384479 | -0.062098 | -0.995304 | -11.488033 | -5.978134 | -11.899368 | 36.7916667 | 2.252968 | [illegible] |
| 36.7152778 | 1.558643 | 1.769207 | 1.526269 | 1.068605 | 1.648049 | 3.489675 | 1.336994 | 2.554853 | 0.48531 | -0.241861 | 10.421806 | -2.613704 | -3.680338 | 4.046026 | 2.329058 | 1.31382 | -0.094729 | -2.055603 | -0.991017 | 2.889393 | 2.640224 | -6.983385 | 1.77572 | 0.006893 | 36.8055556 | -0.142511 | [illegible] |
| 36.7291667 | 1.427798 | 1.615947 | 1.42947 | 1.288316 | 1.018528 | 4.229717 | 0.16188 | 1.070632 | 0.634867 | -0.754454 | 4.836714 | -0.333505 | -4.684837 | 3.439641 | 1.632422 | 0.892649 | 0.089187 | -0.357239 | 2.248051 | 4.730753 | 3.413557 | 7.568865 | 6.312987 | 7.425142 | 36.8194444 | -0.688433 | [illegible] |
| 36.7430556 | -0.003941 | -0.033392 | 0.247842 | 0.293667 | 0.385726 | 0.999224 | 0.406832 | 0.735583 | 0.976366 | 0.350692 | 1.645447 | 0.271513 | -1.519487 | 1.23134 | 0.094746 | 0.441395 | -0.278833 | 0.519314 | 4.479858 | 4.78099 | -1.292093 | 12.2081 | 3.802583 | 5.826593 | 36.8333333 | 0.928441 | [illegible] |
| 36.7569444 | -1.174176 | -1.037008 | -0.063203 | -0.154817 | 0.137838 | 0.307425 | 0.902157 | 0.753977 | 1.03944 | 1.69027 | -0.726566 | 0.298792 | 1.780614 | -1.19995 | -4.433129 | -2.396745 | -1.078803 | -2.607025 | 2.850172 | 0.148002 | -4.063599 | 4.95233 | 1.638702 | -7.300504 | 36.8472222 | 3.043175 | [illegible] |
| 36.7708333 | -2.977647 | -1.570663 | -0.224824 | -0.806887 | 0.099907 | -0.097675 | 0.318661 | 1.142313 | 1.056067 | 2.256395 | -2.280356 | -0.551791 | 3.047082 | -2.799914 | -5.616995 | -5.454844 | -1.474992 | -4.648721 | -0.270603 | -5.254372 | -5.18565 | -2.532173 | 0.517595 | -22.77221 | 36.8611111 | 1.489607 | [illegible] |
| 36.7847222 | -3.37902 | -1.255134 | -1.338733 | -1.39321 | 0.069487 | -0.730827 | -0.762496 | -0.735371 | 0.666823 | 2.270356 | -7.712166 | -4.873314 | 0.549864 | -5.930281 | -4.336447 | 0.075475 | -0.63719 | -1.145357 | -4.361172 | -7.344821 | -0.153126 | -11.974942 | 1.903306 | -17.483346 | 36.875 | -4.14774 | [illegible] |
| 36.7986111 | -2.577868 | 0.142311 | -1.322315 | -1.399707 | -0.160513 | -0.474241 | -0.639813 | -1.880512 | 0.096841 | 0.768409 | -11.569515 | -6.85115 | 1.060252 | -5.808475 | 1.980402 | 7.269809 | -0.092952 | 3.036028 | -6.036114 | -2.662212 | 9.412407 | -14.937006 | 4.141111 | 1.45384 | 36.8888889 | -6.995267 | [illegible] |
| 36.8125 | -1.232985 | 0.28452 | -0.761458 | -0.635149 | 0.356402 | -0.467671 | 0.399992 | -1.379341 | -0.556209 | -0.658714 | -6.231497 | -4.889693 | -0.159705 | -0.428437 | 3.874004 | 5.815864 | 0.223567 | 2.04105 | -2.640873 | 1.23049 | 4.173676 | -4.878713 | 6.120185 | 13.046281 | 36.9027778 | -4.957502 | [illegible] |
| 36.8263889 | -0.049155 | 0.074255 | 0.288441 | -0.307308 | 0.851256 | -1.375206 | 1.839085 | 0.808363 | -1.450333 | -1.888651 | -0.337944 | -3.998074 | -2.235875 | 2.226119 | 1.2353 | 2.154443 | -0.090034 | -2.302768 | 2.940029 | 2.049853 | -1.878715 | 8.191467 | 2.690967 | 2.625453 | 36.9166667 | -3.093373 | [illegible] |
| 36.8402778 | 0.150999 | -0.083433 | 1.491019 | 0.907898 | 0.777159 | -1.695869 | 0.052137 | -0.989017 | -2.681893 | -3.978011 | 2.361124 | -2.069083 | -1.332031 | 3.464972 | -1.234054 | -1.928945 | -2.231639 | -4.83702 | 3.91118 | -2.315072 | -4.600197 | 11.289778 | -0.995974 | -5.36523 | 36.9305556 | -3.961119 | [illegible] |
| 36.8541667 | -0.612671 | -0.545317 | 0.882956 | 1.173071 | 1.344905 | -0.643723 | -2.278379 | -4.358281 | -2.254669 | -2.354257 | 1.550356 | -0.312047 | -2.562189 | 4.836577 | -0.592756 | -1.34152 | -3.0514 | 2.013963 | 2.911591 | -4.886298 | -8.025318 | 6.93926 | -3.697037 | -2.759698 | 36.9444444 | -2.481964 | [illegible] |
| 36.8680556 | -2.313986 | -0.640469 | -0.596924 | -0.743831 | -1.196365 | -1.243963 | -3.860205 | -5.291188 | -1.122904 | -0.335333 | -2.663801 | -0.633935 | -2.488788 | 3.998583 | -0.153508 | 0.3033 | -1.973433 | 2.651233 | 1.025757 | -0.63216 | 0.367775 | 4.7357 | -1.248699 | -0.782754 | 36.9583333 | -0.982585 | [illegible] |
| 36.8819444 | -3.737427 | -0.632004 | -2.074237 | -2.855823 | -1.530565 | -1.031112 | -3.758931 | -2.185441 | -0.631254 | 0.17403 | -7.829528 | -4.923308 | -4.523055 | -0.038752 | -1.704697 | -2.191198 | -1.582388 | -2.793598 | -0.766589 | 2.486945 | 8.393989 | -2.808961 | -0.386184 | -4.264459 | 36.9722222 | -0.559784 | [illegible] |
| 36.8958333 | -4.932889 | -1.353878 | -3.538039 | -5.068468 | -3.169678 | -1.279312 | -3.376294 | 0.564931 | -1.336 | -1.20368 | -10.548181 | -9.27256 | -5.527256 | -4.134152 | -4.27535 | -3.984604 | -1.574371 | -8.024689 | -3.011858 | -0.462379 | -0.604424 | -8.759808 | -0.922791 | -18.24605 | 36.9861111 | -2.468835 | [illegible] |
| 36.9097222 | -5.093311 | -3.258044 | -4.359642 | -5.636182 | -3.026178 | -2.193978 | -2.401136 | -0.99003 | -2.113282 | -3.649563 | -8.427494 | -8.803092 | -2.652102 | -2.478389 | -6.448432 | -4.706326 | -2.529632 | -7.416146 | -5.023694 | -5.603962 | -8.850673 | -12.008288 | -4.256403 | -24.125386 | 37 | -5.481475 | [illegible] |
| 36.9236111 | -4.416712 | -4.108949 | -2.308958 | -3.198176 | -0.541582 | -2.840182 | -1.683888 | -2.442011 | -3.77669 | -5.736891 | -7.090761 | -8.016724 | -0.150298 | -2.708409 | -1.565029 | 0.39008 | -2.901228 | -2.378426 | -6.257632 | -10.808131 | 0.683047 | -15.758006 | -11.117799 | -18.006873 | 37.0138889 | -6.97662 | [illegible] |
| 36.9375 | -4.694117 | -5.372755 | -1.422754 | -1.040885 | -0.785261 | -4.190261 | -1.299487 | -3.29005 | -5.345617 | -5.527829 | -15.572844 | -12.487405 | -1.707654 | -4.520562 | -1.560632 | 1.674137 | -4.569705 | 0.924572 | -10.98243 | -16.447698 | -1.326938 | -9.162865 | -14.035763 | -26.113308 | 37.0277778 | -8.524974 | [illegible] |
| 36.9513889 | -4.382138 | -6.678003 | -1.747586 | -1.380541 | -3.203393 | -3.441477 | -3.808877 | -2.863995 | -6.131749 | -5.682553 | -25.136705 | -17.194938 | -4.433616 | -6.945601 | -9.050425 | -9.777367 | -11.580803 | -6.914217 | -13.404295 | -4.329718 | -3.703669 | -2.614631 | -11.740982 | -46.190005 | 37.0416667 | -10.037674 | [illegible] |
| 36.9652778 | -4.239599 | -5.69163 | -1.372789 | -2.180543 | -5.173089 | -4.350116 | -4.964443 | -2.957516 | -5.658124 | -5.929715 | -25.472894 | -13.612593 | -4.849452 | -7.666164 | -10.816967 | -10.85387 | -17.159642 | -10.269147 | -14.074288 | -7.296787 | -8.8704 | -4.665511 | -8.303064 | -53.030554 | 37.0555556 | -11.145586 | [illegible] |
| 36.9791667 | -3.630853 | -3.987698 | -1.51603 | -3.282595 | -6.718372 | -5.643888 | -5.001982 | -3.050419 | -3.142696 | -4.452935 | -16.548651 | -11.845018 | -2.46541 | -8.940786 | -10.561616 | -9.157561 | -14.516432 | -8.742974 | -9.630958 | -6.19286 | -6.399633 | -9.243962 | -8.04879 | -42.521806 | 37.0694444 | -11.714143 | [illegible] |
| 36.9930556 | -2.645769 | -2.264891 | -1.840737 | -3.839862 | -4.914843 | -3.611123 | -3.845739 | -2.361865 | -2.016715 | -3.330726 | -9.830375 | -6.459718 | 1.114804 | -8.307393 | -8.48747 | -9.246251 | -14.301232 | -7.204751 | -3.954217 | -0.977971 | 0.061843 | -9.88564 | -8.492869 | -25.995315 | 37.0833333 | -11.31421 | [illegible] |
| 37.0069444 | -1.493388 | -0.769426 | -1.955845 | -3.370767 | -3.109182 | -1.549882 | -2.561678 | -1.284725 | -1.866059 | -2.82358 | -6.181656 | -4.512025 | 2.248986 | -3.262488 | -2.607798 | -4.584182 | -8.164743 | -3.302545 | 0.148389 | -0.412941 | -2.245031 | -10.744299 | -7.320471 | -9.756116 | 37.0972222 | -9.51377 | [illegible] |
| 37.0208333 | -0.852647 | 0.216228 | -1.416166 | -2.240617 | -2.748026 | -0.823313 | -1.871614 | -0.615837 | -1.658096 | -2.72118 | -3.893869 | -3.597556 | 3.15821 | 2.039997 | 0.900462 | -1.388822 | -0.97947 | 0.102588 | 3.261633 | 3.095007 | 0.386982 | -10.015899 | -5.026669 | 2.262193 | 37.1111111 | -6.158228 | [illegible] |
| 37.0347222 | -0.63795 | 0.511414 | -1.021125 | -0.945094 | -2.300613 | -0.548828 | -2.21995 | -0.962042 | -1.60622 | -3.144546 | -2.946548 | -2.723491 | 1.838245 | 1.931684 | 1.697083 | -1.090433 | 1.263828 | 0.387586 | -0.73304 | 2.89593 | 2.324968 | -6.193823 | -3.238076 | 4.269077 | 37.125 | -2.916397 | [illegible] |
| 37.0486111 | -0.659446 | 0.376999 | -0.812624 | -0.461918 | -1.912129 | -0.320394 | -2.522224 | -1.345892 | -1.728729 | -3.408008 | -2.388182 | -5.182506 | -2.612023 | -0.27664 | 0.177518 | -2.486496 | -0.488133 | -2.176326 | -3.144811 | -0.823413 | 0.407656 | -5.42735 | -1.966129 | -0.765355 | 37.1388889 | -0.385701 | [illegible] |
| 37.0625 | -0.63926 | 0.230203 | -0.588952 | -0.473452 | -1.346499 | -0.275053 | -2.188521 | -1.317052 | -1.437409 | -2.675795 | -1.512639 | -6.120403 | -4.553733 | -2.649244 | -2.412145 | -4.156387 | -2.564878 | -4.676523 | -3.792377 | -6.665001 | -3.262672 | -3.62832 | -1.632756 | -5.342062 | 37.1527778 | 2.015182 | [illegible] |
| 37.0763889 | -0.59404 | 0.119508 | -0.448363 | -0.695996 | -0.890576 | -0.308798 | -1.496818 | -0.988032 | -1.038535 | -1.779125 | -0.789919 | -6.663862 | -5.294216 | -4.226135 | -4.730118 | -5.67248 | -3.659175 | -6.416174 | -7.433705 | -10.002407 | -5.580839 | -2.325587 | -1.51156 | -7.364129 | 37.1666667 | 4.105218 | [illegible] |
| 37.0902778 | -0.563671 | -0.179137 | -0.524676 | -1.092521 | -0.60096 | -0.27039 | -0.902716 | -0.742478 | -0.896218 | -1.513432 | -0.573129 | -7.319225 | -6.244515 | -5.512594 | -6.255648 | -6.611311 | -3.161937 | -6.126881 | -11.685907 | -9.239349 | -4.005852 | -2.086846 | -1.60888 | -8.318184 | 37.1805556 | 4.581709 | [illegible] |
| 37.1041667 | -0.593515 | -0.647018 | -0.529865 | -1.100216 | -0.259343 | -0.160242 | -0.761551 | -0.556391 | -1.042283 | -1.676696 | -0.556934 | -6.368559 | -5.745821 | -5.093141 | -6.778303 | -6.790516 | -3.525319 | -5.093663 | -10.678914 | -6.271367 | -2.23626 | -2.791806 | -1.562425 | -8.220148 | 37.1944444 | 3.103633 | [illegible] |
| 37.1180556 | -0.641375 | -1.103255 | -0.387903 | -0.205808 | -0.161414 | -0.372258 | -1.037416 | -0.568247 | -1.255554 | -2.065697 | -0.751092 | -4.953097 | -4.356288 | -4.794766 | -6.484108 | -5.403466 | -5.535262 | -5.022232 | -13.46573 | -4.915469 | -1.601152 | -4.222574 | -1.991898 | -7.89992 | 37.2083333 | 1.007483 | [illegible] |
| 37.1319444 | -0.770559 | -1.44675 | -0.269562 | 0.358976 | -0.668538 | -1.141101 | -1.383324 | -0.685957 | -1.537261 | -2.446211 | -1.294993 | -4.125546 | -3.600854 | -4.078892 | -4.669967 | -2.488954 | -6.59711 | -4.000431 | -9.874062 | -3.089795 | -1.577172 | -4.783162 | -2.017881 | -6.484784 | 37.2222222 | -0.792677 | [illegible] |
| 37.1458333 | -1.063812 | -1.604336 | -0.278678 | -0.245254 | -1.817723 | -2.291879 | -1.445709 | -0.613459 | -1.542932 | -2.439845 | -1.888837 | -2.394572 | -1.871229 | -2.607164 | -2.768392 | -0.564305 | -5.28729 | -1.984304 | -4.766154 | -1.078589 | -1.007118 | -2.801098 | -0.987606 | -3.671296 | 37.2361111 | -1.215301 | [illegible] |
| 37.1597222 | -1.313658 | -1.52969 | -0.347087 | -0.932656 | -3.132291 | -3.36693 | -1.528013 | -0.595717 | -1.188583 | -2.09818 | -2.133593 | -0.443121 | -0.209431 | -0.825115 | -0.51531 | -0.135419 | -3.118729 | -0.277471 | -1.480337 | 1.394065 | 0.6483 | 0.26097 | 0.324909 | -0.80159 | 37.25 | -0.910188 | [illegible] |
| 37.1736111 | -1.424088 | -1.354434 | -0.56997 | -1.434916 | -4.154627 | -3.826002 | -1.559383 | -0.586998 | -0.769263 | -1.472306 | -2.227844 | 0.174206 | -0.025171 | -0.195747 | 0.528788 | -0.218606 | -1.264374 | 1.335052 | -0.483248 | 2.647825 | 2.273571 | 1.724217 | 1.336708 | 1.177965 | 37.2638889 | -0.361585 | [illegible] |
| 37.1875 | -1.347084 | -1.054834 | -0.781575 | -1.373109 | -4.668274 | -3.613069 | -1.650779 | -0.625153 | -0.407604 | -0.781251 | -2.202939 | 0.374813 | -0.345608 | -0.170875 | 1.168388 | 0.425514 | 0.80822 | 1.492784 | -1.117026 | 2.899155 | 3.478235 | 1.201106 | 1.692459 | 2.297613 | 37.2777778 | 0.602325 | [illegible] |
| 37.2013889 | -1.342707 | -0.724968 | -0.616926 | -0.953481 | -4.974483 | -2.966568 | -1.569927 | -0.579588 | -0.242716 | -0.304094 | -2.426699 | 1.043586 | -0.093837 | 0.557972 | 2.343247 | 1.73589 | 3.048977 | 1.363329 | -1.324711 | 3.041437 | 4.20283 | 0.026913 | 1.934151 | 2.841149 | 37.2916667 | 1.529043 | [illegible] |
| 37.2152778 | -1.450111 | -0.544283 | -0.706376 | -0.416135 | -5.204231 | -2.107608 | -1.526706 | -0.452937 | -0.332596 | -0.091919 | -2.884477 | 1.924065 | 0.140192 | 1.876466 | 3.65534 | 2.911918 | 4.517486 | 1.316498 | 0.95726 | 3.887604 | 4.402714 | 0.181504 | 2.59595 | 3.265315 | 37.3055556 | 2.268584 | [illegible] |
| 37.2291667 | -1.448123 | -0.316633 | -0.873653 | -0.615708 | -4.246089 | -1.280565 | -1.141611 | -0.174561 | -0.414483 | 0.000105 | -2.864685 | 2.890403 | 0.796239 | 3.099909 | 4.161679 | 2.886267 | 5.298545 | 2.112584 | 6.053474 | 6.065664 | 4.489377 | 1.367051 | 3.364933 | 4.069148 | 37.3194444 | 2.490896 | [illegible] |
| 37.2430556 | -1.101905 | 0.146803 | -0.689233 | -0.597468 | -2.103099 | -0.263115 | -0.698248 | -0.110586 | -0.351084 | 0.256852 | -2.006922 | 3.947952 | 1.893007 | 3.583498 | 4.431842 | 2.75804 | 6.421401 | 4.160641 | 10.077685 | 7.686537 | 4.742937 | 2.593462 | 3.513597 | 5.076446 | 37.3333333 | 2.719808 | [illegible] |
| 37.2569444 | -0.711831 | 0.66348 | -0.359432 | -0.585553 | 0.136226 | 0.992544 | -0.242388 | -0.242073 | 0.012482 | 0.837231 | -0.755586 | 4.864094 | 3.045654 | 2.984948 | 4.220467 | 3.485496 | 8.200996 | 6.732116 | 9.694843 | 8.044483 | 5.347265 | 3.529982 | 2.682195 | 5.950526 | 37.3472222 | 3.720792 | [illegible] |
| 37.2708333 | -0.326078 | 1.061429 | -0.088834 | -0.374814 | 2.526074 | 2.335683 | 0.113003 | -0.144465 | 0.531924 | 1.523408 | 0.496532 | 4.57717 | 2.896583 | 2.408515 | 3.984947 | 3.996433 | 9.996915 | 8.814281 | 9.28397 | 7.976704 | 6.375346 | 3.993462 | 2.208375 | 6.553184 | 37.3611111 | 4.887443 | [illegible] |
| 37.2847222 | 0.010911 | 1.314583 | 0.170184 | 0.283708 | 3.980309 | 2.972172 | 0.378025 | 0.225289 | 1.169386 | 2.056064 | 1.337255 | 4.126325 | 2.390473 | 3.301259 | 4.387522 | 4.296865 | 9.80278 | 9.394724 | 7.833083 | 7.577423 | 7.489153 | 4.580603 | 2.433628 | 6.881856 | 37.375 | 4.888132 | [illegible] |
| 37.2986111 | 0.338275 | 1.589918 | 0.377839 | 1.146592 | 4.408062 | 3.381696 | 0.654952 | 0.541251 | 1.385304 | 2.270668 | 1.9323 | 4.568529 | 2.297591 | 4.427378 | 5.185372 | 4.824903 | 8.857785 | 10.320462 | 7.220652 | 5.913402 | 7.289558 | 5.988357 | 3.237372 | 7.417421 | 37.3888889 | 3.707386 | [illegible] |
| 37.3125 | 0.572395 | 1.689769 | 0.428208 | 1.65277 | 4.53468 | 4.098865 | 1.020952 | 0.73296 | 1.526746 | 2.365856 | 2.307565 | 5.491966 | 2.732376 | 5.045306 | 5.886662 | 5.342757 | 9.941872 | 11.537846 | 6.883765 | 4.531801 | 6.876231 | 6.612167 | 3.98566 | 8.448413 | 37.4027778 | 2.275432 | [illegible] |
| 37.3263889 | 0.752842 | 1.522118 | 0.322038 | 1.371322 | 4.831315 | 4.96246 | 1.223302 | 0.892675 | 1.405627 | 2.3191 | 2.553546 | 6.306328 | 3.527252 | 5.408724 | 6.596955 | 5.185229 | 11.227318 | 11.65529 | 6.041129 | 7.14173 | 6.481126 | 6.403922 | 4.058476 | 9.247615 | 37.4166667 | 1.058387 | [illegible] |
| 37.3402778 | 0.884305 | 1.367767 | 0.191305 | 0.893521 | 5.532017 | 5.553673 | 1.148423 | 1.158686 | 1.04112 | 1.979953 | 2.937313 | 7.414191 | 4.599547 | 5.973665 | 7.256615 | 5.608287 | 10.683103 | 10.880111 | 7.014752 | 10.029514 | 6.3357 | 7.228308 | 2.666073 | 9.450779 | 37.4305556 | 1.089243 | [illegible] |
| 37.3541667 | 1.184059 | 1.545838 | 0.330008 | 0.936298 | 7.102564 | 6.555523 | 1.239786 | 1.719748 | 0.759159 | 1.944562 | 4.052408 | 9.075717 | 5.060616 | 6.425667 | 8.256877 | 6.970905 | 10.003164 | 11.040096 | 6.678376 | 7.939834 | 6.794665 | 8.272187 | 1.812108 | 11.204583 | 37.4444444 | 2.163768 | [illegible] |
| 37.3680556 | 1.505103 | 1.987931 | 0.727695 | 1.68449 | 6.38632 | 5.441393 | 1.99252 | 2.205642 | 0.82296 | 2.480607 | 5.812374 | 9.904696 | 4.648953 | 7.29822 | 9.429192 | 8.729245 | 9.623368 | 11.34928 | 5.626444 | 5.648482 | 7.344353 | 9.286818 | 2.429463 | 14.476382 | 37.4583333 | 2.668299 | [illegible] |
| 37.3819444 | 1.78203 | 2.265716 | 1.075148 | 2.53406 | 3.612117 | 3.470828 | 3.391042 | 2.574011 | 1.085029 | 3.405337 | 7.535239 | 10.823346 | 5.1496 | 7.121914 | 9.818663 | 9.974026 | 8.84778 | 12.637515 | 10.372782 | 6.311022 | 6.108332 | 10.709806 | 4.611817 | 15.934347 | 37.4722222 | 2.618283 | [illegible] |
| 37.3958333 | 2.044449 | 2.552472 | 1.24402 | 2.984878 | 4.374795 | 4.247474 | 3.286472 | 2.885893 | 1.381296 | 4.477716 | 8.277233 | 10.142026 | 5.129333 | 4.243465 | 7.92924 | 8.993742 | 6.19753 | 11.500535 | 9.876989 | 8.085551 | 4.38683 | 12.256049 | 7.314243 | 15.596132 | 37.4861111 | 3.122034 | [illegible] |
| 37.4097222 | 2.469759 | 2.807381 | 1.295898 | 2.977103 | 4.242984 | 4.9321 | 3.47165 | 3.336915 | 1.887732 | 5.387424 | 7.404047 | 10.770995 | 4.903344 | 2.954331 | 4.636222 | 5.901465 | 4.993994 | 9.784882 | 4.192324 | 5.013596 | 4.474498 | 11.320043 | 4.432257 | 15.190382 | 37.5 | 3.975231 | [illegible] |
| 37.4236111 | 2.563332 | 2.809571 | 1.280218 | 2.231498 | 2.81377 | 4.370651 | 2.866504 | 3.533639 | 2.572274 | 5.364285 | 5.854386 | 11.124648 | 4.298919 | 1.896006 | 1.844176 | 2.289972 | 5.721625 | 9.363776 | 0.974339 | 5.405615 | 7.399823 | 8.839728 | 1.885225 | 15.618255 | 37.5138889 | 5.550476 | [illegible] |
| 37.4375 | 2.513277 | 3.298001 | 1.94177 | 1.035978 | 2.232597 | 2.930068 | 3.289638 | 3.496446 | 4.399273 | 6.483528 | 9.910199 | 12.165324 | 4.631656 | 2.732441 | 3.886693 | 5.164073 | 7.534929 | 12.401926 | 1.879557 | 9.056506 | 12.541067 | 8.16014 | 5.048009 | 21.088481 | 37.5277778 | 9.959868 | [illegible] |
| 37.4513889 | 3.117723 | 4.719683 | 3.608844 | 0.130787 | 4.344404 | 2.916452 | 4.178311 | 2.356572 | 6.604834 | 9.177306 | 26.985275 | 15.419165 | 6.389293 | 4.867086 | 9.920765 | 11.503292 | 9.722396 | 16.247158 | 6.892275 | 8.294134 | 11.155649 | 8.519086 | 9.861082 | 34.650546 | 37.5416667 | 17.801825 | [illegible] |
| 37.4652778 | 3.766609 | 4.589838 | 4.469278 | 0.02346 | 5.392812 | 2.358558 | 4.027992 | 0.620088 | 6.72748 | 8.150277 | 36.458239 | 15.151623 | 5.60275 | 6.428307 | 15.485538 | 17.383938 | 8.633918 | 14.932499 | 9.396074 | 11.236614 | 6.72755 | 16.01528 | 8.469278 | 41.721821 | 37.5555556 | 19.056264 | [illegible] |
| 37.4791667 | 4.763346 | 2.200275 | 2.8926 | 1.20993 | 1.873939 | -0.716123 | 1.079045 | -1.196901 | 2.926917 | 3.11164 | 17.677483 | 3.315931 | 2.498601 | 2.473453 | 9.414207 | 14.227298 | 3.923817 | 5.565833 | 4.57996 | 3.2533 | 2.943869 | 5.679276 | 4.55052 | 20.716387 | 37.5694444 | 10.278165 | [illegible] |
| 37.4930556 | 2.348077 | 1.252225 | 2.294323 | 2.184182 | -2.307314 | -2.126118 | -0.437177 | -0.40255 | 0.505092 | 0.862203 | 2.937754 | -8.896002 | 0.905365 | 0.302661 | 3.697921 | 7.426074 | 0.377857 | 3.84355 | 1.004545 | -0.803277 | 6.788468 | 1.455746 | 0.558686 | 7.270315 | 37.5833333 | 3.134663 | [illegible] |
| 37.5069444 | 1.725611 | 1.659584 | 3.273188 | 2.850138 | -1.411154 | -1.417472 | 2.453189 | 2.216327 | 0.563059 | 1.925836 | 5.927963 | -6.694044 | 2.131914 | 0.493459 | 2.32204 | 4.506517 | -0.305201 | 3.522199 | 0.158386 | 1.965232 | 8.224667 | -2.584198 | 3.383611 | 13.786415 | 37.5972222 | 1.753439 | [illegible] |
| 37.5208333 | 2.579607 | 2.091151 | 3.72362 | 2.143653 | 1.196679 | 1.067465 | 4.995039 | 3.955432 | 1.123641 | 3.317753 | 10.279687 | 1.35302 | -0.545779 | 2.545454 | 0.900945 | 2.631616 | 0.616395 | 1.785149 | -2.191577 | 2.684814 | 4.500114 | -2.286057 | 6.394022 | 21.153866 | 37.6111111 | 2.013749 | [illegible] |
| 37.5347222 | 3.480151 | 1.936954 | 4.180963 | 2.17692 | 1.074522 | 0.948716 | 3.945926 | 4.505451 | 1.510293 | 3.263727 | 10.391356 | 7.754103 | 5.53574 | 4.49978 | 0.882856 | 2.51623 | 1.468127 | 1.92994 | -2.870418 | 3.353323 | 2.23545 | -1.554315 | 7.177624 | 21.084358 | 37.625 | 1.697471 | [illegible] |
| 37.5486111 | 3.865524 | 1.489784 | 2.995947 | 1.931601 | 1.264961 | 1.222104 | 1.802775 | 1.983908 | 1.26667 | 1.443355 | 11.204686 | 9.795201 | 6.465399 | 4.621856 | 1.71973 | 6.630698 | 1.115651 | 0.763496 | -0.719885 | 3.103291 | 4.099808 | 10.644354 | 3.953802 | 17.883991 | 37.6388889 | 3.261955 | [illegible] |
| 37.5625 | 3.529102 | 1.066705 | 1.435596 | 1.193605 | 0.875636 | 1.942525 | 0.830428 | -0.262869 | 0.662651 | -1.66074 | 10.127244 | 11.040356 | 6.187113 | 4.398536 | 2.298194 | 6.009931 | 1.110403 | 0.940274 | 3.055836 | 2.968587 | 0.787624 | 15.974576 | 3.183453 | 15.629151 | 37.6527778 | 6.910704 | [illegible] |
| 37.5763889 | 3.746719 | 0.585055 | 0.257143 | -0.456063 | 0.128293 | 0.885908 | -0.660742 | -1.744338 | 0.628219 | -4.156722 | 4.178552 | 7.338104 | 6.848636 | 3.25594 | 3.023314 | 5.1476 | 1.311244 | 0.322621 | 4.981721 | 2.695237 | 1.929014 | 4.524673 | -1.087914 | 14.947345 | 37.6666667 | 10.127716 | [illegible] |
| 37.5902778 | 3.004203 | 0.432803 | -0.894305 | -1.386997 | -1.367738 | -1.573037 | -1.102394 | -1.277691 | -0.459762 | -4.428417 | -3.598192 | 4.645349 | 6.552977 | 1.846782 | 2.513916 | 3.505642 | 1.781011 | 0.420696 | 3.922812 | 3.242013 | 1.782437 | -2.53739 | -1.188175 | 9.139563 | 37.6805556 | 8.430097 | [illegible] |
| 37.6041667 | 2.086884 | 0.527554 | -0.036506 | -0.34659 | -2.026157 | -2.07957 | -0.852245 | 0.390147 | -0.028559 | -2.971314 | -13.221044 | 1.220386 | 3.462416 | 1.276132 | 1.920254 | 1.862726 | 1.700132 | 0.383967 | 1.243664 | 2.418027 | -0.800317 | 6.789911 | 1.464681 | 4.120166 | 37.6944444 | 1.146353 | [illegible] |
| 37.6180556 | 0.63058 | 0.263887 | 0.346333 | 1.03346 | 0.033437 | 3.00164 | 0.49529 | 1.092898 | 1.792064 | 1.439996 | -5.768126 | 0.556298 | 0.438724 | 2.488768 | 2.197688 | 0.148285 | 1.052492 | 0.379362 | 0.658324 | 0.552529 | -0.999328 | 11.523192 | 2.514522 | 3.23034 | 37.7083333 | -5.419043 | [illegible] |
| 37.6319444 | 0.094187 | -0.072922 | 0.65023 | 1.200667 | 1.658797 | 3.453163 | 1.906836 | 2.132097 | 2.450295 | 4.896298 | 5.525671 | 2.802106 | -1.078605 | 3.761424 | 1.498847 | -0.10626 | 0.843312 | 1.98452 | 0.689201 | -0.175602 | -0.208122 | 12.809582 | 3.345106 | 7.558083 | 37.7222222 | -5.923719 | [illegible] |
| 37.6458333 | -0.230995 | 0.553 | 0.945966 | 1.551014 | 1.02796 | 1.176968 | 2.42623 | 1.927959 | 0.908491 | 3.416353 | 9.996013 | 5.8509 | 1.956347 | 3.959308 | 1.842509 | 0.274935 | 0.812352 | 3.275469 | 1.023376 | -0.703015 | 2.495947 | 4.076816 | 1.344083 | 11.769366 | 37.7361111 | -2.979641 | [illegible] |
| 37.6597222 | 0.024414 | 0.436536 | 1.647029 | 1.248484 | -0.896222 | -0.62874 | 1.03221 | 0.713186 | 0.145317 | 0.648249 | 6.614053 | 3.257341 | 1.56032 | 2.546625 | 1.352069 | 0.405523 | 0.533803 | 3.609287 | 1.591804 | -1.419379 | 3.050849 | -0.482963 | -2.0479 | 5.781502 | 37.75 | 0.498368 | [illegible] |
| 37.6736111 | -0.299984 | 0.238111 | 0.852421 | 1.350504 | -2.822711 | -3.278348 | -0.864572 | -0.829564 | -0.196364 | -2.286654 | -0.481846 | -3.481137 | -2.300244 | -0.790687 | -0.266354 | -0.433574 | 0.041996 | 0.409995 | 1.621604 | -0.65577 | 1.660393 | -2.048825 | -1.97625 | -7.742165 | 37.7638889 | 3.846559 | [illegible] |
| 37.6875 | 0.219271 | 0.359852 | 0.806333 | 0.647628 | -2.507108 | -1.526998 | -2.868342 | -4.486622 | -0.873463 | -3.169971 | -7.201006 | -6.017293 | -3.821629 | -3.362421 | -1.54039 | -1.444852 | 0.120935 | -1.207758 | 1.370372 | 1.207419 | -0.003904 | -4.267974 | -8.558956 | -16.296545 | 37.7777778 | 4.565585 | [illegible] |
| 37.7013889 | -0.4786 | 0.385532 | 0.295901 | -0.42705 | -2.390379 | -0.665564 | -1.699395 | -2.609062 | -0.570959 | 0.056249 | -6.572094 | -6.499386 | -0.011402 | -4.309444 | -0.908367 | 1.014027 | 1.194036 | -0.738575 | 0.841193 | 2.764166 | -1.129673 | -0.327694 | -3.721688 | -14.795421 | 37.7916667 | 2.097376 | [illegible] |
| 37.7152778 | -1.048382 | 0.013959 | 0.527307 | -0.436131 | -1.210669 | 0.531757 | -0.611508 | -0.510219 | 0.758971 | 3.726153 | -4.659526 | -10.633641 | -0.6195 | -6.330474 | -0.509396 | 2.840792 | 1.533764 | 2.48128 | -0.230882 | 3.934727 | -3.638813 | 2.769328 | 0.180285 | -2.691267 | 37.8055556 | -0.602292 | [illegible] |
| 37.7291667 | -1.121487 | -0.201439 | 0.085691 | 0.029927 | -2.04354 | 0.489191 | -1.236171 | 0.645503 | 1.342825 | 3.300044 | -4.815919 | -10.5671 | -0.495915 | -3.791761 | -0.263331 | 3.031753 | 1.11835 | 2.052016 | -1.227402 | 0.320487 | -6.581373 | 3.920136 | 5.102256 | 3.03755 | 37.8194444 | -2.314257 | [illegible] |
| 37.7430556 | -0.987866 | -0.395205 | -0.56386 | -0.395585 | -1.444647 | 0.495824 | -1.692392 | -0.858321 | 1.12009 | 2.070783 | -4.253148 | -5.711914 | -1.555164 | -0.407099 | -0.313377 | -0.616334 | 0.890808 | 1.123754 | -1.638869 | -1.425933 | -9.868169 | -0.39669 | 4.593783 | -2.558732 | 37.8333333 | -2.676911 | [illegible] |
| 37.7569444 | -0.852009 | -0.250997 | 0.265435 | 1.06868 | 1.446274 | 2.995443 | 0.056818 | -1.540428 | 0.961393 | -0.089963 | 2.176664 | -0.286732 | -4.11106 | 2.878458 | -0.023409 | -4.21796 | 0.866023 | -1.540651 | -1.564736 | -2.425588 | -4.703462 | -6.285574 | 3.258199 | -12.311067 | 37.8472222 | -4.698314 | [illegible] |
| 37.7708333 | -0.86712 | -0.196709 | 1.909331 | 2.328293 | 3.551299 | 4.176664 | 1.465937 | -1.028414 | 0.374076 | -2.415299 | 7.558849 | 2.706828 | -3.426601 | 3.323595 | 1.198753 | -4.017011 | 0.543027 | -3.226681 | -1.943487 | -2.322612 | -2.305891 | -10.276582 | -1.371391 | -15.516354 | 37.8611111 | -7.36439 | [illegible] |
| 37.7847222 | -1.538863 | -0.701958 | 2.742106 | 2.185054 | 3.15619 | 3.734611 | 1.165998 | -0.626891 | -0.221235 | -2.819085 | 8.866942 | 5.401925 | 0.988866 | 2.505793 | 1.310777 | 1.415626 | -1.076995 | -4.732113 | -2.891305 | -1.631861 | 2.944979 | -10.084995 | -2.056056 | -7.057007 | 37.875 | -11.029248 | [illegible] |
| 37.7986111 | -2.141777 | -1.255802 | 0.696859 | 1.643594 | 1.515739 | 3.145632 | 1.676676 | 1.590141 | 0.308284 | 0.611078 | 7.104964 | 5.271201 | 3.985407 | 1.40192 | 1.000386 | 4.725633 | -3.360502 | -5.081754 | -2.363799 | 0.136392 | 7.194546 | -4.649837 | -1.044823 | -2.247433 | 37.8888889 | -10.927532 | [illegible] |
| 37.8125 | -2.280443 | -1.524576 | -0.38485 | 0.324048 | 0.389372 | 0.543214 | 1.657965 | 2.735749 | 1.032898 | 4.004501 | 1.669941 | 0.439416 | 0.993193 | -0.952578 | -0.284144 | 1.664808 | -2.557963 | -3.465781 | 0.695895 | 1.403031 | 6.480423 | -3.661299 | -0.53521 | -3.617253 | 37.9027778 | -3.75798 | [illegible] |
| 37.8263889 | -2.122197 | -1.732653 | -0.914991 | -1.727268 | -1.38881 | -2.071025 | 1.40679 | 1.972747 | 1.502262 | 4.067545 | -8.463331 | -6.018273 | -1.458197 | -4.00171 | -1.415581 | -1.940525 | 1.581784 | 2.691863 | 2.518176 | 0.561721 | 0.200825 | 1.082606 | -2.04796 | -11.686218 | 37.9166667 | 2.841483 | [illegible] |
| 37.8402778 | -2.29162 | -2.588549 | -1.503243 | -2.56726 | -2.74579 | -0.908621 | 0.109456 | -0.021611 | 0.588844 | 1.427325 | -18.41514 | -8.953226 | -1.738111 | -6.238242 | -2.805425 | -3.203121 | 3.617994 | 2.743748 | 3.136931 | -1.015724 | -11.301658 | 5.117115 | -2.972259 | -11.136764 | 37.9305556 | 10.018195 | [illegible] |
| 37.8541667 | -2.026142 | -0.477643 | -1.90782 | -3.804727 | -4.254509 | -0.960839 | -2.195017 | -2.751909 | -1.434959 | -2.800414 | -20.057422 | -5.14209 | -0.784323 | -5.276082 | -3.389735 | -2.933882 | 0.499384 | 1.360371 | 1.396561 | -0.703971 | -5.246354 | 10.143828 | -1.324317 | -5.176476 | 37.9444444 | 9.973233 | [illegible] |
| 37.8680556 | [illegible] | [illegible] | [illegible] | [illegible] | [illegible] | [illegible] | [illegible] | [illegible] | [illegible] | [illegible] | [illegible] | [illegible] | [illegible] | [illegible] | [illegible] | [illegible] | [illegible] | [illegible] | [illegible] | [illegible] | [illegible] | [illegible] | [illegible] | [illegible] | 37.9583333 | [illegible] | [illegible] |
| 37.8819444 | [illegible] | [illegible] | [illegible] | [illegible] | [illegible] | [illegible] | [illegible] | [illegible] | [illegible] | [illegible] | [illegible] | [illegible] | [illegible] | [illegible] | [illegible] | [illegible] | [illegible] | [illegible] | [illegible] | [illegible] | [illegible] | [illegible] | [illegible] | [illegible] | 37.9722222 | [illegible] | [illegible] |
| 37.8958333 | [illegible] | [illegible] | [illegible] | [illegible] | [illegible] | [illegible] | [illegible] | [illegible] | [illegible] | [illegible] | [illegible] | [illegible] | [illegible] | [illegible] | [illegible] | [illegible] | [illegible] | [illegible] | [illegible] | [illegible] | [illegible] | [illegible] | [illegible] | [illegible] | 37.9861111 | [illegible] | [illegible] |
| 37.9097222 | [illegible] | [illegible] | [illegible] | [illegible] | [illegible] | [illegible] | [illegible] | [illegible] | [illegible] | [illegible] | [illegible] | [illegible] | [illegible] | [illegible] | [illegible] | [illegible] | [illegible] | [illegible] | [illegible] | [illegible] | [illegible] | [illegible] | [illegible] | [illegible] | 38 | [illegible] | [illegible] |
| 37.9236111 | [illegible] | [illegible] | [illegible] | [illegible] | [illegible] | [illegible] | [illegible] | [illegible] | [illegible] | [illegible] | [illegible] | [illegible] | [illegible] | [illegible] | [illegible] | [illegible] | [illegible] | [illegible] | [illegible] | [illegible] | [illegible] | [illegible] | [illegible] | [illegible] | 38.0138889 | [illegible] | [illegible] |

**Supplementary Table S7 (j)** Direct motion in 100 mM NaCl stressed Tomato and their respective controls under dimG day/night lighting conditions

| Time | dimGControl1 | dimGControl2 | dimGControl3 | dimGControl4 | dimGControl5 | dimGControl6 | dimGControl7 | dimGControl8 | dimGControl9 | dimGControl10 | dimGControl11 | dimGControl12 | dimGControlTank | dimGStress1 | dimGStress2 | dimGStress3 | dimGStress4 | dimGStress5 | dimGStress6 | dimGStress7 | dimGStress8 | dimGStress9 | dimGStress10 | dimGStressTank | | Time | Mean ControldimG | Mean StressdimG |
|---|---|---|---|---|---|---|---|---|---|---|---|---|---|---|---|---|---|---|---|---|---|---|---|---|---|---|---|---|
| 18 | -0.912584 | 0.632298 | 0.104055 | -0.083241 | -0.002849 | -0.32583 | -0.760588 | 0.000602 | -0.505864 | 0.117068 | -0.17306 | -1.83946 | -2.423506 | -0.338732 | -0.30295 | -0.69029 | 0.053185 | -0.005687 | -0.093674 | 0.003617 | 0.075817 | -0.420271 | -1.002942 | 0.789062 | | 18.09722222 | 0.258951 | 0.49336 |
| 18.013889 | -0.939992 | 0.162588 | -0.187539 | -0.037269 | -0.11235 | -0.365116 | -0.547296 | -0.699342 | -0.683746 | 0.113634 | -0.350953 | -1.306459 | -0.325102 | -0.395435 | -0.469676 | -0.929854 | -0.227241 | -0.227416 | -0.492923 | -0.070019 | -0.071607 | -0.390599 | -0.567425 | 0.022439 | | 18.11111111 | 0.063093 | 0.370954 |
| 18.027778 | -0.997027 | 0.038475 | -0.413183 | -0.107944 | -0.190857 | -0.446541 | -0.575839 | -1.207009 | -0.801889 | 0.104882 | -0.525071 | -0.335303 | -0.175343 | -0.46398 | -0.602534 | -0.86094 | -0.354667 | -0.340772 | -0.682225 | -0.12295 | -0.127105 | -0.533231 | -0.507067 | -0.2233 | | 18.125 | -0.04064 | 0.444188 |
| 18.041667 | -0.842602 | 0.00557 | -0.50053 | -0.236479 | -0.234449 | -0.528717 | -0.643809 | -1.392434 | -0.776654 | 0.074261 | -0.613302 | -0.052843 | -0.3181 | -0.435708 | -0.643058 | -0.650162 | -0.36014 | -0.34326 | -0.720916 | -0.108135 | -0.120142 | -0.626076 | -0.488009 | -0.207498 | | 18.13888889 | -0.075982 | 0.278917 |
| 18.055556 | -0.578463 | -0.001516 | -0.462717 | -0.311908 | -0.240082 | -0.527971 | -0.560552 | -1.199938 | -0.6238 | 0.038262 | -0.459138 | 0.001993 | -0.427388 | -0.342177 | -0.636603 | -0.470252 | -0.337727 | -0.346168 | -0.692934 | -0.126003 | -0.13357 | -0.610766 | -0.423174 | -0.174029 | | 18.15277778 | -0.093308 | 0.019811 |
| 18.069444 | -0.398455 | -0.031316 | -0.410907 | -0.330561 | -0.194732 | -0.463555 | -0.422889 | -0.875322 | -0.434386 | 0.004775 | -0.282718 | -0.020141 | -0.38839 | -0.251075 | -0.561971 | -0.338372 | -0.319979 | -0.364372 | -0.681408 | -0.166851 | -0.178396 | -0.542703 | -0.349551 | -0.202998 | | 18.16666667 | -0.076685 | -0.050764 |
| 18.083333 | -0.278169 | -0.082392 | -0.3285 | -0.246317 | -0.100114 | -0.376031 | -0.309833 | -0.604007 | -0.298275 | -0.009838 | -0.213196 | -0.058811 | -0.310518 | -0.171501 | -0.426399 | -0.231799 | -0.263897 | -0.330566 | -0.583755 | -0.176287 | -0.193233 | -0.426652 | -0.229723 | -0.23028 | | 18.18055556 | -0.063397 | 0.058439 |
| 18.097222 | -0.14108 | -0.118981 | -0.181737 | -0.122948 | -0.024695 | -0.272558 | -0.198889 | -0.369584 | -0.214073 | 0.000631 | -0.22157 | -0.073121 | -0.263685 | -0.128674 | -0.311894 | -0.156752 | -0.21138 | -0.284674 | -0.497952 | -0.163694 | -0.173875 | -0.306123 | -0.135832 | -0.18472 | | 18.19444444 | -0.103236 | 0.13249 |
| 18.111111 | -0.076946 | -0.146579 | -0.143215 | -0.054886 | 0.006367 | -0.236876 | -0.141638 | -0.232548 | -0.15135 | -0.000568 | -0.218712 | -0.059666 | -0.152831 | -0.111521 | -0.249146 | -0.111642 | -0.162285 | -0.234153 | -0.414372 | -0.146174 | -0.13388 | -0.231071 | -0.117904 | -0.059843 | | 18.20833333 | -0.133982 | 0.153171 |
| 18.125 | -0.172019 | -0.183295 | -0.179823 | -0.052928 | 0.005259 | -0.269826 | -0.165859 | -0.216586 | -0.112732 | -0.01432 | -0.158208 | 0.018015 | 0.079729 | -0.088108 | -0.19919 | -0.085946 | -0.126903 | -0.198968 | -0.352345 | -0.126567 | -0.094312 | -0.161225 | -0.098186 | 0.016221 | | 18.22222222 | -0.13025 | 0.10139 |
| 18.138889 | -0.3718 | -0.267977 | -0.259819 | -0.093524 | -0.018528 | -0.338486 | -0.243821 | -0.27507 | -0.112867 | -0.027383 | -0.132331 | 0.06575 | 0.124378 | -0.060346 | -0.162361 | -0.099462 | -0.137824 | -0.190834 | -0.333662 | -0.125399 | -0.078711 | -0.11037 | -0.056693 | -0.0304 | | 18.23611111 | -0.143075 | -0.096466 |
| 18.152778 | -0.527074 | -0.271297 | -0.311474 | -0.128016 | -0.031038 | -0.369802 | -0.294115 | -0.252965 | -0.124138 | -0.011283 | -0.125474 | -0.047323 | -0.202892 | -0.06926 | -0.16784 | -0.127034 | -0.159844 | -0.198284 | -0.330476 | -0.133207 | -0.091676 | -0.106038 | -0.048411 | -0.122923 | | 18.25 | -0.160497 | -0.226049 |
| 18.166667 | -0.606334 | -0.19185 | -0.262192 | -0.098301 | 0.009327 | -0.306864 | -0.267264 | -0.132397 | -0.082473 | 0.023928 | -0.057699 | -0.117419 | -0.388572 | -0.096642 | -0.192332 | -0.125224 | -0.146558 | -0.17315 | -0.282373 | -0.113471 | -0.102382 | -0.122919 | -0.06808 | -0.134173 | | 18.26388889 | -0.16437 | -0.14036 |
| 18.180556 | -0.718909 | -0.168841 | -0.271823 | -0.101307 | 0.022405 | -0.295199 | -0.279972 | -0.076714 | -0.038473 | 0.016234 | -0.004645 | -0.0667 | -0.322195 | -0.087655 | -0.182845 | -0.12409 | -0.132497 | -0.151488 | -0.230637 | -0.08063 | -0.088365 | -0.113875 | -0.060127 | -0.057486 | | 18.27777778 | -0.164828 | -0.045195 |
| 18.194444 | -0.83969 | -0.20974 | -0.379082 | -0.167575 | -0.03033 | -0.383059 | -0.414298 | -0.115294 | -0.038747 | -0.029655 | 0.003748 | -0.05645 | -0.347666 | -0.050218 | -0.142043 | -0.141956 | -0.136331 | -0.146837 | -0.222115 | -0.0596 | -0.065301 | -0.082407 | -0.029956 | -0.000695 | | 18.29166667 | -0.121251 | -0.037484 |
| 18.208333 | -0.804409 | -0.218286 | -0.37355 | -0.184731 | -0.064032 | -0.466149 | -0.55503 | -0.154227 | -0.062572 | -0.054438 | -0.023572 | -0.069798 | -0.404941 | -0.019919 | -0.102809 | -0.144562 | -0.118233 | -0.134358 | -0.207358 | -0.041007 | -0.042311 | -0.057106 | -0.006824 | -0.004648 | | 18.30555556 | -0.035904 | -0.064816 |
| 18.222222 | -0.709871 | -0.196441 | -0.297179 | -0.142803 | -0.050881 | -0.48483 | -0.584812 | -0.178083 | -0.097326 | -0.051635 | -0.07773 | -0.063952 | -0.381321 | -0.023056 | -0.096921 | -0.152802 | -0.110647 | -0.125257 | -0.191312 | -0.045087 | -0.038729 | -0.054856 | -0.003611 | -0.05993 | | 18.31944444 | -0.006961 | -0.126301 |
| 18.236111 | -0.629263 | -0.169714 | -0.239654 | -0.099444 | -0.040842 | -0.430766 | -0.451244 | -0.179434 | -0.132318 | -0.055649 | -0.116489 | -0.05801 | -0.278218 | -0.069999 | -0.130189 | -0.180045 | -0.148736 | -0.13472 | -0.199607 | -0.073882 | -0.073816 | -0.092966 | -0.042267 | -0.141649 | | 18.33333333 | -0.078423 | -0.214082 |
| 18.25 | -0.543793 | -0.122866 | -0.14528 | -0.039773 | -0.012724 | -0.310872 | -0.245566 | -0.130459 | -0.119355 | -0.049744 | -0.082972 | -0.009781 | -0.120391 | -0.086517 | -0.127613 | -0.169517 | -0.153857 | -0.119777 | -0.187242 | -0.071301 | -0.085034 | -0.108387 | -0.074051 | -0.130739 | | 18.34722222 | -0.140056 | -0.185336 |
| 18.263889 | -0.427637 | -0.065355 | -0.094232 | 0.009994 | 0.018546 | -0.19831 | -0.091183 | -0.088287 | -0.071776 | -0.023582 | -0.032006 | 0.033946 | -0.055893 | -0.041168 | -0.075659 | -0.130633 | -0.102416 | -0.082219 | -0.132123 | -0.044585 | -0.047661 | -0.073563 | -0.052351 | -0.042551 | | 18.36111111 | -0.082072 | -0.015583 |
| 18.277778 | -0.348353 | -0.02761 | -0.123726 | 0.02324 | 0.011784 | -0.133284 | -0.021702 | -0.092022 | -0.044578 | -0.014583 | -0.040187 | -0.007348 | -0.152193 | -0.012364 | -0.051735 | -0.111876 | -0.066444 | -0.053955 | -0.070333 | -0.032749 | -0.02022 | -0.058881 | -0.035226 | -0.013802 | | 18.375 | -0.000731 | 0.120726 |
| 18.291667 | -0.314684 | -0.011638 | -0.201802 | -0.020809 | -0.018256 | -0.119628 | -0.063179 | -0.107769 | -0.051042 | -0.034159 | -0.080765 | -0.07671 | -0.242015 | -0.017623 | -0.055515 | -0.109152 | -0.058434 | -0.047304 | -0.03858 | -0.034701 | -0.018772 | -0.062526 | -0.040347 | -0.020841 | | 18.38888889 | -0.005837 | 0.17907 |
| 18.305556 | -0.285366 | -0.021343 | -0.230983 | -0.095699 | -0.057601 | -0.137436 | -0.154566 | -0.111093 | -0.0722 | -0.058259 | -0.104477 | -0.094604 | -0.229178 | -0.037761 | -0.061938 | -0.109295 | -0.064802 | -0.058135 | -0.055492 | -0.048627 | -0.033544 | -0.058169 | -0.044726 | -0.044377 | | 18.40277778 | -0.055133 | 0.183937 |
| 18.319444 | -0.221727 | -0.012422 | -0.130951 | -0.079449 | -0.04077 | -0.095818 | -0.155589 | -0.077194 | -0.055194 | -0.033405 | -0.073461 | -0.025374 | -0.116035 | -0.044371 | -0.039366 | -0.093375 | -0.059239 | -0.052806 | -0.055919 | -0.049766 | -0.040429 | -0.045498 | -0.026868 | -0.096608 | | 18.41666667 | -0.09778 | 0.106592 |
| 18.333333 | -0.156954 | 0.025253 | -0.005105 | 0.019739 | 0.023496 | -0.010538 | -0.058107 | -0.00645 | 0.004889 | 0.021427 | -0.034619 | 0.067695 | 0.043752 | -0.030349 | 0.010784 | -0.059422 | -0.034043 | -0.018173 | -0.008584 | -0.033113 | -0.027474 | -0.0312 | -0.003825 | -0.100241 | | 18.43055556 | -0.118544 | 0.010911 |
| 18.347222 | -0.179599 | 0.044612 | 0.008606 | 0.07267 | 0.05246 | 0.024151 | 0.013933 | 0.03991 | 0.042954 | 0.037134 | -0.043342 | 0.060987 | 0.047109 | -0.021218 | 0.041637 | -0.039892 | -0.02116 | 0.018241 | 0.038443 | -0.026623 | -0.015377 | -0.029414 | -0.004856 | -0.054691 | | 18.44444444 | -0.127871 | -0.049199 |
| 18.361111 | -0.288373 | 0.050335 | -0.026827 | 0.054862 | 0.015376 | 0.008125 | 0.026522 | 0.020519 | 0.022117 | 0.011754 | -0.07462 | -0.034749 | -0.089181 | -0.026276 | 0.042067 | -0.035218 | -0.016786 | 0.034543 | 0.049084 | -0.031398 | -0.017919 | -0.042297 | -0.01592 | -0.046041 | | 18.45833333 | -0.185379 | -0.207719 |
| 18.375 | -0.47195 | -0.034333 | -0.103311 | -0.000135 | -0.056754 | -0.052865 | -0.030686 | -0.10487 | -0.082777 | -0.065459 | -0.143962 | -0.175671 | -0.141196 | -0.051909 | 0.042562 | -0.028755 | -0.021321 | 0.032133 | 0.00676 | -0.045559 | -0.05558 | -0.044245 | -0.018633 | -0.163656 | | 18.47222222 | -0.402793 | -0.604676 |
| 18.388889 | -0.650609 | -0.452492 | -0.342008 | 0.045259 | 0.018706 | -0.020126 | -0.102527 | -0.385178 | -0.213439 | -0.132913 | -0.040639 | -0.164207 | 0.503103 | -0.033656 | 0.158668 | -0.01674 | -0.102913 | -0.066405 | -0.160087 | -0.083572 | -0.190839 | 0.022481 | 0.085315 | -0.76589 | | 18.48611111 | -1.029221 | -2.392149 |
| 18.402778 | -0.715835 | -1.186389 | -0.592513 | 0.217342 | 0.289547 | 0.165962 | 0.11694 | -0.702 | -0.2166 | 0.038793 | 0.540786 | 0.748324 | 3.861349 | 0.100838 | 0.33319 | -0.049542 | -0.318284 | -0.253054 | -0.404946 | -0.122999 | -0.369408 | 0.132919 | 0.678798 | -1.895939 | | 18.5 | -2.093219 | -7.271394 |
| 18.416667 | -0.591133 | -1.906313 | -0.519321 | 0.246085 | 0.078228 | 0.382077 | 0.410793 | -0.46564 | -0.067951 | 0.237516 | 1.068192 | 2.478238 | 8.040137 | 0.051069 | 0.055501 | -0.138278 | -0.447676 | -0.264104 | -0.449095 | -0.114074 | -0.350456 | 0.189736 | 1.437728 | -2.559835 | | 18.51388889 | -3.767556 | -11.909655 |
| 18.430556 | -0.353326 | -0.222515 | -0.041732 | -0.097159 | -0.332822 | -0.078459 | 0.269561 | -0.095076 | 0.123496 | 0.023882 | 0.004759 | 1.955065 | 3.925487 | -0.066764 | -0.332791 | -0.130058 | -0.199526 | 0.020988 | -0.095732 | -0.070279 | -0.080252 | -0.019918 | 0.771348 | -0.714351 | | 18.52777778 | -1.86802 | -3.016824 |
| 18.444444 | -0.701177 | 1.051025 | -0.423216 | -1.013847 | -0.883143 | -1.320723 | -0.735998 | -0.307346 | -0.48275 | -0.785907 | -1.612098 | -2.34696 | -6.460823 | -0.191697 | -0.228881 | -0.277441 | 0.18593 | 0.097503 | 0.129236 | -0.173724 | 0.170068 | -0.284881 | -0.894781 | 1.847029 | | 18.54166667 | 2.750943 | 8.862467 |
| 18.458333 | -1.631878 | -0.374124 | -1.985075 | -2.086069 | -1.640616 | -2.675462 | -2.114488 | -1.399105 | -1.575886 | -1.654384 | -2.440937 | -4.202063 | -7.902136 | -0.300553 | -0.115744 | -0.572926 | 0.158313 | -0.282324 | -0.15648 | -0.358133 | 0.123443 | -0.27216 | -1.230425 | 2.0803 | | 18.55555556 | 2.858122 | 6.381064 |
| 18.472222 | -1.935354 | -1.564446 | -2.68372 | -2.251749 | -1.634879 | -3.006813 | -2.201401 | -2.322749 | -1.982315 | -1.850548 | -1.746542 | -2.379438 | -4.905797 | -0.280379 | -0.280762 | -0.79296 | -0.086315 | -0.643067 | -0.573439 | -0.336166 | -0.111112 | -0.173103 | -0.391413 | 0.655085 | | 18.56944444 | 1.003003 | 0.165567 |
| 18.486111 | -1.16697 | -1.357674 | -1.80275 | -1.55939 | -1.164384 | -2.384289 | -1.850802 | -1.789229 | -1.365526 | -1.499252 | -1.051145 | -0.879175 | -2.09439 | -0.319343 | -0.427771 | -0.618174 | -0.136334 | -0.501146 | -0.763343 | -0.108735 | -0.202375 | -0.178519 | -0.060909 | -0.077779 | | 18.58333333 | 0.12875 | -1.266633 |
| 18.5 | -0.207845 | -0.621723 | -0.856007 | -0.884556 | -0.46401 | -1.711034 | -1.434301 | -0.637952 | -0.513088 | -0.78104 | -0.417927 | -0.213928 | -0.844916 | -0.282966 | -0.310914 | -0.33361 | -0.068611 | -0.336915 | -0.771225 | 0.020163 | -0.106432 | -0.134896 | -0.037705 | -0.005852 | | 18.59722222 | 0.129212 | -0.39405 |
| 18.513889 | 0.539055 | -0.18548 | -0.417651 | -0.437726 | -0.082053 | -1.079541 | -1.045138 | -0.047588 | -0.07969 | -0.267346 | -0.089144 | 0.126279 | -0.326037 | -0.221303 | -0.127815 | -0.193367 | -0.063675 | -0.257214 | -0.610987 | -0.013572 | -0.035881 | -0.082057 | -0.019916 | 0.030659 | | 18.61111111 | 0.142903 | 0.090378 |
| 18.527778 | 1.301917 | -0.00565 | -0.184214 | -0.175671 | 0.073047 | -0.552267 | -0.569327 | -0.046327 | -0.033312 | -0.099076 | 0.019148 | 0.211818 | -0.101955 | -0.189449 | -0.035266 | -0.135892 | -0.076799 | -0.169581 | -0.373136 | -0.08417 | -0.038279 | -0.077594 | -0.037933 | -0.025354 | | 18.625 | 0.10924 | 0.262236 |
| 18.541667 | 1.406335 | 0.113313 | 0.011496 | -0.025565 | 0.134695 | -0.239174 | -0.356935 | 0.045142 | -0.027423 | -0.061862 | 0.07818 | 0.213384 | -0.009435 | -0.121887 | 0.016792 | -0.043308 | -0.054767 | -0.114853 | -0.188734 | -0.094438 | -0.024953 | -0.053713 | -0.036228 | -0.015336 | | 18.63888889 | 0.088364 | 0.421193 |
| 18.555556 | 1.121237 | 0.201531 | 0.135589 | 0.048554 | 0.191315 | -0.198023 | -0.450031 | 0.186384 | 0.079084 | 0.006125 | 0.148981 | 0.240805 | 0.042234 | -0.052783 | 0.059051 | 0.074505 | -0.035714 | -0.098576 | -0.117945 | -0.071389 | -0.009776 | -0.031992 | -0.023667 | 0.026939 | | 18.65277778 | 0.06215 | 0.586478 |
| 18.569444 | 0.837965 | 0.213876 | 0.127464 | 0.042023 | 0.225118 | -0.293555 | -0.583413 | 0.258681 | 0.253034 | 0.134737 | 0.22532 | 0.320463 | 0.069934 | -0.044676 | 0.070588 | 0.215222 | -0.0518 | -0.083495 | -0.098897 | -0.045669 | -0.008634 | -0.044259 | -0.038217 | 0.058661 | | 18.66666667 | 0.002931 | 0.658139 |
| 18.583333 | 0.622049 | 0.189176 | 0.079713 | 0.014176 | 0.253036 | -0.276834 | -0.533119 | 0.250421 | 0.440653 | 0.28199 | 0.276485 | 0.412849 | 0.112016 | -0.048981 | 0.069081 | 0.409448 | -0.04471 | -0.041899 | -0.043829 | -0.008268 | 0.022007 | -0.044887 | -0.031483 | 0.092403 | | 18.68055556 | -0.033653 | 0.647881 |
| 18.597222 | 0.528425 | 0.23115 | 0.065838 | 0.016254 | 0.274843 | -0.06541 | -0.10977 | 0.208422 | 0.614122 | 0.430874 | 0.285469 | 0.460967 | 0.238181 | -0.010022 | 0.07253 | 0.520766 | -0.005911 | -0.014142 | 0.033579 | 0.035135 | 0.102651 | 0.00359 | 0.02741 | 0.114843 | | 18.69444444 | -0.012165 | 0.688558 |
| 18.611111 | 0.517047 | 0.359971 | 0.167407 | 0.111056 | 0.310755 | 0.300152 | 0.552263 | 0.248213 | 0.797772 | 0.532129 | 0.306057 | 0.470904 | 0.452772 | -0.01342 | -0.014604 | 0.538682 | -0.006892 | -0.057399 | -0.001362 | 0.062944 | 0.135017 | 0.021771 | 0.049105 | 0.077852 | | 18.70833333 | 0.001214 | 0.818061 |
| 18.625 | 0.46763 | 0.48338 | 0.367568 | 0.290404 | 0.423315 | 0.726409 | 1.475417 | 0.44101 | 0.98193 | 0.629264 | 0.402147 | 0.509157 | 0.660617 | -0.056493 | -0.147252 | 0.46869 | -0.048431 | -0.182069 | -0.231308 | 0.087565 | 0.053309 | 0.011107 | 0.043907 | 0.027156 | | 18.72222222 | 0.000344 | 0.979269 |
| 18.638889 | 0.324154 | 0.526557 | 0.457355 | 0.420961 | 0.52014 | 1.178838 | 3.026955 | 0.56432 | 1.005913 | 0.488554 | 0.561334 | 0.72684 | 0.556179 | -0.065179 | -0.186159 | 0.389063 | -0.048837 | -0.329532 | -0.556175 | 0.090812 | -0.045341 | 0.029049 | 0.053588 | 0.028709 | | 18.73611111 | 0.021082 | 1.020883 |
| 18.652778 | 0.153407 | 0.471891 | 0.406847 | 0.400203 | 0.481877 | 1.13341 | 2.58502 | 0.506276 | 0.828308 | 0.548081 | 0.447645 | 0.512474 | 0.543216 | -0.090577 | -0.23631 | 0.334309 | 0.040337 | -0.458912 | -0.917527 | 0.085007 | -0.142256 | 0.000147 | -0.00163 | 0.05674 | | 18.75 | 0.015962 | 0.802715 |
| 18.666667 | 0.024747 | 0.363939 | 0.361587 | 0.337168 | 0.383998 | 0.777119 | 1.467443 | 0.396223 | 0.562205 | 0.292736 | 0.347308 | 0.395455 | 0.394376 | -0.151285 | -0.338468 | 0.332161 | 0.209077 | -0.533105 | -1.173679 | 0.086033 | -0.144112 | -0.076335 | -0.079476 | 0.088927 | | 18.76388889 | -0.03391 | 0.420537 |
| 18.680556 | 0.160958 | 0.281828 | 0.309782 | 0.292678 | 0.299594 | 0.40202 | 0.664685 | 0.249301 | 0.256706 | -0.033838 | 0.251787 | 0.272403 | 0.251859 | -0.187362 | -0.366129 | 0.332503 | 0.299741 | -0.562753 | -1.189066 | 0.039708 | -0.102539 | -0.129072 | -0.084145 | 0.13026 | | 18.77777778 | -0.052554 | 0.147758 |
| 18.694444 | 0.419251 | 0.204924 | 0.178141 | 0.198304 | 0.206635 | 0.039446 | -0.002169 | 0.037263 | -0.044586 | -0.285383 | 0.158127 | 0.178561 | 0.236491 | -0.262496 | -0.29516 | 0.186131 | 0.160858 | -0.465741 | -0.827029 | -0.039745 | -0.069136 | -0.179123 | -0.058284 | 0.135741 | | 18.79166667 | -0.039104 | -0.011066 |
| 18.708333 | 0.604233 | 0.118754 | 0.066833 | 0.094838 | 0.139786 | -0.216353 | -0.527619 | -0.115853 | -0.122782 | -0.294579 | 0.114047 | 0.154885 | 0.175546 | -0.316384 | -0.251266 | 0.021319 | -0.007235 | -0.311451 | -0.478315 | -0.058626 | -0.052143 | -0.29811 | -0.083738 | 0.05852 | | 18.80555556 | -0.065218 | -0.148238 |
| 18.722222 | 0.658634 | 0.048368 | 0.116016 | 0.064455 | 0.135311 | -0.344193 | -1.132121 | -0.134917 | -0.007701 | -0.120548 | 0.11903 | 0.184349 | 0.172746 | -0.29014 | -0.23078 | -0.079966 | -0.168147 | -0.238046 | -0.238327 | -0.024745 | -0.043672 | -0.469862 | -0.140487 | -0.099413 | | 18.81944444 | -0.087874 | -0.130509 |
| 18.736111 | 0.682406 | -0.006892 | 0.203181 | 0.086975 | 0.139413 | -0.370354 | -1.812941 | -0.096253 | 0.167702 | 0.080819 | 0.08836 | 0.201543 | 0.23015 | -0.267921 | -0.190937 | -0.143737 | -0.28856 | -0.27839 | -0.151588 | -0.010224 | -0.029398 | -0.53373 | -0.160739 | -0.212329 | | 18.83333333 | -0.056013 | 0.011823 |
| 18.75 | 0.829214 | -0.078724 | 0.148586 | 0.091216 | 0.098225 | -0.197563 | -1.205526 | -0.115163 | 0.260435 | 0.233556 | 0.010004 | 0.236333 | 0.353382 | -0.230734 | -0.132155 | -0.118144 | -0.280149 | -0.299101 | -0.06439 | 0.017172 | 0.056545 | -0.454463 | -0.140974 | -0.146349 | | 18.84722222 | -0.028452 | 0.005231 |
| 18.763889 | 0.960152 | -0.154046 | 0.00339 | 0.059773 | 0.01709 | -0.118834 | -0.448587 | -0.137275 | 0.235605 | 0.308352 | -0.072788 | 0.275252 | 0.40866 | -0.178064 | -0.06813 | -0.025018 | -0.167328 | -0.255347 | 0.12653 | 0.06462 | 0.138945 | -0.324196 | -0.104092 | 0.016266 | | 18.86111111 | -0.026558 | -0.216761 |
| 18.777778 | 0.713386 | -0.169846 | -0.023227 | 0.033093 | -0.060311 | -0.0944 | -0.1327 | -0.045755 | 0.203151 | 0.339693 | -0.126366 | 0.256471 | 0.242582 | -0.116357 | -0.046056 | 0.086149 | -0.059055 | -0.214606 | 0.358121 | 0.07851 | 0.128306 | -0.229842 | -0.090587 | 0.090855 | | 18.875 | -0.004741 | -0.398142 |
| 18.791667 | 0.355304 | -0.120763 | 0.110455 | 0.067457 | -0.084117 | 0.069768 | 0.273075 | 0.099564 | 0.170495 | 0.269897 | -0.106272 | 0.120376 | 0.035959 | -0.096697 | -0.04883 | 0.168762 | 0.004659 | -0.095751 | 0.463963 | 0.071946 | 0.080889 | -0.190776 | -0.103288 | 0.088458 | | 18.88888889 | 0.029858 | -0.362656 |
| 18.805556 | 0.332977 | -0.05621 | 0.2677 | 0.172447 | -0.060666 | 0.377724 | 1.158887 | 0.244702 | 0.082656 | 0.095351 | -0.054478 | 0.002157 | 0.059719 | -0.061378 | -0.000949 | 0.226047 | 0.062554 | 0.065142 | 0.484276 | 0.067034 | 0.058394 | -0.117244 | -0.099645 | 0.093614 | | 18.90277778 | 0.031054 | -0.207667 |
| 18.819444 | 0.689652 | 0.005303 | 0.39785 | 0.300326 | -0.037378 | 0.647289 | 1.912511 | 0.328294 | -0.001928 | -0.036241 | -0.003535 | -0.113428 | 0.266717 | 0.007243 | 0.097471 | 0.287759 | 0.14614 | 0.17964 | 0.573384 | 0.073343 | 0.067887 | 0.000863 | -0.061153 | 0.132444 | | 18.91666667 | 0.0177 | -0.061554 |
| 18.833333 | 0.649263 | 0.092901 | 0.385037 | 0.409401 | -0.040854 | 0.686182 | 1.518273 | 0.329026 | -0.086787 | -0.109199 | 0.029725 | -0.17463 | 0.293365 | 0.088031 | 0.211204 | 0.321137 | 0.21093 | 0.215736 | 0.653284 | 0.080463 | 0.089004 | 0.117325 | -0.018154 | 0.16641 | | 18.93055556 | 0.018066 | -0.007477 |
| 18.847222 | -0.067272 | 0.194412 | 0.277485 | 0.350239 | -0.057113 | 0.690397 | 1.120609 | 0.270302 | -0.181191 | -0.155634 | 0.004336 | -0.178752 | -0.010021 | 0.160717 | 0.273859 | 0.308175 | 0.220356 | 0.232258 | 0.792221 | 0.05783 | 0.080111 | 0.201315 | -0.010351 | 0.134118 | | 18.94444444 | -0.002126 | -0.067422 |
| 18.861111 | -0.456957 | 0.313082 | 0.164122 | 0.242049 | -0.050597 | 0.725993 | 0.852205 | 0.297295 | -0.194001 | -0.147319 | -0.053966 | -0.198766 | -0.233615 | 0.226203 | 0.253423 | 0.260764 | 0.19711 | 0.328902 | 0.908677 | 0.013324 | 0.02626 | 0.252672 | -0.045404 | 0.04156 | | 18.95833333 | -0.029607 | -0.130687 |
| 18.875 | 0.084971 | 0.459458 | 0.167646 | 0.195398 | 0.001251 | 0.788222 | 1.174912 | 0.537954 | -0.098059 | -0.121584 | -0.055294 | -0.177929 | 0.065875 | 0.183931 | 0.231217 | 0.176174 | 0.17174 | 0.43737 | 1.018441 | 0.028606 | -0.026262 | 0.297998 | -0.0654 | -0.021328 | | 18.97222222 | 0.027279 | -0.051505 |
| 18.888889 | 0.998501 | 0.506449 | 0.243259 | 0.223248 | 0.096639 | 0.711781 | 0.664632 | 0.675348 | 0.014212 | -0.139887 | -0.017767 | -0.15129 | 0.287774 | 0.425252 | 0.280749 | 0.209447 | 0.152958 | 0.469557 | 1.014509 | 0.014895 | -0.033959 | 0.228005 | -0.04828 | -0.012484 | | 18.98611111 | -0.003062 | 0.130599 |
| 18.902778 | 1.576354 | 0.451287 | 0.327487 | 0.237617 | 0.144921 | 0.617951 | 0.467052 | 0.582999 | 0.066624 | -0.157084 | 0.006568 | -0.170874 | 0.487291 | 0.464905 | 0.308605 | 0.170848 | 0.102996 | 0.410035 | 0.84463 | 0.006269 | -0.036926 | 0.136057 | -0.044519 | 0.014333 | | 19 | 0.00287 | 0.269533 |
| 18.916667 | 0.802698 | 0.341105 | 0.315033 | 0.201656 | 0.123075 | 0.691049 | 0.303724 | 0.42938 | 0.051059 | 0.09874 | 0.038684 | -0.096767 | 0.921759 | 0.499565 | 0.267324 | 0.116476 | 0.039705 | 0.295651 | 0.660604 | -0.019462 | -0.063955 | 0.077613 | -0.008531 | -0.014812 | | 19.01388889 | -0.06067 | 0.083328 |
| 18.930556 | -0.286168 | -0.117705 | 0.098836 | 0.168664 | -0.045522 | 0.6479 | 0.381805 | 0.184776 | 0.040957 | 0.236311 | 0.196015 | 0.730194 | 3.768116 | 0.464917 | 0.205035 | 0.019925 | -0.079925 | 0.152342 | 0.399967 | -0.052651 | -0.153922 | 0.166128 | 0.324585 | -0.392901 | | 19.02777778 | -0.844132 | -2.033511 |
| 18.944444 | -1.084993 | -0.770179 | -0.112502 | 0.155742 | -0.279675 | 0.535952 | 0.576306 | -0.004112 | -0.015826 | 0.278766 | 0.54357 | 1.675033 | 6.272372 | 0.467525 | 0.167654 | -0.048148 | -0.25065 | 0.003807 | 0.113452 | -0.072663 | -0.300346 | 0.370582 | 0.928594 | -1.161143 | | 19.04166667 | -2.364306 | -6.89362 |
| 18.958333 | -1.126307 | -0.220265 | 0.124442 | 0.040018 | -0.167376 | 0.184367 | -0.19205 | 0.071002 | -0.305288 | -0.215189 | 0.239902 | -0.359237 | -0.183732 | 0.288536 | 0.200985 | 0.027756 | -0.243659 | -0.038119 | -0.020125 | -0.025851 | -0.268428 | 0.234969 | 0.482627 | -0.826108 | | 19.05555556 | -1.610217 | -7.715458 |
| 18.972222 | -0.238872 | 1.429958 | 0.393953 | -0.115661 | 0.329181 | -0.023416 | -1.041913 | 0.064921 | -0.521199 | -0.354242 | -0.718231 | -2.889315 | -11.057575 | -0.172788 | 0.294894 | 0.093698 | 0.015823 | 0.053942 | 0.055136 | 0.030373 | 0.020726 | -0.325399 | -1.082884 | 0.902819 | | 19.06944444 | 1.826071 | 7.318583 |
| 18.986111 | -0.145079 | 1.136295 | 0.352298 | -0.226315 | 0.044096 | -0.110381 | -0.839815 | -0.13666 | -0.433908 | -0.209572 | -0.789084 | -1.68723 | -5.391152 | -0.420042 | 0.121294 | -0.132055 | 0.091095 | 0.03751 | 0.095533 | 0.055996 | 0.196997 | -0.569685 | -1.548221 | 1.474613 | | 19.08333333 | 2.702155 | 6.865038 |
| 19 | -0.352337 | 0.301989 | 0.094189 | -0.14032 | -0.246292 | -0.086784 | -0.406932 | -0.513479 | -0.343991 | 0.004877 | -0.370426 | -0.007274 | 2.105581 | -0.31571 | -0.092274 | -0.472701 | -0.069104 | -0.037424 | -0.05089 | 0.030215 | 0.106598 | -0.341394 | -0.729265 | 0.559518 | | 19.09722222 | 1.04342 | 2.537986 |
| 19.013889 | -0.352328 | 0.106851 | -0.029636 | -0.176983 | -0.460125 | -0.223818 | -0.554563 | -0.663825 | -0.379258 | 0.050865 | -0.350816 | -0.081389 | 2.202304 | -0.208944 | -0.166564 | -0.519199 | -0.135888 | -0.007069 | -0.113097 | -0.012226 | 0.004755 | -0.190882 | -0.260016 | 0.007039 | | 19.11111111 | 0.094233 | 0.432359 |
| 19.027778 | -0.315412 | 0.056398 | -0.100636 | -0.250817 | -0.443323 | -0.451013 | -0.918946 | -0.677291 | -0.394788 | -0.011325 | -0.587506 | -0.515674 | 0.291158 | -0.199291 | -0.274608 | -0.418831 | -0.129311 | 0.058405 | -0.070307 | -0.060052 | -0.05127 | -0.151338 | -0.206893 | -0.072081 | | 19.125 | -0.045058 | 0.503715 |
| 19.041667 | -0.312116 | -0.031059 | -0.1475 | -0.303022 | -0.446124 | -0.608906 | -1.114309 | -0.619688 | -0.318496 | -0.029523 | -0.499089 | -0.445788 | -0.307355 | -0.206723 | -0.417567 | -0.368459 | -0.180311 | 0.00356 | -0.066609 | -0.15792 | -0.133339 | -0.153531 | -0.178455 | -0.164519 | | 19.13888889 | -0.0466 | 0.590848 |
| 19.055556 | -0.318842 | -0.096531 | -0.16244 | -0.311473 | -0.38014 | -0.640034 | -1.130678 | -0.537664 | -0.249909 | -0.066254 | -0.247042 | -0.524231 | -1.073071 | -0.182875 | -0.490163 | -0.311089 | -0.223022 | -0.060103 | -0.079678 | -0.213663 | -0.187386 | -0.158298 | -0.140575 | -0.255263 | | 19.15277778 | -0.071082 | -0.147644 |
| 19.069444 | -0.359327 | -0.231595 | -0.204903 | -0.313406 | -0.260751 | -0.672375 | -1.103975 | -0.529361 | -0.261879 | -0.134893 | -0.600254 | -0.650383 | -1.643232 | -0.132897 | -0.524097 | -0.215162 | -0.17437 | -0.049686 | -0.054846 | -0.17552 | -0.175657 | -0.127852 | -0.091527 | -0.208911 | | 19.16666667 | -0.101019 | -0.70625 |
| 19.083333 | -0.410555 | -0.446475 | -0.279825 | -0.279025 | -0.211906 | -0.60982 | -0.913934 | -0.528623 | -0.261566 | -0.132768 | -0.423991 | -0.191581 | -0.213314 | -0.099671 | -0.407816 | -0.115495 | -0.08463 | -0.02935 | -0.026114 | -0.111558 | -0.127877 | -0.084328 | -0.065471 | -0.081091 | | 19.18055556 | -0.107032 | -0.092023 |
| 19.097222 | -0.379165 | -0.480002 | -0.273188 | -0.200651 | -0.147126 | -0.474579 | -0.620961 | -0.397376 | -0.179451 | -0.086051 | -0.193852 | -0.218915 | 1.218997 | -0.050913 | -0.272331 | -0.012838 | -0.011961 | -0.037586 | -0.030817 | -0.077636 | -0.085299 | -0.054539 | -0.069639 | -0.017527 | | 19.19444444 | -0.033665 | 0.505208 |
| 19.111111 | -0.313361 | -0.394727 | -0.192634 | -0.1536 | -0.092083 | -0.343609 | -0.471418 | -0.320303 | -0.197753 | -0.110242 | -0.255413 | -0.056967 | 0.109142 | -0.101452 | -0.191945 | -0.106375 | -0.056778 | -0.0602 | -0.062545 | -0.077786 | -0.068051 | -0.050839 | -0.079208 | -0.042965 | | 19.20833333 | -0.017631 | -0.026901 |
| 19.125 | -0.316102 | -0.352351 | -0.165175 | -0.163494 | -0.092338 | -0.3304 | -0.413885 | -0.275637 | -0.262468 | -0.182685 | -0.427971 | -0.499196 | -1.181261 | -0.10274 | -0.174298 | -0.128187 | -0.06961 | -0.076318 | -0.090669 | -0.077423 | -0.063041 | -0.067085 | -0.088344 | -0.067304 | | 19.22222222 | -0.079069 | -0.63061 |
| 19.138889 | -0.371887 | -0.375182 | -0.221207 | -0.186393 | -0.130475 | -0.315728 | -0.318415 | -0.339685 | -0.299065 | -0.190588 | -0.412599 | -0.233045 | -0.211794 | -0.096928 | -0.156832 | -0.117328 | -0.051047 | -0.068024 | -0.087703 | -0.069719 | -0.051643 | -0.07906 | -0.098407 | -0.042829 | | 19.23611111 | -0.041719 | -0.409815 |
| 19.152778 | -0.388527 | -0.379516 | -0.256992 | -0.192387 | -0.14567 | -0.281679 | -0.22271 | -0.365844 | -0.27259 | -0.157723 | -0.303922 | 0.144016 | 0.903257 | -0.072282 | -0.044737 | -0.090643 | -0.028923 | -0.049153 | -0.063904 | -0.048572 | -0.029558 | -0.072452 | -0.090696 | -0.012512 | | 19.25 | -0.00049 | -0.135101 |
| 19.166667 | -0.372552 | -0.378855 | -0.270164 | -0.199762 | -0.131837 | -0.284411 | -0.19947 | -0.324819 | -0.24475 | -0.167132 | -0.285494 | 0.037891 | 0.46735 | -0.049287 | 0.244235 | -0.059393 | -0.035805 | -0.047969 | -0.05184 | -0.047939 | -0.015852 | -0.060272 | -0.073701 | -0.003461 | | 19.26388889 | -0.000444 | -0.102718 |
| 19.180556 | -0.325389 | -0.327692 | -0.263701 | -0.178134 | -0.084471 | -0.270423 | -0.175456 | -0.26502 | -0.212199 | -0.149124 | -0.282922 | -0.096569 | -0.121243 | -0.050372 | 0.487192 | -0.025077 | -0.053779 | -0.060679 | -0.053361 | -0.062788 | -0.012075 | -0.052261 | -0.065961 | -0.00347 | | 19.27777778 | -0.009474 | -0.039601 |
| 19.194444 | -0.253587 | -0.277685 | -0.200066 | -0.111629 | -0.009233 | -0.204908 | -0.103096 | -0.190449 | -0.152333 | -0.098955 | -0.238636 | -0.056364 | -0.154943 | -0.071871 | 0.374021 | -0.035407 | -0.069672 | -0.059228 | -0.051035 | -0.063977 | -0.022285 | -0.052033 | -0.061307 | -0.038378 | | 19.29166667 | -0.039828 | -0.037488 |
| 19.208333 | -0.27212 | -0.306278 | -0.204012 | -0.109444 | -0.011055 | -0.21553 | -0.104966 | -0.180255 | -0.153669 | -0.114928 | -0.223997 | -0.06064 | -0.119155 | -0.073987 | 0.132288 | -0.055772 | -0.080784 | -0.043964 | -0.045906 | -0.050521 | -0.025638 | -0.03959 | -0.044173 | -0.055216 | | 19.30555556 | -0.053566 | -0.164435 |
| 19.222222 | -0.384516 | -0.424673 | -0.328977 | -0.202826 | -0.097947 | -0.333357 | -0.211719 | -0.263546 | -0.240353 | -0.20726 | -0.287242 | -0.162243 | -0.318408 | -0.048428 | 0.018588 | -0.073098 | -0.073721 | -0.040247 | -0.043774 | -0.039792 | -0.009838 | -0.015807 | -0.022189 | -0.040303 | | 19.31944444 | -0.033281 | -0.338006 |
| 19.236111 | -0.455277 | -0.496525 | -0.382176 | -0.277955 | -0.151177 | -0.43257 | -0.334933 | -0.313809 | -0.308375 | -0.311495 | -0.394912 | -0.384248 | -1.072961 | -0.042155 | -0.027727 | -0.084067 | -0.072555 | -0.046732 | -0.048803 | -0.038528 | -0.011945 | -0.016993 | -0.012593 | -0.05247 | | 19.33333333 | -0.015487 | -0.457252 |
| 19.25 | -0.420475 | -0.452428 | -0.318323 | -0.259736 | -0.140809 | -0.426704 | -0.360647 | -0.271003 | -0.287628 | -0.316331 | -0.396631 | -0.433271 | -1.525439 | -0.034127 | -0.029721 | -0.076045 | -0.061143 | -0.030772 | -0.036337 | -0.026292 | -0.007292 | -0.016418 | 0.001723 | -0.059359 | | 19.34722222 | 0.023681 | -0.440724 |
| 19.263889 | -0.309956 | -0.304145 | -0.25182 | -0.15922 | -0.066646 | -0.306399 | -0.199482 | -0.197798 | -0.176436 | -0.162903 | -0.20594 | -0.011123 | -0.34486 | -0.011358 | -0.004438 | -0.039111 | -0.019042 | 0.006725 | -0.000483 | -0.00132 | 0.024596 | 0.00074 | 0.011343 | -0.023396 | | 19.36111111 | 0.040502 | -0.270382 |
| 19.277778 | -0.154154 | -0.083039 | -0.131311 | -0.045561 | 0.040086 | -0.168252 | -0.04649 | -0.080265 | -0.065974 | -0.014573 | -0.071248 | 0.186784 | 0.149141 | -0.002441 | -0.002342 | -0.005309 | 0.004901 | 0.017445 | 0.010131 | 0.009924 | 0.039268 | 0.012278 | 0.011534 | 0.003714 | | 19.375 | 0.011503 | -0.011135 |
| 19.291667 | -0.039316 | 0.064597 | -0.009069 | 0.009996 | 0.092361 | -0.093433 | -0.059644 | 0.020573 | -0.016503 | 0.013318 | -0.095501 | -0.114041 | -0.778317 | -0.006635 | -0.004588 | 0.000201 | -0.008459 | -0.006064 | -0.011208 | 0.000377 | 0.032001 | 0.018574 | 0.011141 | 0.012249 | | 19.38888889 | 0.007196 | 0.209356 |
| 19.305556 | -0.057874 | -0.018966 | -0.048505 | -0.001411 | 0.04616 | -0.027102 | -0.068762 | -0.043754 | -0.006713 | -0.008764 | -0.042428 | 0.057464 | 0.046863 | -0.045636 | -0.038328 | -0.024521 | -0.041147 | -0.039449 | -0.042229 | -0.027747 | -0.003982 | -0.015937 | -0.021257 | -0.007194 | | 19.40277778 | 0.024159 | 0.236409 |
| 19.319444 | -0.127047 | -0.163782 | -0.135939 | -0.035363 | -0.019073 | 0.012142 | -0.040858 | -0.162618 | -0.029138 | -0.028581 | 0.024658 | 0.431124 | 1.375129 | -0.077436 | -0.069401 | -0.053053 | -0.069472 | -0.055318 | -0.058683 | -0.050128 | -0.042668 | -0.058279 | -0.044512 | -0.056181 | | 19.41666667 | 0.048862 | 0.086557 |
| 19.333333 | -0.105735 | -0.134052 | -0.09938 | -0.027698 | 0.002197 | 0.020975 | -0.036027 | -0.149635 | -0.04597 | -0.023304 | 0.005445 | 0.348755 | 0.895137 | -0.050259 | -0.041196 | -0.042633 | -0.053282 | -0.034396 | -0.030787 | -0.032449 | -0.027255 | -0.041002 | -0.021343 | -0.068391 | | 19.43055556 | 0.04771 | -0.060144 |
| 19.347222 | -0.040217 | -0.0156 | -0.011136 | 0.009713 | 0.062151 | 0.041003 | -0.038647 | -0.067665 | -0.019237 | -0.007213 | -0.022655 | 0.156199 | 0.015083 | -0.008275 | 0.006834 | 0.010764 | 0.009347 | 0.004589 | 0.013283 | 0.006086 | 0.012476 | -0.00264 | 0.016102 | -0.009399 | | 19.44444444 | 0.005731 | -0.10612 |
| 19.361111 | -0.00207 | 0.073615 | 0.043344 | 0.018119 | 0.081462 | 0.027588 | -0.082321 | -0.019029 | -0.012652 | -0.031847 | -0.089273 | -0.112735 | -0.770576 | 0.021019 | 0.053029 | 0.059488 | 0.06393 | 0.02237 | 0.032023 | 0.029231 | 0.035199 | 0.015545 | 0.046796 | 0.06186 | | 19.45833333 | -0.076408 | -0.129306 |
| 19.375 | -0.009049 | 0.124945 | 0.0301 | -0.006333 | 0.045275 | -0.028195 | -0.139071 | -0.028083 | -0.019805 | -0.074281 | -0.145611 | -0.322286 | -1.229603 | 0.018145 | 0.070722 | 0.049956 | 0.061421 | 0.006077 | 0.0171 | 0.018691 | 0.028009 | 0.006279 | 0.040818 | 0.063053 | | 19.47222222 | -0.211747 | -0.202895 |
| 19.388889 | -0.074376 | 0.157911 | -0.074195 | -0.037779 | 0.00342 | -0.046384 | -0.097689 | -0.093801 | 0.013135 | -0.052906 | -0.04022 | 0.022989 | -0.043605 | -0.008445 | 0.069309 | 0.017094 | 0.020222 | -0.012125 | 0.000822 | 0.002347 | 0.00417 | -0.007715 | 0.025689 | -0.003622 | | 19.48611111 | -0.291959 | -0.359274 |
| 19.402778 | -0.178792 | 0.141196 | -0.200392 | -0.092237 | -0.017506 | -0.050625 | -0.045297 | -0.177144 | 0.009488 | -0.050335 | 0.0383 | 0.376853 | 1.125066 | -0.003774 | 0.113825 | 0.02371 | 0.009384 | 0.005564 | 0.017275 | 0.008489 | 0.005132 | 0.007296 | 0.043296 | -0.017332 | | 19.5 | -0.217398 | -0.657126 |
| 19.416667 | -0.239804 | 0.077753 | -0.212324 | -0.135862 | -0.008149 | -0.029484 | -0.012664 | -0.210417 | -0.06644 | -0.137996 | -0.037577 | 0.219501 | 0.616913 | 0.01557 | 0.158428 | 0.044487 | 0.01981 | 0.040646 | 0.043247 | 0.023089 | 0.017612 | 0.022343 | 0.058217 | 0.043459 | | 19.51388889 | -0.075669 | -0.898918 |
| 19.430556 | -0.159233 | 0.113769 | -0.113838 | -0.063846 | 0.083085 | 0.091166 | 0.148135 | -0.130963 | -0.040222 | -0.115982 | 0.008663 | 0.052566 | 0.012557 | 0.029204 | 0.146759 | 0.021857 | 0.004614 | 0.035346 | 0.040416 | 0.014422 | 0.018046 | 0.03705 | 0.085225 | 0.033626 | | 19.52777778 | -0.007376 | -0.74033 |
| 19.444444 | -0.137804 | 0.106183 | -0.315555 | -0.120458 | 0.090642 | -0.006179 | 0.182168 | -0.175314 | -0.033661 | -0.067755 | 0.134343 | -0.047033 | -0.247784 | 0.060934 | 0.159133 | -0.164512 | -0.073912 | -0.069761 | -0.012791 | -0.07328 | 0.02213 | 0.121384 | 0.208866 | -0.037257 | | 19.54166667 | -0.06437 | -0.375654 |
| 19.458333 | -0.416276 | -0.219754 | -1.02732 | -0.544172 | -0.195172 | -0.533878 | -0.210494 | -0.558765 | -0.270887 | -0.295909 | 0.134086 | -0.418132 | -0.805929 | 0.054288 | 0.209291 | -0.53912 | -0.216681 | -0.236111 | -0.106233 | -0.255836 | -0.013336 | 0.214597 | 0.364967 | -0.098962 | | 19.55555556 | -0.211603 | -0.297376 |
| 19.472222 | -0.6749 | -0.502748 | -1.230679 | -0.80247 | -0.357838 | -0.766525 | -0.55456 | -0.808231 | -0.41544 | -0.604213 | 0.127849 | -0.770662 | -1.054654 | -0.035561 | 0.195883 | -0.72747 | -0.275706 | -0.275536 | -0.132854 | -0.347357 | -0.134007 | 0.136835 | 0.303919 | -0.190218 | | 19.56944444 | -0.25827 | -0.530399 |
| 19.486111 | -0.524032 | -0.165151 | -0.480351 | -0.576173 | -0.331913 | -0.236042 | -0.398523 | -0.546837 | -0.159698 | -0.557704 | 0.251934 | -0.607758 | -0.607516 | -0.103193 | 0.191332 | -0.50352 | -0.147941 | -0.013872 | -0.021208 | -0.124729 | -0.1823 | 0.00982 | 0.112272 | -0.189043 | | 19.58333333 | -0.134017 | -0.497301 |
| 19.5 | -0.196677 | 0.408903 | 0.11381 | -0.142559 | -0.048159 | 0.227786 | -0.052101 | -0.093608 | 0.127816 | -0.215135 | 0.467598 | -0.080121 | -0.080672 | -0.063054 | 0.231329 | -0.203615 | -0.011436 | 0.160421 | 0.078955 | 0.046043 | -0.092775 | 0.034161 | 0.035057 | -0.035999 | | 19.59722222 | -0.007757 | -0.068521 |
| 19.513889 | 0.011005 | 0.530901 | 0.280123 | 0.124126 | 0.169792 | 0.30578 | 0.14977 | 0.191887 | 0.245262 | 0.051506 | 0.541738 | 0.396092 | 0.095703 | -0.048318 | 0.161626 | -0.05399 | -0.022528 | 0.156036 | 0.055509 | 0.085792 | -0.02695 | 0.065774 | 0.02266 | 0.049701 | | 19.61111111 | 0.022227 | 0.247722 |
| 19.527778 | 0.134152 | 0.412893 | 0.304923 | 0.205807 | 0.244267 | 0.317113 | 0.237369 | 0.251165 | 0.244917 | 0.161094 | 0.533521 | 0.678975 | 0.110651 | -0.054173 | 0.060593 | 0.105569 | -0.097778 | 0.09599 | 0.021445 | 0.055404 | -0.030707 | 0.025314 | 0.012196 | 0.025458 | | 19.625 | 0.010261 | 0.300491 |
| 19.541667 | 0.20918 | 0.327162 | 0.311638 | 0.236412 | 0.255094 | 0.310643 | 0.321559 | 0.214609 | 0.227045 | 0.210571 | 0.48861 | 0.920062 | 0.153032 | 0.034816 | 0.008392 | 0.383628 | -0.128749 | 0.103028 | 0.138548 | 0.027568 | -0.022968 | 0.000377 | 0.025801 | 0.00316 | | 19.63888889 | 0.001845 | 0.238475 |
| 19.555556 | 0.217757 | 0.239617 | 0.298444 | 0.243076 | 0.247622 | 0.259313 | 0.352393 | 0.193761 | 0.205447 | 0.217207 | 0.42466 | 1.069777 | 0.198907 | 0.186327 | -0.014748 | 0.790352 | -0.105697 | 0.220905 | 0.478784 | 0.043451 | 0.007001 | 0.03057 | 0.056711 | 0.012138 | | 19.65277778 | -0.008945 | 0.195747 |
| 19.569444 | 0.182368 | 0.15828 | 0.264213 | 0.237722 | 0.233174 | 0.163418 | 0.261982 | 0.218177 | 0.201287 | 0.196437 | 0.327697 | 1.146194 | 0.179225 | 0.337883 | 0.074078 | 1.18842 | 0.004572 | 0.42352 | 1.009237 | 0.091094 | 0.056578 | 0.100255 | 0.13222 | 0.066214 | | 19.66666667 | -0.017448 | 0.204411 |
| 19.583333 | 0.142095 | 0.068972 | 0.243211 | 0.236731 | 0.223164 | 0.059215 | 0.06044 | 0.299732 | 0.19807 | 0.173494 | 0.239819 | 1.057609 | 0.108587 | 0.495024 | 0.154723 | 1.697386 | 0.193385 | 0.659796 | 1.666425 | 0.093234 | 0.101621 | 0.216978 | 0.207384 | 0.139731 | | 19.68055556 | -0.007369 | 0.258195 |
| 19.597222 | 0.109115 | -0.011284 | 0.232504 | 0.226971 | 0.200918 | 0.002147 | -0.236856 | 0.4382 | 0.179836 | 0.132082 | 0.151378 | 0.889088 | 0.055216 | 0.75748 | 0.253084 | 2.0171 | 0.323121 | 0.900367 | 2.347119 | 0.06068 | 0.087818 | 0.408263 | 0.299332 | 0.155712 | | 19.69444444 | -0.001096 | 0.327535 |
| 19.611111 | 0.118778 | -0.01002 | 0.249702 | 0.217356 | 0.186862 | 0.059695 | -0.472102 | 0.606263 | 0.174794 | 0.089188 | 0.084103 | 0.763196 | 0.034709 | 1.08471 | 0.302471 | 1.98276 | 0.329157 | 0.927412 | 2.500022 | -0.033584 | 0.021199 | 0.681648 | 0.44739 | 0.107432 | | 19.70833333 | 0.013747 | 0.333664 |
| 19.625 | 0.172939 | 0.052855 | 0.29009 | 0.23225 | 0.209648 | 0.262293 | -0.423733 | 0.740565 | 0.201071 | 0.082008 | 0.099657 | 0.640054 | 0.005471 | 1.395054 | 0.268517 | 1.635336 | 0.445269 | 0.665399 | 1.722762 | 0.01792 | 0.049251 | 0.965909 | 0.690011 | 0.13676 | | 19.72222222 | 0.033701 | 0.273696 |
| 19.638889 | 0.249347 | 0.128677 | 0.379435 | 0.313605 | 0.279585 | 0.683236 | 0.073963 | 0.797137 | 0.301361 | 0.114681 | 0.187266 | 0.521847 | -0.008934 | 1.377259 | 0.250019 | 1.32775 | 0.638648 | 0.412527 | 1.030401 | 0.141422 | 0.147631 | 1.000503 | 0.837425 | 0.246737 | | 19.73611111 | 0.036988 | 0.213378 |
| 19.652778 | 0.317166 | 0.25381 | 0.486216 | 0.424996 | 0.350471 | 1.113071 | 0.715597 | 0.934071 | 0.413083 | 0.160341 | 0.260423 | 0.362692 | 0.210133 | 1.134863 | 0.231217 | 1.072328 | 0.876198 | 0.214048 | 0.46382 | 0.193104 | 0.182874 | 0.810957 | 0.746294 | 0.26112 | | 19.75 | 0.017826 | 0.234585 |
| 19.666667 | 0.329968 | 0.422551 | 0.632888 | 0.440397 | 0.405494 | 1.938215 | 3.434097 | 0.581353 | 0.458237 | 0.169207 | 0.289732 | 0.233689 | 0.131813 | 0.735692 | 0.19765 | 0.861277 | 1.011117 | 0.034855 | -0.038178 | 0.138661 | 0.133766 | 0.556806 | 0.518441 | 0.126476 | | 19.76388889 | -0.048496 | 0.234937 |
| 19.680556 | 0.281397 | 0.48423 | 0.852087 | 0.397869 | 0.350651 | 1.816902 | 3.541088 | 0.447459 | 0.434073 | 0.123039 | 0.279023 | 0.144829 | 0.10744 | 0.378215 | 0.217159 | 0.570962 | 0.8491 | -0.048811 | -0.299737 | 0.10639 | 0.068066 | 0.365599 | 0.332902 | 0.041544 | | 19.77777778 | -0.069116 | 0.142701 |
| 19.694444 | 0.215574 | 0.392373 | 0.476149 | 0.336357 | 0.257491 | 1.190149 | 2.39013 | 0.314746 | 0.344566 | 0.07906 | 0.264214 | 0.101578 | 0.202232 | 0.095945 | 0.292568 | 0.343917 | 0.517666 | -0.001038 | -0.313236 | 0.115579 | 0.059045 | 0.264992 | 0.23462 | 0.076494 | | 19.79166667 | -0.003939 | 0.040962 |
| 19.708333 | 0.138536 | 0.177987 | 0.243926 | 0.273697 | 0.160143 | 0.56998 | 1.104094 | 0.208677 | 0.21116 | 0.027446 | 0.192245 | 0.074299 | 0.241927 | -0.10457 | 0.296912 | 0.16047 | 0.159961 | 0.119773 | 0.039519 | 0.078851 | 0.022885 | 0.127654 | 0.122444 | 0.055052 | | 19.80555556 | 0.038959 | 0.014961 |
| 19.722222 | 0.105599 | -0.052156 | 0.241441 | 0.292093 | 0.175593 | 0.132941 | 0.172238 | 0.157777 | 0.121024 | -0.004486 | 0.149789 | 0.05162 | 0.256796 | -0.131256 | 0.28066 | 0.01974 | -0.116914 | 0.257194 | 0.426997 | 0.017147 | -0.017829 | -0.022705 | 0.018255 | -0.063739 | | 19.81944444 | 0.056925 | 0.101766 |
| 19.736111 | 0.15207 | -0.081501 | 0.309089 | 0.307229 | 0.221058 | -0.141943 | -0.547655 | 0.143914 | 0.081966 | 0.010613 | 0.14189 | 0.038467 | 0.299756 | 0.041593 | 0.296863 | -0.202718 | -0.259587 | 0.339337 | 0.762725 | -0.00239 | 0.01644 | -0.053314 | 0.009376 | -0.134103 | | 19.83333333 | 0.061619 | 0.268796 |
| 19.75 | 0.207264 | 0.058769 | 0.272863 | 0.256606 | 0.214266 | -0.429251 | -1.700976 | 0.128427 | 0.051694 | 0.029372 | 0.13031 | 0.009213 | 0.191349 | 0.168578 | 0.344927 | -0.419547 | -0.321053 | 0.187944 | 0.389163 | 0.009284 | 0.077445 | 0.016538 | 0.031681 | -0.081867 | | 19.84722222 | 0.039536 | 0.387841 |
| 19.763889 | 0.251133 | 0.180572 | 0.19488 | 0.176192 | 0.163053 | -0.723865 | -3.26606 | 0.149895 | 0.051367 | 0.030962 | 0.12034 | -0.000348 | 0.093963 | 0.086247 | 0.292494 | -0.452562 | -0.261087 | -0.052377 | -0.116484 | 0.028347 | 0.070634 | 0.075582 | -0.00426 | 0.01811 | | 19.86111111 | 0.036828 | 0.377537 |
| 19.777778 | 0.267563 | 0.183381 | 0.179477 | 0.137827 | 0.132662 | -0.471394 | -2.637431 | 0.194844 | 0.090718 | 0.043702 | 0.138916 | 0.018411 | 0.116194 | -0.034274 | 0.199354 | -0.189176 | -0.067369 | -0.170364 | -0.357874 | 0.030914 | 0.011474 | 0.07696 | -0.053404 | 0.081045 | | 19.875 | 0.071064 | 0.278442 |
| 19.791667 | 0.239081 | 0.09745 | 0.140111 | 0.117375 | 0.107873 | 0.077687 | -0.204396 | 0.188113 | 0.113191 | 0.069612 | 0.175343 | 0.031895 | 0.128611 | -0.051313 | 0.16352 | 0.210667 | 0.131987 | -0.15538 | -0.328451 | 0.018112 | -0.000014 | 0.06761 | -0.034405 | 0.116701 | | 19.88888889 | 0.108867 | 0.206421 |
| 19.805556 | 0.193591 | 0.029659 | 0.062662 | 0.077137 | 0.073162 | 0.414414 | 1.516682 | 0.127887 | 0.085261 | 0.066764 | 0.194918 | 0.014356 | 0.056152 | 0.007913 | 0.230543 | 0.39211 | 0.128753 | -0.077589 | -0.173443 | 0.001374 | 0.026359 | 0.05124 | 0.027177 | 0.103274 | | 19.90277778 | 0.137998 | 0.194216 |
| 19.819444 | 0.1594 | -0.001465 | 0.008355 | 0.021988 | 0.02017 | 0.410809 | 1.704298 | 0.07165 | 0.041197 | 0.023447 | 0.17148 | -0.043863 | 0.009233 | -0.010107 | 0.281449 | 0.189676 | -0.038536 | 0.056844 | 0.131292 | -0.015261 | 0.016733 | -0.030739 | 0.021012 | -0.025127 | | 19.91666667 | 0.140732 | 0.156751 |
| 19.833333 | 0.148738 | -0.022182 | 0.017793 | -0.007943 | -0.020055 | 0.247278 | 0.869136 | 0.058669 | 0.035722 | -0.002484 | 0.143189 | -0.088982 | 0.084068 | -0.108004 | 0.255777 | -0.205462 | -0.216718 | 0.127689 | 0.323623 | -0.034904 | -0.026191 | -0.1658 | -0.070642 | -0.137376 | | 19.93055556 | 0.132291 | 0.03788 |
| 19.847222 | 0.137912 | -0.099351 | 0.044295 | 0.000059 | -0.022427 | 0.083801 | 0.179009 | 0.055531 | 0.051728 | 0.005868 | 0.153786 | -0.126714 | 0.140246 | -0.176705 | 0.239489 | -0.54595 | -0.366631 | 0.072495 | 0.191216 | -0.039752 | -0.038121 | -0.247389 | -0.170682 | -0.096189 | | 19.94444444 | 0.114943 | -0.085873 |
| 19.861111 | 0.103363 | -0.285759 | 0.018538 | 0.007925 | -0.017463 | -0.088174 | -0.1361 | 0.013969 | 0.038481 | 0.023829 | 0.144326 | -0.203649 | 0.125481 | -0.239733 | 0.222468 | -0.725157 | -0.475085 | -0.054437 | -0.189166 | -0.036535 | -0.0241 | -0.244965 | -0.263988 | 0.003619 | | 19.95833333 | 0.084304 | -0.184312 |
| 19.875 | 0.088433 | -0.393026 | -0.060671 | -0.021715 | -0.048481 | -0.264757 | -0.306326 | -0.028822 | -0.060409 | 0.016756 | 0.091745 | -0.347179 | 0.138811 | -0.453198 | 0.139709 | -0.749848 | -0.509648 | -0.208329 | -0.573581 | -0.054354 | -0.018991 | -0.243623 | -0.373412 | 0.070716 | | 19.97222222 | 0.043261 | -0.232629 |
| 19.888889 | 0.162862 | -0.242675 | -0.111694 | -0.075154 | -0.102517 | -0.388323 | -0.422059 | 0.098422 | -0.001246 | -0.026021 | 0.035614 | -0.52962 | 0.160093 | -0.793263 | 0.01484 | -0.77778 | -0.525661 | -0.35285 | -0.858892 | -0.024348 | -0.043186 | -0.370065 | -0.563712 | 0.073717 | | 19.98611111 | -0.008206 | -0.260596 |
| 19.902778 | 0.264491 | -0.04352 | -0.106906 | -0.098046 | -0.136597 | -0.416922 | -0.508305 | 0.453936 | 0.077875 | -0.049385 | 0.000057 | -0.681354 | 0.184295 | -1.019962 | -0.0868 | -0.896665 | -0.437559 | -0.365045 | -0.895587 | -0.022567 | -0.090455 | -0.6466 | -0.727403 | -0.040146 | | 20 | -0.015433 | -0.244651 |
| 19.916667 | 0.284972 | -0.016774 | -0.132422 | -0.124792 | -0.142637 | -0.450304 | -0.613249 | 0.415364 | 0.080632 | -0.036473 | -0.041897 | -0.817024 | 0.173838 | -0.898256 | -0.062737 | -1.105864 | -0.478101 | -0.331243 | -0.845895 | -0.099769 | -0.144676 | -1.021904 | -0.852397 | -0.252069 | | 20.01388889 | 0.065395 | -0.100514 |
| 19.930556 | 0.157691 | -0.245041 | -0.248584 | -0.243202 | -0.213524 | -0.566927 | -0.769033 | -0.153671 | -0.148926 | -0.04464 | -0.097835 | -1.01975 | 0.098402 | -0.671404 | -0.025457 | -1.354383 | -0.592767 | -0.328297 | -0.914377 | -0.150628 | -0.190817 | -1.248079 | -0.787608 | -0.394961 | | 20.02777778 | 0.203405 | 0.021031 |
| 19.944444 | -0.037256 | -0.453911 | -0.42166 | -0.454225 | -0.387064 | -0.753943 | -0.983452 | -0.680454 | -0.476326 | -0.169508 | -0.128834 | -0.966013 | 0.120169 | -0.541792 | -0.066237 | -1.434738 | -0.744671 | -0.264956 | -0.864556 | -0.13509 | -0.186689 | -0.923314 | -0.657653 | -0.286141 | | 20.04166667 | 0.274551 | -0.311562 |
| 19.958333 | -0.21817 | -0.434839 | -0.463855 | -0.567614 | -0.494721 | -0.858926 | -1.077429 | -1.067537 | -0.802437 | -0.314987 | -0.138586 | -0.677845 | 0.26635 | -0.581973 | -0.099126 | -1.187969 | -0.554108 | -0.197061 | -0.638958 | -0.010792 | -0.125596 | -0.426115 | -0.514326 | -0.091889 | | 20.05555556 | 0.231268 | -1.070254 |
| 19.972222 | -0.228475 | -0.237454 | -0.324462 | -0.425751 | -0.381865 | -0.701567 | -0.879619 | -1.103172 | -0.873272 | -0.263099 | -0.148919 | -0.456025 | 0.310676 | -0.677208 | -0.192659 | -0.976374 | -0.245998 | -0.241642 | -0.633129 | 0.034728 | -0.070002 | -0.183434 | -0.424834 | 0.065997 | | 20.06944444 | 0.175895 | -1.16481 |
| 19.986111 | -0.080224 | -0.046011 | -0.150922 | -0.185163 | -0.13426 | -0.395365 | -0.510279 | -0.901312 | -0.682368 | -0.057171 | -0.293005 | -0.467426 | 0.121237 | -0.657424 | -0.24836 | -0.967366 | -0.083962 | -0.463435 | -0.932008 | -0.001198 | -0.029446 | -0.159357 | -0.351907 | 0.135549 | | 20.08333333 | 0.119487 | -0.385942 |
| 20 | 0.014335 | -0.002437 | -0.138583 | -0.071361 | -0.014182 | -0.250989 | -0.270518 | -0.674552 | -0.521262 | 0.057753 | -0.556619 | -0.526573 | -0.056106 | -0.514679 | -0.24352 | -0.870465 | -0.028934 | -0.555835 | -1.074281 | -0.023206 | -0.035094 | -0.169087 | -0.298079 | 0.145282 | | 20.09722222 | 0.062887 | 0.114886 |
| 20.013889 | 0.017805 | -0.026172 | -0.207475 | -0.087093 | -0.025703 | -0.273289 | -0.206249 | -0.50721 | -0.486131 | 0.059864 | -0.622073 | -0.495052 | -0.127608 | -0.305889 | -0.166679 | -0.612835 | -0.004729 | -0.498369 | -0.895462 | 0.008713 | -0.037271 | -0.152832 | -0.232754 | 0.079575 | | 20.11111111 | 0.068447 | 0.271135 |
| 20.027778 | -0.017618 | -0.059384 | -0.268879 | -0.109729 | -0.050986 | -0.357455 | -0.187119 | -0.422967 | -0.441111 | 0.045995 | -0.48761 | -0.363568 | -0.178835 | -0.154266 | -0.100251 | -0.481356 | 0.000221 | -0.342699 | -0.632571 | 0.018114 | -0.039457 | -0.139424 | -0.190161 | -0.010019 | | 20.125 | 0.091832 | 0.287885 |
| 20.041667 | -0.063701 | -0.09071 | -0.295382 | -0.115445 | -0.060765 | -0.411728 | -0.165277 | -0.394613 | -0.395873 | 0.039826 | -0.373195 | -0.256371 | -0.207762 | -0.04021 | -0.034664 | -0.481041 | -0.063412 | -0.202133 | -0.287614 | -0.019784 | -0.042646 | -0.124669 | -0.133908 | -0.057105 | | 20.13888889 | 0.062506 | 0.098726 |
| 20.055556 | -0.103719 | -0.074315 | -0.235919 | -0.100757 | -0.05922 | -0.367249 | -0.129827 | -0.35606 | -0.303888 | 0.042843 | -0.29196 | -0.134962 | -0.150424 | 0.036177 | -0.003932 | -0.460097 | -0.145547 | -0.143894 | -0.155445 | -0.072552 | -0.035303 | -0.075953 | -0.069647 | -0.027903 | | 20.15277778 | 0.00719 | -0.043871 |
| 20.069444 | -0.164433 | -0.059001 | -0.145674 | -0.083847 | -0.041108 | -0.272352 | -0.111666 | -0.316046 | -0.204751 | 0.040565 | -0.205795 | -0.016314 | -0.069871 | 0.07236 | -0.003932 | -0.326597 | -0.148418 | -0.161599 | -0.226494 | -0.077128 | -0.025166 | -0.035977 | -0.033793 | 0.046645 | | 20.16666667 | -0.025999 | -0.08579 |
| 20.083333 | -0.228832 | -0.103966 | -0.120007 | -0.100585 | -0.039534 | -0.23106 | -0.103164 | -0.296485 | -0.1726 | 0.007003 | -0.172268 | 0.041265 | -0.039005 | 0.057023 | -0.015569 | -0.093533 | -0.009832 | -0.15876 | -0.277306 | -0.038059 | -0.004811 | -0.026793 | -0.018988 | 0.122347 | | 20.18055556 | -0.045945 | -0.085867 |
| 20.097222 | -0.267739 | -0.165449 | -0.146933 | -0.14299 | -0.068189 | -0.291087 | -0.122907 | -0.267985 | -0.181163 | -0.034952 | -0.194276 | 0.002577 | -0.020279 | -0.012281 | -0.028334 | 0.066998 | 0.155894 | -0.121663 | -0.20188 | -0.006727 | -0.002562 | -0.0538 | -0.001848 | 0.133785 | | 20.19444444 | -0.045866 | -0.104822 |
| 20.111111 | -0.249379 | -0.161993 | -0.152397 | -0.144121 | -0.070048 | -0.324693 | -0.135719 | -0.206218 | -0.153091 | -0.027794 | -0.191003 | -0.046839 | -0.022788 | -0.025968 | -0.020743 | 0.012233 | 0.114055 | -0.083527 | -0.140555 | -0.012821 | -0.000928 | -0.076246 | 0.027295 | 0.092424 | | 20.20833333 | -0.040279 | -0.013702 |
| 20.125 | -0.19996 | -0.114242 | -0.153209 | -0.114729 | -0.050213 | -0.343282 | -0.144259 | -0.155733 | -0.096271 | 0.004749 | -0.176776 | -0.026611 | -0.116935 | -0.031891 | 0.009867 | 0.012249 | 0.200052 | -0.073588 | -0.09759 | -0.047907 | 0.010027 | -0.081408 | 0.035493 | 0.067598 | | 20.22222222 | 0.057513 | -0.125776 |
| 20.138889 | -0.192831 | -0.108644 | -0.174893 | -0.104213 | -0.05864 | -0.322116 | -0.137672 | -0.180004 | -0.08254 | -0.000095 | -0.197443 | -0.080501 | -0.169681 | -0.050816 | -0.029083 | -0.041907 | 0.10831 | -0.124218 | -0.168286 | -0.107966 | -0.021431 | -0.131185 | -0.030384 | 0.014649 | | 20.23611111 | 0.015434 | -0.108844 |
| 20.152778 | -0.209476 | -0.126674 | -0.139809 | -0.099225 | -0.080418 | -0.28285 | -0.14642 | -0.267674 | -0.143696 | -0.039501 | -0.24631 | -0.112146 | -0.154618 | -0.07989 | -0.081941 | -0.031898 | 0.040246 | -0.175649 | -0.251764 | -0.133095 | -0.059311 | -0.198765 | -0.104852 | -0.053872 | | 20.25 | -0.041461 | 0.105052 |
| 20.166667 | -0.211044 | -0.150051 | -0.097465 | -0.097299 | -0.093321 | -0.237544 | -0.164669 | -0.379334 | -0.227508 | -0.060445 | -0.280169 | -0.090203 | -0.102071 | -0.04121 | -0.057618 | 0.045416 | 0.032666 | -0.17833 | -0.269594 | -0.118369 | -0.040917 | -0.208313 | -0.115337 | -0.044962 | | 20.26388889 | -0.034321 | 0.290877 |
| 20.180556 | -0.237378 | -0.195347 | -0.13547 | -0.113016 | -0.096221 | -0.208654 | -0.169683 | -0.441271 | -0.254241 | -0.040514 | -0.230592 | 0.005381 | -0.022926 | 0.007552 | -0.013592 | 0.148722 | 0.057935 | -0.162534 | -0.206565 | -0.11376 | -0.02265 | -0.201998 | -0.10608 | -0.011355 | | 20.27777778 | -0.038331 | 0.127406 |
| 20.194444 | -0.298684 | -0.266497 | -0.203228 | -0.144117 | -0.105425 | -0.219356 | -0.176297 | -0.392316 | -0.21159 | -0.036886 | -0.204818 | 0.067165 | -0.027974 | 0.021801 | -0.007078 | 0.259408 | 0.071115 | -0.124417 | -0.091569 | -0.113655 | -0.03117 | -0.206769 | -0.093089 | -0.007155 | | 20.29166667 | -0.077243 | 0.046432 |
| 20.208333 | -0.353836 | -0.300479 | -0.240339 | -0.179581 | -0.118201 | -0.25942 | -0.210816 | -0.303857 | -0.159688 | -0.083227 | -0.245621 | 0.035716 | -0.17538 | 0.041356 | -0.013859 | 0.351854 | 0.069516 | -0.082099 | 0.016397 | -0.09174 | -0.040319 | -0.176038 | -0.061038 | 0.011614 | | 20.30555556 | -0.070444 | 0.139112 |
| 20.222222 | -0.348176 | -0.260869 | -0.207047 | -0.15693 | -0.081073 | -0.242611 | -0.20606 | -0.199701 | -0.095587 | -0.089077 | -0.296234 | 0.007881 | -0.272241 | 0.060279 | -0.017676 | 0.387723 | 0.058488 | -0.07482 | 0.024517 | -0.05589 | -0.034064 | -0.09878 | -0.024884 | 0.052678 | | 20.31944444 | -0.023923 | -0.204378 |
| 20.236111 | -0.305898 | -0.197784 | -0.139667 | -0.096562 | -0.023718 | -0.194326 | -0.173367 | -0.116795 | -0.052869 | -0.066028 | -0.311823 | 0.030537 | -0.209079 | 0.067372 | -0.015116 | 0.398459 | 0.05624 | -0.07627 | -0.000118 | -0.019297 | -0.020402 | -0.018218 | 0.022477 | 0.05699 | | 20.33333333 | 0.000783 | -0.392725 |
| 20.25 | -0.319122 | -0.231737 | -0.153325 | -0.122981 | -0.058508 | -0.236506 | -0.217911 | -0.162681 | -0.119674 | -0.131807 | -0.341751 | 0.013762 | -0.16041 | 0.072987 | -0.008776 | 0.401576 | 0.082959 | -0.040689 | 0.037676 | 0.002585 | -0.015909 | 0.03974 | 0.063927 | 0.006976 | | 20.34722222 | 0.01562 | 0.089824 |
| 20.263889 | -0.354902 | -0.328862 | -0.217411 | -0.210142 | -0.15478 | -0.31954 | -0.317641 | -0.260233 | -0.212105 | -0.233476 | -0.295551 | -0.056367 | -0.23748 | 0.089938 | -0.016026 | 0.38117 | 0.0955 | 0.013467 | 0.108471 | 0.007529 | -0.02478 | 0.095638 | 0.086435 | -0.03248 | | 20.36111111 | 0.041918 | 0.240753 |
| 20.277778 | -0.331706 | -0.358547 | -0.233497 | -0.233481 | -0.192302 | -0.328487 | -0.357828 | -0.264491 | -0.207542 | -0.251684 | -0.159722 | -0.077339 | -0.310804 | 0.109985 | -0.036575 | 0.367677 | 0.091123 | 0.052146 | 0.164061 | 0.019362 | -0.024717 | 0.180651 | 0.134849 | -0.020707 | | 20.375 | 0.021955 | -0.004642 |
| 20.291667 | -0.256246 | -0.276977 | -0.174182 | -0.166145 | -0.141313 | -0.241805 | -0.304309 | -0.152893 | -0.124943 | -0.204007 | -0.052565 | -0.044337 | -0.27079 | 0.126565 | -0.045597 | 0.396902 | 0.08266 | 0.084835 | 0.210287 | 0.044338 | -0.00567 | 0.268285 | 0.191552 | 0.008711 | | 20.38888889 | -0.029555 | -0.131982 |
| 20.305556 | -0.189931 | -0.158364 | -0.097182 | -0.084758 | -0.075577 | -0.139091 | -0.194947 | -0.000606 | -0.048078 | -0.170941 | -0.027612 | -0.02334 | -0.207963 | 0.149111 | -0.051894 | 0.366796 | 0.038481 | 0.112643 | 0.269371 | 0.050479 | -0.000152 | 0.282124 | 0.202358 | -0.011765 | | 20.40277778 | -0.044956 | 0.138822 |
| 20.319444 | -0.170497 | -0.112968 | -0.098341 | -0.077568 | -0.078636 | -0.119099 | -0.130482 | 0.08409 | -0.038933 | -0.179671 | -0.035237 | -0.015423 | -0.234952 | 0.220507 | -0.041695 | 0.327608 | -0.003891 | 0.147456 | 0.34903 | 0.050098 | -0.003692 | 0.296029 | 0.233839 | -0.048904 | | 20.41666667 | -0.02818 | 0.142881 |
| 20.333333 | -0.2421 | -0.191863 | -0.227713 | -0.192617 | -0.201182 | -0.221285 | -0.192385 | 0.013214 | -0.122191 | -0.285276 | -0.099777 | -0.076986 | -0.329414 | 0.327613 | -0.001464 | 0.455315 | 0.030089 | 0.20498 | 0.433474 | 0.064299 | 0.021301 | 0.369893 | 0.357849 | -0.022556 | | 20.43055556 | -0.035536 | -0.106352 |
| 20.347222 | -0.362455 | -0.328174 | -0.338375 | -0.320418 | -0.355657 | -0.323452 | -0.243085 | -0.102134 | -0.207338 | -0.403698 | -0.188803 | -0.205501 | -0.389044 | 0.329937 | -0.00653 | 0.731746 | 0.147115 | 0.270928 | 0.517851 | 0.067962 | 0.03302 | 0.429938 | 0.443849 | 0.015052 | | 20.44444444 | -0.109276 | -0.341073 |
| 20.361111 | -0.286034 | -0.267366 | -0.268416 | -0.266265 | -0.303926 | -0.212694 | -0.11852 | 0.079879 | -0.066609 | -0.402324 | -0.078319 | -0.188273 | -0.275235 | 0.269237 | -0.023472 | 0.988151 | 0.342213 | 0.361892 | 0.668791 | 0.080851 | 0.026804 | 0.494884 | 0.488626 | 0.010674 | | 20.45833333 | -0.20346 | -0.338654 |
| 20.375 | 0.000937 | 0.010721 | 0.009355 | -0.010423 | -0.066945 | 0.038401 | 0.143801 | 0.49332 | 0.23298 | -0.205418 | 0.236822 | -0.030039 | -0.099978 | 0.338461 | 0.032432 | 1.253113 | 0.556108 | 0.471194 | 0.850539 | 0.122743 | 0.058263 | 0.646231 | 0.587696 | -0.062397 | | 20.47222222 | -0.285915 | -0.278085 |
| 20.388889 | 0.260279 | 0.226457 | 0.212808 | 0.211119 | 0.159549 | 0.226489 | 0.278936 | 0.813408 | 0.471594 | -0.080501 | 0.525817 | 0.081008 | -0.309043 | 0.529965 | 0.112528 | 1.597152 | 0.790874 | 0.559917 | 0.912237 | 0.188054 | 0.127774 | 0.789207 | 0.744798 | -0.133785 | | 20.48611111 | -0.259974 | -0.144005 |
| 20.402778 | 0.44965 | 0.399147 | 0.315285 | 0.361247 | 0.311764 | 0.361 | 0.398883 | 1.039332 | 0.615072 | -0.040356 | 0.664109 | 0.273191 | -0.520721 | 0.709502 | 0.185761 | 1.824187 | 0.924937 | 0.585137 | 0.895032 | 0.24213 | 0.21554 | 0.888162 | 0.884964 | -0.084689 | | 20.5 | -0.121467 | 0.212274 |
| 20.416667 | 0.613786 | 0.508611 | 0.313471 | 0.380033 | 0.34391 | 0.489 | 0.678057 | 1.246416 | 0.765512 | 0.000208 | 0.653909 | 0.765321 | 0.258092 | 0.819487 | 0.229574 | 1.855216 | 0.889689 | 0.596584 | 0.989384 | 0.26987 | 0.24676 | 1.061982 | 0.994997 | 0.066242 | | 20.51388889 | -0.040973 | 0.424422 |
| 20.430556 | 0.685469 | 0.473584 | 0.167269 | 0.272912 | 0.258809 | 0.498114 | 0.884361 | 1.491467 | 0.927115 | -0.046582 | 0.499698 | 1.059131 | 0.980673 | 0.943298 | 0.258305 | 1.832656 | 0.846335 | 0.61192 | 1.176576 | 0.207627 | 0.270144 | 1.330846 | 1.171722 | 0.229934 | | 20.52777778 | -0.049471 | 0.194779 |
| 20.444444 | 0.520806 | 0.026041 | -0.380898 | -0.13106 | -0.063152 | 0.047538 | 0.57035 | 1.299376 | 0.830325 | -0.387686 | 0.104111 | 0.495125 | 0.187318 | 1.228801 | 0.282024 | 1.693591 | 0.725978 | 0.539653 | 1.368705 | 0.097576 | 0.287028 | 1.494023 | 1.465811 | 0.202261 | | 20.54166667 | -0.117773 | -0.327261 |
| 20.458333 | -0.031715 | -0.625246 | -1.341983 | -0.777071 | -0.567698 | -0.819209 | -0.353072 | 0.576169 | 0.291787 | -0.957326 | -0.514866 | -0.386701 | -1.177 | 1.512749 | 0.24413 | 1.545518 | 0.487987 | 0.38071 | 1.227848 | -0.081934 | 0.216051 | 1.414381 | 1.649907 | -0.006333 | | 20.55555556 | -0.22497 | -0.793478 |
| 20.472222 | 0.025708 | -0.546072 | -1.320384 | -0.791808 | -0.586705 | -0.854907 | -0.741959 | 0.436309 | 0.061831 | -1.05447 | -0.464647 | -0.378521 | -1.293033 | 1.474346 | 0.313336 | 1.71522 | 0.447687 | 0.330839 | 1.026957 | -0.061194 | 0.109683 | 1.057084 | 1.355896 | -0.180019 | | 20.56944444 | -0.216804 | -0.535567 |
| 20.486111 | 0.979146 | 0.424891 | -0.086527 | -0.052945 | -0.013982 | 0.280907 | -0.294737 | 1.10279 | 0.458466 | -0.51617 | 0.404695 | 0.322358 | -0.303384 | 1.503983 | 0.441384 | 2.087939 | 0.466707 | 0.522699 | 0.980818 | 0.112721 | 0.074891 | 0.942837 | 0.990986 | -0.054525 | | 20.58333333 | -0.084302 | 0.24716 |
| 20.5 | 1.870099 | 1.115917 | 0.501631 | 0.538527 | 0.337084 | 1.338427 | 0.161275 | 1.589068 | 0.946244 | -0.082706 | 1.142399 | 0.639509 | 0.264581 | 1.374515 | 0.528956 | 2.369663 | 0.431054 | 0.652341 | 1.137552 | 0.190799 | 0.132118 | 1.328213 | 1.190071 | 0.302456 | | 20.59722222 | -0.047436 | 0.618067 |
| 20.513889 | 1.812779 | 1.036969 | 0.39228 | 0.572683 | 0.361703 | 1.51708 | 0.48882 | 1.600576 | 1.274758 | -0.060199 | 1.339589 | 0.597602 | 0.00577 | 0.999492 | 0.577596 | 1.822968 | 0.374239 | 0.184203 | 0.034337 | 0.141651 | 0.115333 | 1.669829 | 1.5111 | 0.397544 | | 20.61111111 | -0.144556 | 0.293745 |
| 20.527778 | 1.207337 | 0.807608 | 0.342108 | 0.50369 | 0.324256 | 1.340991 | 1.062781 | 1.376635 | 1.54736 | -0.197287 | 1.012533 | 0.873872 | -0.295021 | 0.666288 | 0.5131 | 0.854498 | 0.121783 | -0.035326 | -0.219529 | 0.078745 | 0.177618 | 1.164883 | 1.270236 | -0.005101 | | 20.625 | -0.205641 | -0.139072 |
| 20.541667 | 0.44903 | 0.507042 | 0.497355 | 0.472692 | 0.35262 | 1.018171 | 1.62979 | 0.911307 | 1.134813 | -0.23321 | 0.346226 | 0.899178 | -0.307524 | 0.414931 | 0.312741 | 0.308041 | -0.008263 | 0.062127 | 0.110619 | 0.180743 | 0.100476 | -0.24466 | 0.196841 | -0.35421 | | 20.63888889 | -0.169258 | -0.294983 |
| 20.555556 | 0.339352 | 0.205032 | 0.798958 | 0.296544 | 0.361149 | 0.898484 | 2.08014 | 0.47947 | 0.174877 | -0.160574 | -0.133298 | 0.407764 | -0.097592 | 0.494534 | 0.492709 | 0.178423 | 0.104091 | 0.197574 | 0.274149 | 0.166966 | 0.029203 | -0.420164 | -0.056752 | -0.314188 | | 20.65277778 | -0.106502 | -0.211366 |
| 20.569444 | 0.549996 | 0.126098 | 1.347776 | 0.126936 | 0.115799 | 0.628063 | 1.618137 | 0.240281 | -0.254748 | -0.107344 | -0.059044 | 0.068613 | 0.14824 | 0.555074 | 0.606037 | 0.881725 | 0.397835 | 0.367505 | 0.876062 | 0.13008 | 0.065266 | 0.238612 | 0.257319 | 0.038525 | | 20.66666667 | -0.062215 | -0.068578 |
| 20.583333 | 0.388204 | 0.391799 | 1.155955 | 0.13149 | 0.130584 | 0.290786 | 0.393375 | 0.279362 | 0.136144 | -0.095713 | 0.476882 | 0.103201 | 0.291441 | 0.468328 | 0.568669 | 1.284144 | 0.400552 | 0.15628 | 0.232216 | 0.064825 | 0.083499 | 0.49679 | 0.520827 | 0.20821 | | 20.68055556 | -0.041867 | -0.007297 |
| 20.597222 | 0.223811 | 0.625659 | 0.783347 | 0.151283 | -0.137227 | 0.185084 | -0.144341 | 0.614043 | 0.558522 | -0.109812 | 0.730594 | 0.30891 | 0.271935 | 0.25953 | 0.430591 | 0.908103 | 0.215966 | 0.153406 | 0.205242 | 0.105785 | 0.078104 | 0.128638 | 0.289349 | 0.12613 | | 20.69444444 | -0.041696 | -0.051947 |
| 20.611111 | 0.264558 | 0.470469 | 0.509134 | 0.156958 | -0.087849 | 0.263631 | 0.142256 | 0.561765 | 0.526912 | -0.126361 | 0.506533 | 0.723089 | 0.212525 | 0.281664 | 0.591362 | 0.621353 | 0.092179 | -0.001039 | -0.036057 | 0.150065 | 0.057919 | 0.205187 | 0.031184 | 0.189338 | | 20.70833333 | -0.026668 | -0.112771 |
| 20.625 | 0.50134 | 0.113459 | 0.532046 | 0.149956 | 0.038478 | 0.451662 | 0.074837 | 0.287281 | 0.186284 | -0.110513 | 0.066687 | 0.853347 | 0.122579 | 0.473639 | 1.000263 | 0.799094 | 0.271633 | 0.215723 | 0.628927 | 0.08881 | 0.094905 | 0.802947 | 0.1307 | 0.323586 | | 20.72222222 | -0.004237 | -0.08651 |
| 20.638889 | 0.320782 | 0.096584 | 0.604346 | 0.123166 | 0.02548 | 0.182317 | 0.307904 | 0.682135 | 0.288158 | -0.047334 | -0.113973 | 0.746461 | 0.020029 | 0.112735 | 0.569089 | 0.77585 | 0.341318 | 0.081228 | 0.097402 | -0.010199 | 0.045647 | 0.473292 | 0.115798 | 0.219239 | | 20.73611111 | -0.005182 | 0.028356 |
| 20.652778 | -0.146955 | 0.3824 | 0.388572 | 0.041867 | -0.122199 | -0.019632 | 0.628517 | 0.848066 | 0.394218 | 0.029944 | 0.108258 | 0.371424 | -0.075915 | -0.274769 | 0.008509 | -0.070476 | 0.093196 | 0.224032 | 0.449187 | -0.061377 | -0.028745 | -0.234476 | -0.185781 | -0.090213 | | 20.75 | -0.016658 | 0.125003 |
| 20.666667 | -0.117163 | 0.455621 | 0.148622 | -0.006195 | -0.063403 | 0.124961 | 0.487643 | 0.446094 | 0.072396 | 0.026381 | 0.28901 | 0.025893 | -0.058644 | -0.20012 | 0.029533 | -0.998288 | -0.312158 | 0.102384 | 0.170373 | 0.004835 | -0.023691 | -0.374992 | -0.351678 | -0.292477 | | 20.76388889 | -0.017764 | 0.194747 |
| 20.680556 | 0.652104 | 0.175318 | 0.348442 | 0.052084 | -0.052126 | 0.189907 | 0.126912 | 0.011015 | -0.365238 | -0.003641 | 0.204088 | 0.25896 | 0.067447 | 0.132843 | 0.468333 | -0.991435 | -0.172513 | 0.038533 | 0.164079 | -0.027656 | 0.00144 | 0.048493 | -0.225068 | -0.214515 | | 20.77777778 | 0.00006 | 0.212983 |
| 20.694444 | 1.605316 | 0.028161 | 0.679177 | 0.25288 | 0.015443 | 0.435683 | 0.265185 | -0.172114 | -0.205842 | -0.016454 | -0.018093 | 0.149124 | 0.193829 | 0.357974 | 0.745963 | -0.11606 | 0.378557 | 0.055695 | 0.103523 | -0.08607 | 0.052915 | 0.026659 | -0.16989 | -0.007245 | | 20.79166667 | 0.036866 | 0.225254 |
| 20.708333 | 0.68647 | 0.160599 | 0.563205 | 0.340596 | 0.117226 | 0.483888 | 0.401615 | 0.143001 | 0.48062 | 0.046791 | -0.130972 | 0.382251 | 0.169383 | 0.05558 | 0.399533 | 0.511738 | 0.429233 | 0.243856 | 0.457722 | -0.059257 | 0.004332 | -0.492367 | -0.438831 | 0.119675 | | 20.80555556 | 0.077426 | 0.269389 |
| 20.722222 | -1.18714 | 0.219923 | 0.187568 | 0.28004 | 0.123401 | 0.195341 | 0.185458 | 0.329691 | 0.729073 | 0.091099 | -0.143542 | 0.200019 | -0.070021 | -0.358456 | -0.176571 | 0.086557 | -0.005996 | 0.384529 | 0.803418 | 0.028764 | -0.058589 | -0.498849 | -0.583829 | 0.08241 | | 20.81944444 | 0.11423 | 0.325807 |
| 20.736111 | -1.513127 | 0.132666 | 0.013273 | 0.177595 | 0.051252 | 0.15261 | 0.219203 | 0.031473 | 0.113622 | -0.050608 | -0.128677 | 0.003778 | -0.13986 | -0.330752 | -0.145932 | -0.516414 | -0.111308 | -0.011864 | 0.039871 | -0.049557 | -0.077018 | 0.173884 | -0.199194 | -0.102081 | | 20.83333333 | 0.143594 | 0.34818 |
| 20.75 | 0.199802 | -0.162097 | 0.001264 | 0.097036 | -0.011184 | 0.139421 | 0.025897 | -0.397031 | -0.65479 | -0.112581 | -0.098601 | -0.162026 | 0.029277 | 0.133883 | 0.540087 | -0.72334 | -0.136872 | -0.543625 | -1.17947 | -0.185572 | -0.069906 | 0.367398 | 0.273077 | -0.340508 | | 20.84722222 | 0.11994 | 0.280725 |
| 20.763889 | 1.980653 | 0.080162 | 0.044957 | -0.001271 | -0.045466 | 0.17079 | -0.075312 | -0.45394 | -0.531527 | -0.044517 | -0.030909 | -0.168063 | 0.169443 | 0.339159 | 0.867809 | -0.1636 | -0.078807 | -0.375653 | -0.948209 | -0.219129 | -0.133754 | -0.154957 | 0.205204 | -0.197408 | | 20.86111111 | 0.074174 | 0.09251 |
| 20.777778 | 1.226743 | 0.129449 | 0.065229 | -0.101956 | -0.198565 | 0.105507 | -0.173383 | 0.313426 | 0.543887 | 0.047928 | 0.086023 | 0.15082 | 0.092781 | 0.031709 | 0.25094 | 0.726889 | 0.23729 | 0.020763 | 0.049703 | -0.223808 | -0.19668 | -0.179453 | -0.033138 | 0.259557 | | 20.875 | 0.054218 | -0.121253 |
| 20.791667 | -0.568489 | -0.053955 | -0.017809 | -0.190703 | -0.231022 | 0.00373 | -0.111199 | 1.297669 | 1.661433 | 0.045231 | 0.138119 | 0.305569 | -0.163841 | -0.353191 | -0.399535 | 0.572143 | 0.179951 | 0.22618 | 0.666412 | -0.122434 | -0.141793 | 0.131925 | -0.083373 | 0.322002 | | 20.88888889 | 0.046649 | -0.23753 |
| 20.805556 | -1.893125 | -0.510737 | -0.258477 | -0.294392 | -0.199104 | 0.001306 | 0.10925 | 0.855818 | 1.168439 | -0.109108 | -0.056916 | -0.139178 | -0.331488 | -0.314207 | -0.121843 | -0.319172 | -0.071355 | -0.187493 | -0.346574 | -0.146141 | -0.08975 | 0.165647 | 0.203236 | -0.188498 | | 20.90277778 | 0.073217 | -0.338695 |
| 20.819444 | -0.963733 | -0.958905 | -0.430218 | -0.333628 | -0.243298 | 0.009862 | 0.286373 | -0.423149 | -0.230854 | -0.217912 | -0.359777 | -0.667544 | -0.220161 | -0.071986 | 0.472971 | -1.025441 | -0.301259 | -0.558969 | -1.406786 | -0.214303 | -0.098881 | -0.320963 | 0.245115 | -0.709316 | | 20.91666667 | 0.08657 | -0.486623 |
| 20.833333 | 0.664434 | 0.084745 | -0.436788 | -0.328547 | -0.241432 | -0.262556 | -0.060687 | -0.886132 | -0.906063 | -0.270861 | -0.459056 | -0.593779 | -0.100269 | -0.119939 | 0.384085 | -0.901936 | -0.338652 | 0.007279 | -0.138951 | -0.096926 | -0.103252 | -0.435624 | -0.129909 | -0.597149 | | 20.93055556 | 0.069989 | -0.636794 |
| 20.847222 | 0.502422 | 0.098758 | -0.334977 | -0.381166 | -0.355367 | -0.657517 | -0.612849 | -0.521823 | -0.486059 | -0.380107 | -0.225971 | -0.238198 | -0.112795 | -0.48552 | -0.419971 | -0.736872 | -0.14776 | 0.200324 | 0.481419 | -0.037611 | -0.067678 | -0.10314 | -0.590285 | 0.005519 | | 20.94444444 | 0.048809 | -0.707898 |
| 20.861111 | -0.721589 | 0.099098 | -0.253777 | -0.540331 | -0.439498 | -0.622083 | -0.812687 | -0.268979 | -0.076738 | -0.279628 | -0.072376 | -0.243765 | -0.104868 | -0.506042 | -0.525363 | -0.692658 | -0.264739 | 0.133101 | 0.37591 | 0.051332 | -0.059526 | -0.512434 | -0.949769 | 0.259025 | | 20.95833333 | 0.011883 | -0.678655 |
| 20.875 | -1.538306 | -0.681661 | -0.292024 | -0.568478 | -0.324048 | -0.371095 | -0.689782 | -0.771484 | -0.381045 | -0.202083 | -0.302699 | -0.609338 | 0.027644 | -0.435975 | -0.33217 | -1.441439 | -0.438186 | -0.465784 | -1.202547 | -0.049087 | -0.162742 | -0.989795 | -0.617315 | -0.076162 | | 20.97222222 | 0.011567 | -0.577518 |
| 20.888889 | -1.010624 | -0.933338 | -0.52774 | -0.503747 | -0.242438 | -0.282576 | -0.81331 | -1.376101 | -1.323737 | 0.000559 | -0.748729 | -0.568433 | 0.002213 | -0.638524 | -0.348614 | -2.6259 | -0.5725 | -0.405943 | -1.097583 | -0.070907 | -0.199039 | -0.873295 | -0.412455 | -0.406217 | | 20.98611111 | 0.060151 | -0.404612 |
| 20.902778 | -0.270131 | -0.309025 | -0.661161 | -0.495582 | -0.263882 | -0.575794 | -1.022264 | -1.490327 | -1.720798 | 0.1346 | -0.877376 | -0.430411 | -0.208822 | -1.031258 | -0.658528 | -3.046417 | -0.622164 | -0.285211 | -0.80088 | -0.09924 | -0.195252 | -0.449504 | -0.781068 | -0.420397 | | 21 | 0.097002 | -0.249493 |
| 20.916667 | -0.475615 | -0.048589 | -0.686596 | -0.618807 | -0.34687 | -0.979678 | -1.273179 | -1.118347 | -1.419066 | -0.267111 | -0.739465 | -0.513041 | -0.462817 | -1.226712 | -0.945408 | -1.903427 | -0.648968 | -0.403927 | -1.0186 | -0.1226 | -0.168108 | -0.673514 | -1.43493 | -0.128219 | | 21.01388889 | 0.13989 | -0.10365 |
| 20.930556 | -1.446793 | -0.263732 | -0.733665 | -0.693986 | -0.325425 | -0.941327 | -1.421817 | -0.999396 | -1.023953 | -0.10009 | -0.710982 | -0.686621 | -0.560993 | -1.147284 | -1.098817 | -0.815022 | -0.707521 | -0.655465 | -1.554711 | -0.136414 | -0.261778 | -1.352288 | -1.502654 | 0.147538 | | 21.02777778 | 0.23897 | 0.014763 |
| 20.944444 | -1.798978 | -0.67421 | -0.606809 | -0.578498 | -0.239173 | -0.698466 | -1.445283 | -1.360053 | -1.144253 | 0.037268 | -0.76238 | -0.936128 | -0.495239 | -0.785583 | -0.810108 | -1.579374 | -0.893055 | -1.013854 | -2.379107 | -0.060051 | -0.31293 | -1.447173 | -1.090742 | 0.103522 | | 21.04166667 | 0.314076 | -0.098288 |
| 20.958333 | -1.429877 | -0.599731 | -0.347534 | -0.453189 | -0.276132 | -0.487299 | -1.242085 | -1.387036 | -1.381876 | -0.099444 | -0.904298 | -0.735713 | -0.328556 | -0.713482 | -0.878077 | -2.442314 | -0.724884 | -0.642725 | -1.518778 | 0.088971 | -0.228676 | -0.892954 | -0.842637 | 0.11678 | | 21.05555556 | 0.276559 | -0.440554 |
| 20.972222 | -1.072556 | -0.100647 | -0.296035 | -0.354777 | -0.315551 | -0.364072 | -0.866108 | -1.15779 | -1.284733 | -0.046674 | -1.092362 | -0.742068 | 0.07538 | -0.870957 | -1.259195 | -1.926916 | -0.251894 | -0.365328 | -0.793169 | 0.089991 | -0.07758 | -0.406668 | -0.828429 | 0.242149 | | 21.06944444 | 0.223088 | -0.457649 |
| 20.986111 | -1.091644 | 0.461796 | -0.562862 | -0.370616 | -0.28986 | -0.396239 | -0.345145 | -1.082356 | -1.329466 | 0.061302 | -1.147027 | -0.895954 | 0.420842 | -0.973539 | -1.547618 | -0.975749 | -0.064809 | -0.405322 | -0.757214 | -0.093125 | -0.072607 | -0.175895 | -0.776644 | 0.254704 | | 21.08333333 | 0.157398 | -0.090353 |
| 21 | -1.286175 | 0.209792 | -0.909023 | -0.39079 | -0.19725 | -0.621261 | -0.213534 | -1.153179 | -1.212034 | 0.058526 | -0.938272 | -1.04834 | 0.314035 | -0.627448 | -0.940198 | -0.403765 | -0.152018 | -0.527412 | -0.917671 | -0.19299 | -0.173219 | -0.102742 | -0.53745 | 0.142622 | | 21.09722222 | 0.068333 | 0.111149 |
| 21.013889 | -1.335846 | -0.807975 | -0.969462 | -0.440269 | -0.223362 | -0.888427 | -0.47125 | -1.05191 | -0.941015 | -0.041862 | -0.637794 | -0.989398 | -0.039391 | -0.381188 | -0.414586 | -0.559541 | -0.296508 | -0.531969 | -0.973999 | -0.196567 | -0.181811 | -0.234968 | -0.385167 | -0.09325 | | 21.11111111 | 0.039317 | 0.139987 |
| 21.027778 | -1.117598 | -0.999292 | -0.644248 | -0.449129 | -0.306836 | -0.854048 | -0.716325 | -1.037921 | -0.745224 | -0.111618 | -0.484709 | -0.707048 | -0.219141 | -0.355885 | -0.325923 | -0.946935 | -0.479209 | -0.561267 | -1.132064 | -0.262355 | -0.186221 | -0.545127 | -0.460625 | -0.233213 | | 21.125 | 0.044424 | 0.228201 |
| 21.041667 | -0.671078 | -0.470533 | -0.264126 | -0.390463 | -0.340352 | -0.733013 | -0.846984 | -0.87458 | -0.660223 | -0.103172 | -0.50322 | -0.423402 | -0.236339 | -0.24088 | -0.37099 | -0.915932 | -0.561074 | -0.602918 | -1.351909 | -0.322321 | -0.185547 | -0.623796 | -0.51965 | -0.064127 | | 21.13888889 | 0.04635 | 0.282808 |
| 21.055556 | -0.371885 | -0.190053 | -0.176631 | -0.319343 | -0.286632 | -0.68272 | -0.967335 | -0.676786 | -0.583262 | -0.064631 | -0.506474 | -0.239743 | -0.241609 | -0.053766 | -0.278244 | -0.808942 | -0.559722 | -0.548264 | -1.186689 | -0.340641 | -0.124008 | -0.40502 | -0.437706 | 0.147274 | | 21.15277778 | 0.03684 | 0.185608 |
| 21.069444 | -0.381623 | -0.300573 | -0.286174 | -0.276961 | -0.239099 | -0.661055 | -0.930832 | -0.522667 | -0.522084 | -0.048245 | -0.44119 | -0.160162 | -0.304579 | -0.074276 | -0.271052 | -0.992169 | -0.648488 | -0.418673 | -0.799415 | -0.238343 | -0.078219 | -0.277625 | -0.309112 | 0.099042 | | 21.16666667 | 0.020488 | -0.049122 |
| 21.083333 | -0.478842 | -0.448109 | -0.353349 | -0.265999 | -0.236597 | -0.577784 | -0.715546 | -0.483081 | -0.440969 | -0.063594 | -0.435945 | -0.165272 | -0.305338 | -0.171131 | -0.346583 | -1.062627 | -0.713081 | -0.352771 | -0.603879 | -0.106408 | -0.074822 | -0.32381 | -0.279357 | -0.025734 | | 21.18055556 | 0.020056 | -0.22068 |
| 21.097222 | -0.452207 | -0.290591 | -0.390332 | -0.271779 | -0.26291 | -0.503854 | -0.483337 | -0.462289 | -0.378876 | -0.08164 | -0.4847 | -0.219171 | -0.244346 | -0.073195 | -0.287024 | -0.713952 | -0.640992 | -0.305868 | -0.603313 | -0.002531 | -0.049916 | -0.25477 | -0.25619 | -0.002358 | | 21.19444444 | 0.003878 | -0.215061 |
| 21.111111 | -0.323683 | -0.050007 | -0.389449 | -0.256883 | -0.250653 | -0.441746 | -0.345395 | -0.433574 | -0.343273 | -0.084143 | -0.447961 | -0.284945 | -0.265663 | 0.054417 | -0.158702 | -0.276986 | -0.411687 | -0.323213 | -0.600193 | 0.036367 | -0.027849 | -0.097242 | -0.20754 | 0.077097 | | 21.20833333 | -0.009936 | -0.162155 |
| 21.125 | -0.209071 | -0.029504 | -0.299973 | -0.181058 | -0.158031 | -0.376191 | -0.297706 | -0.358631 | -0.259087 | -0.045308 | -0.330797 | -0.349217 | -0.221362 | 0.037531 | -0.104219 | -0.050864 | -0.182237 | -0.274597 | -0.457876 | 0.032747 | -0.034672 | -0.032208 | -0.147958 | 0.064898 | | 21.22222222 | 0.055616 | -0.037289 |
| 21.138889 | -0.159622 | -0.167081 | -0.191301 | -0.056277 | 0.000043 | -0.279487 | -0.201526 | -0.178744 | -0.082847 | 0.013487 | -0.13622 | -0.271939 | -0.056744 | -0.046532 | -0.096657 | 0.058439 | -0.100468 | -0.245568 | -0.341856 | 0.031672 | -0.033325 | -0.009463 | -0.020995 | -0.135329 | | 21.23611111 | 0.144851 | 0.120677 |
| 21.152778 | -0.101259 | -0.158147 | -0.089957 | 0.057317 | 0.123637 | -0.117328 | -0.085009 | 0.002169 | 0.066542 | 0.07721 | 0.024986 | -0.055293 | -0.001265 | -0.022053 | -0.077929 | -0.085843 | -0.471107 | -0.177917 | -0.255232 | 0.029514 | -0.016138 | 0.070466 | 0.107733 | -0.359347 | | 21.25 | 0.176514 | 0.179746 |
| 21.166667 | 0.016333 | 0.104379 | 0.007697 | 0.119277 | 0.16041 | 0.030486 | -0.053335 | 0.098341 | 0.136293 | 0.124924 | 0.124835 | 0.082136 | 0.015174 | 0.132151 | 0.01559 | -0.063397 | -0.67168 | -0.072053 | -0.056399 | 0.01612 | 0.007126 | 0.162112 | 0.150198 | -0.142919 | | 21.26388889 | 0.12686 | 0.119929 |
| 21.180556 | 0.112664 | 0.359684 | 0.07054 | 0.105976 | 0.13089 | 0.075694 | -0.086437 | 0.156042 | 0.175966 | 0.099817 | 0.161142 | 0.091149 | 0.120247 | 0.239195 | 0.151688 | 0.201623 | -0.250998 | 0.03379 | 0.147994 | 0.028531 | 0.034716 | 0.199329 | 0.111274 | 0.28225 | | 21.27777778 | 0.041134 | 0.054559 |
| 21.194444 | 0.105931 | 0.346297 | 0.047169 | 0.062499 | 0.108151 | 0.035236 | -0.075827 | 0.217348 | 0.204227 | 0.051606 | 0.121097 | 0.072089 | 0.143403 | 0.232886 | 0.247949 | 0.396533 | 0.017066 | 0.088993 | 0.205703 | 0.098007 | 0.068058 | 0.185463 | 0.082317 | 0.352794 | | 21.29166667 | -0.032133 | 0.078177 |
| 21.208333 | 0.060852 | 0.16734 | 0.004194 | 0.065276 | 0.138017 | 0.074844 | 0.034832 | 0.288082 | 0.210321 | 0.061714 | 0.144285 | 0.120769 | 0.046981 | 0.24032 | 0.270807 | 0.668902 | 0.467221 | 0.098404 | 0.161166 | 0.163222 | 0.089698 | 0.182821 | 0.117473 | 0.176943 | | 21.30555556 | -0.039648 | 0.089587 |
| 21.222222 | 0.067932 | 0.135911 | 0.040177 | 0.080523 | 0.147384 | 0.198337 | 0.18799 | 0.270974 | 0.180305 | 0.073892 | 0.467127 | 0.240121 | -0.038498 | 0.28912 | 0.281825 | 0.745144 | 0.735086 | 0.130575 | 0.183092 | 0.150812 | 0.084692 | 0.230122 | 0.175466 | 0.030128 | | 21.31944444 | -0.01822 | -0.009632 |
| 21.236111 | 0.195165 | 0.309617 | 0.104875 | 0.093546 | 0.14039 | 0.273378 | 0.317092 | 0.244017 | 0.225087 | 0.076633 | 0.634254 | 0.290669 | 0.023862 | 0.336325 | 0.322378 | 0.64655 | 0.604849 | 0.160143 | 0.26904 | 0.140674 | 0.117316 | 0.295506 | 0.233131 | 0.006172 | | 21.33333333 | 0.003517 | 0.009072 |
| 21.25 | 0.391738 | 0.460243 | 0.191335 | 0.158721 | 0.187254 | 0.285049 | 0.459001 | 0.391885 | 0.373228 | 0.127127 | 0.439716 | 0.249746 | 0.217301 | 0.317231 | 0.394134 | 0.710571 | 0.580536 | 0.154202 | 0.235445 | 0.151028 | 0.177783 | 0.276824 | 0.224092 | 0.049666 | | 21.34722222 | 0.06572 | 0.241482 |
| 21.263889 | 0.477893 | 0.401116 | 0.242244 | 0.213615 | 0.21715 | 0.315955 | 0.589808 | 0.573698 | 0.511499 | 0.155884 | 0.472976 | 0.270855 | 0.320209 | 0.268191 | 0.50044 | 0.772031 | 0.606782 | 0.139403 | 0.177537 | 0.140028 | 0.17278 | 0.224523 | 0.158303 | 0.030526 | | 21.36111111 | 0.118693 | 0.380137 |
| 21.277778 | 0.443963 | 0.354899 | 0.240757 | 0.221066 | 0.180553 | 0.388336 | 0.674217 | 0.586253 | 0.569576 | 0.126053 | 0.650516 | 0.28648 | 0.262463 | 0.287386 | 0.552017 | 0.709825 | 0.555005 | 0.162244 | 0.248407 | 0.129768 | 0.128504 | 0.273517 | 0.20474 | 0.027606 | | 21.375 | 0.124168 | 0.330665 |

| | | | | | | | | | |
|---|---|---|---|---|---|---|---|---|---|
| 21.291667 | 0.408437 | 0.415786 | 0.256182 | 0.227755 | 0.167482 | 0.444241 | 0.742039 | 0.519717 | [illegible] |
| 21.305556 | 0.377981 | 0.354531 | 0.306659 | 0.230248 | 0.201259 | 0.463669 | 0.71909 | 0.51487 | [illegible] |
| 21.319444 | 0.415481 | 0.247822 | 0.340438 | 0.242808 | 0.252947 | 0.486309 | 0.675639 | 0.522252 | [illegible] |
| 21.333333 | 0.521509 | 0.31185 | 0.339391 | 0.295639 | 0.310685 | 0.545188 | 0.567242 | 0.498167 | [illegible] |
| 21.347222 | 0.58989 | 0.370132 | 0.329408 | 0.336234 | 0.331451 | 0.554467 | 0.462002 | 0.614273 | [illegible] |
| 21.361111 | 0.7211 | 0.34316 | 0.483382 | 0.444532 | 0.414181 | 0.632027 | 0.558863 | 0.922175 | [illegible] |
| 21.375 | 0.936791 | 0.448594 | 0.808976 | 0.681237 | 0.610185 | 0.888223 | 0.969156 | 1.166541 | [illegible] |
| 21.388889 | 1.10769 | 0.615106 | 0.98036 | 0.862384 | 0.747363 | 1.087337 | 1.239847 | 1.278073 | [illegible] |
| 21.402778 | 1.251396 | 0.630225 | 0.957386 | 0.873191 | 0.741372 | 1.070377 | 1.183426 | 1.5765 | [illegible] |
| 21.416667 | 1.479782 | 0.567733 | 1.084728 | 0.9419 | 0.78281 | 1.164133 | 1.307668 | 2.088242 | [illegible] |
| 21.430556 | 1.739453 | 0.694369 | 1.322202 | 1.0929 | 0.901844 | 1.399954 | 1.620112 | 2.241158 | [illegible] |
| 21.444444 | 1.612354 | 0.841427 | 1.124656 | 1.011596 | 0.910458 | 1.302895 | 1.300089 | 1.937534 | [illegible] |
| 21.458333 | 1.15735 | 0.734348 | 0.313912 | 0.386242 | 0.331894 | 0.616284 | 0.506416 | 1.425384 | [illegible] |
| 21.472222 | 1.081458 | 0.60345 | 0.309336 | 0.026512 | -0.221984 | 0.349318 | 0.5096 | 1.565096 | [illegible] |
| 21.486111 | 1.89098 | 1.017417 | 1.326257 | 0.665639 | -0.207654 | 1.052561 | 1.71224 | 2.164491 | [illegible] |
| 21.5 | 2.500347 | 1.546403 | 1.629057 | 1.046739 | -0.009054 | 1.785101 | 1.226485 | 1.528228 | [illegible] |
| 21.513889 | 1.484658 | 0.979872 | 0.601568 | 0.543504 | 0.805088 | 0.984477 | 0.721276 | 0.856521 | [illegible] |
| 21.527778 | 0.724479 | 0.496267 | 0.340343 | 0.567955 | 0.903143 | 0.257696 | 0.148955 | 0.630474 | [illegible] |
| 21.541667 | 0.482771 | 0.213278 | 0.313273 | 0.56581 | 0.484501 | 0.346353 | 0.528715 | 0.463729 | [illegible] |
| 21.555556 | 0.361389 | 0.095478 | 0.285105 | 0.35904 | 0.184763 | 0.474713 | 0.780373 | 0.317606 | [illegible] |
| 21.569444 | 0.135478 | 0.217932 | 0.337751 | 0.170347 | -0.143703 | 0.314076 | 0.493536 | 0.470075 | [illegible] |
| 21.583333 | 0.427177 | 0.545606 | 0.336882 | 0.106056 | -0.164381 | 0.05157 | -0.118635 | 0.542415 | [illegible] |
| [illegible] | [illegible] | [illegible] | [illegible] | [illegible] | [illegible] | [illegible] | [illegible] | [illegible] | [illegible] |

| | | | | | | | | | | | | | | | |
|---|---|---|---|---|---|---|---|---|---|---|---|---|---|---|---|
| 24.611111 | 0.167457 | 0.750663 | 1.067529 | 1.128593 | 0.148773 | -0.359929 | 0.293443 | -0.366112 | 0.613001 | -0.169558 | 0.228267 | 0.617669 | 0.708484 | 1.051848 | 1.998285 |
| 24.625 | 0.003386 | -0.490563 | 0.560027 | 0.148334 | -0.131037 | 0.231688 | 0.547937 | -0.149638 | 0.274883 | -0.176646 | 0.150249 | -0.055294 | 0.55116 | 0.976274 | 1.721862 |
| 24.638889 | 0.319397 | -0.446516 | 0.200048 | -0.329803 | -0.271671 | 0.73062 | 0.170167 | 0.375411 | -0.673649 | 0.36766 | 0.378008 | -0.405777 | -0.151743 | 0.784281 | 1.514676 |
| 24.652778 | 0.509782 | 0.309407 | -0.889545 | -0.900872 | 0.145264 | 0.53624 | -0.225564 | 0.047812 | -0.776991 | 0.580183 | 0.672757 | -0.194995 | -0.724828 | 0.566349 | 1.391952 |
| 24.666667 | -0.123623 | -0.403018 | -0.283988 | 0.140054 | 0.266803 | -0.195833 | -0.648871 | -0.188308 | 0.217325 | 0.71436 | 0.520436 | -0.592751 | -0.392977 | 0.425364 | 1.512376 |
| 24.680556 | -0.443578 | -0.32732 | 0.120552 | 0.453104 | 0.61973 | 0.002686 | 0.547879 | 0.190116 | 0.94341 | 0.290261 | 0.384115 | -0.354679 | 0.525173 | 0.371538 | 1.443475 |
| 24.694444 | 0.34383 | 0.667878 | 0.443057 | 0.494046 | 0.577734 | 0.197392 | 0.379581 | 0.022559 | 1.105095 | 0.751013 | -1.004665 | 1.00914 | 0.636355 | 0.343158 | 1.112488 |
| 24.708333 | 1.319734 | 2.122552 | 0.647258 | 0.059443 | -0.434161 | 0.146096 | -0.532855 | -0.368687 | -0.841025 | 0.105568 | -1.557619 | 0.036685 | -0.291388 | 0.341975 | 0.736734 |
| 24.722222 | 0.667145 | -0.293458 | -0.145387 | -0.13983 | -0.024936 | -0.355191 | -0.535706 | -0.630996 | -2.63246 | -1.026862 | -0.403044 | -1.382533 | -0.989558 | 0.357425 | 0.399936 |
| 24.736111 | -0.706617 | -1.784445 | -0.227429 | 0.288463 | 0.48606 | -0.853077 | -0.689911 | 0.524521 | -1.510711 | -2.04549 | 0.555633 | -1.498254 | -0.736961 | 0.372116 | 0.153752 |
| 24.75 | -1.140801 | -1.49433 | 0.218212 | 0.650306 | 0.131164 | 0.552819 | 0.965363 | 1.193383 | 0.951968 | -0.628112 | 1.43224 | 0.681274 | 0.158678 | 0.398775 | 0.236748 |
| 24.763889 | 0.178662 | 1.222825 | 0.22076 | 0.114958 | 0.15404 | 0.715031 | 0.819707 | -0.541831 | 1.026835 | 0.110601 | -2.231689 | 0.526168 | 0.212327 | 0.446513 | 0.550266 |
| 24.777778 | 0.896857 | 0.903701 | 0.099518 | -0.512959 | -0.645385 | -0.151617 | 0.155792 | -1.385552 | -0.771921 | 0.317709 | -2.255278 | 0.721888 | -0.374848 | 0.429673 | 0.892636 |
| 24.791667 | 0.209693 | -0.955918 | -0.354379 | -0.69124 | -0.866299 | -0.193901 | -0.223612 | -0.605518 | -1.827995 | 0.323678 | 0.639152 | -1.139965 | -0.617132 | 0.330464 | 0.903415 |
| 24.805556 | -1.305211 | -2.001661 | -0.104974 | -0.020712 | -0.704324 | -0.185998 | -0.750624 | 1.045221 | 0.250897 | -0.27825 | 1.658248 | -0.765591 | -0.507898 | 0.220226 | 0.38449 |
| 24.819444 | -1.217876 | -0.700425 | -0.080482 | 0.044025 | -0.589357 | 1.083002 | 0.742838 | 0.68648 | 1.102538 | -0.89063 | -0.036475 | 1.368687 | -1.061456 | 0.080847 | -0.359578 |
| 24.833333 | -0.120305 | 0.684033 | -0.096144 | -0.358799 | -0.467735 | 0.582447 | -0.36546 | -0.691633 | -0.238126 | -0.823172 | -0.08484 | 0.830329 | -0.530304 | -0.10208 | -0.983645 |
| 24.847222 | -0.604533 | -0.731791 | -1.026026 | -1.095556 | 0.017269 | -0.785883 | -0.768574 | -1.96155 | -1.783108 | -0.096236 | 1.297371 | 1.485648 | -1.111602 | -0.268159 | -1.138596 |
| 24.861111 | -2.024832 | -2.441126 | -0.824317 | -0.759253 | 0.083962 | -1.150697 | -0.095508 | -1.782267 | -1.853664 | -0.023467 | 1.177693 | 0.682569 | -2.165716 | -0.335628 | -1.132276 |
| 24.875 | -2.122896 | -1.746691 | -0.372174 | 0.043598 | -0.452552 | -0.74577 | -0.458538 | -0.338823 | -0.778027 | -1.080129 | -0.318632 | -1.163522 | -1.822634 | -0.308242 | -1.239441 |
| 24.888889 | -1.37308 | -0.101172 | 0.006063 | 0.116479 | -0.762344 | -0.579025 | -2.144229 | -0.45852 | 0.120067 | -1.549788 | -2.222731 | 0.912068 | -1.676392 | -0.41991 | -1.638692 |
| 24.902778 | -2.157739 | -1.303472 | 0.282079 | -0.134641 | -0.709309 | -1.173245 | -3.980037 | -1.98632 | -0.857387 | -1.230232 | -1.49188 | -1.035934 | -1.800964 | -0.704933 | -2.150181 |
| 24.916667 | -2.544157 | -2.326051 | -1.34402 | -1.248331 | -0.959271 | -1.842186 | -3.117204 | -3.401899 | -1.72675 | -1.435495 | 0.79207 | -2.82403 | -2.320594 | -1.007373 | -2.204309 |
| 24.930556 | -1.77576 | -2.444627 | -3.012704 | -2.072423 | -1.748839 | -1.965707 | -1.510113 | -3.540747 | -2.262645 | -2.236072 | -0.518771 | -1.730693 | -3.026274 | -1.0463 | -2.018747 |
| 24.944444 | -1.437943 | -1.48938 | -3.189736 | -2.281652 | -1.862946 | -1.1277 | -0.434064 | -2.923116 | -2.687202 | -2.773121 | -3.577546 | -1.138362 | -3.227993 | -0.732491 | -1.485857 |
| 24.958333 | -2.19628 | -1.797389 | -2.135182 | -2.292549 | -2.122928 | -1.065241 | -1.171342 | -2.232018 | -2.906391 | -2.799453 | -4.113305 | -1.122873 | -2.388266 | -0.277493 | -0.696963 |
| 24.972222 | -3.415815 | -2.329182 | -1.162774 | -1.884217 | -1.762511 | -1.495593 | -2.746684 | -1.947593 | -2.436398 | -1.918848 | -1.782674 | -1.246438 | -1.580373 | -0.260123 | -0.518827 |
| 24.986111 | -3.735937 | -1.971994 | -0.900607 | -1.422961 | -1.155609 | -1.884885 | -3.919688 | -1.644735 | -1.496016 | -1.08283 | -0.057181 | -1.0861 | -1.278911 | -0.51394 | -1.018479 |
| 25 | -1.625272 | -1.102149 | -1.140739 | -1.08816 | -0.591019 | -1.906297 | -4.147011 | -1.227199 | -1.047983 | -0.44318 | -0.274676 | -1.04316 | -1.089799 | -0.481709 | -1.113071 |
| 25.013889 | -1.161209 | -1.171074 | -1.337768 | -1.181337 | -0.471885 | -2.039102 | -4.163464 | -1.126034 | -1.34131 | -0.758974 | -1.709724 | -1.322159 | -1.446787 | -0.337787 | -0.878022 |
| 25.027778 | -2.128156 | -2.318488 | -1.571855 | -1.813175 | -1.083845 | -2.537973 | -4.316145 | -1.584474 | -2.168843 | -2.011761 | -3.078524 | -1.73659 | -2.233412 | -0.363975 | -0.875516 |
| 25.041667 | -3.086854 | -2.632396 | -1.971589 | -2.131583 | -1.751115 | -2.637366 | -3.643311 | -2.137461 | -2.647742 | -2.744934 | -3.081696 | -1.798754 | -2.565248 | -0.52558 | -1.091721 |
| 25.055556 | -2.877348 | -2.04896 | -2.294255 | -1.751916 | -1.650856 | -1.834257 | -2.28437 | -2.105505 | -2.162223 | -2.085378 | -2.122119 | -1.445345 | -2.093805 | -0.622488 | -1.189526 |
| 25.069444 | -1.735368 | -1.442864 | -2.239158 | -1.225088 | -1.13021 | -1.182174 | -1.314105 | -1.731732 | -1.501747 | -1.138926 | -1.592744 | -1.110508 | -1.51233 | -0.635345 | -1.123504 |
| 25.083333 | -0.825304 | -1.007903 | -1.663534 | -0.971424 | -0.827544 | -0.980762 | -0.827699 | -1.307534 | -1.166803 | -0.7033 | -1.420066 | -0.928309 | -1.126659 | -0.646059 | -1.059589 |
| 25.097222 | -0.588654 | -0.40644 | -1.235023 | -0.957399 | -0.834182 | -1.070723 | -0.833511 | -0.946739 | -1.061795 | -0.659431 | -1.400579 | -0.849712 | -0.99426 | -0.672942 | -1.106656 |
| 25.111111 | -1.066294 | -0.289753 | -1.088073 | -0.924473 | -0.845504 | -1.198846 | -1.075668 | -0.783606 | -0.886188 | -0.616874 | -1.40124 | -0.778777 | -1.034516 | -0.748926 | -1.235806 |
| 25.125 | -1.443878 | -0.720636 | -0.98152 | -0.711574 | -0.644819 | -1.135854 | -1.171798 | -0.82895 | -0.684574 | -0.480418 | -1.236507 | -0.706319 | -1.026886 | -0.838795 | -1.342817 |
| 25.138889 | -1.165169 | -1.012959 | -0.728034 | -0.477492 | -0.44468 | -1.009004 | -1.111705 | -0.876772 | -0.638118 | -0.393908 | -0.98428 | -0.6787 | -0.852514 | -0.971672 | -1.430435 |
| 25.152778 | -0.674105 | -0.757412 | -0.461946 | -0.44647 | -0.479947 | -0.912892 | -0.956133 | -0.803634 | -0.615425 | -0.373335 | -0.814235 | -0.685833 | -0.624413 | -1.043168 | -1.335305 |
| 25.166667 | -0.37459 | -0.378982 | -0.470738 | -0.594601 | -0.537456 | -0.811378 | -0.696363 | -0.640582 | -0.469395 | -0.359783 | -0.788136 | -0.652667 | -0.522632 | -0.93869 | -1.137148 |
| 25.180556 | -0.441377 | -0.456739 | -0.755333 | -0.618107 | -0.390139 | -0.712098 | -0.516147 | -0.567271 | -0.360229 | -0.371496 | -0.83595 | -0.619717 | -0.593342 | -0.748799 | -0.828139 |
| 25.194444 | -0.5498 | -0.622332 | -0.903509 | -0.464897 | -0.245775 | -0.685308 | -0.506526 | -0.630508 | -0.395032 | -0.396038 | -0.868725 | -0.623473 | -0.647259 | -0.609835 | -0.564435 |
| 25.208333 | -0.436093 | -0.461321 | -0.731255 | -0.405595 | -0.310433 | -0.649381 | -0.545136 | -0.620724 | -0.460891 | -0.368427 | -0.71462 | -0.59387 | -0.584027 | -0.571391 | -0.41625 |
| 25.222222 | -0.234506 | -0.246746 | -0.481444 | -0.43994 | -0.359847 | -0.473189 | -0.408439 | -0.390446 | -0.456947 | -0.319981 | -0.377619 | -0.449395 | -0.447678 | -0.508654 | -0.327017 |
| 25.236111 | -0.160103 | -0.187753 | -0.337857 | -0.304508 | -0.179674 | -0.186789 | -0.072361 | -0.141699 | -0.333478 | -0.259191 | -0.15074 | -0.260611 | -0.30364 | -0.349249 | -0.170403 |
| 25.25 | -0.157127 | -0.03385 | -0.230598 | 0.00033 | 0.05285 | -0.016959 | 0.173213 | -0.042656 | -0.188046 | -0.161828 | -0.074009 | -0.150306 | -0.199308 | -0.166607 | 0.081471 |
| 25.263889 | -0.150415 | 0.171212 | -0.064422 | 0.118808 | 0.054845 | -0.036077 | 0.104952 | 0.01102 | -0.134364 | -0.077859 | 0.099788 | -0.084495 | -0.157567 | -0.094632 | 0.220999 |
| 25.277778 | -0.239345 | 0.027218 | 0.065926 | -0.031108 | -0.096843 | -0.096001 | -0.105447 | -0.020259 | -0.129243 | -0.086713 | 0.282299 | -0.060355 | -0.144953 | -0.053246 | 0.215969 |
| 25.291667 | -0.220202 | -0.141015 | 0.008804 | -0.080618 | -0.074583 | -0.028854 | -0.078373 | -0.093949 | -0.074165 | -0.110951 | 0.324137 | -0.054454 | -0.078743 | 0.176345 | 0.303102 |
| 25.305556 | 0.122925 | 0.044775 | -0.042144 | 0.112412 | 0.096177 | 0.155589 | 0.224469 | -0.048603 | 0.019741 | -0.044217 | 0.345357 | 0.033677 | 0.068681 | 0.539454 | 0.577886 |
| 25.319444 | 0.457312 | 0.279422 | 0.160306 | 0.251485 | 0.081764 | 0.291435 | 0.527225 | 0.073471 | 0.09202 | 0.042751 | 0.394015 | 0.127332 | 0.206214 | 0.781862 | 0.884767 |
| 25.333333 | 0.514878 | 0.371204 | 0.55474 | 0.290447 | 0.022966 | 0.43068 | 0.785779 | 0.197895 | 0.218232 | 0.14004 | 0.497214 | 0.192127 | 0.313644 | 0.811483 | 1.048002 |
| 25.347222 | 0.480414 | 0.571283 | 0.73093 | 0.485582 | 0.236454 | 0.636924 | 1.107786 | 0.26798 | 0.400609 | 0.265962 | 0.577829 | 0.300568 | 0.424606 | 0.827485 | 1.107193 |
| 25.361111 | 0.688285 | 0.993982 | 0.68243 | 0.788518 | 0.559695 | 0.820874 | 1.472405 | 0.365026 | 0.531322 | 0.375776 | 0.651551 | 0.421219 | 0.593154 | 1.144153 | 1.515718 |
| 25.375 | 1.177415 | 1.378001 | 0.90753 | 1.072839 | 0.716948 | 1.009577 | 1.846033 | 0.701551 | 0.756115 | 0.57437 | 0.91495 | 0.60262 | 0.913328 | 1.700462 | 2.133885 |
| 25.388889 | 1.778789 | 1.615569 | 1.581639 | 1.425459 | 0.846467 | 1.302866 | 2.216318 | 1.221043 | 1.302349 | 0.958002 | 1.383327 | 1.039401 | 1.395034 | 2.007244 | 2.760995 |
| 25.402778 | 2.376175 | 1.993954 | 2.13051 | 1.87279 | 1.36944 | 1.680678 | 2.542082 | 1.725228 | 2.153122 | 1.573909 | 2.030311 | 1.611511 | 1.936561 | 2.141036 | 3.100135 |
| 25.416667 | 3.128565 | 2.744718 | 2.310611 | 2.358068 | 2.06596 | 2.072597 | 3.037593 | 2.293323 | 2.976617 | 2.284815 | 2.732736 | 2.150884 | 2.603695 | 1.97323 | 3.194481 |
| 25.430556 | 3.89977 | 3.393835 | 2.708201 | 2.764352 | 2.318547 | 2.512795 | 3.993837 | 3.084514 | 3.710402 | 3.101336 | 3.339888 | 2.590949 | 3.330118 | 2.030111 | 4.48558 |
| 25.444444 | 4.189271 | 3.611964 | 3.506485 | 2.94059 | 2.039636 | 2.893671 | 5.000992 | 3.835848 | 4.221794 | 3.273015 | 3.674586 | 2.528079 | 3.415456 | 2.000514 | 4.501515 |
| 25.458333 | 3.652024 | 3.254686 | 3.69056 | 2.469481 | 1.485554 | 2.378116 | 3.917705 | 3.50709 | 4.259657 | 2.469741 | 3.492859 | 2.488559 | 2.471854 | 1.40843 | 2.254885 |
| 25.472222 | 2.979801 | 2.746119 | 3.098463 | 1.829295 | 1.274269 | 1.651921 | 1.969126 | 2.958346 | 3.929478 | 1.57936 | 3.477336 | 2.929387 | 1.874636 | 0.94607 | 0.744642 |
| 25.486111 | 2.989544 | 2.758377 | 2.567646 | 1.85849 | 1.4871 | 1.652811 | 1.055101 | 3.196867 | 3.609009 | 1.700283 | 4.042591 | 3.253499 | 2.666064 | 1.286394 | 1.21476 |
| 25.5 | 2.668996 | 2.547874 | 2.279581 | 2.169264 | 1.521911 | 1.892682 | 1.241381 | 3.306235 | 3.26038 | 3.138698 | 4.78837 | 2.664369 | 3.807545 | 1.811867 | 1.826852 |
| 25.513889 | 1.965344 | 2.958845 | 2.406865 | 2.221931 | 1.363034 | 1.996198 | 2.321513 | 3.070649 | 3.15614 | 3.60977 | 4.500935 | 2.115305 | 4.468273 | 1.773214 | 2.189556 |
| 25.527778 | 2.703011 | 4.74477 | 2.971805 | 2.687765 | 1.106009 | 2.436999 | 3.486248 | 2.042344 | 2.73595 | 3.387983 | 2.569522 | 2.177402 | 4.289419 | 0.95415 | 1.882839 |
| 25.541667 | 3.066178 | 5.670417 | 2.358944 | 2.057419 | 1.051994 | 2.690437 | 3.628081 | 0.734877 | 1.873283 | 1.048889 | 1.03685 | 1.934095 | 3.409452 | 0.174832 | 0.737619 |
| 25.555556 | 2.403068 | 3.923637 | 1.03821 | 0.457367 | 0.666428 | 0.666429 | 0.846826 | 1.053865 | 2.383425 | -0.2261 | -0.157534 | -0.068729 | 1.151975 | -0.480142 | -0.927694 |
| 25.569444 | 0.9131 | 0.544547 | 0.390548 | 0.070678 | 0.294408 | -0.464152 | -0.771175 | 0.789474 | 0.790385 | 0.01789 | -0.117983 | -1.785075 | -0.743601 | 0.232934 | -0.024851 |
| 25.583333 | 0.32055 | -0.401292 | 0.457214 | 0.468217 | 0.16696 | -0.098977 | 0.040522 | 0.614985 | -0.048465 | 0.812615 | 0.552922 | -2.863934 | -0.073055 | 1.286368 | 1.957764 |
| 25.597222 | 0.502344 | 1.051233 | 0.943845 | 1.436545 | 0.357363 | 0.531049 | 1.3367 | 0.500382 | 0.829838 | 1.51924 | 1.098833 | -0.82229 | 1.530196 | 1.265284 | 2.307547 |
| 25.611111 | 0.540531 | 1.608806 | 0.993955 | 1.260659 | 0.202436 | 1.500188 | 2.514598 | 0.133919 | 0.67049 | 0.593058 | 0.10127 | 1.280632 | 1.602248 | 0.469091 | 0.389749 |
| 25.625 | -0.244344 | 0.011133 | 0.489625 | 0.274604 | -0.280892 | 0.283712 | 0.072103 | 0.916811 | 0.579545 | -1.297906 | -0.735894 | -0.387171 | 0.216947 | -0.444626 | -0.755424 |
| 25.638889 | -0.236551 | -0.771168 | 0.022094 | -0.210284 | -0.026541 | -1.259778 | -2.27928 | 0.433035 | -0.787488 | -1.263073 | -0.617006 | -2.134635 | -2.016452 | -0.463526 | -1.117382 |
| 25.652778 | -0.194817 | -1.214866 | -0.383427 | -0.448641 | -0.219477 | -1.073676 | -1.568257 | -0.621758 | -2.066069 | 0.225387 | -0.123905 | -2.864194 | -1.858737 | 0.091822 | -0.035224 |
| 25.666667 | 0.243121 | 0.234387 | 0.094865 | 0.102411 | 0.215381 | 0.206291 | -0.124626 | -0.873659 | -1.19196 | 1.438037 | 0.481612 | -1.357471 | -0.552945 | 0.893704 | 1.62117 |
| 25.680556 | 0.104032 | 1.048874 | 0.044157 | 0.358172 | 0.052731 | 1.037267 | 0.845911 | -0.431478 | -0.183119 | 1.241105 | 0.680278 | -0.986134 | 0.147297 | 1.129612 | 0.66784 |
| 25.694444 | -0.466105 | -0.189664 | 0.153872 | 0.250514 | -0.00517 | 0.466798 | 1.057971 | 1.014098 | 1.529452 | 1.001819 | 1.68857 | 1.174584 | 0.799348 | 0.563181 | 0.452594 |
| 25.708333 | 0.292991 | 0.174335 | 0.319928 | -0.062099 | -0.089605 | -0.787711 | -1.098887 | 1.270875 | 1.638002 | 0.989288 | 1.22457 | 2.559296 | 0.875803 | -0.120158 | -0.068588 |

| | | | | | | | | | |
|---|---|---|---|---|---|---|---|---|---|
| 27.930556 | -1.385129 | -2.493917 | 0.384409 | -0.193154 | -1.786612 | -1.947468 | -2.687682 | -2.76048 | -4.159368 |
| 27.944444 | -2.140148 | -2.119334 | -1.131017 | -1.619508 | -2.416498 | -1.107709 | -1.782667 | -4.95102 | -6.175604 |
| 27.958333 | -1.659976 | -1.651047 | -3.139882 | -3.741857 | -2.651002 | -1.751161 | -2.079632 | -4.277141 | -5.129501 |
| 27.972222 | -1.675896 | -2.115277 | -3.724367 | -4.088112 | -3.605951 | -2.27177 | -1.512566 | -2.533071 | -2.822495 |
| 27.986111 | -1.824602 | -1.955563 | -3.751732 | -3.332055 | -2.765908 | -1.77744 | -0.967779 | -0.855363 | -0.935594 |
| 28 | -1.253525 | -0.468279 | -3.06164 | -1.654221 | -0.247712 | -0.686619 | -0.505164 | 0.215822 | 0.061614 |
| 28.013889 | -1.346797 | -0.380938 | -2.194318 | -0.850426 | 0.922677 | -0.333442 | -0.568495 | 0.270838 | 0.158932 |
| 28.027778 | -1.810714 | -1.787474 | -2.602828 | -2.081286 | -0.597612 | -0.808843 | -1.341641 | -0.767254 | -0.92237 |
| 28.041667 | -2.344998 | -3.2426 | -3.439606 | -3.3081 | -2.604319 | -1.814805 | -2.301702 | -1.623042 | -2.051641 |
| 28.055556 | -2.461105 | -3.148505 | -3.852729 | -3.269928 | -3.348695 | -2.067047 | -2.536457 | -1.537915 | -1.984481 |
| 28.069444 | -1.774676 | -2.030874 | -3.227118 | -2.089463 | -2.19918 | -1.913104 | -2.242968 | -1.250436 | -1.460807 |
| 28.083333 | -1.189739 | -1.264073 | -2.500081 | -1.281799 | -1.051302 | -1.466773 | -1.58046 | -1.222673 | -1.084577 |
| 28.097222 | -0.94913 | -1.085275 | -2.068186 | -1.305687 | -1.071619 | -0.896937 | -0.856649 | -1.188469 | -1.126146 |
| 28.111111 | -0.760743 | -0.911263 | -1.872898 | -1.540614 | -1.55194 | -0.658162 | -0.586276 | -0.904016 | -1.372628 |
| 28.125 | -0.596864 | -0.628287 | -1.665487 | -1.177234 | -1.015759 | -0.812157 | -0.838216 | -0.752006 | -1.280301 |
| 28.138889 | -0.615896 | -0.415103 | -1.358122 | -0.783104 | -0.335753 | -0.897742 | -1.183161 | -0.829905 | -0.870187 |
| 28.152778 | -0.637054 | -0.329315 | -0.942502 | -0.803938 | -0.31333 | -0.642545 | -0.996182 | -0.832151 | -0.570919 |
| 28.166667 | -0.503599 | -0.230576 | -0.499053 | -0.822195 | -0.56388 | -0.360991 | -0.565494 | -0.588907 | -0.544177 |
| 28.180556 | -0.373676 | -0.150265 | -0.224037 | -0.576235 | -0.461554 | -0.336044 | -0.415043 | -0.364253 | -0.525885 |
| 28.194444 | -0.443245 | -0.22675 | -0.151455 | -0.296999 | -0.177019 | -0.335303 | -0.457025 | -0.305366 | -0.428751 |
| 28.208333 | -0.657922 | -0.409019 | -0.10238 | -0.185063 | -0.063226 | -0.175809 | -0.274576 | -0.236588 | -0.324808 |
| 28.222222 | -0.742862 | -0.439511 | 0.01179 | -0.066931 | -0.056808 | -0.071466 | -0.031698 | -0.041541 | -0.179118 |

| | | | | | | | | | | | | | | | | | | | | | | | | | | | | |
|---|---|---|---|---|---|---|---|---|---|---|---|---|---|---|---|---|---|---|---|---|---|---|---|---|---|---|---|---|
| 31.25 | -0.89675 | 0.268298 | 0.292673 | 0.255795 | 0.56453 | 0.280516 | 0.396997 | 1.072956 | 0.449068 | 0.105684 | -0.299434 | 0.12554 | 0.510537 | -1.640617 | -0.740606 | -0.64093 | -0.090908 | -0.168484 | -0.300566 | -0.702325 | -0.375844 | -0.38047 | -0.310527 | -0.609787 | | 31.34722222 | 4.388785 | 0.756095 |
| 31.263889 | -0.122951 | 0.437347 | 0.569544 | 0.614431 | 0.443646 | 0.725717 | 0.382643 | 0.98237 | 0.944455 | 0.377017 | 0.342202 | 0.091253 | 1.322902 | -0.59618 | -0.051078 | -0.189113 | -0.003595 | -0.088403 | -0.174303 | -0.466555 | -0.152966 | -0.201103 | -0.143365 | -0.049761 | | 31.36111111 | 3.773801 | 0.434622 |
| 31.277778 | 0.954704 | 0.773869 | 1.028644 | 0.841958 | 0.612679 | 0.686755 | 0.421067 | 0.503113 | 1.111855 | 0.936603 | 0.65145 | 0.055523 | 1.609773 | 0.681503 | 0.569639 | 0.110511 | -0.01519 | -0.020019 | -0.123038 | -0.22187 | 0.07128 | -0.043077 | -0.078451 | 0.266212 | | 31.375 | 1.549985 | 0.410192 |
| 31.291667 | 1.822087 | 1.377252 | 1.499018 | 0.910069 | 1.006708 | 0.63082 | 0.682488 | 0.739597 | 1.185489 | 1.115857 | 0.71245 | 0.384622 | 1.649132 | 1.953405 | 1.013172 | 0.263528 | 0.009944 | 0.071652 | -0.002595 | -0.004937 | 0.275057 | 0.051987 | -0.045669 | 0.484004 | | 31.38888889 | -1.448155 | 0.583686 |
| 31.305556 | 2.112187 | 1.809498 | 1.800967 | 1.095607 | 1.305508 | 0.981795 | 1.079065 | 1.59251 | 1.526971 | 1.070118 | 0.887108 | 0.973862 | 1.937273 | 2.513437 | 1.269246 | 0.468327 | 0.136703 | 0.210839 | 0.291862 | 0.117923 | 0.40768 | 0.102821 | 0.072412 | 0.766154 | | 31.40277778 | -2.144878 | 0.756707 |
| 31.319444 | 1.955927 | 1.914704 | 2.224905 | 1.513793 | 1.569928 | 1.379068 | 1.225296 | 1.838208 | 2.252513 | 1.405685 | 1.000814 | 1.631923 | 2.512434 | 2.775202 | 1.818596 | 0.779506 | 0.37955 | 0.455536 | 0.719853 | 0.231365 | 0.526948 | 0.256774 | 0.410744 | 1.174793 | | 31.41666667 | -0.230978 | 0.776154 |
| 31.333333 | 1.746856 | 2.022062 | 2.67423 | 1.999129 | 1.961469 | 1.541938 | 1.538871 | 1.765512 | 2.822014 | 1.999679 | 1.157248 | 2.425924 | 3.058125 | 3.306574 | 2.566349 | 1.116888 | 0.743055 | 0.740458 | 1.12511 | 0.494436 | 0.776216 | 0.620349 | 0.884552 | 1.873049 | | 31.43055556 | 2.963924 | 0.761328 |
| 31.347222 | 1.859478 | 2.238886 | 3.155958 | 2.335382 | 2.283317 | 1.829558 | 2.17938 | 1.967125 | 3.090195 | 2.325225 | 1.507604 | 3.296646 | 3.608163 | 4.105111 | 3.185713 | 1.307663 | 1.102992 | 0.91093 | 1.33025 | 0.883802 | 1.170458 | 0.987283 | 1.211722 | 2.53674 | | 31.44444444 | 5.0604 | 0.776643 |
| 31.361111 | 2.343619 | 2.731641 | 3.723858 | 2.884347 | 2.799265 | 2.581189 | 2.913873 | 2.5551 | 3.656191 | 2.817856 | 2.279191 | 3.822437 | 4.55162 | 4.524699 | 3.727685 | 1.526738 | 1.506744 | 1.033002 | 1.428711 | 1.283021 | 1.629698 | 1.206061 | 1.40113 | 2.887446 | | 31.45833333 | 4.522083 | 0.740028 |
| 31.375 | 3.037881 | 3.705337 | 4.774626 | 3.952084 | 3.73958 | 3.611931 | 3.741069 | 3.037839 | 4.488323 | 3.805108 | 2.95398 | 4.275875 | 5.960005 | 5.048197 | 4.413065 | 1.927064 | 1.933842 | 1.369548 | 1.838175 | 1.863267 | 2.163291 | 1.463898 | 1.891105 | 3.271667 | | 31.47222222 | 2.491862 | 0.888369 |
| 31.388889 | 3.943167 | 4.811393 | 6.397878 | 5.79739 | 4.8527 | 4.389106 | 4.488268 | 3.36576 | 5.336051 | 5.148257 | 3.78405 | 4.553754 | 7.727419 | 5.614139 | 5.372602 | 2.523874 | 2.444419 | 1.987451 | 2.781851 | 2.645716 | 2.772146 | 1.923511 | 2.619348 | 4.529128 | | 31.48611111 | 0.987383 | 1.300825 |
| 31.402778 | 4.912207 | 5.825031 | 7.510215 | 7.590611 | 5.671042 | 4.374383 | 4.505494 | 3.709123 | 6.330828 | 6.018912 | 5.002555 | 4.703939 | 9.459664 | 6.845005 | 6.64793 | 3.125052 | 3.325786 | 2.36948 | 3.634888 | 3.152878 | 3.328485 | 2.364988 | 3.22852 | 6.332047 | | 31.5 | 0.926992 | 1.530511 |
| 31.416667 | 5.490625 | 6.682095 | 7.24295 | 7.468664 | 5.530381 | 4.197912 | 4.919676 | 4.322282 | 6.725273 | 6.719114 | 6.348123 | 5.290486 | 10.869273 | 8.147069 | 8.35501 | 3.543 | 3.97336 | 2.338523 | 3.913132 | 3.397851 | 3.717381 | 2.782766 | 3.774435 | 8.049885 | | 31.51388889 | 2.283175 | 1.525502 |
| 31.430556 | 5.269907 | 7.274382 | 6.746746 | 6.869022 | 5.445936 | 4.746125 | 5.97903 | 4.922931 | 6.766237 | 7.264749 | 7.284641 | 6.053569 | 11.063353 | 8.634502 | 10.21222 | 4.049605 | 4.427174 | 2.066064 | 3.777771 | 3.51311 | 4.226031 | 3.272778 | 4.33182 | 9.086188 | | 31.52777778 | 4.062317 | 1.407558 |
| 31.444444 | 3.793279 | 6.417788 | 5.537901 | 5.785722 | 6.172574 | 4.95657 | 5.467825 | 5.352759 | 7.055037 | 7.071772 | 7.764217 | 5.837171 | 10.674027 | 7.668878 | 10.742596 | 4.158966 | 4.62994 | 1.68295 | 3.635884 | 3.768781 | 4.788277 | 3.535513 | 4.939311 | 9.951984 | | 31.54166667 | 5.074501 | 1.279341 |
| 31.458333 | 3.246997 | 4.8479 | 3.202663 | 3.546735 | 3.943971 | 3.16606 | 2.914904 | 4.844052 | 6.217094 | 5.026831 | 6.189239 | 3.749757 | 6.919664 | 4.934861 | 6.952608 | 4.253795 | 3.68944 | 1.631354 | 3.465495 | 3.477219 | 4.569915 | 3.635009 | 4.723413 | 9.561666 | | 31.55555556 | 4.708018 | 0.87052 |
| 31.472222 | -0.208617 | 1.152169 | -0.00896 | 0.93693 | -0.101655 | -0.019724 | -0.36383 | 2.821756 | 2.563102 | 0.432778 | 1.654859 | 0.707303 | -0.966755 | -1.590941 | -1.16172 | 1.87534 | 1.713348 | 1.459924 | 3.07728 | 2.909157 | 2.913472 | 2.907398 | 3.036501 | 3.80044 | | 31.56944444 | 3.207424 | 0.263871 |
| 31.486111 | -2.176991 | -0.79915 | -2.0782 | -0.851347 | -1.274315 | -1.257398 | -0.443138 | 1.815965 | -0.222492 | -1.430508 | -2.083292 | -0.228177 | -3.505612 | -6.162828 | -6.444982 | -1.031918 | 0.605218 | 1.353339 | 2.707679 | 1.798176 | 0.177241 | 1.056634 | 1.177374 | -3.967615 | | 31.58333333 | -0.246137 | 0.004406 |
| 31.5 | 0.257147 | 1.529944 | 0.46883 | 0.260625 | 0.206846 | 0.882805 | 2.604141 | 3.0376 | 2.087732 | 2.060511 | 0.914857 | 1.845411 | 4.740083 | -4.801232 | -3.961099 | -1.322739 | 0.131927 | 0.867556 | 1.711011 | 1.112602 | -0.106406 | -0.082 | 0.349498 | -5.933791 | | 31.59722222 | -2.728486 | 0.1251 |
| 31.513889 | 3.817278 | 5.661354 | 1.74936 | 3.533331 | 2.718815 | 4.18807 | 5.23032 | 4.394736 | 5.481231 | 4.911681 | 5.328703 | 2.195445 | 14.907538 | -0.828573 | 0.268262 | -0.973489 | 0.507995 | 0.618759 | 0.840471 | 0.949324 | 1.291217 | -0.726893 | 0.138541 | -0.507617 | | 31.61111111 | -1.991589 | -0.114627 |
| 31.527778 | 4.560833 | 8.593902 | 4.289953 | 4.618421 | 3.424508 | 4.297111 | 4.763894 | 4.582079 | 8.175526 | 8.178749 | 7.145933 | 2.722413 | 20.279793 | 2.5661 | 3.472496 | 0.451149 | 0.844972 | 0.715774 | 0.932558 | 0.497682 | 2.113032 | -0.722027 | 0.03722 | 5.854749 | | 31.625 | 0.142015 | -0.53389 |
| 31.541667 | 2.728369 | 6.879908 | 2.610866 | 3.553959 | 3.937228 | 3.288391 | 2.914642 | 2.999781 | 7.570445 | 6.717999 | 5.425455 | 1.903263 | 14.052263 | 4.412155 | 5.250712 | 1.490021 | 0.997134 | 0.588601 | 0.953826 | 0.471824 | 1.899822 | 0.31198 | 0.803328 | 9.203241 | | 31.63888889 | 2.851679 | -0.763736 |
| 31.555556 | 2.181655 | 5.558964 | 1.444676 | 1.572655 | 2.806269 | 2.307903 | 1.287378 | 1.184017 | 3.749424 | 1.978017 | 2.684494 | -0.699559 | 2.13178 | 3.381738 | 5.511194 | 1.820624 | 1.184738 | 0.193861 | 0.482732 | 0.51953 | 1.965256 | 1.381919 | 1.408534 | 7.084031 | | 31.65277778 | 4.594795 | -0.744942 |
| 31.569444 | 1.198283 | 2.484001 | 0.667466 | -0.782885 | -1.591787 | 0.649074 | -0.159446 | -1.339566 | -0.903762 | -3.182816 | 2.205487 | -0.770912 | -4.527768 | 2.191309 | 4.53989 | 2.756666 | 0.94872 | 0.149925 | -0.127884 | 0.679525 | 0.598557 | 2.231736 | 2.745115 | 4.684377 | | 31.66666667 | 4.093244 | -0.301688 |
| 31.583333 | 0.417547 | -0.548839 | 0.339637 | -1.237684 | -1.680537 | -0.263844 | -0.152471 | -1.434889 | -3.055814 | -4.531712 | 1.267304 | 0.010844 | -3.704589 | 0.647523 | 1.894355 | 2.358578 | 0.425304 | 0.189003 | 0.178943 | 0.650123 | -1.156524 | 2.004526 | 2.450705 | 1.949375 | | 31.68055556 | 3.177323 | 0.182588 |
| 31.597222 | 0.060933 | -0.76038 | 0.322832 | -0.859357 | -1.804614 | -0.578162 | -0.506247 | 1.18829 | -0.656113 | -0.35781 | -0.787662 | 2.445626 | 1.718167 | -0.478288 | -0.852084 | 0.879132 | 0.164125 | 0.385505 | 1.072051 | 1.098754 | 0.278471 | 1.583207 | 1.347797 | 0.094262 | | 31.69444444 | 2.226656 | 0.089992 |
| 31.611111 | -0.223201 | 0.696677 | 0.486964 | 0.632061 | -0.25903 | -0.113403 | -0.212827 | 0.661635 | 1.917916 | 3.015354 | -0.77269 | 2.204475 | 5.478099 | -2.38124 | -3.639862 | -0.460346 | 0.080168 | 0.437399 | 0.891301 | 1.207523 | 1.320768 | 0.3523 | 0.50745 | -2.899357 | | 31.70833333 | 1.330336 | -0.336559 |
| 31.625 | -0.354329 | 0.471712 | 0.549232 | 2.097481 | 1.344062 | 0.075631 | 0.857561 | -0.650937 | 1.942472 | 3.283557 | 0.227356 | 1.272831 | 3.555641 | -2.191461 | -2.986205 | -1.541456 | -0.211509 | -0.488862 | -0.419188 | 0.413247 | 0.971863 | -0.361897 | -0.313991 | -4.663376 | | 31.72222222 | 0.78686 | -0.610712 |
| 31.638889 | -0.412853 | -0.151885 | -0.067042 | 0.157178 | 0.686493 | -0.229598 | -0.11497 | 0.330146 | 0.84232 | 1.191496 | 2.663004 | 0.006063 | -2.227804 | -1.330377 | -0.98169 | -1.62174 | 0.316408 | -0.220431 | -0.575481 | -0.648798 | 0.272175 | -1.196796 | -1.42521 | -3.684846 | | 31.73611111 | -0.525784 | -0.611402 |
| 31.652778 | -0.396089 | -1.991653 | -0.348641 | -1.238197 | -1.855037 | -0.368046 | -0.768974 | 1.804705 | 1.207767 | -2.242582 | 2.92276 | 0.283056 | -6.747472 | 1.006626 | 2.281197 | -0.62356 | 0.906214 | -0.129436 | -0.858808 | -1.198645 | 0.697998 | -1.385782 | -0.895862 | -1.277075 | | 31.75 | -0.658584 | -0.595697 |
| 31.666667 | -0.155981 | -2.251968 | 0.311637 | -1.43384 | -2.430683 | -0.237892 | -0.822257 | 1.331872 | -0.629779 | -1.034563 | -1.130991 | 1.414494 | -5.380424 | 2.502004 | 4.818748 | 1.599246 | 1.255001 | -0.146415 | -0.469457 | -0.980302 | 0.443662 | -0.460342 | 0.780691 | 1.643851 | | 31.76388889 | -0.480755 | -0.595787 |
| [illegible] | [illegible] | [illegible] | [illegible] | [illegible] | [illegible] | [illegible] | [illegible] | [illegible] | [illegible] | [illegible] | [illegible] | [illegible] | [illegible] | [illegible] | [illegible] | [illegible] | [illegible] | [illegible] | [illegible] | [illegible] | [illegible] | [illegible] | [illegible] | [illegible] | | [illegible] | [illegible] | [illegible] |

| | | | | | | | | | | | | | | | | | | | | | | | | | | | | |
|---|---|---|---|---|---|---|---|---|---|---|---|---|---|---|---|---|---|---|---|---|---|---|---|---|---|---|---|---|
| 34.569444 | 1.129615 | 3.911527 | 2.360222 | 3.798691 | 3.982751 | 1.588273 | 3.114673 | 4.112648 | 3.194596 | -1.277655 | 8.73324 | 2.449617 | 6.452117 | 6.088251 | 5.045141 | -0.408917 | 0.210016 | 0.297188 | 0.687036 | 1.143304 | 0.766533 | 1.205505 | 0.393377 | 6.445702 | | 34.66666667 | -1.10393 | 0.802522 |
| 34.583333 | 0.850384 | 1.25865 | 2.346945 | 3.451801 | 2.215223 | 0.62478 | 0.772454 | 2.318568 | -0.848258 | -1.990167 | 6.167792 | 0.717015 | 1.989811 | 4.511204 | 4.521702 | 0.001204 | 0.13228 | 0.176797 | 0.343132 | 0.7011 | 0.865401 | 0.833313 | -0.614179 | 1.286858 | | 34.68055556 | 3.01991 | 0.721836 |
| 34.597222 | 0.442971 | -0.643281 | 1.775389 | 2.366826 | 1.494787 | 0.068915 | -1.219345 | -0.074195 | -0.585251 | -0.229329 | 1.477425 | 0.41945 | 0.709101 | 0.863091 | -0.498213 | 0.849347 | 0.001034 | 0.237447 | 0.048311 | -0.403475 | 0.335546 | 0.244316 | -1.943059 | -1.811346 | | 34.69444444 | 5.941992 | 0.841704 |
| 34.611111 | 0.506142 | -0.547085 | 0.791869 | 0.383001 | 0.526714 | 0.377851 | -1.813268 | -2.847927 | 0.403069 | 0.230048 | -4.092272 | -2.034782 | -1.873833 | -2.713011 | -3.390032 | -0.024698 | -0.428024 | 0.002205 | -0.093812 | -1.285152 | 0.380926 | 0.147152 | -0.729197 | -2.990275 | | 34.70833333 | 4.492715 | 0.397296 |
| 34.625 | 0.913286 | 1.275232 | -0.033559 | -1.680497 | -1.086367 | 0.631561 | -0.673729 | -2.755569 | 0.216572 | -2.313699 | -5.276965 | -1.213316 | -1.937521 | -4.267383 | -1.679473 | 0.459121 | -0.255791 | 0.323942 | 0.710241 | -1.09853 | -1.322015 | 0.644953 | 1.823919 | -3.1709 | | 34.72222222 | -0.221142 | -0.280004 |
| 34.638889 | 0.667967 | 1.755989 | -1.145206 | -3.02512 | -3.519553 | -0.070515 | 0.528233 | 0.623044 | 1.4954 | 0.301127 | -3.251828 | 0.769251 | 0.078449 | -4.762829 | -2.376634 | 1.131799 | 0.458765 | 0.446222 | 0.168145 | 0.436852 | -0.611942 | 0.504894 | 3.074051 | 0.62916 | | 34.73611111 | -3.084296 | -0.702204 |
| 34.652778 | -0.333935 | -0.103403 | -0.822739 | -1.713442 | -2.933682 | -1.49428 | 1.055709 | 2.361667 | 1.252681 | 1.043814 | -1.210651 | 1.160591 | 1.040461 | -4.332716 | -1.199616 | 1.894951 | 1.531818 | 0.309396 | 0.730113 | 1.438758 | 1.19778 | -0.026937 | 2.549375 | 3.552546 | | 34.75 | -0.741768 | -0.67246 |
| 34.666667 | -1.765715 | -2.618257 | 0.46427 | 0.283503 | -1.777661 | -1.625427 | 0.090321 | 1.184026 | -1.466702 | 2.140651 | 1.125035 | -0.969599 | -2.12016 | -0.032276 | -0.340663 | 2.197452 | 1.779763 | -0.07551 | 0.576296 | 1.220204 | 3.428726 | -0.680884 | 0.331997 | 2.771903 | | 34.76388889 | 3.574236 | -0.437479 |
| 34.680556 | -3.990925 | -5.704123 | 1.093268 | 2.022386 | 1.05559 | -0.441674 | 0.229449 | -0.088426 | -0.231559 | 0.364105 | 3.142945 | -3.110938 | -5.337384 | 3.506087 | 0.878901 | 1.211998 | 1.621488 | -0.624032 | -0.090922 | 0.099663 | 1.81484 | -1.139814 | -0.571269 | 1.397147 | | 34.77777778 | 6.467291 | -0.208458 |
| 34.694444 | -6.23375 | -9.171139 | 0.07686 | 1.232287 | 1.022658 | -0.001246 | -1.975749 | -1.503829 | 0.740789 | -0.826869 | 0.577744 | -2.429072 | -5.455843 | 5.186549 | 0.657808 | -0.035363 | 0.46417 | -0.514951 | -0.01464 | -0.538999 | 0.269135 | -0.973704 | -0.973602 | 0.375274 | | 34.79166667 | 8.326622 | -0.11456 |
| 34.708333 | -4.443576 | -7.180187 | -1.794806 | -1.079177 | -2.600676 | 1.060952 | -1.121799 | -2.217932 | -1.219671 | -3.618828 | -1.496356 | -1.809759 | -6.408333 | 3.324633 | 1.435054 | -1.868111 | -0.779392 | -0.177077 | -0.019556 | 0.117386 | 0.303153 | 0.259119 | -0.084594 | -0.405326 | | 34.80555556 | 5.166945 | -0.260912 |
| 34.722222 | -2.477349 | -5.22685 | -5.139895 | -2.421894 | -2.780777 | 0.639621 | 0.111538 | -1.841738 | -2.87286 | -4.896985 | -2.267796 | 0.197051 | -5.574331 | -0.403154 | 0.984168 | -2.838544 | -1.327336 | 0.217523 | 0.795008 | 0.514744 | 0.059059 | 1.425793 | 0.231793 | -1.378797 | | 34.81944444 | 1.5563 | -0.262925 |
| 34.736111 | 0.874431 | -0.054076 | -5.38273 | -3.149395 | -0.471441 | -1.224653 | 0.994223 | -2.594033 | -2.64369 | -0.793043 | -5.280067 | 2.686515 | -5.268682 | -1.798917 | -0.392978 | -1.942248 | -0.709028 | 0.497192 | 0.599212 | 0.658541 | -1.011686 | 1.606436 | 1.271078 | -2.282845 | | 34.83333333 | -1.129979 | -0.321207 |
| 34.75 | 2.53938 | 2.890391 | -1.707317 | -1.233093 | 1.248341 | -1.028762 | 0.764722 | -3.039296 | -1.731753 | 2.262344 | -5.254474 | 0.574635 | -2.16005 | -0.115405 | 0.254938 | -0.009165 | -0.199714 | 0.418155 | 0.296283 | 0.055172 | -1.105226 | 0.767858 | 0.187206 | -1.507667 | | 34.84722222 | -3.84707 | -0.531301 |
| 34.763889 | -0.642133 | -0.727679 | 2.462129 | 0.877917 | -0.454301 | 0.534909 | 0.25785 | 1.967457 | 1.259799 | 0.982813 | -6.15184 | -5.998543 | 0.700475 | 0.484375 | 1.002738 | 0.47791 | -0.655392 | -0.358323 | -1.331417 | -1.667783 | -2.087766 | -0.091816 | -0.759429 | -2.690819 | | 34.86111111 | -4.999066 | -0.922753 |
| 34.777778 | -4.682299 | -4.439367 | 4.179703 | 1.173964 | -3.136057 | 1.432994 | -0.077018 | 3.78292 | 2.443588 | -1.911698 | -7.163084 | -2.514371 | -0.684567 | 0.250132 | 0.772745 | 0.241029 | -0.943727 | -0.717 | -1.692006 | -2.089669 | -1.673039 | -0.890583 | -1.110916 | -3.98791 | | 34.875 | -3.40161 | -1.374753 |
| 34.791667 | -4.628984 | -4.303475 | 1.24388 | -0.125083 | -2.621351 | 1.936902 | 0.214604 | 3.002566 | -0.079184 | -3.017741 | 0.248947 | 2.448678 | -2.477123 | -1.489154 | -0.10615 | 0.084172 | -1.200732 | -0.017518 | 0.078694 | -1.776881 | -1.008757 | -1.194483 | -0.116351 | -2.793354 | | 34.88888889 | -0.76362 | -1.793687 |
| 34.805556 | -2.064294 | -1.923196 | -0.106248 | -0.648723 | -0.821994 | 1.991788 | 0.790241 | 2.137561 | -1.124517 | -1.636536 | 8.100932 | -4.016166 | -0.146665 | -2.780556 | -1.908916 | 1.044031 | 0.026147 | 0.354753 | 2.064659 | -1.352593 | -0.516207 | -0.906133 | 0.625222 | -0.526539 | | 34.90277778 | 0.37405 | -2.544179 |
| 34.819444 | -0.859099 | -0.555431 | -2.387706 | -0.974179 | 0.511984 | 0.849412 | 2.085319 | 0.3846 | -1.499925 | -0.500123 | 12.214995 | 1.816163 | 2.495425 | -4.12897 | -3.100699 | 1.427456 | 1.171054 | 0.722245 | 1.549676 | 0.490607 | 0.394614 | -0.830585 | 1.595946 | 1.226554 | | 34.91666667 | 0.595648 | -2.965316 |
| 34.833333 | -1.016922 | 0.568985 | -5.53245 | -1.044584 | 3.080358 | -0.737828 | 1.021531 | -2.619642 | -5.429692 | -2.618189 | 4.957982 | 2.940605 | -4.477788 | -4.627927 | -3.082365 | 0.054777 | -0.021322 | 0.138135 | -0.076438 | 1.394656 | 0.538211 | -1.42854 | 1.567924 | -1.160981 | | 34.93055556 | -0.512538 | -2.97511 |
| 34.847222 | -4.946914 | -1.796327 | -5.066118 | -1.791427 | 0.059073 | -2.782396 | -2.753086 | -6.874185 | -10.483789 | -2.682931 | -9.078135 | 1.183397 | -15.939153 | -5.131113 | -3.748024 | -1.572675 | -0.858277 | -0.73806 | -2.516769 | 0.561569 | 0.576549 | -1.492695 | -0.856738 | -6.374454 | | 34.94444444 | -3.112764 | -2.906785 |
| 34.861111 | -3.74283 | -2.203675 | -1.059934 | -2.062707 | -2.004163 | -3.86326 | -4.945569 | -5.080444 | -7.563848 | -1.33869 | -12.63716 | -1.621867 | -13.907353 | -5.530479 | -4.431382 | -2.319599 | -1.527487 | -0.857024 | -2.435499 | -1.143476 | 0.224528 | -1.229679 | -1.173588 | -10.318894 | | 34.95833333 | -6.489951 | -2.682148 |
| 34.875 | -1.470999 | -2.441902 | 3.375586 | -1.388403 | -3.028837 | -4.158092 | -5.016549 | 1.165946 | -1.917976 | -0.646067 | -0.95723 | -4.754166 | -6.47043 | -5.726281 | -4.407404 | -2.248449 | -2.137626 | 0.166059 | -0.555834 | -2.523054 | -0.392885 | -1.084783 | -2.286068 | -10.153537 | | 34.97222222 | -8.762678 | -2.430736 |
| 34.888889 | -2.650558 | -3.266701 | 3.081879 | -0.331927 | -2.543066 | -3.405395 | -3.727428 | 3.906176 | 2.429289 | -0.111551 | 0.355914 | -4.989212 | -1.666153 | -6.750383 | -4.02226 | -2.711725 | -2.247554 | -0.900862 | 0.587071 | -2.245063 | -1.056367 | -1.728034 | -2.386462 | -7.583607 | | 34.98611111 | -8.169484 | -2.148624 |
| 34.902778 | -7.916461 | -5.429138 | -0.664516 | -0.443442 | -0.806893 | -2.930153 | -2.82672 | 0.370602 | 1.491951 | -0.426642 | -2.595868 | -2.886193 | -3.141349 | -6.035086 | -2.463437 | -2.132046 | -1.884237 | 0.067891 | 0.959547 | -3.436701 | -2.978679 | -1.713298 | -2.875081 | -5.604921 | | 35 | -5.760156 | -1.871811 |
| 34.916667 | -13.223741 | -5.697486 | -4.859814 | -2.683611 | -1.27935 | -4.158033 | -4.353963 | -2.877236 | -1.785556 | -1.449431 | -7.713917 | -0.684605 | -10.790627 | -3.955837 | -1.88608 | -0.741875 | -0.488256 | 0.475583 | -0.063628 | -3.953291 | -3.848562 | -1.965126 | -3.512595 | -4.459988 | | 35.01388889 | -2.984709 | -1.49039 |
| 34.930556 | -11.19745 | -5.708989 | -6.516413 | -5.477745 | -2.120962 | -6.337928 | -7.986034 | -5.836973 | -5.919918 | -3.593232 | -9.890788 | -2.554808 | -17.728018 | -1.737023 | -2.52487 | -0.771084 | -0.871106 | -0.418528 | -2.357095 | -2.165578 | -2.979072 | -1.951771 | -3.576178 | -5.053957 | | 35.02777778 | -1.57073 | -0.990457 |
| 34.944444 | -8.277198 | -7.07688 | -7.611692 | -7.91052 | -4.375739 | -10.081841 | -10.684591 | -6.6141 | -7.572508 | -5.169197 | -11.480914 | -5.5808 | -19.766509 | -1.889014 | -3.982021 | -1.769804 | -2.343157 | -2.809529 | -4.513826 | 0.070597 | -2.339774 | -1.994601 | -3.315761 | -10.129467 | | 35.04166667 | -2.131693 | -0.754244 |
| 34.958333 | -6.438313 | -7.436759 | -6.857283 | -7.856295 | -5.517244 | -11.314509 | -9.01996 | -5.984864 | -5.892853 | -4.900885 | -10.134942 | -7.666898 | -17.744394 | -4.342728 | -5.009635 | -3.074074 | -3.292099 | -4.638085 | -7.408346 | -0.863041 | -3.304874 | -2.309839 | -3.457038 | -14.463628 | | 35.05555556 | -5.148354 | -1.095231 |
| 34.972222 | -7.808944 | -4.495104 | -6.870698 | -4.053183 | -1.581788 | -10.326087 | -5.511955 | -2.555686 | -4.533243 | -4.181733 | -11.270375 | -6.843711 | -12.622142 | -6.187369 | -4.69092 | -3.061968 | -3.310291 | -4.784697 | -7.326859 | -1.808949 | -3.496522 | -2.640275 | -3.092749 | -13.862785 | | 35.06944444 | -8.672078 | -1.659599 |
| 34.986111 | -10.779941 | -2.345231 | -7.412068 | -3.42678 | -1.04496 | -9.405465 | -1.49679 | -1.262227 | -3.647033 | -2.759122 | -0.742619 | -4.173202 | -8.242482 | -4.739702 | -3.236879 | -2.477636 | -2.510126 | -3.978454 | -4.813961 | -2.088968 | -2.678451 | -2.362346 | -2.725454 | -7.98199 | | 35.08333333 | -10.09945 | -1.440897 |
| 35 | -7.729194 | -1.712588 | -6.953682 | -6.578565 | -3.168819 | -8.223959 | -0.809842 | -1.00235 | -2.582807 | -1.649391 | -1.566037 | -2.918997 | -7.449927 | -3.430682 | -1.715689 | -1.912692 | -1.599787 | -2.198784 | -1.570397 | -1.796707 | -1.822601 | -1.84217 | -2.168425 | -3.215742 | | 35.09722222 | -8.483471 | -0.456763 |
| 35.013889 | -5.597638 | -2.892781 | -7.132762 | -8.560768 | -3.443503 | -6.012894 | -1.131824 | -2.467716 | -2.888873 | -1.349417 | -3.755518 | -2.906396 | -8.787482 | -2.363375 | -0.874356 | -1.561895 | -1.04897 | -1.213704 | 0.270663 | -1.787828 | -1.64907 | -1.336005 | -1.560036 | -1.273873 | | 35.11111111 | -5.315838 | 0.152679 |
| 35.027778 | -5.757035 | -2.386203 | -3.608219 | -1.808451 | -0.964598 | -3.086122 | -1.008275 | -3.872235 | -2.66312 | -1.156077 | -3.446167 | -2.807803 | -4.964311 | -1.512175 | -1.048768 | -1.294745 | -0.743712 | -0.974653 | 0.366078 | -1.74663 | -1.582542 | -1.050501 | -1.201353 | -1.003087 | | 35.125 | -2.844371 | 0.286124 |
| 35.041667 | -7.469202 | -2.173144 | -0.834247 | 1.508366 | 0.849135 | -0.003686 | 0.08644 | -2.896692 | -2.181271 | -1.578957 | -2.10326 | -3.21885 | -2.125902 | -1.084174 | -1.245476 | -1.047618 | -0.723267 | -0.704614 | -0.072761 | -1.542977 | -1.424684 | -0.976485 | -1.07171 | -1.33574 | | 35.13888889 | -1.148574 | 0.099091 |
| 35.055556 | -8.351592 | -2.855825 | -2.12257 | -0.524435 | -0.633146 | 0.423631 | -0.150058 | -2.793988 | -2.793908 | -2.072983 | -2.96533 | -2.970905 | -2.423065 | -1.094135 | -1.08539 | -0.778757 | -0.672396 | -0.337671 | -0.066447 | -1.474699 | -1.431139 | -0.850905 | -1.020692 | -1.370576 | | 35.15277778 | 0.505255 | -0.494151 |
| 35.069444 | -8.114233 | -3.749545 | -4.031236 | -4.068671 | -3.726269 | -0.956812 | -1.073362 | -2.218844 | -3.20707 | -2.62064 | -4.398843 | -2.725972 | -4.260739 | -1.198319 | -0.91874 | -0.672796 | -0.65129 | -0.072218 | 0.070321 | -1.478646 | -1.546629 | -0.675956 | -0.874276 | -1.09068 | | 35.16666667 | 2.152693 | -0.97102 |
| 35.083333 | -6.023204 | -3.256607 | -5.14099 | -6.029127 | -5.298342 | -3.017472 | -1.633261 | -1.969219 | -3.423241 | -3.000654 | -5.25177 | -2.530174 | -5.861686 | -1.303626 | -1.108565 | -0.685479 | -0.631458 | -0.072963 | 0.003072 | -1.371697 | -1.551689 | -0.710371 | -0.813833 | -1.174218 | | 35.18055556 | 3.047182 | -0.678421 |
| 35.097222 | -3.852014 | -2.394213 | -3.673122 | -6.22973 | -4.341554 | -5.038275 | -1.984084 | -2.234111 | -2.101572 | -2.786669 | -4.530192 | -2.355062 | -5.884568 | -1.362432 | -1.493497 | -0.58588 | -0.450004 | -0.25299 | -0.302824 | -1.293433 | -1.505863 | -0.887047 | -0.877099 | -1.446929 | | 35.19444444 | 3.070732 | 0.238691 |
| 35.111111 | -2.097313 | -1.29058 | -2.022539 | -5.99392 | -2.982681 | -6.004893 | -1.495579 | 0.592626 | 0.186301 | -1.651993 | -2.929132 | -1.959905 | -4.692668 | -1.371271 | -1.570707 | -0.489415 | -0.448219 | -0.572621 | -0.77928 | -1.503624 | -1.736958 | -1.04123 | -1.045411 | -1.539703 | | 35.20833333 | 2.324708 | 1.158361 |
| 35.125 | -0.794033 | -0.204666 | -1.665962 | -4.158237 | -1.731637 | -5.419574 | -0.689826 | 0.586362 | 0.625988 | -0.776235 | -2.155117 | -1.93944 | -3.115849 | -1.317707 | -1.288417 | -0.577641 | -0.935288 | -1.035121 | -1.322516 | -1.641907 | -1.901856 | -1.127916 | -1.237032 | -1.509005 | | 35.22222222 | 1.122725 | 1.804279 |
| 35.138889 | 0.905646 | 0.926708 | -1.493829 | -2.096892 | -0.816908 | -3.88645 | -0.597854 | -2.296398 | -0.428311 | -0.100707 | -2.598196 | -1.932462 | -2.151015 | -1.269875 | -0.810617 | -0.659534 | -1.458954 | -1.407743 | -1.734107 | -1.60879 | -1.902017 | -1.069238 | -1.332948 | -1.284052 | | 35.23611111 | -0.083495 | 1.658759 |
| 35.152778 | 2.694355 | 1.162123 | -1.364944 | -1.156559 | -0.361 | -3.042004 | -0.711902 | -3.540559 | -0.899512 | 0.270775 | -2.148835 | -1.194809 | -1.464102 | -1.209395 | -0.275223 | -0.557317 | -1.394494 | -1.31421 | -1.61882 | -1.620255 | -1.922397 | -0.942153 | -1.08419 | -0.85018 | | 35.25 | -1.166463 | 0.704156 |
| 35.166667 | 4.354302 | 1.225525 | 0.114327 | 0.783554 | 1.122483 | -1.035667 | 0.095468 | -0.44972 | -0.449592 | 0.448648 | -0.628378 | -0.125825 | 0.259887 | -1.199917 | -0.064605 | -0.64159 | -1.033394 | -0.78511 | -1.029622 | -1.657891 | -1.879055 | -0.748776 | -0.562202 | -0.628689 | | 35.26388889 | -0.470458 | -0.513237 |
| 35.180556 | 4.693323 | 0.947066 | 0.908879 | 2.333206 | 2.557119 | 1.251797 | 1.25774 | 2.100124 | 0.880926 | 0.731689 | 1.056188 | 0.918101 | 2.030086 | -1.061985 | 0.284522 | -0.730297 | -0.778326 | -0.068701 | -0.302507 | -1.655027 | -1.641227 | -0.58255 | -0.013332 | -0.489293 | | 35.27777778 | 2.230508 | -0.705332 |
| 35.194444 | 4.187718 | 0.592531 | 0.576291 | 1.97251 | 2.404092 | 2.784151 | 1.494308 | 2.292296 | 1.888701 | 1.038046 | 1.585732 | 1.582667 | 2.279062 | -1.05093 | 0.659819 | -0.588233 | -0.706485 | 0.382736 | 0.195873 | -1.431688 | -1.181736 | -0.461369 | 0.293025 | -0.160022 | | 35.29166667 | 3.994666 | 0.179076 |
| 35.208333 | 3.88469 | 0.645358 | 0.446019 | 0.987155 | 1.865109 | 2.729113 | 1.16405 | 1.849899 | 1.88247 | 1.112542 | 1.626766 | 1.665353 | 1.705361 | -0.8478 | 1.243673 | -0.290205 | -0.537958 | 0.613686 | 0.478463 | -1.096381 | -0.764312 | -0.28074 | 0.441911 | 0.408188 | | 35.30555556 | 3.680862 | 1.145415 |
| 35.222222 | 3.754237 | 0.394621 | 0.620412 | 1.047797 | 2.082218 | 2.100414 | 1.032469 | 1.624838 | 1.386234 | 1.133234 | 1.849356 | 1.519791 | 1.430025 | -0.419793 | 1.826295 | -0.265798 | -0.408207 | 0.815524 | 0.690405 | -0.893613 | -0.577398 | 0.093615 | 0.573318 | 0.785709 | | 35.31944444 | 1.878912 | 1.421689 |
| 35.236111 | 2.310895 | 0.045397 | 0.856013 | 1.768868 | 2.581498 | 2.065669 | 1.06479 | 0.933119 | 0.727768 | 1.370043 | 2.035558 | 1.635555 | 1.609109 | -0.068696 | 1.85189 | -0.561899 | -0.372897 | 0.89654 | 0.807163 | -0.732269 | -0.444483 | 0.28903 | 0.679088 | 0.675389 | | 35.33333333 | 0.309014 | 0.800764 |
| 35.25 | 0.271383 | 0.147119 | 0.471317 | 1.798527 | 2.33674 | 3.245946 | 0.971003 | 0.715002 | 1.167122 | 2.231132 | 2.034634 | 1.799323 | 1.756746 | -0.444301 | 1.094993 | -0.56963 | -0.29597 | 0.837623 | 0.860112 | -0.54948 | -0.183608 | 0.167026 | 0.613913 | 0.215525 | | 35.34722222 | 0.203285 | 0.038414 |
| 35.263889 | -0.405411 | 1.248525 | 0.816389 | 1.937517 | 2.141779 | 5.846334 | 1.555337 | 2.148396 | 2.820173 | 3.034495 | 2.476347 | 1.91453 | 2.369002 | -0.561062 | 0.674512 | -0.413311 | 0.04299 | 0.927058 | 1.091681 | -0.277958 | 0.170535 | -0.026601 | 0.507268 | 0.030209 | | 35.36111111 | 1.450582 | 0.588577 |
| 35.277778 | 0.453126 | 2.309568 | 1.720017 | 3.206181 | 2.40809 | 9.590811 | 2.546461 | 2.74297 | 3.199609 | 3.314488 | 3.676107 | 1.936853 | 3.594656 | -0.114105 | 1.025353 | -0.085151 | 0.434517 | 1.572152 | 1.797721 | 0.142674 | 0.555081 | 0.025661 | 0.478895 | 0.620863 | | 35.375 | 3.363388 | 1.449178 |
| 35.291667 | 1.780135 | 2.089727 | 2.246846 | 3.969196 | 2.439696 | 10.533051 | 3.348435 | 2.718004 | 2.729185 | 3.477047 | 4.86965 | 2.078078 | 4.634731 | 0.587453 | 1.499477 | 0.135386 | 0.65464 | 2.299554 | 2.732371 | 0.415301 | 0.770361 | 0.137917 | 0.403133 | 1.24511 | | 35.38888889 | 4.483599 | 1.862721 |
| 35.305556 | 3.221312 | 1.635036 | 2.511719 | 3.647288 | 2.132913 | 8.022785 | 3.221067 | 1.076791 | 2.315351 | 4.009183 | 5.111912 | 2.478525 | 4.683387 | 1.154089 | 1.372261 | 0.305423 | 0.798923 | 2.868407 | 3.595394 | 0.464358 | 0.865626 | 0.148948 | 0.323525 | 1.414374 | | 35.40277778 | 3.843499 | 1.754822 |

| | | | | | | | | | | | | | | | | | | | | | | | | | | | | |
|---|---|---|---|---|---|---|---|---|---|---|---|---|---|---|---|---|---|---|---|---|---|---|---|---|---|---|---|---|
| 37.888889 | 1.484999 | -7.498397 | 0.51427 | 0.122602 | -2.76106 | -1.879043 | -1.123768 | -0.443772 | -2.812192 | -5.733143 | -5.46204 | -3.016174 | -12.129541 | -1.595081 | -0.25209 | -0.59053 | -1.03534 | -1.360948 | -3.177501 | -4.038222 | -1.714596 | -4.479727 | -5.677631 | -7.346539 | | 37.98611112 | -4.631411 | -1.455759 |
| 37.902778 | -2.272488 | -10.121343 | -0.833626 | -2.039263 | -5.386098 | -1.411722 | -3.866164 | -1.239183 | -9.499935 | -10.447705 | -6.377131 | -3.533878 | -22.860094 | -2.427089 | -3.072176 | 0.054804 | -0.710935 | 0.183976 | -0.828982 | -4.608636 | -1.174574 | -2.153533 | -2.640893 | -4.865954 | | 38.00000001 | -7.589501 | -1.336154 |
| 37.916667 | -11.968421 | -10.665056 | -3.211413 | -6.60445 | -8.01705 | -1.524654 | -4.189276 | -4.564 | -12.194333 | -11.566422 | -7.376033 | -4.072867 | -26.457559 | -3.579741 | -5.726833 | -0.792244 | -0.566795 | -1.864666 | -2.455498 | -4.841726 | -2.256021 | -0.072938 | 0.624983 | -6.229523 | | 38.0138889 | -8.444878 | -1.07209 |

**Supplementary Table S7 (k)** Mean day-to-day motion rate in Control and Stress tanks of Tomato measured under dimGday/night and RGBday dimGnight, calculated using either 24 h motion integration or 22 h motion integration excluding day/night transitions.

| Row | dimG22Control | dimG22Stress | RGB22Stress | RGB22Control | RGBStress | RGBControl | dimGControl | dimGStress |
|---|---|---|---|---|---|---|---|---|
| 18.5 | 0.411659945 | 0.083862023 | 0.2475837 | 0.632238543 | 1.442081257 | 2.998852565 | 0.404396759 | 0.085667411 |
| 19.5 | 0.160611025 | 0.126969294 | 0.337891244 | 0.423781964 | 1.341851244 | 2.825103819 | 0.224511035 | 0.141608168 |
| 20.5 | 0.306309136 | 0.163212343 | 0.296915559 | 0.63195615 | 0.72921816 | 3.388928366 | 0.345430893 | 0.179864273 |
| 21.5 | 0.3925127 | 0.256798167 | 0.408886872 | 0.674854783 | 0.61103435 | 3.642023584 | 0.494981961 | 0.322058956 |
| 22.5 | 0.643896555 | 0.418982574 | 0.699636734 | 0.857272302 | 1.025030359 | 3.515162406 | 0.748682408 | 0.5390435 |
| 23.5 | 0.800534679 | 0.587836934 | 0.8815553 | 0.948219253 | 1.268116011 | 3.368079724 | 1.025945132 | 0.653165183 |
| 24.5 | 1.100500891 | 0.792721948 | 1.283797168 | 1.175972969 | 2.152471791 | 3.10891236 | 1.307576125 | 0.92849965 |
| 25.5 | 1.370302604 | 1.001983273 | 1.586795794 | 1.451482799 | 2.68381231 | 3.029718842 | 1.758498587 | 1.186534492 |
| 26.5 | 1.820803282 | 1.185933263 | 1.800456887 | 1.766586636 | 3.427382952 | 2.897103732 | 2.248888225 | 1.454863888 |
| 27.5 | 2.387114027 | 1.378535838 | 1.934155466 | 2.325413981 | 4.027186533 | 3.227347867 | 2.837375299 | 1.757900756 |
| 28.5 | 2.876462196 | 1.727779803 | 2.09666563 | 2.849641585 | 4.490020228 | 3.460953144 | 3.438940756 | 2.25855075 |
| 29.5 | 3.022420914 | 2.025312483 | 2.544953737 | 2.968206257 | 5.457992186 | 3.697678266 | 3.698421272 | 2.650331749 |
| 30.5 | 3.39414198 | 2.495530336 | 3.426847006 | 3.435964894 | 6.539776254 | 4.7009595 | 4.177569515 | 3.248608875 |
| 31.5 | 3.880419497 | 2.188172703 | 2.635851689 | 4.105890838 | 6.095868223 | 5.816788415 | 4.772278779 | 3.111141469 |
| 32.5 | 4.326852482 | 2.821033089 | 3.462356469 | 4.744484125 | 6.536949157 | 7.458432676 | 5.639095321 | 3.805350476 |
| 33.5 | 4.603641195 | 3.001492765 | 3.428504573 | 4.900690708 | 6.388311272 | 8.274959018 | 6.137448277 | 4.110794109 |
| 34.5 | 5.544745285 | 3.397404272 | 3.024776193 | 5.909639524 | 5.272445312 | 9.882655266 | 7.359947067 | 4.394971139 |
| 35.5 | 6.93563027 | 3.461039892 | 3.358689533 | 7.205522214 | 5.728062561 | 10.91237843 | 8.887767979 | 4.575205872 |
| 36.5 | 7.461745542 | 3.343566187 | 4.308073136 | 7.16235349 | 5.781283935 | 9.362239534 | 8.698898686 | 4.167020971 |

**Supplementary Table S7 (l)** Mean day-to-day motion rate of individual Tomato plants under dimGday/night and RGBday dimGnight. Data represent means of 9–10 plants

| Interval | RGBMeanControl | RGBSEMControl | RGBMeanStress | RGBMStress | dimGMeanControl | dimGSEMControl | dimGMeanStress | dimGSEMStress |
|---|---|---|---|---|---|---|---|---|
| 18.5 | 0.330086984 | 0.051125966 | 0.340512478 | 0.056839297 | 0.268859961 | 0.038375869 | 0.207764348 | 0.037077777 |
| 19.5 | 0.40209644 | 0.045969353 | 0.4309264 | 0.081045225 | 0.360932706 | 0.040814785 | 0.371078959 | 0.070974056 |
| 20.5 | 0.54059503 | 0.059474136 | 0.542169894 | 0.082565638 | 0.54035336 | 0.054905837 | 0.532916132 | 0.075920116 |
| 21.5 | 0.498262687 | 0.049106899 | 0.630202006 | 0.083784601 | 0.473690579 | 0.038427714 | 0.611817547 | 0.070562049 |
| 22.5 | 0.684357387 | 0.076755927 | 0.768334047 | 0.086835431 | 0.746559779 | 0.063987173 | 0.790691671 | 0.077145023 |
| 23.5 | 0.869432993 | 0.068349351 | 0.753295092 | 0.092441857 | 0.976811375 | 0.064342704 | 0.752273425 | 0.085375273 |
| 24.5 | 1.073363276 | 0.071808229 | 0.911785954 | 0.095419105 | 1.14628576 | 0.067106863 | 0.894490812 | 0.083416637 |
| 25.5 | 1.265269504 | 0.053567906 | 1.009621174 | 0.128098219 | 1.336409702 | 0.070161201 | 0.960144801 | 0.116065022 |
| 26.5 | 1.369540643 | 0.076175802 | 1.108278691 | 0.121397328 | 1.430376148 | 0.077259884 | 1.056608366 | 0.106008835 |
| 27.5 | 1.602816709 | 0.080924512 | 1.20518364 | 0.129092343 | 1.664769652 | 0.08007946 | 1.150598666 | 0.127536042 |
| 28.5 | 1.739481384 | 0.093151706 | 1.333628577 | 0.14519965 | 1.780230397 | 0.091836449 | 1.244744535 | 0.138103595 |
| 29.5 | 1.809899499 | 0.09233038 | 1.381658083 | 0.163872588 | 1.859444323 | 0.084994543 | 1.31003894 | 0.182889392 |
| 30.5 | 2.043669278 | 0.120432772 | 1.587735104 | 0.22675308 | 2.03606315 | 0.114200031 | 1.504065594 | 0.228420128 |
| 31.5 | 2.35187113 | 0.124902595 | 1.663124268 | 0.242163131 | 2.276110462 | 0.106887054 | 1.56302429 | 0.236033219 |
| 32.5 | 2.891150445 | 0.180956988 | 1.820386478 | 0.304812953 | 2.835898599 | 0.176584283 | 1.706357187 | 0.295163393 |
| 33.5 | 3.141275204 | 0.291462532 | 1.715157257 | 0.239582992 | 3.053675755 | 0.272549358 | 1.615010176 | 0.228489507 |
| 34.5 | 3.889159075 | 0.296531508 | 1.655780055 | 0.130215654 | 3.726381279 | 0.301721152 | 1.537006631 | 0.132574385 |
| 35.5 | 4.315266162 | 0.342251561 | 1.805492013 | 0.163377647 | 4.162676849 | 0.322094133 | 1.725211361 | 0.166759693 |
| 36.5 | 3.713967897 | 0.226805959 | 1.58710932 | 0.164364602 | 3.569032709 | 0.219197884 | 1.552341315 | 0.159783026 |
